# Conserving genetic diversity during climate change: Niche marginality and discrepant monitoring capacity in Europe

**DOI:** 10.1101/2023.03.24.533448

**Authors:** Peter B. Pearman, Olivier Broennimann, Tamer Albayrak, Paulo Célio Alves, Laura D. Bertola, Aleksandra Biedrzycka, Elena Buzan, Vlatka Cubric-Curik, Ancuta Fedorca, José A. Godoy, Christina Hvilsom, Peter Klinga, Maciej K. Konopiński, Alexander Kopatz, Linda Laikre, Margarida Lopez Fernandez, Joachim Mergeay, Charalambos Neophytou, Snæbjörn Pálsson, Ivan Paz-Vinas, Diana Posledovich, Barbora Rolečková, Dainis Ruņģis, Gernot Segelbacher, Katja Kavčič Sonnenschein, Henrik Thurfjell, Sabrina Träger, Cristiano Vernesi, Carles Vilà, Marjana Westergren, Frank E. Zachos, Antoine Guisan, Michael Bruford

## Abstract

Genetic monitoring of populations currently attracts interest in the context of the Convention on Biological Diversity but needs long-term planning and investments. Genetic diversity has been largely neglected in biodiversity monitoring, and when addressed is treated separately, detached from other conservation issues, such as habitat alteration due to climate change. Genetic monitoring supports the conservation and management of fisheries, game, and threatened populations. It also can contribute to the assessment of predicted and realized impacts of climate change, and their management. We report the first accounting of genetic monitoring efforts among countries in Europe (their ‘genetic monitoring capacity’, GMC) to determine where GMC suggests the combination of national infrastructure, political support and resources for continued and expanded monitoring. Overlaying GMC with areas where species ranges approach current and future climate niche limits (i.e., niche marginality) helps identify whether GMC coincides with anticipated climate change effects on biodiversity. Our analysis suggests that country area extent, financial resources, and conservation policy influence GMC, high values of which inconsistently match joint species patterns of climate niche marginality. Populations at niche margins likely hold genetic diversity that is important to adaptation to changing climate, and our results illuminate the need in Europe for expanded genetic monitoring across the climate gradients occupied by species, a need arguably greatest in southeastern European countries.

---

Maintenance of wild population genetic diversity is an important component of the Convention on Biodiversity (CBD) ^1^, but it has received little international attention until recently^1–4^, reducing our ability to monitor and manage wild populations to sustain population genetic diversity^5^. The resulting urgent need for expanded monitoring of population genetic diversity (PGD) motivates development of globally implementable indicators of genetic diversity^6–9^, some of which are included in the recently-adopted CBD Kunming-Montreal Global Biodiversity Framework^3,10^. But while ongoing anthropogenic loss of PGD is being documented^11–13^, efforts to detect climate change effects on PGD are taxonomically and geographically limited^14,15^, and absent from international biodiversity agreements. Populations in extreme climatic conditions, such as those near their climatic niche margins, are particularly relevant to species potential for adaptation to changing climate^16^. Nonetheless, multi-species patterns of populations near to niche margins, as indicators of adaptive potential and, thus, possible PGD monitoring sites, remain unidentified. This calls for improved accounting of the relationship between species limits along environmental gradients and associated PGD^17,18^.

Species populations close to their environmental niche margin may differ genetically from those at the niche center, and influence the course of adaptation to changing environment^19,20^. Evidence shows that populations at niche margins toward stressful environmental extremes are locally adapted^21^, having distinguishable genetic architecture independent of their geographic position within the species range^22^. Populations near warm/dry niche limits likely hold important adaptive genetic diversity^22–24^ that can reduce predicted range loss^18,25^, and contribute to adaptation in environmentally central populations^26^ to warming, drying climate, despite greater gene flow from niche center to these marginal populations^27^. Nonetheless, genetic diversity held in marginal populations may be endangered when geneflow to environmentally central areas is impeded^28^. These results suggest that global genetic monitoring frameworks^10^ need to anticipate climate impacts, collect samples across entire climate gradients, and evaluate the contributions of marginal populations to genetic diversity and adaptive potential^29^. However, no accounting of recent and historical PGD monitoring exists, leaving us ignorant of taxonomic, national, and geographic trends in monitoring effort, thus hampering our capacity to detect PGD and adaptive potential under climate change threat. Yet, even without such accounting, known PDG monitoring efforts suggest notable resources, infrastructure, and political support and can serve as an index of current and potential future ‘genetic monitoring capacity’ (GMC).

Here, we aim to fill this gap by asking: (1) How is GMC distributed across Europe and on which taxa has PGD monitoring focused? (2) Which factors explain among-country variation in GMC? (3) How will countries differ in climate change exposure of threatened species? Finally, (4) How does GMC coincide with anticipated impacts of climate change on habitat suitability for populations? Using evidence of monitoring from the peer reviewed and technical literature, we examine how countries in the European Commission’s Cooperation in Science and Technology (COST) program^30^ demonstrate GMC for purposes of biodiversity conservation and management. We explain variation in GMC in relation to two fundamental characteristics of countries, per capita Gross Domestic Product (GDP) and area extent. We then compare GMC to multi-species indicators of niche marginality and declining environmental conditions due to climate change. We use climate and biological data to stratify species ranges into climatic niche centrality and marginality areas. We then estimate impacts of climate change on the future geographic distribution of conditions near climatic niche margins^31^, range-wide and for four groups of species selected for recognized and potential conservation and management interest (amphibians, large birds, carnivorans, and forest trees). We estimate how climate change impacts on these species distribute among European countries, as indicated by present and future patterns of climate niche marginality, and compare it to the distribution of GMC among countries.

## Results

Between 22.11.2019 and 31.12.2021, we received 480 submissions of candidate monitoring projects from conservation geneticists, practitioners and stakeholders. We evaluated these for validity as Category II genetic monitoring^32^, which report temporally separate assessments of genetic diversity metrics of one or more populations of a species. We focus here exclusively on this type of genetic monitoring because it directly tracks PGD over time, while we also recognize that other genetic monitoring, including genetic assessments and sample identification programs, are also highly relevant to conservation, but address other questions. We found 38 additional candidate Category II monitoring projects through a structured search of the Web of Science. Of the total 518 candidates, we identify 103 as valid Category II monitoring projects, the vast majority of which report sampled populations from one (84) or two (14) countries. We tally international and transboundary projects separately by country, and we document a total of 151 national-level projects of Category II genetic monitoring. We find Category II monitoring in 31 of 38 COST countries that were full members at the beginning of data solicitation (Fig 1a, b).

**Figure 1.**
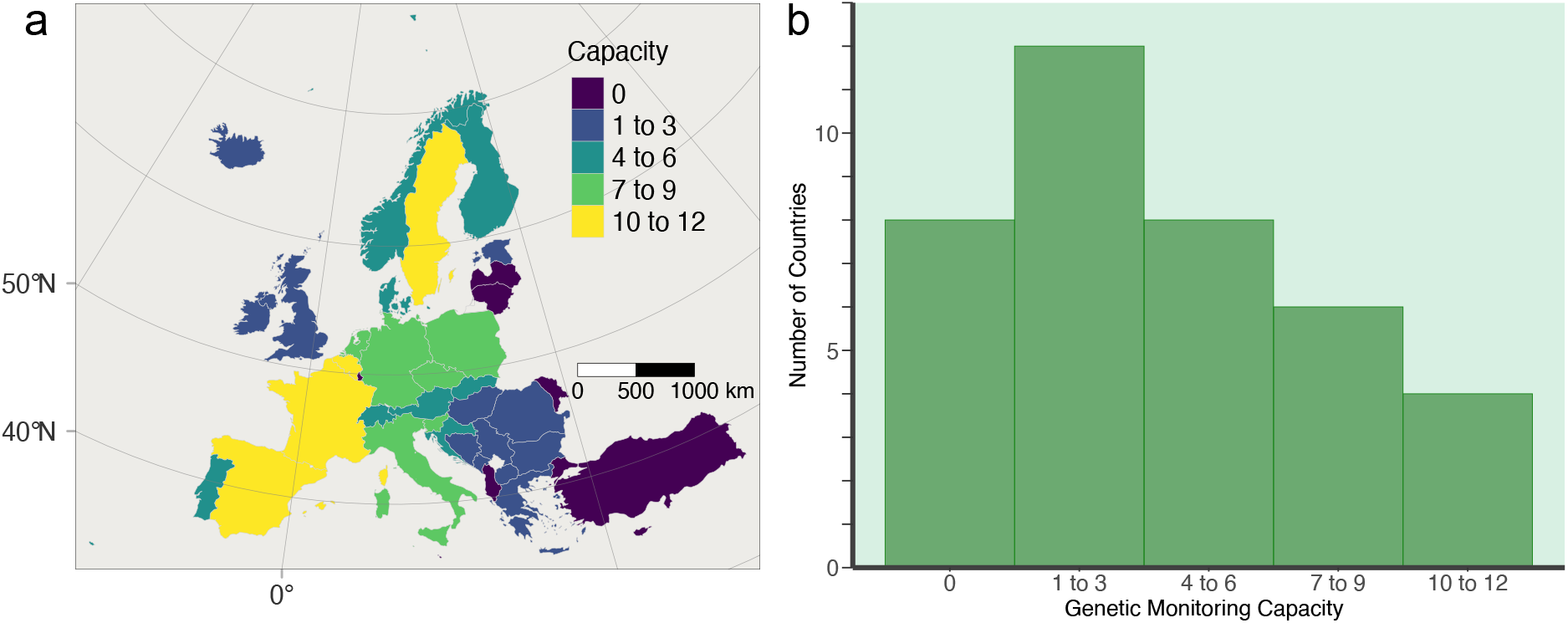
Documented programs to monitor population genetic diversity for conservation and management in COST member countries, as an indicator of genetic monitoring capacity, up to 31.12.2021. The geographic distribution of monitoring capacity to countries, as a tally across all domestic and wild terrestrial and marine species (a) indicates that countries with relatively high capacity for monitoring are found in both northern and southern Europe. COST countries in southeastern Europe present generally low genetic monitoring capacity. The distribution of programs to countries (b) shows that most countries have established six or fewer monitoring programs.

### Genetic monitoring capacity--GMC

To understand the patterns of GMC among countries, we examine the variation in the tally of Category II PGD monitoring projects among COST countries, and partition this indicator of GMC to taxonomic and functional groups. We find that GMC is not uniquely attributable to the geographic location of countries, although we generally find few PGD monitoring projects in southeastern Europe. European countries with few PGD monitoring projects (three or fewer) occur across a range of latitudes and present no striking north-south pattern (Fig. 1a, b). Countries with high GMC appear in both northern and southern Europe (Fig. 1a). We document a maximum of 12 projects for Belgium and Sweden, and 11 projects for Spain and France (Fig. 1a). We find no GMC in eight countries (Fig. 1b), including ones as geographically and economically disparate as Turkey and Luxemburg. Nonetheless, a majority of countries (31 of 38) demonstrated some GMC. This pattern is robust to the exclusive consideration of terrestrial wild species (i.e. exclusion of programs monitoring fish, marine species, and domesticated/captive populations; Extended Data Fig. 1, Appendix S1, Supplementary Materials).

The GMC of COST countries varies greatly by taxonomic and functional groups. For example, while many amphibians are of recognized conservation concern, only two European countries demonstrate GMC for amphibians (Belgium and Spain, Fig. 2a). Many more countries (9) have monitored PGD in at least one bird species (Fig. 2b, Extended Data Fig. 2b). Approximately half of COST countries (17) have monitored PGD in one or more large carnivorans (Fig. 2c, Extended Data Fig. 2c), although certain carnivorans are absent from some COST countries (Extended Data Fig. 3a-c). In contrast, while all COST countries have tree species, less than one quarter of COST countries (7) have monitored PGD in at least one of these species (Fig. 2d, Extended Data Fig. 2d).

**Figure 2.**
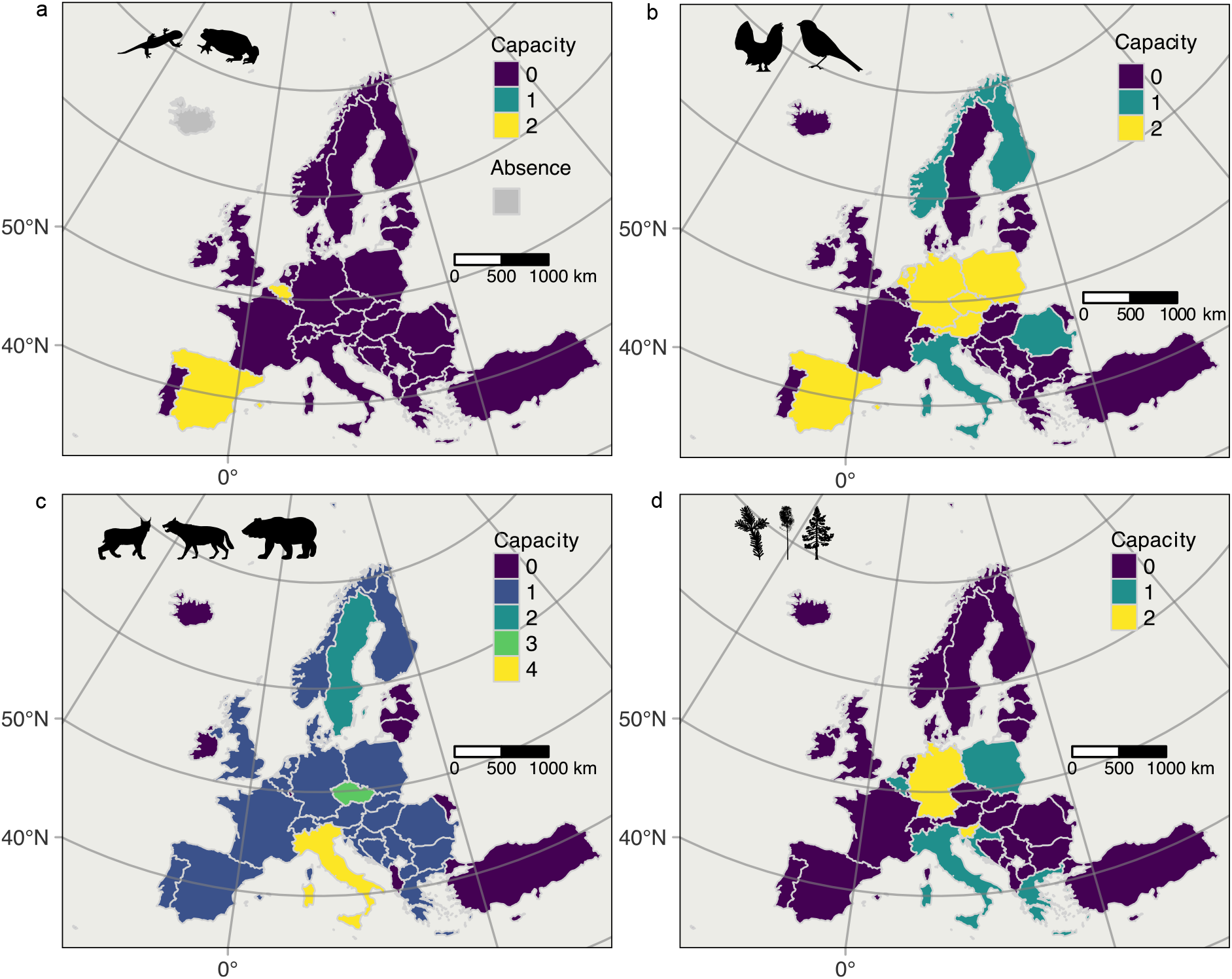
Geographic distribution of genetic monitoring capacity for population genetic diversity, for purposes of conservation or management, among COST full-member countries, showing the tally of programs for amphibians (a), birds (b), carnivorans (c), and forest trees (d). Programs included here are consistent with requirements for Category II monitoring, and they offer documentation of multiple estimates over time of at least one index of genetic diversity. Few countries have genetic monitoring capacity for amphibians, while most countries have established at least one program for a carnivoran species.

Inspection of the data on PGD monitoring programs for COST countries, especially the lower number of projects documented from countries in southeastern Europe, led us to ask whether fundamental geographical and economic data are consistent with this variation. We examine the relationship between GMC and both land area and recent GDP, and we present generalized linear models to test the form and significance of the relationships. Turkey is by far the largest COST country by area, and with almost 784,600 km^2^, it is 42% larger than the next largest country, France (excluding its overseas territories). With no documented PGD monitoring, Turkey is an outlier for its size and absence of GMC, and is an influential observation in statistical analysis. Omitting Turkey, other COST countries demonstrate that larger countries tend to have higher GMC (Fig. 3a, neg. binomial regression, P=0.02). In contrast, intermediate GDP is associated with greater GMC (Fig. 3b, binomial regression, GDP quadratic term P=0.003; model pseudo-R^2^ = 0.47; Appendix S2, Supplemental Materials). Substantial residual variation remains, with Finland, the United Kingdom, and Norway having fewer projects than expected, and Belgium and Sweden more projects, in relation to both size and GDP (Fig 3b). The negative quadratic relationship of GMC with GDP remains statistically significant despite the potential omission of data from any single potential outlier or extreme value.

**Figure 3.**
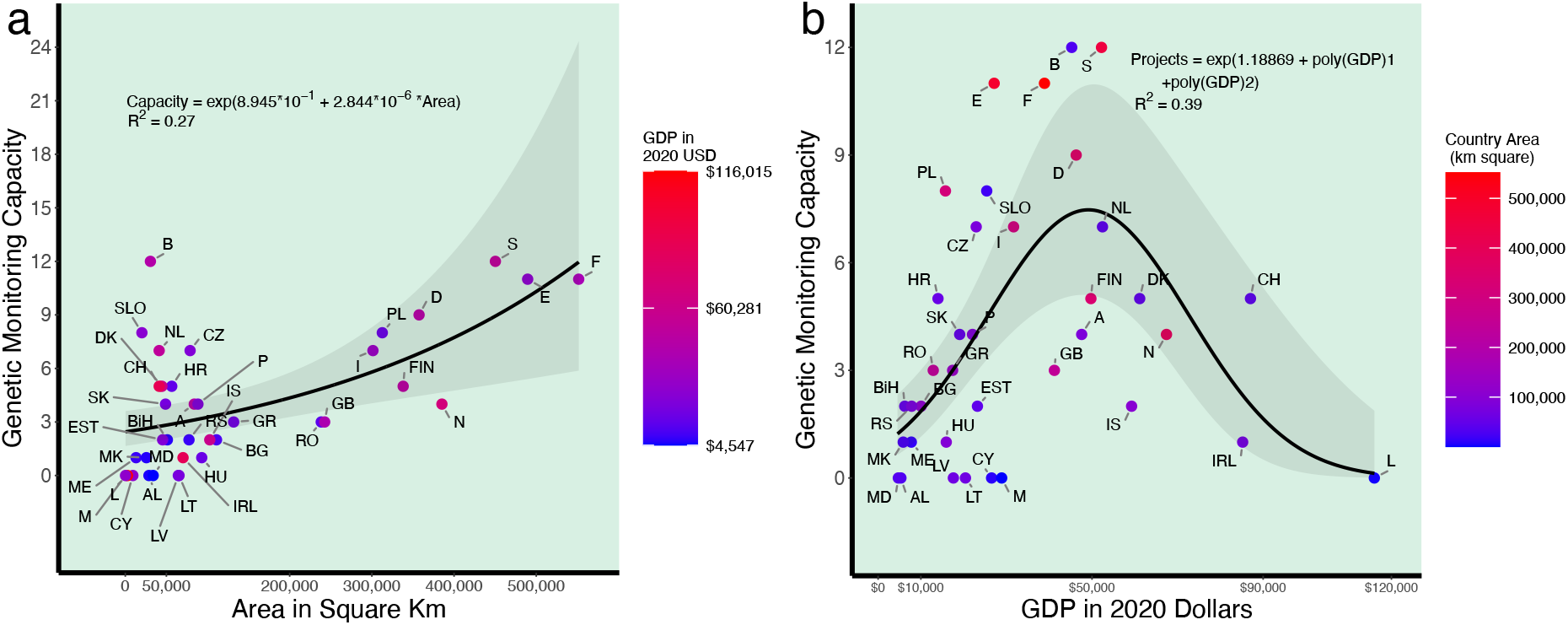
Generalized linear models for the genetic monitoring capacity of COST full-member countries, represented by international postal codes, as a function of (a) area extent and (b) average per capita gross domestic product (GDP). Equations of the lines are shown, along with 95% confidence intervals in shading. Models were fit as negative binomial distribution with the log link function. Model fit is given as Veall-Zimmermann R^2^. Turkey is of substantially greater geographic extent than the displayed countries, yet has no documented genetic monitoring capacity and is omitted as an outlier and influential observation. Both the linear area term and the quadratic GDP term are significant in the multiple generalized model (area: P < 0.018; GDP quadratic: P<0.002; see online Methods for details). A significant quadratic term remains upon omission of any one of the three high-GDP countries.

### Joint environmental niche marginality framework

To integrate PGD monitoring into a framework for addressing climate change impacts, we evaluate the relationship between GMC and expected climate change effects on species climatic niche marginality at the national level. Countries with a relatively large GMC should be well prepared to evaluate climate impacts on genetic diversity. These countries have much relevant infrastructure (i.e., genetic laboratories) and some aspects of adapting monitoring programs to detect effects of climate change are technically simple, such as expanding sampling to cover climate gradients. In contrast, countries with relatively little GMC and substantial predicted, climate-driven decline in habitat suitability likely present opportunities for focused development of PGD monitoring capacity, to help predict, evaluate and manage climate impacts on important populations. We calculated for each species separately an index of climate niche marginality^31^, based on variation in climate across the entire species range, as an indicator of marginally suitable habitat for the species (but not necessarily for other species). Areas at niche margins within species ranges are those areas that coincide with the most marginal 25% of climatic conditions within the species global range, while areas in the rest of the species range experience core climatic conditions. We present the joint geographic distribution of niche marginality across a total of 185 species, spread across amphibians (44 Anura, 26 Caudata), large birds (16 species in the Accipitridae, Anatidae, Gallidae and Otididae), carnivorans (eight species), and forest trees (91 species), which are of current or potential future conservation or management interest (Extended Data Table 1).

Species vary in range size and geographic location, with the result that current and future distributions of niche marginality conditions for groups of species are diverse and complex (Appendices S3-S6, Supplementary Materials). Patterns of current joint niche marginality vary greatly among the four study groups, with foci of joint niche marginality in the Iberian Peninsula (amphibians, large birds and forest trees), central Turkey (large birds), coastal areas in southeastern Europe (forest trees), and the Carpathian Mountains (amphibians, forest trees; Fig. 4). Increases and decreases in the total number of study species with populations at niche margins vary broadly across COST countries (compare Figs. 5a, b). Assuming that species climatic niches remain stable in time, we predict decreases in the number of species with marginal habitat in France, Italy, and Turkey, but increases in Bulgaria, Hungary, and Poland. Spatiotemporal trends in niche marginality in the four groups of species also differ substantially among COST countries (Extended data Fig. 4, Appendices S3-S6, Supplementary Materials). For example, many amphibian species are endemic to Europe or nearly so (Appendix S3, Supplementary Materials). European endemic amphibian species inhabit areas at niche margins in both the northern, higher (cool) and southern, lower (warm) portions of their ranges (Appendix S3, Supplementary Materials). This is also true for a group of large European birds (Appendix S4, Supplementary Materials).

**Figure 4.**
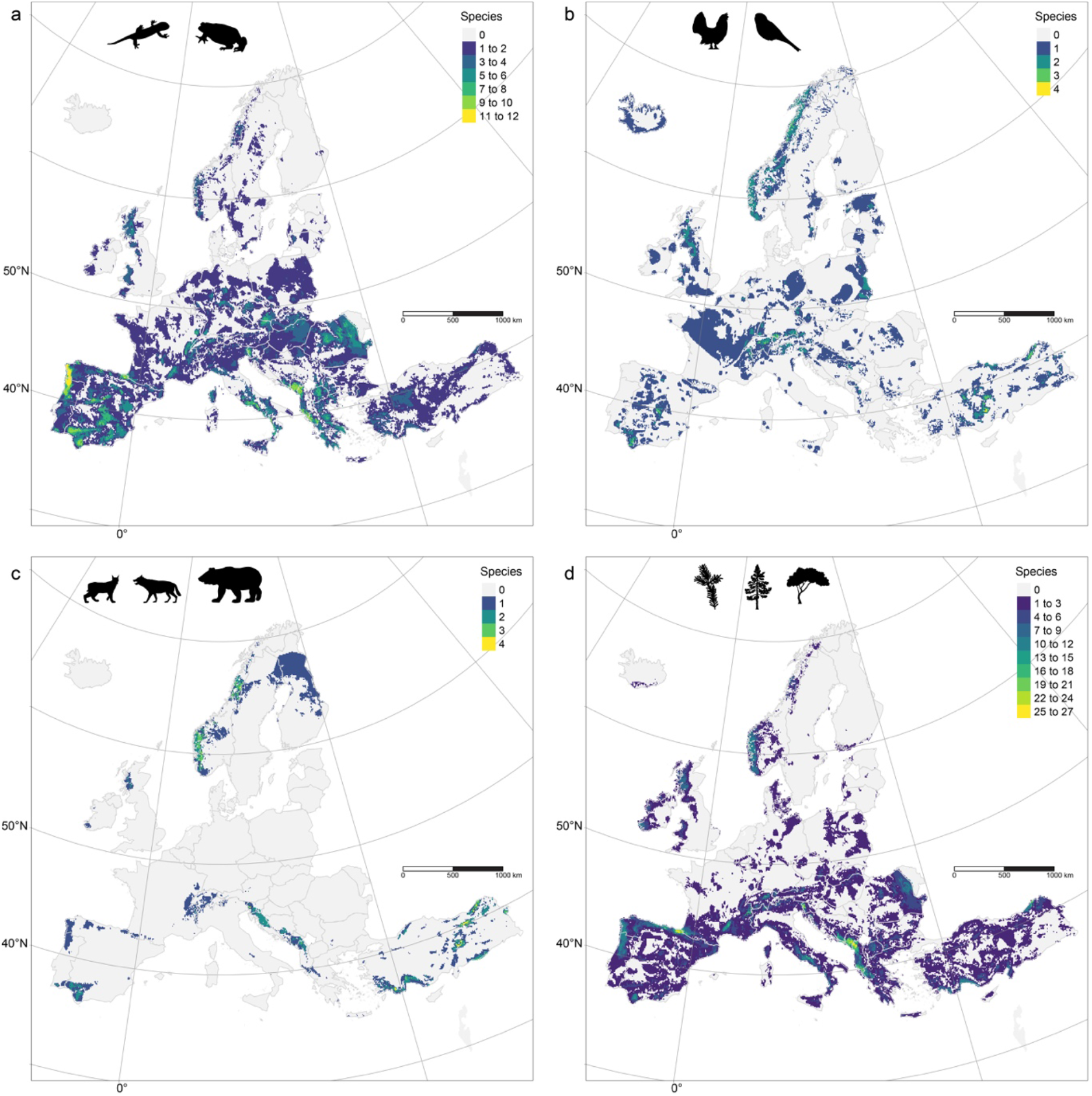
Current joint marginality of four groups of species. The colored areas represent the tally of species that have marginal climatic conditions in each pixel, for (a) amphibians, (b) a collection of relatively large birds, (c) large European carnivorans and (d) a set of forest tree species. Pixels with marginal niche conditions are among the 25% most climatically marginal across the global range of each species. Pixels are aggregated to 100 km^2^ to improve visualization, and the highest value within this area is displayed.

Comparison of current and future distributions of niche marginality in individual species often indicates increasing area of environmental marginality, but not always in the southern portion of species ranges (Appendices S3, S4, and S6, Supplementary Materials). In contrast, species with only a small portion of their range in COST countries, such as wolverine (*Gulo gulo*) and brown bear (*Ursus arctos*), show little change in distribution of habitat at climatic niche margins in COST countries (Appendix S5, Supplementary Materials). Across the four groups, the number of species with habitat at niche margins in each country is similar between current and future time periods (Fig. 5a, b). Nonetheless, we predict that future numbers of species with habitat at climatic niche margins will decline in some countries, while increasing in others. For example, the number of species of amphibians with climate conditions at niche margins increases in central Europe (Extended Data Fig. 4a, b) but the number of large bird species with niche margin conditions decreases in this region, as well as in France and Italy (Extended Data Fig. 4c, d). We predict that the number of carnivorans that experience climates at niche margins decreases in some Nordic countries and in Spain (Extended Data Fig. 4e, f), providing no evidence of a north-south trend in changing niche marginality in Europe for this taxon. The data suggest that the number of tree species experiencing niche margin conditions will increase in some countries in southeastern Europe, decrease in others, and decrease in both Spain and Italy (Extended Data Fig. 4g, h). Regional trends in niche marginality are also visible at the pixel level, at which national trends are more difficult to visualize (Extended Data Figure 5).

**Figure 5.**
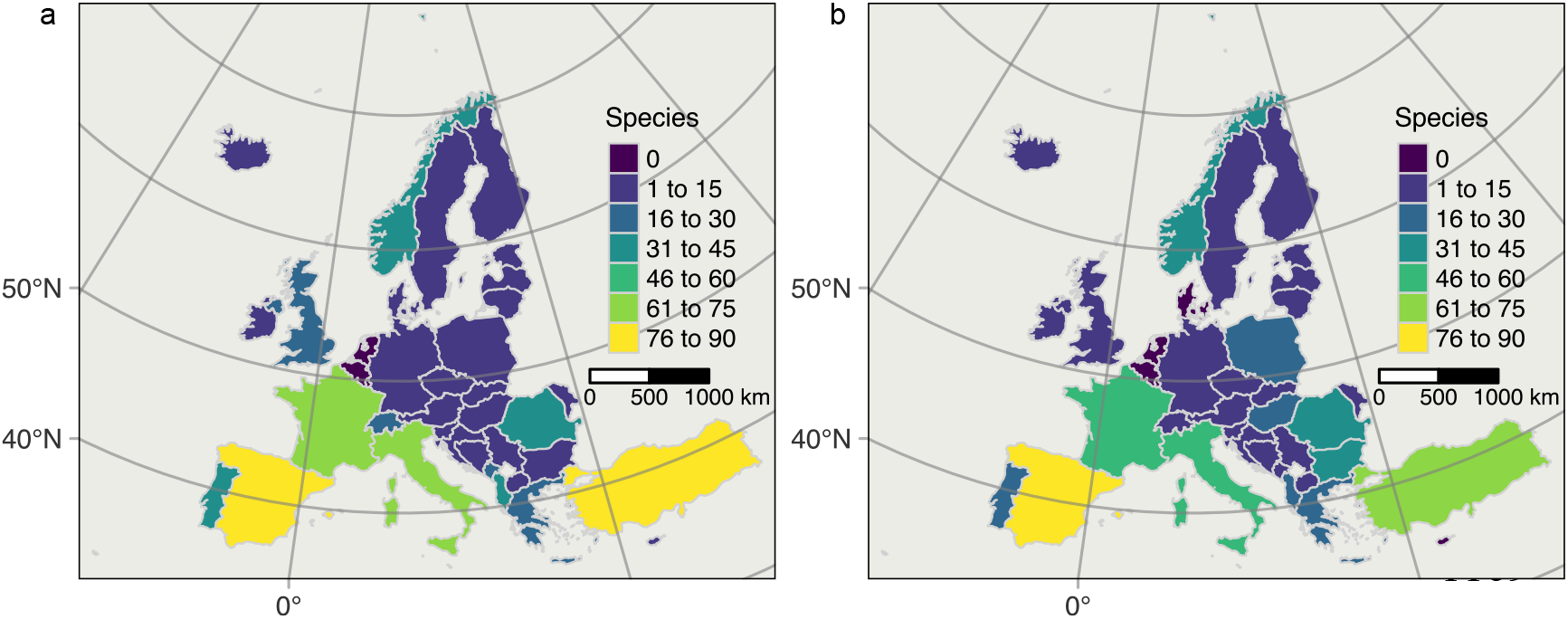
The number of species with marginal niche situations currently (a) and predicted on the basis of climate averaged over the interval 2041-2070 (b). A species is included in the tally for marginal species in a country whenever the country has at least five percent of the total niche marginal pixels for the species across all COST countries. This tally includes selected species of amphibians, large birds, large carnivorans and forest trees. As presented here, forest trees drive the differences among countries because this is the largest group, with 91 species.

Future numbers of species with habitats at niche margins and GMC vary greatly among countries, but show no linear relationship (Fig. 6). Generally, we predict countries with larger geographical extent have more species with niche margin habitat, and have greater GMC, with the exception of Turkey. This country rivals Spain in likely having many species with niche margin habitat in the future, but lacks documented GMC (Fig. 6).

**Figure 6.**
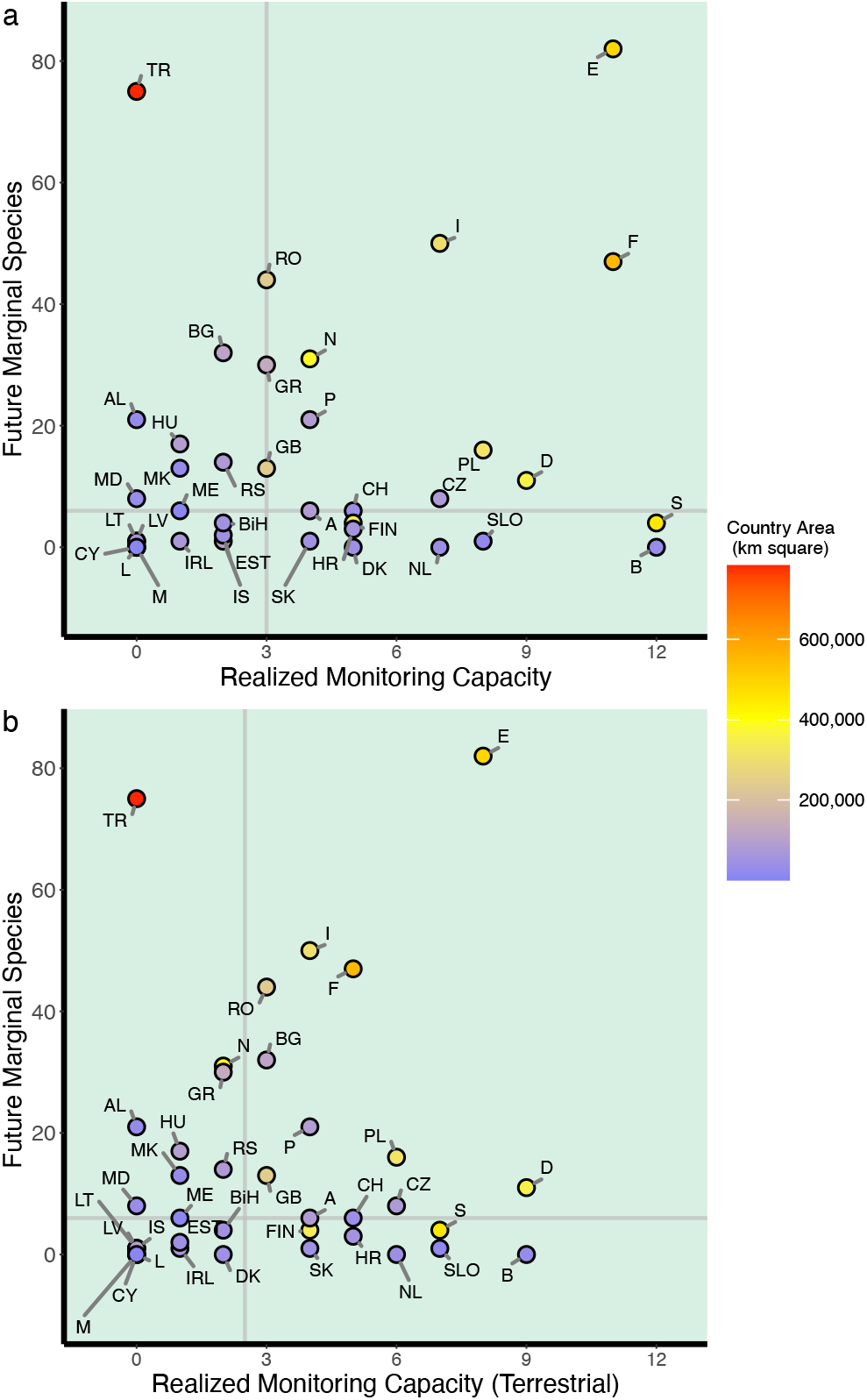
The relationship between genetic monitoring capacity and the number of species with marginal climatic niche conditions as of the years 2041-2070, showing data for all Category II monitoring as an indicator of genetic monitoring capacity at the national level (a) and programs monitoring amphibian, avian, carnivoran, and plant species only (b). Countries are indicated by postal codes. Marginal species include all species chosen for calculation of marginality, including non-troglobite amphibians, a collection of large birds, selected large carnivorans, and a set of forest trees. No general linear trends exist, although there is substantial variation both in numbers of species in marginal niche situations and in genetic monitoring capacity of countries.

## Discussion

Contrary to our expectations, areal extent of countries does not generally account for variation in GMC. Only by excluding Turkey as an outlier do we observe a positive relationship between country area and GMC. Turkey produces population genetic research but is not a member of the EU. The reporting requirements of the EU Habitat and Birds Directives may successfully promote the use of Category II genetic monitoring. In contrast, and in line with our expectations, countries with relatively low per capita GDP generally had lower GMC. However, it appears that countries with intermediate GDP have on average the highest GMC. Countries with high GDP are in many cases relatively small (Fig. 3), and many factors conceivably influence the establishment of monitoring programs, regardless of country size or per capita GDP. Extensive exploration of country characteristics that influence the establishment of PGD and other monitoring programs is beyond the scope of this paper, but could be explored in future research.

Our data collection on genetic monitoring projects was designed to capture reports in the scientific literature as well as unpublished and unreleased technical documents. Our results likely represent the distribution of such monitoring programs and GMC in Europe, up until the end of 2021. Still, a small number of projects may have been missed, as when the criteria for Category II monitoring were met by December 2021, but reports or papers were not emitted until late in 2022. Our estimate of GMC is also lower than it would have been had we not held rigorously to the requirements for Category II monitoring. For example, we do not include or analyze Category I monitoring projects, which address the detection or identification of individuals, populations and species, and do not monitor PGD^32^. Further, our standards of documentation, such as not including studies based only on personal communications, likely excludes a few monitoring efforts. The absence of publicly available documentation with sufficient project description would essentially mean that programs are unannounced, or confidential, and not evaluable by third parties. For example, we exclude some unpublished efforts to develop and test genomic markers prior to the establishment of actual monitoring of populations. Finally, some published genetic assessments present sufficient data to serve as genetic baselines (see^33^), but without a clear declaration of the establishment of a monitoring program, would not be included in our tally.

The monitoring programs we report here generally focus on detecting changes in population diversity of neutral nuclear marker loci and of mitochondrial DNA (haplotypes). These loci are not likely directly involved with adaptation to climate. The studies minimally report allelic or haplotype diversity and none are specifically designed to detect genetic response to climate change or deteriorating environment per se. Nonetheless, climate change can affect both species distributions and PGD^34,35^ and, thus, needs to be accounted for in the design of monitoring projects. Genetic characteristics of populations at environmental niche margins could make them critical resources for managing the impacts of climate change, such as through translocation programs^36,37^(but see^38^). However, monitoring neutral genetic markers and indicators of effective population size alone is unlikely to provide representative data on the ability of populations to adapt to changing environments, such as caused by ongoing climate change, because of weak correlation between population genetic marker loci and specific genetic variants affecting functional traits that confer adaptation to environment^39–41^. Nonetheless, GMC and genetic monitoring using marker loci is suggestive of the future capacity of countries to conduct monitoring of genetic diversity related to predicted or observed climate change response of species. The technological capacity and financial resources relevant to PGD monitoring are likely highly relevant to efforts to monitor populations at functional loci. Diversity at functional loci, combined with neutral loci and demographic information, may provide improved empirical indicators of potential for resilient adaptive responses to changing climate^39,42,43^. Finally, substantial national activity in additional types of population genetic and evolutionary research may also reflect potential national responses to international initiatives for expanded genetic monitoring^10^.

The geography of monitoring efforts to date does not align well with the distribution of decreasing environmental suitability due to changing climate niche marginality. The geographic distribution of GMC suggests that monitoring capacity is not adequately distributed to detect effects of climate change on genetic diversity, degree of adaptation, or developing vulnerability to climate change effects. In particular, efforts to increase capacity for genetic monitoring could emphasize eastern and southeastern COST countries, where the number of species in areas at their climatic niche margins is relatively high currently and expected to remain so in the future (Fig. 5, Extended Data Fig. 4), and GMC for terrestrial species is sparse (Extended Data Fig. 1). Baseline genetic assessments are needed in geographic areas, such as the Iberian Peninsula for amphibians and southeastern Europe for forest trees (Extended Data Fig. 4, Appendix S3, S6, Supplementary Materials), where multiple species will experience environmental deterioration due to rapidly changing climate. These geographic areas differ substantially depending on the taxonomic group under consideration (Fig. 5). Future efforts to monitor genetic diversity of all kinds need greater political and financial support in order to focus on areas where species will increasingly experience niche margin for climate and other environmental conditions, where adaptive genetic variation needs to be maintained, and loss of diversity due to low effective population size needs to be avoided. These efforts will complement approaches that predict climate change effects relative to the distribution of adaptive genetic variation^29^.

To address the importance of environmental gradients to the conservation of genetic diversity, we distinguish here between populations that are geographically peripheral with regard to a range centroid and populations that are environmentally marginal, occurring towards the edge of their realized environmental niche. Relative geographic position can present little relationship to variation at functional loci, while relative environmental niche marginality of populations can predict the amount of variation at these loci^41^. Establishment, adaptation, and persistence of populations at environmental niche margins may depend on the steepness of environmental gradients, rates of gene flow from non-marginal populations, and stochastic processes^20,44^. Monitoring studies that employ both neutral and functional loci, and are designed to span environmental gradients to include populations from both core and marginal niche situations, will help elucidate generalities in how genetic diversity and adaptive potential vary across species ranges. Our results here based on a joint niche marginality approach indicate that for various groups of species, the Iberian Peninsula, the eastern Adriatic coast, central Turkey, and the Carpathian Mountains can serve as foci for international, cooperative monitoring programs that anticipate the effects of climate change by establishing genetic baselines that include populations in these areas. Monitoring multiple species with populations in areas of high joint niche marginality may help to identify similar genetic responses to environmental decline among species, much as exists for life history traits^45^, with the potential to develop genetic indicator species.

Our results indicate that the number of species with climatic conditions at niche margins will likely decrease in some southern European countries, for example trees in Italy and France (Fig. 5). This counter-intuitive pattern is the result of both methodological and biological factors. First, we account for all types of climatic niche marginality, not just for warm edge marginality. Countries in southern Europe have species with cold marginal conditions at upper elevation limits, and these areas face rapid warming, loss of cold marginal conditions, and substantial predicted changes in species distributions^46^. Tallies of marginal populations may decrease when climate change causes leading edge populations to newly experience core climatic conditions, or causes trailing edge populations to experience conditions outside of the species niche. Second, our approach also only reports changes in niche marginality within current species ranges in Europe. Species for which the distribution of populations at climatic niche margins in Europe changes little may experience substantial changes elsewhere. We leave examination of these patterns for future studies that take focal-species approaches. Further, range expansion with climate change will result in the influx of species into areas with newly suitable climate on leading range edges^16^. Future studies can refine predictions for climatic conditions and niche marginality in the context of specific goals for genetic monitoring and population management.

Populations at environmental niche margins, although often substantially locally adapted, have been found repeatedly less fully locally adapted than those in more central situations within the niche^21,41^. Additionally, populations towards species warm niche margins (i.e., trailing edges) may be relatively isolated from one another^16^. For these reasons, populations at niche margins can together present valuable genetic diversity at loci associated with local adaptation to climate that is not present or rare in populations more centrally located within the niche^41^. The relationship of this variability to the efficacy of adaptation to changing climate is complex (see Appendix 7 Extended Discussion, Supplementary Materials). Nonetheless, detecting any loss of genetic diversity in niche margin populations should be a priority, and if detected should likely trigger management response. To inform management in this way, monitoring projects need to span entire environmental gradients as occupied by species, in order to sample relevant genetic variation in niche marginal populations. Genetic samples from such prospective monitoring designs will be well suited for evaluating PGD and adaptive capacity of populations, and designing appropriate management strategies^47^. The present study suggests that populations towards the warm/dry, retreating niche margins are geographically clustered in Europe, which indicates the need to promote and develop monitoring capacity in countries with low GMC and high joint niche marginality (Figs. 4-6; Extended Data Fig. 5).

## Online Methods

We compare data on genetic monitoring capacity (GMC) and climatic niche marginality to address whether historical effort and experience in PGD monitoring at a national scale correspond to the anticipated impacts of climate change on environmental suitability for ensembles of wild species. We call this approach a ‘joint species niche marginality framework’, to express how areas of marginal conditions within the niches of multiple species coincide geographically, and we use it to propose taxonomic and geographic foci for future programs of genetic monitoring. Generally, we anticipate that larger, high-GDP countries will have conducted a greater number of monitoring programs than smaller and less wealthy ones. To address our four guiding questions, we report results from a comprehensive survey of the scientific literature, as represented in the Web of Science Core Collection of journals, with use of a simple, inclusive search string of relevant terms. We also collect references and documentation of unpublished monitoring programs by using professional networks to comprehensively access the gray literature, including governmental and non-governmental reports and web pages in national languages. We focus our analysis exclusively on monitoring programs that report repeated measures of PGD indicators (Category II programs^32^), and we exclude genetic assessments, which lack temporal replication, from consideration. We compile and summarize these data by country to address the geographic and taxonomic distribution of monitoring projects as an indicator of GMC. We then assemble groups of species of current or potential conservation interest based on taxonomic and functional characteristics and predict changes in their environmental niche marginality within their current range by using the range-wide occurrence of species, range polygons, and digital layers that express current climate and projected changes^31^.

### Distribution of genetic monitoring capacity in Europe

The gray literature: Beginning in October 2019 we began to solicit submission of published and unpublished (grey literature) materials documenting genetic monitoring programs, projects, and activities (forward, ‘projects’). We used social media and e-mail to contact the extended network of relationships centered on participants in the COST Action ‘Genomic Biodiversity Knowledge for Resilient Ecosystems (G-BiKE, https://www.cost.eu/actions/CA18134/), a Europe-wide effort to improve and promote the use of genetic and genomic methods for supporting delivery of ecosystem services. We directly contacted colleagues, government officials and non-governmental agency (NGO) representatives in their home countries to identify and solicit information on past and on-going projects. Submission of information was open to this broad community of scientists, policy makers and stakeholders, and was structured by variables describing each project, organized in an on-line spreadsheet (Appendix S8, Supplementary Materials). We labored to follow leads and make direct contacts in order to obtain internal documents and unreleased private reports. We collected all available documentation in the form of web documents and their URLs, white papers, internal and released reports, and published papers that were associated with, and substantiated, each submitted project. Solicitation and submission of information continued until 31 December 2021. We focused our data collection efforts exclusively on COST Full Member countries (here forward, COST countries) except for Ukraine, due to its inclusion in the COST program after the start of data collection. Submitted projects that did not sample populations in at least one COST country were excluded from subsequent data aggregation and analyses.

Project submissions required independent evaluation because no consistently applied definition of ‘genetic monitoring’ was evident upon inspection of the submissions. We developed standardized criteria for judging the validity of projects to monitor population genetic diversity by following a published definition of genetic monitoring^32^ and by defining a decision tree (Online Methods Fig. 1). Each submitted project was assigned using computer-generated pseudo-random numbers to two of 14 evaluators, who sought additional information in national languages as needed through web search and personal inquiries. Pairs of evaluators examined projects independently from one another. When the evaluators disagreed on project validity, the evaluators attempted to reach consensus. Persistent disagreements were mediated by two co-authors (PBP and MB). Written documentation, broadly defined, was required for positive decision on project validity, thereby excluding projects only reported by personal communication, e-mail, or lacking documentation (Online Methods Fig. 1). Valid monitoring projects included those that acquired and analyzed genotype data from the same populations or identical locations, at two or more time points at least one year or one generation apart, whichever was longer. Additionally, candidate projects needed to explicitly declare the goal of informing management and/or conservation policy and activities (Online Methods Fig. 1). Genetic assessments, i.e., projects lacking temporal replication, also known as ‘snapshot’ studies^32^, were excluded, as were projects with no clearly stated motivation to inform management, conservation policy or activity. This excluded studies on pathogens and disease vectors, as well as studies focused on questions clearly restricted to the field of population biology and without explicit conservation motivation. Several criteria permitted inclusion of monitoring projects that had not yet collected initial data (Online Methods Fig. 1).

**Online Methods Figure 1.**
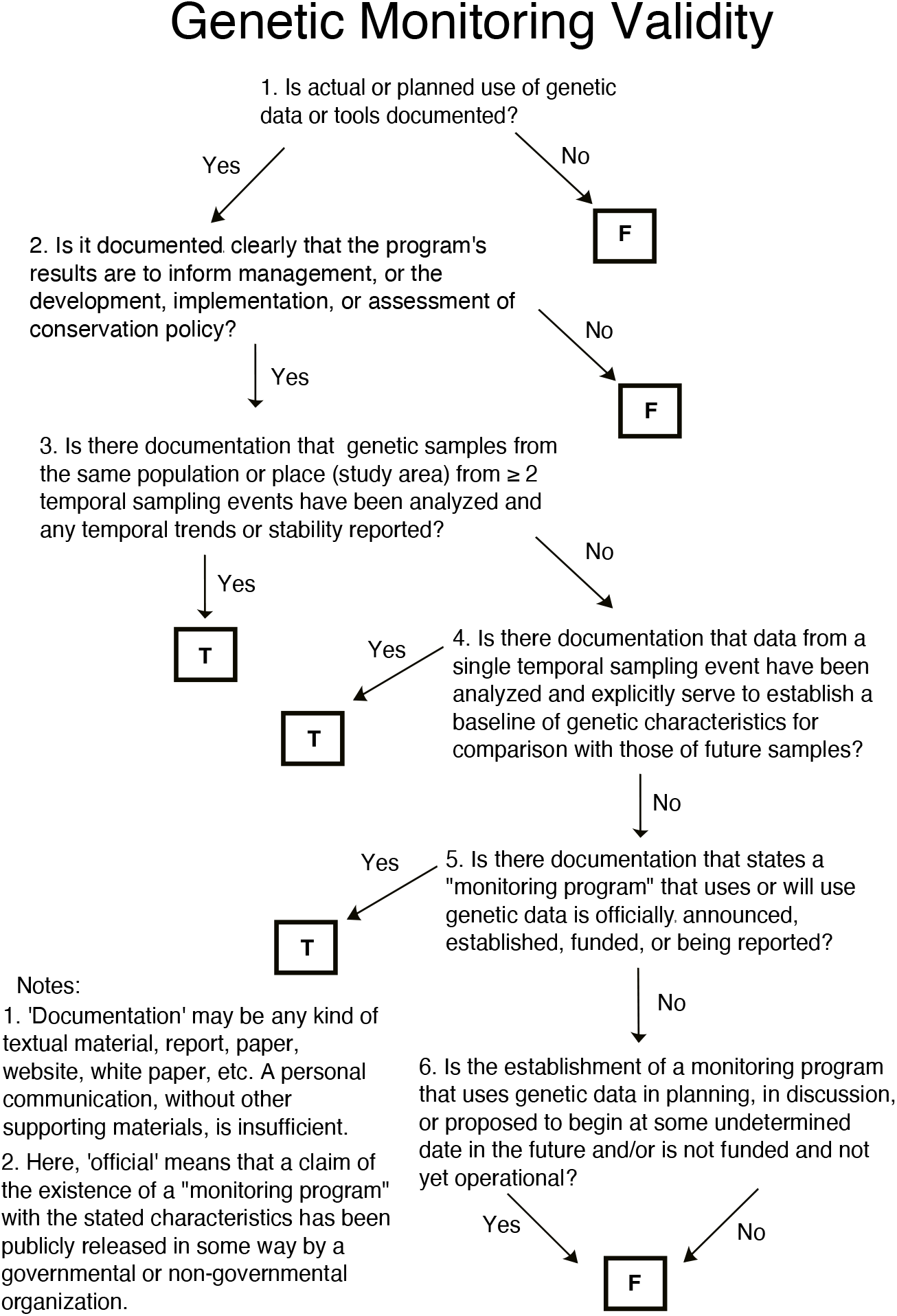
A flow chart for guiding decisions on the validity of projects as constituting genetic monitoring, given a wide range of potential documentation, originating in government reports, web documents, and the peer-reviewed literature.

A second round of evaluation classified valid monitoring projects into two groups. We distinguished between Category I projects that collected genotype or haplotype data for species and individual identification, and Category II projects that reported at least one index of population genetic diversity, such as number of alleles, observed or expected heterozygosity, etc. ^32^. The use of genetic data from archived samples or collections to establish an initial temporal reference for focal populations was acceptable, as long as the populations were strictly identical. Certain problems were presented by projects that evaluated changes in genetic diversity in re-introduced populations and those receiving introduced individuals to support levels of population genetic diversity (i.e., genetic support or assisted gene flow)^36^. For validity of these studies as Category II monitoring, a baseline sample was needed from the population of individuals initially chosen for re-introduction, or repeat temporal samples from the focal, reintroduced or supported population itself. We excluded projects comparing genetic diversity in contemporary samples to that from the original or putative source populations when these were only sampled after (re-) introductions, due to the potential for sampling bias. As in the initial evaluation of validity, both evaluators needed to express a consensus concerning the type of monitoring (Category I or II) that was conducted.

The scientific literature: We also conducted a separate survey of the peer-reviewed scientific literature to identify projects monitoring genetic diversity. On 1 December 2021, one co-author (PBP) conducted a search of all Web of Science (WoS) collections with the search string “Topic: ‘genetic population diversity monitoring’ NOT ‘cell’ NOT ‘virus’ NOT ‘medical’. Citations were then filtered to come only from the following journal categories: Agriculture, Agronomy, Dairy Animal Science, Biodiversity Conservation, Marine Freshwater Biology, Ecology, Entomology, Environmental Sciences, Evolutionary Biology, Fisheries, Forestry, Genetics and Heredity, Horticulture, Multidisciplinary, Multidisciplinary Sciences, Ornithology, Plant Sciences, and Zoology. Other strategies, such as additionally restricting the search to COST countries, resulted in the omission of studies that qualified as Category II monitoring in Europe. One co-author (PBP) scored all collected citations for being conducted in COST countries and either Category I or II monitoring. Each of these candidate studies was re-examined independently by one of four additional co-authors (DR, EB, AK, FEZ), to both evaluate the initial assessment and to identify redundancy within the original list of validated projects. Confirmed, non-redundant cases were then added to the list of monitoring projects. Ad hoc repetition of the WoS search to identify additional studies published in late 2021 and efforts to obtain documentation of specific unpublished projects, produced before the end of 2021, continued during the first four months of 2022.

We focused on Category II monitoring studies because of their relevance to mandates to conserve genetic diversity, and we carefully tallied these studies by country, and by taxonomic and additional groupings (Appendix 8, Supplementary Materials). We considered submitted projects that monitored particular single species in a country as distinct projects when different populations were studied by different research groups, institutes, or organizations. We also considered projects conducted by a single research group but having more than one focal species as distinct. Projects addressing different focal populations of a single species, analyzed as exclusive, distinct sets of populations by a single research group, were also counted as distinct projects. Nonetheless, publications that presented analyses of repeated samples from a single set of populations, and were extensions of original studies, and used the original published data in establishing temporal trajectories of genetic diversity, were not counted as separate projects regardless of author identity. Analyses of samples by contract laboratories, in a separate country from that of the study population(s), research group or monitoring organization, did not qualify the project to count toward the tally of projects for that separate country, unless of course at least one sampled population came from that country. In multi-country projects generally, samples for genetic analysis needed to be physically collected within a country for a project to count towards the tally of projects in that country. This meant that potentially a project was assigned (tallied) only to a subset of participating countries that were the sources of genetic samples. Projects reporting a temporal trajectory of genetic diversity in captive or domestic populations needed to employ genetic analysis of repeated samples and not rely exclusively on estimates of genetic diversity or change thereof that were obtained from pedigree analysis of breeding records. Because some projects sampled populations in more than one country, we defined the ‘genetic monitoring capacity’ of a country, GMC, as the tally of Category II monitoring projects obtaining genetic data from within the country. We determined the geographic distribution of GMC for focal taxonomic and functional species groups by mapping GMC for each group in each COST country and examining the frequency distribution of GMC among countries. We focus our analyses exclusively on Category II monitoring studies and will address Category I studies in a future publication.

### Climate niche marginality in Europe

Focal species-- We defined four divergent groups of species for examination of current and future geographical patterns of climatic conditions. Our objective was to construct groups with membership that exceeded the scope of current genetic monitoring efforts and which, because of taxon identity or life history traits, are either currently of conservation interest or could conceivably become of interest as climate change proceeds. Thus, while many of the species may be on national Red Lists in European countries, this was not a requirement for inclusion. We also did not attempt to comprehensively include species of conservation interest. We explicitly disregarded membership on Red Lists and European Union (EU) Directives as criteria because of the varying completeness, taxonomic resolution, and criteria for species inclusion of national Red Lists across Europe made it impossible to implement a single standard. Additionally, COST countries could not be assumed to place uniform emphasis on Red Lists as a foundation for conservation, management, or future development of monitoring programs. Further, not all COST countries are members of the EU and subject to the Directives. We developed lists of focal taxa to include: (1) most native European Amphibia (44 Anura. 26 Caudata), because of their recognized sensitivity to climate change. We excluded cave dwelling amphibians because of their limited exposure to terrestrial climate; (2) sixteen species of large birds, representing the Accipitridae, Anatidae, Gallidae, and Otididae, because size is related to extinction probability in birds globally^48^; (3) a set of eight relatively large carnivorans because of their general economic, ecological and cultural importance, and (4) a set of 91 species of forest trees (64 Magnoliopsida, 27 Pinopsida), because of the general economic and cultural importance of trees (Extended Data Table 1). Global range maps for each focal species were retrieved as polygons from the data portal of the International Union for the Conservation of Nature (IUCN) ^49^, and species occurrence data from the Global Biodiversity Information Facility^50–53^. We then defined species distributions as the pixels occupied by the species according to the IUCN range maps. We further refined species distributions within range polygons by filtering out pixels that corresponded to the CORINE Land Cover 2018 habitats classes^54^ that were not intersected at least once by occurrences of the corresponding species in question. This removed urban areas and other habitat/land cover types for which we found no evidence of occupation by species in the occurrence data.

Marginality calculations-- We used the worldwide 19 bioclimatic variables from the Chelsa database of global climate values at 30 arcsec resolution (http://chelsa-climate.org^55^) to calibrate principal component scores (PCA). We defined a working environmental space consisting of the first two PCA axes. This space summarized the main climatic gradients present on Earth (75.7% of variation explained). We rasterized IUCN species range maps at 30 arcsec resolution, extracted bioclimatic values for every occupied pixel (after filtering with CORINE 2018), and projected these values to the global climate space to generate species scores^56^. Using these species scores, we delineated the niche margins of each species by kernel density estimation (i.e. the 0.99 quantile)^31,56^. These niche margins delineated the boundaries of the climatic conditions currently occupied by the species throughout their global ranges. Finally, we calculated a standardized metric of climate marginality for each pixel of each species distribution, based on the multivariate distance to the niche margins, using the approach of Broennimann et al. ^31^. The marginality metric for each species varies from 0 to 1, with values of 0 indicating that the climatic conditions in the pixel are at the center of the niche, and values of 1 indicating that conditions are at the niche margin. In order to provide synthetic niche marginality maps for each species, we considered that pixels with the 25 percent most marginal conditions for a species, determined globally, constituted climatically marginal areas for the species, while the rest of pixels within the species niche constituted the core of the species environmental distribution. Notably, niche marginal situations could occur in geographically central or peripheral areas of the species range.

To map the future distribution of marginality of the climate niches of species, we updated the climatic values of pixels corresponding to the species distributions in the study area using a Shared Socioeconomic Pathway scenario, SSP5-8.5, for a 30-year future time period, 2041-2070. which we extracted from the Chelsa database Vers. 2.1^57^. We recalculated the marginality metric for each species in each pixel, and produced maps of species future niche marginality. This entails the assumption that the climate niches of species do not change substantially over this time frame (i.e., exhibit niche stability^58,59^). Multispecies marginality maps were produced for each species group by stacking the species maps and calculating maps of the number of species in marginal conditions of their climate niche for each pixel, at present and in the future. We compared maps of current and future niche marginality to identify pixels in which we estimated populations of species will shift into climatically marginal niche conditions in the future.

To facilitate comparison of GMC to the predicted effects of climate change on species niche situations at the country level, we converted species maps of niche marginality to country tallies of species with marginal niche conditions and tallied change over time. For each COST country, we obtained a shapefile of country boundaries at 10 m resolution from the Natural Earth website (www.naturalearthdata.com). We excluded overseas territories and regions of European countries, i.e. islands and areas outside of a rectangular bounding box defined by −25° W, 57° W, 29.1° N, 73° N. This excluded, for example, the Canary Islands (Spain), Svalbard (Norway), and French Guiana (France). We implemented a threshold for counting a species as having climatically marginal niche conditions in a country by requiring that at least 5% of the number of niche margin pixels in COST countries be within the country. This prevented countries from accruing species at niche margins because of just a few marginal pixels. We used the R package “tmap” ^60^ to map the number of PGD monitoring projects in each country, the number of marginal species in each focal taxonomic group currently and in the future, and the predicted number of species that newly experience niche margin conditions within a country as an index of the change in niche marginality We plotted future joint niche marginality against country tallies of PGD monitoring programs to visualize the relationship between national capacity for PGD monitoring and geographic foci of future climatic niche margin conditions.

### Statistical analyses

We compared GMC among countries by modeling the number of Category II monitoring projects as a function of two broadly applicable indicators. We used country area as an example indicator of the physical aspects of countries, and we estimated land area of COST countries in continental Europe, the Mediterranean and Baltic islands, and in Asia using the R package ‘sf’^61^. While many more physical aspects could be explored, a comprehensive study of physical aspects of COST countries is beyond the scope of the present paper. We also chose per capita Gross Domestic Product (GDP) as an example indicator of economic activity and available resources, one which is available for all COST countries. Data on GDP in 2020 U. S. Dollars were obtained from an authoritative on-line source^62^, the most recent year for which data from all COST countries was available. The relationship of monitoring capacity with many other social and economic indicators could be explored, but we leave this as well for future analyses. Based on inspection of scatter plots, country area entered models as a first order effect while GDP entered as a second order orthogonal polynomial.

We used a Generalized Linear Model (GLM) framework to analyze country counts of PGD monitoring projects. Models were fit with functions from the R packages ‘stats’, ‘MASS’ and ‘hermite’^63,64^. Outlier and influential data points were identified with leverage statistics and by inspection. We quantified model explanatory capacity with the Veall-Zimmermann pseudo-R^2^ ^65^ calculated on deviance residuals, and used model likelihoods and *χ*^2^ statistics to compare models during model development. We modeled the data with Poisson, negative binomial and Hermite regressions and based statistical decisions on negative binomial models because of a significant reduction in over-dispersion of residuals in comparison with the Poisson model, and no additional improvement provided by the Hermite model (Appendix 2, Supplementary Materials). We examined negative-binomial GLM model residuals for small-scale spatial autocorrelation (SAC) using a randomization test of the significance of Moran’s I (H0: I=0), at successive intervals of 300 km between country centroids, using the ‘correlog’ function in the R package ‘ncf” ^66^. We did not address large scale spatial structure (>1500 km). Because SAC can bias tests of significance of model effects when analyzing spatial data, we removed SAC from GLM residuals by first constructing spatial eigenvectors (Moran’s Eigenvector Maps) with function ‘mem’ from the R package ‘adespatial’^67^ and a regional distance network among country centroids constructed with functions ‘dnearneigh’ and ‘nb2listw’ in the R package ‘spdep’^68^ (Appendix 2, Supplementary Materials). Eigenvectors with positive eigenvalues were included as additional linear terms (regardless of statistical significance) in GLM models. We added eigenvectors until p-values of the randomization test of Moran’s I, calculated on model residuals at intervals to 1500 km, equaled or exceeded 0.05 after rounding. Although significance levels were reduced by the addition of spatial eigenvectors, decisions concerning statistical significance of model terms were not affected.

**Table 1.**
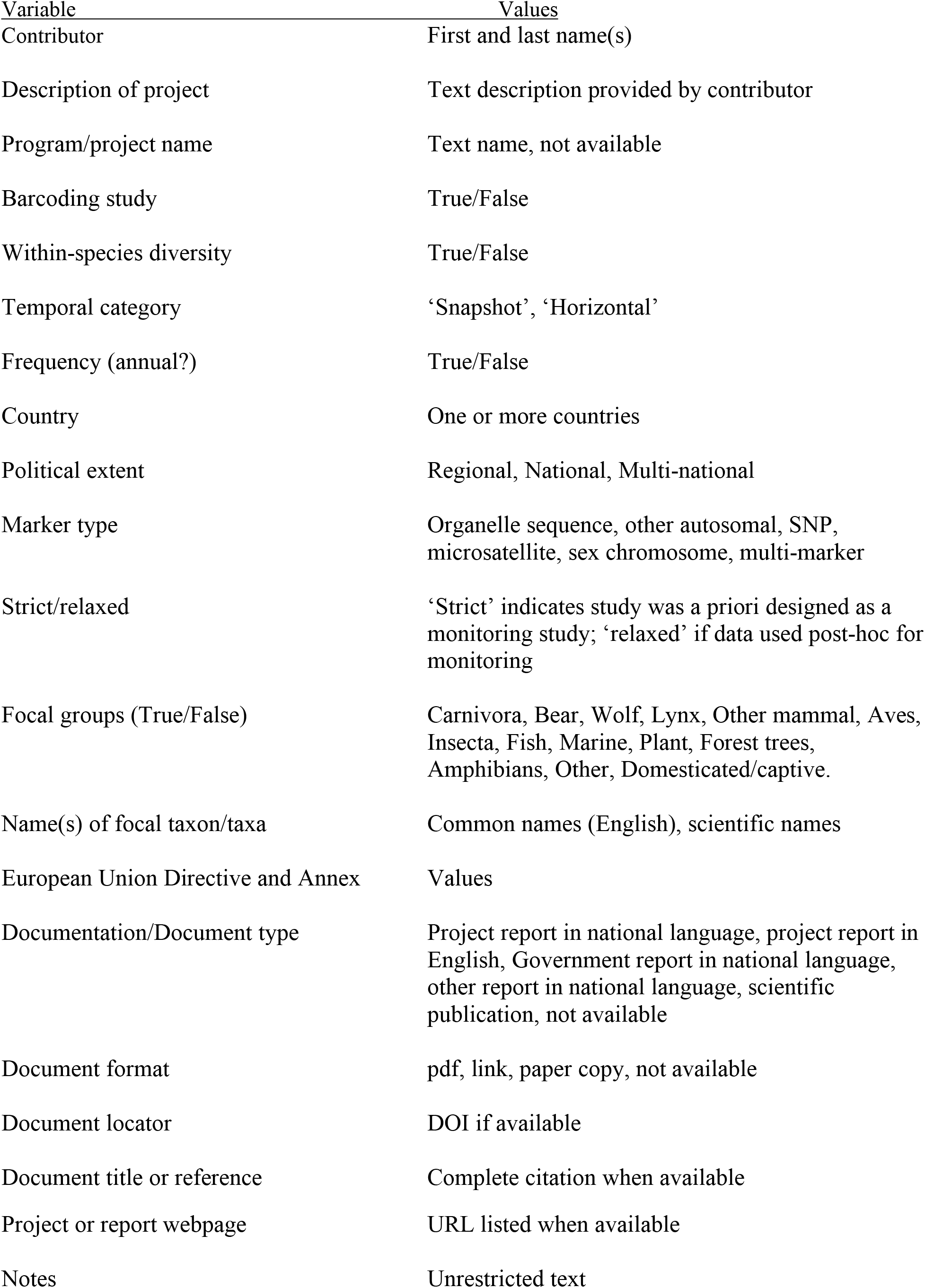
Requested information to characterize submitted monitoring projects/programs.

## Acknowledgements

This article/publication is based upon work from COST Action GBiKE, CA 18134, supported by COST (European Cooperation in Science and Technology), www.cost.eu.

IP-V is supported by the U.S. Geological Survey Powell Center for Synthesis and Analysis. BR was supported by INTER-EXCELLENCE – INTER-COST (LTC20021). DP and LL were supported by grants to LL from the Swedish Research Council Formas (grant 2020-01290) and the Swedish Research Council (grant 2019-05503).

## Author Contributions

MB and PBP conceived of the study. PBP conducted a search of published literature, organized and mapped monitoring project data, and conducted the statistical analyses. OB conducted the niche marginality analysis and mapping with support from AG. PBP, PCA, LDB, AB, EB, VCC, JAG, CH, PK, MKK, AK, CN, DP, BR, DR, ST, CV, and MB evaluated submitted monitoring projects. PBP wrote the first draft and all co-authors participated in discussions and contributed to the writing.

## Data Availability

The raw data on submitted candidate monitoring projects and a variable indicating their validity as Category II monitoring are available for download at this link: raw data withheld pending publication

## Literature Cited

1 McOwen, C. J. et al. Sufficiency and Suitability of Global Biodiversity Indicators for Monitoring Progress to 2020 Targets. Conservation Letters 9, 489–494 (2016). https://doi.org:10.1111/conl.112329

2 Laikre, L. et al. Post-2020 goals overlook genetic diversity. Science 367, 1083 (2020). https://doi.org:10.1126/science.abb2748

3 Hoban, S. et al. Genetic diversity goals and targets have improved, but remain insufficient for clear implementation of the post-2020 global biodiversity framework. Conservation genetics (Print), 1–11 (2023). https://doi.org:10.1007/s10592-022-01492-0

4 Convention on Biodiversity. Decision adopted by the Conference of the Parties to the Convention on Biological Diversity. Decision XV/4. Kunming-Montreal Global Biodiversity Framework., (2022).

5 Ette, J.-S. & Geburek, T. Why European biodiversity reporting is not reliable. Ambio 50, 929–941 (2021). https://doi.org:10.1007/s13280-020-01415-8

6 O’Brien, D. et al. Bringing together approaches to reporting on within species genetic diversity. Journal of Applied Ecology 59, 2227–2233 (2022). https://doi.org:10.1111/1365-2664.14225

7 Hoban, S. et al. Comparative evaluation of potential indicators and temporal sampling protocols for monitoring genetic erosion. Evolutionary Applications 7, 984–998 (2014). https://doi.org:10.1111/eva.12197

8 Hoban, S. et al. Genetic diversity targets and indicators in the CBD post-2020 Global Biodiversity Framework must be improved. Biological Conservation 248(2020). https://doi.org:10.1016/j.biocon.2020.108654

9 Pereira, H. M. et al. Essential Biodiversity Variables. Science 339, 277–278 (2013). https://doi.org:10.1126/science.1229931

10 Convention on Biodiversity. Monitoring framework for the Kunming-Montreal Global Biodiversity Framework. Decision XV/5. Kunming-Montreal Global Biodiversity Framework. (2022).

11 Leigh, D. M., Hendry, A. P., Vazquez-Dominguez, E. & Friesen, V. L. Estimated six per cent loss of genetic variation in wild populations since the industrial revolution. Evolutionary Applications 12, 1505–1512 (2019). https://doi.org:10.1111/eva.12810

12 Miraldo, A. et al. An Anthropocene map of genetic diversity. Science 353, 1532–1535 (2016). https://doi.org:10.1126/science.aaf4381

13 Exposito-Alonso, M. et al. Genetic diversity loss in the Anthropocene. Science 377, 1431–1435 (2022). https://doi.org:10.1126/science.abn5642

14 Merilä, J. & Hendry, A. P. Climate change, adaptation, and phenotypic plasticity: the problem and the evidence. Evolutionary Applications 7, 1–14 (2014). https://doi.org:10.1111/eva.12137

15 Purvis, A. M., Z.; Obura, D.; Ichii, K.; Willis, K.; Chettri, N.; Dulloo, E.; Hendry, A.; Gabrielyan, B.; Gutt, J.; Jacob, U.; Keskin, E.; Niamir, A.; Öztürk, B.; Salimov, R; Jaureguiberry, P. in Global Assessment Report of the Intergovernmental Science-Policy Platform on Biodiversity and Ecosystem Services (ed E. S.; Settele Brondízio, J.; Díaz, S.; Ngo, H. T.) Ch. Chapter 2.2, (IPBES Secretariat, 2019).

16 Hampe, A. & Petit, R. J. Conserving biodiversity under climate change: the rear edge matters. Ecology Letters 8, 461–467 (2005). https://doi.org:10.1111/j.1461-0248.2005.00739.x

17 Carvalho, S. B., Torres, J., Tarroso, P. & Velo-Anton, G. Genes on the edge: A framework to detect genetic diversity imperiled by climate change. Global Change Biology 25, 4034–4047 (2019). https://doi.org:10.1111/gcb.14740

18 Razgour, O. et al. Considering adaptive genetic variation in climate change vulnerability assessment reduces species range loss projections. Proceedings of the National Academy of Sciences of the United States of America 116, 10418–10423 (2019). https://doi.org:10.1073/pnas.1820663116

19 Bridle, J. R. & Vines, T. H. Limits to evolution at range margins: when and why does adaptation fail? Trends in Ecology & Evolution 22, 140–147 (2007). https://doi.org:10.1016/j.tree.2006.11.002

20 Kawecki, T. J. Adaptation to Marginal Habitats. Annual Review of Ecology Evolution and Systematics 39, 321–342 (2008). https://doi.org:10.1146/annurev.ecolsys.38.091206.095622

21 Rehfeldt, G. E., Ying, C. C., Spittlehouse, D. L. & Hamilton, D. A. Genetic responses to climate in *Pinus contorta*: Niche breadth, climate change, and reforestation. Ecological Monographs 69, 375–407 (1999). https://doi.org:10.1890/0012-9615(1999)069[0375:grtcip]2.0.co;2

22 Bontrager, M. et al. Adaptation across geographic ranges is consistent with strong selection in marginal climates and legacies of range expansion. Evolution 75, 1316–1333 (2021). https://doi.org:10.1111/evo.14231

23 Carnaval, A. C., Hickerson, M. J., Haddad, C. F. B., Rodrigues, M. T. & Moritz, C. Stability Predicts Genetic Diversity in the Brazilian Atlantic Forest Hotspot. Science 323, 785–789 (2009). https://doi.org:10.1126/science.1166955

24 Hewitt, G. M. Genetic consequences of climatic oscillations in the Quaternary. Philosophical Transactions of the Royal Society B-Biological Sciences 359, 183–195 (2004). https://doi.org:10.1098/rstb.2003.1388

25 Nadeau, C. P. & Urban, M. C. Eco-evolution on the edge during climate change. Ecography 42, 1280–1297 (2019). https://doi.org:10.1111/ecog.04404

26 Aitken, S. N., Yeaman, S., Holliday, J. A., Wang, T. L. & Curtis-McLane, S. Adaptation, migration or extirpation: climate change outcomes for tree populations. Evolutionary Applications 1, 95–111 (2008). https://doi.org:10.1111/j.1752-4571.2007.00013.x

27 Holliday, J. A., Suren, H. & Aitken, S. N. Divergent selection and heterogeneous migration rates across the range of Sitka spruce (*Picea sitchensis*). Proceedings of the Royal Society B-Biological Sciences 279, 1675–1683 (2012). https://doi.org:10.1098/rspb.2011.1805

28 Wessely, J. et al. Climate warming may increase the frequency of cold-adapted haplotypes in alpine plants. Nature Climate Change 12, 77-+ (2022). https://doi.org:10.1038/s41558-021-01255-8

29 Flanagan, S. P., Forester, B. R., Latch, E. K., Aitken, S. N. & Hoban, S. Guidelines for planning genomic assessment and monitoring of locally adaptive variation to inform species conservation. Evolutionary Applications 11, 1035–1052 (2018). https://doi.org:10.1111/eva.12569

30 COST. <https://www.cost.eu/about/members/> (2023).

31 Broennimann, O. et al. Distance to native climatic niche margins explains establishment success of alien mammals. Nature Communications 12 (2021). https://doi.org:10.1038/s41467-021-22693-0

32 Schwartz, M. K., Luikart, G. & Waples, R. S. Genetic monitoring as a promising tool for conservation and management. Trends in Ecology & Evolution 22, 25–33 (2007). https://doi.org:10.1016/j.tree.2006.08.009

33 Menguellueoglu, D., Fickel, J., Hofer, H. & Foerster, D. W. Non-invasive faecal sampling reveals spatial organization and improves measures of genetic diversity for the conservation assessment of territorial species: Caucasian lynx as a case species. Plos One 14(2019). https://doi.org:10.1371/journal.pone.0216549

34 Rubidge, E. M. et al. Climate-induced range contraction drives genetic erosion in an alpine mammal. Nature Climate Change 2, 285–288 (2012). https://doi.org:10.1038/nclimate1415

35 Yang, D. S., Conroy, C. J. & Moritz, C. Contrasting responses of Peromyscus mice of Yosemite National Park to recent climate change. Global Change Biology 17, 2559–2566 (2011). https://doi.org:10.1111/j.1365-2486.2011.02394.x

36 Aitken, S. N. & Whitlock, M. C. in Annual Review of Ecology, Evolution, and Systematics, Vol 44 Vol. 44 Annual Review of Ecology Evolution and Systematics (ed D. J. Futuyma) 367–388 (2013).

37 Hoegh-Guldberg, O. et al. Assisted colonization and rapid climate change. Science 321, 345–346 (2008). https://doi.org:10.1126/science.1157897

38 Van Daele, F., Honnay, O. & De Kort, H. Genomic analyses point to a low evolutionary potential of prospective source populations for assisted migration in a forest herb. Evolutionary Applications 15, 1859–1874 (2022). https://doi.org:10.1111/eva.13485

39 Bonin, A., Nicole, F., Pompanon, F., Miaud, C. & Taberlet, P. Population adaptive index: a new method to help measure intraspecific genetic diversity and prioritize populations for conservation. Conservation Biology 21, 697–708 (2007). https://doi.org:10.1111/j.1523-1739.2007.00685.x

40 Willi, Y., Van Buskirk, J. & Hoffmann, A. A. Limits to the adaptive potential of small populations. Annual Review of Ecology Evolution and Systematics 37, 433–458 (2006). https://doi.org:10.1146/annurev.ecolsys.37.091305.110145

41 Dauphin, B. et al. Disentangling the effects of geographic peripherality and habitat suitability on neutral and adaptive genetic variation in Swiss stone pine. Molecular Ecology 29, 1972–1989 (2020). https://doi.org:10.1111/mec.15467

42 Hoffmann, A. A. & Sgro, C. M. Climate change and evolutionary adaptation. Nature 470, 479–485 (2011). https://doi.org:10.1038/nature09670

43 Hansen, M. M., Olivieri, I., Waller, D. M., Nielsen, E. E. & Ge, M. W. G. Monitoring adaptive genetic responses to environmental change. Molecular Ecology 21, 1311–1329 (2012). https://doi.org:10.1111/j.1365-294X.2011.05463.x

44 Bridle, J. R., Polechova, J., Kawata, M. & Butlin, R. K. Why is adaptation prevented at ecological margins? New insights from individual-based simulations. Ecology Letters 13, 485–494 (2010). https://doi.org:10.1111/j.1461-0248.2010.01442.x

45 Ellegren, H. & Galtier, N. Determinants of genetic diversity. Nature Reviews Genetics 17, 422–433 (2016). https://doi.org:10.1038/nrg.2016.58

46 Engler, R. et al. 21st century climate change threatens mountain flora unequally across Europe. Global Change Biology 17, 2330–2341 (2011). https://doi.org:10.1111/j.1365-2486.2010.02393.x

47 Thurman, L. L. et al. Persist in place or shift in space? Evaluating the adaptive capacity of species to climate change. Frontiers in Ecology and the Environment 18, 520–528 (2020). https://doi.org:10.1002/fee.2253

48 Hughes, E. C., Edwards, D. P. & Thomas, G. H. The homogenization of avian morphological and phylogenetic diversity under the global extinction crisis. Current Biology 32, 3830–3837 (2022). https://doi.org:10.1016/j.cub.2022.06.018

49 IUCN. (2022).

50 GBIF.org. GBIF Occurrence Download https://doi.org/10.15468/dl.8skxjd, 2021).

51 GBIF.org. GBIF Occurrence Download https://doi.org/10.15468/dl.guf53c, 2021).

52 GBIF.org. GBIF Occurrence Download https://doi.org/10.15468/dl.z88kj8, 2021).

53 GBIF.org. GBIF Occurrence Download https://doi.org/10.15468/dl.zyhtqq, 2021).

54 Copernicus Land Monitoring Service. (European Environment Agency, 2018).

55 Karger, D. N. et al. Climatologies at high resolution for the Earth land surface areas. Scientific Data 4, 170122 (2017).

56 Broennimann, O. et al. Measuring ecological niche overlap from occurrence and spatial environmental data. Global Ecology and Biogeography 21, 481–497 (2012). https://doi.org:10.1111/j.1466-8238.2011.00698.x

57 Karger, D. N. et al. Data Descriptor: Climatologies at high resolution for the earth’s land surface areas. Scientific Data 4 (2017). https://doi.org:10.1038/sdata.2017.122

58 Guisan, A., Petitpierre, B., Broennimann, O., Daehler, C. & Kueffer, C. Unifying niche shift studies: insights from biological invasions. Trends in Ecology & Evolution 29, 260–269 (2014). https://doi.org:10.1016/j.tree.2014.02.009

59 Pearman, P. B., Guisan, A., Broennimann, O. & Randin, C. F. Niche dynamics in space and time. Trends in Ecology & Evolution 23, 149–158 (2008). https://doi.org:10.1016/j.tree.2007.11.005

60 Tennekes, M. tmap: Thematic Maps in R. Journal of Statistical Software 84, 1–39 (2018). https://doi.org:10.18637/jss.v084.i06

61 Pebesma, E. Simple Features for R: Standardized Support for Spatial Vector Data. The R Journal 10, 439–446 (2018).

62 The World Bank. DataBank. World Development Indicators, <https://data.worldbank.org/indicator/NY.GDP.PCAP.CD> (2022).

63 Moriña, D., Higueras, M., Puig, P. & Oliveira, M. hermite: Generalized Hermite Distribution. R package version 1.1.2. R Journal 7, 263–274 (2018).

64 Venables, W. N. & Ripley, B. D. Modern Applied Statistics with S. Fourth Edition. (Springer, 2002).

65 Veall, M. R. & Zimmermann, K. F. Evaluating pseudo-R^2^s for binary probit models,. Quality & Quantity 28, 151–164 (1994). https://doi.org:10.1007/bf01102759

66 Bjornstad, O. N. ncf: Spatial Covariance Functions. v. 1.3-2. (2022).

67 Dray, S. et al. adespatial: Multivariate Multiscale Spatial Analysis. v. 0.3-16. (2022).

68 Bivand, R. S., Pebesma, E. & Gomez-Rubio, V. Applied spatial data analysis with R, Second edition. (Springer, 2013).

