## Extended Data for "Conserving genetic diversity during climate change: Niche marginality and discrepant monitoring capacity in Europe"

### Extended Data Figures and Table

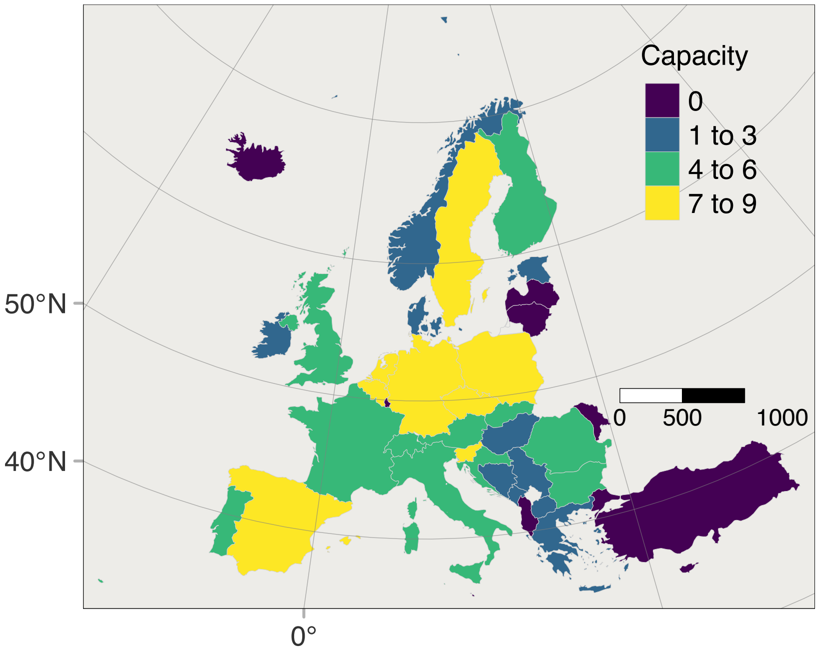


Extended Data Figure 1. Terrestrial population genetic diversity monitoring programs, reflecting genetic monitoring capacity, in each COST Full-Member country, up to 31.12.2021. The data include projects monitoring amphibians, birds, carnivorans and trees, but exclude programs/projects that monitored fish, marine species and domesticated/captive species.

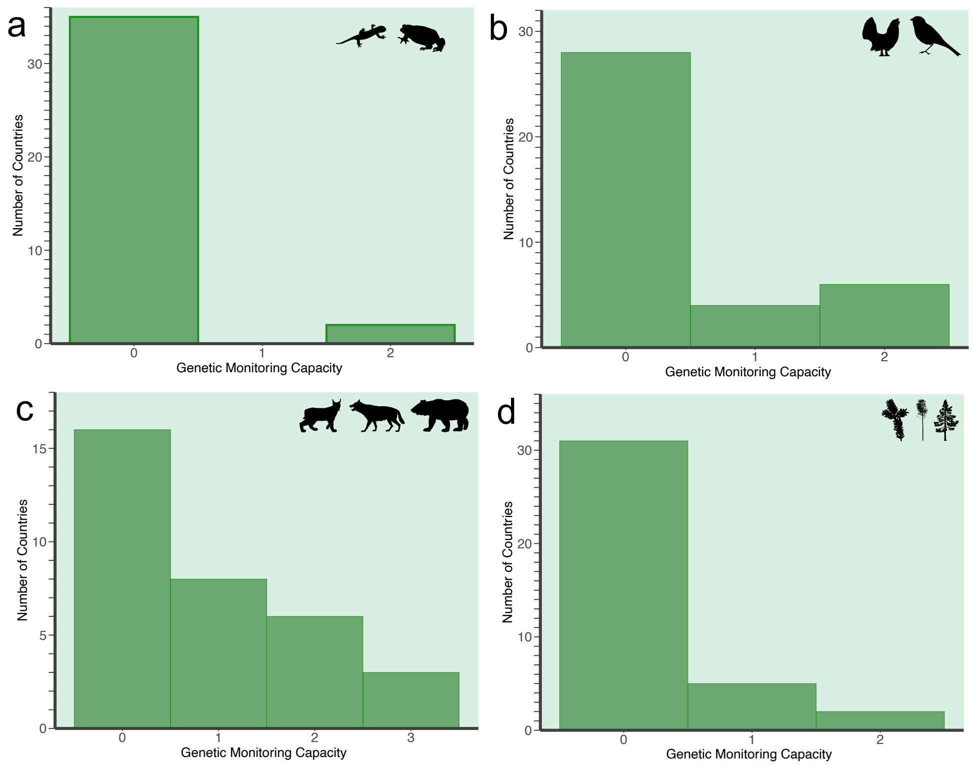


Extended Data Figure 2. Frequency Distribution of Genetic Monitoring Capacity among COST Full-Member Countries. The data are for amphibians (a), birds (b), large European carnivorans (c), and forest trees (d). They represent all projects and programs using genetic data and reporting data on genetic diversity, from at least two time points, thus qualifying as Category II monitoring of population genetic diversity. Attribution for all silhouettes in

this paper:
https://www.phylopic.org/permalinks/ec1bdb9ee275ab1dc5a68e090a169a96c06c559e516989215bb5d881696db666

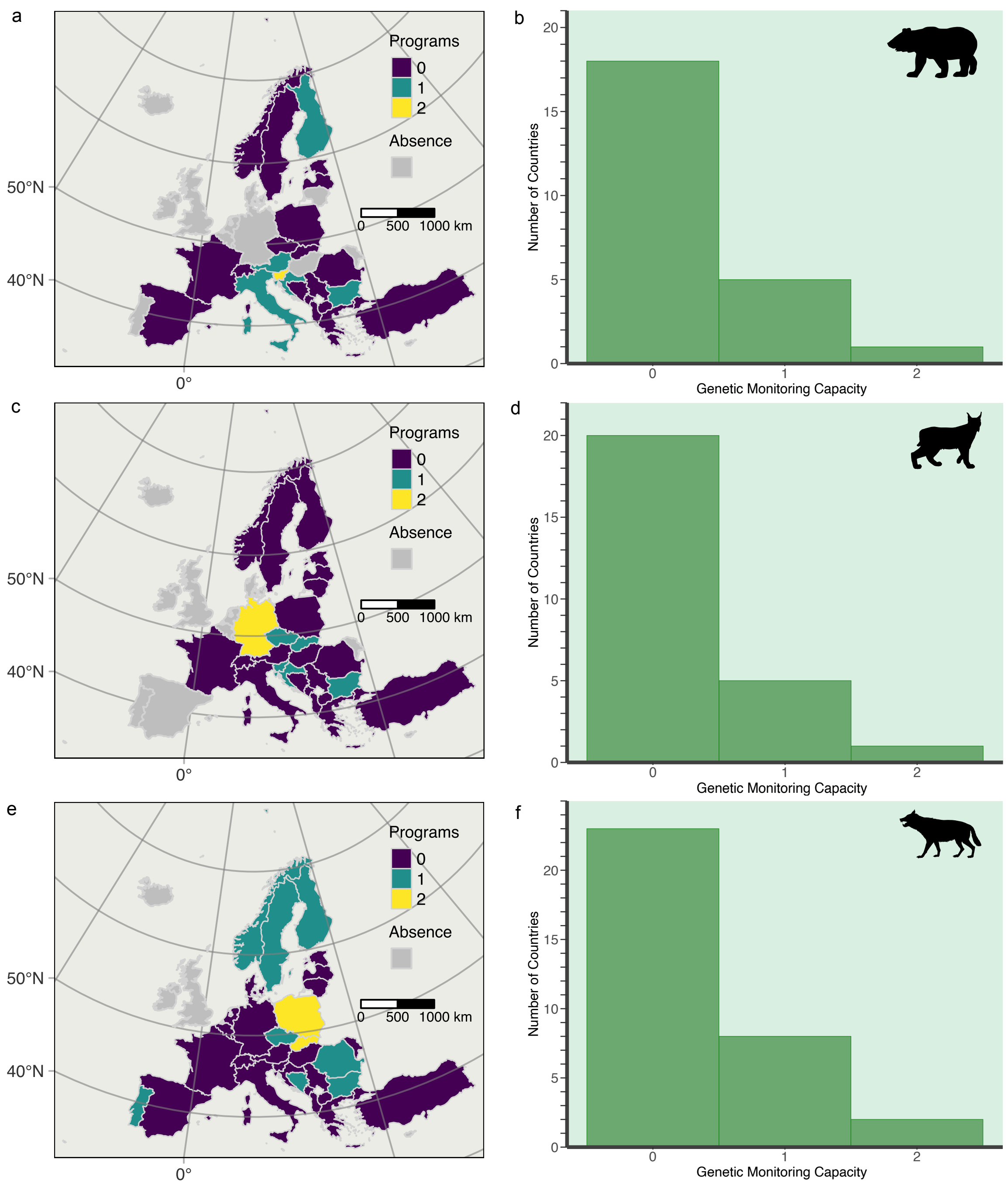


Extended Data Figure 3. Distribution and frequency of monitoring programs for three carnivorans in COST Full-Member Countries. The data are for Eurasian brown bear (*Ursus arctos*; a, b), Eurasian lynx (*Lynx lynx*; c, d), and Eurasian wolf (*Canis lupus*; e, f). Tallies are of identifiably distinct monitoring projects and programs that report Category II monitoring of population genetic diversity.

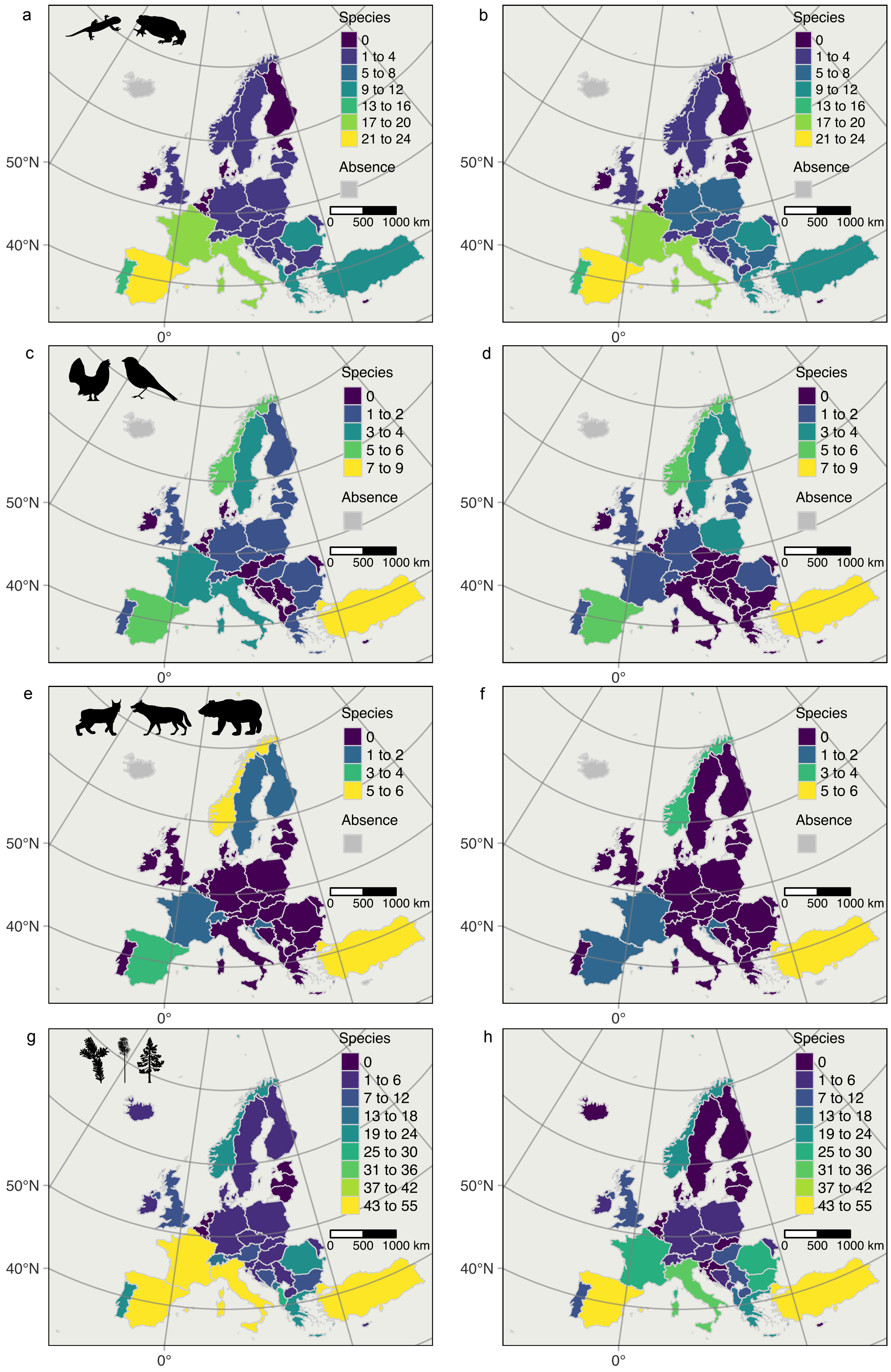


Extended Data Figure 4. Number of species with climatic niche margin conditions in each COST country at two times, current (left column) and future (right column). The number of species with estimated range falling into areas with climatic niche marginality are shown tallied by country for amphibians (a, b), a selection of large birds (c, d), several large carnivorans (e, f), and forest trees (g, h). The pattern of changing joint niche marginality differs among species groups.  An increasing number of populations at niche margins occurs in central and eastern Europe. Niche margin populations of birds increase broadly across Europe. Populations of large carnivorans increasingly experience marginal niche conditions in Turkey. Populations of forest trees increasingly experience niche marginality conditions in the Balkan states and decreasingly in France, Italy and Switzerland.  Niche marginality here does not distinguish between warm edge and cold edge populations.  Thus, mountainous countries may show decreasing numbers of species here because some populations may no longer experience cold niche margin conditions in the future.

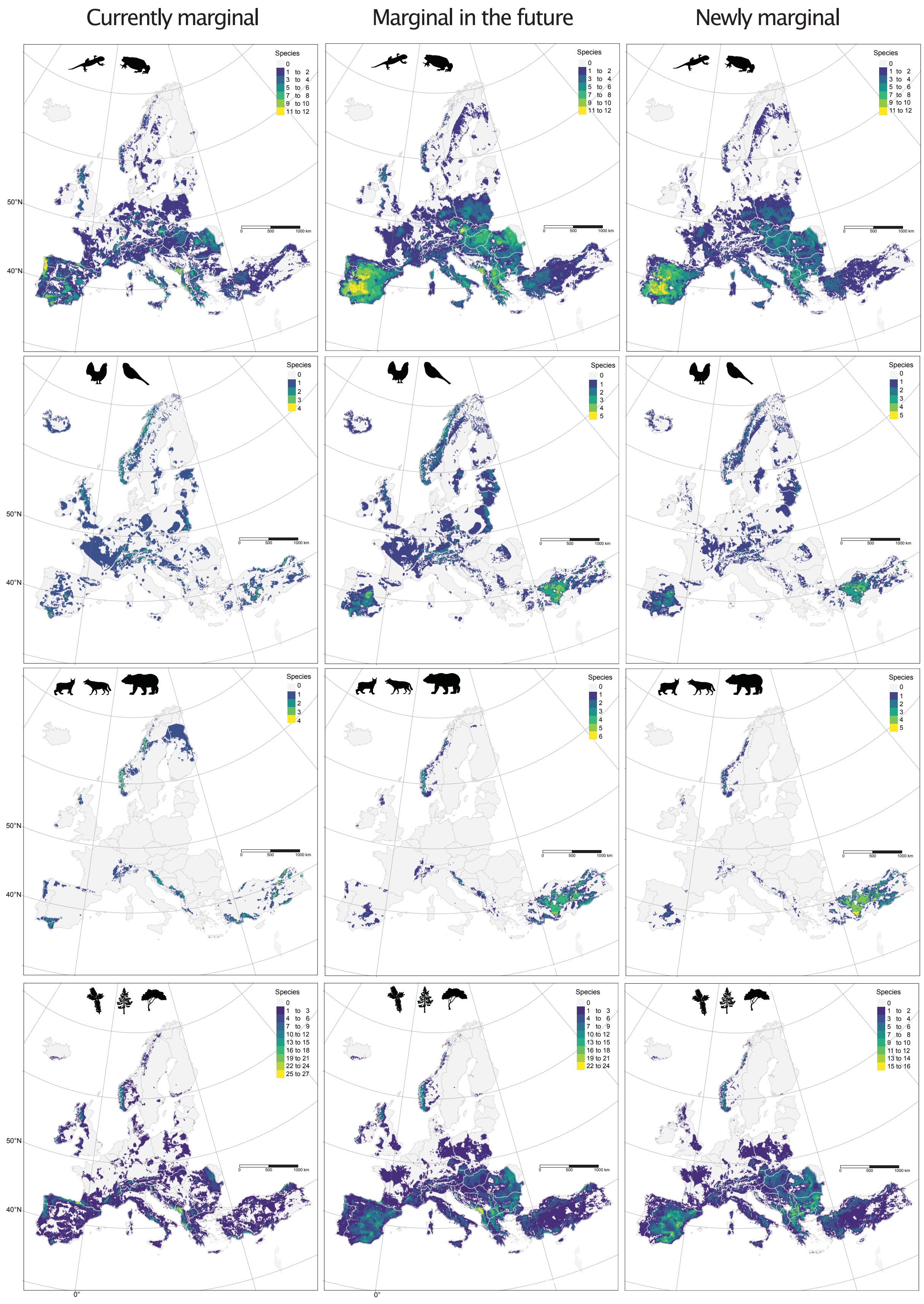


Extended Data Figure 5. Numbers of species with conditions at climatic niche margins for COST full member countries (as of 1 January 2020). Original data are at the level of 1 $km^2$ pixels, but here the data are aggregated to 10 km x 10 km pixels to improve visualization. The highest value within this 100 $km^2$ area is displayed. Current values (left column), future values (middle), and change in numbers of species (right) are shown for (top to bottom) amphibians, a selection of large birds, several large carnivorans, and a collection of forest tree species. Future marginality of species was calculated with climate data averaged over a

30-year period, 2041-2070. The right column (change) presents in each pixel the number of focal species predicted newly to experience marginal conditions during the interval between current and future and, thus, represents habitat degradation due to climate change. Calculation of change in number of species with marginal niche conditions was based on species maps of core and marginal areas (Appendices S2-S5, Supplemental Materials). Areas that because of climate change are no longer within the niche of a species are not included in the calculation of change in marginal/core status. Pixel aggregation and displayed values are as above. The areas of habitat degradation (right column) and future niche marginality patterns (middle column) largely coincide (this also holds for unaggregated pixels, not shown). See Extended Data Table 1 for species identities. See on-line Methods for details on the calculation of climatic niche marginality for species and the filtering of pixels with a CORINE land cover layer. Values shown here and tallies by country (Extended Data Fig. 4) can differ because of range differences among the species and the requirement that within a country and for each species, the pixel tally exceed five percent of the total number of marginal pixels in COST countries for the species to be counted towards the country total.

Extended Data Table 1. Species of current or potential future conservation or management interest and used in this study.

| Group | Subgroup | G-BIKE_name | GBIF_name | IUCN_name | English_name |
|---|---|---|---|---|---|
| amphibians | Anura | *Alytes cisternasii* | *Alytes cisternasii* | *Alytes cisternasii* | Iberian midwife toad |
| amphibians | Anura | *Alytes dickhilleni* | *Alytes dickhilleni* | *Alytes dickhilleni* | Betic Midwife Toad |
| amphibians | Anura | *Alytes obstetricans* | *Alytes obstetricans* | *Alytes obstetricans* | Midwife toad |
| amphibians | Anura | *Bombina bombina* | *Bombina bombina* | *Bombina bombina* | European fire-bellied toad |
| amphibians | Anura | *Bombina pachypus* | *Bombina pachypus* | *Bombina pachypus* | Appenine Yellow-bellied Toad |
| amphibians | Anura | *Bombina variegata* | *Bombina variegata* | *Bombina variegata* | Yellow-bellied toad |
| amphibians | Anura | *Bufo bufo* | *Bufo bufo* | *Bufo bufo* | Common European Toad |
| amphibians | Anura | *Discoglossus galganoi* | *Discoglossus galganoi* | *Discoglossus galganoi* | Iberian painted frog |
| amphibians | Anura | *Discoglossus montalentii* | *Discoglossus montalentii* | *Discoglossus montalentii* | Corsica Painted Frog |
| amphibians | Anura | *Discoglossus pictus* | *Discoglossus pictus* | *Discoglossus pictus* | Painted Frog |
| amphibians | Anura | *Discoglossus sardus* | *Discoglossus sardus* | *Discoglossus sardus* | Sardinia painted frog |
| amphibians | Anura | *Epidalea calamita* | *Epidalea calamita* | *Epidalea calamita* | Natterjack toad |
| amphibians | Anura | *Hyla arborea* | *Hyla arborea* | *Hyla arborea* | European tree frog |
| amphibians | Anura | *Hyla intermedia* | *Hyla intermedia* | *Hyla intermedia* | Italian Tree Frog |
| amphibians | Anura | *Hyla meridionalis* | *Hyla meridionalis* | *Hyla meridionalis* | Mediterranean Treefrog |
| amphibians | Anura | *Hyla sarda* | *Hyla sarda* | *Hyla sarda* | Sardinian Tree Frog |
| amphibians | Anura | *Hyla savignyi* | *Hyla savignyi* | *Hyla savignyi* | Savigny's treefrog |
| amphibians | Anura | *Pelobates cultripes* | *Pelobates cultripes* | *Pelobates cultripes* | western spadefoot |
| amphibians | Anura | *Pelobates fuscus* | *Pelobates fuscus* | *Pelobates fuscus* | Common Eurasian spadefoot toad |
| amphibians | Anura | *Pelobates syriacus* | *Pelobates syriacus* | *Pelobates syriacus* | Syrian spadefoot toad |
| amphibians | Anura | *Pelodytes ibericus* | *Pelodytes ibericus* | *Pelodytes ibericus* | Iberian spadefoot toad |
| amphibians | Anura | *Pelodytes punctatus* | *Pelodytes punctatus* | *Pelodytes punctatus* | Parsley frog |
| amphibians | Anura | *Pelophylax bedriagae* | *Pelophylax bedriagae* | *Pelophylax bedriagae* | Bedriaga's Frog |
| amphibians | Anura | *Pelophylax bergeri* | *Pelophylax bergeri* | *Pelophylax bergeri* | Italian Pool Frog |
| amphibians | Anura | *Pelophylax cerigensis* | *Pelophylax cerigensis* | *Pelophylax cerigensis* | Karpathos Frog |
| amphibians | Anura | *Pelophylax cretensis* | *Pelophylax cretensis* | *Pelophylax cretensis* | Cretan Frog |
| amphibians | Anura | *Pelophylax epeiroticus* | *Pelophylax epeiroticus* | *Pelophylax epeiroticus* | Epirus Water Frog |
| amphibians | Anura | *Pelophylax kurtmuelleri* | *Pelophylax kurtmuelleri* | *Pelophylax kurtmuelleri* | Balkan Water Frog |

| | | | | | |
|---|---|---|---|---|---|
| amphibians | Anura | *Pelophylax lessonae* | *Pelophylax lessonae* | *Pelophylax lessonae* | Pool Frog |
| amphibians | Anura | *Pelophylax perezi* | *Pelophylax perezi* | *Pelophylax perezi* | Perez's Frog |
| amphibians | Anura | *Pelophylax ridibundus* | *Pelophylax ridibundus* | *Pelophylax ridibundus* | Marsh frog |
| amphibians | Anura | *Pelophylax shqipericus* | *Pelophylax shqipericus* | *Pelophylax shqipericus* | Albanian Water Frog |
| amphibians | Anura | *Pseudepidalea balearica* | *Bufotes balearicus* | *Bufotes balearicus* | Balearic green toad |
| amphibians | Anura | *Pseudepidalea sicula* | *Bufotes siculus* | *Bufotes siculus* | African Green Toad |
| amphibians | Anura | *Pseudepidalea variabilis* | *Bufotes variabilis* | *Bufotes variabilis* | Varying Toad |
| amphibians | Anura | *Pseudepidalea viridis* | *Bufotes viridis* | *Bufotes viridis* | Green Toad |
| amphibians | Anura | *Rana arvalis* | *Rana arvalis* | *Rana arvalis* | Moor frog |
| amphibians | Anura | *Rana dalmatina* | *Rana dalmatina* | *Rana dalmatina* | Agile frog |
| amphibians | Anura | *Rana graeca* | *Rana graeca* | *Rana graeca* | Stream frog |
| amphibians | Anura | *Rana iberica* | *Rana iberica* | *Rana iberica* | Iberian frog |
| amphibians | Anura | *Rana italica* | *Rana italica* | *Rana italica* | Italian stream frog |
| amphibians | Anura | *Rana latastei* | *Rana latastei* | *Rana latastei* | Italian agile frog |
| amphibians | Anura | *Rana pyrenaica* | *Rana pyrenaica* | *Rana pyrenaica* | Pyrenean Frog |
| amphibians | Anura | *Rana temporaria* | *Rana temporaria* | *Rana temporaria* | Common frog |
| amphibians | Caudata | *Calotriton arnoldi* | *Calotriton arnoldi* | *Calotriton arnoldi* | Montseny brook newt |
| amphibians | Caudata | *Calotriton asper* | *Calotriton asper* | *Calotriton asper* | Pyrenean Brook Salamander |
| amphibians | Caudata | *Chioglossa lusitanica* | *Chioglossa lusitanica* | *Chioglossa lusitanica* | Gold-striped salamander |
| amphibians | Caudata | *Euproctus montanus* | *Euproctus montanus* | *Euproctus montanus* | Corsican mountain newt |
| amphibians | Caudata | *Euproctus platycephalus* | *Euproctus platycephalus* | *Euproctus platycephalus* | Sardinian mountain newt |
| amphibians | Caudata | *Lissotriton boscai* | *Lissotriton boscai* | *Lissotriton boscai* | Bosca's Newt |
| amphibians | Caudata | *Lissotriton helveticus* | *Lissotriton helveticus* | *Lissotriton helveticus* | Palmate Newt |
| amphibians | Caudata | *Lissotriton italicus* | *Lissotriton italicus* | *Lissotriton italicus* | Italian Newt |
| amphibians | Caudata | *Lissotriton montandoni* | *Lissotriton montandoni* | *Lissotriton montandoni* | Carpathian Newt |
| amphibians | Caudata | *Lissotriton vulgaris* | *Lissotriton vulgaris* | *Lissotriton vulgaris* | Smooth Newt |
| amphibians | Caudata | *Lyciasalamandra helverseni* | *Lyciasalamandra helverseni* | *Lyciasalamandra helverseni* | Karpathos salamander |
| amphibians | Caudata | *Lyciasalamandra luschani* | *Lyciasalamandra luschani* | *Lyciasalamandra luschani* | Luschan's salamander |
| amphibians | Caudata | *Mesotriton alpestris* | *Ichthyosaura alpestris* | *Ichthyosaura alpestris* | Alpine Newt |
| amphibians | Caudata | *Pleurodeles waltl* | *Pleurodeles waltl* | *Pleurodeles waltl* | Iberian ribbed newt |
| amphibians | Caudata | *Salamandra atra* | *Salamandra atra* | *Salamandra atra* | Alpine salamander |
| amphibians | Caudata | *Salamandra corsica* | *Salamandra corsica* | *Salamandra corsica* | Corsican Fire Salamander |
| amphibians | Caudata | *Salamandra lanzai* | *Salamandra lanzai* | *Salamandra lanzai* | Lanza's Alpine salamander |

| | | | | | |
|---|---|---|---|---|---|
| amphibians | Caudata | *Salamandra salamandra* | *Salamandra salamandra* | *Salamandra salamandra* | Fire salamander |
| amphibians | Caudata | *Salamandrina perspicillata* | *Salamandrina perspicillata* | *Salamandrina perspicillata* | Northern spectacled salamander |
| amphibians | Caudata | *Salamandrina terdigitata* | *Salamandrina terdigitata* | *Salamandrina terdigitata* | Southern spectacled salamander |
| amphibians | Caudata | *Triturus carnifex* | *Triturus carnifex* | *Triturus carnifex* | Italian crested newt |
| amphibians | Caudata | *Triturus cristatus* | *Triturus cristatus* | *Triturus cristatus* | Crested newt |
| amphibians | Caudata | *Triturus dobrogicus* | *Triturus dobrogicus* | *Triturus dobrogicus* | Danube Crested Newt |
| amphibians | Caudata | *Triturus karelinii* | *Triturus karelinii* | *Triturus karelinii* | Southern Crested Newt |
| amphibians | Caudata | *Triturus marmoratus* | *Triturus marmoratus* | *Triturus marmoratus* | Marbled newt |
| amphibians | Caudata | *Triturus pygmaeus* | *Triturus pygmaeus* | *Triturus pygmaeus* | Southern Marbled Newt |
| large birds | Acciptridae | *Aquila adalberti* | *Aquila adalberti* | *Aquila adalberti* | Spanish Imperial Eagle |
| large birds | Acciptridae | *Aquila heliaca* | *Aquila heliaca* | *Aquila heliaca* | Eastern Imperial Eagle |
| large birds | Acciptridae | *Clanga clanga* | *Clanga clanga* | *Clanga clanga* | Greater Spotted Eagle |
| large birds | Acciptridae | *Gypaetus barbatus* | *Gypaetus barbatus* | *Gypaetus barbatus* | Bearded Vulture |
| large birds | Acciptridae | *Neophron percnopterus* | *Neophron percnopterus* | *Neophron percnopterus* | Egyptian Vulture |
| large birds | Anatidae | *Anser erythropus* | *Anser erythropus* | *Anser erythropus* | Lesser White-fronted Goose |
| large birds | Anatidae | *Aythya ferina* | *Aythya ferina* | *Aythya ferina* | Common Pochard |
| large birds | Anatidae | *Clangula hyemalis* | *Clangula hyemalis* | *Clangula hyemalis* | Long-tailed Duck |
| large birds | Anatidae | *Marmaronetta angustirostris* | *Marmaronetta angustirostris* | *Marmaronetta angustirostris* | Marbled Teal |
| large birds | Anatidae | *Melanitta fusca* | *Melanitta fusca* | *Melanitta fusca* | Velvet Scoter |
| large birds | Anatidae | *Oxyura leucocephala* | *Oxyura leucocephala* | *Oxyura leucocephala* | White-headed Duck |
| large birds | Anatidae | *Polysticta stelleri* | *Polysticta stelleri* | *Polysticta stelleri* | Steller's Eider |
| large birds | Gallidae | *Lyrurus mlokosiewiczi* | *Lyrurus mlokosiewiczi* | *Lyrurus mlokosiewiczi* | Caucasian Grouse |
| large birds | Gallidae | *Lyrurus tetrix* | *Lyrurus tetrix* | *Lyrurus tetrix* | Black Grouse |
| large birds | Gallidae | *Tetrao urogallus* | *Tetrao urogallus* | *Tetrao urogallus* | Western Capercaillie |
| large birds | Otididae | *Otis tarda* | *Otis tarda* | *Otis tarda* | Great Bustard |
| carnivorans | Canidae | *Canis aureus* | *Canis aureus* | *Canis aureus* | Golden Jackal |
| carnivorans | Canidae | *Canis lupus* | *Canis lupus* | *Canis lupus* | Wolf |
| carnivorans | Mustelidae | *Gulo gulo* | *Gulo gulo* | *Gulo gulo* | Wolverine |
| carnivorans | Mustelidae | *Lutra lutra* | *Lutra lutra* | *Lutra lutra* | Eurasian Otter |
| carnivorans | Felidae | *Lynx lynx* | *Lynx lynx* | *Lynx lynx* | Eurasian Lynx |
| carnivorans | Felidae | *Lynx pardinus* | *Lynx pardinus* | *Lynx pardinus* | Iberian Lynx |
| carnivorans | Mustelidae | *Meles meles* | *Meles meles* | *Meles meles* | European Badger |
| carnivorans | Ursidae | *Ursus arctos* | *Ursus arctos* | *Ursus arctos* | Eurasian brown Bear |

| | | | |
|---|---|---|---|
| trees | Magnoliopsida | *Acer campestre* | Field maple |
| trees | Magnoliopsida | *Acer monspessulanum* | Montepelier maple |
| trees | Magnoliopsida | *Acer platanoides* | Norway maple |
| trees | Magnoliopsida | *Acer pseudoplatanus* | Sycamore |
| trees | Magnoliopsida | *Aesculus hippocastanum* | horse chestnut |
| trees | Magnoliopsida | *Alnus cordata* | Italian alder |
| trees | Magnoliopsida | *Alnus glutinosa* | Black alder |
| trees | Magnoliopsida | *Alnus incana* | Grey alder |
| trees | Magnoliopsida | *Alnus viridis* | Green alder |
| trees | Magnoliopsida | *Arbutus unedo* | strawberry tree |
| trees | Magnoliopsida | *Betula pendula* | Silver birch |
| trees | Magnoliopsida | *Betula pubescens* | Downy birch |
| trees | Magnoliopsida | *Buxus sempervirens* | common box |
| trees | Magnoliopsida | *Carpinus betulus* | European hornbeam |
| trees | Magnoliopsida | *Carpinus orientalis* | Oriental hornbeam |
| trees | Magnoliopsida | *Castanea sativa* | Chestnut |
| trees | Magnoliopsida | *Celtis australis* | European nettle tree |
| trees | Magnoliopsida | *Cornus mas* | Cornelian cherry |
| trees | Magnoliopsida | *Cornus sanguinea* | Common dogwood |
| trees | Magnoliopsida | *Corylus avellana* | Common hazel |
| trees | Magnoliopsida | *Euonymus europaeus* | common spindle |
| trees | Magnoliopsida | *Fagus sylvatica* | European beech |
| trees | Magnoliopsida | *Frangula alnus* | Glossy buckthorn |
| trees | Magnoliopsida | *Fraxinus angustifolia* | Narrow-leaved ash |
| trees | Magnoliopsida | *Fraxinus excelsior* | Common ash |
| trees | Magnoliopsida | *Fraxinus ornus* | manna ash |
| trees | Magnoliopsida | *Ilex aquifolium* | Common holly |
| trees | Magnoliopsida | *Juglans regia* | Common walnut |
| trees | Magnoliopsida | *Juniperus phoenicea* | Phoenicean juniper |
| trees | Magnoliopsida | *Juniperus thurifera* | Spanish juniper |
| trees | Magnoliopsida | *Liquidambar orientalis* | Oriental sweet gum |
| trees | Magnoliopsida | *Olea europaea* | European olive |
| trees | Magnoliopsida | *Ostrya carpinifolia* | Hop hornbeam |

| | | | |
|---|---|---|---|
| trees | Magnoliopsida | *Platanus orientalis* | Oriental plane |
| trees | Magnoliopsida | *Populus alba* | White poplar |
| trees | Magnoliopsida | *Populus nigra* | European black poplar |
| trees | Magnoliopsida | *Populus tremula* | Eurasian aspen |
| trees | Magnoliopsida | *Prunus avium* | Wild cherry |
| trees | Magnoliopsida | *Prunus padus* | Bird cherry |
| trees | Magnoliopsida | *Prunus spinosa* | Blackthorn |
| trees | Magnoliopsida | *Quercus cerris* | Turkey oak |
| trees | Magnoliopsida | *Quercus coccifera* | Kermes oak |
| trees | Magnoliopsida | *Quercus faginea* | Portuguese oak |
| trees | Magnoliopsida | *Quercus frainetto* | Hungarian oak |
| trees | Magnoliopsida | *Quercus ilex* | Holm oak |
| trees | Magnoliopsida | *Quercus petraea* | Sessile oak |
| trees | Magnoliopsida | *Quercus pubescens* | Pubescent oak |
| trees | Magnoliopsida | *Quercus pyrenaica* | Pyrenean oak |
| trees | Magnoliopsida | *Quercus robur* | Pedunculate oak |
| trees | Magnoliopsida | *Quercus suber* | Cork oak |
| trees | Magnoliopsida | *Quercus trojana* | Macedonian oak |
| trees | Magnoliopsida | *Salix alba* | White willow |
| trees | Magnoliopsida | *Salix caprea* | Goat willow |
| trees | Magnoliopsida | *Sambucus nigra* | black elder |
| trees | Magnoliopsida | *Sorbus aria* | common whitebeam |
| trees | Magnoliopsida | *Sorbus aucuparia* | European mountain ash |
| trees | Magnoliopsida | *Sorbus domestica* | Service tree |
| trees | Magnoliopsida | *Sorbus torminalis* | Wild service tree |
| trees | Magnoliopsida | *Tilia cordata* | Small-leaved lime |
| trees | Magnoliopsida | *Tilia platyphyllos* | Large-leaved lime |
| trees | Magnoliopsida | *Tilia tomentosa* | Silver lime |
| trees | Magnoliopsida | *Ulmus glabra* | Wych elm |
| trees | Magnoliopsida | *Ulmus laevis* | European white elm |
| trees | Magnoliopsida | *Ulmus minor* | Field elm |
| trees | Pinopsida | *Abies alba* | Silver fir |
| trees | Pinopsida | *Abies borisii-regis* | King Boris fir |

| | | | |
|---|---|---|---|
| trees | Pinopsida | *Abies cephalonica* | Grecian fir |
| trees | Pinopsida | *Abies cilicica* | Cilician fir |
| trees | Pinopsida | *Abies nebrodensis* | Sicilian fir |
| trees | Pinopsida | *Abies nordmanniana* | Caucasian fir |
| trees | Pinopsida | *Abies numidica* | Algerian fir |
| trees | Pinopsida | *Abies pinsapo* | Spanish fir |
| trees | Pinopsida | *Cedrus libani* | Cedar of Lebanon |
| trees | Pinopsida | *Cupressus sempervirens* | Italian cypress |
| trees | Pinopsida | *Juniperus communis* | Common juniper |
| trees | Pinopsida | *Juniperus excelsa* | Greek juniper |
| trees | Pinopsida | *Juniperus oxycedrus* | Prickly juniper |
| trees | Pinopsida | *Larix decidua* | European larch |
| trees | Pinopsida | *Picea abies* | Norway spruce |
| trees | Pinopsida | *Picea omorika* | Serbian spruce |
| trees | Pinopsida | *Pinus brutia* | Brutia pine |
| trees | Pinopsida | *Pinus cembra* | Swiss stone pine |
| trees | Pinopsida | *Pinus halepensis* | Aleppo pine |
| trees | Pinopsida | *Pinus heldreichii* | Bosnian pine |
| trees | Pinopsida | *Pinus mugo* | Mountain pine |
| trees | Pinopsida | *Pinus nigra* | European black pine |
| trees | Pinopsida | *Pinus peuce* | Macedonian pine |
| trees | Pinopsida | *Pinus pinaster* | Maritime pine |
| trees | Pinopsida | *Pinus pinea* | Stone pine |
| trees | Pinopsida | *Pinus sylvestris* | Scots pine |
| trees | Pinopsida | *Taxus baccata* | Common yew |
