## Appendix S1 for "Conserving genetic diversity during climate change: Niche marginality and discrepant monitoring capacity in Europe"

Appendix S1. Projects monitoring population genetic diversity in captive/domesticated, fish, and marine species.

The following figures present maps of country-level tallies of the number of population genetic diversity monitoring projects in three categories: captive, fish, and marine species.  Marine fish may appear in both ‘fish’ and ‘marine’ figures.

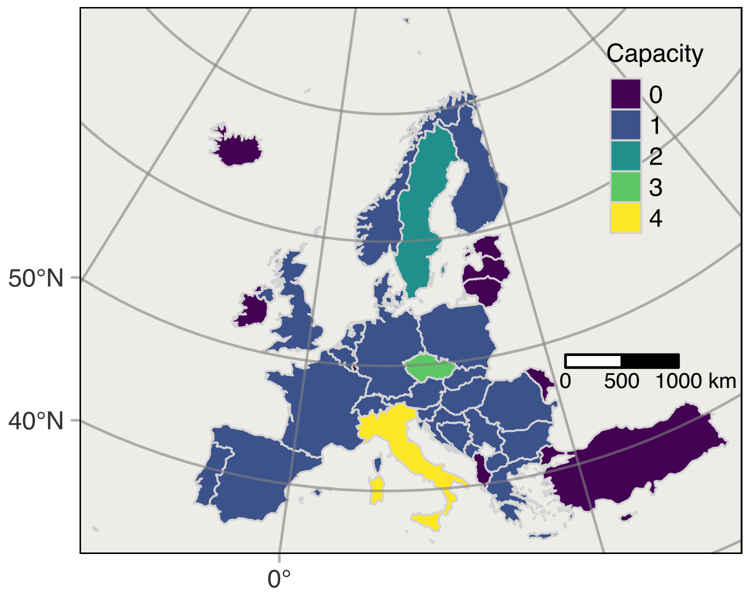


Number of projects monitoring population genetic diversity in captive populations and domesticated species.

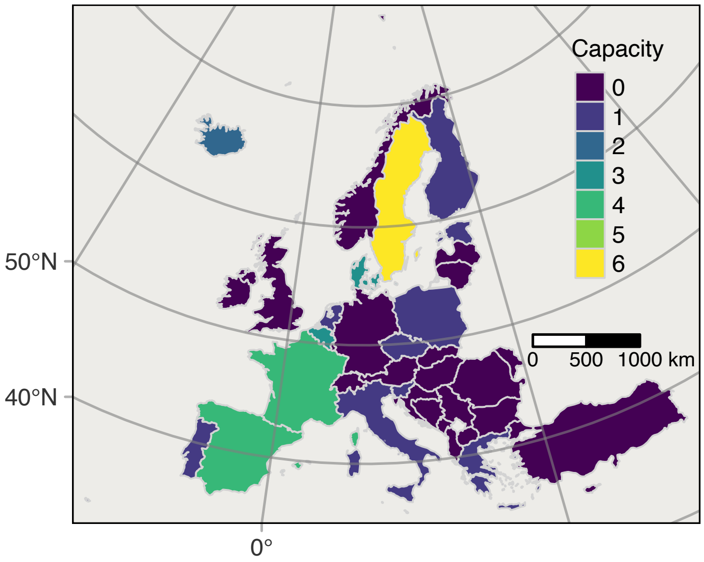


Number of projects monitoring population genetic diversity of fish species, including both freshwater, marine and migratory (e.g., anadromous) populations.

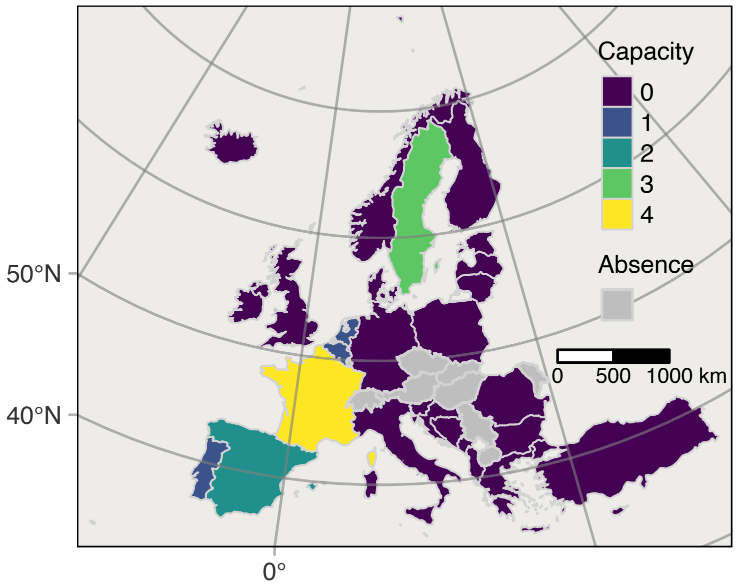


Number of projects monitoring population genetic diversity of marine organisms, including marine fish species.
