## Appendix S2 for "Conserving genetic diversity during climate change: Niche marginality and discrepant monitoring capacity in Europe"

### Monitoring by Country

Peter B. Pearman

2/02/2023

**This script runs a bunch of regressions on GDP and Country area.**

```
library(tidyverse)
```

```
## -- Attaching packages --------------------------------------- tidyverse 1.3.2 --
## v ggplot2 3.4.0     v purrr   1.0.1
## v tibble  3.1.8     v dplyr   1.1.0
## v tidyr   1.3.0     v stringr 1.5.0
## v readr   2.1.3     v forcats 1.0.0
## -- Conflicts ------------------------------------------ tidyverse_conflicts() --
## x dplyr::filter() masks stats::filter()
## x dplyr::lag()    masks stats::lag()
```

```
library(magrittr)
```

```
##
## Attaching package: 'magrittr'
##
## The following object is masked from 'package:purrr':
##
##     set_names
##
## The following object is masked from 'package:tidyr':
##
##     extract
```

```
library(car)
```

```
## Loading required package: carData
##
## Attaching package: 'car'
##
## The following object is masked from 'package:dplyr':
##
##     recode
##
## The following object is masked from 'package:purrr':
##
##     some
```

```
library(ggfortify)
library(ggrepel)
library(scales)
```

```
##
## Attaching package: 'scales'
##
## The following object is masked from 'package:purrr':
##
##     discard
##
## The following object is masked from 'package:readr':
##
##     col_factor
```

```
library(pscl)
```

```
## Classes and Methods for R developed in the
## Political Science Computational Laboratory
## Department of Political Science
## Stanford University
## Simon Jackman
## hurdle and zeroinfl functions by Achim Zeileis
```

```
library(MASS)
```

```
##
## Attaching package: 'MASS'
##
## The following object is masked from 'package:dplyr':
##
##     select
```

```
library(sandwich)
library(DescTools)
```

```
##
## Attaching package: 'DescTools'
##
## The following object is masked from 'package:car':
##
##     Recode
```

```
library(cowplot)
```

#### PGD monitoring programs, T cases

```
rm(list=ls())
# GDP from https://data.worldbank.org/indicator/NY.GDP.PCAP.CD, accessed 24.05.2022
# Country area in Europe from https://en.wikipedia.org/wiki/List_of_European_countries_by_area
# accessed 24.05.2022, which includes only the area of countries in continental Europe.


data.a <- read_csv('country_area_GDP.csv')
```

```
## Rows: 39 Columns: 11
## -- Column specification --------------------------------------------------------
## Delimiter: ","
## chr (6): POSTAL, NAME, NAME_EN, code, name, region
```

```
## dbl (5): ntotal_studies, area_kmsq, per_capita_GDP_CIA, year, Per_cap_GDP_Wo...
##
## i Use `spec()` to retrieve the full column specification for this data.
## i Specify the column types or set `show_col_types = FALSE` to quiet this message.
```

```
data.a <- as_tibble(data.a) %>%
  dplyr::select(POSTAL,area_kmsq,per_capita_GDP_CIA,year,Per_cap_GDP_WoBa_2020dollars)
#data.b <- as_tibble(read_csv('./country_data3_w_U.csv')) %>%
#  arrange(NAME)
data.b <- as_tibble(read_csv('./country_data3_PGD_only.csv')) %>%
  arrange(NAME)
```

```
## Rows: 38 Columns: 23
## -- Column specification --------------------------------------------------------
## Delimiter: ","
## chr  (6): POSTAL, NAME, NAME_EN, code, name, region
## dbl (17): new_Area, ncarn, nbear, nwolf, nlynx, nom, nbird, ninsect, nfish, ...
##
## i Use `spec()` to retrieve the full column specification for this data.
## i Specify the column types or set `show_col_types = FALSE` to quiet this message.
```

```
data1 <- left_join(data.b,data.a,by='POSTAL')

data1 %<>% filter(!NAME %in% c("Liechtenstein")) %>%
  mutate(GDP=Per_cap_GDP_WoBa_2020dollars) %>%
  mutate(Area=as.numeric(area_kmsq), Area_sq = Area*Area,
         lnGDP = log10(GDP), GDP_sq = GDP^2,
         lnArea=log10(area_kmsq),
         lnAreasq=lnArea*lnArea,lnStudies=log10(ntotal_studies+1)) %>%
  dplyr::select(-one_of('Per_cap_GDP_WoBa_2020dollars')) %>%
  mutate(area3 = area_kmsq/1000,GDP5=GDP/10^5,GDPsq5=GDP_sq/10^5) %>%
  arrange(NAME)
```

```
mod <- glm(ntotal_studies ~ area_kmsq +poly(GDP,2,raw=TRUE), family = poisson(link = "log"),data=data1)

summary(mod)
```

```
##
## Call:
## glm(formula = ntotal_studies ~ area_kmsq + poly(GDP, 2, raw = TRUE),
##     family = poisson(link = "log"), data = data1)
##
## Deviance Residuals:
##      Min        1Q    Median        3Q       Max
## -2.78259  -1.19696  -0.00034   0.56841   2.28067
##
## Coefficients:
##                             Estimate Std. Error z value Pr(>|z|)
## (Intercept)               -2.910e-01  2.926e-01  -0.994  0.32001
## area_kmsq                  1.165e-06  4.202e-07   2.773  0.00555 **
## poly(GDP, 2, raw = TRUE)1  8.262e-05  1.551e-05   5.327 1.00e-07 ***
## poly(GDP, 2, raw = TRUE)2 -8.483e-10  1.853e-10  -4.578 4.69e-06 ***
## ---
## Signif. codes:  0 '***' 0.001 '**' 0.01 '*' 0.05 '.' 0.1 ' ' 1
##
## (Dispersion parameter for poisson family taken to be 1)
```

```
##
##     Null deviance: 138.313  on 37  degrees of freedom
## Residual deviance:  79.678  on 34  degrees of freedom
## AIC: 185.24
##
## Number of Fisher Scoring iterations: 5
```

The Poisson regression doesn't fit very well. Still strongly over-dispersed even when using quasipoisson()

```
yhat <- predict(mod,type="response")
z <- data1$ntotal_studies/sqrt(yhat)
cat("overdispersion ratio is ", sum(z^2)/(nrow(data1)-3),"\n")
```

```
## overdispersion ratio is  6.253132
```

```
cat("p-value of the overdispersion test is ", pchisq(sum(z^2),df=nrow(data1)-3))
```

```
## p-value of the overdispersion test is  1
```

```
ggplot(data1, aes(ntotal_studies)) +
  geom_histogram()
```

```
## `stat_bin()` using `bins = 30`. Pick better value with `binwidth`.
```

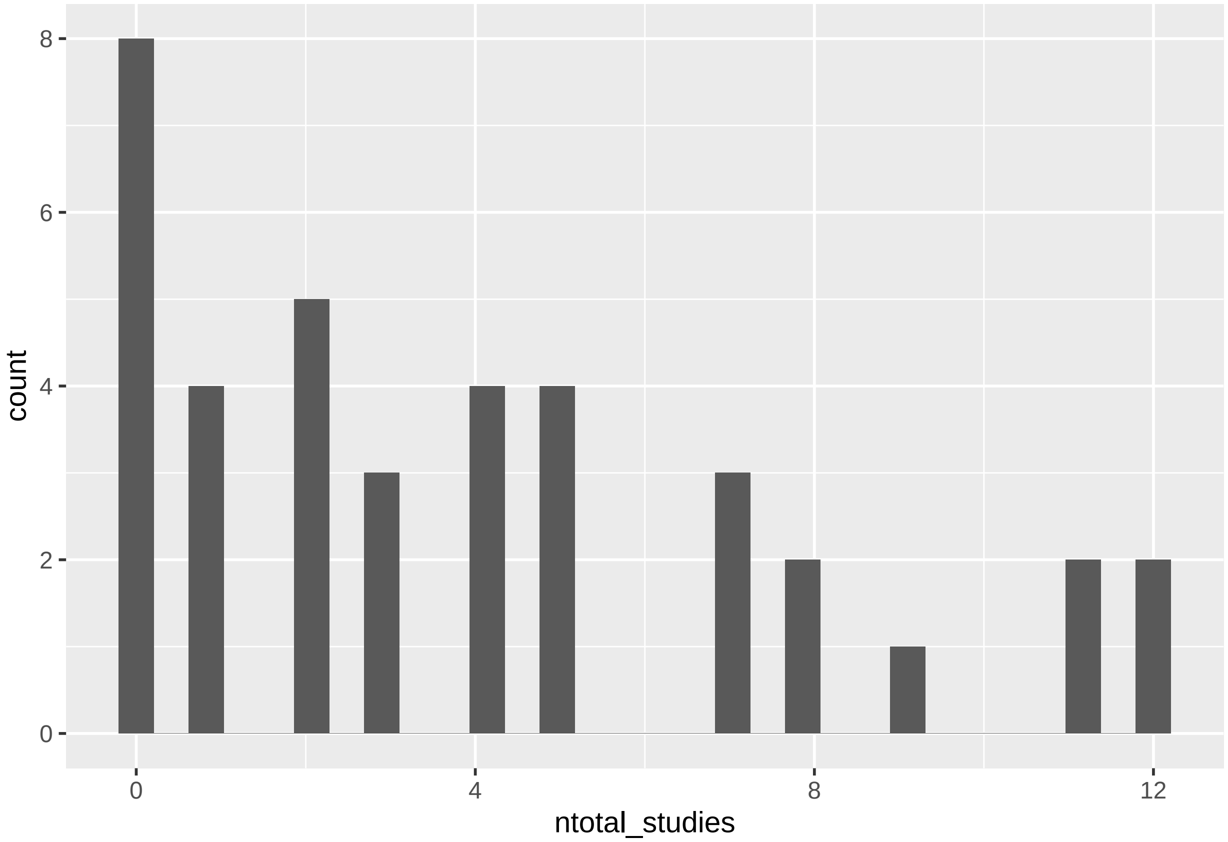


```
library(pscl)
library(MASS)
```

```
summary(mod1 <- zeroinfl(ntotal_studies ~ area3+poly(GDP5,2,raw=TRUE) | GDP5  , data=data1))
```

```
##
## Call:
```

```
## zeroinfl(formula = ntotal_studies ~ area3 + poly(GDP5, 2, raw = TRUE) |
##     GDP5, data = data1)
##
## Pearson residuals:
##      Min       1Q   Median       3Q      Max
## -1.56645 -0.83096  0.02589  0.47442  2.58156
##
## Count model coefficients (poisson with log link):
##                              Estimate Std. Error z value Pr(>|z|)
## (Intercept)                 0.2606055  0.3040176   0.857 0.391331
## area3                       0.0012112  0.0005104   2.373 0.017635 *
## poly(GDP5, 2, raw = TRUE)1  6.5387622  1.7143453   3.814 0.000137 ***
## poly(GDP5, 2, raw = TRUE)2 -7.2422049  2.0059015  -3.610 0.000306 ***
##
## Zero-inflation model coefficients (binomial with logit link):
##             Estimate Std. Error z value Pr(>|z|)
## (Intercept)  -0.2384     0.8711  -0.274    0.784
## GDP5         -5.8264     3.7993  -1.534    0.125
## ---
## Signif. codes:  0 '***' 0.001 '**' 0.01 '*' 0.05 '.' 0.1 ' ' 1
##
## Number of iterations in BFGS optimization: 14
## Log-likelihood: -80.79 on 6 Df
```

```
summary(mod2 <- glm.nb(formula=ntotal_studies ~ area3+poly(GDP,2),data=data1))
```

```
##
## Call:
## glm.nb(formula = ntotal_studies ~ area3 + poly(GDP, 2), data = data1,
##     init.theta = 3.600571833, link = log)
##
## Deviance Residuals:
##      Min        1Q    Median        3Q       Max
## -2.29740  -0.83022  -0.02857   0.40634   1.58077
##
## Coefficients:
##                Estimate Std. Error z value Pr(>|z|)
## (Intercept)    0.9699799  0.1774861   5.465 4.63e-08 ***
## area3          0.0010654  0.0006637   1.605 0.108468
## poly(GDP, 2)1  0.2172634  1.0930483   0.199 0.842444
## poly(GDP, 2)2 -4.4938641  1.2497818  -3.596 0.000323 ***
## ---
## Signif. codes:  0 '***' 0.001 '**' 0.01 '*' 0.05 '.' 0.1 ' ' 1
##
## (Dispersion parameter for Negative Binomial(3.6006) family taken to be 1)
##
##     Null deviance: 75.499  on 37  degrees of freedom
## Residual deviance: 46.250  on 34  degrees of freedom
## AIC: 178.57
##
## Number of Fisher Scoring iterations: 1
##
##
##               Theta:  3.60
##           Std. Err.:  1.98
```

```
##
##  2 x log-likelihood:  -168.569
```

The negative binomial regression is a significantly better fit than the poisson regression __

```
pchisq(2 * (logLik(mod2) - logLik(mod)), df = 1, lower.tail = FALSE)
```

```
## 'log Lik.' 0.003240003 (df=5)
```

```
yhat <- predict(mod2,type="response")
z <- data1$ntotal_studies/sqrt(yhat)
cat("The sum of the squared standardized residuals is ",sum(z^2),"\n")
```

```
## The sum of the squared standardized residuals is  218.3741
```

```
cat("The expected value for the sum of the squared standardized residuals is",nrow(data1)-3,"\n")
```

```
## The expected value for the sum of the squared standardized residuals is 35
```

```
cat("overdispersion ratio is ", sum(z^2)/(nrow(data1)-3),"\n")
```

```
## overdispersion ratio is  6.239261
```

```
cat("p-value of the overdispersion test is ", pchisq(sum(z^2),df=nrow(data1)-3), "\n")
```

```
## p-value of the overdispersion test is  1
```

```
cat("The factor to correct regression standard errors is ",sqrt(sum(z^2)/(nrow(data1)-3)))
```

```
## The factor to correct regression standard errors is  2.497851
```

```
cor.fac <- sqrt(sum(z^2)/(nrow(data1)-3))
```

Calculate robust standard errors using the package 'sandwich' Confidence using robust standard errors shows only intercept and GDP^2 are significant

```
cov.mod2 <- vcovHC(mod2, type="HC0")
std.err <- sqrt(diag(cov.mod2)) #* cor.fac      # Application of the correction factor results in non-s
r.est <- cbind(Estimate= coef(mod2), "Robust SE" = std.err,
"Pr(>|z|)" = 2 * pnorm(abs(coef(mod2)/std.err), lower.tail=FALSE),
LL = coef(mod2) - 1.96 * std.err,
UL = coef(mod2) + 1.96 * std.err)

r.est
```

```
##                    Estimate    Robust SE     Pr(>|z|)            LL           UL
## (Intercept)    0.969979898 0.1800348466 7.135147e-08  0.6171115982  1.322848197
## area3          0.001065388 0.0007064293 1.315211e-01 -0.0003192137  0.002449989
## poly(GDP, 2)1  0.217263363 1.2155320253 8.581424e-01 -2.1651794063  2.599706133
## poly(GDP, 2)2 -4.493864086 1.2811215316 4.519002e-04 -7.0048622875 -1.982865884
```

###Remove Turkey for subsequent analyses below Here is the result without Turkey. Area and GDP^2 are significant

```
data1a <- data1 %>%
  filter(NAME != "Turkey")
summary(mod2a <- glm.nb(formula=ntotal_studies ~ area3+poly(GDP,2),data=data1a))
```

```
##
## Call:
## glm.nb(formula = ntotal_studies ~ area3 + poly(GDP, 2), data = data1a,
##     init.theta = 4.973684807, link = log)
```

```
##
## Deviance Residuals:
##      Min        1Q    Median        3Q       Max
## -2.29009  -0.95022   0.00816   0.40982   1.74260
##
## Coefficients:
##                 Estimate Std. Error z value Pr(>|z|)
## (Intercept)    0.9212916  0.1699262   5.422  5.9e-08 ***
## area3          0.0018005  0.0007446   2.418  0.01561 *
## poly(GDP, 2)1 -0.0011630  1.0060392  -0.001  0.99908
## poly(GDP, 2)2 -3.6952140  1.2018836  -3.075  0.00211 **
## ---
## Signif. codes:  0 '***' 0.001 '**' 0.01 '*' 0.05 '.' 0.1 ' ' 1
##
## (Dispersion parameter for Negative Binomial(4.9737) family taken to be 1)
##
##     Null deviance: 79.286  on 36  degrees of freedom
## Residual deviance: 45.266  on 33  degrees of freedom
## AIC: 172.63
##
## Number of Fisher Scoring iterations: 1
##
##
##               Theta:  4.97
##           Std. Err.:  3.18
##
##  2 x log-likelihood:  -162.629
```

Fit poisson regression w/o Turkey

```
mod_p <- glm(ntotal_studies ~ area3 +poly(GDP,2), family = poisson(link = "log"),data=data1a)

summary(mod_p)
```

```
##
## Call:
## glm(formula = ntotal_studies ~ area3 + poly(GDP, 2), family = poisson(link = "log"),
##     data = data1a)
##
## Deviance Residuals:
##      Min        1Q    Median        3Q       Max
## -2.64015  -1.27320  -0.05605   0.55690   2.74836
##
## Coefficients:
##                 Estimate Std. Error z value Pr(>|z|)
## (Intercept)    0.9267872  0.1311376   7.067 1.58e-12 ***
## area3          0.0017953  0.0004923   3.647 0.000265 ***
## poly(GDP, 2)1 -0.2667098  0.8313079  -0.321 0.748338
## poly(GDP, 2)2 -3.6187803  1.0009075  -3.615 0.000300 ***
## ---
## Signif. codes:  0 '***' 0.001 '**' 0.01 '*' 0.05 '.' 0.1 ' ' 1
##
## (Dispersion parameter for poisson family taken to be 1)
##
##     Null deviance: 130.259  on 36  degrees of freedom
```

```
## Residual deviance:  70.467  on 33  degrees of freedom
## AIC: 176.02
##
## Number of Fisher Scoring iterations: 5
```

The negative binomial regression is a significantly better fit than the poisson regression _

```
pchisq(2 * (logLik(mod2a) - logLik(mod_p)), df = 1, lower.tail = FALSE)
```

```
## 'log Lik.' 0.02020007 (df=5)
```

Without Turkey, Area effect is significant. Analysis is done using Area as number of 1000s of sq KM's.

```
summary(mod2b <- glm.nb(formula=ntotal_studies ~ area3,data=data1a))
```

```
##
## Call:
## glm.nb(formula = ntotal_studies ~ area3, data = data1a, init.theta = 2.499916087,
##     link = log)
##
## Deviance Residuals:
##     Min       1Q   Median       3Q      Max
## -1.9742  -0.8628  -0.3508   0.4037   2.4822
##
## Coefficients:
##              Estimate Std. Error z value Pr(>|z|)
## (Intercept) 0.8942072  0.1915754   4.668 3.05e-06 ***
## area3       0.0028772  0.0008297   3.468 0.000525 ***
## ---
## Signif. codes:  0 '***' 0.001 '**' 0.01 '*' 0.05 '.' 0.1 ' ' 1
##
## (Dispersion parameter for Negative Binomial(2.4999) family taken to be 1)
##
##     Null deviance: 59.646  on 36  degrees of freedom
## Residual deviance: 46.190  on 35  degrees of freedom
## AIC: 181.09
##
## Number of Fisher Scoring iterations: 1
##
##
##               Theta:  2.50
##           Std. Err.:  1.20
##
##  2 x log-likelihood:  -175.091
```

```
pseudo_Rsq <- PseudoR2(mod2b,which="all")
pseudo_Rsq
```

```
##        McFadden     McFaddenAdj        CoxSnell      Nagelkerke   AldrichNelson
##      0.05711110      0.03557052      0.24921043      0.25086928      0.22277570
## VeallZimmermann           Efron McKelveyZavoina            Tjur             AIC
##      0.26716383      0.42328436              NA              NA    181.09074093
##             BIC          logLik         logLik0              G2
##    185.92349467    -87.54537047    -92.84802309     10.60530525
```

###Hermite regression Area and the squared GDP are also significant in Hermite regression

```
library(hermite)
```

```
## Loading required package: maxLik

## Loading required package: miscTools

##
## Please cite the 'maxLik' package as:
## Henningsen, Arne and Toomet, Ott (2011). maxLik: A package for maximum likelihood estimation in R. Co
##
## If you have questions, suggestions, or comments regarding the 'maxLik' package, please use a forum or
## https://r-forge.r-project.org/projects/maxlik/
```

```
mod3 <- glm.hermite(ntotal_studies ~ area3+poly(GDP5,2),
                    data = data1a,
                    link = "log",
                    m=2)
```

```
## Warning in model.matrix.default(mt, mf, contrasts): non-list contrasts argument
## ignored
```

```
summary(mod3)
```

```
## Call:
## glm.hermite(formula = ntotal_studies ~ area3 + poly(GDP5, 2),
##     data = data1a, link = "log", m = 2)
##
## Deviance Residuals:
##         Min          1Q      Median          3Q         Max
## -11.8978147  -3.9602302  -2.2468776   0.4910989 539.9487000
##
## Coefficients:
##                        Estimate   Std. Error    z value      p-value
## (Intercept)         0.925858476 0.1688239374  5.4841659 4.154249e-08
## area3               0.001867181 0.0006397753  2.9184943 3.517263e-03
## poly(GDP5, 2)1     -0.108320017 1.0147630645 -0.1067441 9.149920e-01
## poly(GDP5, 2)2     -3.375445747 1.2376000803 -2.7274124 6.383321e-03
## dispersion.index    1.671424842 0.1589042420  8.3612580 1.916526e-03
## order               2.000000000           NA         NA           NA
## (Likelihood ratio test against Poisson is reported by *z value* for *dispersion.index*)
##
## AIC:  169.7343
```

The Hermite model fits better than the Poisson model.

```
pchisq(2 * (mod3$loglik - logLik(mod_p)), df = 1, lower.tail = FALSE)
```

```
## 'log Lik.' 0.003987192 (df=4)
```

The Hermite model does not fit the data better than the Negative Binomial model

```
pchisq(2 * (mod3$loglik - logLik(mod2a)), df = 1, lower.tail = FALSE)
```

```
## 'log Lik.' 0.0888483 (df=5)
```

The Hermite regression residuals are overdispersed compared to the Poisson expectation

```
yhat <- mod3$fitted.values
z <- data1a$ntotal_studies/sqrt(yhat)
cat("The sum of the squared standardized residuals is ",sum(z^2),"\n")
```

```
## The sum of the squared standardized residuals is  213.7942
```

```
cat("The expected value for the sum of the squared standardized residuals is",nrow(data1a)-3,"\n")
```

```
## The expected value for the sum of the squared standardized residuals is 34
```

```
cat("overdispersion ratio is ", sum(z^2)/(nrow(data1)-3),"\n")
```

```
## overdispersion ratio is  6.108405
```

```
cat("p-value of the overdispersion test is ", pchisq(sum(z^2),df=nrow(data1a)-3), "\n")
```

```
## p-value of the overdispersion test is  1
```

```
cat("The factor to correct regression standard errors is ",sqrt(sum(z^2)/(nrow(data1a)-3)))
```

```
## The factor to correct regression standard errors is  2.507601
```

```
cor.fac <- sqrt(sum(z^2)/(nrow(data1a)-3))
```

###Autoplot() plots showing influence of Turkey mod2 was created on data1, which included Turkey

```
data1 <- data1%>%
  mutate(nb.residuals=residuals(mod2,type="pearson"),predicted = mod2$fitted.values)
```

Here is why it's a good idea to remove Turkey. It's a high leverage data point.

```
(autoplot(mod2))
```

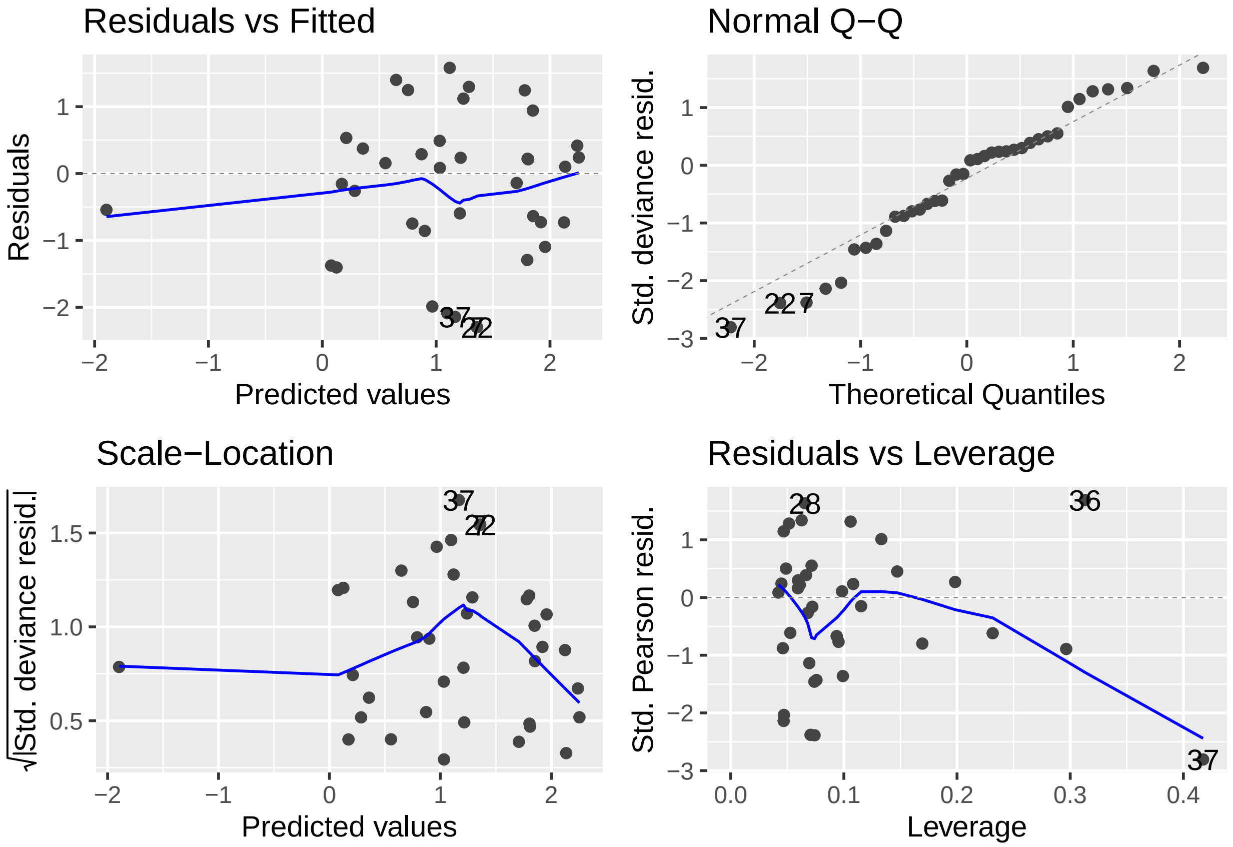


```
cooks_2 <- cooks.distance(mod2)
sample_size <- nrow(data1)
```

```
plot(cooks_2, pch="*", cex=2, main="Influential Obs by Cooks distance")  # plot cook's distance
abline(h = 4/sample_size, col="red")  # add cutoff line
text(x=1:length(cooks_2)+1, y=cooks_2, labels=ifelse(cooks_2>4/sample_size, names(cooks_2),""), col="re
```

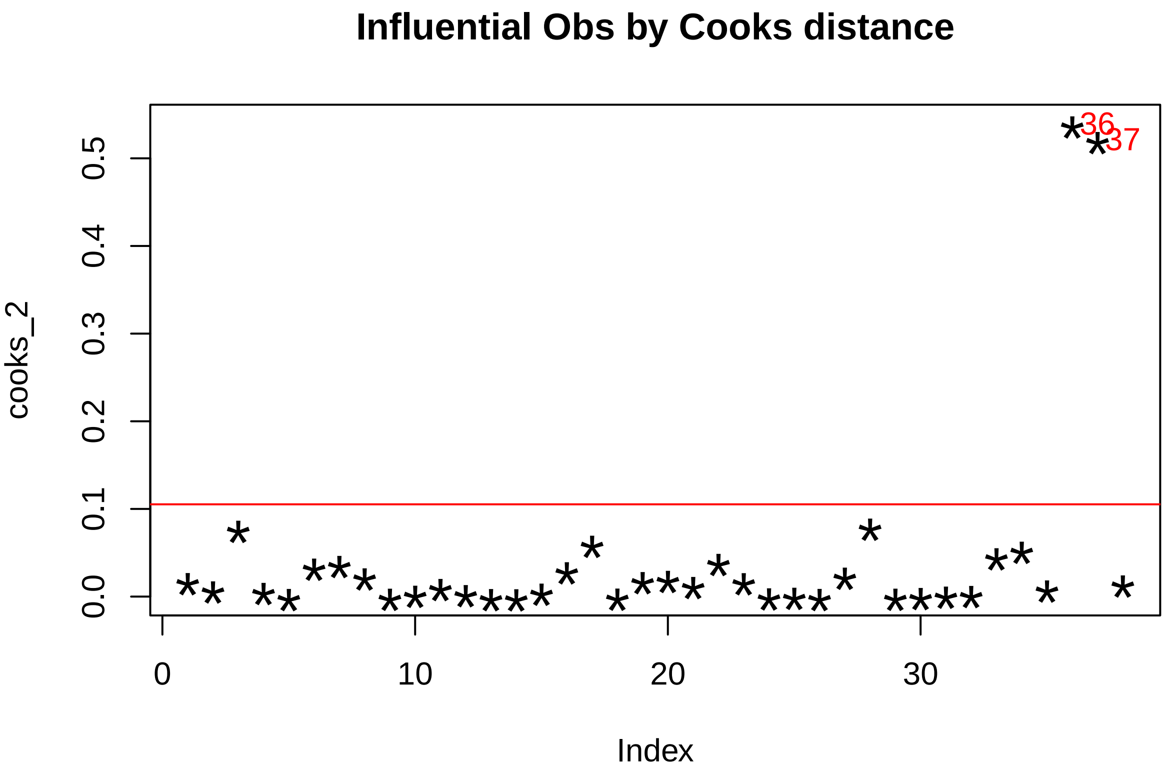


```
#summary(mod2a <- glm.nb(formula=ntotal_studies ~ area_kmsq+poly(GDP,2),data=data1))
```

Including Turkey, data1

```
plot1 <- ggplot(data1,aes(x=lnArea,y=ntotal_studies,color=GDP5)) +
  geom_point(size=3) +
  scale_color_gradient(low='blue',high='red',
                       limits=round(c(min(data1$lnGDP),max(data1$lnGDP)),digits = 2),
                       breaks=round(c(min(data1$lnGDP), sum(range(data1$lnGDP))/2 ,max(data1$lnGDP)),di
  labs(x="Log10 Square Kilometers",y="Total Number Monitoring Projects") +
  #guides(color=guide_legend("log10(GDP in USD)"))+
  labs(color="log10(GDP in USD)") +
  scale_x_continuous(breaks = c(2.0,2.5,3.0,3.5,4.0,4.5,5.0,5.5,6.0), limits = c(2.0,6.5)) +
  theme(panel.grid.minor = element_blank(),
    panel.grid.major = element_blank(),
    panel.background = element_rect(fill= alpha('medium sea green', alpha=0.2)),
    axis.line= element_line(size = 0.75, color = "black"),
    axis.text= element_text(size=rel(1.0)),
    axis.title = element_text(size=rel(1.3)),
    legend.title.align = 0) +
  geom_label_repel(aes(label=POSTAL),
    size = 3,
    box.padding   = 0.35,
    point.padding = 0.5,
    label.size=0.1,
    max.overlaps = 20,
```

```
    fill=NA,
    segment.color = 'grey50')
```

```
## Warning: The `size` argument of `element_line()` is deprecated as of ggplot2 3.4.0.
## i Please use the `linewidth` argument instead.
```

```
plot1
```

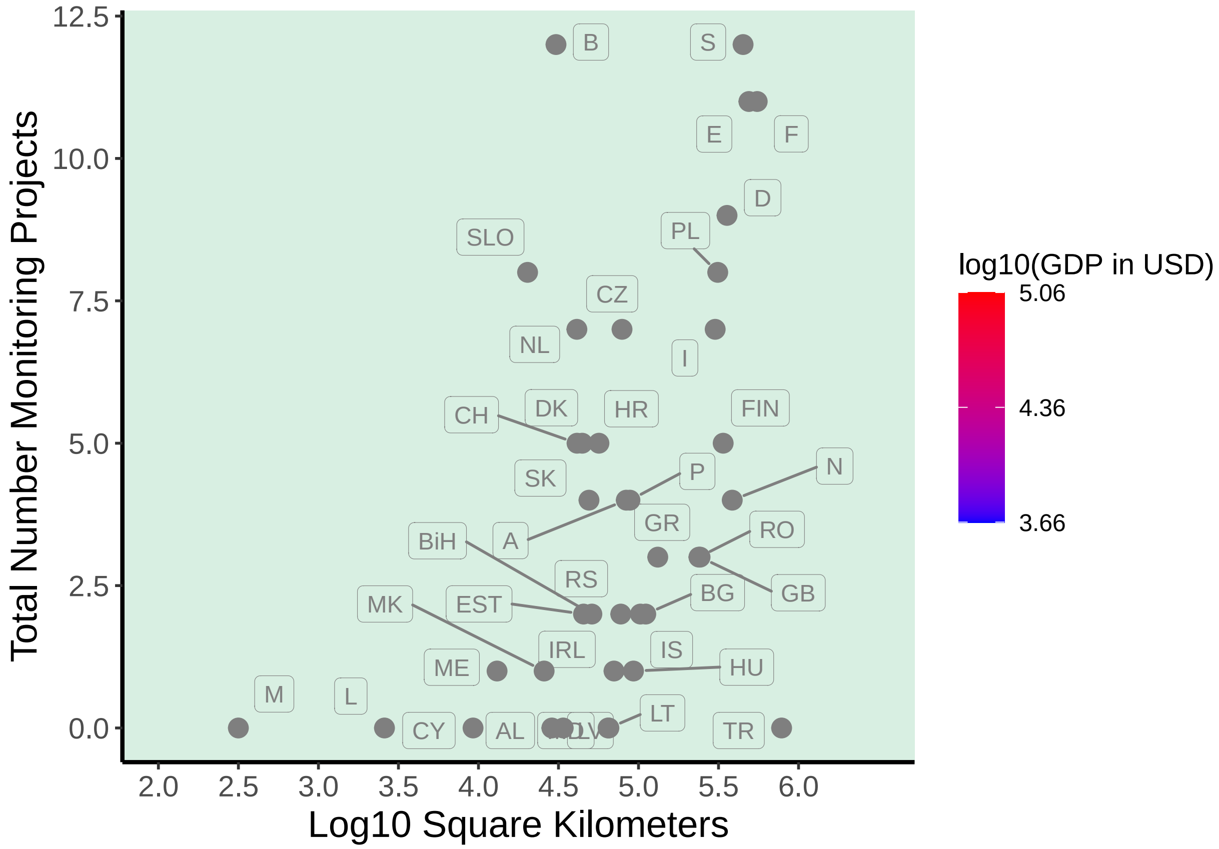


###Scatter plots, data with Turkey

```
# manually jitter RO and GB
#data1a <- data1
#data1a$Area[which(data1a$POSTAL=='GB')] <- data1a$Area[which(data1a$POSTAL=='GB')] + 5000
#data1a$Area[which(data1a$POSTAL=='RO')] <- data1a$Area[which(data1a$POSTAL=='RO')] - 5000


plot1a <- ggplot(data1,aes(x=Area,y=ntotal_studies)) +
  geom_point(aes(color=GDP),size=3) +
  #scale_color_continuous(formatter=comma)*
  scale_color_gradient(low='blue',high='red',
                       limits=c(min(data1$GDP),max(data1$GDP)),
                       breaks=c(min(data1$GDP), sum(range(data1$GDP))/2 ,max(data1$GDP)),
                       labels=scales::label_dollar())+
                       #labels = c(paste("a",100),"b","c"))+
  labs(x="Square Kilometers",y="Total Number Monitoring Projects") +
  #guides(color=guide_legend("log10(GDP in USD)"))+
  labs(color="GDP 2020 in USD") +
  scale_x_continuous(labels = scales::comma)+
```

```
  #scale_x_continuous(breaks = c(2.0,2.5,3.0,3.5,4.0,4.5,5.0,5.5,6.0), limits = c(2.0,6.5)) +
  theme(panel.grid.minor = element_blank(),
    panel.grid.major = element_blank(),
    panel.background = element_rect(fill= alpha('medium sea green', alpha=0.2)),
    axis.line= element_line(size = 0.75, color = "black"),
    axis.text= element_text(size=rel(1.0)),
    axis.title = element_text(size=rel(1.3)),
    legend.title.align = 0) +
  geom_label_repel(aes(label=POSTAL),
    size=2,
    max.overlaps = 19,
    box.padding   = 0.35,
    point.padding = 0.5,
    label.size=0.05,
    min.segment.length = 0.2,
    segment.color = 'grey50')

plot1a
```

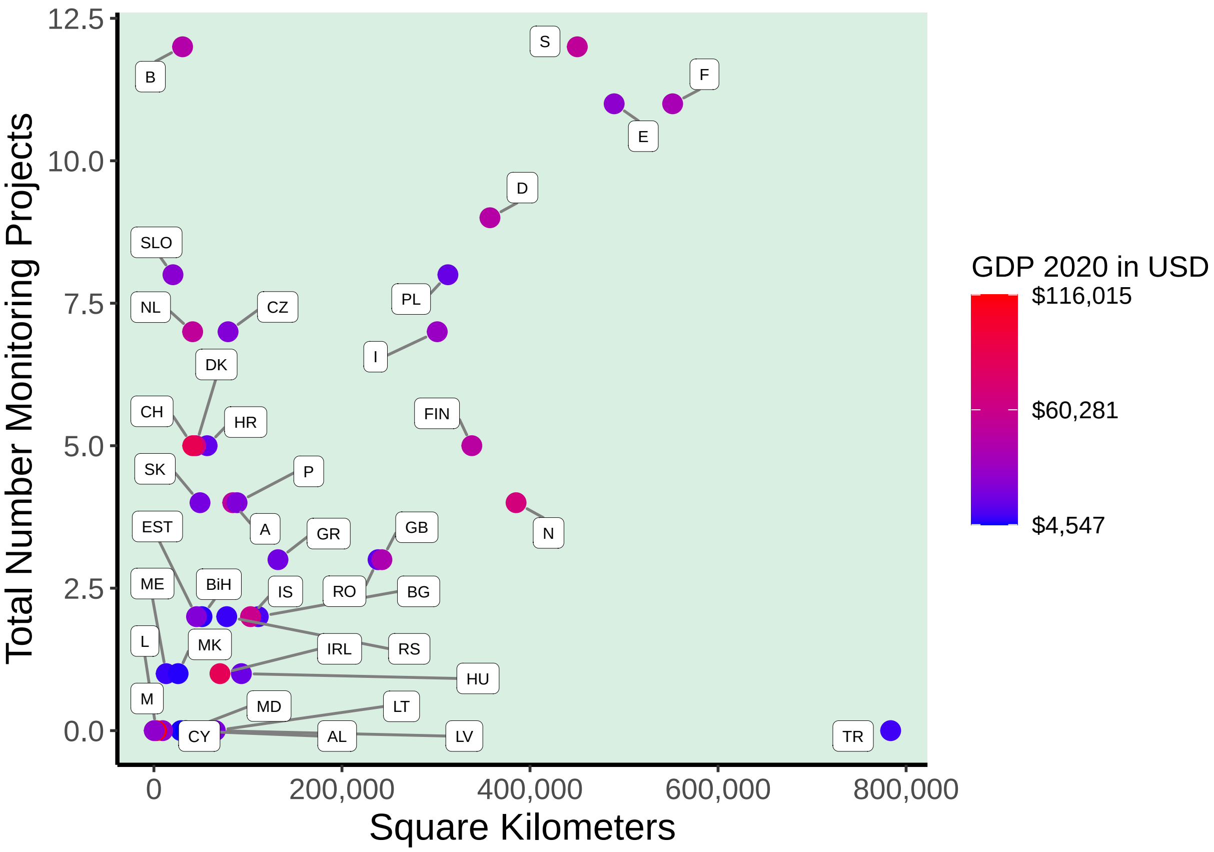


```
plot2 <- ggplot(data1,aes(x=lnGDP, y=ntotal_studies, color=area_kmsq)) +
  geom_point(size=3) +   #position=position_dodge(width = 0.2),
  scale_color_gradient(low='blue',high='red')+
  theme(
    panel.grid.minor = element_blank(),
    panel.grid.major = element_blank(),
    panel.background = element_rect(fill= alpha('medium sea green', alpha=0.2)),
    axis.line= element_line(size = 0.75, color = "black"),
```

```
    axis.text= element_text(size=rel(1.0)),
    axis.title = element_text(size=rel(1.3)))+
  labs(x="GDP in log10 Dollars",y= "Number Genetic Monitoring Projects") +
    geom_label_repel(aes(label=POSTAL),
    size=2,
    max.overlaps = 19,
    box.padding   = 0.35,
    point.padding = 0.5,
    label.size=0.05,
    segment.color = 'grey50')


plot2
```

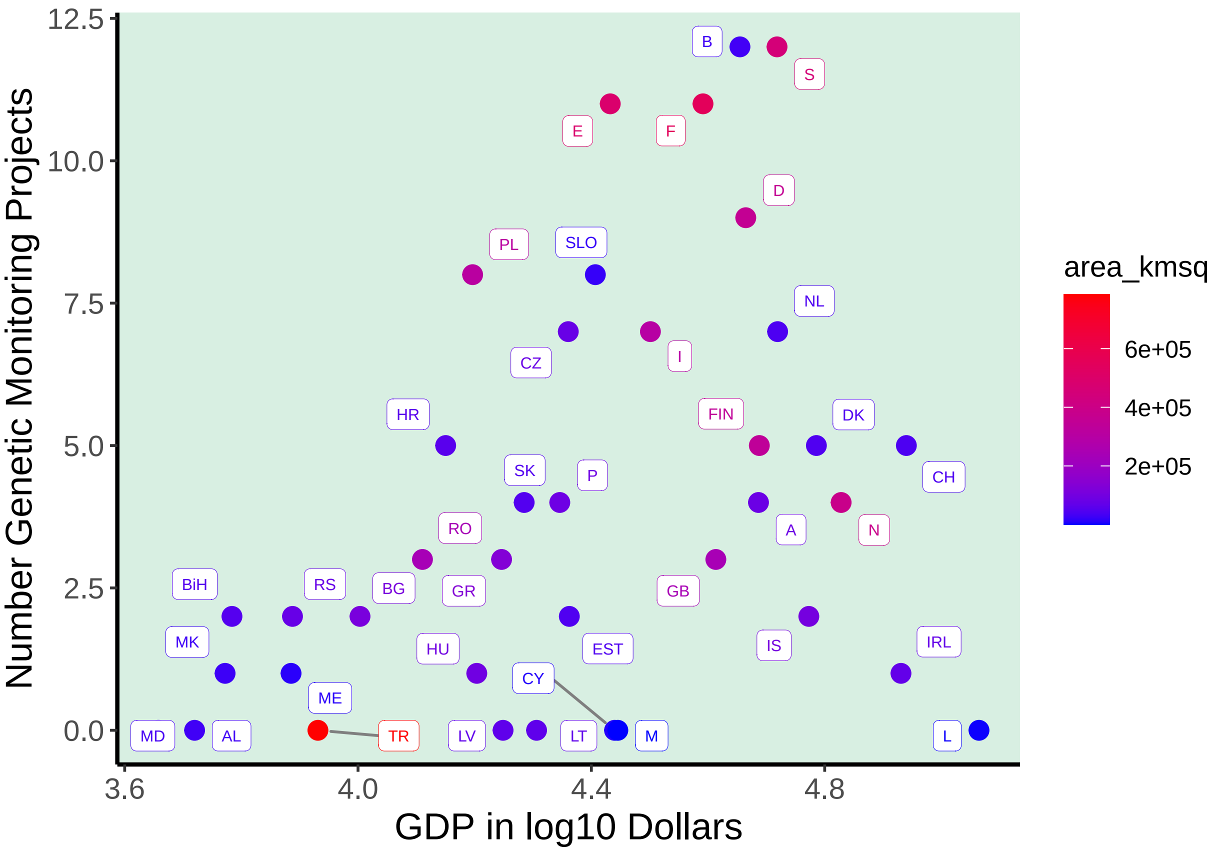


###Re-do nb model with package trending

```
library(ciTools)
```

```
## ciTools version 0.6.1 (C) Institute for Defense Analyses
```

```
library(trending)
```

```
data1a <- data1 %>%
   filter(!NAME=="Turkey")
```

Turkey is not included in data1a. This uses straight sq km values in the variable Area, for easier plotting

```
set.seed(12345)
nb_mod1 <- glm_nb_model(ntotal_studies ~ Area)
```

```
fitted_nb1 <- fit(nb_mod1,data1a)
new_Area <- c(seq(from=min(data1a$Area), to=max(data1a$Area), by=500),max(data1a$Area))
new_data1 <- tibble(Area=as.integer(new_Area),index=1:length(new_Area))
data_nbmod1ci <- predict(fitted_nb1, new_data1, simulate_pi=FALSE,uncertain=FALSE) %>%
  mutate(ntotal_studies=estimate)

get_model(fitted_nb1)
```

```
##
## Call:  MASS::glm.nb(formula = ntotal_studies ~ Area, data = data, init.theta = 2.499916087,
##     link = log)
##
## Coefficients:
## (Intercept)         Area
##   8.942e-01    2.877e-06
##
## Degrees of Freedom: 36 Total (i.e. Null);  35 Residual
## Null Deviance:       59.65
## Residual Deviance: 46.19     AIC: 181.1
```

```
theme_set(theme_grey(base_size = 5))
aspect=0.8
```

The negative binomial regression line is plotted having omitted Turkey as an outlier

```
data1a <- data1a %>%
  mutate(Area_jit = jitter(Area,factor=4))

set.seed(12345)
plot1 <- ggplot(data1a,aes(x=Area_jit,y=ntotal_studies)) +
  geom_point(aes(color=GDP),size=rel(1)) +
  scale_color_gradient(low='blue',high='red',
                       limits=c(min(data1a$GDP),max(data1a$GDP)),
                       breaks=c(min(data1a$GDP), sum(range(data1a$GDP))/2 ,max(data1a$GDP)),
                       labels=scales::label_dollar())+
  geom_ribbon(aes(x=Area,ymin=lower_ci,ymax=upper_ci),data_nbmod1ci,color='gray',alpha=0.1,linetype=0)+
  geom_line(aes(Area,estimate),data_nbmod1ci)+
  labs(x="Area in Square Km",y="Realized Monitoring Capacity") +
  #labs(title = "Country Extent vs. Number Monitoring Projects") +
  #guides(color=guide_legend("GDP in 2020 USD")+
  labs(color="GDP in\n2020 USD") +
  #scale_x_continuous(breaks = c(2.0,2.5,3.0,3.5,4.0,4.5,5.0,5.5,6.0), limits = c(2.0,6.5)) +
  scale_x_continuous(breaks=c(0,50000,200000,300000,400000,500000),limits=c(-50000,570000),labels = scal
  scale_y_continuous(breaks = c(0,3,6,9,12,15,18,21,24), limits = c(-4.0,25)) +
  theme(panel.grid.minor = element_blank(),
    panel.grid.major = element_blank(),
    panel.background = element_rect(fill= alpha('medium sea green', alpha=0.2)),
    axis.line= element_line(size = 0.75, color = "black"),
    axis.text.x= element_text(size=rel(1.3)),
    axis.text.y= element_text(size=rel(1.6)),
    axis.title = element_text(size=rel(1.6)),
    legend.title.align = 0,
    legend.text = element_text(size = rel(1.0)),
    plot.title = element_text(size=rel(1.6),hjust = 0.5)) +
  geom_label_repel(aes(label=POSTAL),
```

```r
    size = rel(1.6),
    max.time = 2.0,
    max.iter = 1000000,
    label.padding = 0.1,
    box.padding   = 0.1,
    point.padding = 0.20,
    label.size=NA,
    max.overlaps = Inf,
    min.segment.length = 0,
    segment.angle = 10,
    fill=NA,
    segment.color = 'grey50',
    segment.size = 0.3)
  #annotate(geom="text", x=-50000, y=21, label="90% quantile", color="red", size=5/.pt)

#equation = expression(paste("Projects = exp(8.945*10"^"-1"," + 2.844*10"^"-6"," *Area)"))

plot1 <- plot1 +
   annotate(geom="text", x=205000, y=21,
            label= 'paste("Capacity = exp(8.945*10"^"-1"," + 2.844*10"^"-6"," *Area)")', parse=TRUE,
size=5/.pt)

plot1 <- plot1 +
   annotate(geom="text", x=44000, y=20,
            label= 'paste("R"^"2"," = 0.27")', parse=TRUE, size=5/.pt)



plot1
```

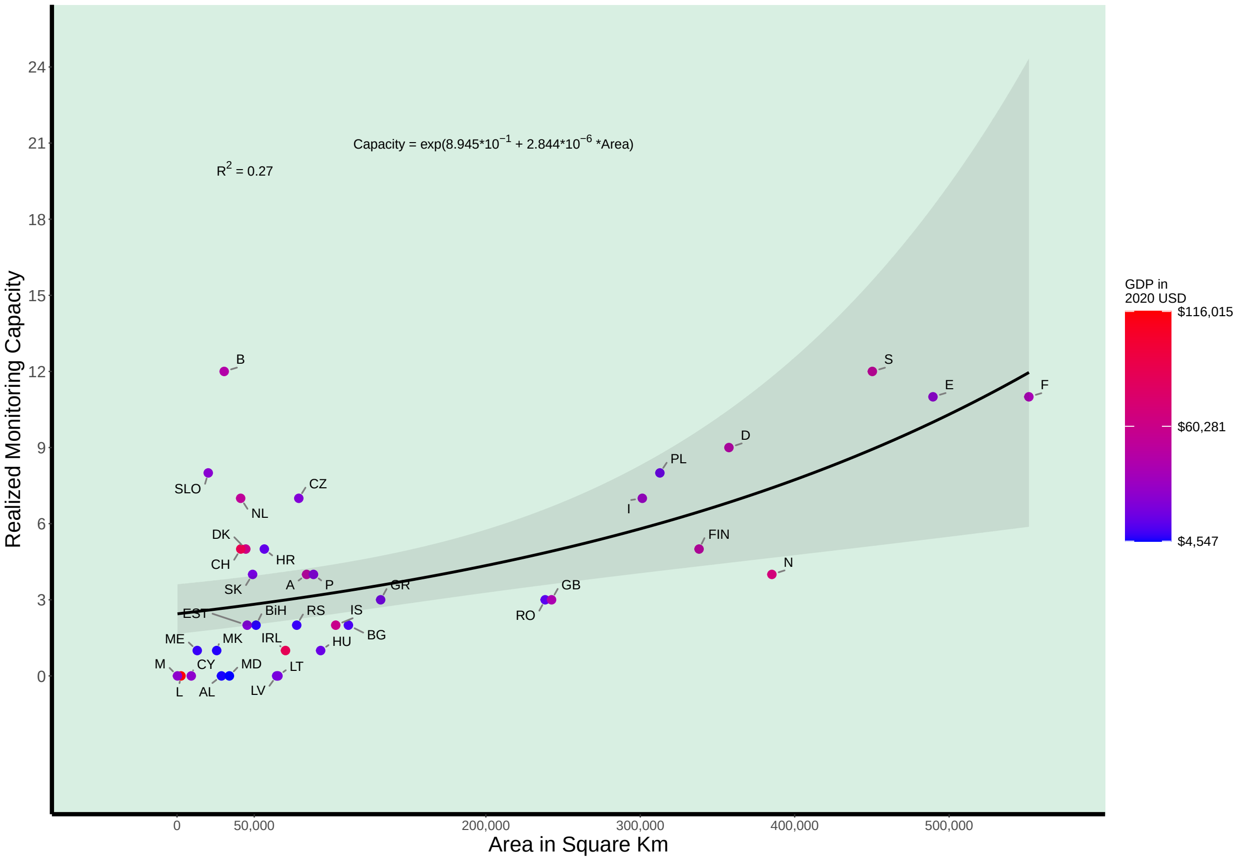


```
#ggsave("PGD_projects_vs_Area_wo_Türkiye_nbreg.pdf", plot=plot1,device = "pdf", width = 8,height=6, uni
ggsave("PGD_projects_vs_Area_wo_Türkiye_nbreg.pdf",plot = plot1,device = "pdf", height = 88*aspect,widt
ggsave("PGD_projects_vs_Area_wo_Türkiye_nbreg.svg",plot = plot1, height = 88*aspect,width = 88,units =
```

```
get_model(fitted_nb1)
```

```
##
## Call:  MASS::glm.nb(formula = ntotal_studies ~ Area, data = data, init.theta = 2.499916087,
##     link = log)
##
## Coefficients:
## (Intercept)         Area
##   8.942e-01    2.877e-06
##
## Degrees of Freedom: 36 Total (i.e. Null);  35 Residual
## Null Deviance:       59.65
## Residual Deviance: 46.19     AIC: 181.1
```

make and plot an nb_model with only polynomial effects of GDP, w/o Turkey

```
set.seed(12345)
polynom <- poly(data1a$GDP,2)
polynom1 <- as_tibble(polynom) %>%
  rename(GDP_orth = 1,GDP_sq_orth = 2)
data1a$GDP_orth <- polynom1$GDP_orth; data1a$GDP_sq_orth <- polynom1$GDP_sq_orth

nb_mod2 <- glm_nb_model(ntotal_studies ~ GDP_orth + GDP_sq_orth)
fitted_nb2 <- fit(nb_mod2,data1a)
```

```r
new_GDP <- as.integer(c(seq(from=min(data1a$GDP), to=max(data1a$GDP), by=500),max(data1a$GDP)))
polynom_pr <- as_tibble(predict(polynom,new_GDP)) %>%
  rename(GDP_orth =1,GDP_sq_orth=2)
#polynom_pr$GDP <- new_GDP
#new_data1 <- tibble(Area=as.integer(new_Area),index=1:length(new_Area))
data_nbmod2ci <- predict(fitted_nb2, polynom_pr, simulate_pi=FALSE,uncertain=FALSE) %>%
  mutate(ntotal_studies=estimate)
data_nbmod2ci$GDP <- new_GDP

data1a <- data1a %>%
  mutate(GDP_jit = jitter(GDP,factor=10))

get_model(fitted_nb2)
```

```
##
## Call:  MASS::glm.nb(formula = ntotal_studies ~ GDP_orth + GDP_sq_orth,
##     data = data, init.theta = 3.106413934, link = log)
##
## Coefficients:
## (Intercept)     GDP_orth  GDP_sq_orth
##     1.19182     -0.07139     -4.80917
##
## Degrees of Freedom: 36 Total (i.e. Null);  34 Residual
## Null Deviance:       65.85
## Residual Deviance: 43.29     AIC: 176.1
```

```r
summary(nb_mod2)
```

```
##             Length Class  Mode
## model_class 1      -none- character
## fit         1      -none- function
```

```r
#use nb_glm to get pseudo R2

summary(nb_mod2c <- glm.nb(formula=ntotal_studies ~ GDP_orth + GDP_sq_orth,data=data1a))
```

```
##
## Call:
## glm.nb(formula = ntotal_studies ~ GDP_orth + GDP_sq_orth, data = data1a,
##     init.theta = 3.106413934, link = log)
##
## Deviance Residuals:
##      Min        1Q    Median        3Q       Max
## -2.43410  -0.80926  -0.08203   0.39578   1.71779
##
## Coefficients:
##             Estimate Std. Error z value Pr(>|z|)
## (Intercept)  1.19182    0.14188   8.400  < 2e-16 ***
## GDP_orth    -0.07139    1.12604  -0.063 0.949445
## GDP_sq_orth -4.80917    1.26813  -3.792 0.000149 ***
## ---
## Signif. codes:  0 '***' 0.001 '**' 0.01 '*' 0.05 '.' 0.1 ' ' 1
##
## (Dispersion parameter for Negative Binomial(3.1064) family taken to be 1)
##
```

```
##     Null deviance: 65.847  on 36  degrees of freedom
## Residual deviance: 43.291  on 34  degrees of freedom
## AIC: 176.14
##
## Number of Fisher Scoring iterations: 1
##
##
##               Theta:  3.11
##           Std. Err.:  1.51
##
##  2 x log-likelihood:  -168.144
```

```
pseudo_Rsq_a <- PseudoR2(nb_mod2c,which="all")
pseudo_Rsq_a
```

```
##        McFadden     McFaddenAdj        CoxSnell      Nagelkerke   AldrichNelson
##      0.09452115      0.06221028      0.37773166      0.38024599      0.32175058
## VeallZimmermann           Efron McKelveyZavoina            Tjur             AIC
##      0.38585949      0.35184420              NA              NA    176.14384322
##             BIC          logLik         logLik0              G2
##    182.58751487    -84.07192161    -92.84802309     17.55220297
```

```
data1a1 <- data1a %>%
   filter(NAME!="Luxembourg")
summary(nb_mod2d <- glm.nb(formula=ntotal_studies ~ GDP_orth + GDP_sq_orth,data=data1a1))
```

```
##
## Call:
## glm.nb(formula = ntotal_studies ~ GDP_orth + GDP_sq_orth, data = data1a1,
##     init.theta = 3.107271634, link = log)
##
## Deviance Residuals:
##      Min        1Q    Median        3Q       Max
## -2.42459  -0.87437  -0.04346   0.47869   1.71361
##
## Coefficients:
##             Estimate Std. Error z value Pr(>|z|)
## (Intercept)  1.20396    0.14461   8.326  < 2e-16 ***
## GDP_orth     0.09358    1.19963   0.078 0.937820
## GDP_sq_orth -4.58991    1.38868  -3.305 0.000949 ***
## ---
## Signif. codes:  0 '***' 0.001 '**' 0.01 '*' 0.05 '.' 0.1 ' ' 1
##
## (Dispersion parameter for Negative Binomial(3.1073) family taken to be 1)
##
##     Null deviance: 60.595  on 35  degrees of freedom
## Residual deviance: 43.008  on 33  degrees of freedom
## AIC: 175.86
##
## Number of Fisher Scoring iterations: 1
##
##
##               Theta:  3.11
##           Std. Err.:  1.51
##
```

```
##  2 x log-likelihood:  -167.856
```

```
pseudo_Rsq_b <- PseudoR2(nb_mod2d,which="all")
pseudo_Rsq_b       #squared term P= 0.0009
```

```
##        McFadden     McFaddenAdj        CoxSnell      Nagelkerke   AldrichNelson
##      0.07675402      0.04375273      0.32133758      0.32340979      0.27934755
## VeallZimmermann           Efron McKelveyZavoina            Tjur             AIC
##      0.33466052      0.32557774              NA              NA    175.85637227
##             BIC          logLik         logLik0              G2
##    182.19044803    -83.92818614    -90.90555230     13.95473233
```

```
data1a2 <- data1a %>%
   filter(NAME!="Switzerland")
summary(nb_mod2e <- glm.nb(formula=ntotal_studies ~ GDP_orth + GDP_sq_orth,data=data1a2))
```

```
##
## Call:
## glm.nb(formula = ntotal_studies ~ GDP_orth + GDP_sq_orth, data = data1a2,
##     init.theta = 3.462178852, link = log)
##
## Deviance Residuals:
##      Min        1Q    Median        3Q       Max
## -2.56062  -0.91411   0.00269   0.42877   1.75204
##
## Coefficients:
##             Estimate Std. Error z value Pr(>|z|)
## (Intercept)    1.033      0.174   5.936 2.92e-09 ***
## GDP_orth      -2.331      1.810  -1.287 0.197945
## GDP_sq_orth   -6.891      1.893  -3.640 0.000272 ***
## ---
## Signif. codes:  0 '***' 0.001 '**' 0.01 '*' 0.05 '.' 0.1 ' ' 1
##
## (Dispersion parameter for Negative Binomial(3.4622) family taken to be 1)
##
##     Null deviance: 68.878  on 35  degrees of freedom
## Residual deviance: 41.601  on 33  degrees of freedom
## AIC: 168.17
##
## Number of Fisher Scoring iterations: 1
##
##
##               Theta:  3.46
##           Std. Err.:  1.80
##
##  2 x log-likelihood:  -160.172
```

```
pseudo_Rsq_c <- PseudoR2(nb_mod2e,which="all")
pseudo_Rsq_c       #squared term P=0.00272
```

```
##        McFadden     McFaddenAdj        CoxSnell      Nagelkerke   AldrichNelson
##      0.11263847      0.07939822      0.43150874      0.43439498      0.36092817
## VeallZimmermann           Efron McKelveyZavoina            Tjur             AIC
##      0.43291225      0.39900896              NA              NA    168.17231437
##             BIC          logLik         logLik0              G2
##    174.50639013    -80.08615719    -90.25200531     20.33169625
```

```r
data1a3 <- data1a %>%
  filter(NAME!="Ireland")
summary(nb_mod2f <- glm.nb(formula=ntotal_studies ~ GDP_orth + GDP_sq_orth,data=data1a3))
```

```
##
## Call:
## glm.nb(formula = ntotal_studies ~ GDP_orth + GDP_sq_orth, data = data1a3,
##     init.theta = 3.106858644, link = log)
##
## Deviance Residuals:
##      Min        1Q    Median        3Q       Max
## -2.42308  -0.85352  -0.05971   0.47153   1.72444
##
## Coefficients:
##             Estimate Std. Error z value Pr(>|z|)
## (Intercept)   1.2211     0.1451   8.415  < 2e-16 ***
## GDP_orth      0.3461     1.2187   0.284 0.776412
## GDP_sq_orth  -4.4721     1.2881  -3.472 0.000517 ***
## ---
## Signif. codes:  0 '***' 0.001 '**' 0.01 '*' 0.05 '.' 0.1 ' ' 1
##
## (Dispersion parameter for Negative Binomial(3.1069) family taken to be 1)
##
##     Null deviance: 64.039  on 35  degrees of freedom
## Residual deviance: 42.555  on 33  degrees of freedom
## AIC: 173.11
##
## Number of Fisher Scoring iterations: 1
##
##
##               Theta:  3.11
##           Std. Err.:  1.52
##
##  2 x log-likelihood:  -165.114
```

```r
pseudo_Rsq_d <- PseudoR2(nb_mod2f,which="all")
pseudo_Rsq_d       #squared term P=0.000517
```

```
##        McFadden     McFaddenAdj        CoxSnell      Nagelkerke   AldrichNelson
##      0.09252332      0.05954687      0.37351045      0.37590990      0.31862619
## VeallZimmermann           Efron McKelveyZavoina            Tjur             AIC
##      0.38166916      0.33501796              NA              NA    173.11357893
##             BIC          logLik         logLik0              G2
##    179.44765468    -82.55678947    -90.97400691     16.83443488
```

```r
#data1b$fitted_11b <- mod11b$fitted.values
#new_data <- data.frame(GDP = seq(from=min(data1b$GDP),to=max(data1b$GDP),by=100))
#new_data$GDP_sq <- new_data$GDP^2
#conf_11b <- as.data.frame(stats::predict(mod11b, new_data, interval='confidence'))
#conf_11b$GDP <- new_data$GDP
#data1b$lwr_11b <- conf_11b$lwr;data1b$upr_11b <- conf_11b$upr

data1a$GDP_jit[which(data1a$POSTAL=='FIN')] <- data1a$GDP[which(data1a$POSTAL=='FIN')] + 1000
data1a$GDP_jit[which(data1a$POSTAL=='A')] <- data1a$GDP[which(data1a$POSTAL=='A')] - 1000
data1a$GDP_jit[which(data1a$POSTAL=='M')] <- data1a$GDP[which(data1a$POSTAL=='M')] + 1000
```

```
data1a$GDP_jit[which(data1a$POSTAL=='CY')] <- data1a$GDP[which(data1a$POSTAL=='CY')] - 1000


set.seed(12345)

plot2a <- ggplot(data1a,aes(x=GDP_jit, y=ntotal_studies)) +
  geom_point(aes(color=Area),size=rel(1)) +   #position=position_dodge(width = 0.2),

  scale_color_gradient(low='blue',high='red', labels = scales::comma)+
  scale_x_continuous(limits=c(0,121000),
        breaks = c(0,10000,50000,90000,120000),labels=scales::dollar_format())+
  scale_y_continuous(limits=c(-2,12.5),breaks=c(0,3,6,9,12))+
  geom_ribbon(aes(x=GDP,ymin=lower_ci,ymax=upper_ci),data_nbmod2ci,color='gray',alpha=0.1, linetype=0,
  geom_line(aes(GDP,ntotal_studies),data_nbmod2ci)+
  theme(
    panel.grid.minor = element_blank(),
    panel.grid.major = element_blank(),
    panel.background = element_rect(fill= alpha('medium sea green', alpha=0.2)),
    legend.text = element_text(size = rel(1.0)),
    axis.line= element_line(size = 0.75, color = "black"),
    axis.text.y= element_text(size=rel(1.6)),
    axis.text.x= element_text(size=rel(1.1)),
    axis.title = element_text(size=rel(1.6)),
    plot.title = element_text(hjust = 0.5, size=rel(1.6)))+
  labs(x="GDP in 2020 Dollars",y= "Realized Monitoring Capacity") +
  #labs(title = "Country per capita GDP vs. Number Monitoring Projects") +
  labs(color="Country Area \n (km square)") +
  geom_label_repel(aes(label=POSTAL),
    max.time=5,
    max.iter = 10000000,
    size = rel(1.6),
    label.padding = 0.1,
    box.padding   = 0.1,
    point.padding = 0.20,
    label.size=NA,
    max.overlaps = 25,
    min.segment.length = 0,
    segment.angle = 90,
    fill=NA,
    segment.color = 'grey50',
    segment.size = 0.3)
equation = expression(paste("Projects = exp(1.18869 + poly(GDP)1\n       +poly(GDP)2)"))
plot2a <- plot2a +
   annotate(geom="text", x=90000, y=11, label=equation,size=5/.pt)

plot2a <- plot2a +
   annotate(geom="text", x=67900, y=10.5,
            label= 'paste("R"^"2"," = 0.39")', parse=TRUE, size=5/.pt)

plot2a
```

```
## Warning in is.na(x): is.na() applied to non-(list or vector) of type
## 'expression'
```

```
## Warning in grid.Call.graphics(C_text, as.graphicsAnnot(x$label), x$x, x$y, :
## font metrics unknown for character 0xa

## Warning in grid.Call.graphics(C_text, as.graphicsAnnot(x$label), x$x, x$y, :
## font metrics unknown for character 0xa
```

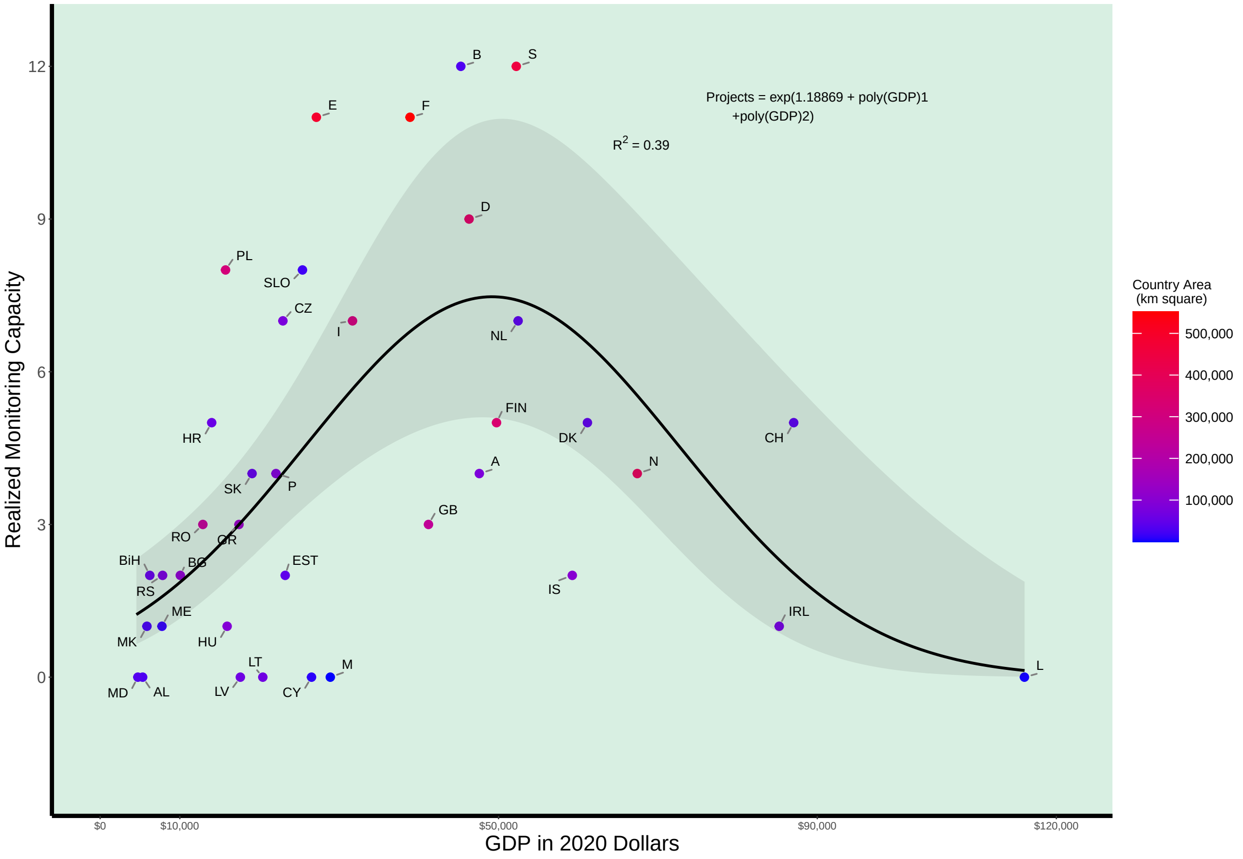


```
#ggsave("PGD_projects_vs_GDP_wo_Turkey_nbreg.pdf", plot=plot2a,device = "pdf", width = 8,height=6, unit
ggsave("PGD_projects_vs_GDP_wo_Türkiye_nbreg.pdf",plot = plot2a, height = 88*aspect,width = 88,units =
```

```
## Warning in is.na(x): is.na() applied to non-(list or vector) of type
## 'expression'

## Warning in is.na(x): font metrics unknown for character 0xa

## Warning in is.na(x): font metrics unknown for character 0xa
```

```
ggsave("PGD_projects_vs_GDP_wo_Türkiye_nbreg.svg",plot = plot2a, height = 88*aspect,width = 88,units =
```

```
## Warning in is.na(x): is.na() applied to non-(list or vector) of type
## 'expression'
```

```
data1a$mod2a_resid <- residuals(mod2a)
data1a$mod_p_resid <- residuals(mod_p)


save(data1a, file="data1a_country_monitoring_regression_Rmd.RData")
```

```
double <- plot_grid(plot1,plot2a,labels = c("a","b"), hjust=-0.5, vjust = 1.2,label_fontface = "plain")
```

```
## Warning in is.na(x): is.na() applied to non-(list or vector) of type
## 'expression'
```

```
ggsave("PGD_projects_vs_GDP_Area_wo_Türkiye_nbreg.pdf",plot = double, height = 88*aspect,width = 2*88,ur
```

```
## Warning in grid.Call.graphics(C_text, as.graphicsAnnot(x$label), x$x, x$y, :
## font metrics unknown for character 0xa
```

```
## Warning in grid.Call.graphics(C_text, as.graphicsAnnot(x$label), x$x, x$y, :
## font metrics unknown for character 0xa
```

```
ggsave("PGD_projects_vs_GDP_Area_wo_Türkiye_nbreg.svg",plot = double, height = 88*aspect,width = 2*88,ur
```

### Spatial Regressions

Peter B. Pearman

5/25/2022

```
library(tidyverse)
```

```
## -- Attaching packages --------------------------------------- tidyverse 1.3.1 --

## v ggplot2 3.3.5     v purrr   0.3.4
## v tibble  3.1.5     v dplyr   1.0.7
## v tidyr   1.1.4     v stringr 1.4.0
## v readr   2.0.2     v forcats 0.5.1

## -- Conflicts ------------------------------------------ tidyverse_conflicts() --
## x dplyr::filter() masks stats::filter()
## x dplyr::lag()    masks stats::lag()
```

```
library(magrittr)
```

```
##
## Attaching package: 'magrittr'

## The following object is masked from 'package:purrr':
##
##     set_names

## The following object is masked from 'package:tidyr':
##
##     extract
```

```
library(car)
```

```
## Loading required package: carData

##
## Attaching package: 'car'

## The following object is masked from 'package:dplyr':
##
##     recode

## The following object is masked from 'package:purrr':
##
##     some
```

```
library(ggfortify)
library(ggrepel)
library(scales)
```

```
##
## Attaching package: 'scales'
```

```
## The following object is masked from 'package:purrr':
##
##     discard

## The following object is masked from 'package:readr':
##
##     col_factor
```

```
library(shapefiles)
```

```
## Loading required package: foreign

##
## Attaching package: 'shapefiles'

## The following objects are masked from 'package:foreign':
##
##     read.dbf, write.dbf
```

```
library(maptools)
```

```
## Loading required package: sp

## Checking rgeos availability: TRUE
## Please note that 'maptools' will be retired by the end of 2023,
## plan transition at your earliest convenience;
## some functionality will be moved to 'sp'.
```

```
library(foreign)
library(sp)
library(rgdal)
```

```
## Please note that rgdal will be retired by the end of 2023,
## plan transition to sf/stars/terra functions using GDAL and PROJ
## at your earliest convenience.
##
## rgdal: version: 1.5-27, (SVN revision 1148)
## Geospatial Data Abstraction Library extensions to R successfully loaded
## Loaded GDAL runtime: GDAL 3.2.1, released 2020/12/29
## Path to GDAL shared files: /Library/Frameworks/R.framework/Versions/4.1/Resources/library/rgdal/gdal
## GDAL binary built with GEOS: TRUE
## Loaded PROJ runtime: Rel. 7.2.1, January 1st, 2021, [PJ_VERSION: 721]
## Path to PROJ shared files: /Library/Frameworks/R.framework/Versions/4.1/Resources/library/rgdal/proj
## PROJ CDN enabled: FALSE
## Linking to sp version:1.4-5
## To mute warnings of possible GDAL/OSR exportToProj4() degradation,
## use options("rgdal_show_exportToProj4_warnings"="none") before loading sp or rgdal.
## Overwritten PROJ_LIB was /Library/Frameworks/R.framework/Versions/4.1/Resources/library/rgdal/proj
```

```
library(raster)
```

```
##
## Attaching package: 'raster'

## The following object is masked from 'package:dplyr':
##
##     select
```

```
library(sf)
```

```
## Linking to GEOS 3.8.1, GDAL 3.2.1, PROJ 7.2.1
```

```
library(mapview)
library(tmap)
```

```
## Registered S3 methods overwritten by 'stars':
##   method             from
##   st_bbox.SpatRaster sf
##   st_crs.SpatRaster  sf
```

```
library(classInt)
library(spdep)
```

```
## Loading required package: spData

## To access larger datasets in this package, install the spDataLarge
## package with: `install.packages('spDataLarge',
## repos='https://nowosad.github.io/drat/', type='source')`
```

```
library(ade4)
```

```
##
## Attaching package: 'ade4'

## The following object is masked from 'package:spdep':
##
##     mstree
```

```
library(adespatial)
```

```
## Registered S3 methods overwritten by 'adegraphics':
##   method         from
##   biplot.dudi    ade4
##   kplot.foucart  ade4
##   kplot.mcoa     ade4
##   kplot.mfa      ade4
##   kplot.pta      ade4
##   kplot.sepan    ade4
##   kplot.statis   ade4
##   scatter.coa    ade4
##   scatter.dudi   ade4
##   scatter.nipals ade4
##   scatter.pco    ade4
##   score.acm      ade4
##   score.mix      ade4
##   score.pca      ade4
##   screeplot.dudi ade4

## Registered S3 method overwritten by 'ape':
##   method   from
##   plot.mst spdep

## Registered S3 methods overwritten by 'adespatial':
##   method             from
##   plot.multispati    adegraphics
##   print.multispati   ade4
##   summary.multispati ade4

##
```

```
## Attaching package: 'adespatial'

## The following object is masked from 'package:ade4':
##
##     multispati
```

```
library(adegraphics)
```

```
##
## Attaching package: 'adegraphics'

## The following objects are masked from 'package:ade4':
##
##     kplotsepan.coa, s.arrow, s.class, s.corcircle, s.distri, s.image,
##     s.label, s.logo, s.match, s.traject, s.value, table.value,
##     triangle.class

## The following object is masked from 'package:raster':
##
##     zoom
```

```
library(deldir)
```

```
## deldir 1.0-6      Nickname: "Mendacious Cosmonaut"

##
##      The syntax of deldir() has had an important change.
##      The arguments have been re-ordered (the first three
##      are now "x, y, z") and some arguments have been
##      eliminated.  The handling of the z ("tags")
##      argument has been improved.
##
##      The "dummy points" facility has been removed.
##      This facility was a historical artefact, was really
##      of no use to anyone, and had hung around much too
##      long.  Since there are no longer any "dummy points",
##      the structure of the value returned by deldir() has
##      changed slightly.  The arguments of plot.deldir()
##      have been adjusted accordingly; e.g. the character
##      string "wpoints" ("which points") has been
##      replaced by the logical scalar "showpoints".
##      The user should consult the help files.
```

```
library(rgeos)
```

```
## rgeos version: 0.5-8, (SVN revision 679)
##  GEOS runtime version: 3.8.1-CAPI-1.13.3
##  Please note that rgeos will be retired by the end of 2023,
## plan transition to sf functions using GEOS at your earliest convenience.
##  Linking to sp version: 1.4-5
##  Polygon checking: TRUE
```

```
library(ncf)
```

```
##
## Attaching package: 'ncf'

## The following object is masked from 'package:tidyr':
##
```

```
##     gather
```

```
library(spatialreg)
```

```
## Loading required package: Matrix

##
## Attaching package: 'Matrix'

## The following objects are masked from 'package:tidyr':
##
##     expand, pack, unpack

##
## Attaching package: 'spatialreg'

## The following objects are masked from 'package:spdep':
##
##     aple, aple.mc, aple.plot, as_dgRMatrix_listw, as_dsCMatrix_I,
##     as_dsCMatrix_IrW, as_dsTMatrix_listw, as.spam.listw, can.be.simmed,
##     cheb_setup, create_WX, do_ldet, eigen_pre_setup, eigen_setup,
##     eigenw, errorsarlm, get.ClusterOption, get.coresOption,
##     get.mcOption, get.VerboseOption, get.ZeroPolicyOption,
##     GMargminImage, GMerrorsar, griffith_sone, gstsls, Hausman.test,
##     impacts, intImpacts, invIrM, invIrW, Jacobian_W, jacobianSetup,
##     l_max, lagmess, lagsarlm, lextrB, lextrS, lextrW, lmSLX, localAple,
##     LU_prepermutate_setup, LU_setup, Matrix_J_setup, Matrix_setup,
##     mcdet_setup, MCMCsamp, ME, mom_calc, mom_calc_int2, moments_setup,
##     powerWeights, sacsarlm, SE_classic_setup, SE_interp_setup,
##     SE_whichMin_setup, set.ClusterOption, set.coresOption,
##     set.mcOption, set.VerboseOption, set.ZeroPolicyOption,
##     similar.listw, spam_setup, spam_update_setup, SpatialFiltering,
##     spautolm, spBreg_err, spBreg_lag, spBreg_sac, stsls,
##     subgraph_eigenw, trW
```

```
library(geosphere)
library(MASS)
```

```
##
## Attaching package: 'MASS'

## The following objects are masked from 'package:raster':
##
##     area, select

## The following object is masked from 'package:dplyr':
##
##     select
```

```
library(pscl)
```

```
## Classes and Methods for R developed in the
## Political Science Computational Laboratory
## Department of Political Science
## Stanford University
## Simon Jackman
## hurdle and zeroinfl functions by Achim Zeileis
```

```
library(ade4)
library(adespatial)
library(DescTools)
```

```
## Registered S3 method overwritten by 'DescTools':
##   method         from
##   reorder.factor gdata

##
## Attaching package: 'DescTools'

## The following object is masked from 'package:car':
##
##     Recode
```

```
rm(list=ls())
# GDP from https://data.worldbank.org/indicator/NY.GDP.PCAP.CD, accessed 24.05.2022
# Country area in Europe from https://en.wikipedia.org/wiki/List_of_European_countries_by_area
# accessed 24.05.2022, which includes only the area of countries in continental Europe.


data.a <- read_csv('country_area_GDP.csv')
```

```
## Rows: 39 Columns: 11

## -- Column specification --------------------------------------------------------
## Delimiter: ","
## chr (6): POSTAL, NAME, NAME_EN, code, name, region
## dbl (5): ntotal_studies, area_kmsq, per_capita_GDP_CIA, year, Per_cap_GDP_Wo...

##
## i Use `spec()` to retrieve the full column specification for this data.
## i Specify the column types or set `show_col_types = FALSE` to quiet this message.
```

```
data.a <- as_tibble(data.a) %>%
  dplyr::select(POSTAL,area_kmsq,per_capita_GDP_CIA,year,Per_cap_GDP_WoBa_2020dollars)
data.b <- as_tibble(read_csv('./country_data3_w_U.csv')) %>%
  arrange(NAME)
```

```
## Rows: 39 Columns: 21

## -- Column specification --------------------------------------------------------
## Delimiter: ","
## chr  (6): POSTAL, NAME, NAME_EN, code, name, region
## dbl (15): ncarn, nbear, nwolf, nlynx, nom, nbird, ninsect, nfish, nmarine, n...

##
## i Use `spec()` to retrieve the full column specification for this data.
## i Specify the column types or set `show_col_types = FALSE` to quiet this message.
```

```
data1 <- left_join(data.b,data.a,by='POSTAL')

data1 %<>% filter(NAME != "Liechtenstein") %>%
  mutate(GDP=Per_cap_GDP_WoBa_2020dollars) %>%
  mutate(Area=as.numeric(area_kmsq), Area_sq = Area*Area,
         lnGDP = log10(GDP), GDP_sq = GDP^2,
         lnArea=log10(area_kmsq),
         lnAreasq=lnArea*lnArea,lnStudies=log10(ntotal_studies+1)) %>%
```

```
  dplyr::select(-one_of('Per_cap_GDP_WoBa_2020dollars')) %>%
  arrange(NAME)
```

Remove Turkey and Lichtenstein

```
earth <- st_read("shapefiles/ne_50m_admin_0_countries_lakes.shp",crs=4326)  #4326 102013 albers_ea: 102
```

```
## Reading layer `ne_50m_admin_0_countries_lakes' from data source
##   `/Users/bgppermp/Google_Drive_EHU/UPV_Research/G-BIKE/R_publication_maps_NEE/shapefiles/ne_50m_adm
##   using driver `ESRI Shapefile'
## Simple feature collection with 241 features and 94 fields
## Geometry type: MULTIPOLYGON
## Dimension:     XY
## Bounding box:  xmin: -180 ymin: -89.99893 xmax: 180 ymax: 83.59961
## Geodetic CRS:  WGS 84
```

```
earth$NAME <- as.character(earth$NAME)
earth$NAME[which(earth$NAME=="Czechia")] <- "Czech Republic"
earth$NAME[which(earth$NAME=="Macedonia")] <- "North Macedonia"

earth_test<-st_intersection(earth, st_set_crs(st_as_sf(as(raster::extent(-25,57,29.1,73), "SpatialPolyg
```

```
## Warning: attribute variables are assumed to be spatially constant throughout all
## geometries
```

```
earth_test1 <- earth_test %>% dplyr::filter(NAME=="Russia") %>%
  st_cast(.,"POLYGON")
```

```
## Warning in st_cast.sf(., "POLYGON"): repeating attributes for all sub-geometries
## for which they may not be constant
```

```
kalingrad <- earth_test1[9,"NAME"]

european_countries <- read_csv("european_countries.csv") %>%
  filter(!(name %in% c("Russia","Ukraine","Israel","Liechtenstein")))
```

```
## Rows: 39 Columns: 4
```

```
## -- Column specification --------------------------------------------------------
## Delimiter: ","
## chr (4): code, POSTAL, name, region
```

```
##
## i Use `spec()` to retrieve the full column specification for this data.
## i Specify the column types or set `show_col_types = FALSE` to quiet this message.
```

```
earth_eu <- earth %>%
  #filter(CONTINENT=="Europe") %>%
  filter(NAME %in% european_countries$name) %>%
  filter(!(NAME %in% c("Russia","Ukraine","Israel","Liechtenstein"))) %>%
  dplyr::select(POSTAL,NAME, NAME_EN)

earth_eu$POSTAL <- as.character(earth_eu$POSTAL)
earth_eu$POSTAL[base::which(earth_eu$NAME=="Israel")] <- "IL"
earth_eu$NAME[base::which(earth_eu$NAME=="Macedonia")] <- "North Macedonia"


# Calculate country area in Europe just below
```

```r
earth_euprog <- st_transform(earth_eu,crs="+proj=lcc +lat_1=43 +lat_2=62 +lat_0=30 +lon_0=10 +x_0=0 +y_
earth_eu2<-st_intersection(earth_eu, st_set_crs(st_as_sf(as(raster::extent(-25,57,28,73), "SpatialPolyg
  mutate(new_Area=units::set_units(st_area(earth_eu),"km^2"))
```

```
## Warning: attribute variables are assumed to be spatially constant throughout all
## geometries
```

```r
earth_eu3 <- st_transform(earth_eu2,crs="+proj=lcc +lat_1=43 +lat_2=62 +lat_0=30 +lon_0=10 +x_0=0 +y_0=
kalingrad <- st_transform(kalingrad,crs="+proj=lcc +lat_1=43 +lat_2=62 +lat_0=30 +lon_0=10 +x_0=0 +y_0=

#plot(earth_eu2)
```

```r
earth_eu2a <- st_transform(earth_eu2,crs = 4326)
earth_eu2b <- earth_eu2a %>%
  filter(!NAME=="Turkey") %>%
  sf::as_Spatial(.)
```

```r
nb_eu <-tri2nb(coordinates(earth_eu2b))
plot(earth_eu2b,border = "gray")
plot(nb_eu, coordinates(earth_eu2b), add = TRUE, pch = 20, col = "red")
```

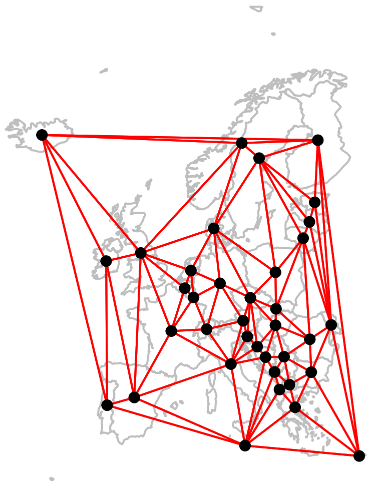

```r
#s.Spatial(earth_eu2b,nb = nb_eu, plabel.cex = 0, pnb.edge.col = 'red')
```

```r
truecentroids <- gCentroid(earth_eu2b, byid = TRUE)
plot(earth_eu2b)
points(coordinates(earth_eu2b),pch=".")
points(truecentroids,pch=2)
```

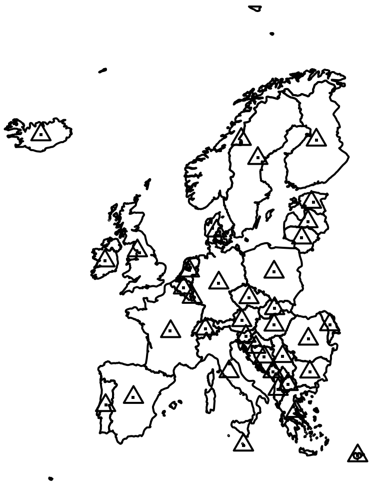
You can use listw.explore to get the code to make nd and listw objects all in one shot with adespatial::listw.explore()

```
xyvals <- coordinates(truecentroids)
x <- truecentroids$x
y <- truecentroids$y
POSTAL <- earth_eu2b$POSTAL
POSTAL_centroids <- tibble(POSTAL,x,y)

nbnear <- dnearneigh(xyvals,0,1500, longlat = TRUE)
distnbnear <- nbdists(nbnear, as.matrix(xyvals))
fdist1 <- lapply(distnbnear, function(x) 1 - x/max(dist(as.matrix(xyvals))))
lwnear <- nb2listw(nbnear, style = 'W', glist = fdist1, zero.policy = TRUE)


nbtri <- tri2nb(xyvals)
distnbtri <- nbdists(nbtri, as.matrix(xyvals))
fdist2 <- lapply(distnbtri, function(x) 1 - x/max(dist(as.matrix(xyvals))))
lwtri <- nb2listw(nbtri, style = 'W', glist = fdist2, zero.policy = TRUE)

nbgab <-graph2nb(gabrielneigh(xyvals), sym = TRUE)
distnbgab <- nbdists(nbgab, as.matrix(xyvals))
fdist3 <- lapply(distnbgab, function(x) 1 - x/max(dist(as.matrix(xyvals))))
lwgab <- nb2listw(nbgab, style = 'W', glist = fdist3, zero.policy = TRUE)

nbrel <-graph2nb(relativeneigh(xyvals), sym = TRUE)
distnbrel <- nbdists(nbrel, as.matrix(xyvals))
fdist4 <- lapply(distnbrel, function(x) 1 - x/max(dist(as.matrix(xyvals))))
lwrel <- nb2listw(nbrel, style = 'W', glist = fdist4, zero.policy = TRUE)

g1 <- s.label(xyvals, nb = nbnear, pnb.edge.col = "red", main = "dnearneigh if 0 < 1500", labels=NULL,pl
g2 <- s.label(xyvals, nb = nbtri, pnb.edge.col = "red", main = "Delaunay", labels=NULL,plot = FALSE)
g3 <- s.label(xyvals, nb = nbgab, pnb.edge.col = "red", main = "Gabriel", labels=NULL,plot = FALSE)
g4 <- s.label(xyvals, nb = nbrel, pnb.edge.col = "red", main = "Relative", labels=NULL,plot = FALSE)


cbindADEg(g1,g2,g3,g4,plot=TRUE)
```

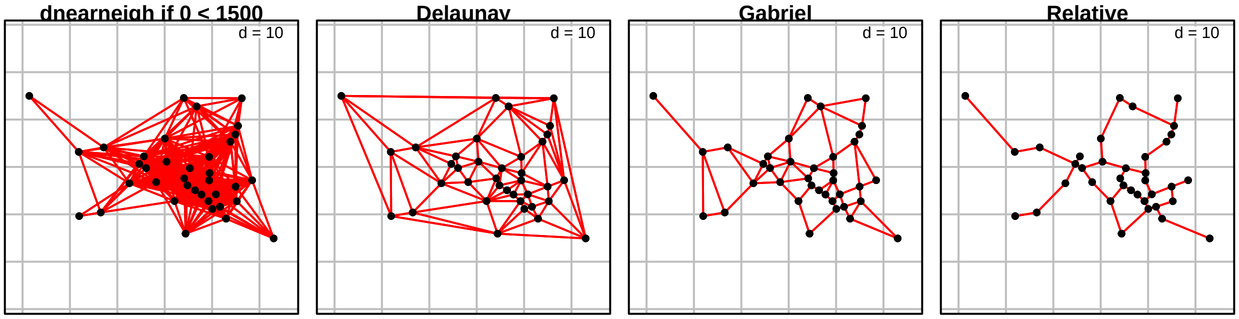


```
# see trellis.par.get() to set parameters for plotting as a trellis plot.
```

###Data preparation

```
data.a <- read_csv('country_area_GDP.csv')
```

```
## Rows: 39 Columns: 11

## -- Column specification --------------------------------------------------------
## Delimiter: ","
## chr (6): POSTAL, NAME, NAME_EN, code, name, region
## dbl (5): ntotal_studies, area_kmsq, per_capita_GDP_CIA, year, Per_cap_GDP_Wo...

##
## i Use `spec()` to retrieve the full column specification for this data.
## i Specify the column types or set `show_col_types = FALSE` to quiet this message.
```

```
data.a <- as_tibble(data.a) %>%
  dplyr::select(POSTAL,area_kmsq,per_capita_GDP_CIA,year,Per_cap_GDP_WoBa_2020dollars)
#data.b <- as_tibble(read_csv('./country_data3_w_U.csv')) %>%
#  arrange(NAME)
data.b <- as_tibble(read_csv('./country_data3_PGD_only.csv')) %>%
  arrange(NAME)
```

```
## Rows: 38 Columns: 23

## -- Column specification --------------------------------------------------------
## Delimiter: ","
## chr  (6): POSTAL, NAME, NAME_EN, code, name, region
## dbl (17): new_Area, ncarn, nbear, nwolf, nlynx, nom, nbird, ninsect, nfish, ...

##
## i Use `spec()` to retrieve the full column specification for this data.
## i Specify the column types or set `show_col_types = FALSE` to quiet this message.
```

```
data1 <- left_join(data.b,data.a,by='POSTAL')

data1 %<>% filter(NAME != "Liechtenstein") %>%
  mutate(GDP=Per_cap_GDP_WoBa_2020dollars) %>%
  mutate(Area=as.numeric(area_kmsq), Area_sq = Area*Area,
         area3 = area_kmsq/1000,
         lnGDP = log10(GDP), GDP_sq = GDP^2,
         lnArea=log10(area_kmsq),
         lnAreasq=lnArea*lnArea,
         lnStudies=log10(ntotal_studies+1)) %>%
  dplyr::select(-one_of('Per_cap_GDP_WoBa_2020dollars')) %>%
  arrange(NAME)
```

**Remove Turkey as an outlier**

**Do regressions as in country_monitoring_regression_PGD_only.Rmd**

```
# Here we drop Turkey because it is clearly an outlier in this analysis
# Then re-run quadratic regression with Area and Area_sq
data1b <- data1 %>%
  filter(POSTAL != "TR")
data1b <- left_join(data1b,POSTAL_centroids,by="POSTAL",copy=FALSE)

data1b <- left_join(data1b,earth_eu2a[c("POSTAL","new_Area")],by="POSTAL",copy=FALSE) %>%
   rename(new_Area = new_Area.y)
```

```
data1b$Area3 <- data1b$new_Area/1000
summary(mod2a <- glm.nb(formula=ntotal_studies ~ new_Area+poly(GDP,2),data=data1b))
```

```
##
## Call:
## glm.nb(formula = ntotal_studies ~ new_Area + poly(GDP, 2), data = data1b,
##     init.theta = 4.641574631, link = log)
##
## Deviance Residuals:
##      Min        1Q    Median        3Q       Max
## -2.27532  -0.93966   0.01741   0.40036   1.69514
##
## Coefficients:
##                 Estimate Std. Error z value Pr(>|z|)
## (Intercept)    9.318e-01  1.685e-01   5.529 3.21e-08 ***
## new_Area       1.689e-06  7.134e-07   2.367  0.01791 *
## poly(GDP, 2)1  1.321e-02  1.014e+00   0.013  0.98961
## poly(GDP, 2)2 -3.684e+00  1.208e+00  -3.050  0.00229 **
## ---
## Signif. codes:  0 '***' 0.001 '**' 0.01 '*' 0.05 '.' 0.1 ' ' 1
##
## (Dispersion parameter for Negative Binomial(4.6416) family taken to be 1)
##
##     Null deviance: 77.242  on 36  degrees of freedom
## Residual deviance: 44.830  on 33  degrees of freedom
## AIC: 172.86
##
## Number of Fisher Scoring iterations: 1
##
##
##               Theta:  4.64
##           Std. Err.:  2.84
##
##  2 x log-likelihood:  -162.86
```

```
set.seed(123456)
cor_mod2a <- correlog(data1b$x, data1b$y, residuals(mod2a), increment = 300, resamp = 100, latlon=TRUE,
```

```
## 10  of  100 20  of  100 30  of  100 40  of  100 50  of  100 60  of  100 70  of  100 80  of  100 90
```

```
plot(cor_mod2a$correlation[1:20],type="s")
```

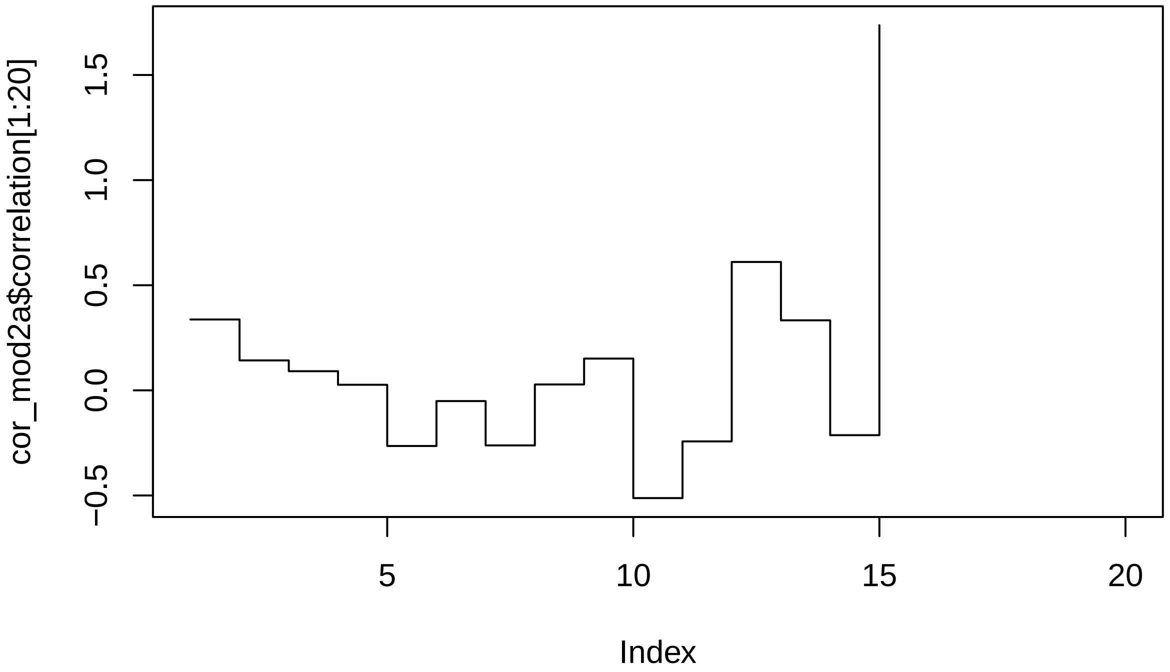


```
print("number of pairs in distance class");(cor_mod2a$nlok)
```

```
## [1] "number of pairs in distance class"
```

```
## NULL
```

```
print("mean distanc in class");(cor_mod2a$mean.of.class)
```

```
## [1] "mean distanc in class"
```

```
##         1         2         3         4         5         6         7         8
##  197.9171  441.9632  760.9924 1045.1570 1367.7932 1652.9023 1940.7598 2245.2692
##         9        10        11        12        13        14        16
## 2533.5246 2848.4211 3151.6718 3443.0474 3720.0842 3916.0736 4754.7293
```

```
print("p-values");(cor_mod2a$p)
```

```
## [1] "p-values"
```

```
##  [1] 0.04950495 0.04950495 0.11881188 0.25742574 0.05940594 0.40594059
##  [7] 0.02970297 0.31683168 0.15841584 0.04950495 0.17821782 0.00990099
## [13] 0.15841584 0.36633663 0.05940594
```

```
print("Moran's I values");(cor_mod2a$correlation)     #[which(cor_for_1$p < 0.05)])
```

```
## [1] "Moran's I values"
```

```
##           1           2           3           4           5           6
##  0.33708329  0.14234475  0.09103260  0.02643232 -0.26466599 -0.05111396
##           7           8           9          10          11          12
## -0.26187315  0.02792414  0.15099876 -0.51249437 -0.24281644  0.61069783
##          13          14          16
##  0.33301854 -0.21336295  1.73688422
```

listw objects from above lwnear lwtri lwgab lwrel

```
mem_near <- mem(lwnear)
mem_tri <- mem(lwtri)
```

```r
mem_gab <- mem(lwgab)
mem_rel <- mem(lwrel)


barplot(attr(mem_near,"values"),
        main = "Eigenvalues of the spatial weighting matrix", cex.main = 0.7)
```

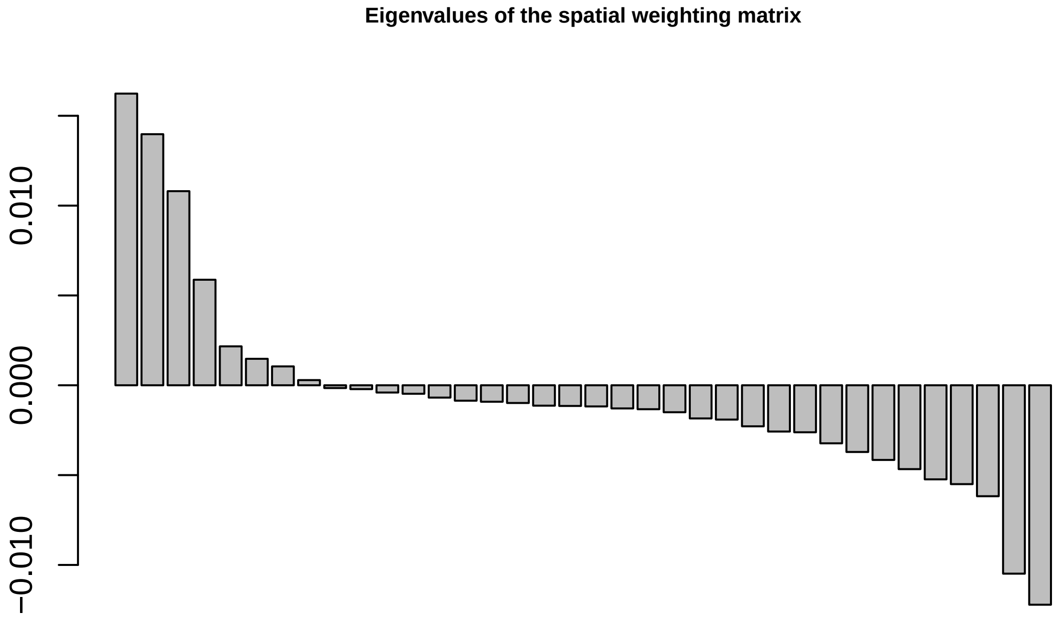


```r
data1c <- cbind(data1b,mem_near)
xnam <- paste0("MEM",1:8)
fmla <- as.formula(paste0("ntotal_studies ~ area3 + poly(GDP,2) +",paste(xnam[c(1,4,8)],collapse = "+")
#fmla <- as.formula("ntotal_studies ~ area3 + poly(GDP,2)")   # +",paste(xnam[1:2],collapse = "+")))
summary(new_mod2a <- glm.nb(formula=fmla, data=data1c))
```

```
##
## Call:
## glm.nb(formula = fmla, data = data1c, init.theta = 4.957131871,
##     link = log)
##
## Deviance Residuals:
##      Min        1Q    Median        3Q       Max
## -2.34334  -0.91110  -0.05347   0.58134   1.87015
##
## Coefficients:
##                   Estimate Std. Error z value Pr(>|z|)
## (Intercept)     0.9009749  0.1753047   5.139 2.76e-07 ***
## area3           0.0017771  0.0007834   2.269  0.02330 *
## poly(GDP, 2)1   0.1541058  1.0379394   0.148  0.88197
## poly(GDP, 2)2  -3.6703095  1.2361137  -2.969  0.00299 **
## MEM1           -0.1400252  0.1429385  -0.980  0.32727
## MEM4            0.0509286  0.1214082   0.419  0.67486
## MEM8           -0.0863815  0.1155839  -0.747  0.45485
## ---
## Signif. codes:  0 '***' 0.001 '**' 0.01 '*' 0.05 '.' 0.1 ' ' 1
##
```

```
## (Dispersion parameter for Negative Binomial(4.9571) family taken to be 1)
##
##     Null deviance: 79.087  on 36  degrees of freedom
## Residual deviance: 44.214  on 30  degrees of freedom
## AIC: 177.3
##
## Number of Fisher Scoring iterations: 1
##
##
##               Theta:  4.96
##           Std. Err.:  3.07
##
##  2 x log-likelihood:  -161.303
```

```
set.seed(123456)
cor_new_mod2a <- correlog(data1b$x, data1b$y, residuals(new_mod2a), increment = 300, resamp = 100, latl
```

```
## 10  of  100 20  of  100 30  of  100 40  of  100 50  of  100 60  of  100 70  of  100 80  of  100 90
```

```
plot(cor_new_mod2a$correlation[1:15],type="s")
```

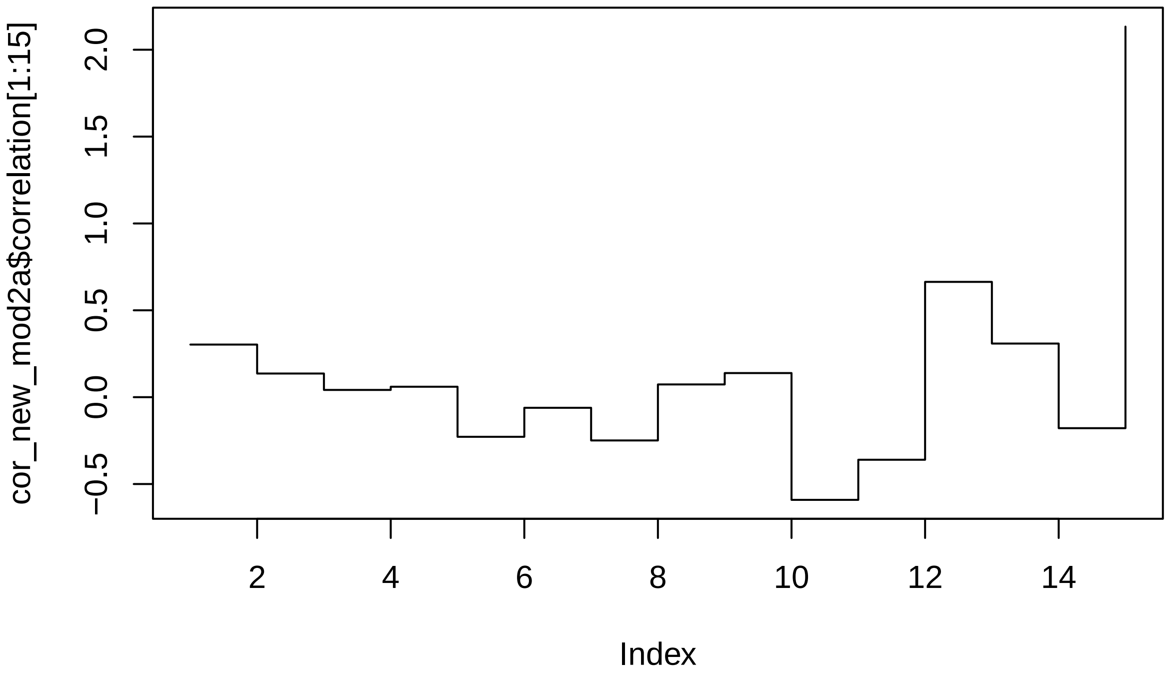


```
print("number of pairs in distance class");(cor_new_mod2a$nlok)
```

```
## [1] "number of pairs in distance class"
```

```
## NULL
```

```
print("mean distanc in class");(cor_new_mod2a$mean.of.class)
```

```
## [1] "mean distanc in class"
```

```
##         1         2         3         4         5         6         7         8
##  197.9171  441.9632  760.9924 1045.1570 1367.7932 1652.9023 1940.7598 2245.2692
##         9        10        11        12        13        14        16
## 2533.5246 2848.4211 3151.6718 3443.0474 3720.0842 3916.0736 4754.7293
```

```r
print("p-values");(cor_new_mod2a$p)
```

```
## [1] "p-values"

##  [1] 0.06930693 0.04950495 0.27722772 0.16831683 0.04950495 0.36633663
##  [7] 0.03960396 0.14851485 0.18811881 0.03960396 0.11881188 0.00990099
## [13] 0.17821782 0.41584158 0.02970297
```

```r
print("Moran's I values");(cor_new_mod2a$correlation)     #[which(cor_for_1$p < 0.05)])
```

```
## [1] "Moran's I values"

##           1           2           3           4           5           6
##  0.30280227  0.13614329  0.04166020  0.06009180 -0.22807966 -0.06110004
##           7           8           9          10          11          12
## -0.24908795  0.07338296  0.13882821 -0.59104639 -0.36026905  0.66374928
##          13          14          16
##  0.30859972 -0.17839661  2.13316065
```

```r
data1b$new_mod2a_resid <- residuals(new_mod2a, type= "deviance")

pseudo_Rsq <- PseudoR2(new_mod2a,which="all")
pseudo_Rsq
```

```
##        McFadden     McFaddenAdj        CoxSnell      Nagelkerke   AldrichNelson
##       0.1293083       0.0537381       0.4766181       0.4798287       0.3929990
## VeallZimmermann           Efron McKelveyZavoina            Tjur             AIC
##       0.4714893       0.4800748              NA              NA     177.3027490
##             BIC          logLik         logLik0              G2
##     190.1900923     -80.6513745     -92.6290855      23.9554220
```

```r
# use Veall & Zimmerman because it best approximates Rsq of OLS
#Veall, M.R., & Zimmermann, K.F. (1992) Evalutating Pseudo-R2's fpr binary probit models. Quality&Quant
# https://search.r-project.org/CRAN/refmans/DescTools/html/PseudoR2.html
```

```r
summary(mod <- glm(formula = fmla, family = poisson(link = "log"),data=data1c))
```

```
##
## Call:
## glm(formula = fmla, family = poisson(link = "log"), data = data1c)
##
## Deviance Residuals:
##     Min       1Q   Median       3Q      Max
## -2.7113  -1.2611  -0.1579   0.5931   2.9342
##
## Coefficients:
##                 Estimate Std. Error z value Pr(>|z|)
## (Intercept)    0.9183731  0.1354328   6.781 1.19e-11 ***
## area3          0.0017309  0.0005336   3.244 0.001180 **
## poly(GDP, 2)1 -0.1290709  0.8393944  -0.154 0.877794
## poly(GDP, 2)2 -3.5640040  1.0023390  -3.556 0.000377 ***
## MEM1          -0.1007204  0.1073557  -0.938 0.348145
## MEM4           0.0804221  0.0886891   0.907 0.364519
## MEM8          -0.0616110  0.0824171  -0.748 0.454730
## ---
## Signif. codes:  0 '***' 0.001 '**' 0.01 '*' 0.05 '.' 0.1 ' ' 1
##
```

```
## (Dispersion parameter for poisson family taken to be 1)
##
##     Null deviance: 130.061  on 36  degrees of freedom
## Residual deviance:  69.553  on 30  degrees of freedom
## AIC: 180.9
##
## Number of Fisher Scoring iterations: 5
```

```
print("Likelihood ratio test (chisq) shows that the negbinomial fits the data better that the poisson")
```

```
## [1] "Likelihood ratio test (chisq) shows that the negbinomial fits the data better that the poisson"
```

```
pchisq(2 * (logLik(new_mod2a) - logLik(mod)), df = 1, lower.tail = FALSE)
```

```
## 'log Lik.' 0.01803628 (df=8)
```

```
library(hermite)
```

```
## Loading required package: maxLik

## Loading required package: miscTools

##
## Please cite the 'maxLik' package as:
## Henningsen, Arne and Toomet, Ott (2011). maxLik: A package for maximum likelihood estimation in R. Co
##
## If you have questions, suggestions, or comments regarding the 'maxLik' package, please use a forum or
## https://r-forge.r-project.org/projects/maxlik/

##
## Attaching package: 'maxLik'

## The following object is masked from 'package:raster':
##
##     maxValue
```

```
mod3 <- glm.hermite(formula=fmla,                          #ntotal_studies ~ area3+poly(GDP5,2),
                    data = data1c,
                    link = "log",
                    m=2)
```

```
## Warning in model.matrix.default(mt, mf, contrasts): non-list contrasts argument
## ignored
```

```
summary(mod3)
```

```
## Call:
## glm.hermite(formula = fmla, data = data1c, link = "log", m = 2)
##
## Deviance Residuals:
##        Min         1Q     Median         3Q        Max
## -12.552321  -5.037995  -2.797650  -1.277825 539.882406
##
## Coefficients:
##                     Estimate   Std. Error     z value      p-value
## (Intercept)       0.924840418 0.1742780316  5.30669534 1.116305e-07
## area3             0.001750807 0.0006948071  2.51984648 1.174060e-02
## poly(GDP, 2)1    -0.023901019 1.0323985310 -0.02315096 9.815299e-01
## poly(GDP, 2)2    -3.393182295 1.2528758879 -2.70831479 6.762585e-03
## MEM1             -0.104610133 0.1372587161 -0.76213836 4.459774e-01
```

```
## MEM4              0.086720205 0.1147186893  0.75593790 4.496864e-01
## MEM8             -0.052688605 0.1066643191 -0.49396654 6.213298e-01
## dispersion.index  1.676111135 0.1648718915  8.10739721 2.204249e-03
## order             2.000000000           NA          NA           NA
## (Likelihood ratio test against Poisson is reported by *z value* for *dispersion.index*)
##
## AIC:  174.8391
```

```
print("Likelihood ratio test (chisq) shows that the spatially corrected hermite does not fit the data be
```

```
## [1] "Likelihood ratio test (chisq) shows that the spatially corrected hermite does not fit the data b
```

```
pchisq(2 * ( mod3$loglik-logLik(new_mod2a)), df = 1, lower.tail = FALSE)
```

```
## 'log Lik.' 0.1165048 (df=8)
```

```
save(data1b,file="data1b_spatial_regressions2_Rmd.RData")
```
