## Appendix S3 for "Conserving genetic diversity during climate change: Niche marginality and discrepant monitoring capacity in Europe"

Appendix S3. Maps of marginality of European amphibian species. The climate marginality of each species is depicted in a series of three figures: raw marginality in which a value of 1 indicates the 99th percentile of the species niche space within its current range (filtered for appropriate habitat with a CORINE land cover layer), the current geographic distribution of areas within the 25% most marginal climatic niche conditions and the remaining core area, and a predicted future distribution of core and marginal areas within the current species range. Please see the Online Methods section for a description of marginality calculations.

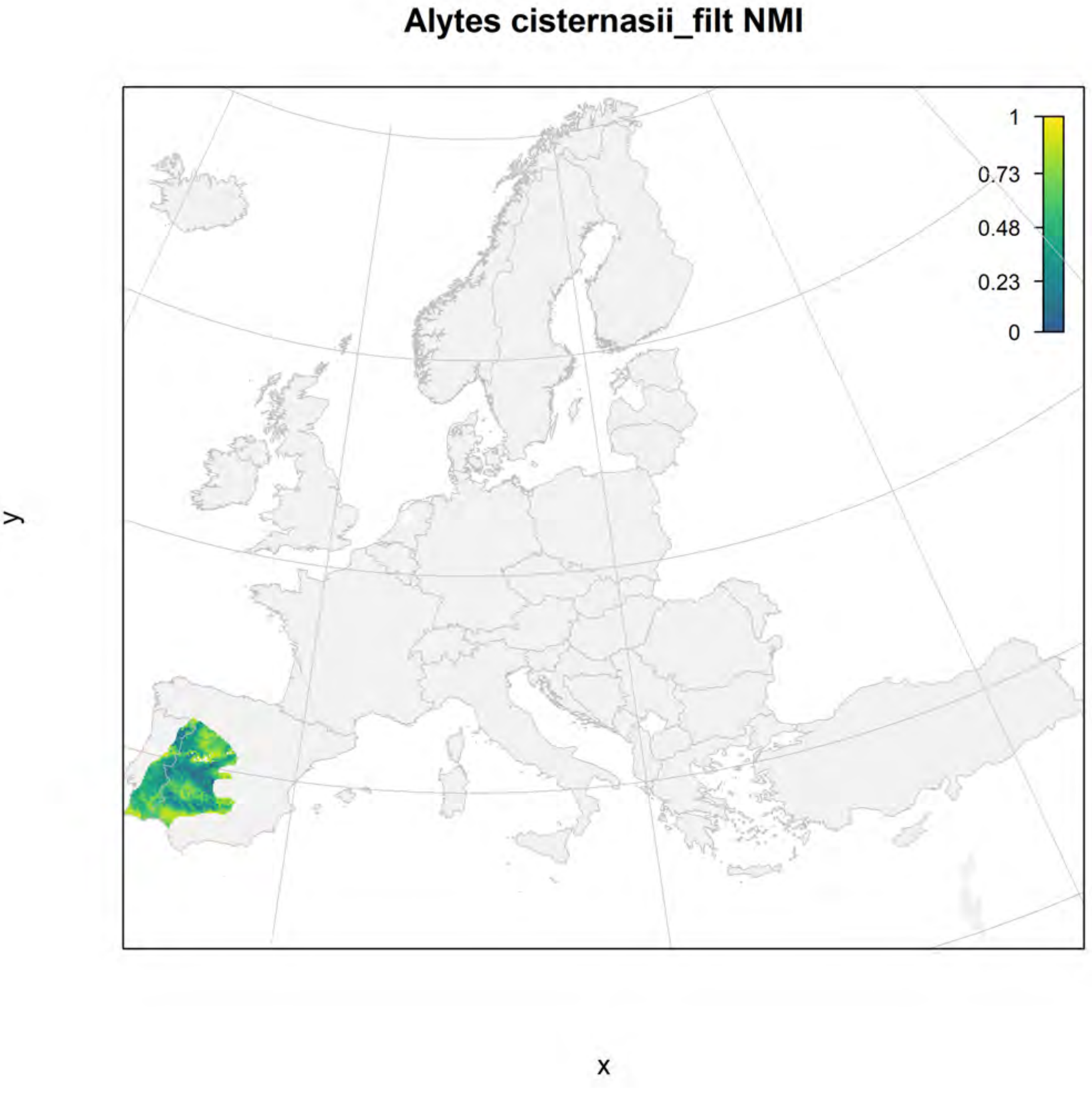

Alytes cisternasii_filt NMI
1
0.73
0.48
0.23
0
y
x

#### Alytes cisternasii_filt current marginality

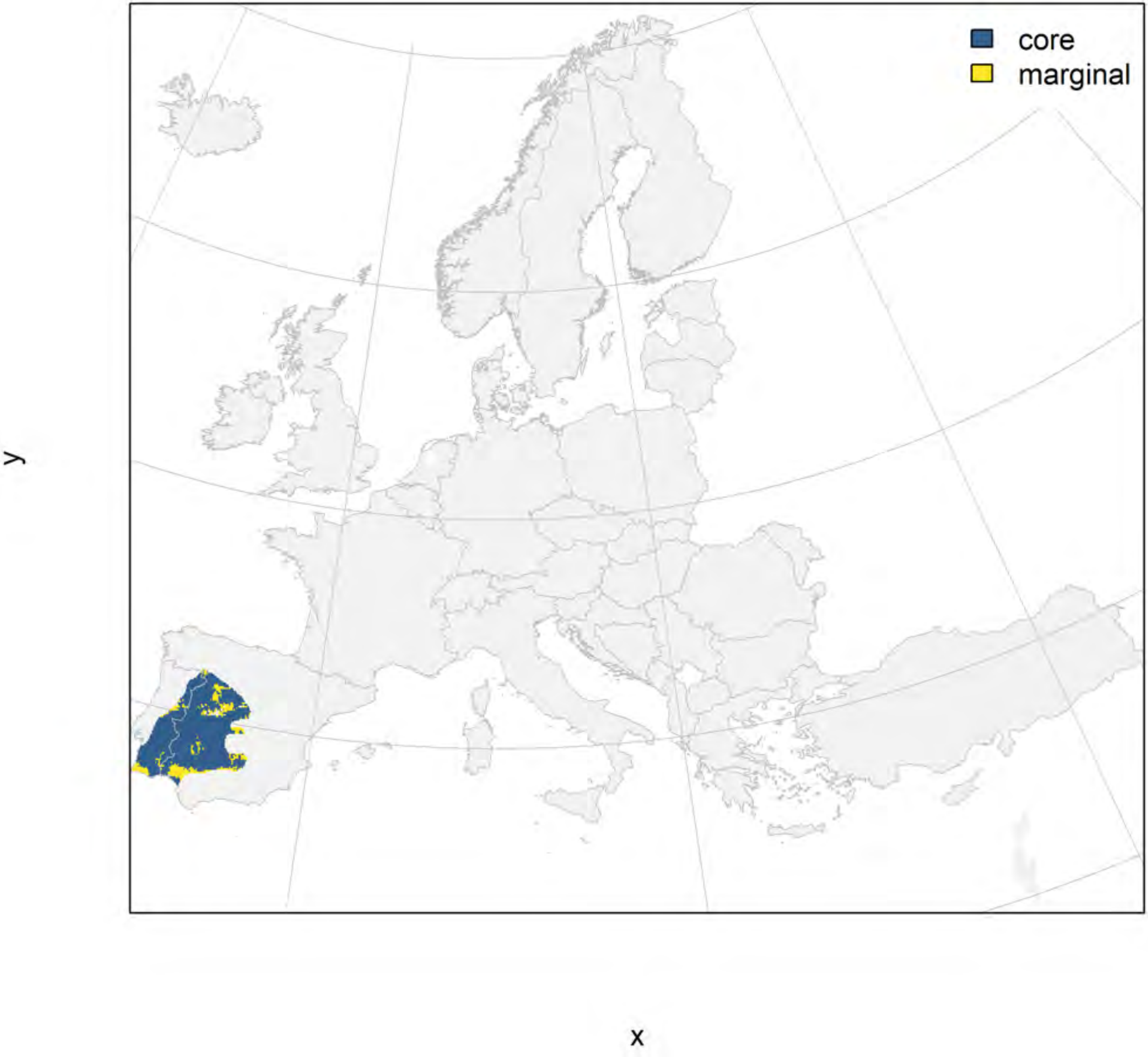

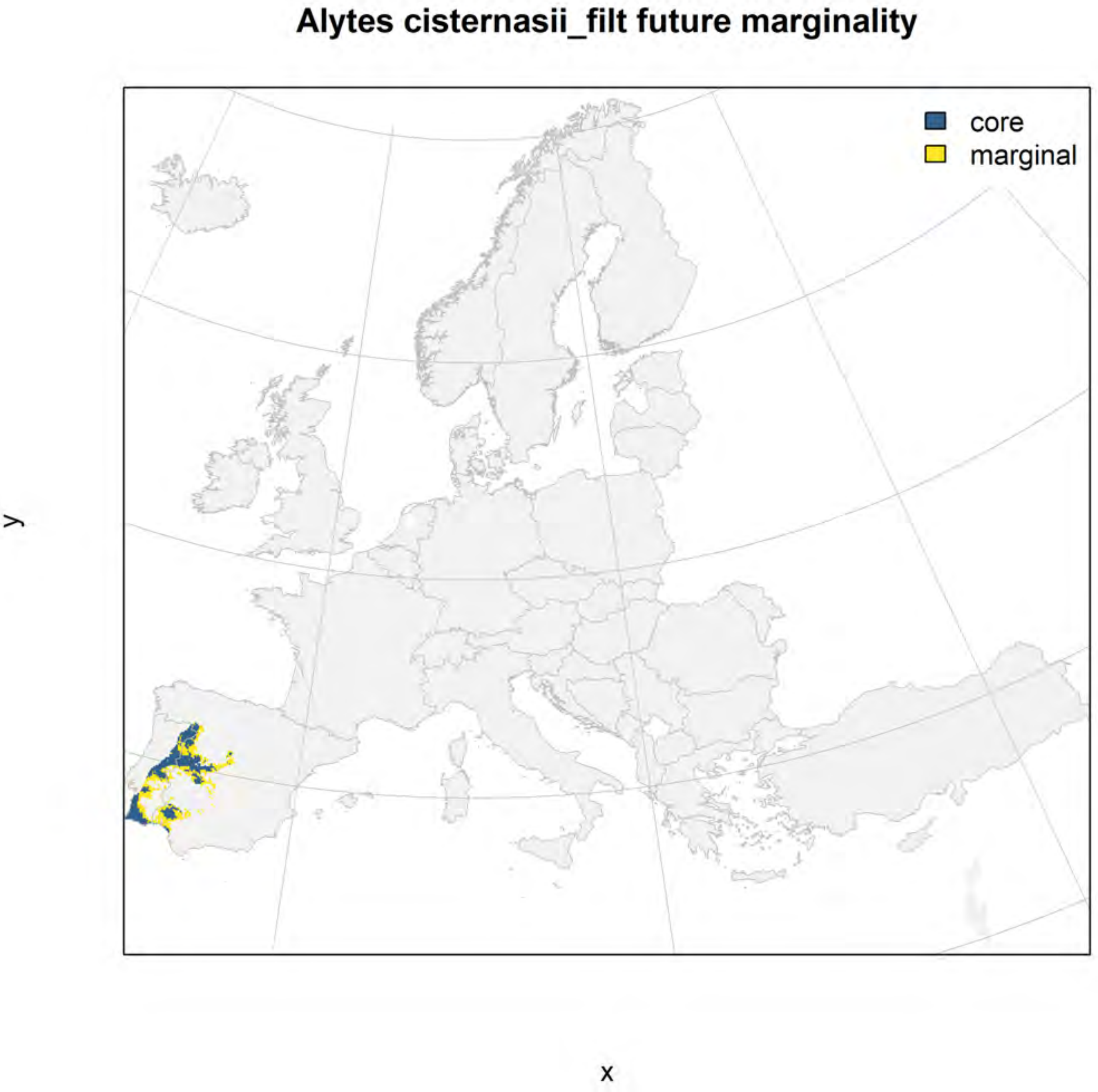
Alytes cisternasii_filt future marginality
core
marginal
y
x

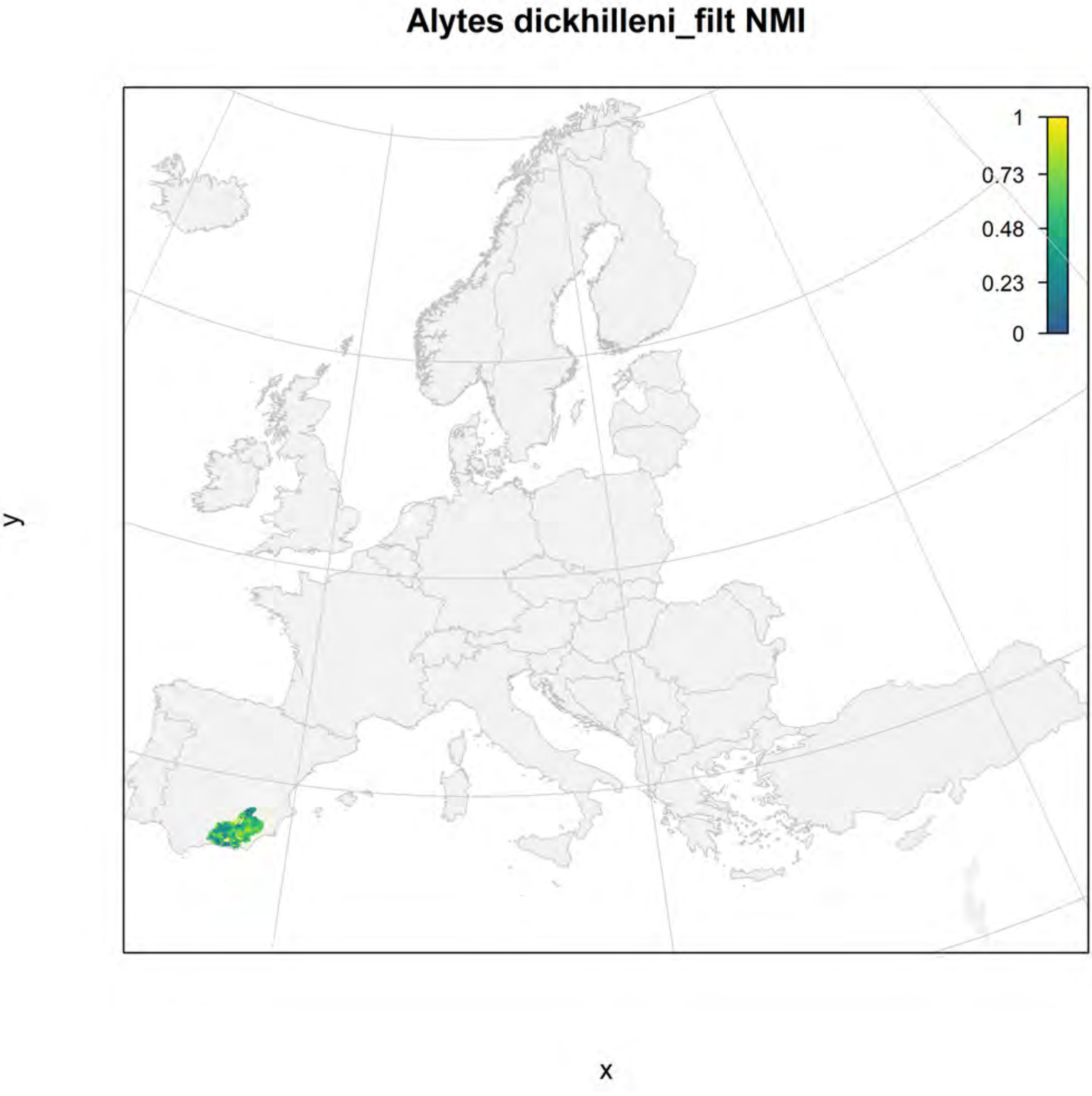
Alytes dickhilleni_filt NMI
1
0.73
0.48
0.23
0
y
x

### Alytes dickhilleni_filt current marginality

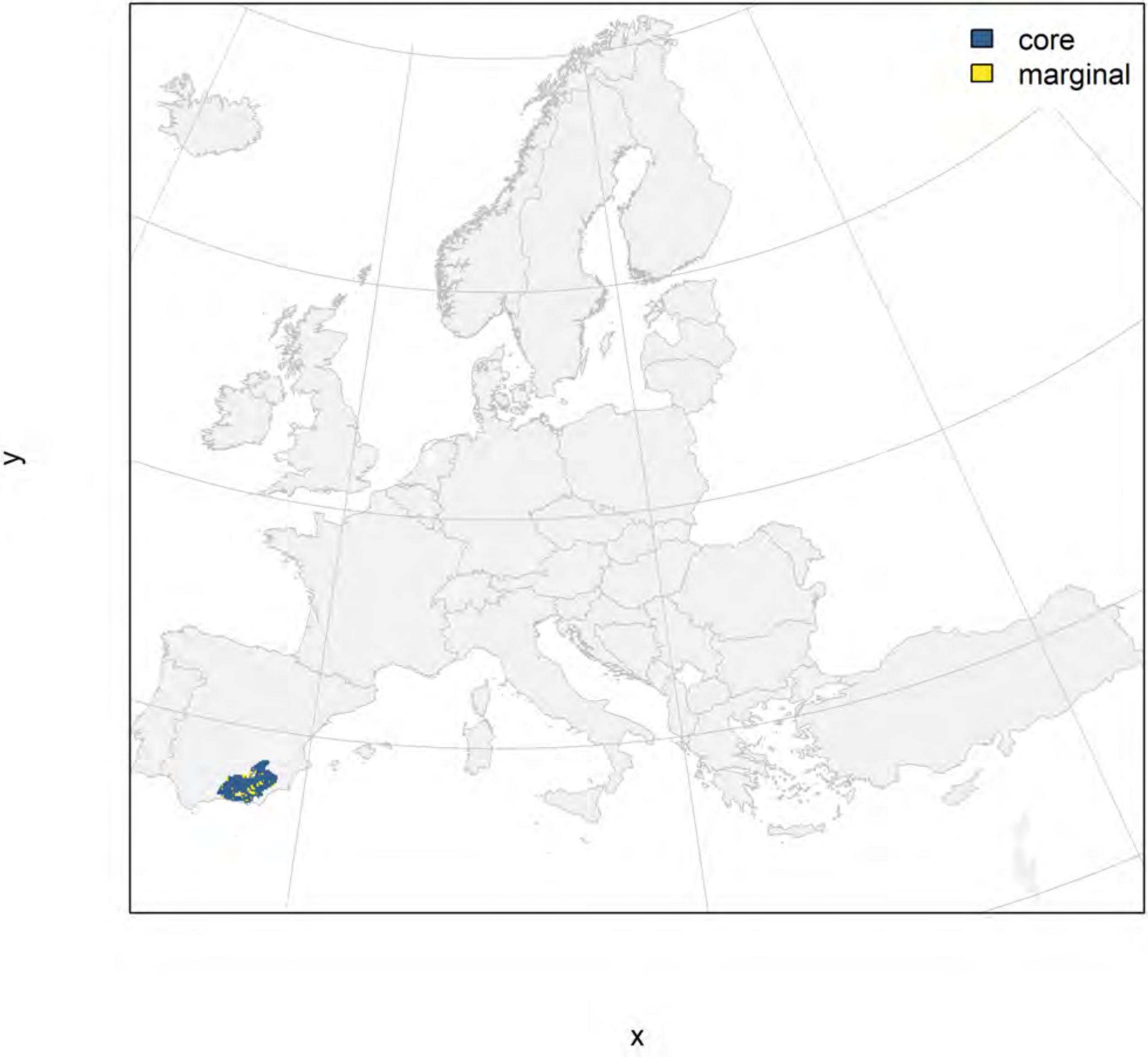

### Alytes dickhilleni_filt future marginality

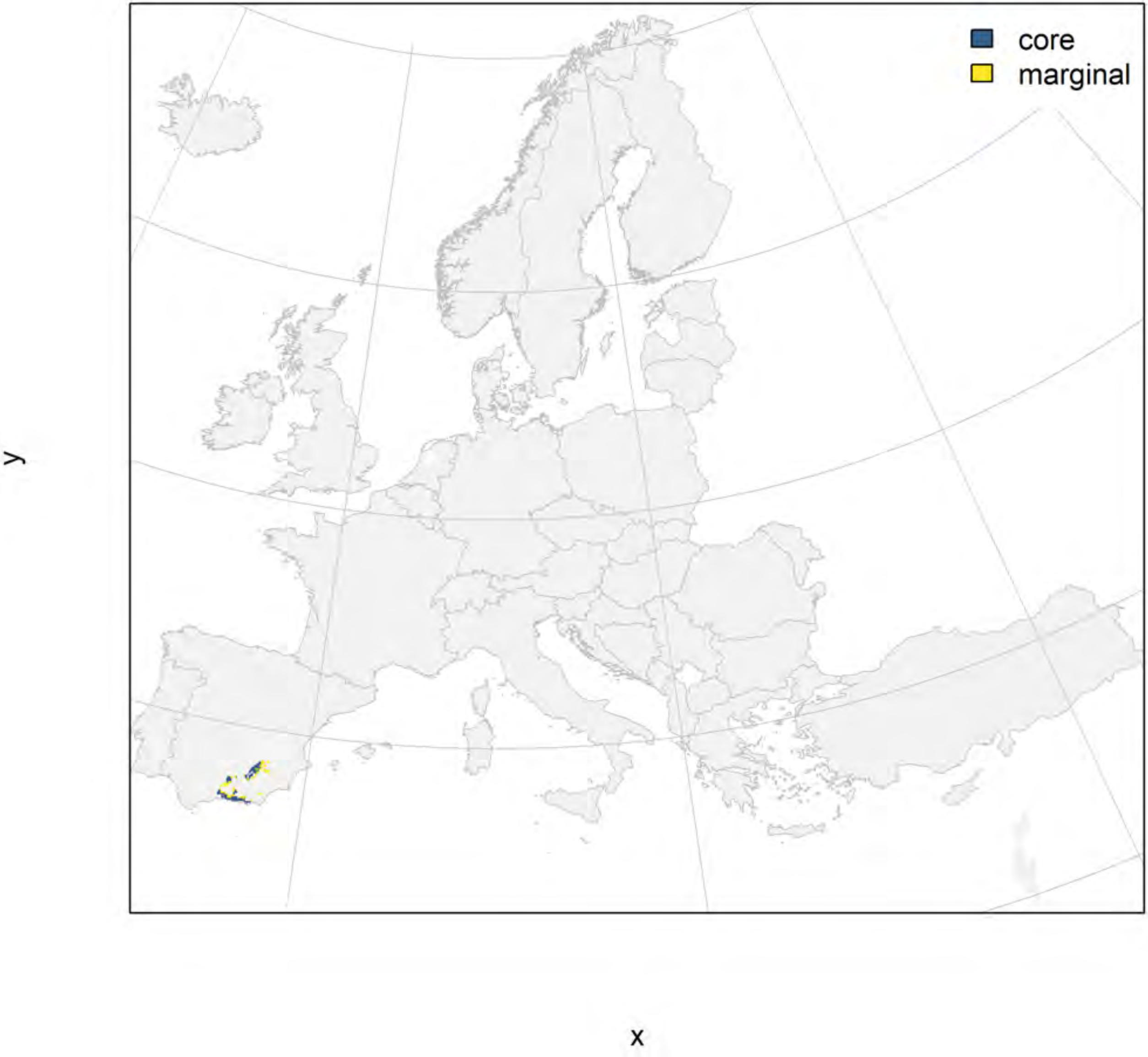

#### Alytes obstetricans_filt NMI

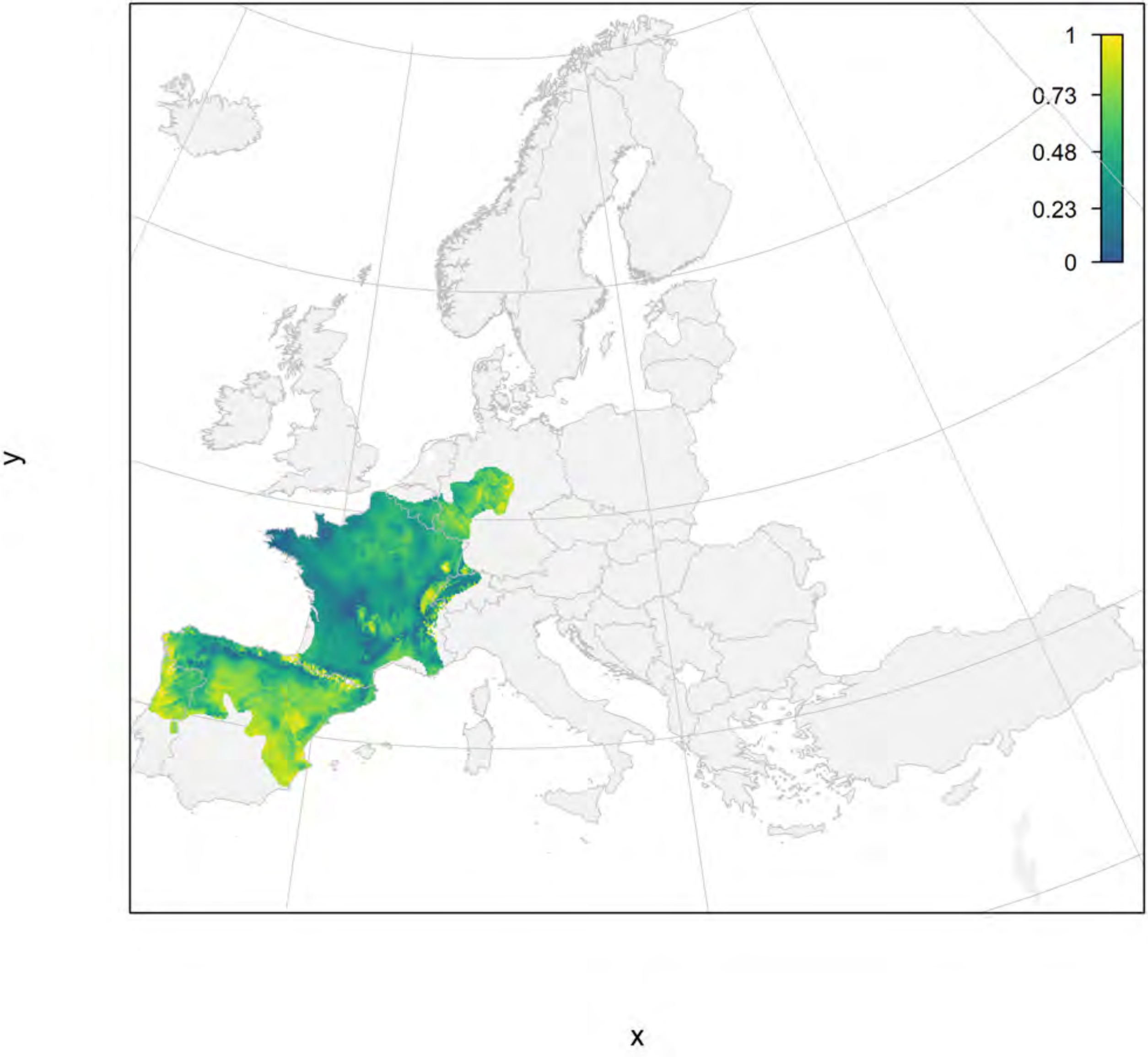

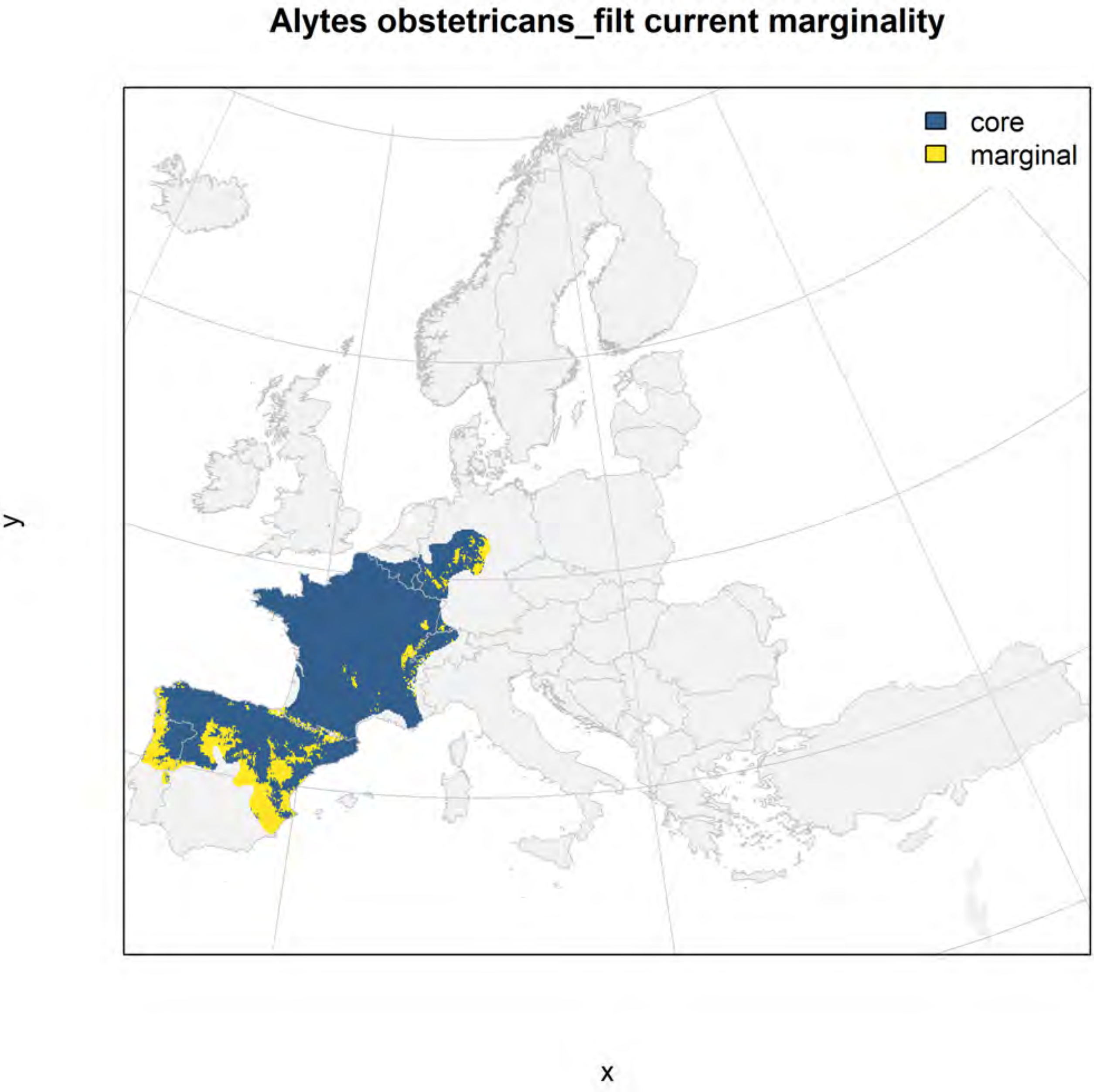
Alytes obstetricans_filt current marginality
core
marginal
y
x

#### Alytes obstetricans_filt future marginality

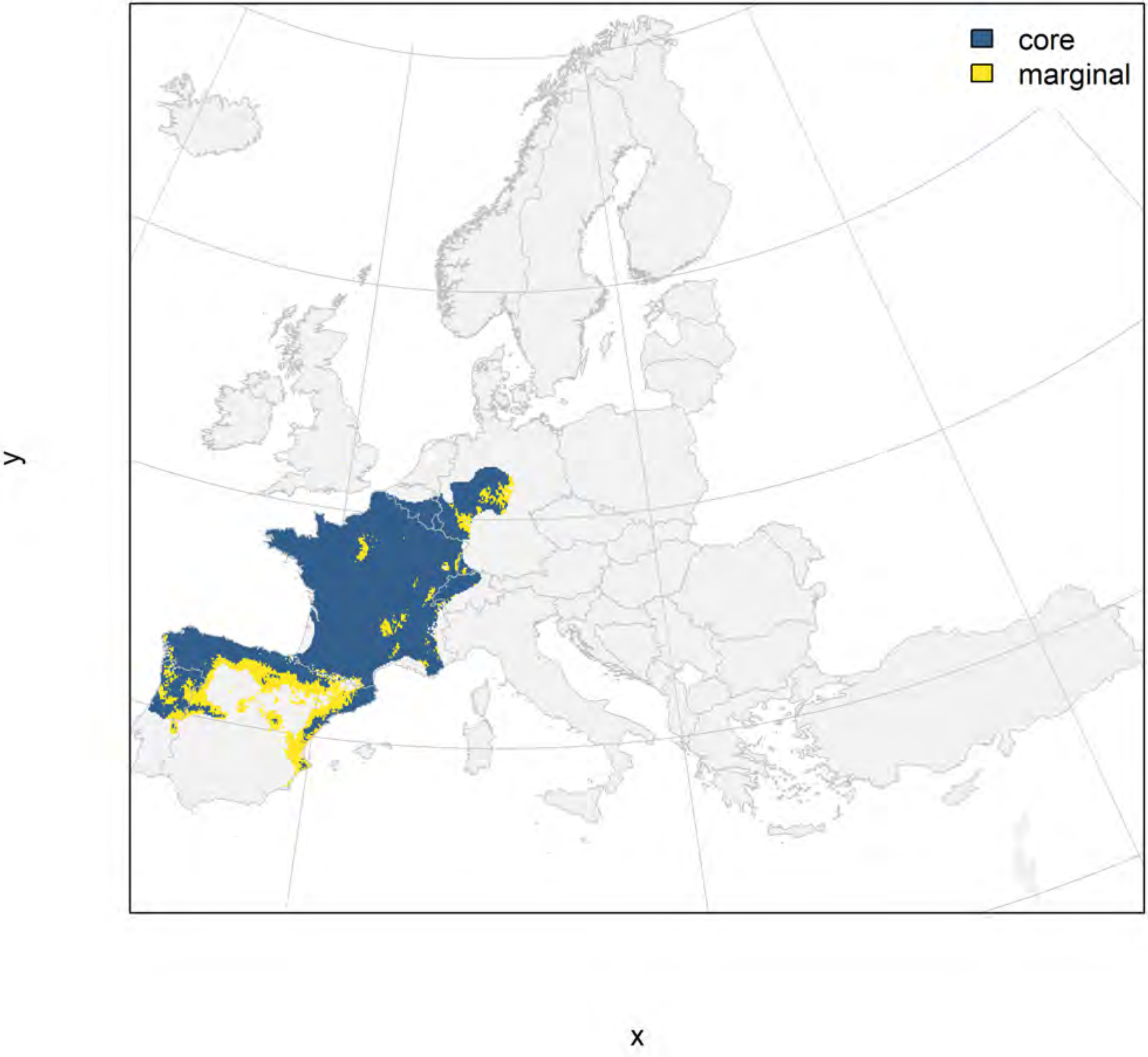

Bombina bombina_filt NMI

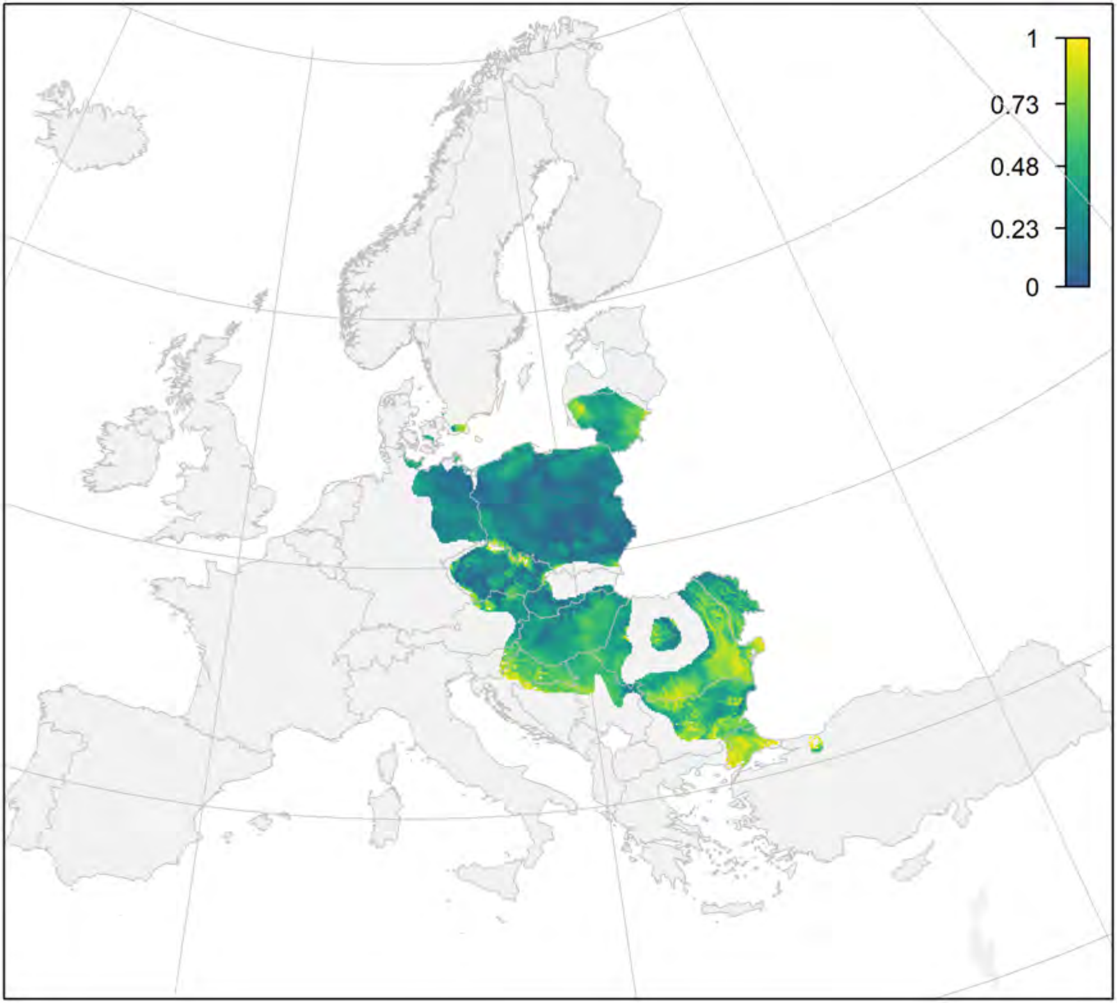


y

x

#### Bombina bombina_filt current marginality

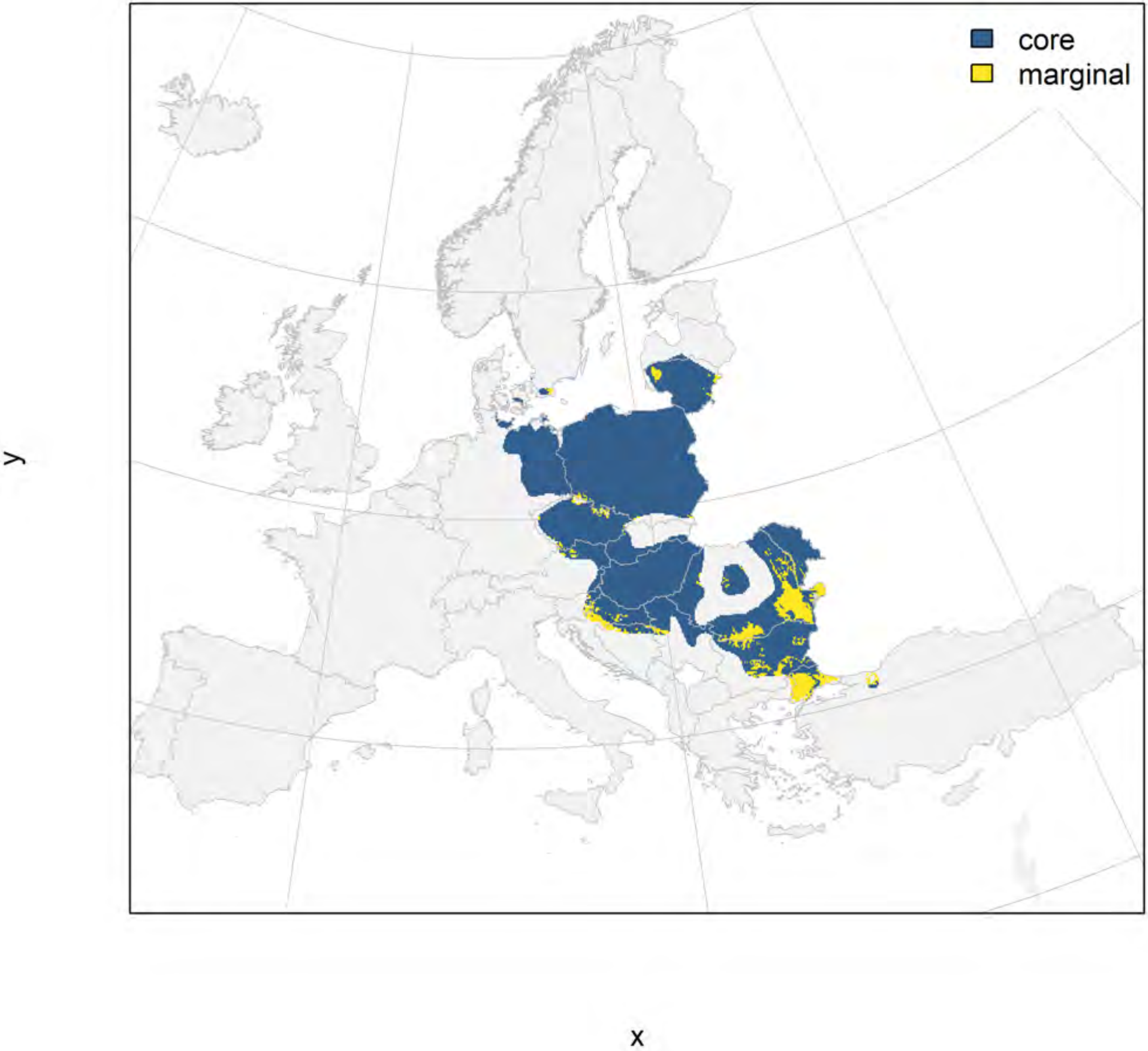

#### Bombina bombina_filt future marginality

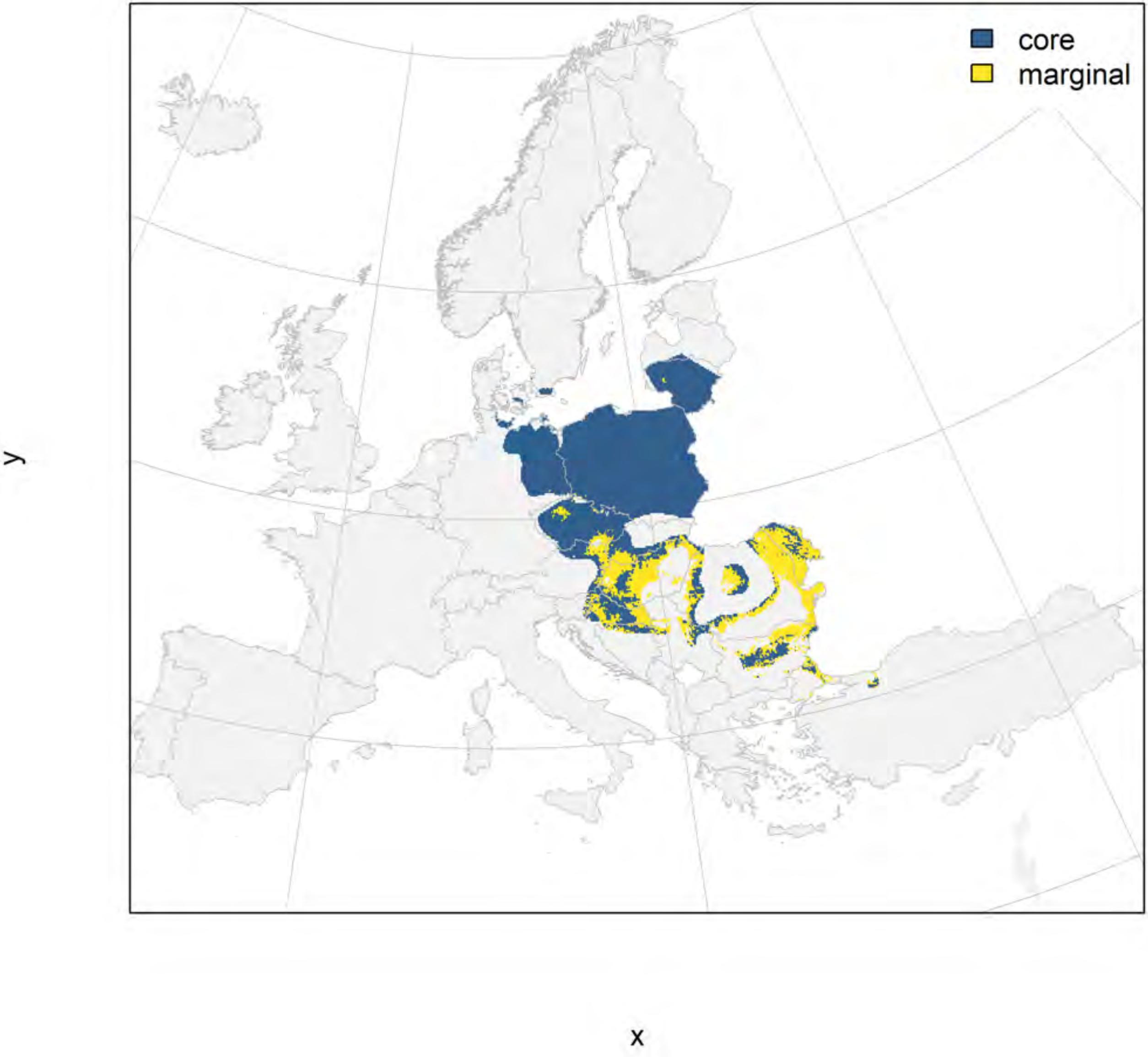

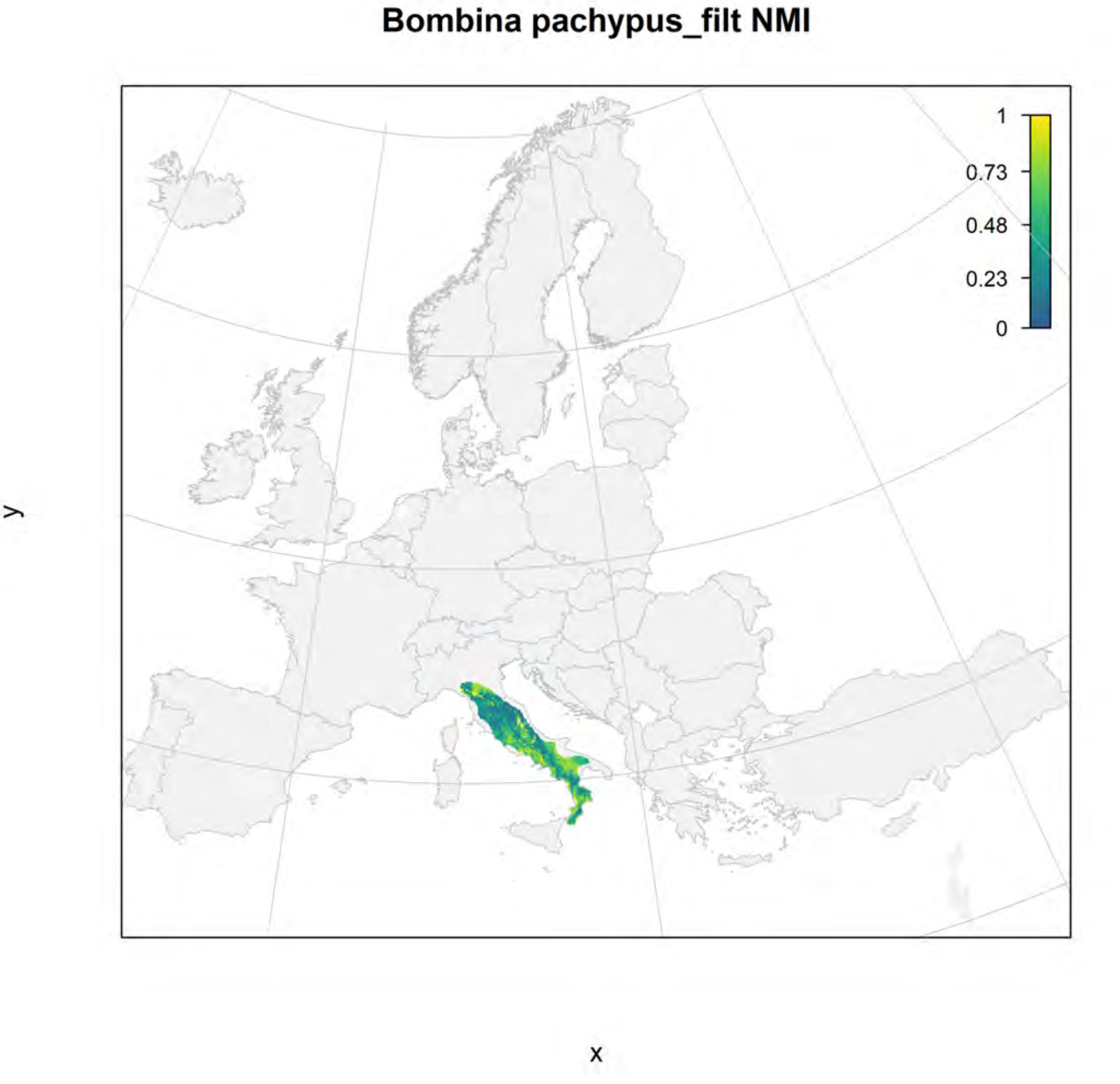
Bombina pachypus_filt NMI
1
0.73
0.48
0.23
0
y
x

#### Bombina pachypus_filt current marginality

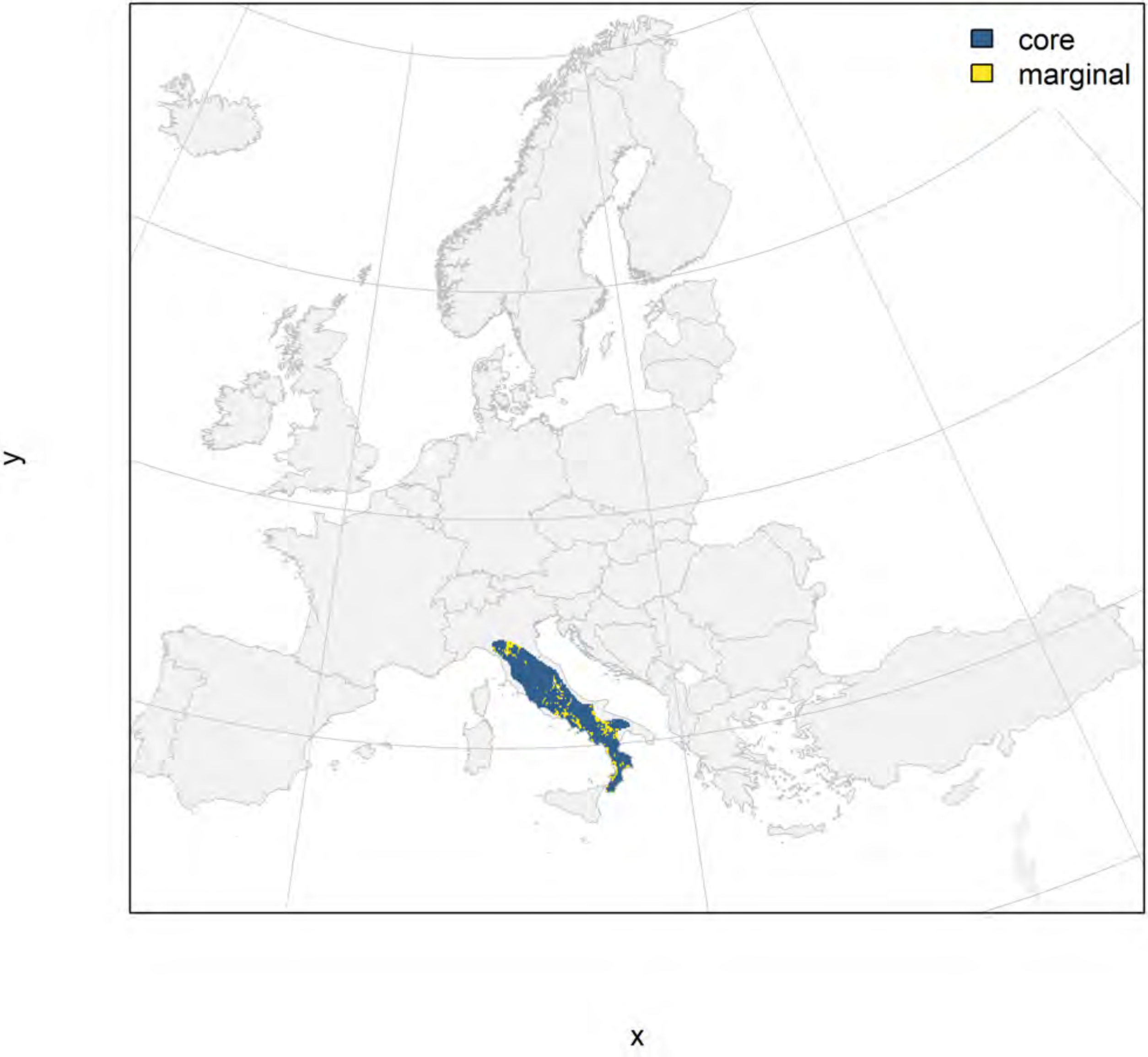

#### Bombina pachypus_filt future marginality

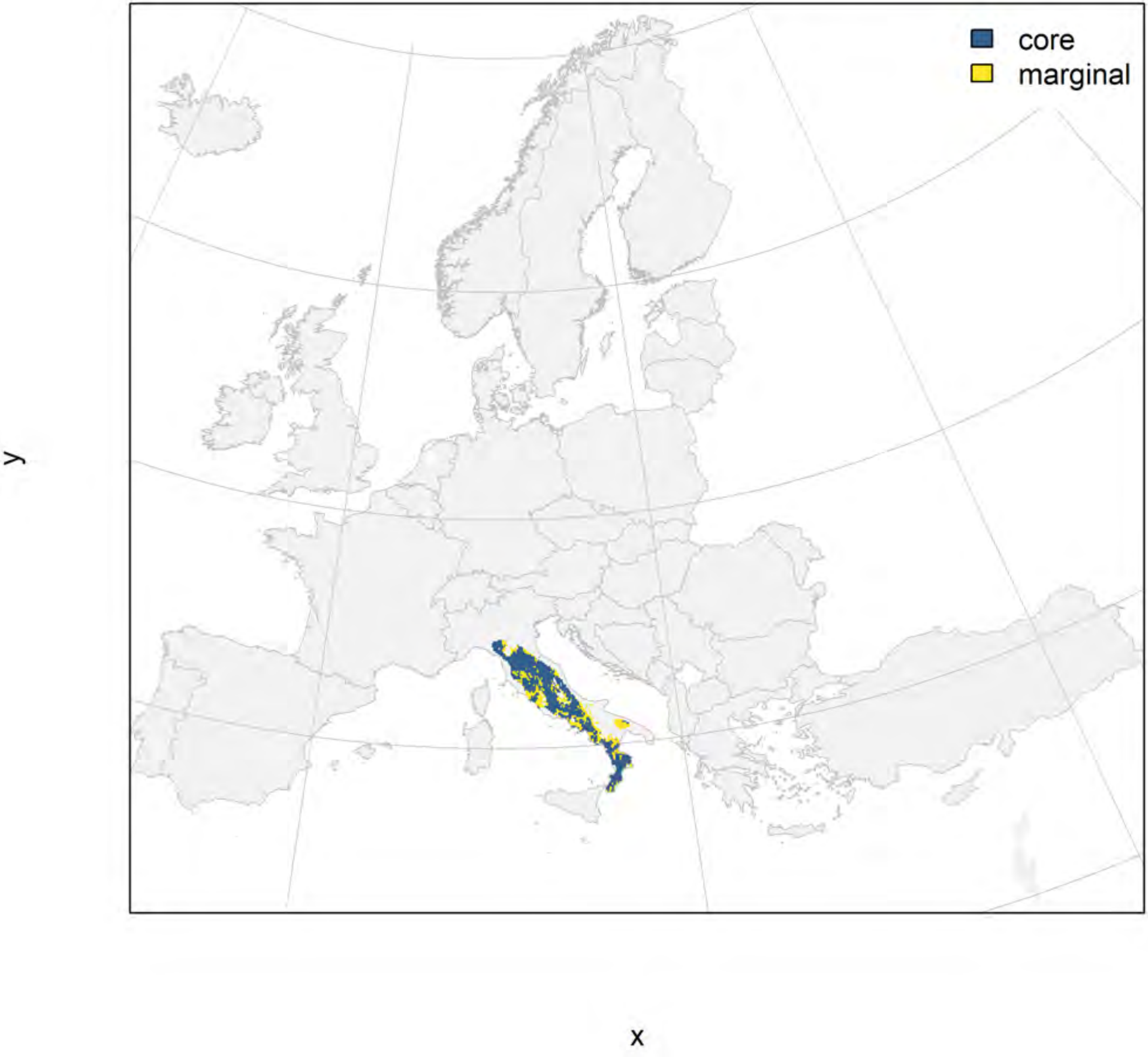

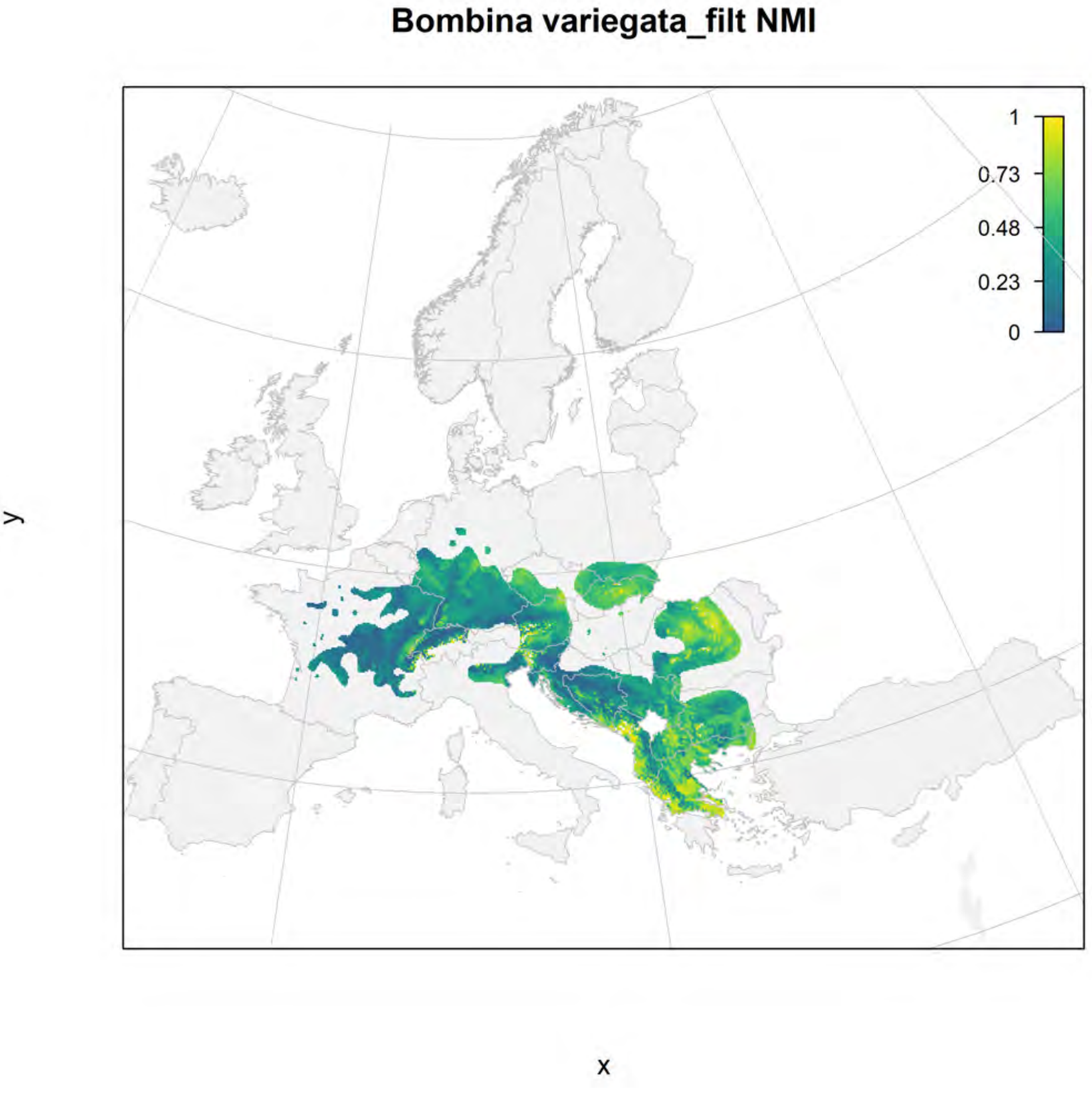

Bombina variegata_filt NMI
1
0.73
0.48
0.23
0
y
x

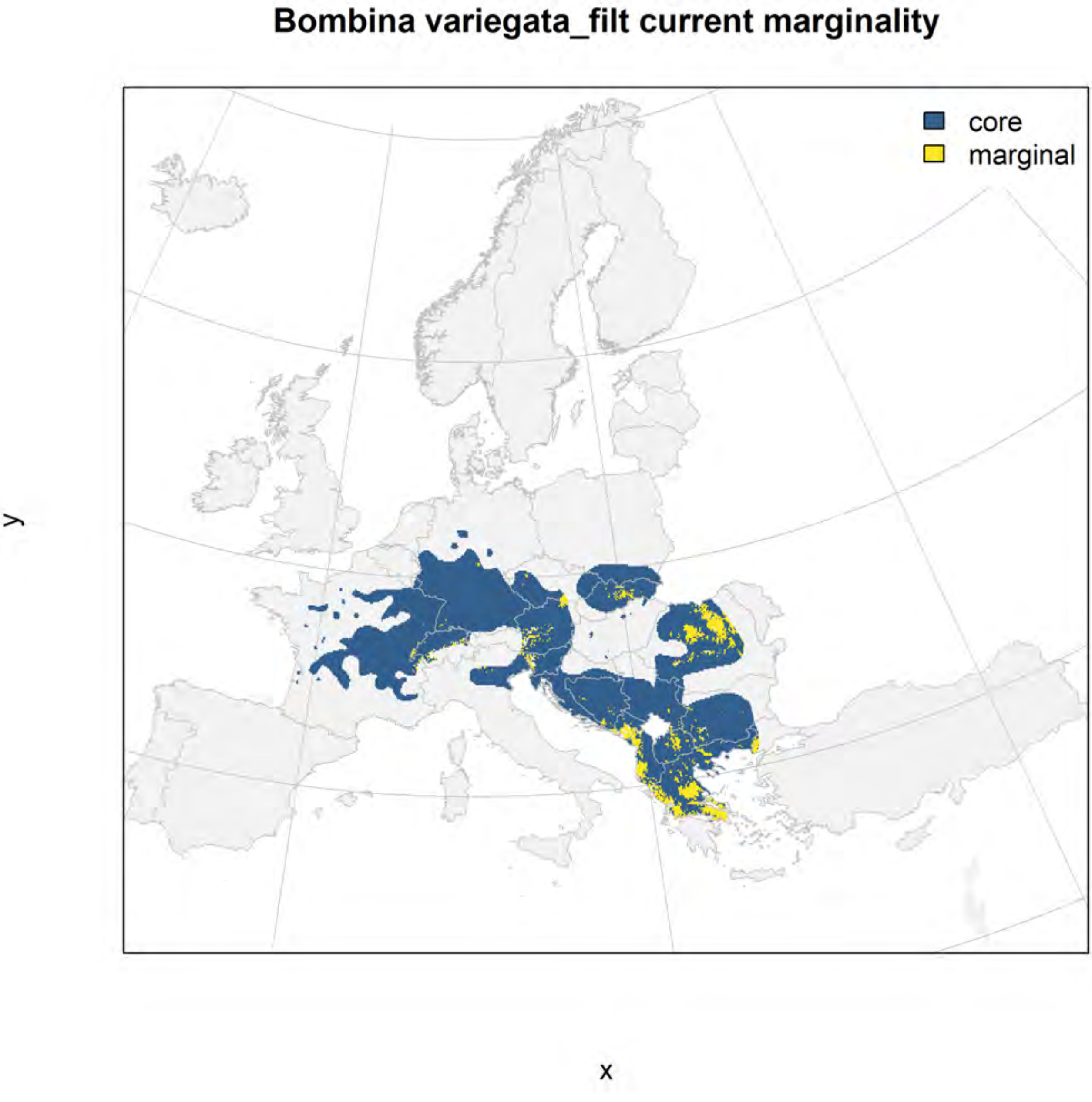
Bombina variegata_filt current marginality
core
marginal
y
x

#### Bombina variegata_filt future marginality

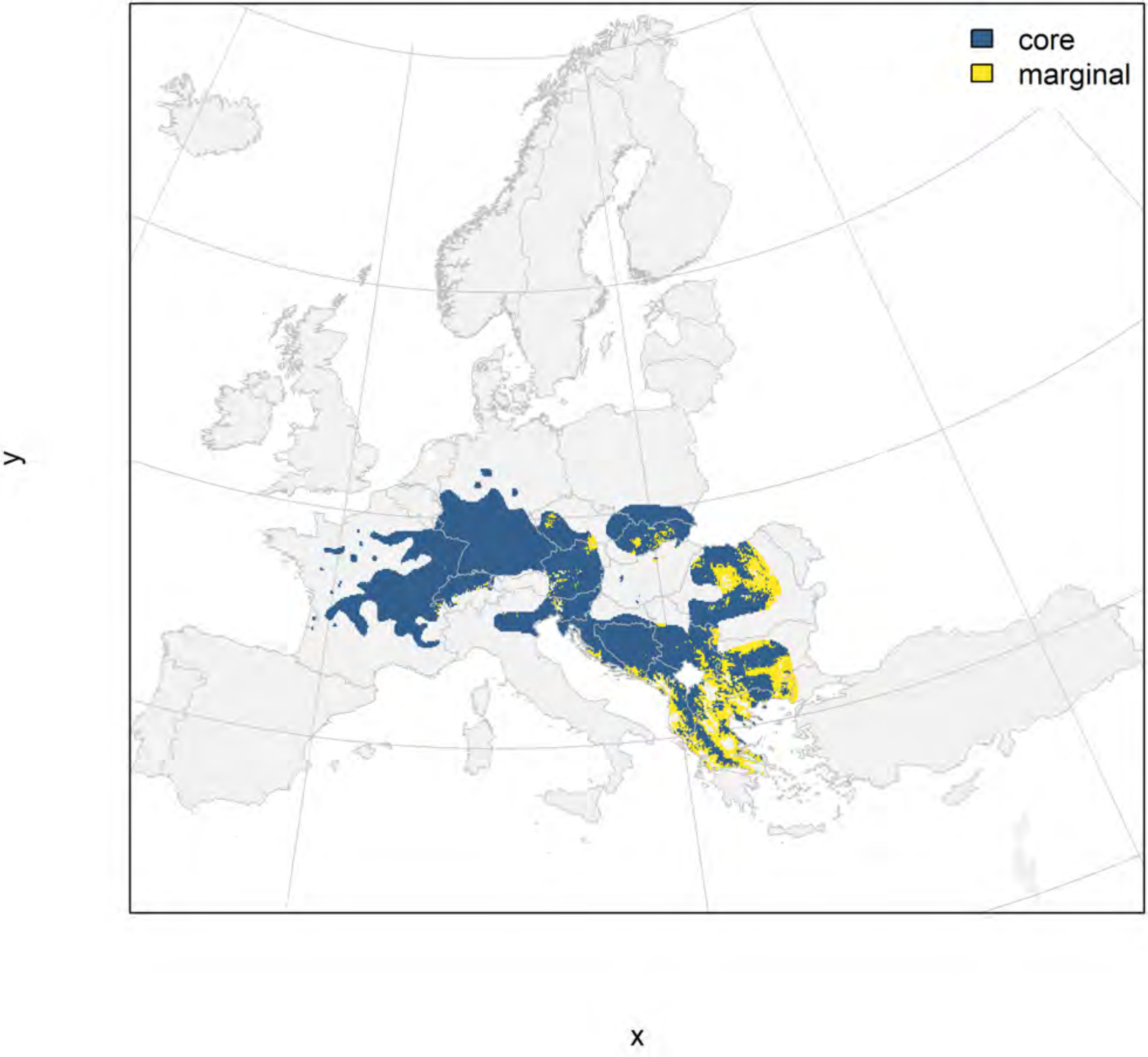

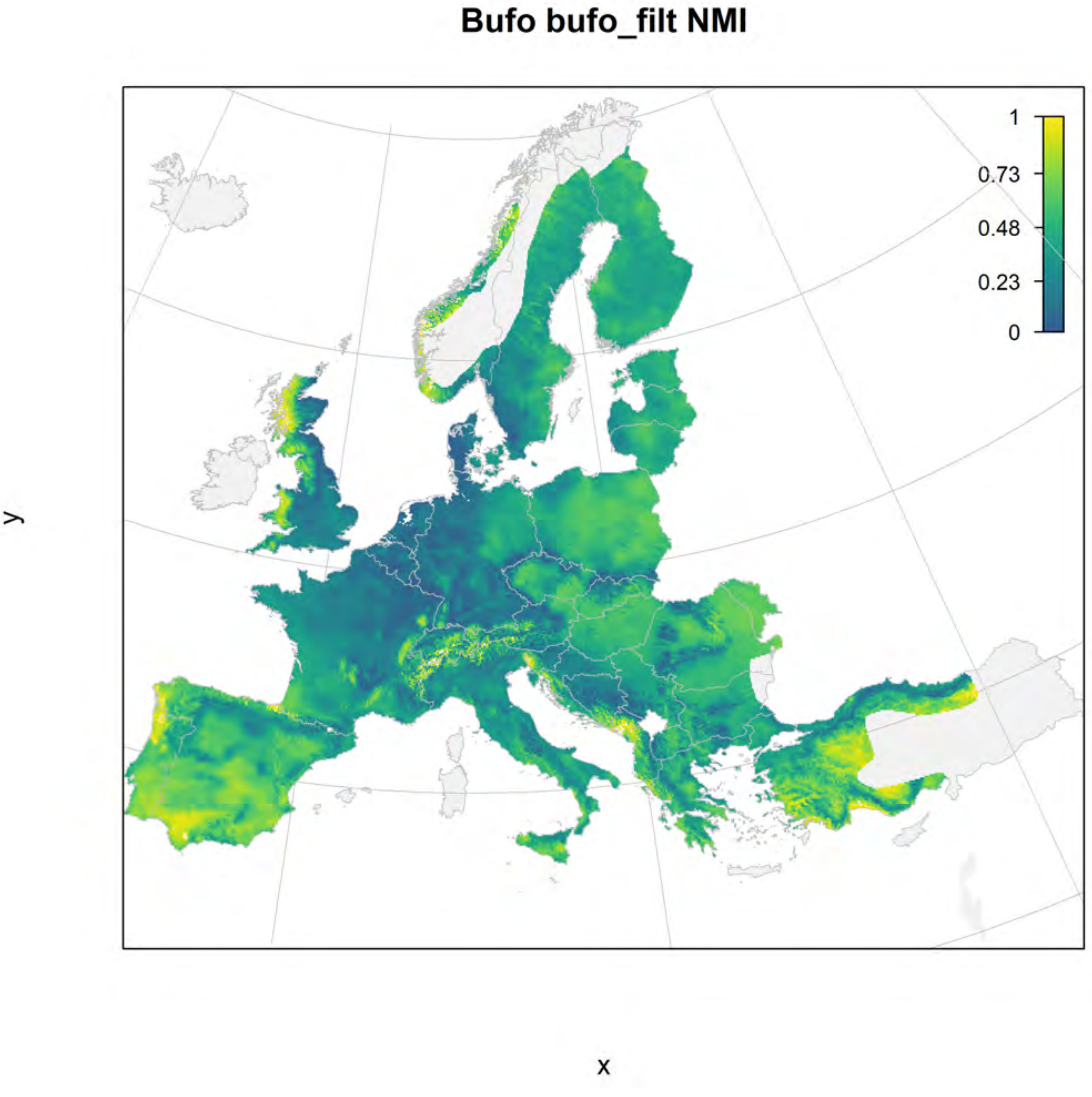
Bufo bufo_filt NMI
1
0.73
0.48
0.23
0
y
x

### Bufo bufo_filt current marginality

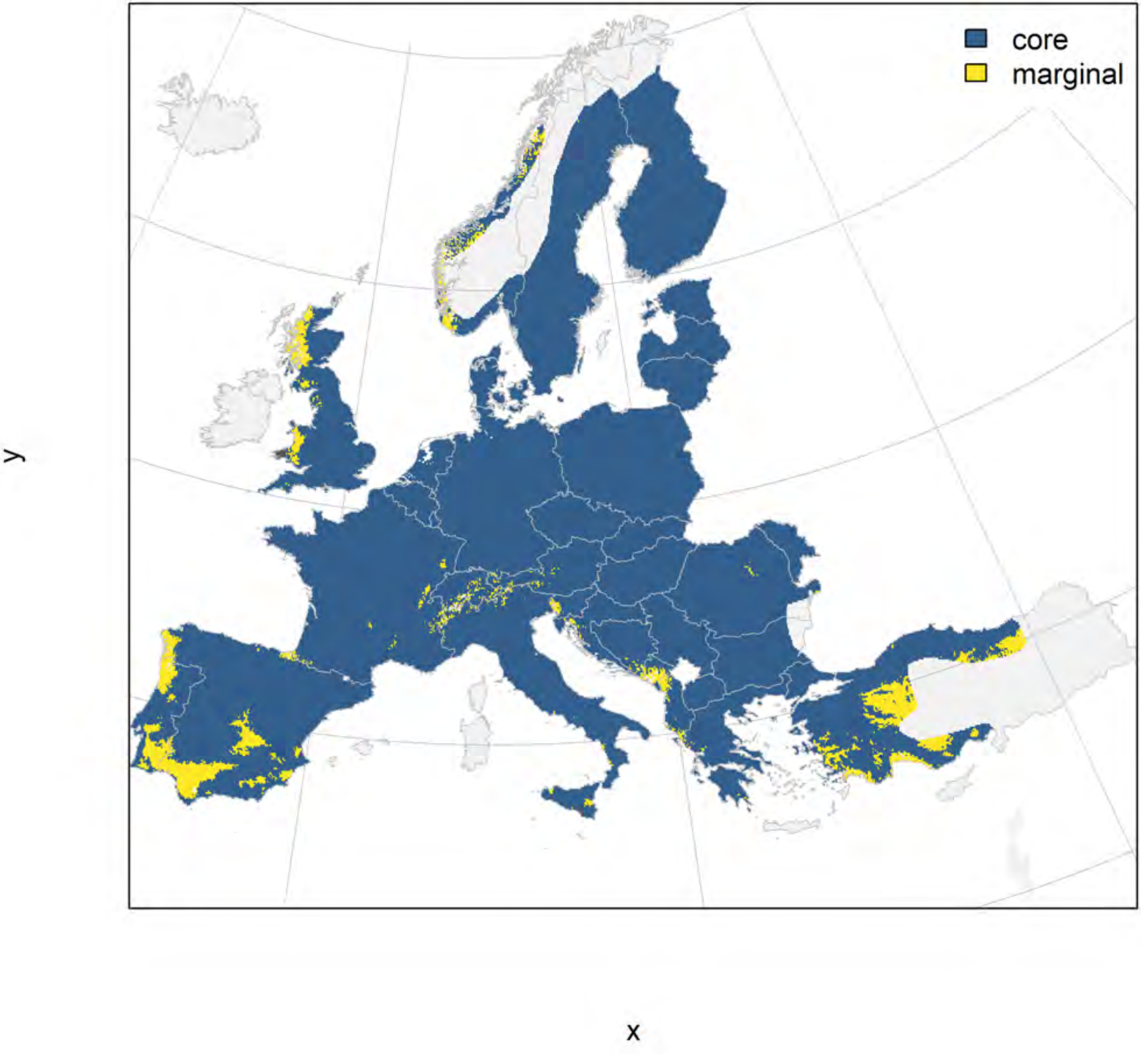

#### Bufo bufo_filt future marginality

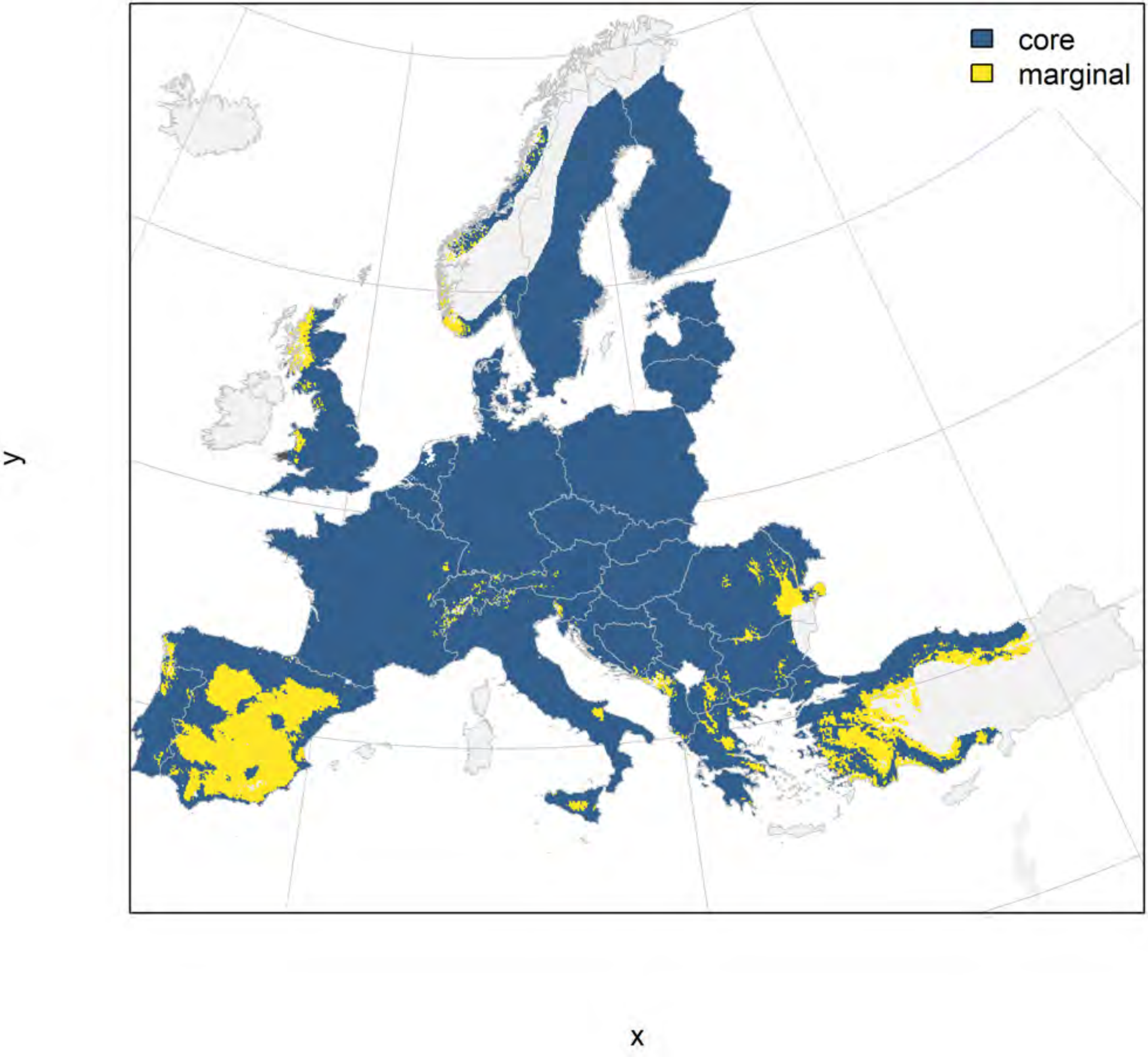

**Discoglossus galganoi_filt NMI**

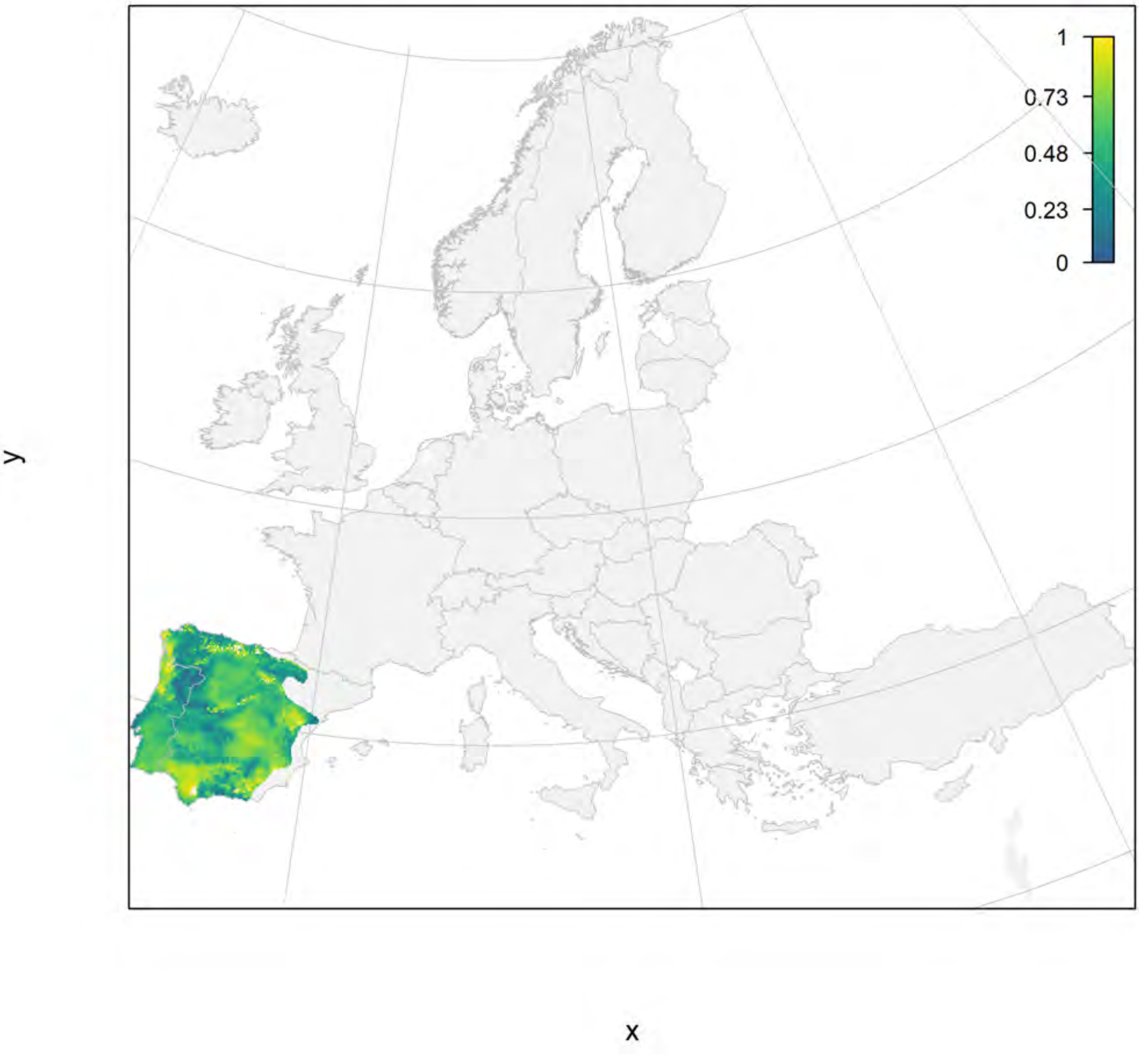

### Discoglossus galganoi_filt current marginality

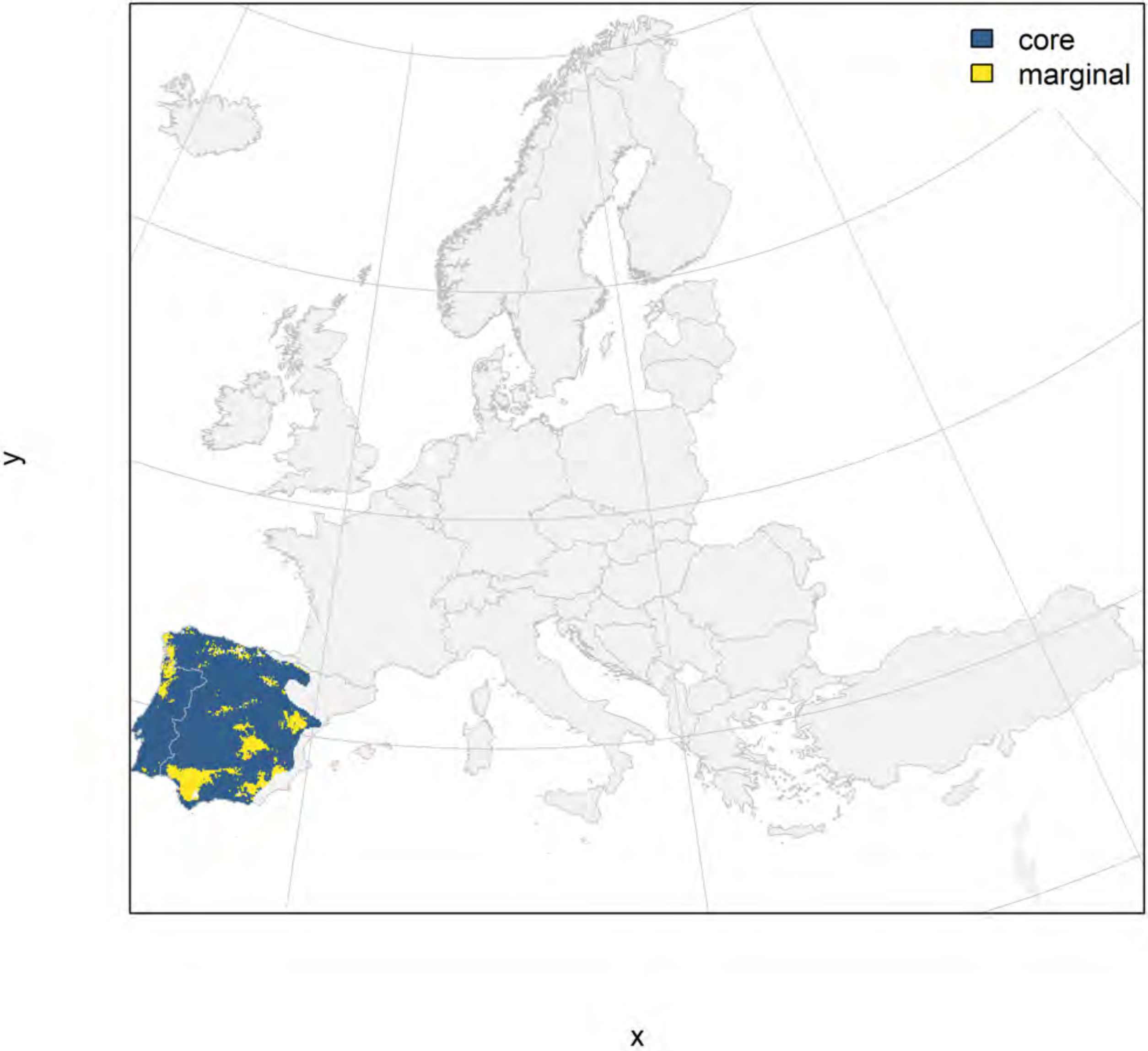

#### Discoglossus galganoi_filt future marginality

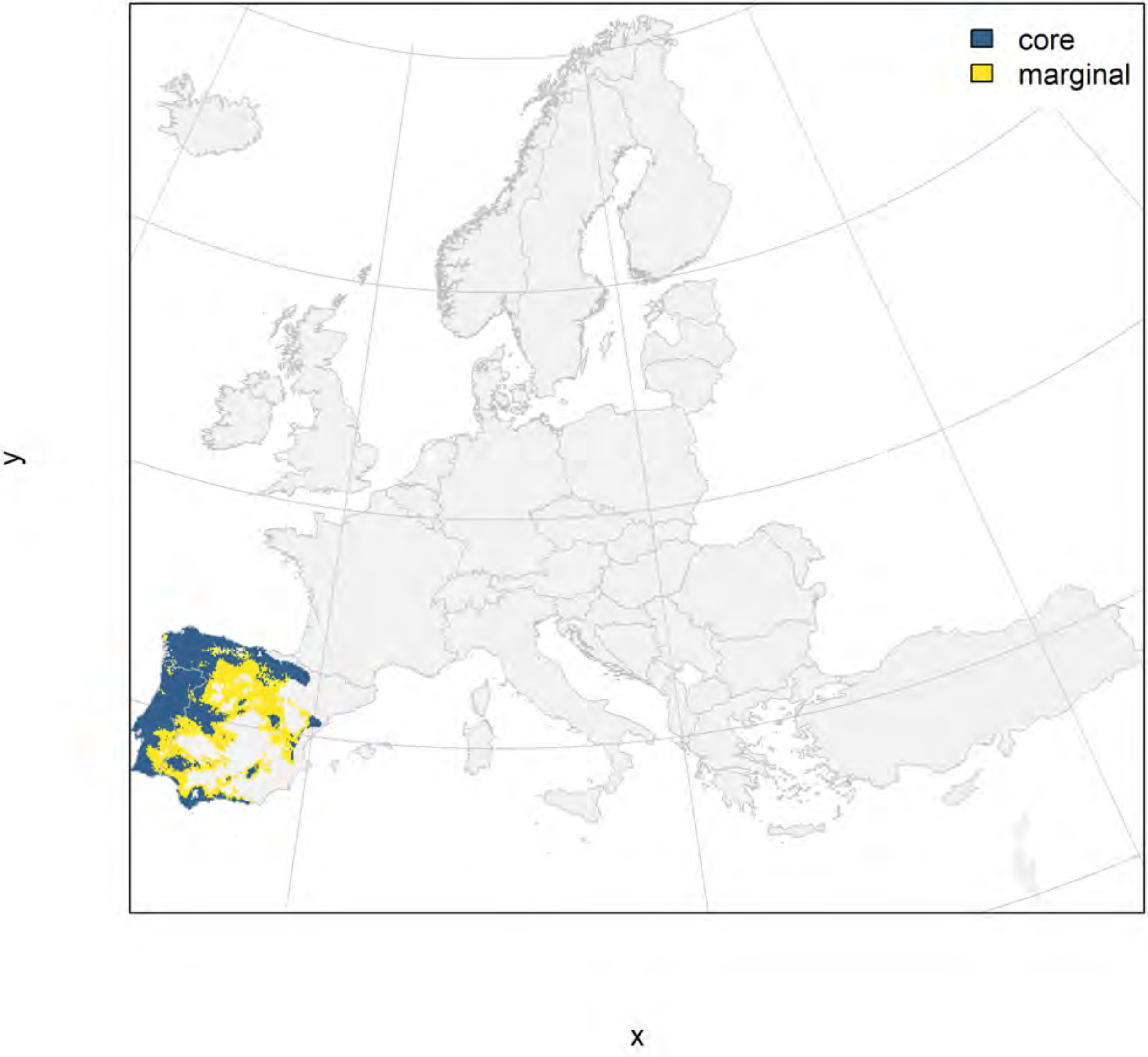

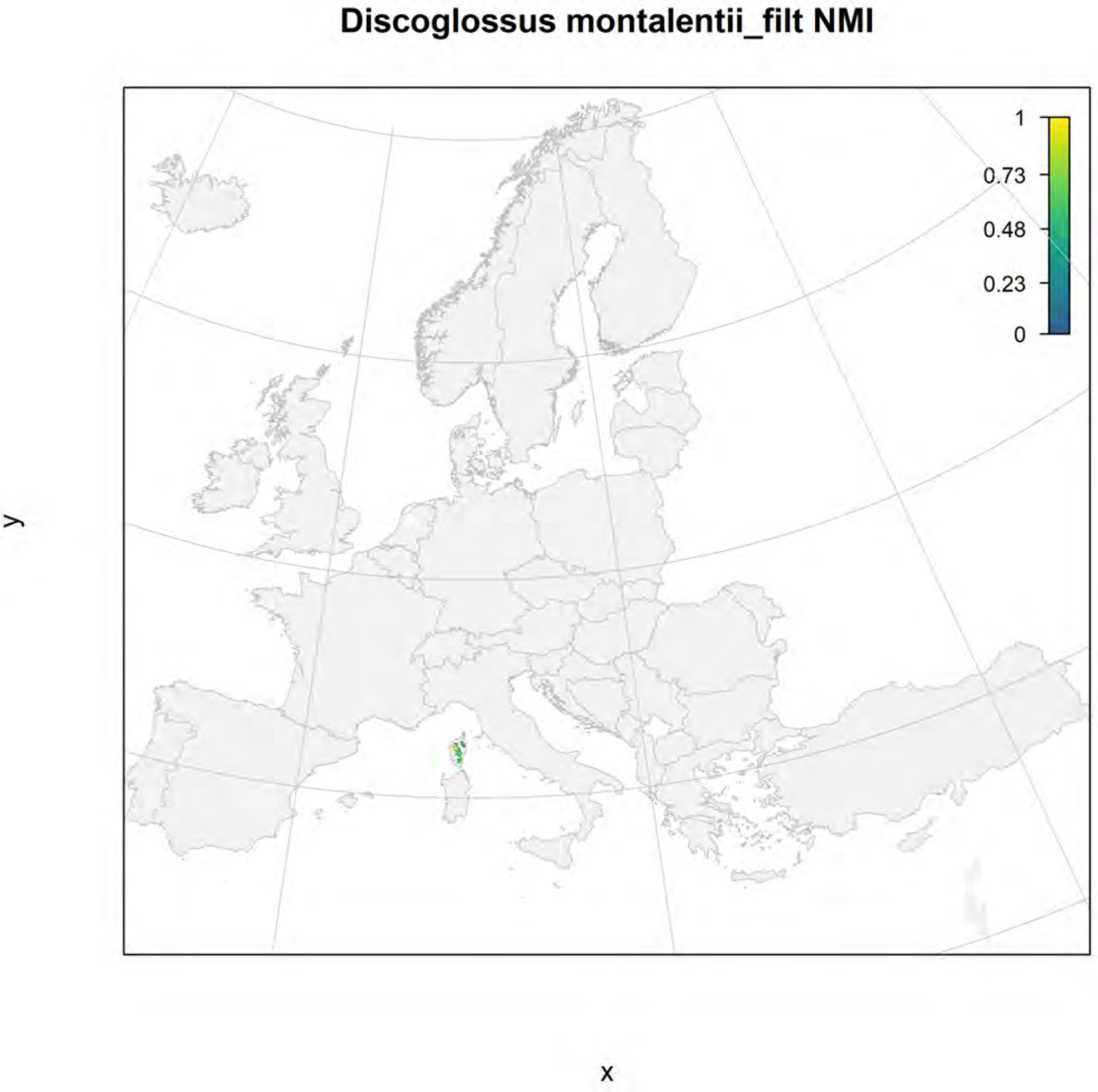
Discoglossus montalentii_filt NMI
1
0.73
0.48
0.23
0
y
x

#### Discoglossus montalentii_filt current marginality

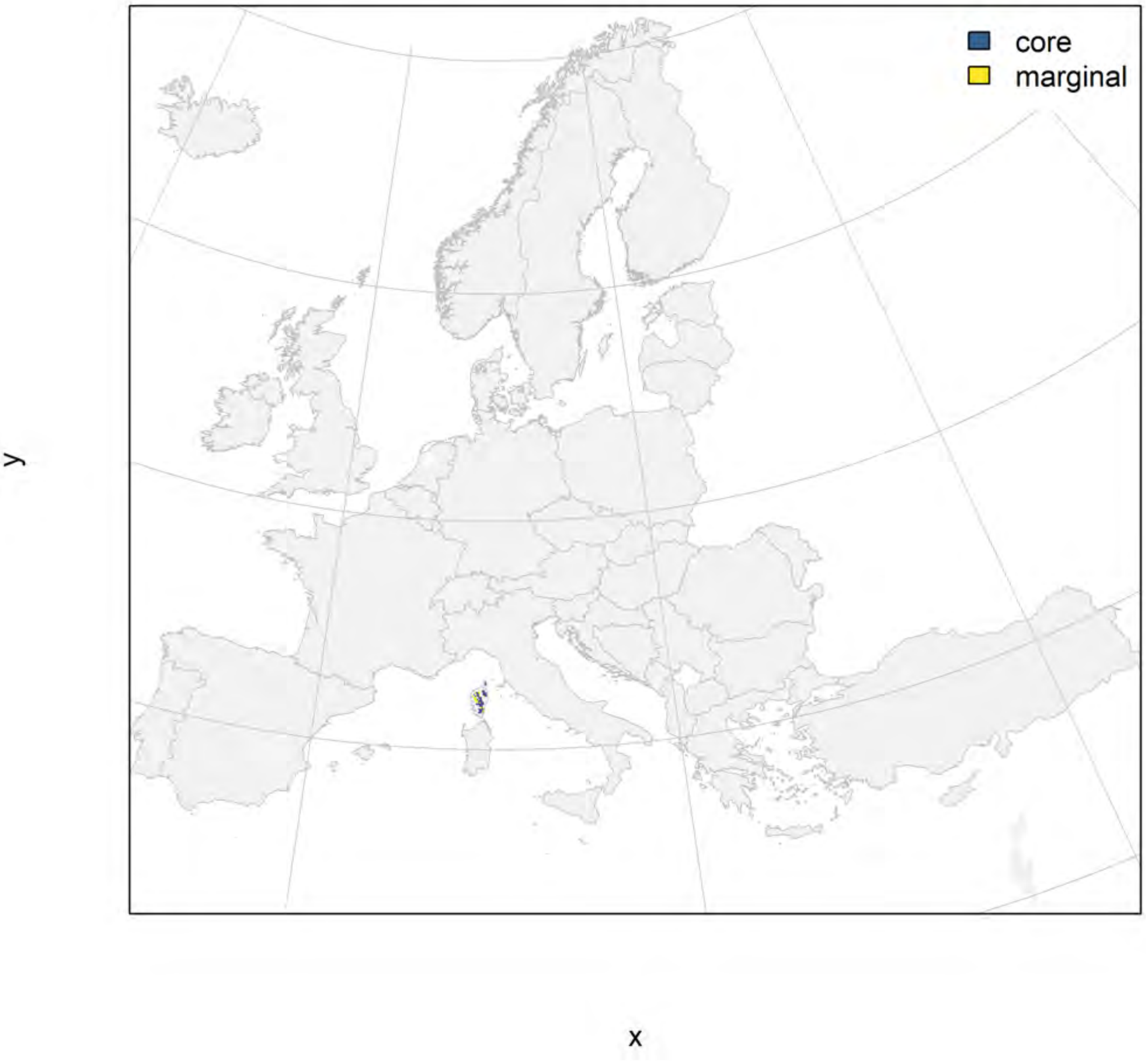

### Discoglossus montalentii_filt future marginality

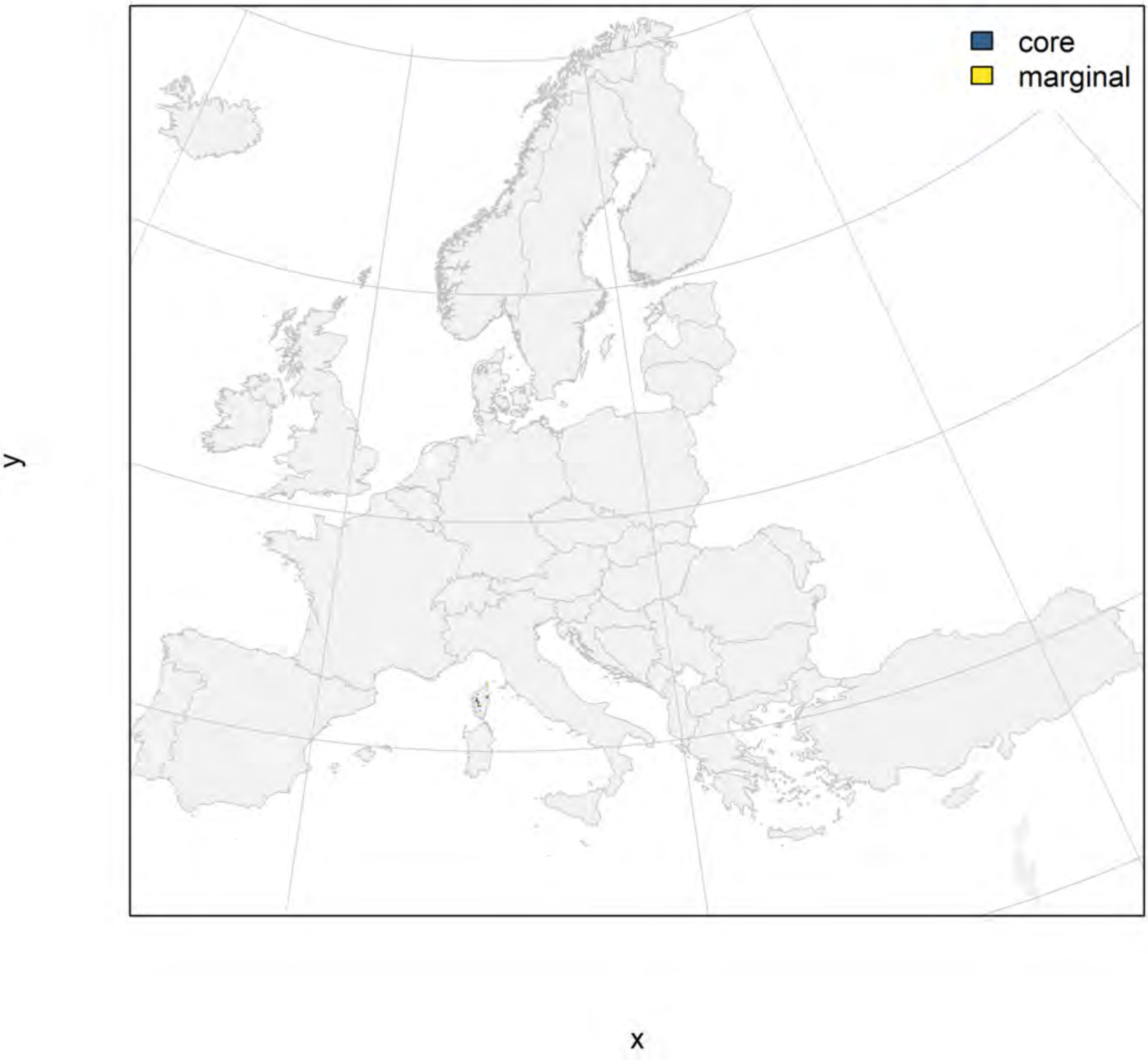

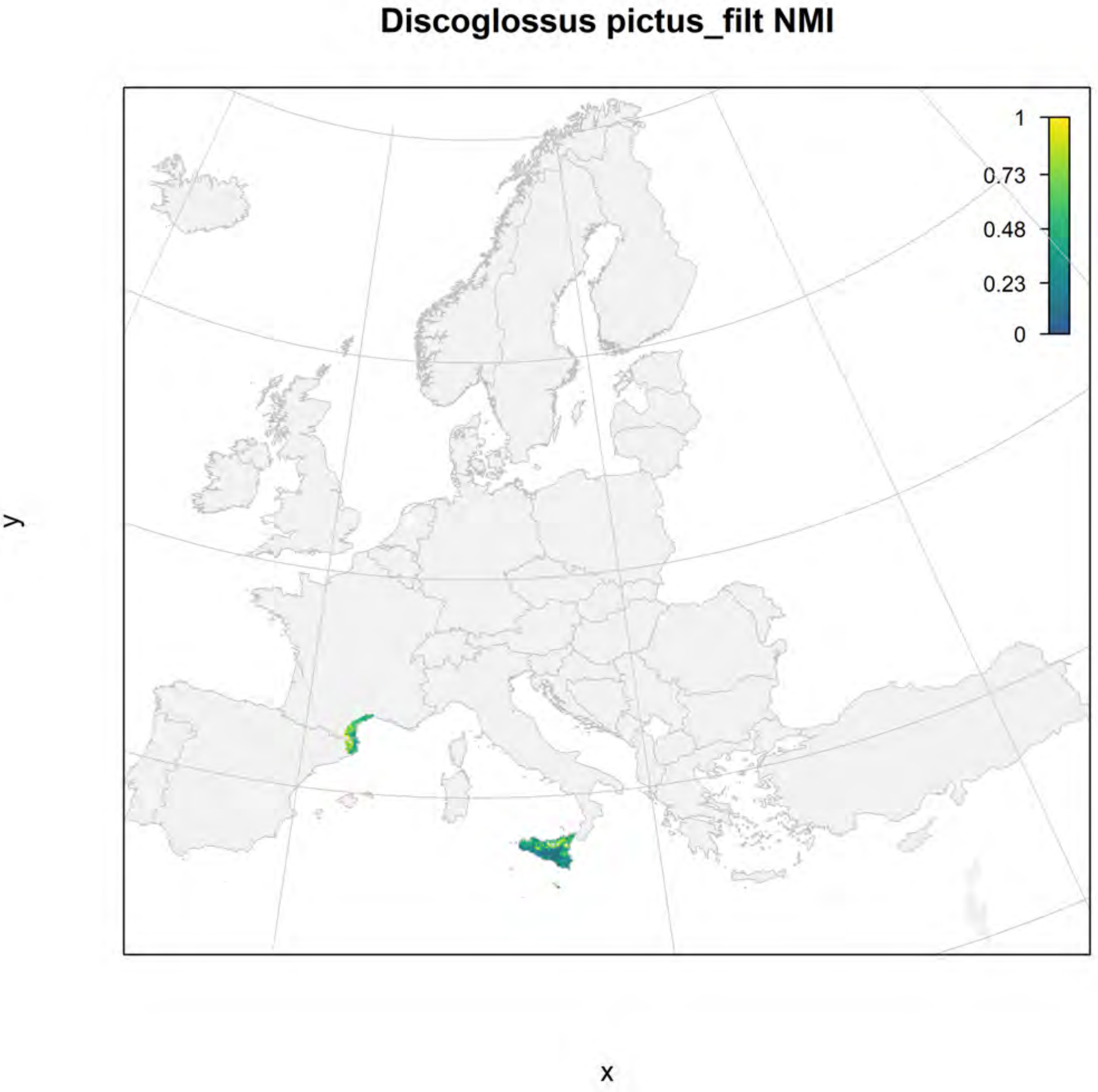
Discoglossus pictus_filt NMI
1
0.73
0.48
0.23
0
y
x

#### Discoglossus pictus_filt current marginality

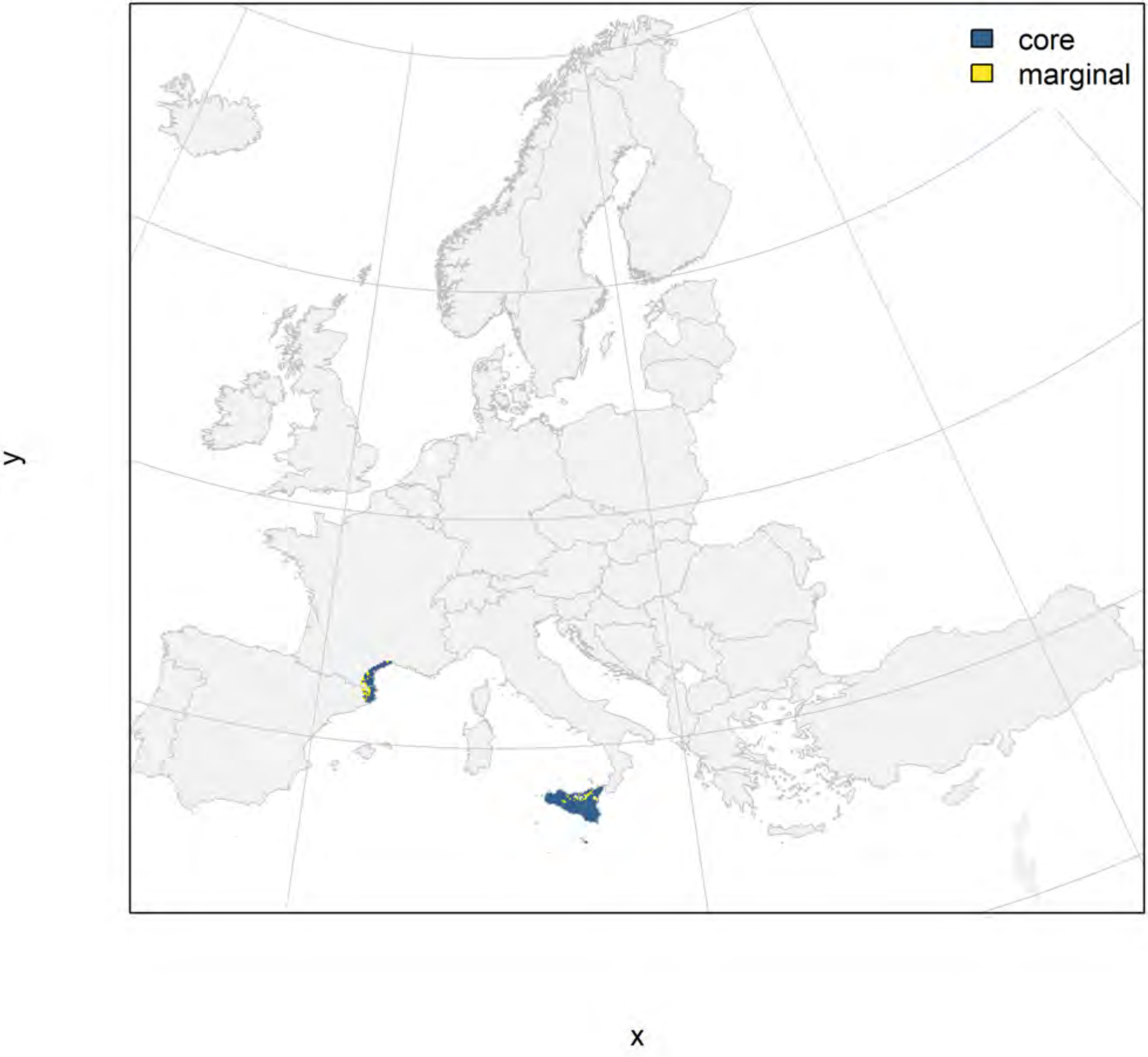

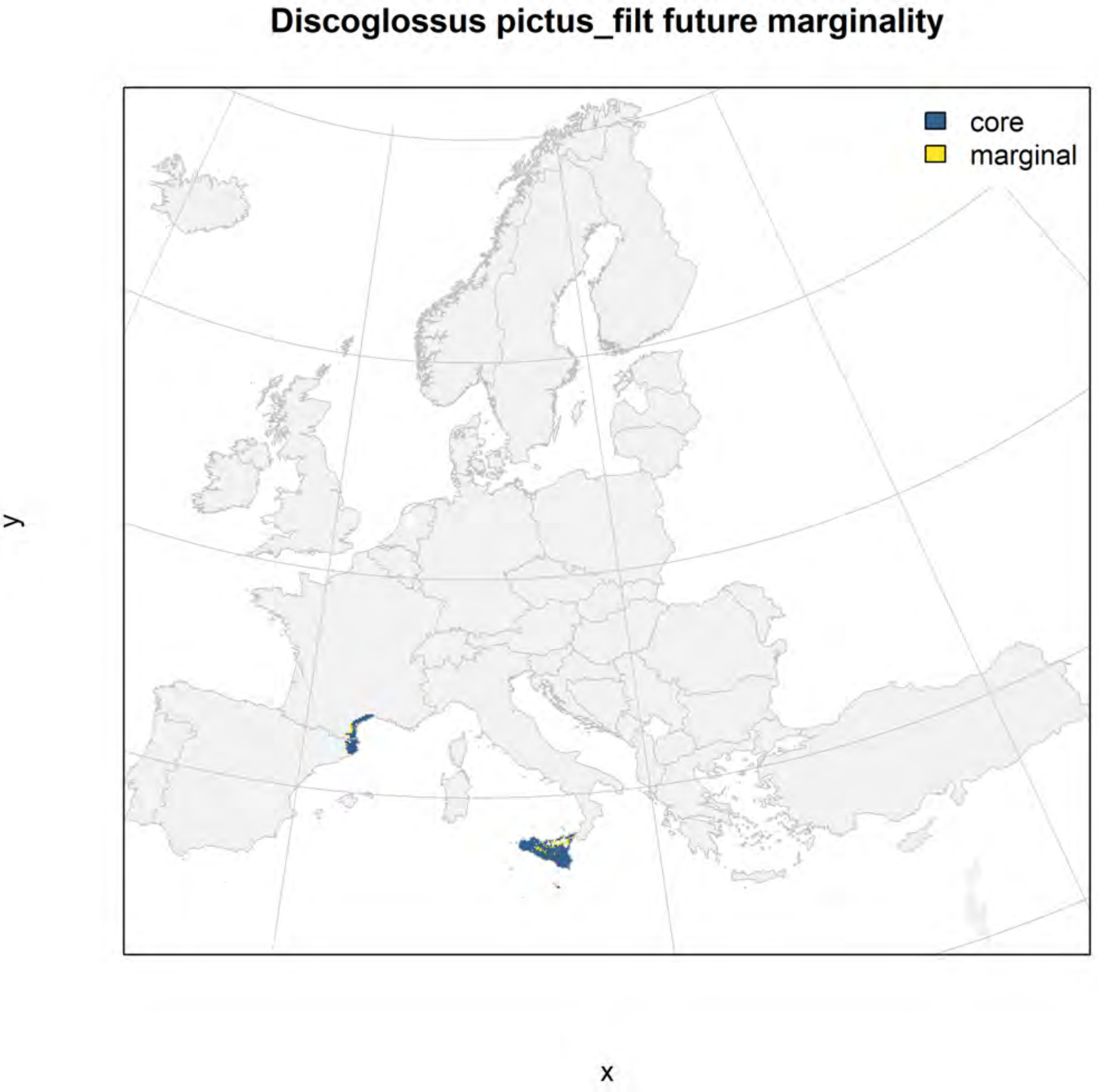
Discoglossus pictus_filt future marginality
core
marginal
y
x

#### Discoglossus sardus_filt NMI

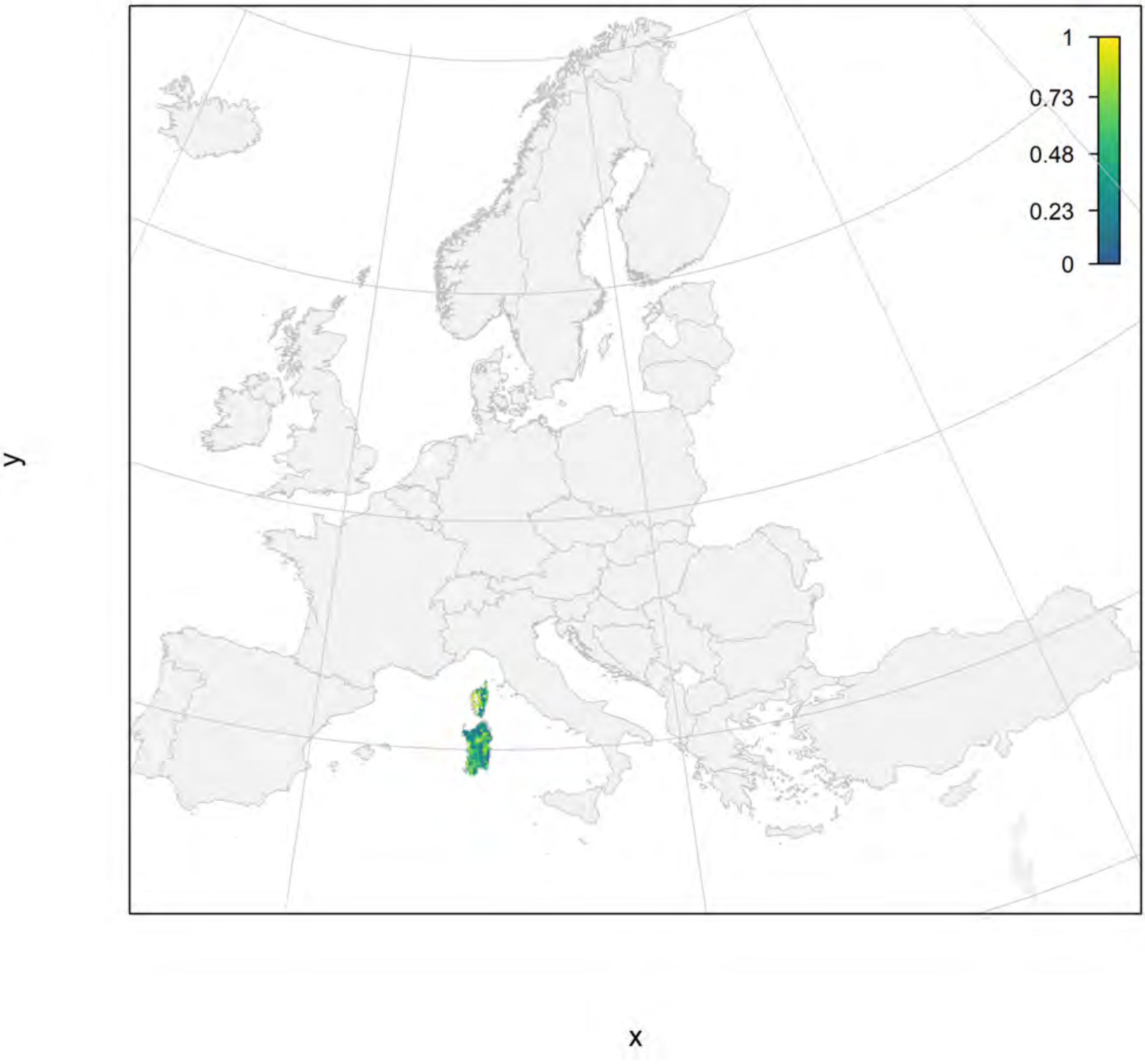

### Discoglossus sardus_filt current marginality

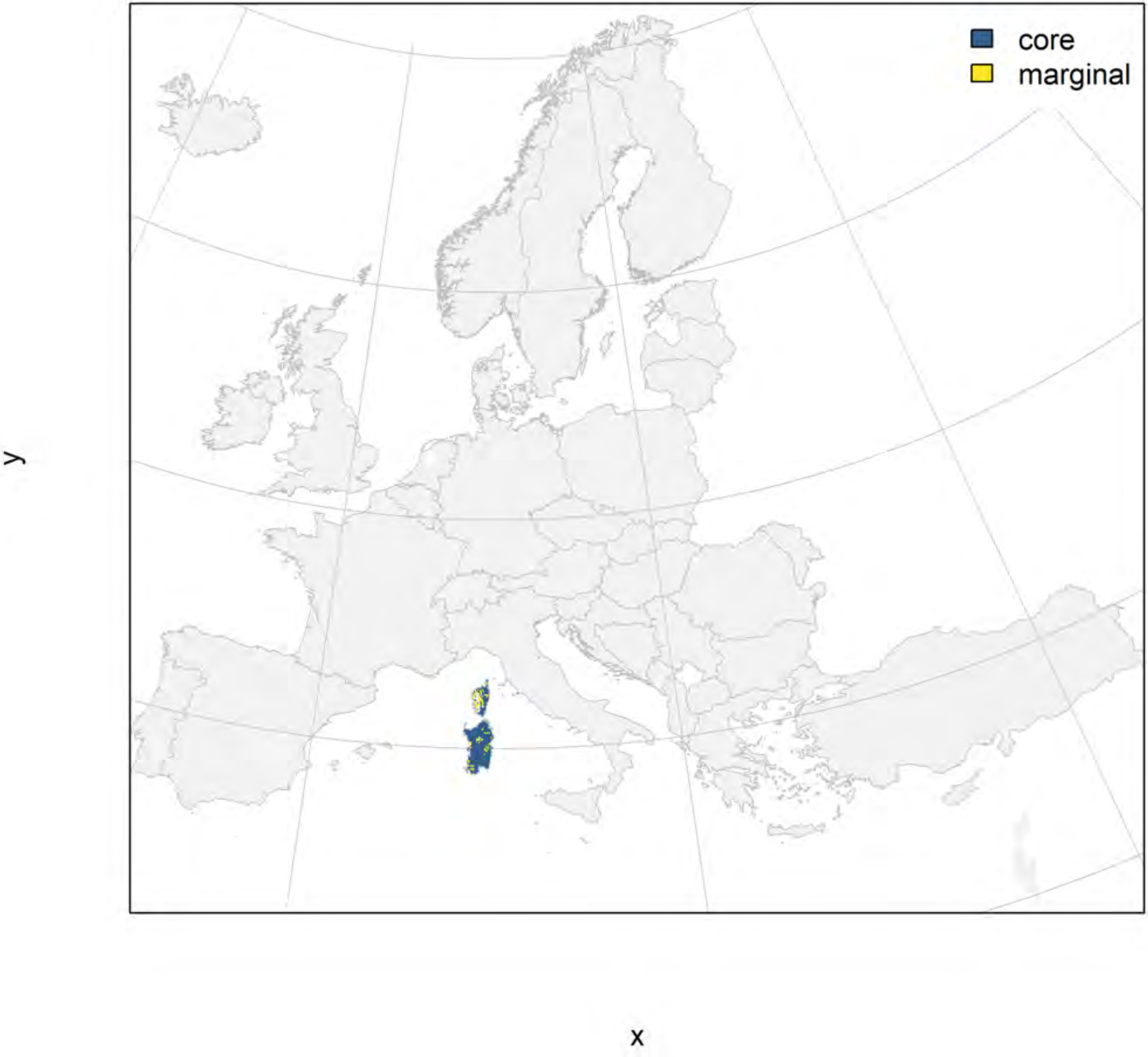

#### Discoglossus sardus_filt future marginality

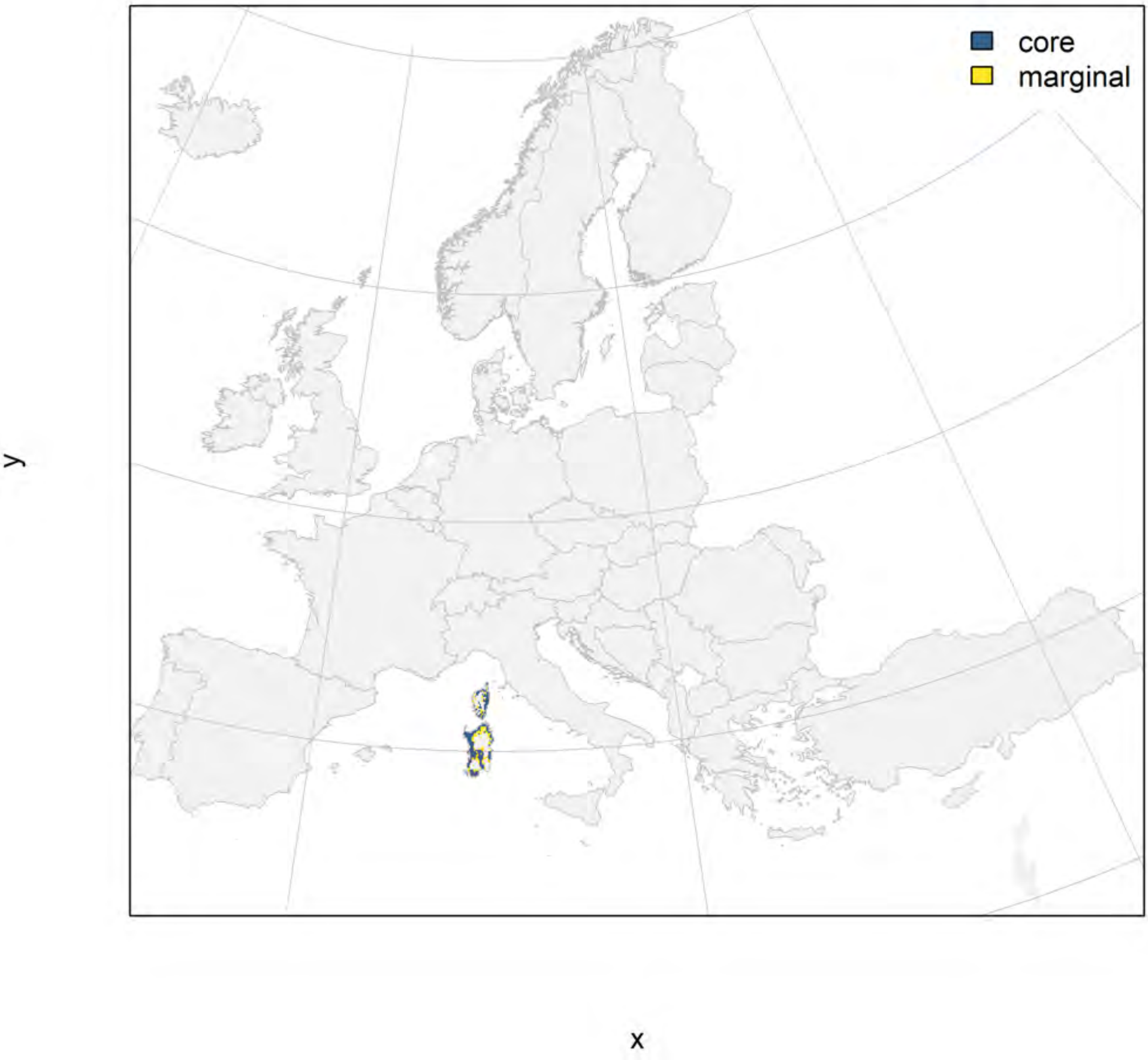

### Epidalea calamita_filt NMI

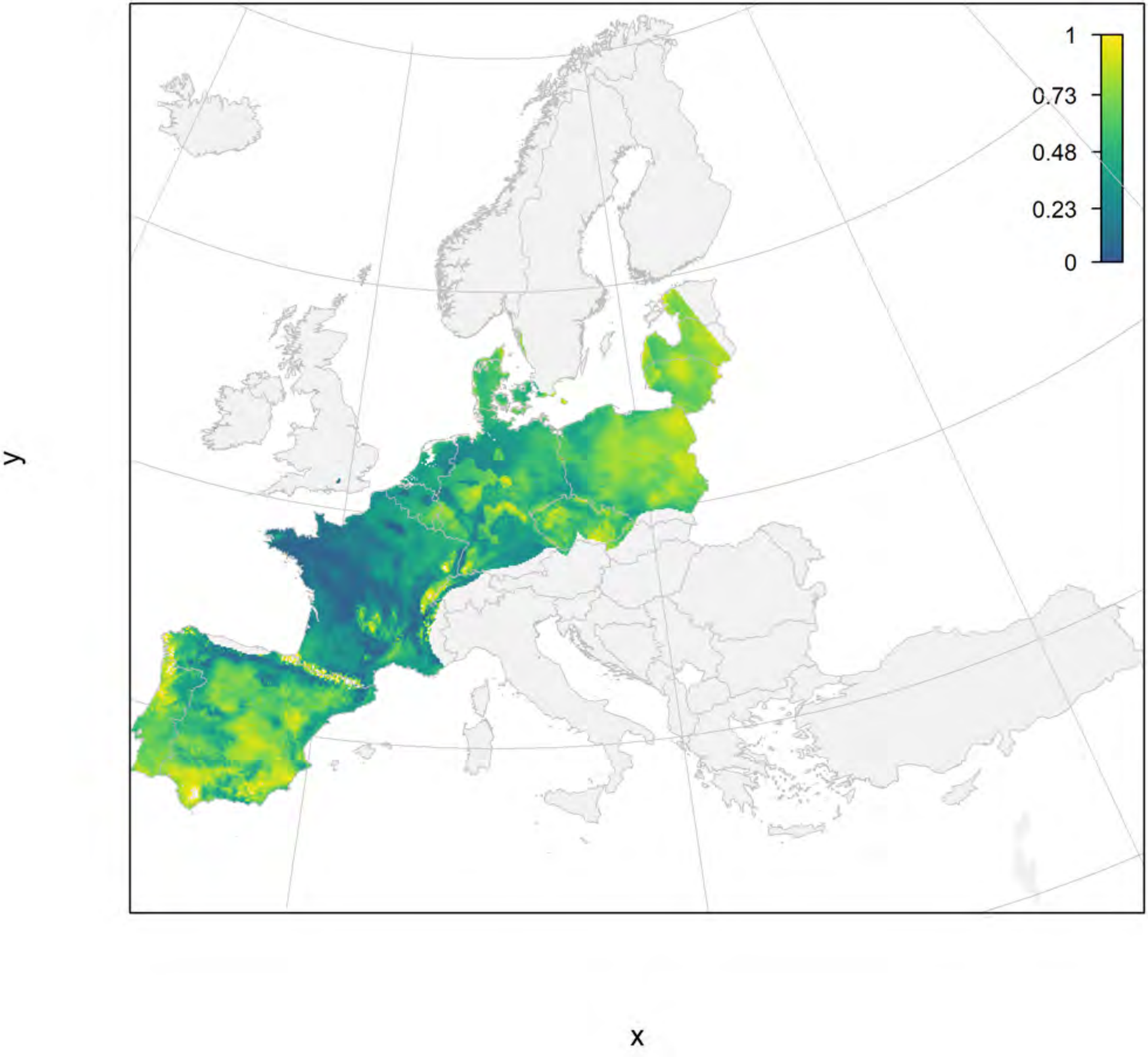

### Epidalea calamita_filt current marginality

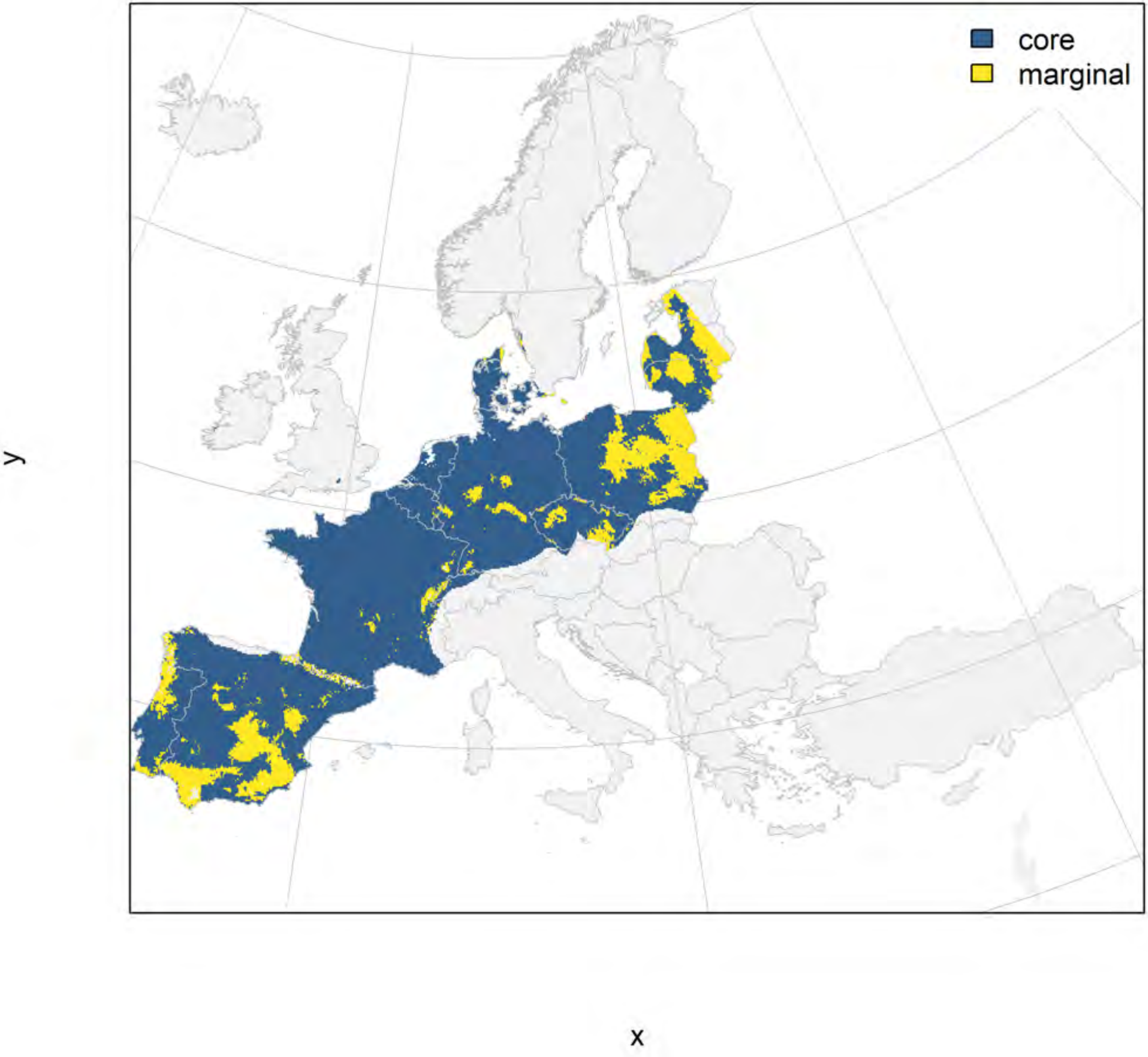

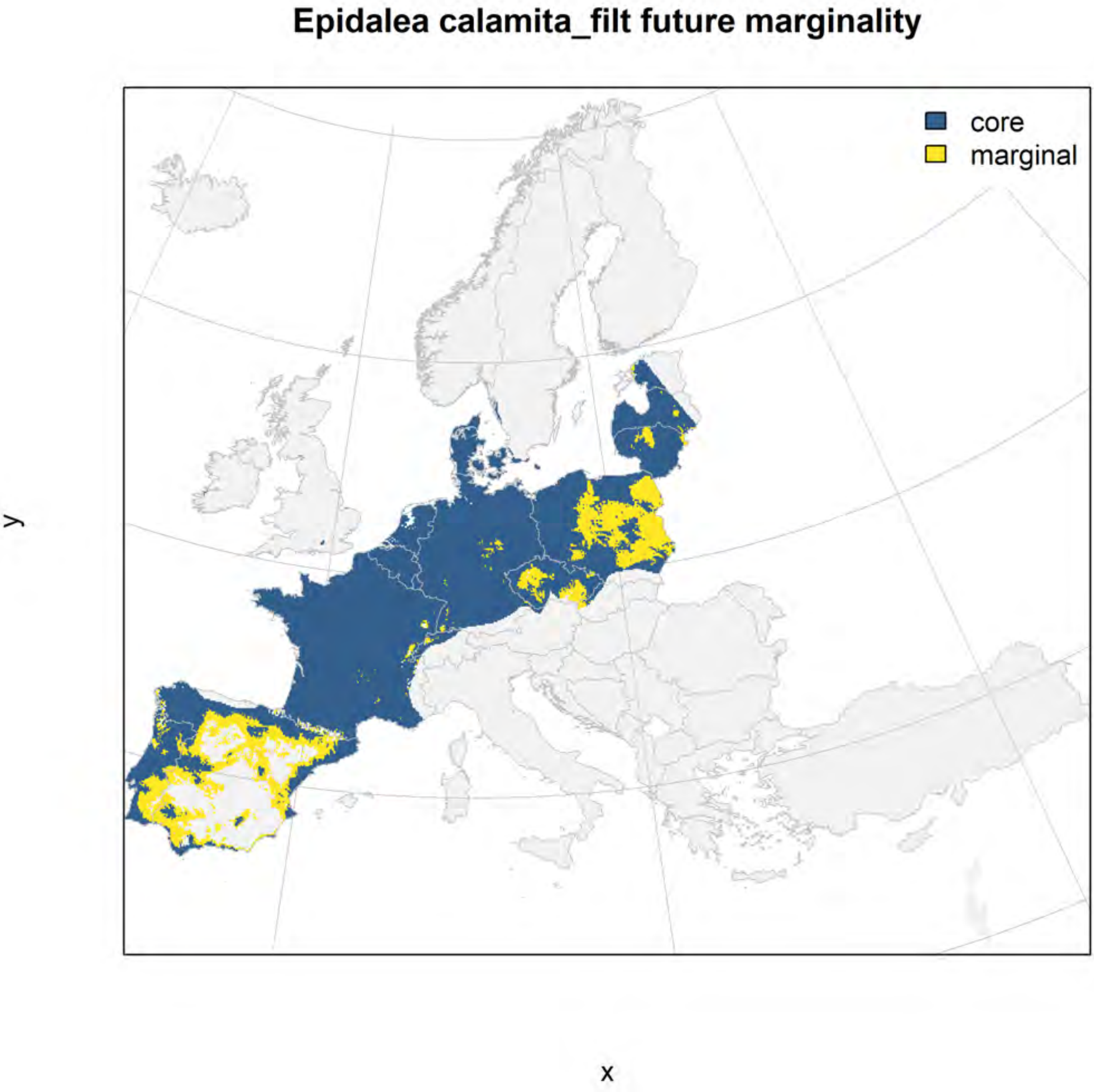
Epidalea calamita_filt future marginality
core
marginal
y
x

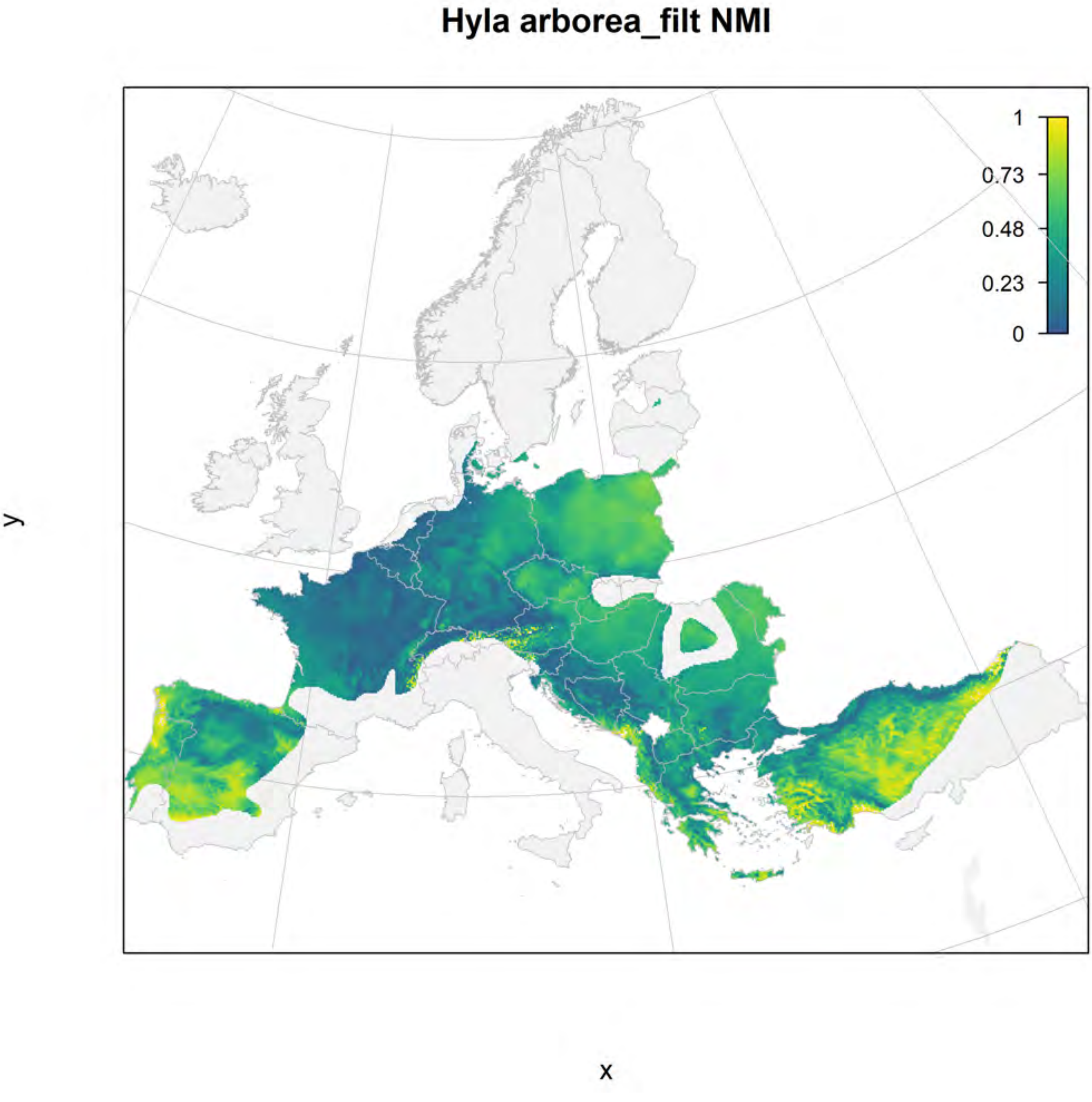

Hyla arborea_filt NMI
1
0.73
0.48
0.23
0
y
x

Hyla arborea_filt current marginality

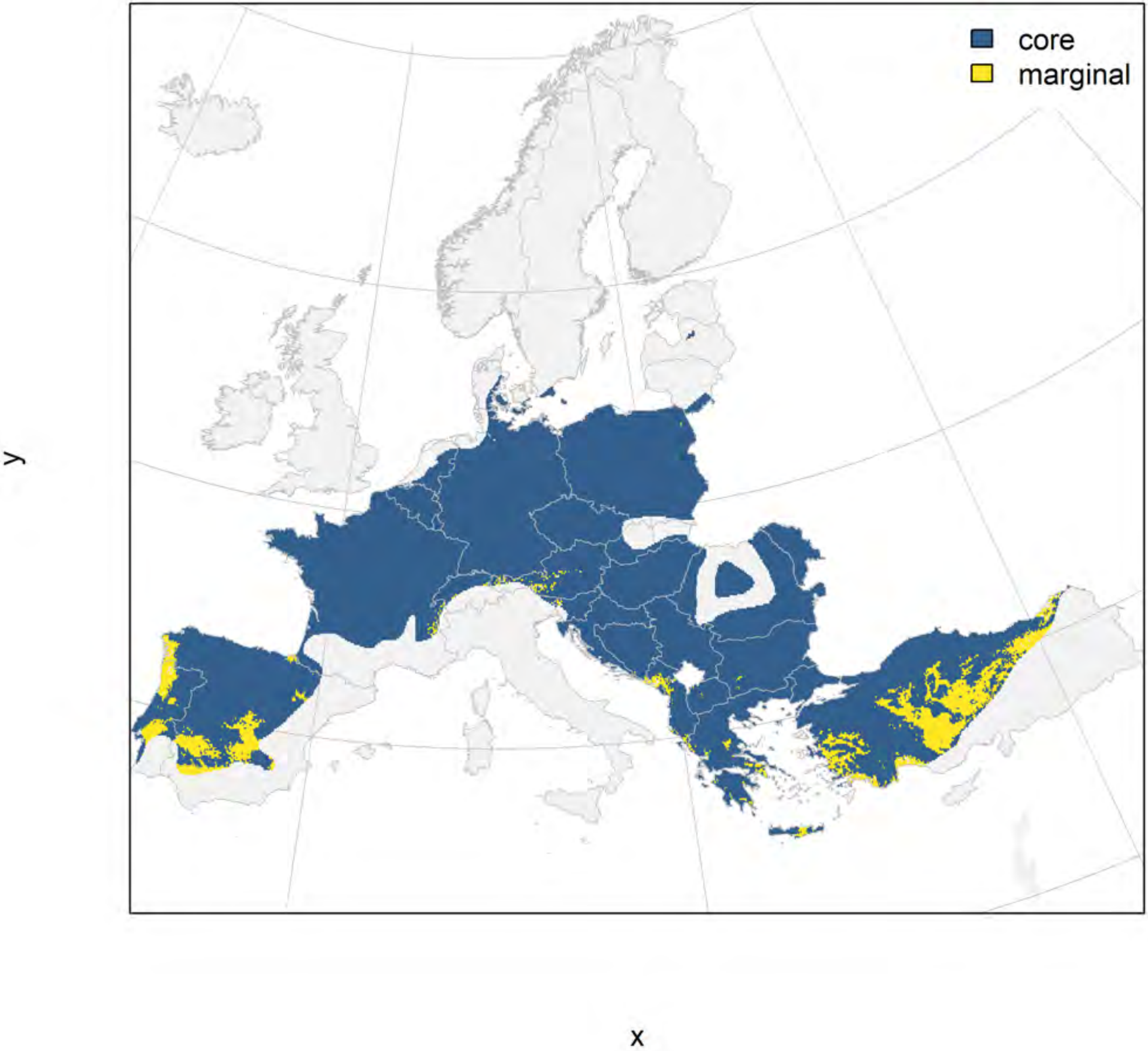

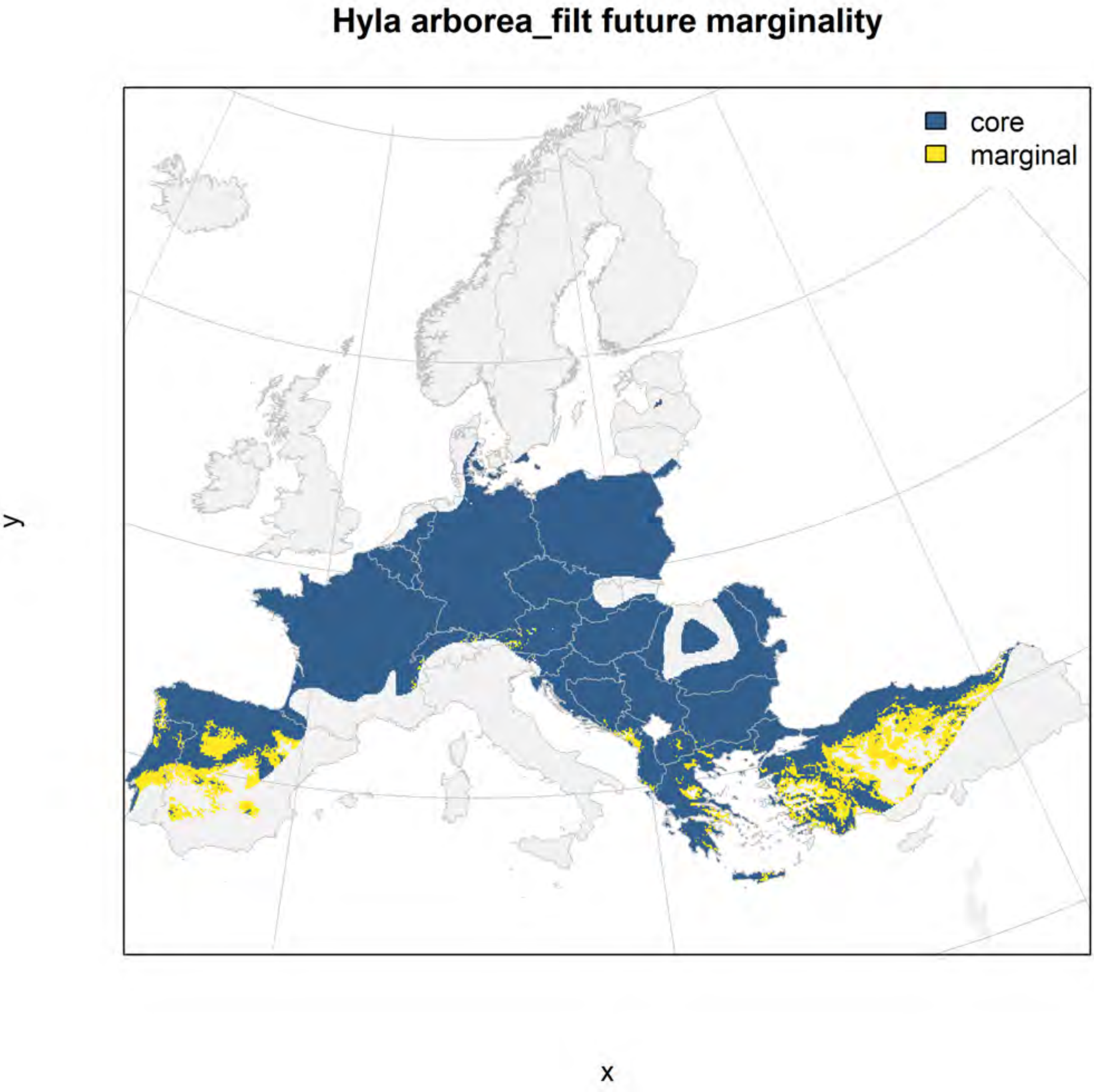

Hyla arborea_filt future marginality
core
marginal
y
x

### Hyla intermedia_filt NMI

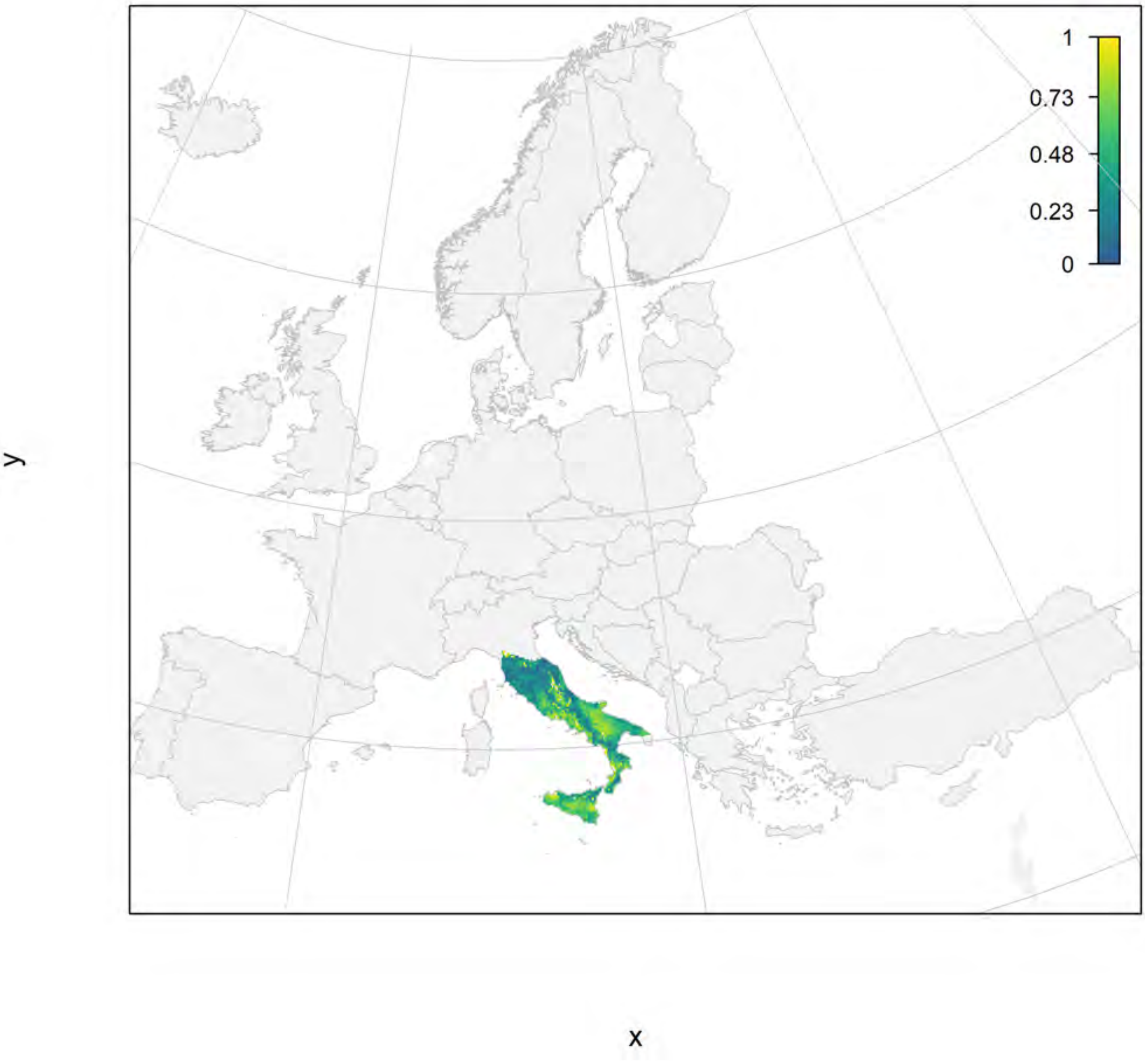

#### Hyla intermedia_filt current marginality

y

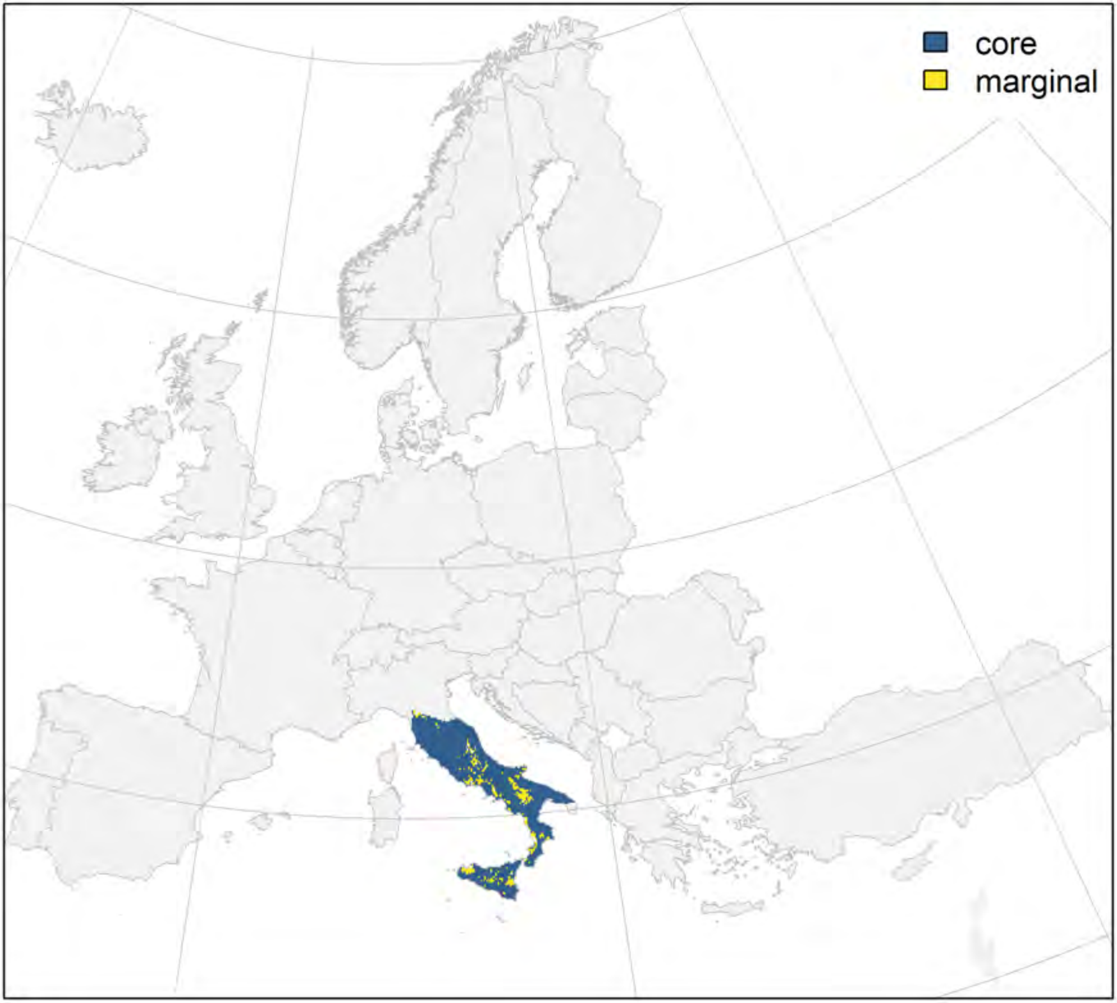


x

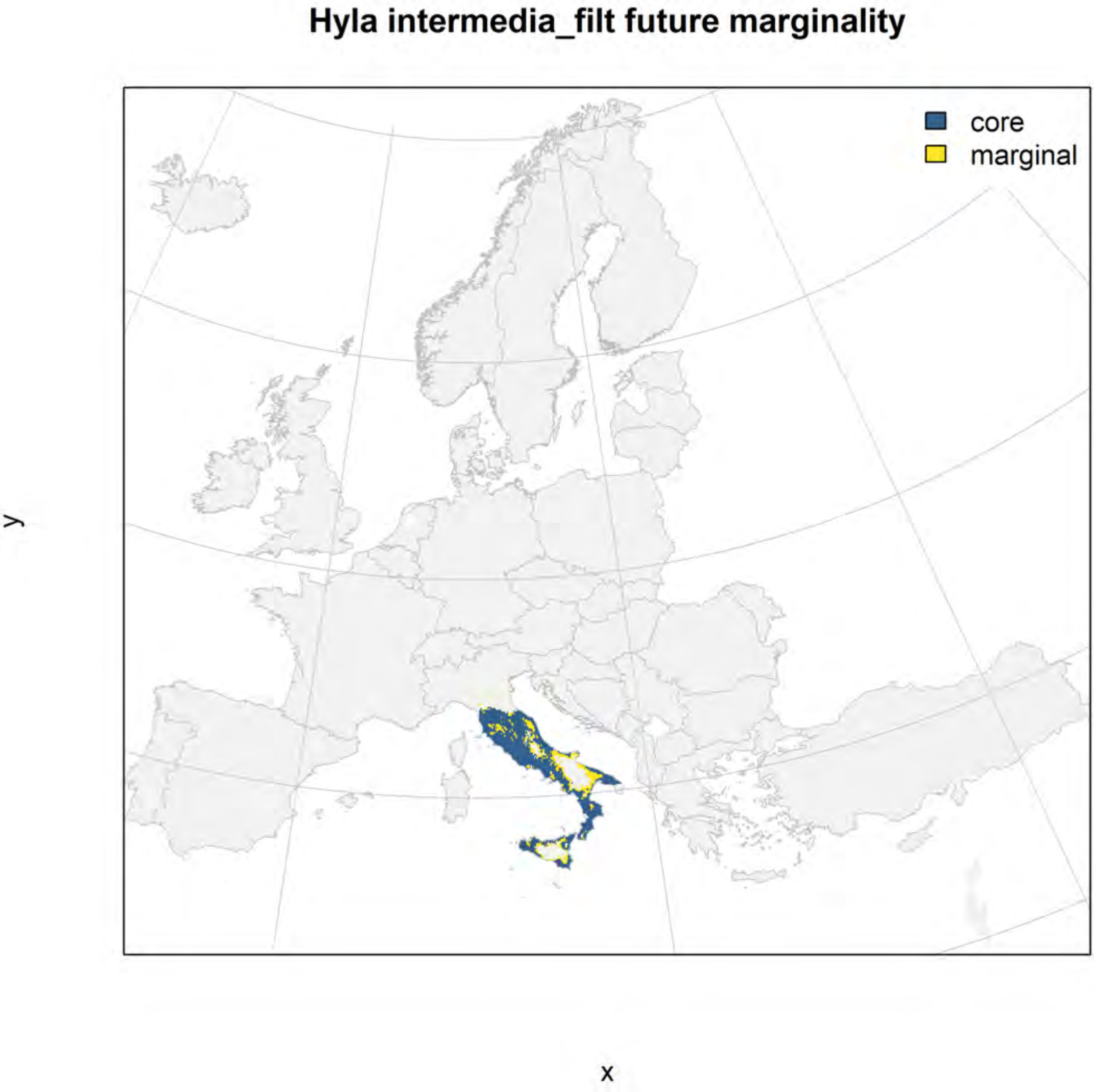
Hyla intermedia_filt future marginality
core
marginal
y
x

#### Hyla meridionalis_filt NMI

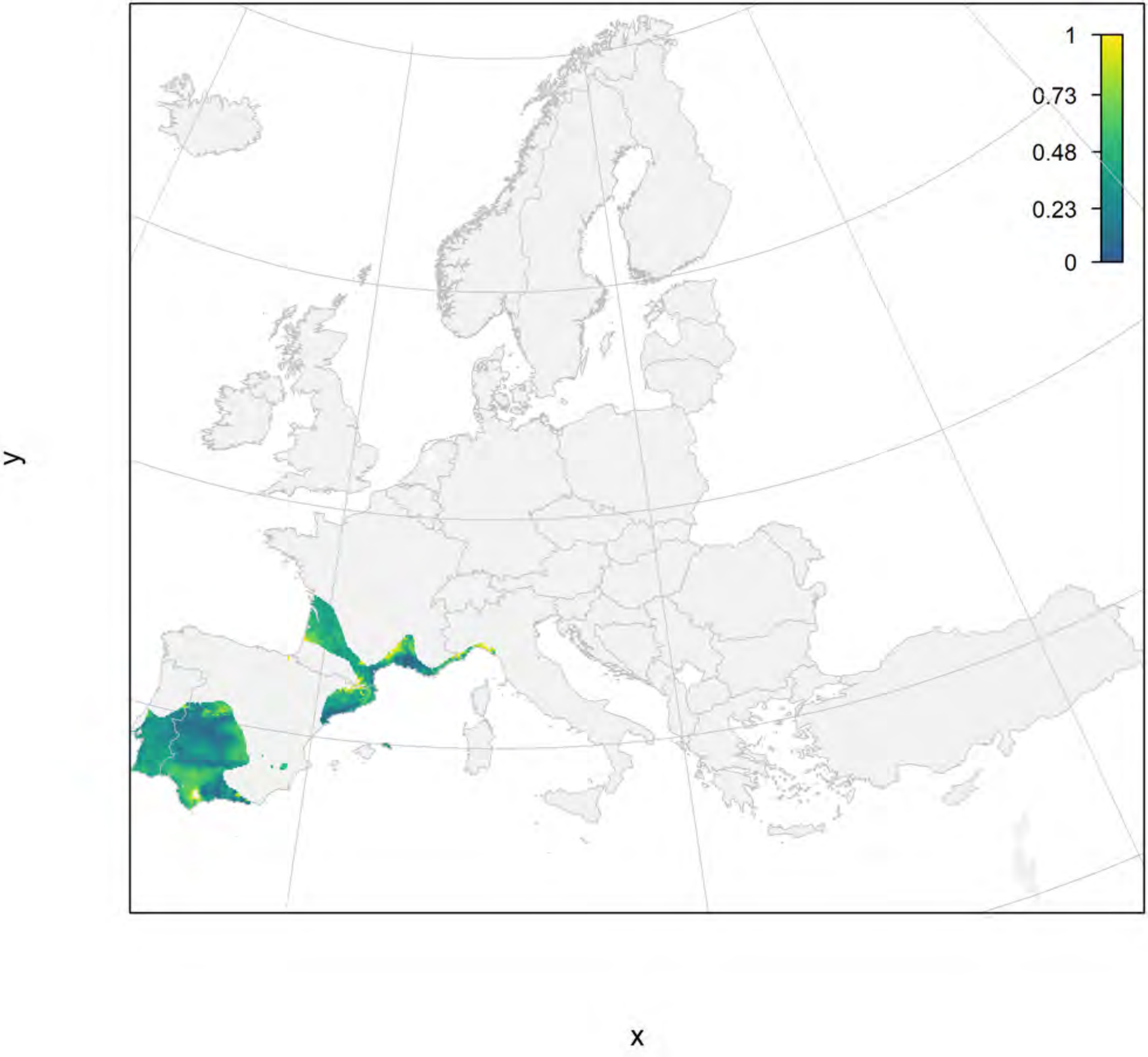

#### Hyla meridionalis_filt current marginality

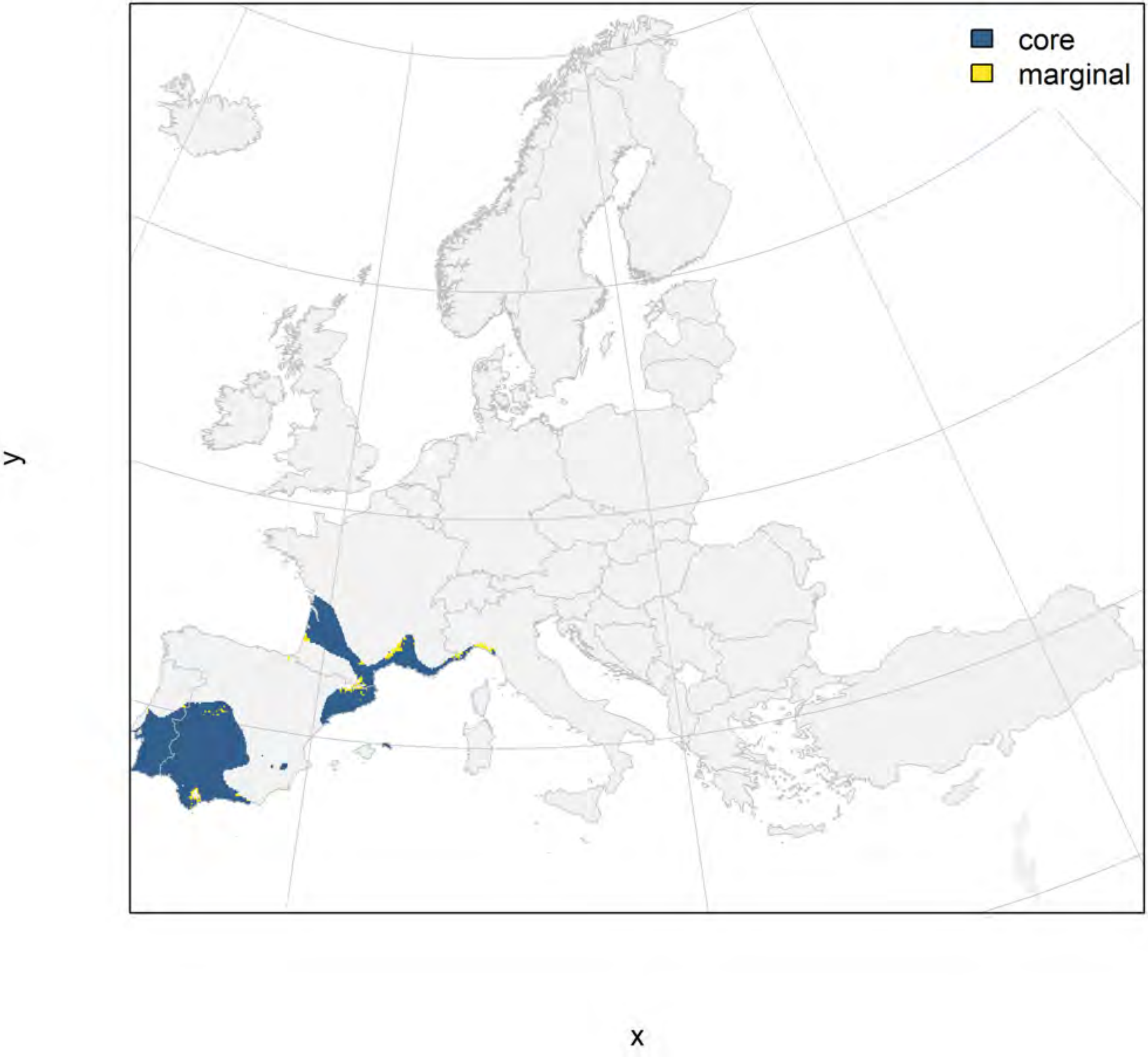

**Hyla meridionalis_filt future marginality**

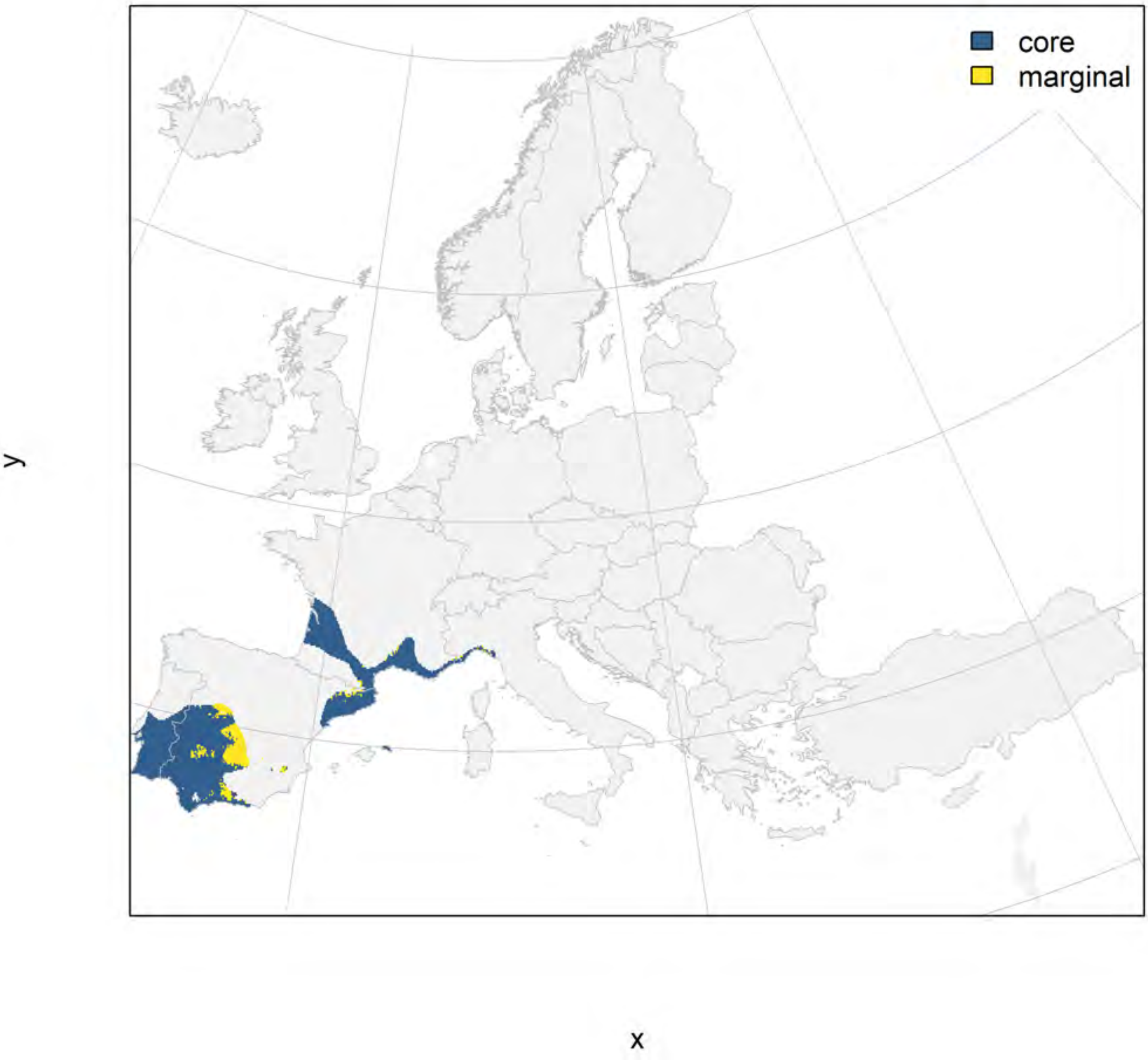

#### Hyla sarda_filt NMI

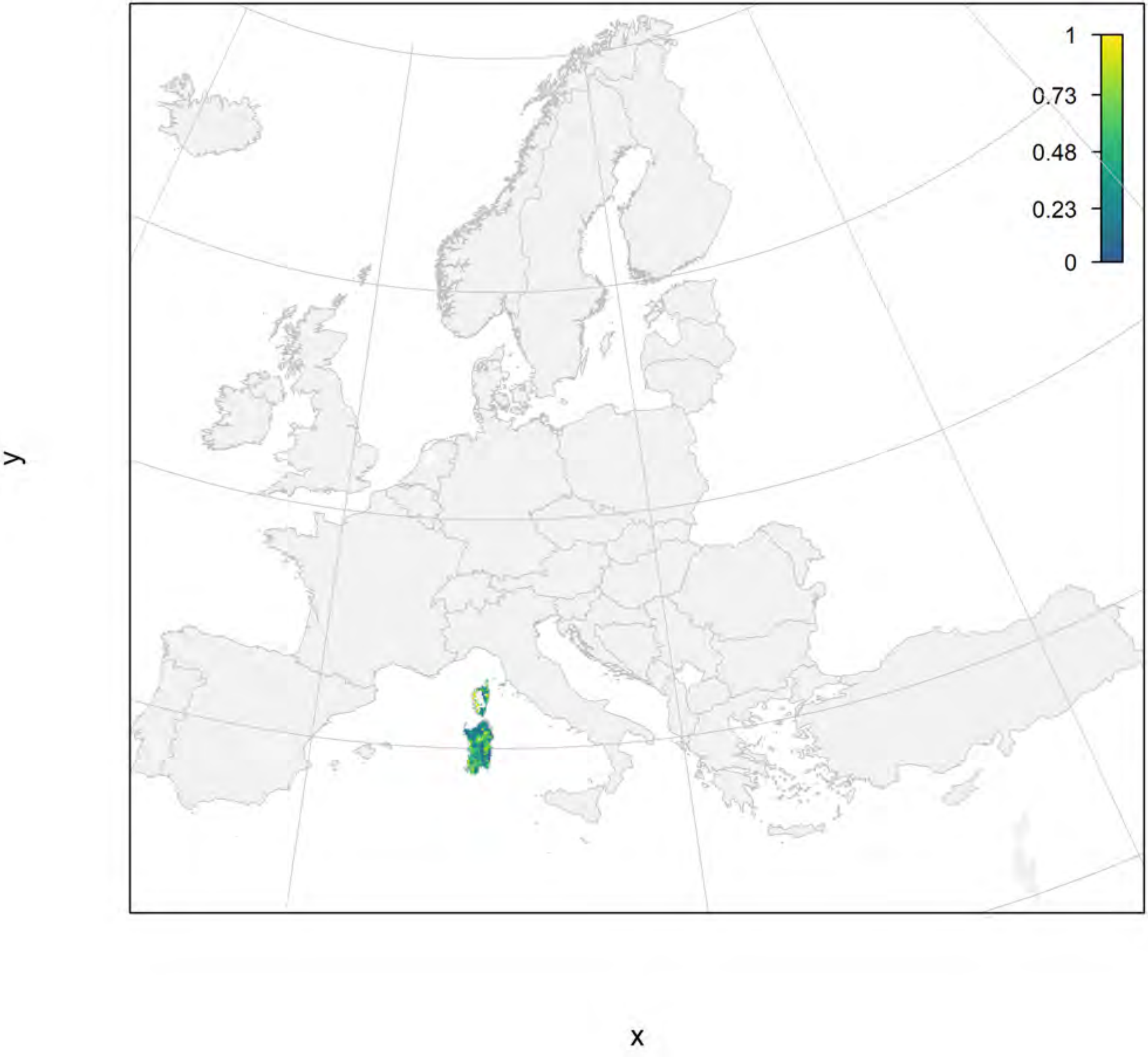

#### Hyla sarda_filt current marginality

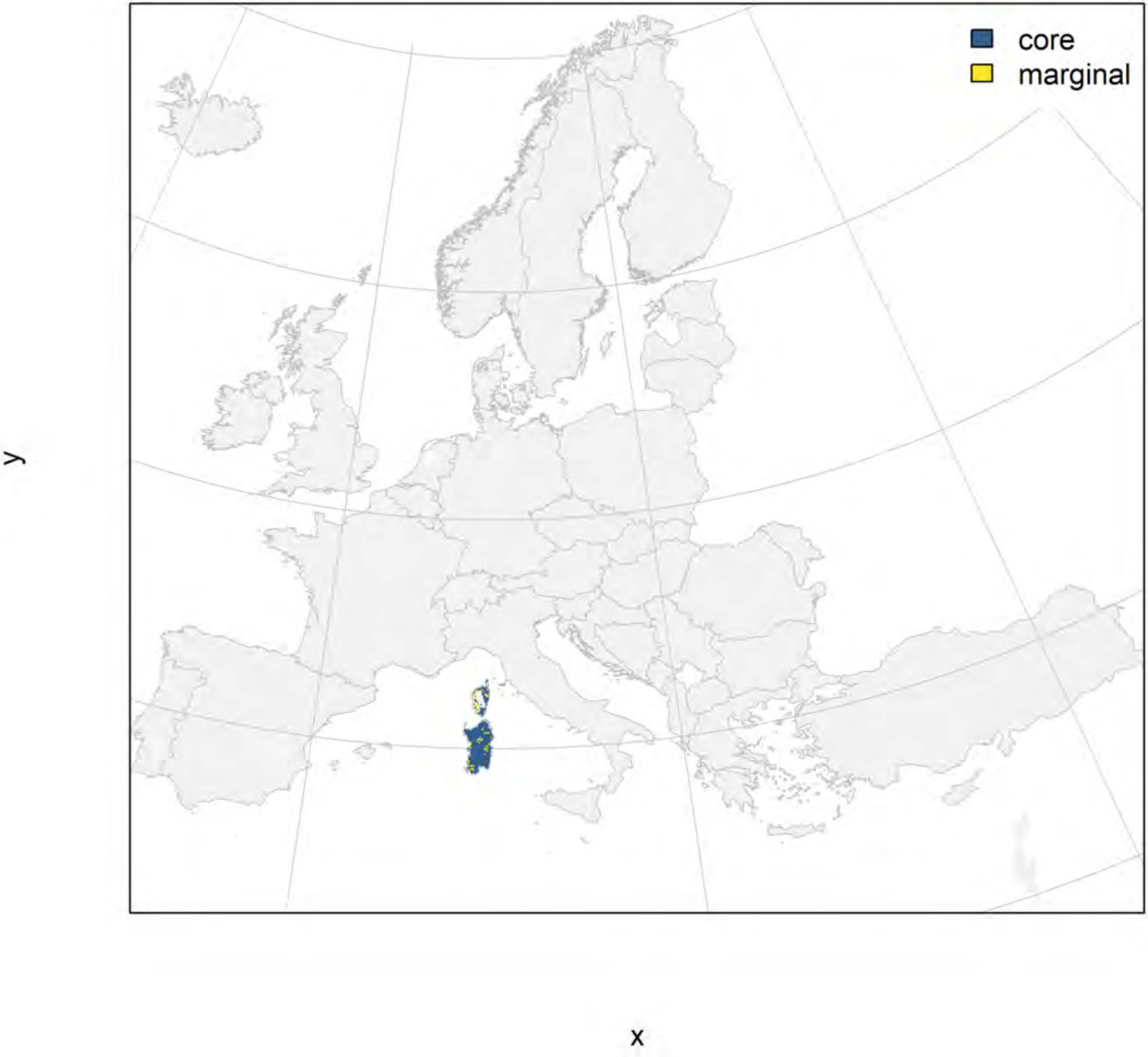

#### Hyla sarda_filt future marginality

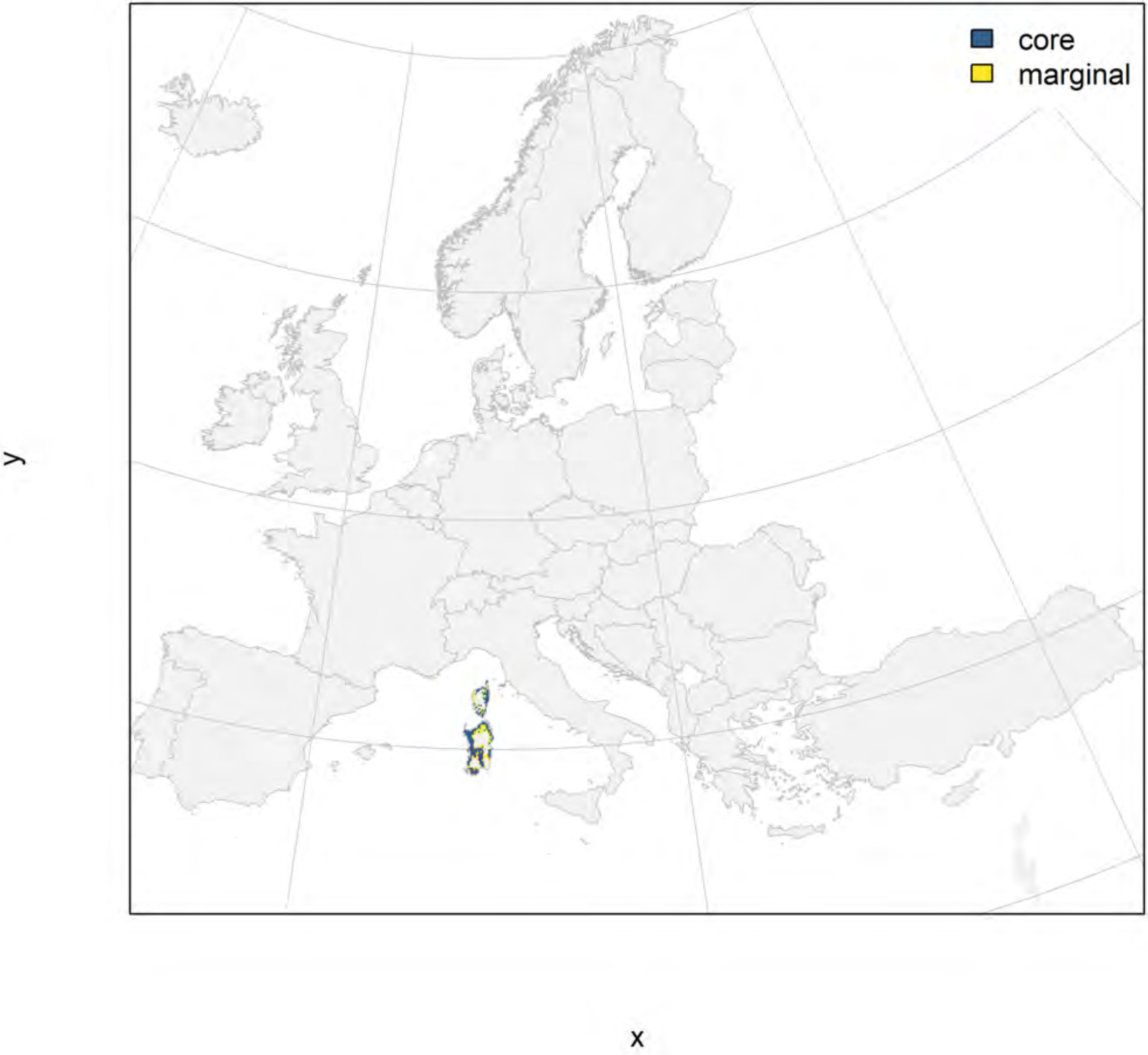

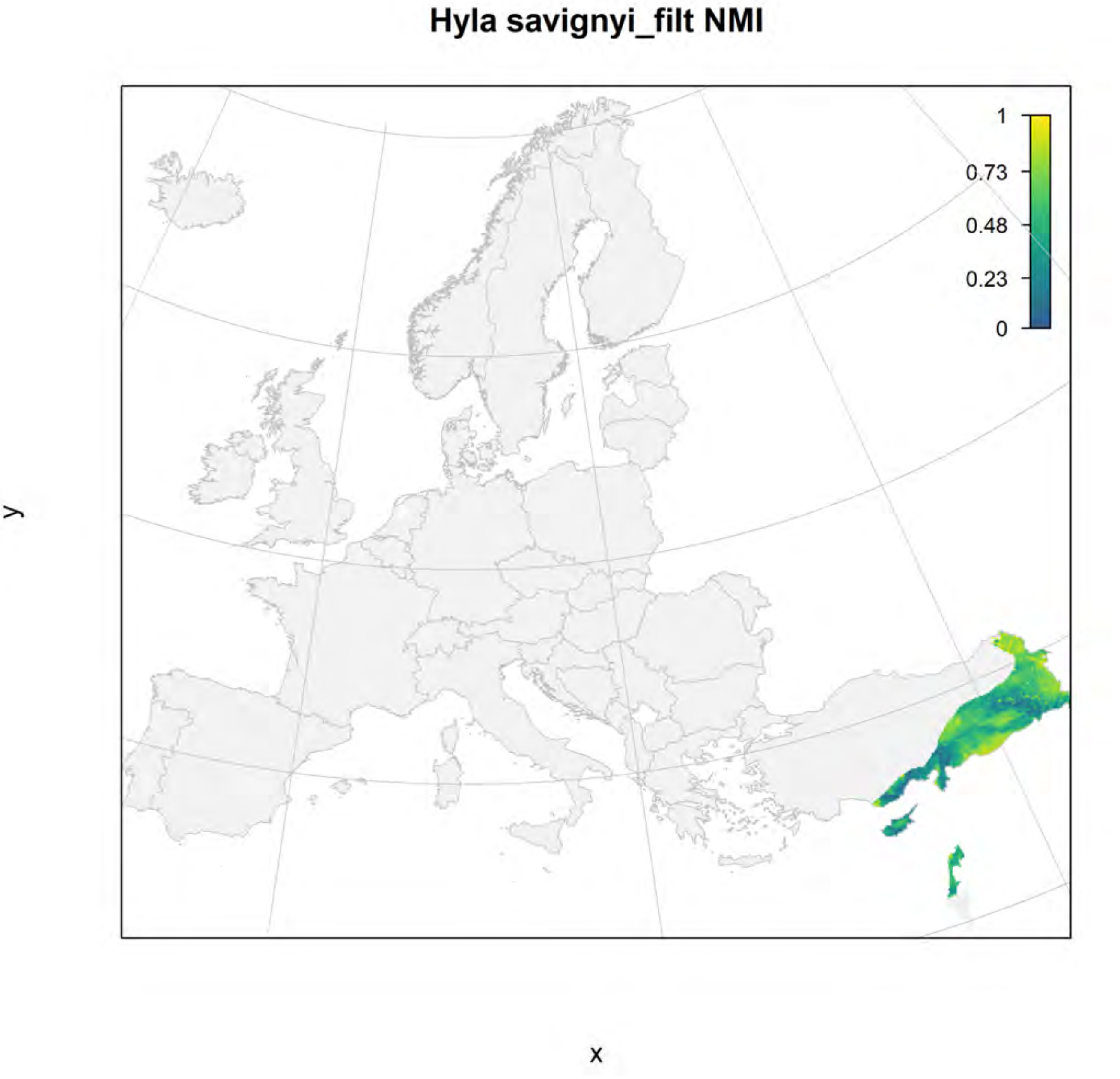
Hyla savignyi_filt NMI
1
0.73
0.48
0.23
0
y
x

#### Hyla savignyi_filt current marginality

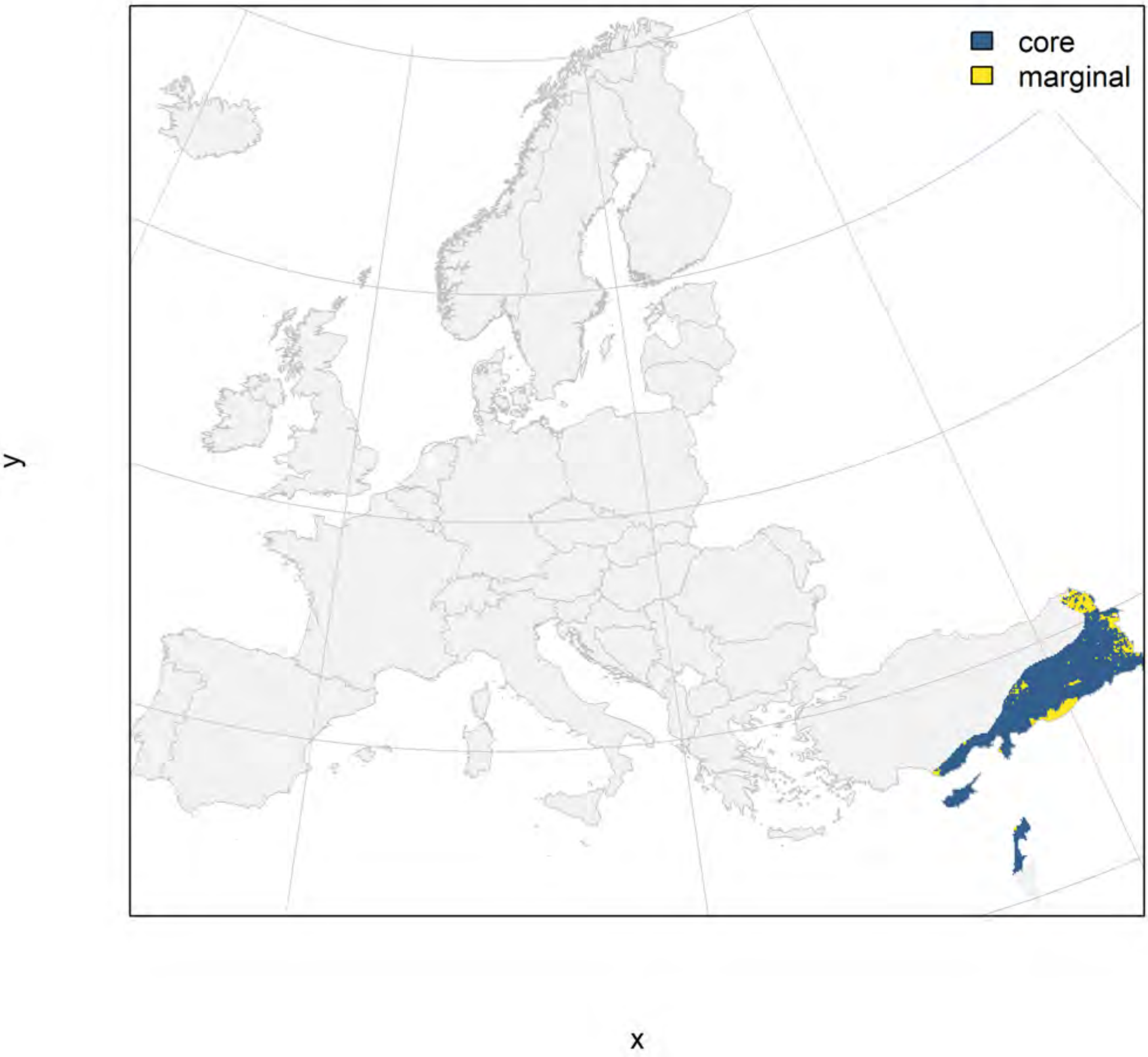

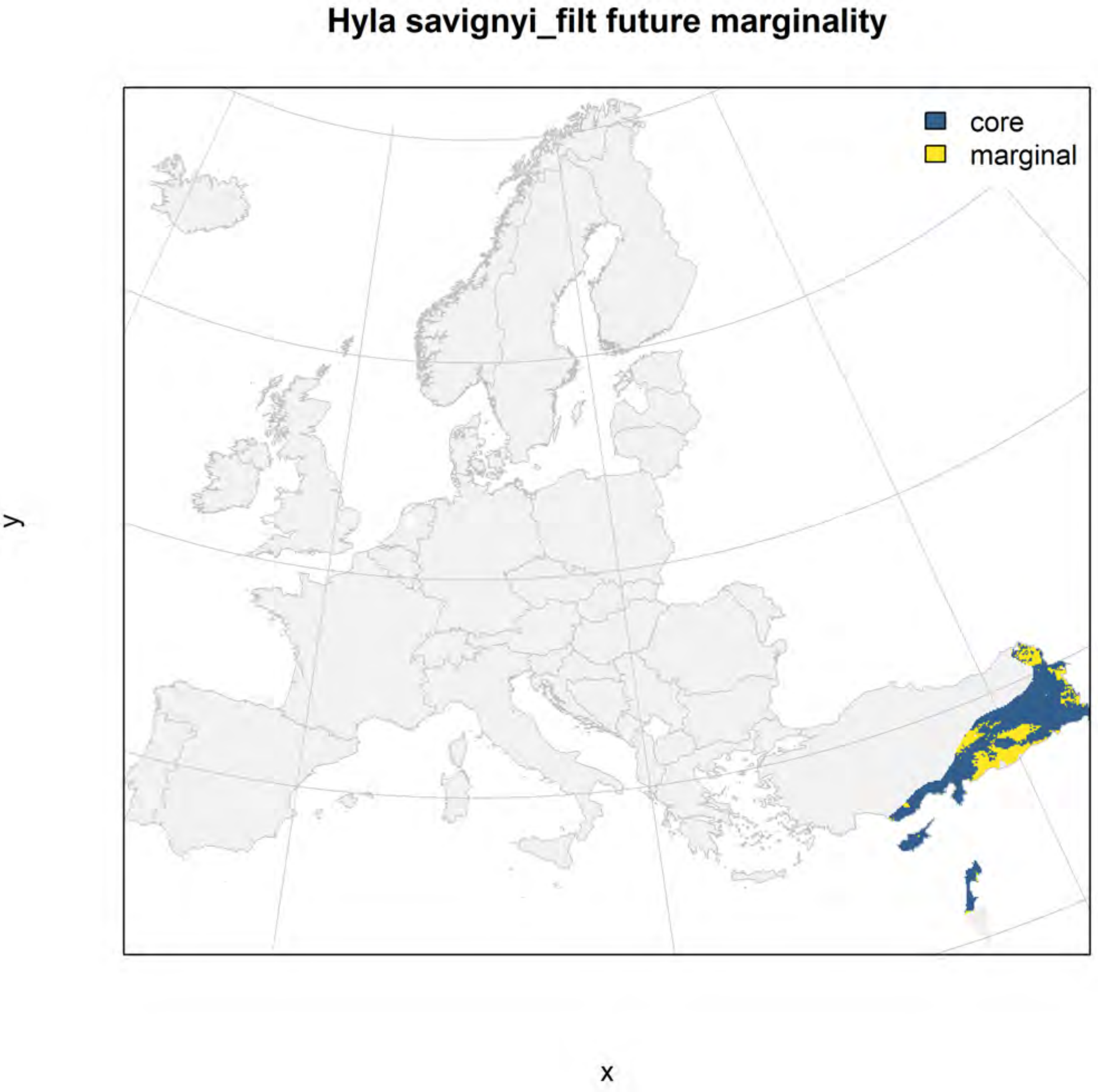
Hyla savignyi_filt future marginality
core
marginal
y
x

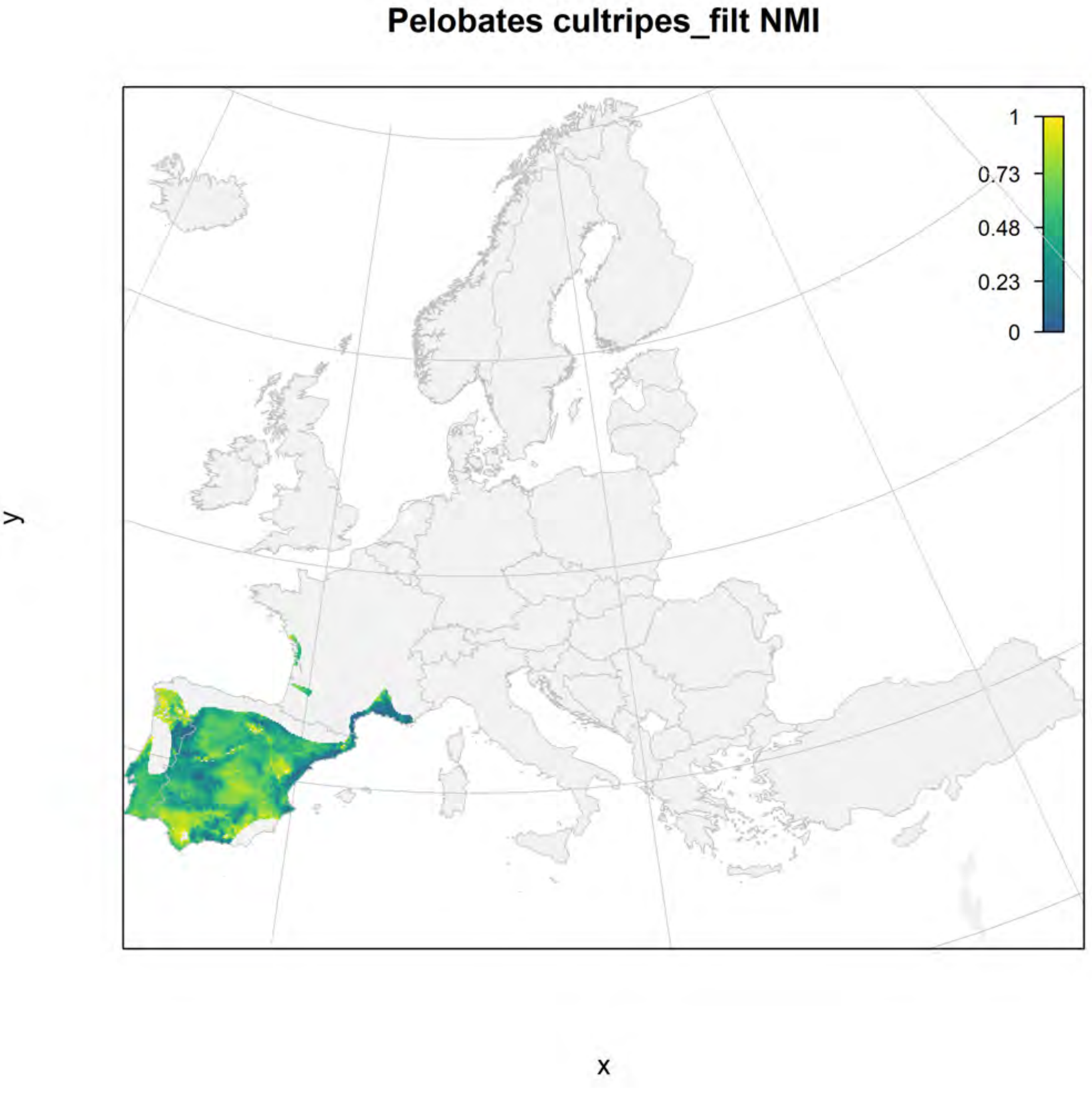

Pelobates cultripes_filt NMI
1
0.73
0.48
0.23
0
y
x

#### Pelobates cultripes_filt current marginality

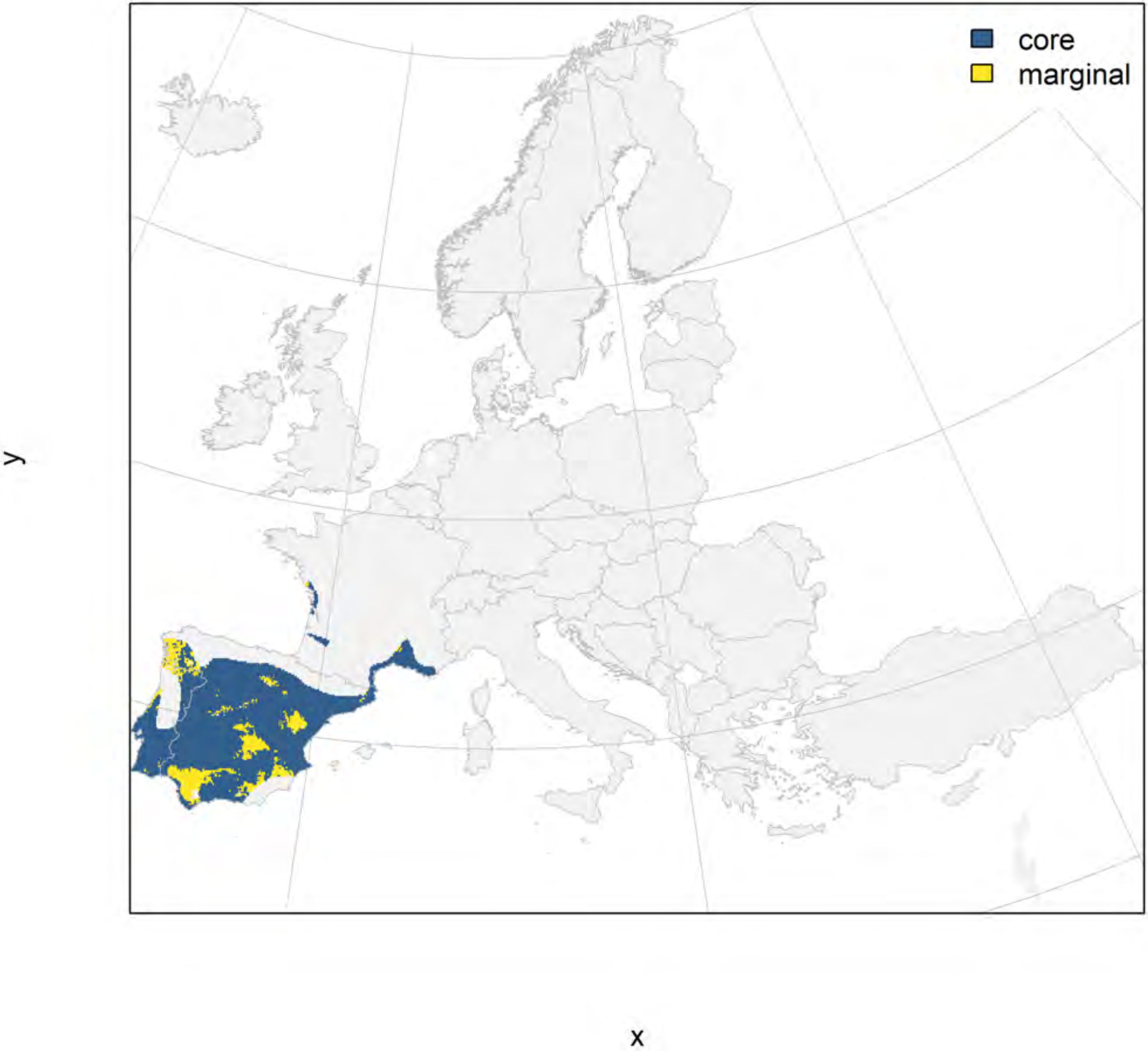

#### Pelobates cultripes_filt future marginality

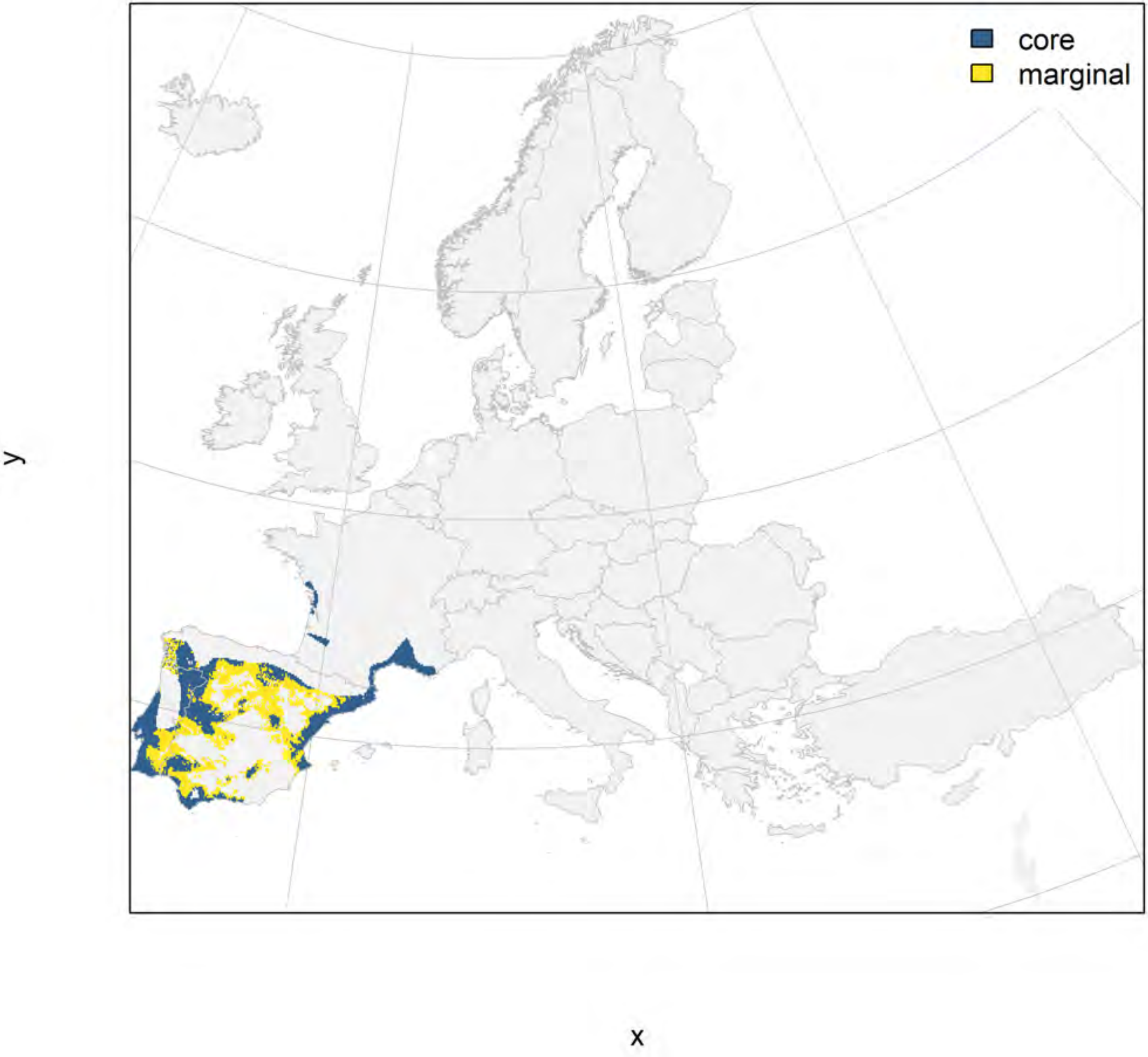

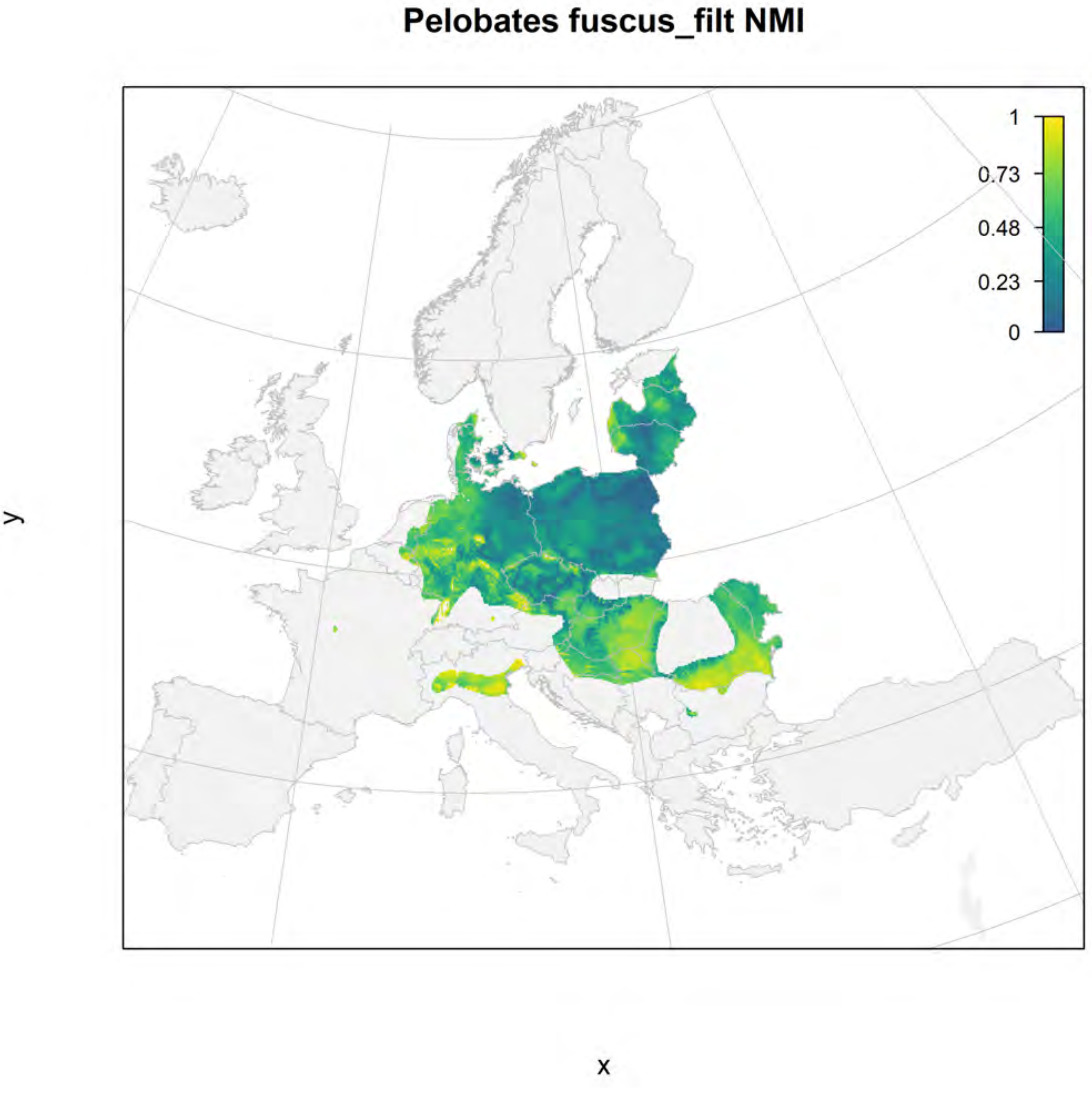

Pelobates fuscus_filt NMI
1
0.73
0.48
0.23
0
y
x

#### Pelobates fuscus_filt current marginality

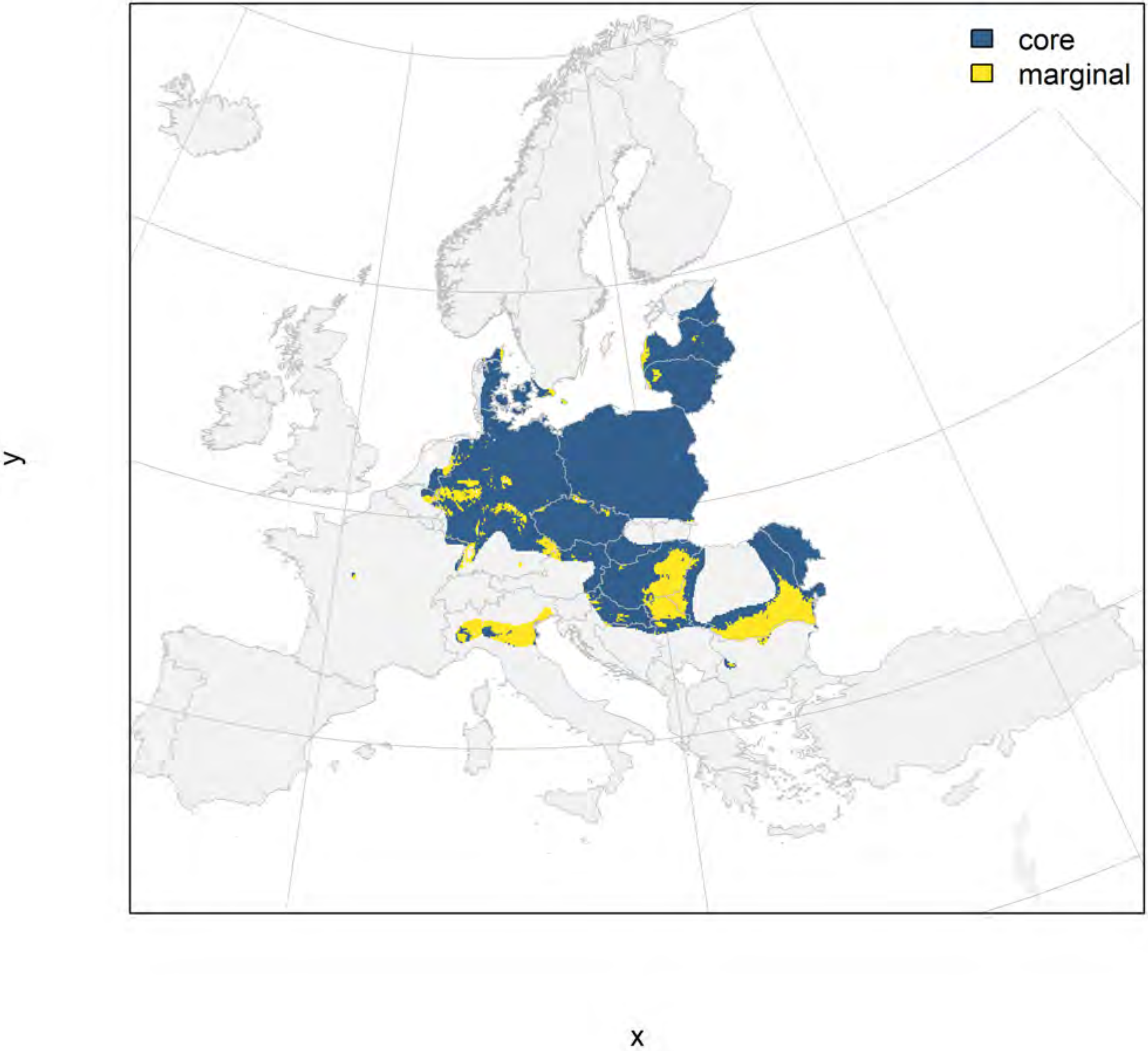

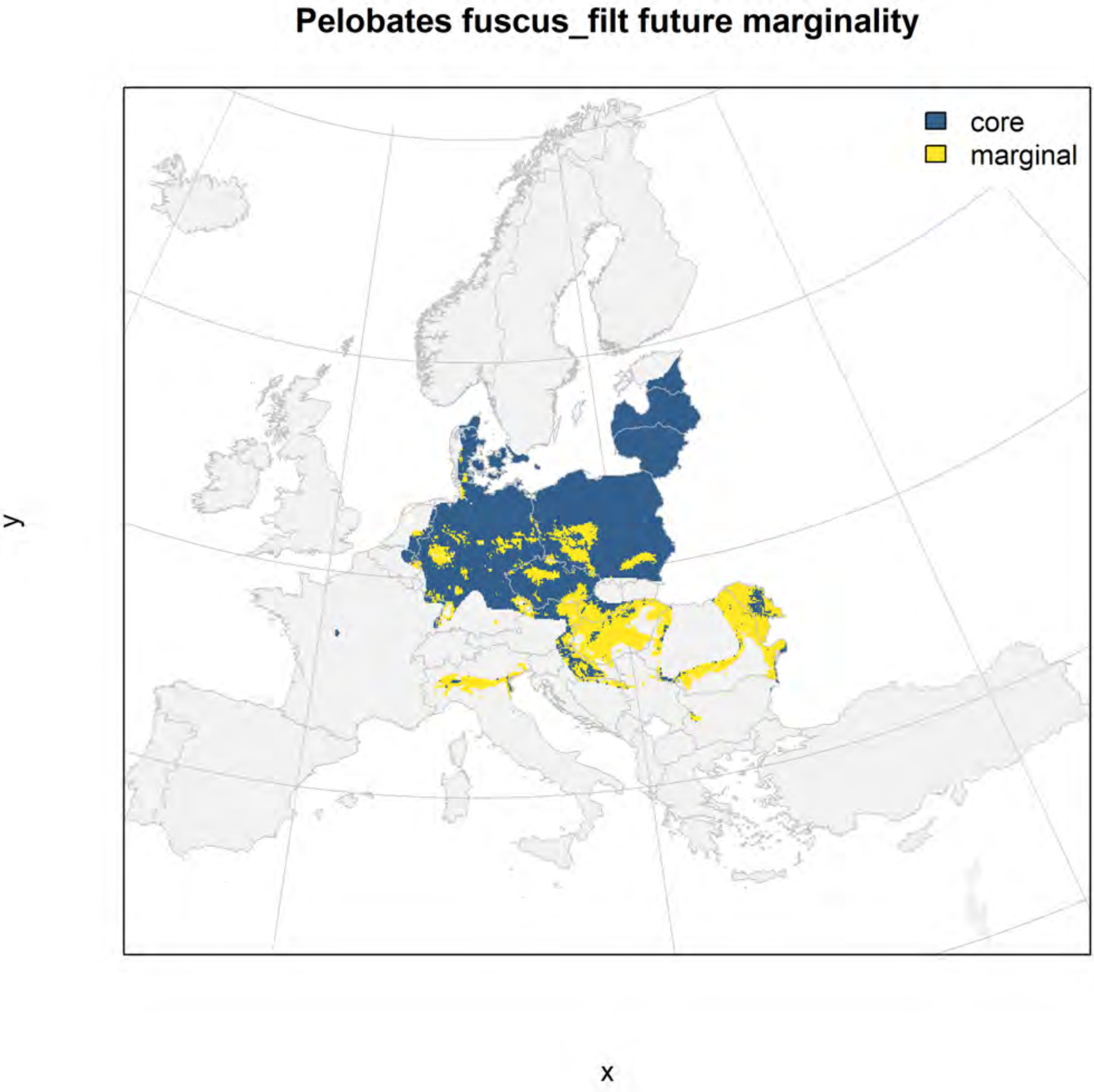
Pelobates fuscus_filt future marginality
core
marginal
y
x

### Pelobates syriacus_filt NMI

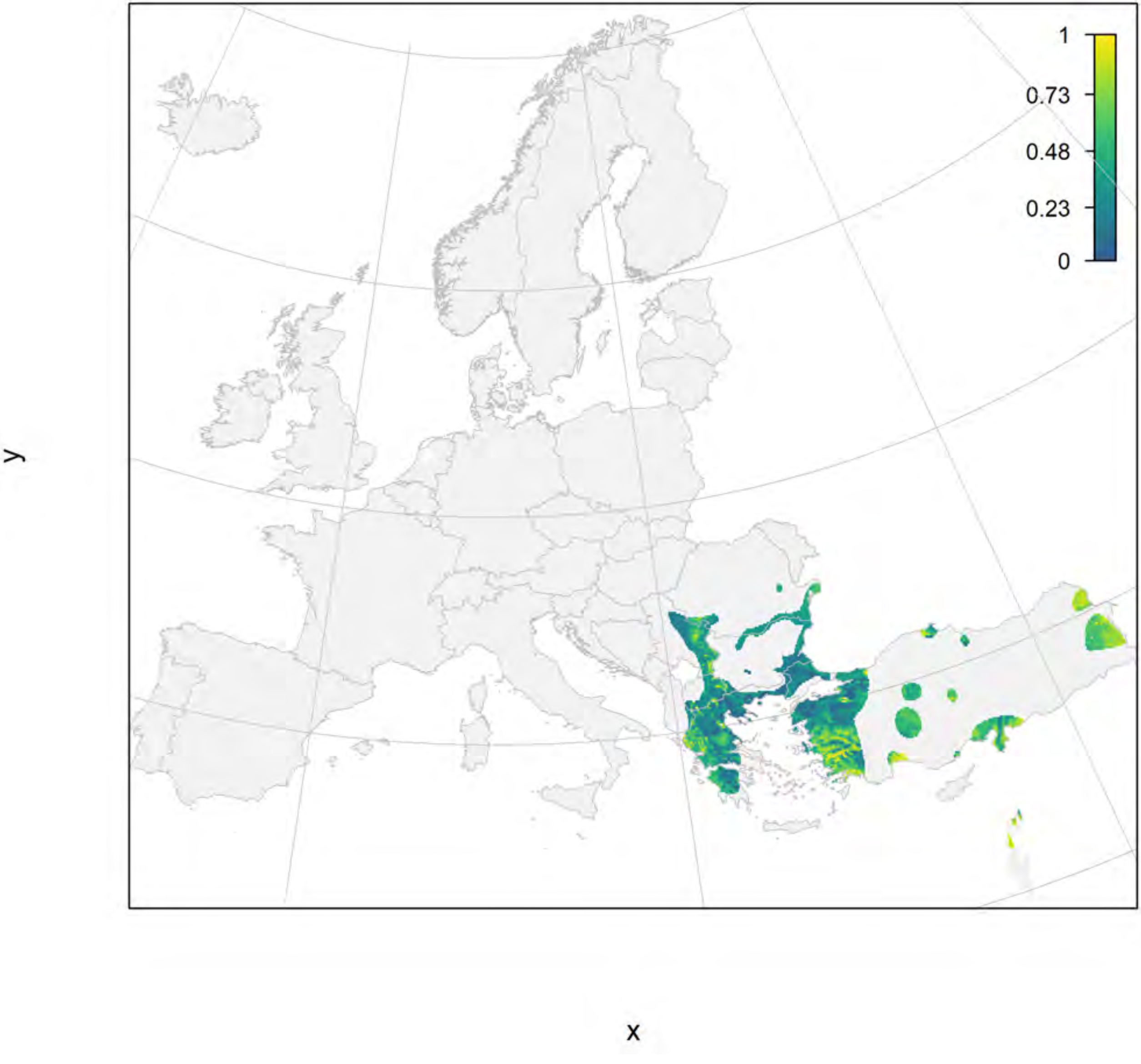

#### Pelobates syriacus_filt current marginality

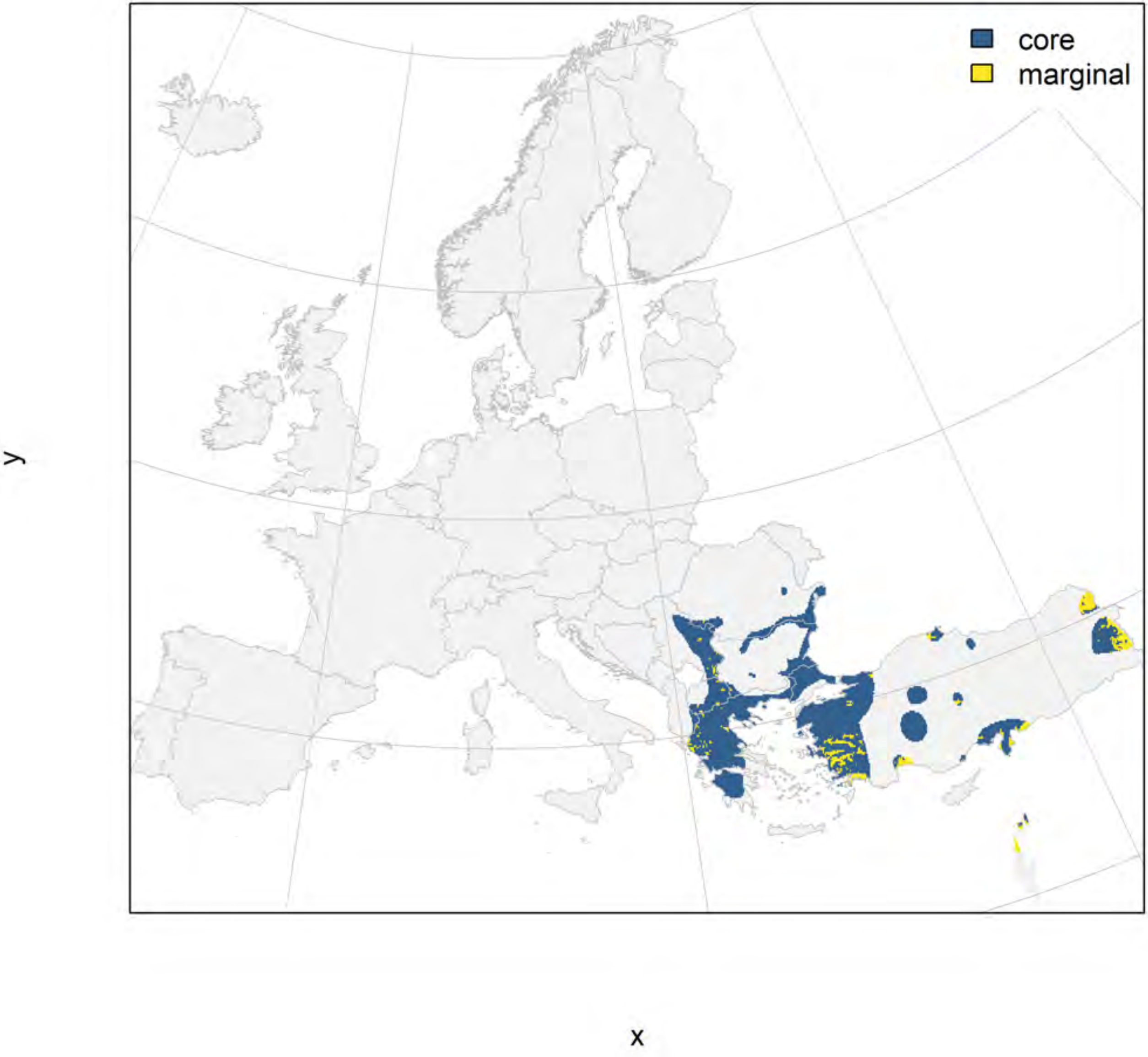

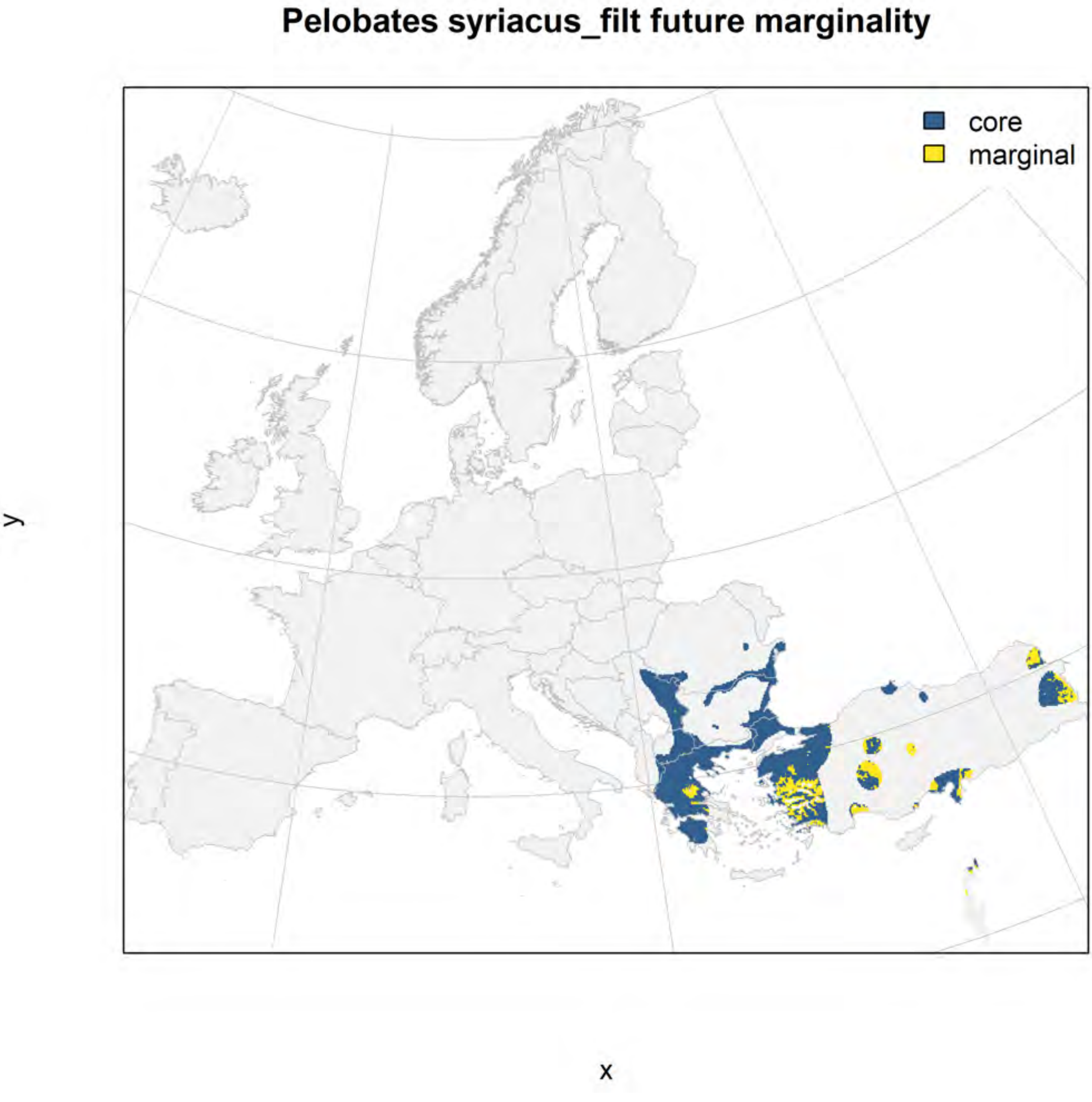
Pelobates syriacus_filt future marginality
core
marginal
y
x

### Pelodytes ibericus_filt NMI

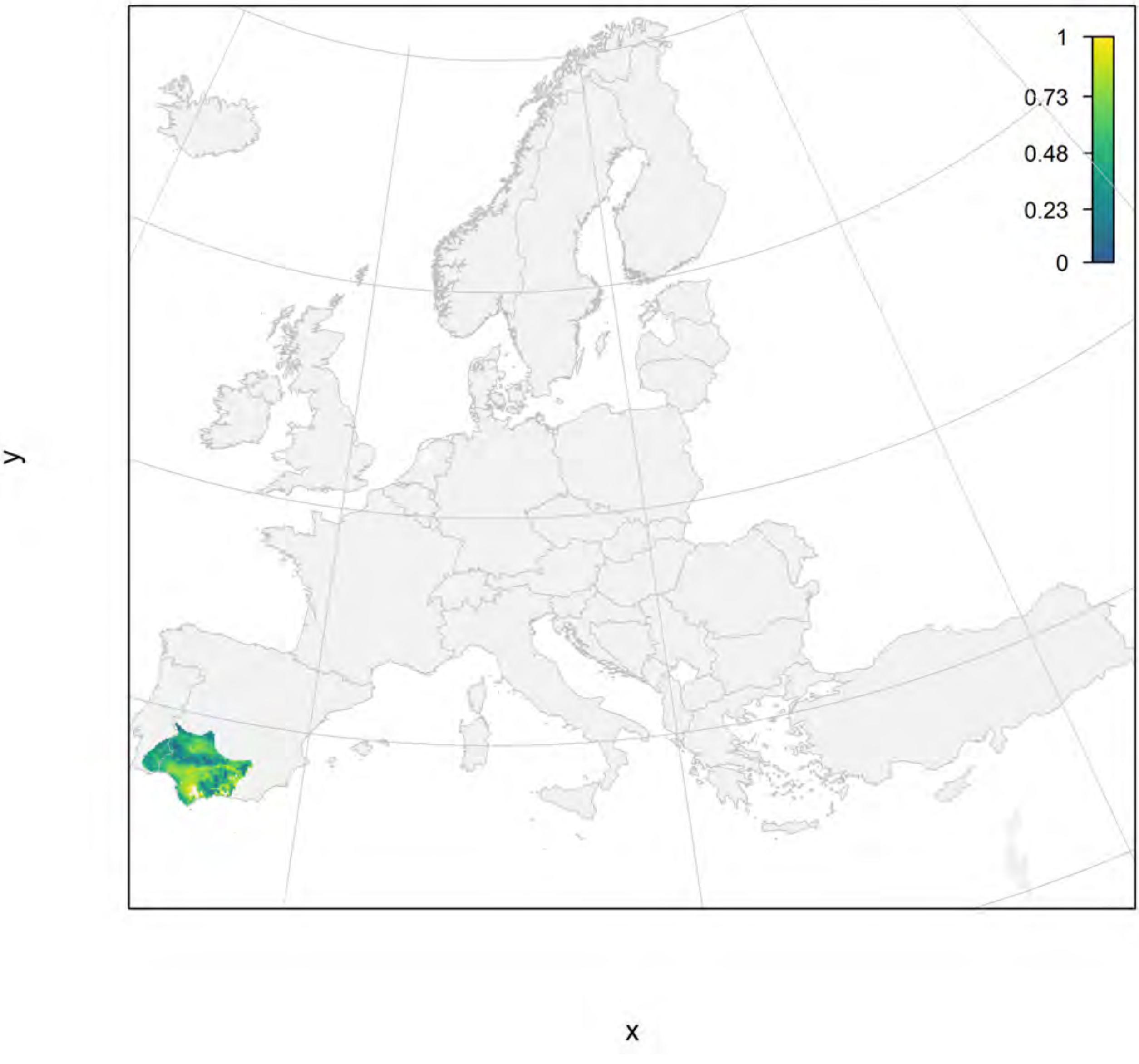

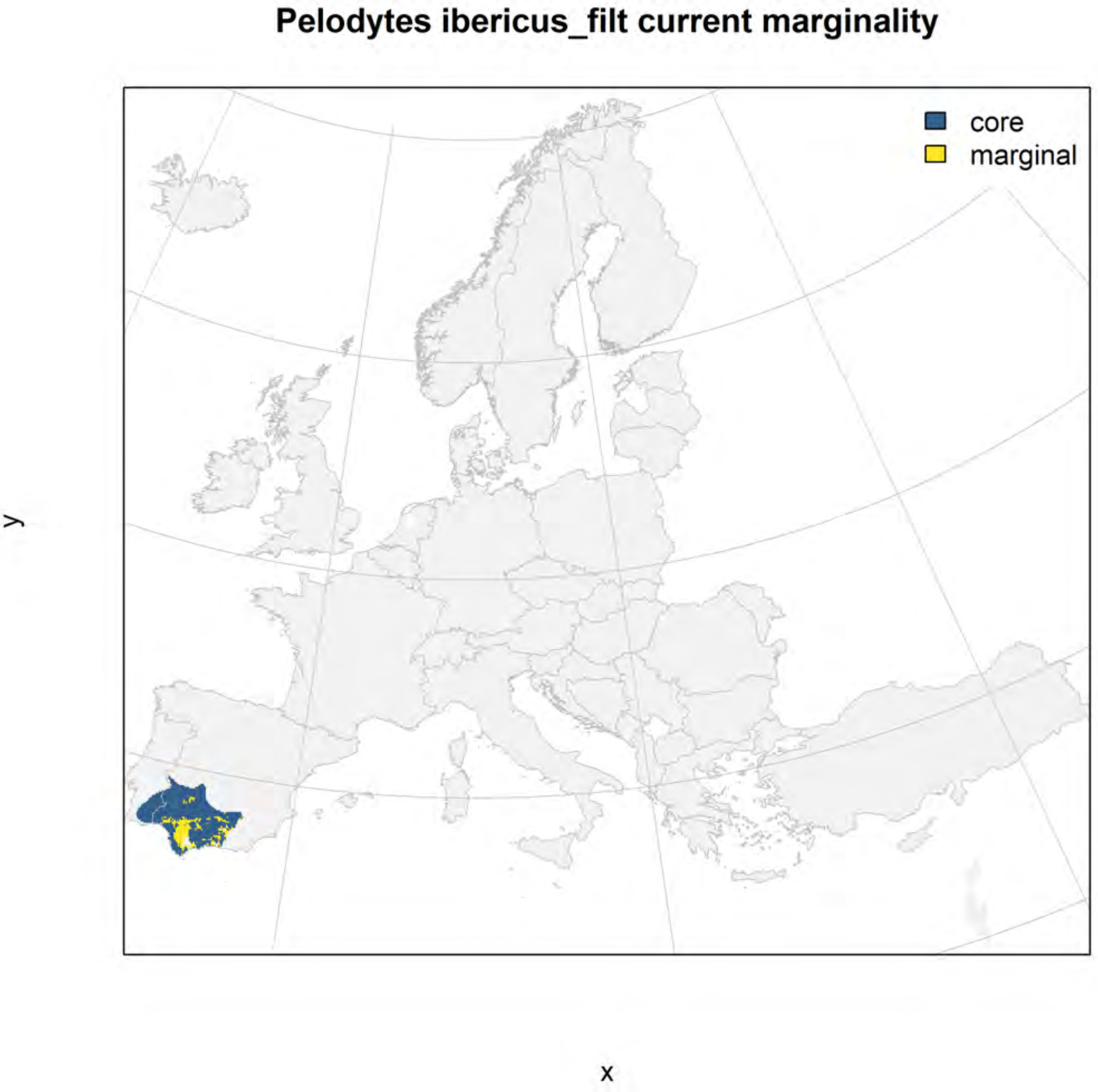
Pelodytes ibericus_filt current marginality
core
marginal
y
x

#### Pelodytes ibericus_filt future marginality

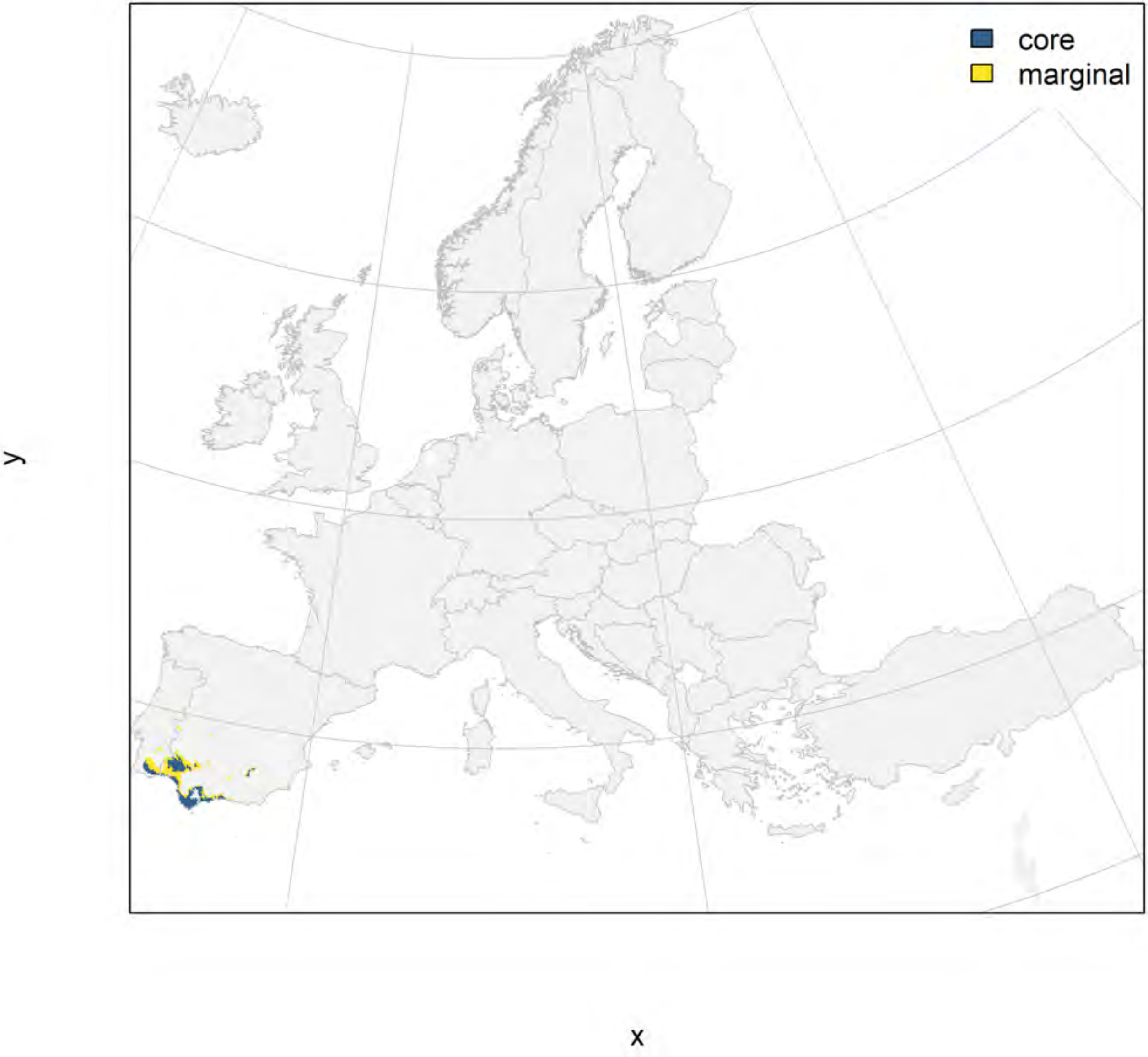

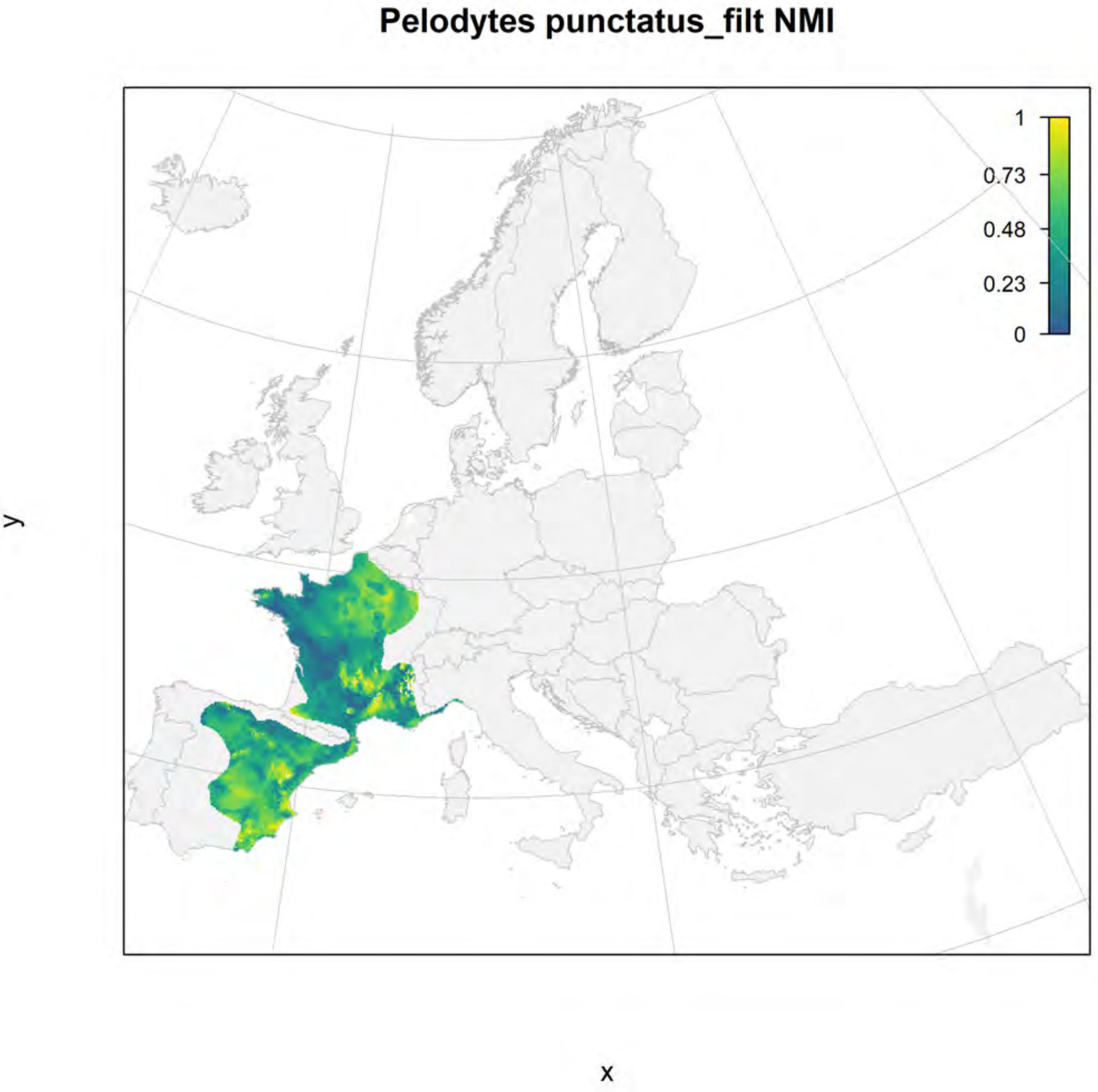
Pelodytes punctatus_filt NMI
1
0.73
0.48
0.23
0
y
x

#### Pelodytes punctatus_filt current marginality

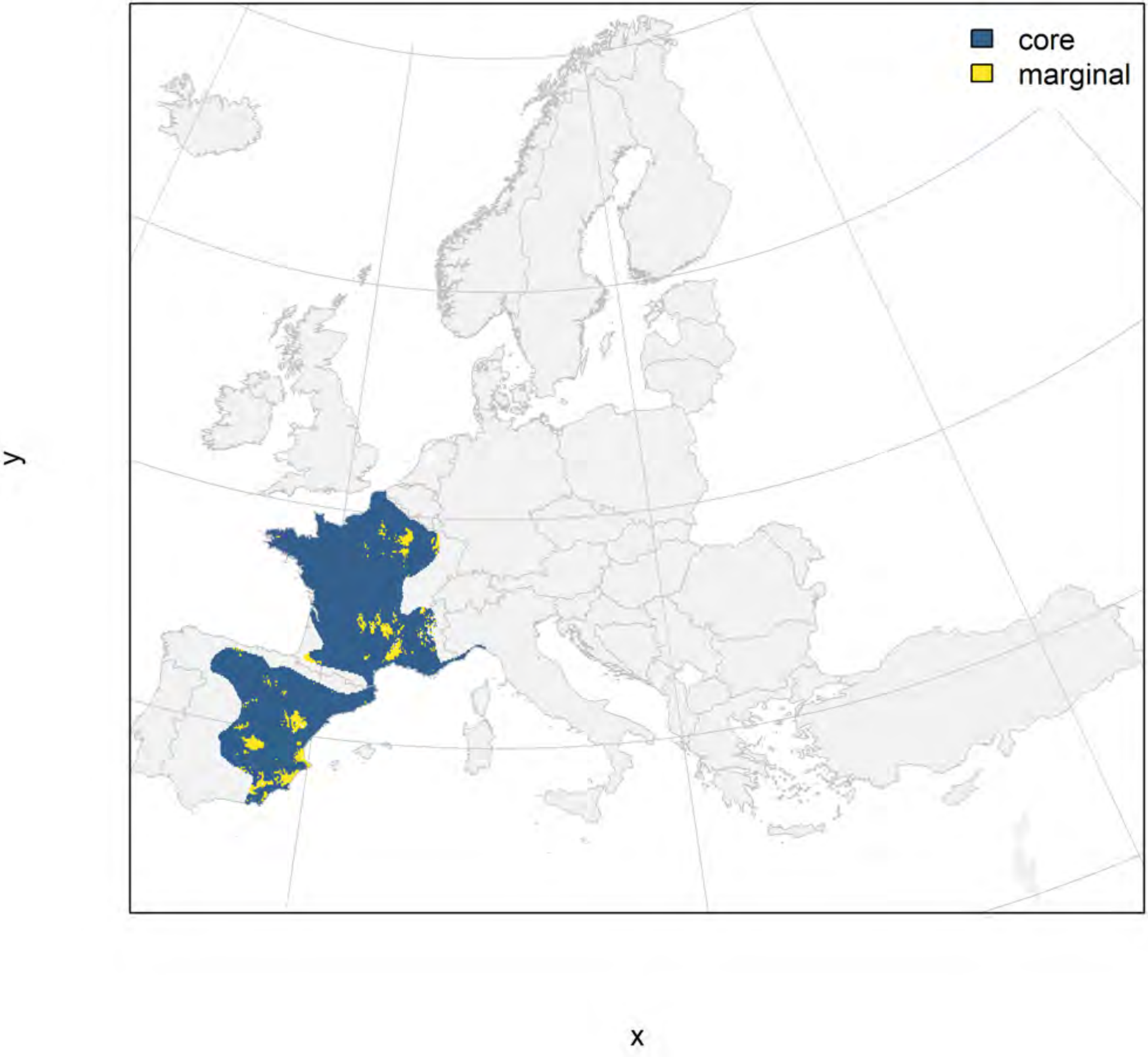

#### Pelodytes punctatus_filt future marginality

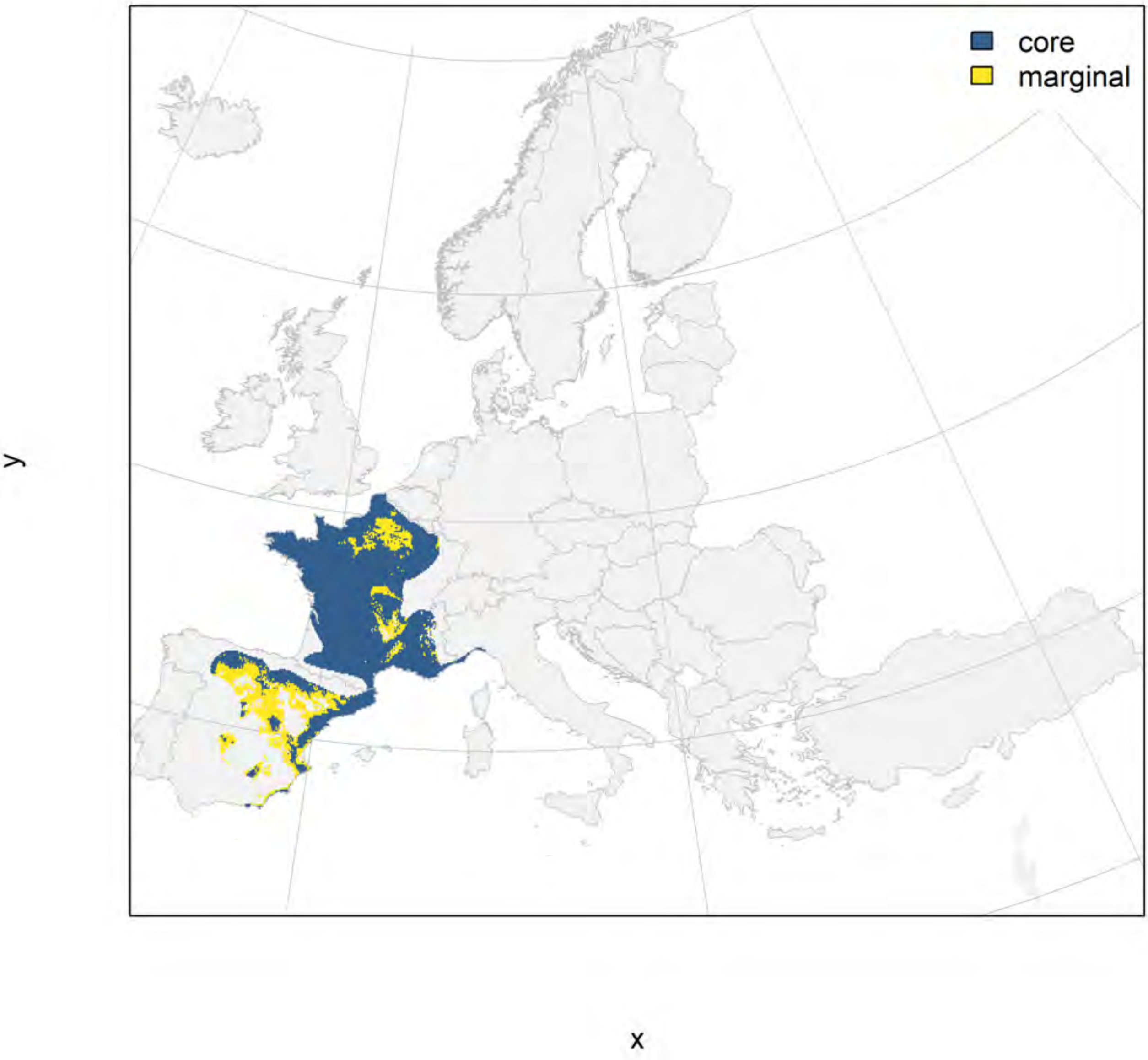

#### Pelophylax bedriagae_filt NMI

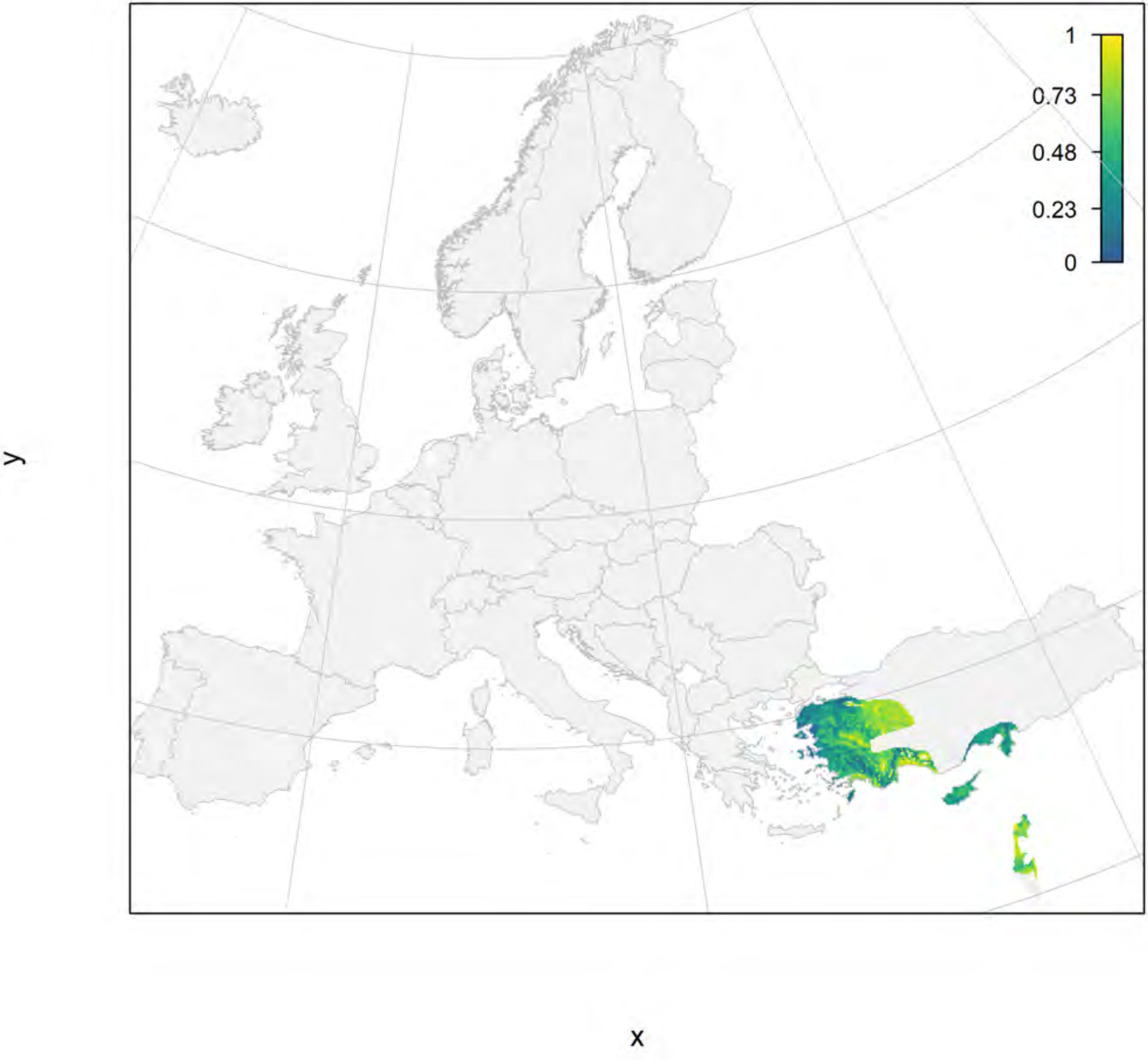

### Pelophylax bedriagae_filt current marginality

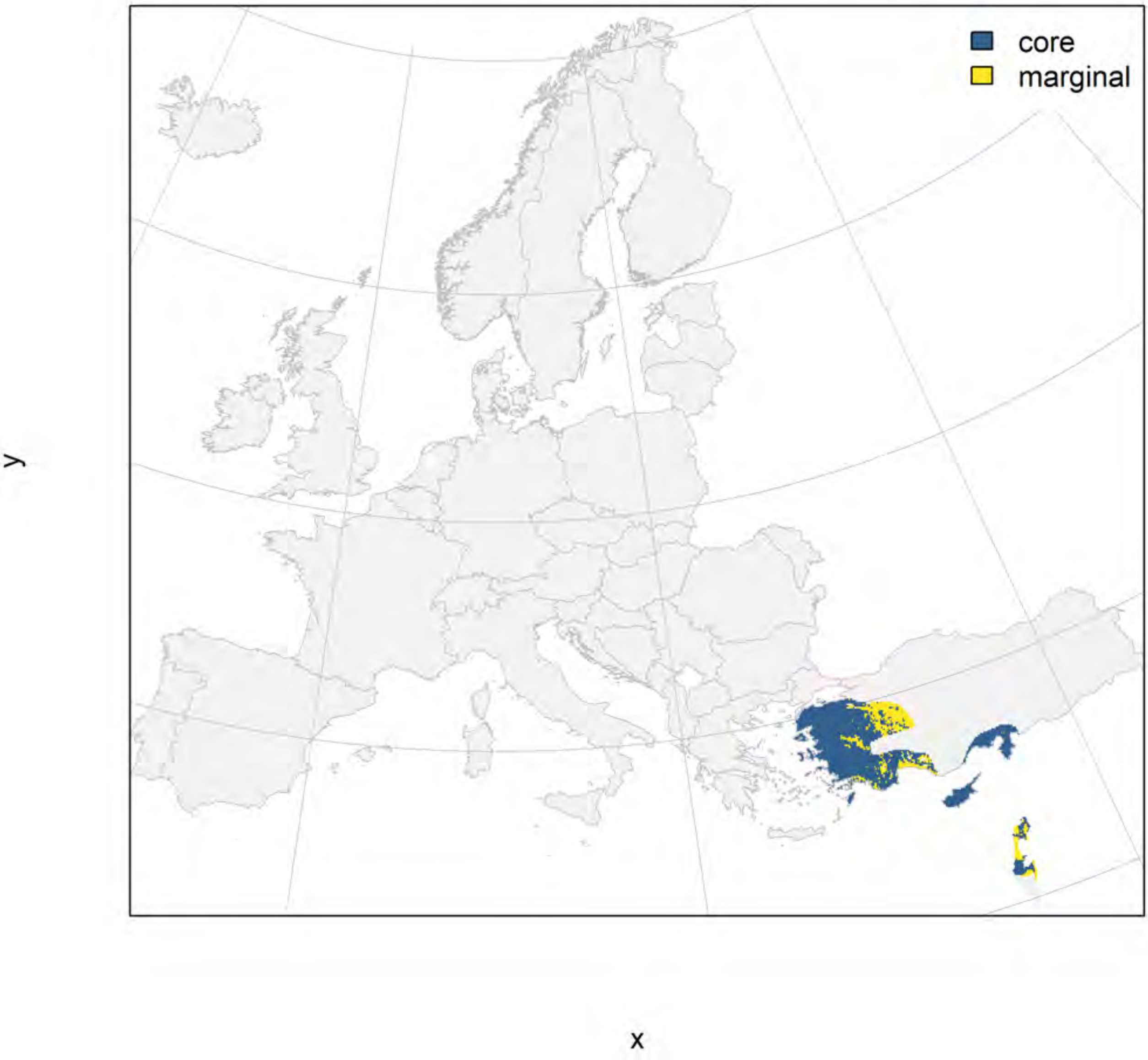

Pelophylax bedriagae_filt future marginality

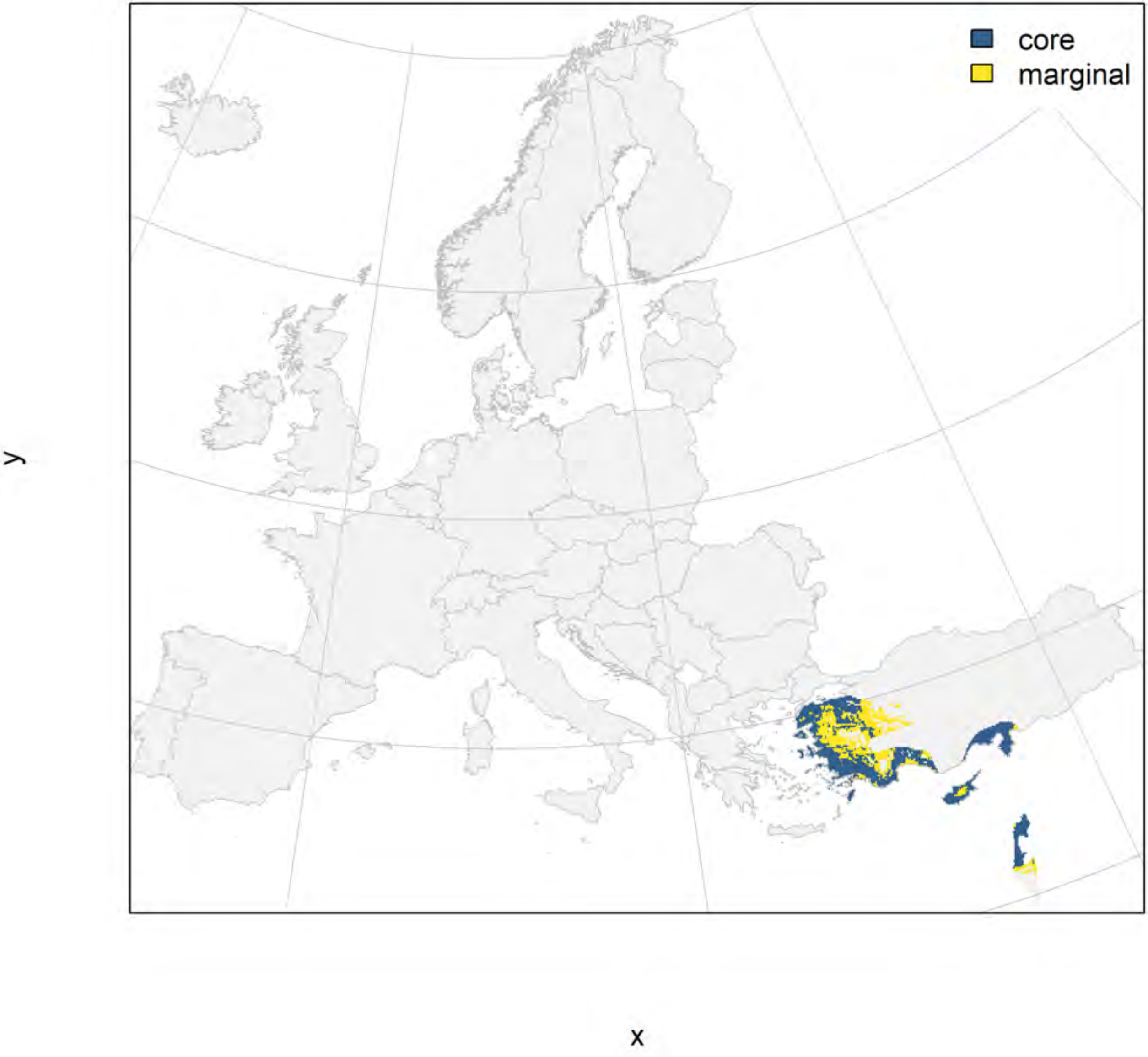

Pelophylax bergeri_filt NMI

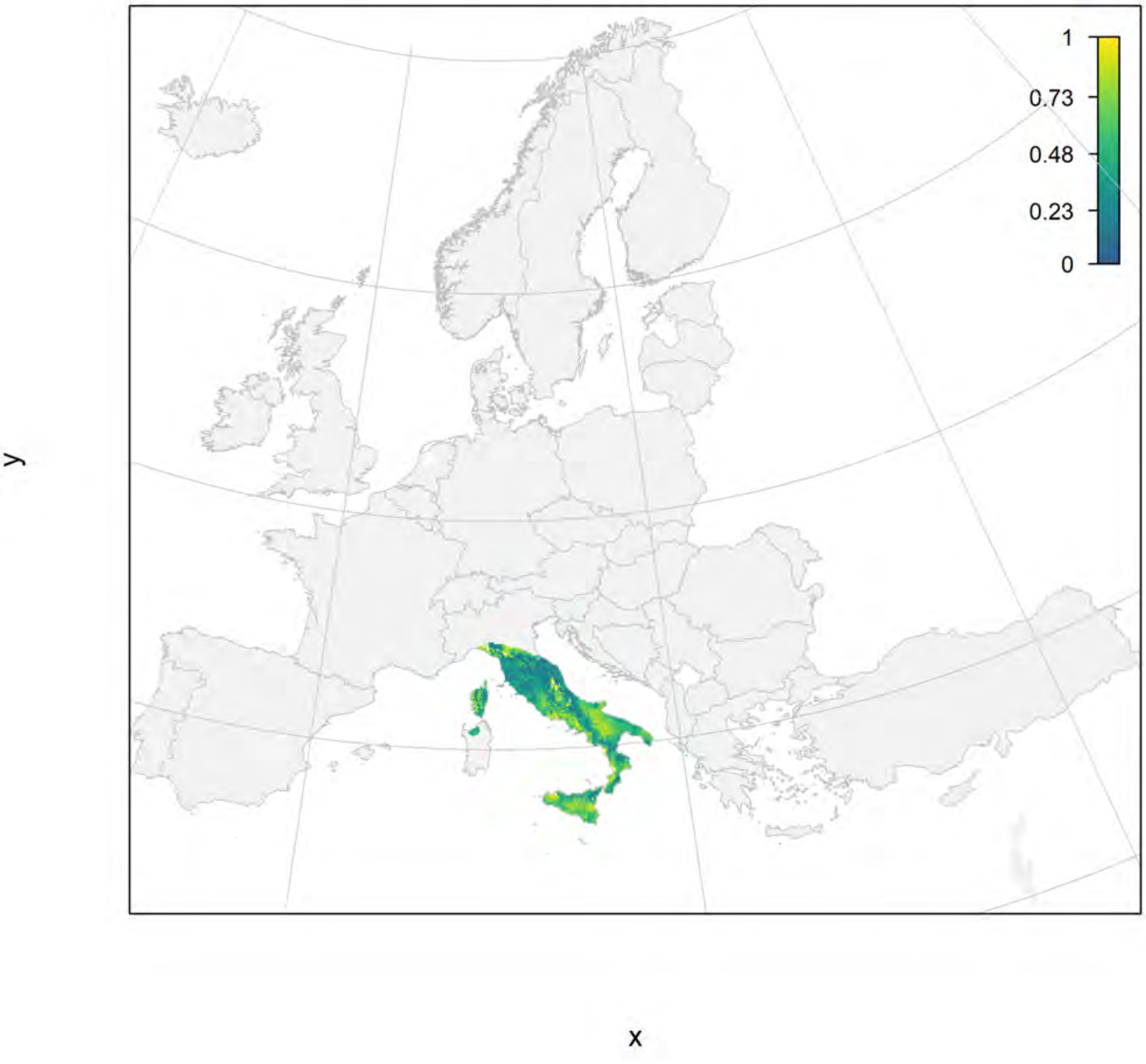

#### Pelophylax bergeri_filt current marginality

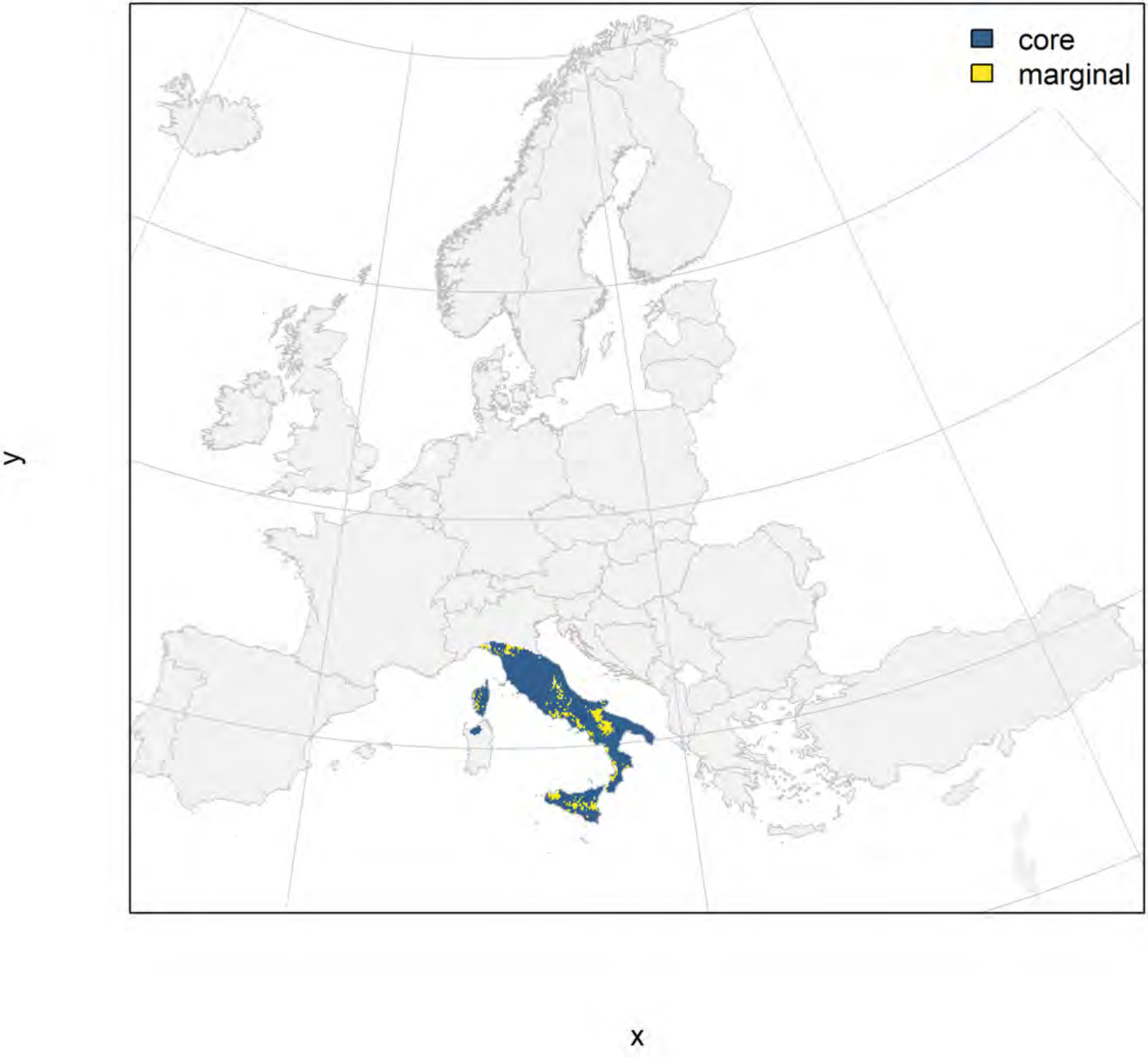

### Pelophylax bergeri_filt future marginality

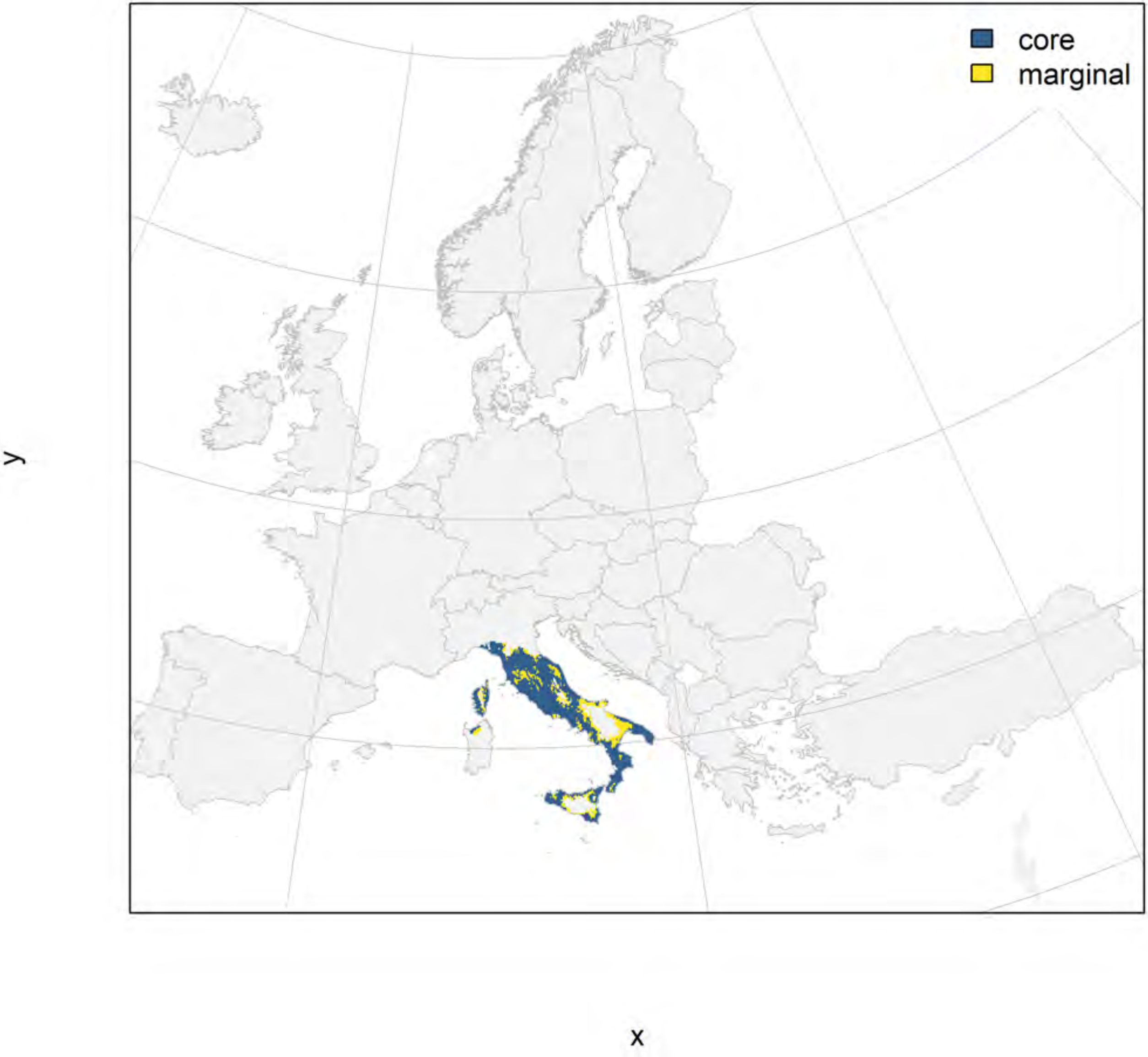

### Pelophylax cretensis_filt NMI

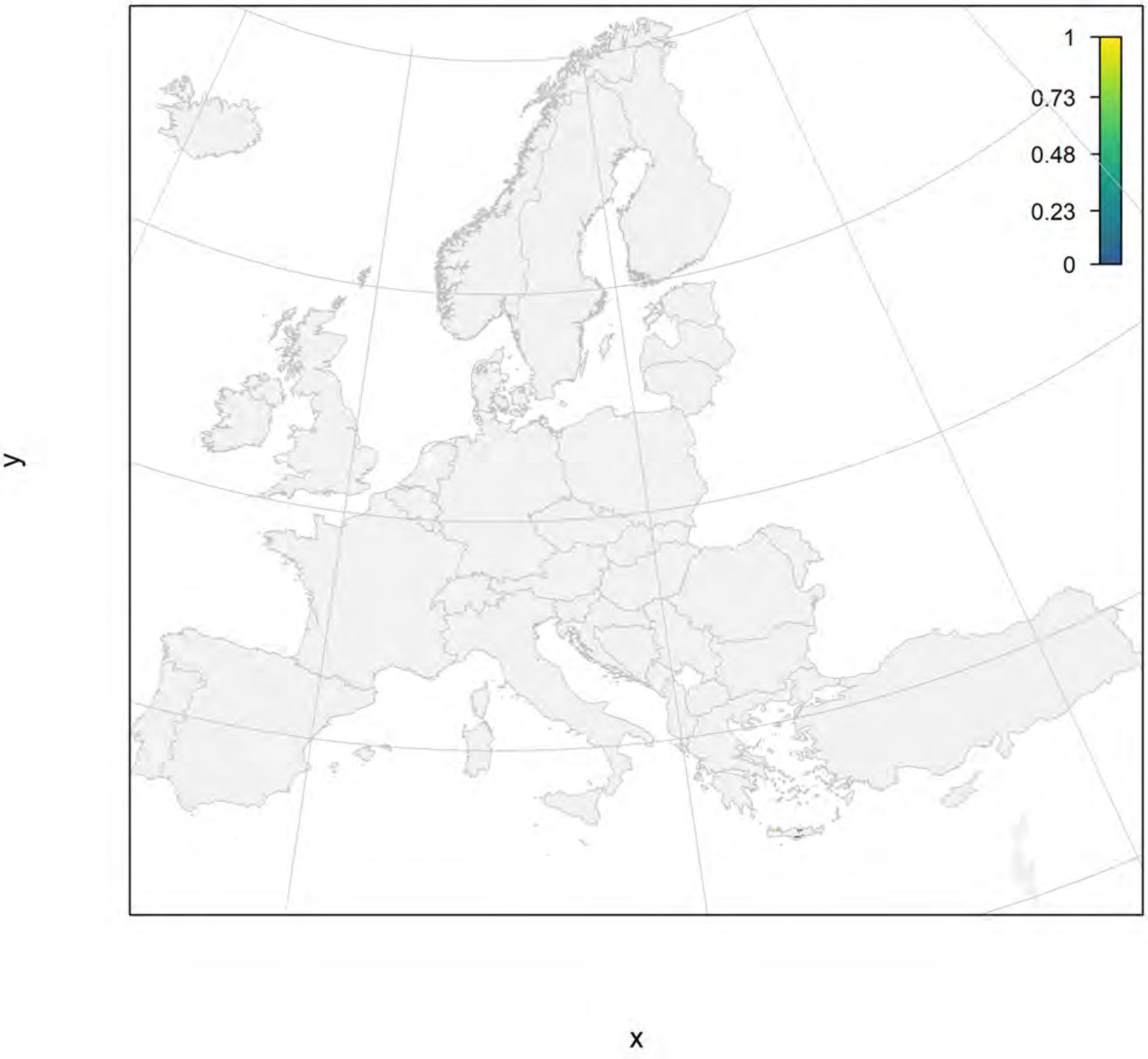

Pelophylax cretensis_filt current marginality

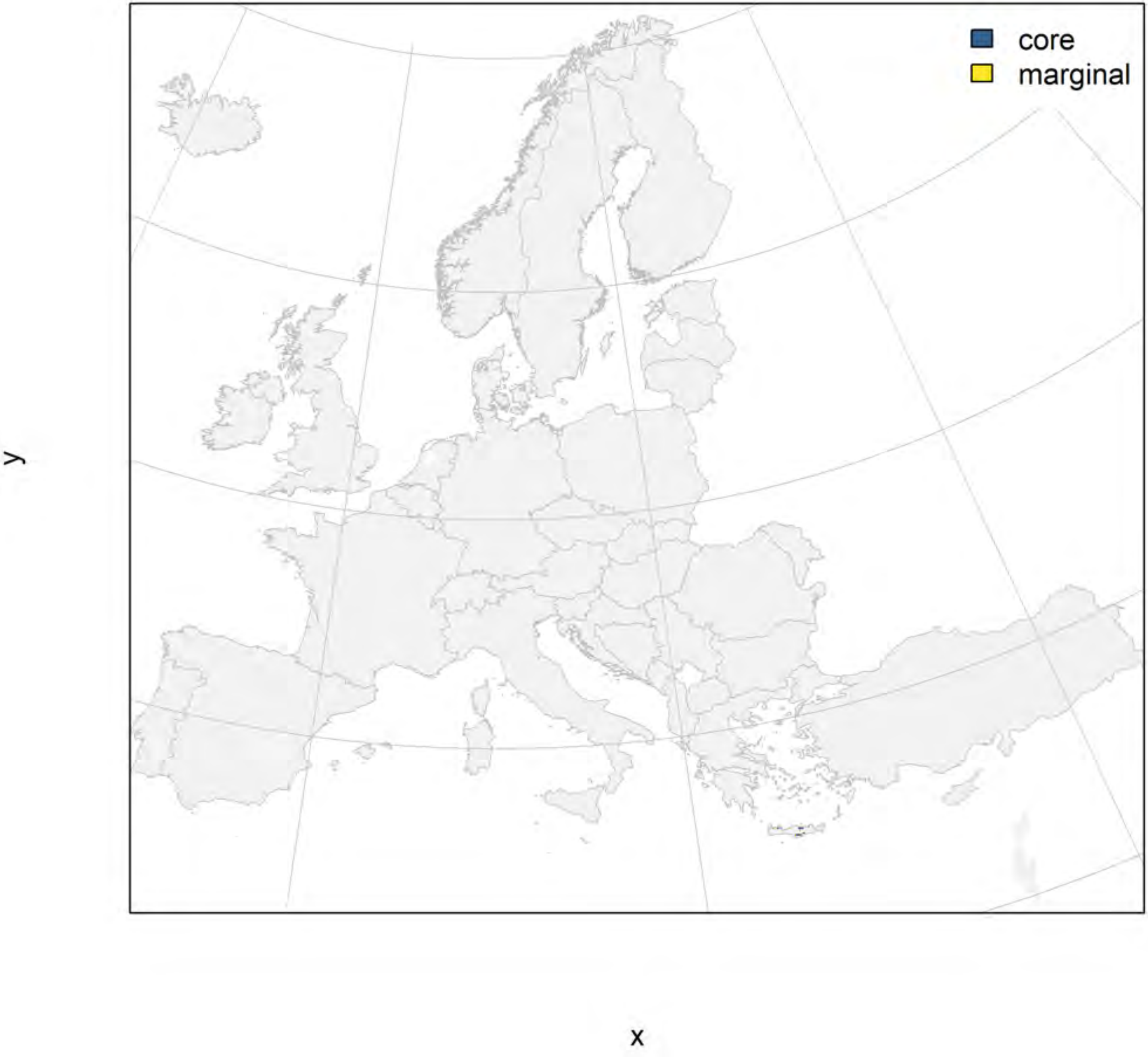

#### Pelophylax cretensis_filt future marginality

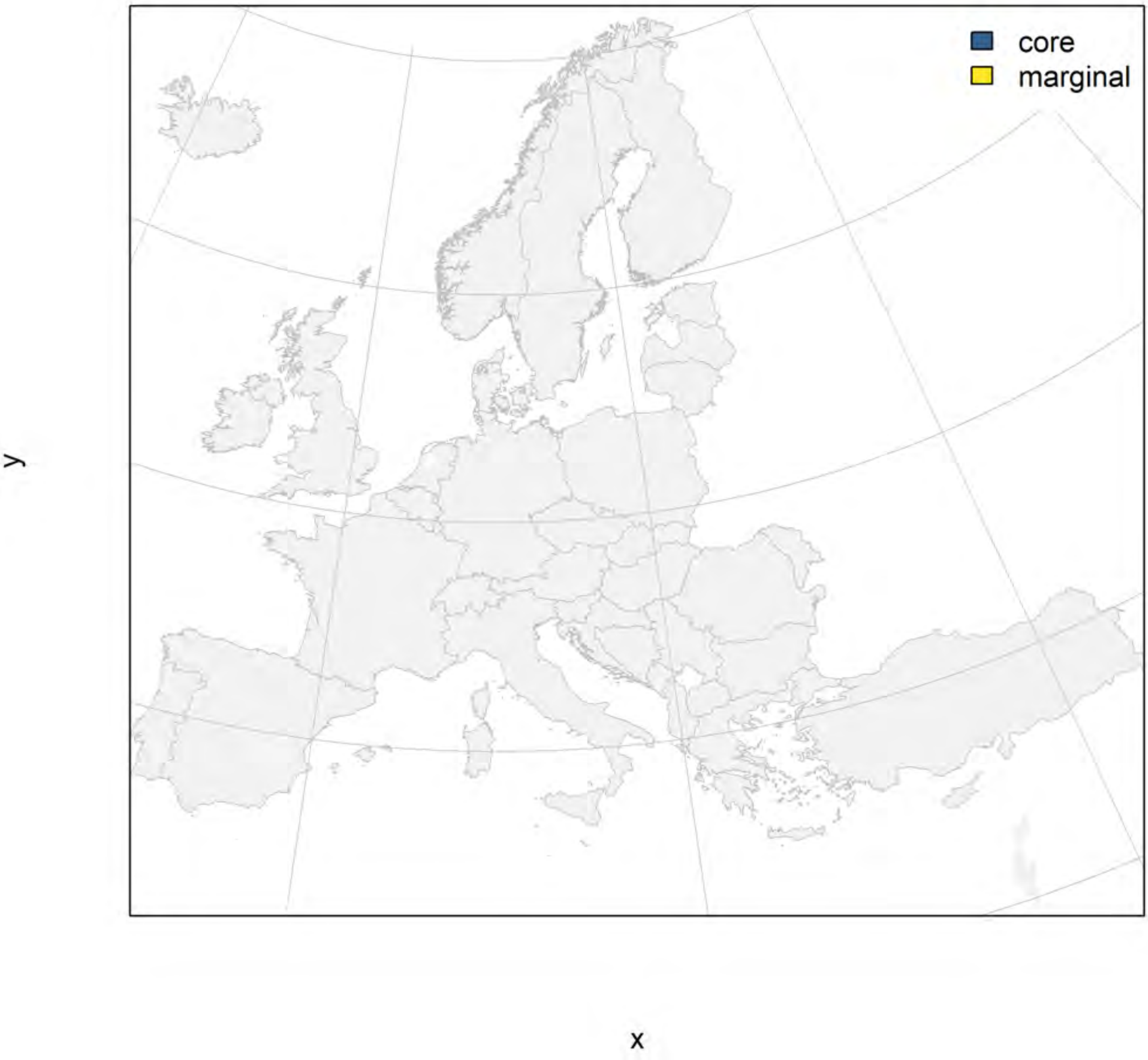

#### Pelophylax epeiroticus_filt NMI

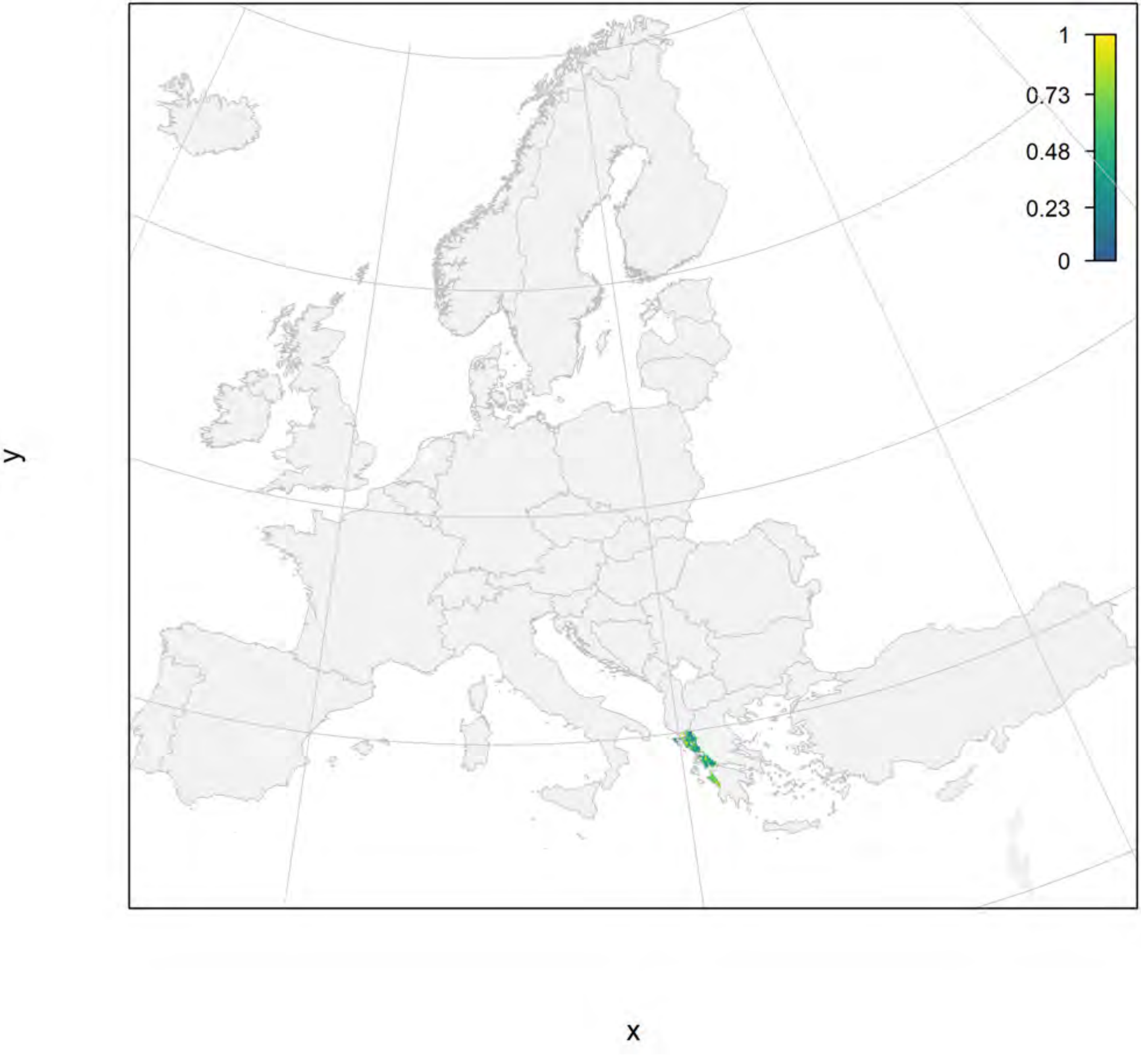

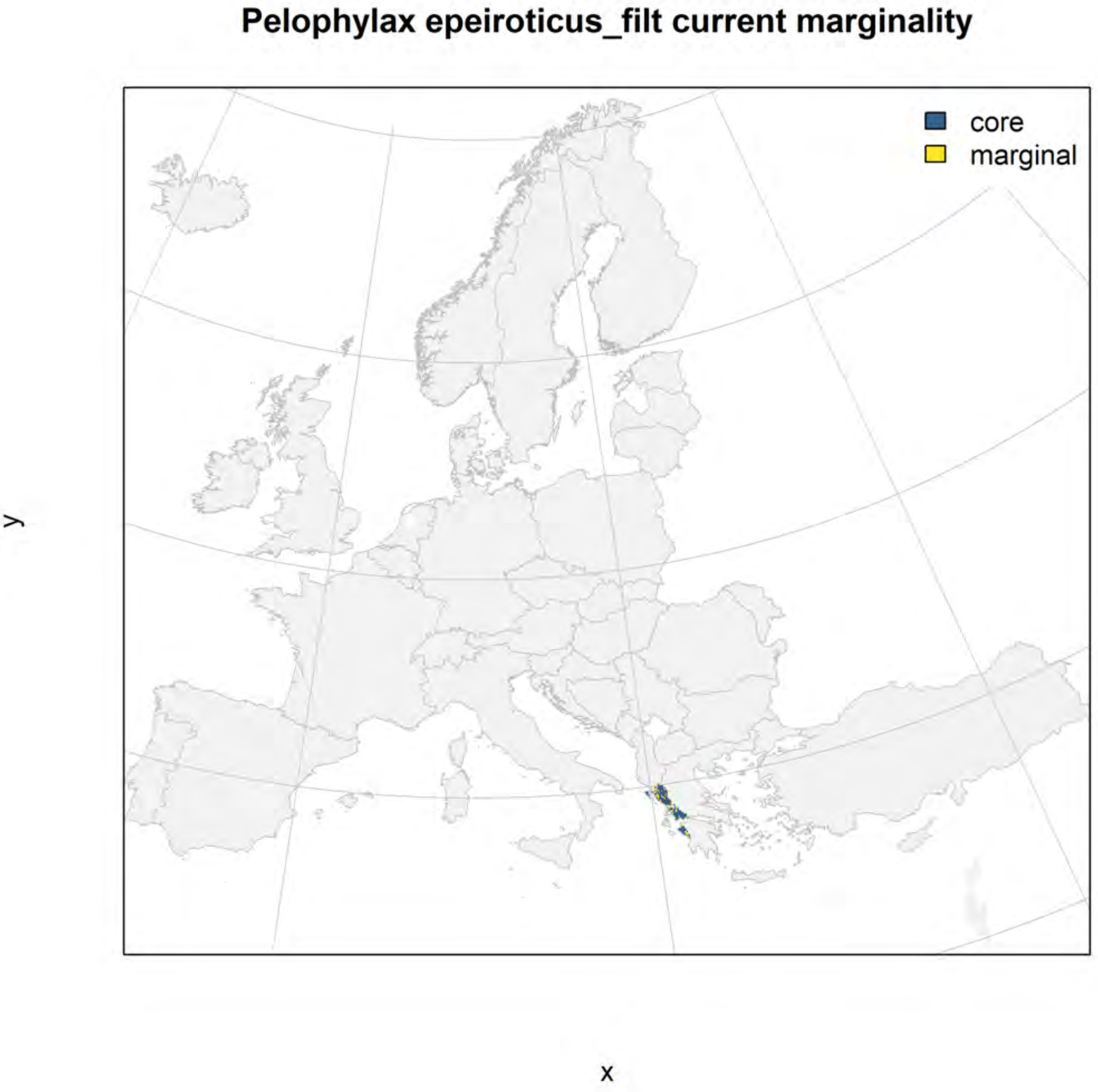
Pelophylax epeiroticus_filt current marginality
core
marginal
y
x

### Pelophylax epeiroticus_filt future marginality

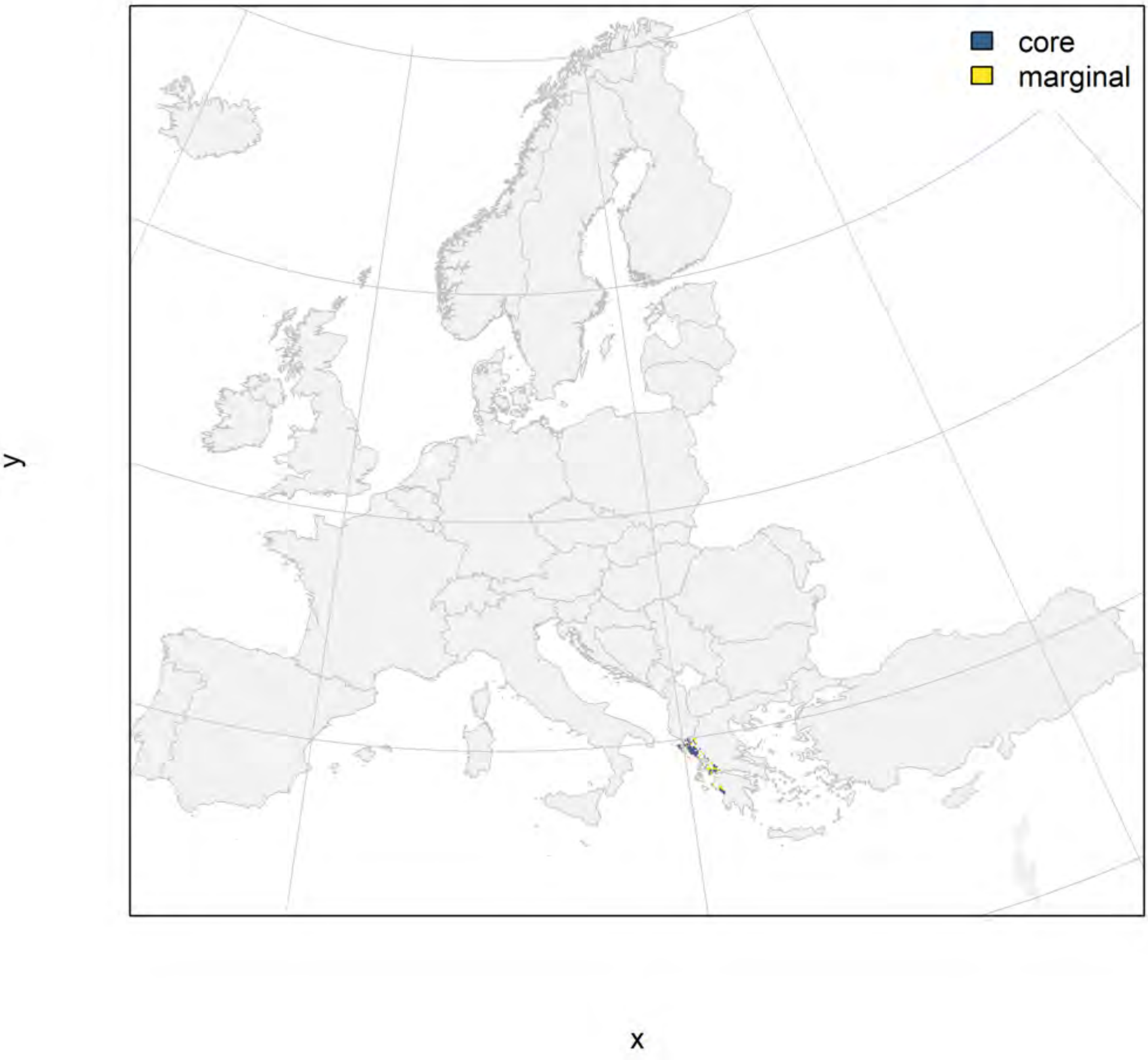

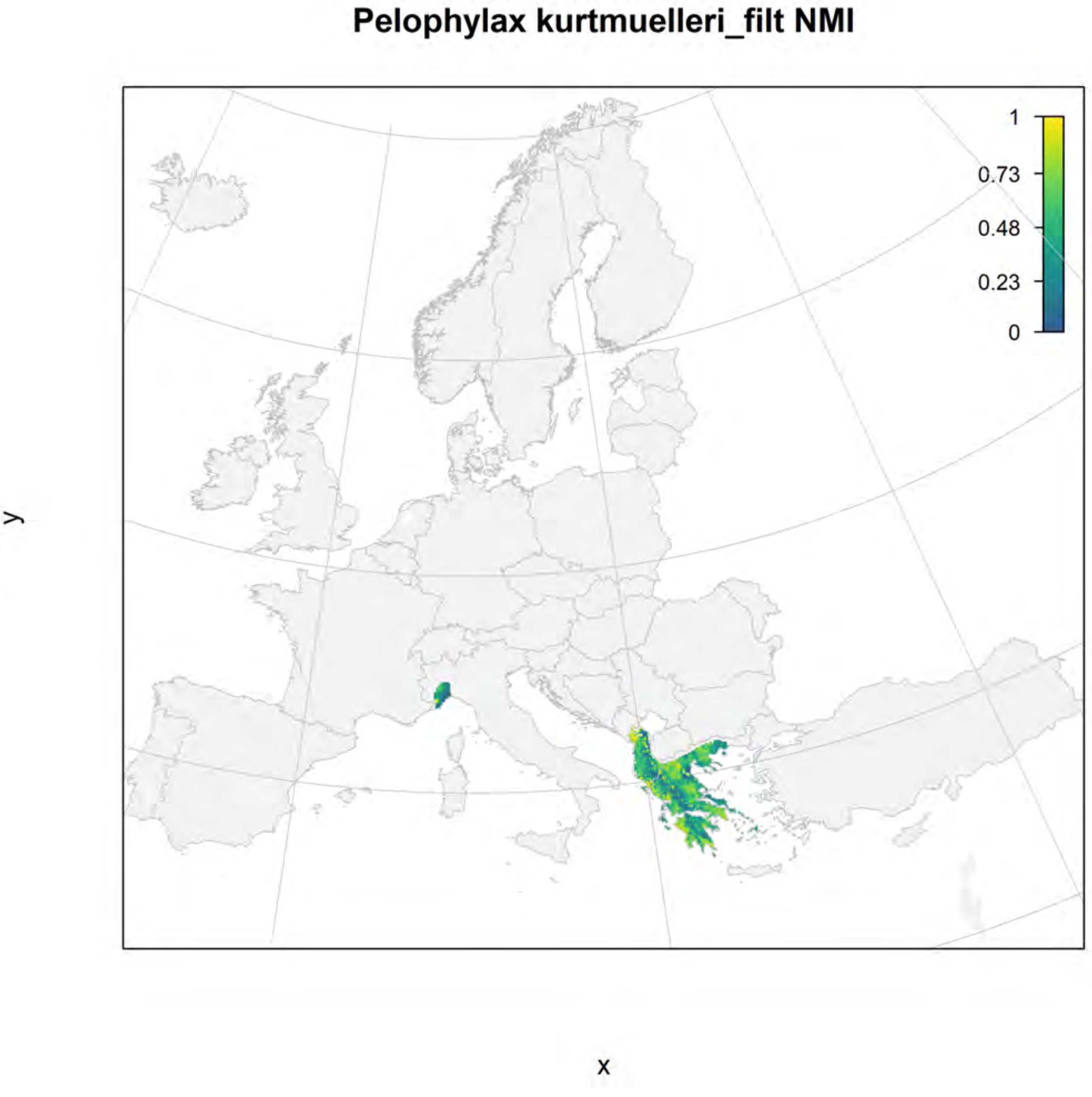
Pelophylax kurtmuelleri_filt NMI
1
0.73
0.48
0.23
0
y
x

#### Pelophylax kurtmuelleri_filt current marginality

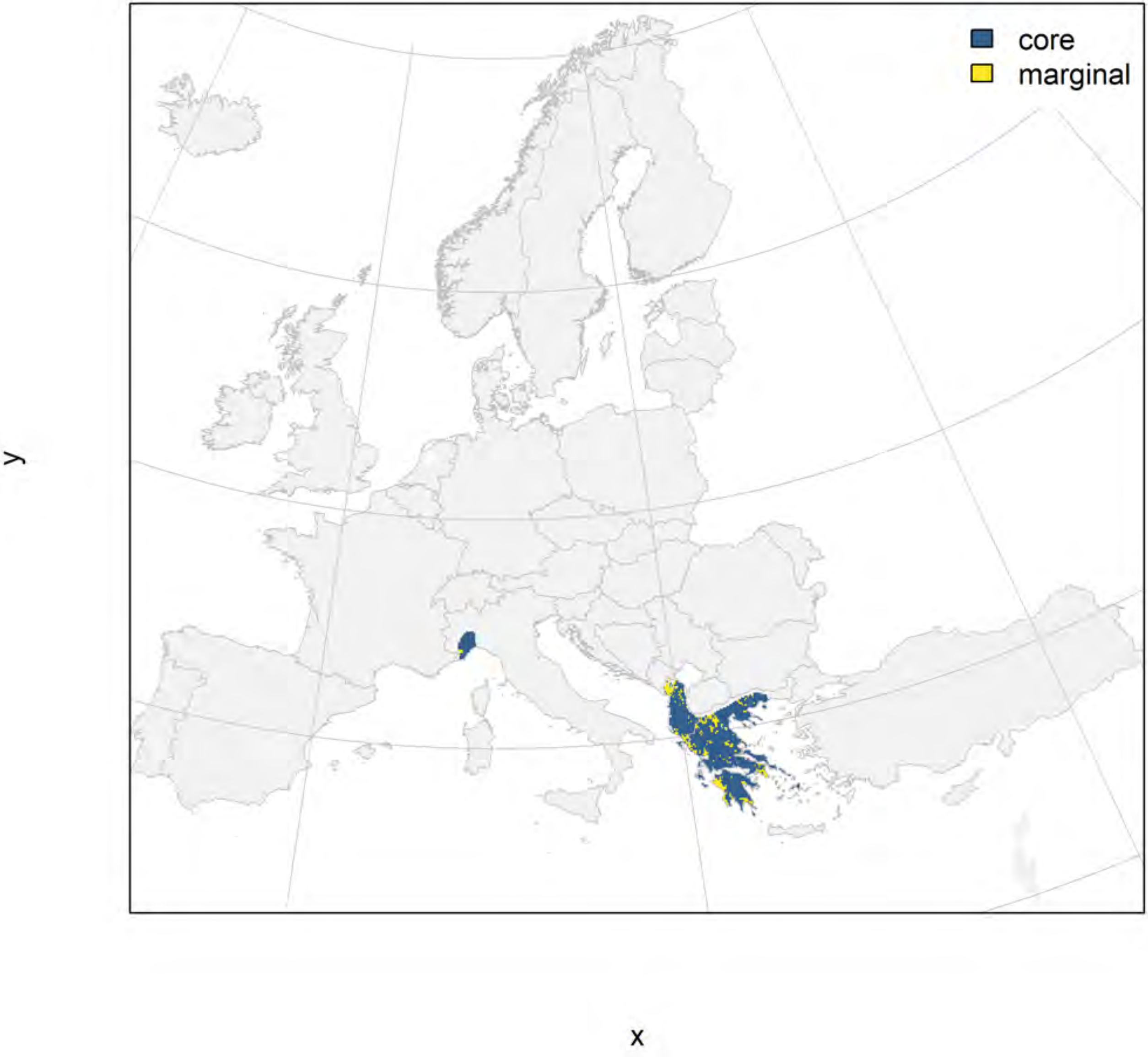

### Pelophylax kurtmuelleri_filt future marginality

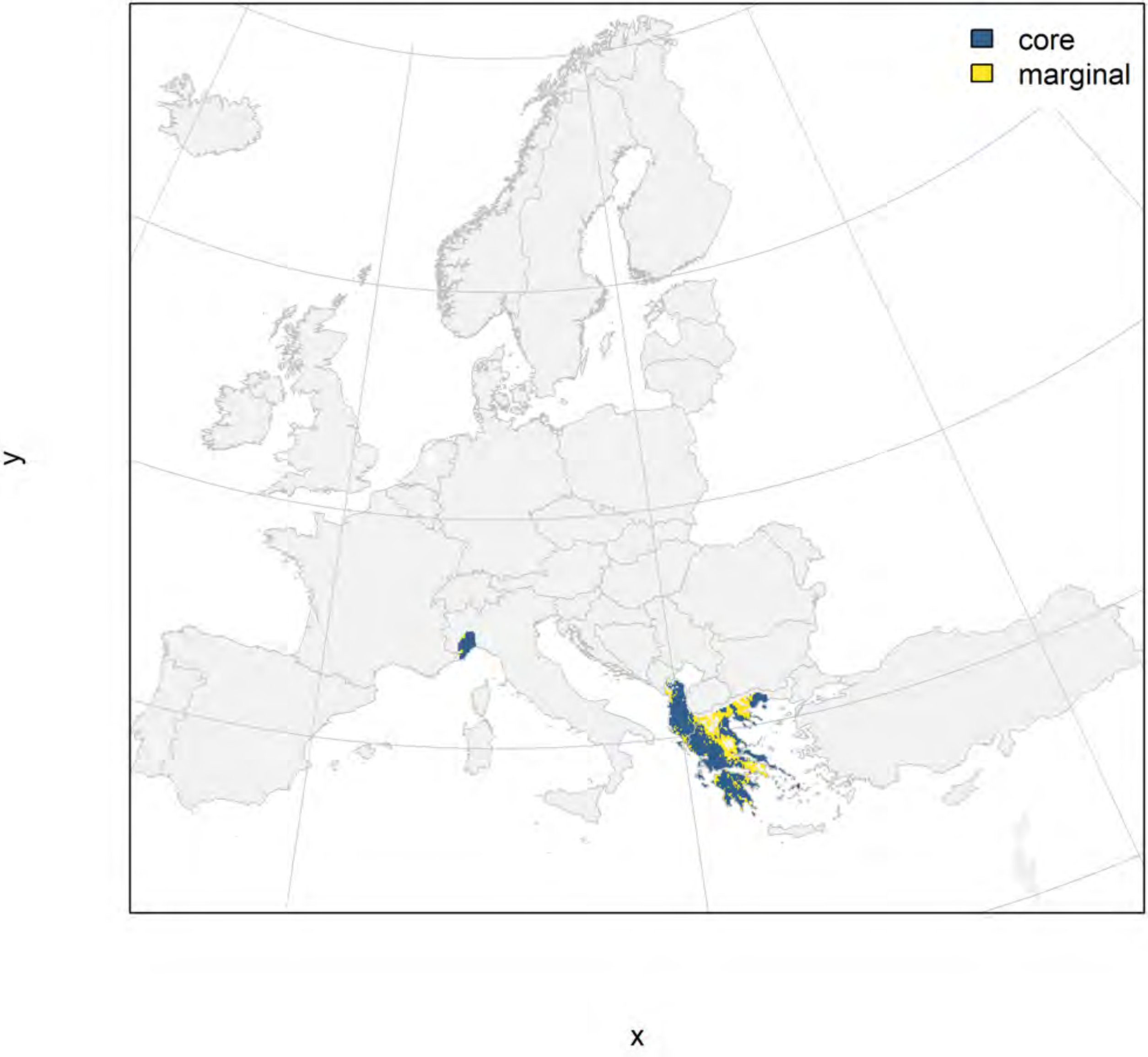

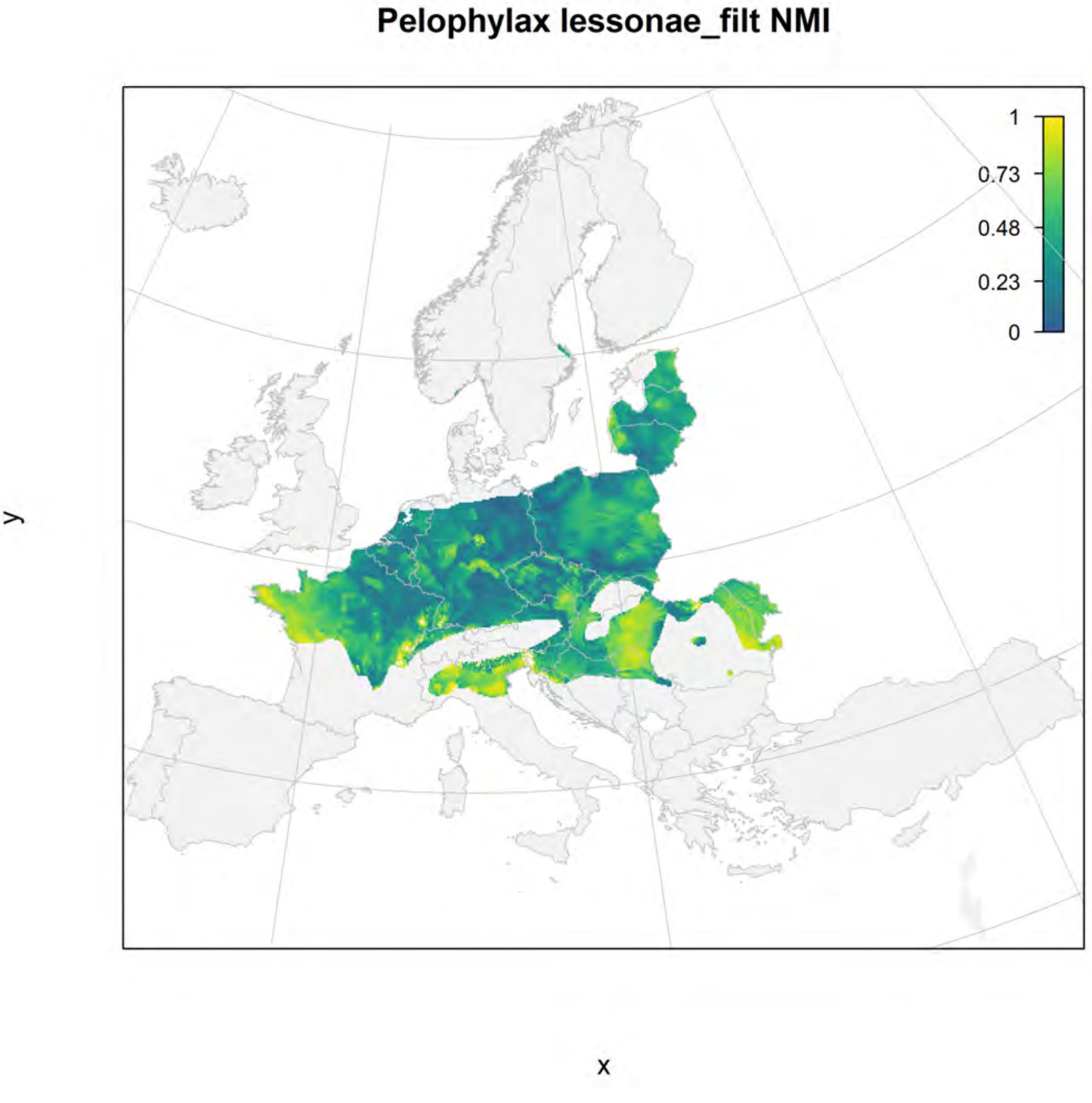
Pelophylax lessonae_filt NMI
1
0.73
0.48
0.23
0
y
x

### Pelophylax lessonae_filt current marginality

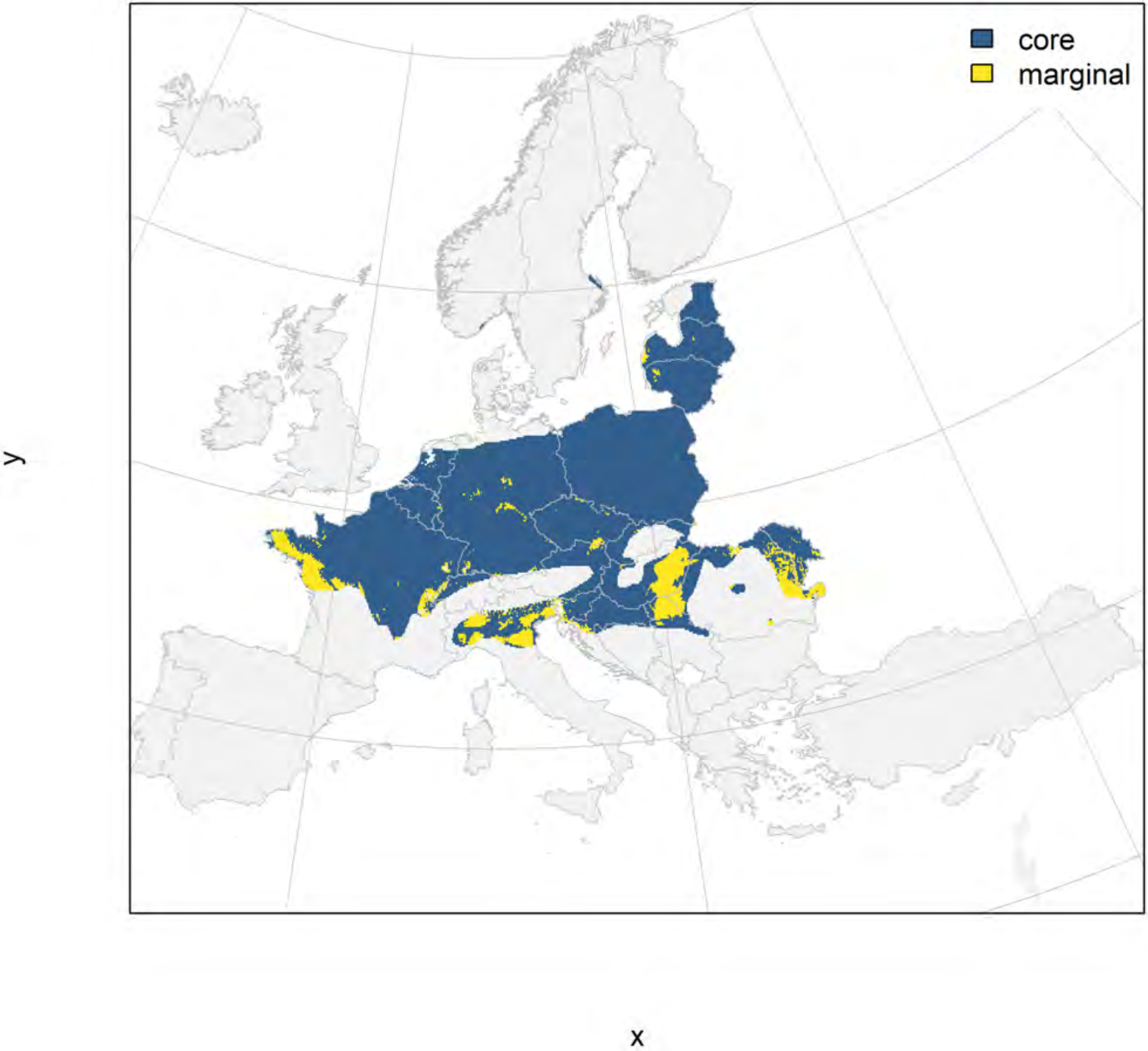

Pelophylax lessonae_filt future marginality

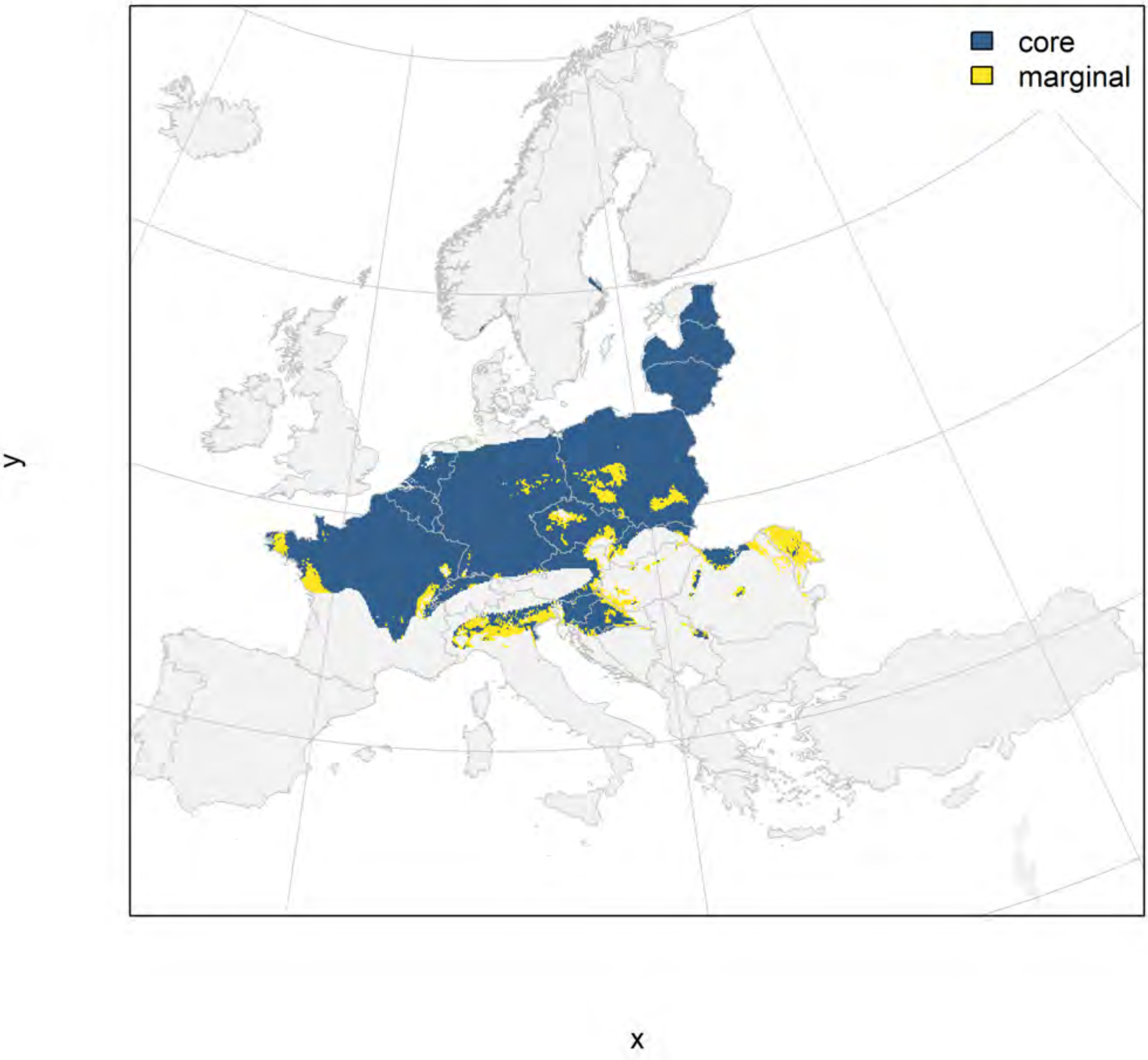

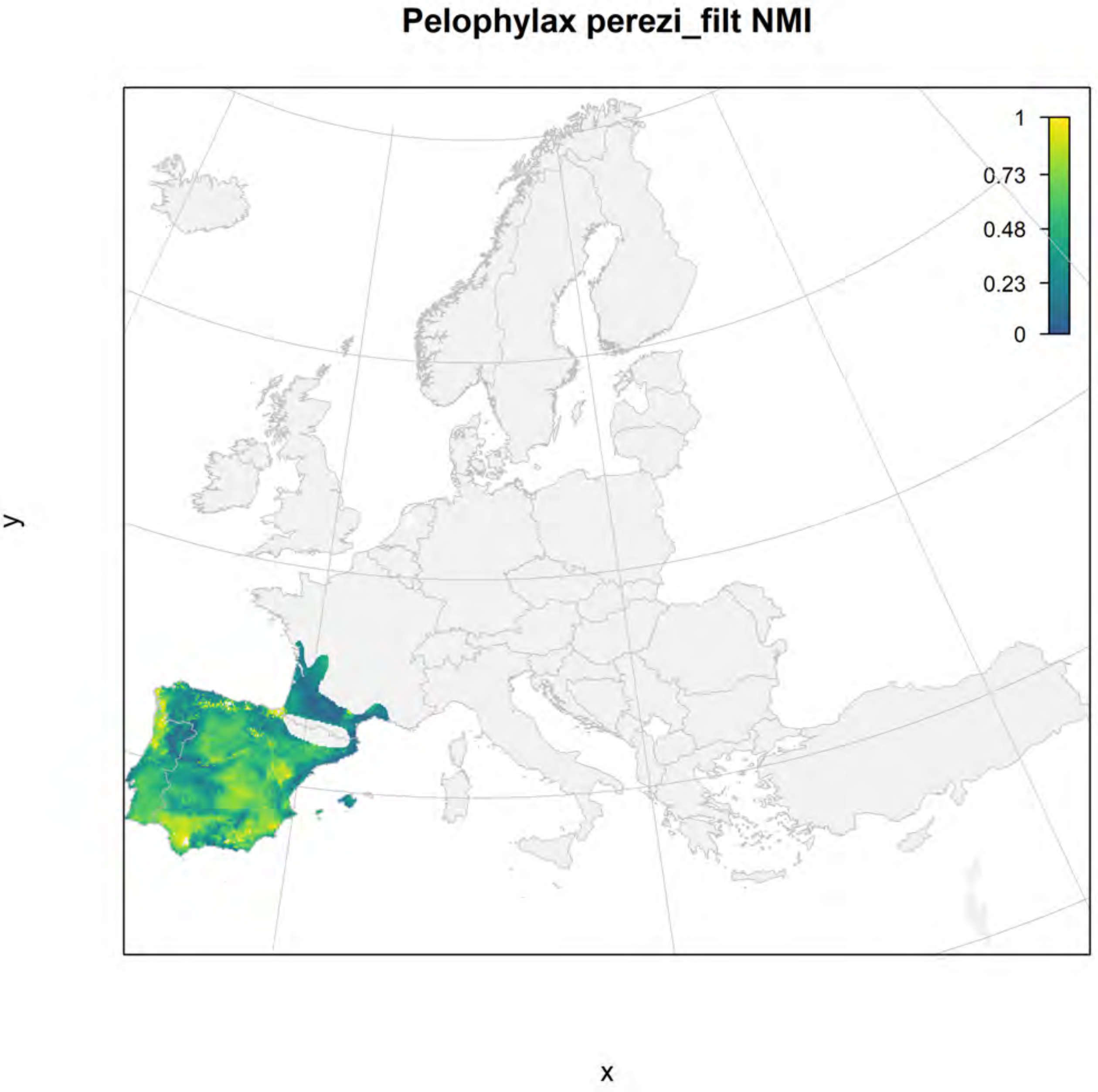
Pelophylax perezi_filt NMI
1
0.73
0.48
0.23
0
y
x

#### Pelophylax perezi_filt current marginality

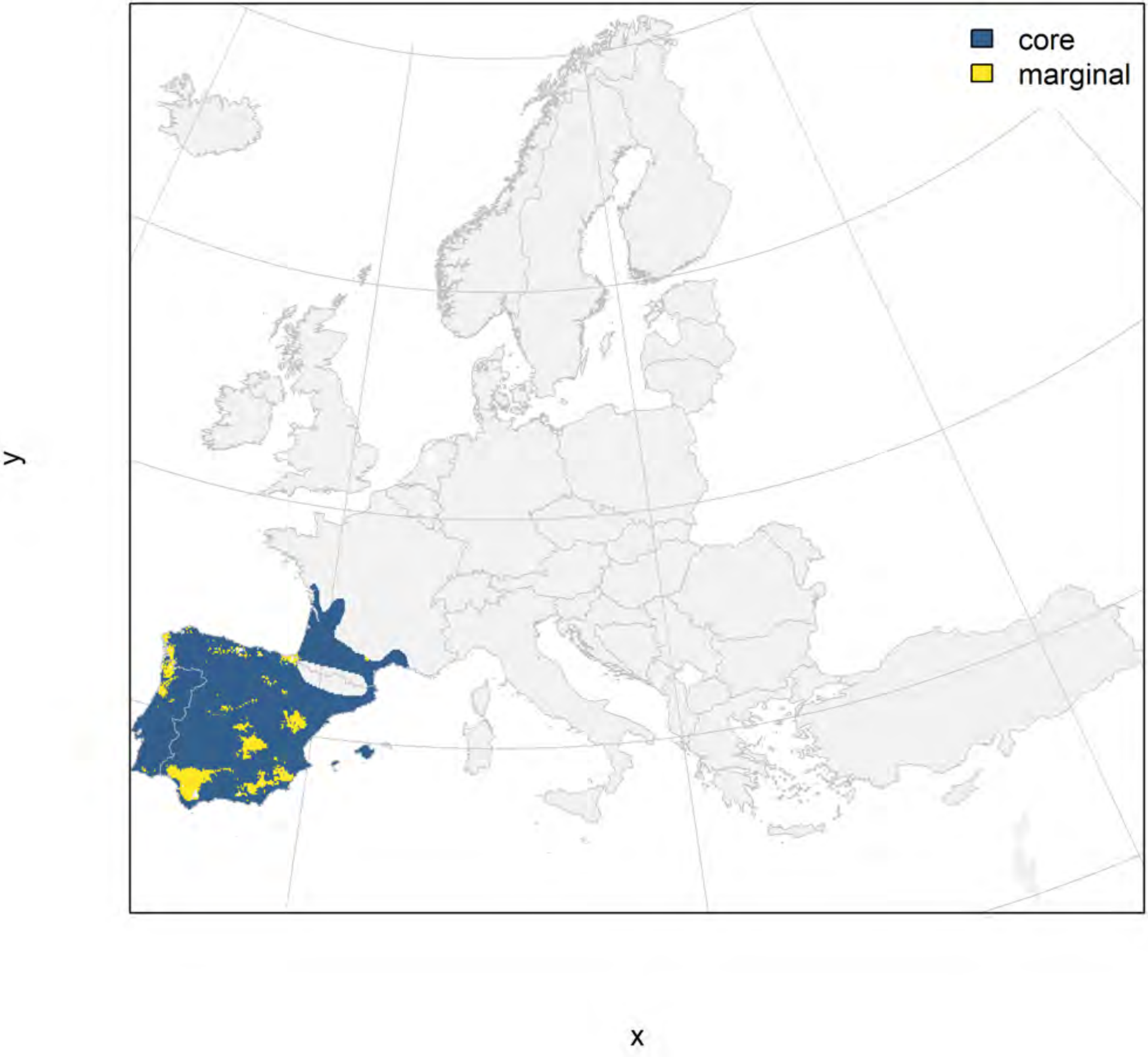

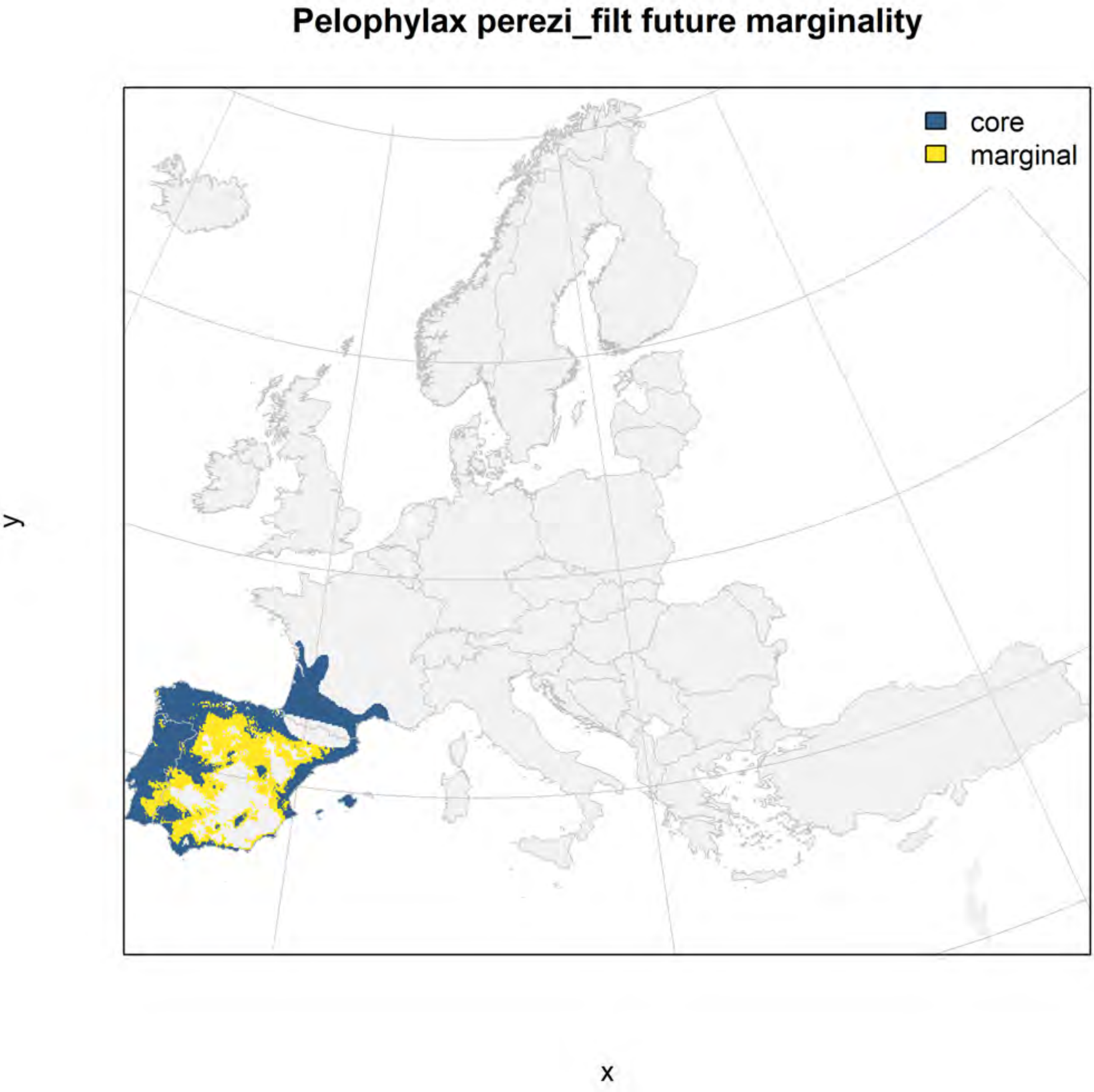
Pelophylax perezi_filt future marginality
core
marginal
y
x

### Pelophylax ridibundus_filt NMI

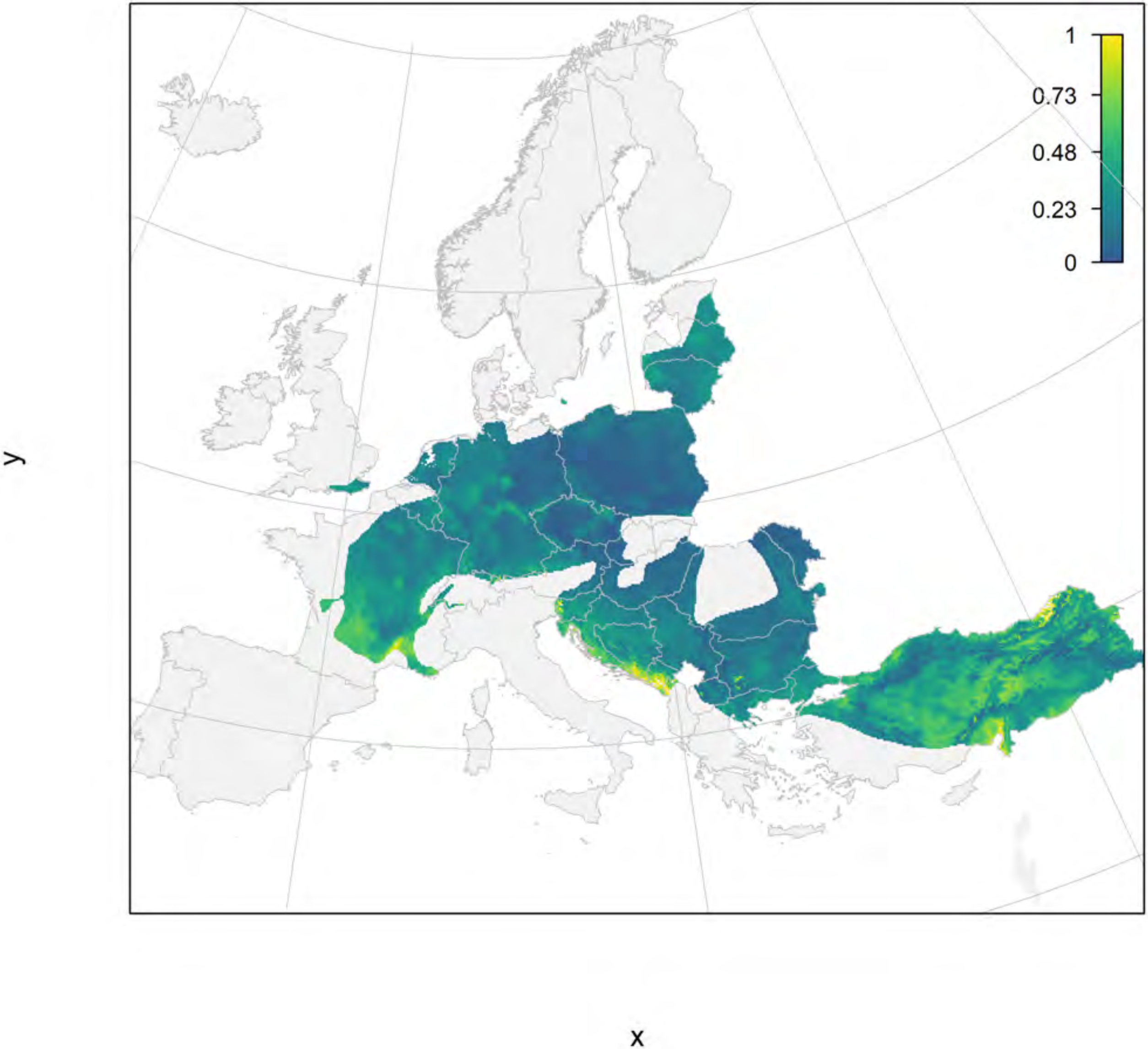

#### Pelophylax ridibundus_filt current marginality

y

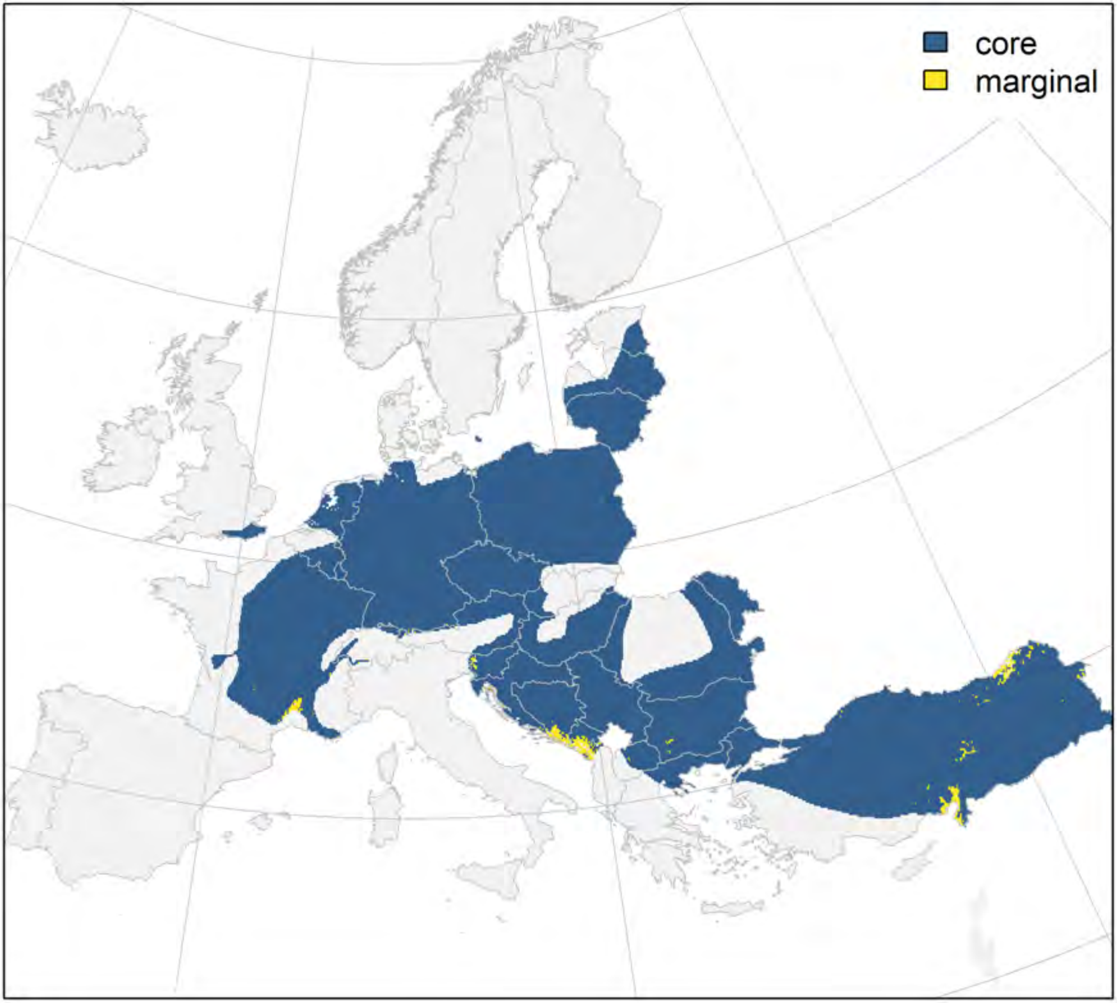

x

### Pelophylax ridibundus_filt future marginality

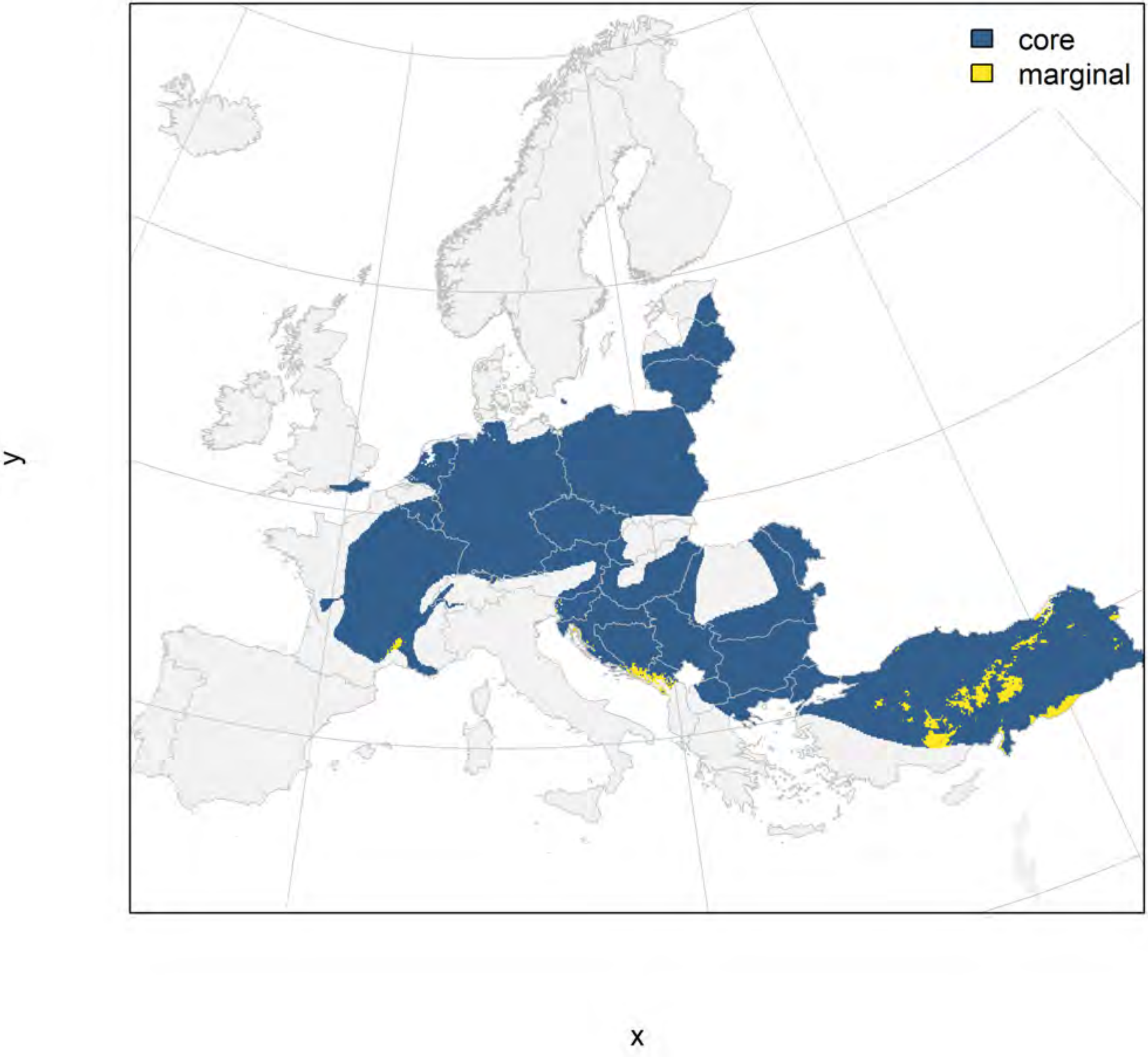

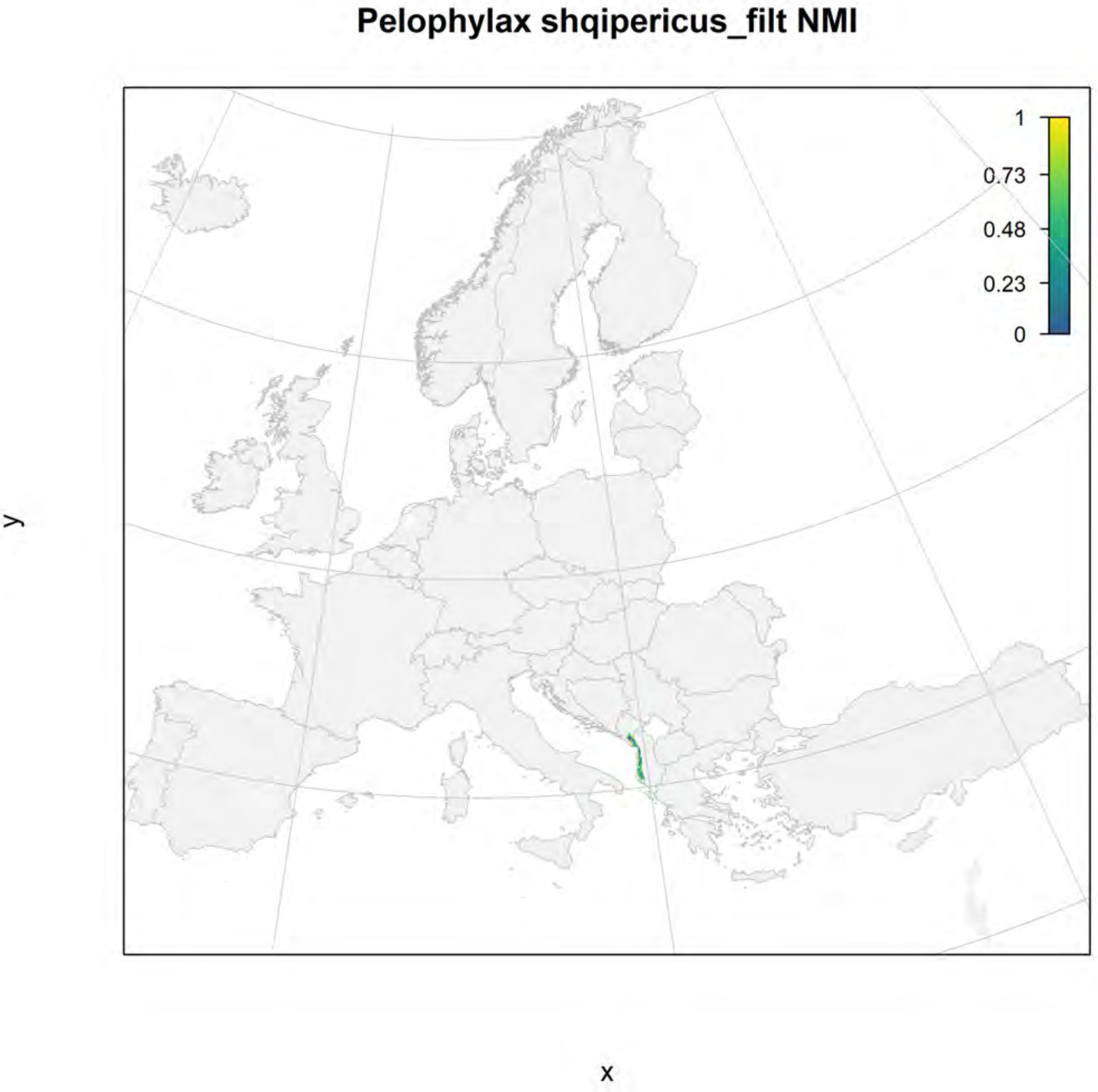

Pelophylax shqipericus_filt NMI
1
0.73
0.48
0.23
0
y
x

#### Pelophylax shqipericus_filt current marginality

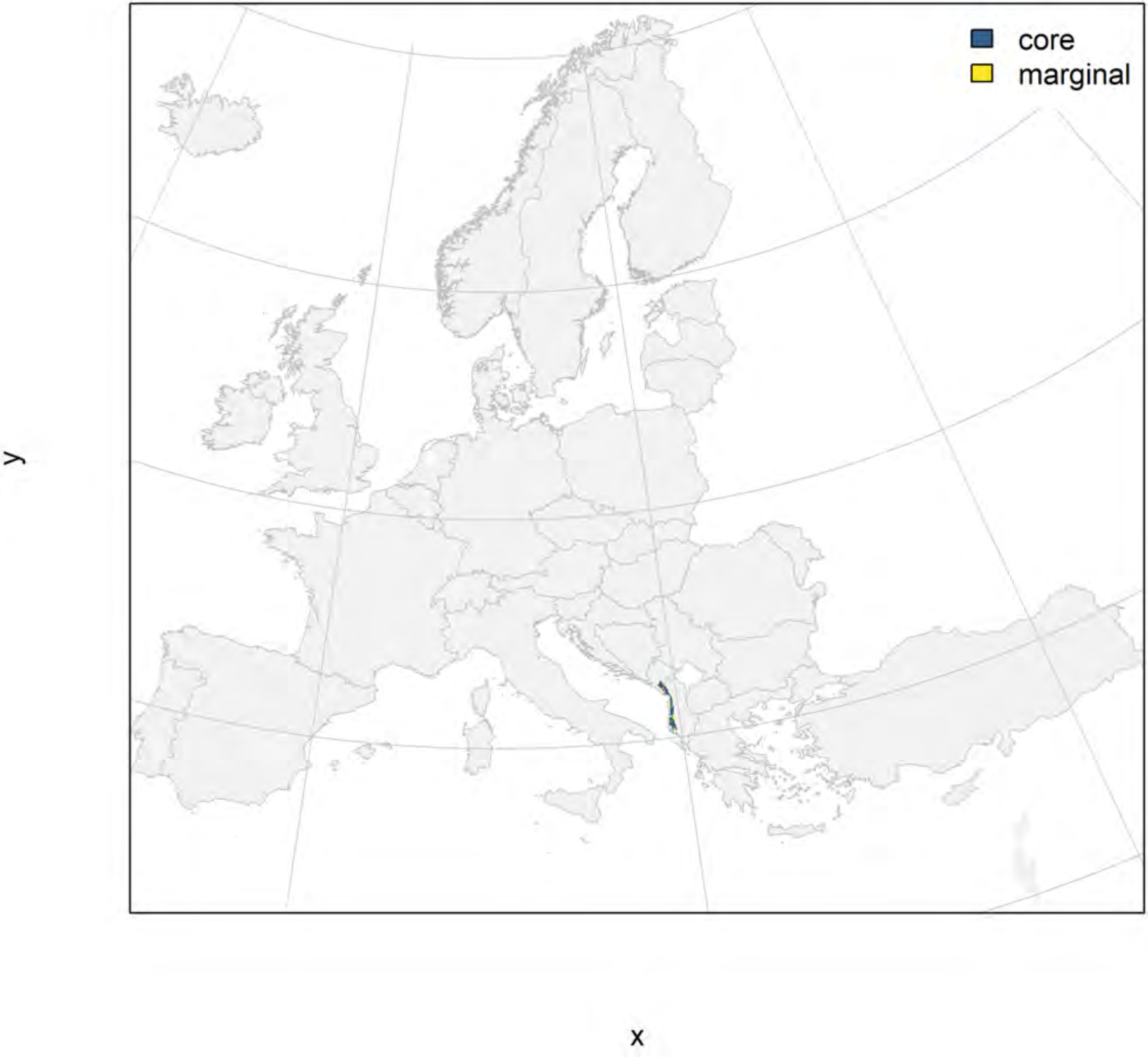

### Pelophylax shqipericus_filt future marginality

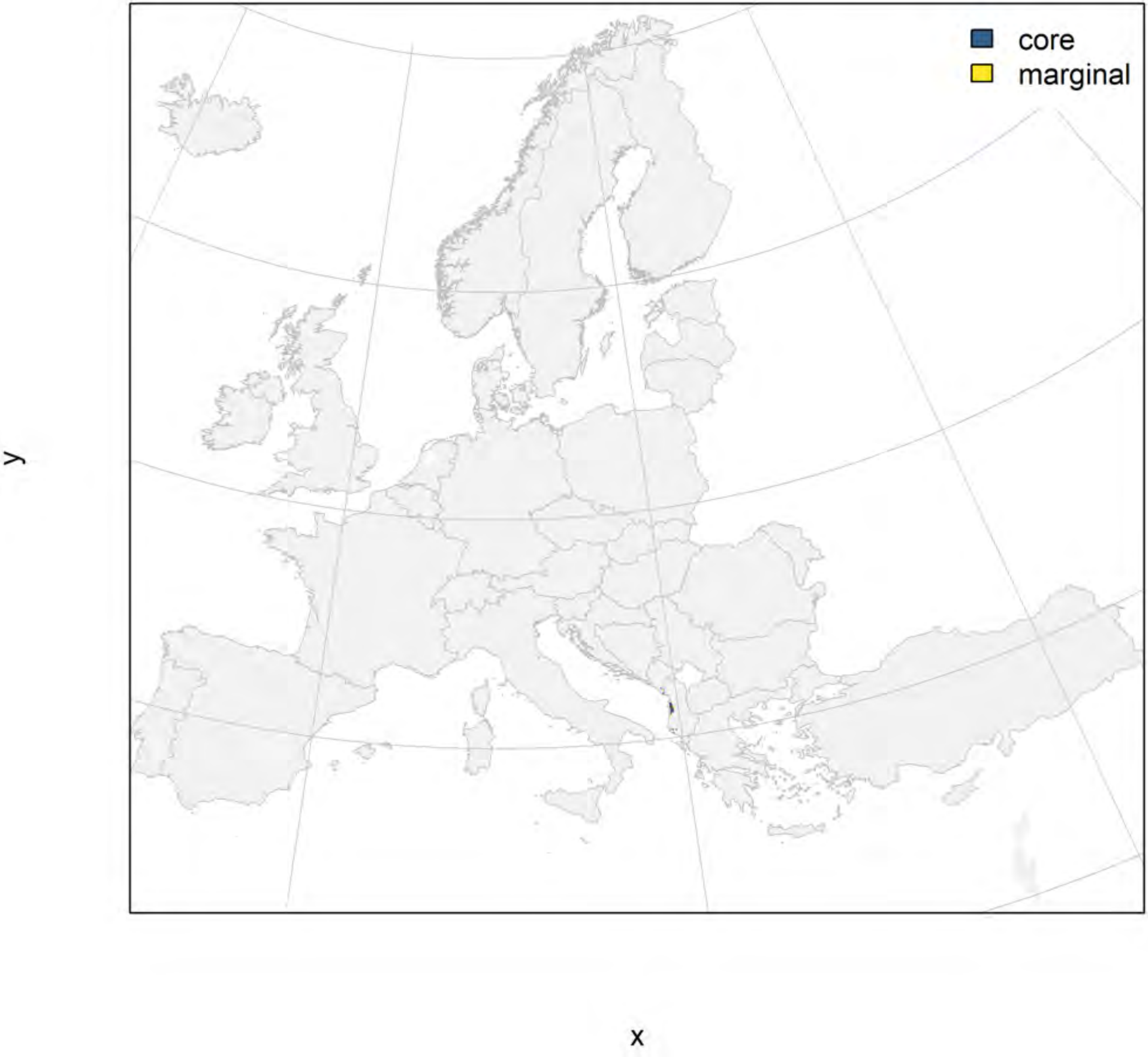

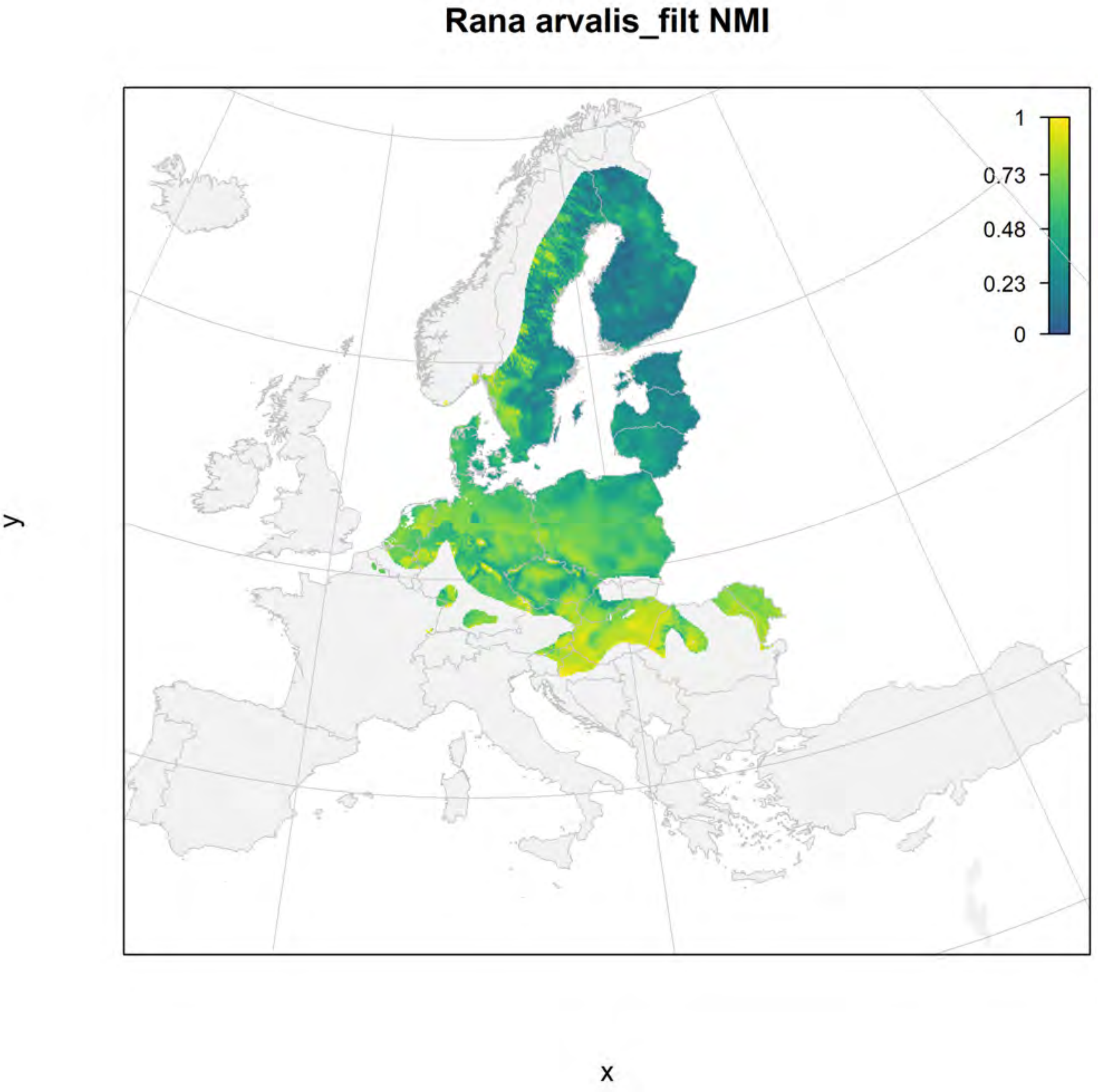

Rana arvalis_filt NMI
1
0.73
0.48
0.23
0
y
x

#### Rana arvalis_filt current marginality

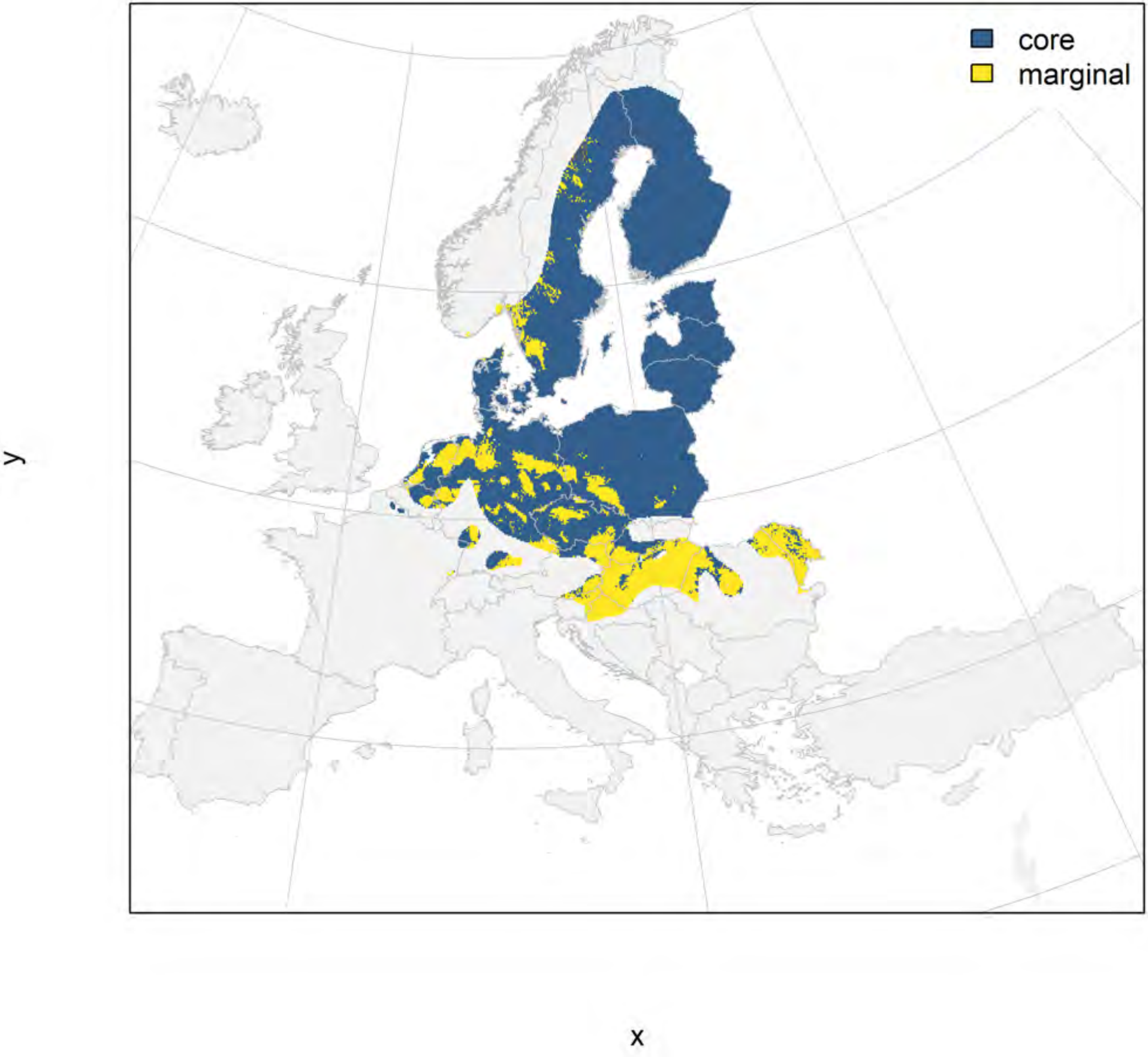

Rana arvalis_filt future marginality

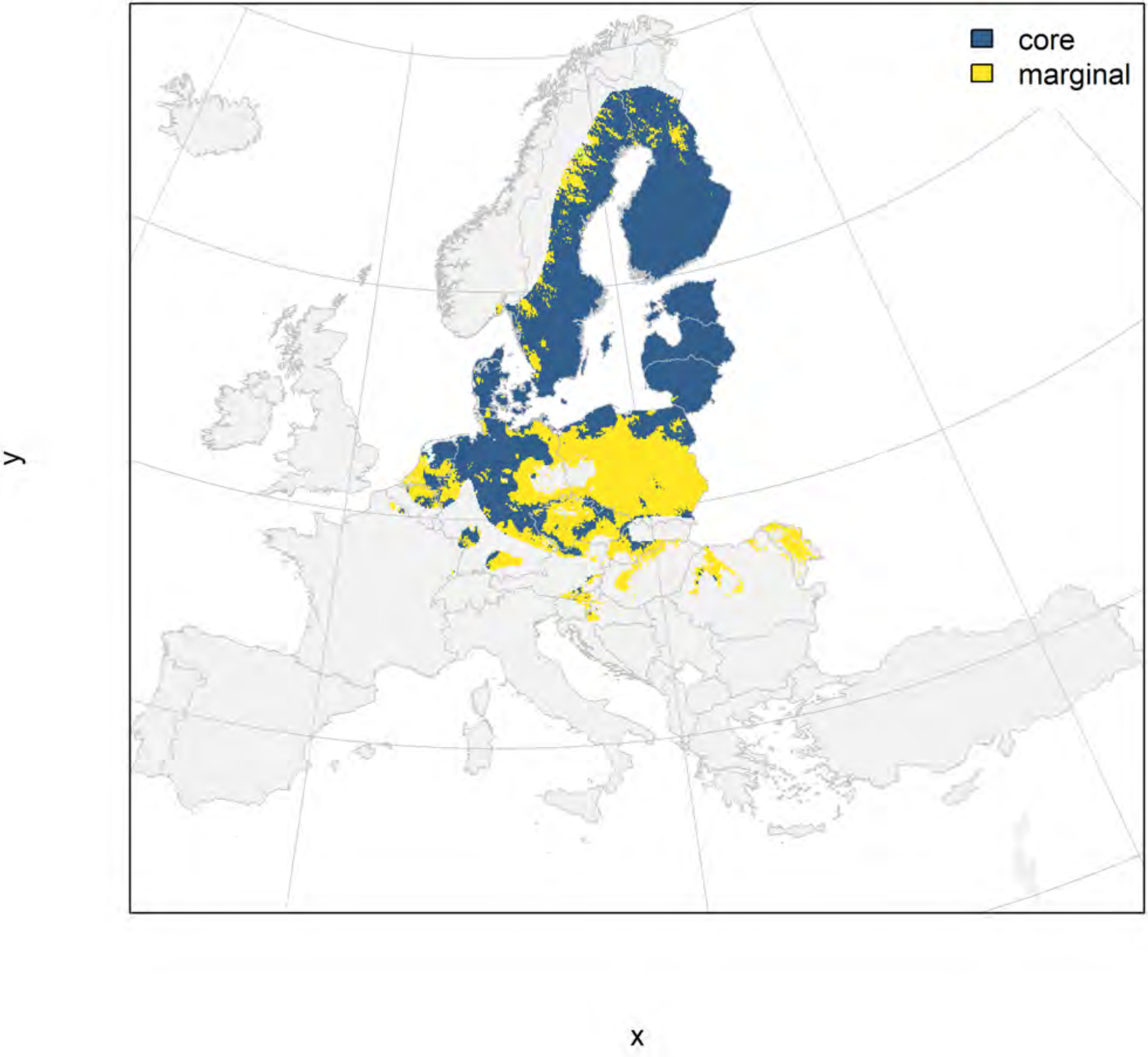

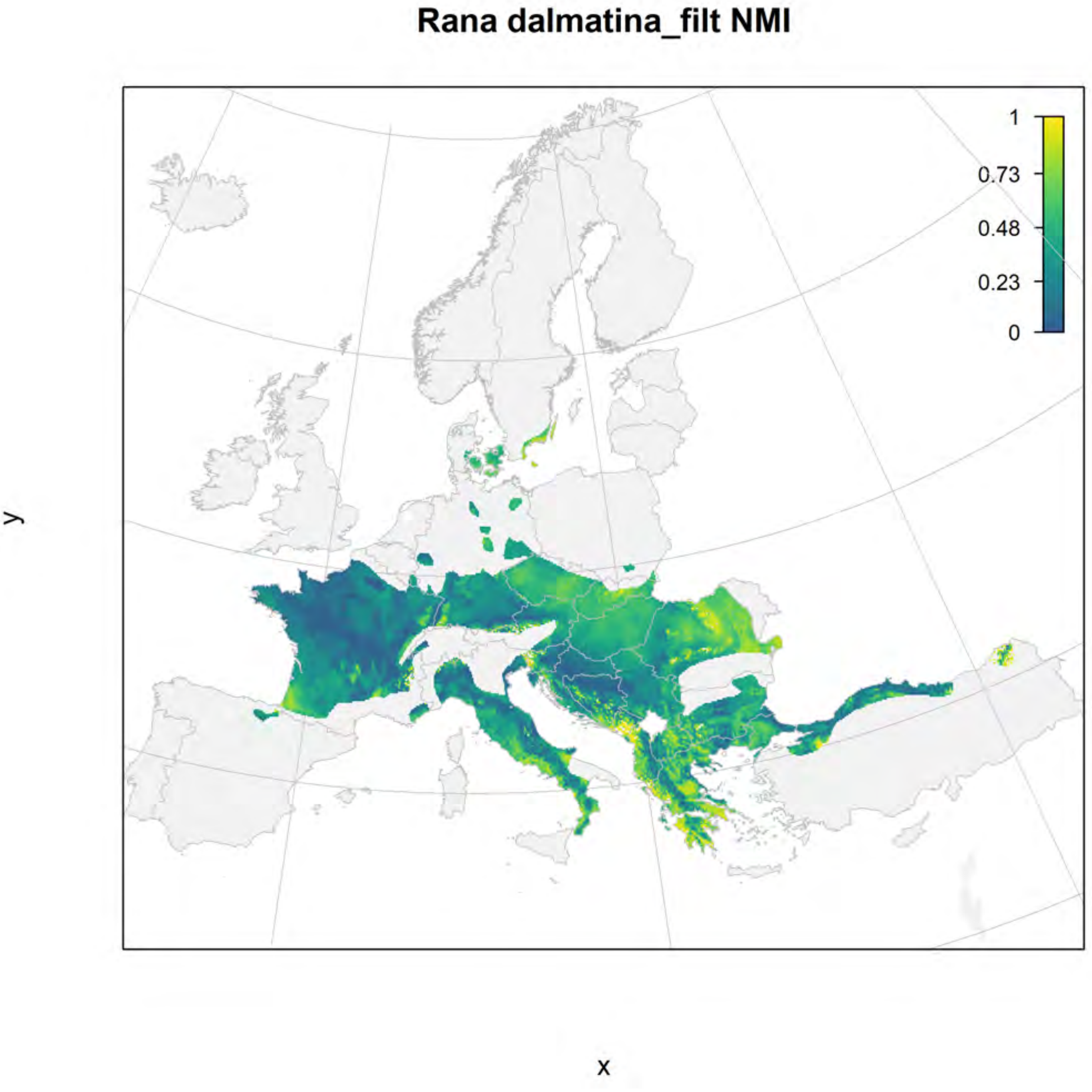
Rana dalmatina_filt NMI
1
0.73
0.48
0.23
0
y
x

#### Rana dalmatina_filt current marginality

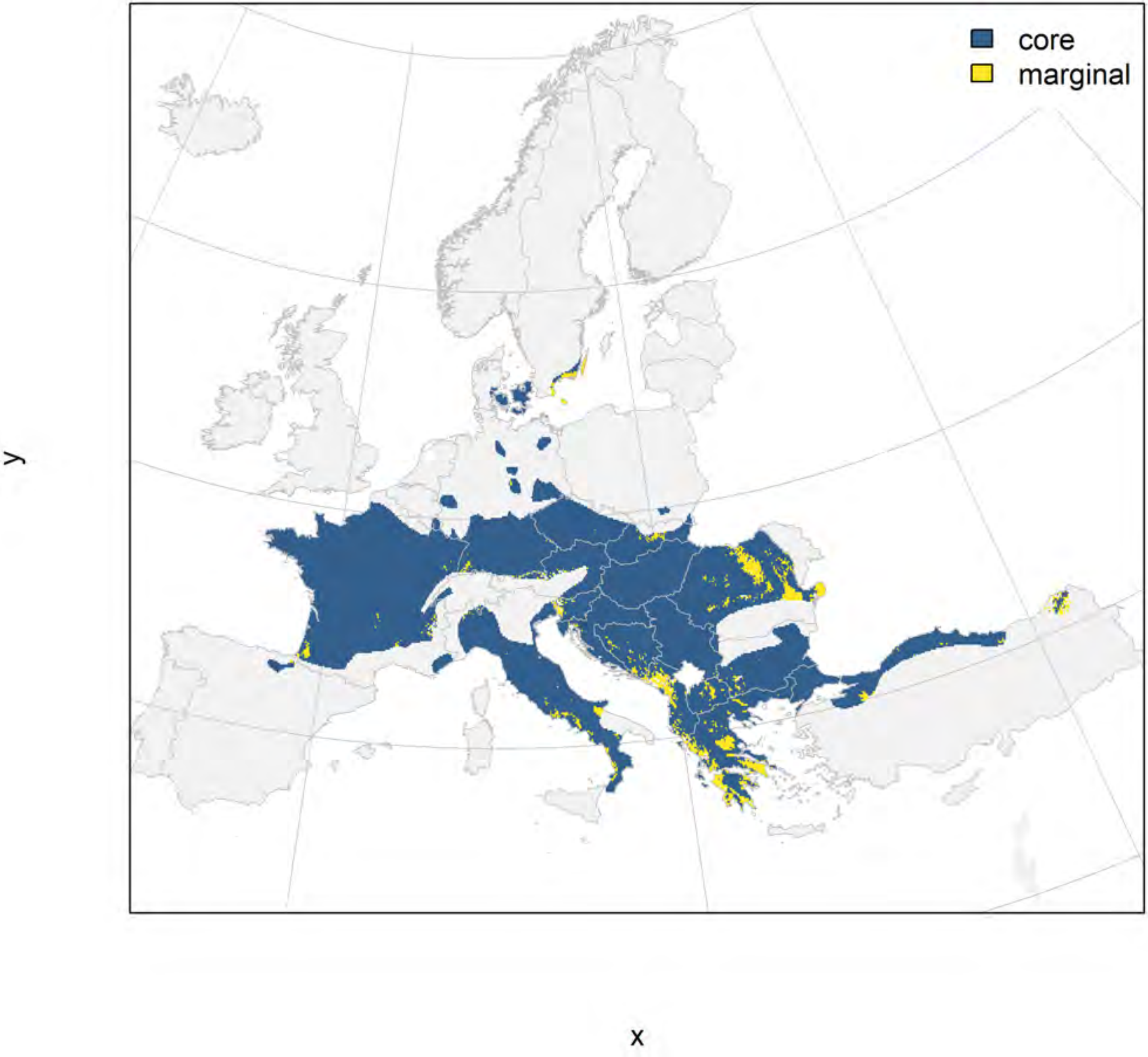

#### Rana dalmatina_filt future marginality

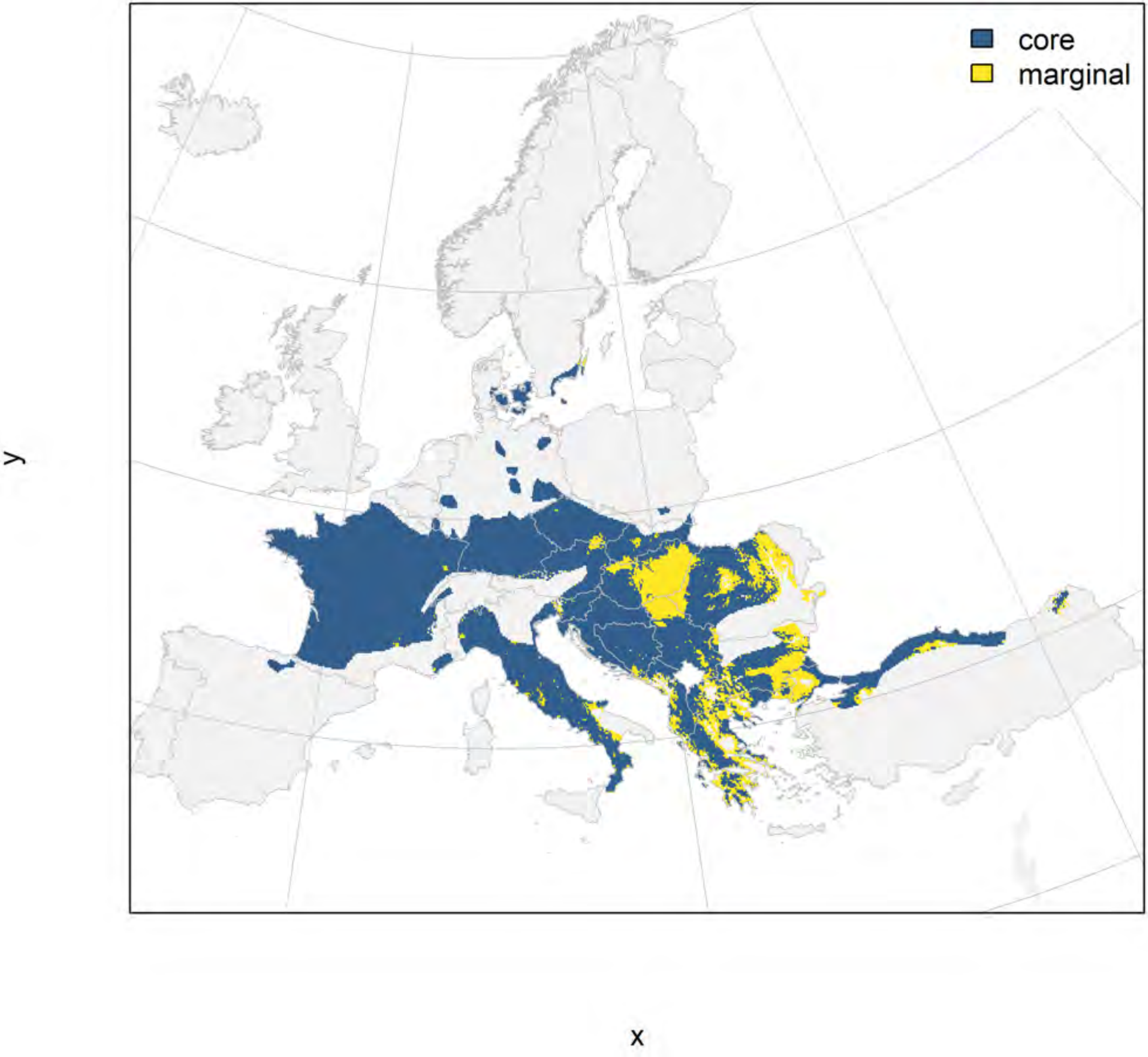

### Rana graeca_filt NMI

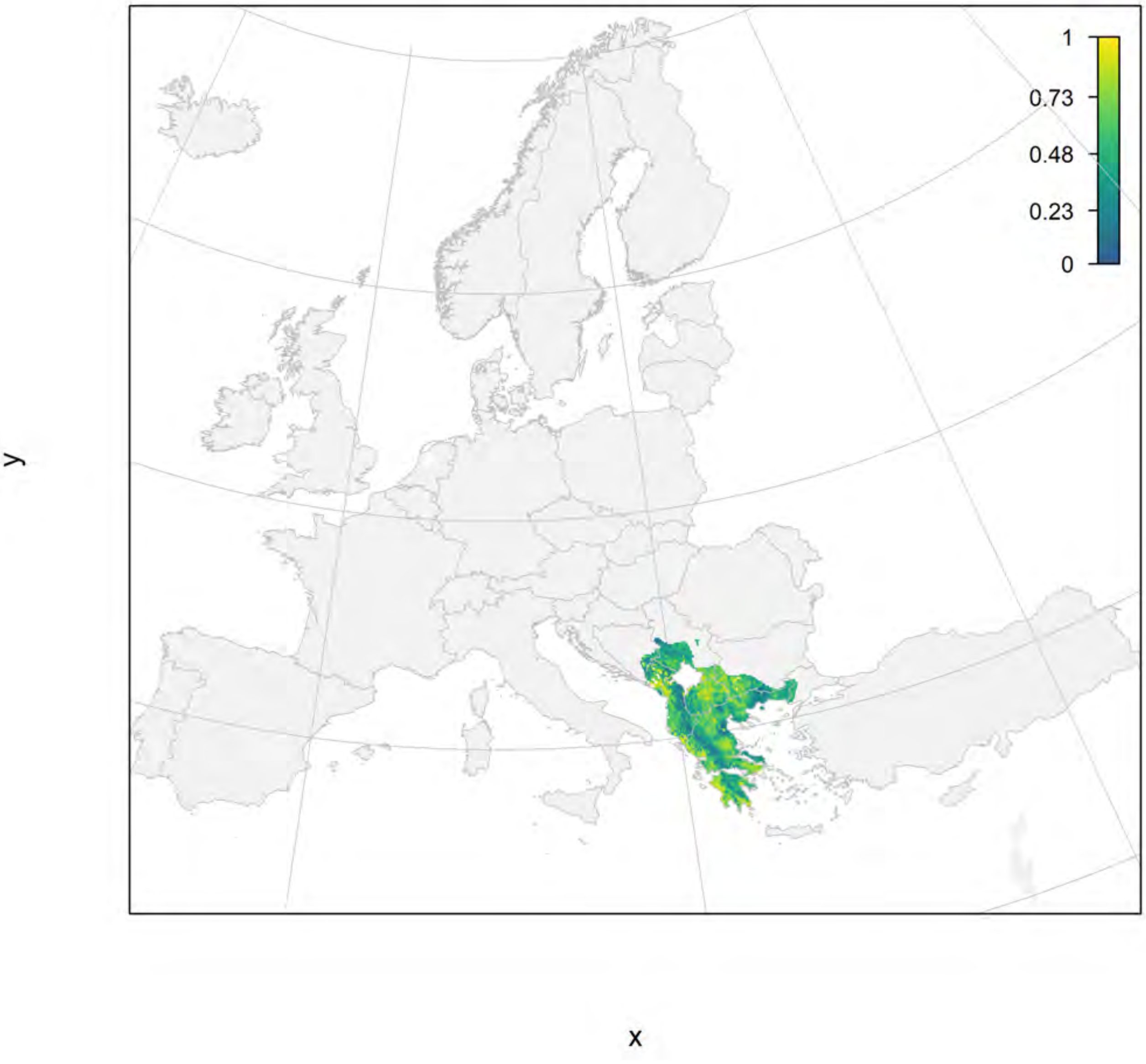

Rana graeca_filt current marginality

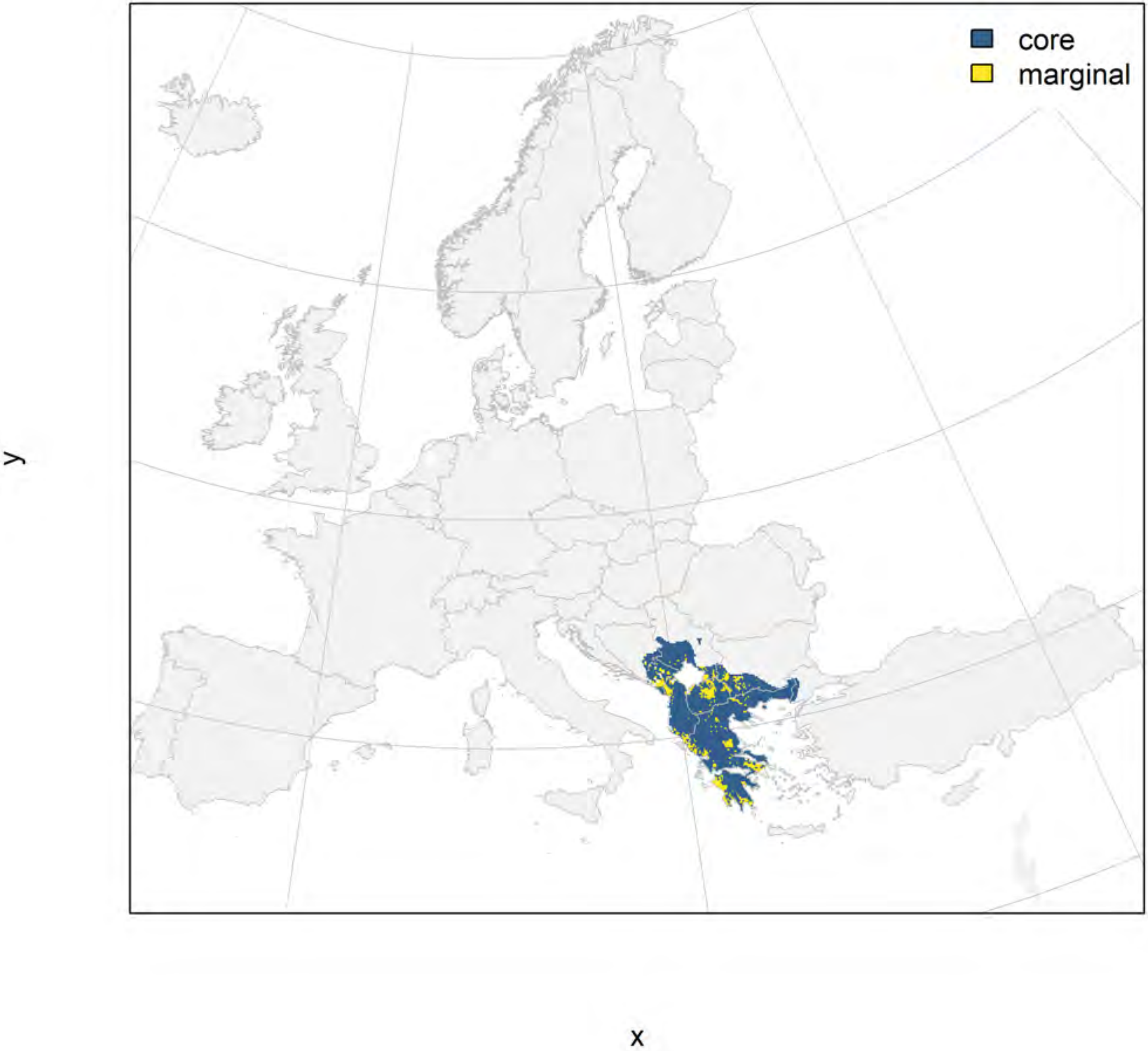

#### Rana graeca_filt future marginality

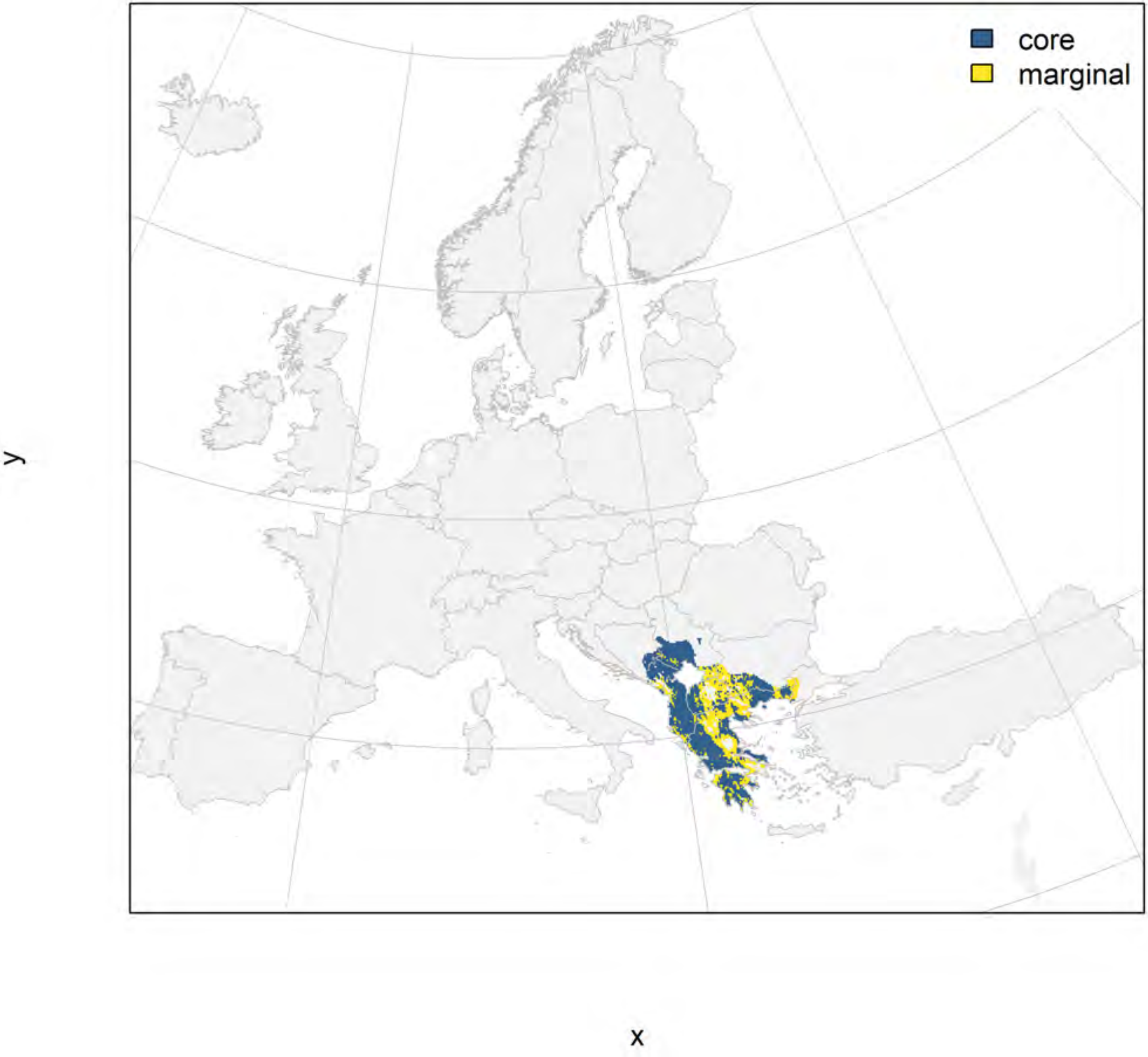

### Rana iberica_filt NMI

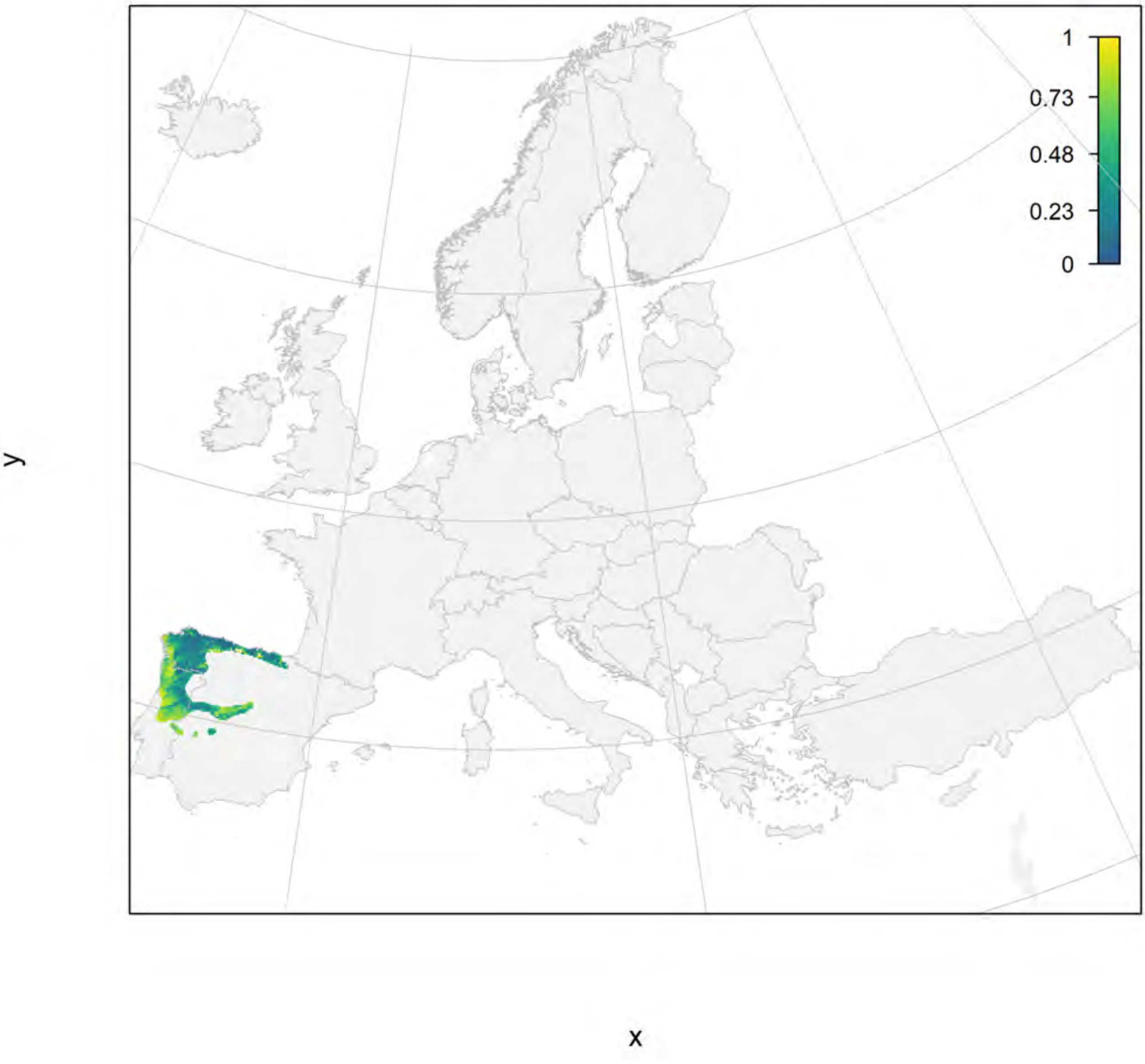

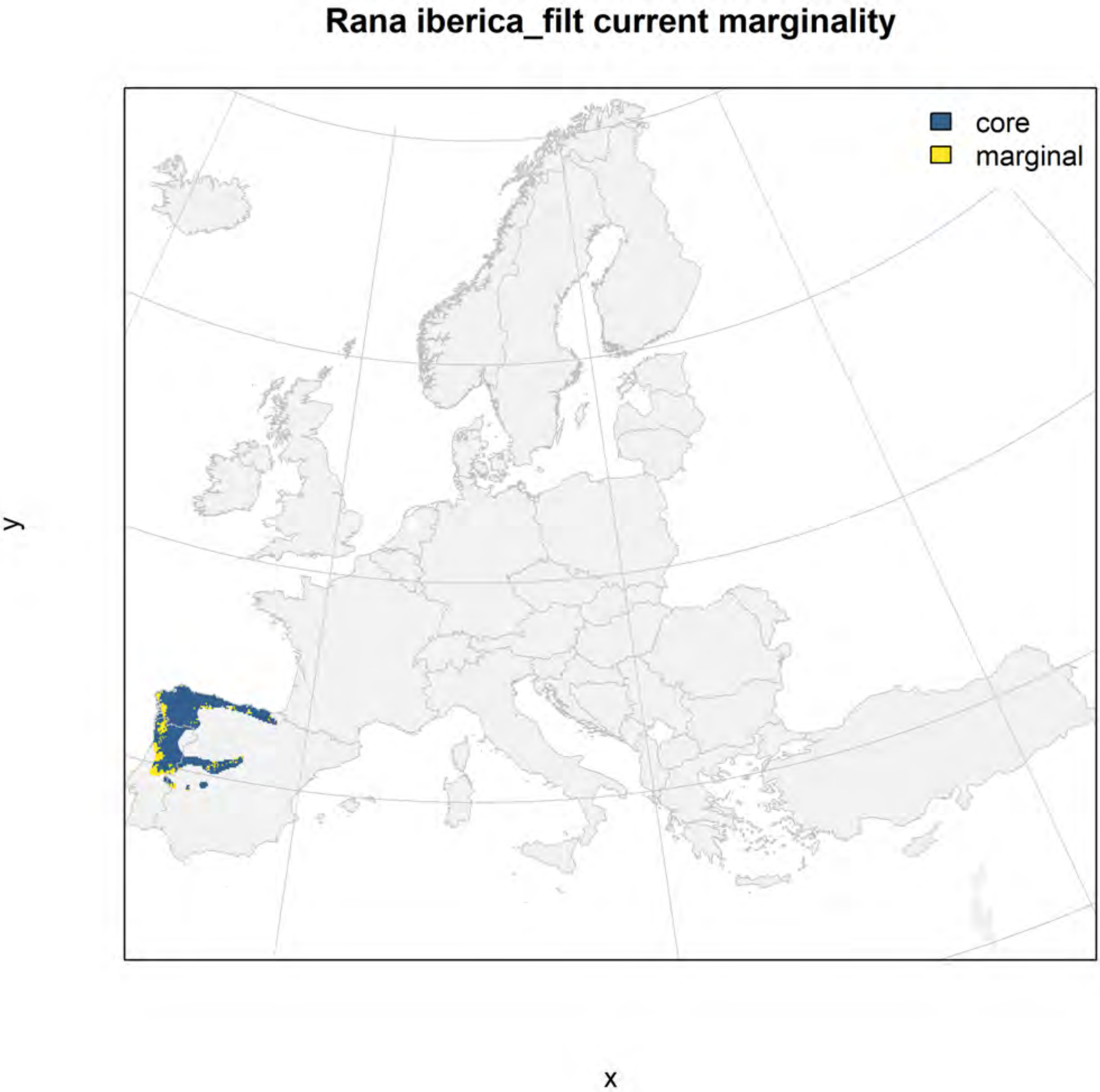

Rana iberica_filt current marginality
core
marginal
y
x

### Rana iberica_filt future marginality

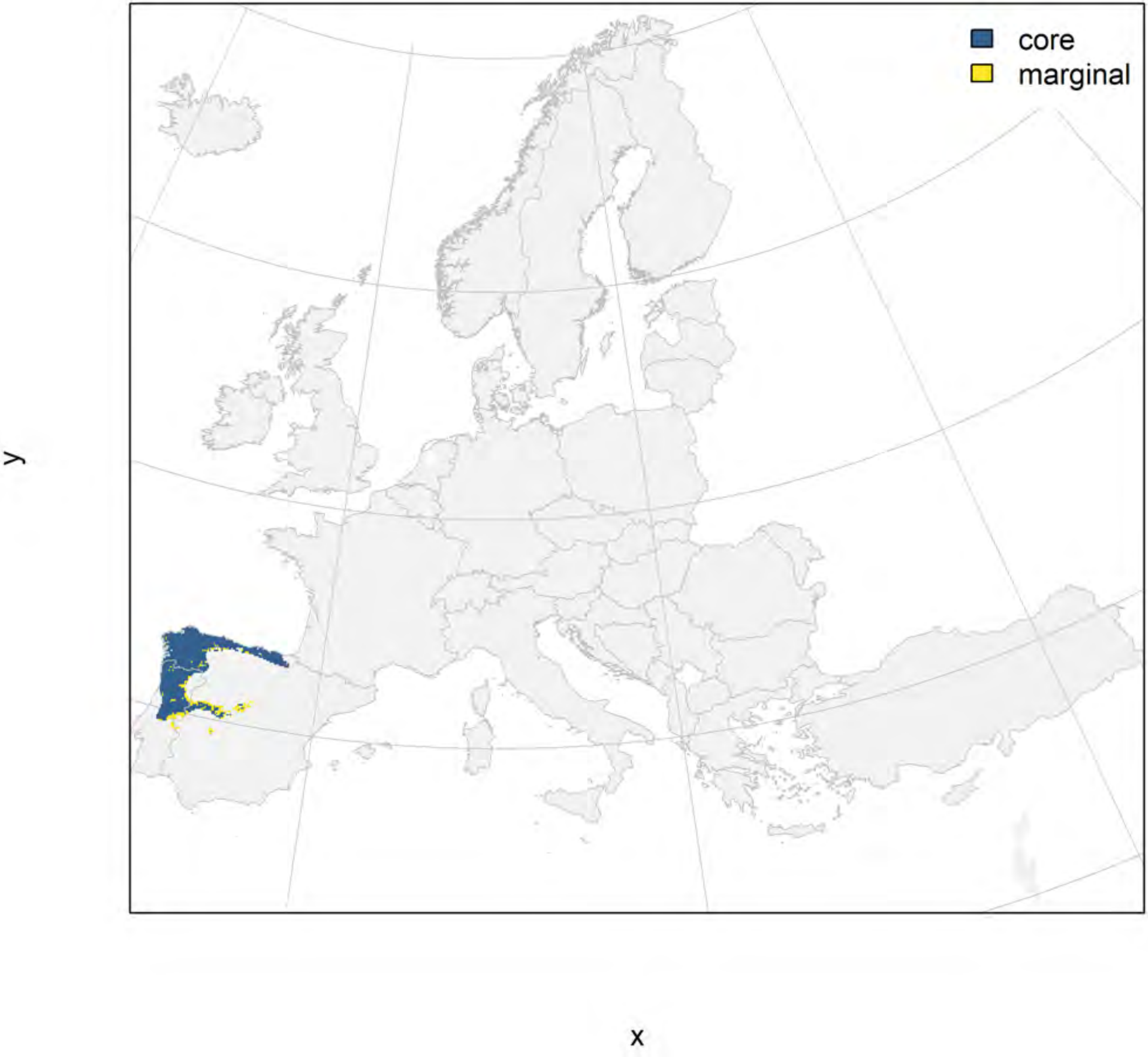

### Rana italica_filt NMI

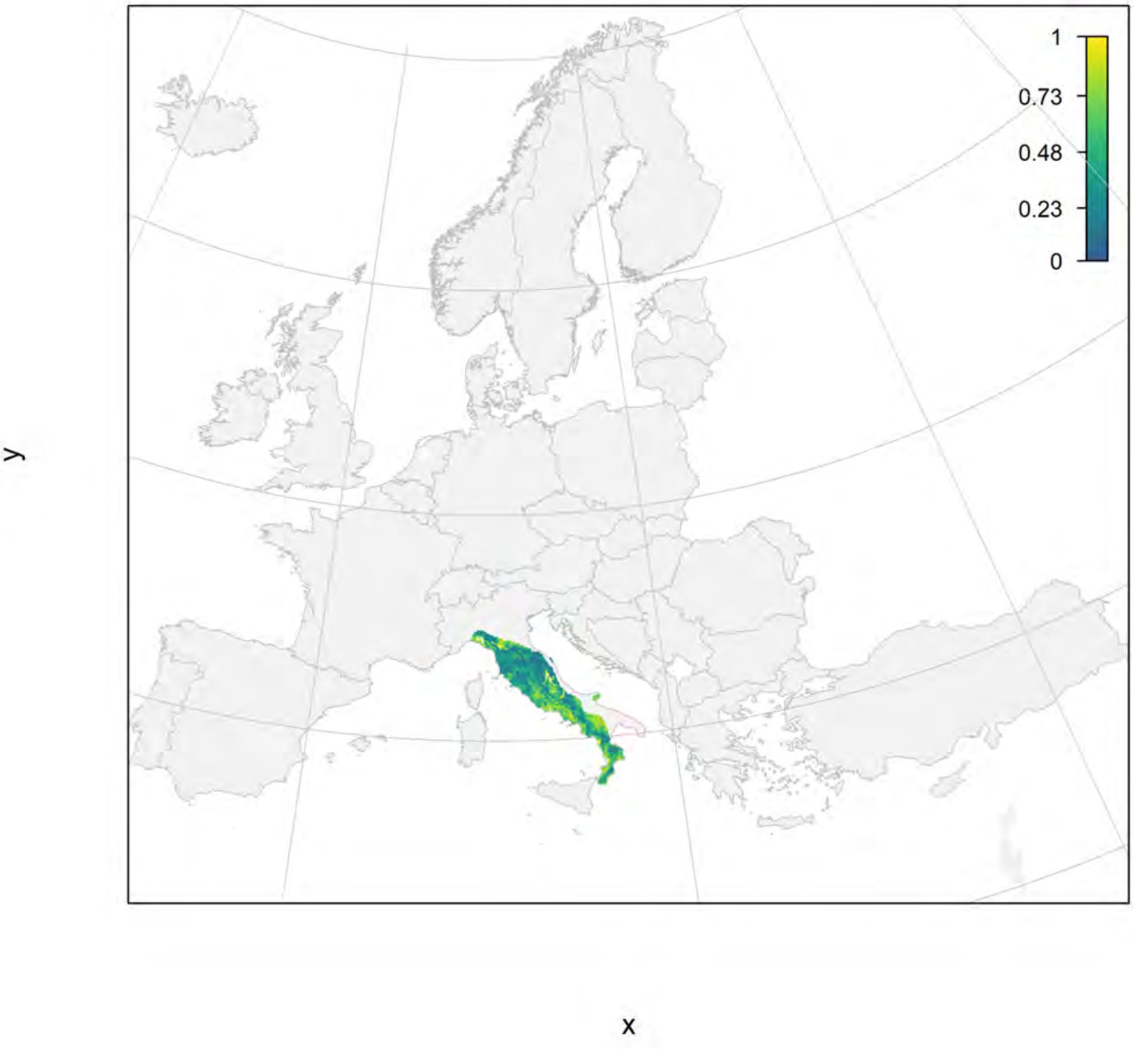

### Rana italica_filt current marginality

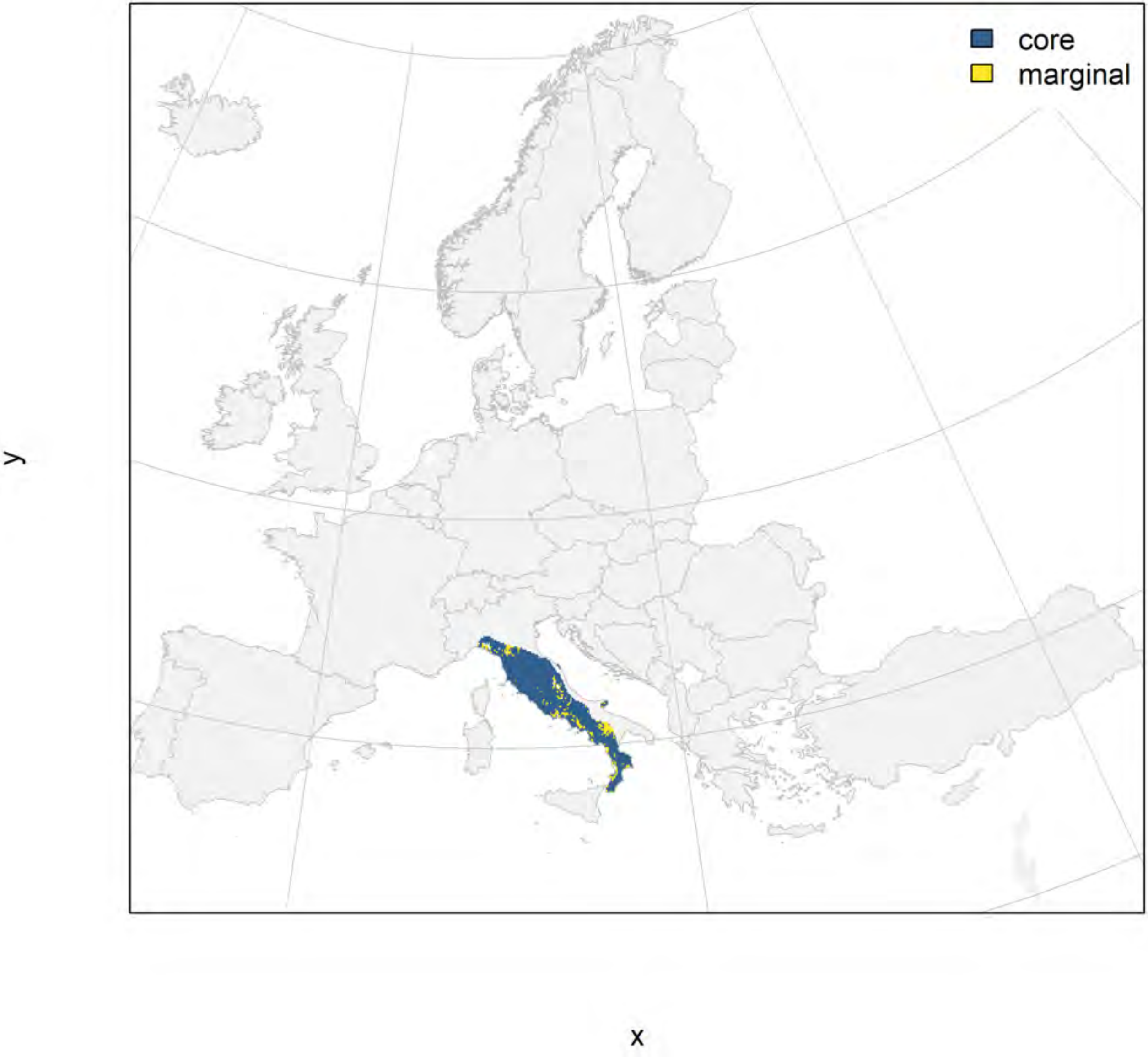

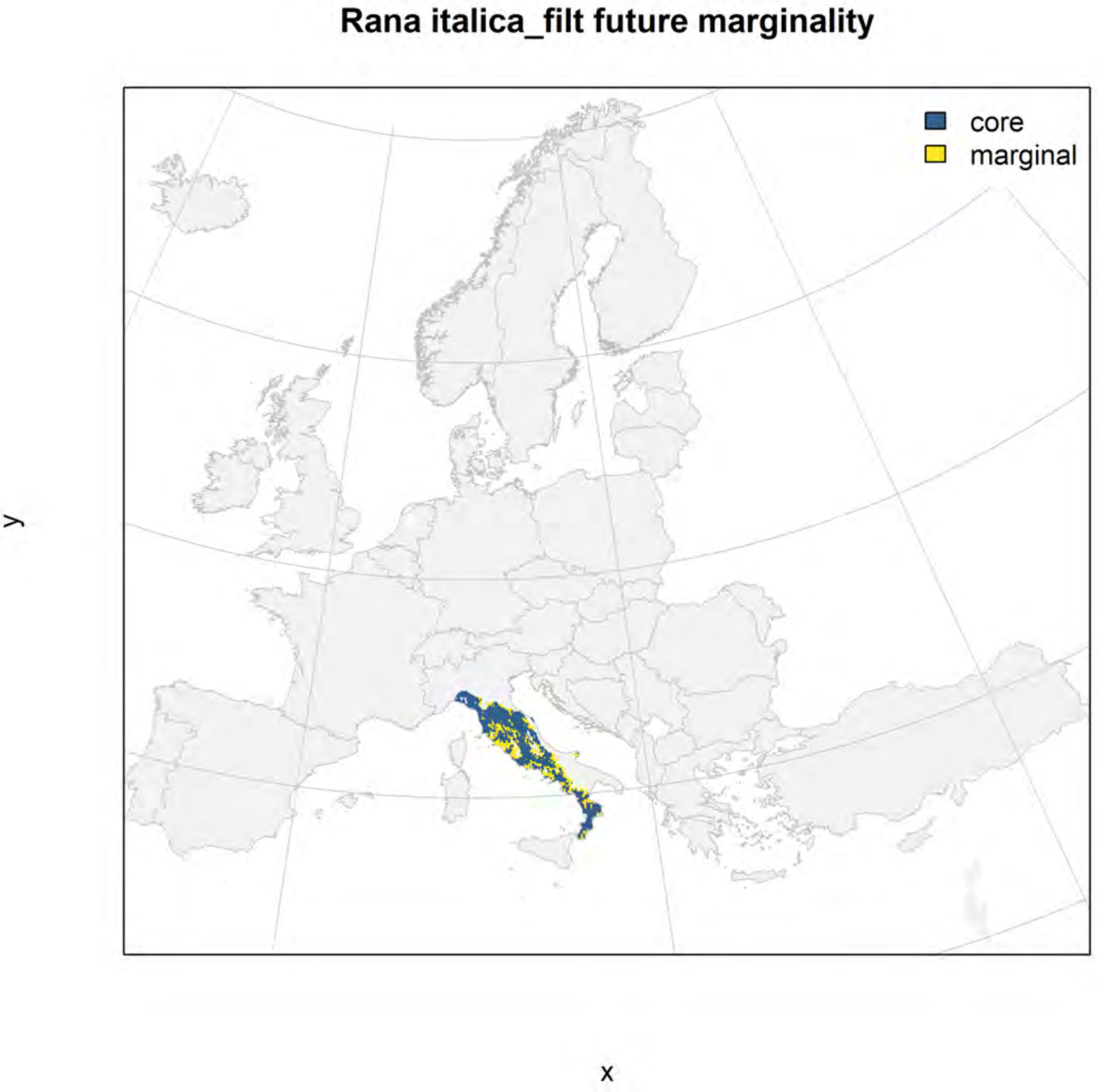

Rana italica_filt future marginality
core
marginal
y
x

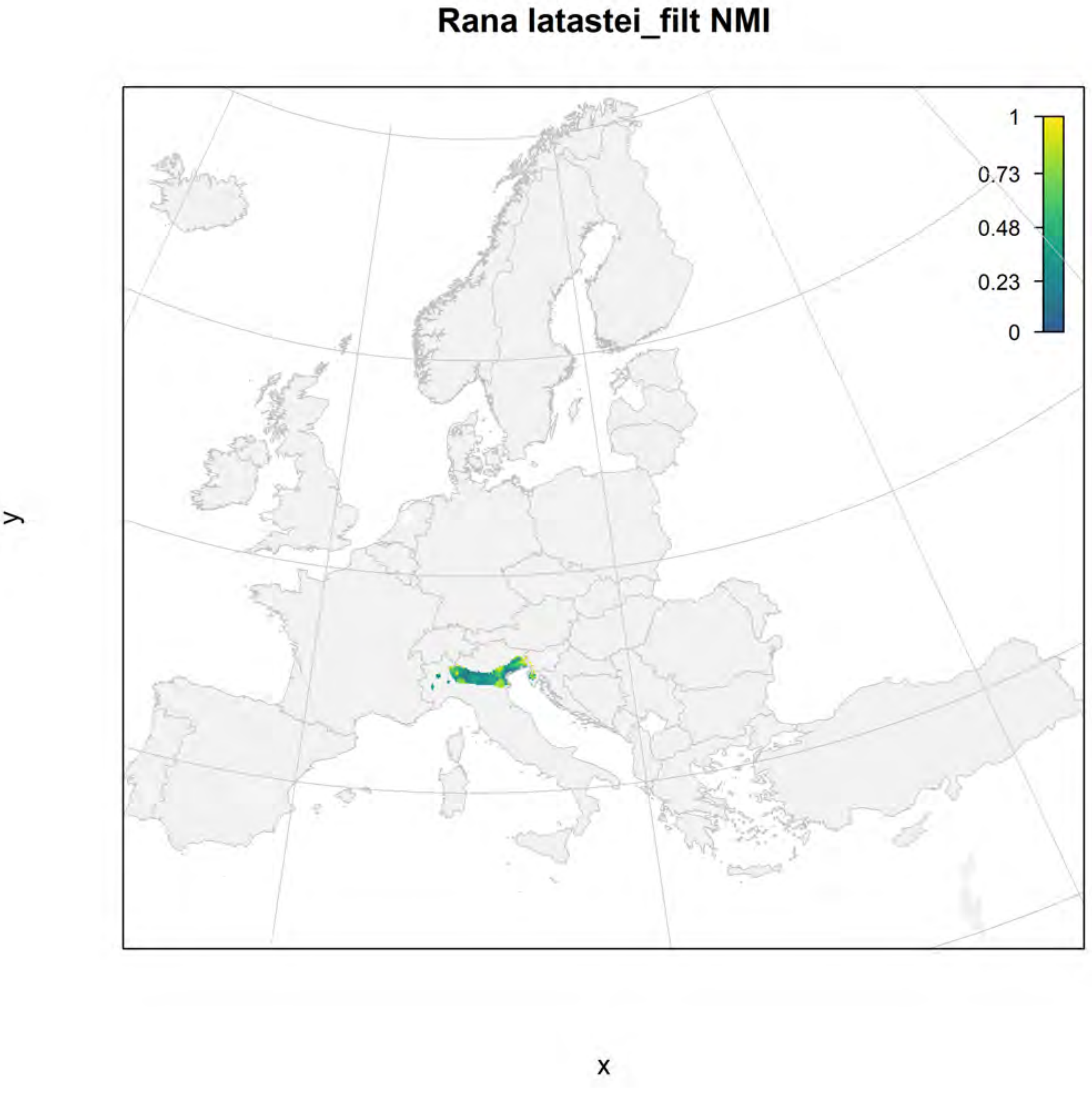

Rana latastei_filt NMI
1
0.73
0.48
0.23
0
y
x

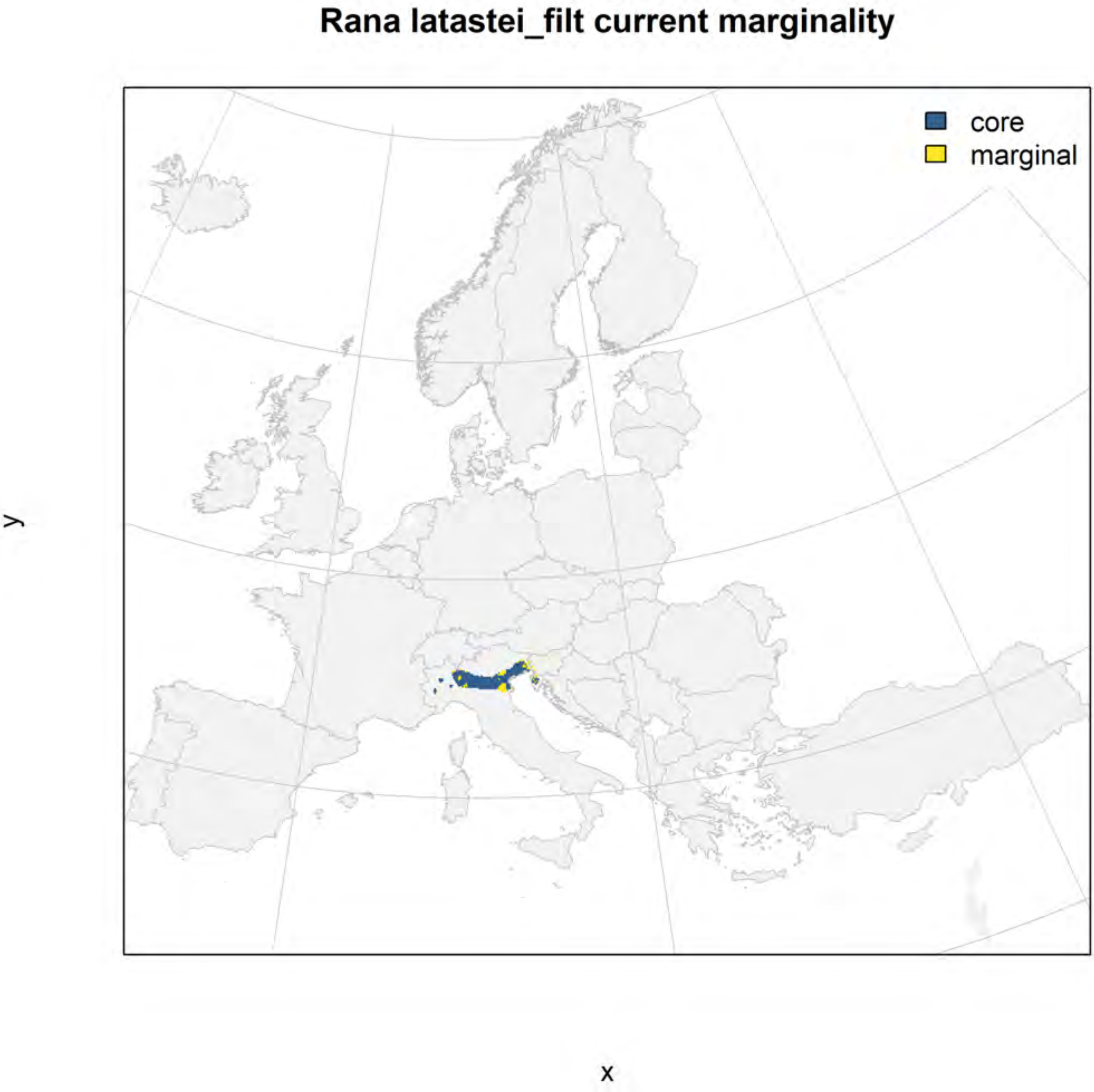
Rana latastei_filt current marginality
core
marginal
y
x

Rana latastei_filt future marginality

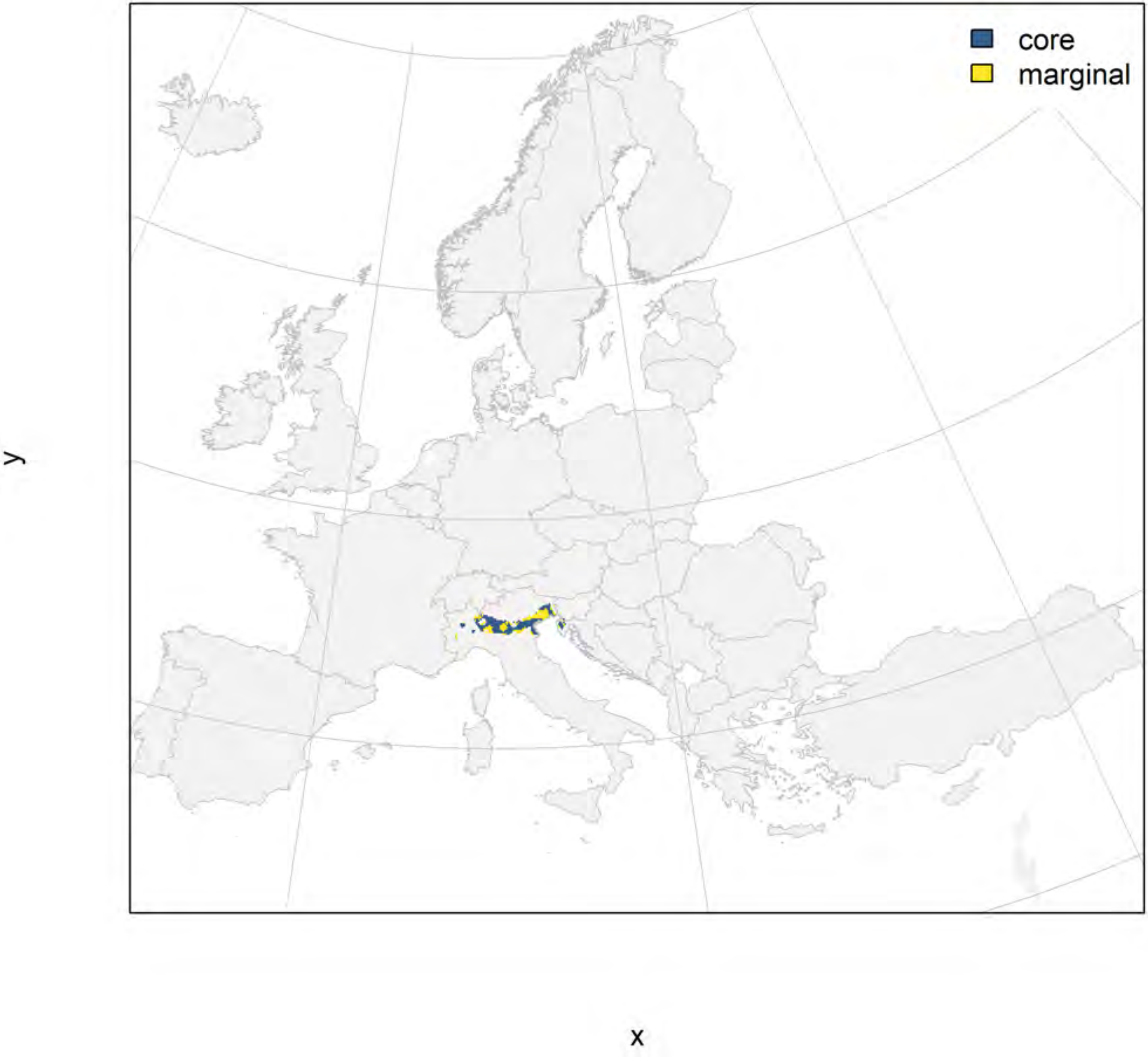

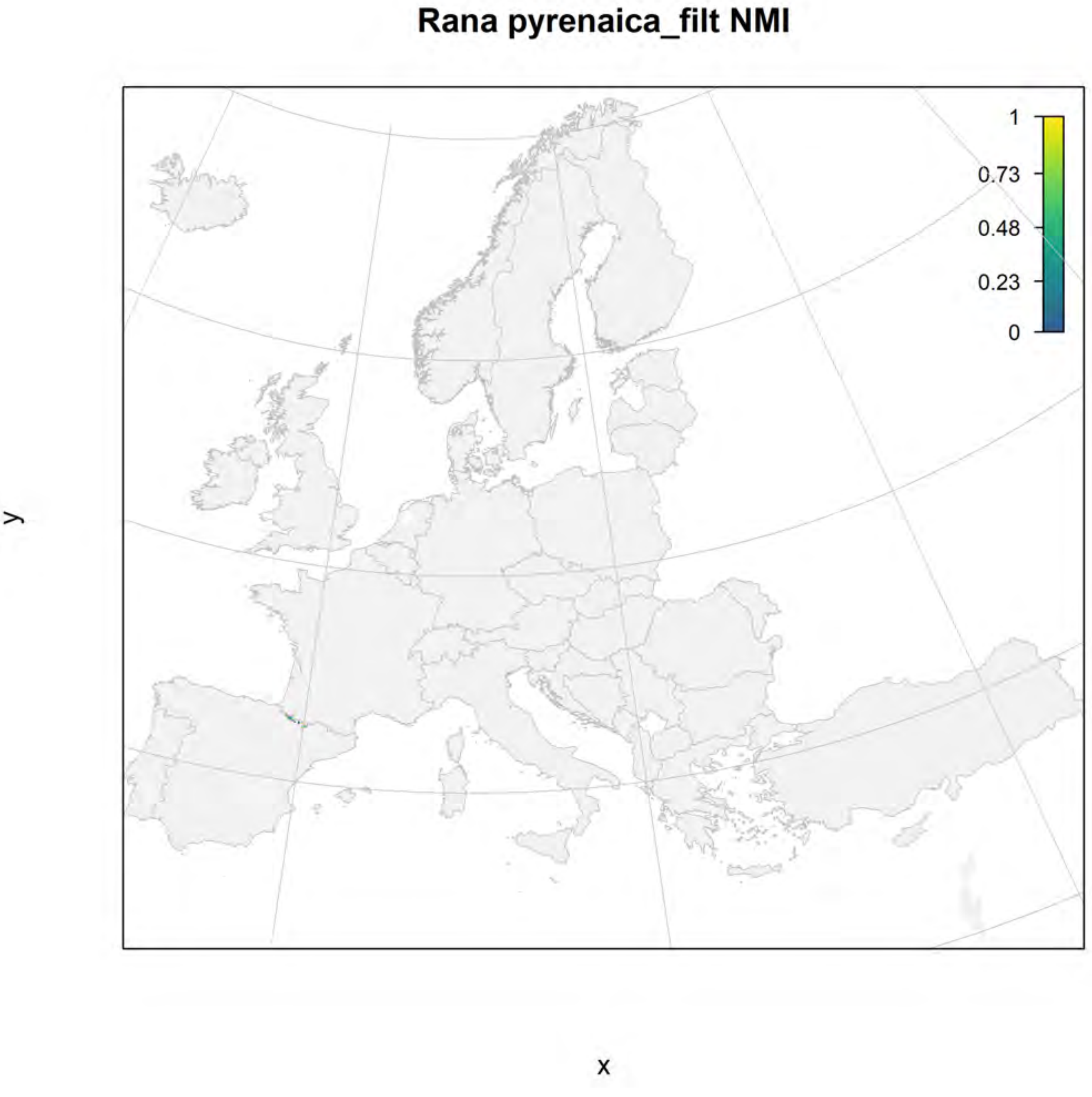

Rana pyrenaica_filt NMI
1
0.73
0.48
0.23
0
y
x

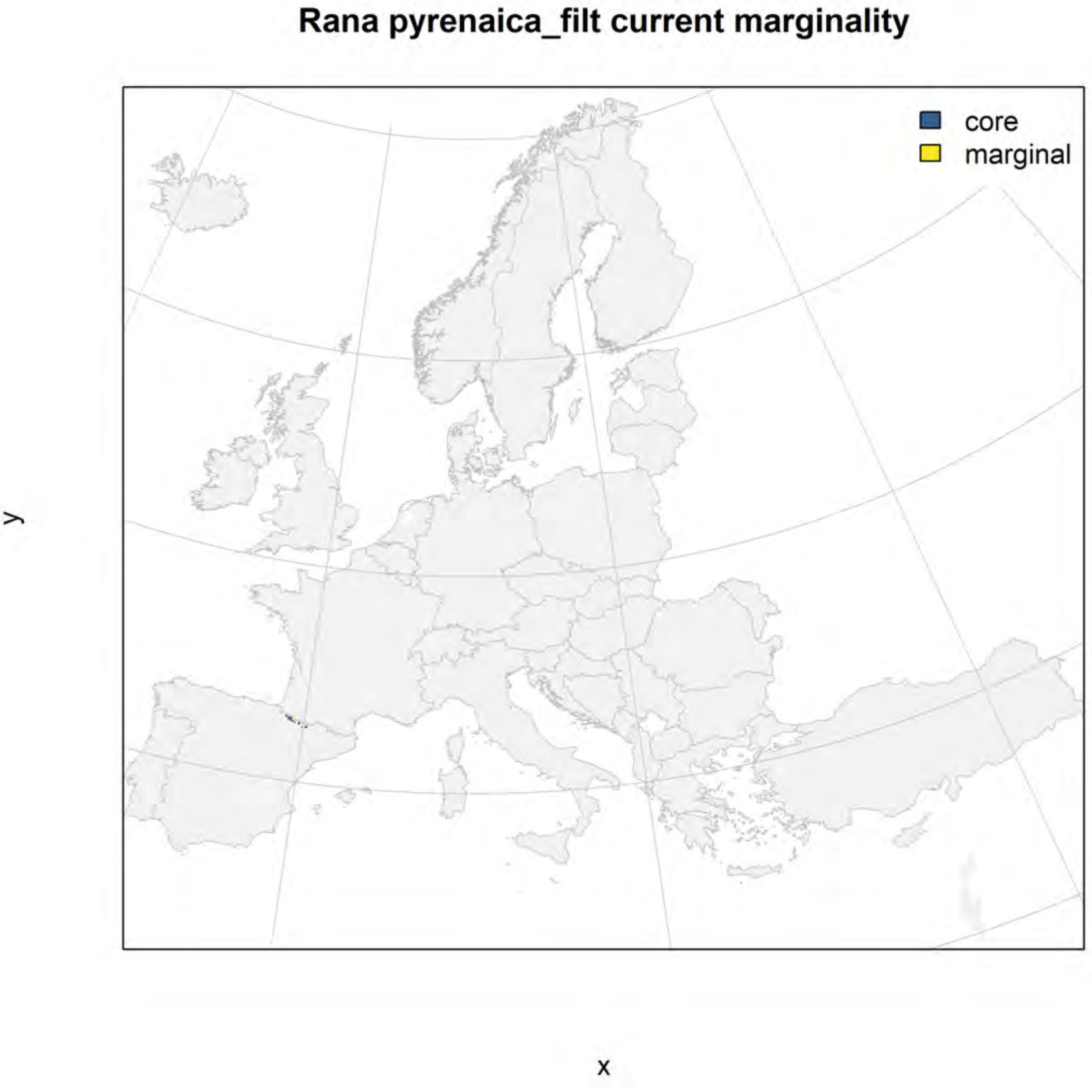
Rana pyrenaica_filt current marginality
core
marginal
y
x

### Rana pyrenaica_filt future marginality

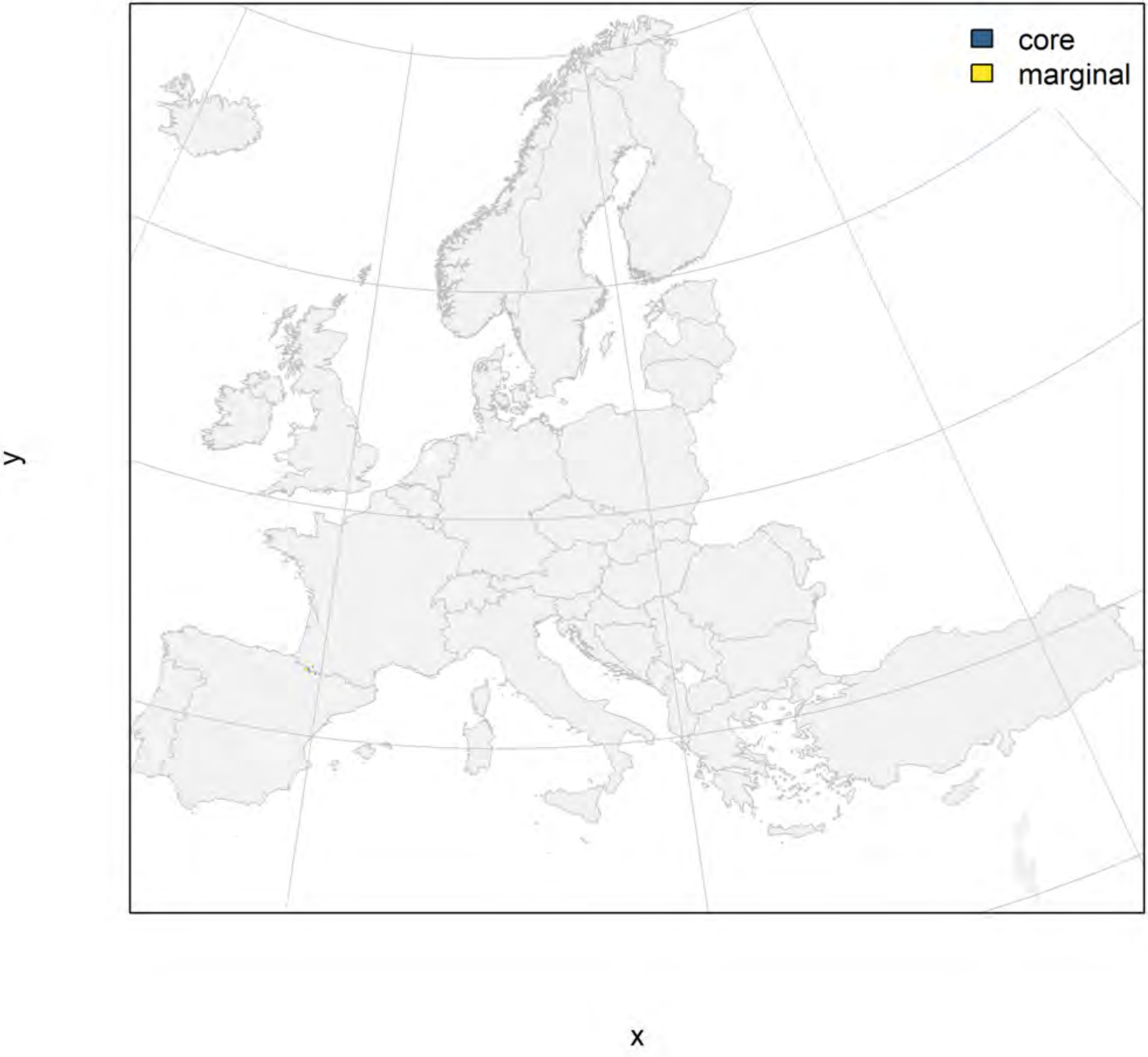

**Rana temporaria_filt NMI**

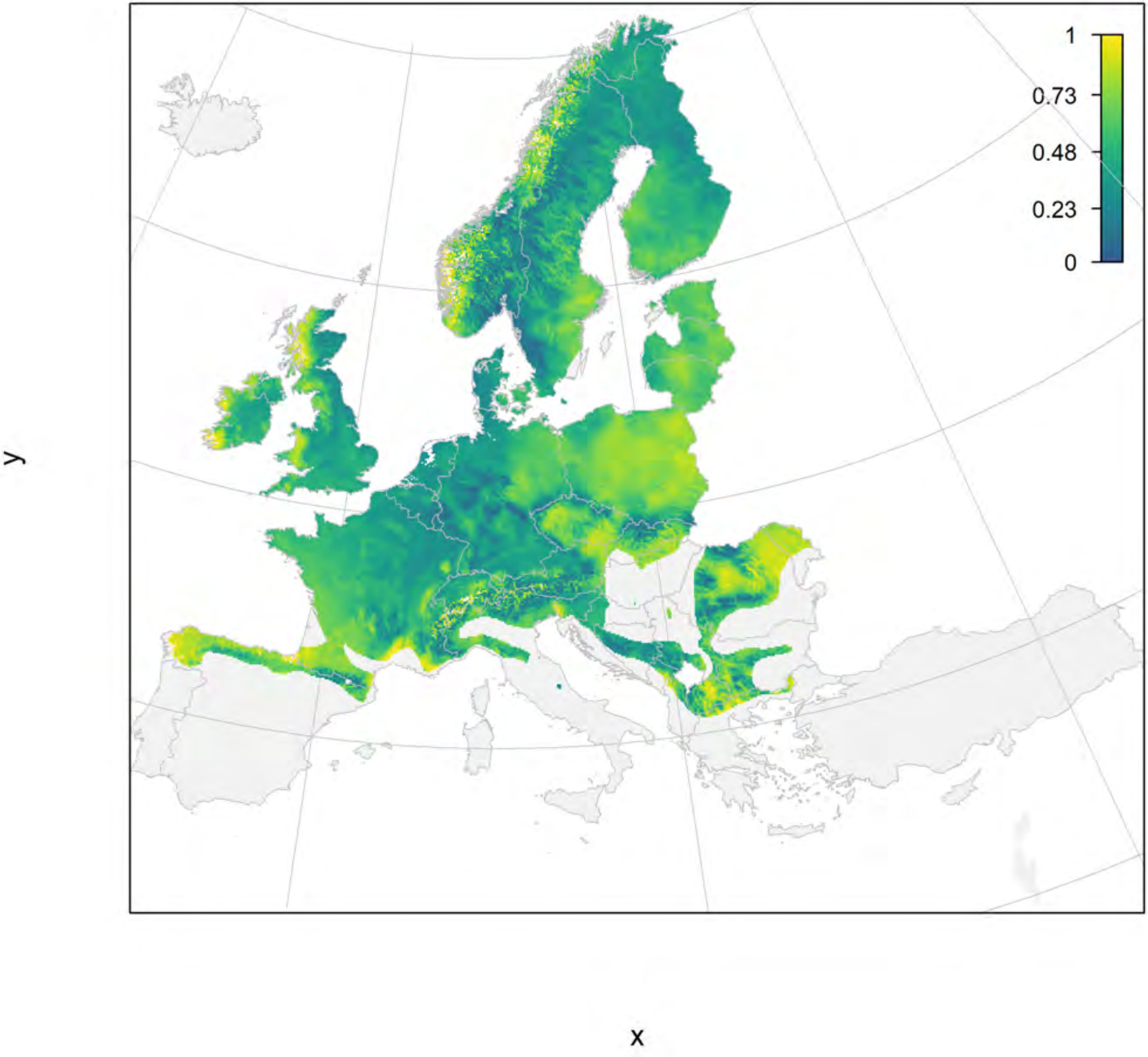

### Rana temporaria_filt current marginality

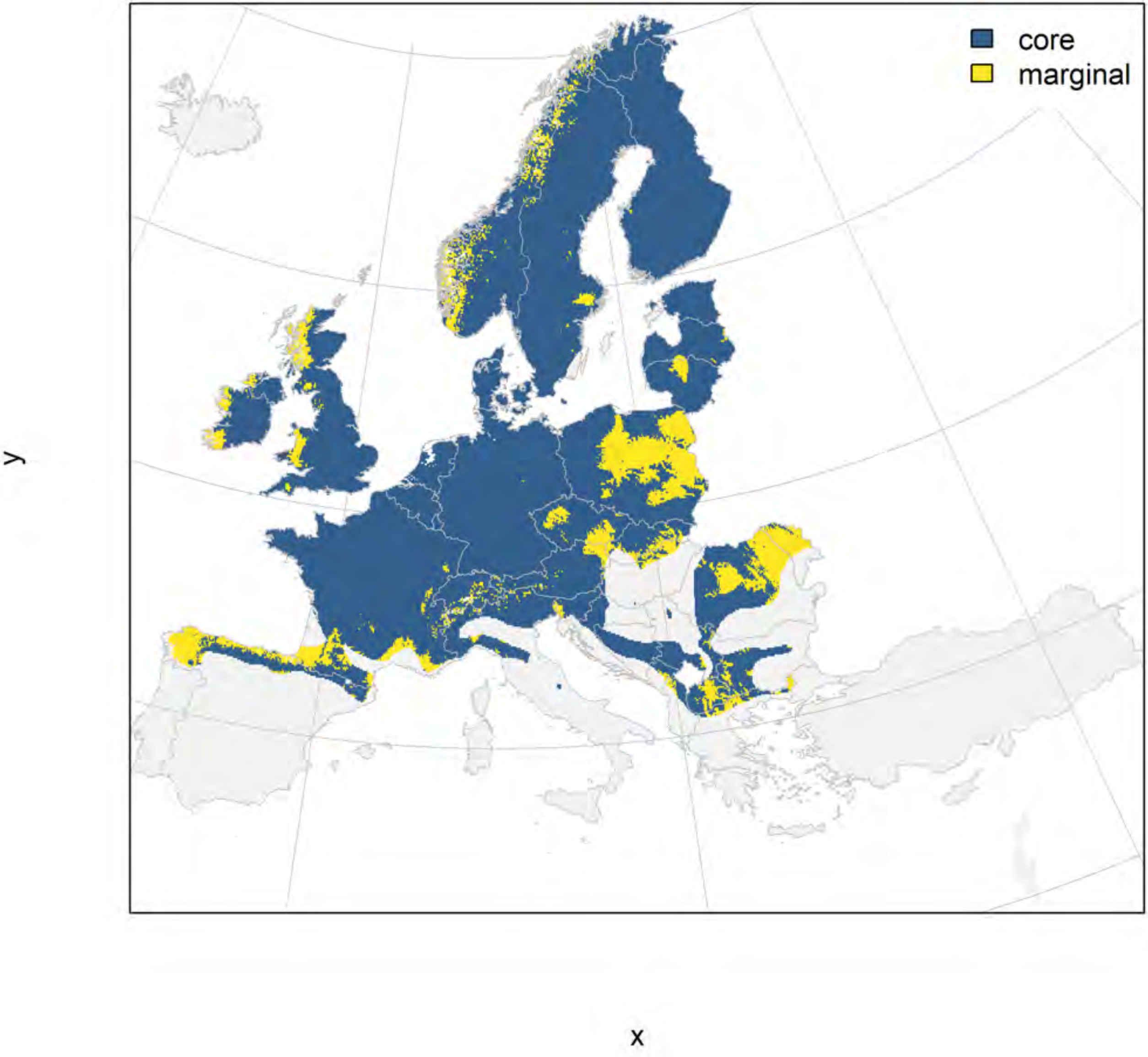

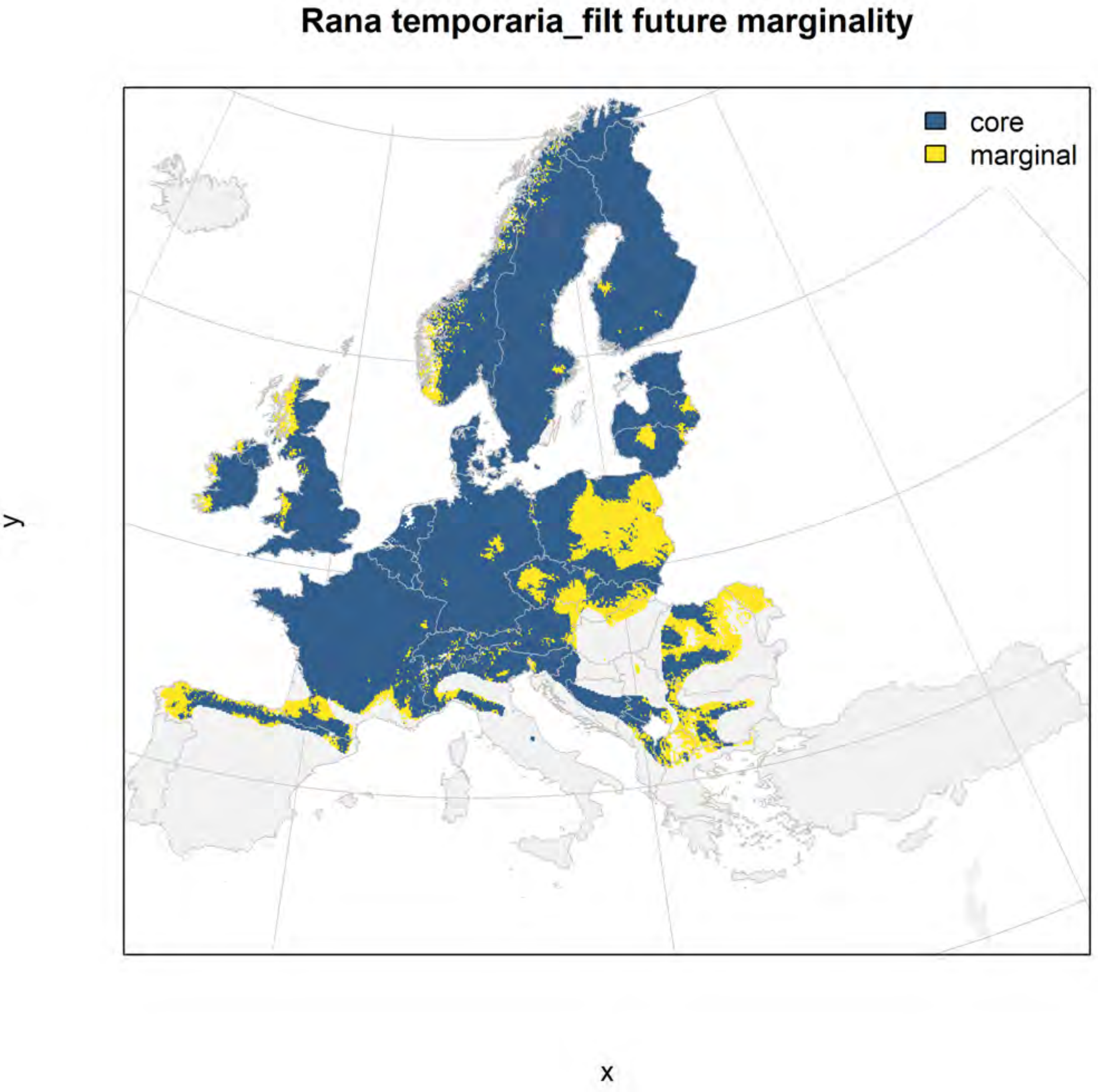
Rana temporaria_filt future marginality
core
marginal
y
x

#### Calotriton asper_filt NMI

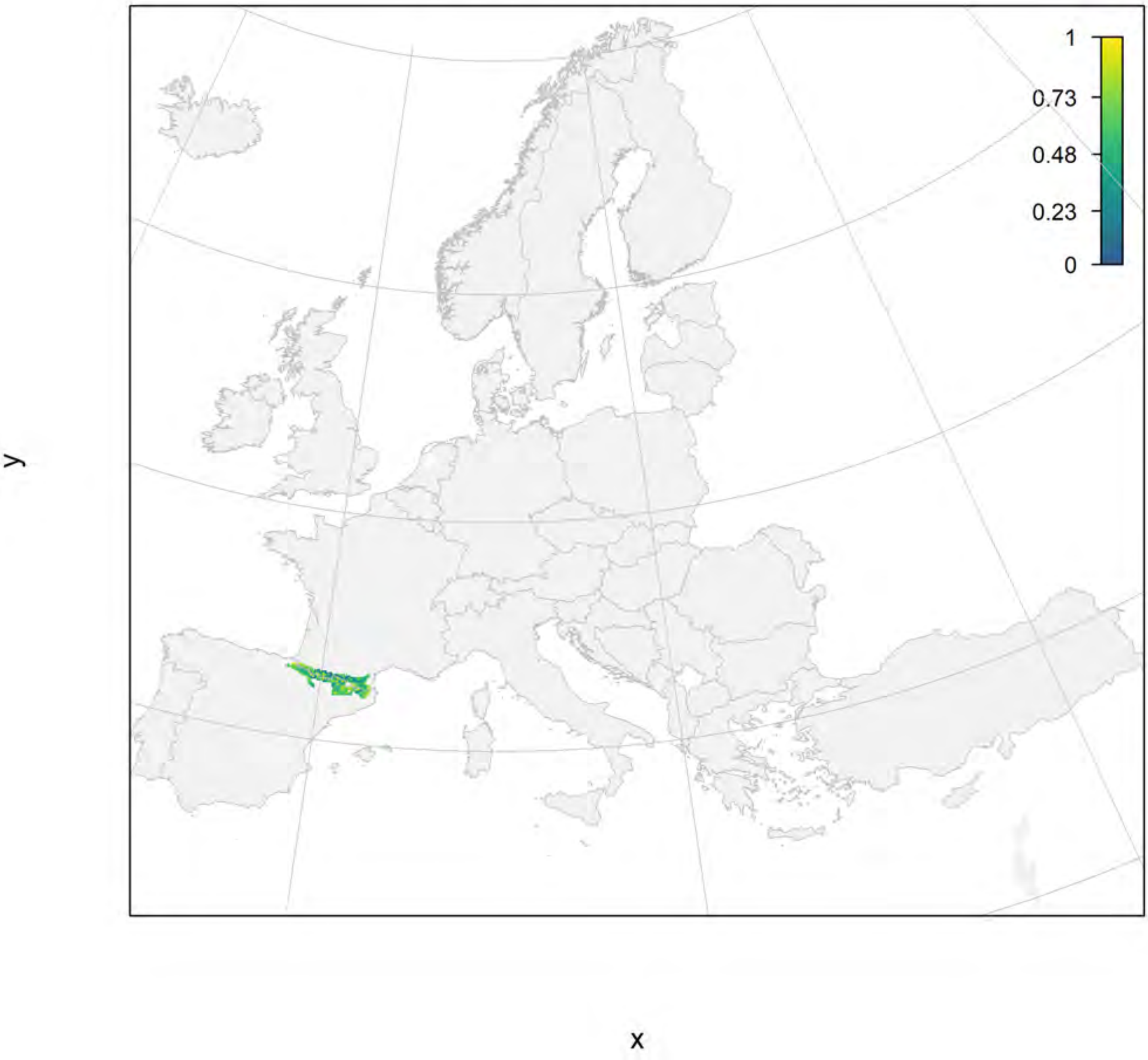

#### Calotriton asper_filt current marginality

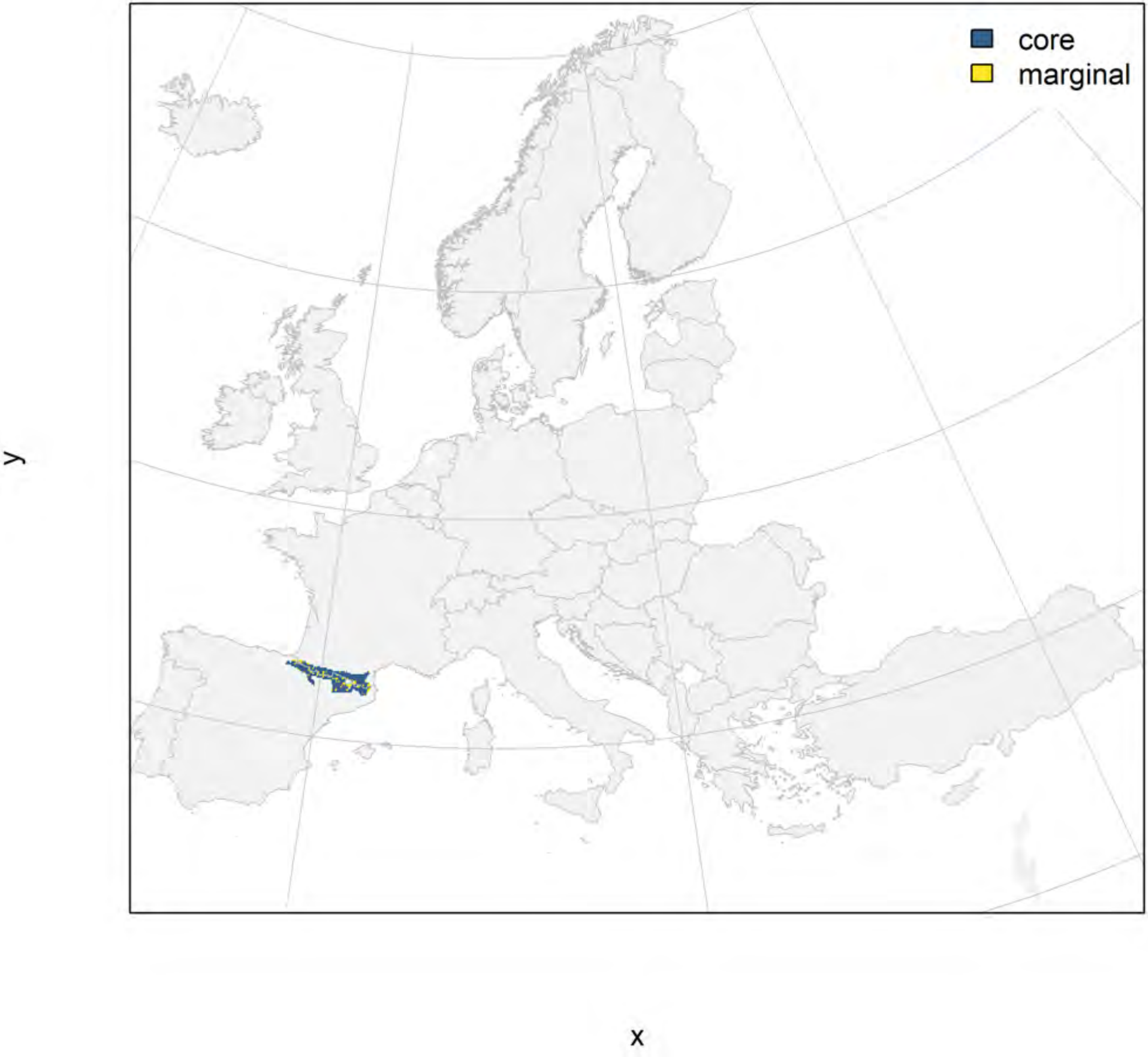

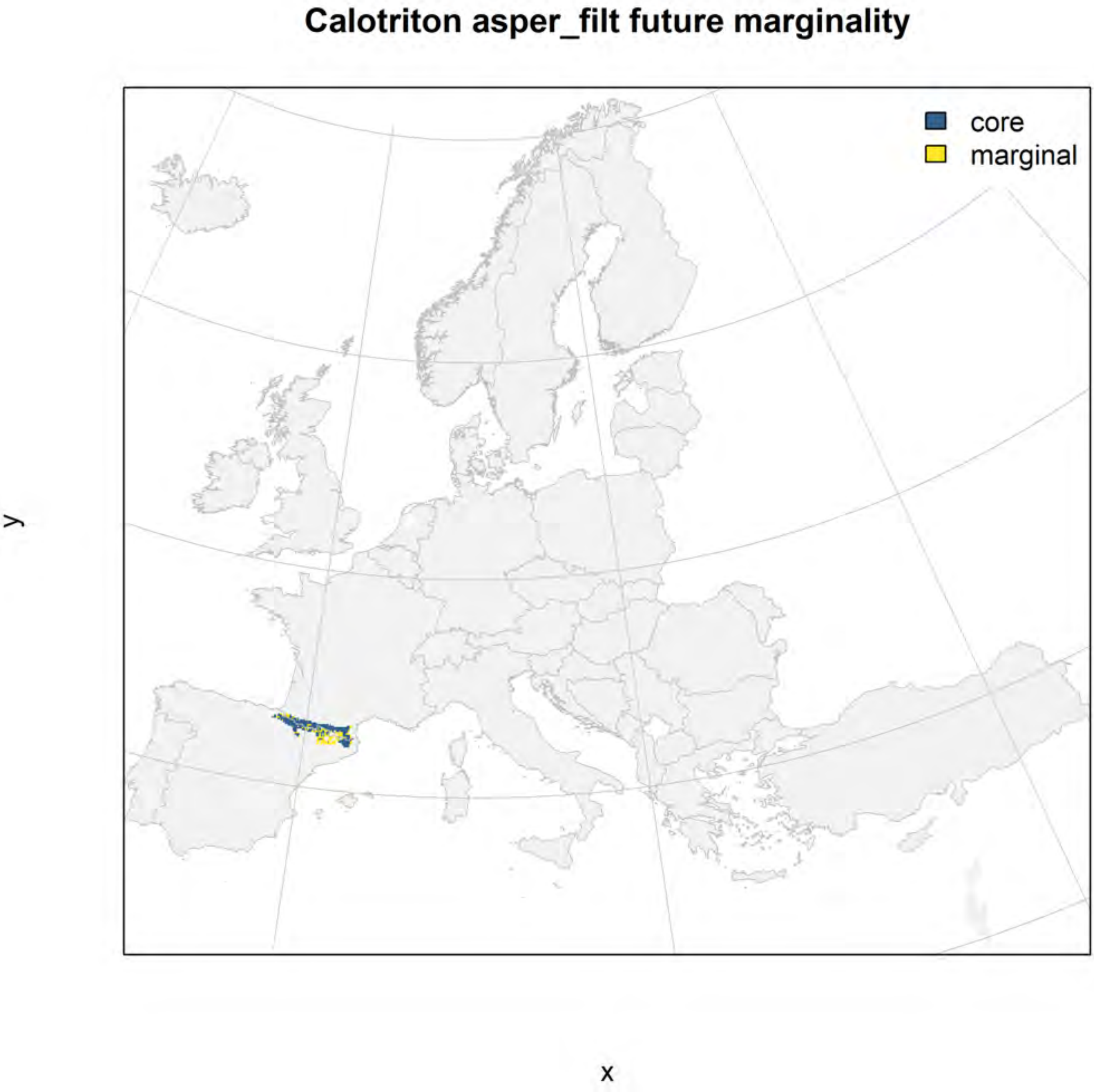

Calotriton asper_filt future marginality
core
marginal
y
x

#### Chioglossa lusitanica_filt NMI

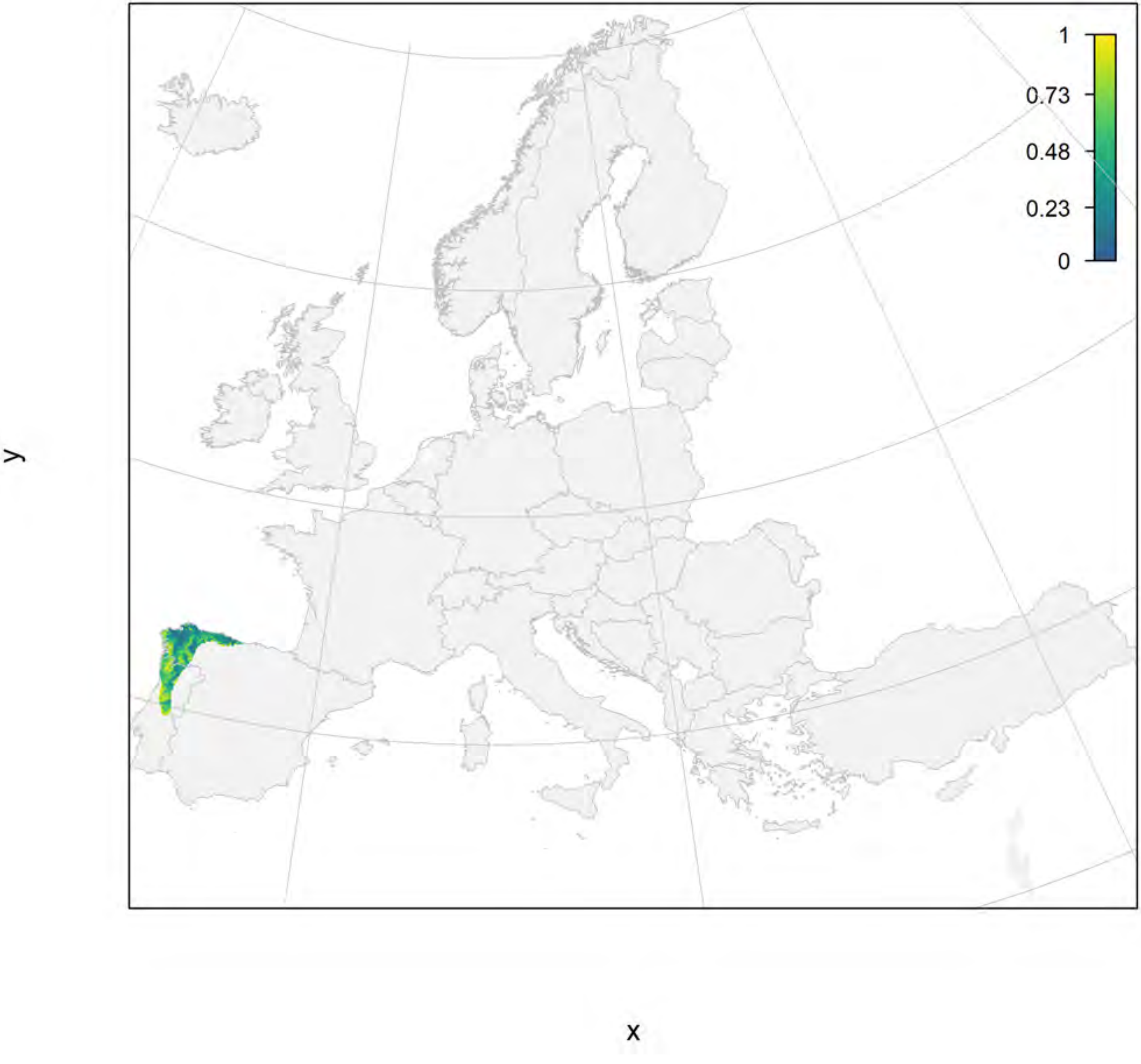

#### Chioglossa lusitanica_filt current marginality

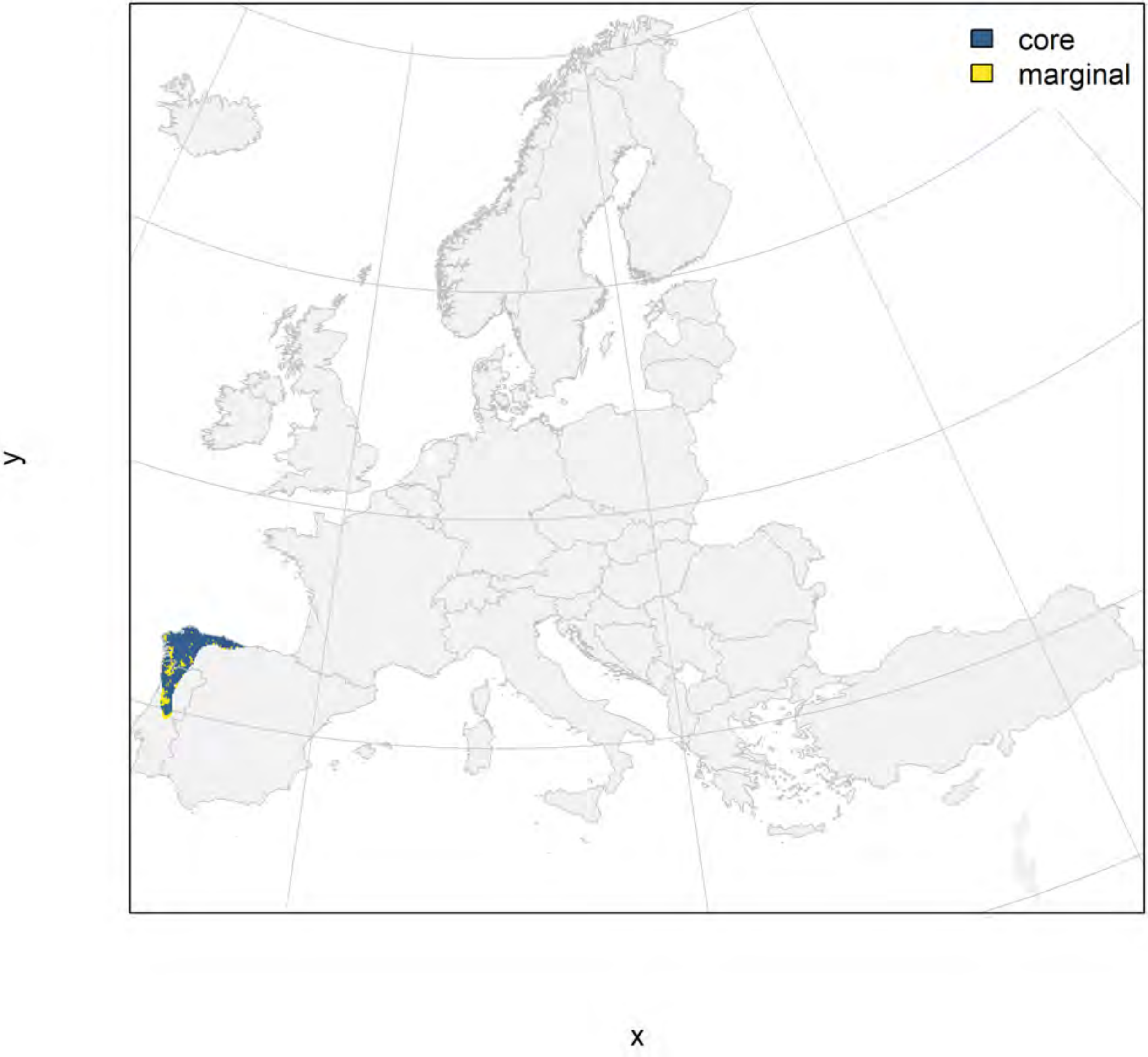

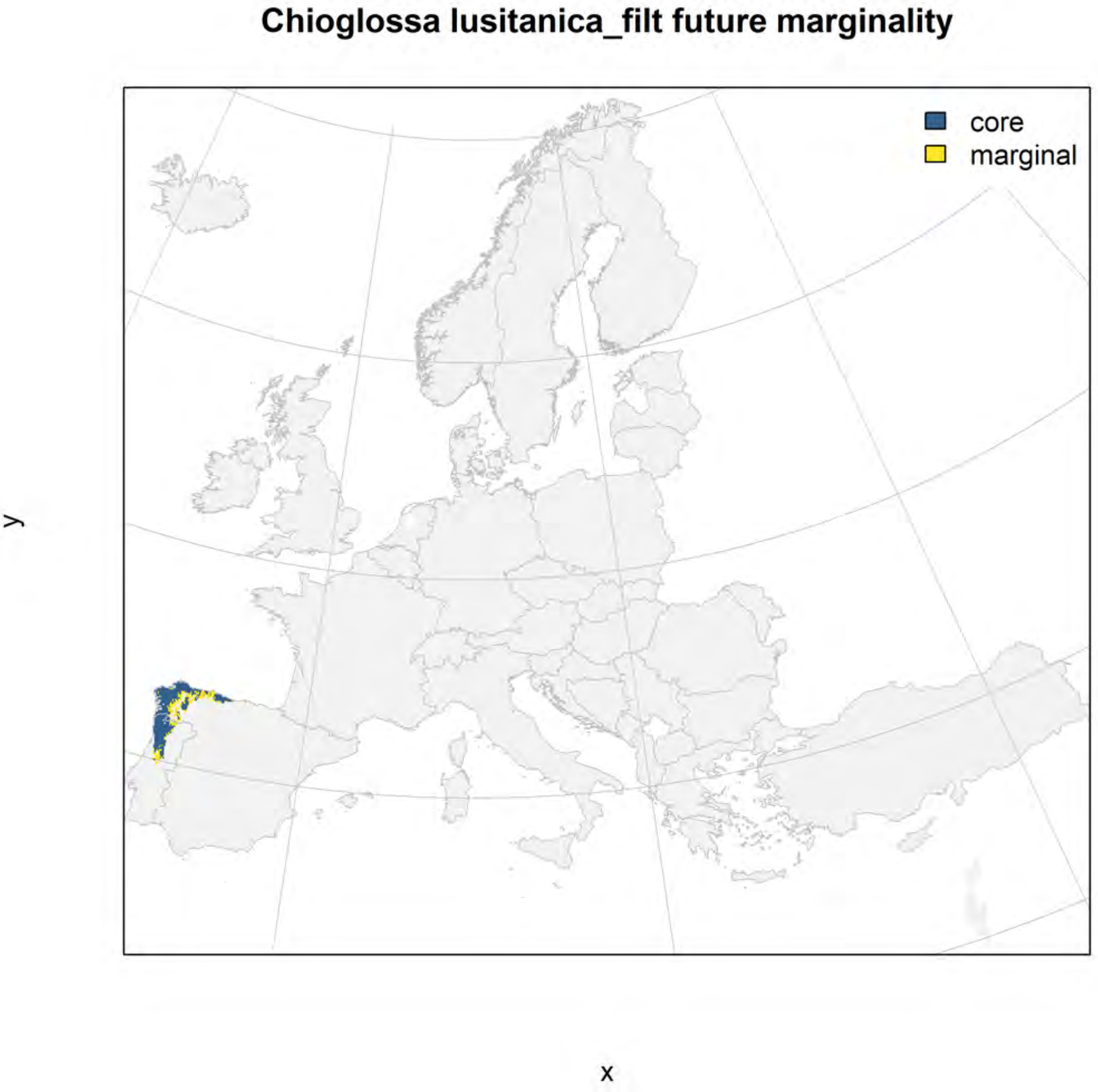
Chioglossa lusitanica_filt future marginality
core
marginal
y
x

### Euproctus montanus_filt NMI

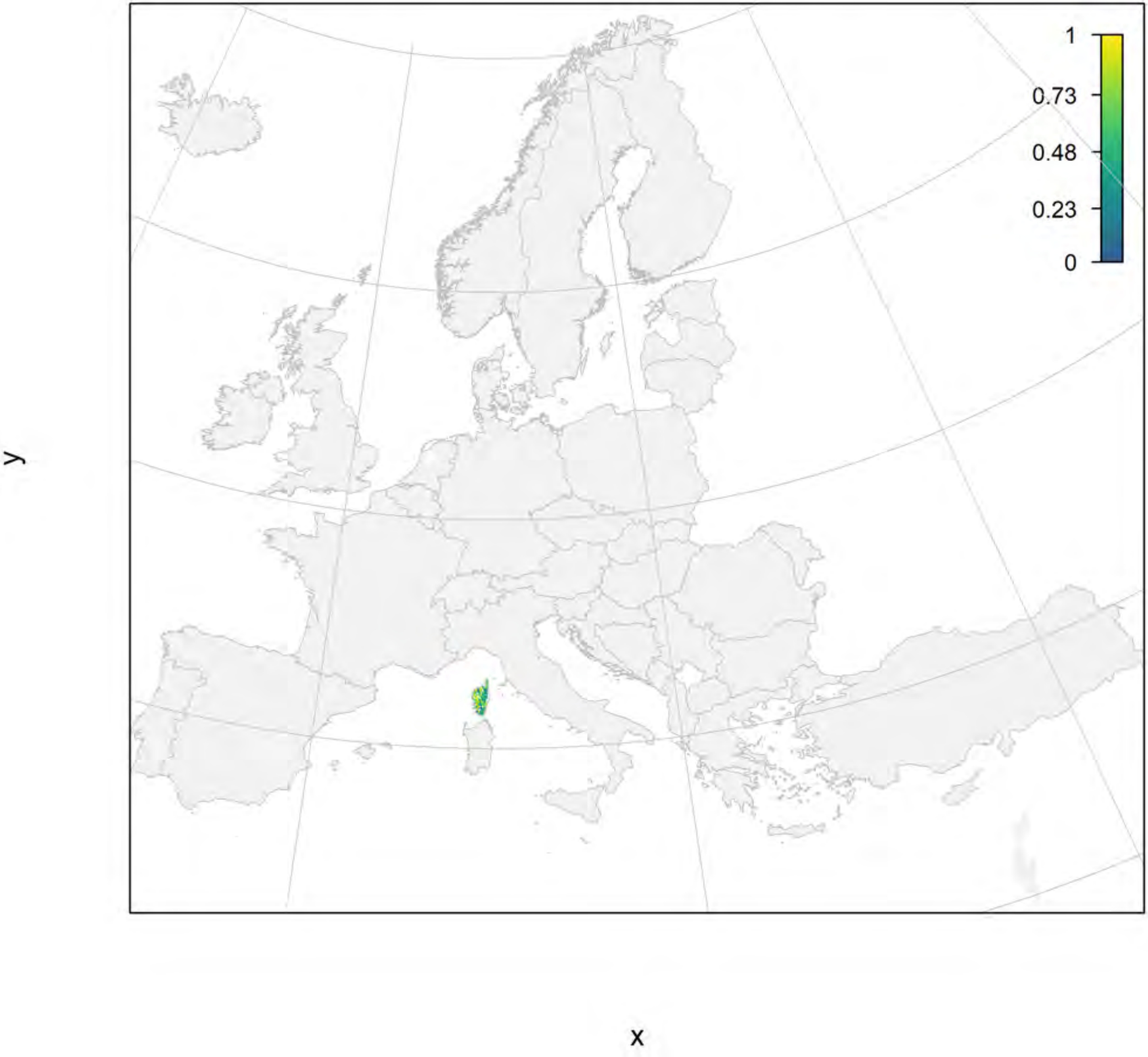

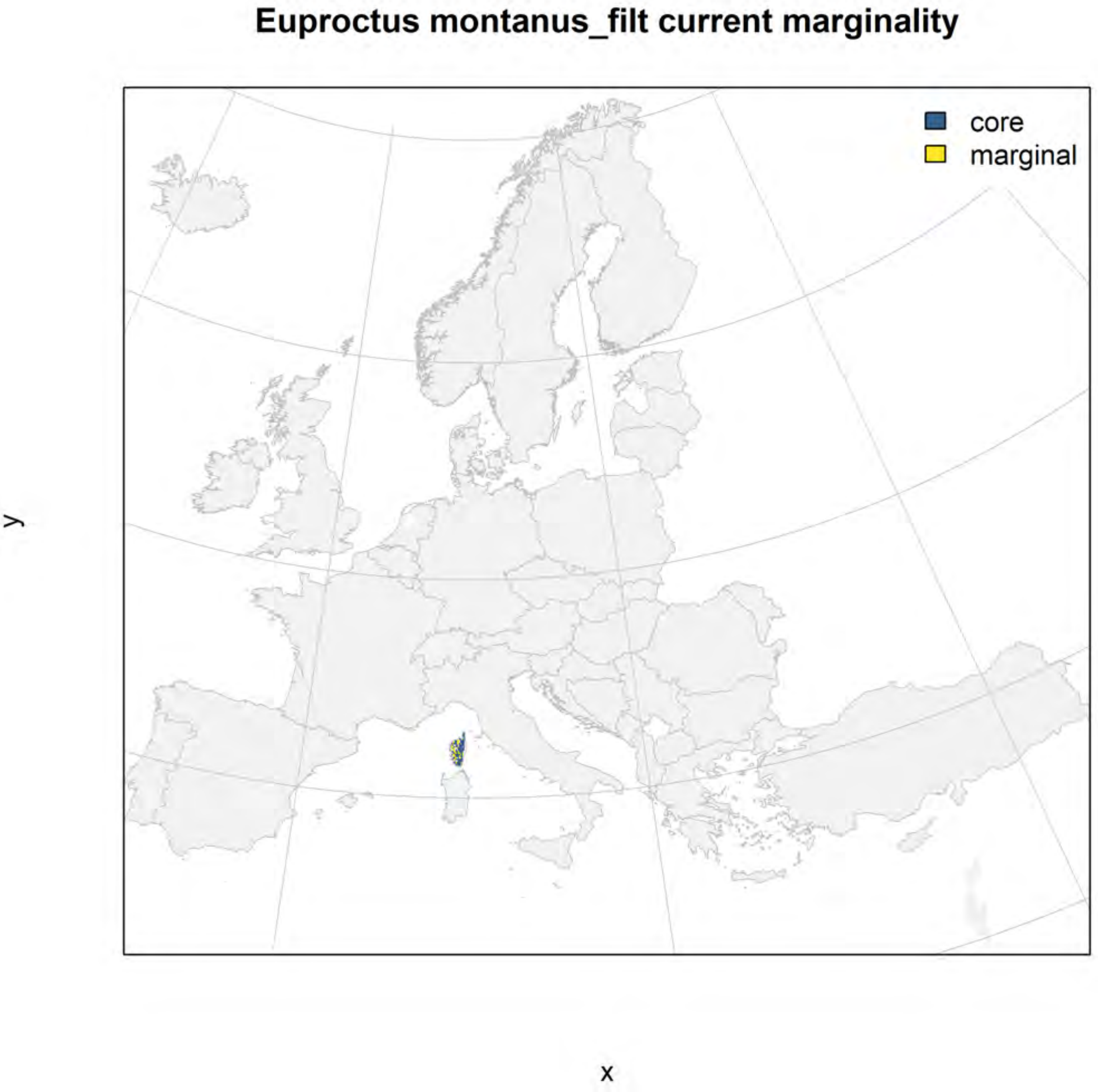
Euproctus montanus_filt current marginality
core
marginal
y
x

### Euproctus montanus_filt future marginality

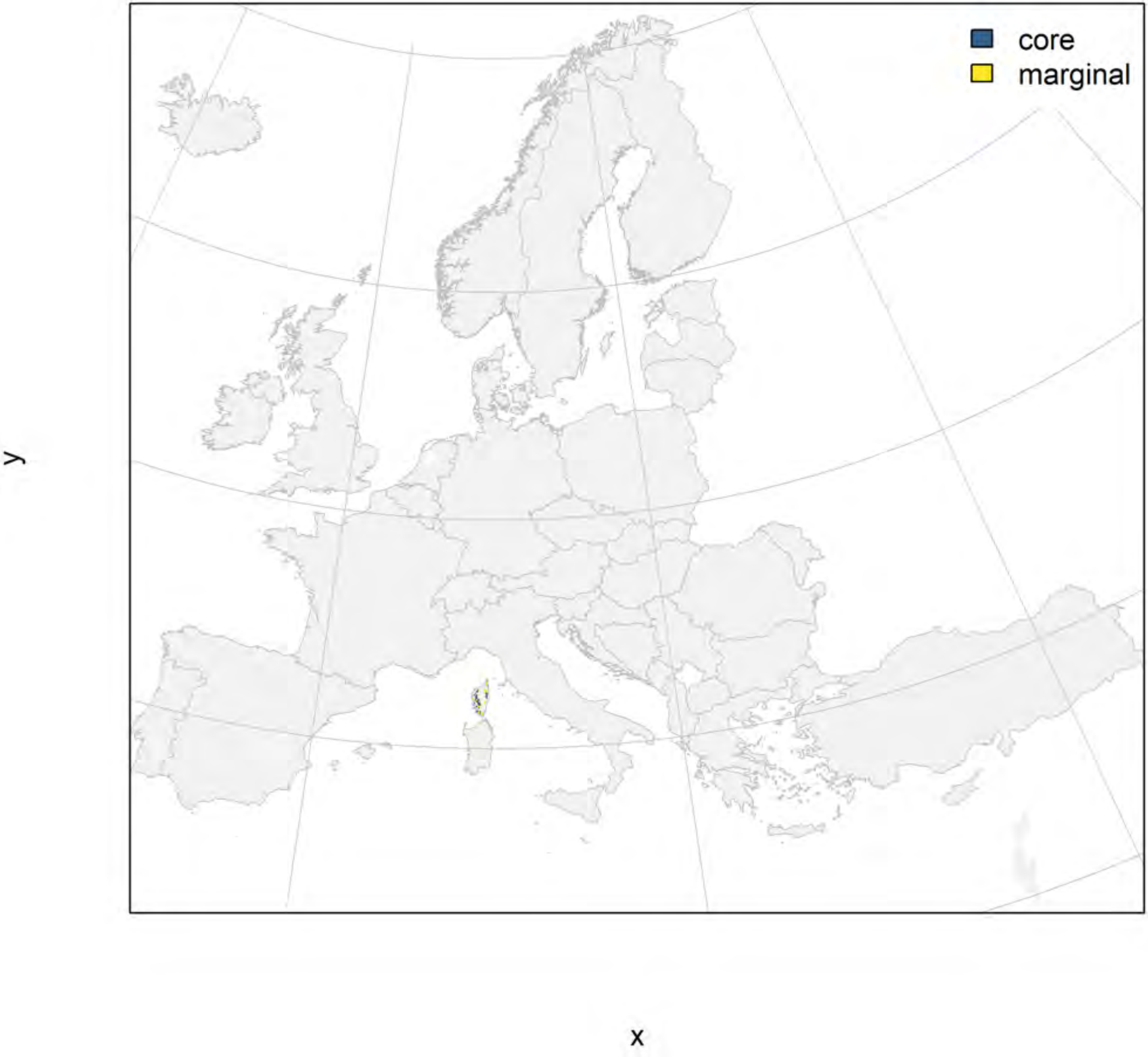

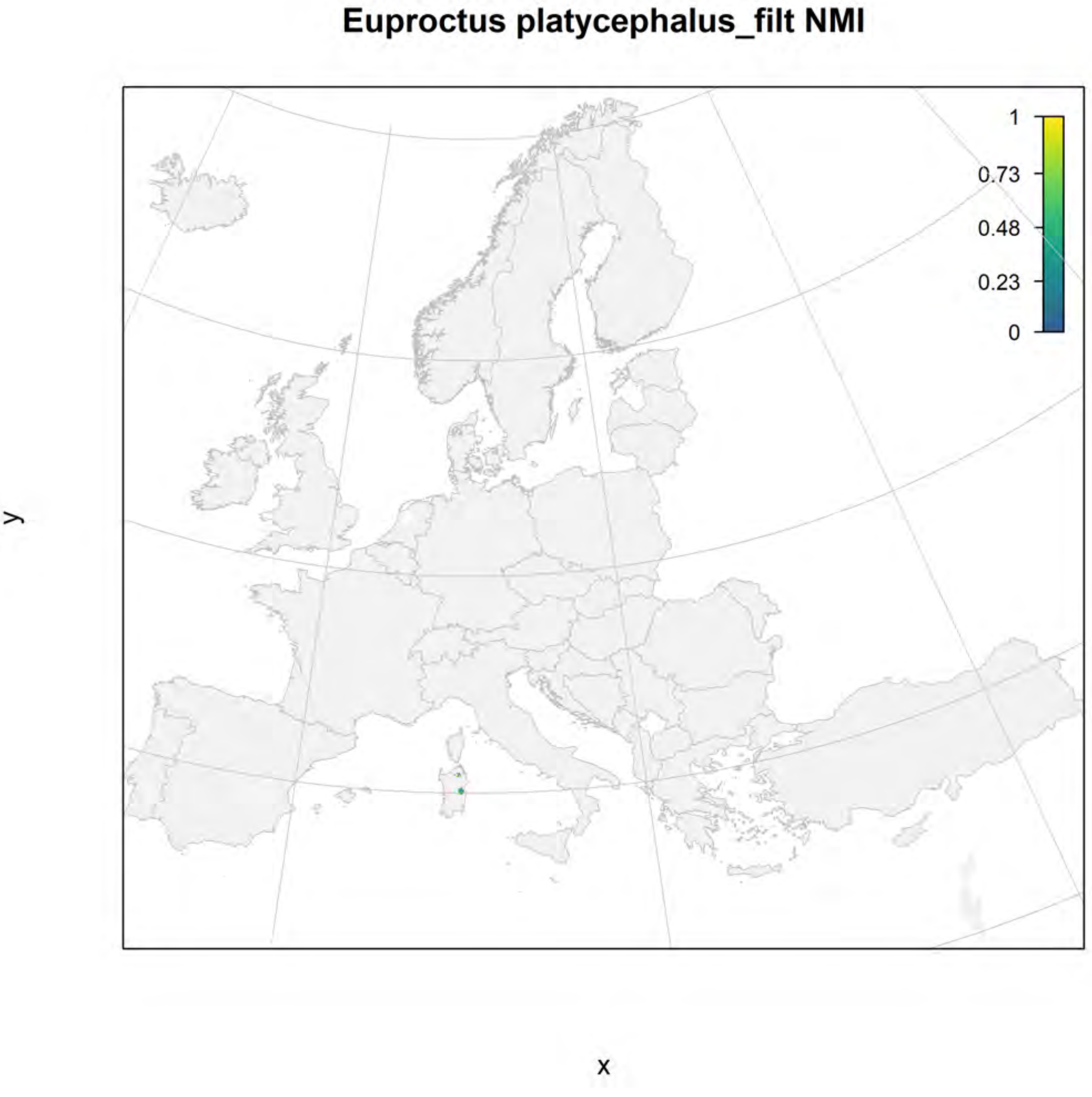
Euproctus platycephalus_filt NMI
1
0.73
0.48
0.23
0
y
x

#### Euproctus platycephalus_filt current marginality

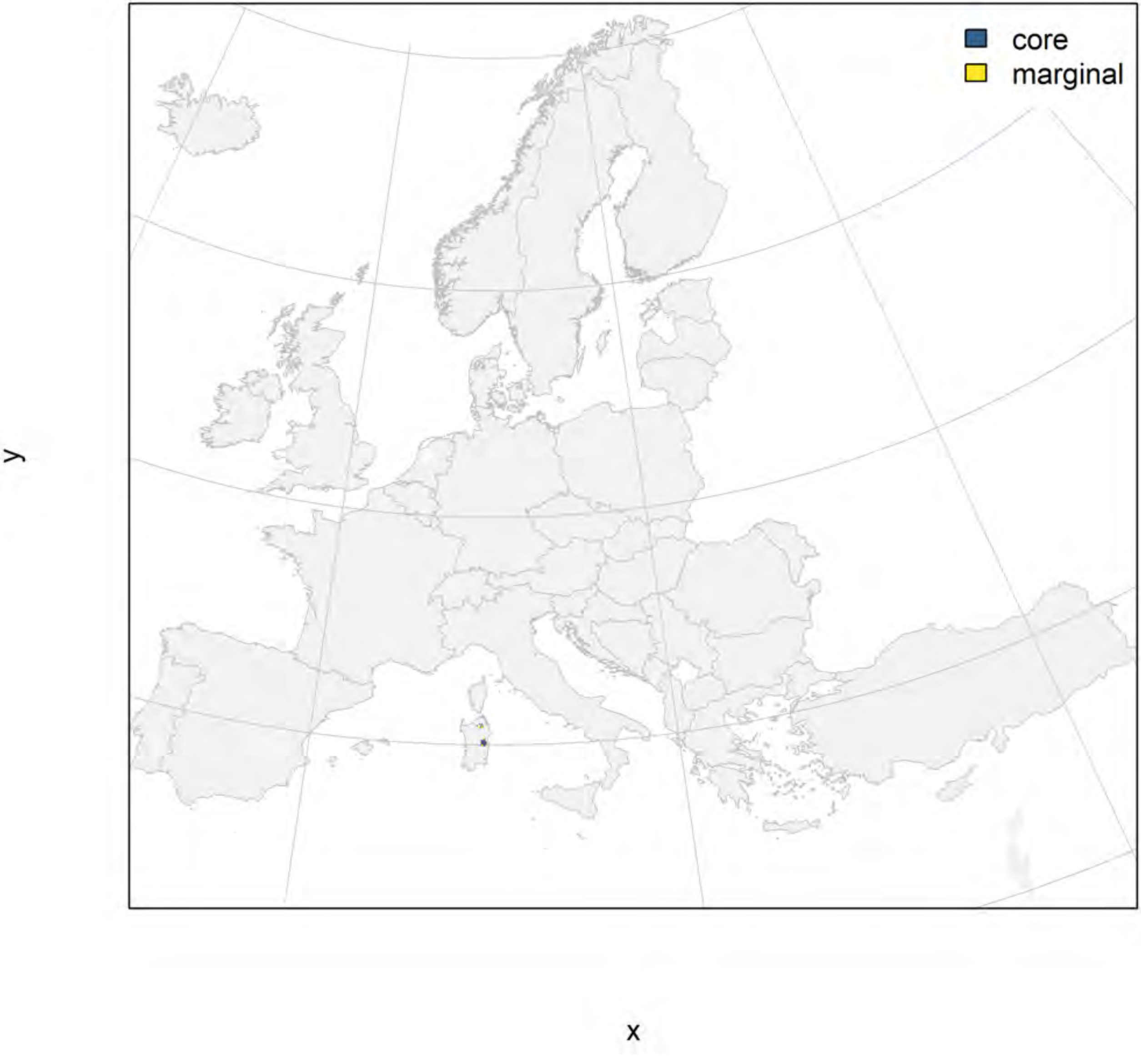

#### Euproctus platycephalus_filt future marginality

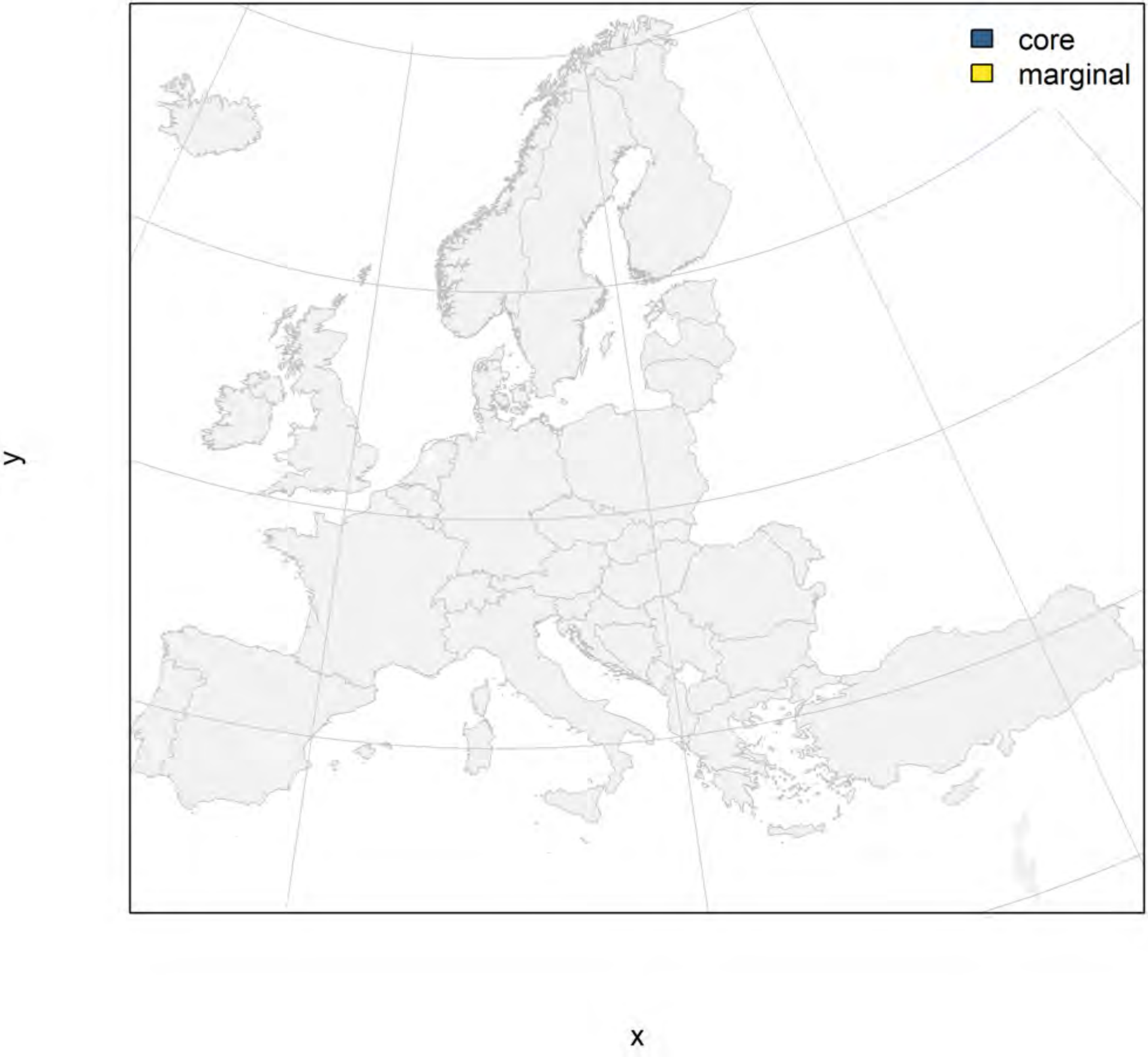

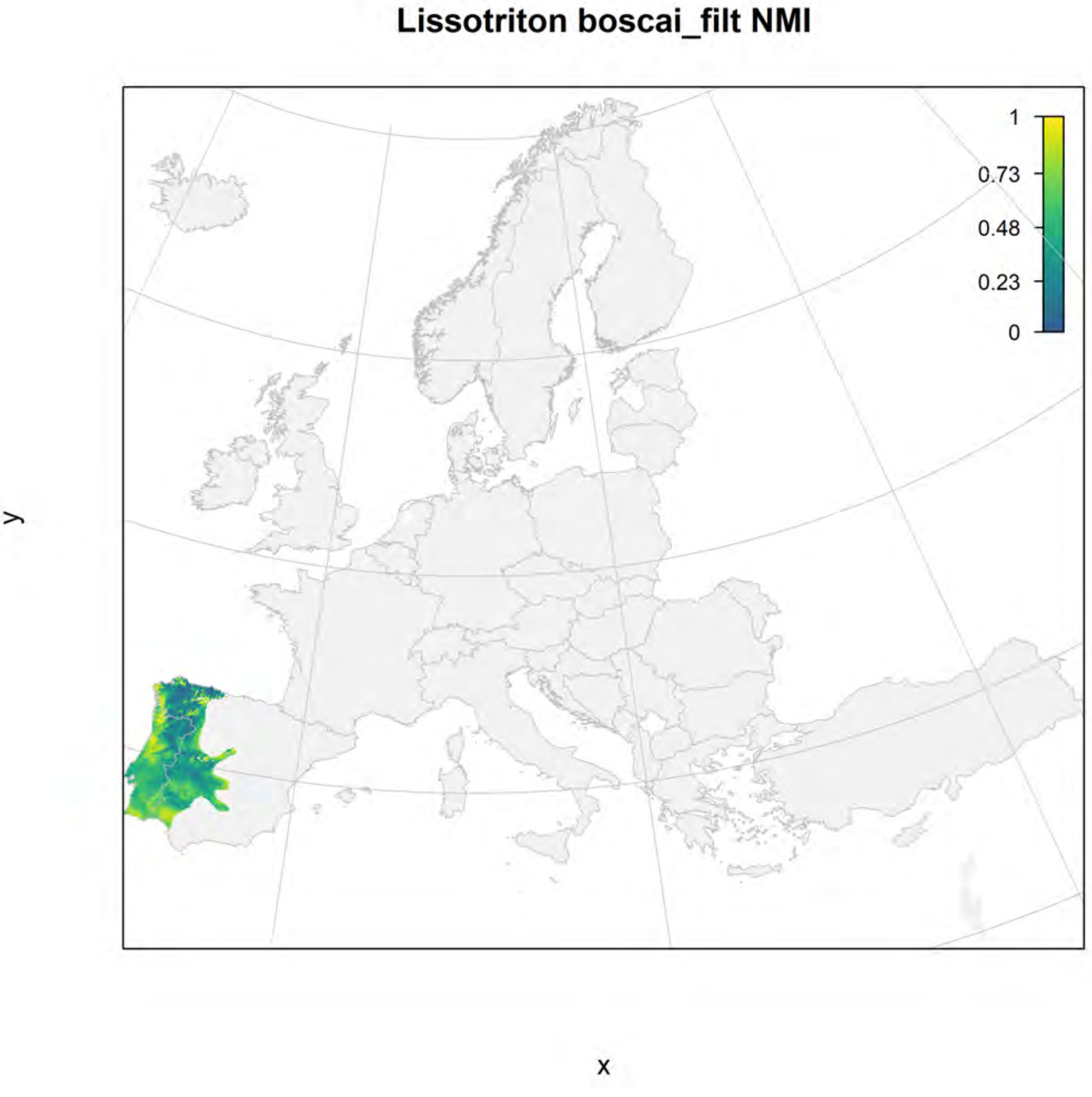

Lissotriton boscai_filt NMI
1
0.73
0.48
0.23
0
y
x

#### Lissotriton boscai_filt current marginality

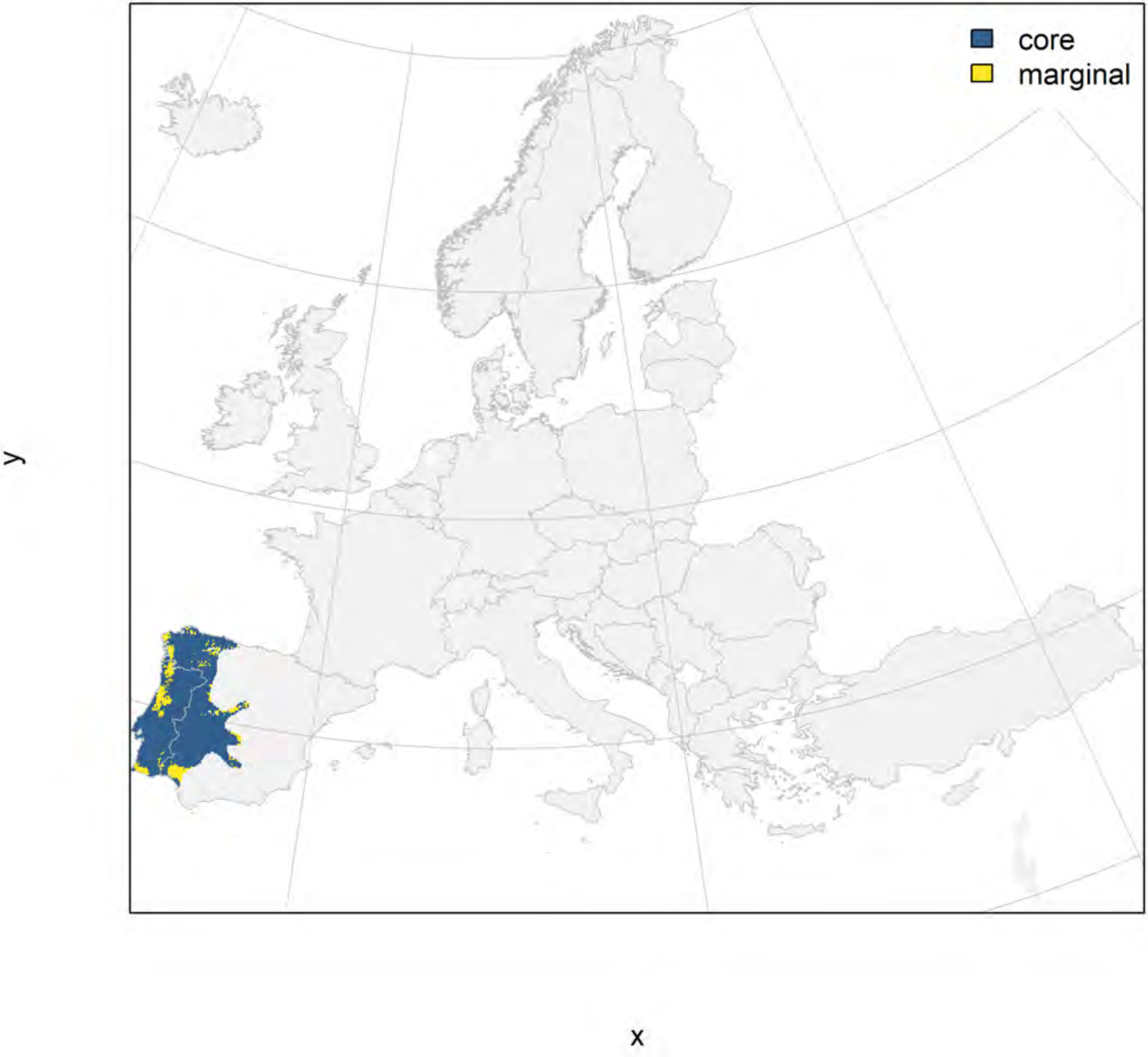

#### Lissotriton boscai_filt future marginality

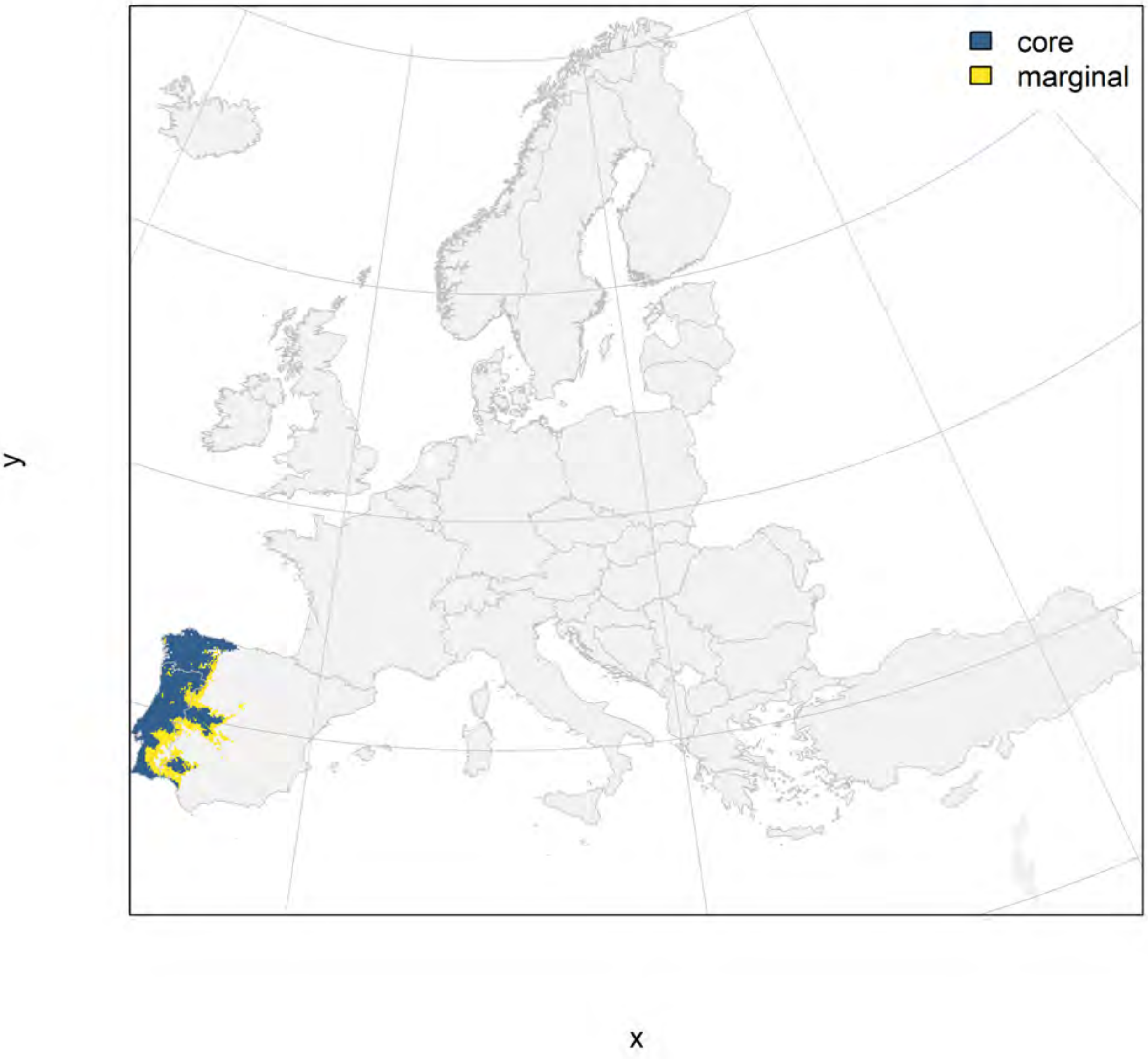

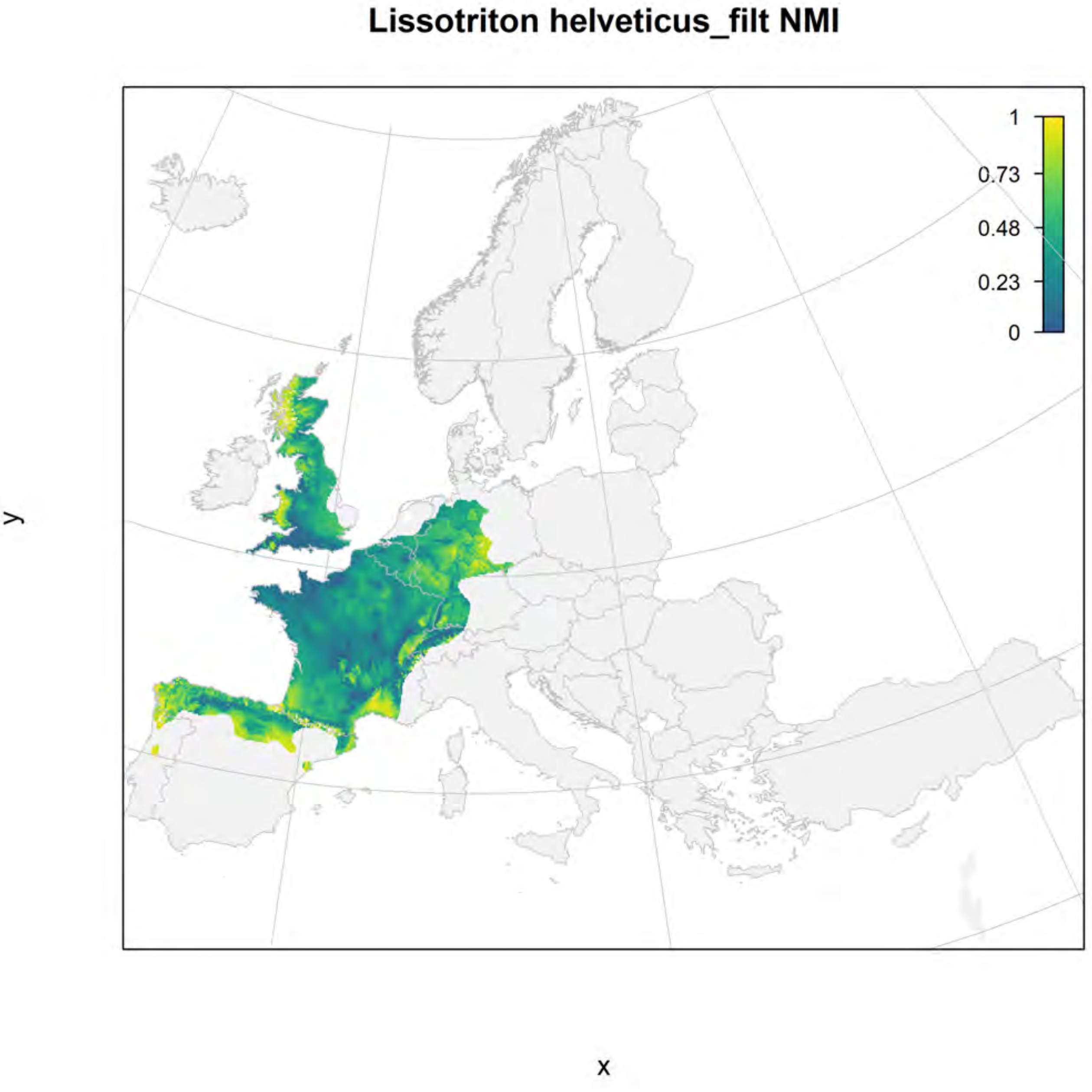
Lissotriton helveticus_filt NMI
1
0.73
0.48
0.23
0
y
x

### Lissotriton helveticus_filt current marginality

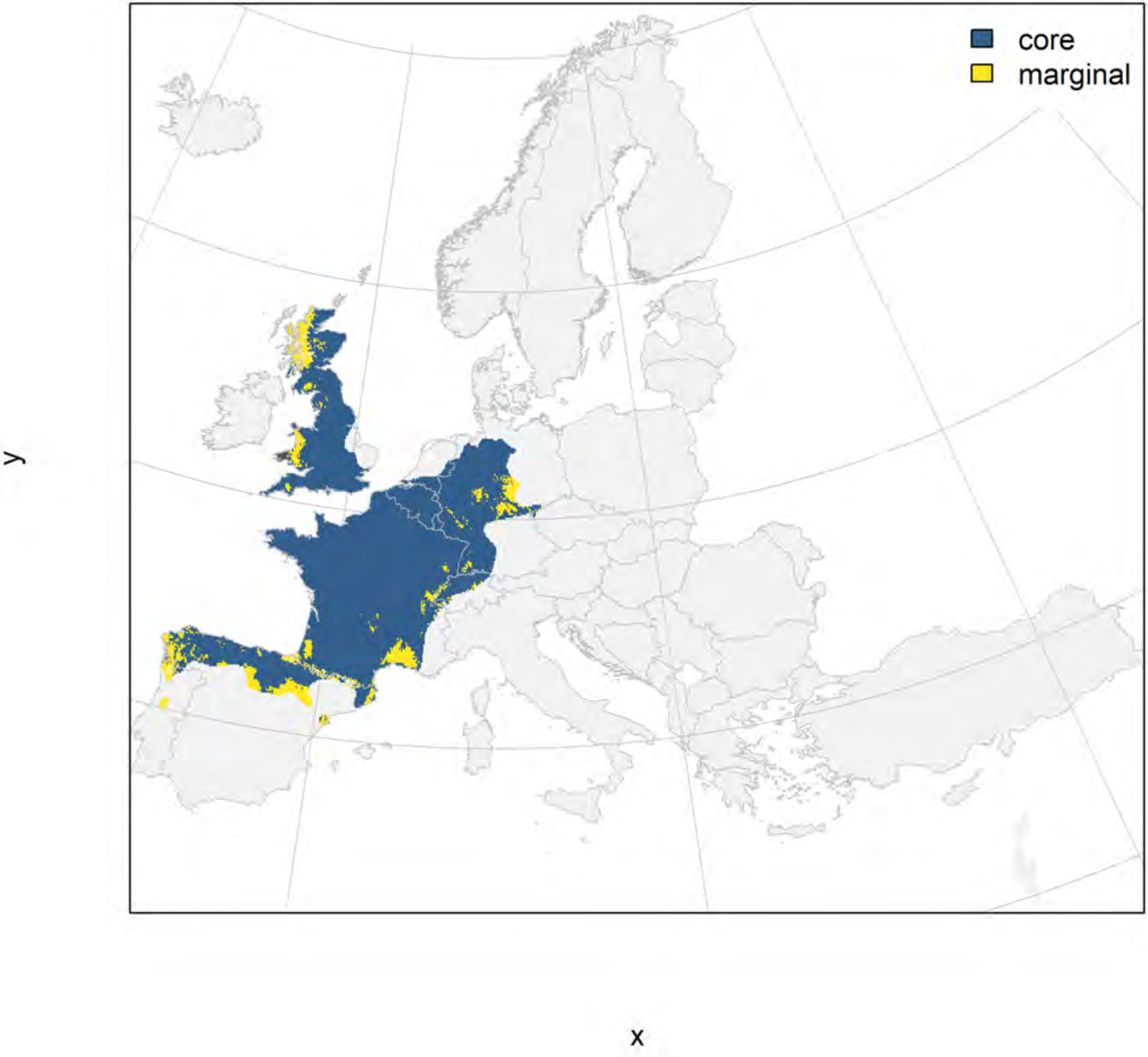

#### Lissotriton helveticus_filt future marginality

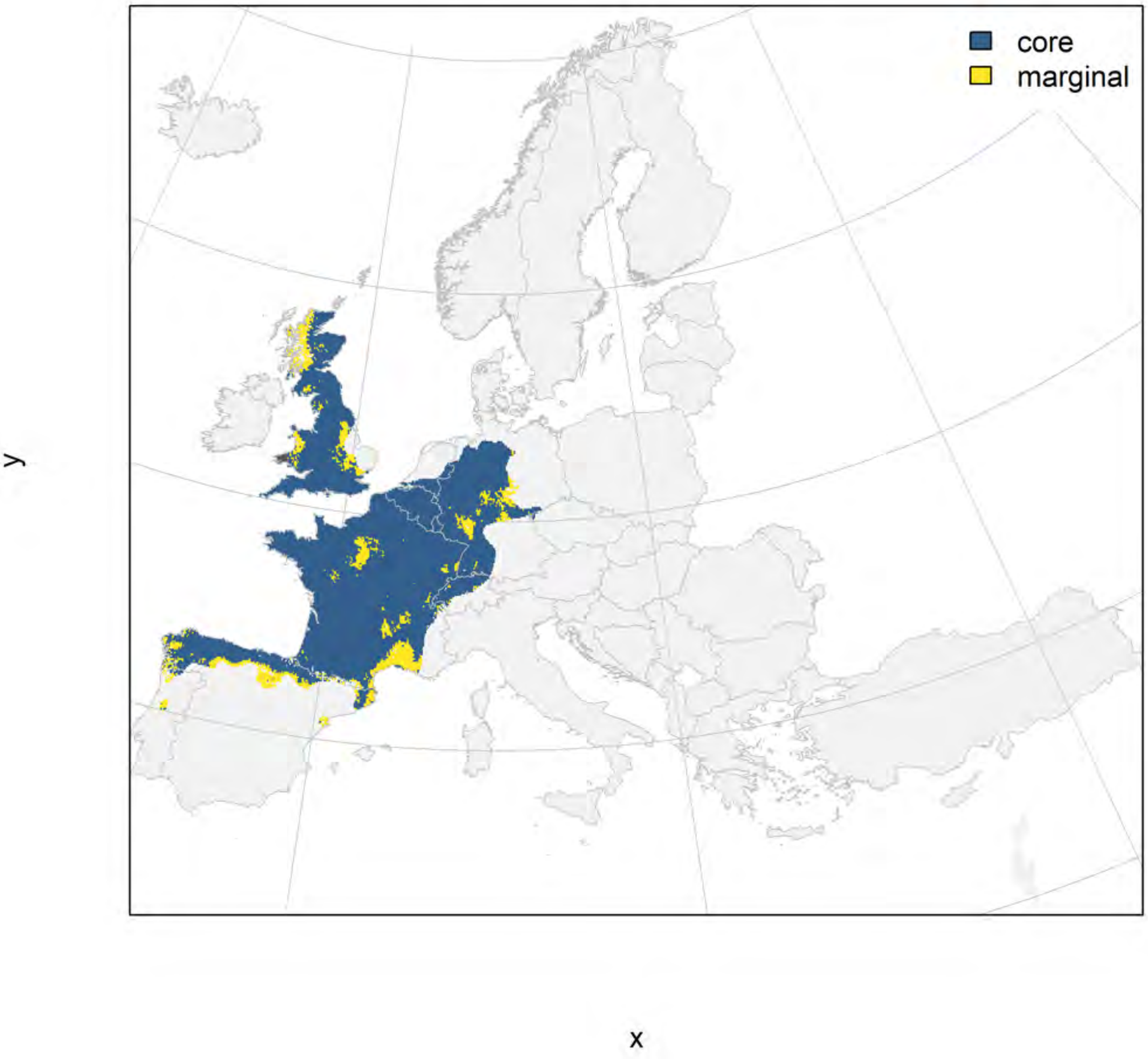

### Lissotriton italicus_filt NMI

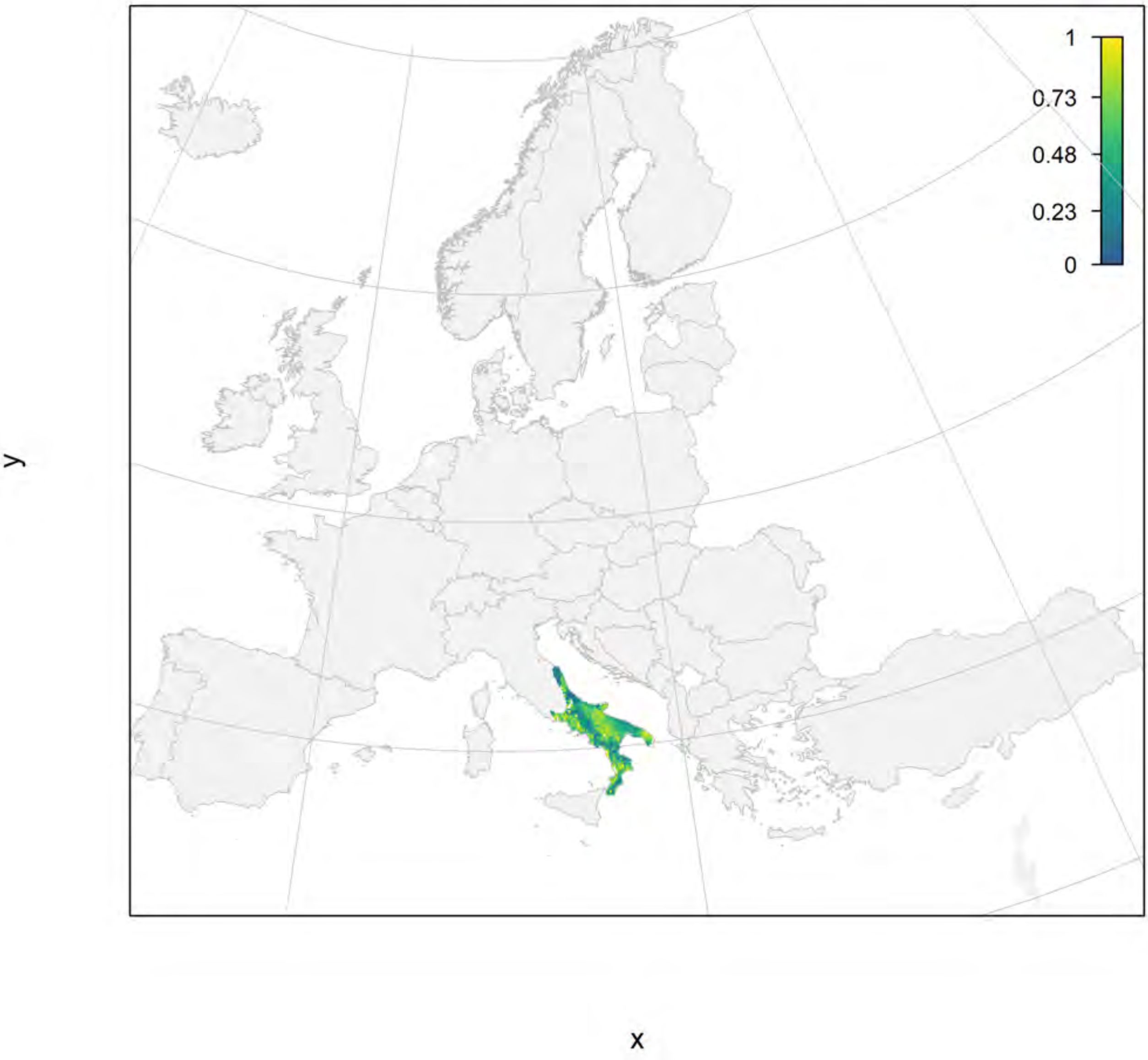

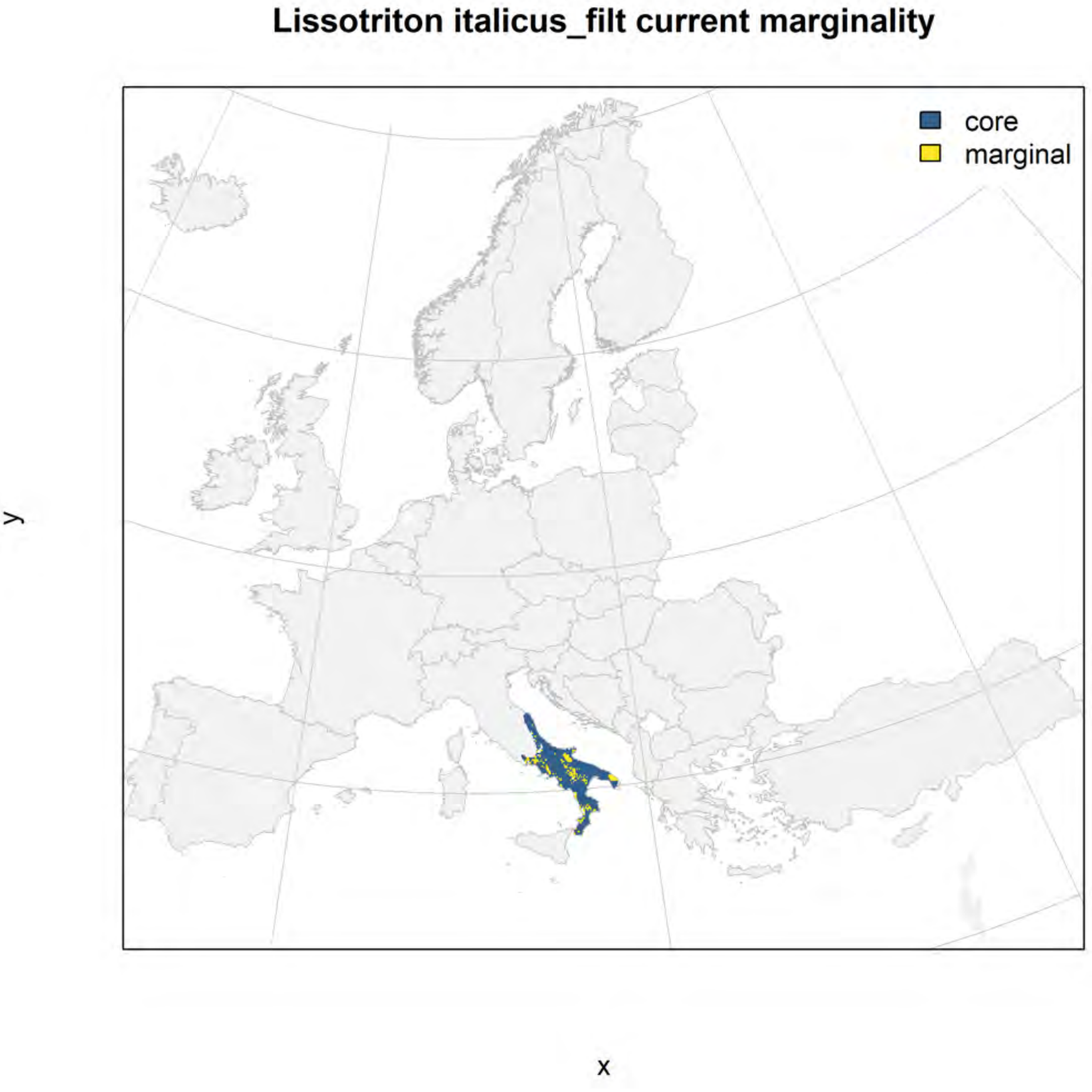
Lissotriton italicus_filt current marginality
core
marginal
y
x

Lissotriton italicus_filt future marginality

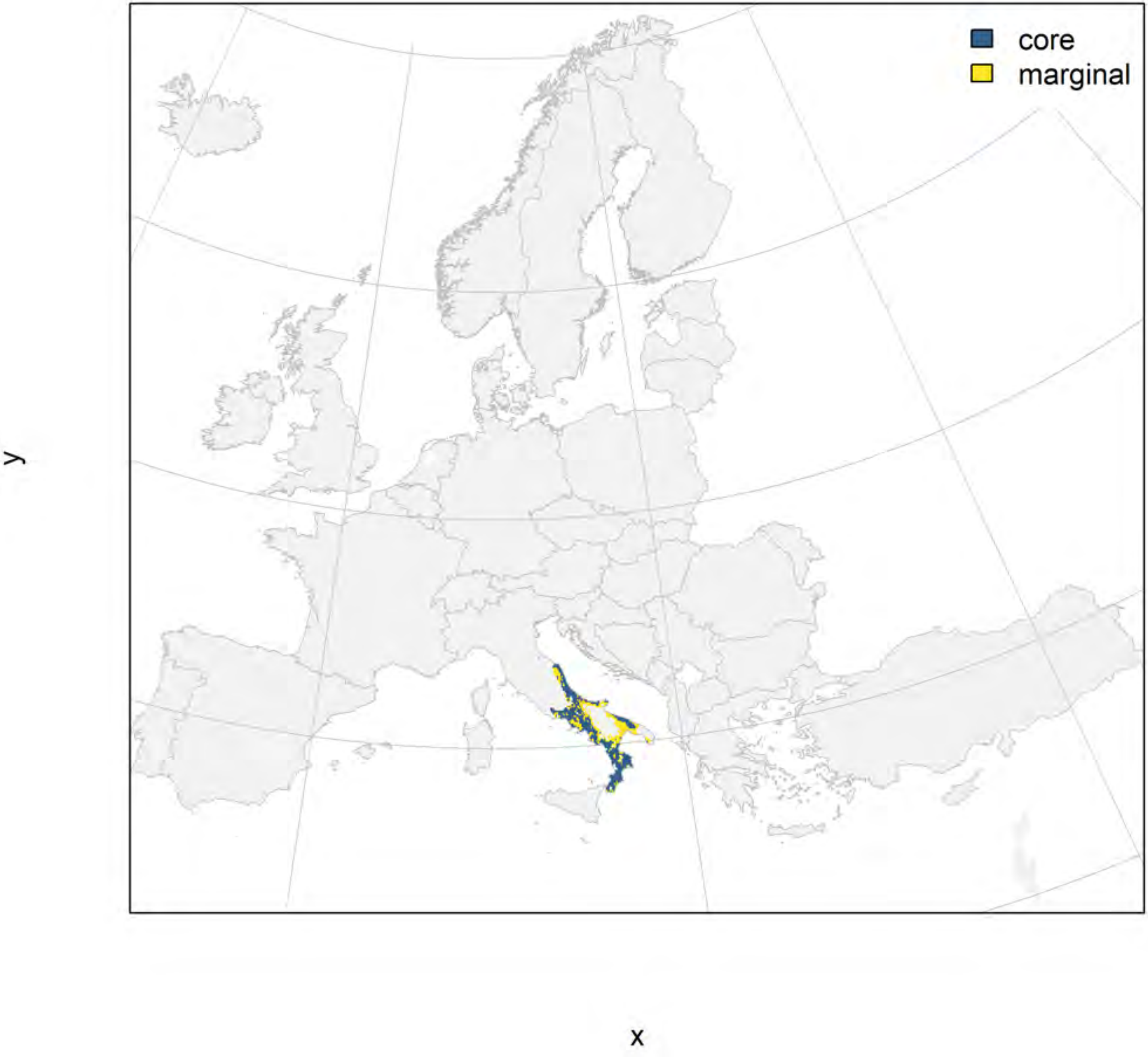

### Lissotriton montandoni_filt NMI

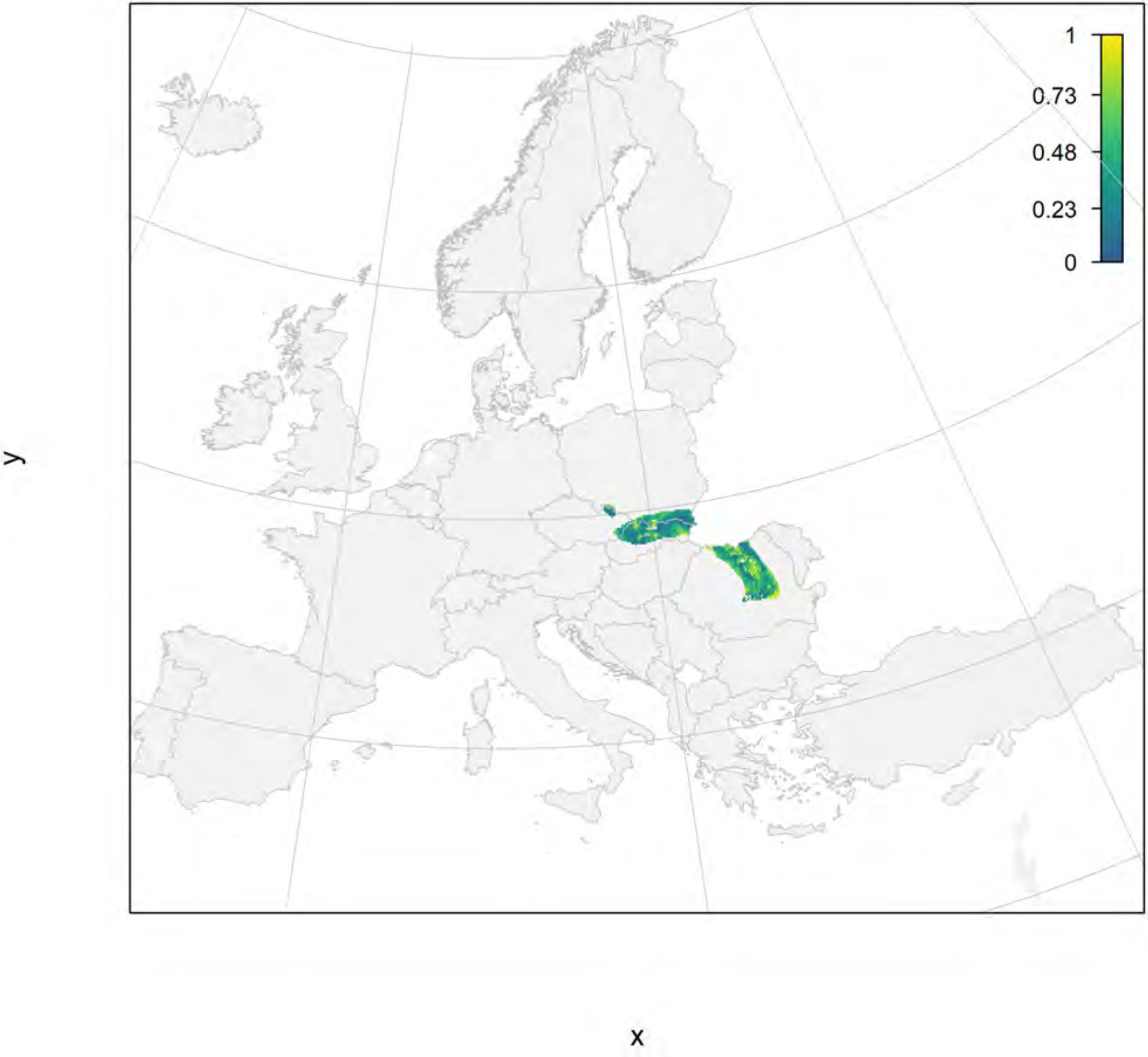

#### Lissotriton montandoni_filt current marginality

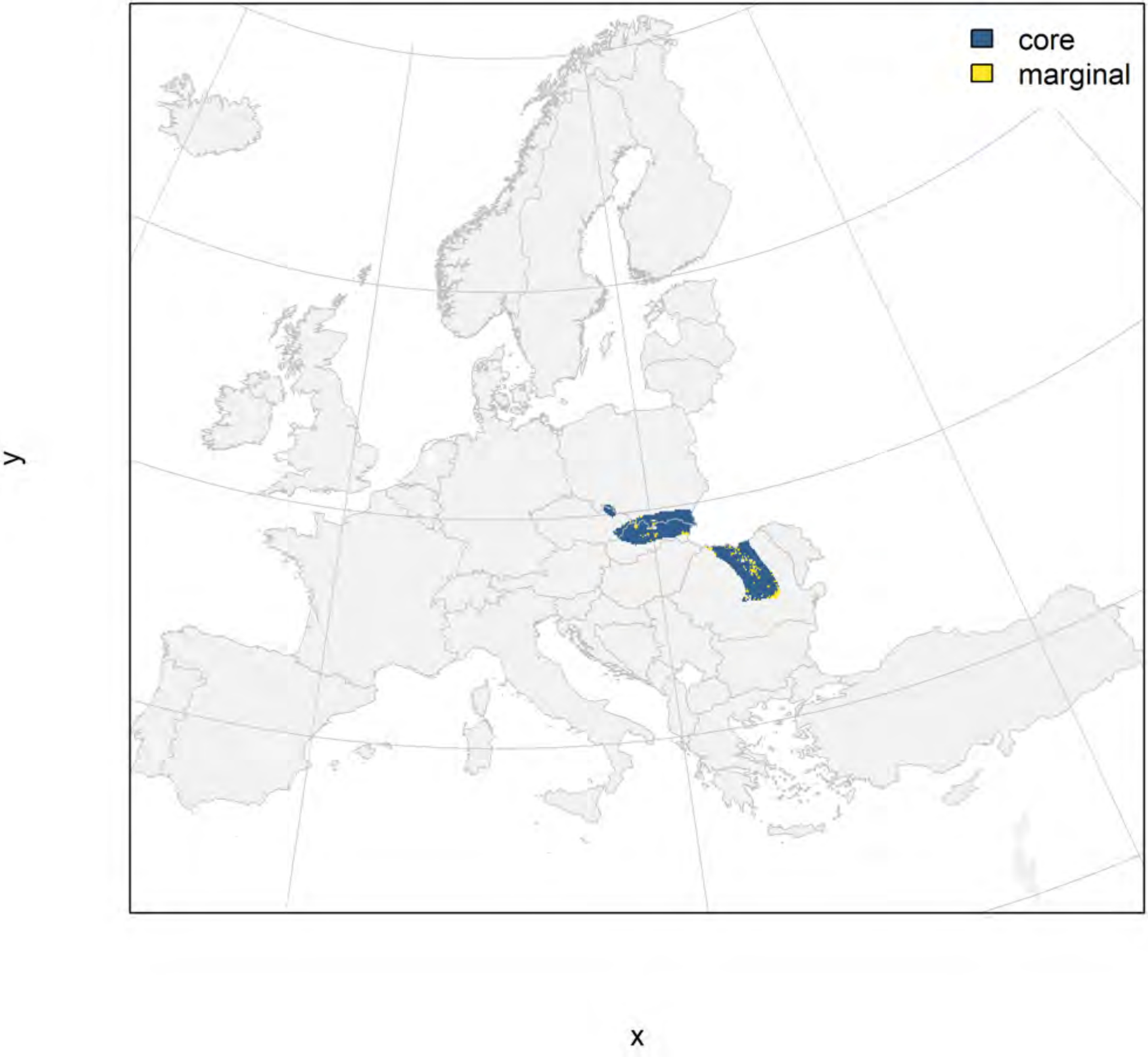

#### Lissotriton montandoni_filt future marginality

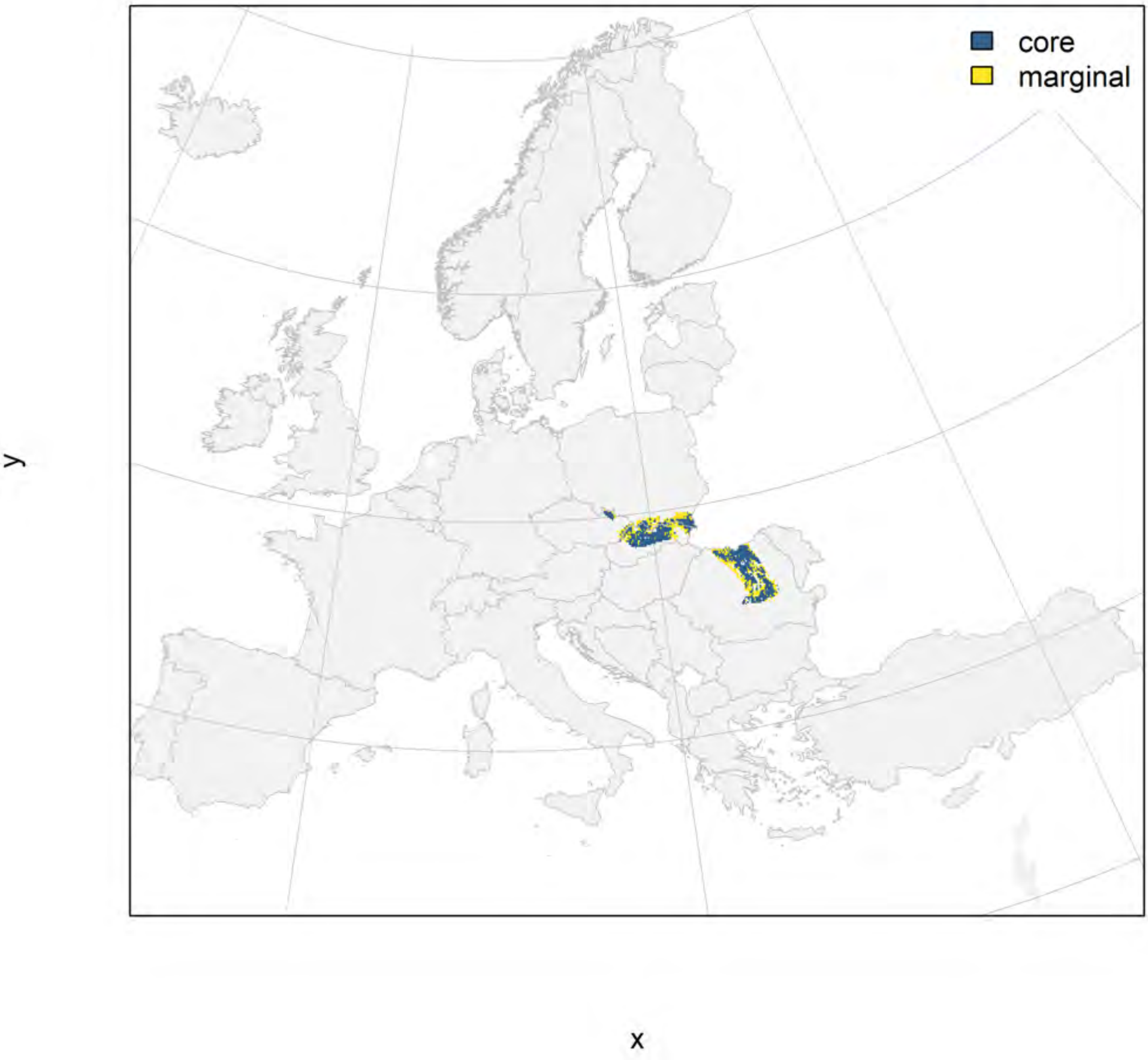

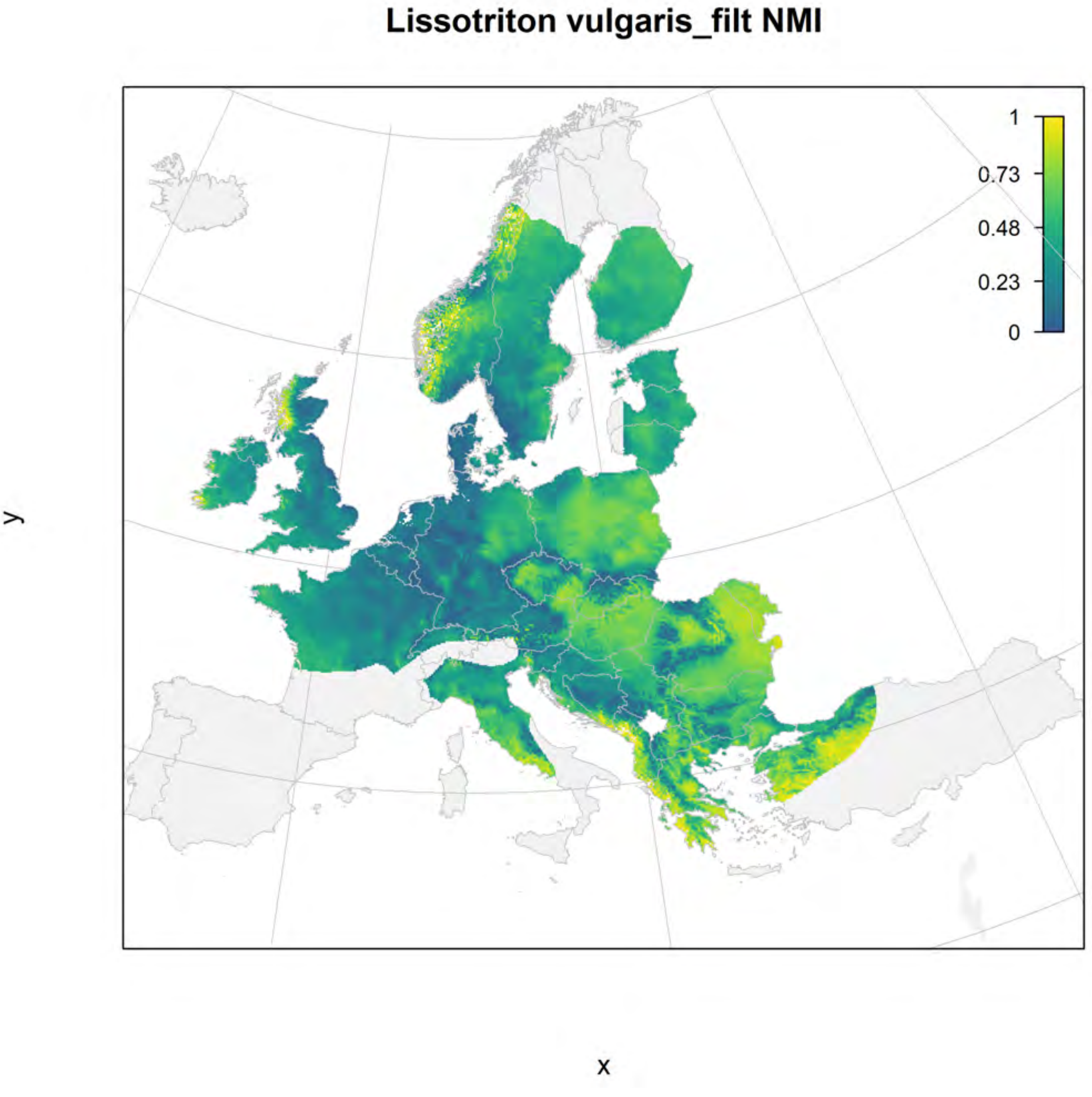
Lissotriton vulgaris_filt NMI
1
0.73
0.48
0.23
0
y
x

### Lissotriton vulgaris_filt current marginality

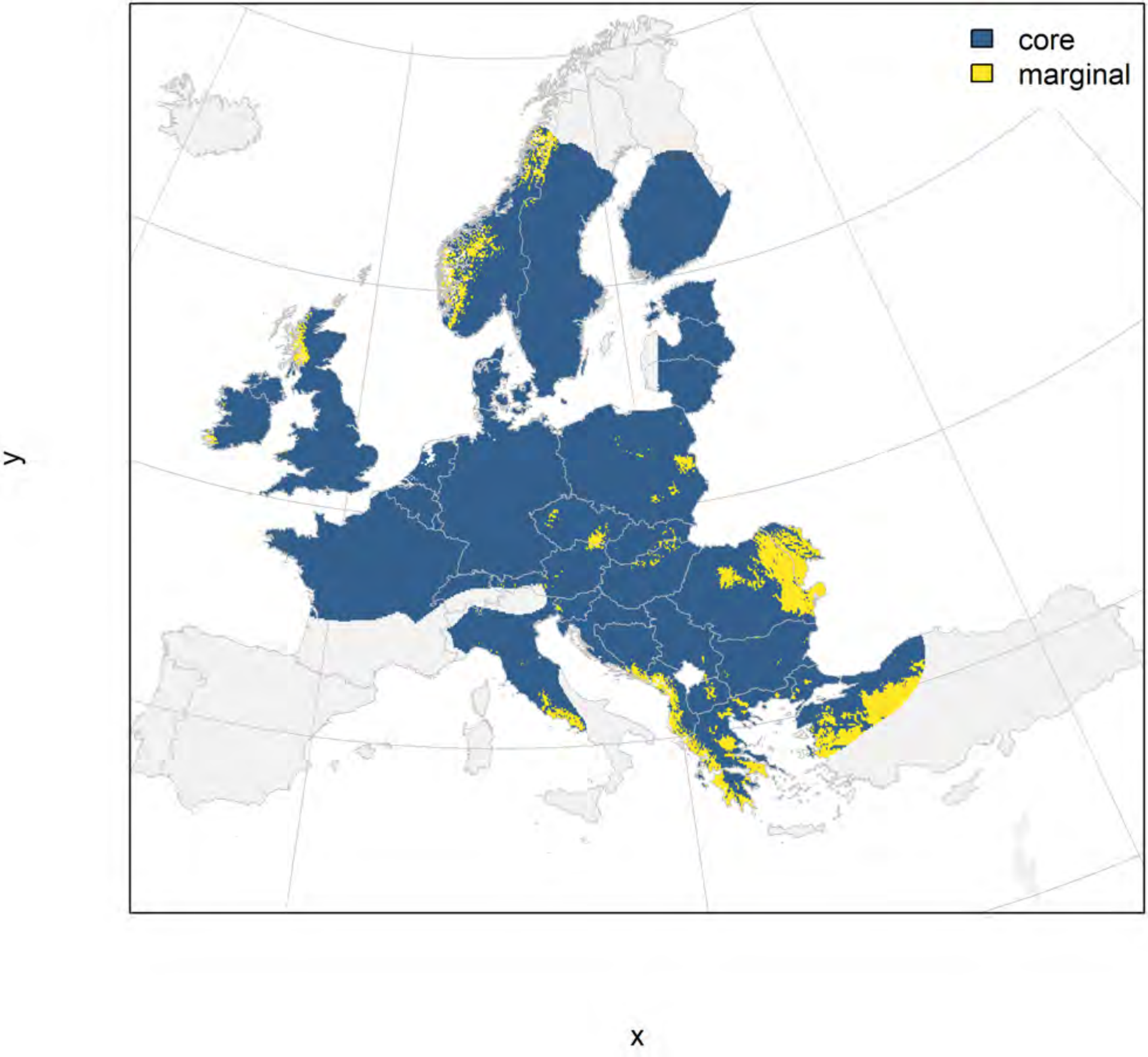

#### Lissotriton vulgaris_filt future marginality

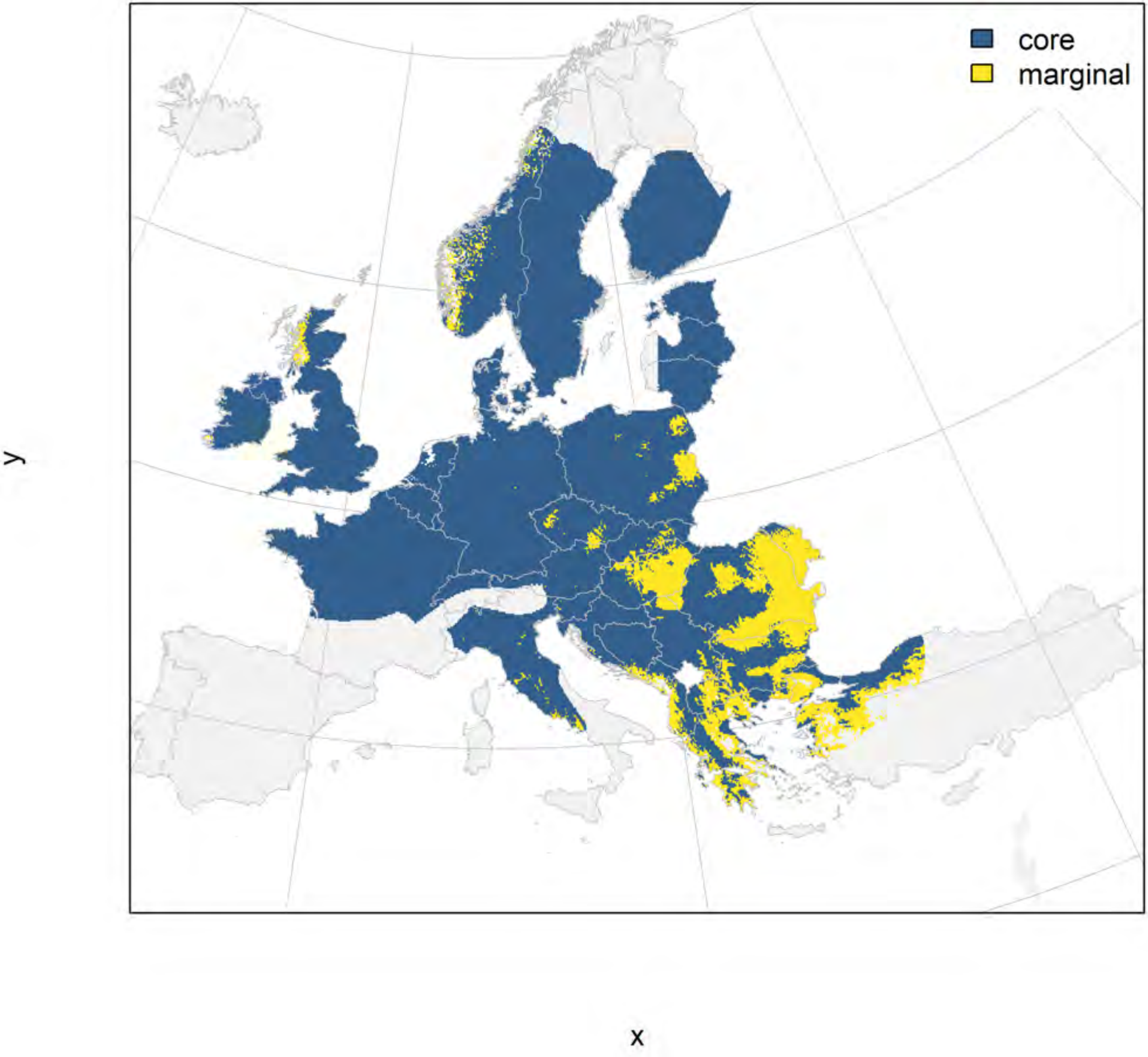

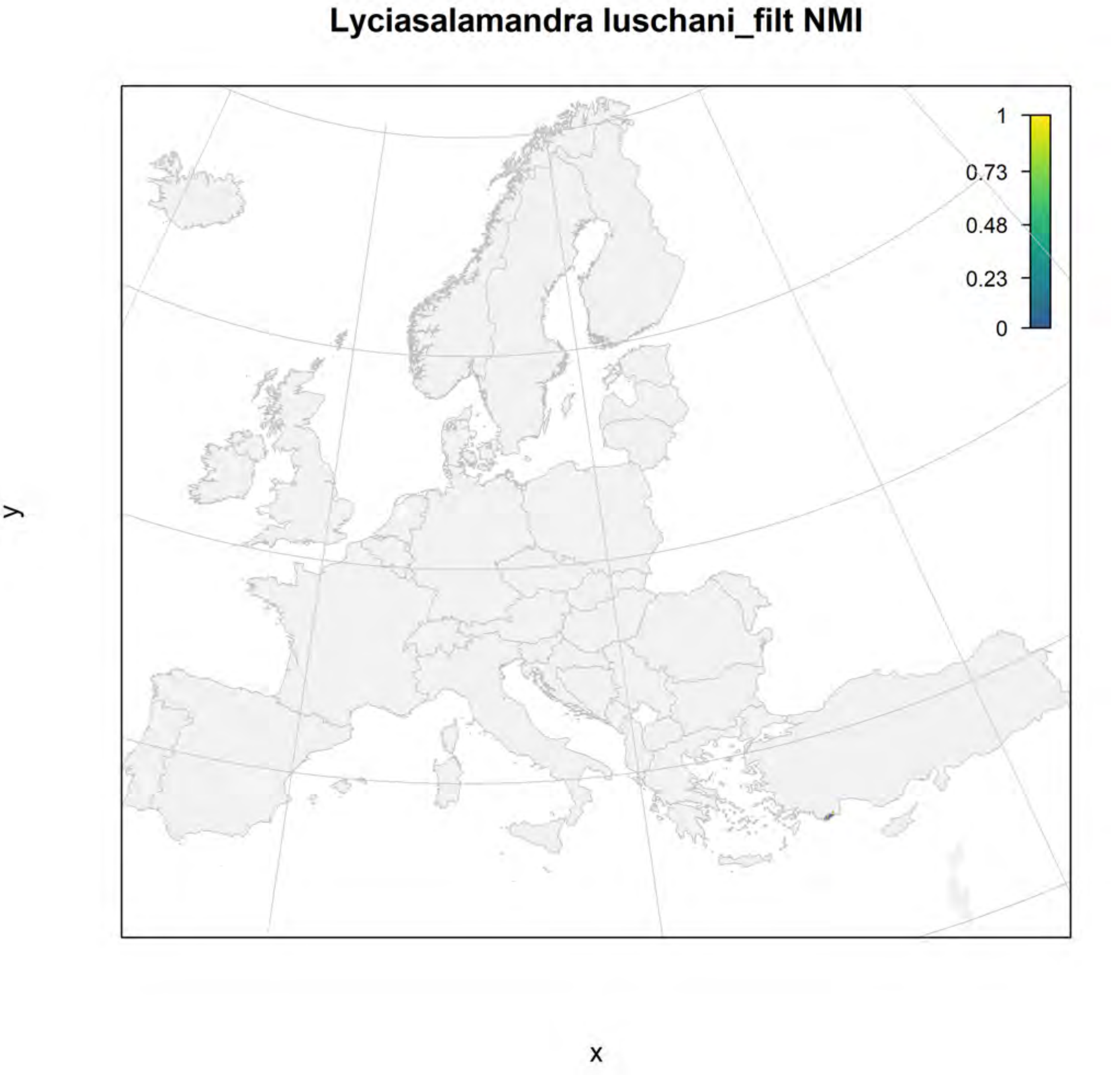
Lyciasalamandra luschani_filt NMI
1
0.73
0.48
0.23
0
y
x

#### Lyciasalamandra luschani_filt current marginality

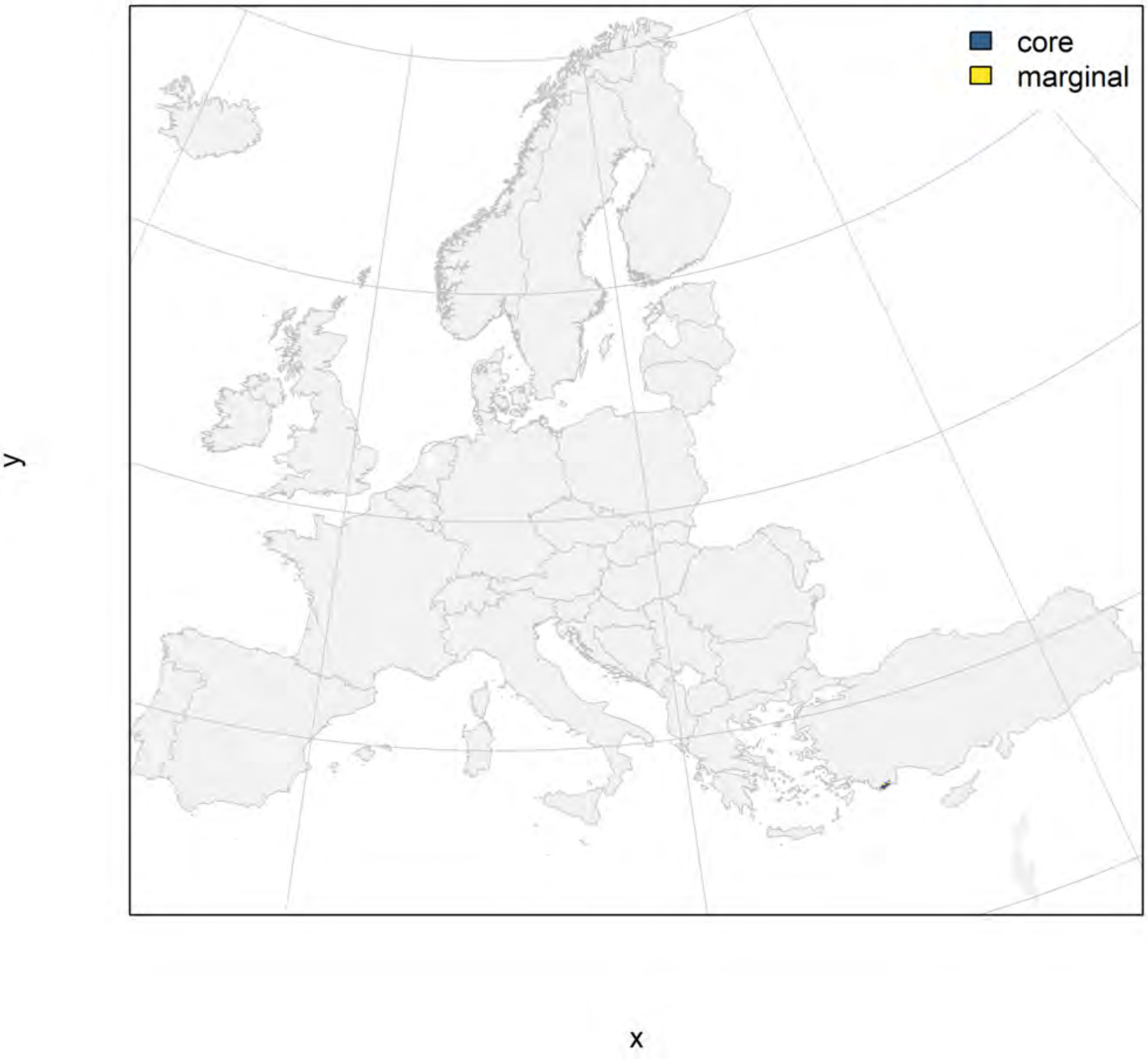

#### Lyciasalamandra luschani_filt future marginality

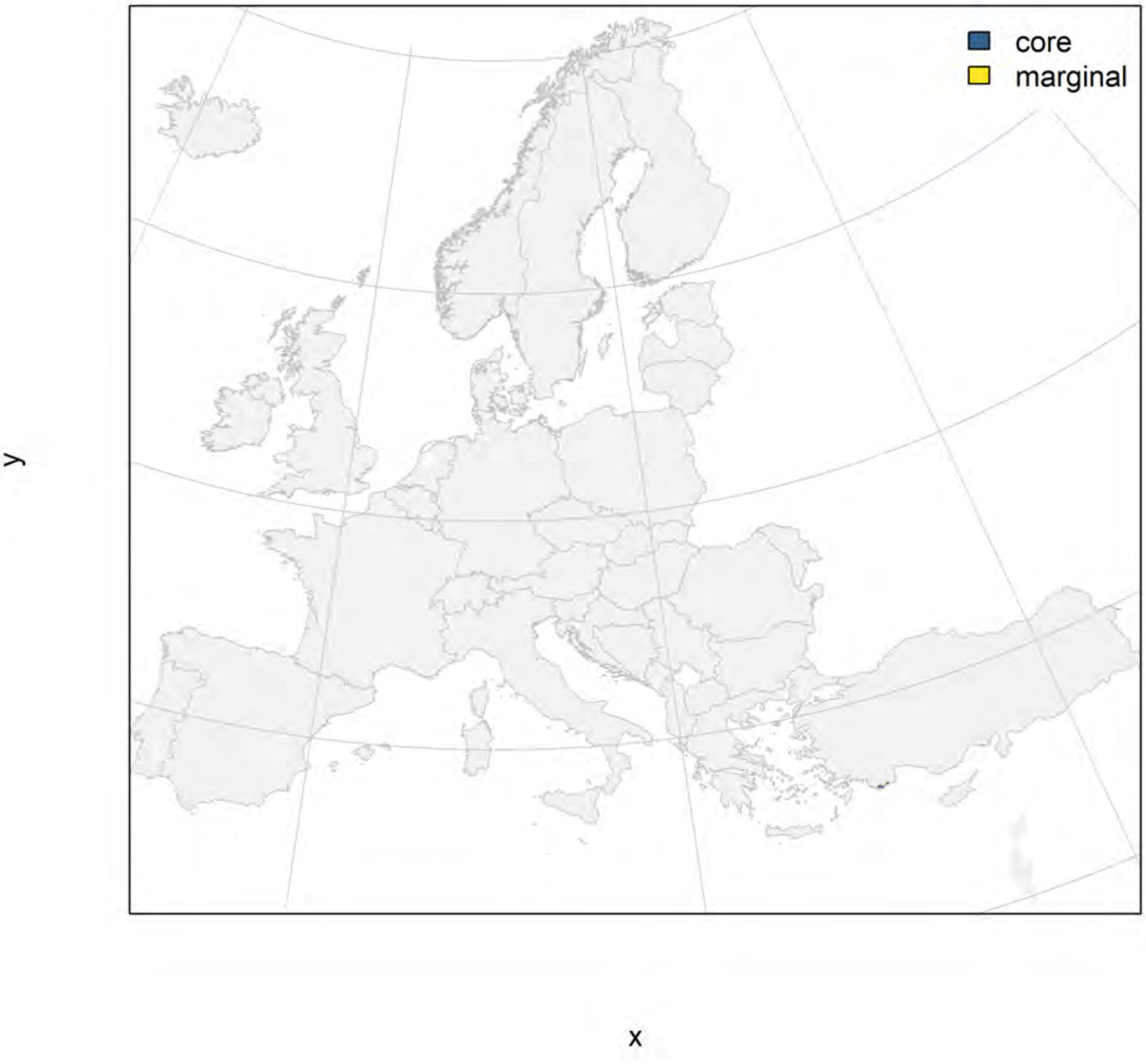

### Pleurodeles waltl_filt NMI

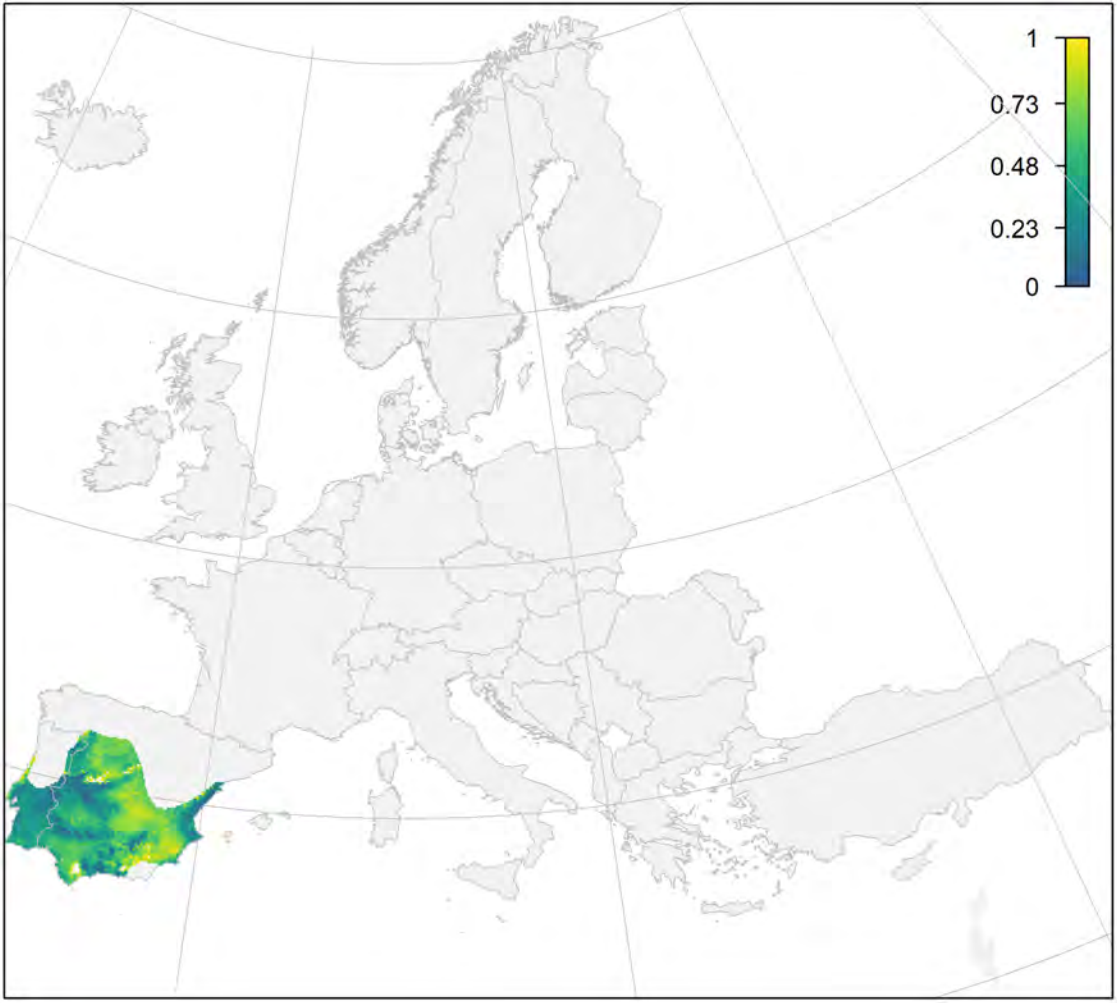


y

x

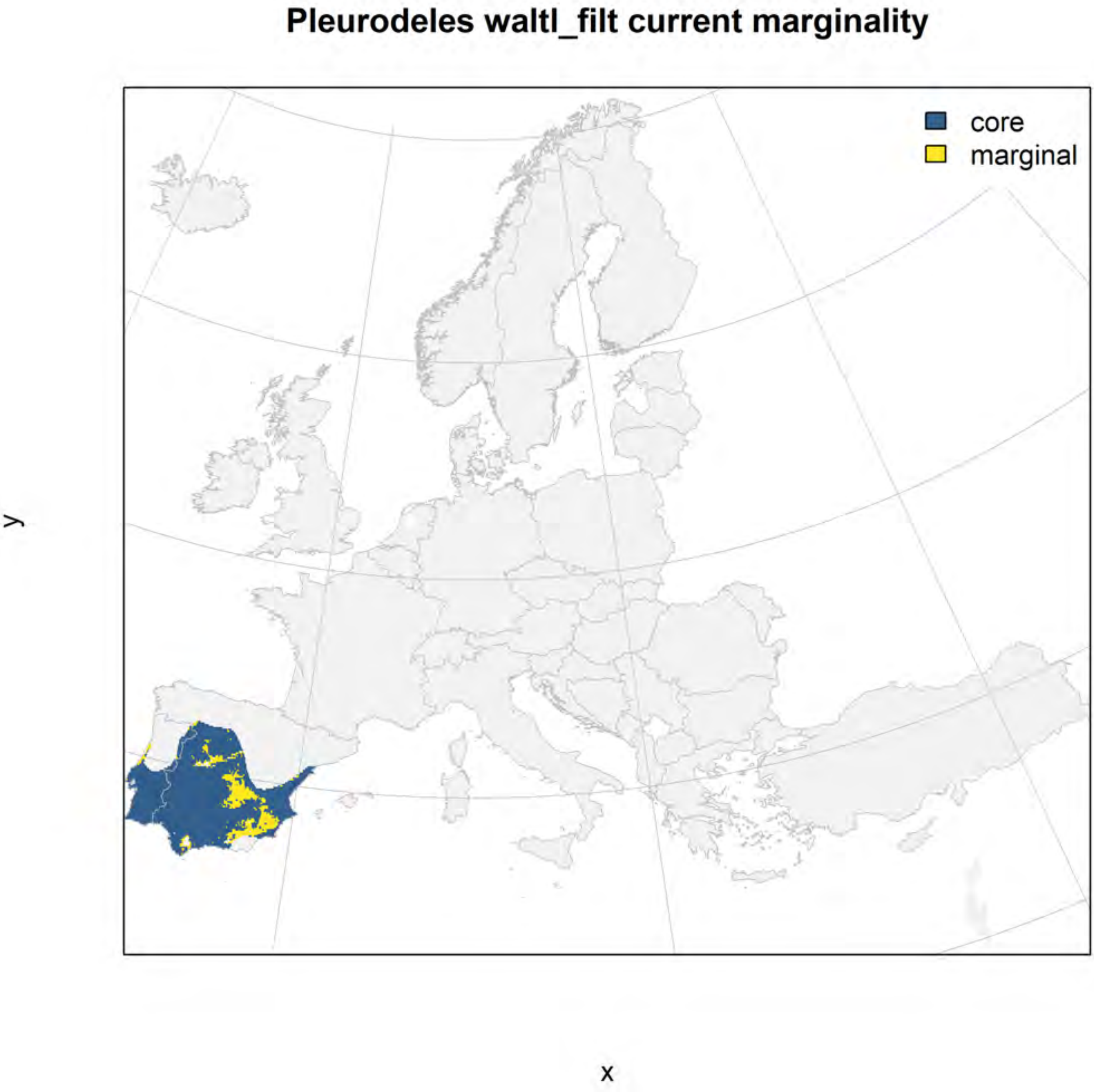
Pleurodeles waltl_filt current marginality
core
marginal
y
x

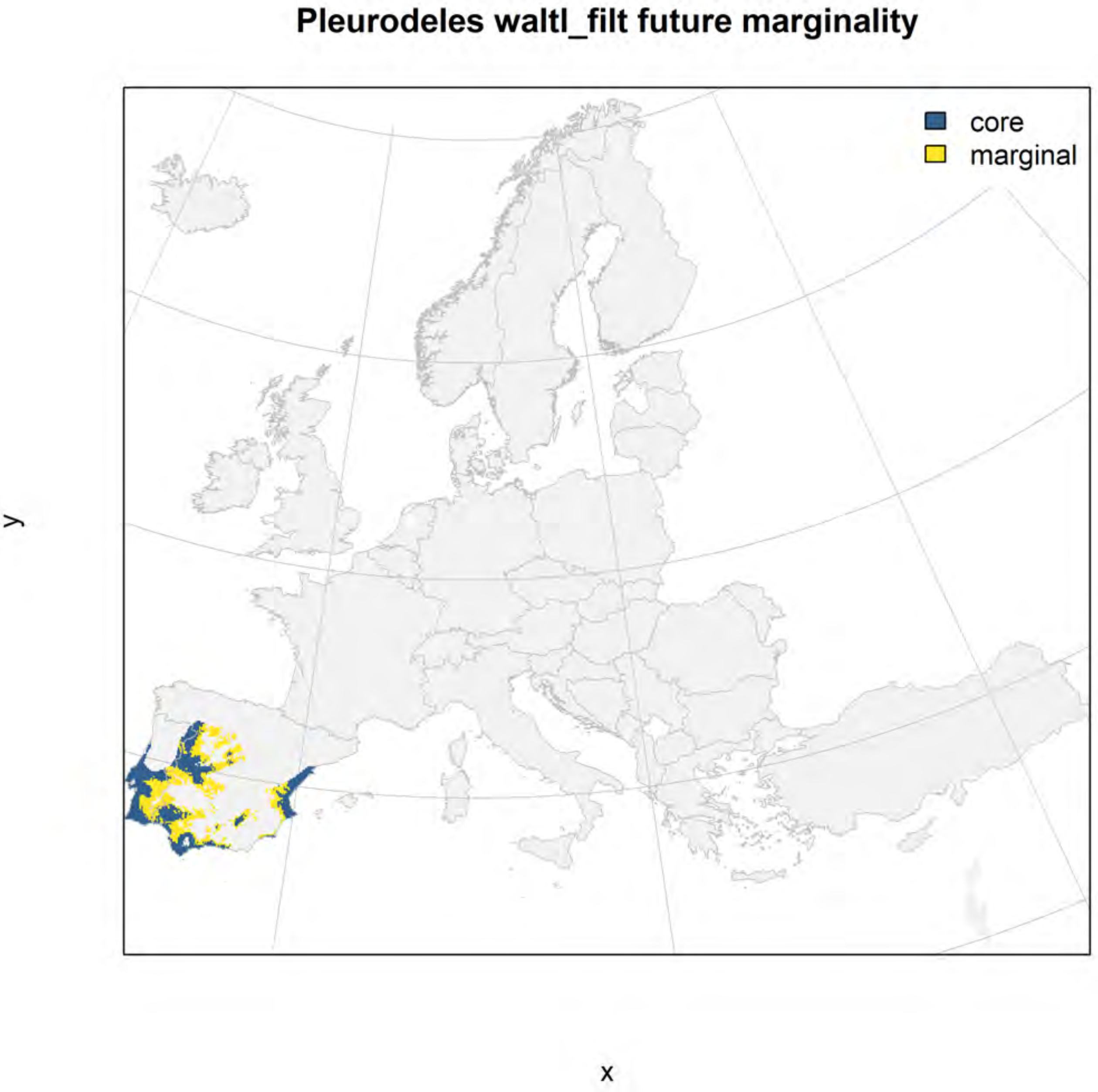
Pleurodeles waltl_filt future marginality
core
marginal
y
x

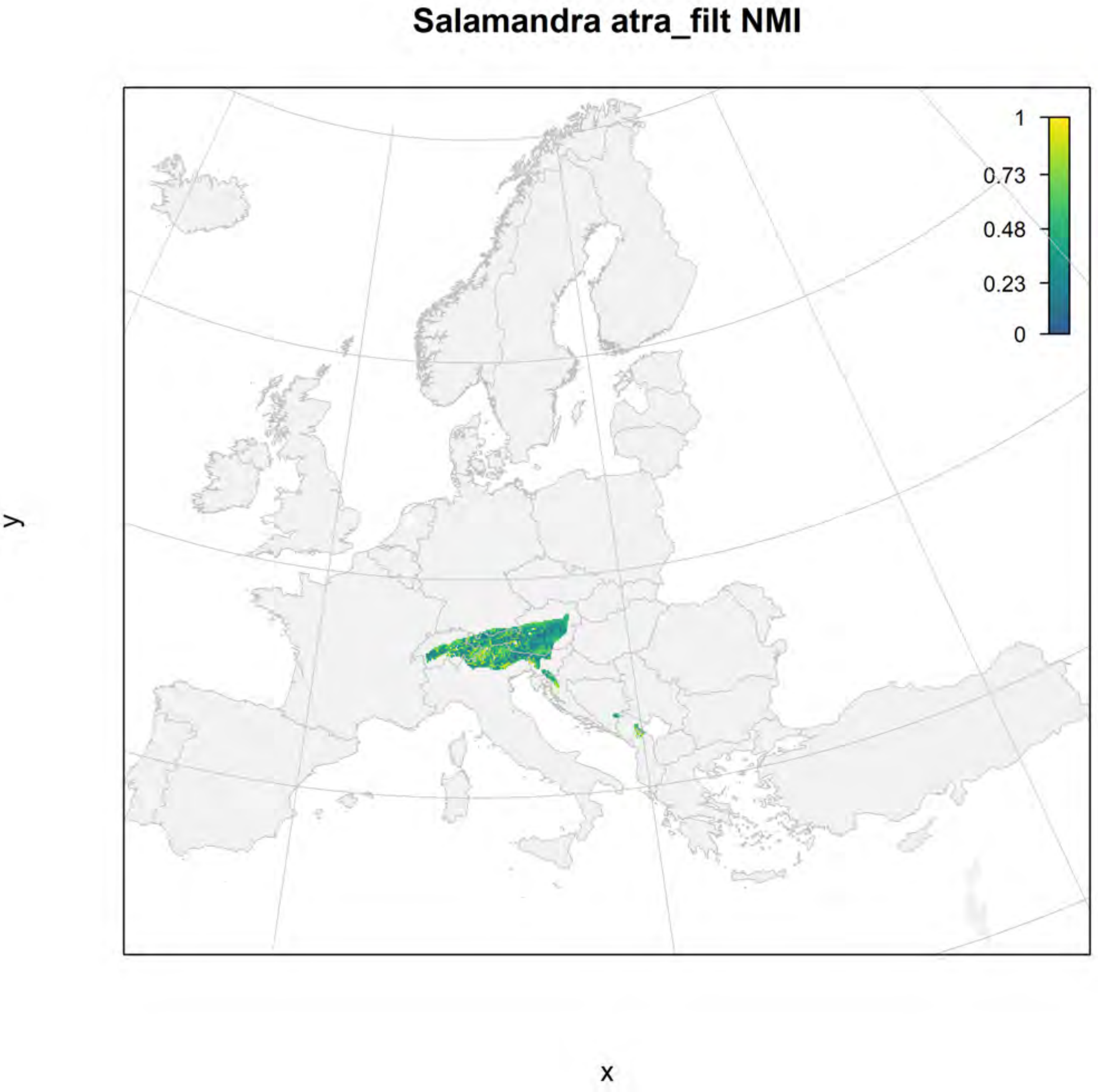

Salamandra atra_filt NMI
1
0.73
0.48
0.23
0
y
x

#### Salamandra atra_filt current marginality

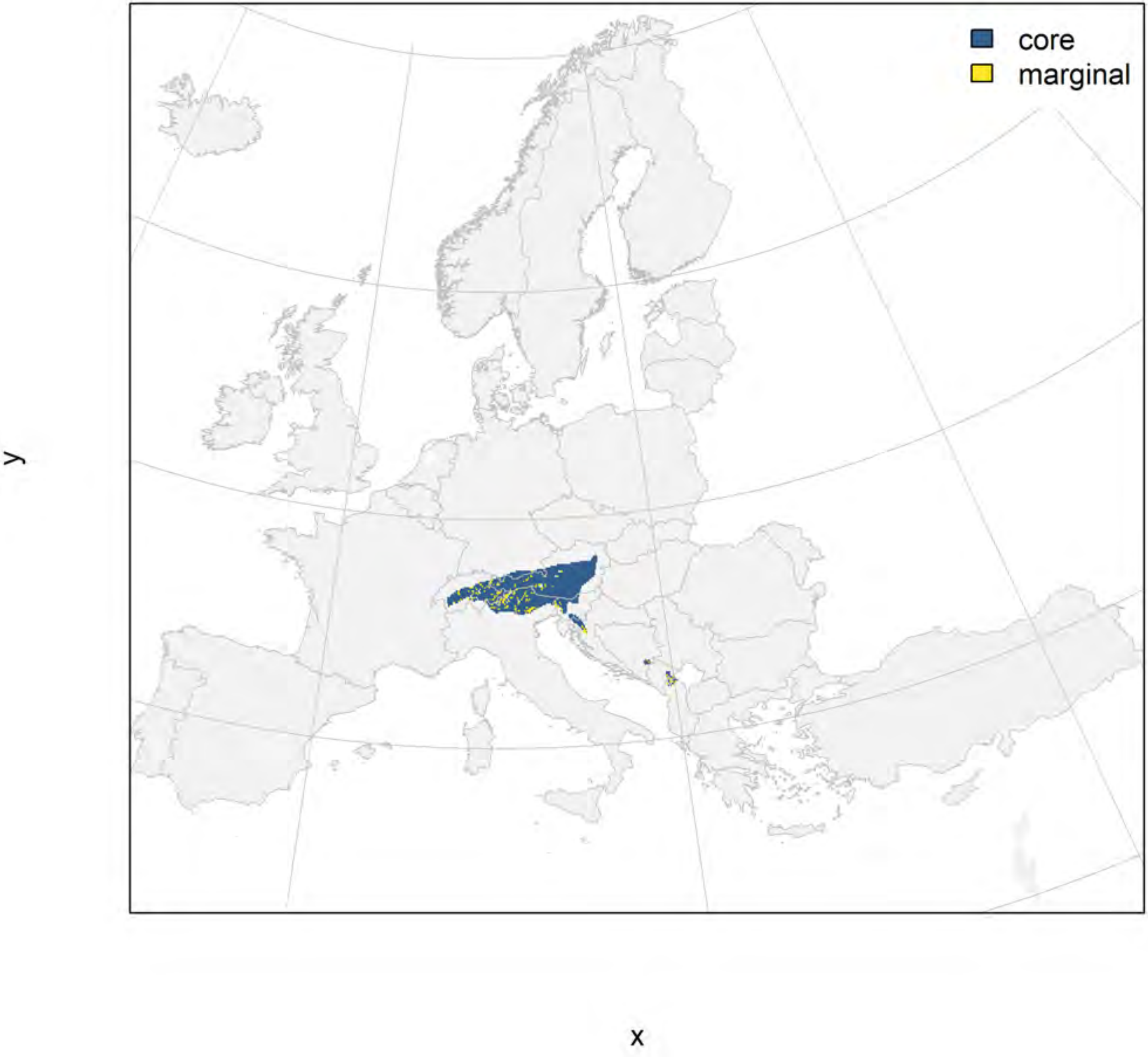

### Salamandra atra_filt future marginality

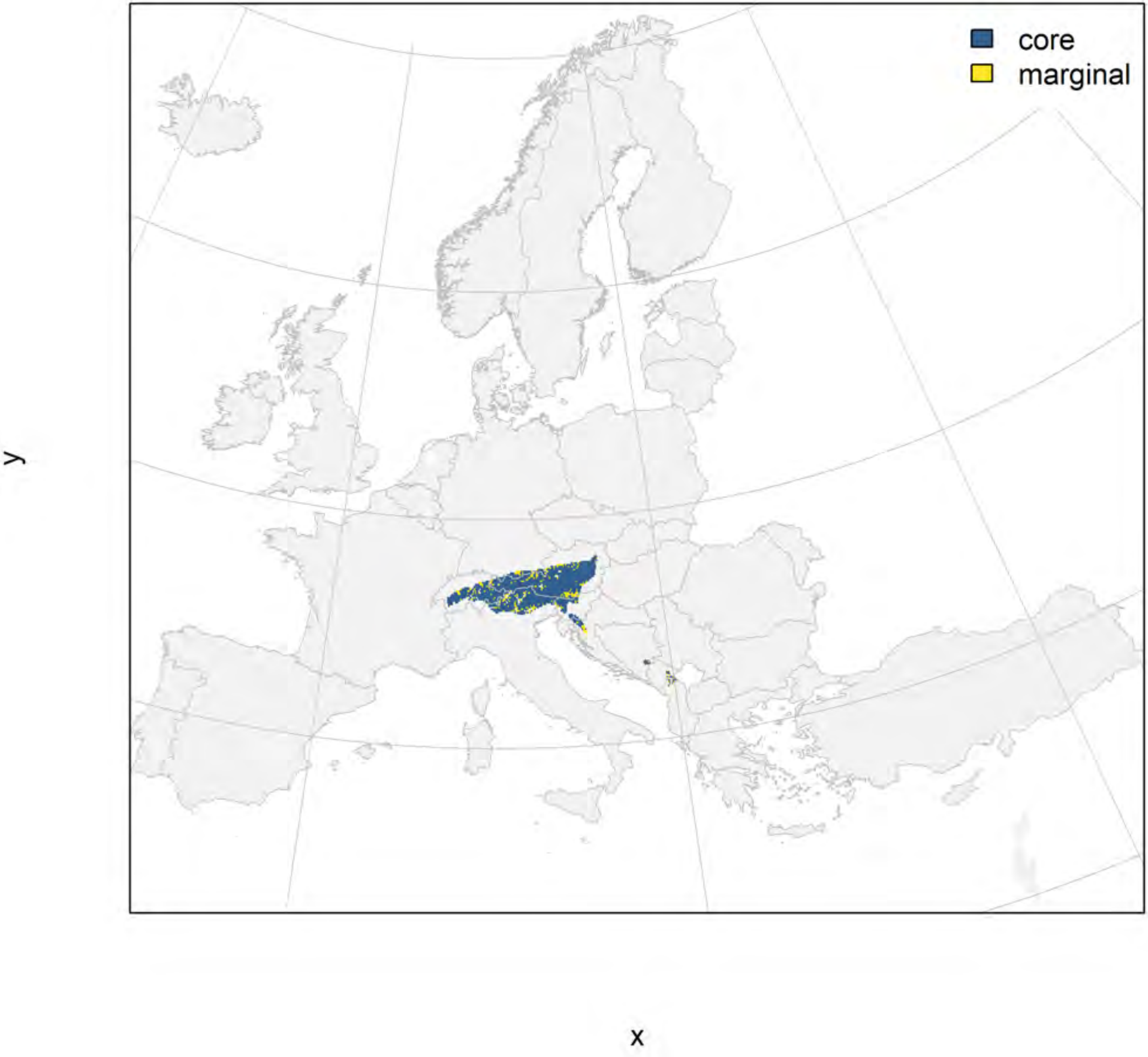

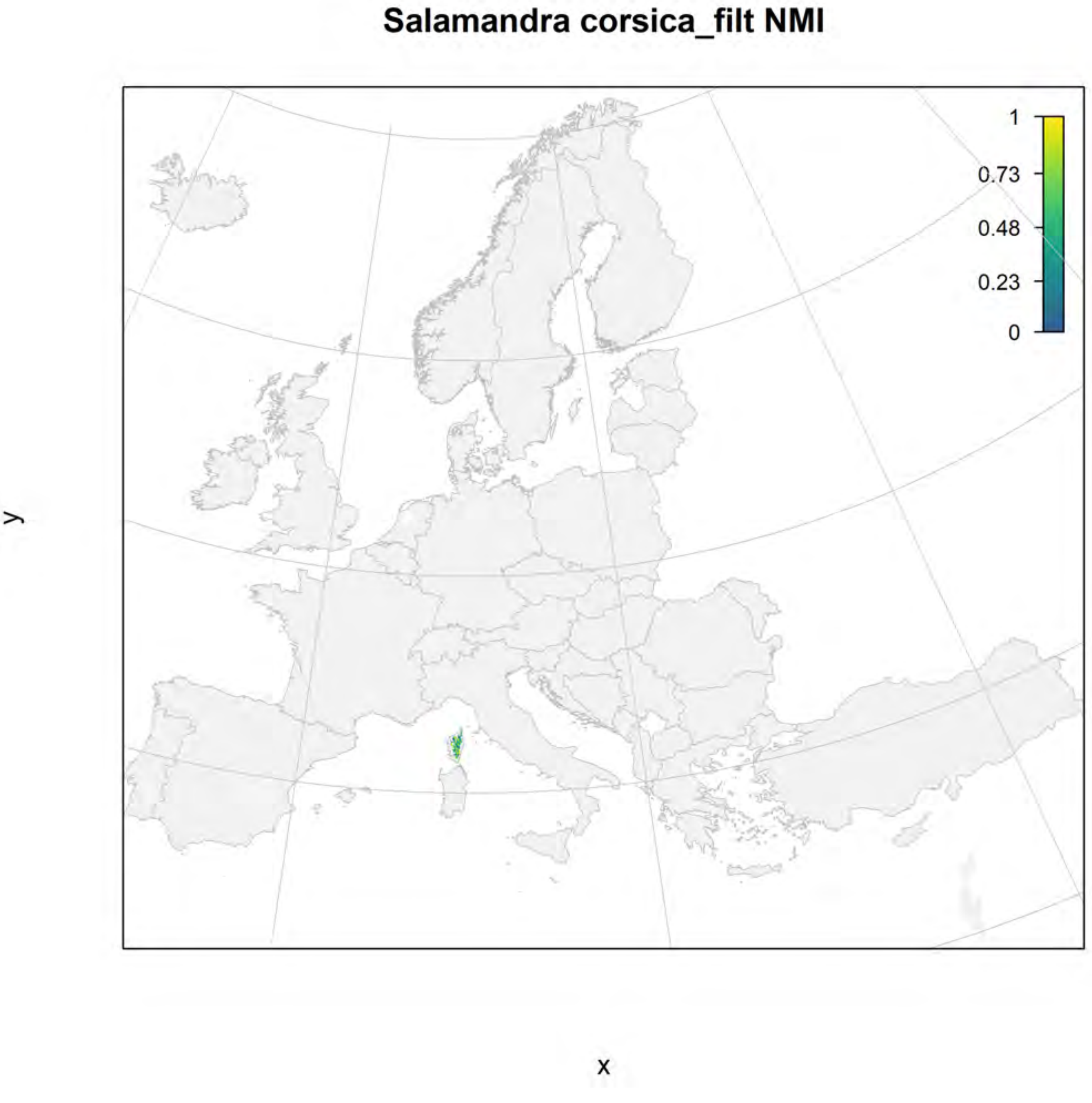
Salamandra corsica_filt NMI
1
0.73
0.48
0.23
0
y
x

#### Salamandra corsica_filt current marginality

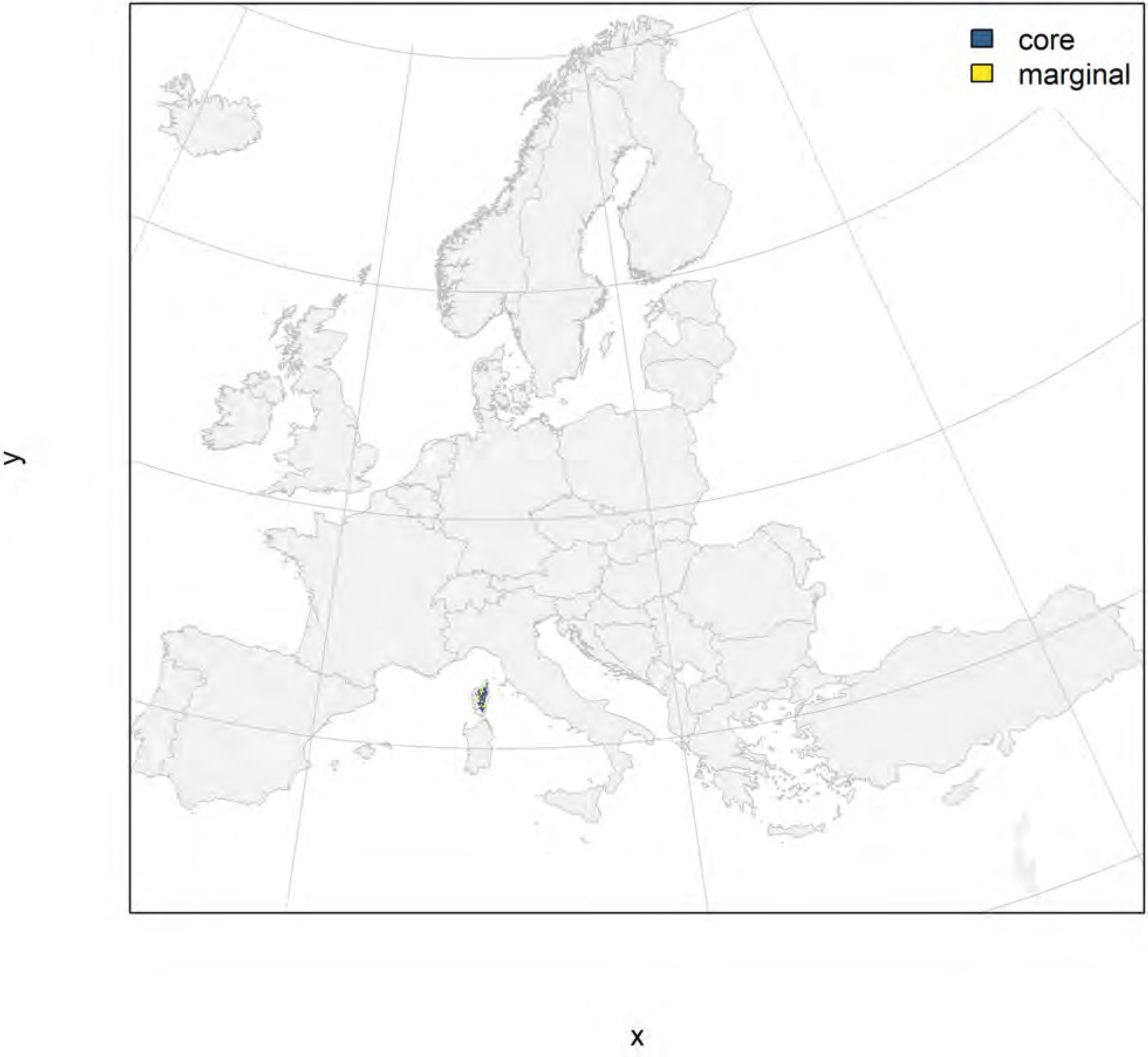

### Salamandra corsica_filt future marginality

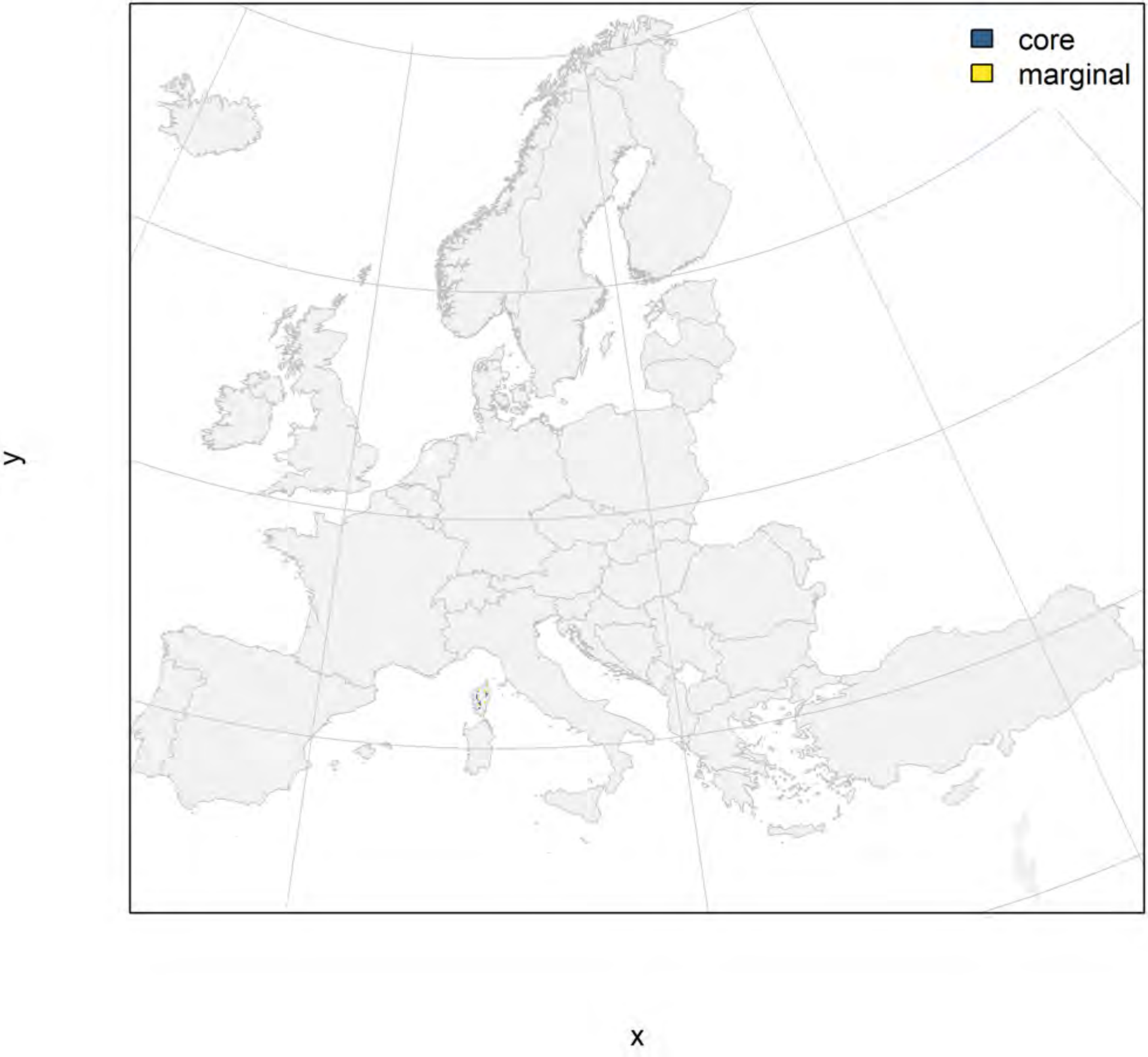

### Salamandra lanzai_filt NMI

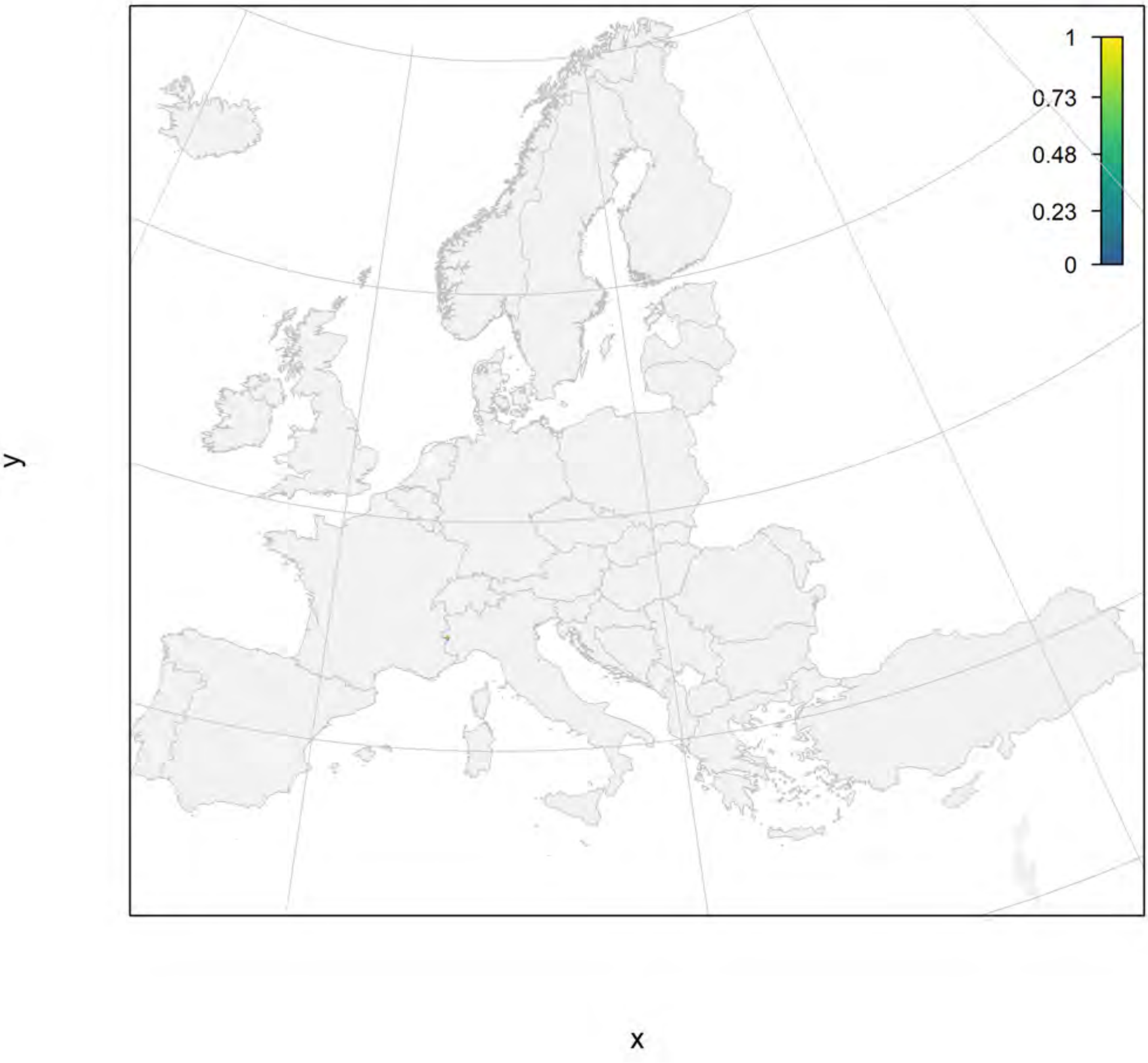

#### Salamandra lanzai_filt current marginality

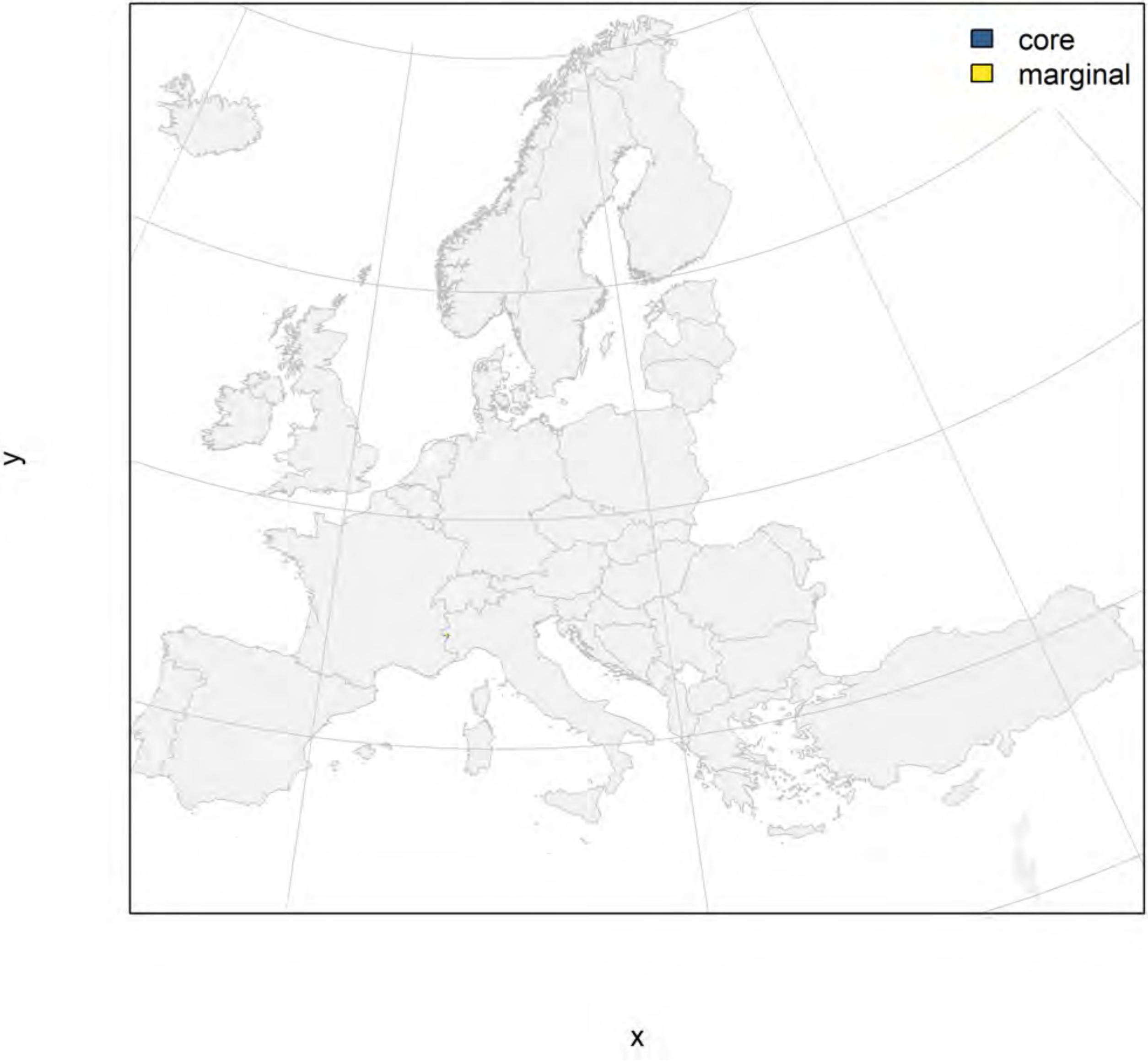

#### Salamandra lanzai_filt future marginality

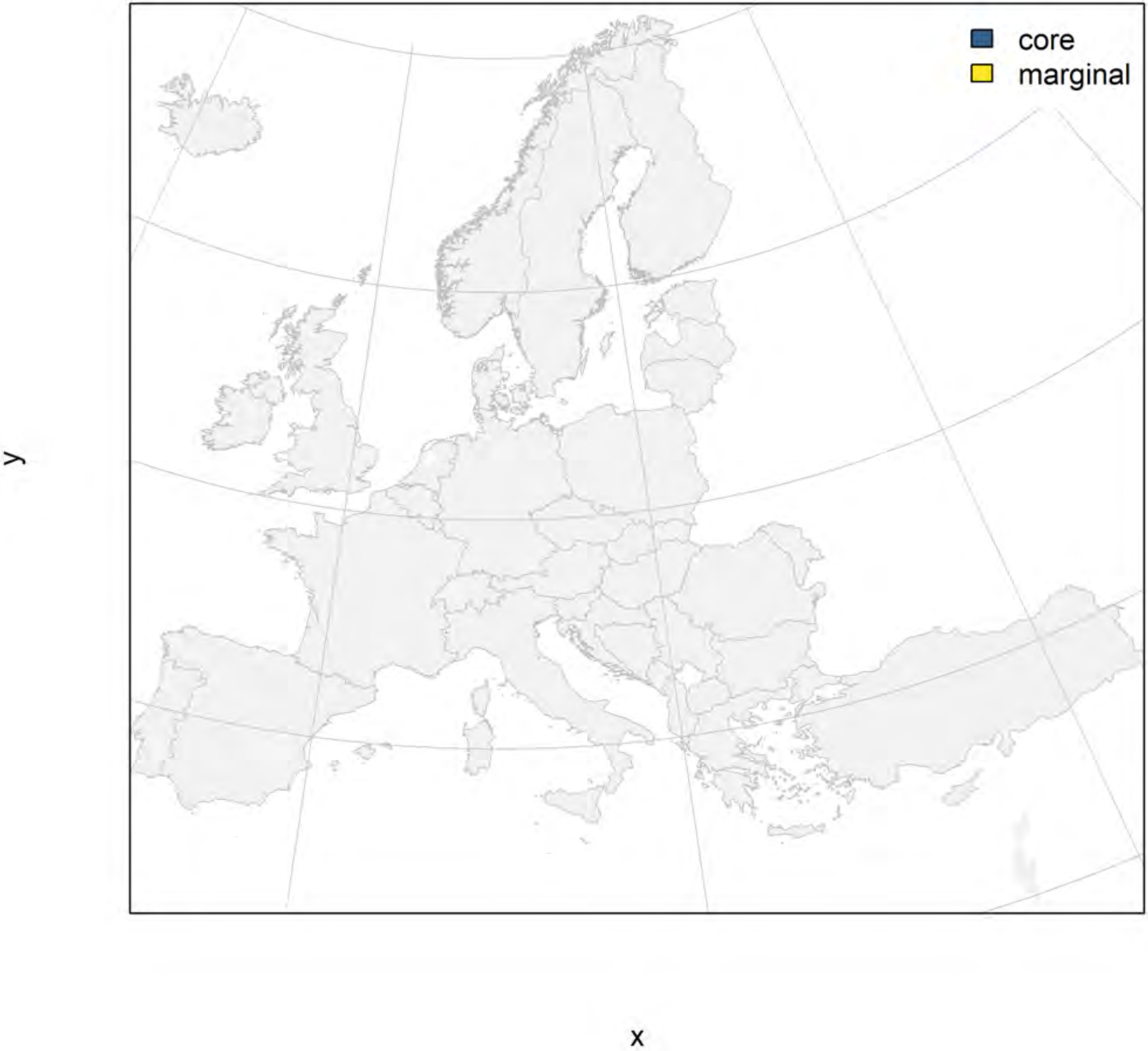

#### Salamandra salamandra_filt NMI

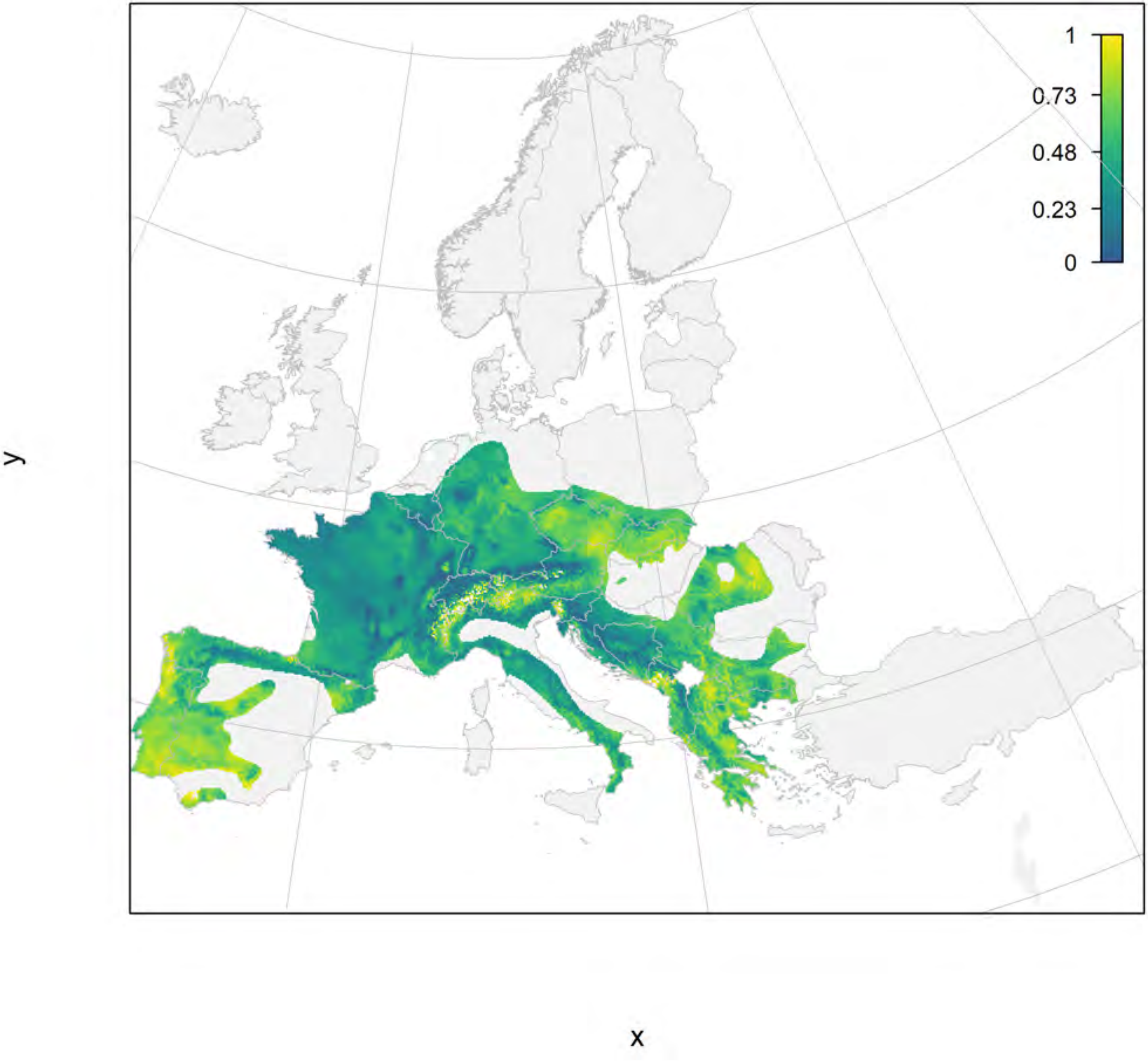

### Salamandra salamandra_filt current marginality

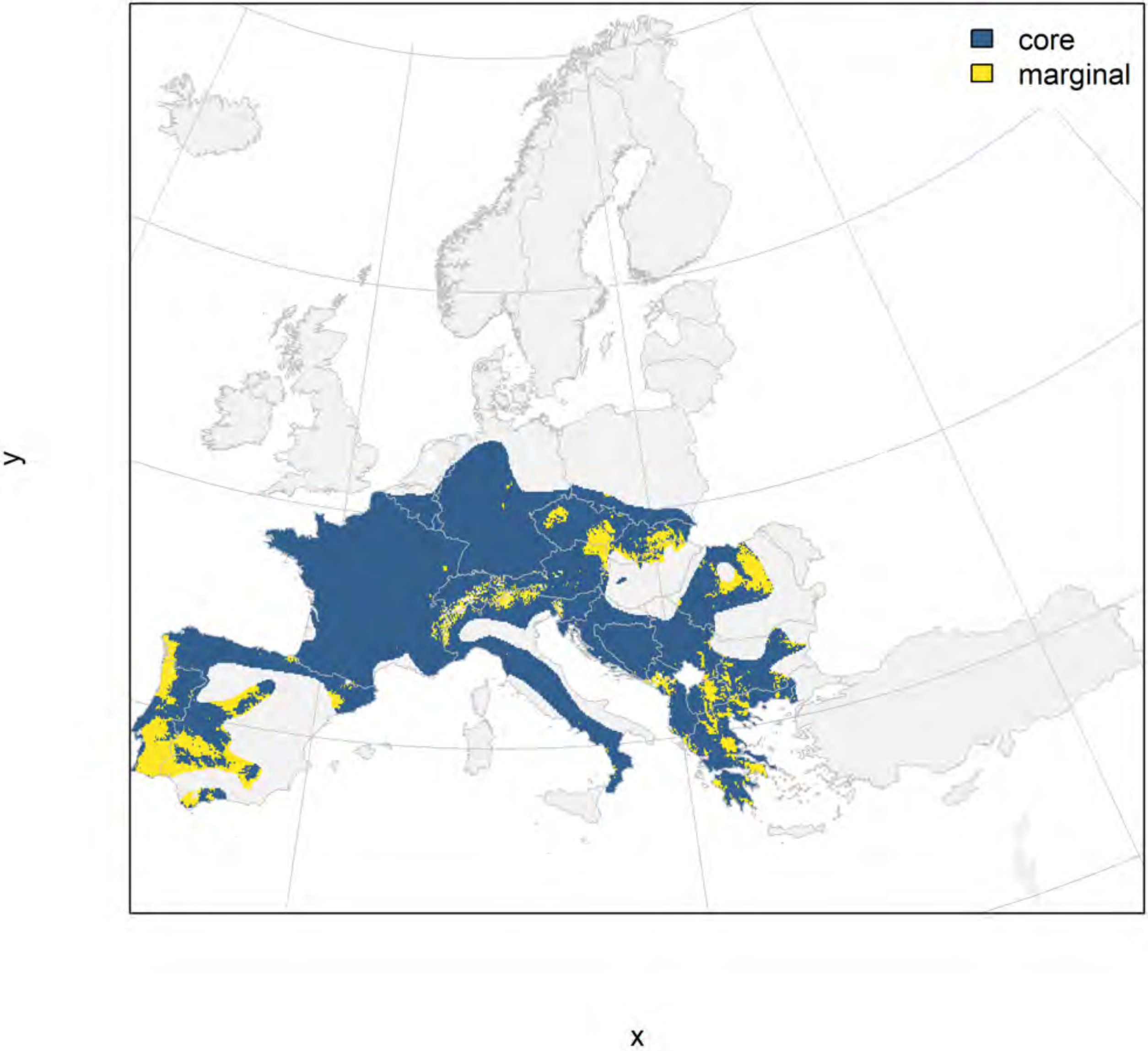

#### Salamandra salamandra_filt future marginality

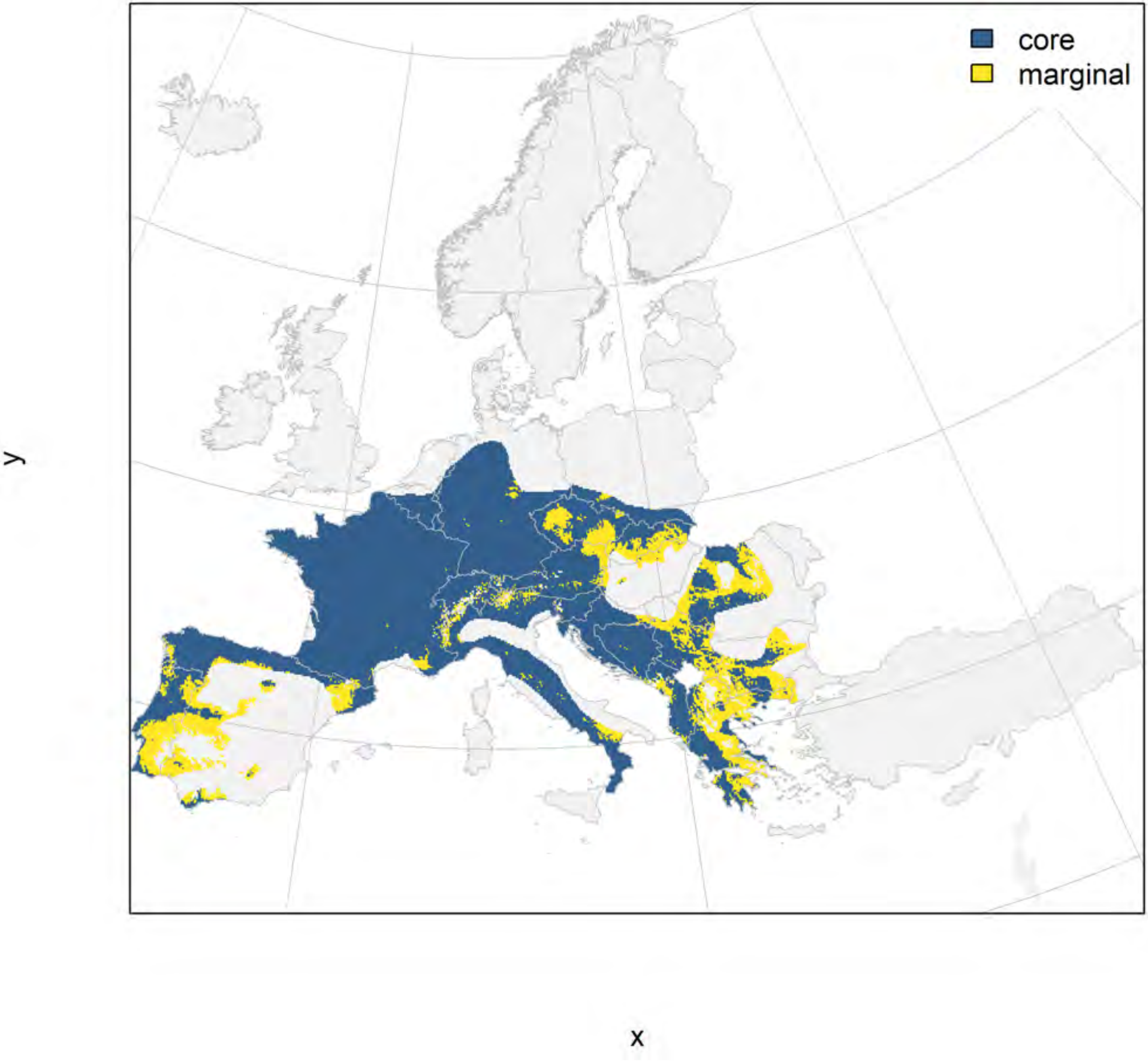

#### Salamandrina perspicillata_filt NMI

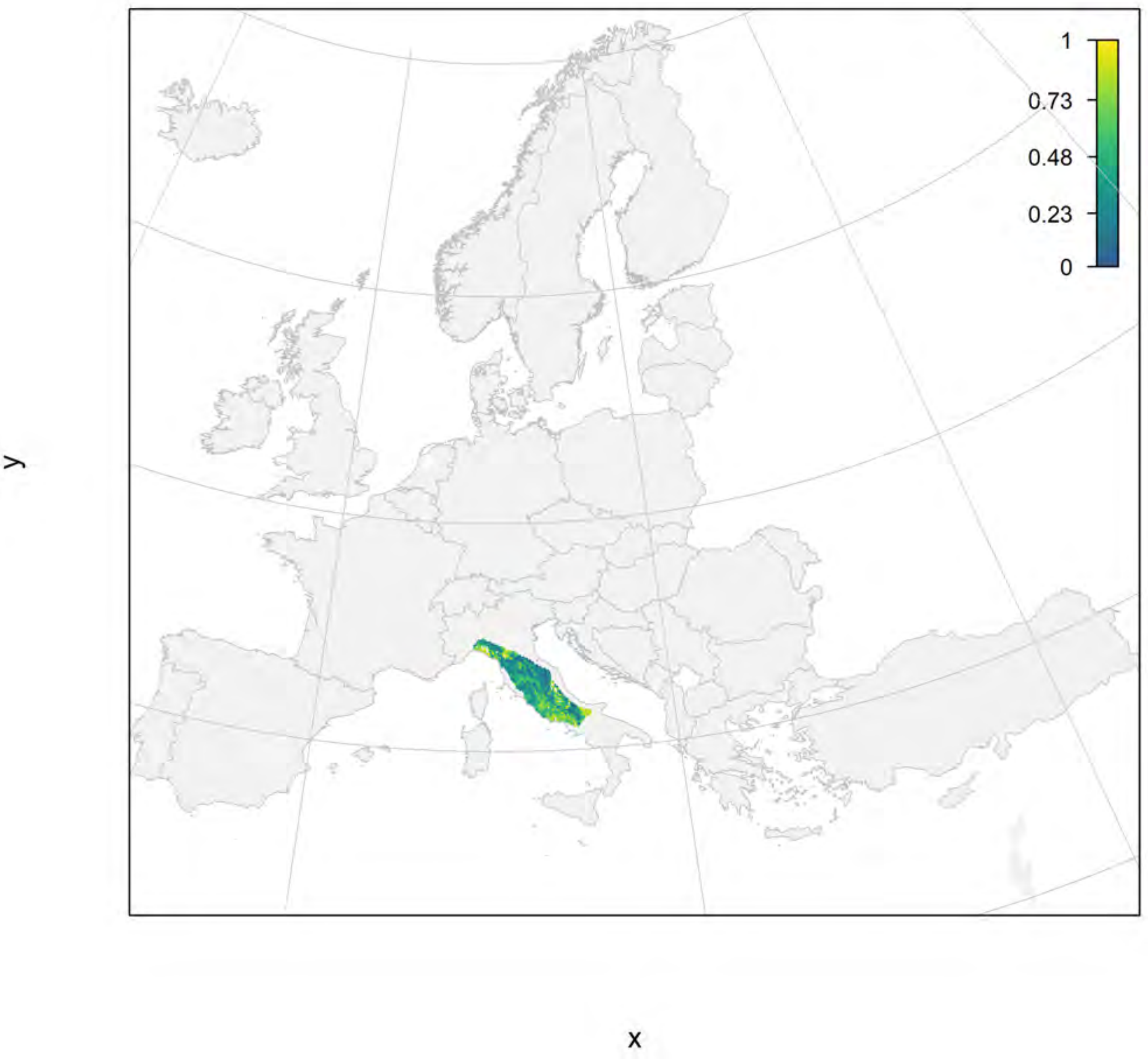

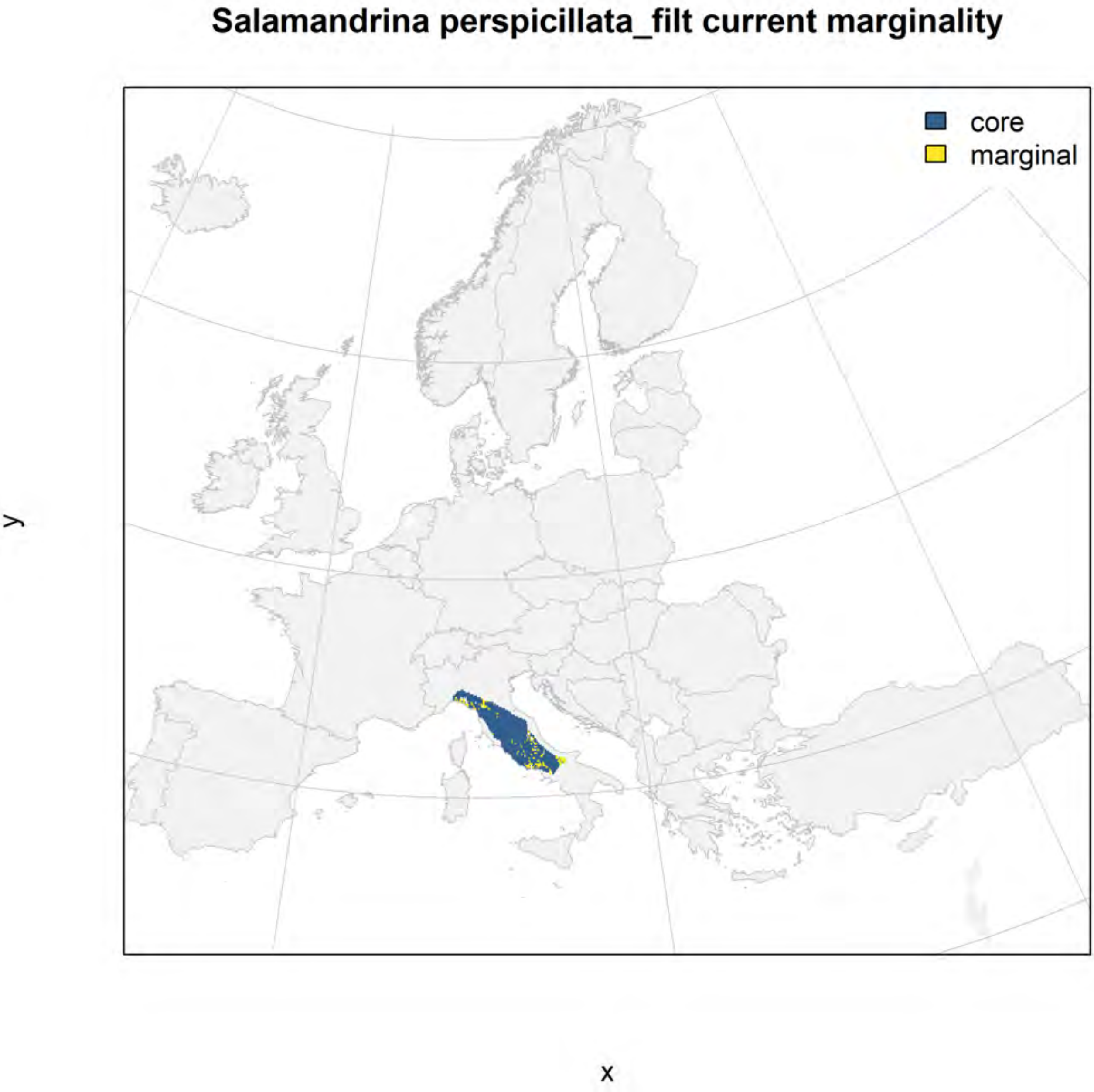
Salamandrina perspicillata_filt current marginality
core
marginal
y
x

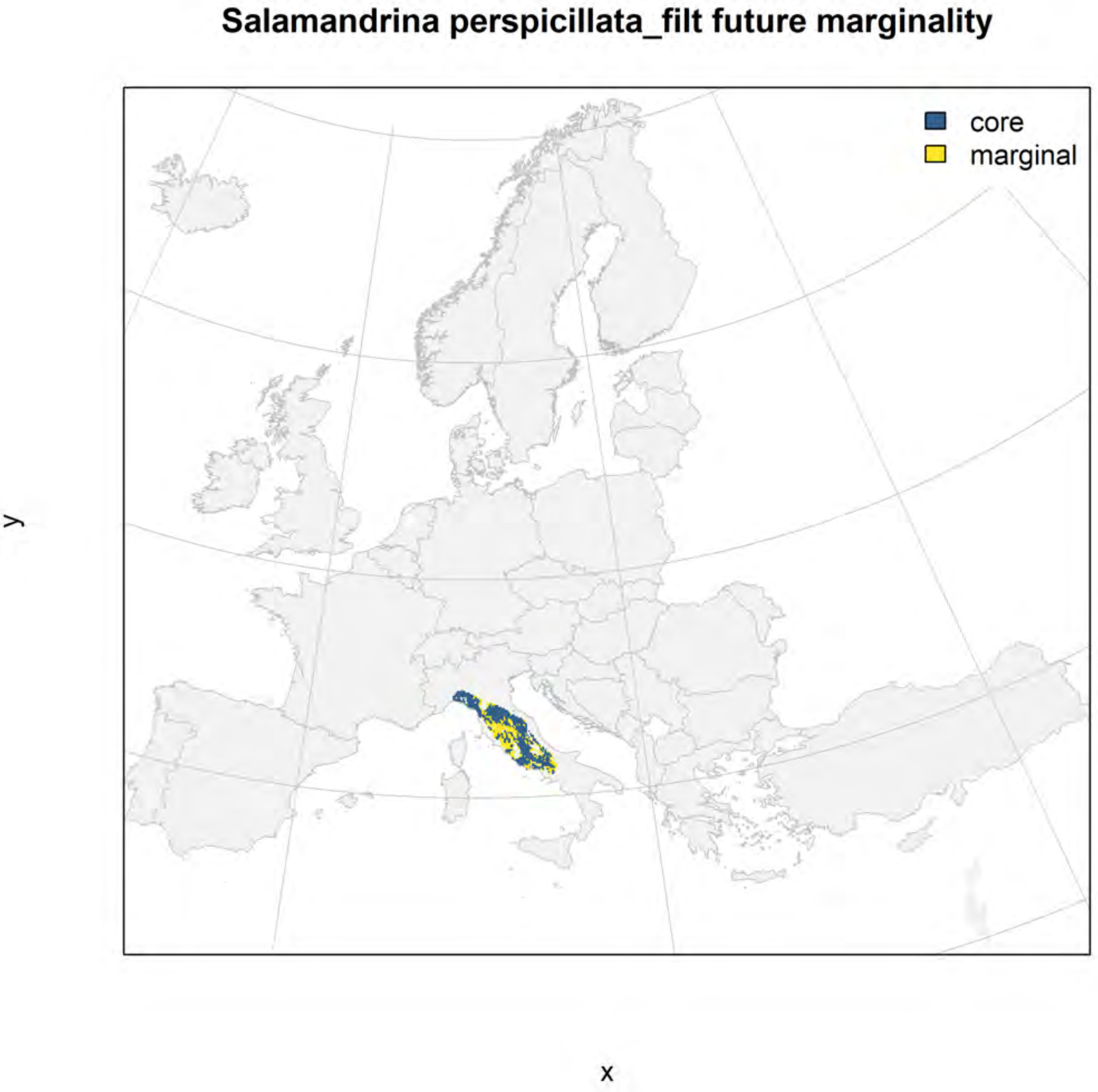
Salamandrina perspicillata_filt future marginality
core
marginal
y
x

#### Salamandrina terdigitata_filt NMI

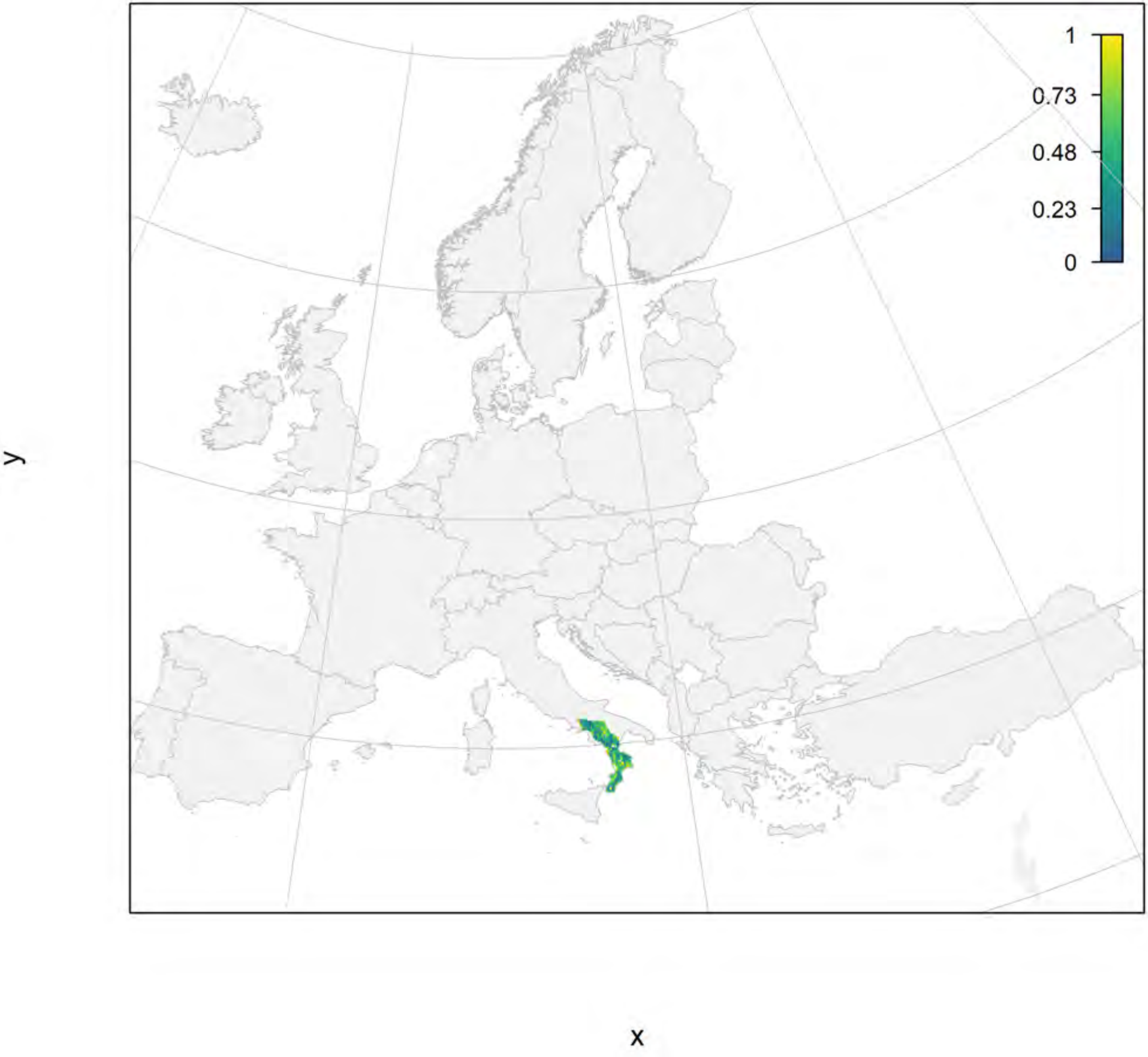

#### Salamandrina terdigitata_filt current marginality

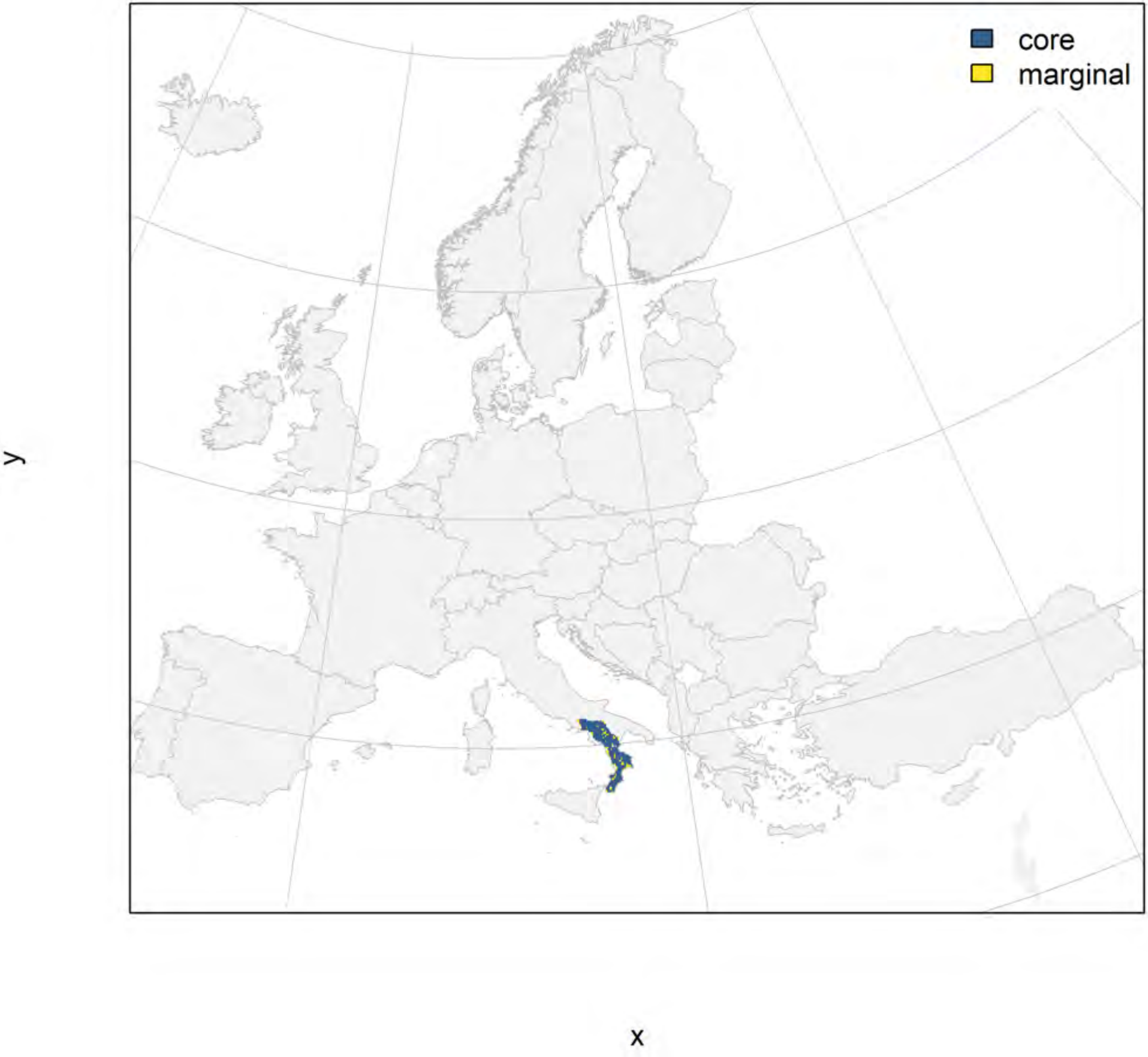

#### Salamandrina terdigitata_filt future marginality

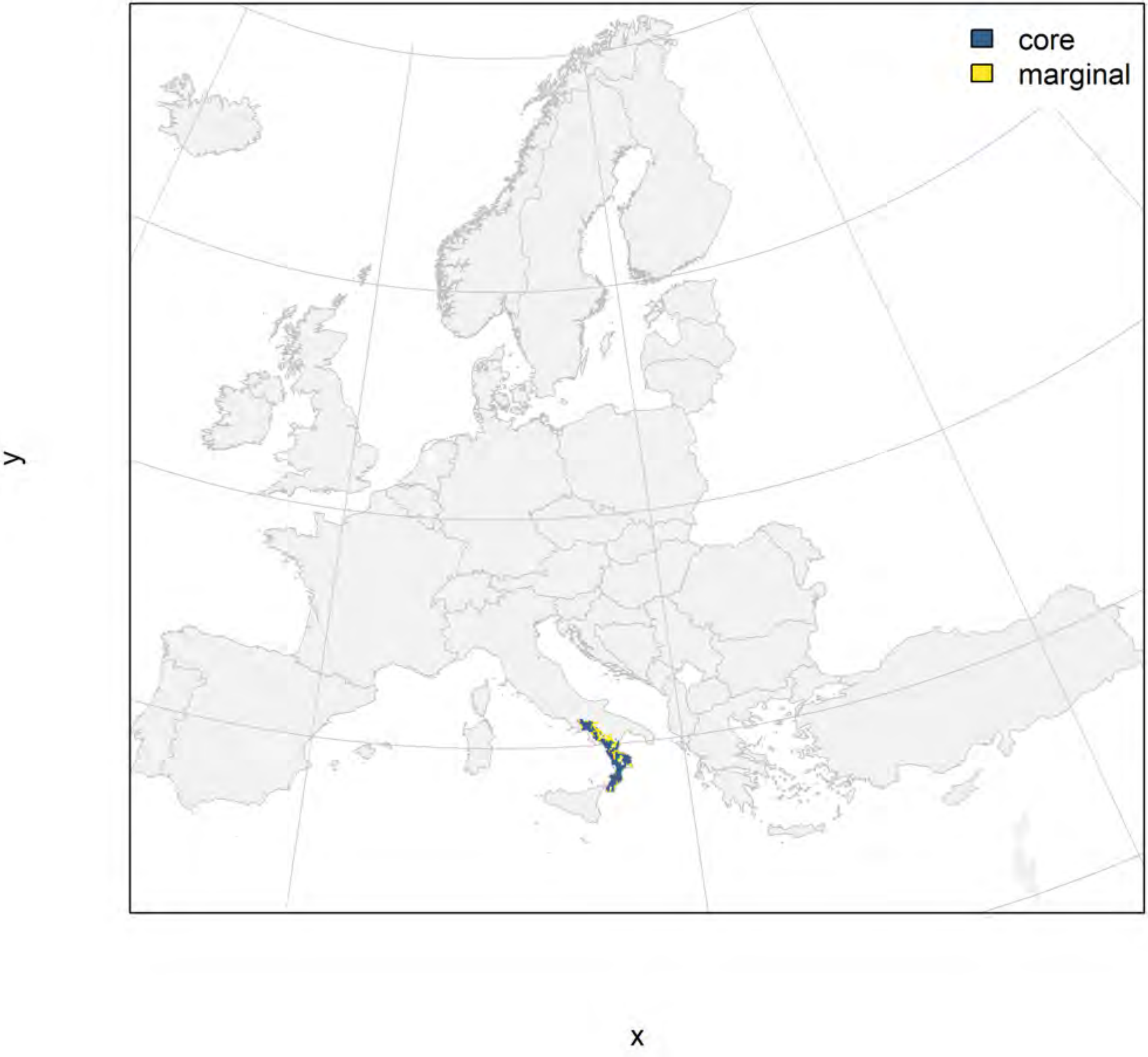

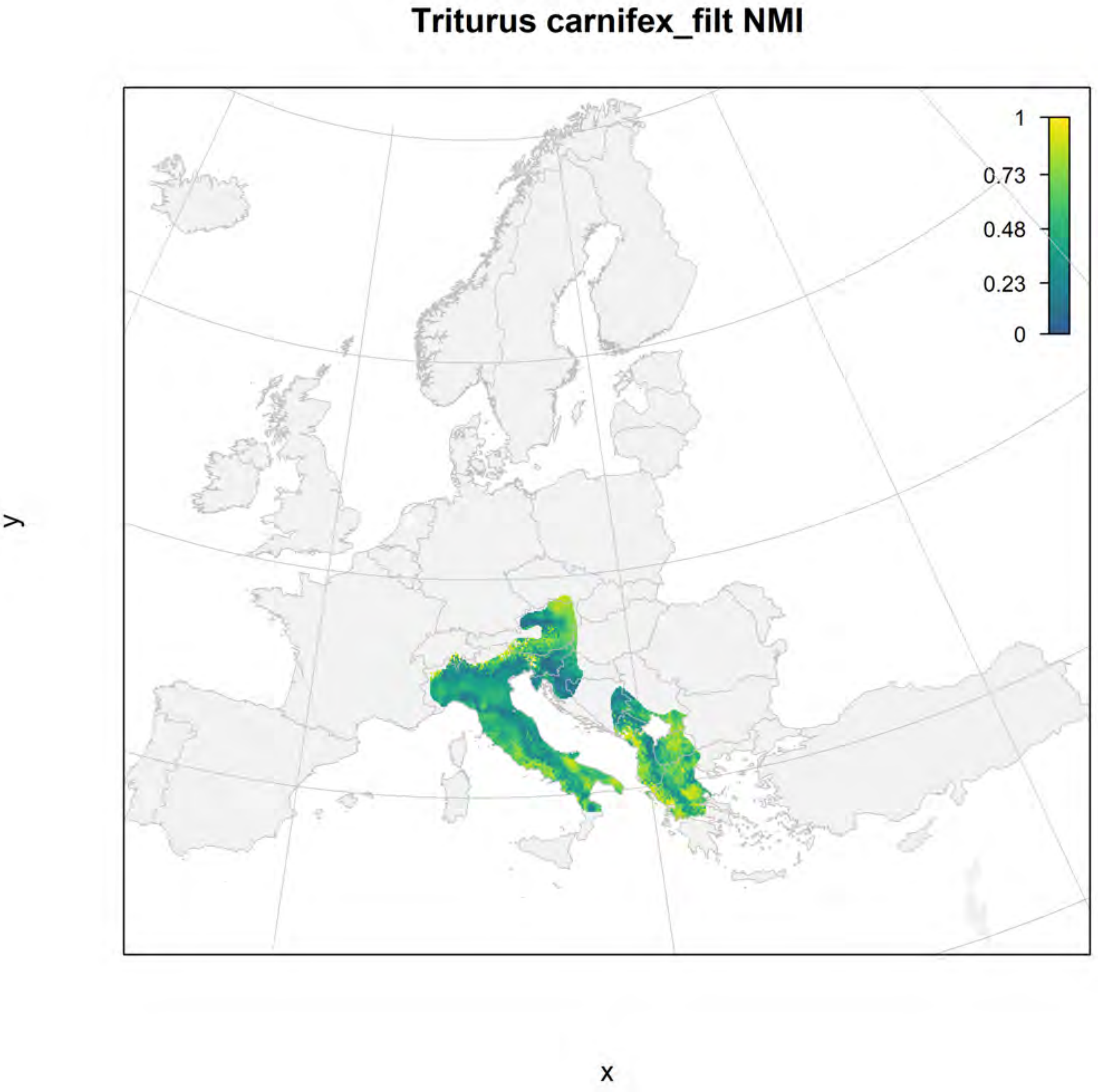
Triturus carnifex_filt NMI
1
0.73
0.48
0.23
0
y
x

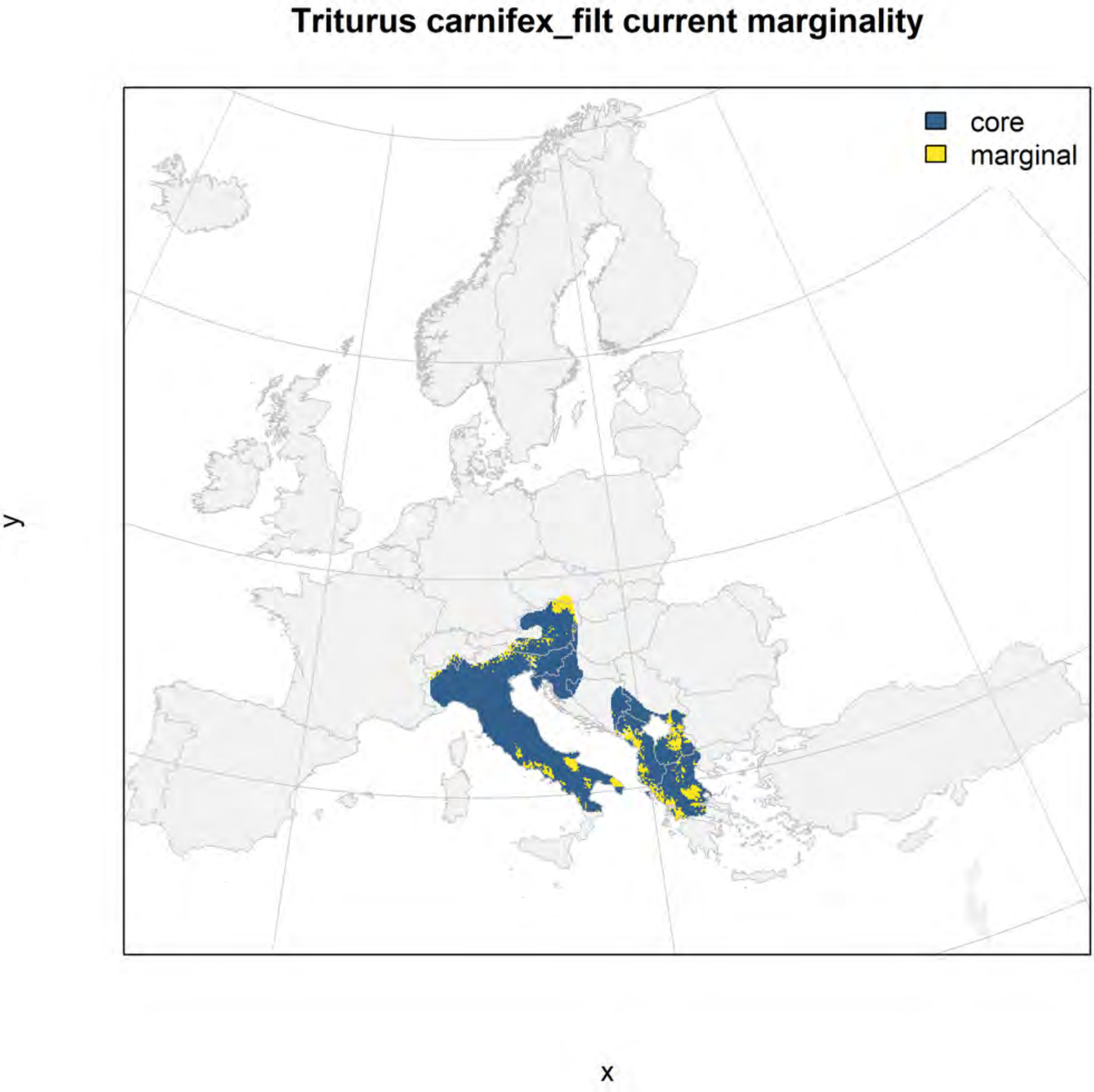
Triturus carnifex_filt current marginality
core
marginal
y
x

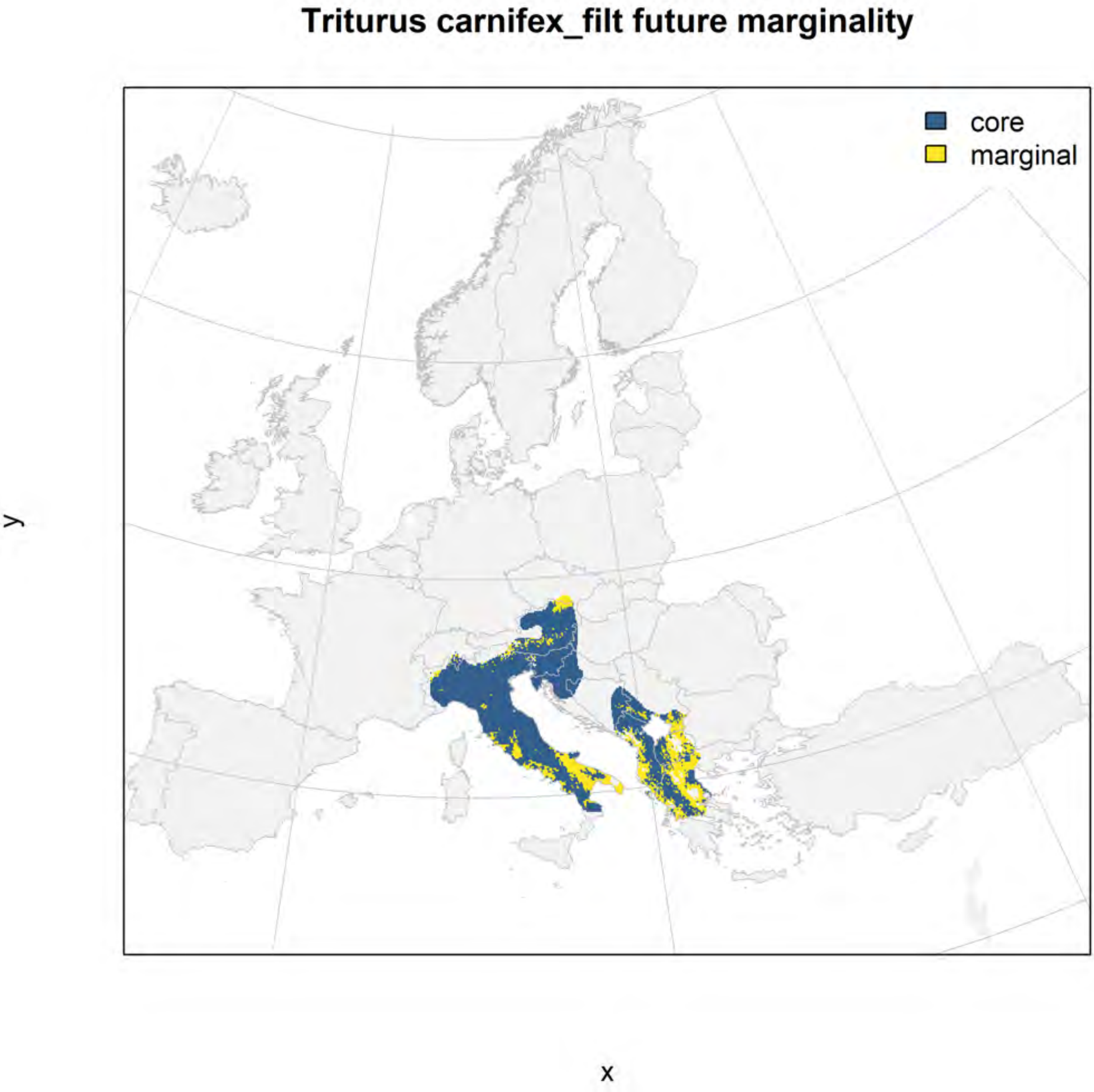

Triturus carnifex_filt future marginality
core
marginal
y
x

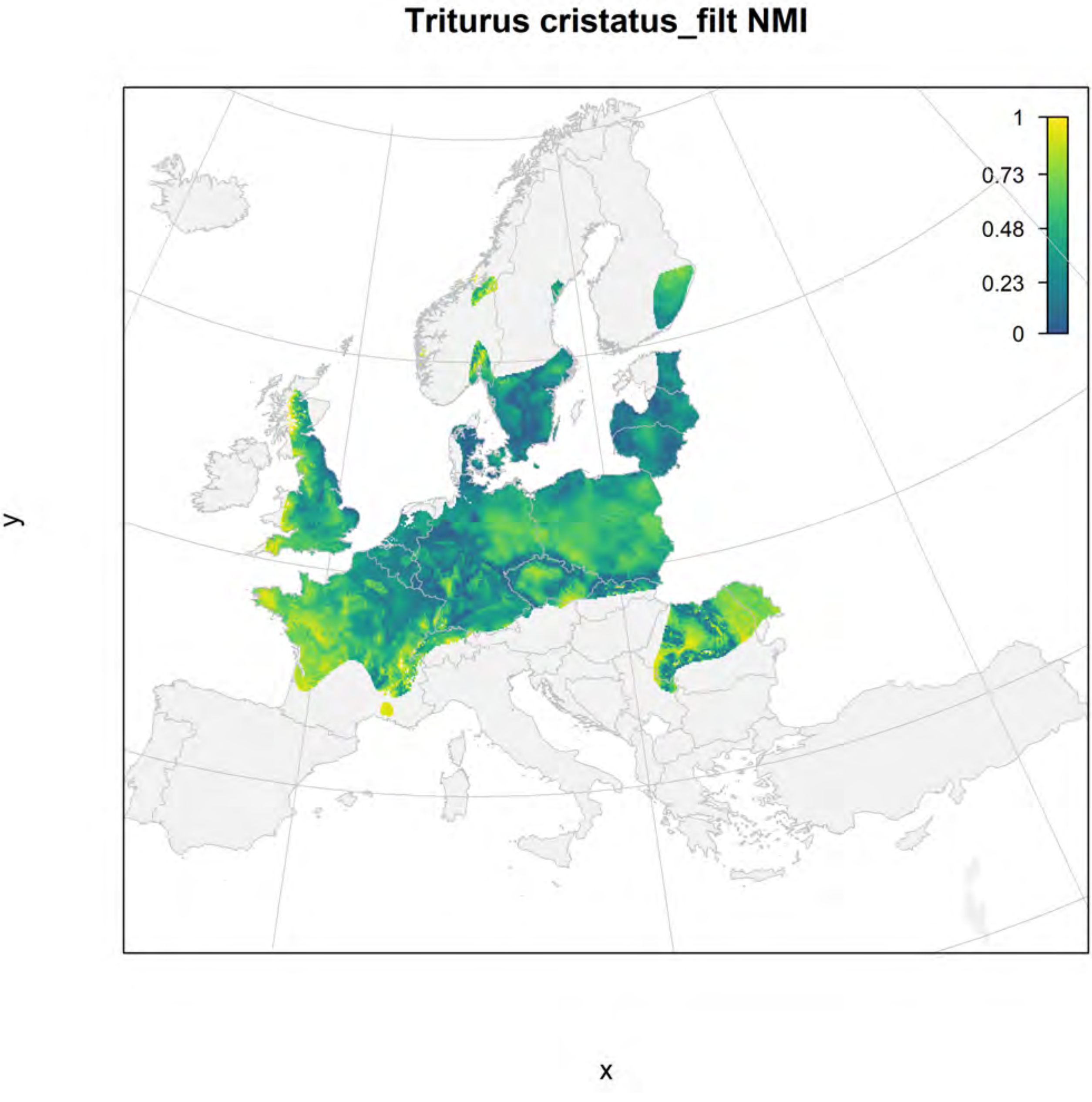
Triturus cristatus_filt NMI
1
0.73
0.48
0.23
0
y
x

#### Triturus cristatus_filt current marginality

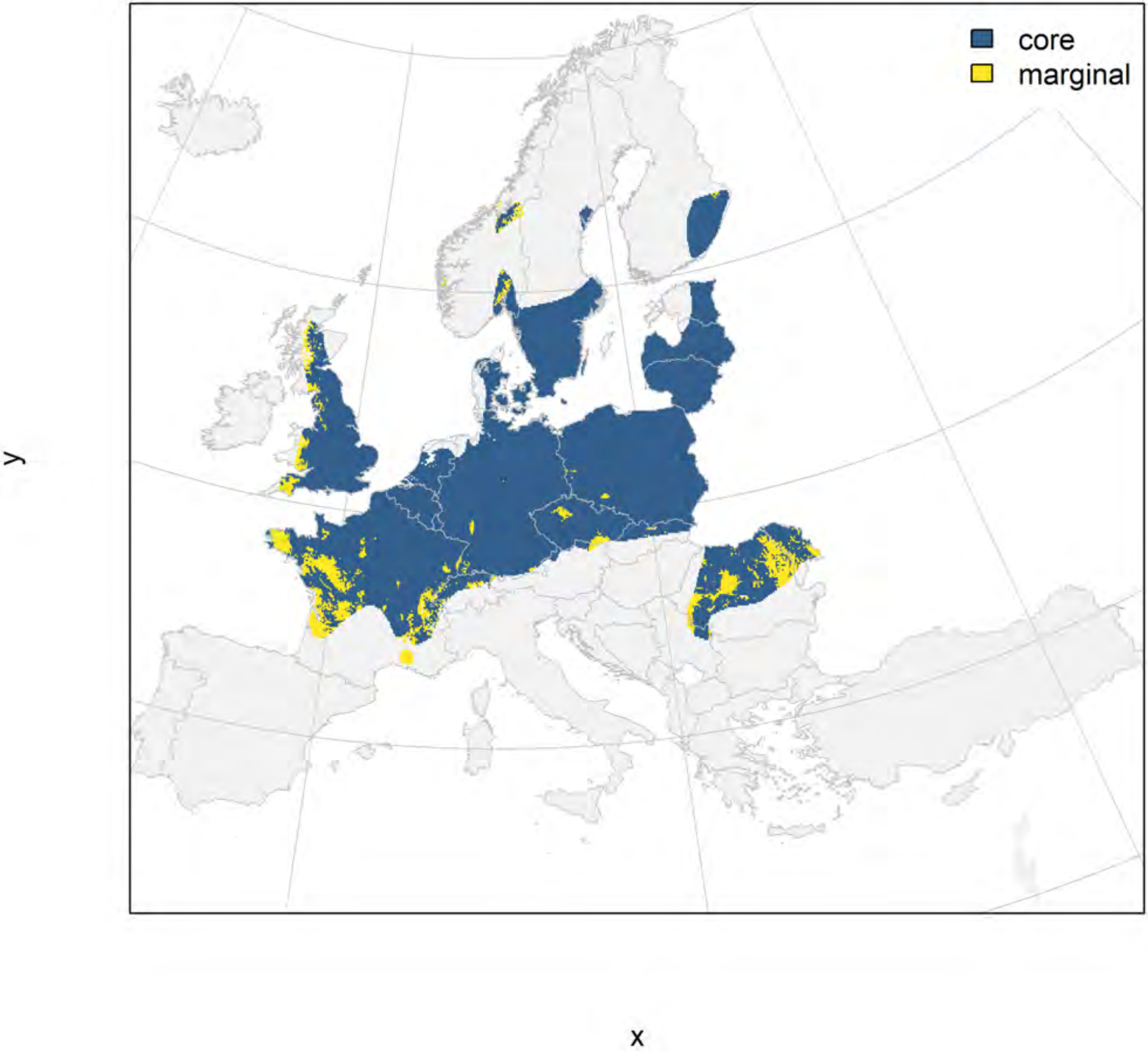

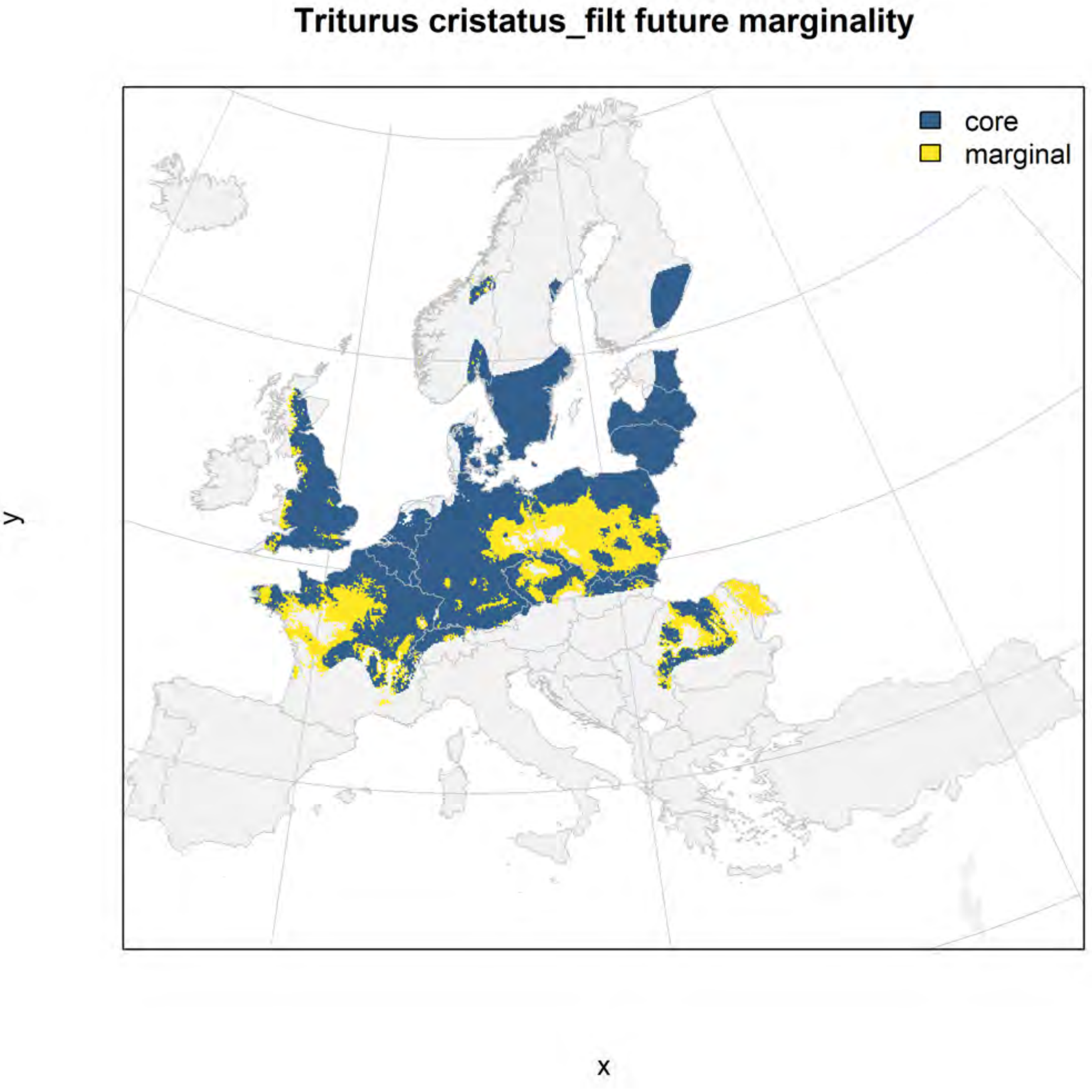
Triturus cristatus_filt future marginality
core
marginal
y
x

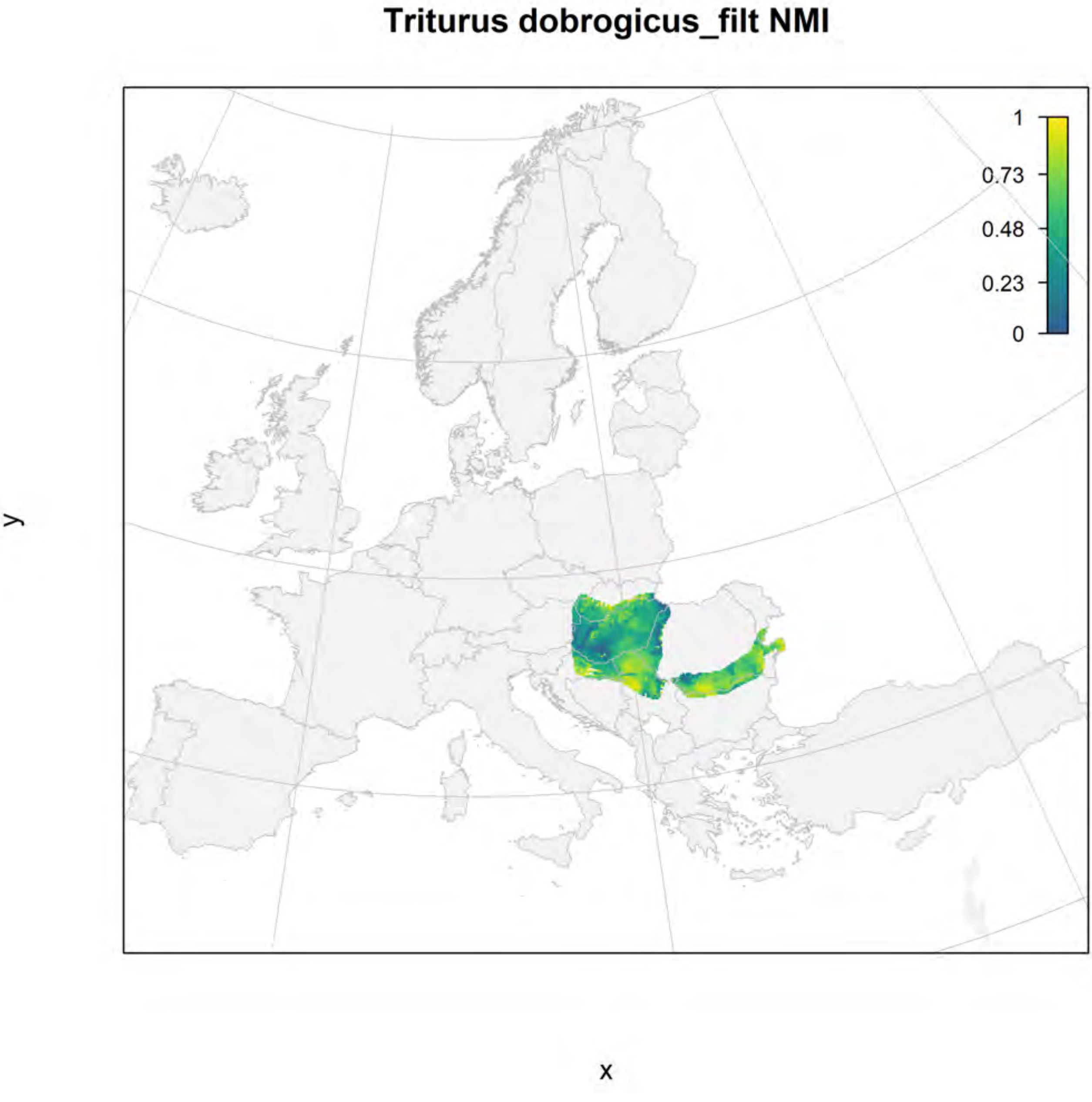
Triturus dobrogicus_filt NMI
1
0.73
0.48
0.23
0
y
x

#### Triturus dobrogicus_filt current marginality

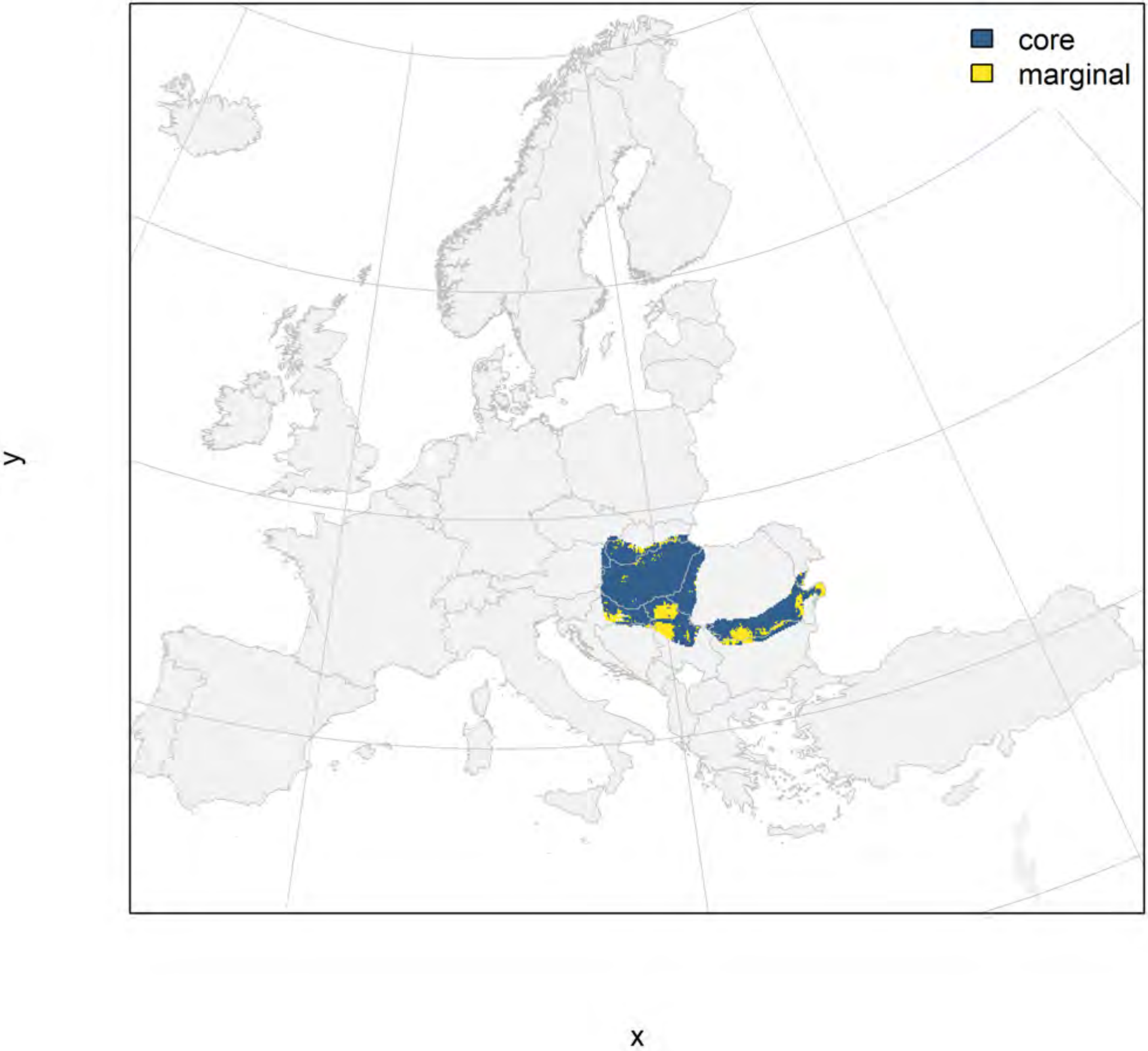

#### Triturus dobrogicus_filt future marginality

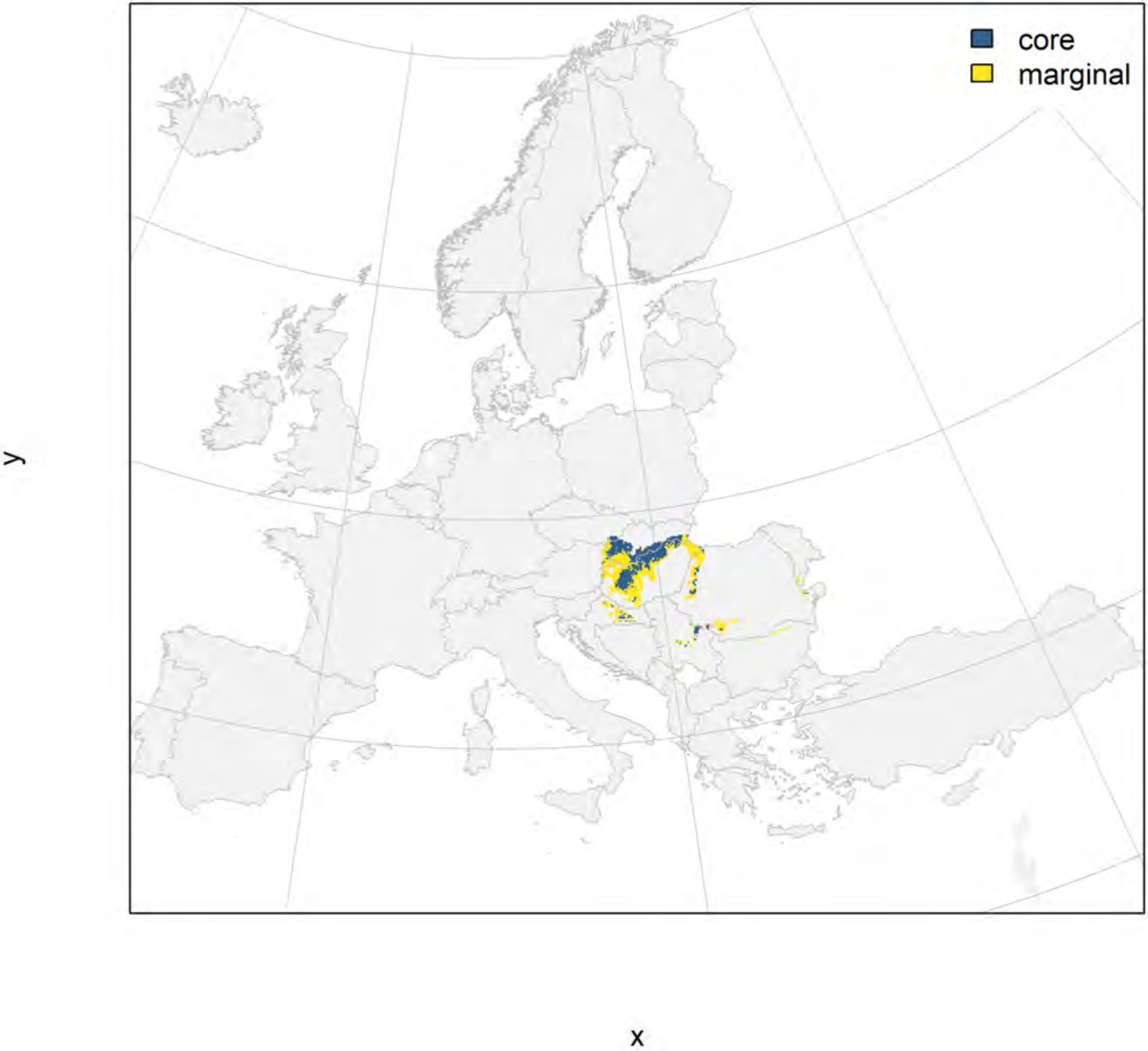

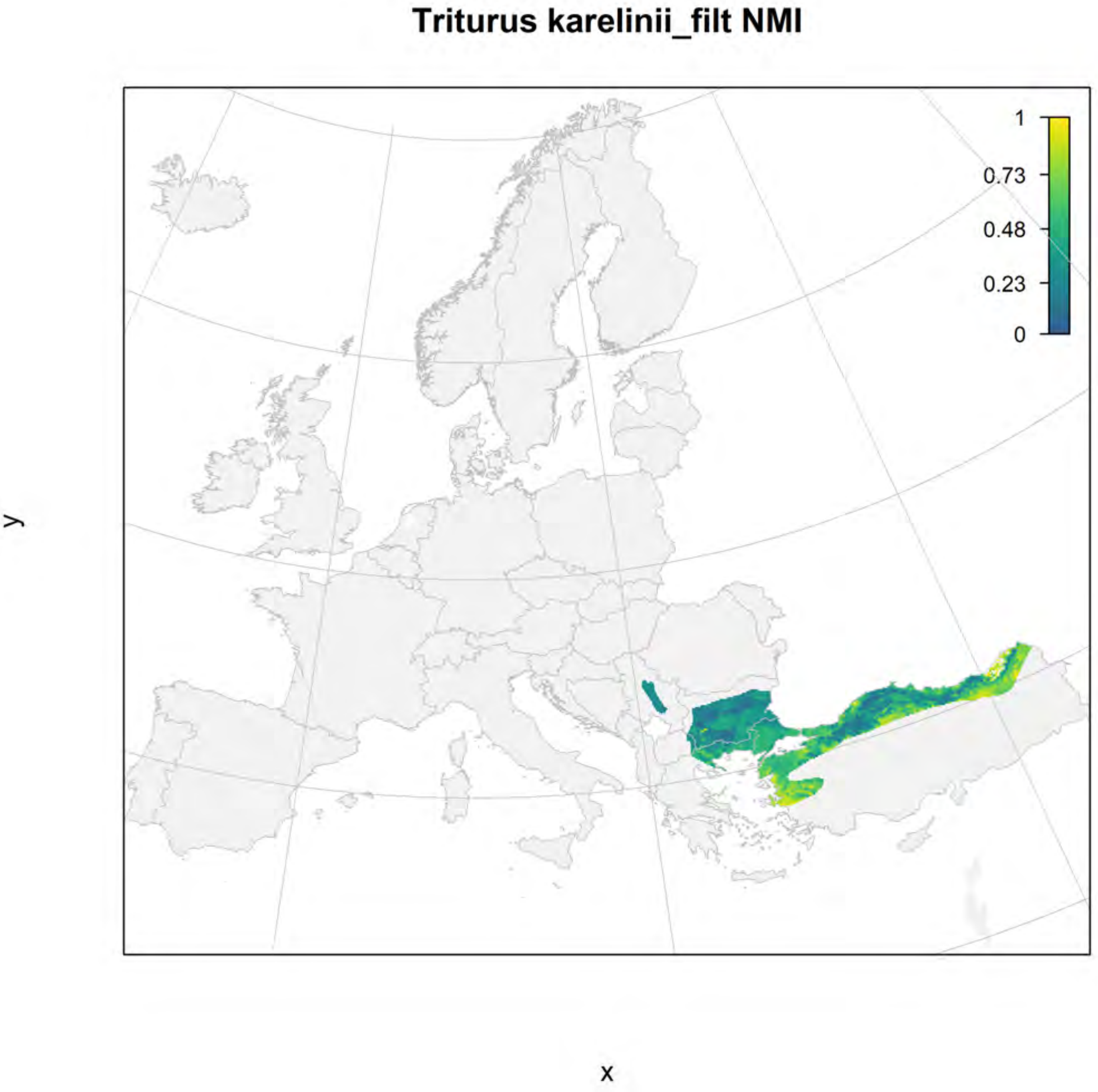
Triturus karelinii_filt NMI
1
0.73
0.48
0.23
0
y
x

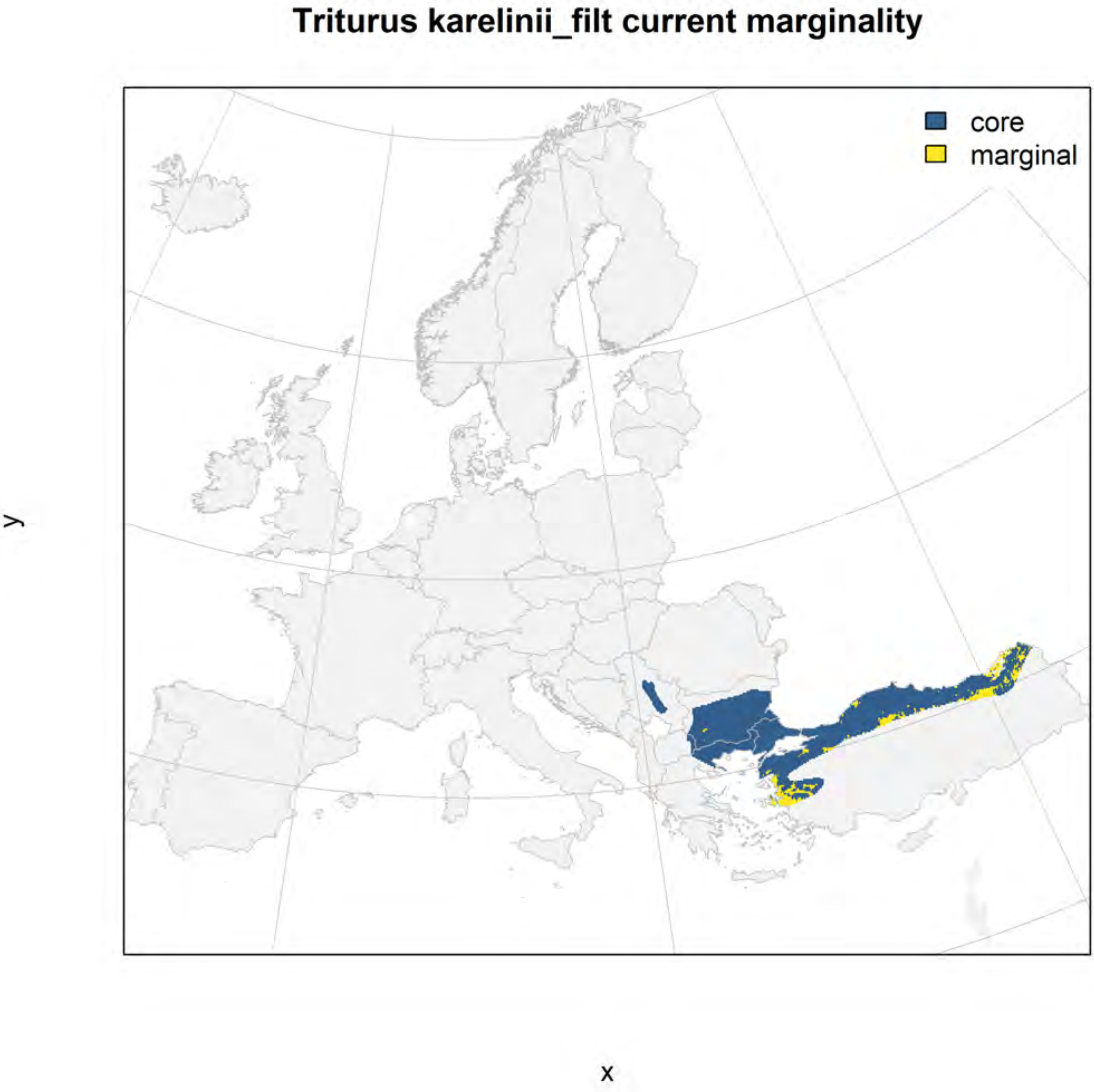
Triturus karelinii_filt current marginality
core
marginal
y
x

#### Triturus karelinii_filt future marginality

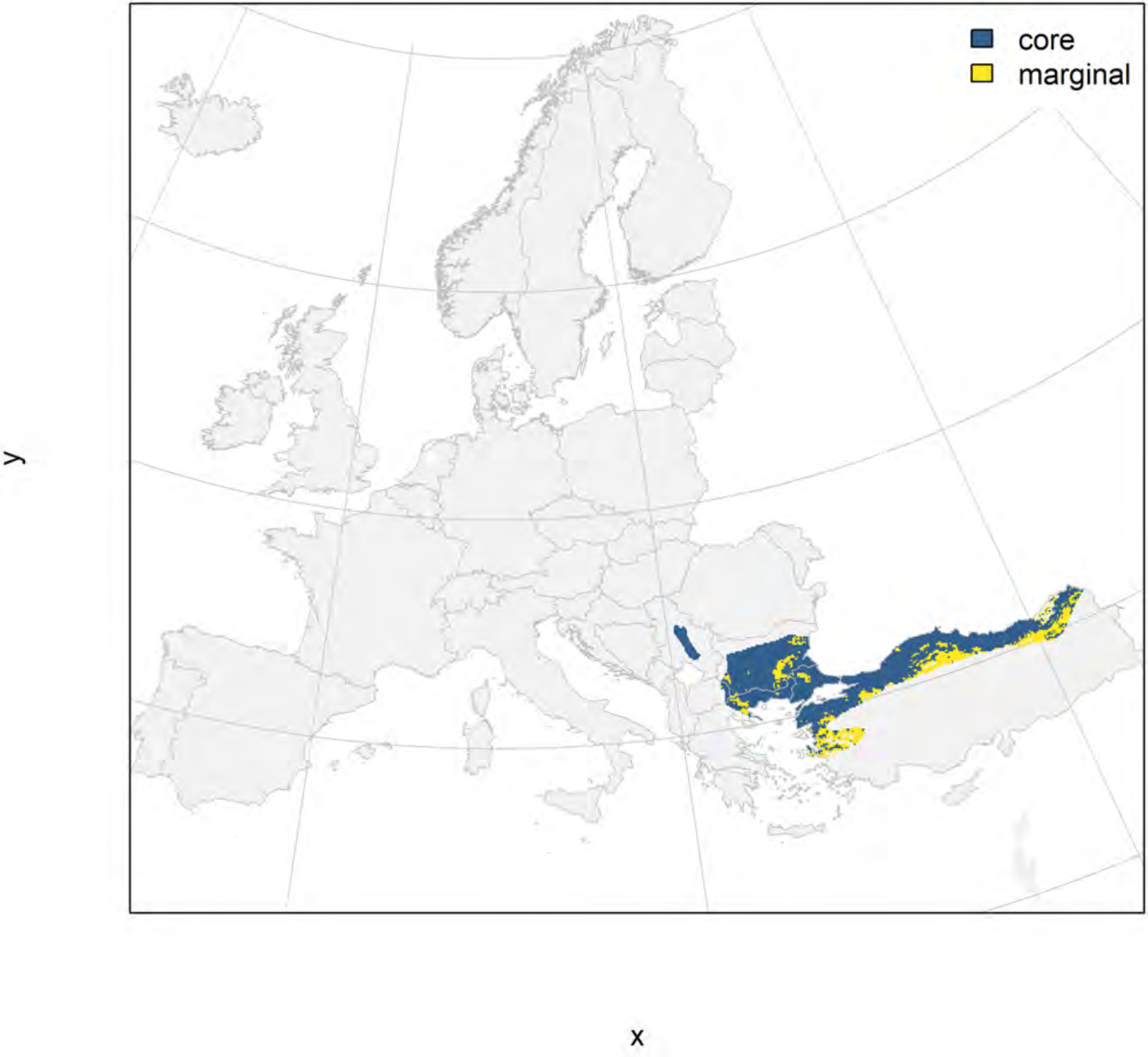

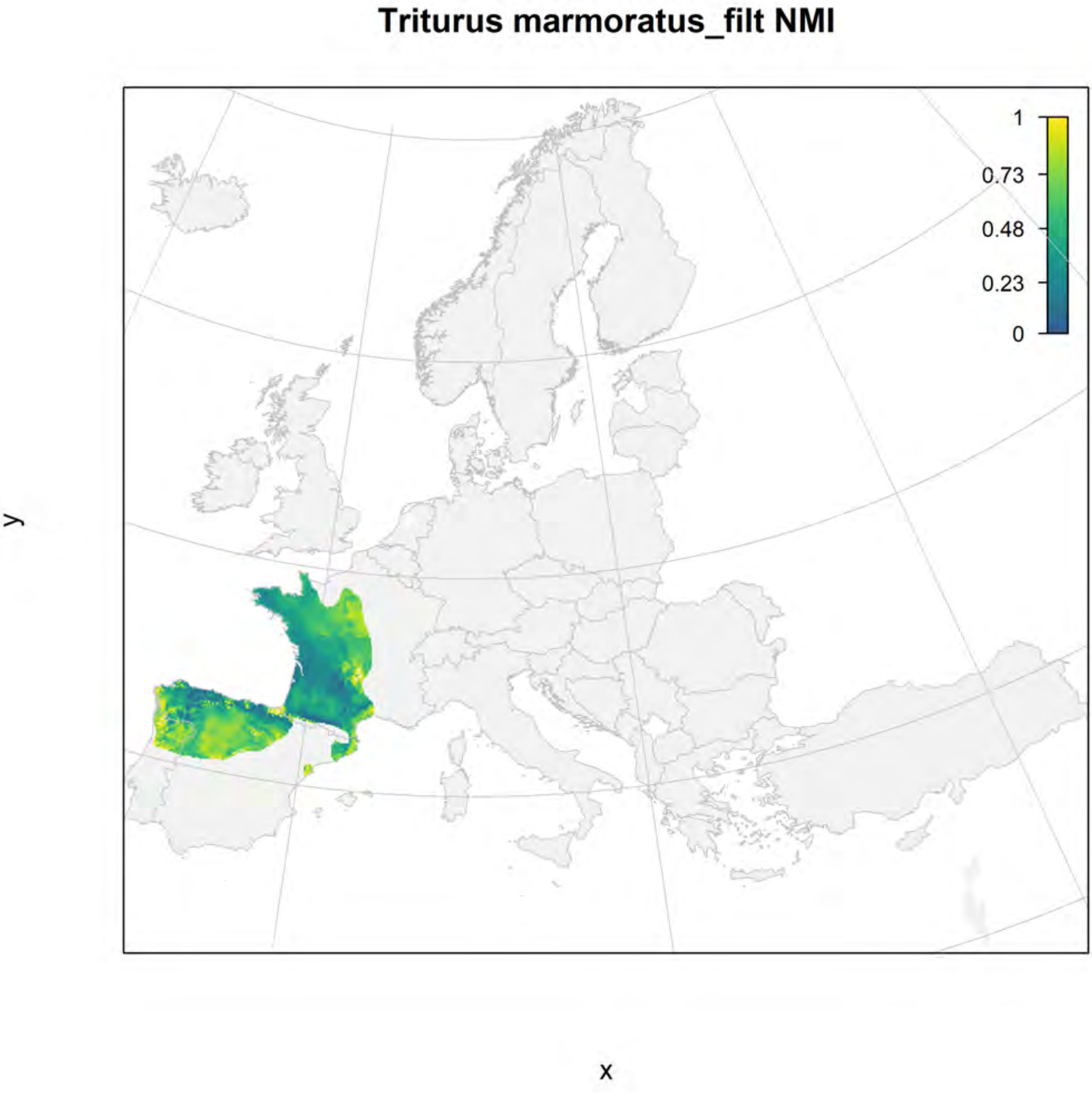
Triturus marmoratus_filt NMI
1
0.73
0.48
0.23
0
y
x

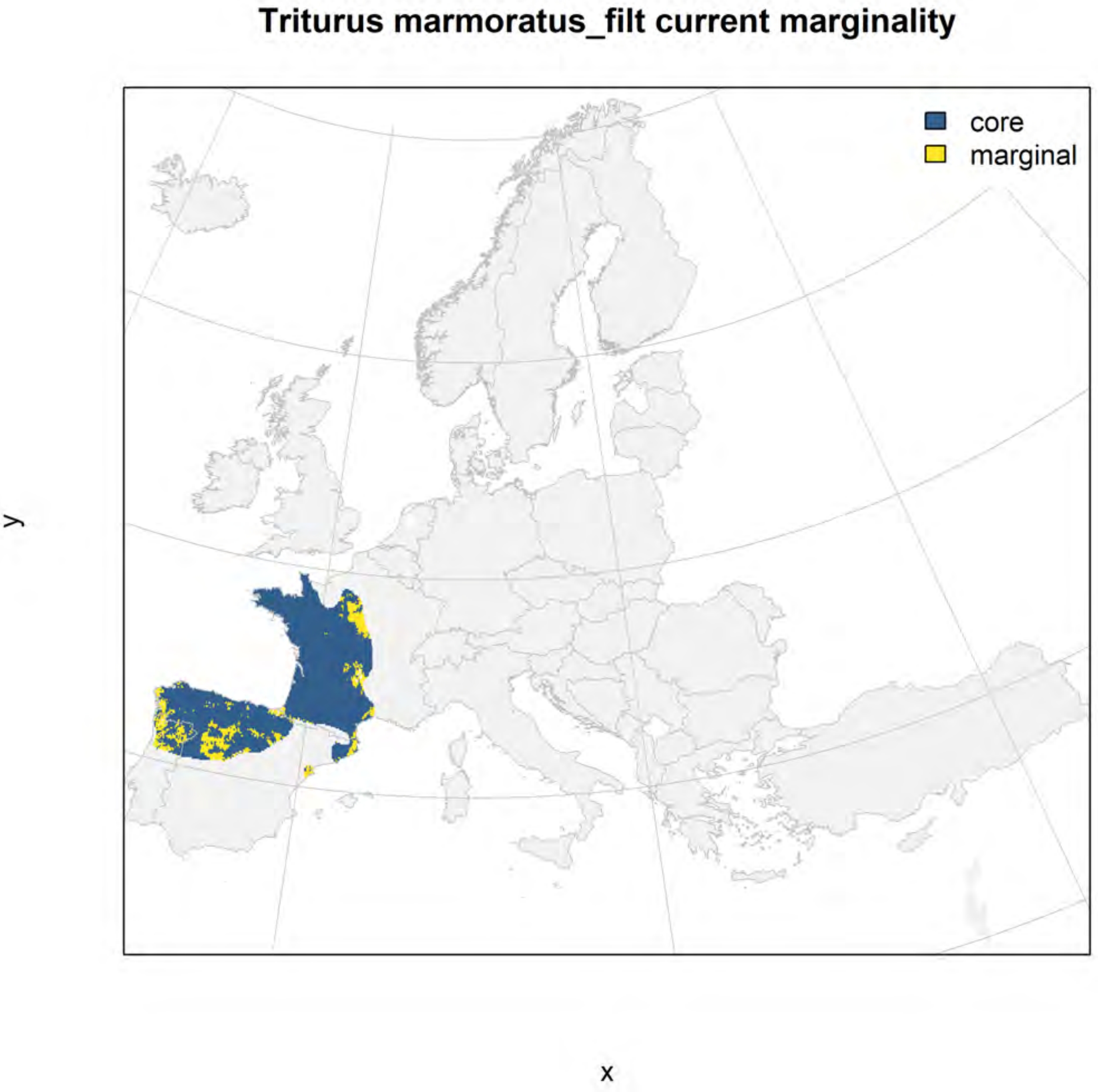
Triturus marmoratus_filt current marginality
core
marginal
y
x

### Triturus marmoratus_filt future marginality

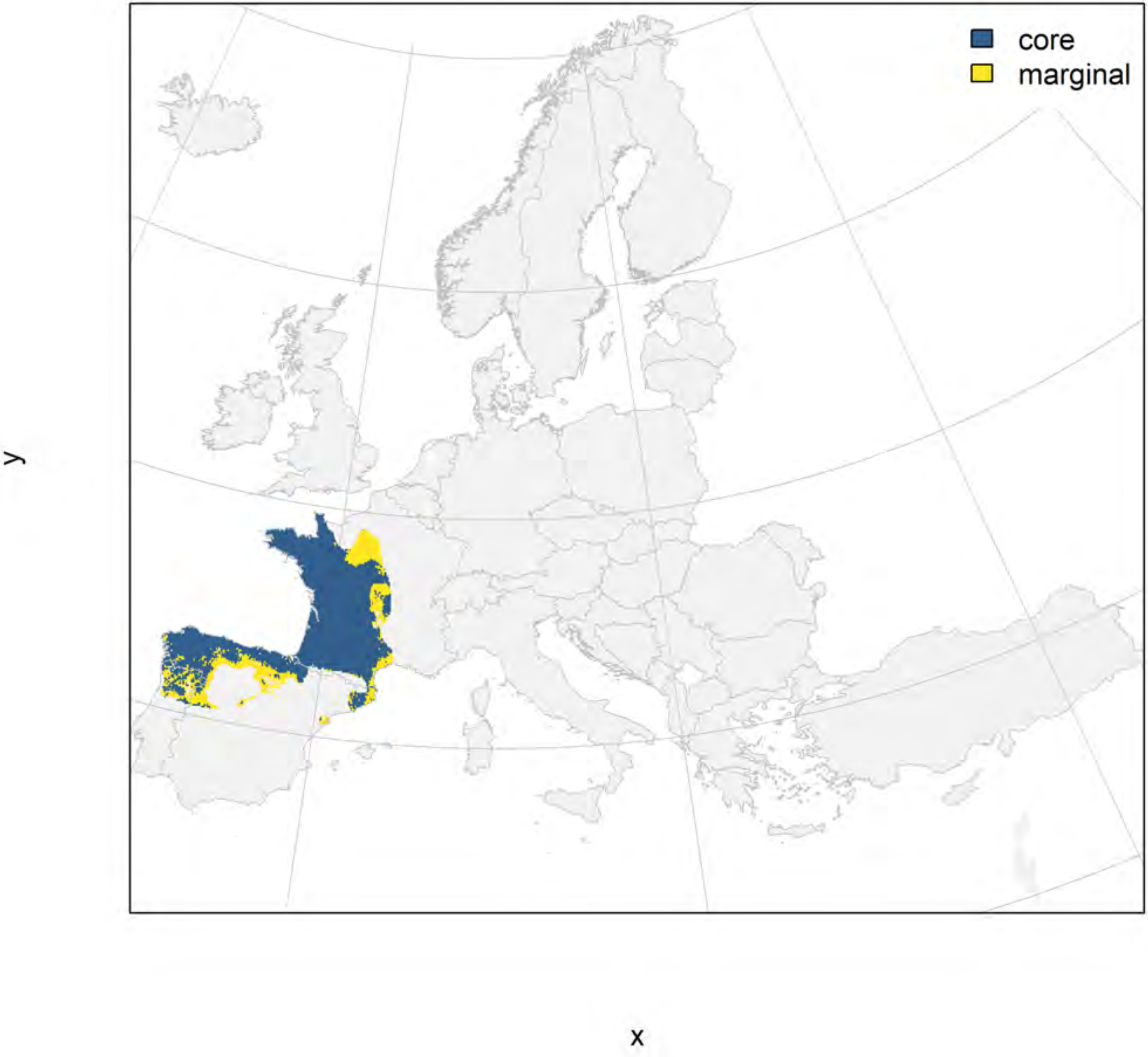

### Triturus pygmaeus_filt NMI

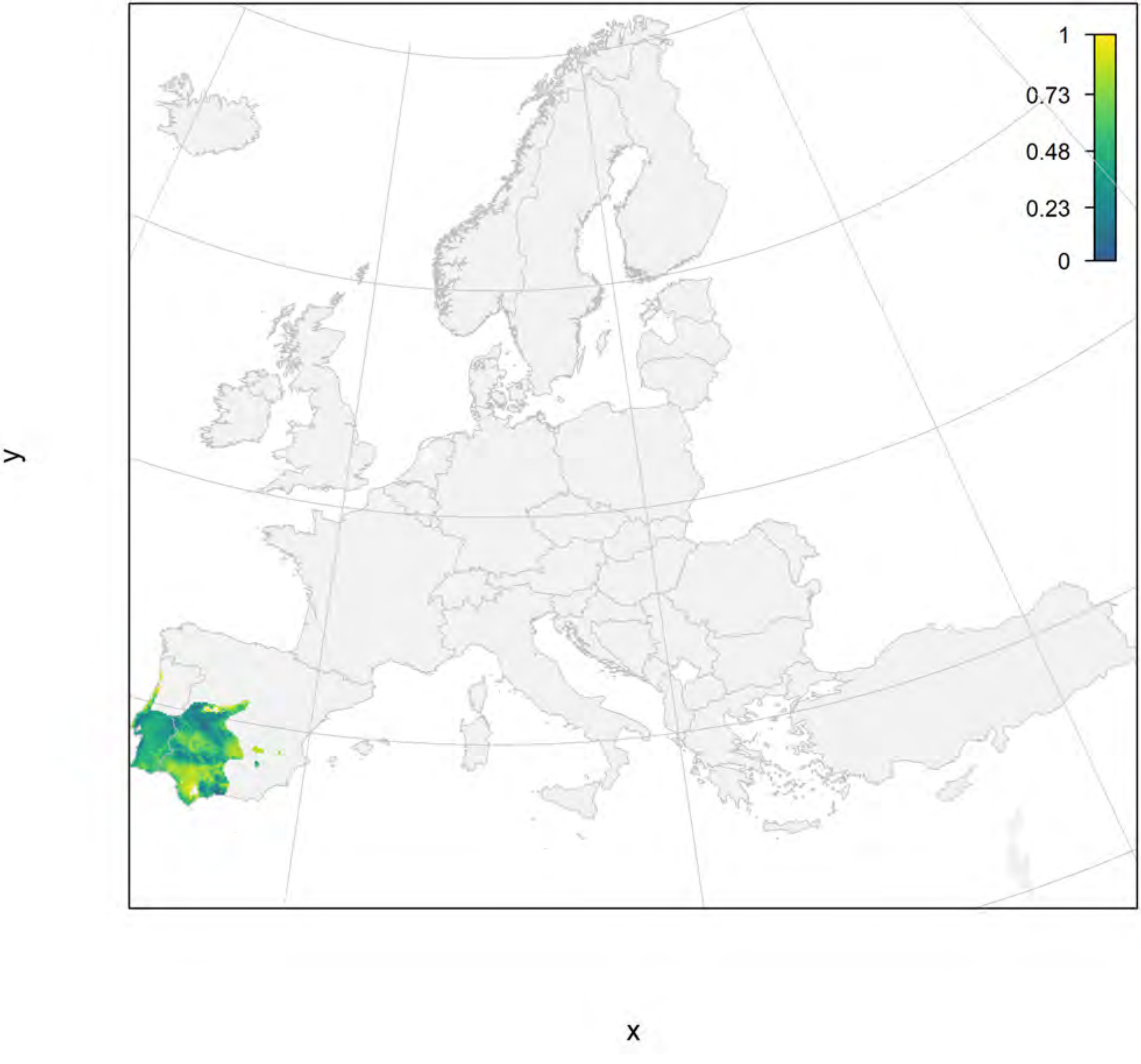

### Triturus pygmaeus_filt current marginality

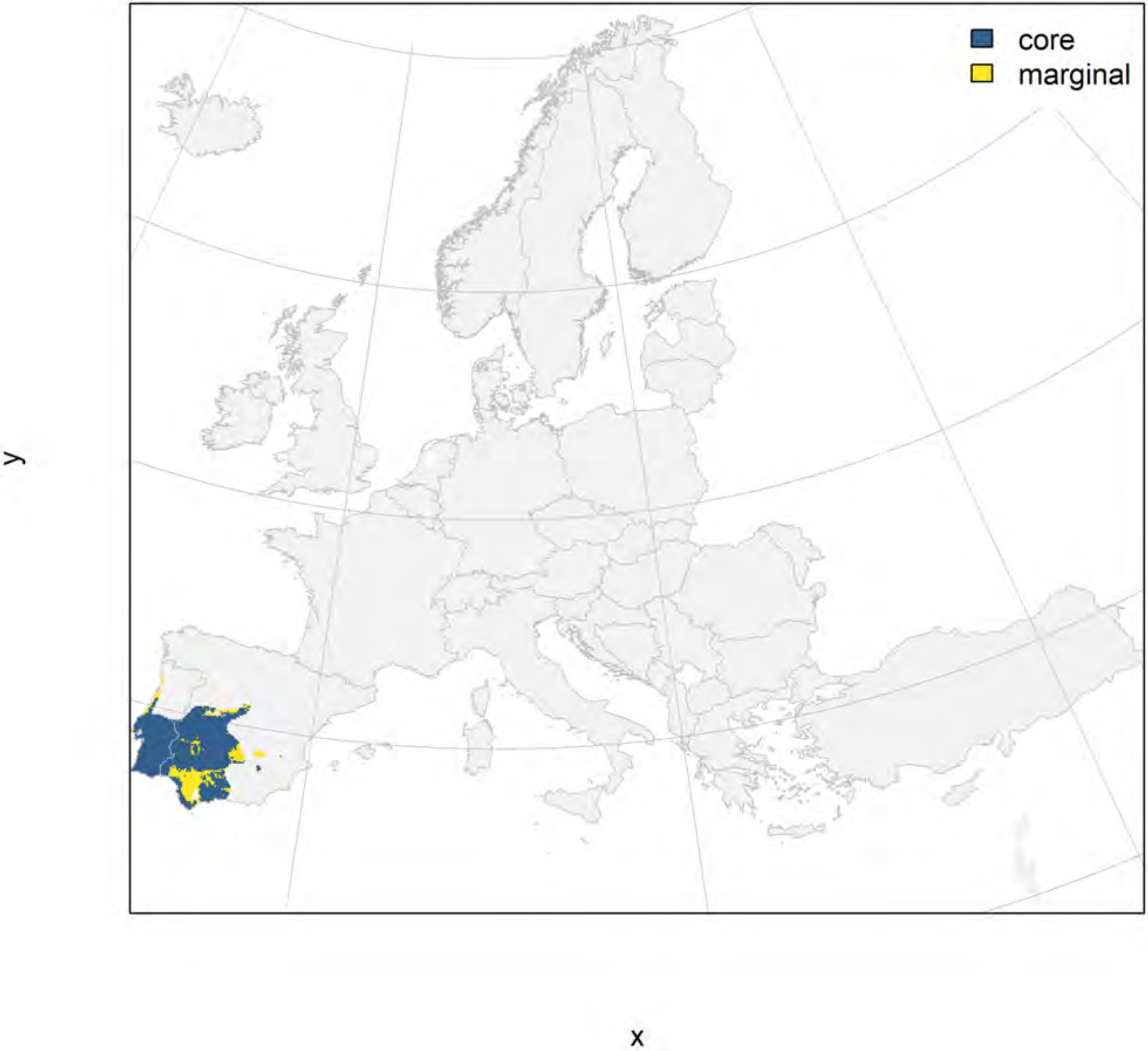

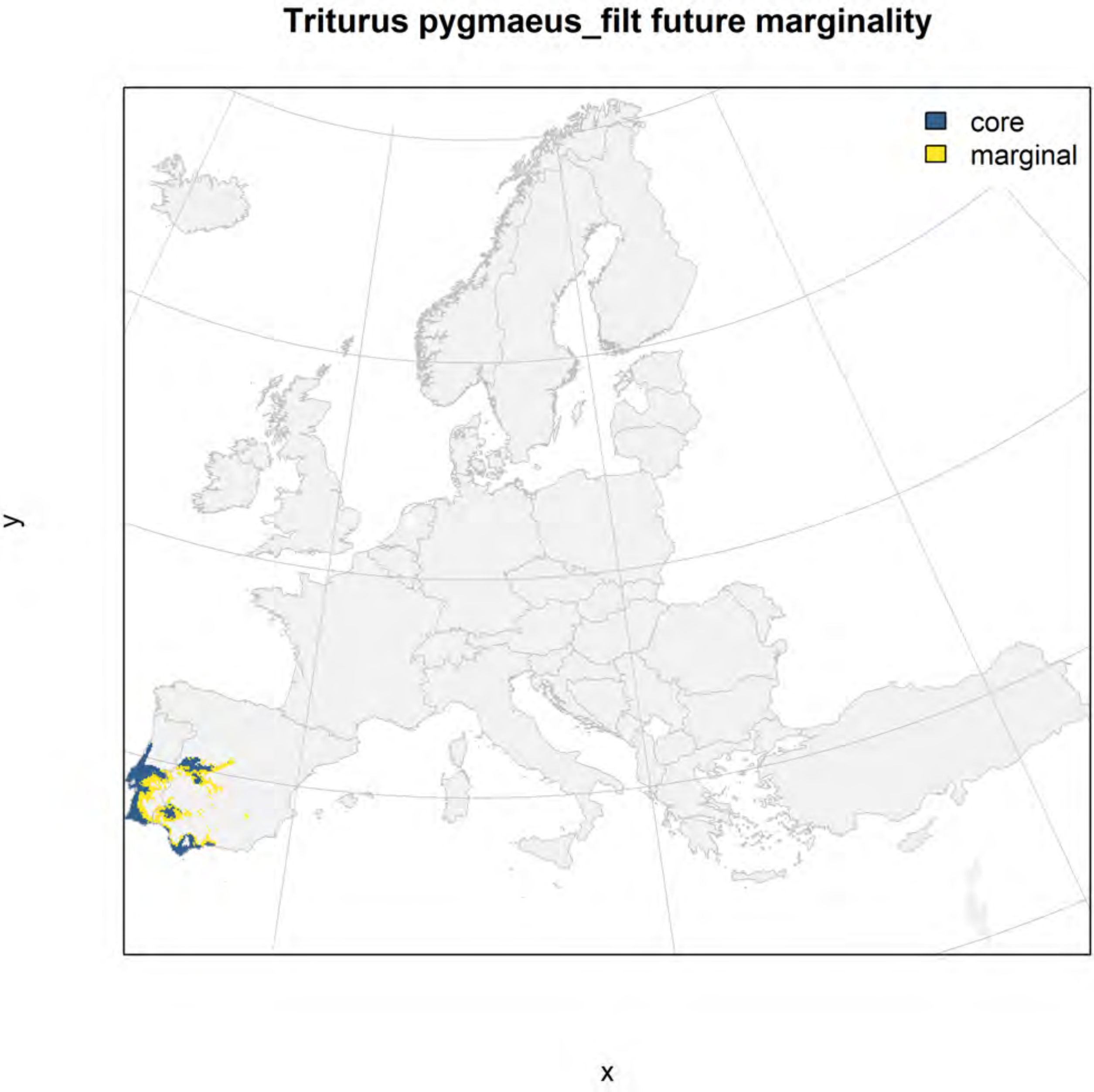
Triturus pygmaeus_filt future marginality
core
marginal
y
x
