## Appendix S4 for "Conserving genetic diversity during climate change: Niche marginality and discrepant monitoring capacity in Europe"

Appendix S4. Maps of marginality of selected species of large European birds. The climate marginality of each species is depicted in a series of three figures: raw marginality in which a value of 1 indicates the 99$^{th}$ percentile of the species niche space within its current range (filtered for appropriate habitat with a CORINE land cover layer), the current geographic distribution of areas within the 25% most marginal climatic niche conditions and the remaining core area, and a predicted future distribution of core and marginal areas within the current species range. Please see the Online Methods section for a description of marginality calculations.

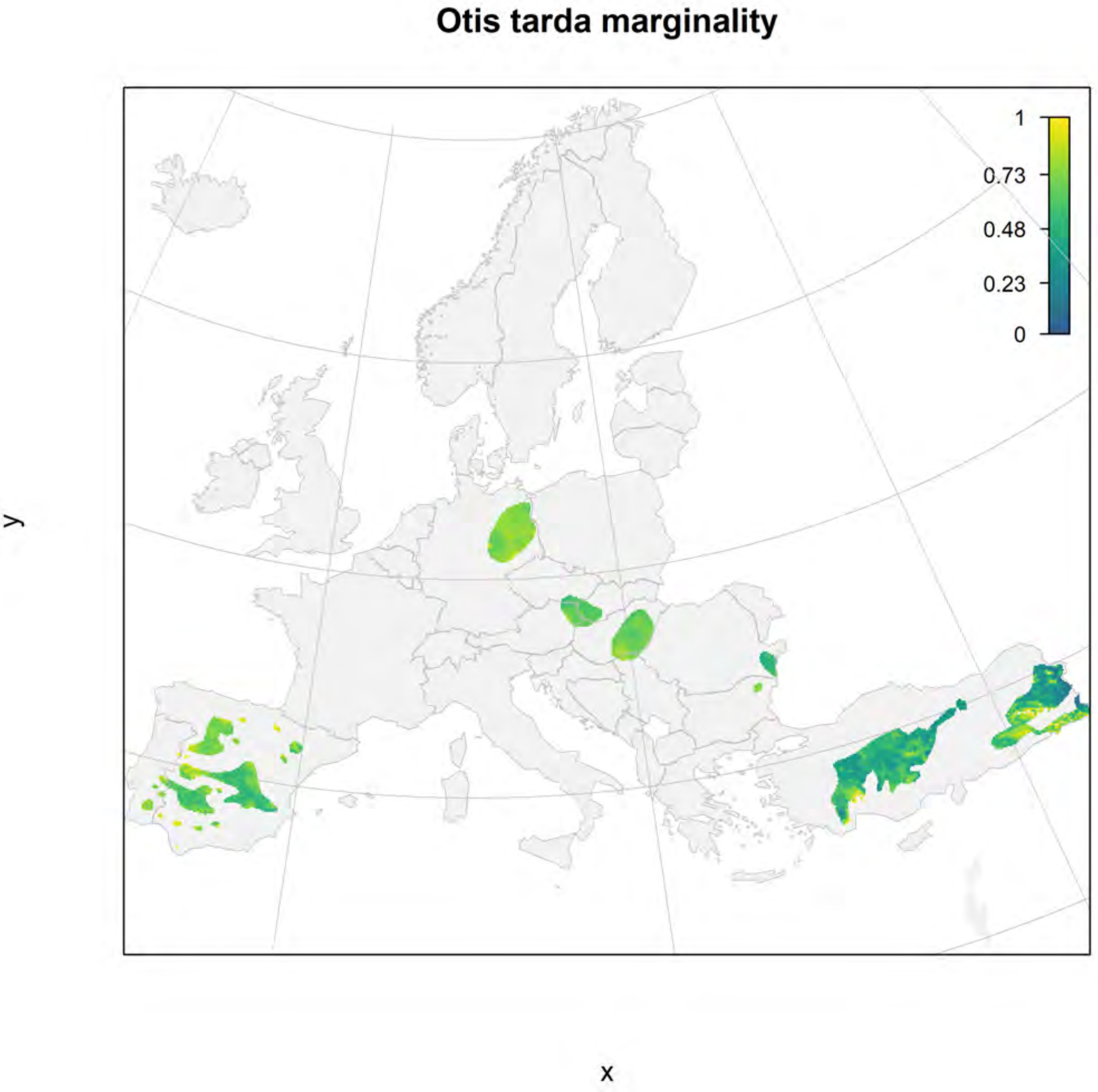
Otis tarda marginality
1
0.73
0.48
0.23
0
y
x

### Otis tarda_filt current marginality

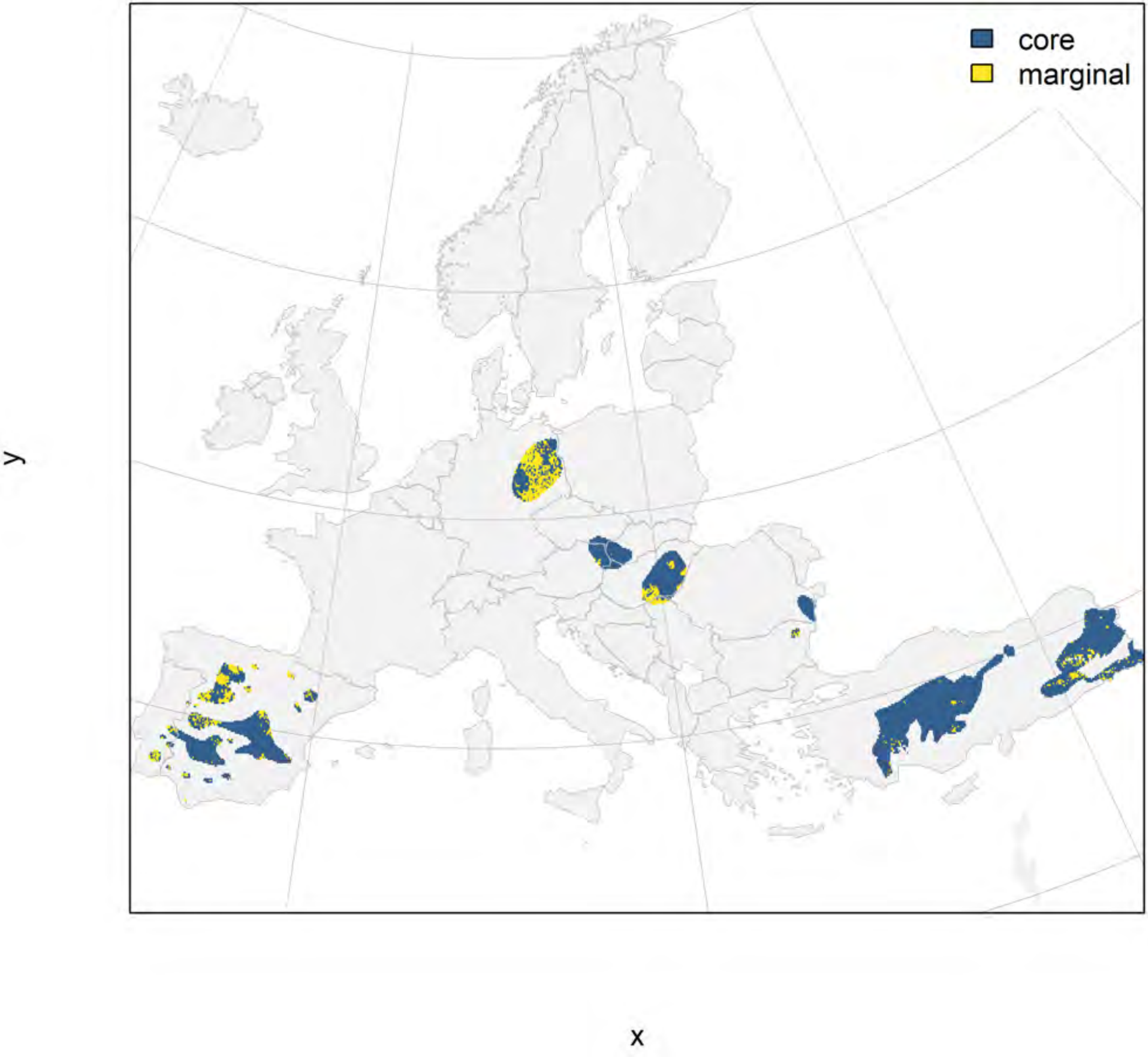

### Otis tarda_filt future marginality

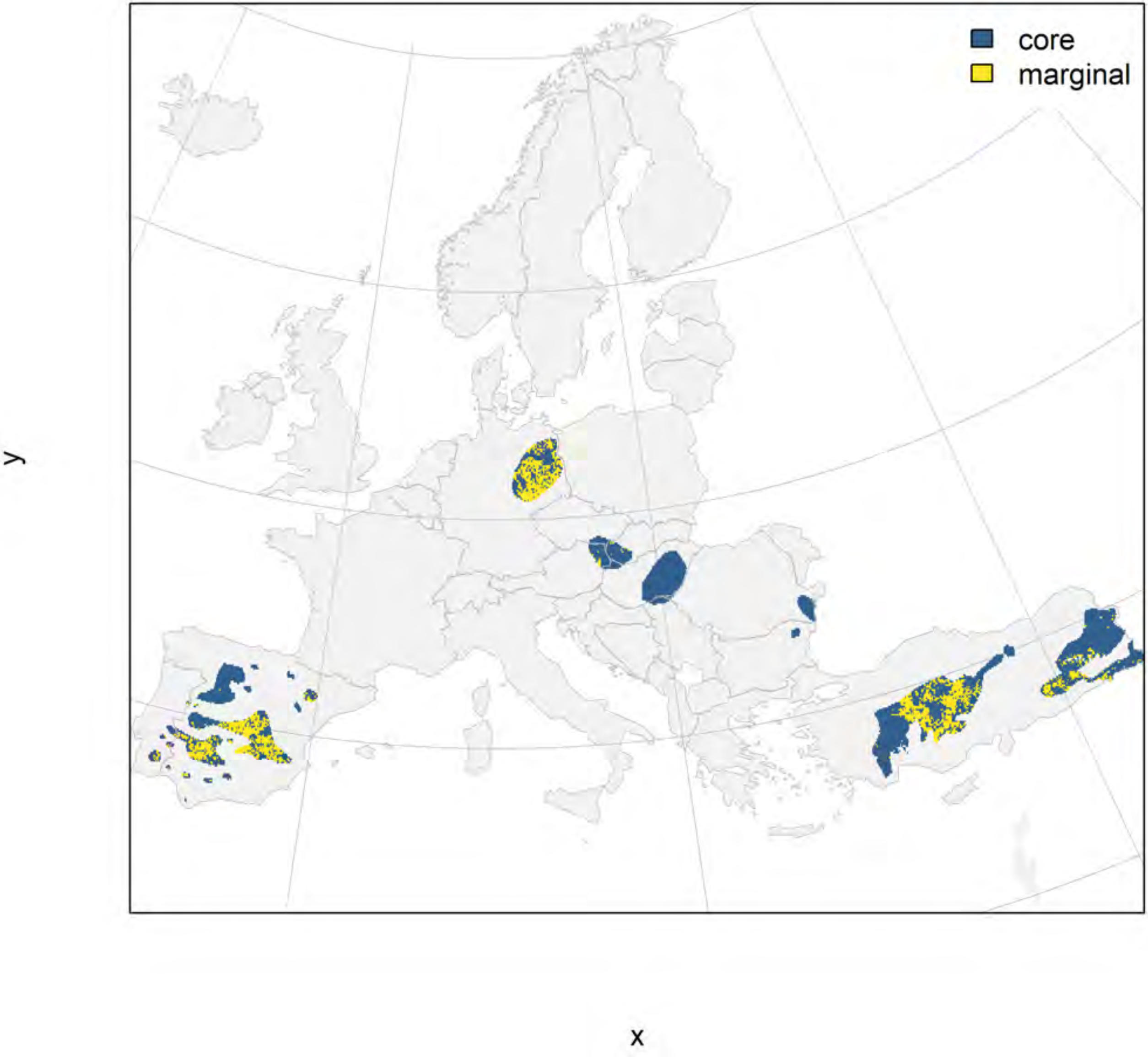

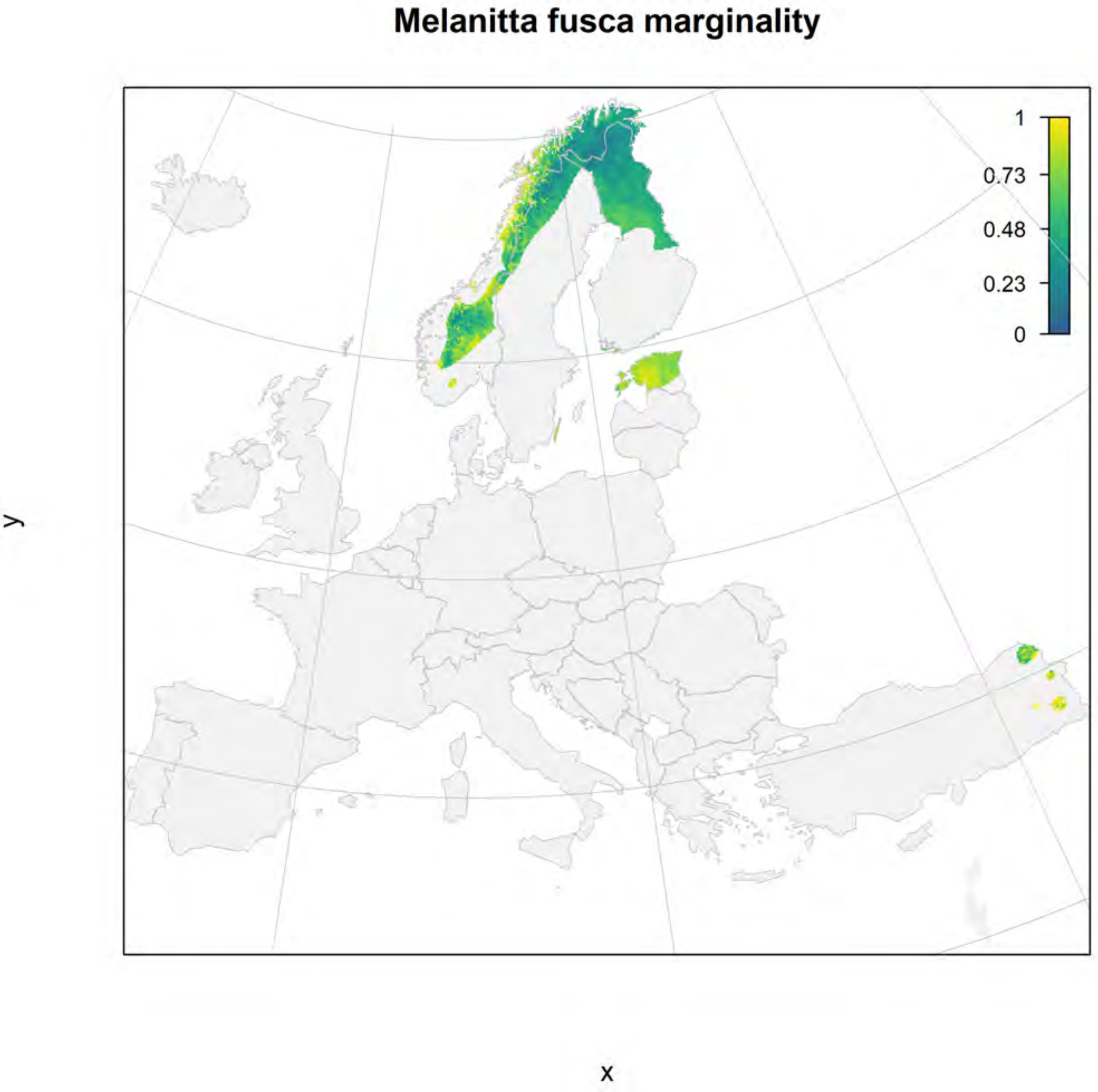
Melanitta fusca marginality
1
0.73
0.48
0.23
0
y
x

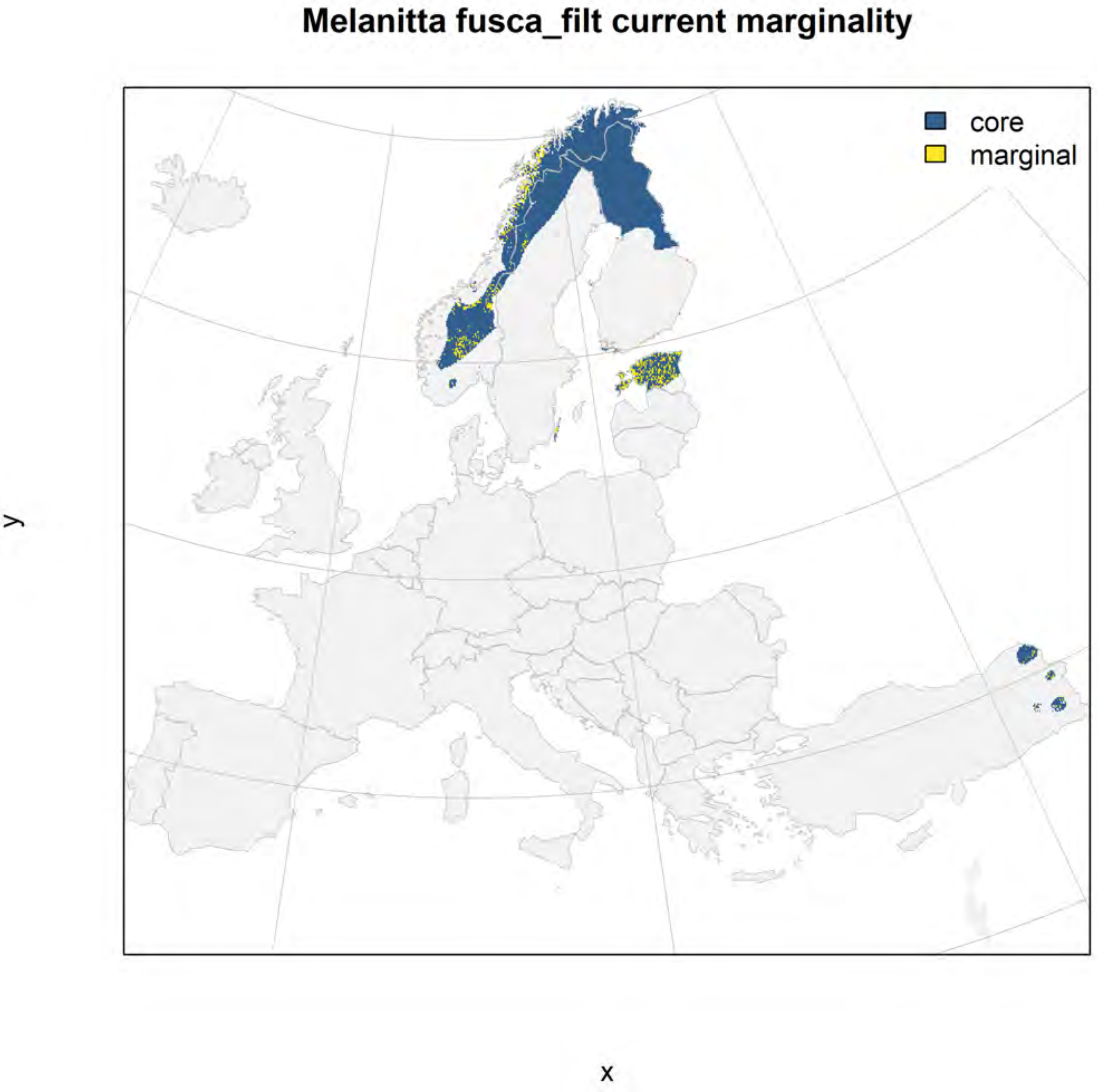

Melanitta fusca_filt current marginality
core
marginal
y
x

### Melanitta fusca_filt future marginality

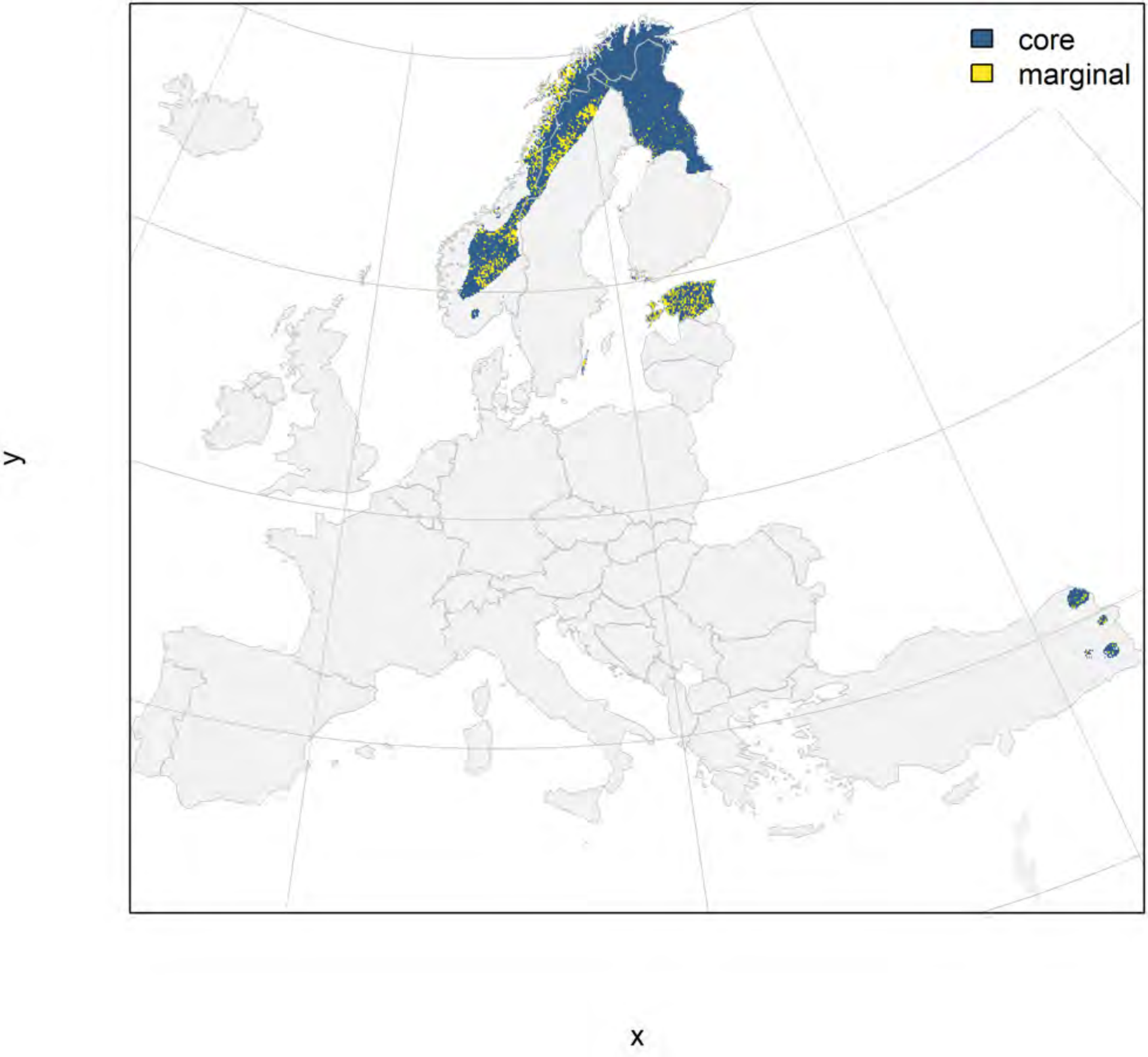

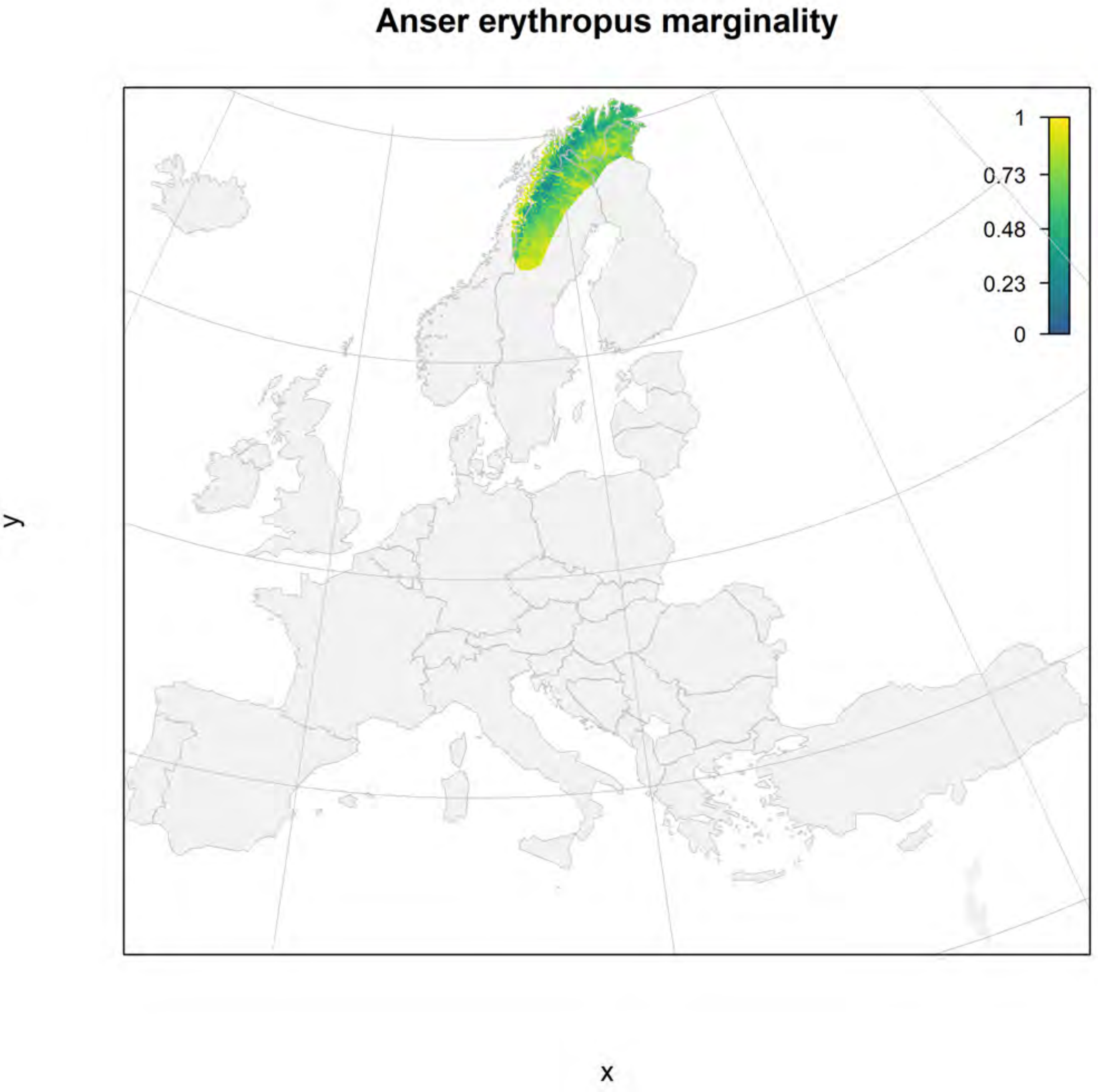
Anser erythropus marginality
1
0.73
0.48
0.23
0
y
x

### Anser erythropus_filt current marginality

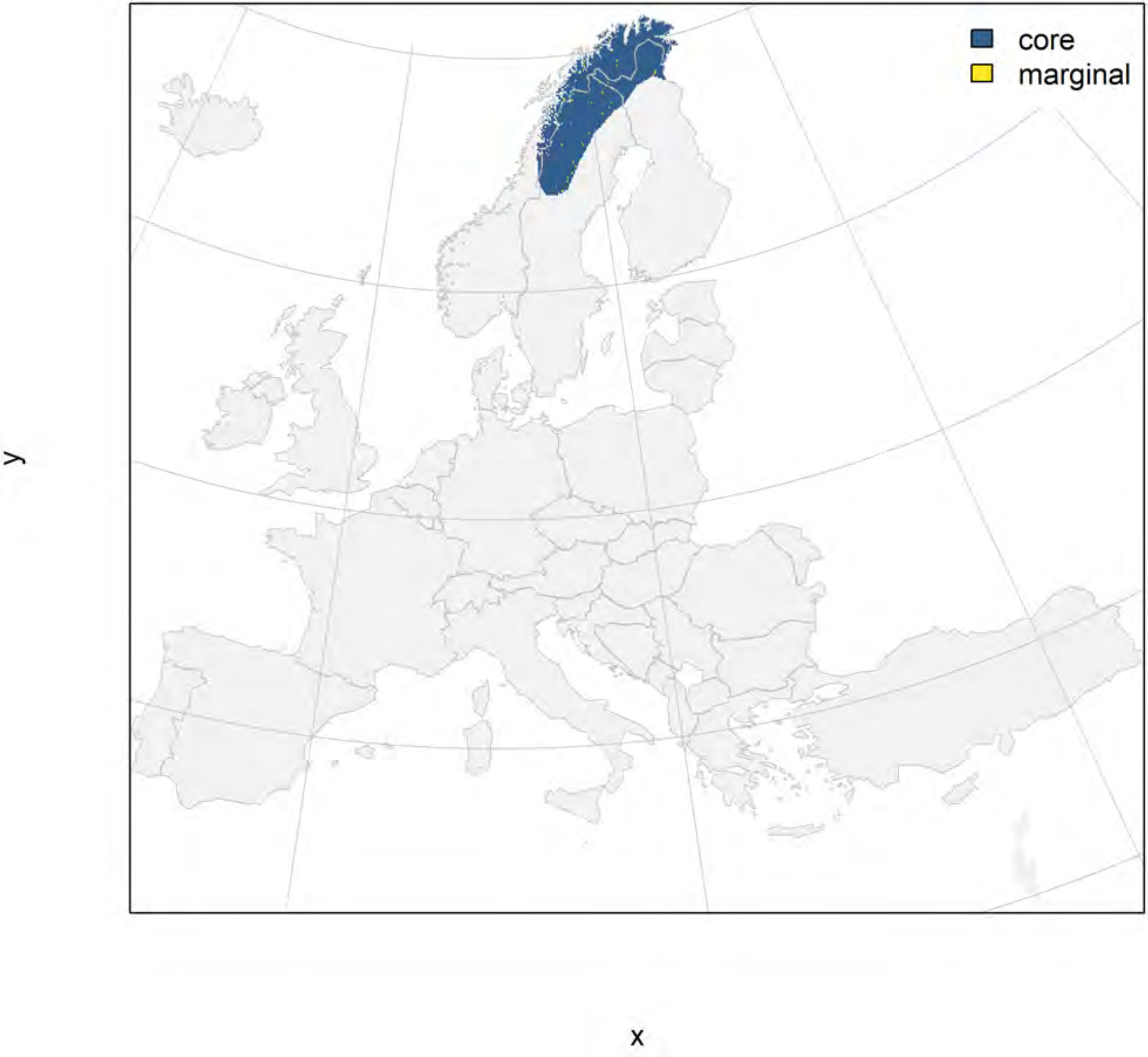

### Anser erythropus_filt future marginality

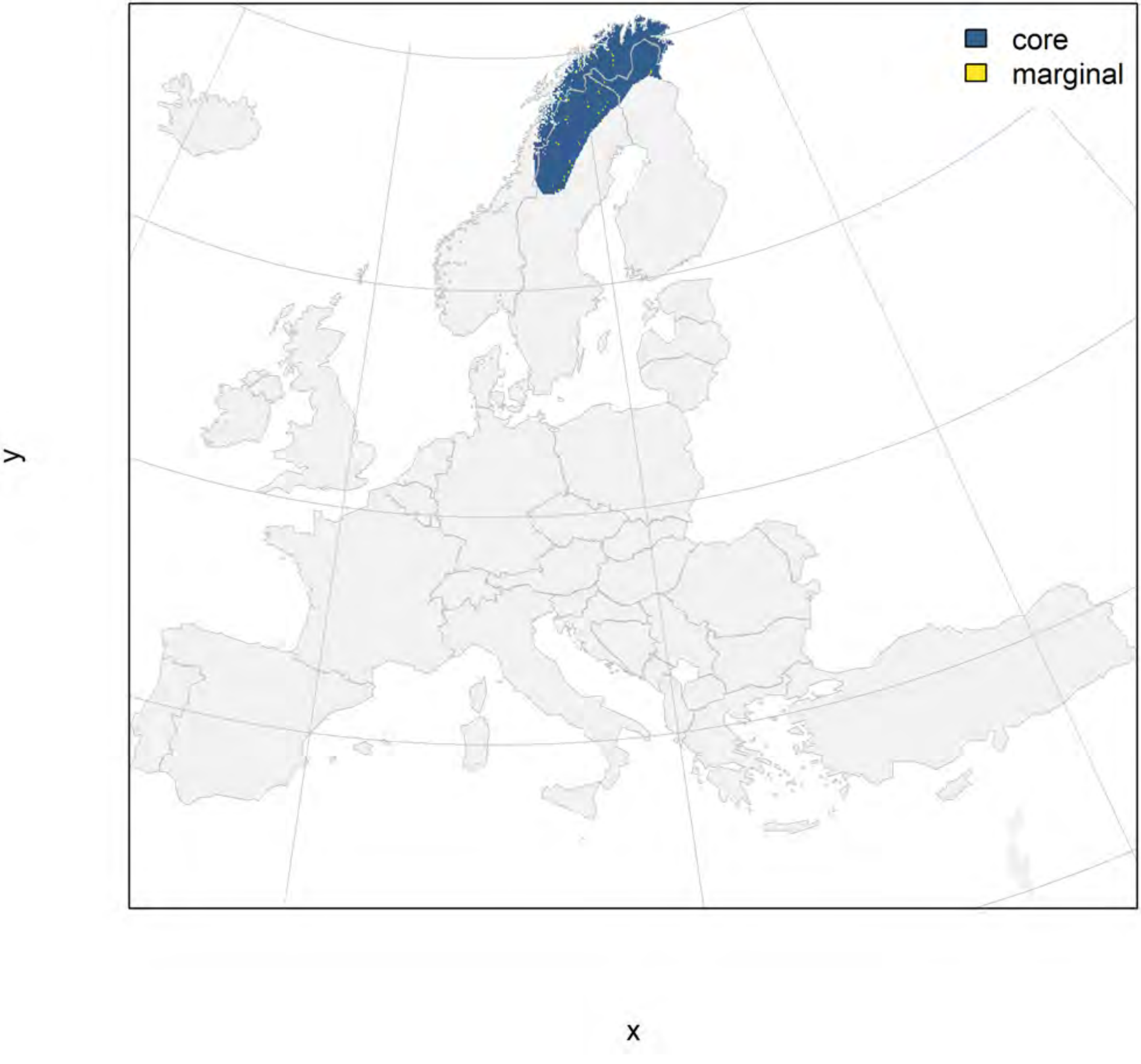

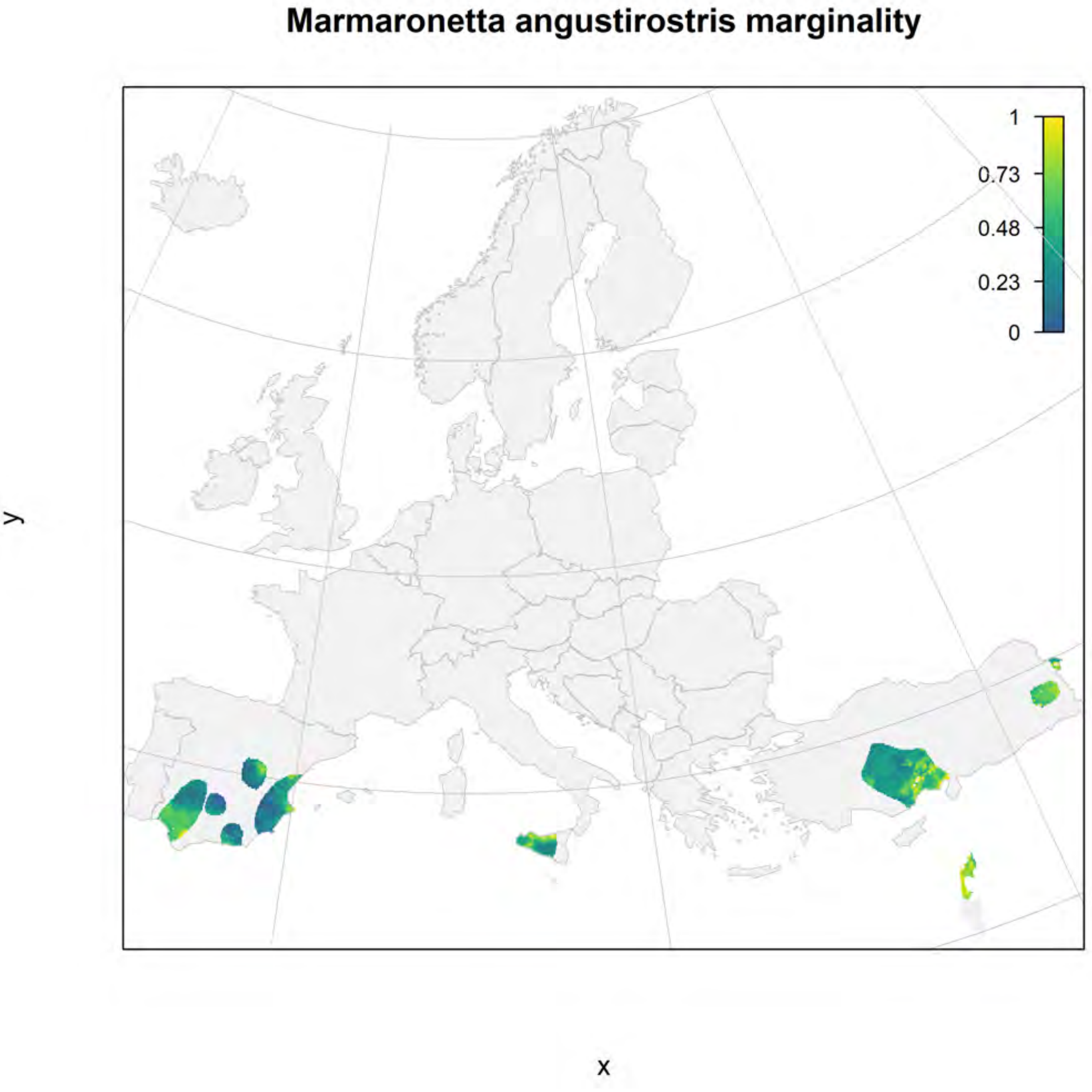

Marmaronetta angustirostris marginality
1
0.73
0.48
0.23
0
y
x

### Marmaronetta angustirostris_filt current marginality

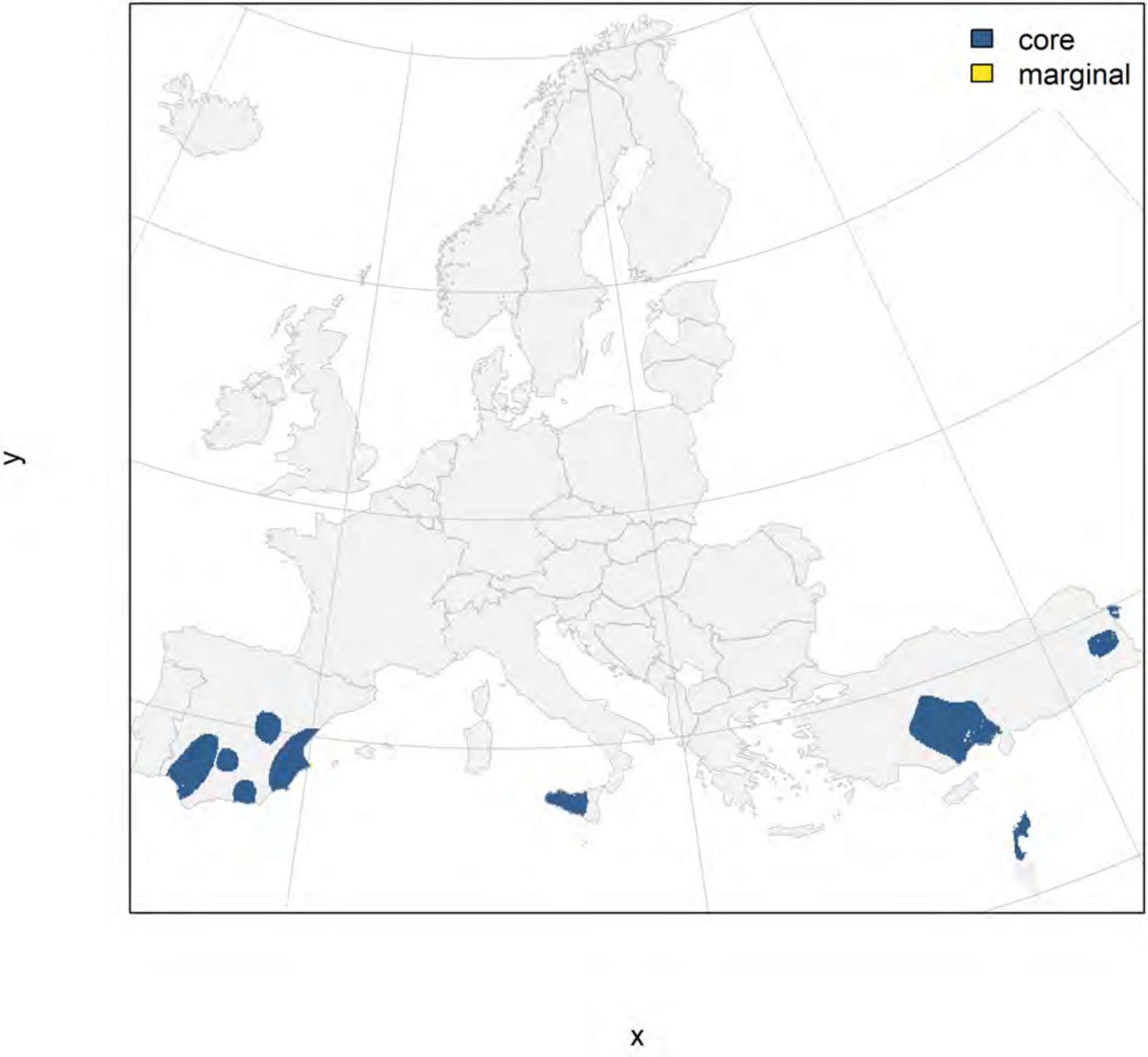

#### Marmaronetta angustirostris_filt future marginality

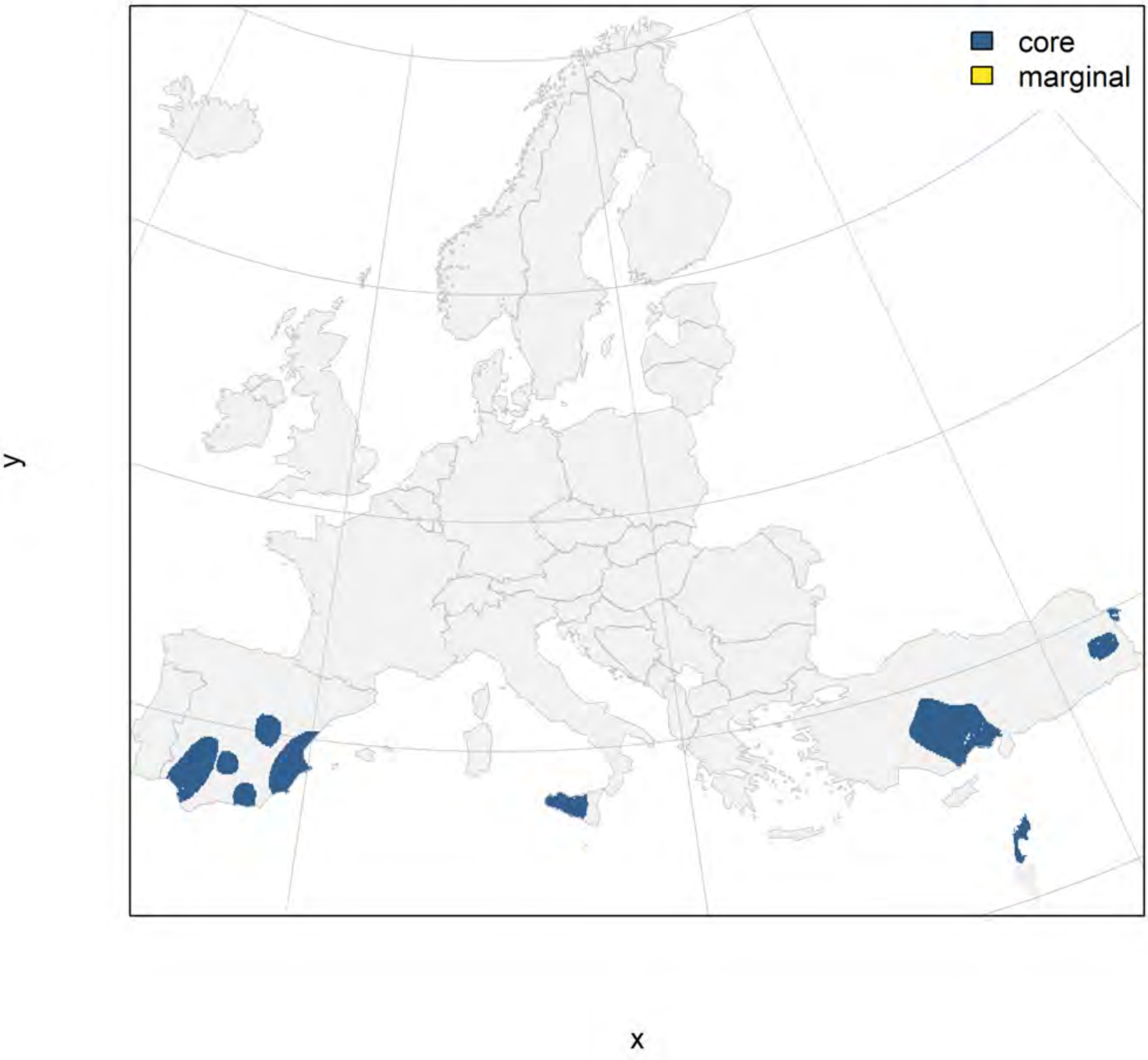

### Polysticta stelleri marginality

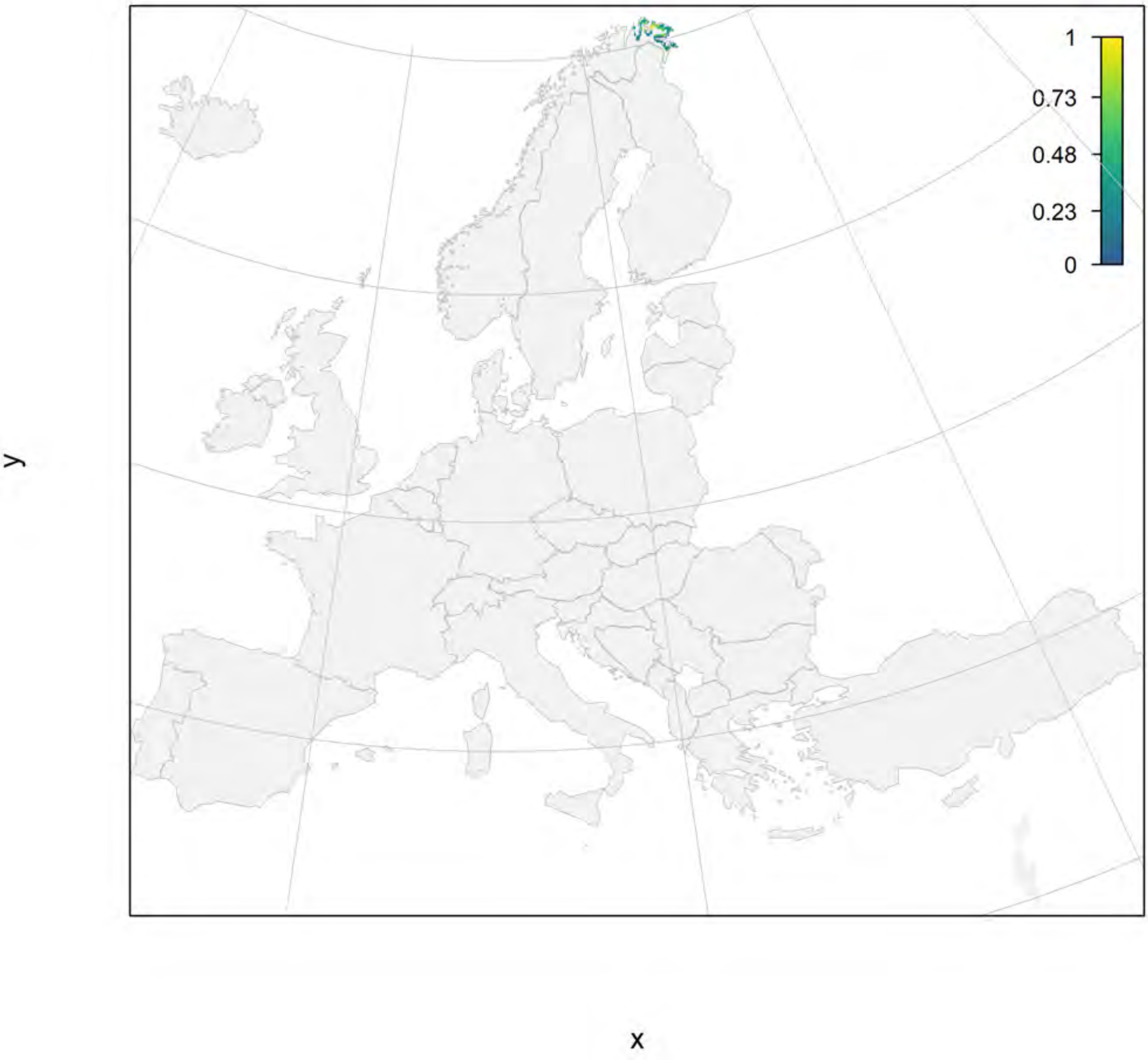

### Polysticta stelleri_filt current marginality

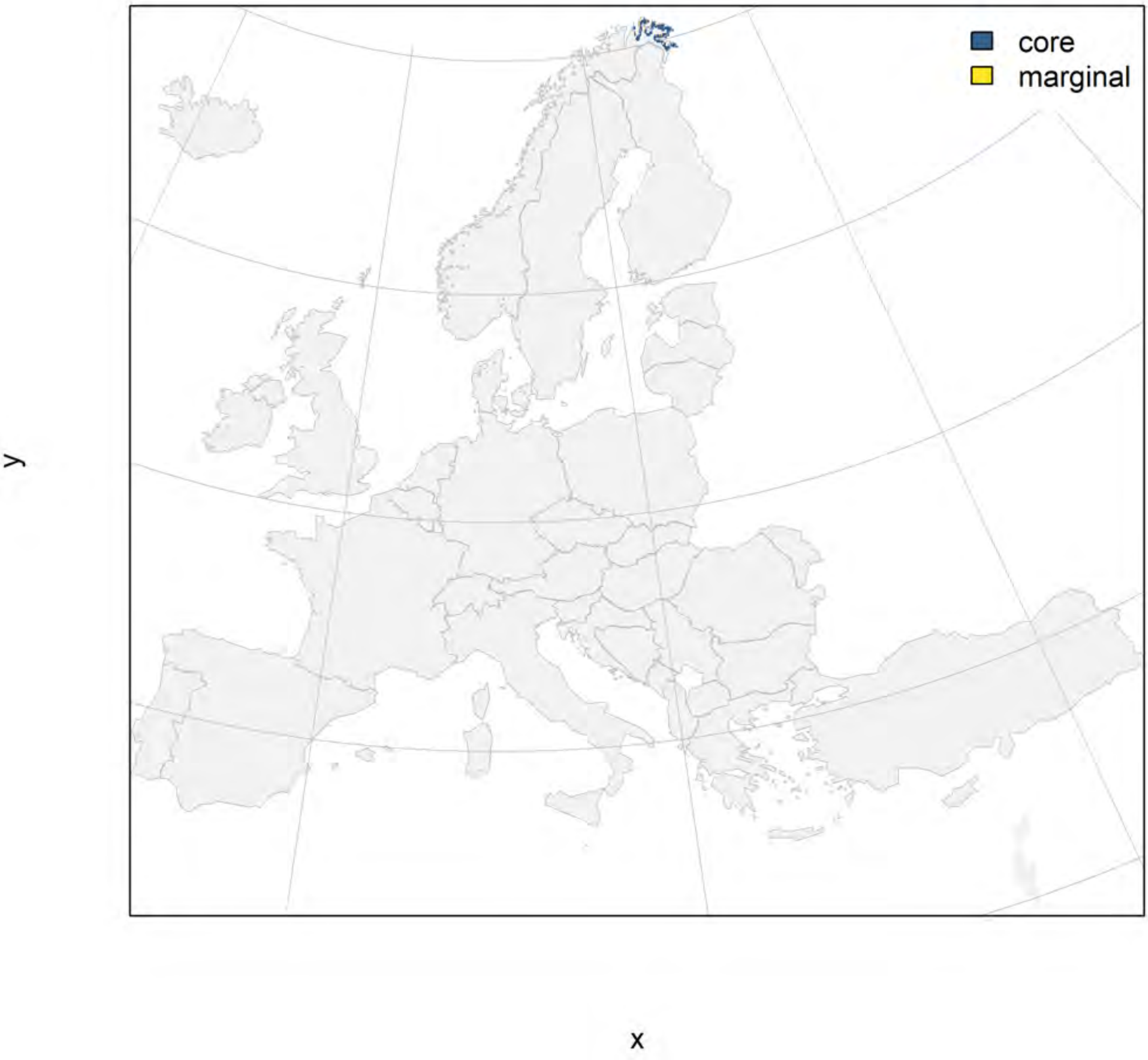

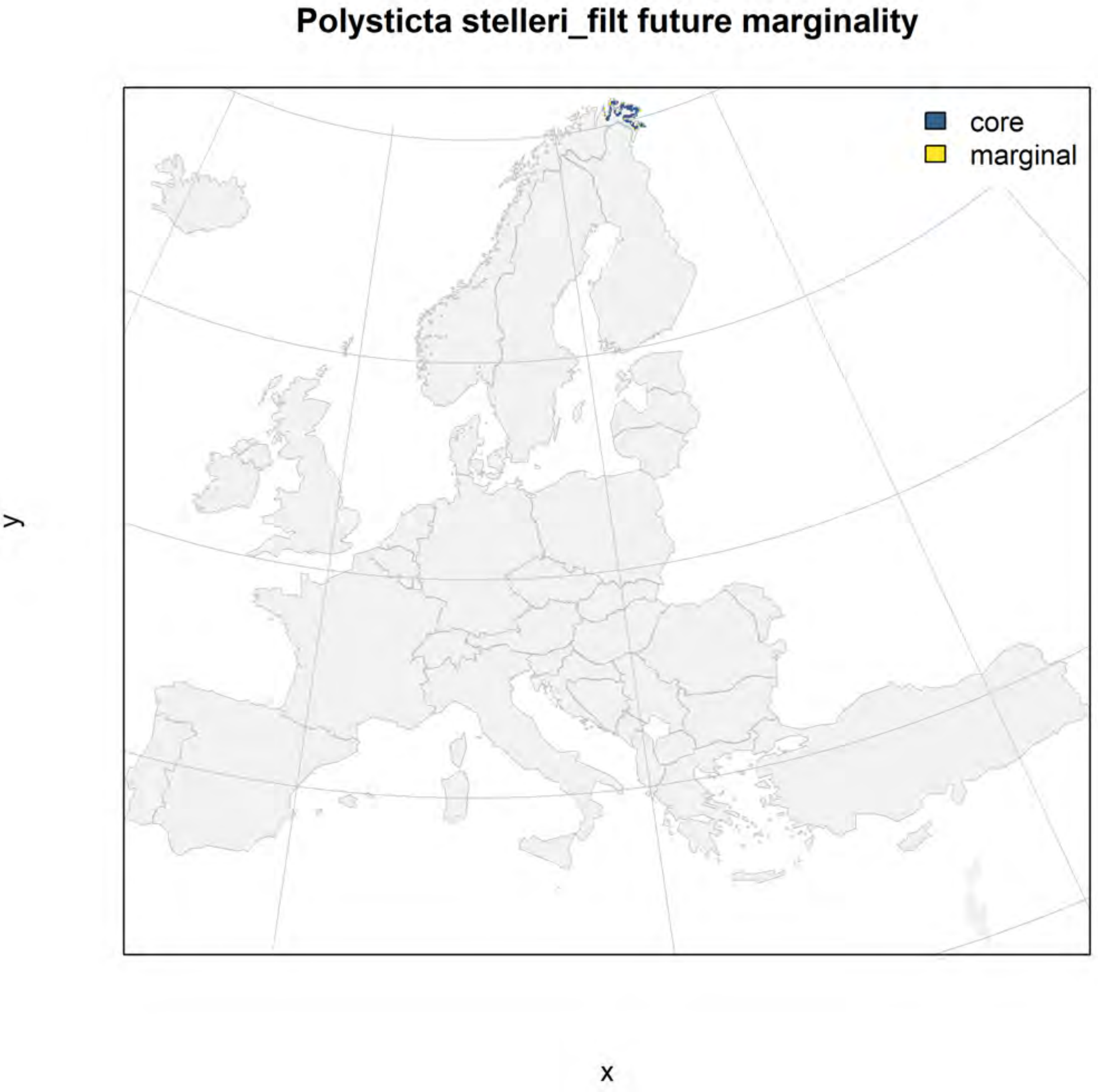

Polysticta stelleri_filt future marginality
core
marginal
y
x

#### Oxyura leucocephala marginality

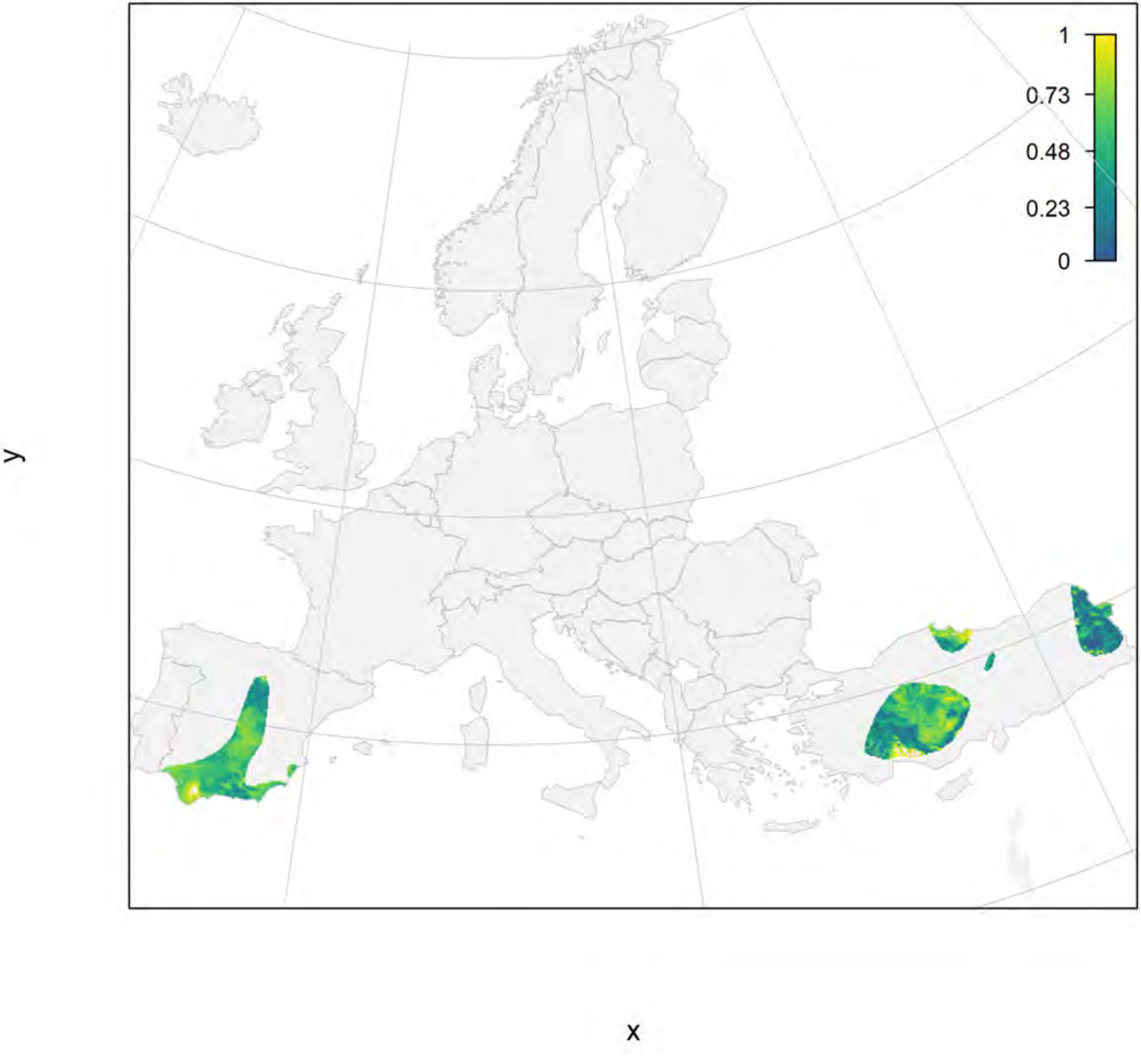

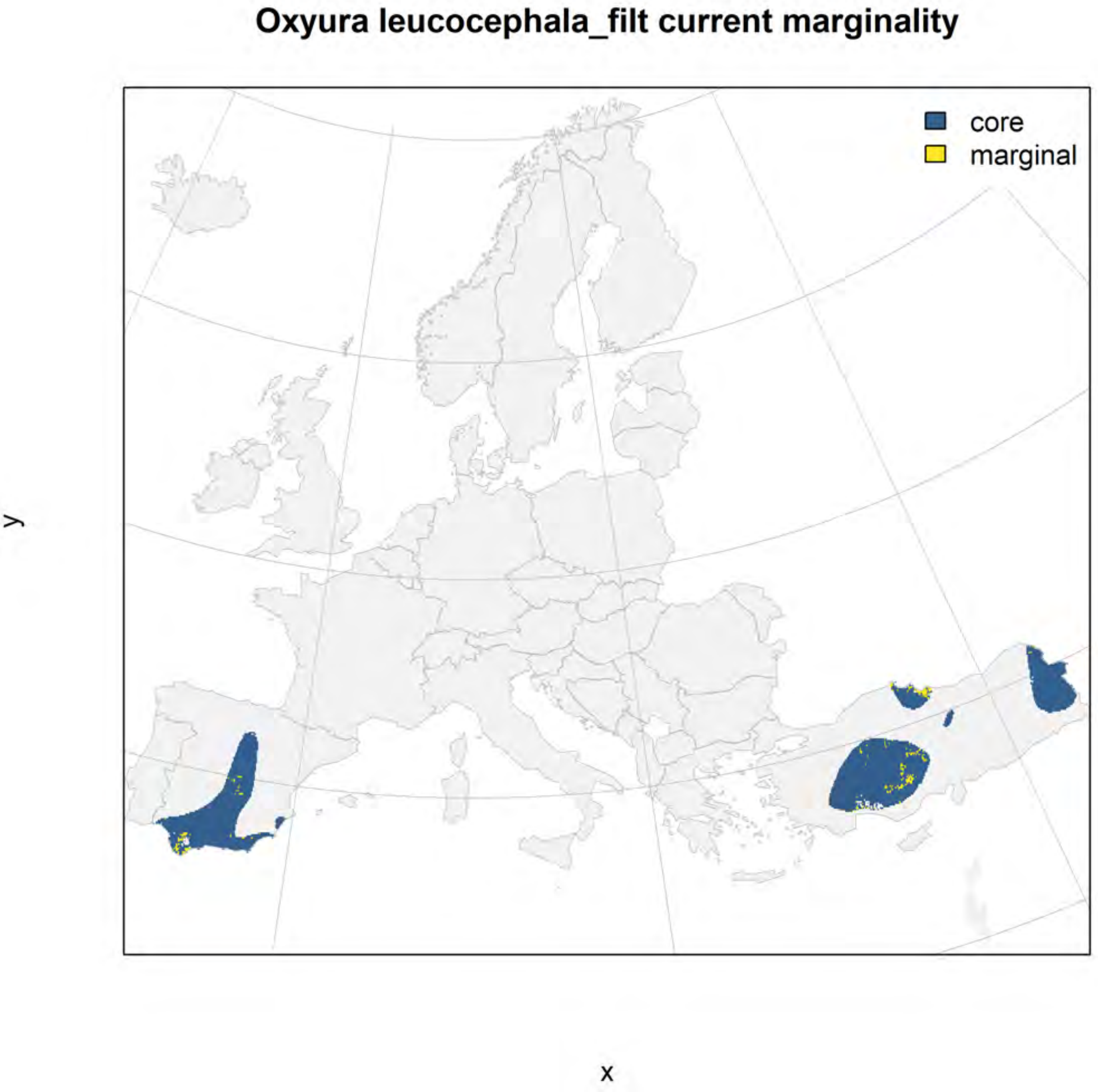
Oxyura leucocephala_filt current marginality
core
marginal
y
x

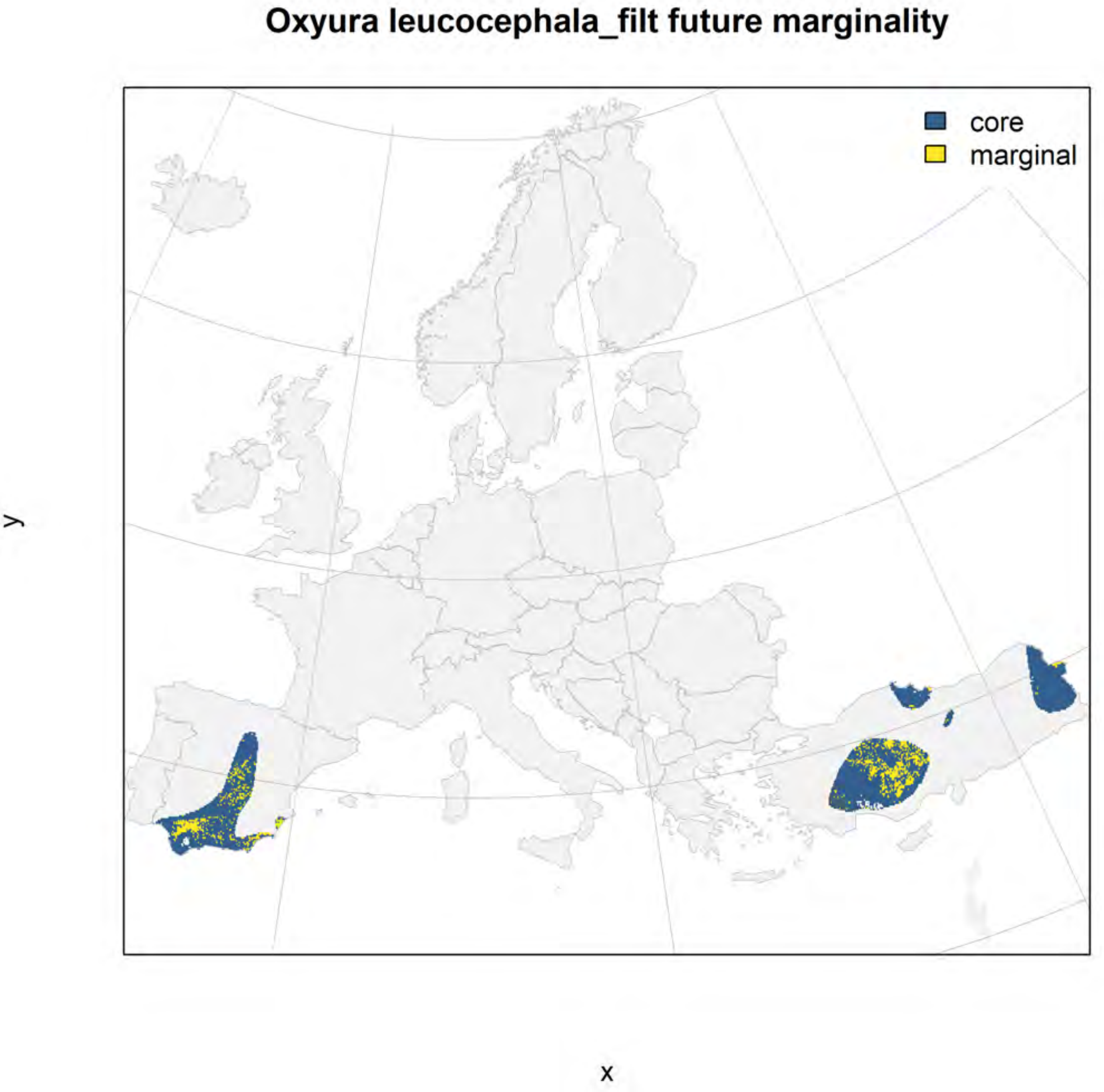
Oxyura leucocephala_filt future marginality
core
marginal
y
x

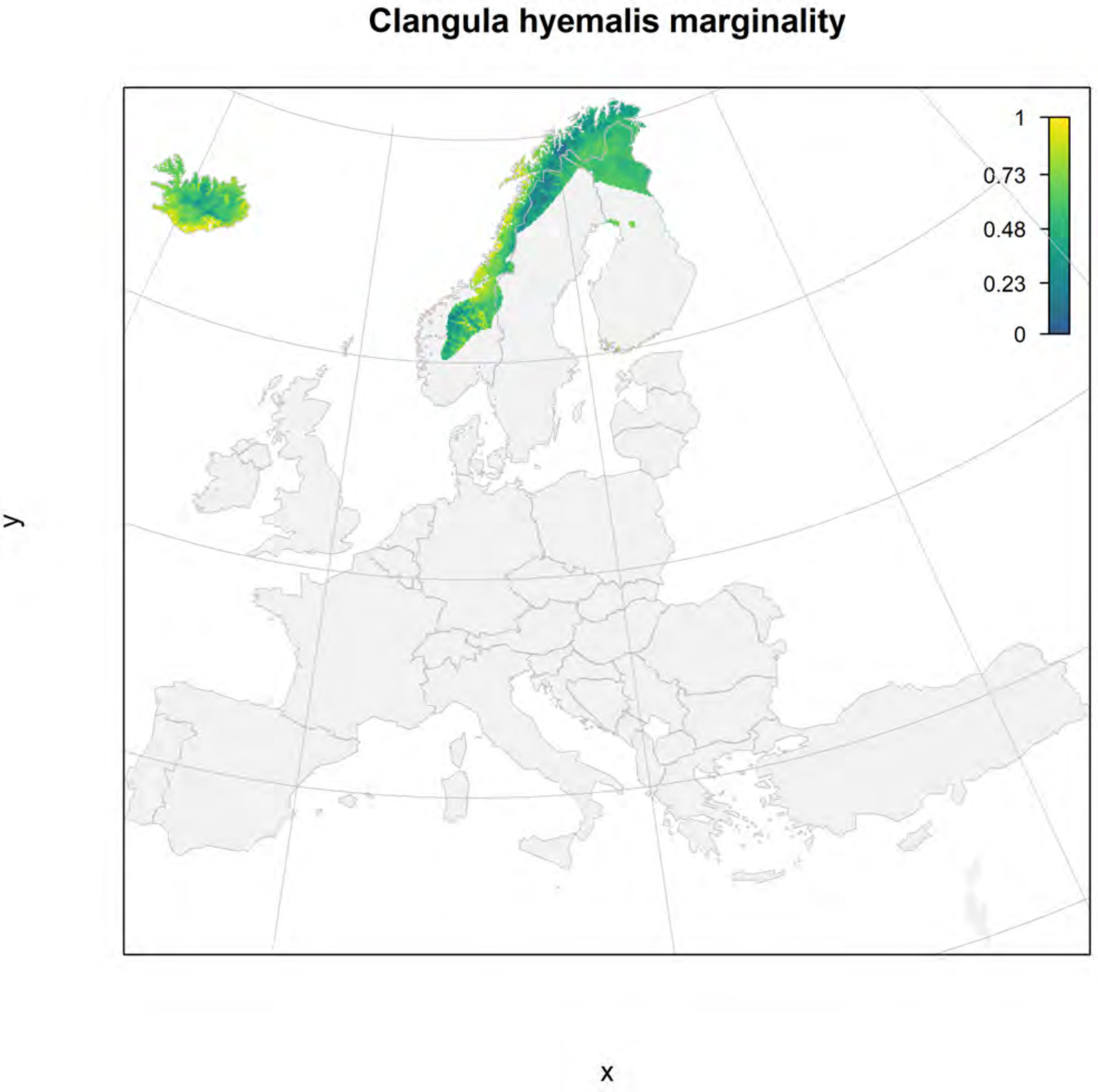
Clangula hyemalis marginality
1
0.73
0.48
0.23
0
y
x

#### Clangula hyemalis_filt current marginality

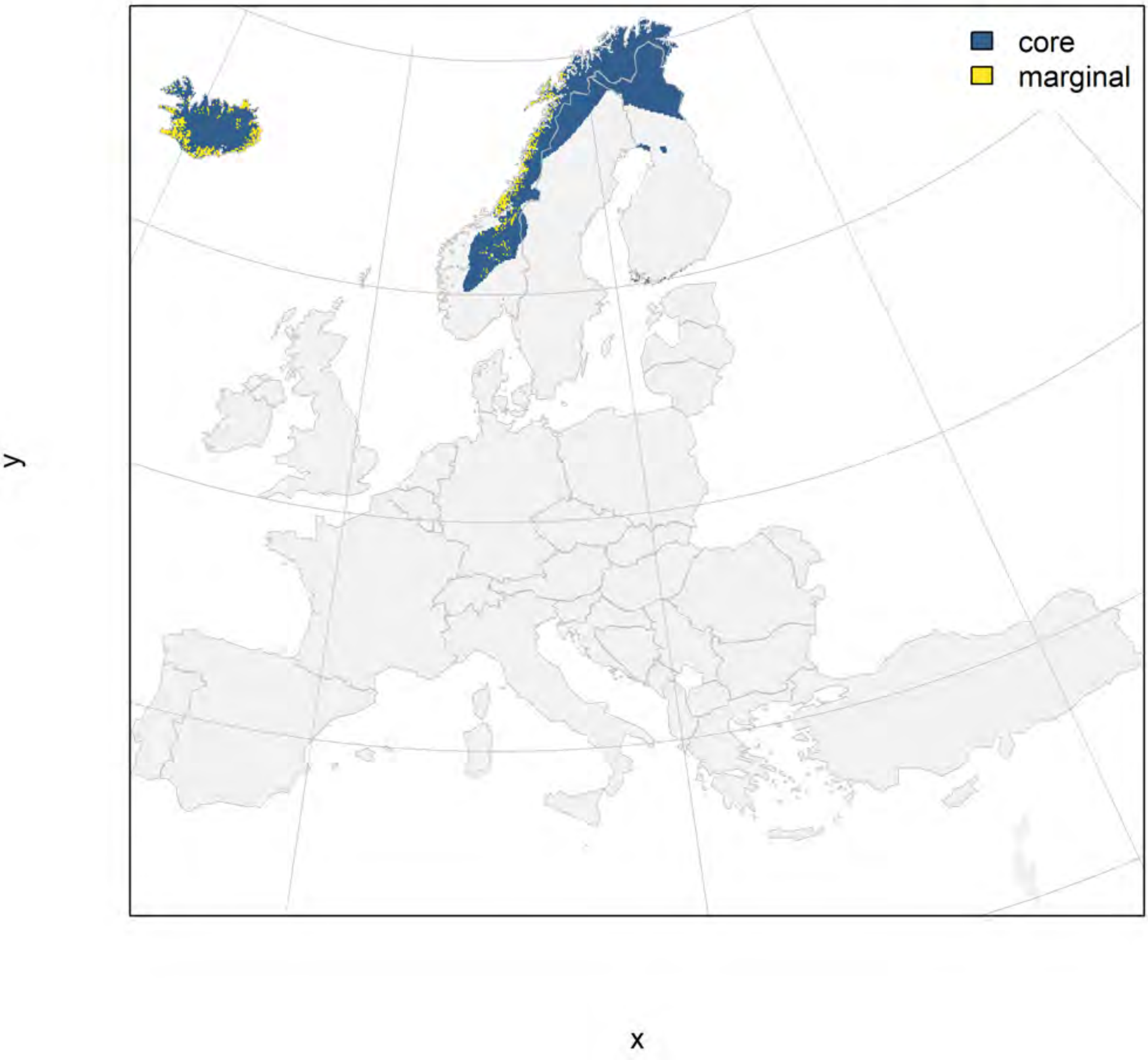

### Clangula hyemalis_filt future marginality

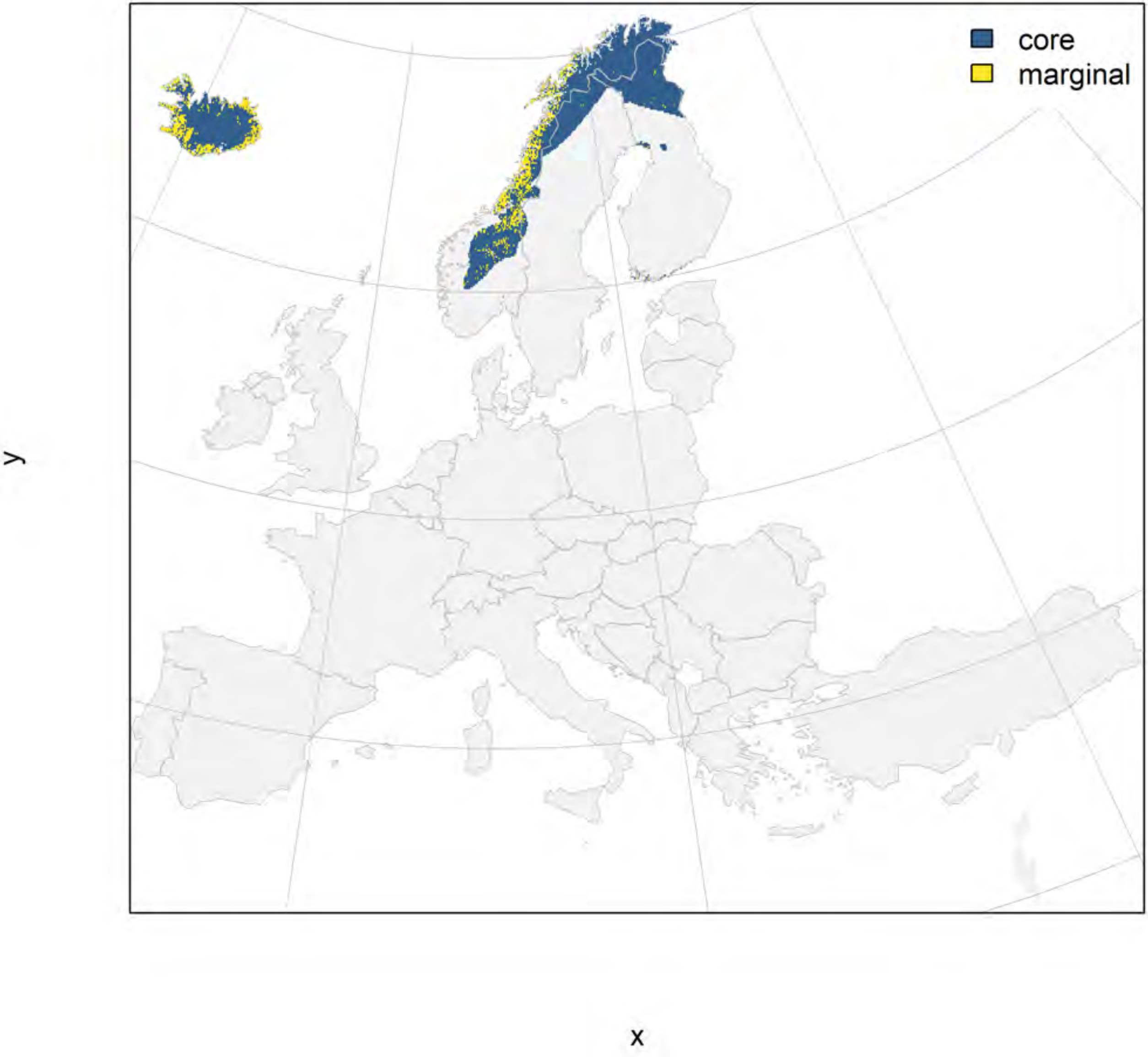

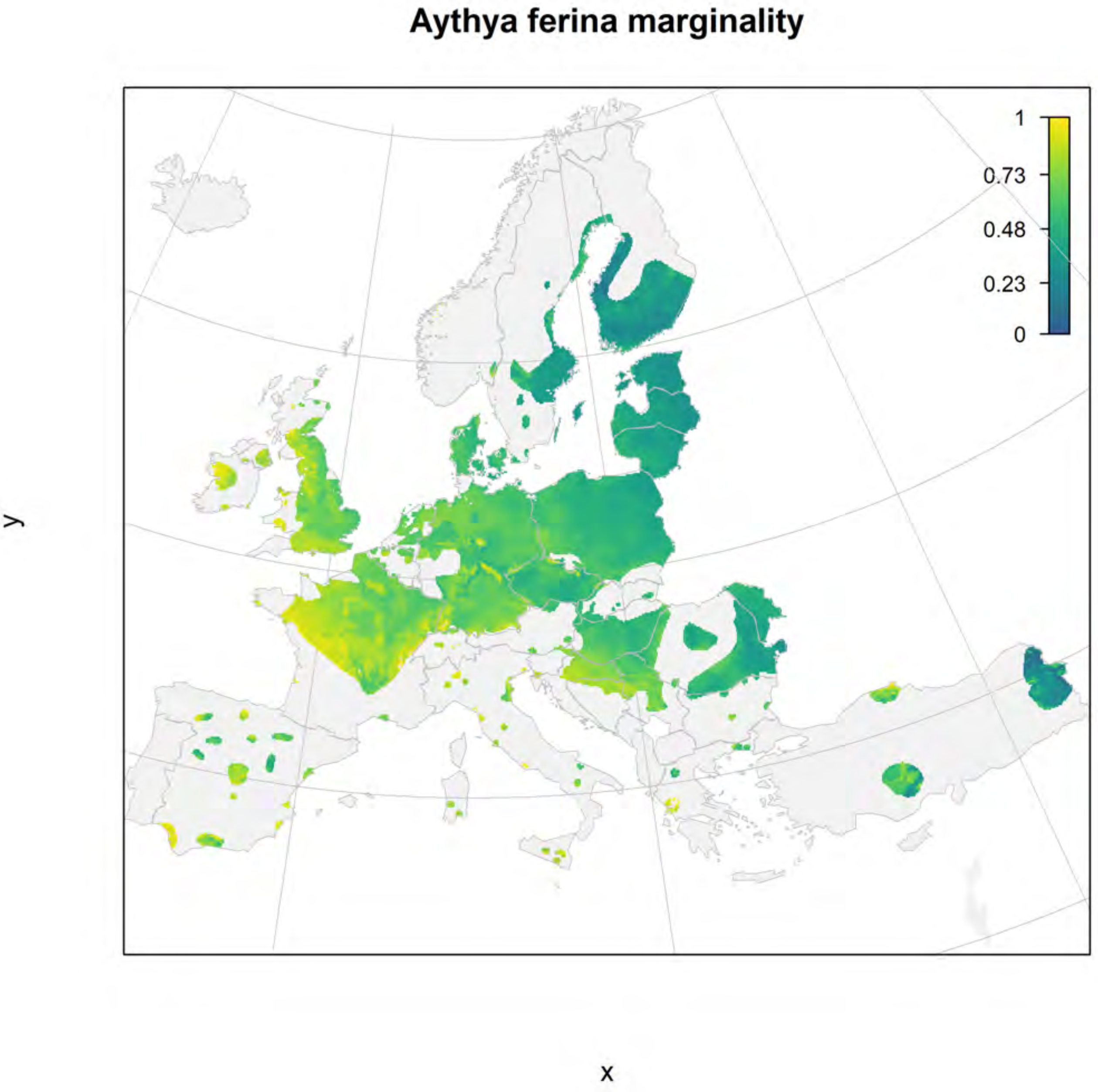

Aythya ferina marginality
1
0.73
0.48
0.23
0
y
x

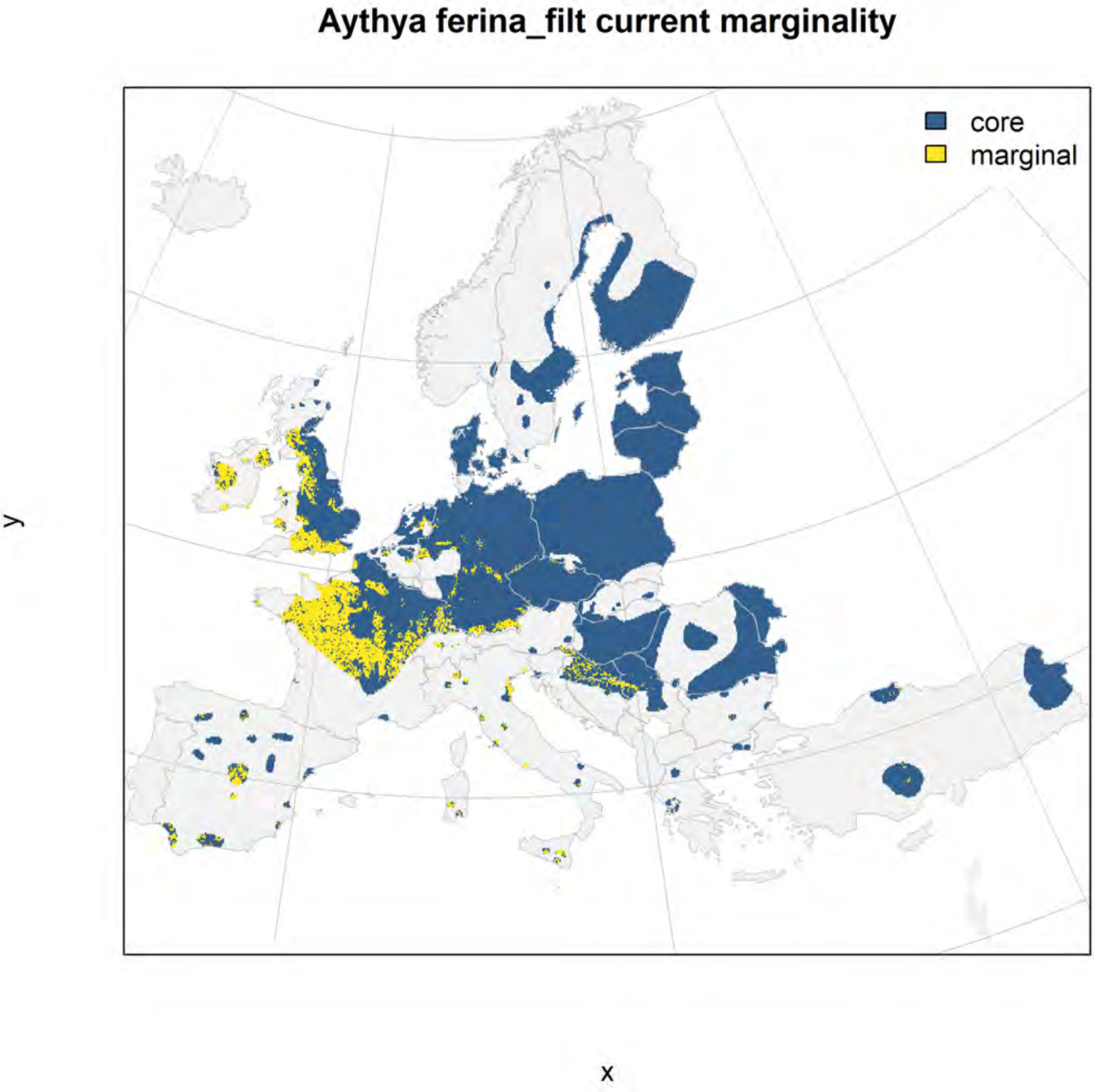
Aythya ferina_filt current marginality
core
marginal
y
x

#### Aythya ferina_filt future marginality

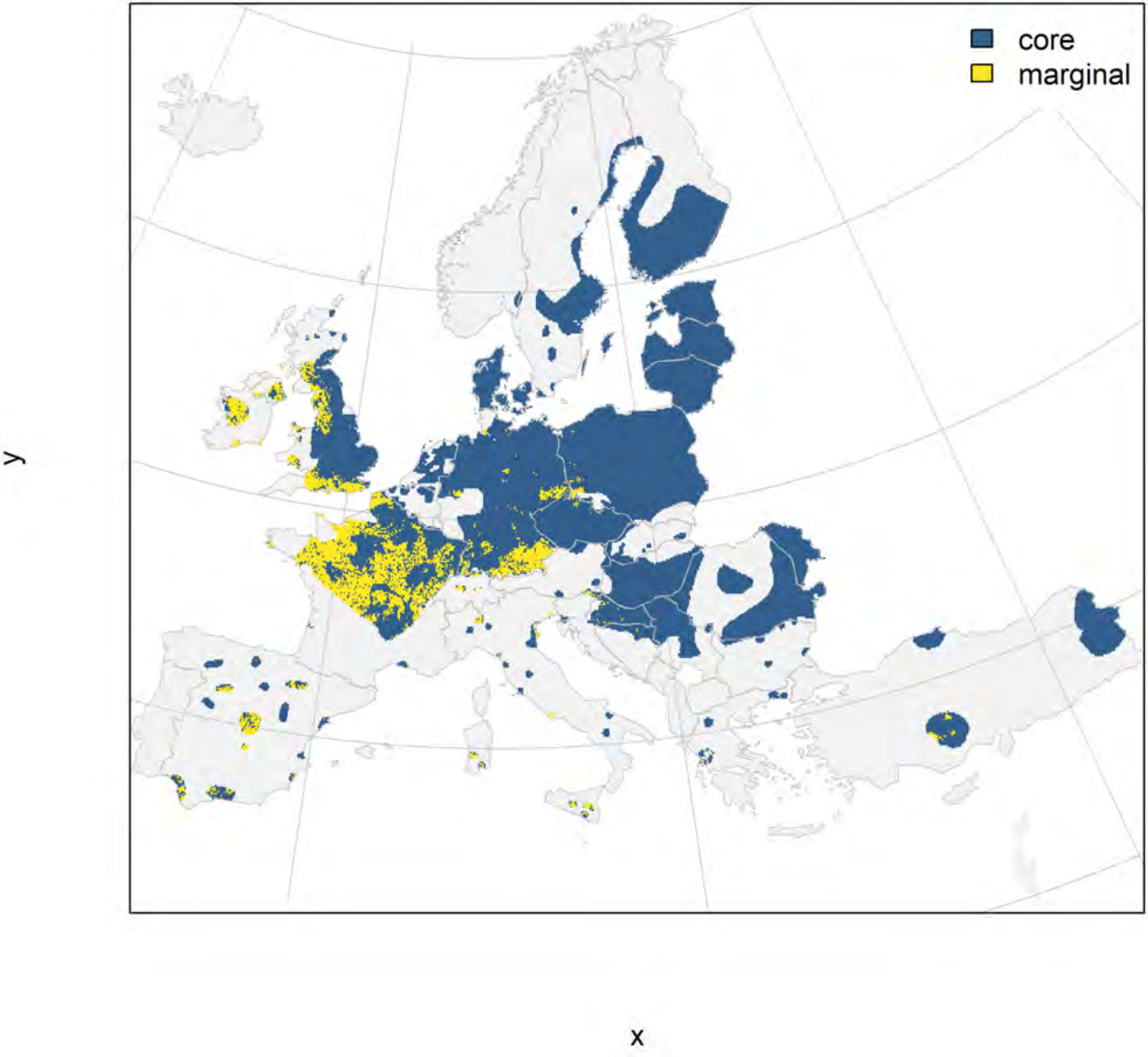

### Tetrao urogallus marginality

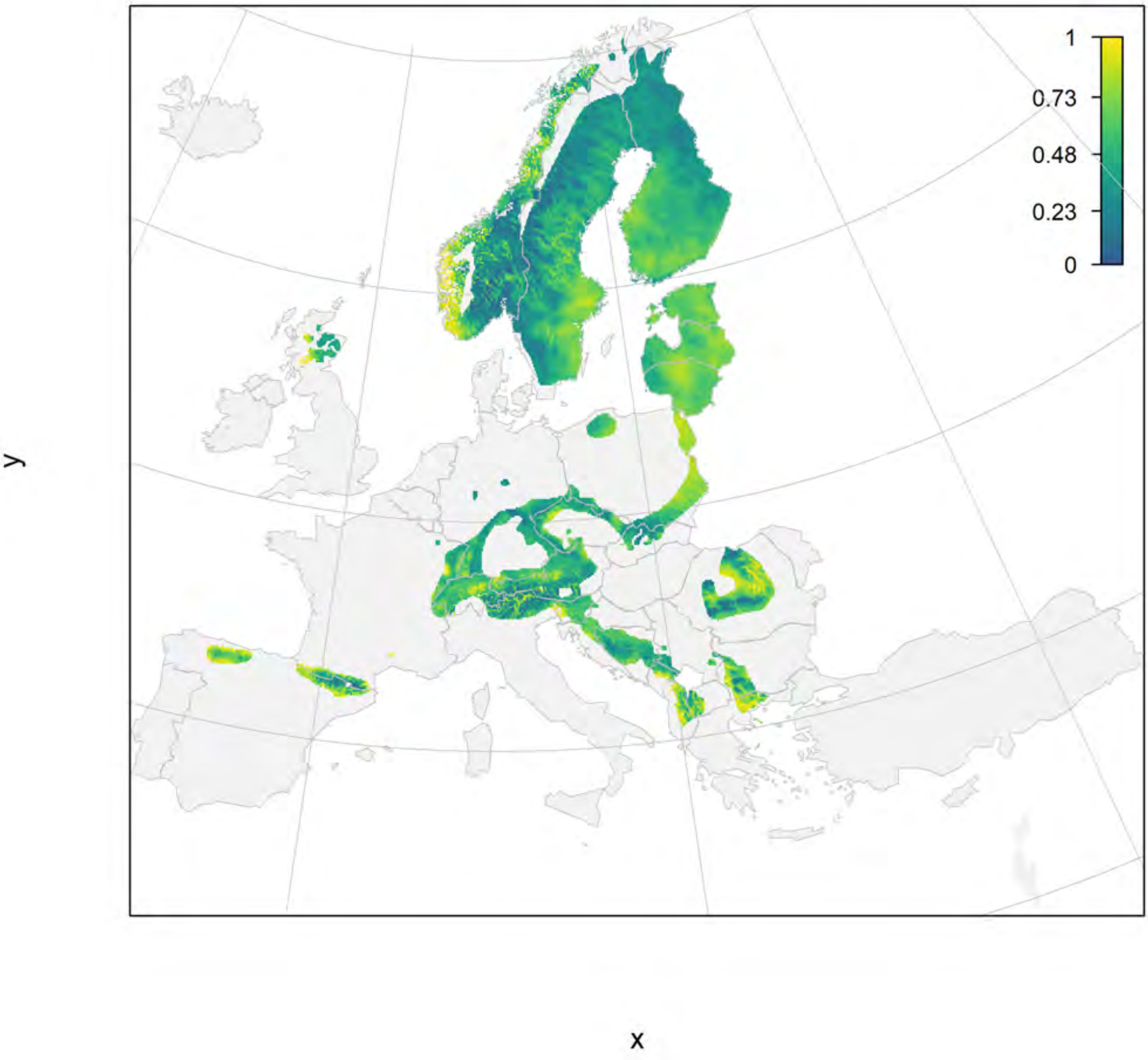

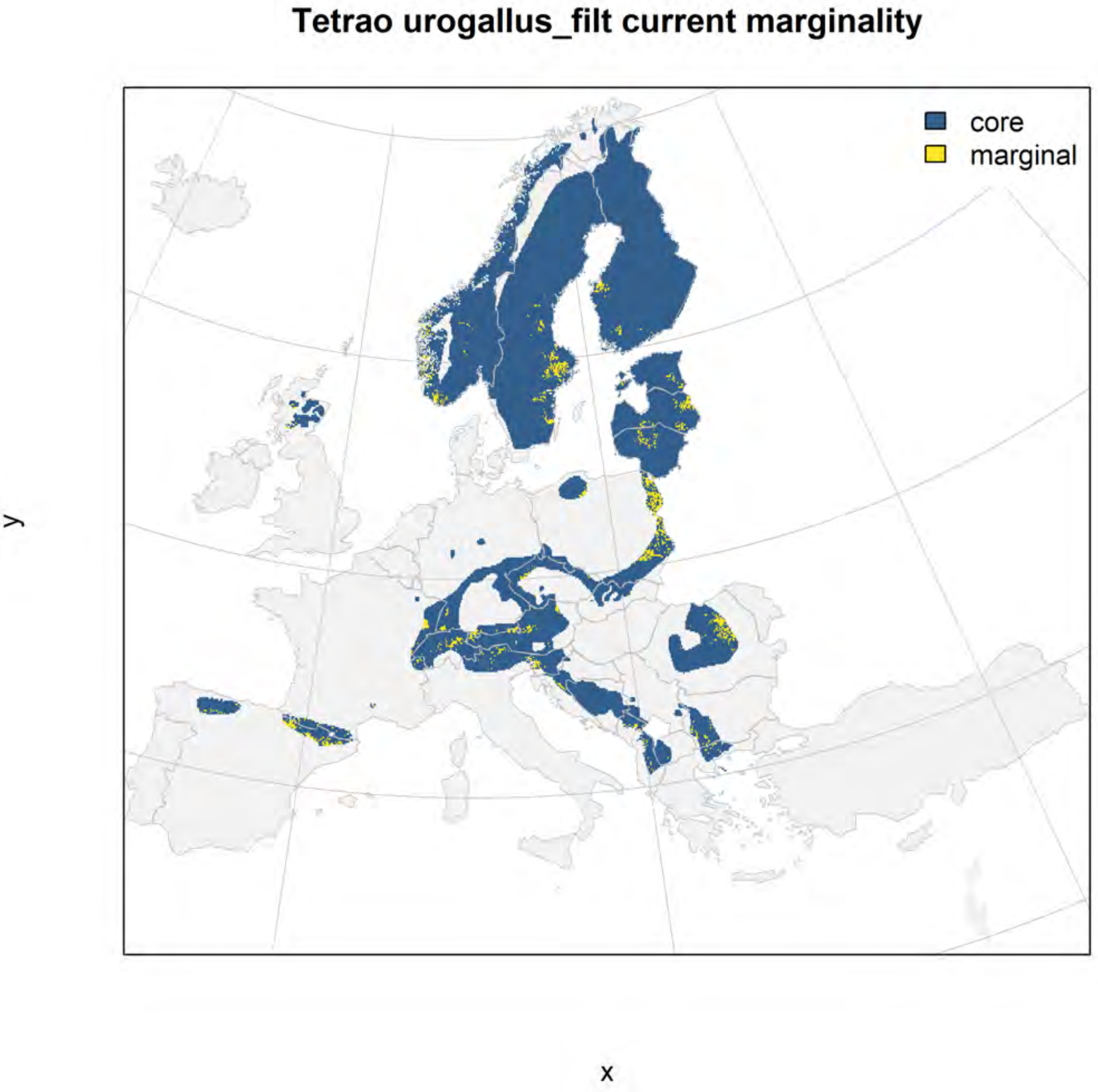

Tetrao urogallus_filt current marginality
core
marginal
y
x

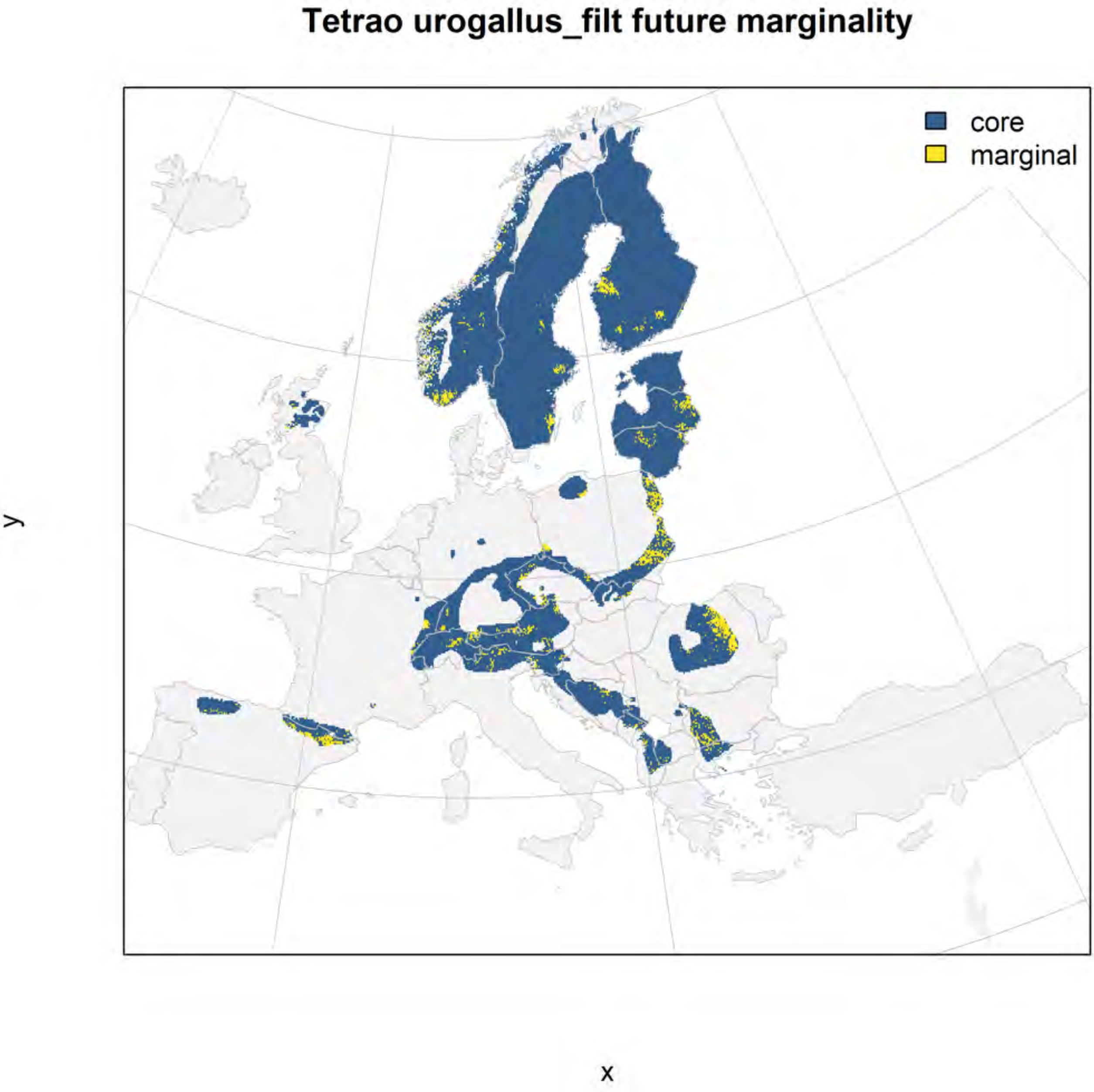
Tetrao urogallus_filt future marginality
core
marginal
y
x

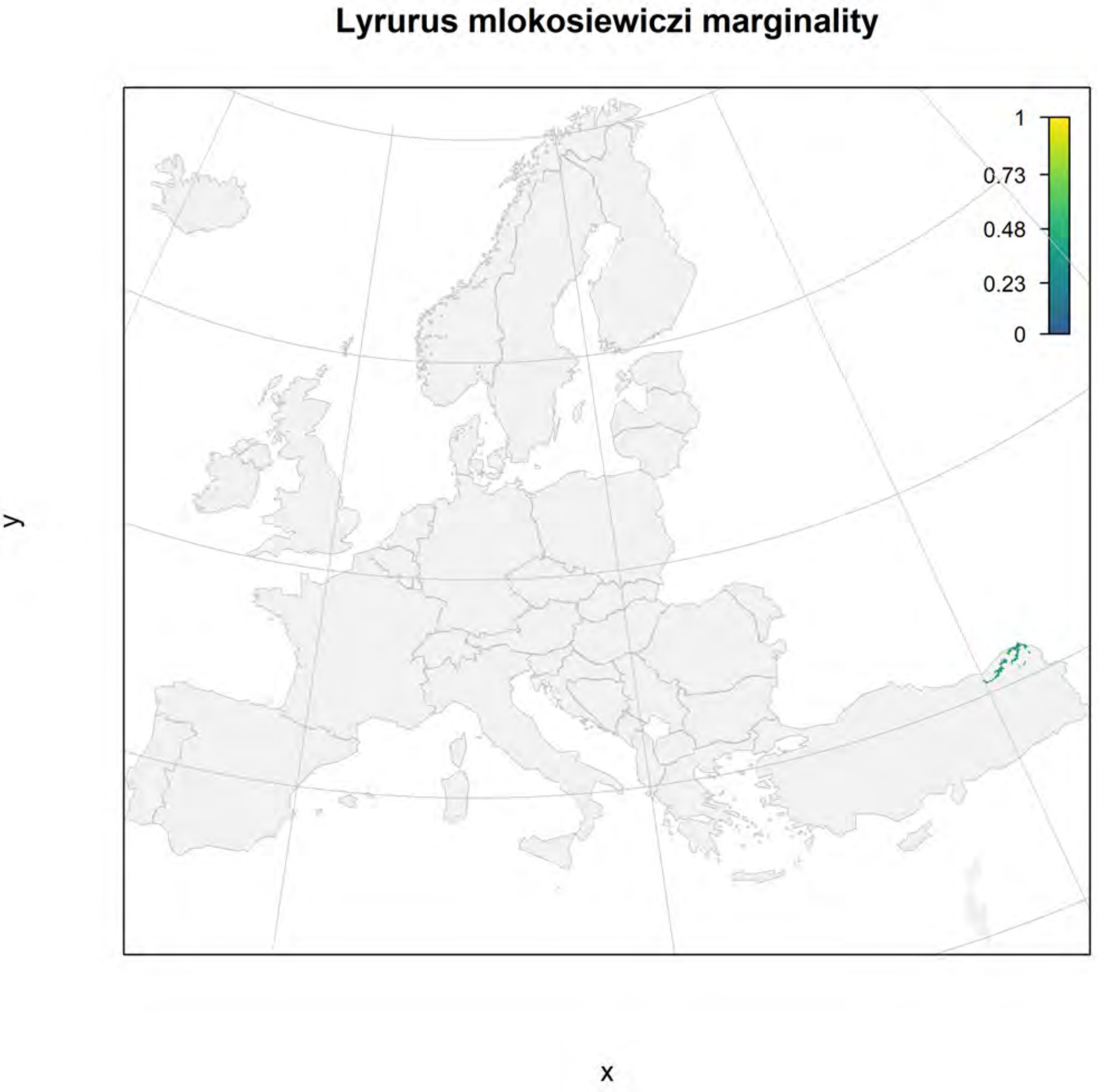
Lyrurus mlokosiewiczi marginality
1
0.73
0.48
0.23
0
y
x

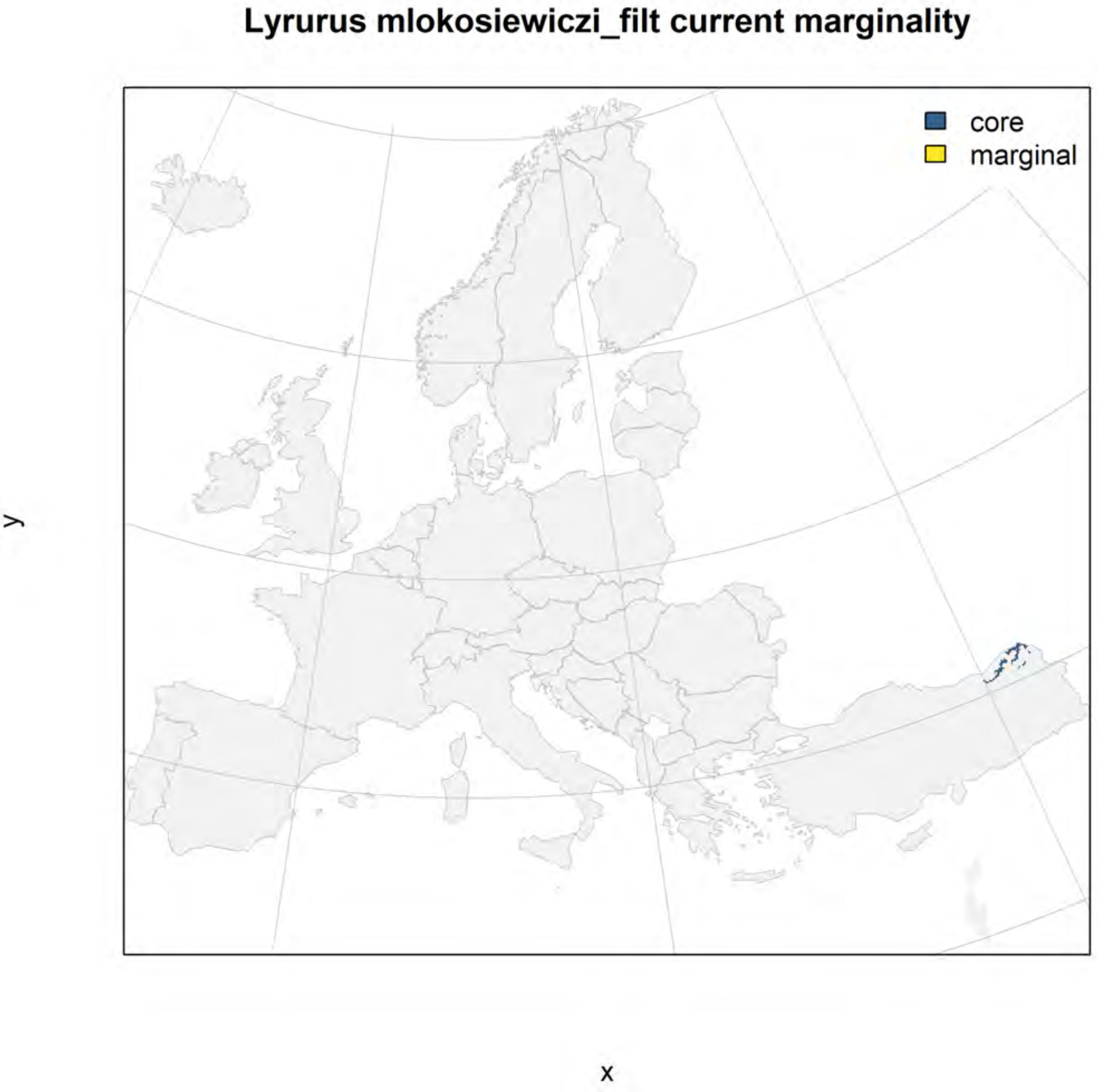

Lyrurus mlokosiewiczi_filt current marginality
core
marginal
y
x

#### Lyrurus mlokosiewiczi_filt future marginality

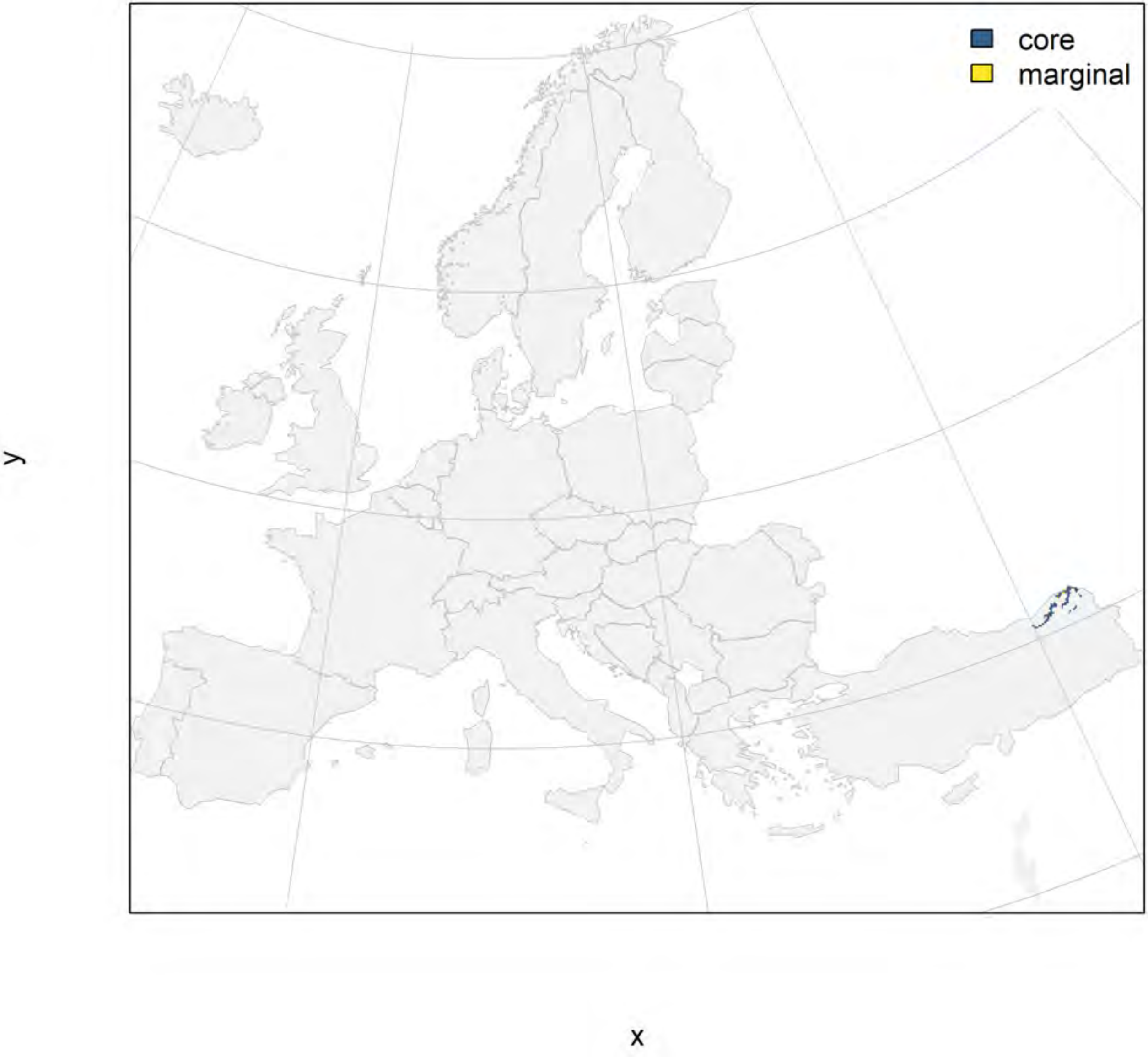

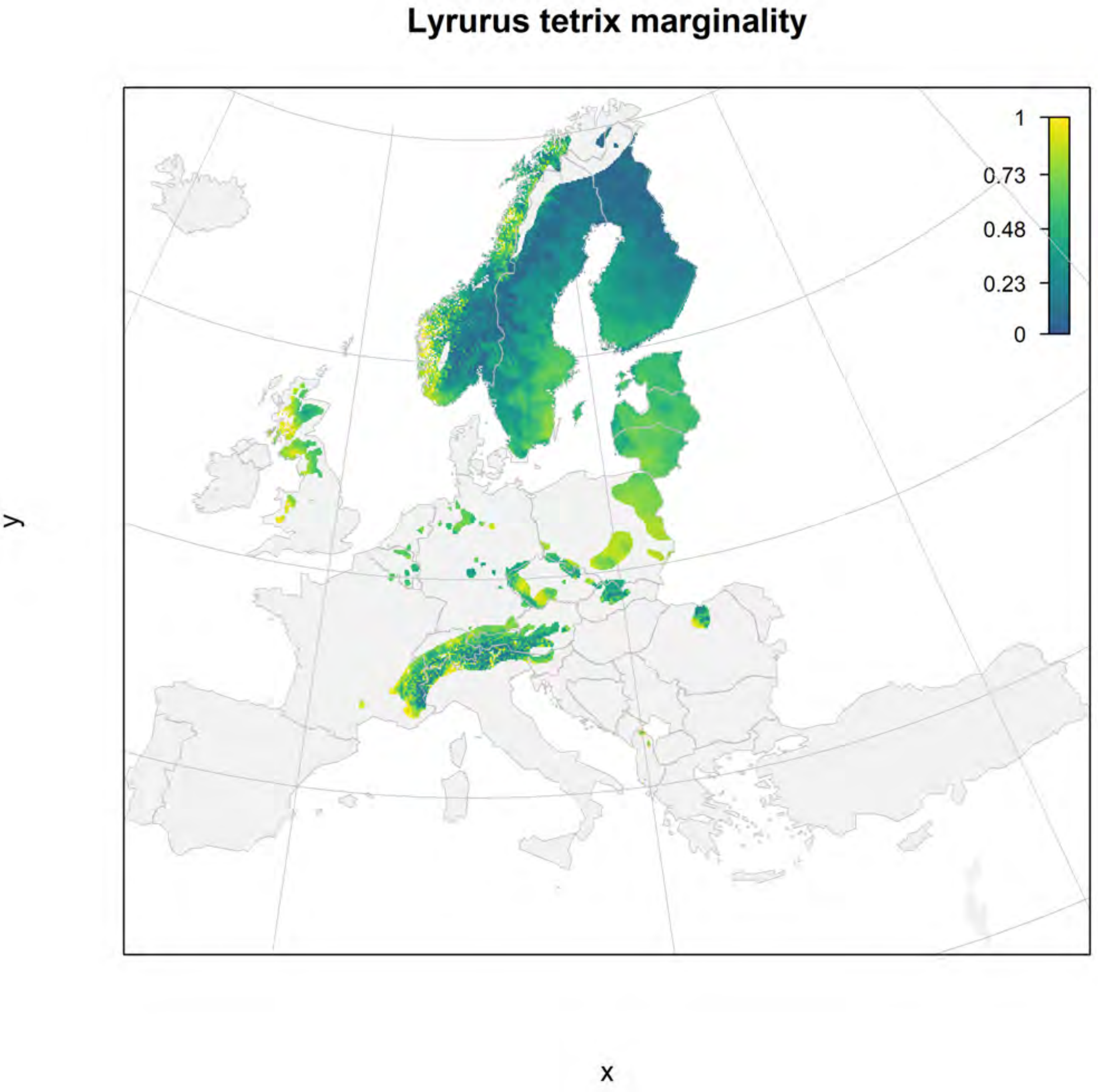

Lyrurus tetrix marginality
1
0.73
0.48
0.23
0
y
x

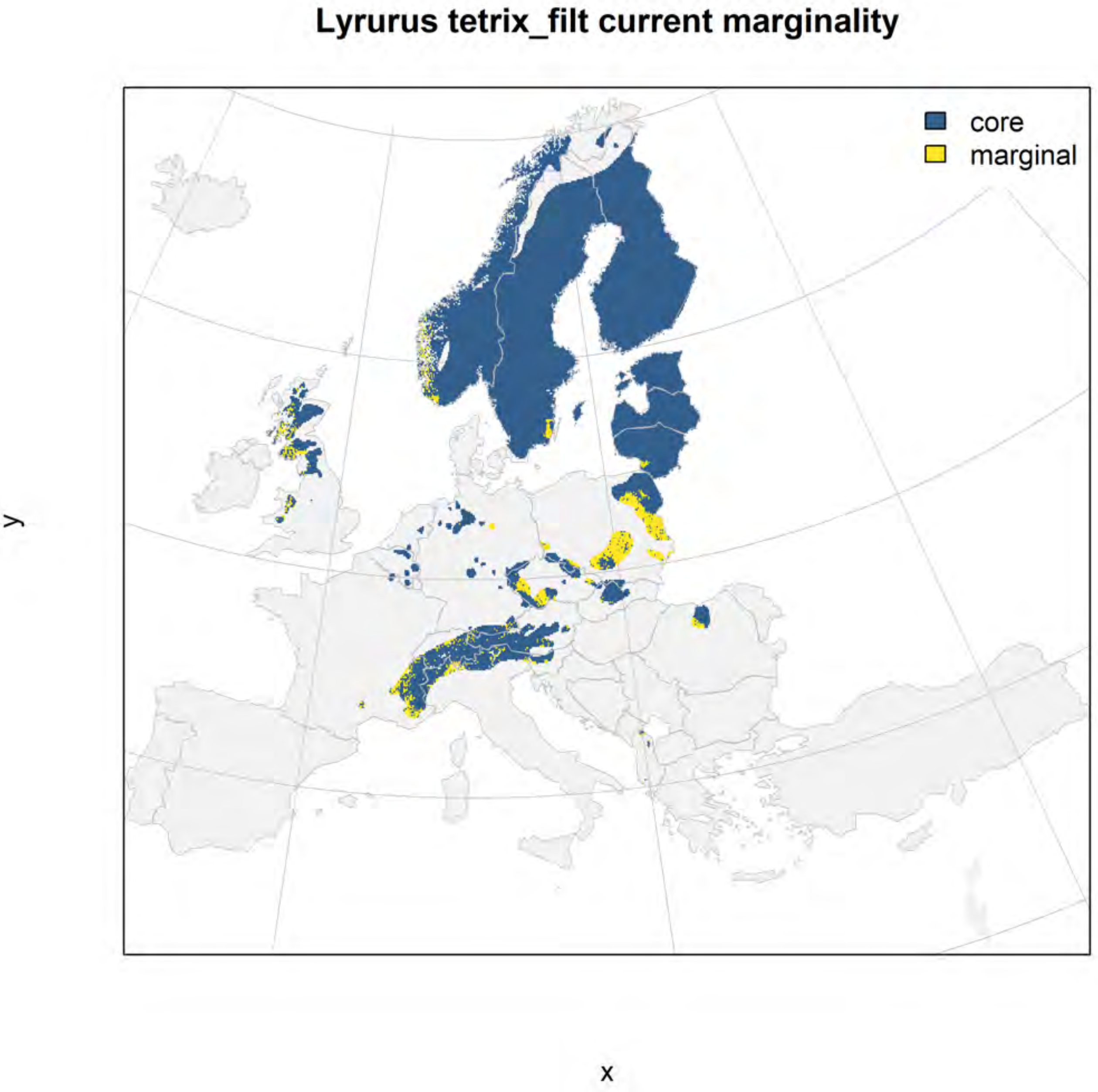
Lyrurus tetrix_filt current marginality
core
marginal
y
x

### Lyrurus tetrix_filt future marginality

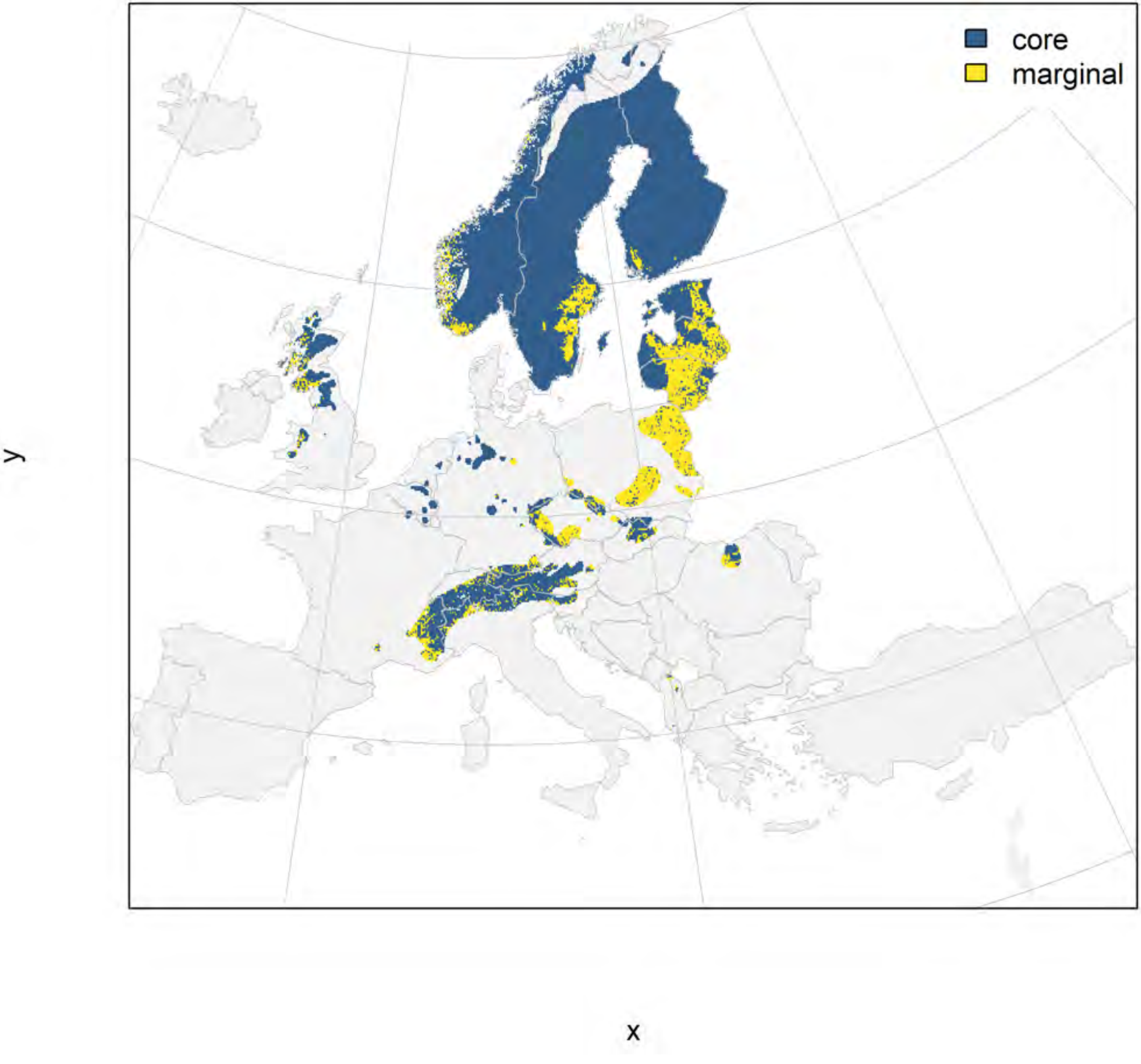

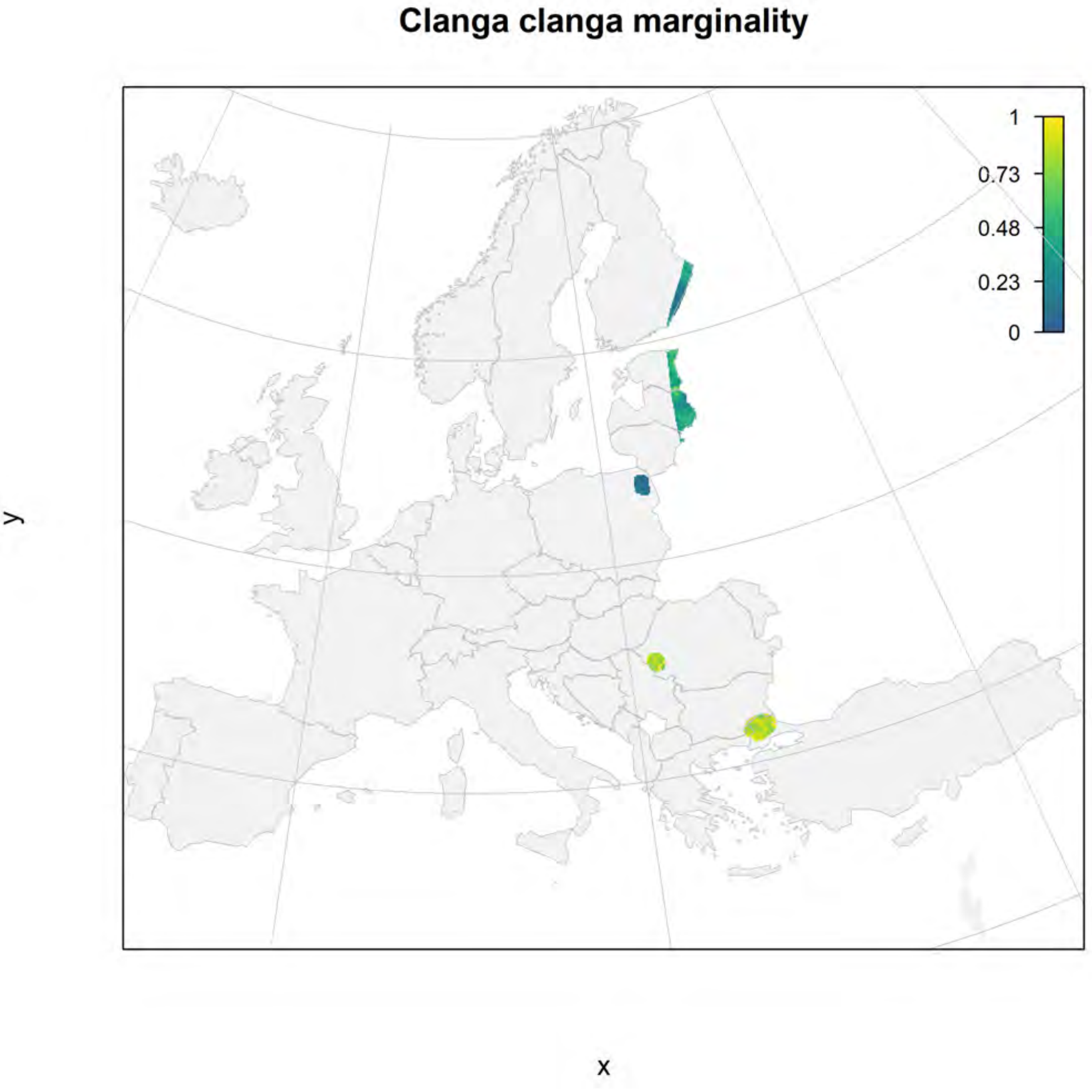
Clanga clanga marginality
1
0.73
0.48
0.23
0
y
x

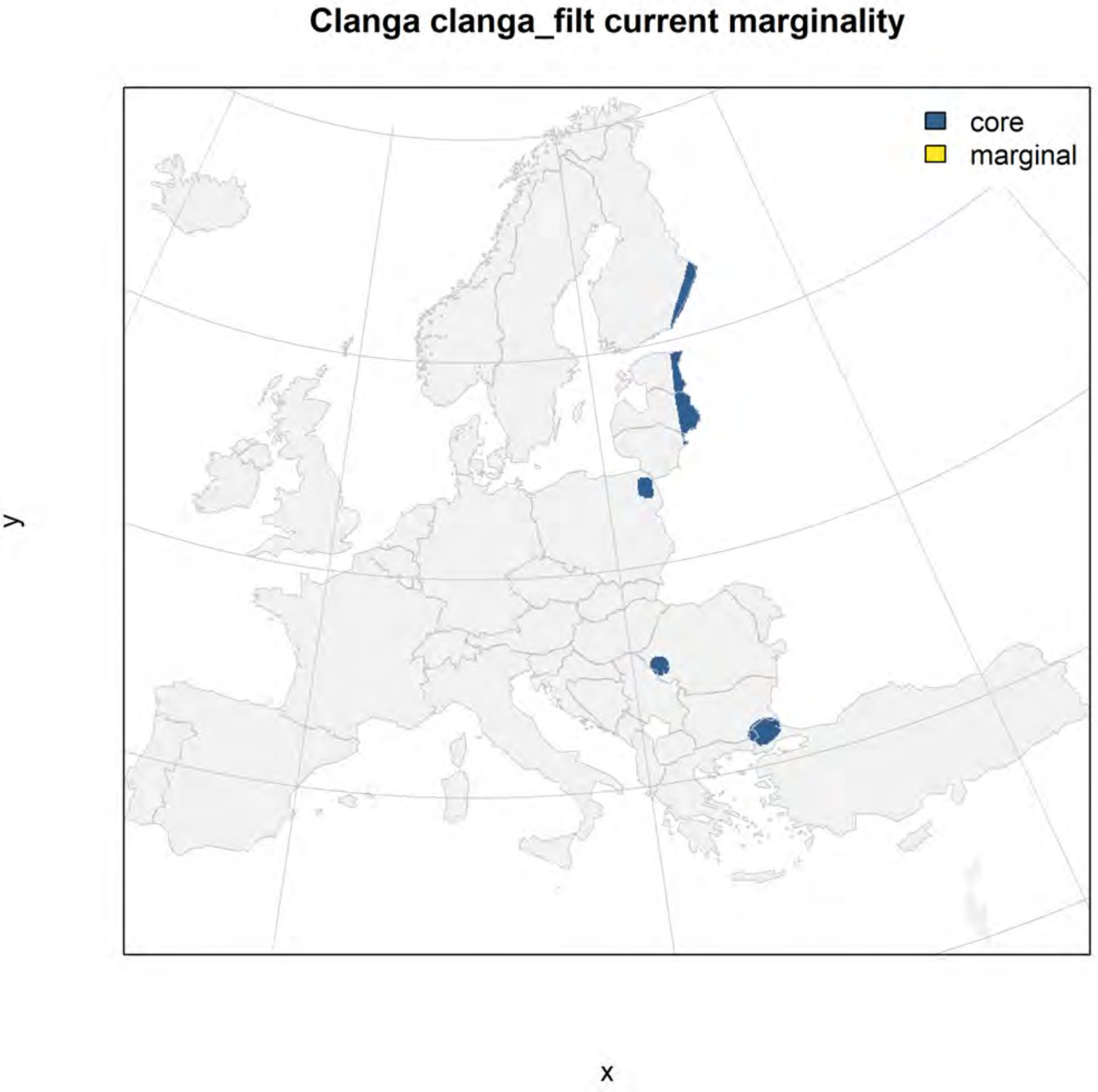
Clanga clanga_filt current marginality
core
marginal
y
x

#### Clanga clanga_filt future marginality

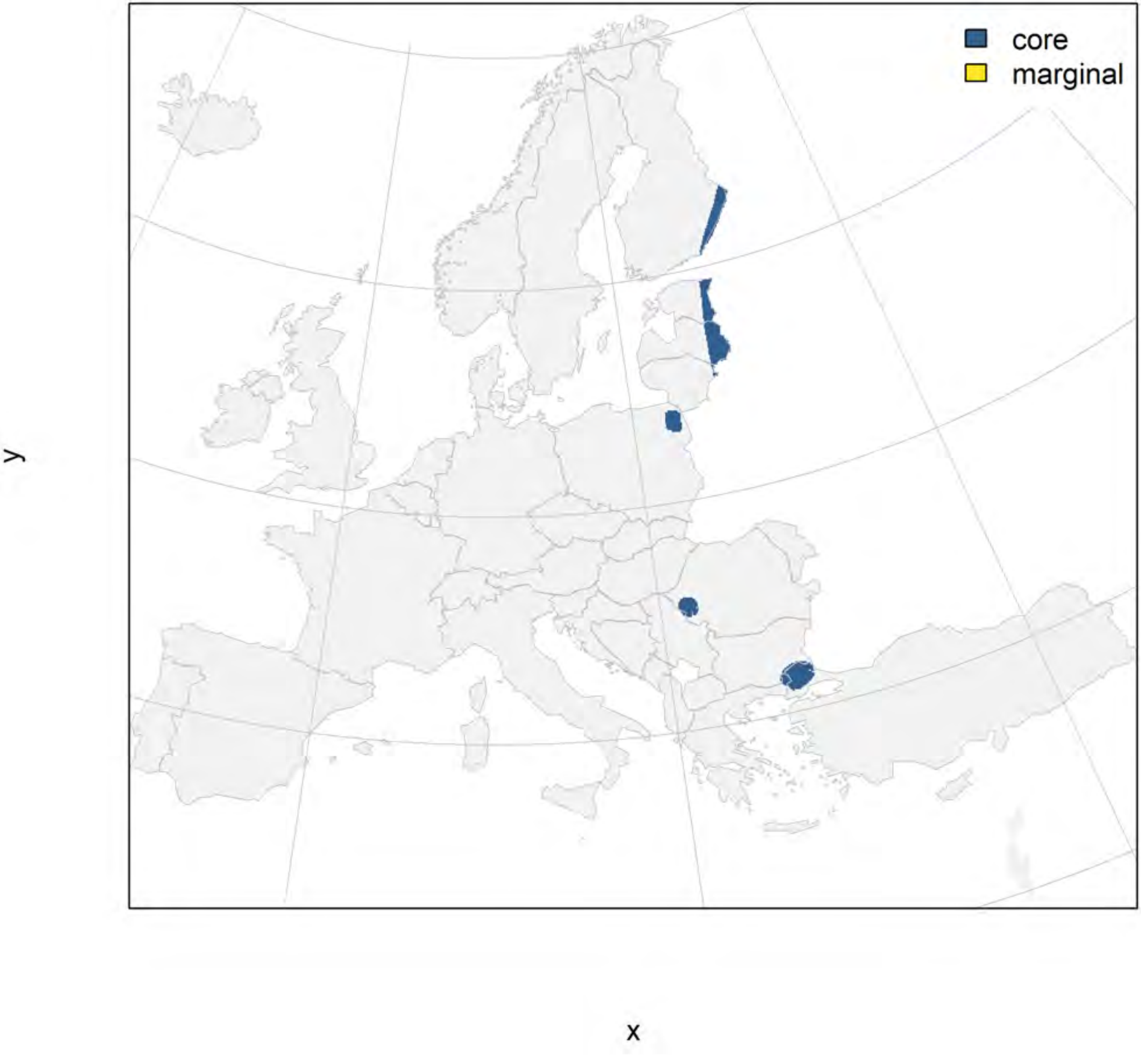

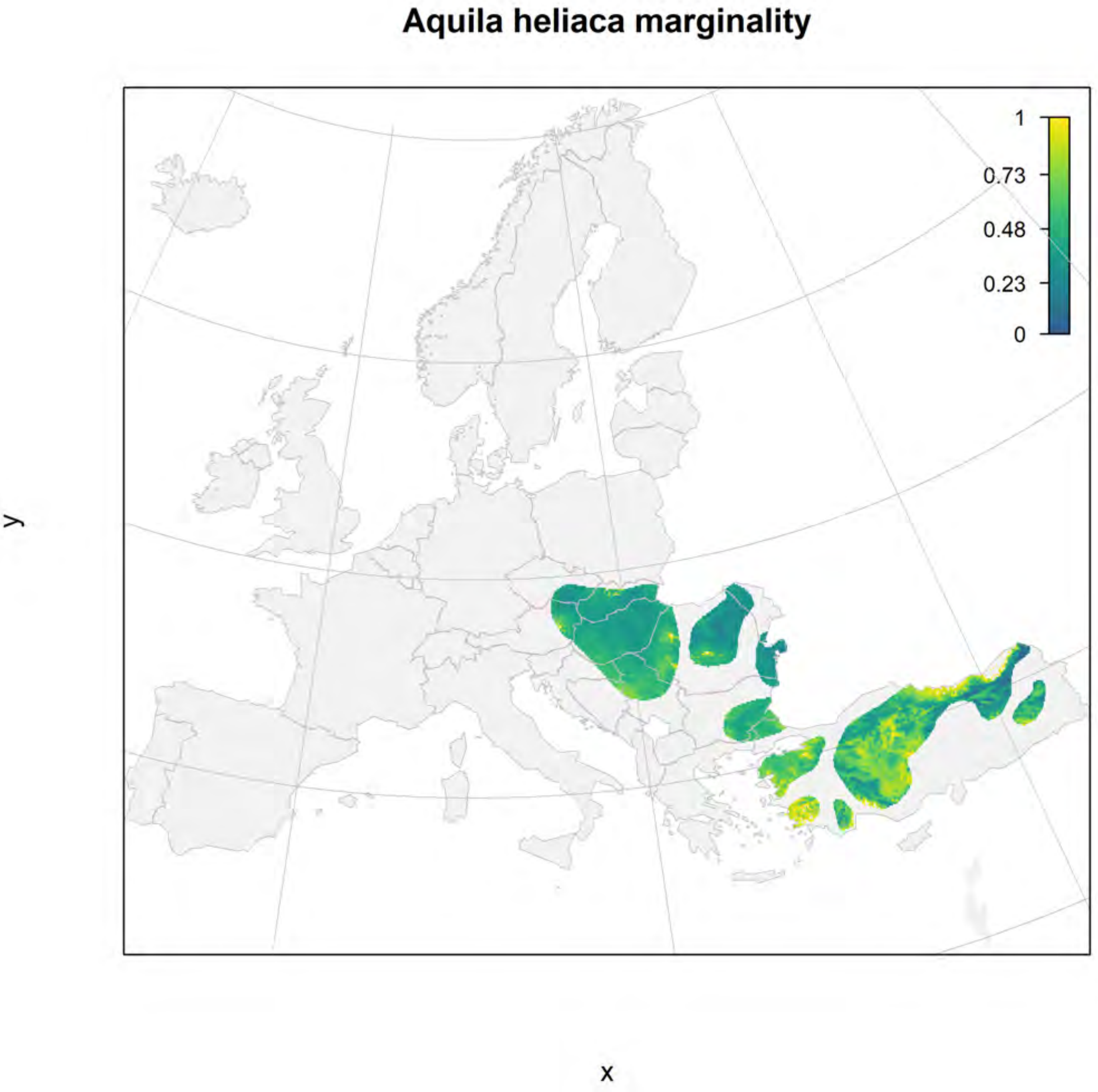
Aquila heliaca marginality
1
0.73
0.48
0.23
0
y
x

#### Aquila heliaca_filt current marginality

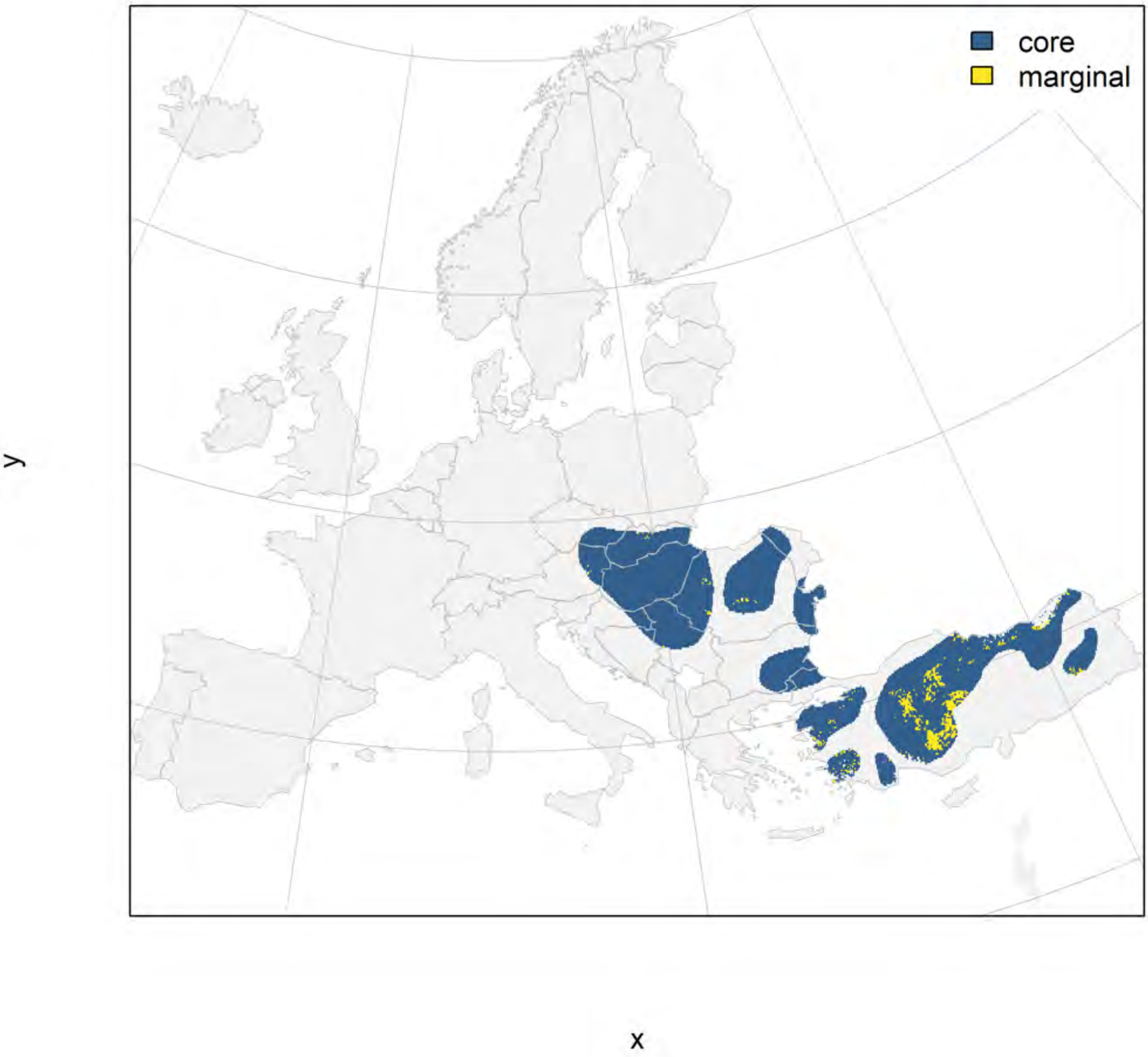

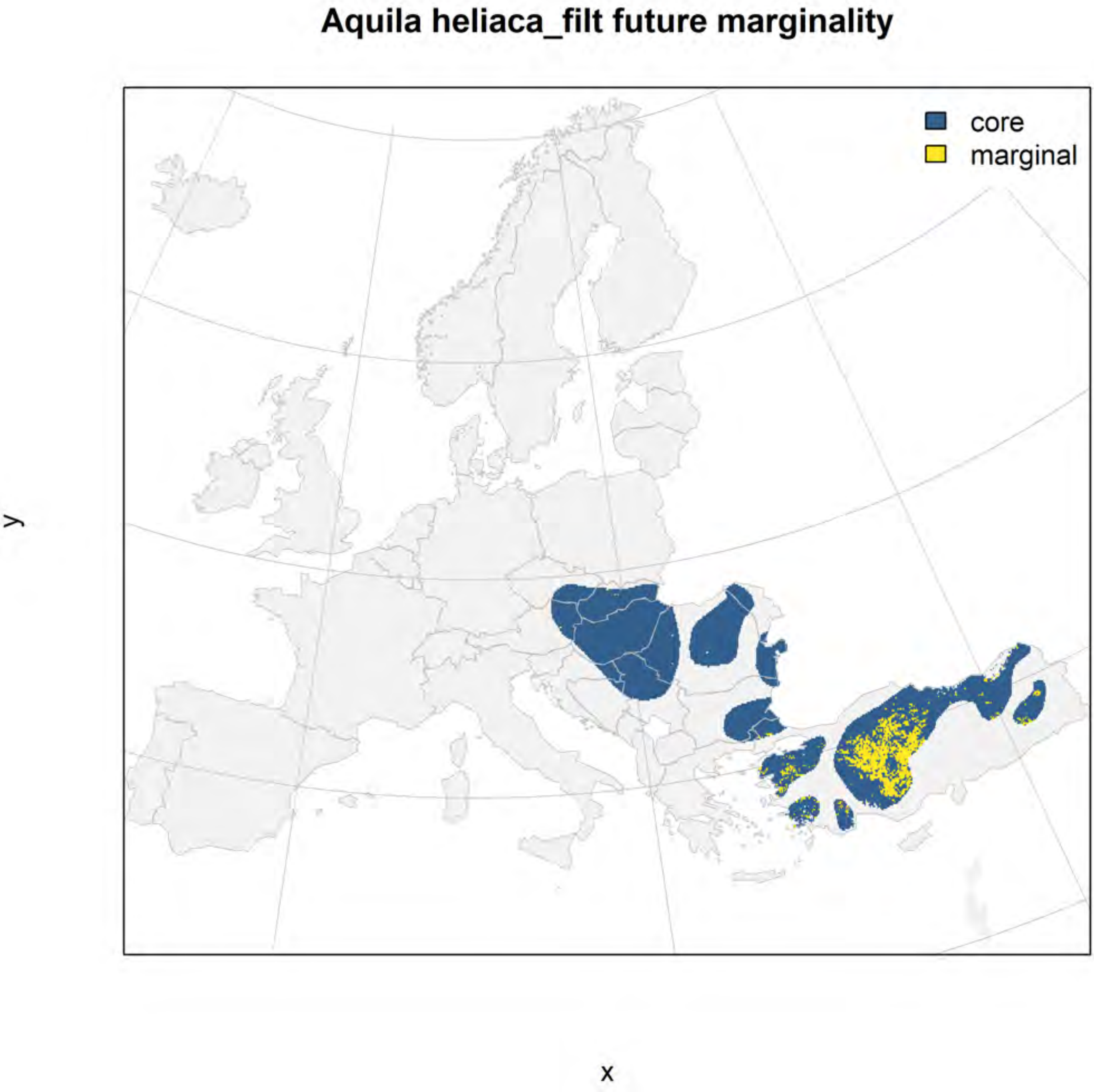
Aquila heliaca_filt future marginality
core
marginal
y
x

### Neophron percnopterus marginality

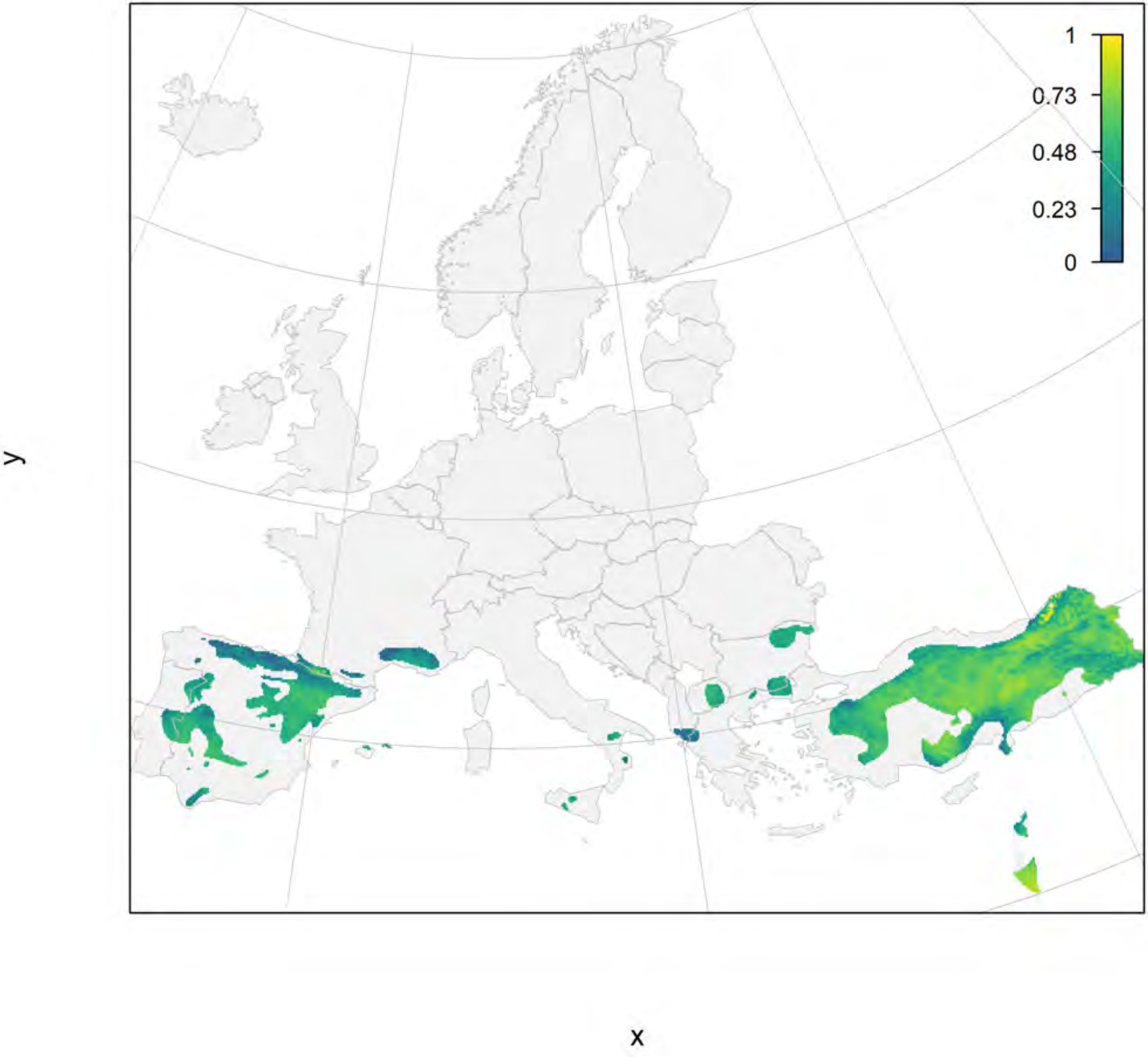

Neophron percnopterus_filt current marginality

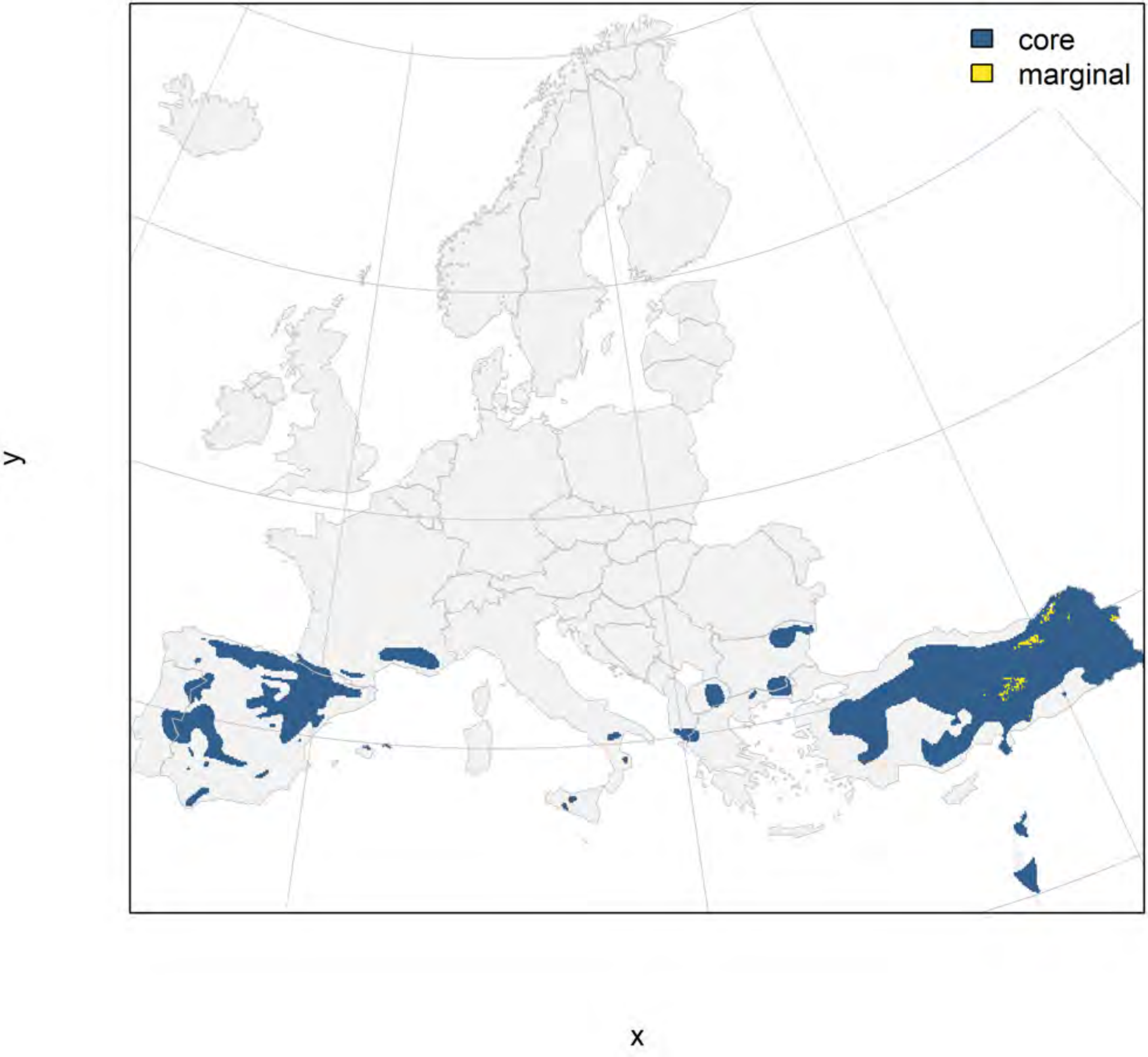

### Neophron percnopterus_filt future marginality

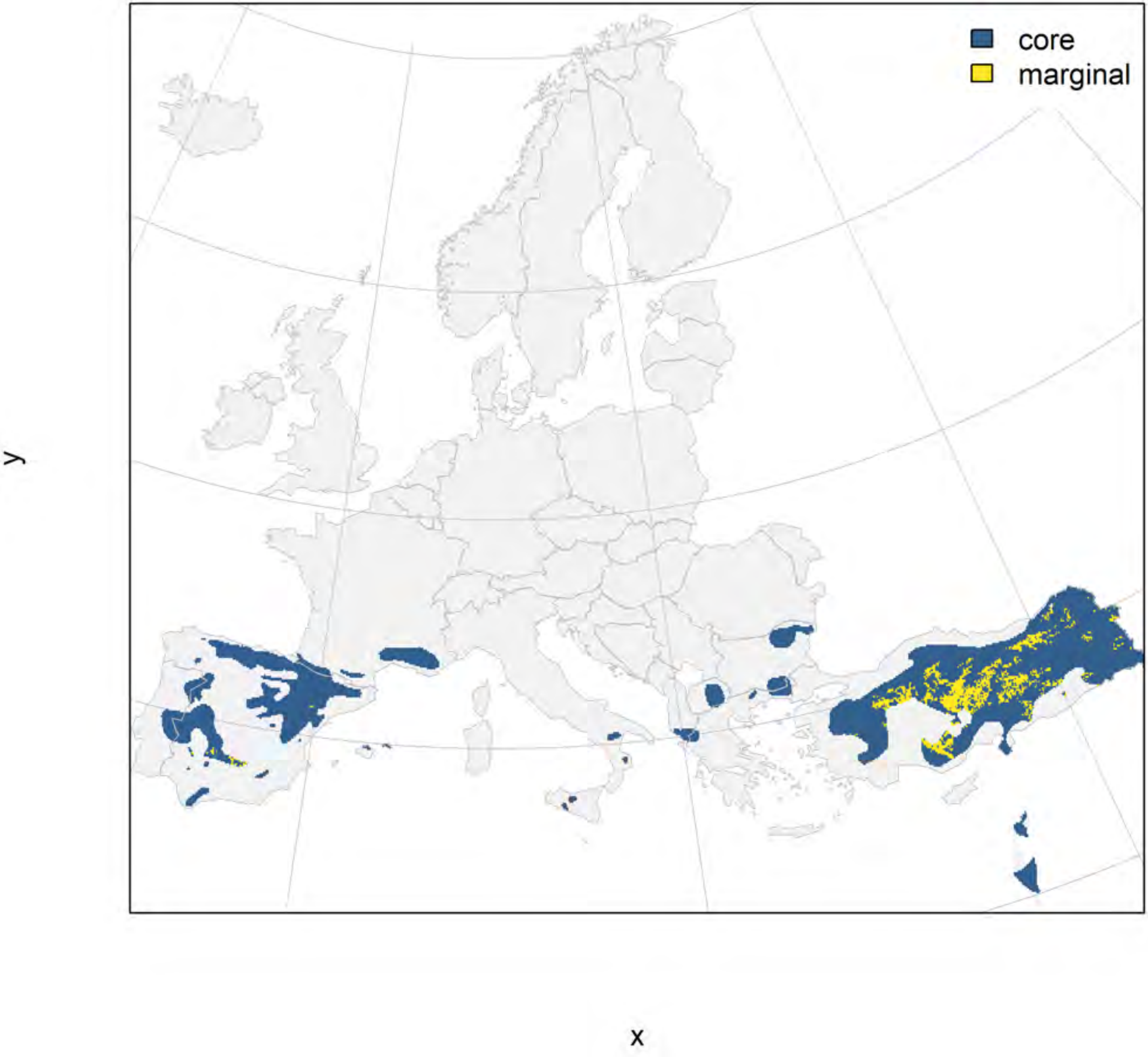

### Aquila adalberti marginality

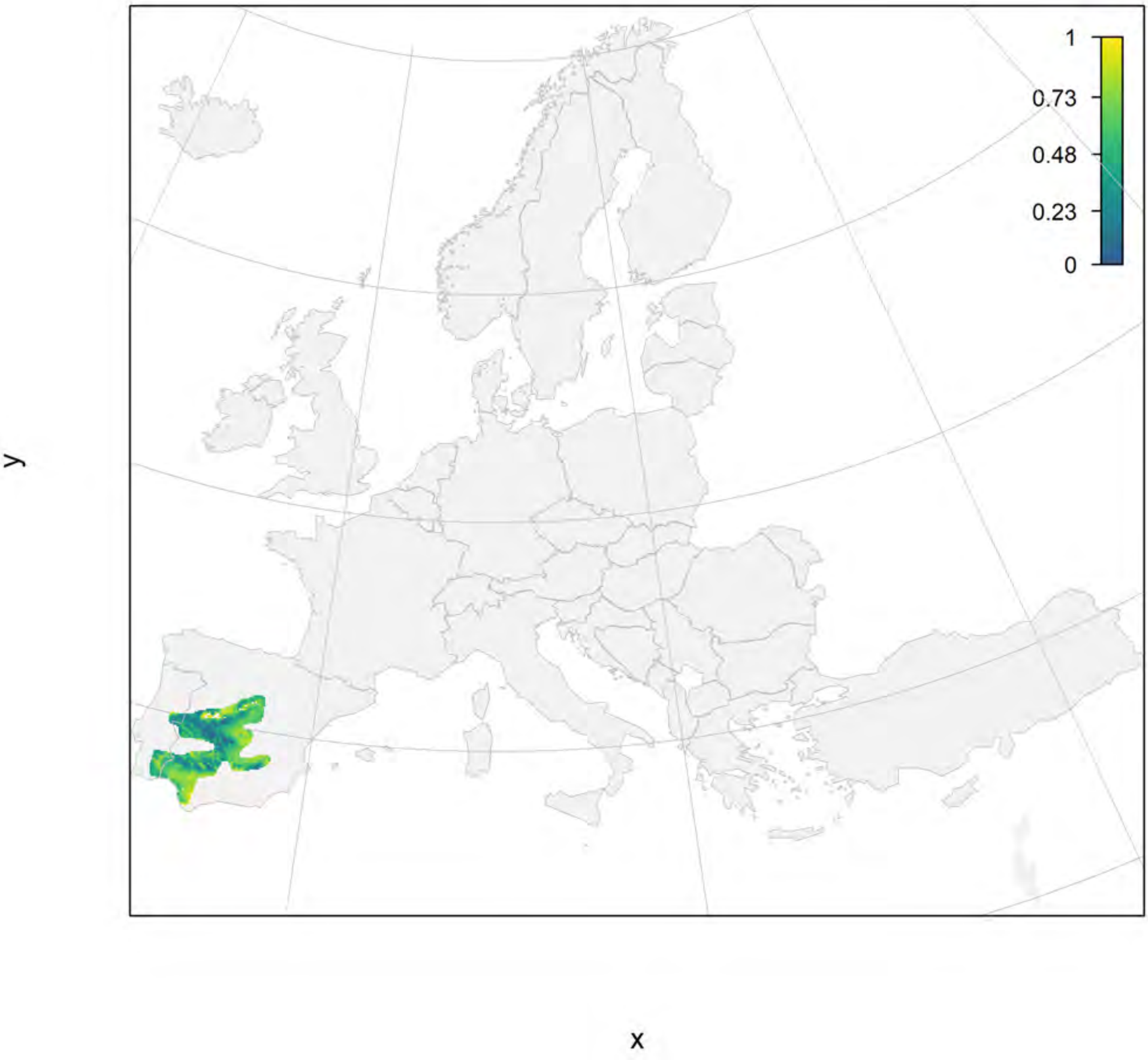

#### Aquila adalberti_filt current marginality

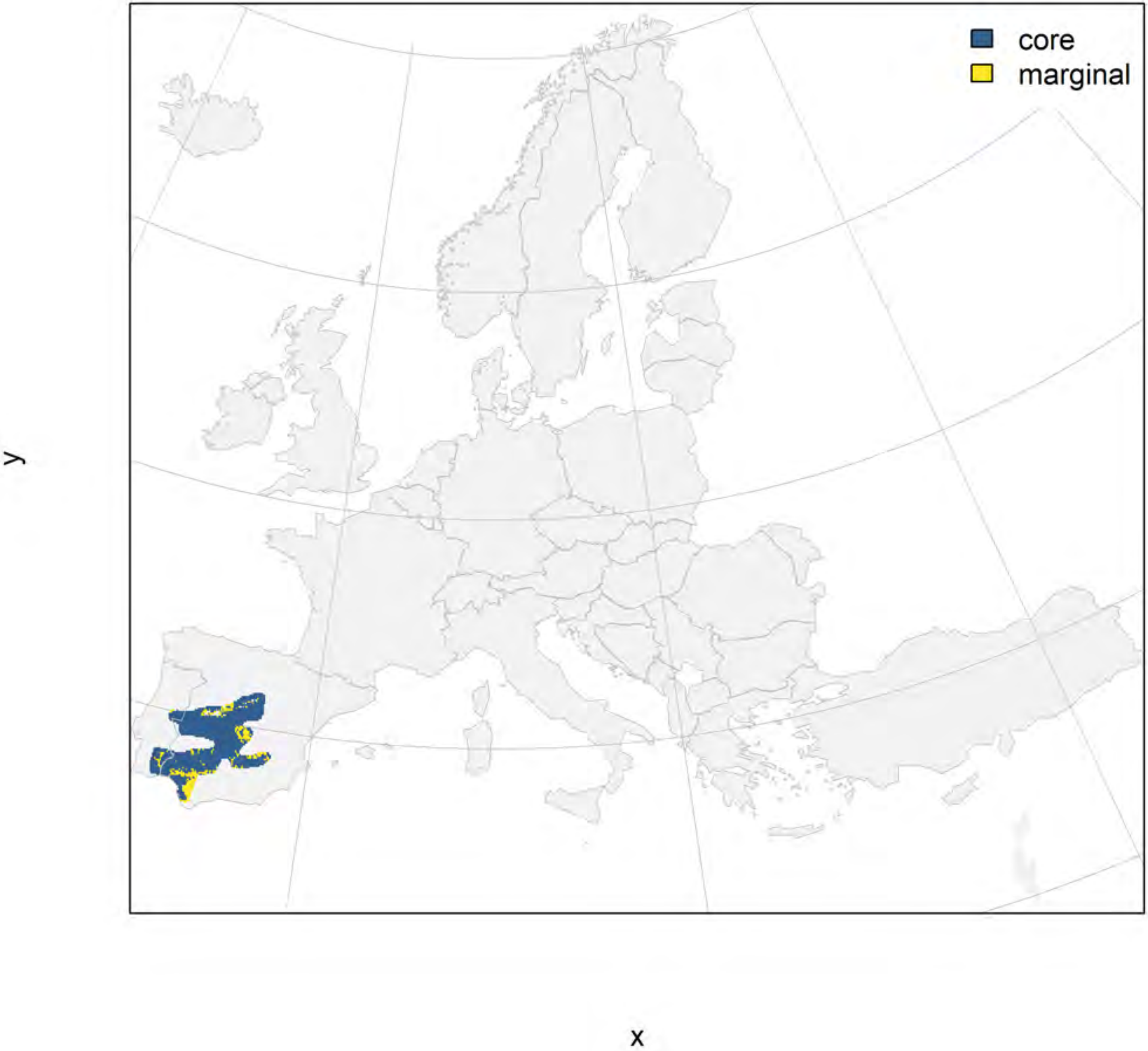

#### Aquila adalberti_filt future marginality

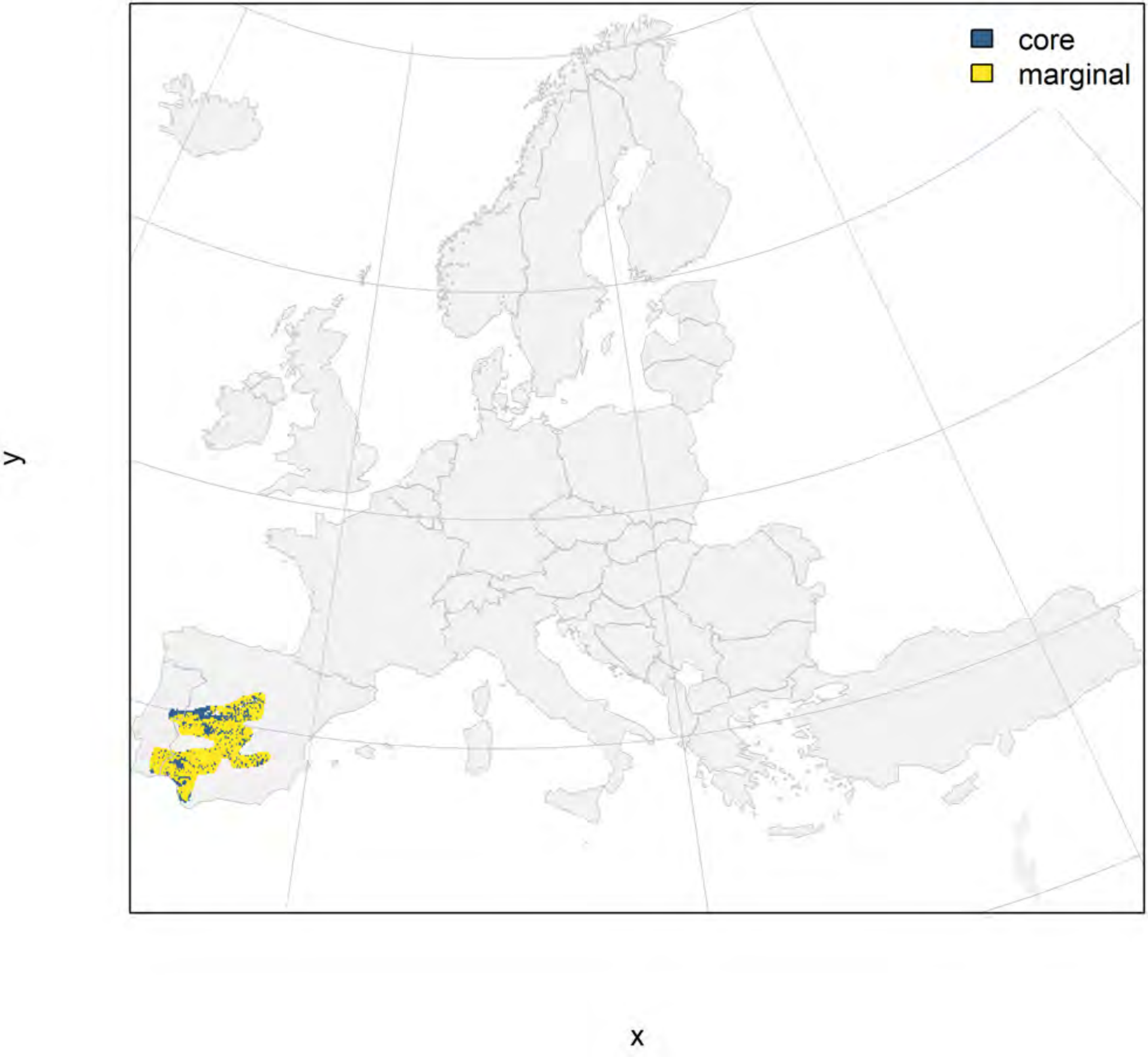

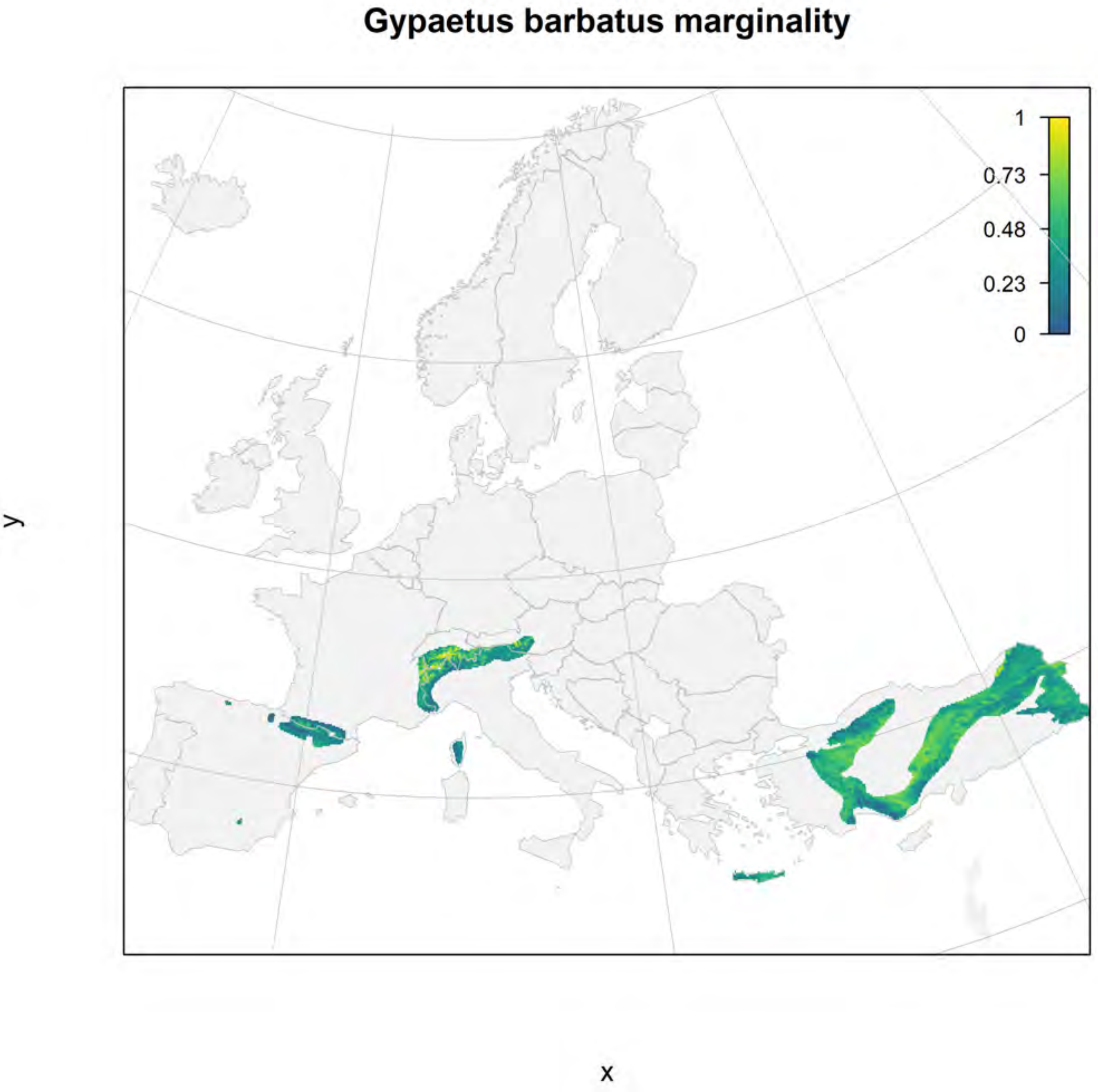

Gypaetus barbatus marginality
1
0.73
0.48
0.23
0
y
x

#### Gypaetus barbatus_filt current marginality

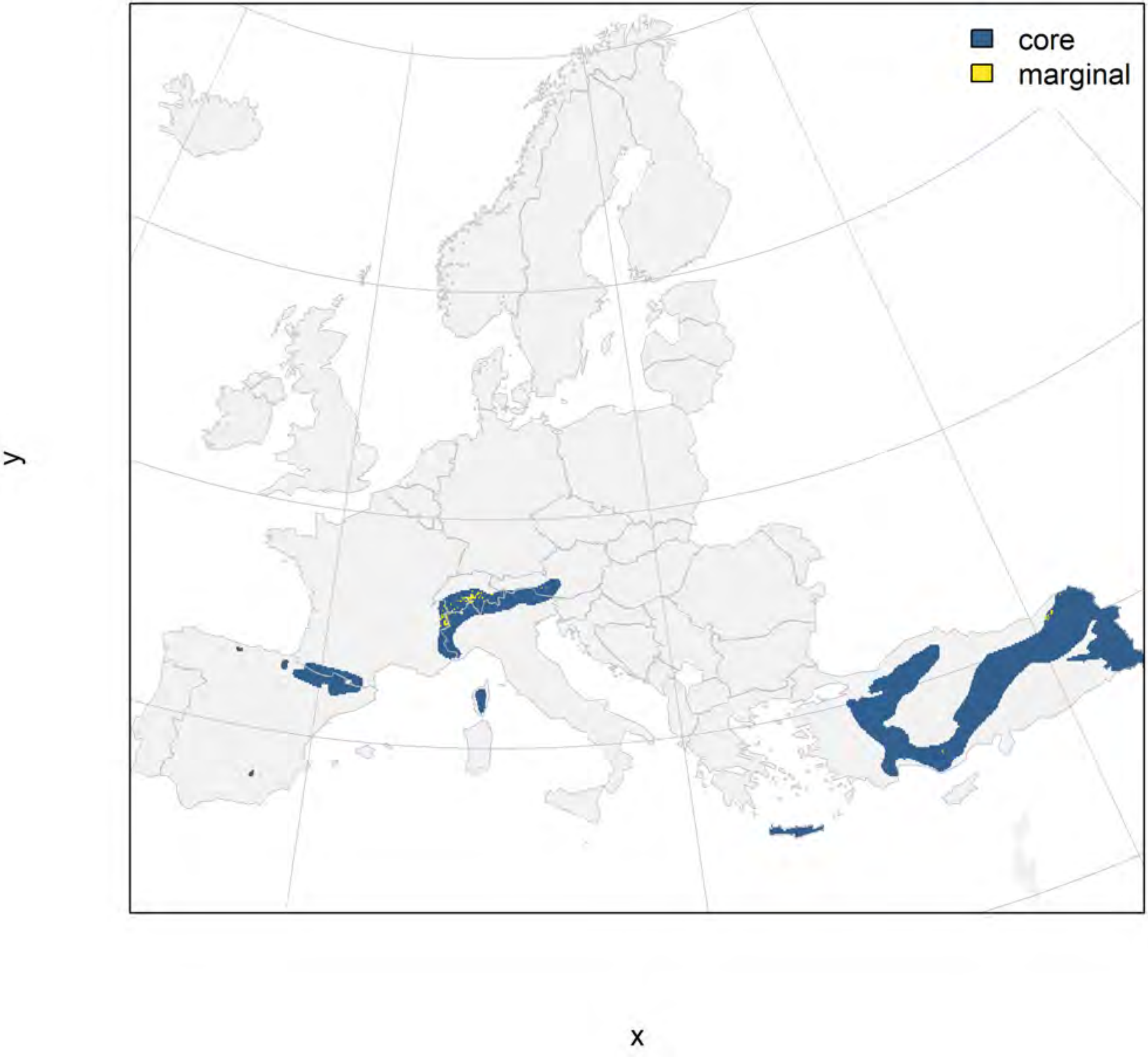

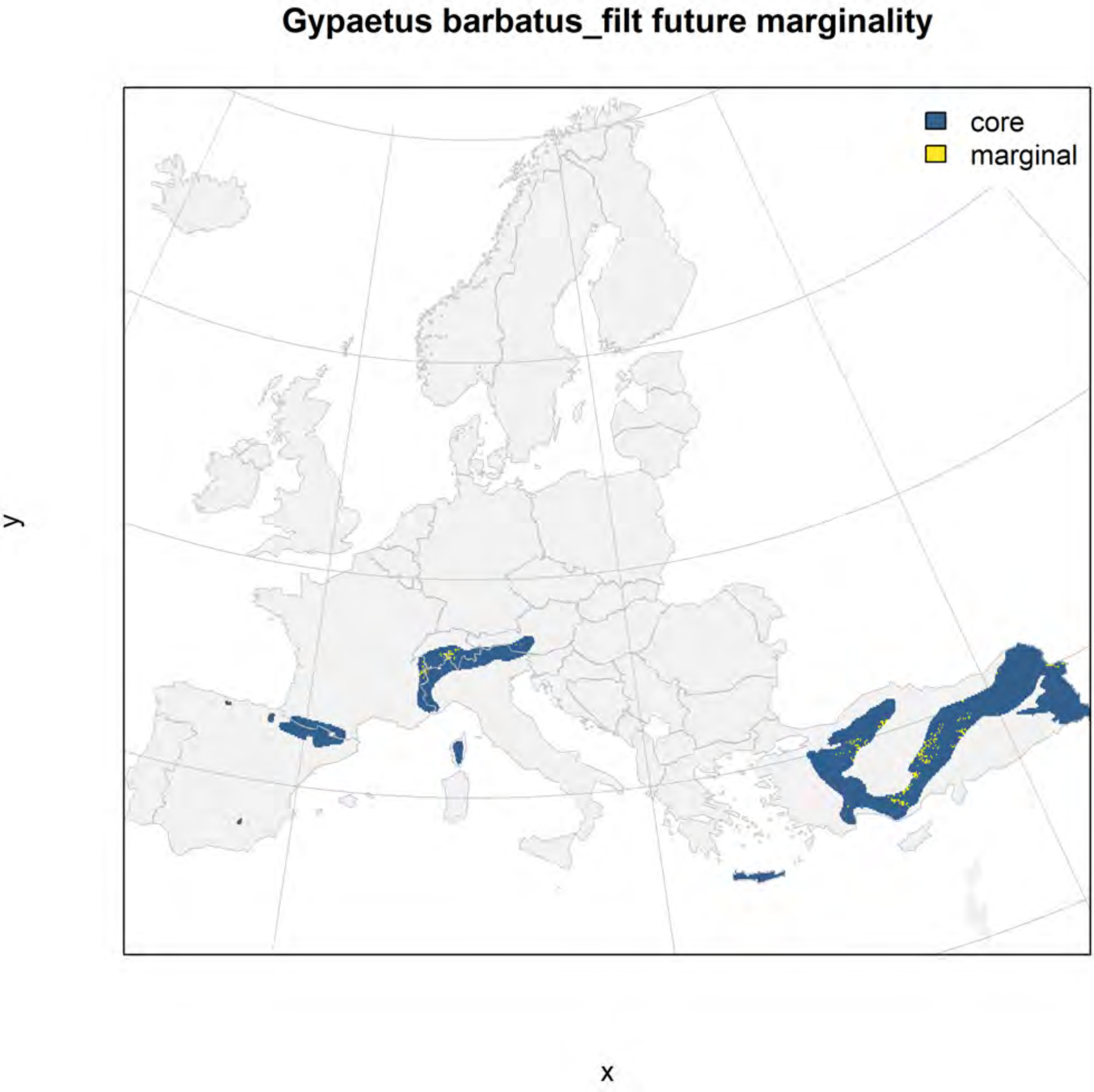
Gypaetus barbatus_filt future marginality
core
marginal
y
x
