## Appendix S5 for "Conserving genetic diversity during climate change: Niche marginality and discrepant monitoring capacity in Europe"

Appendix S5. Maps of marginality of selected species of large European carnivoran species. The climate marginality of each species is depicted in a series of three figures: raw marginality in which a value of 1 indicates the 99$^{th}$ percentile of the species niche space within its current range (filtered for appropriate habitat with a CORINE land cover layer), the current geographic distribution of areas within the 25% most marginal climatic niche conditions and the remaining core area, and a predicted future distribution of core and marginal areas within the current species range. Please see the Online Methods section for a description of marginality calculations.

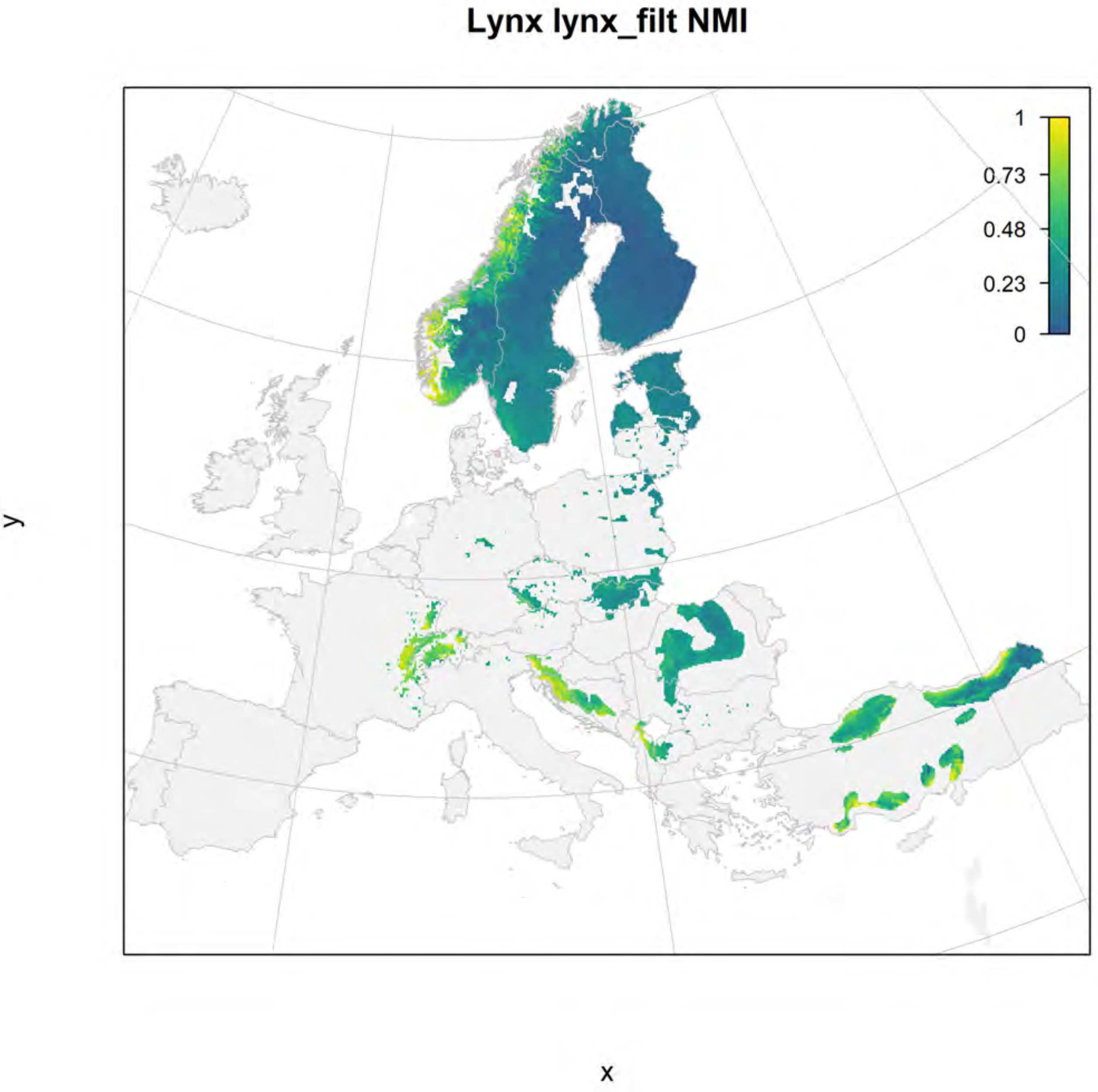
Lynx lynx_filt NMI
1
0.73
0.48
0.23
0
y
x

### Lynx lynx_filt current marginality

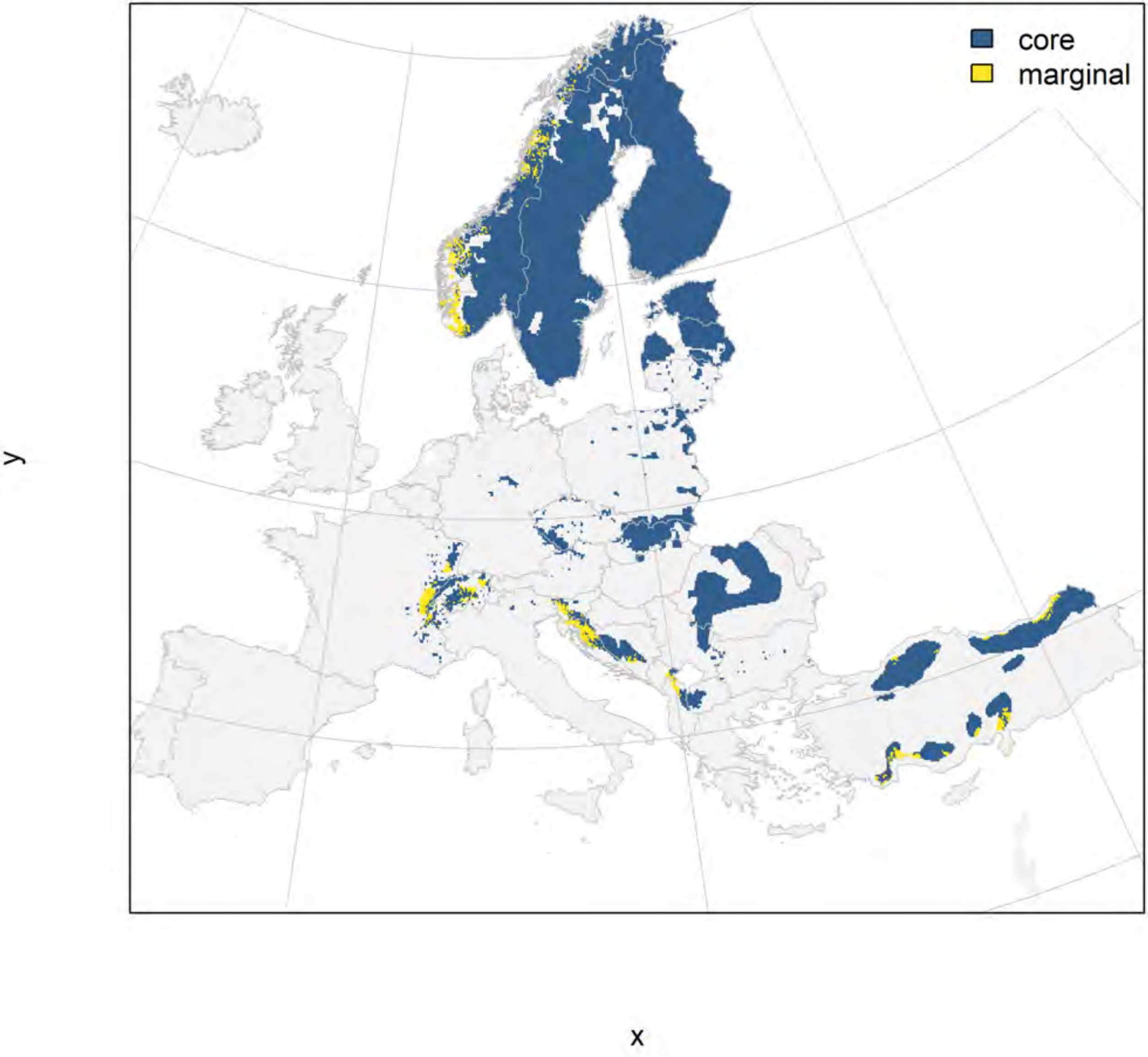

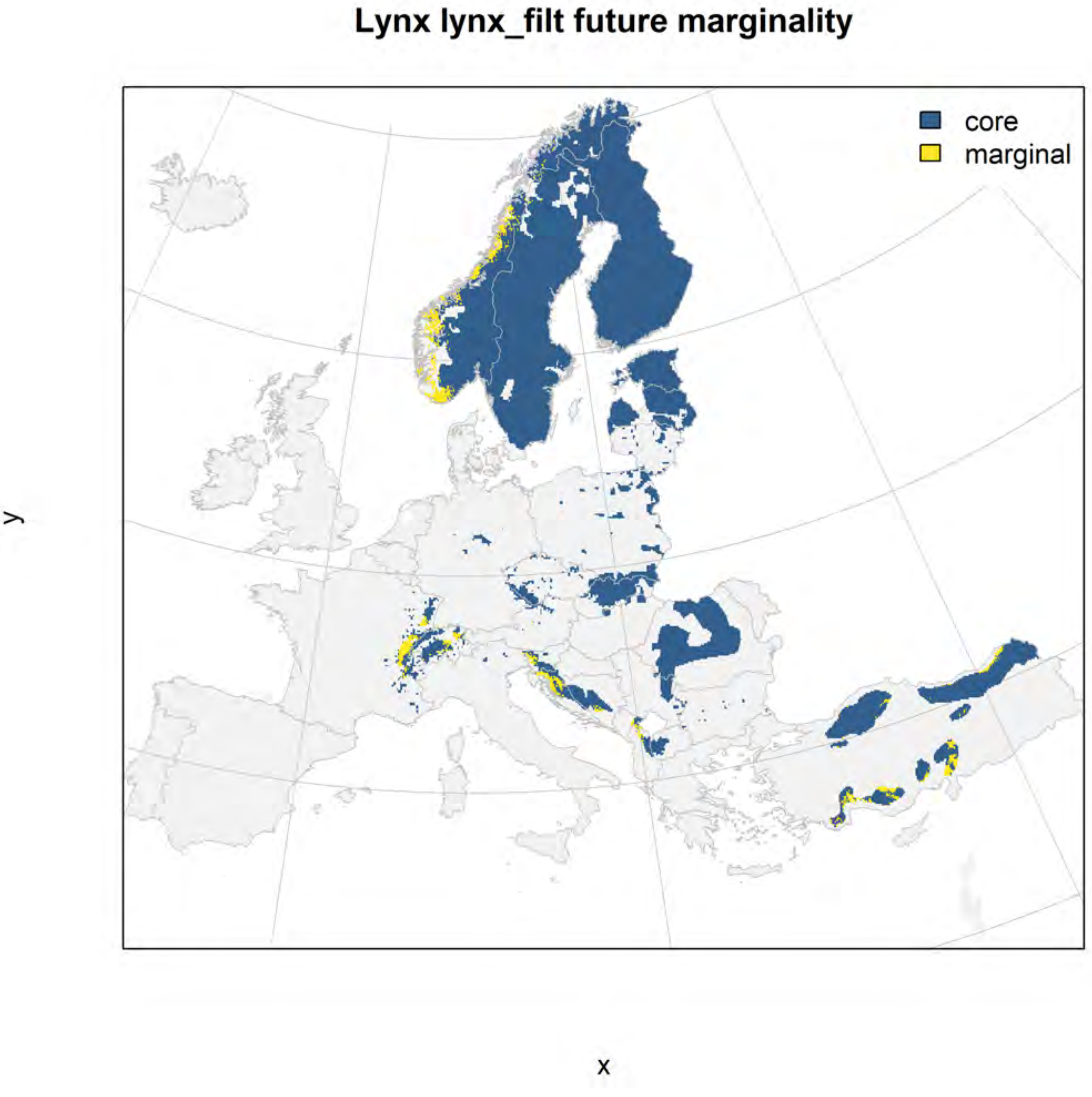
Lynx lynx_filt future marginality
core
marginal
y
x

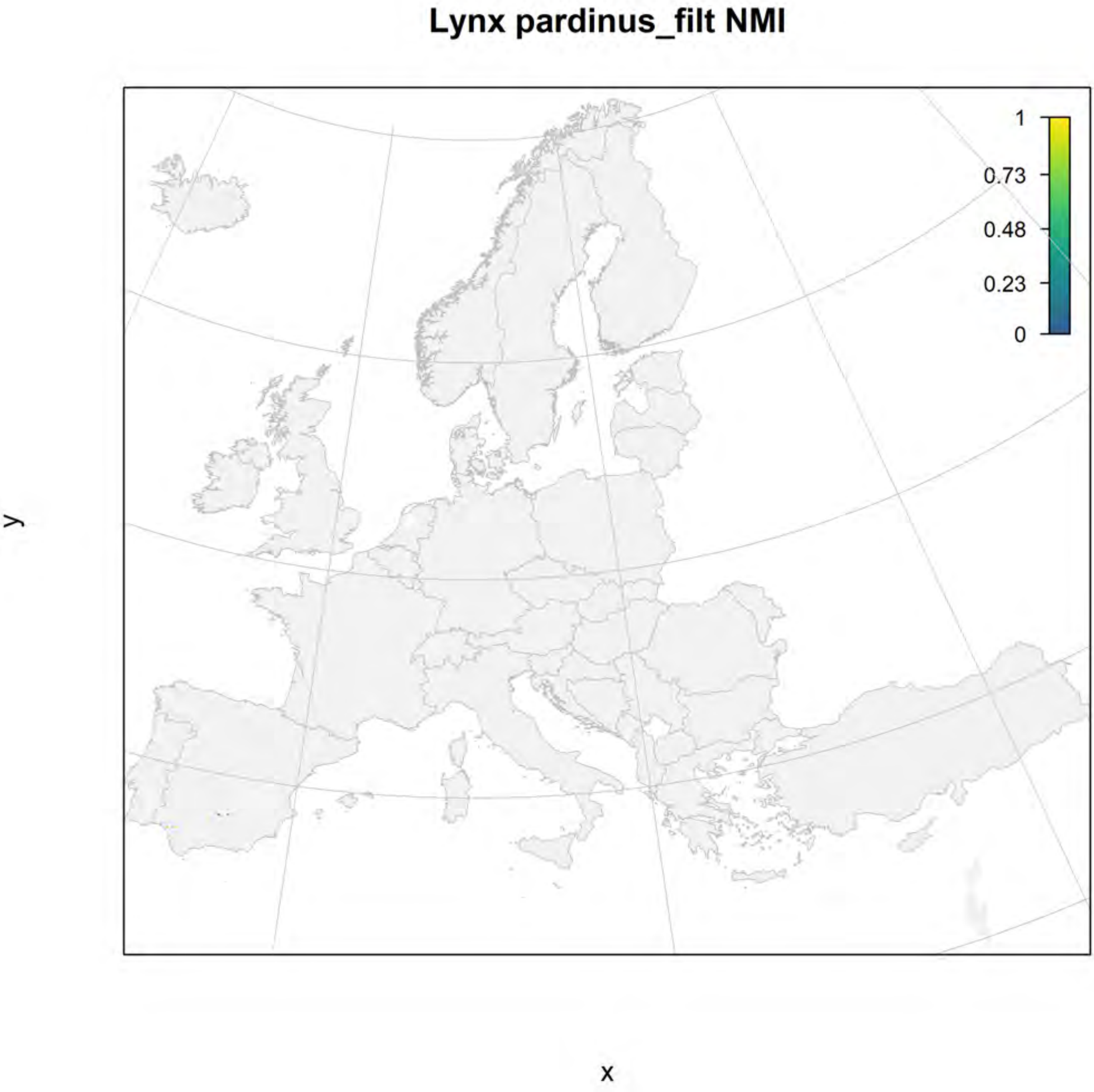

Lynx pardinus_filt NMI
1
0.73
0.48
0.23
0
y
x

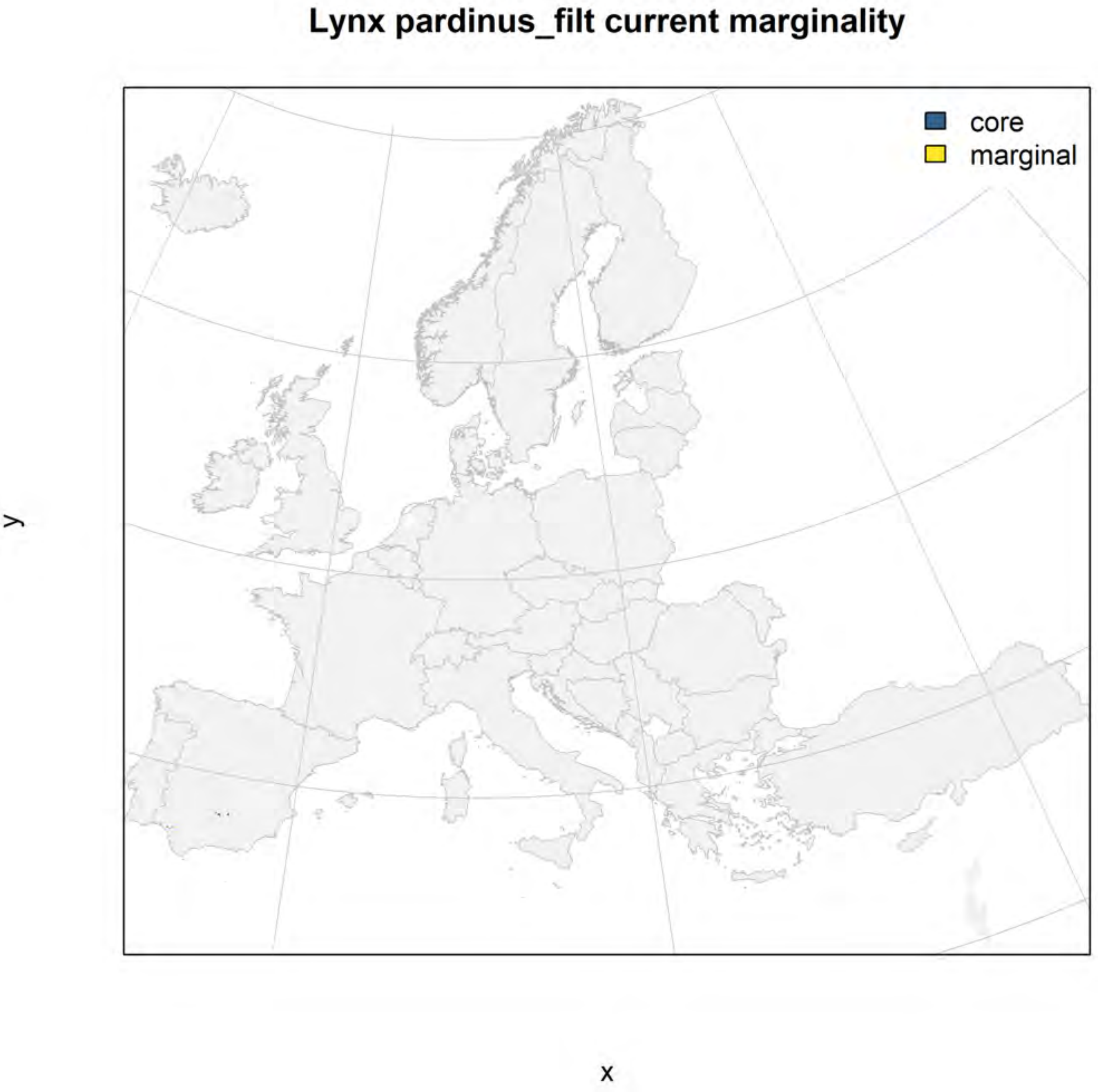

Lynx pardinus_filt current marginality
core
marginal
y
x

### Lynx pardinus_filt future marginality

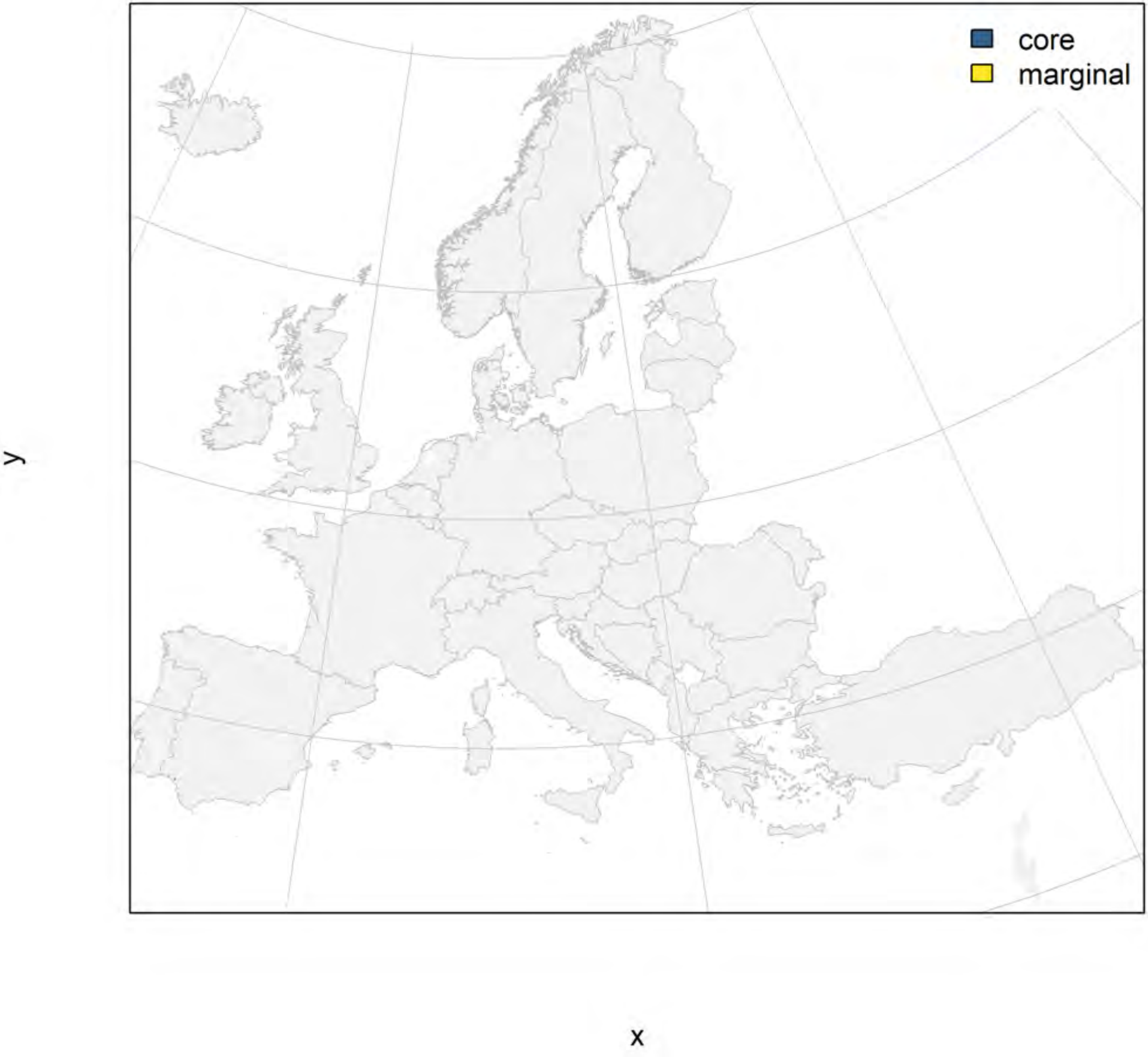

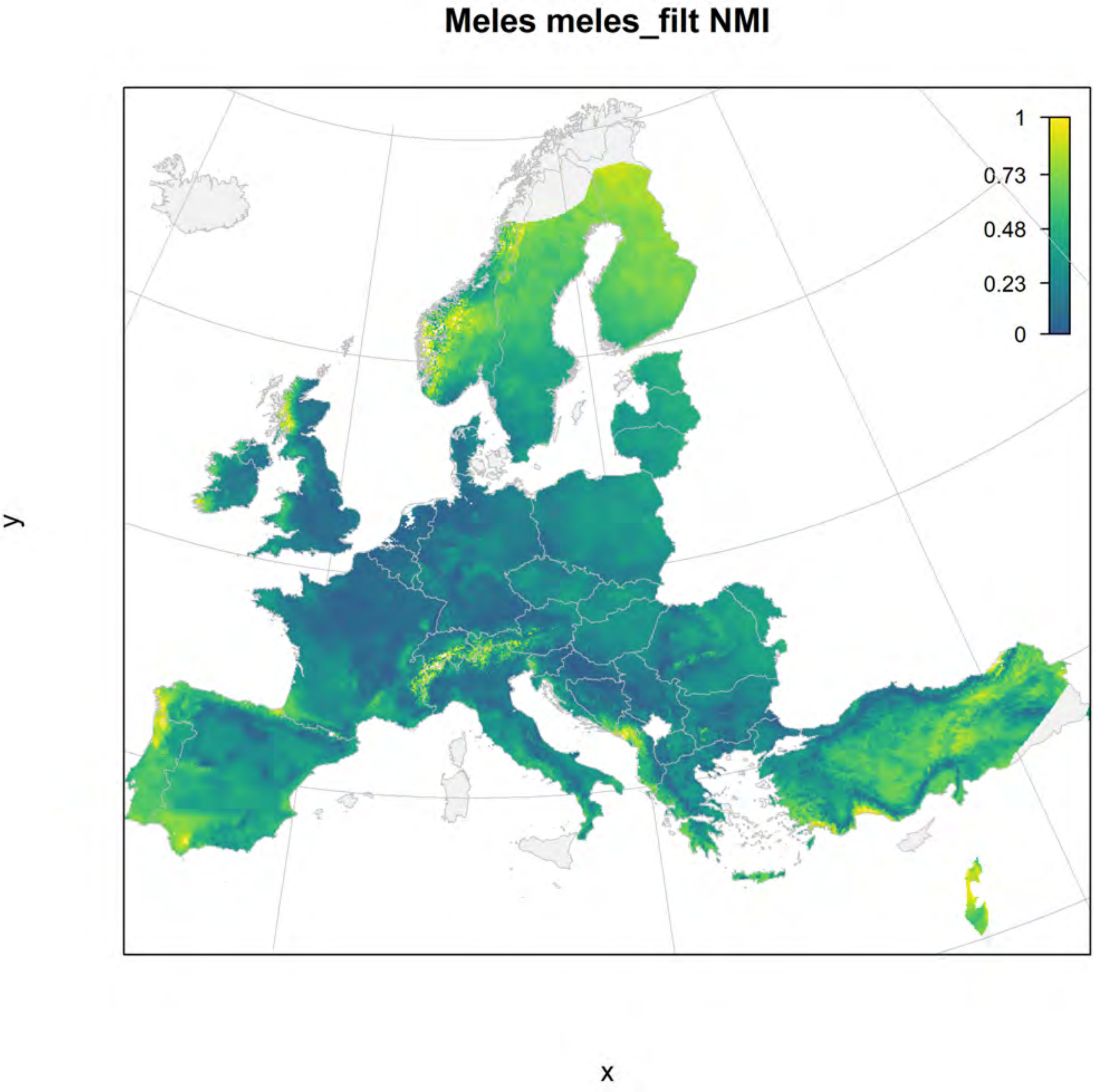
Meles meles_filt NMI
1
0.73
0.48
0.23
0
y
x

### Meles meles_filt current marginality

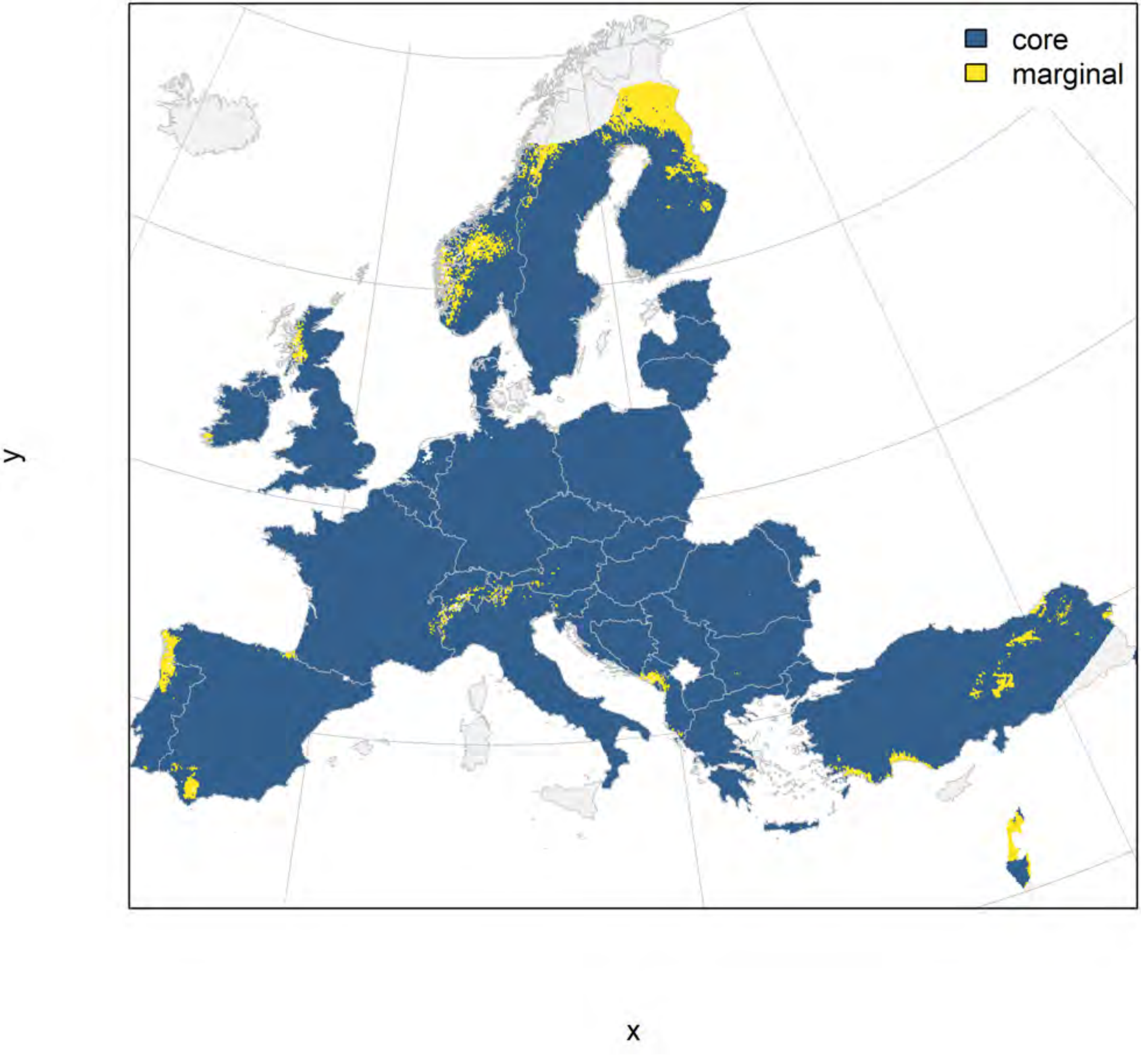

### Meles meles_filt future marginality

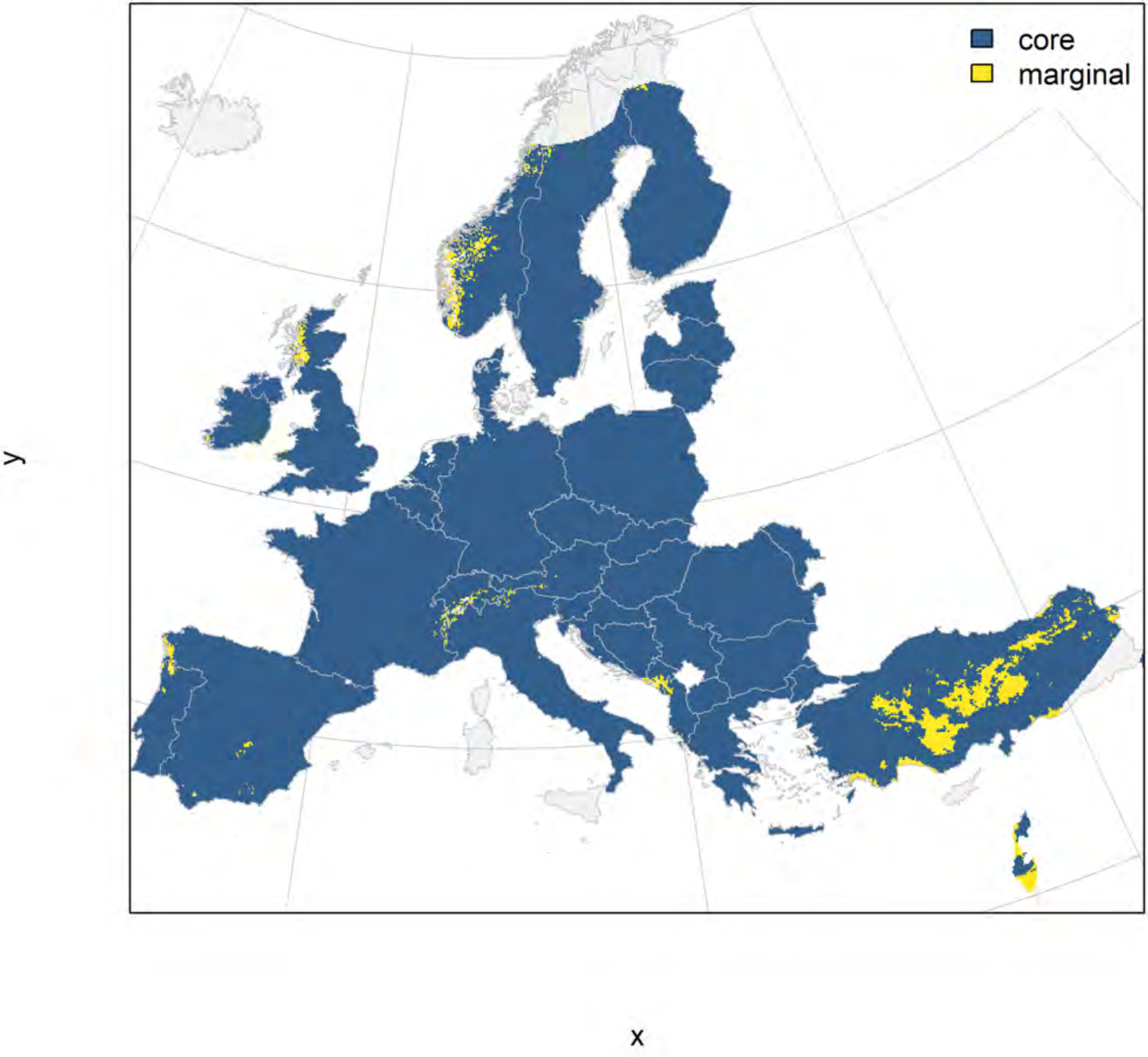

Lutra lutra_filt NMI

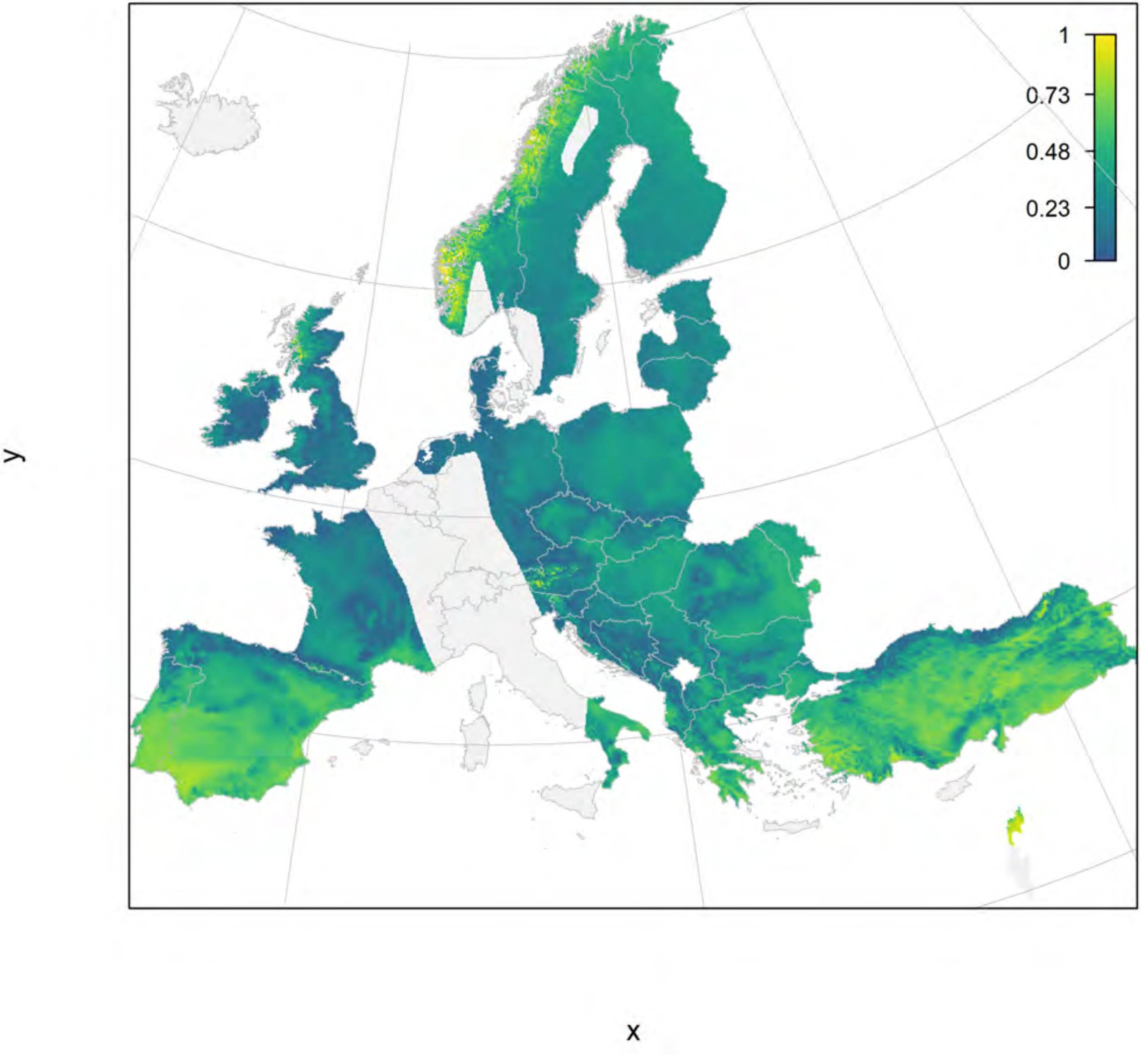

### Lutra lutra_filt current marginality

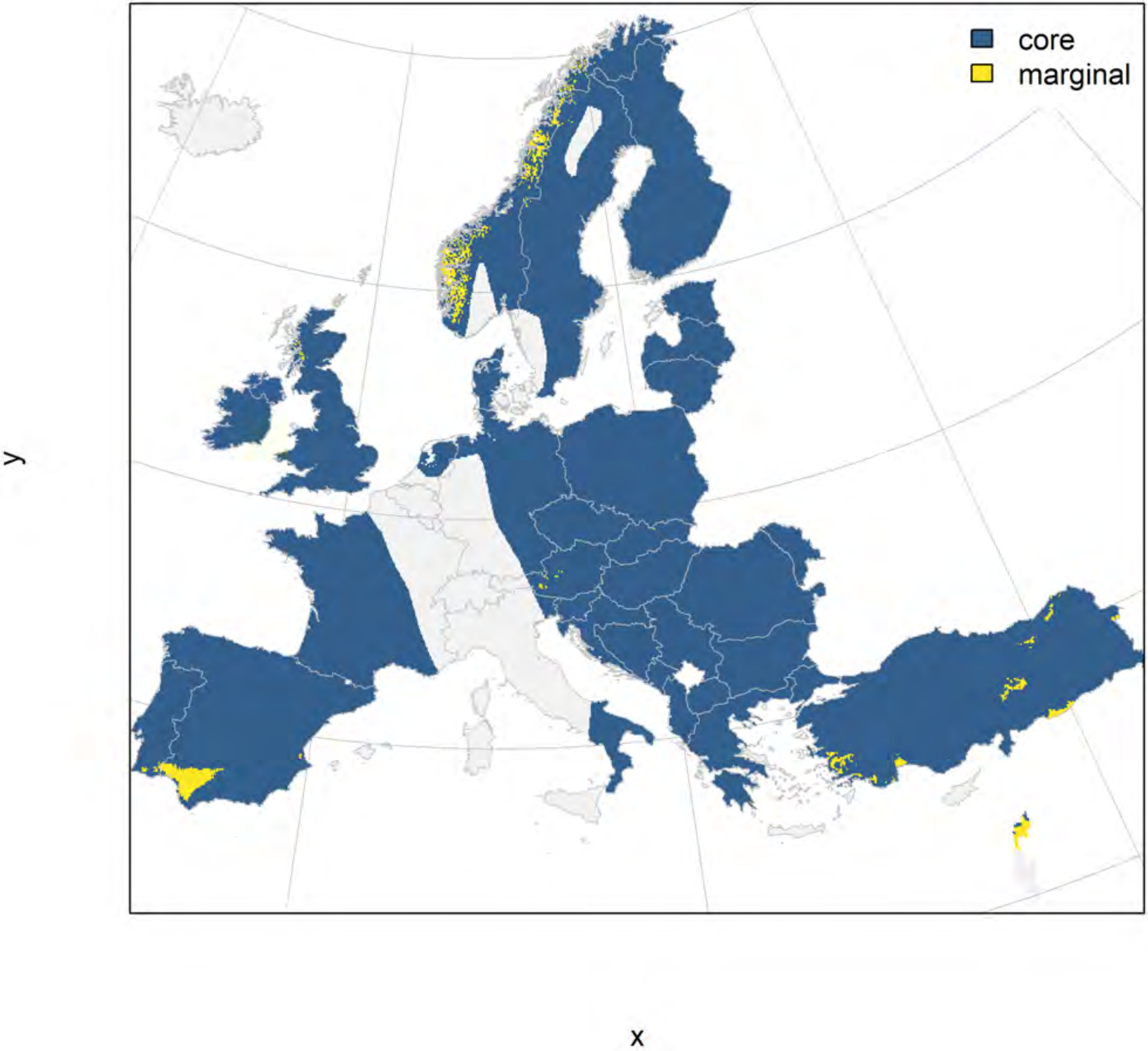

**Lutra lutra_filt future marginality**

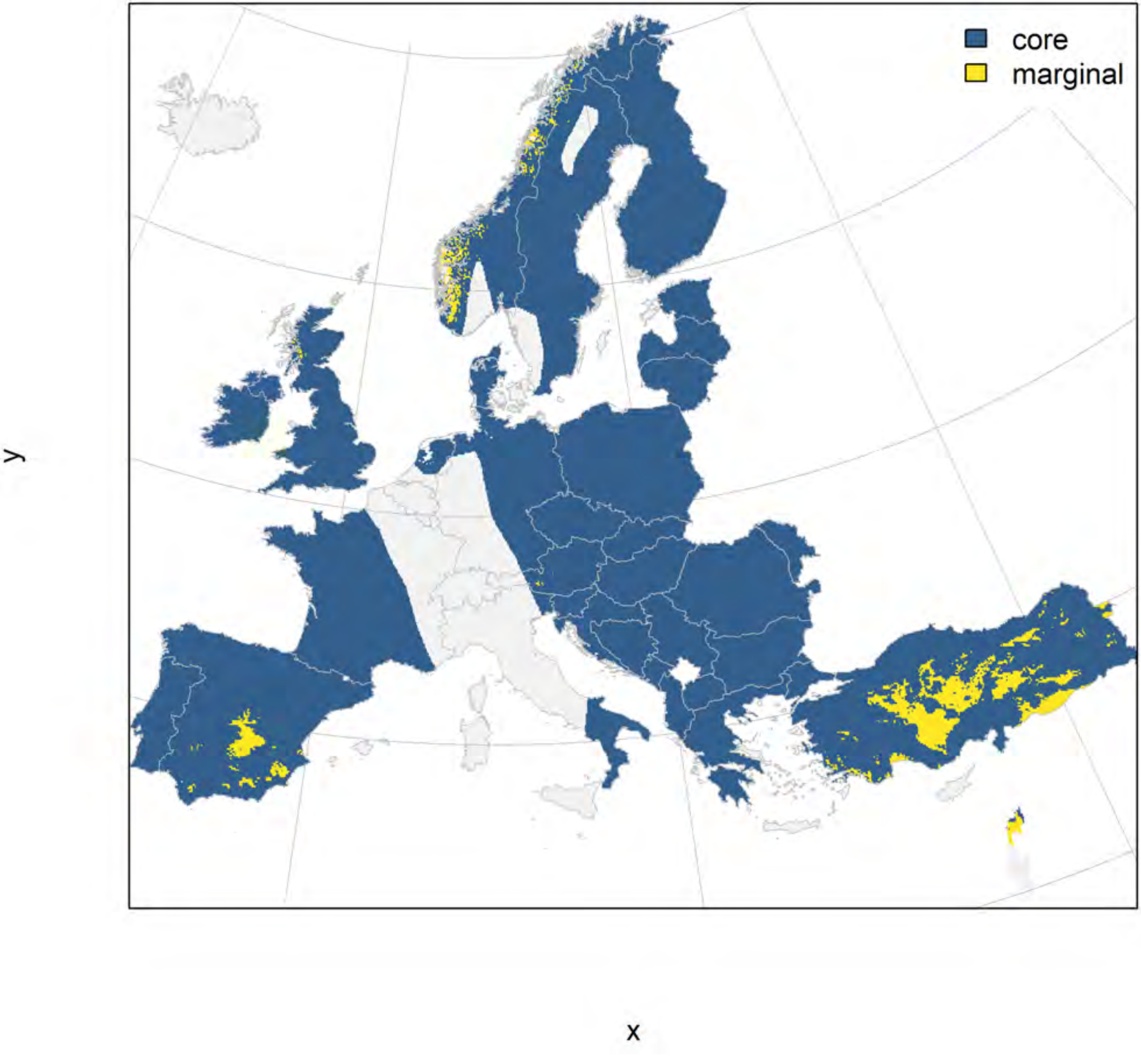

Canis aureus_filt NMI

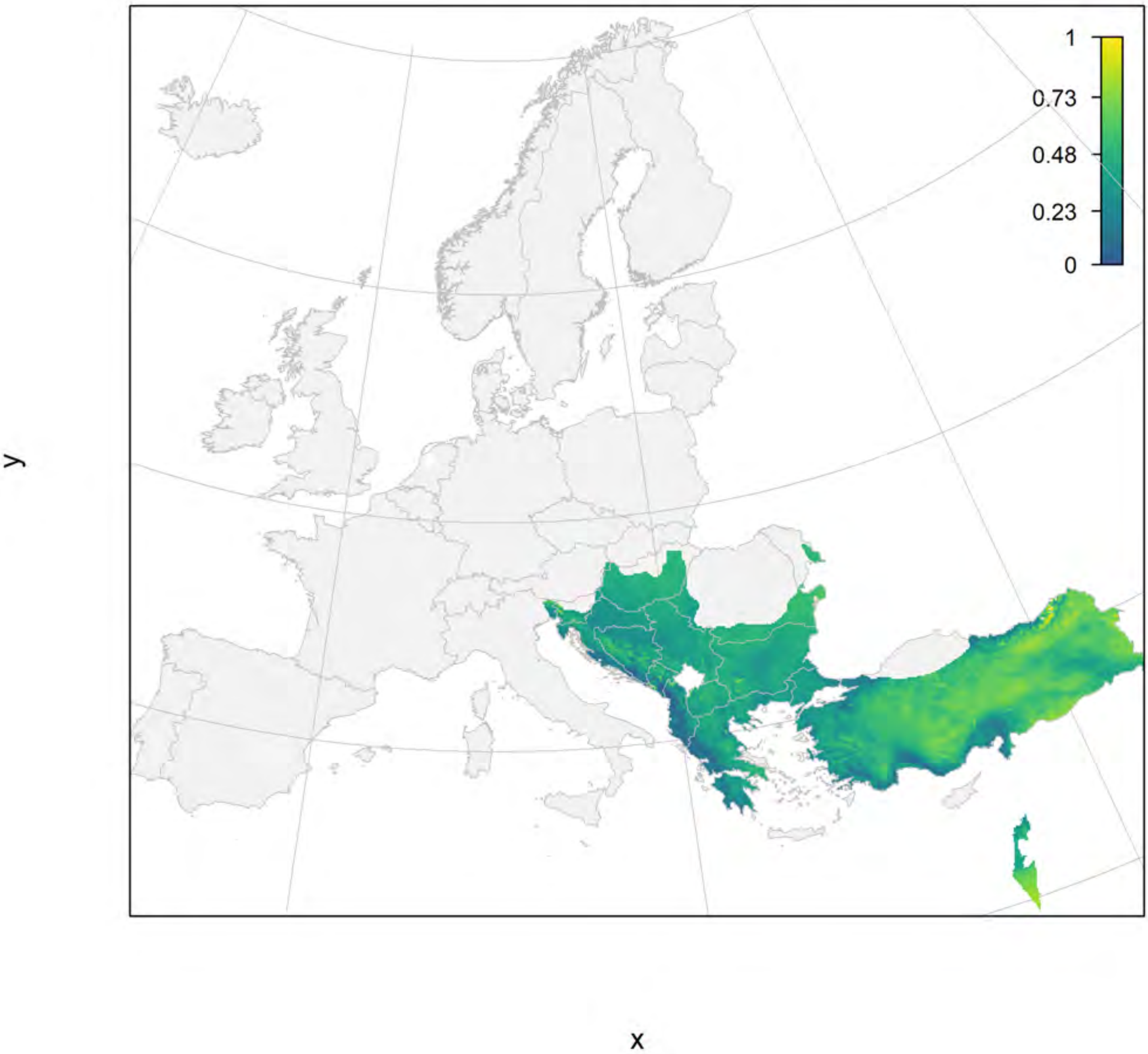

#### Canis aureus_filt current marginality

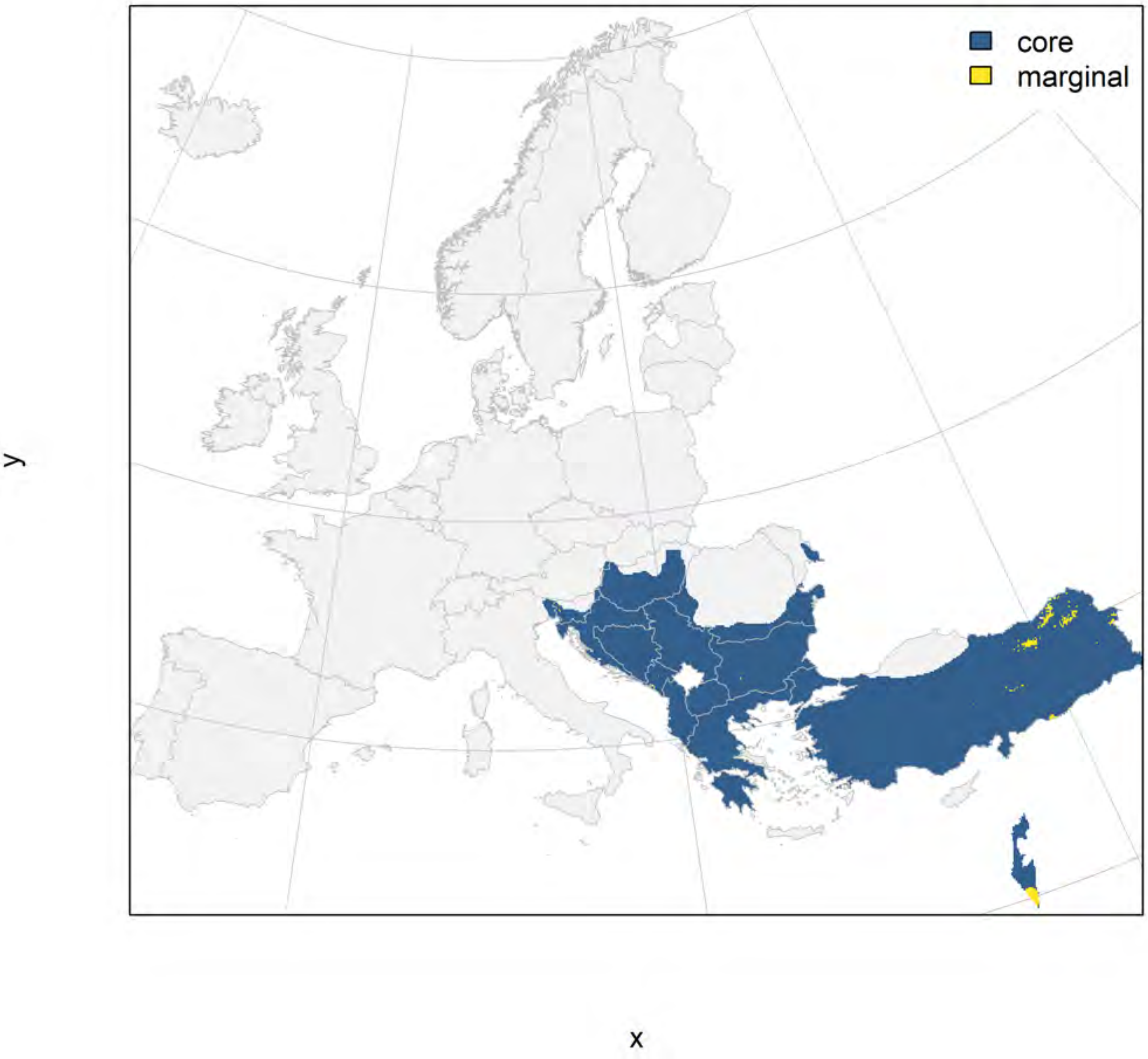

### Canis aureus_filt future marginality

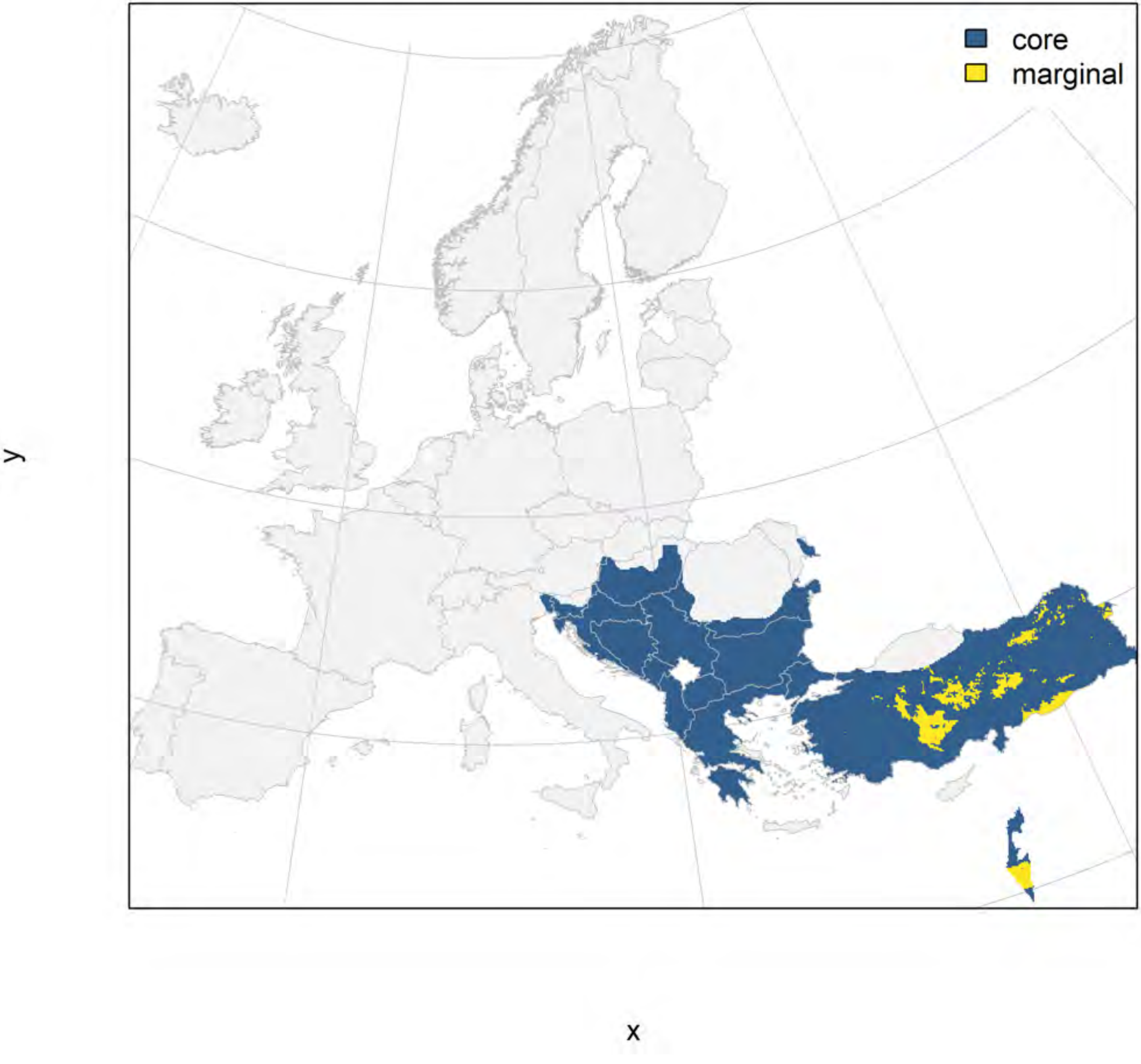

### Canis lupus_filt NMI

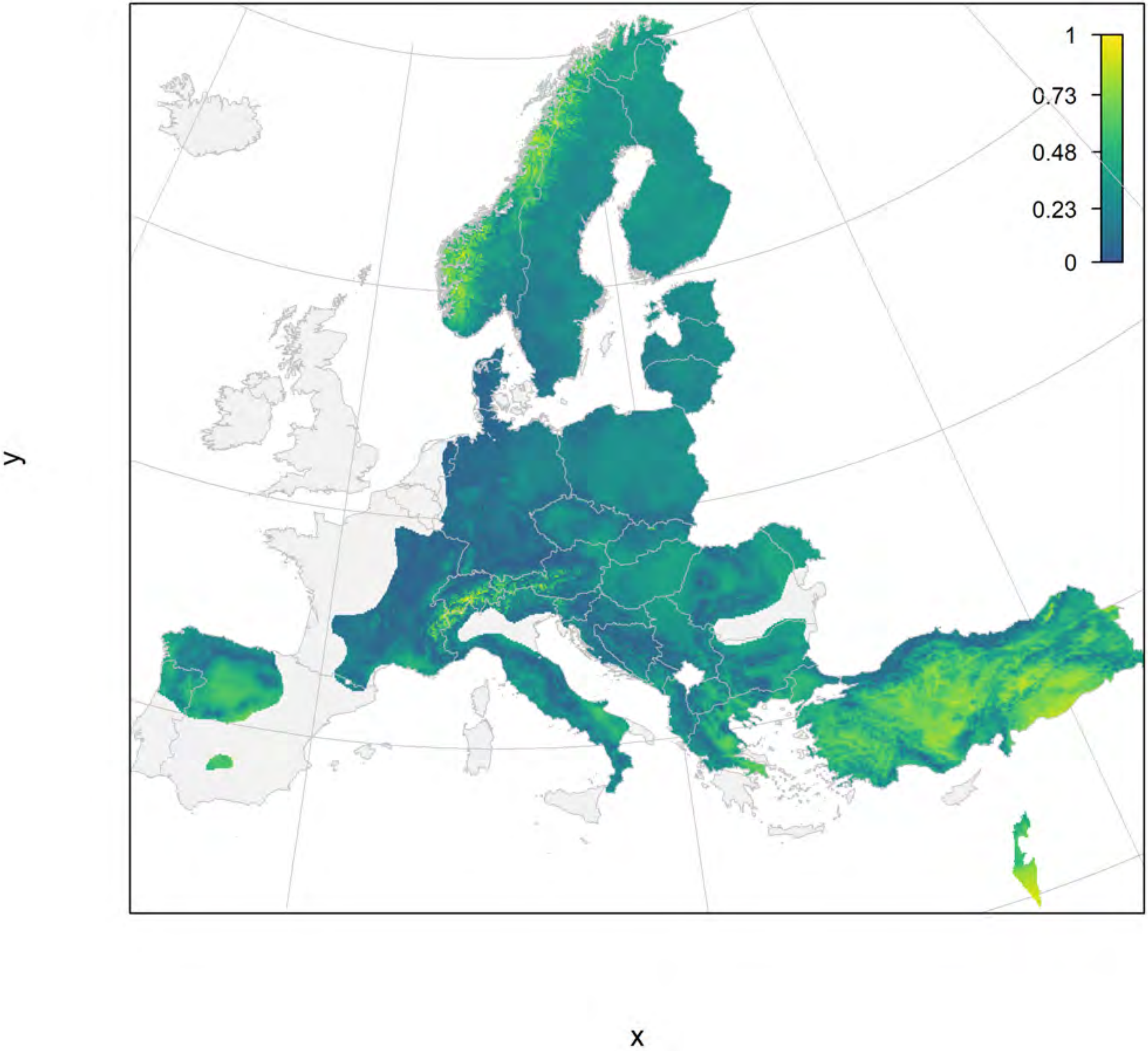

#### Canis lupus_filt current marginality

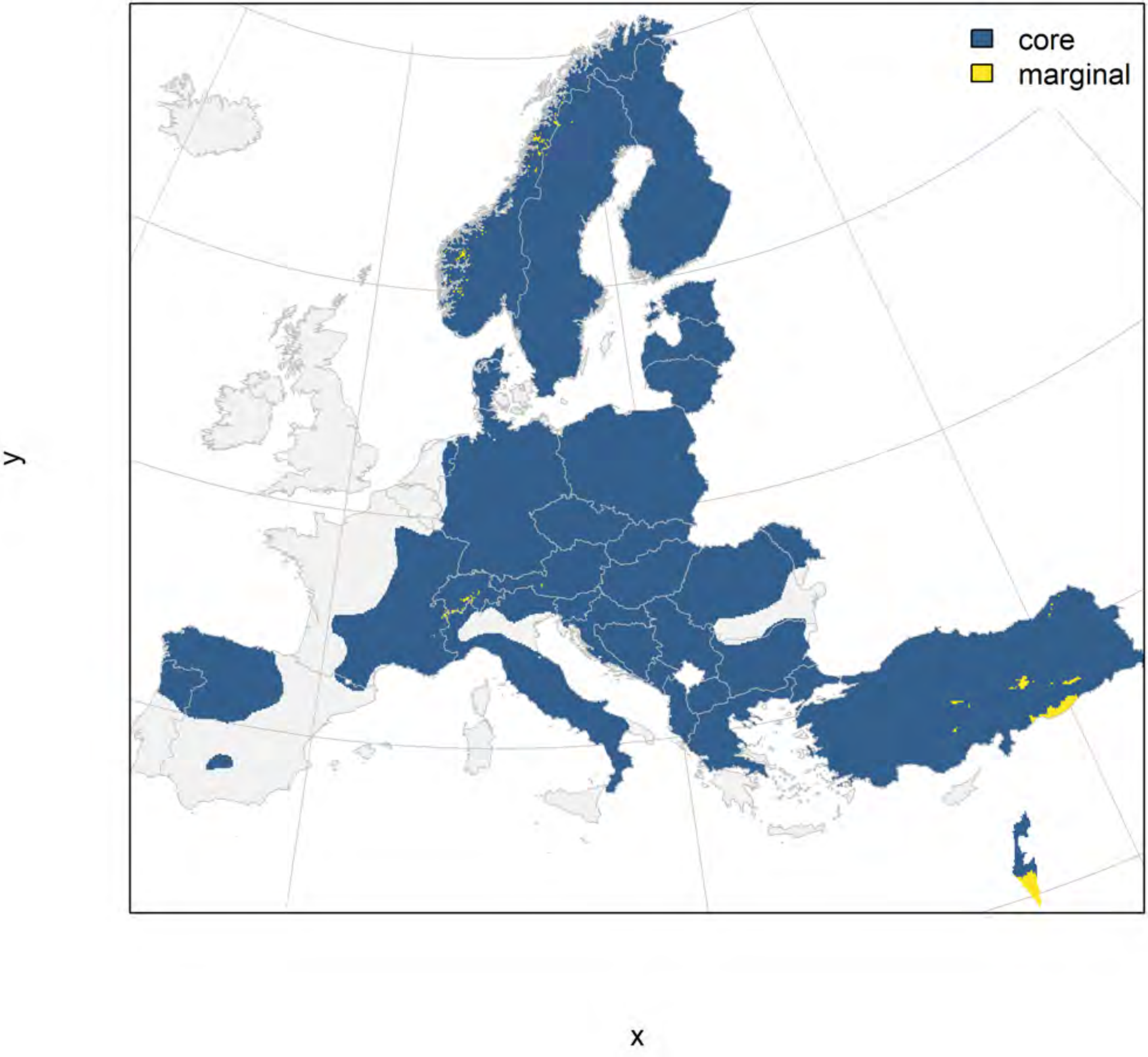

### Canis lupus_filt future marginality

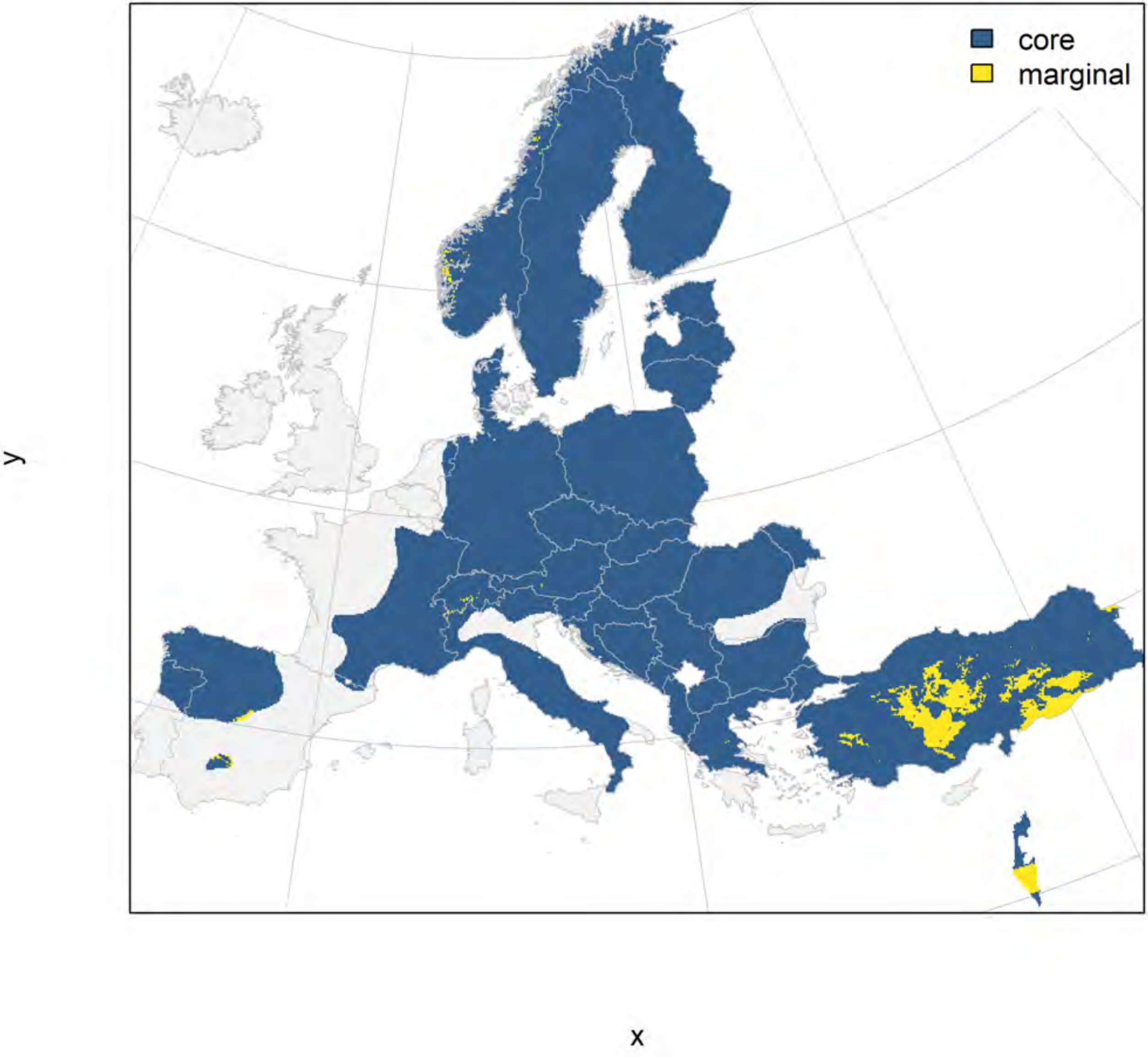

### Gulo gulo_filt NMI

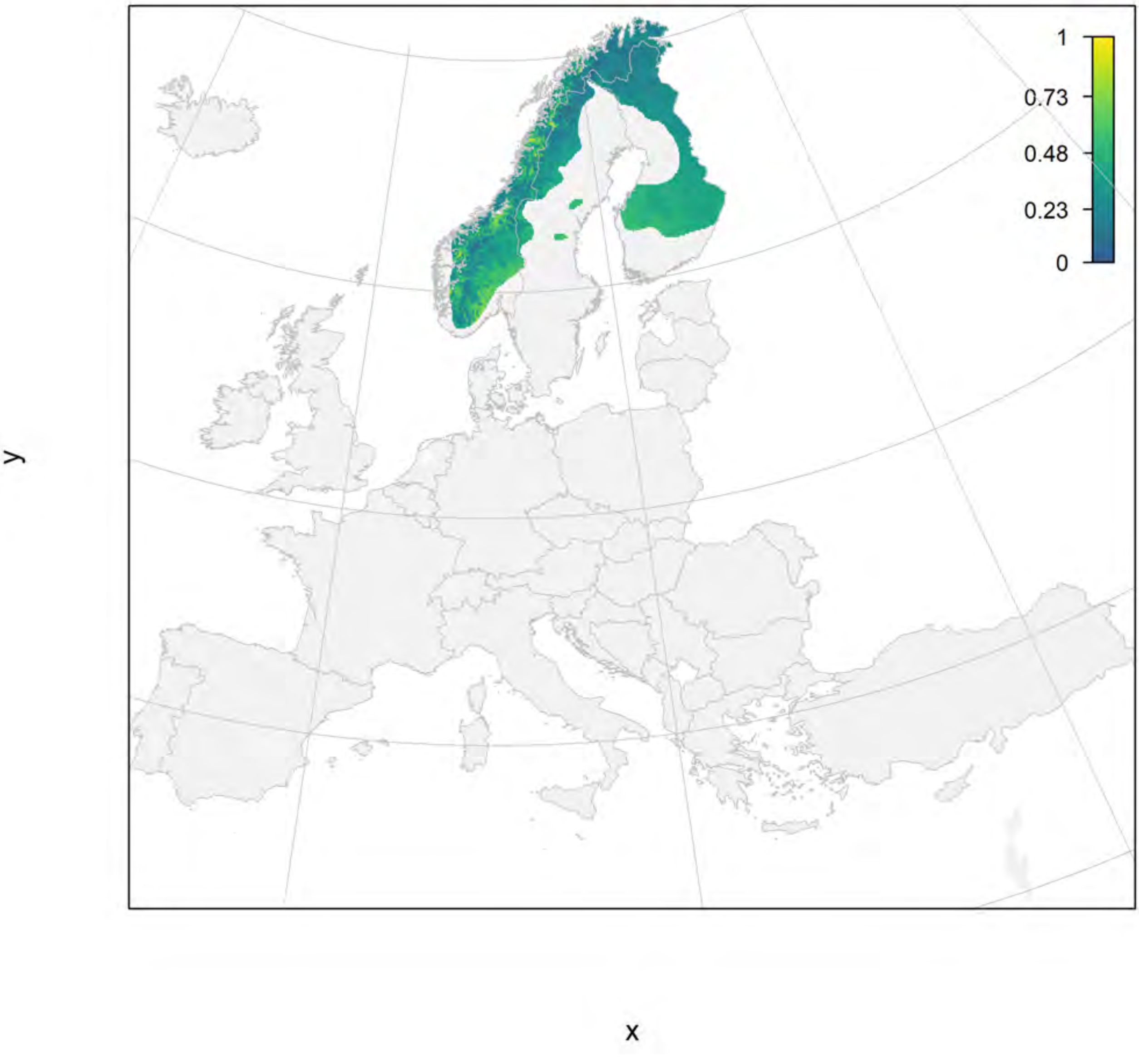

#### Gulo gulo_filt current marginality

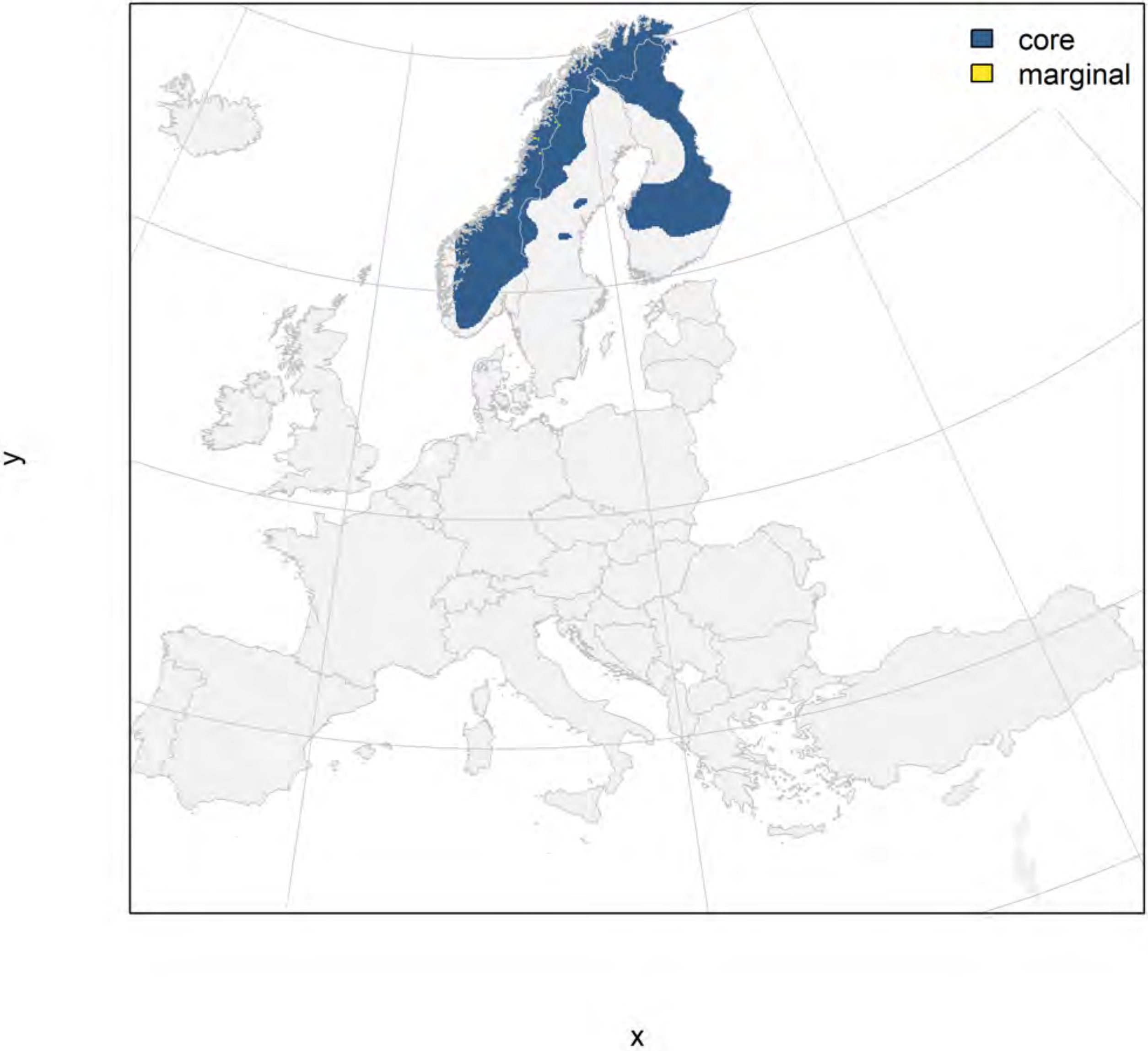

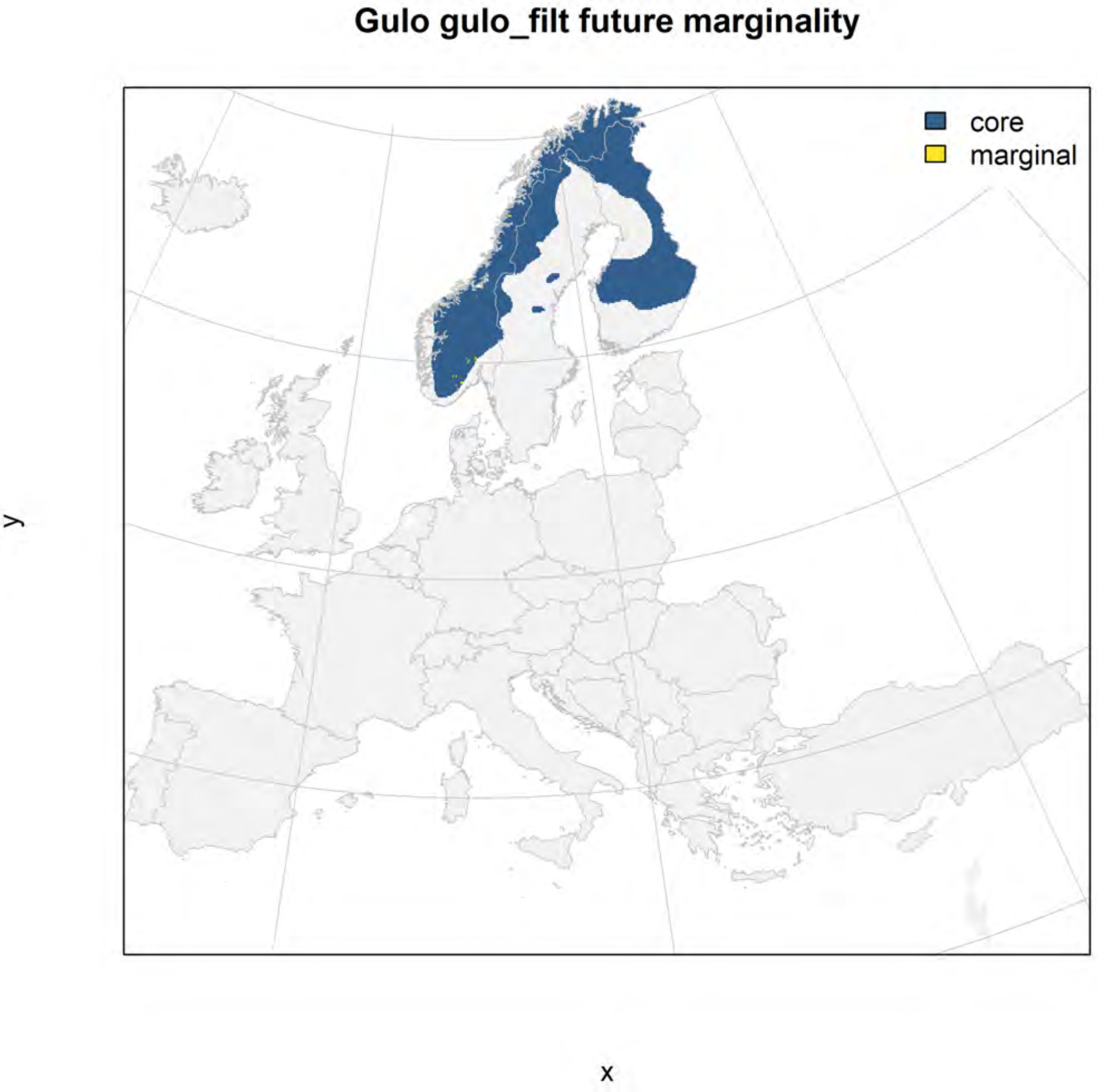
Gulo gulo_filt future marginality
core
marginal
y
x

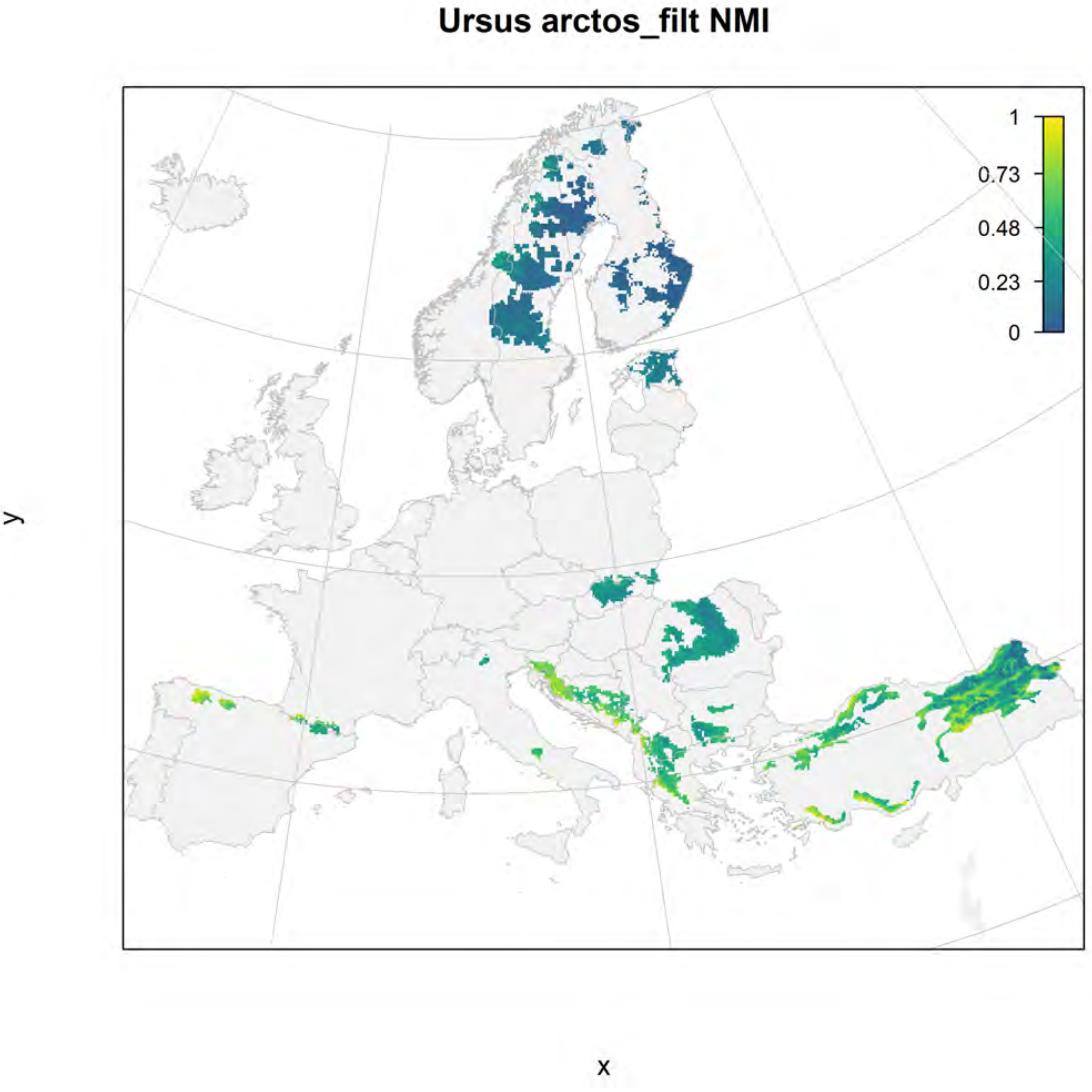
Ursus arctos_filt NMI
1
0.73
0.48
0.23
0
y
x

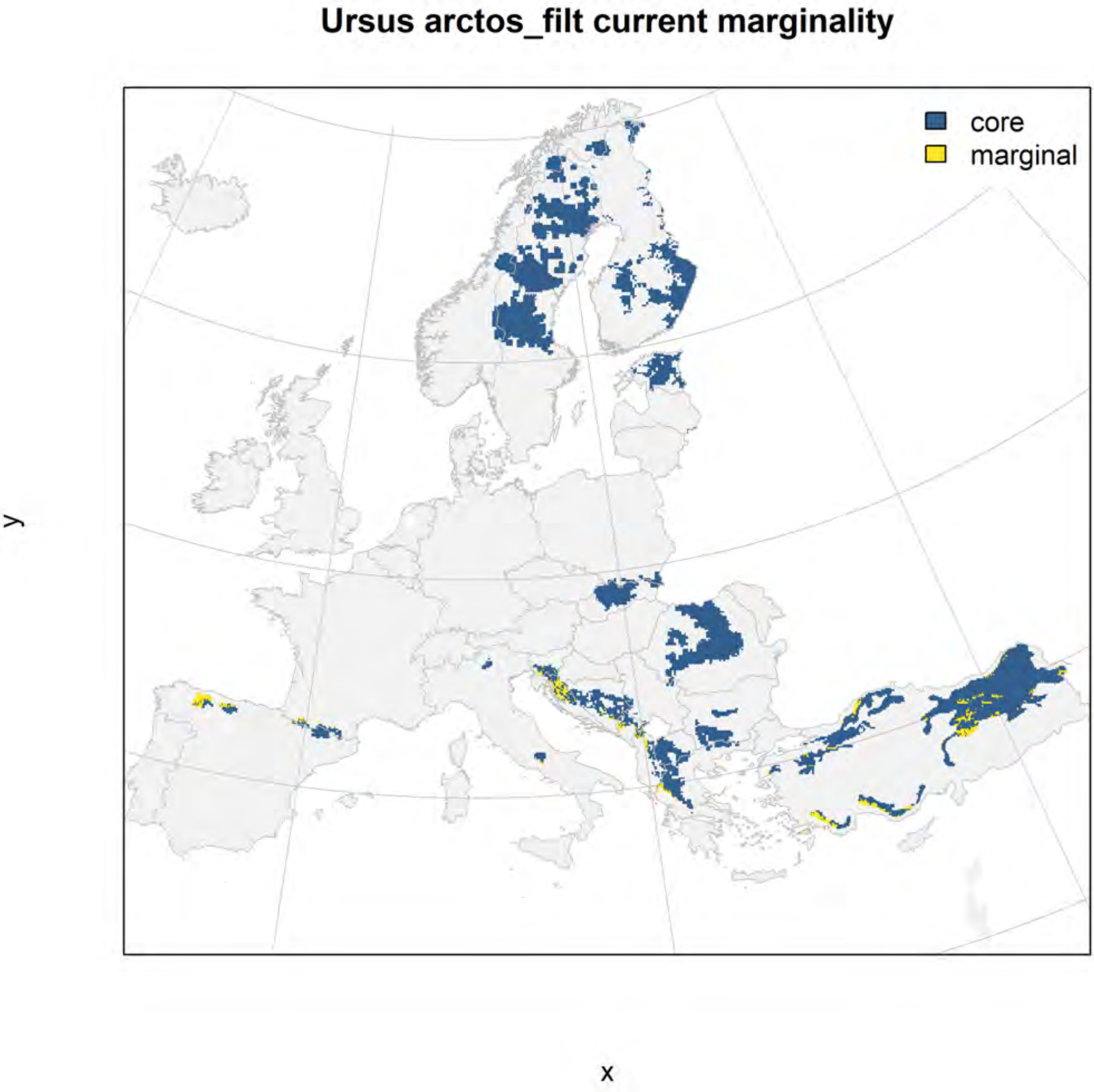
Ursus arctos_filt current marginality
core
marginal
y
x

### Ursus arctos_filt future marginality

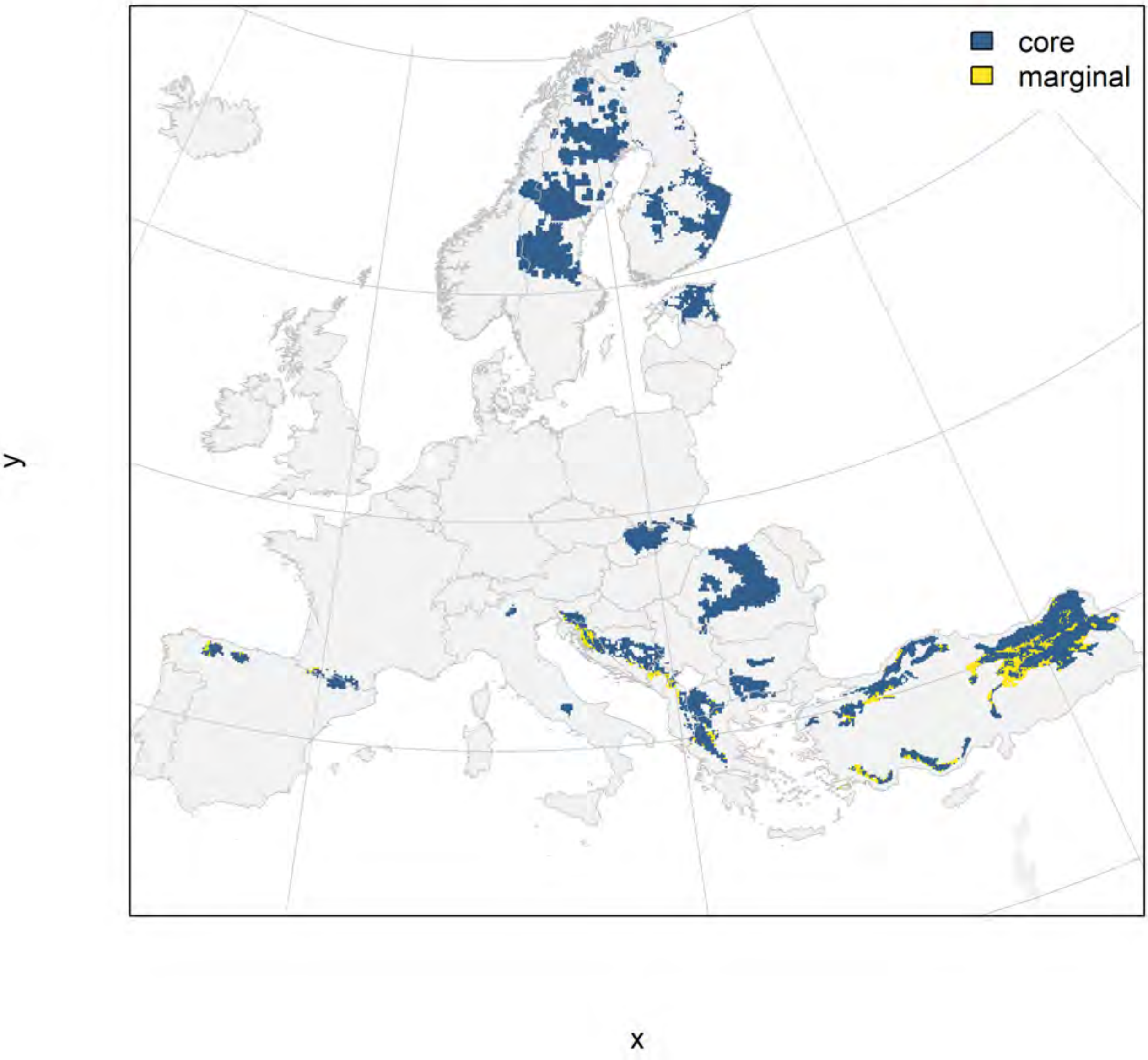
