## Appendix S6 for "Conserving genetic diversity during climate change: Niche marginality and discrepant monitoring capacity in Europe"

Appendix S6. Maps of marginality of selected species of European forest tree species. The climate marginality of each species is depicted in a series of three figures: raw marginality in which a value of 1 indicates the 99th percentile of the species niche space within its current range (filtered for appropriate habitat with a CORINE land cover layer), the current geographic distribution of areas within the 25% most marginal climatic niche conditions and the remaining core area, and a predicted future distribution of core and marginal areas within the current species range. Please see the Online Methods section for a description of marginality calculations.

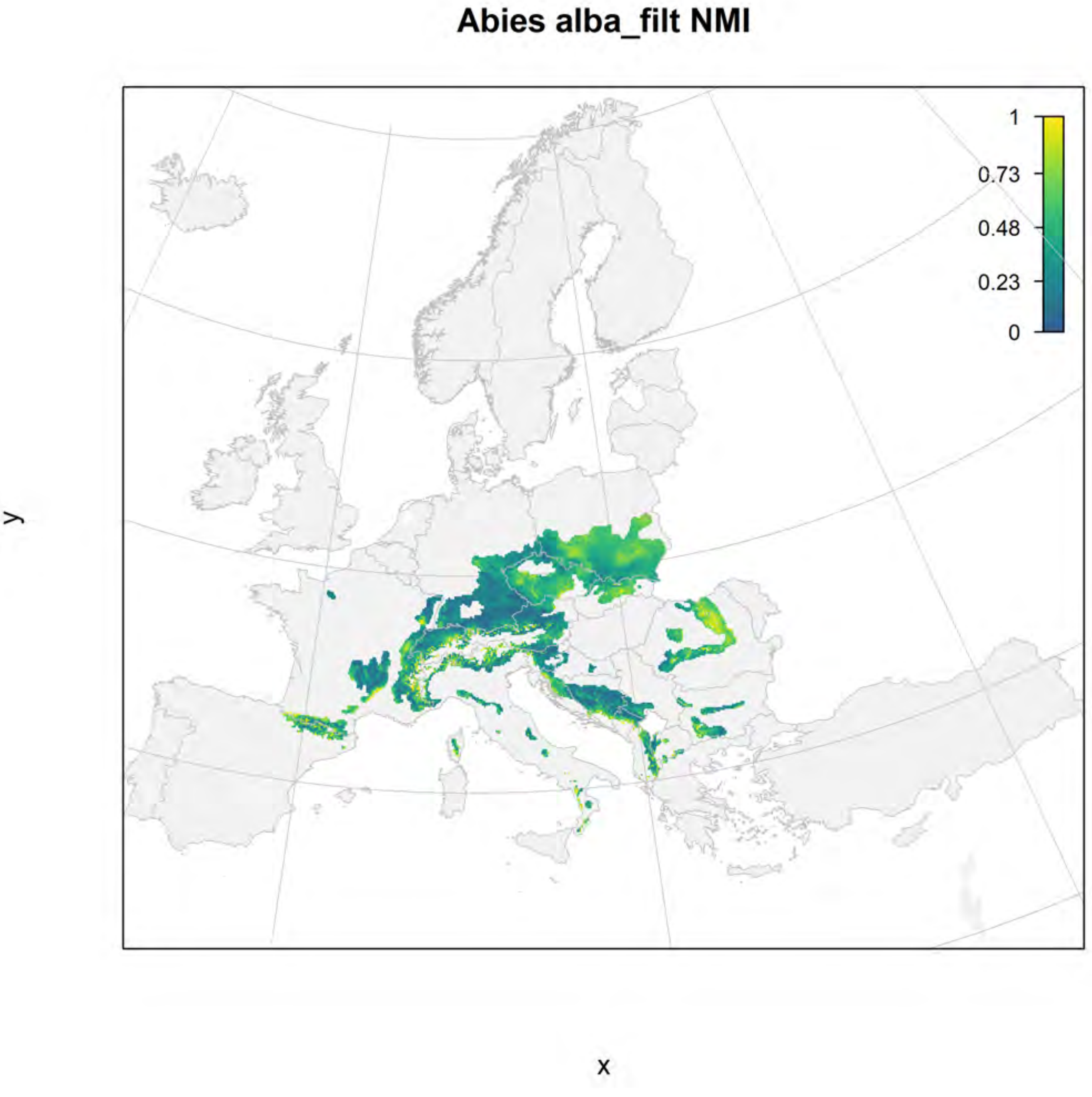

Abies alba_filt NMI
1
0.73
0.48
0.23
0
y
x

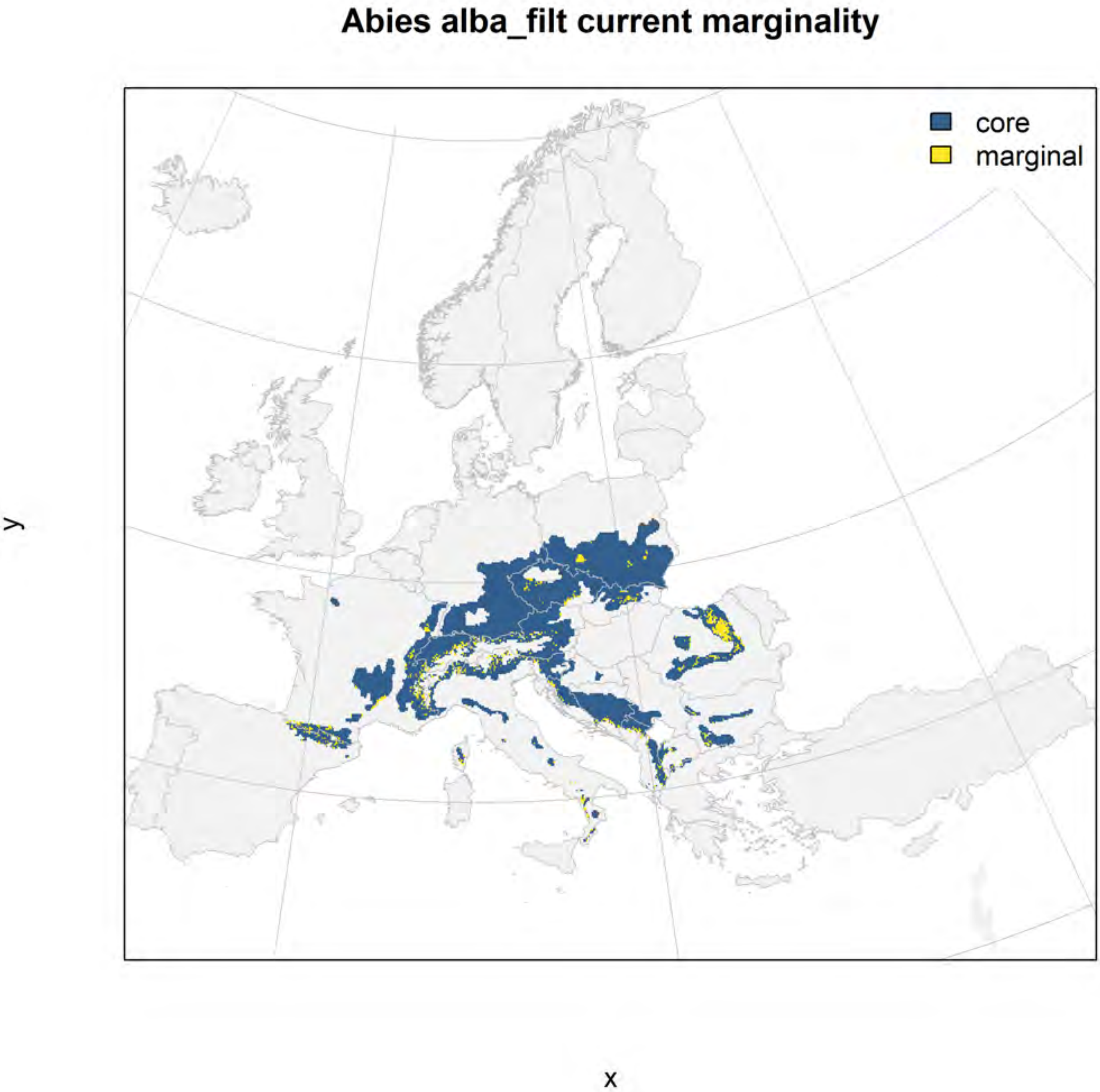
Abies alba_filt current marginality
core
marginal
y
x

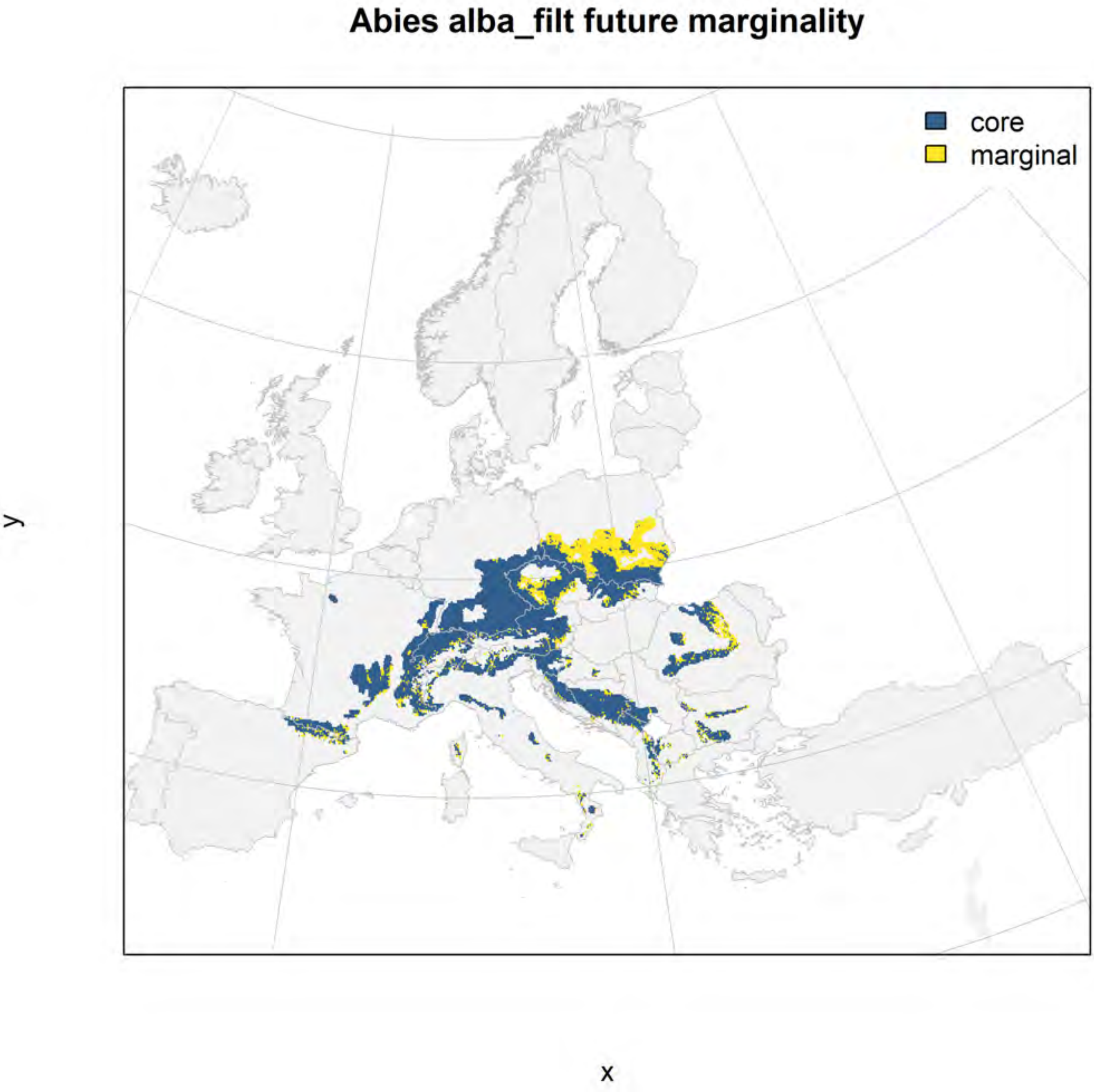
Abies alba_filt future marginality
core
marginal
y
x

#### Abies borisii-regis_filt NMI

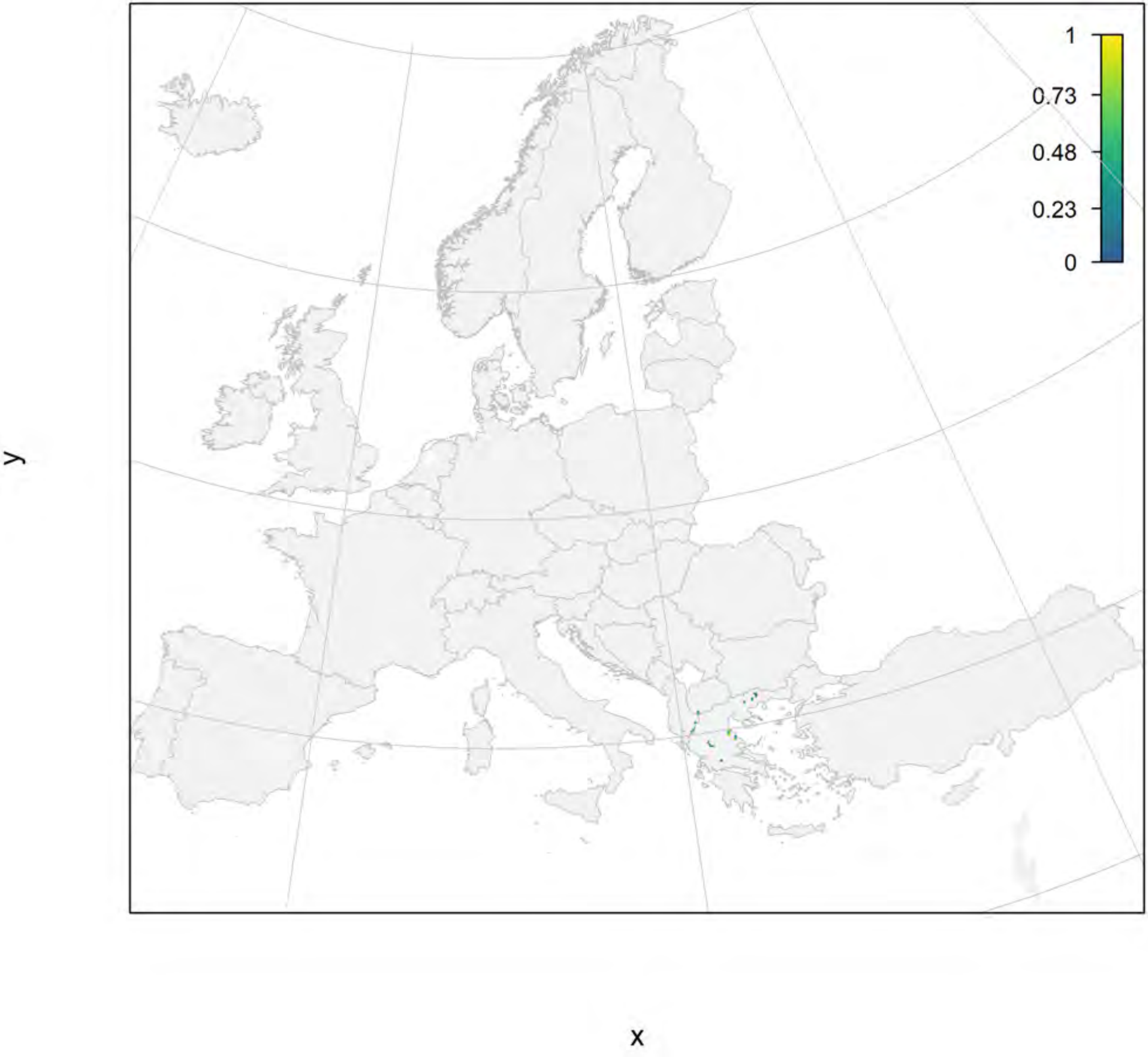

#### Abies borisii-regis_filt current marginality

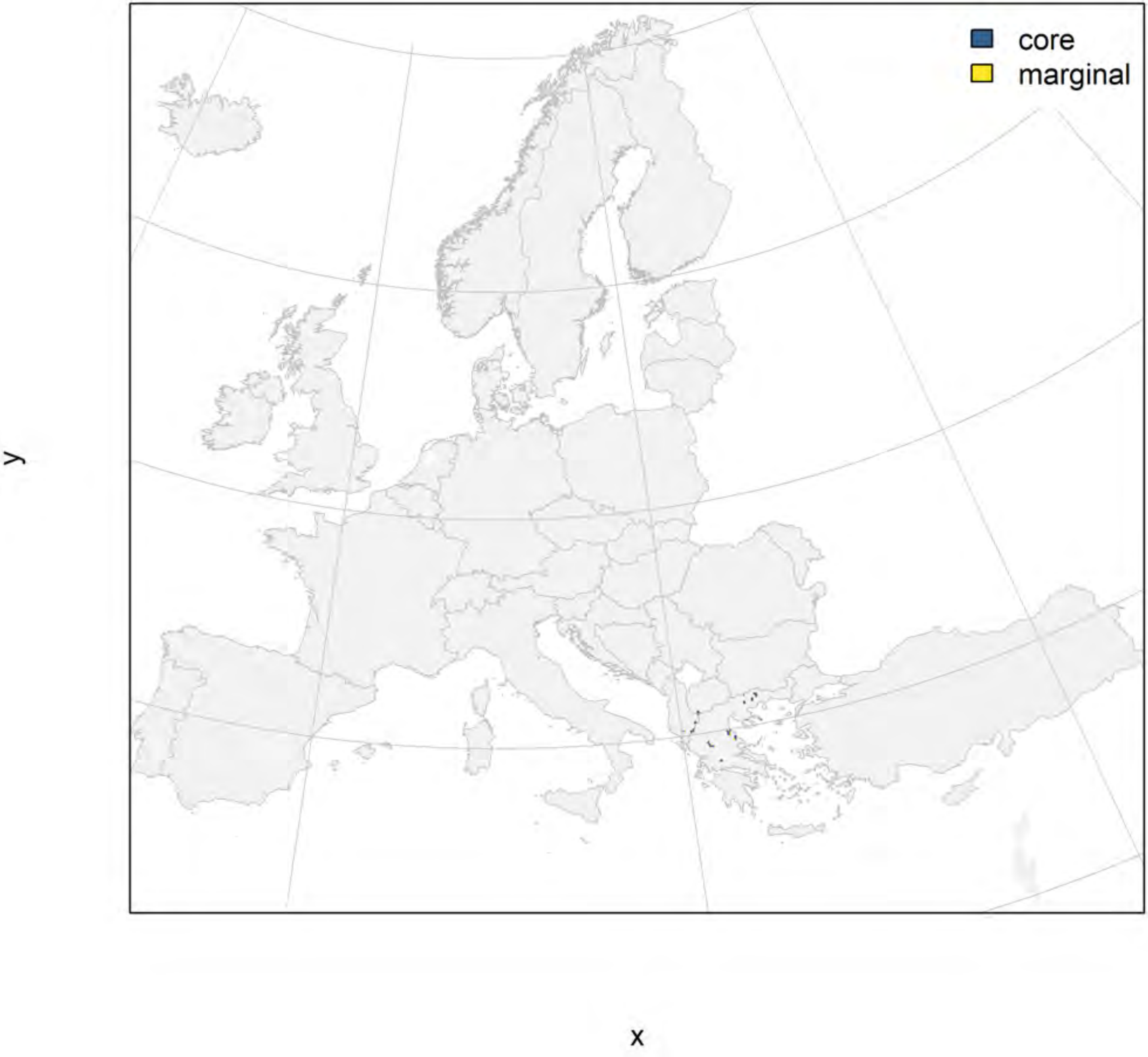

Abies borisii-regis_filt future marginality

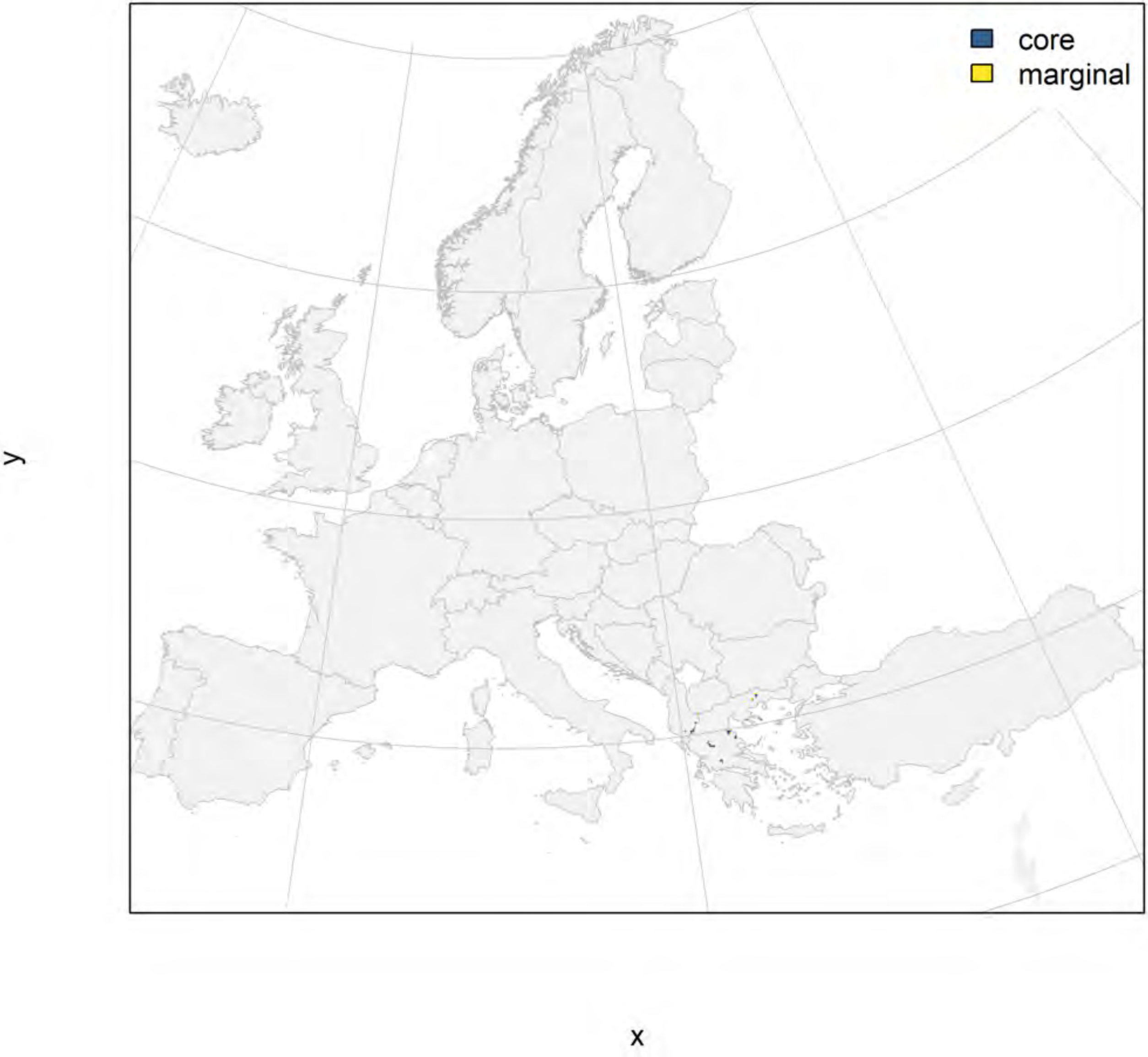

#### Abies cephalonica_filt NMI

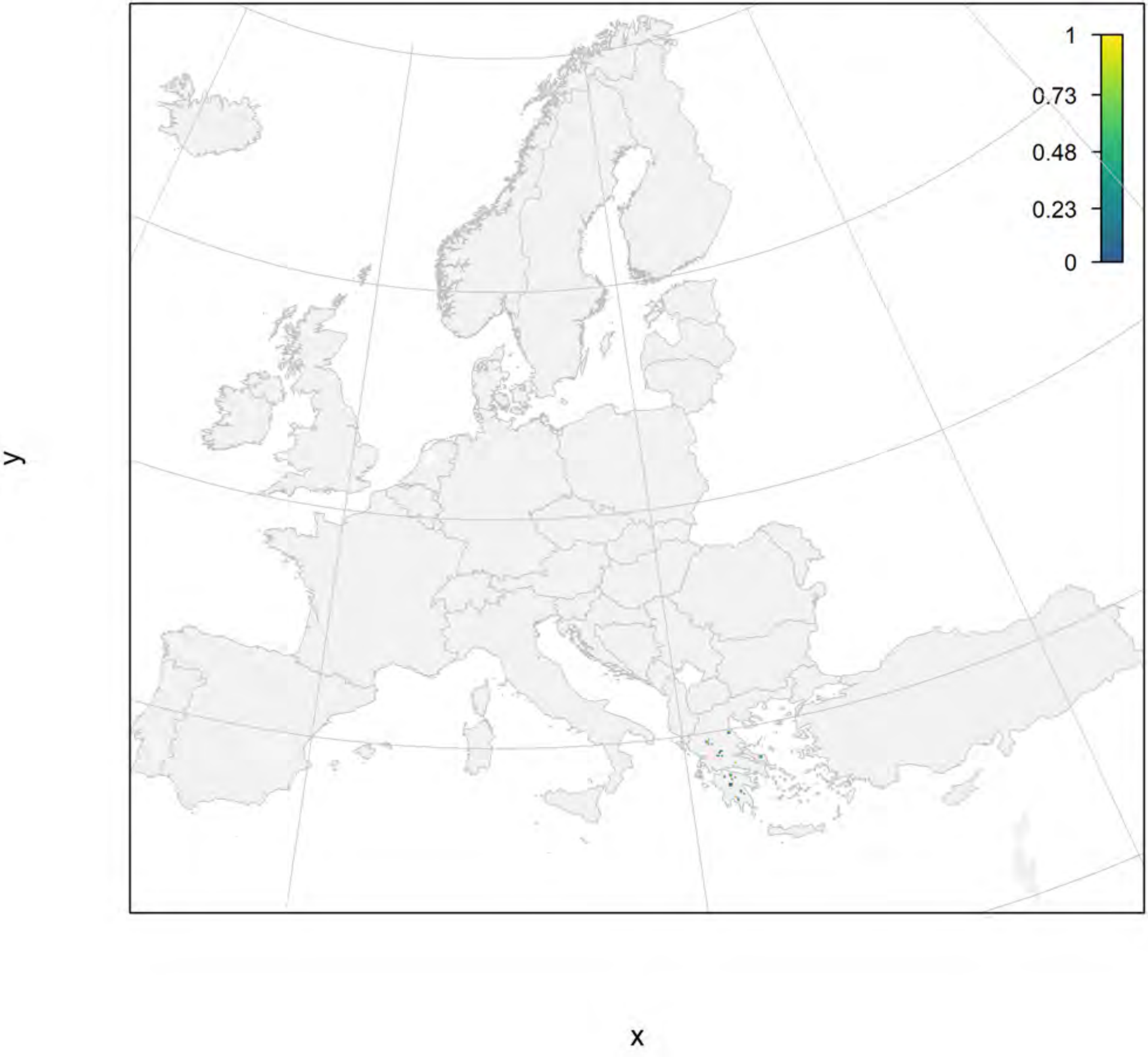

### Abies cephalonica_filt current marginality

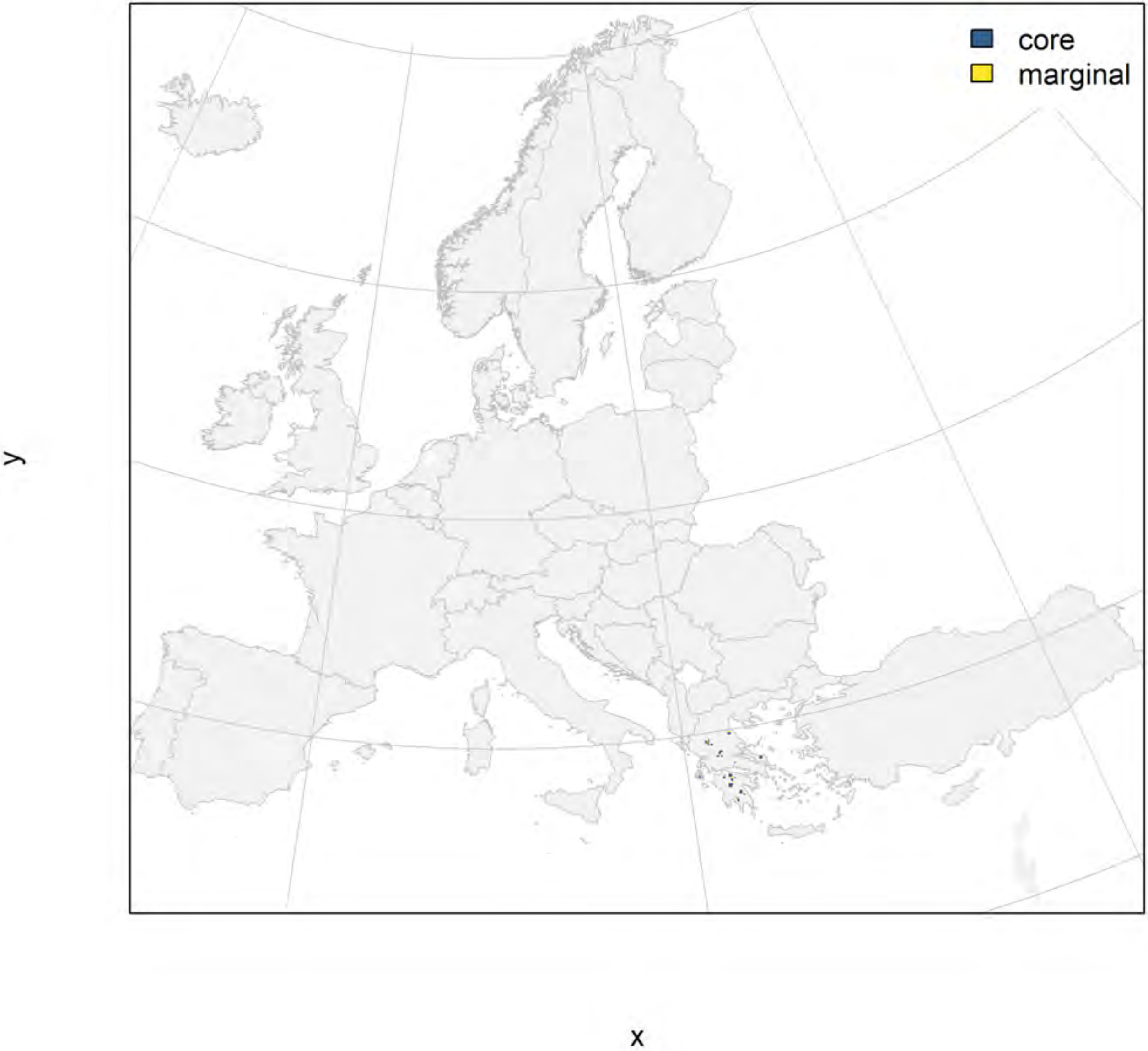

### Abies cephalonica_filt future marginality

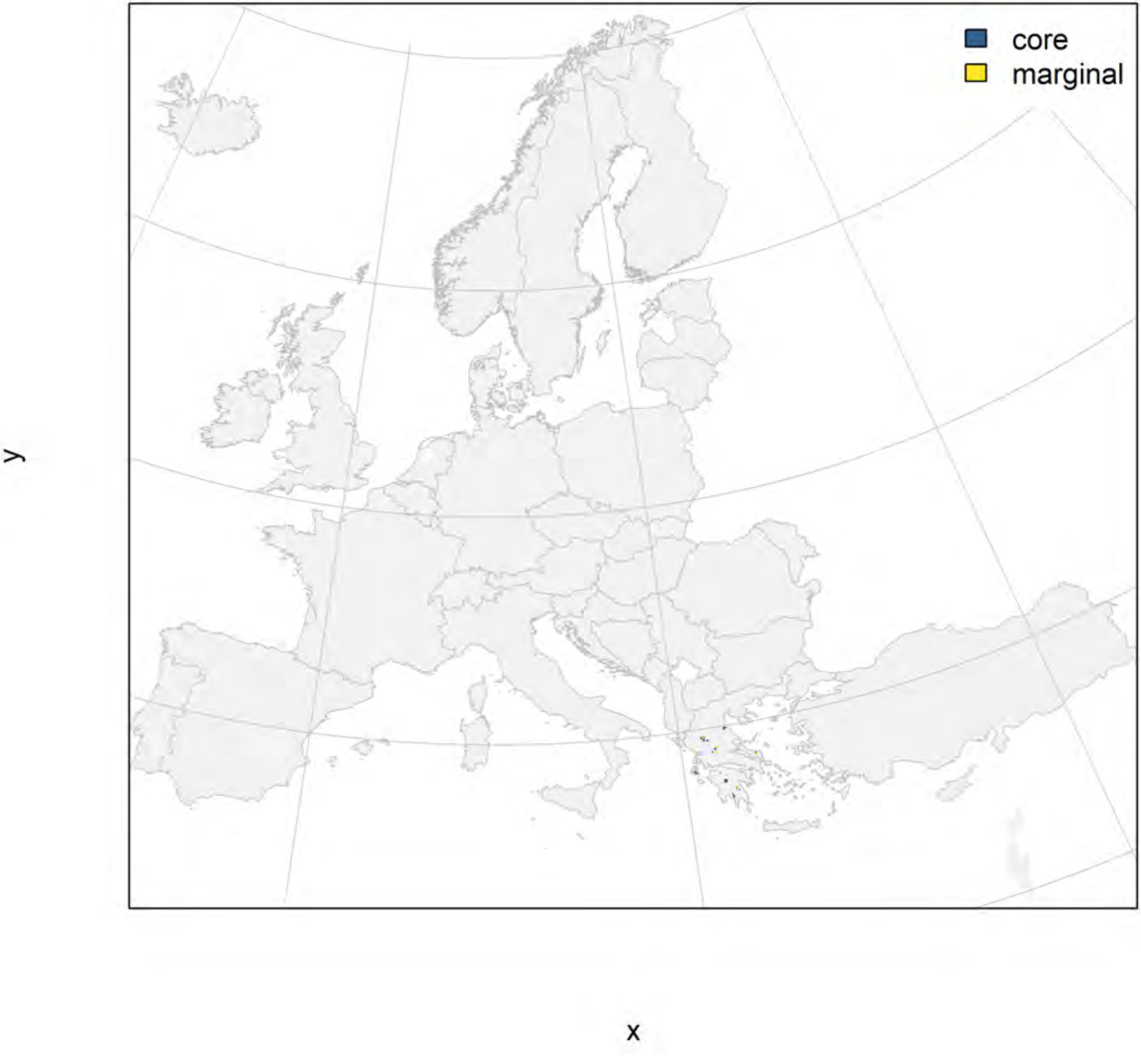

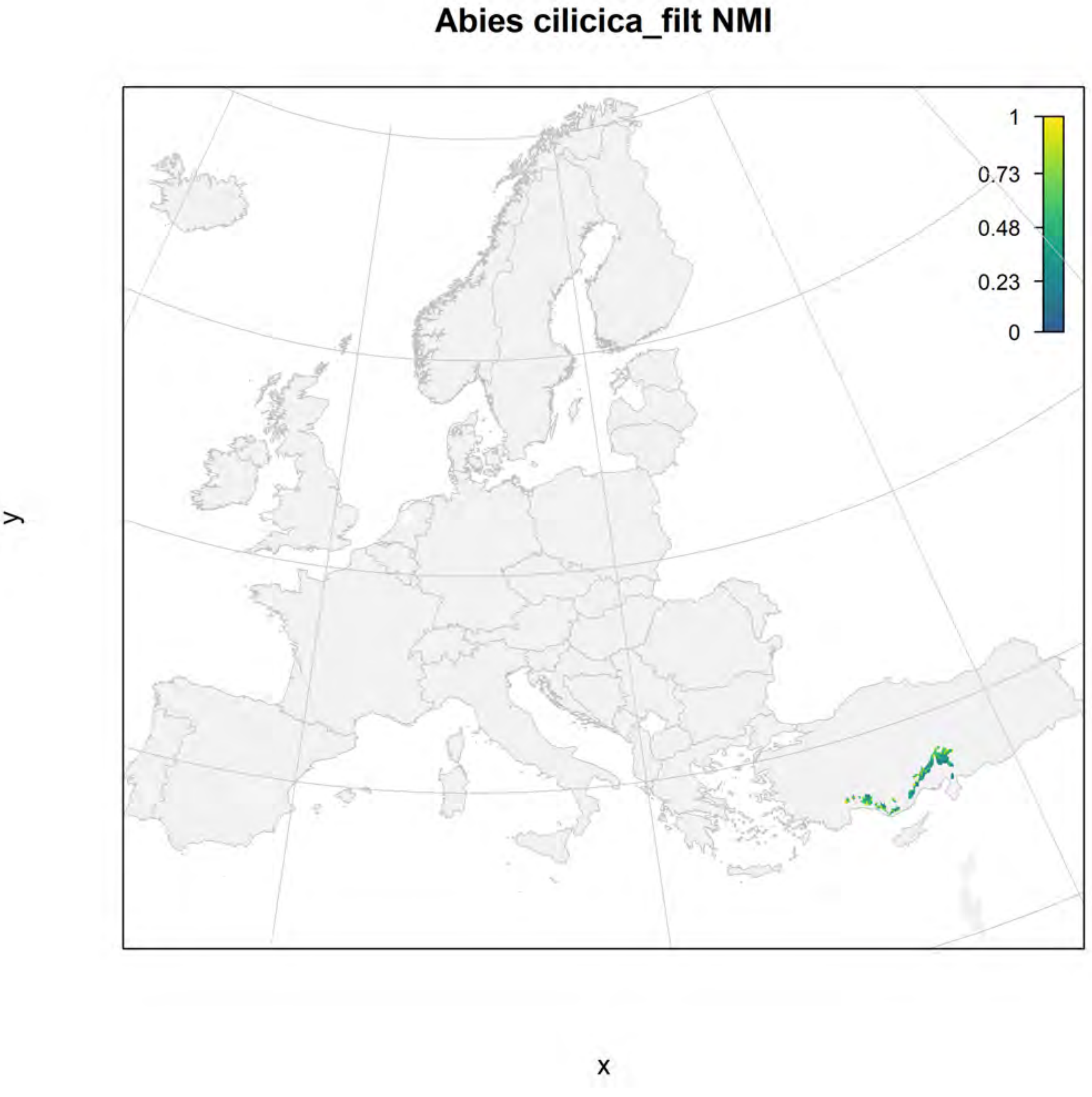
Abies cilicica_filt NMI
1
0.73
0.48
0.23
0
y
x

#### Abies cilicica_filt current marginality

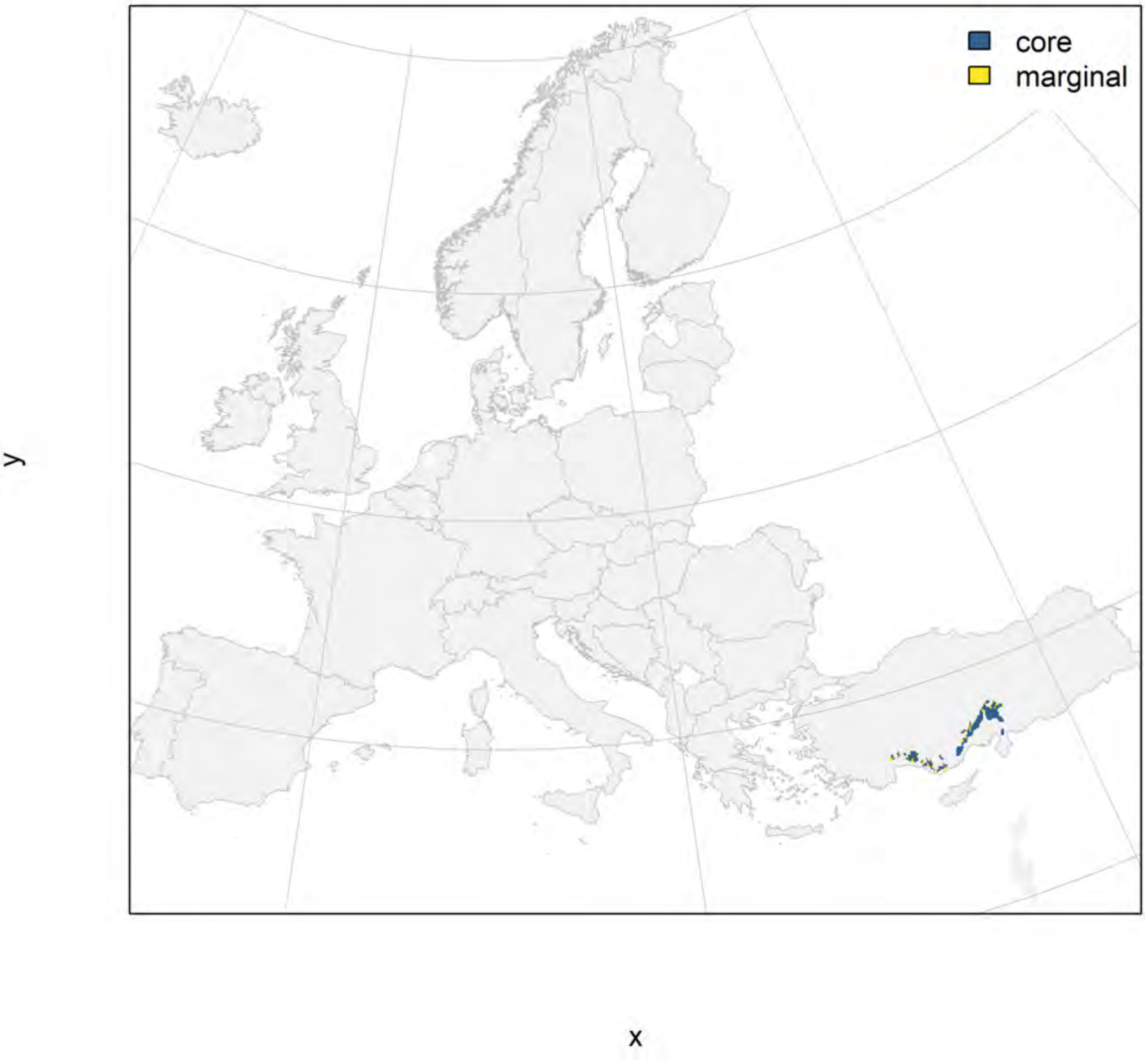

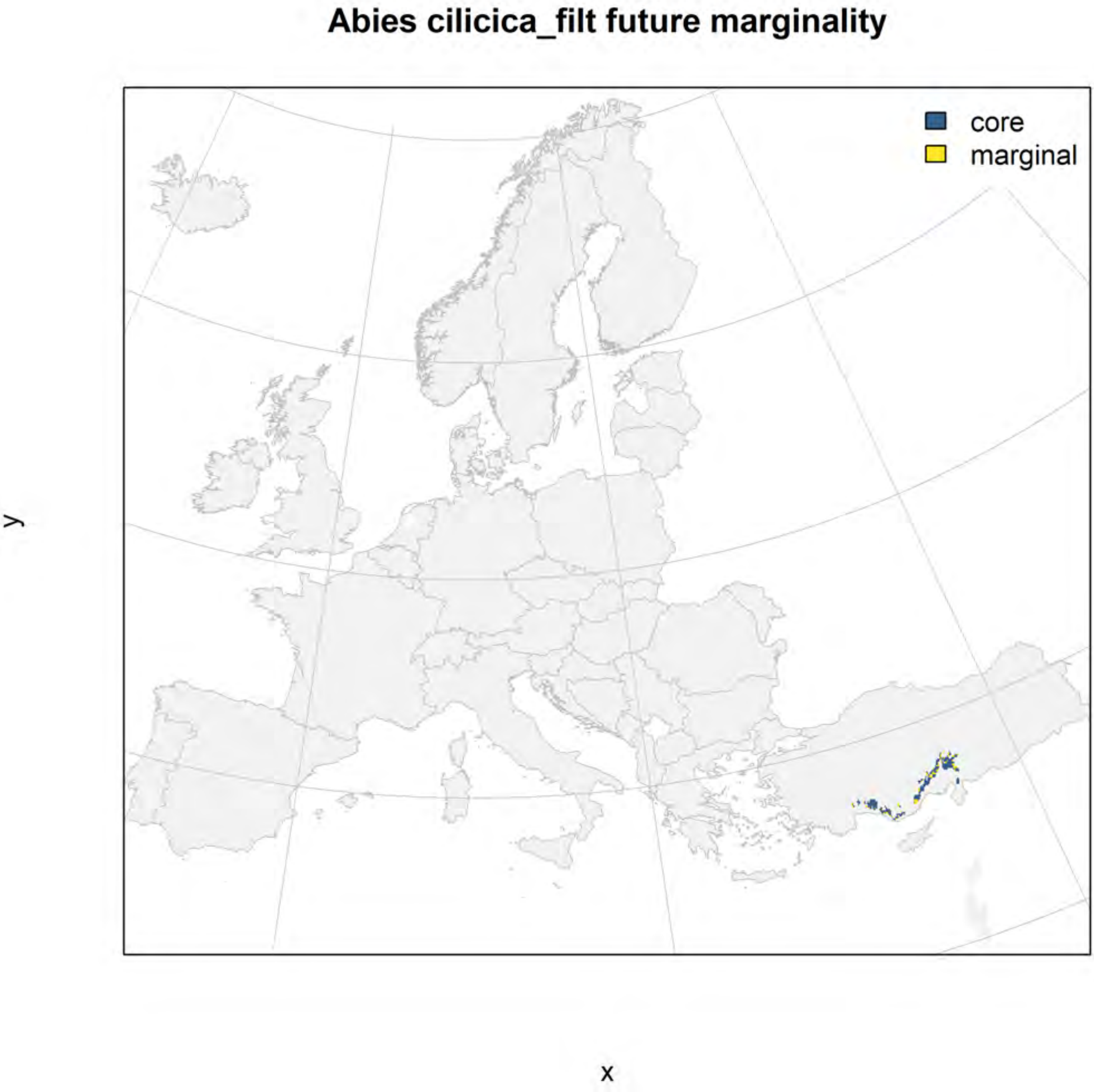
Abies cilicica_filt future marginality
core
marginal
y
x

#### Abies nordmanniana_filt NMI

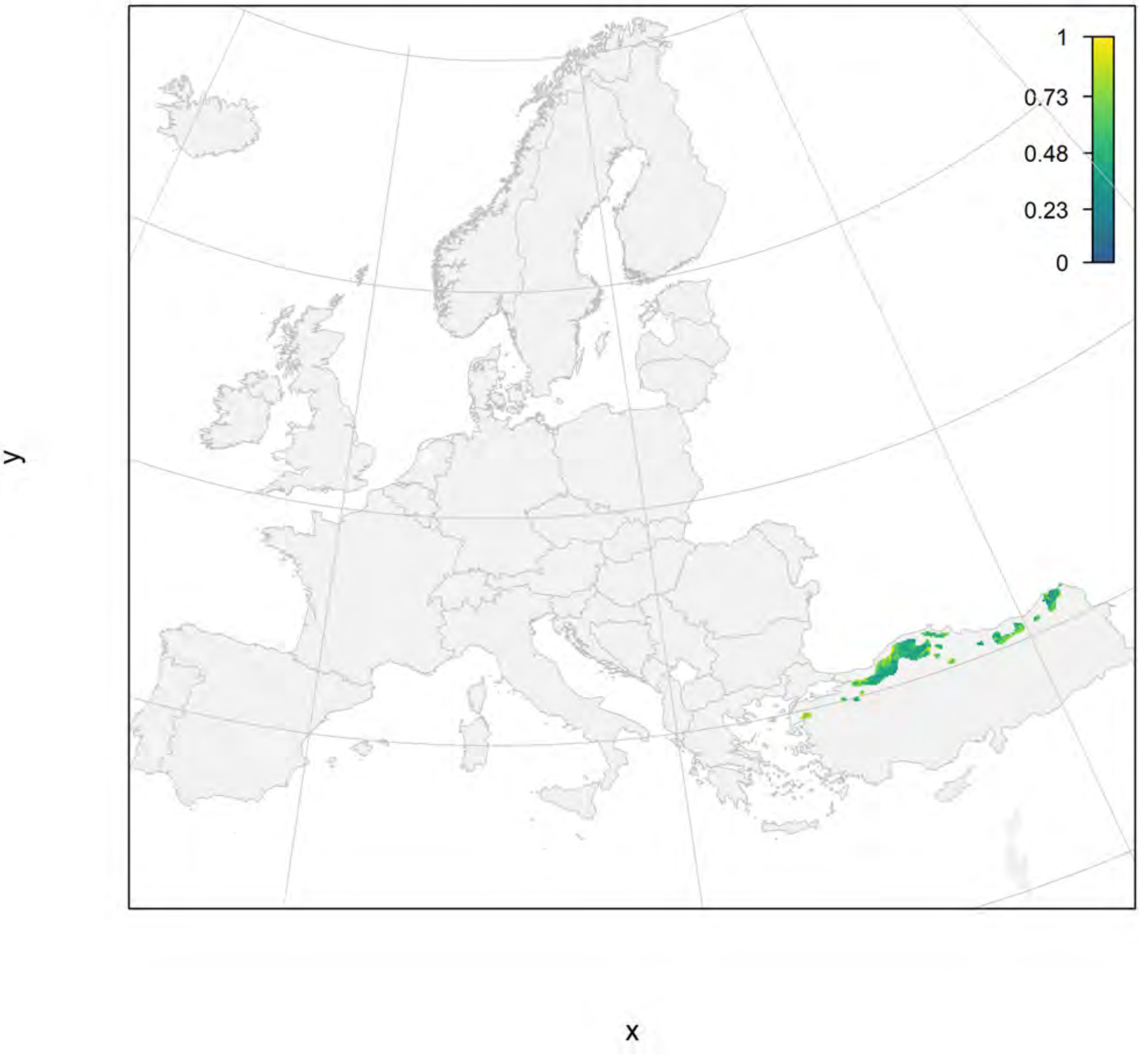

#### Abies nordmanniana_filt current marginality

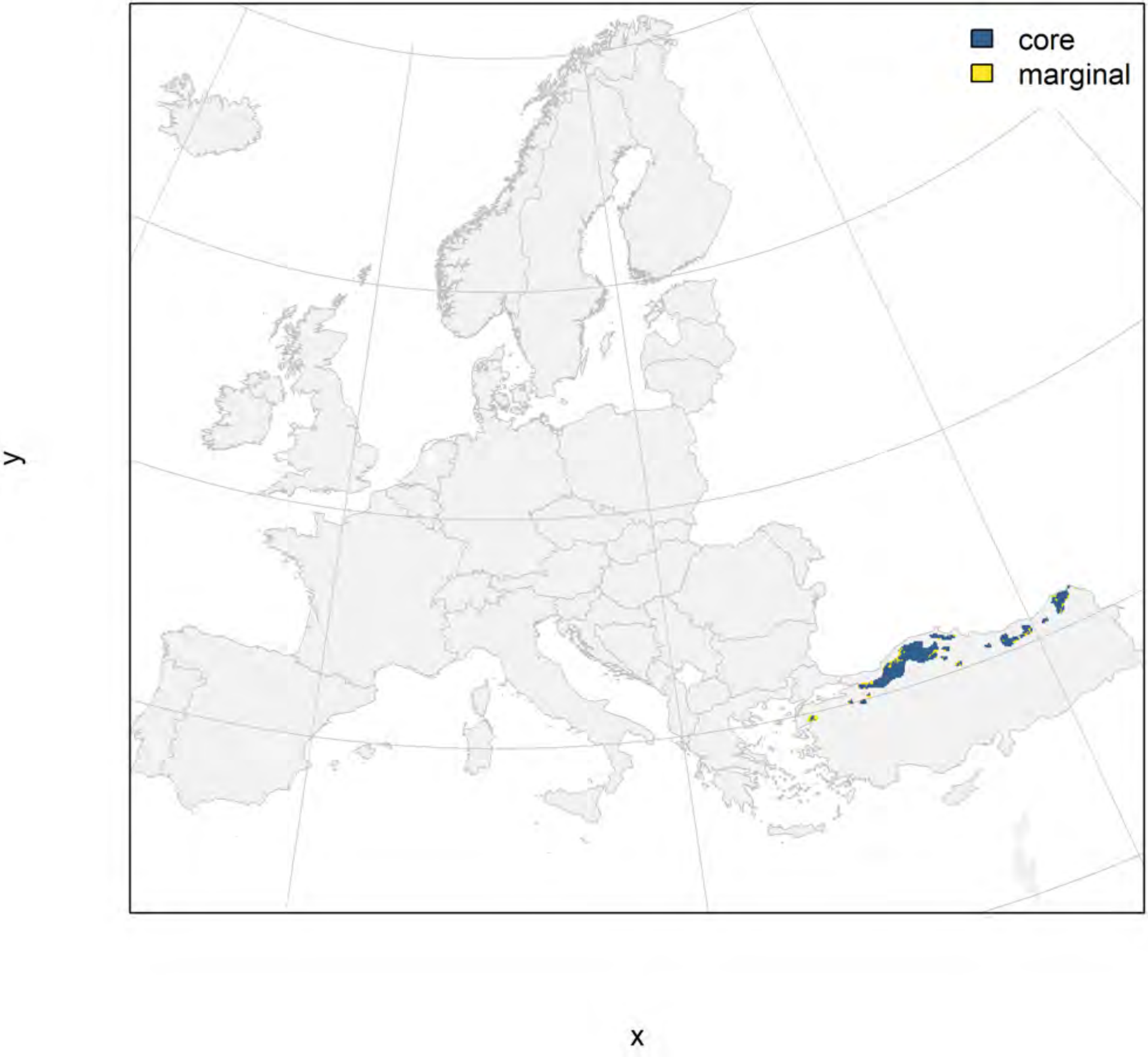

### Abies nordmanniana_filt future marginality

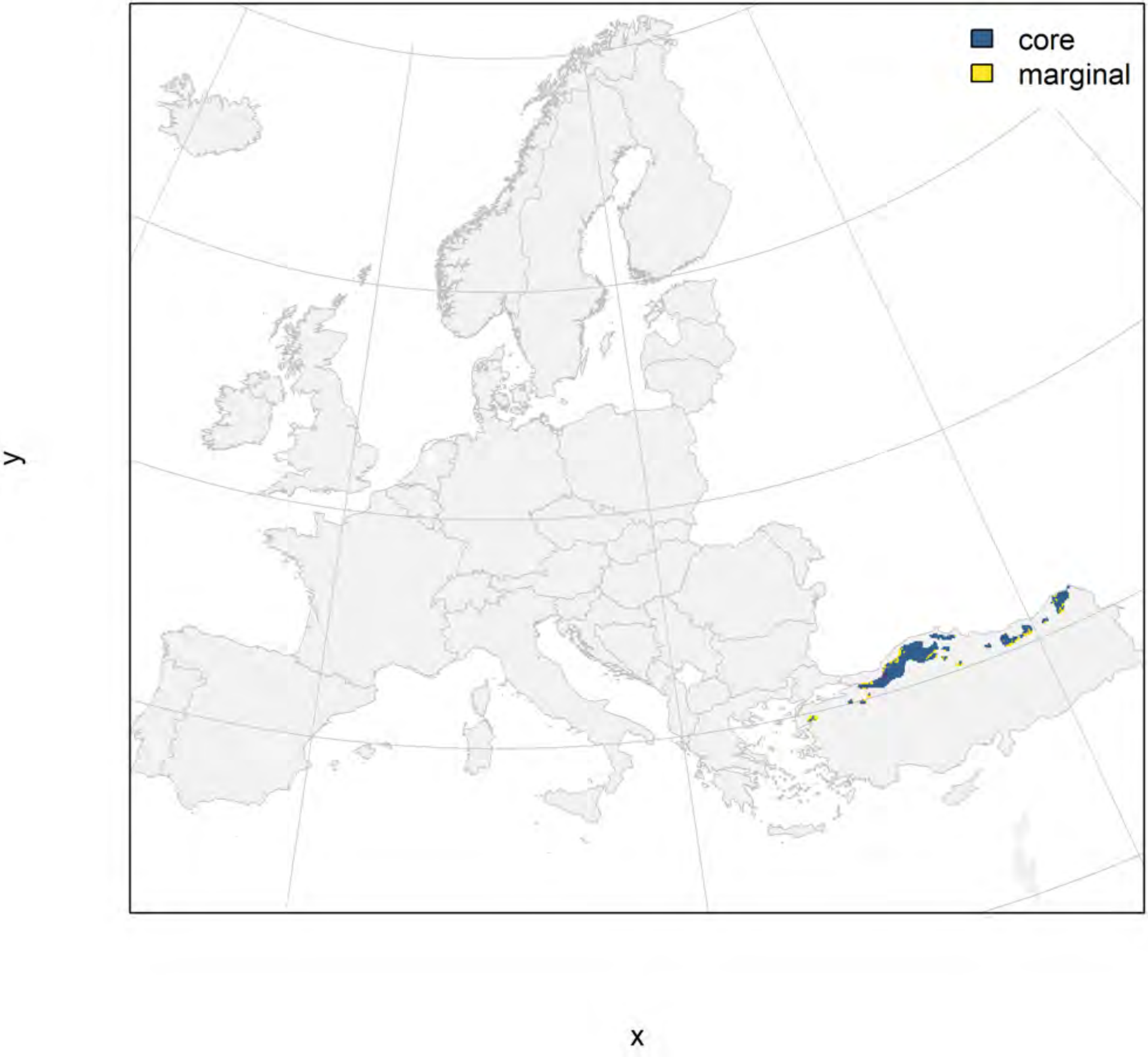

### Abies pinsapo_filt NMI

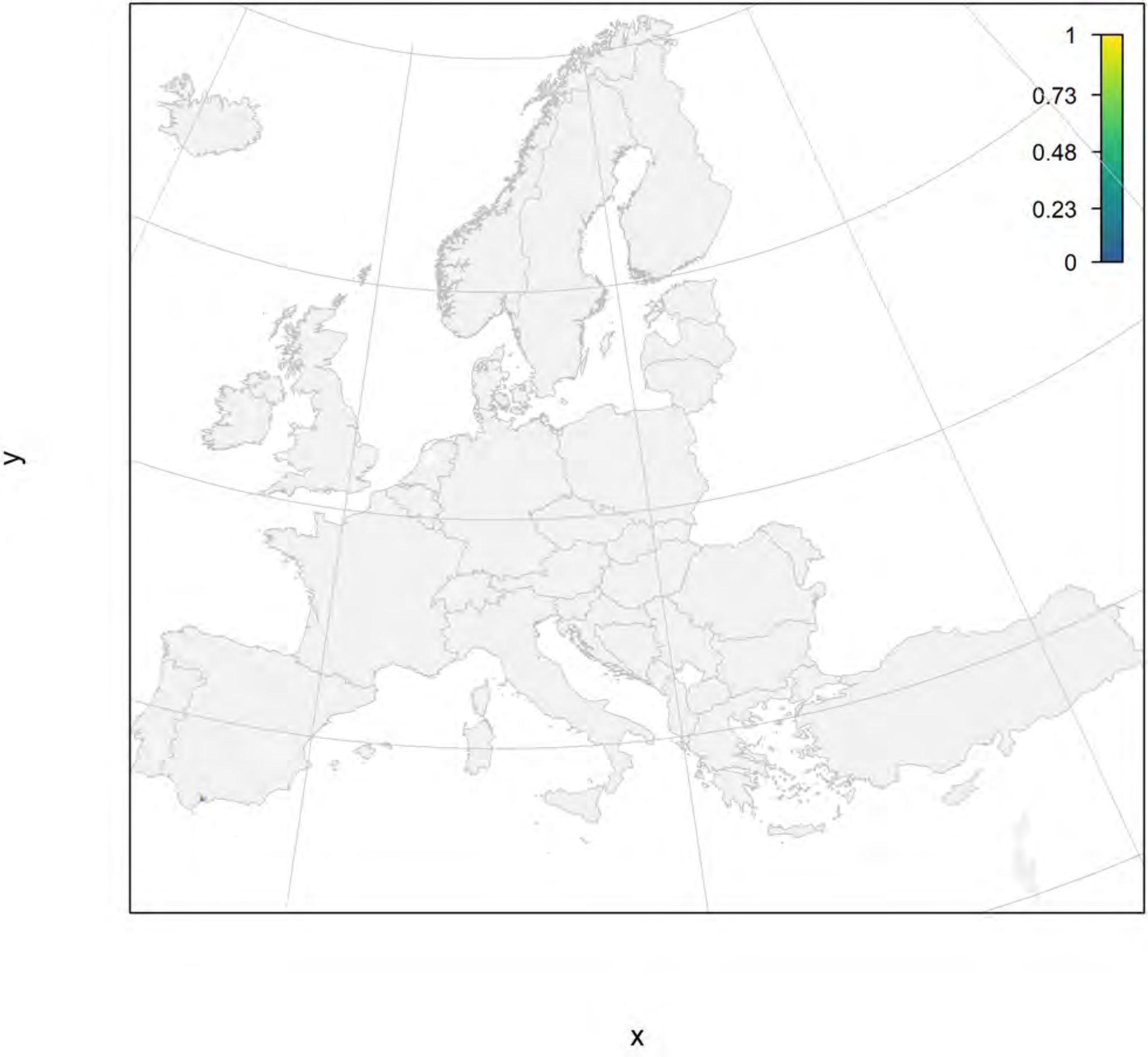

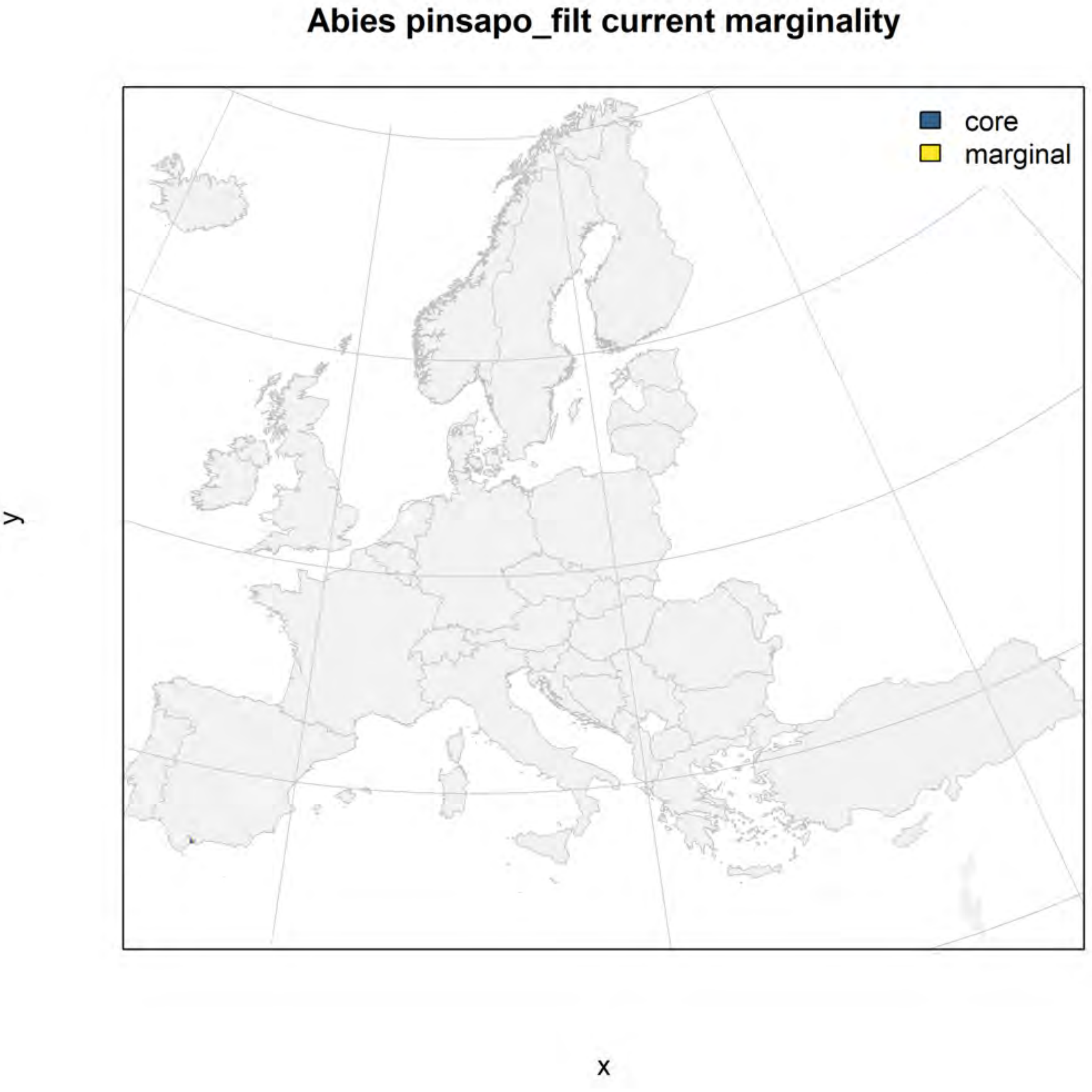
Abies pinsapo_filt current marginality
core
marginal
y
x

#### Abies pinsapo_filt future marginality

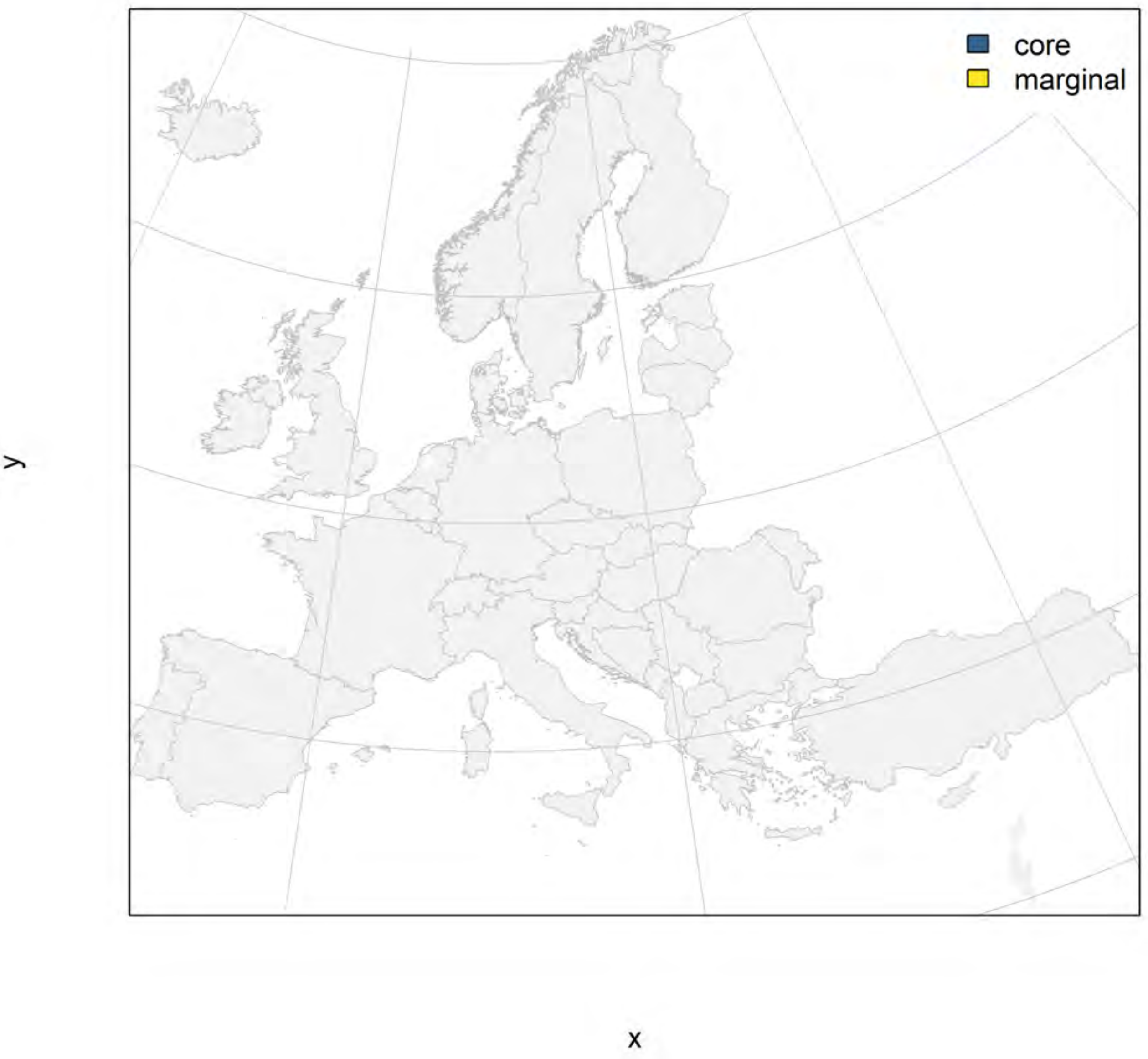

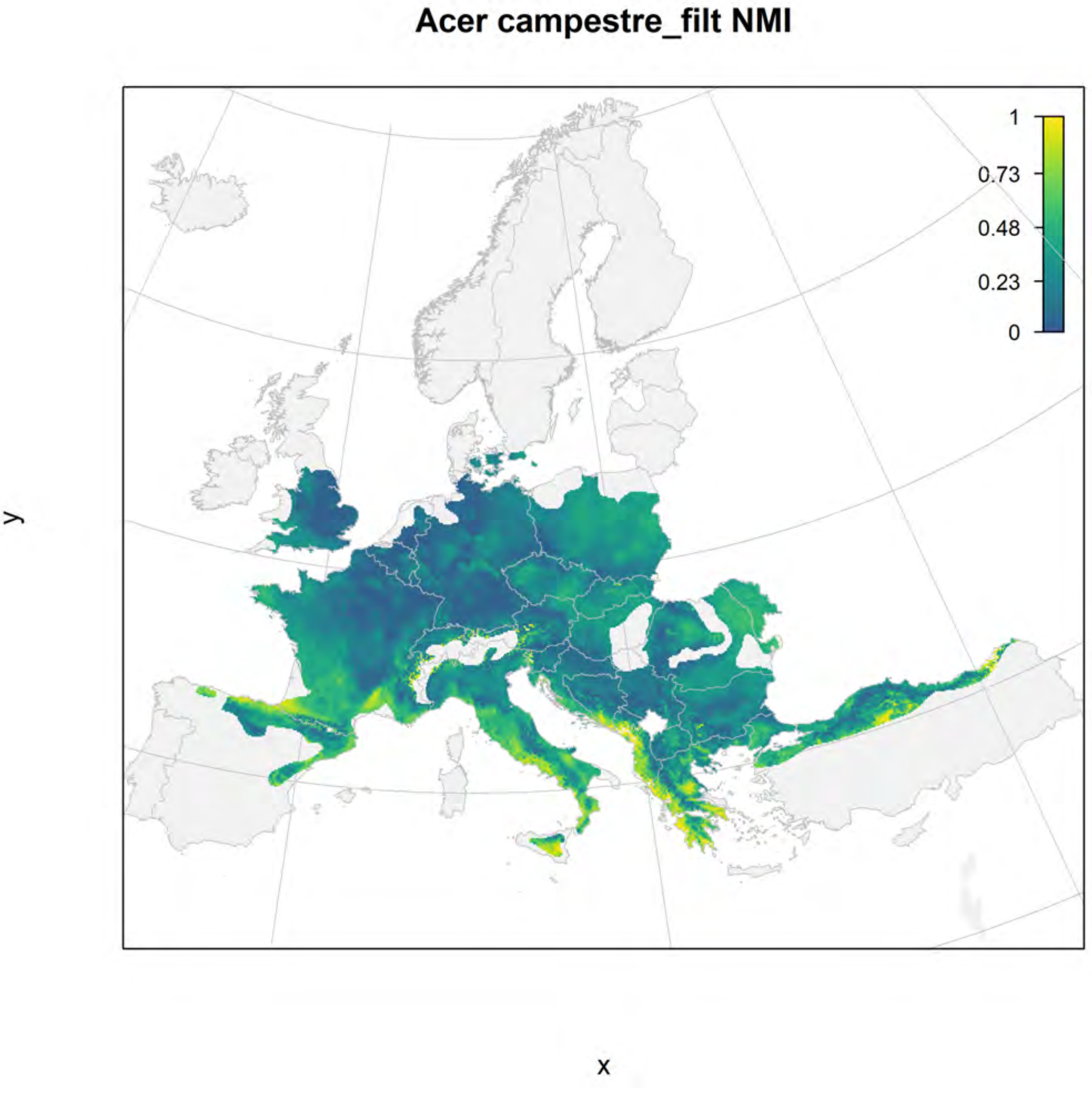
Acer campestre_filt NMI
1
0.73
0.48
0.23
0
y
x

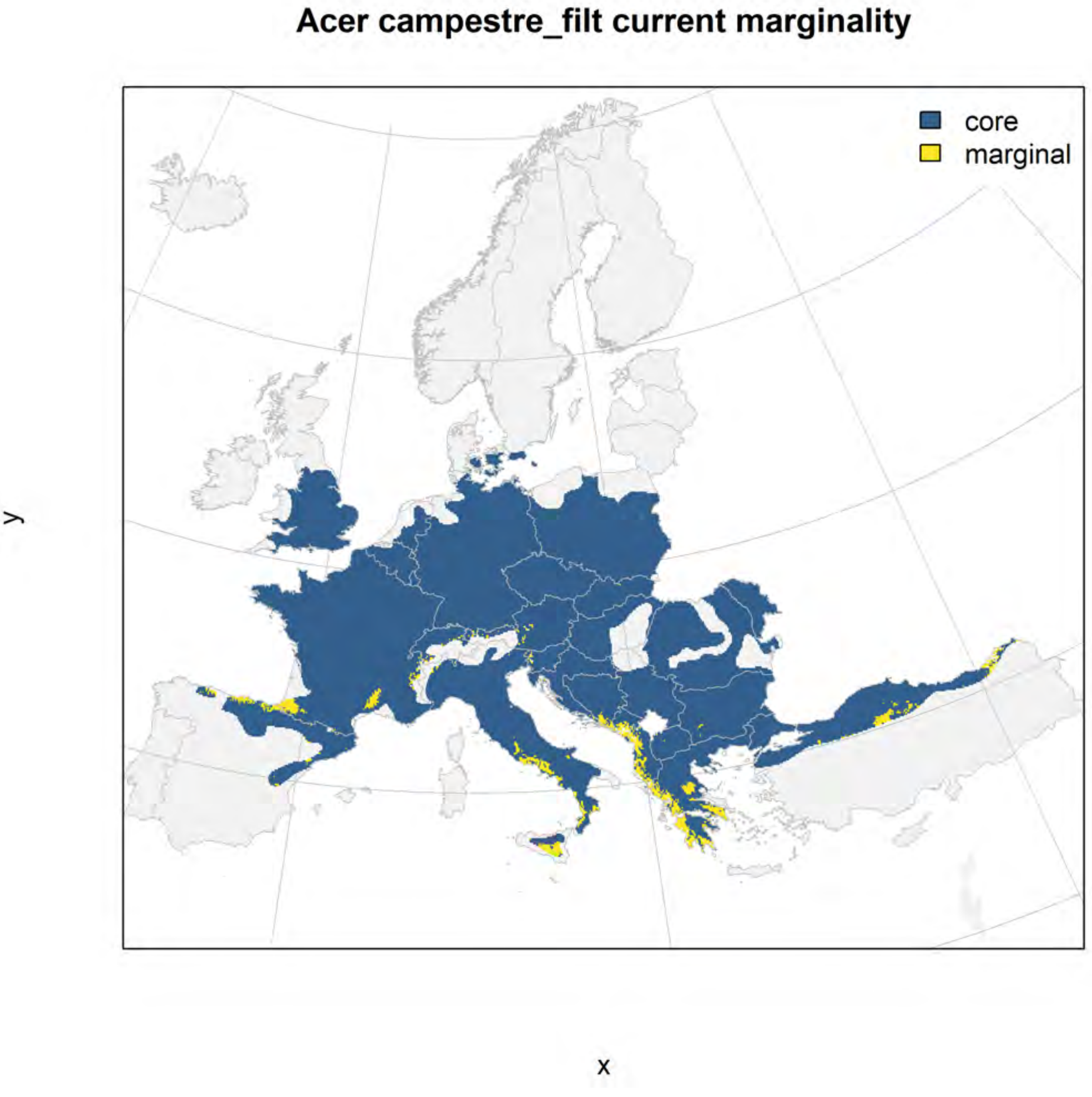
Acer campestre_filt current marginality
core
marginal
y
x

#### Acer campestre_filt future marginality

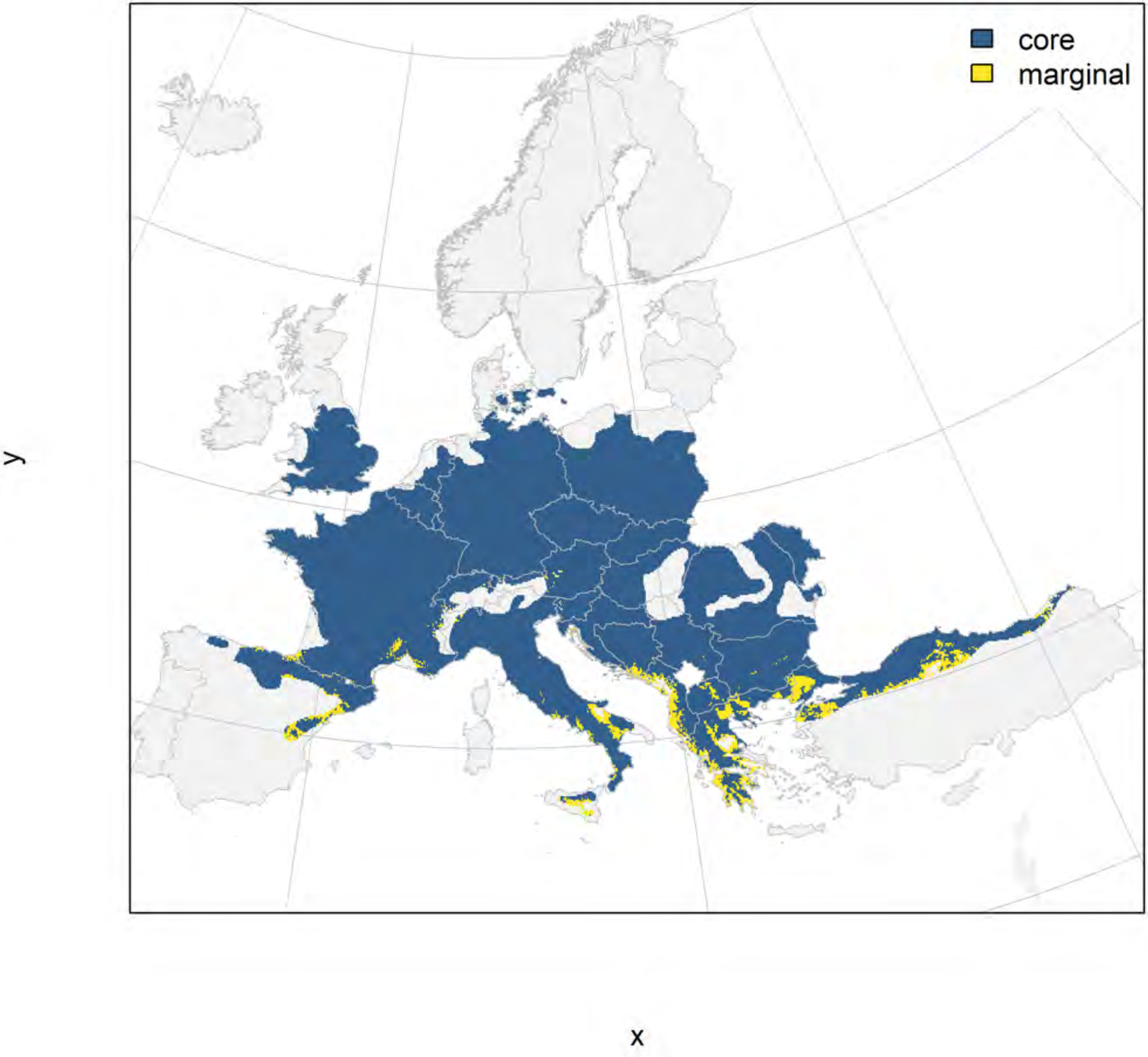

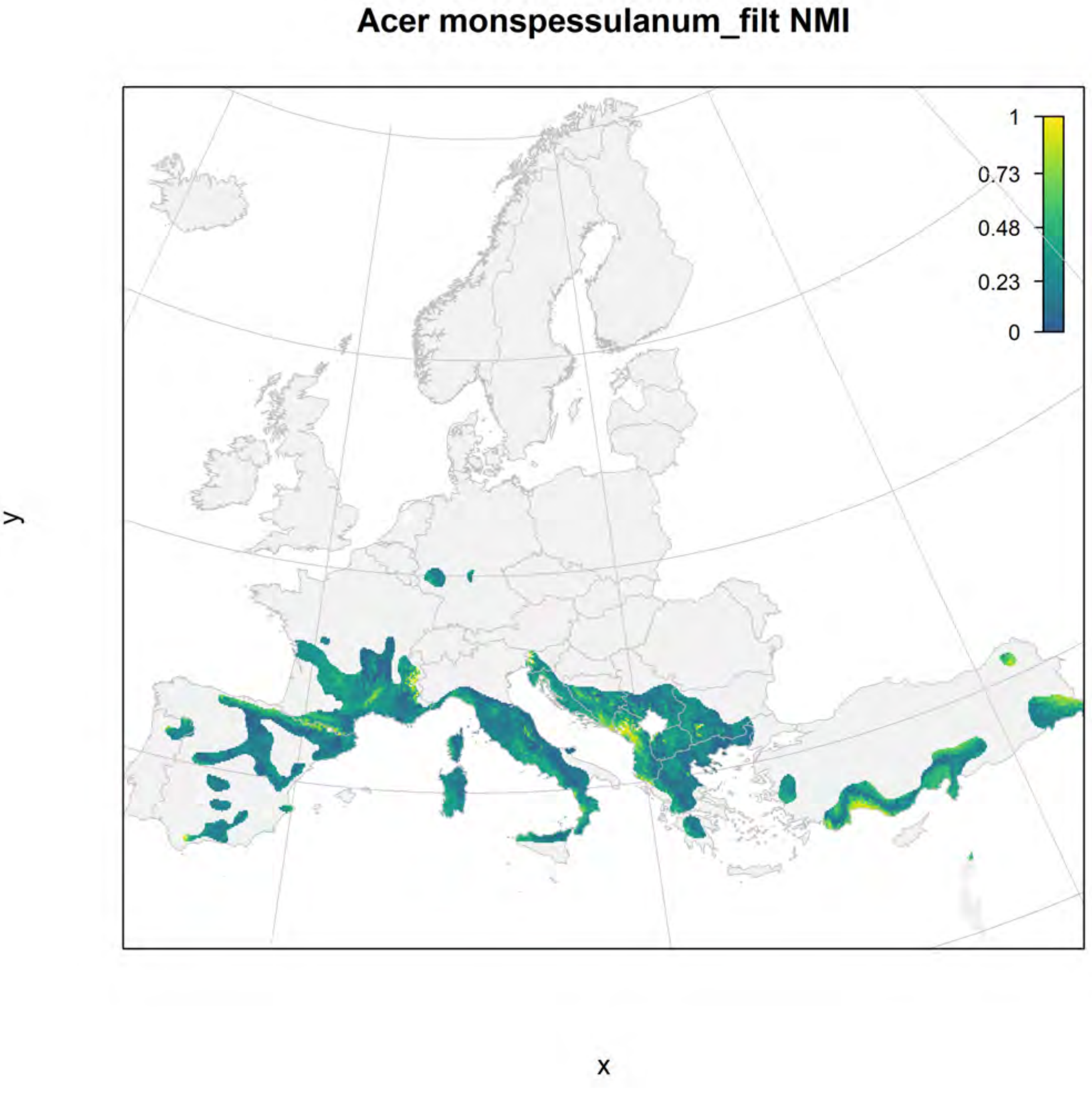
Acer monspessulanum_filt NMI
1
0.73
0.48
0.23
0
y
x

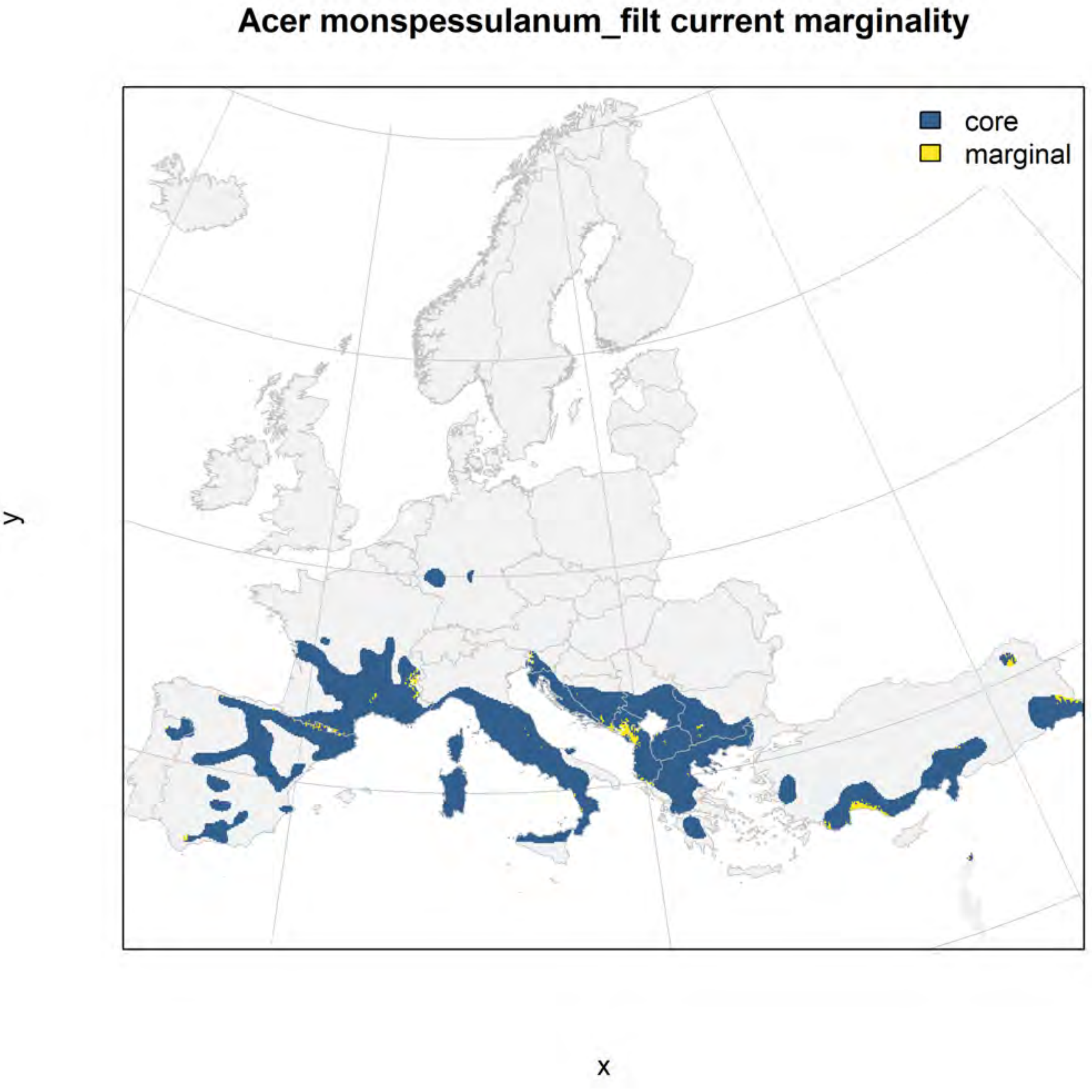
Acer monspessulanum_filt current marginality
core
marginal
y
x

#### Acer monspessulanum_filt future marginality

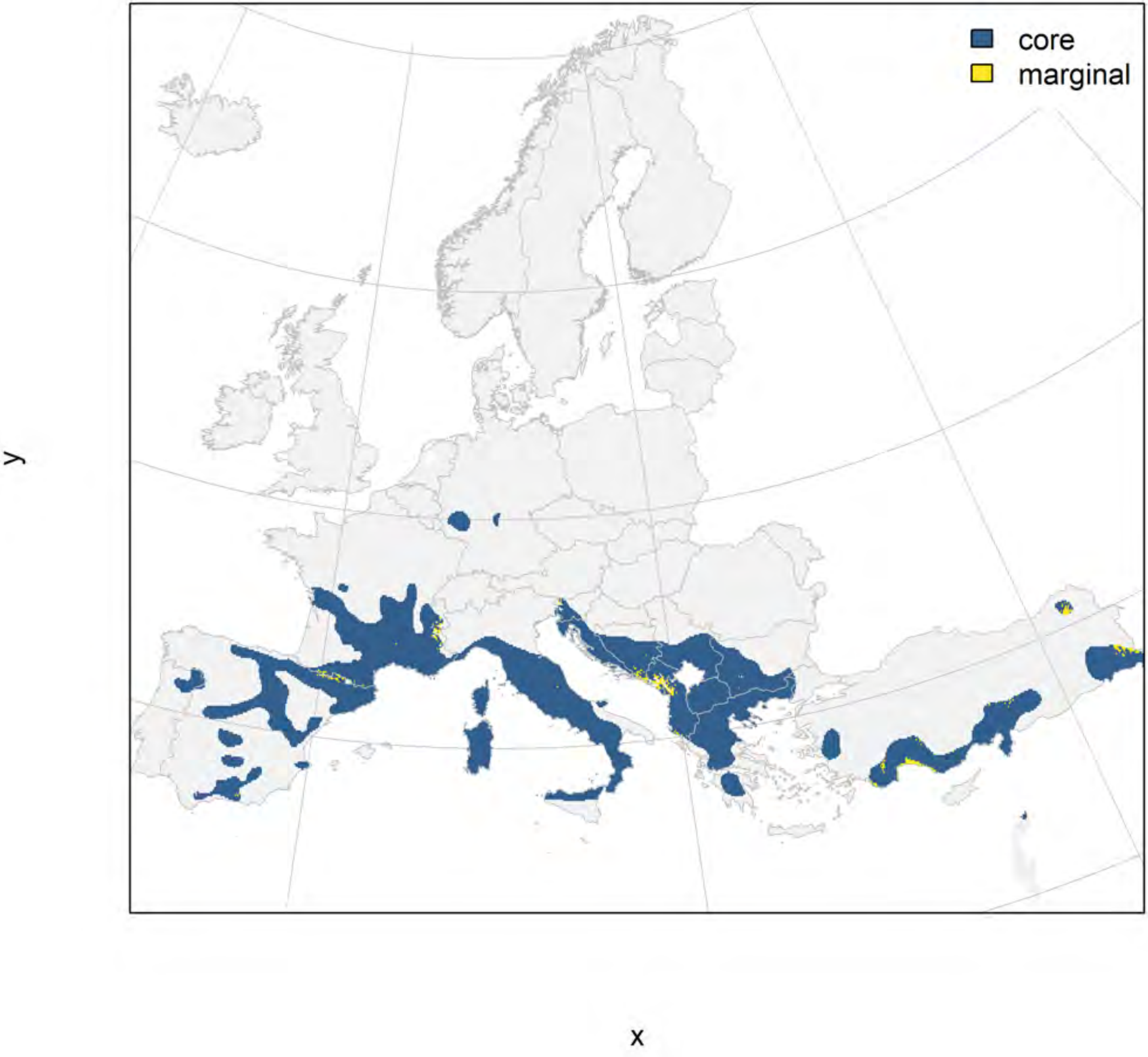

### Acer platanoides_filt NMI

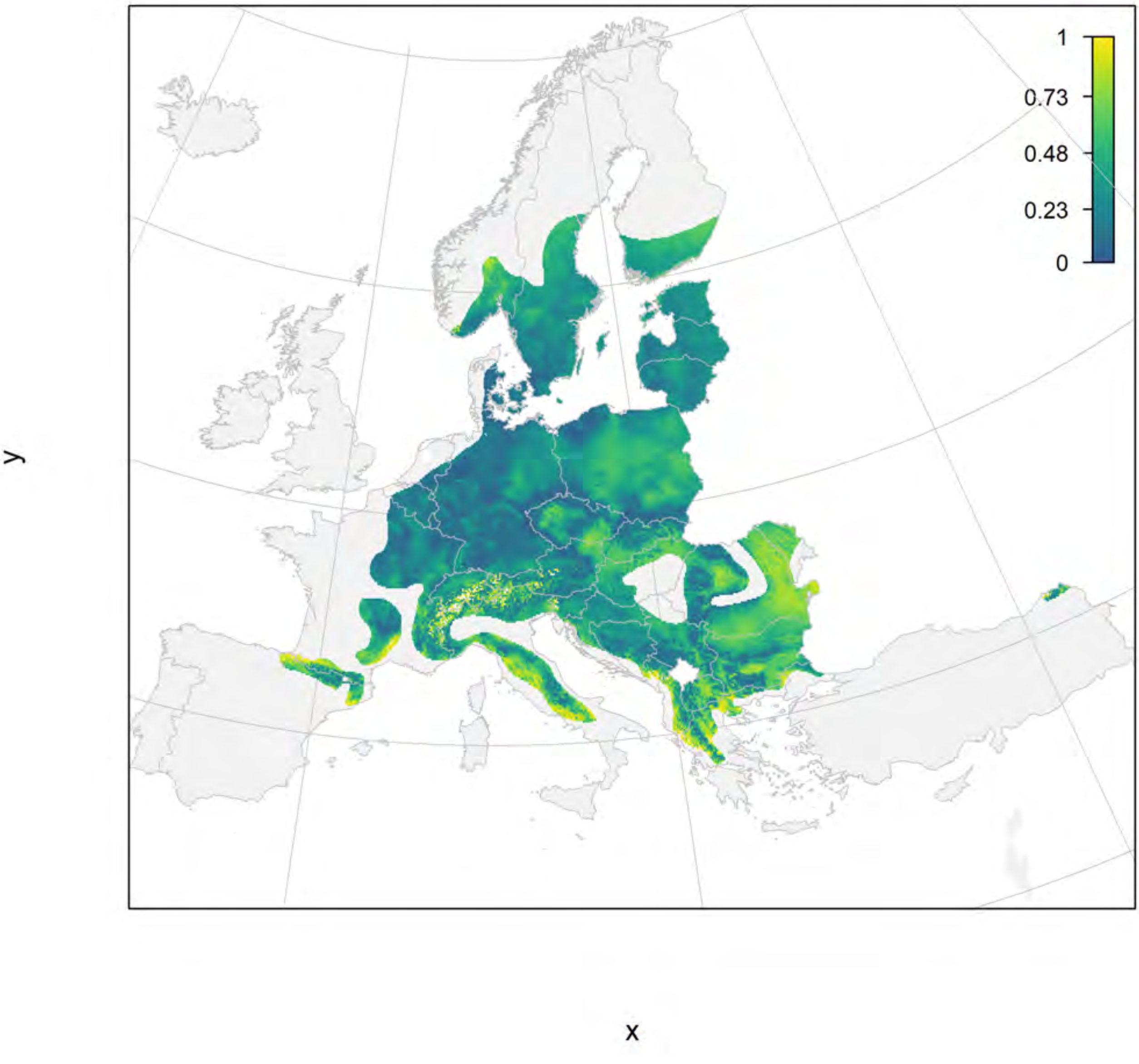

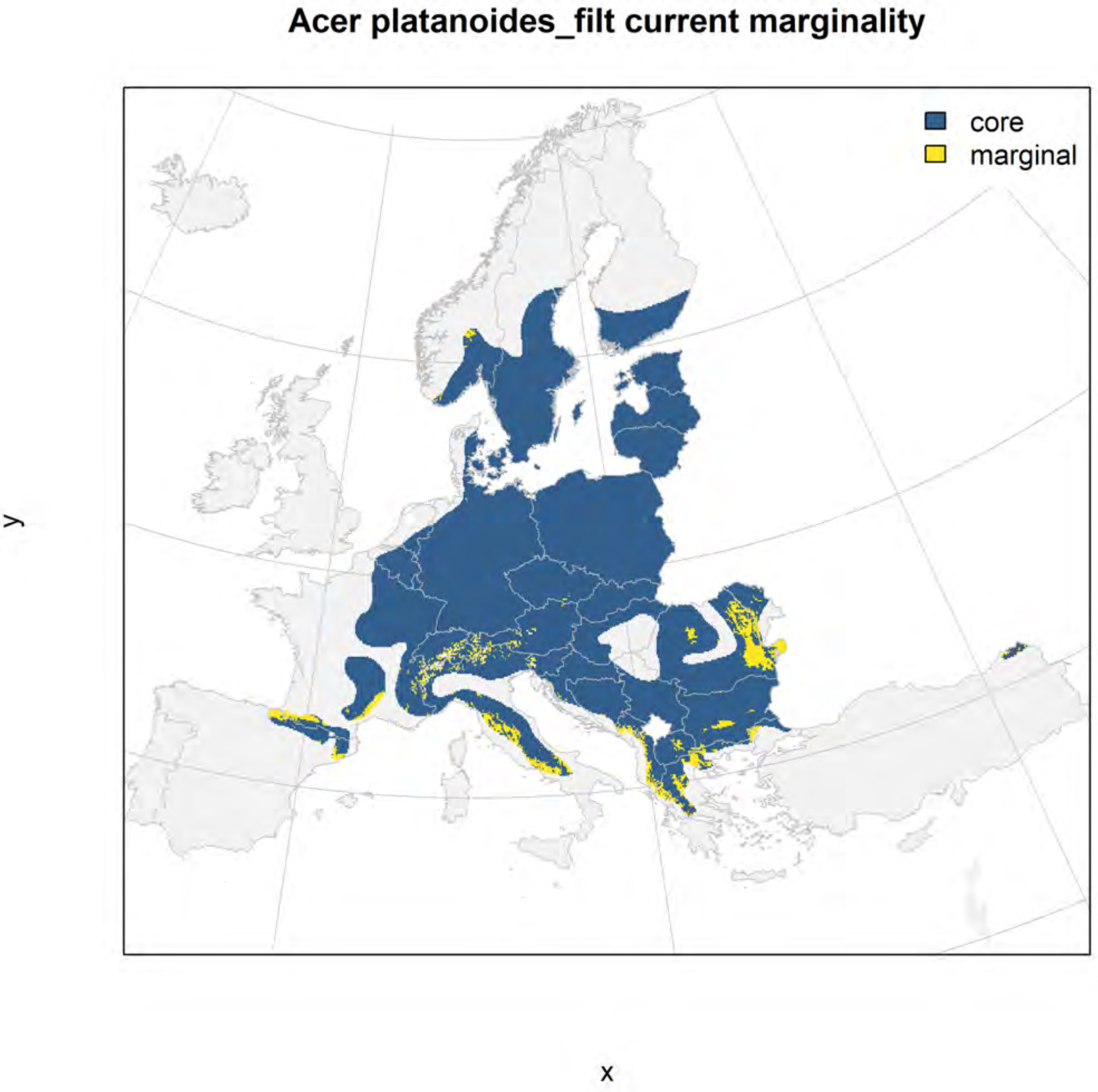
Acer platanoides_filt current marginality
core
marginal
y
x

### Acer platanoides_filt future marginality

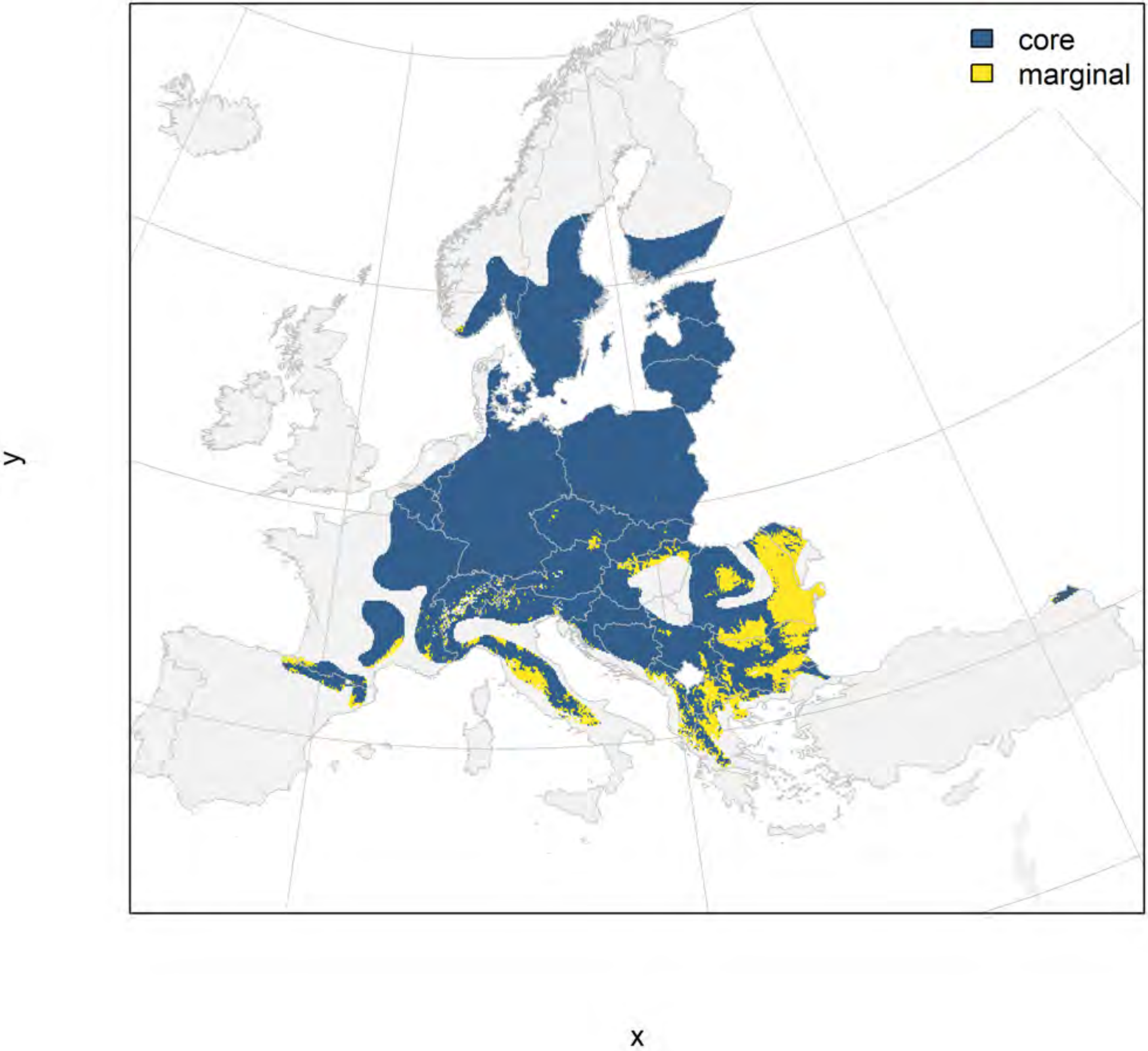

#### Acer pseudoplatanus_filt NMI

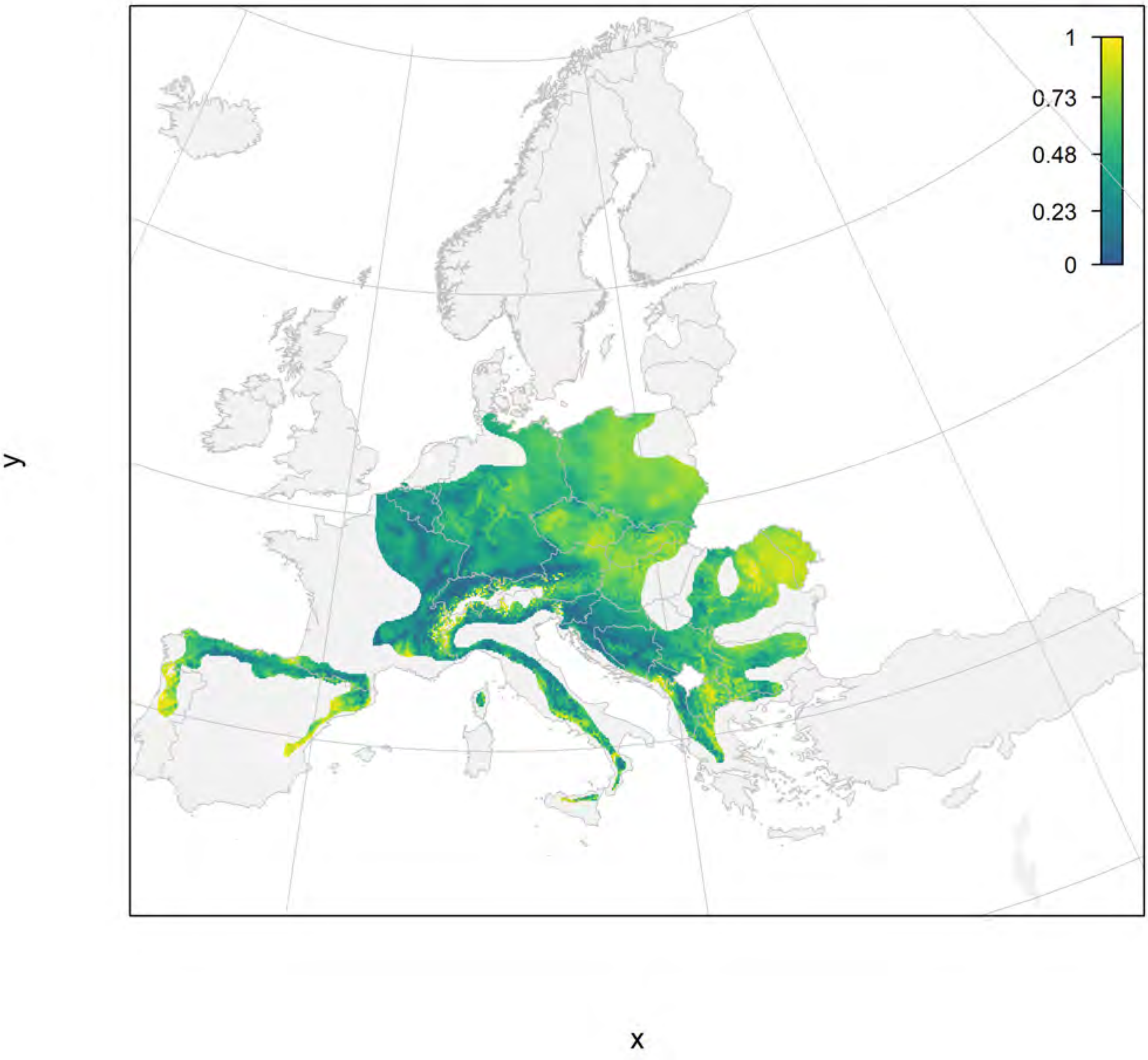

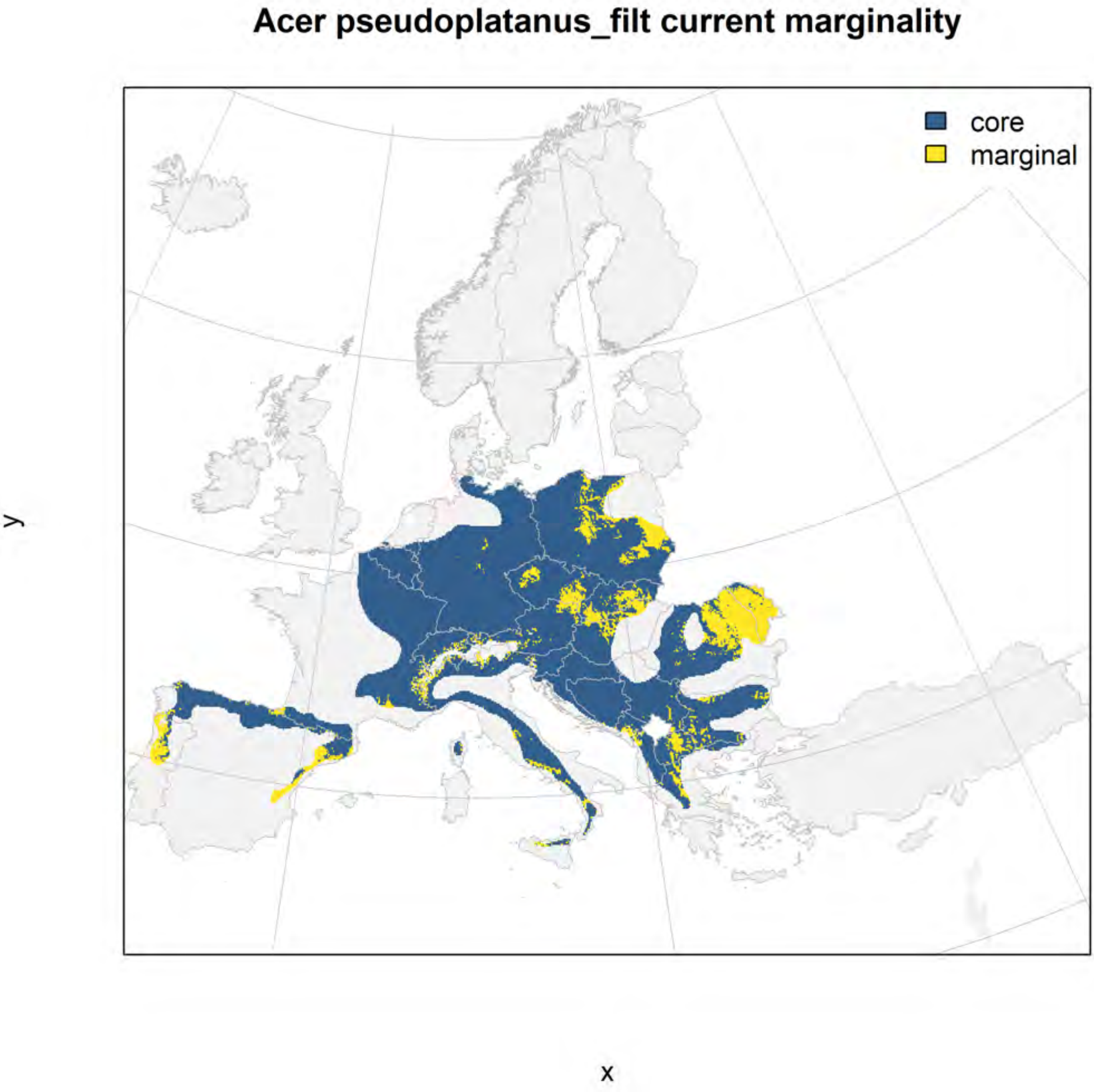
Acer pseudoplatanus_filt current marginality
core
marginal
y
x

### Acer pseudoplatanus_filt future marginality

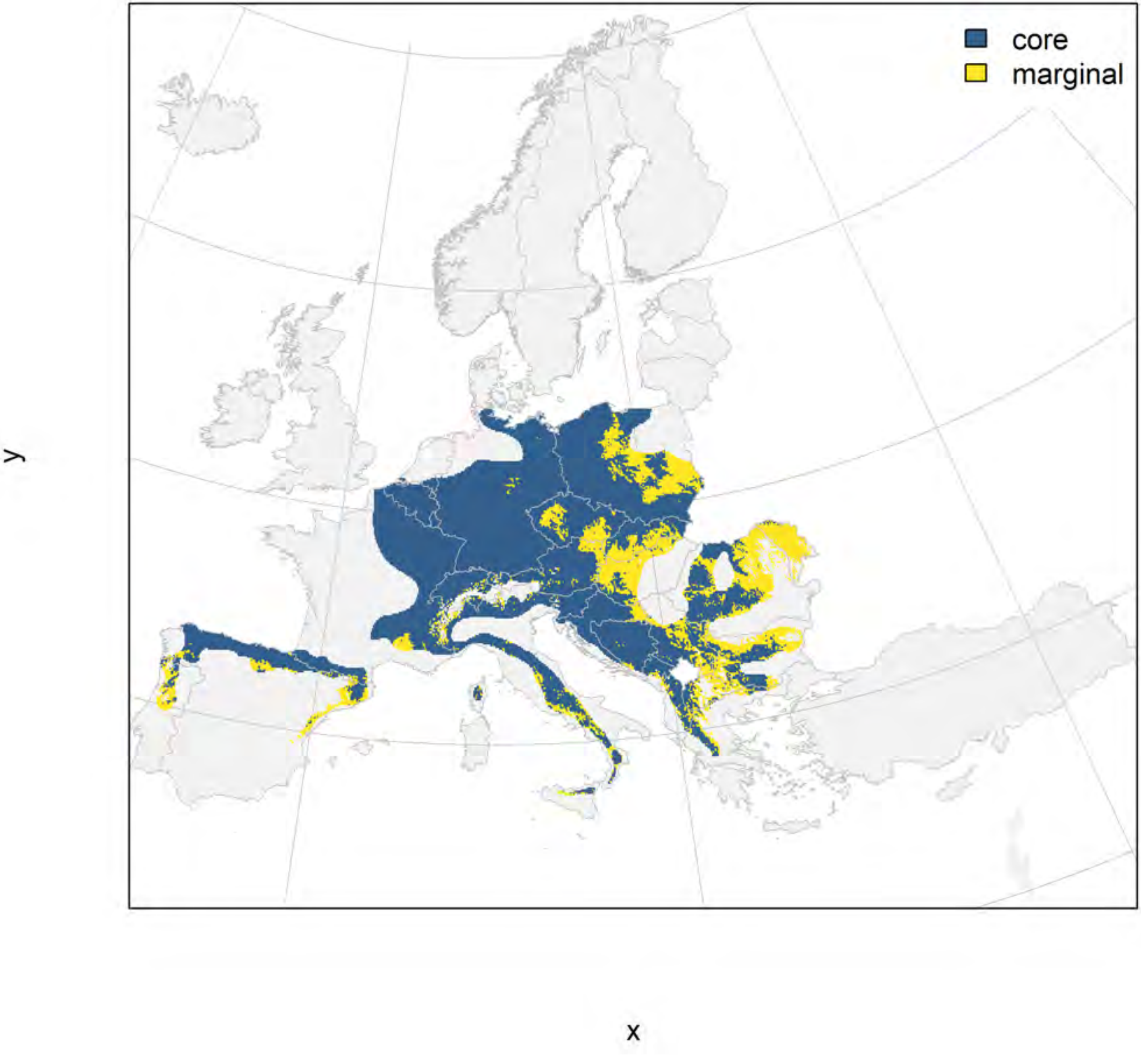

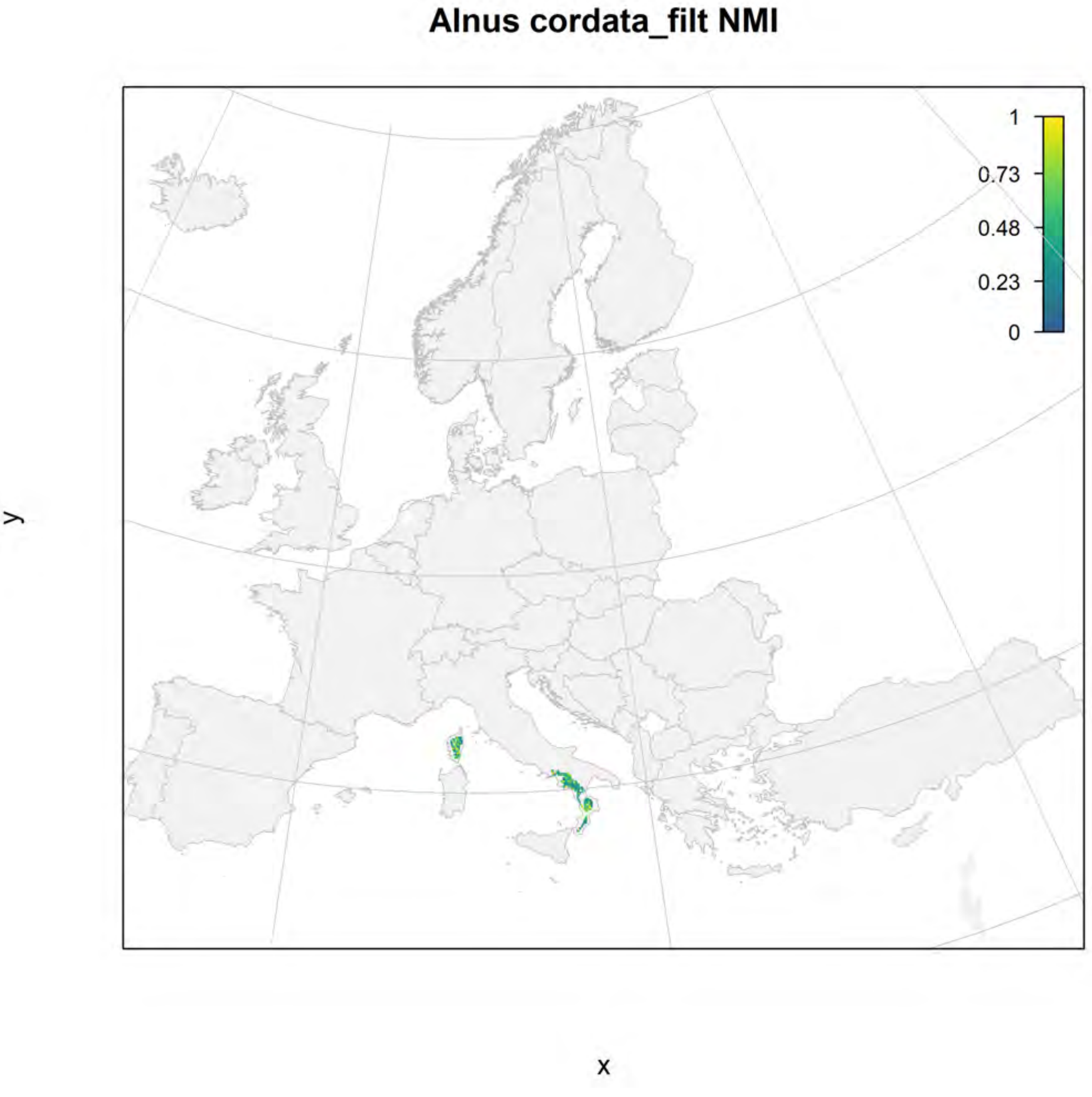

Alnus cordata_filt NMI
1
0.73
0.48
0.23
0
y
x

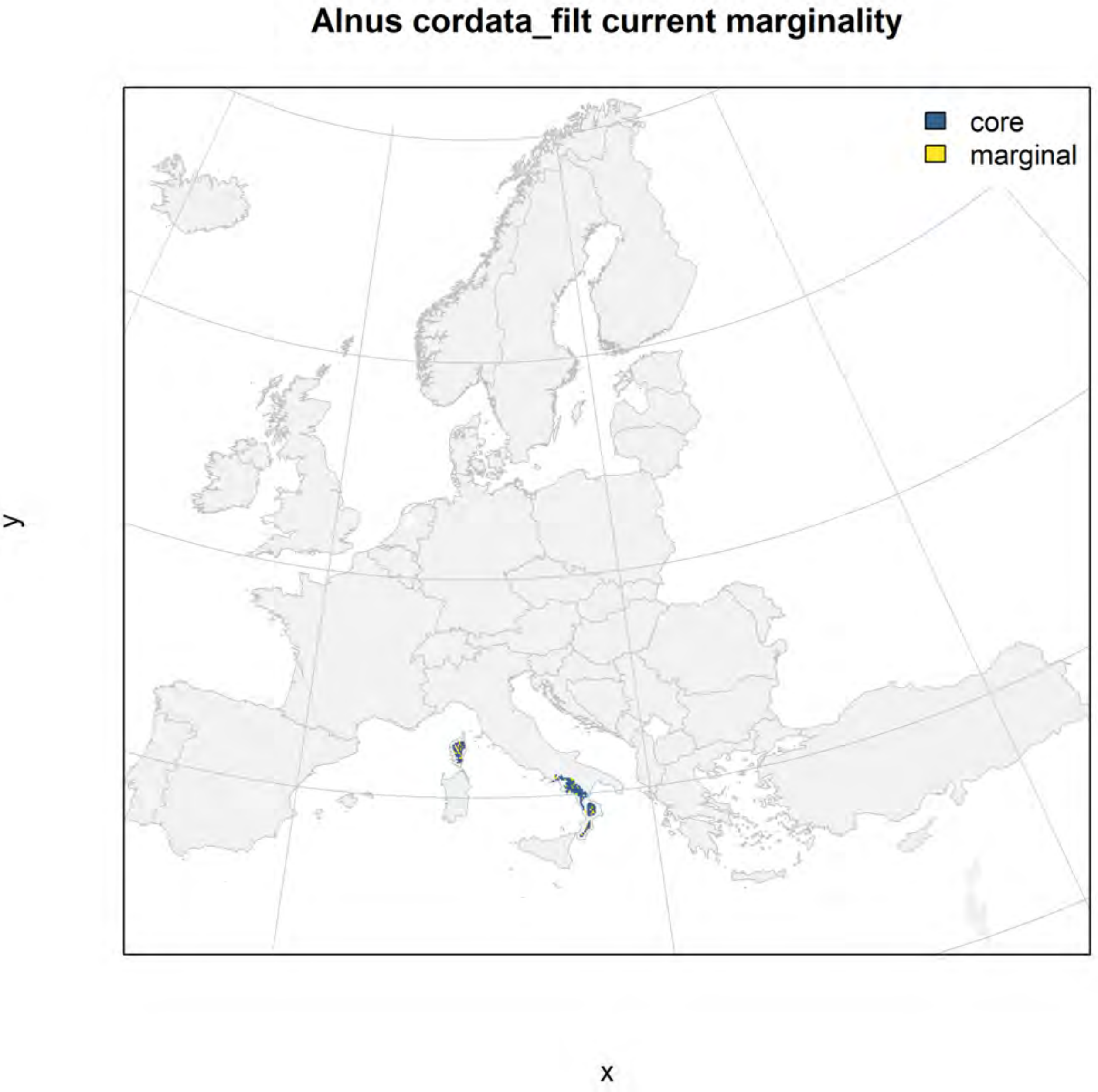
Alnus cordata_filt current marginality
core
marginal
y
x

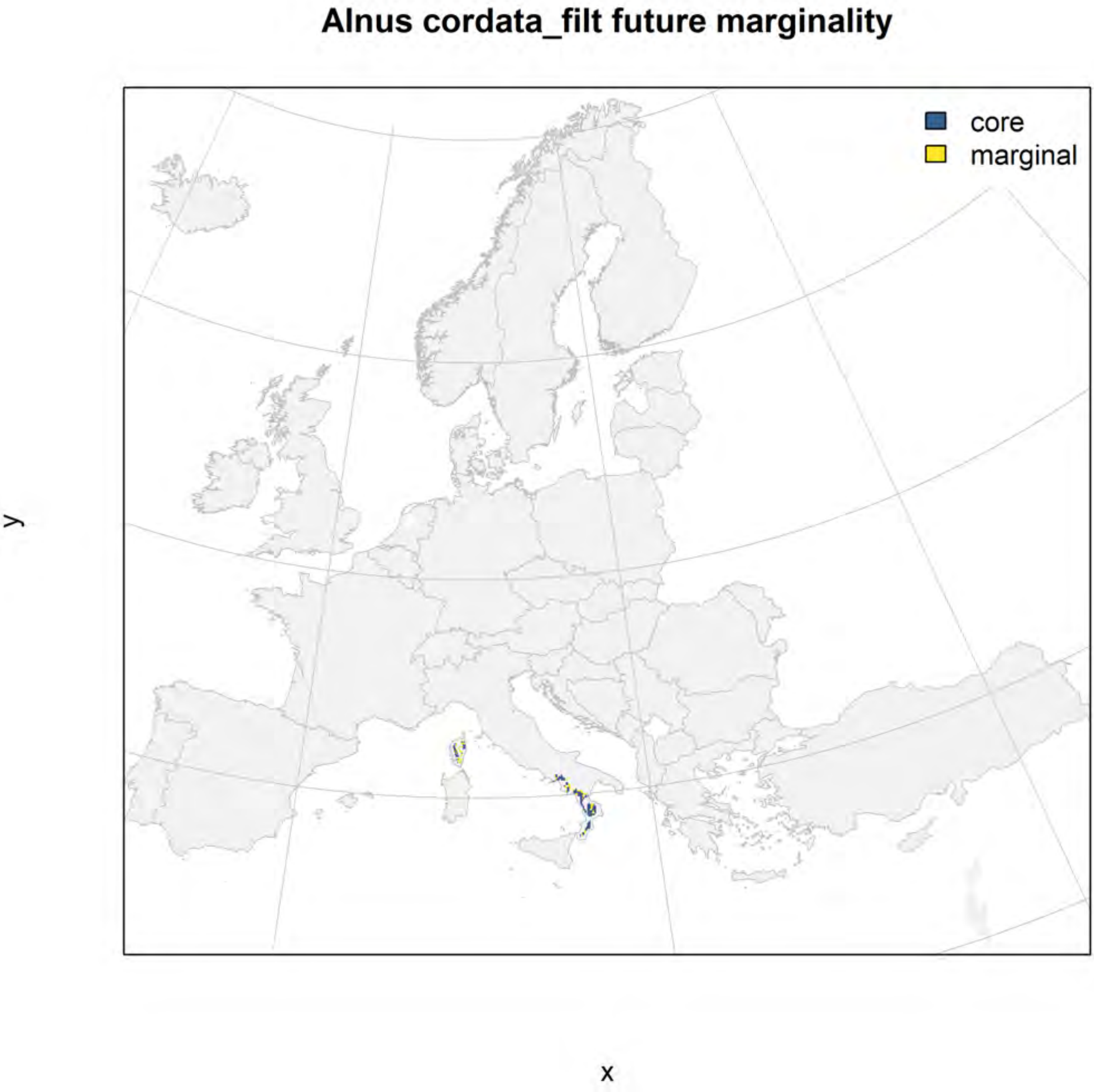
Alnus cordata_filt future marginality
core
marginal
y
x

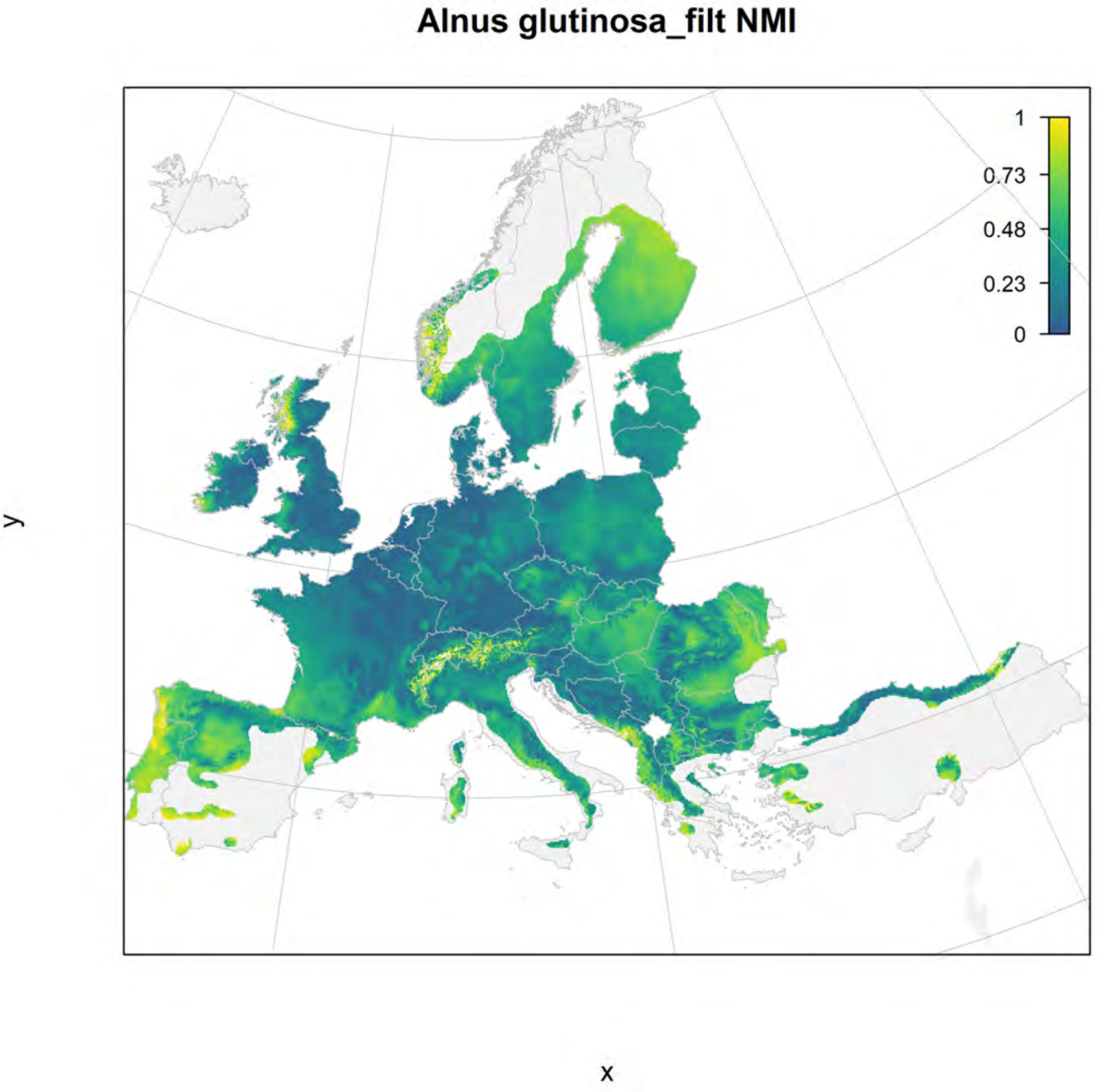

Alnus glutinosa_filt NMI
1
0.73
0.48
0.23
0
y
x

**Alnus glutinosa_filt current marginality**

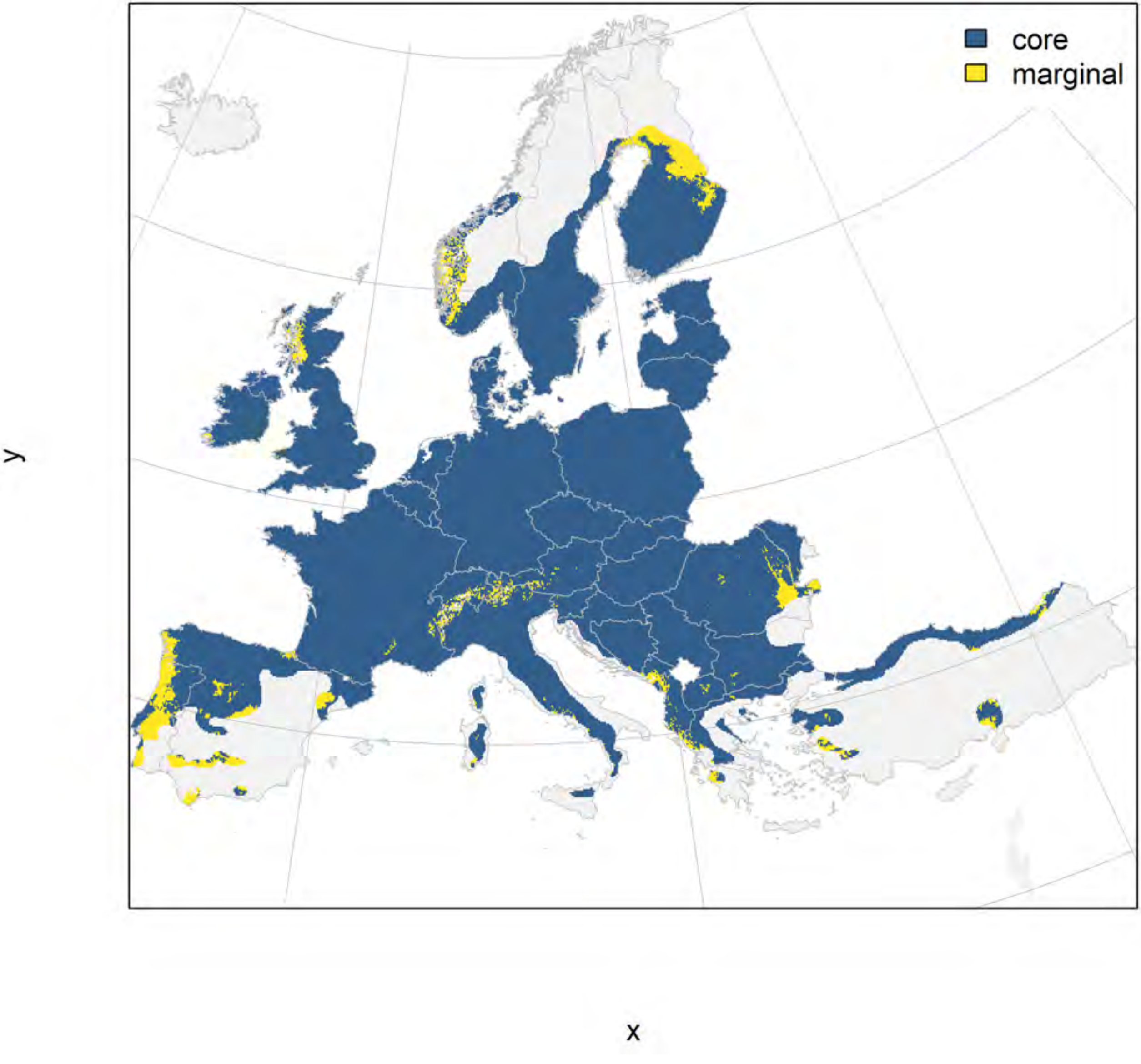

### Alnus glutinosa_filt future marginality

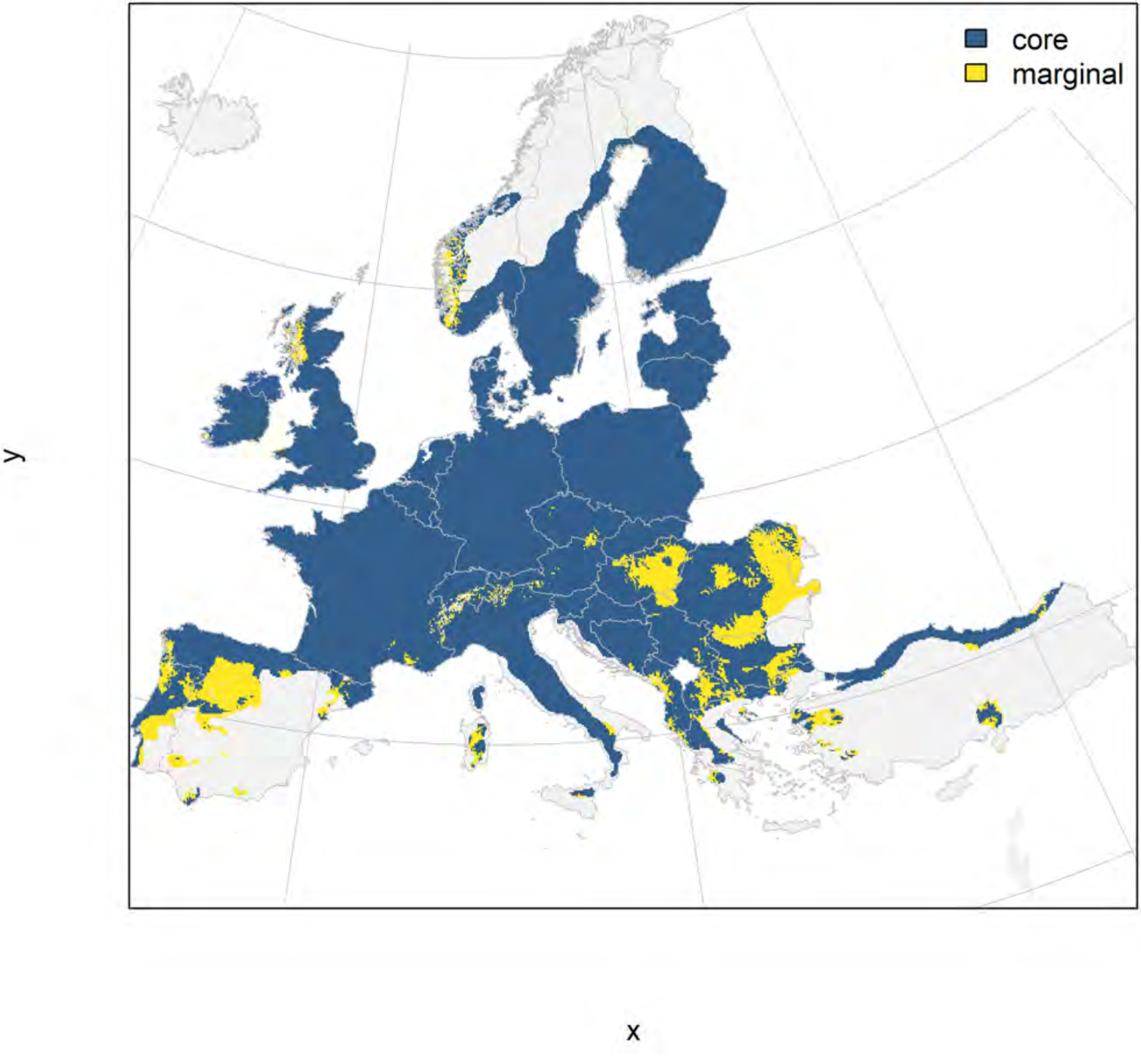

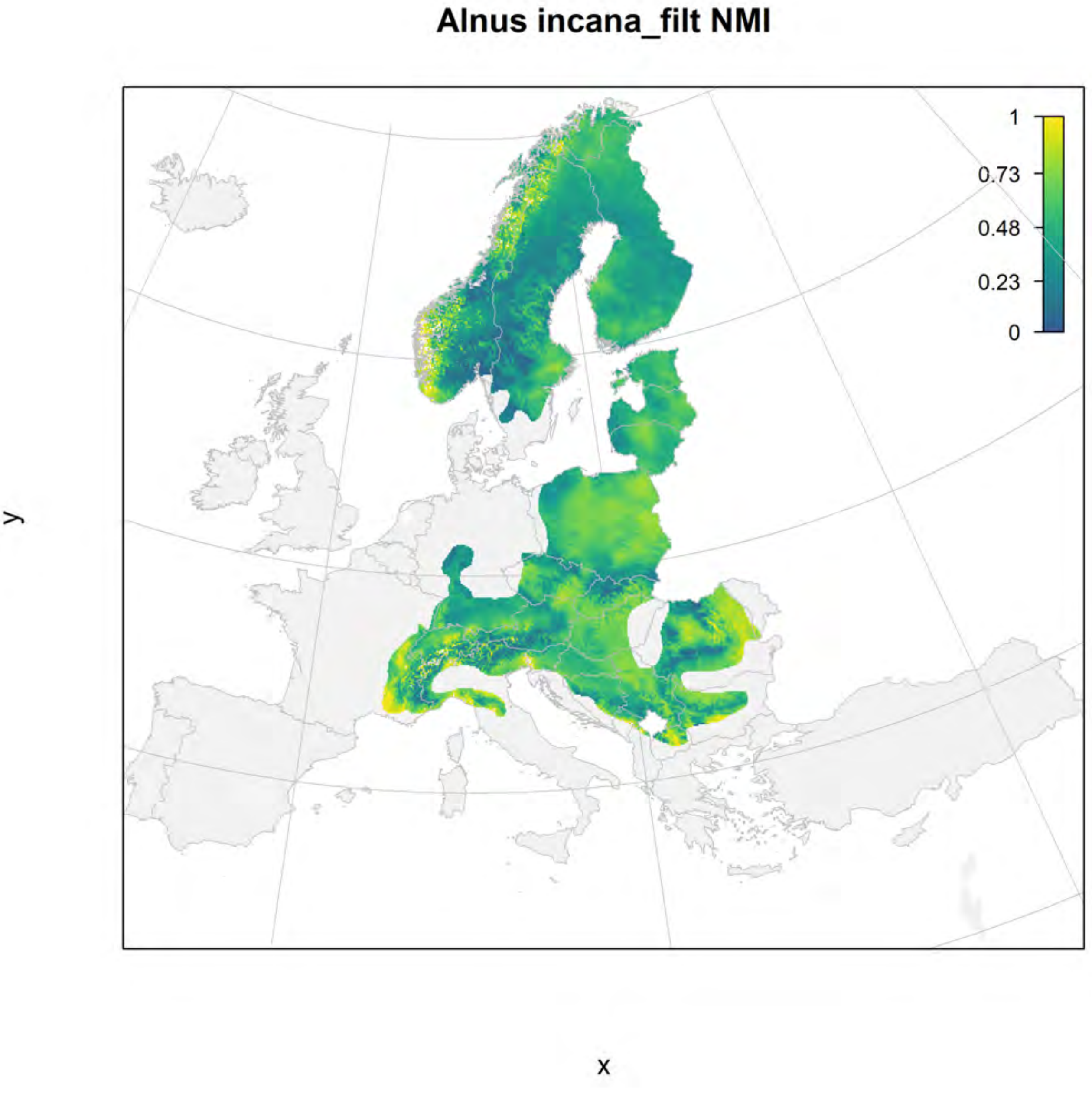
Alnus incana_filt NMI
1
0.73
0.48
0.23
0
y
x

#### Alnus incana_filt current marginality

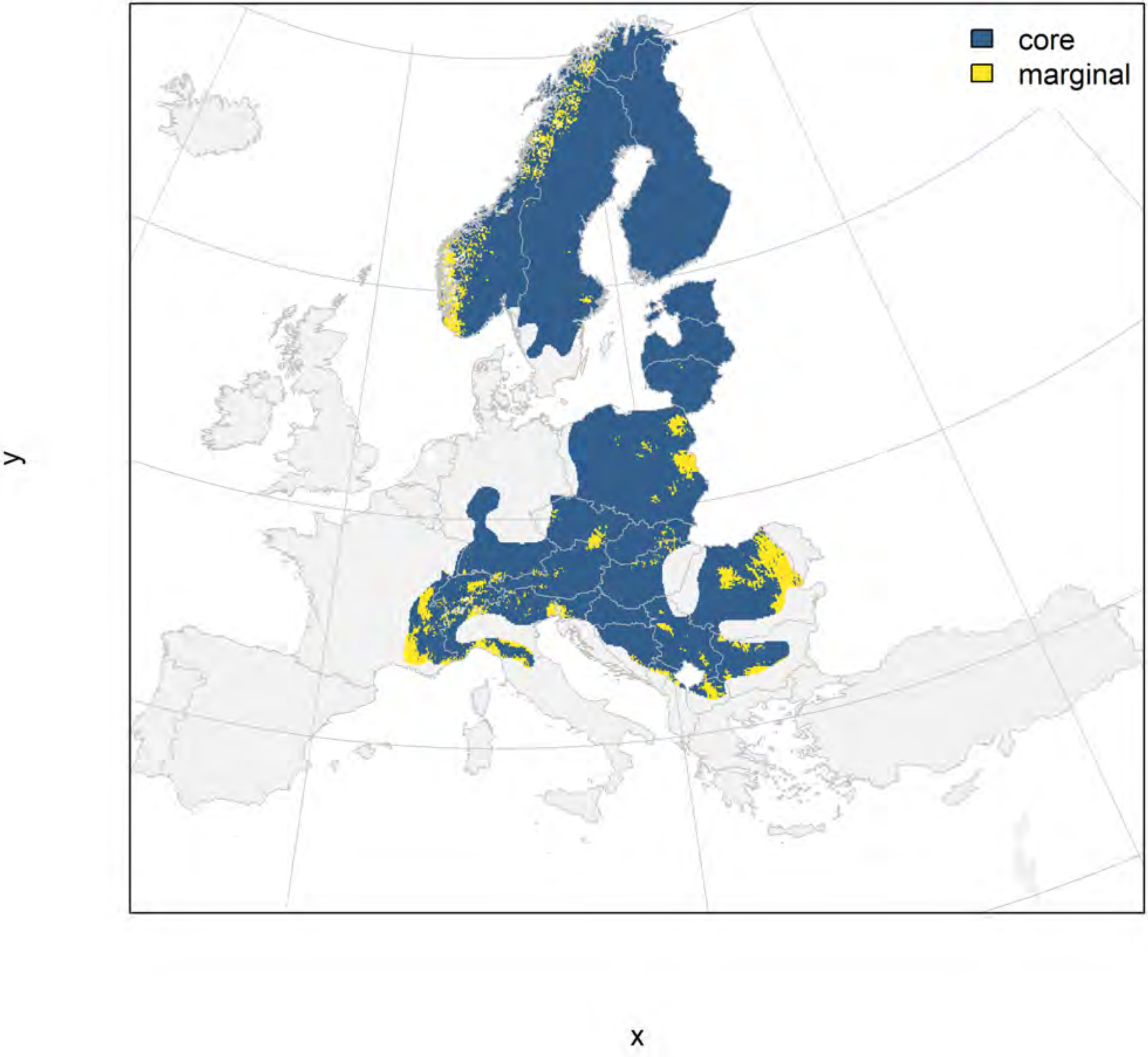

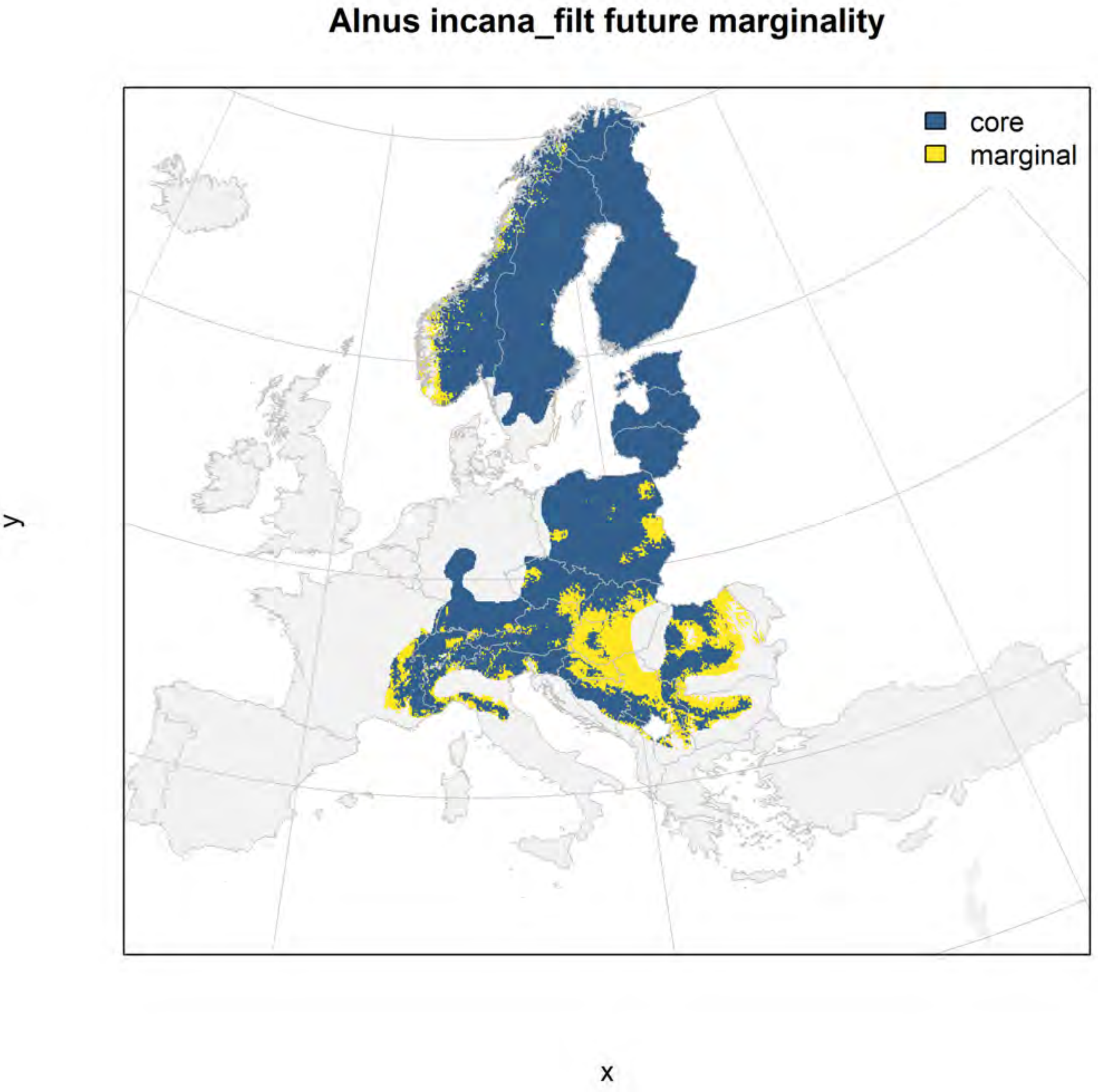
Alnus incana_filt future marginality
core
marginal
y
x

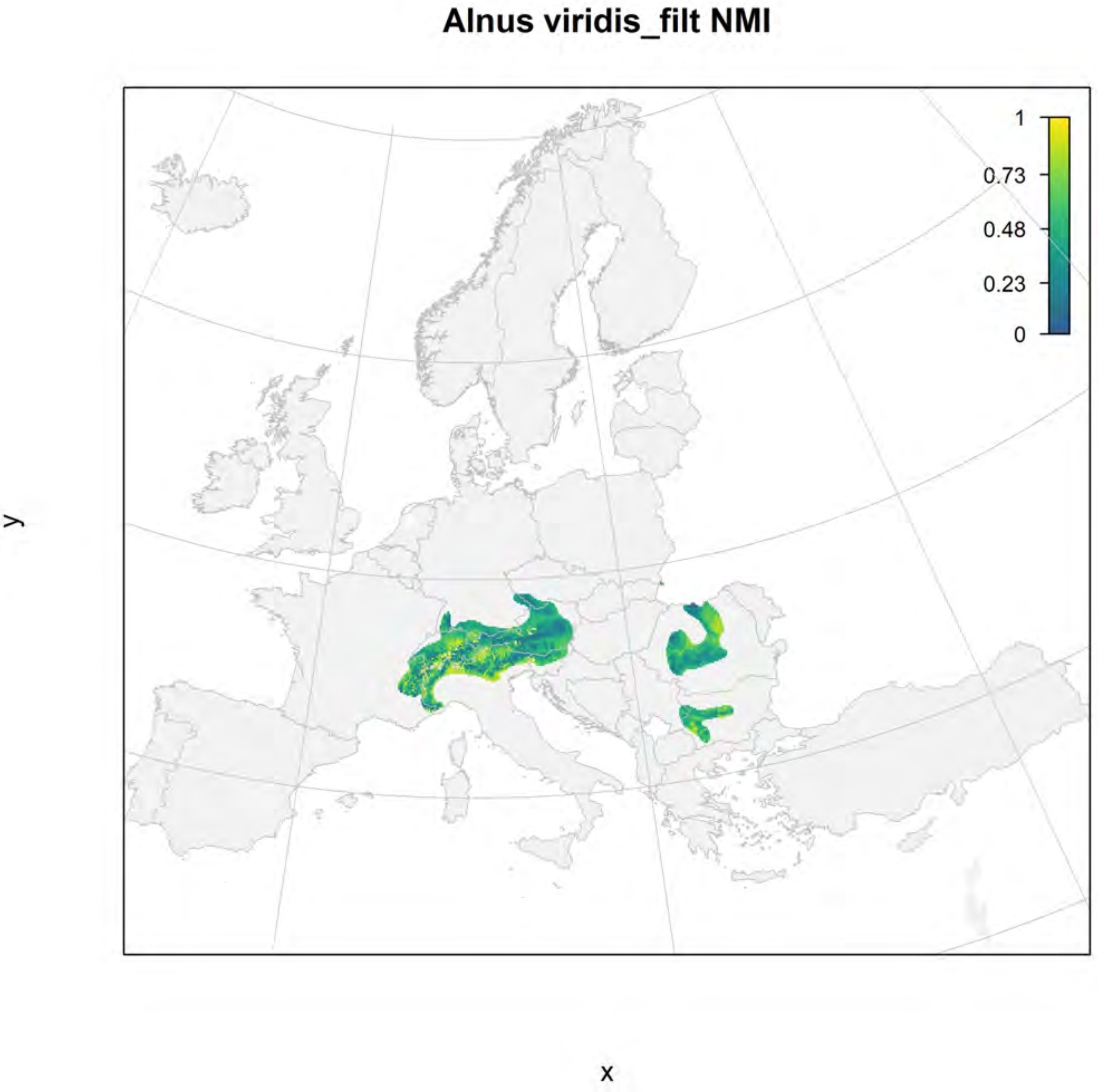
Alnus viridis_filt NMI
1
0.73
0.48
0.23
0
y
x

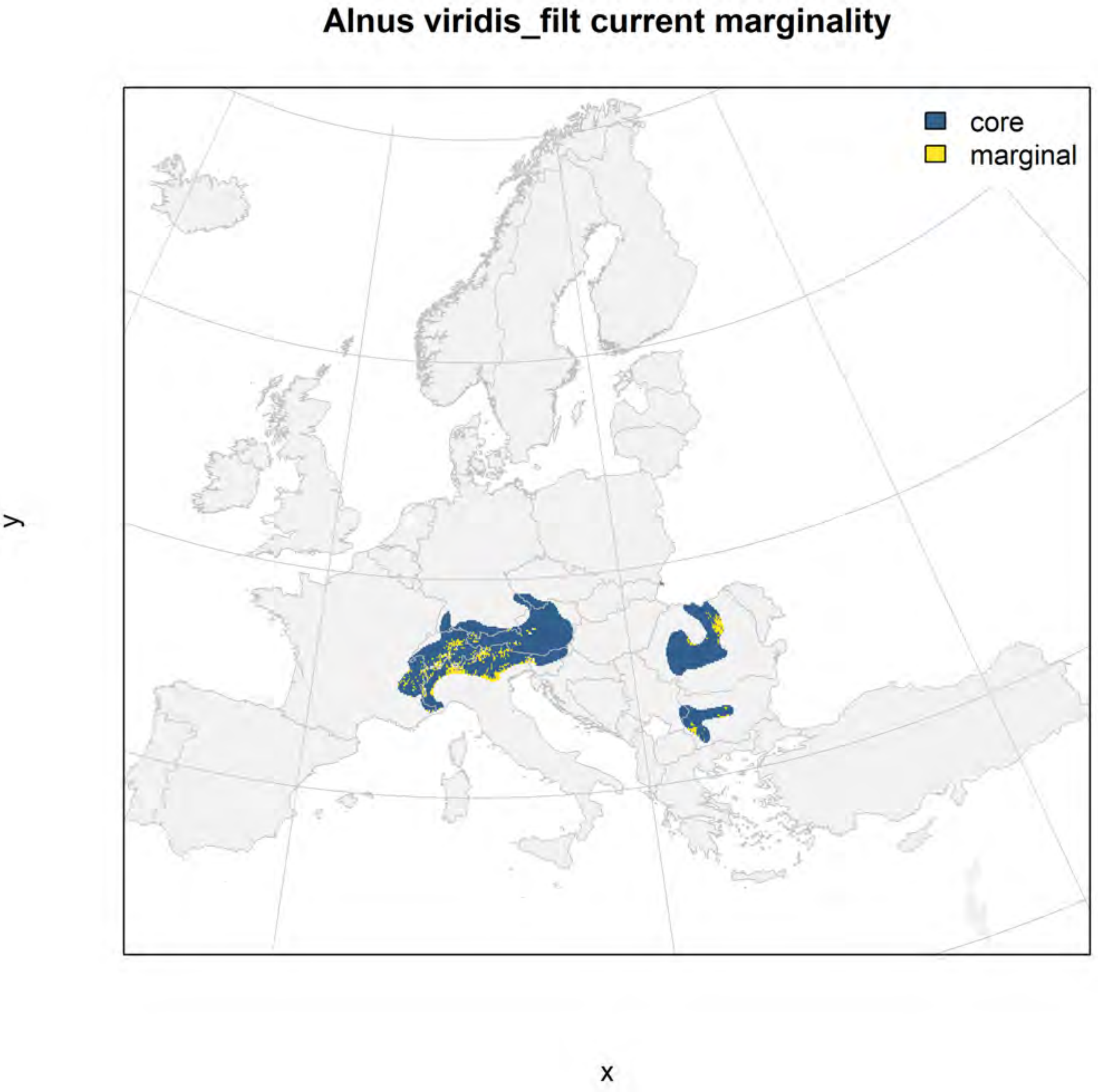
Alnus viridis_filt current marginality
core
marginal
y
x

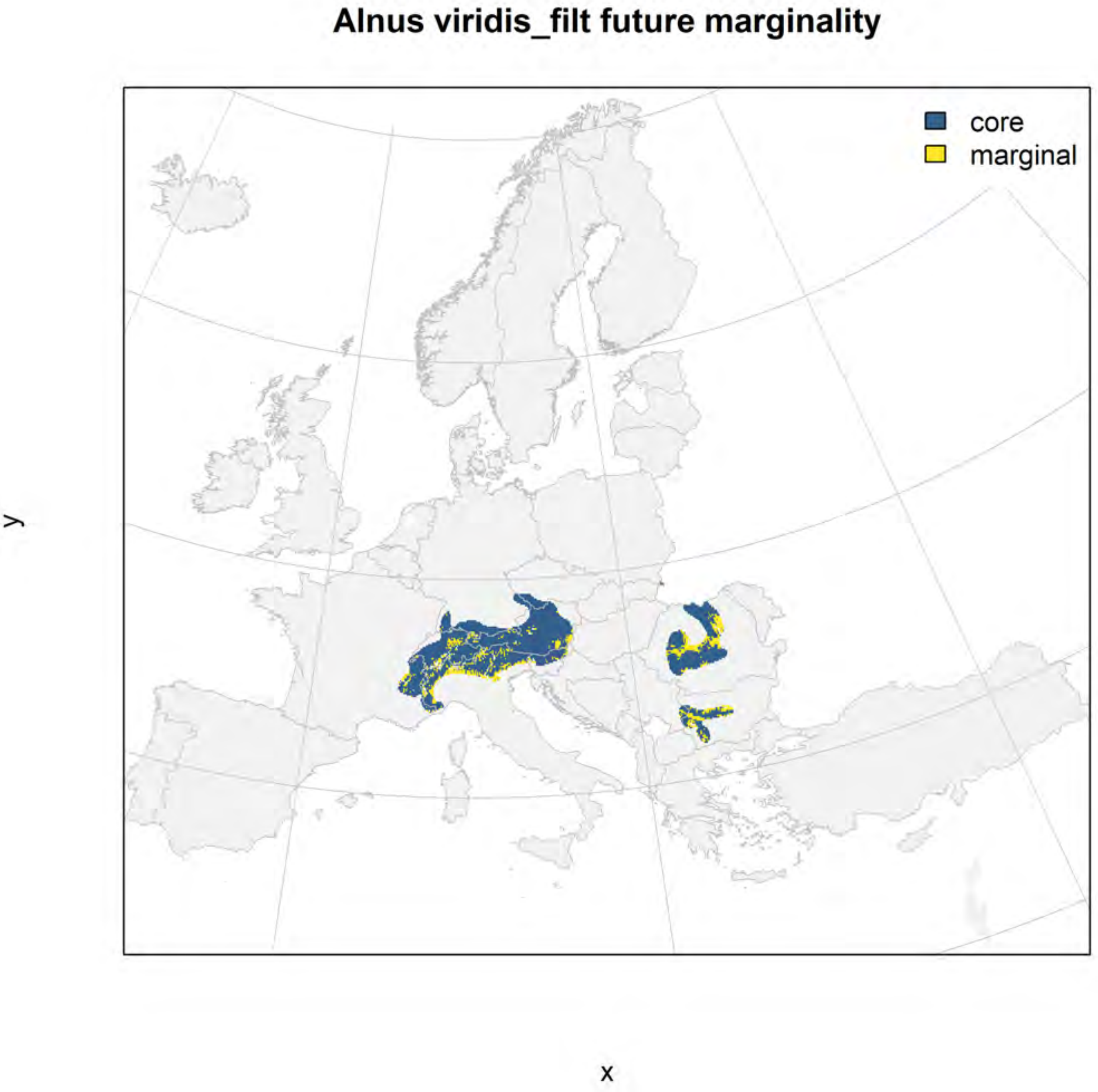
Alnus viridis_filt future marginality
core
marginal
y
x

### Betula pendula_filt NMI

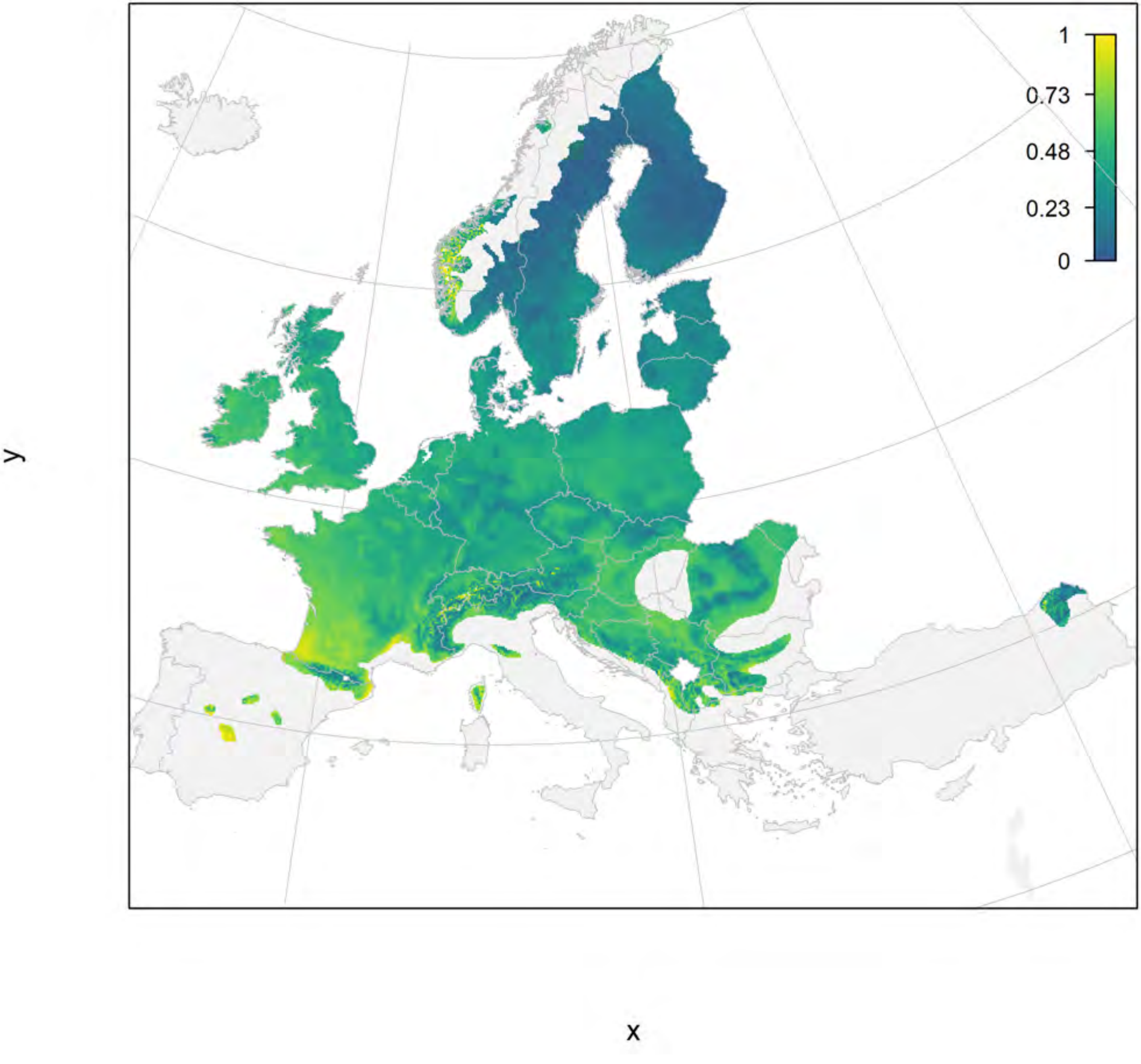

### Betula pendula_filt current marginality

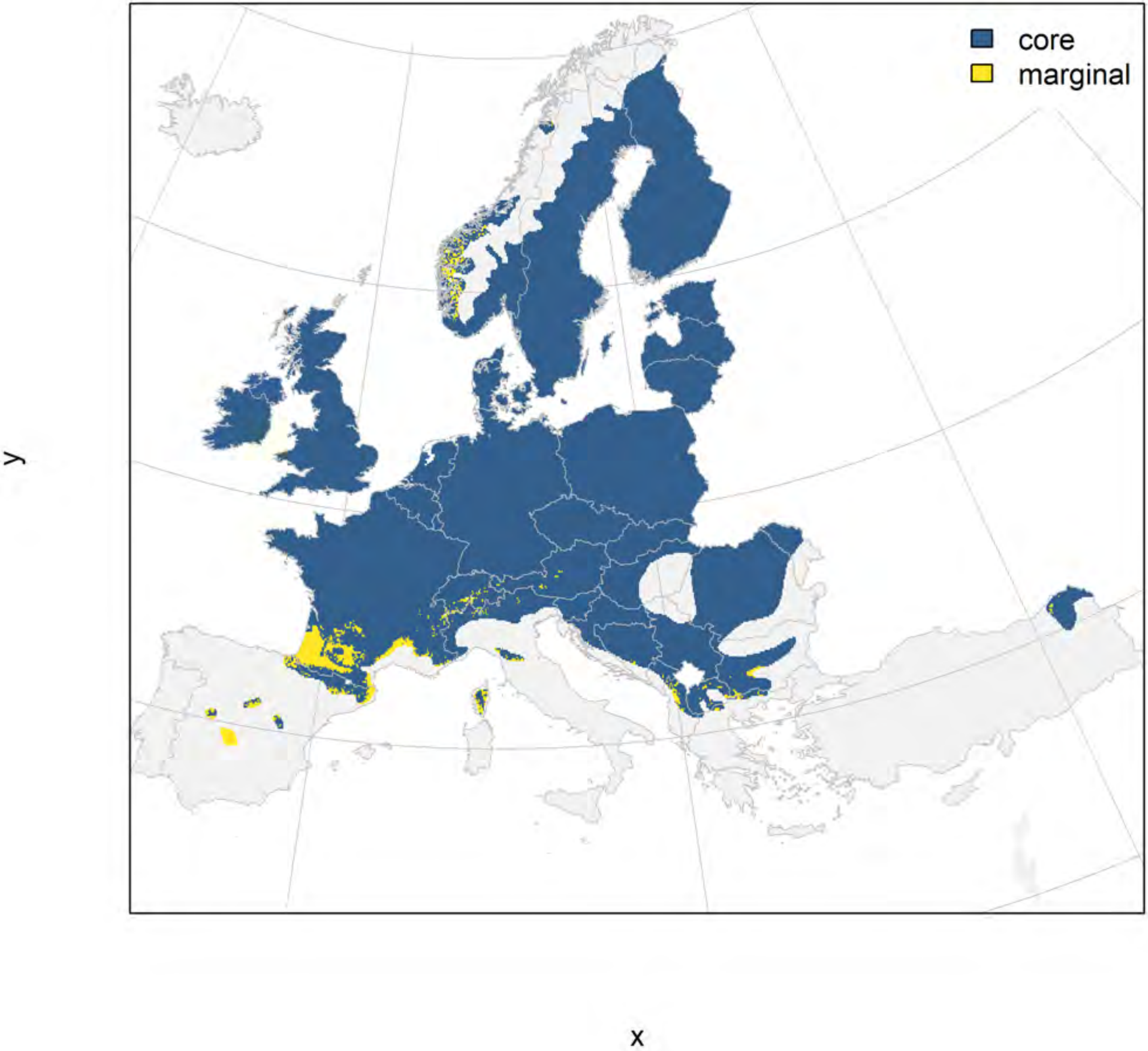

### Betula pendula_filt future marginality

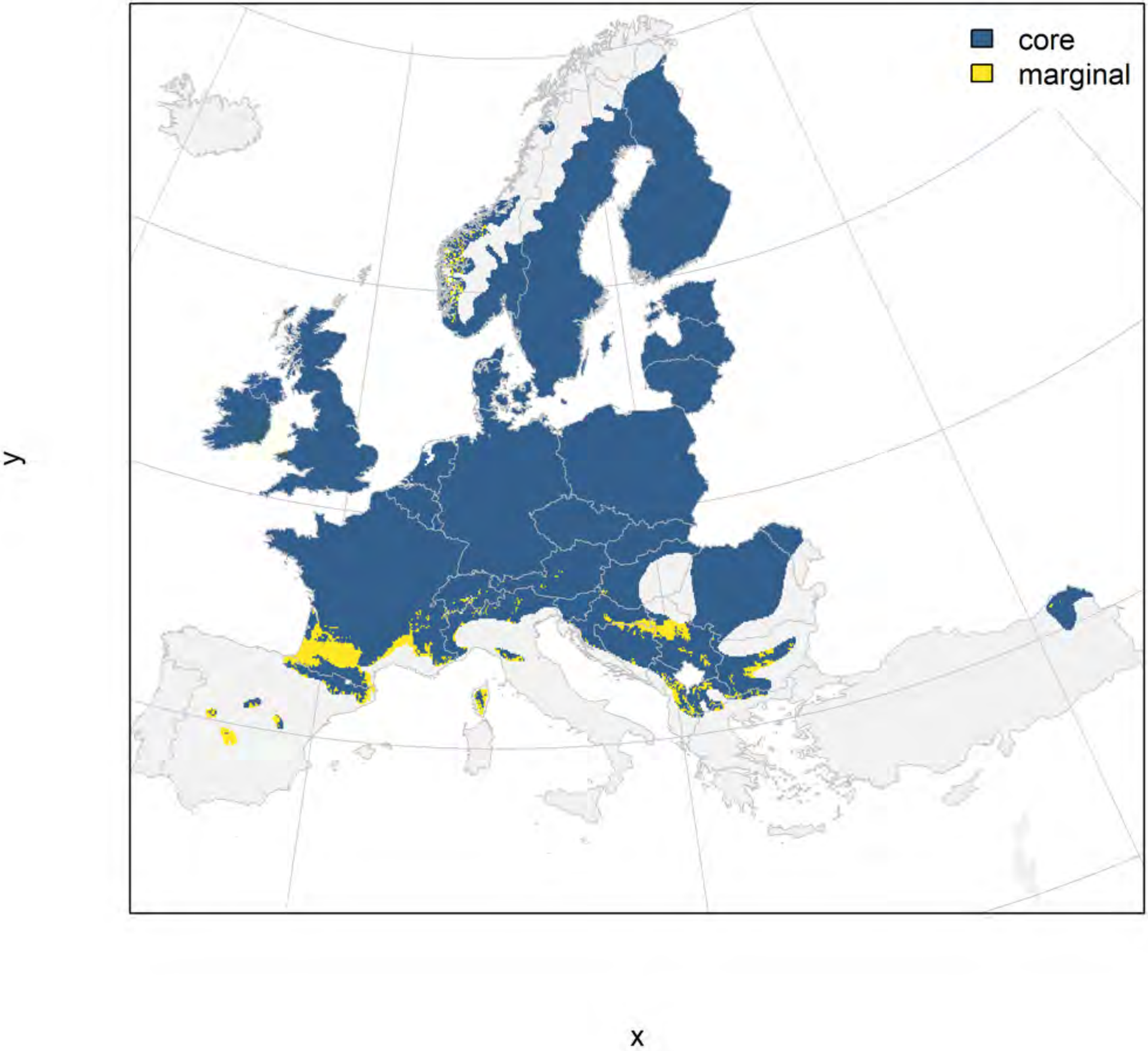

### Betula pubescens_filt NMI

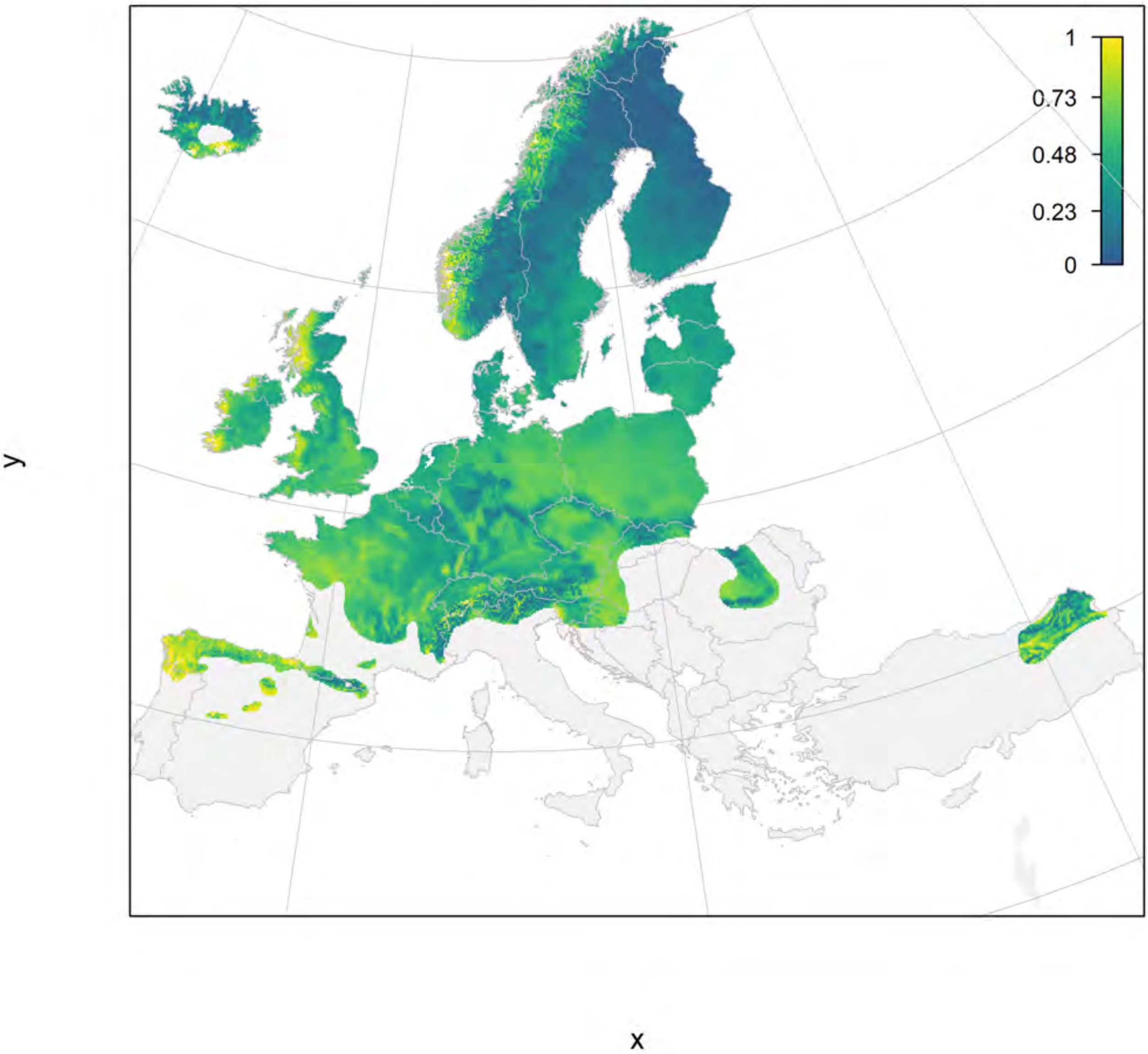

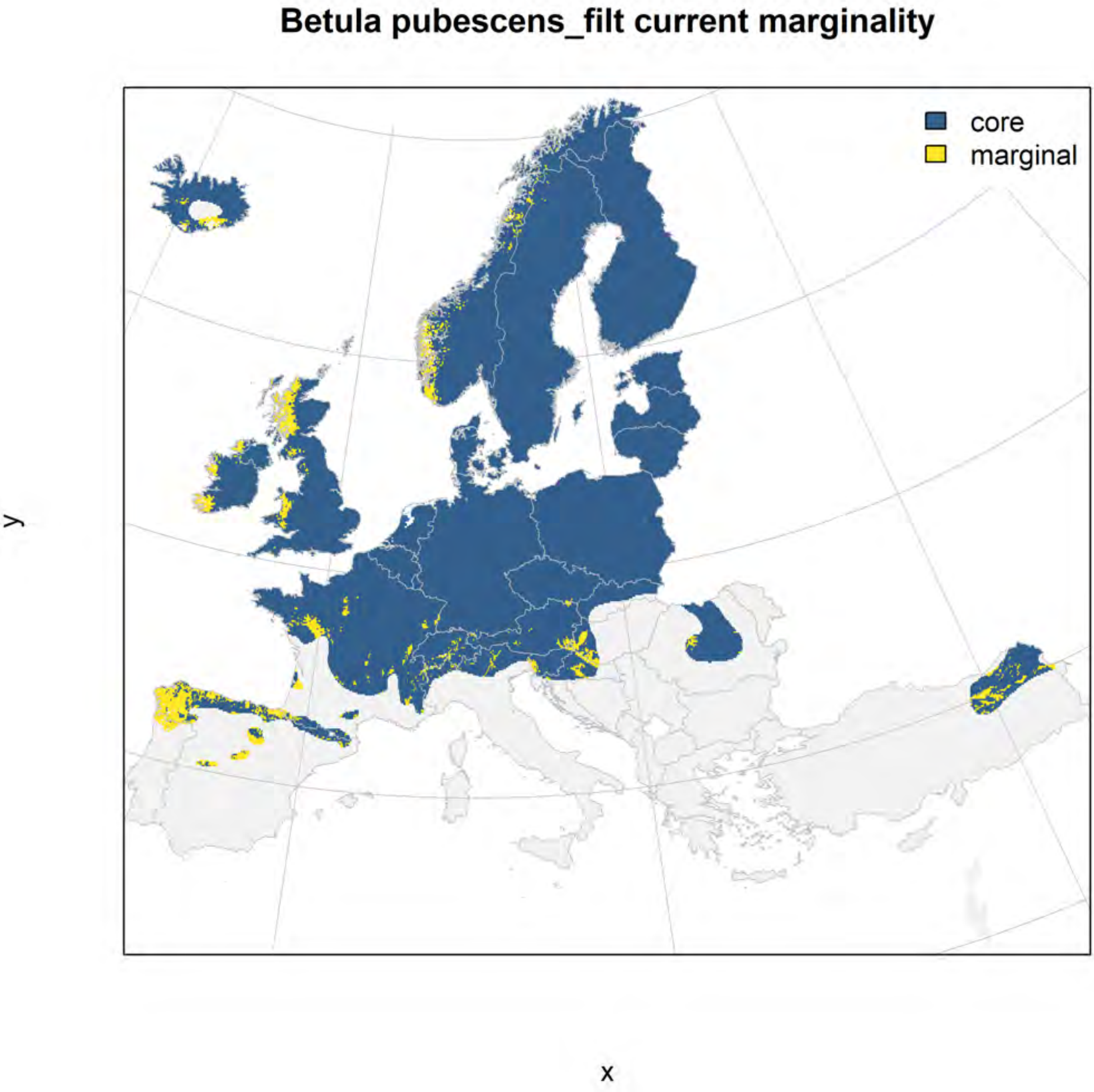
Betula pubescens_filt current marginality
core
marginal
y
x

### Betula pubescens_filt future marginality

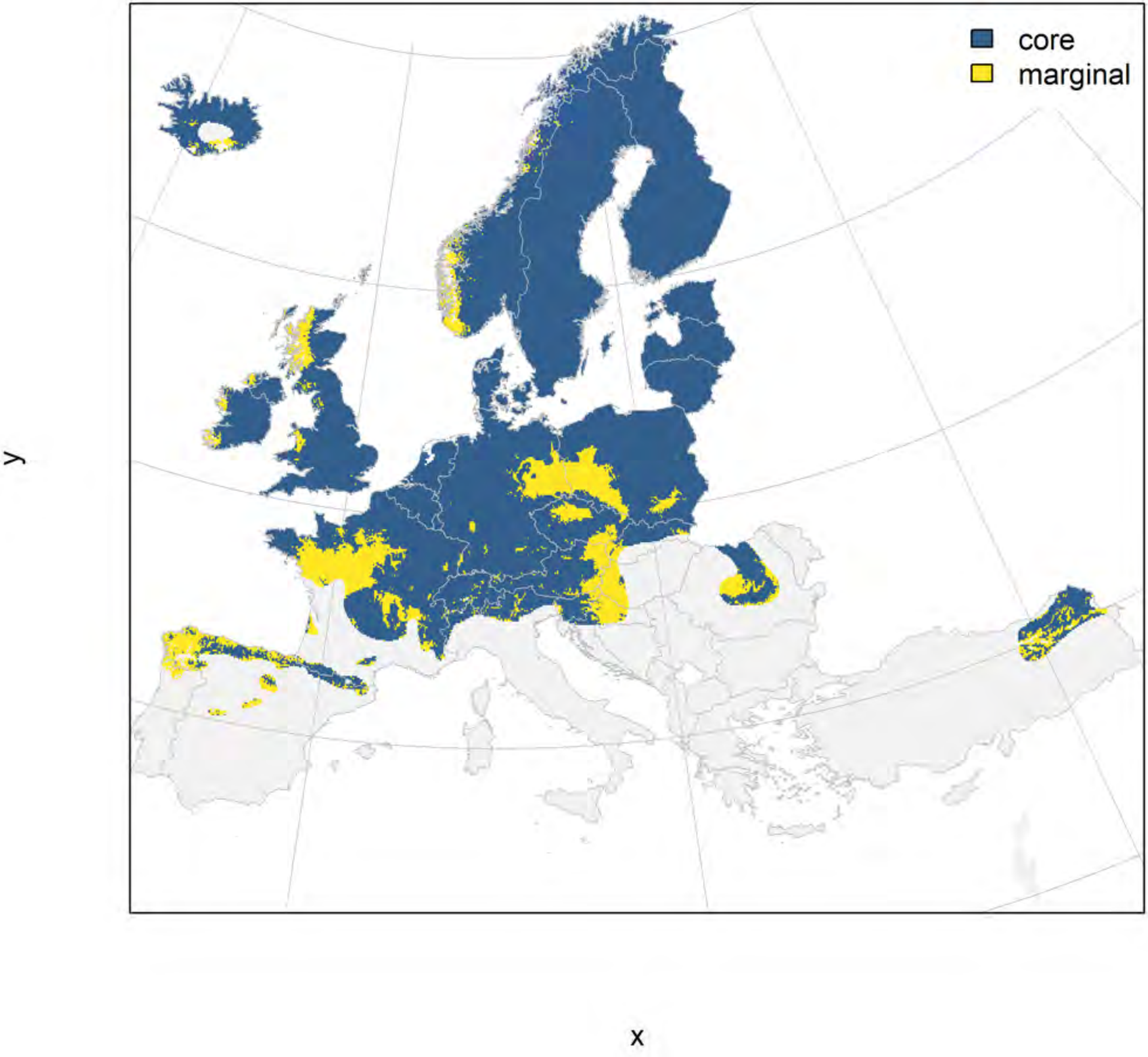

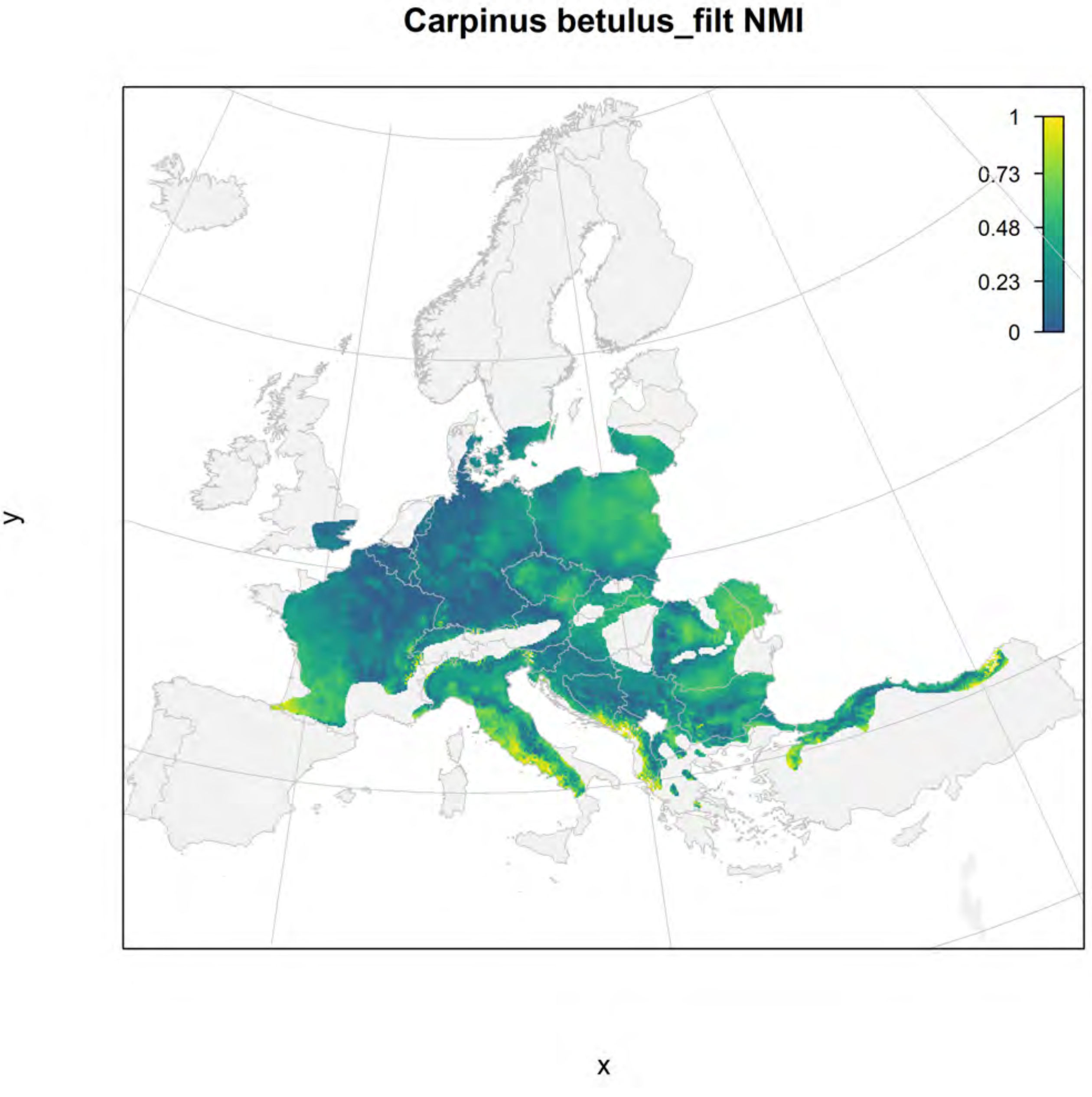
Carpinus betulus_filt NMI
1
0.73
0.48
0.23
0
y
x

#### Carpinus betulus_filt current marginality

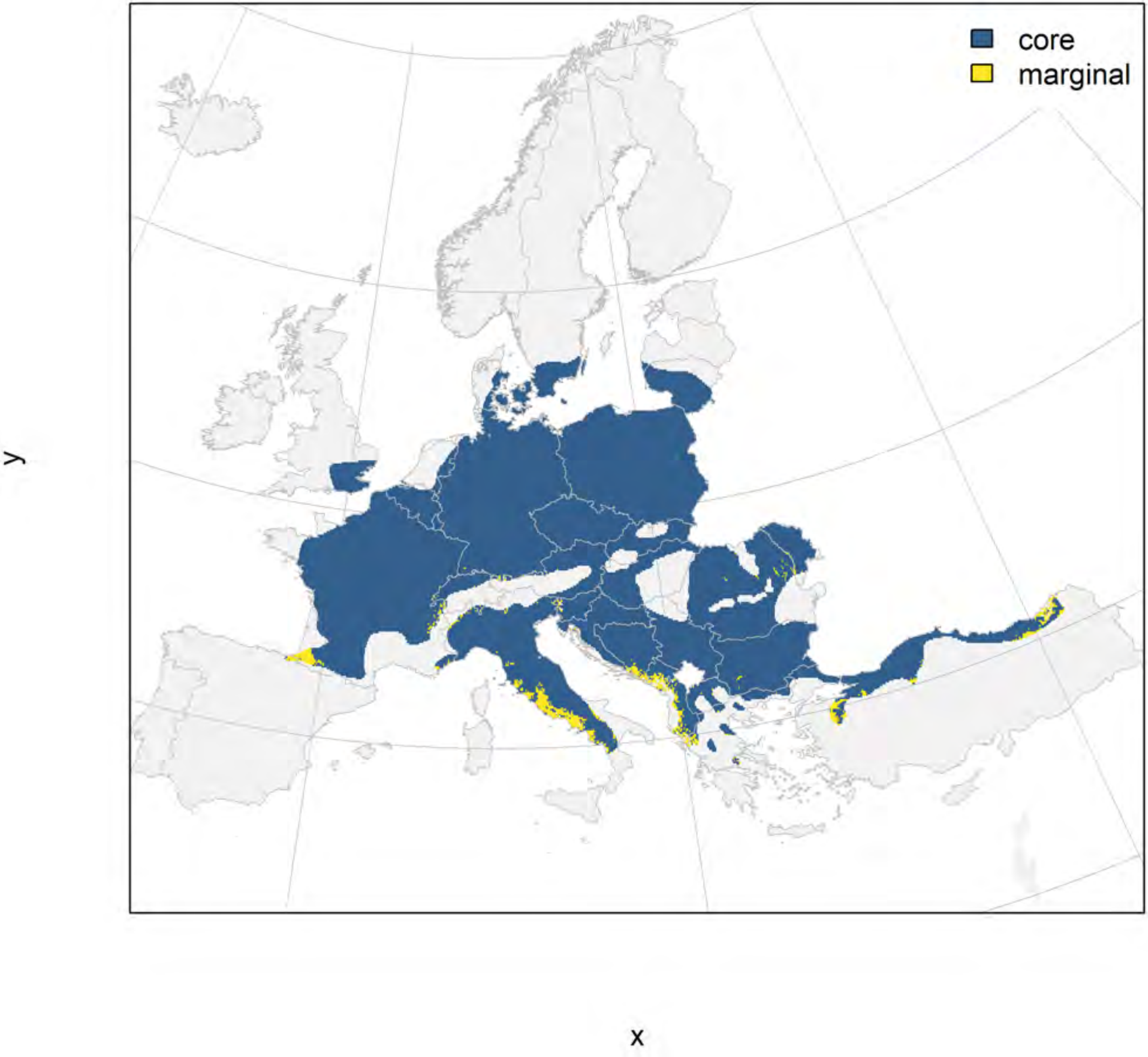

#### Carpinus betulus_filt future marginality

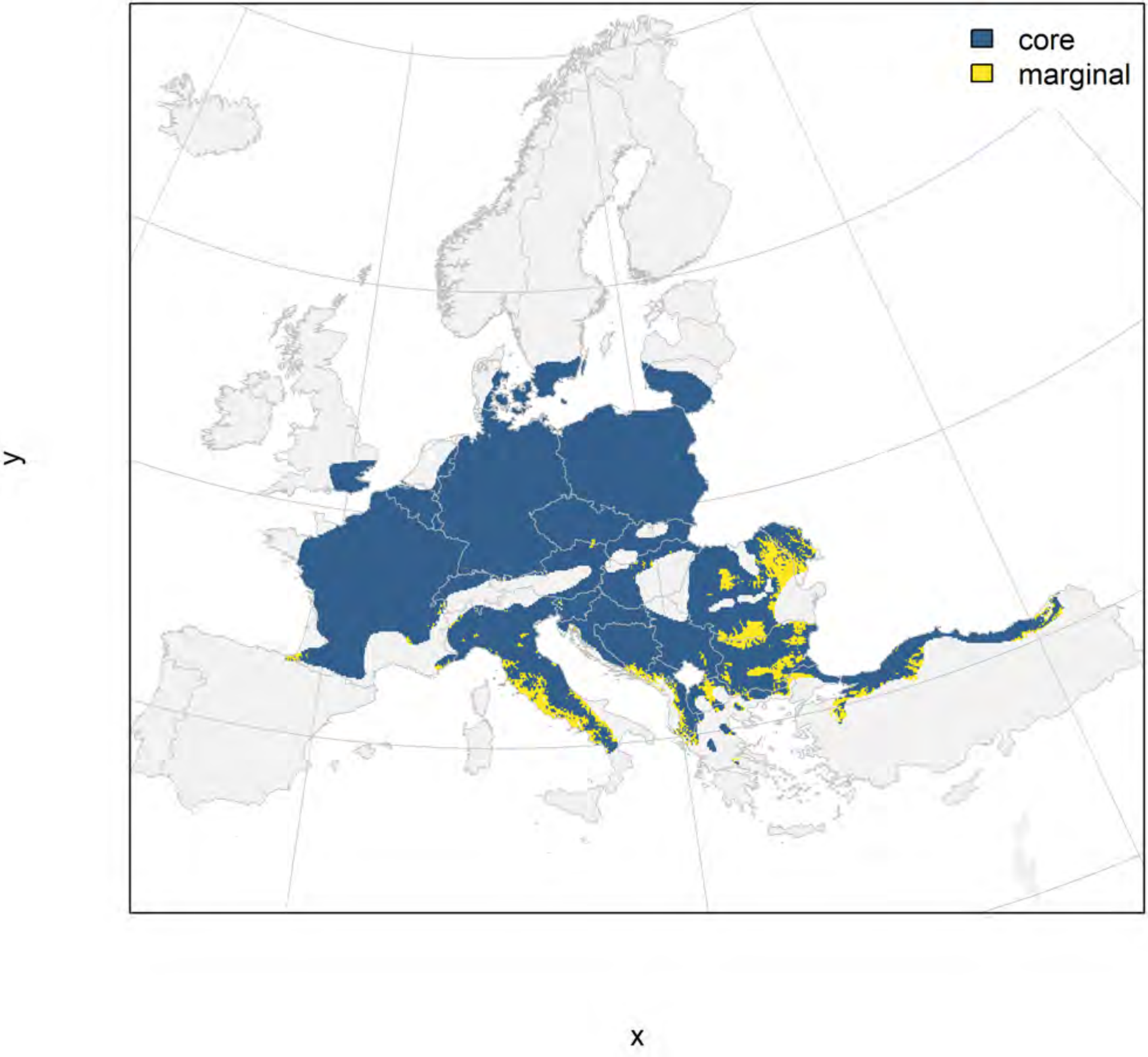

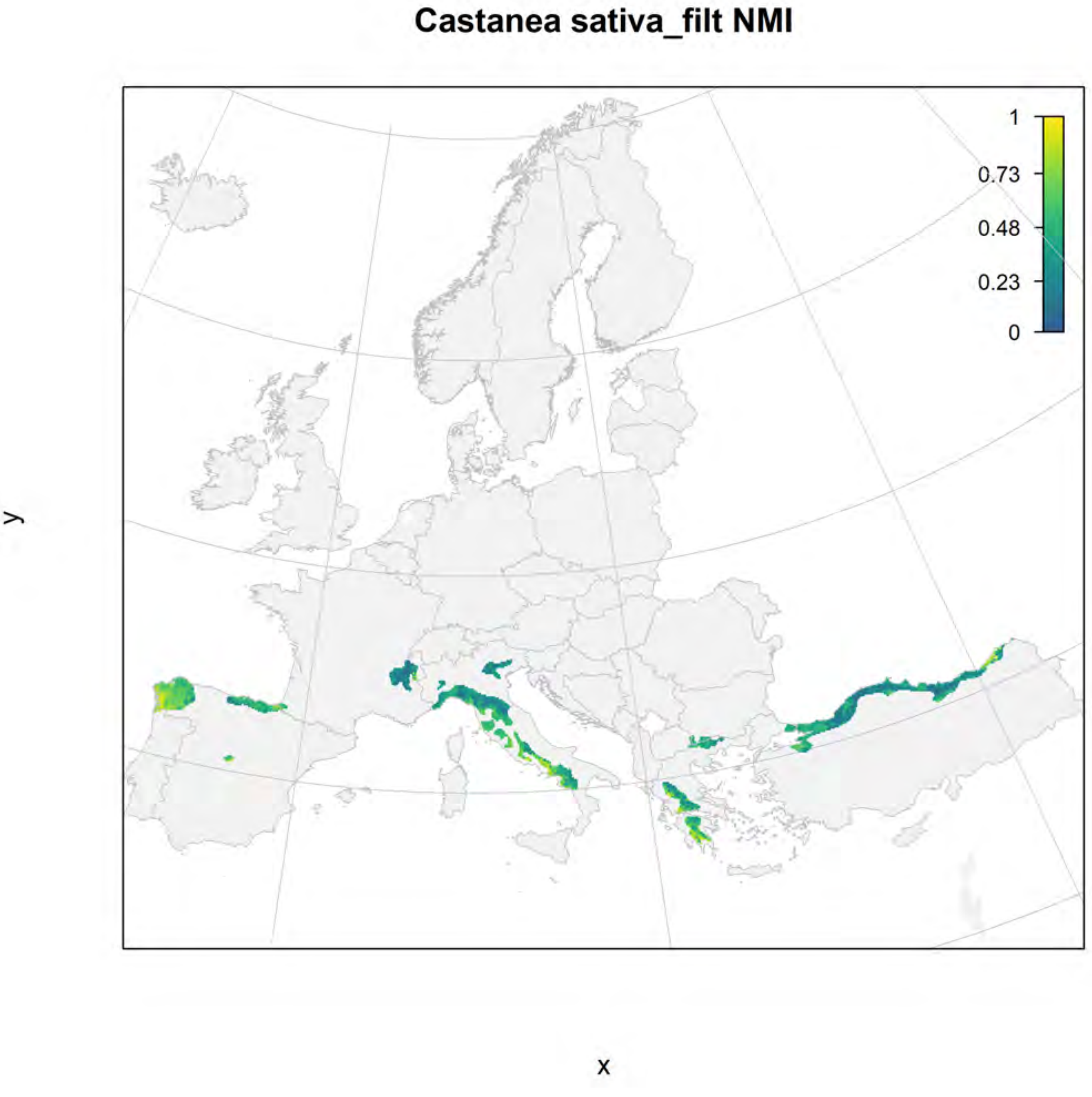
Castanea sativa_filt NMI
1
0.73
0.48
0.23
0
y
x

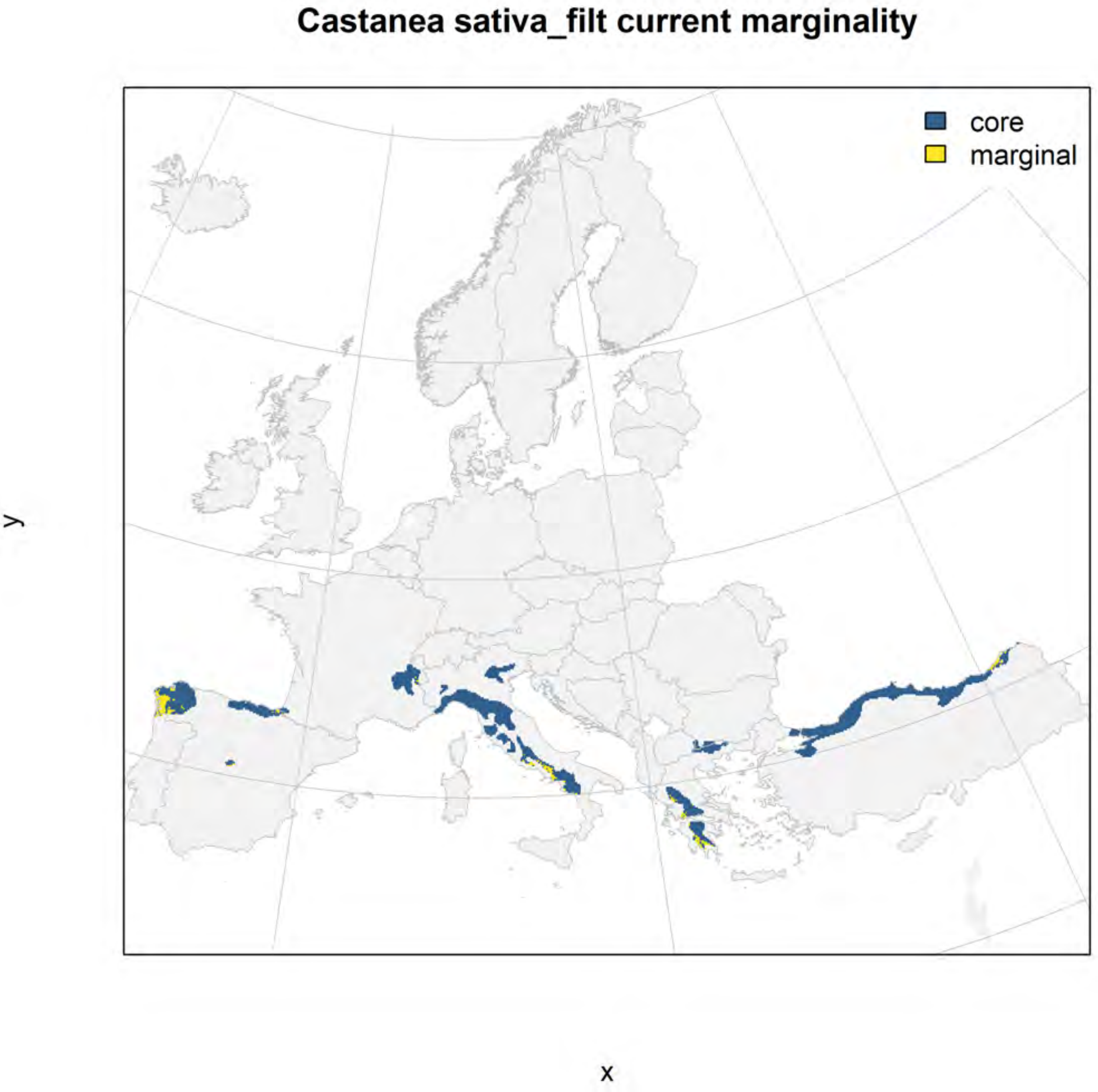
Castanea sativa_filt current marginality
core
marginal
y
x

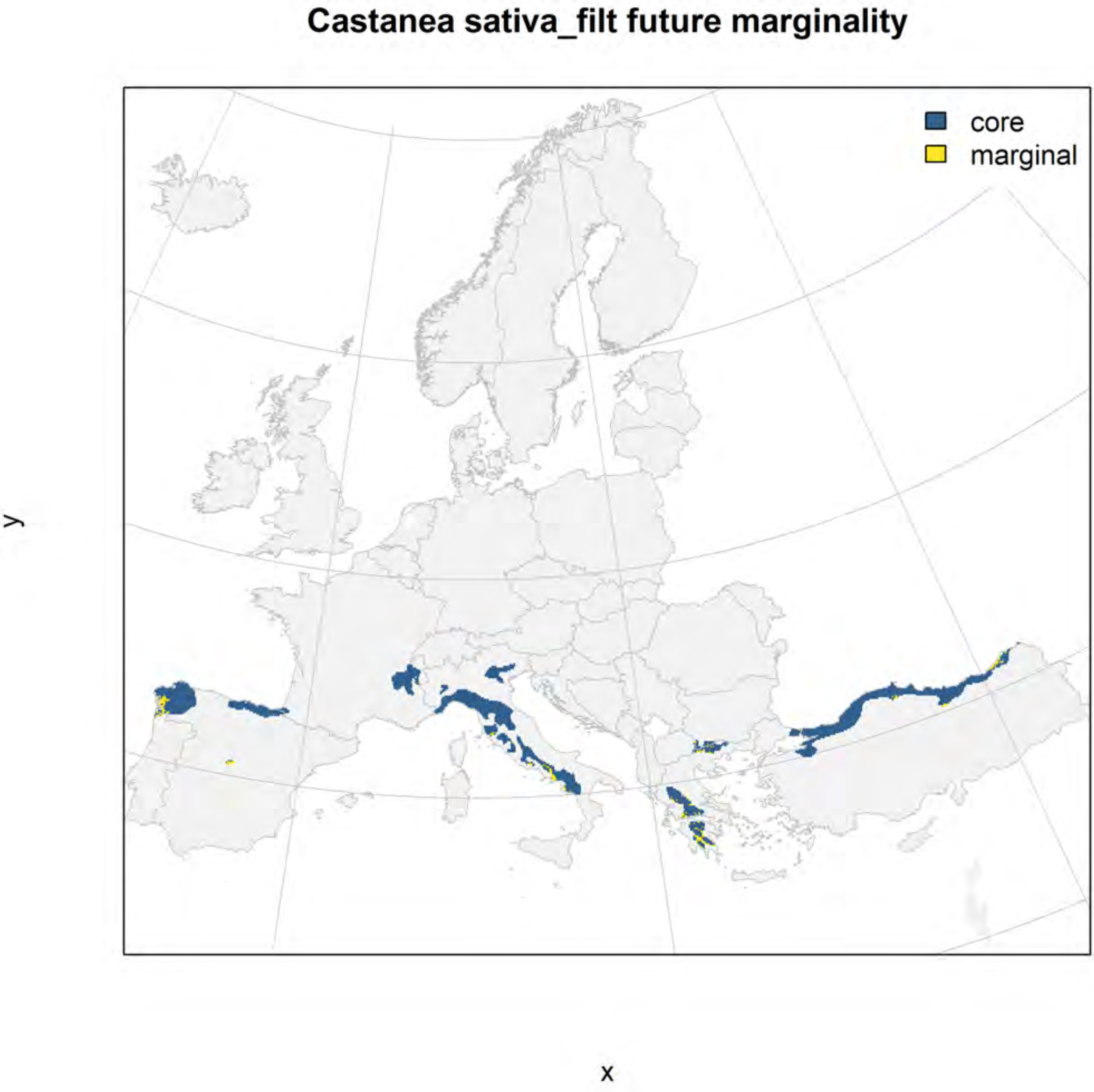
Castanea sativa_filt future marginality
core
marginal
y
x

### Cedrus libani_filt NMI

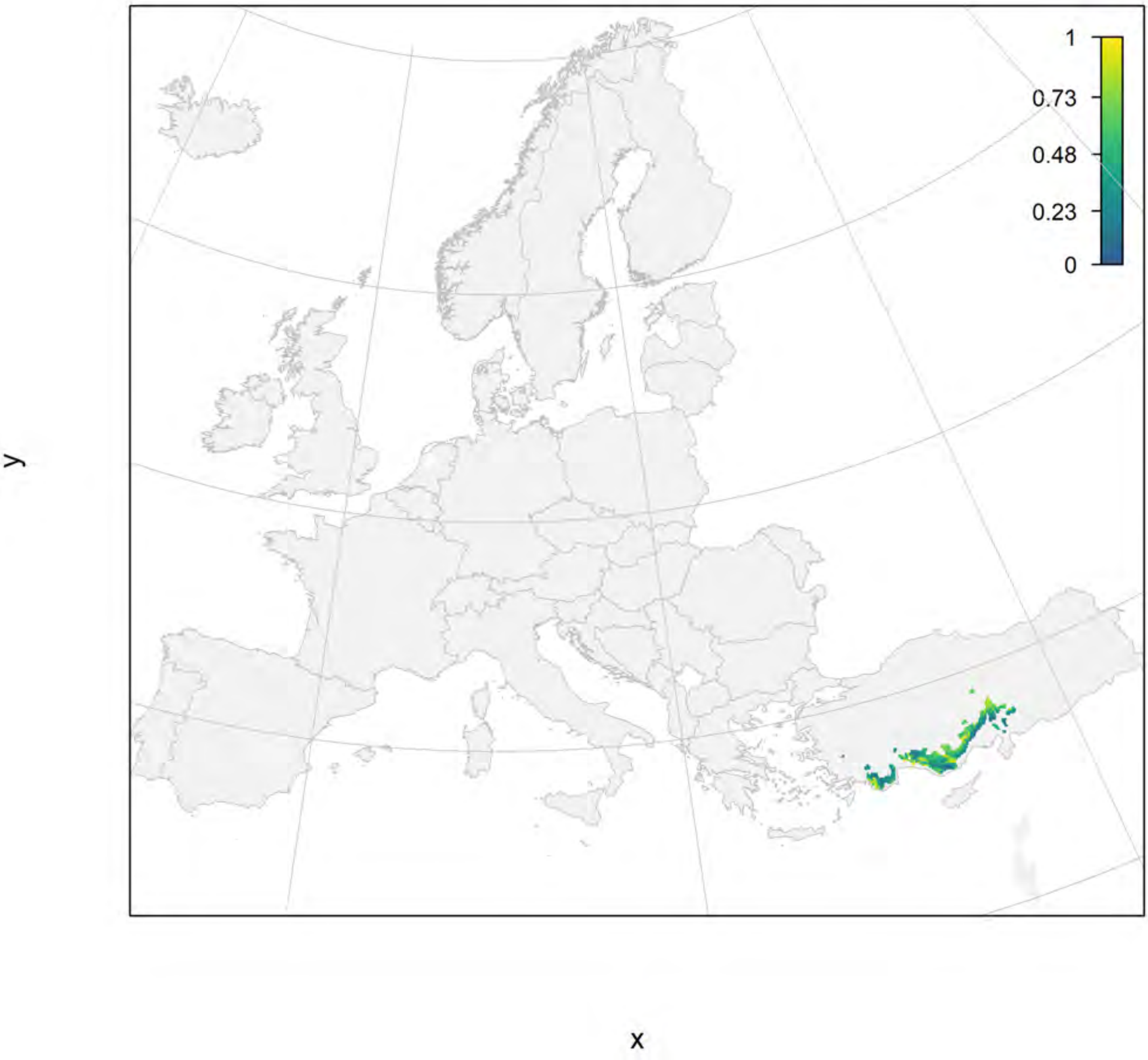

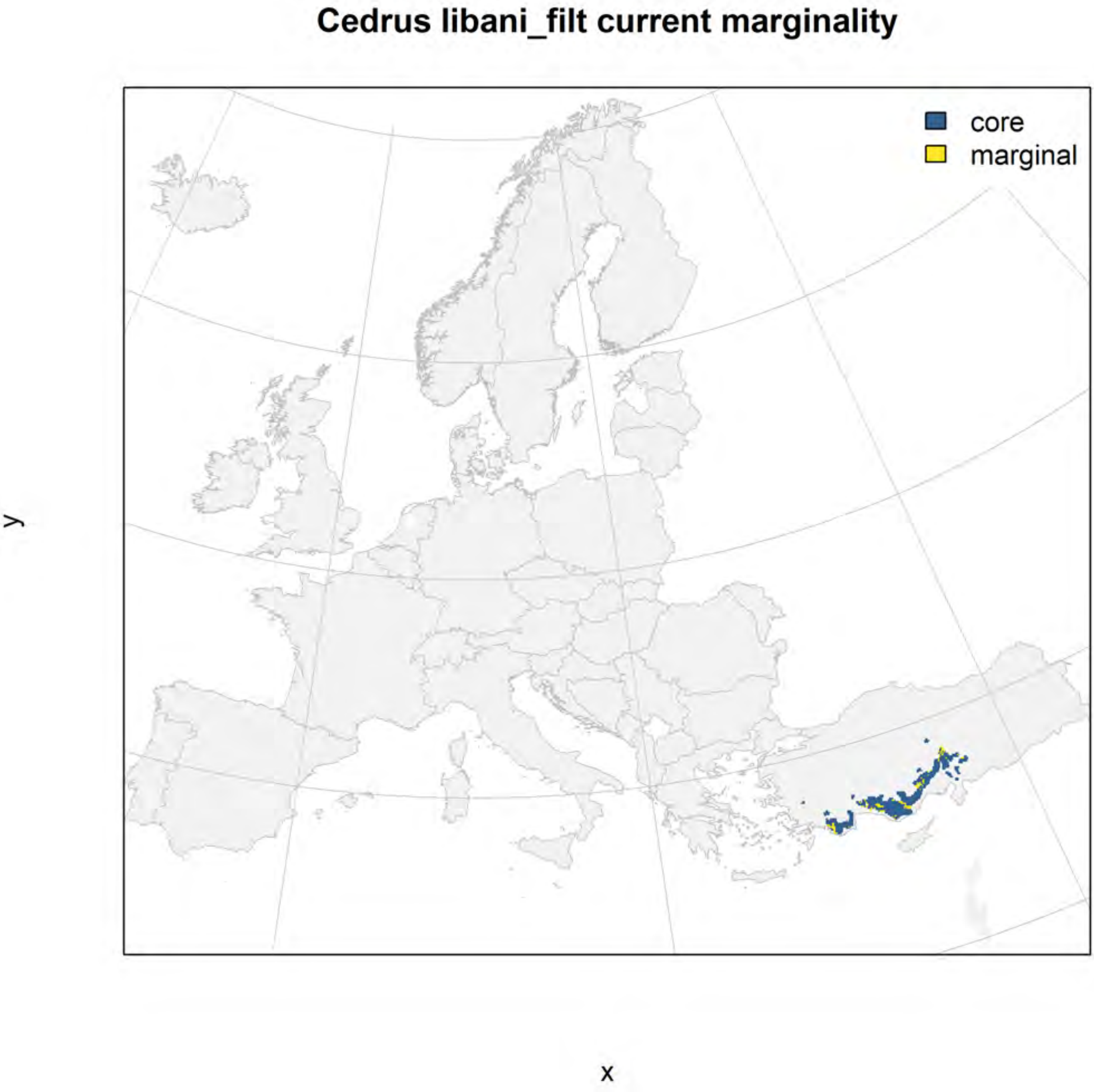
Cedrus libani_filt current marginality
core
marginal
y
x

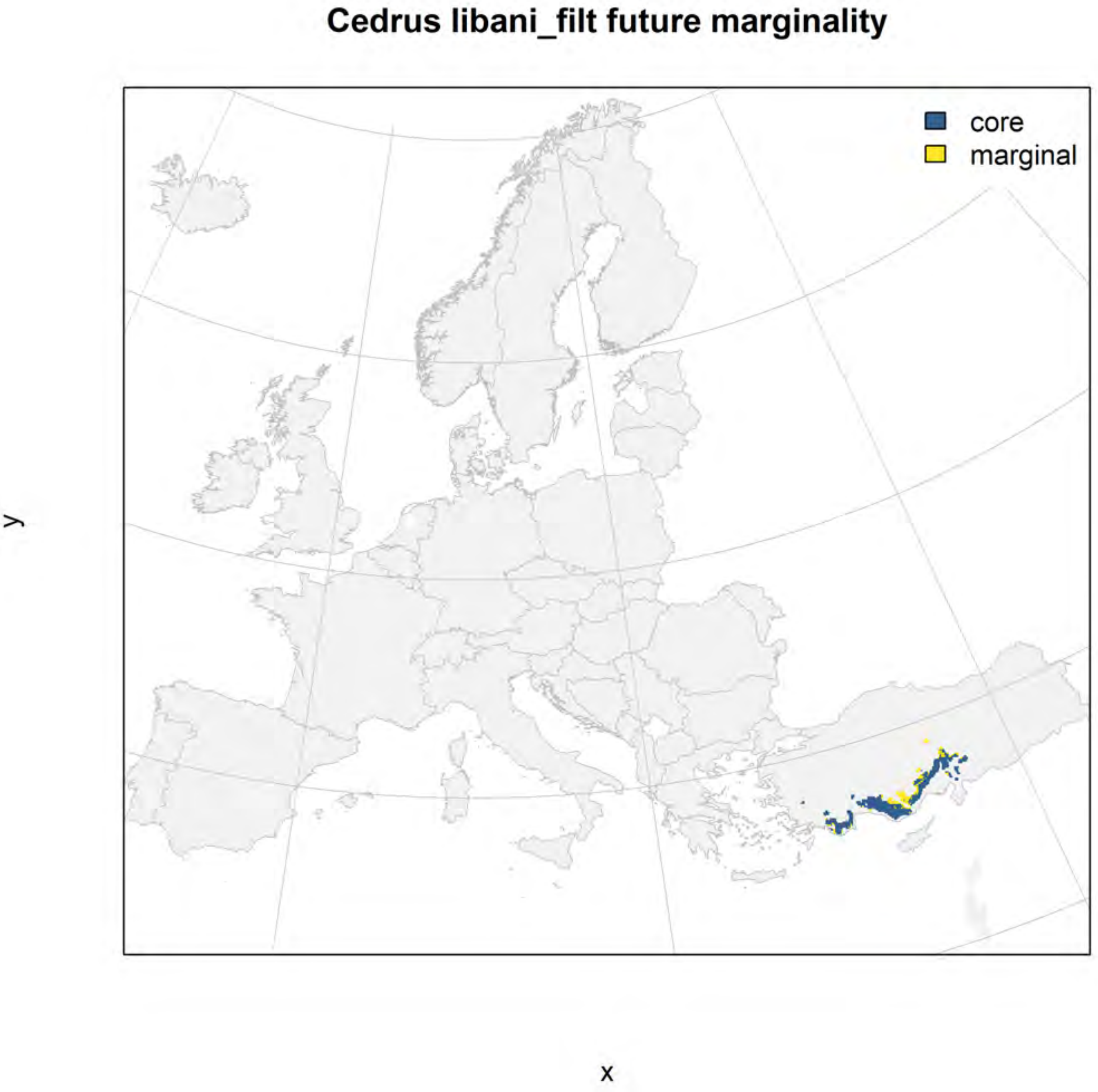
Cedrus libani_filt future marginality
core
marginal
y
x

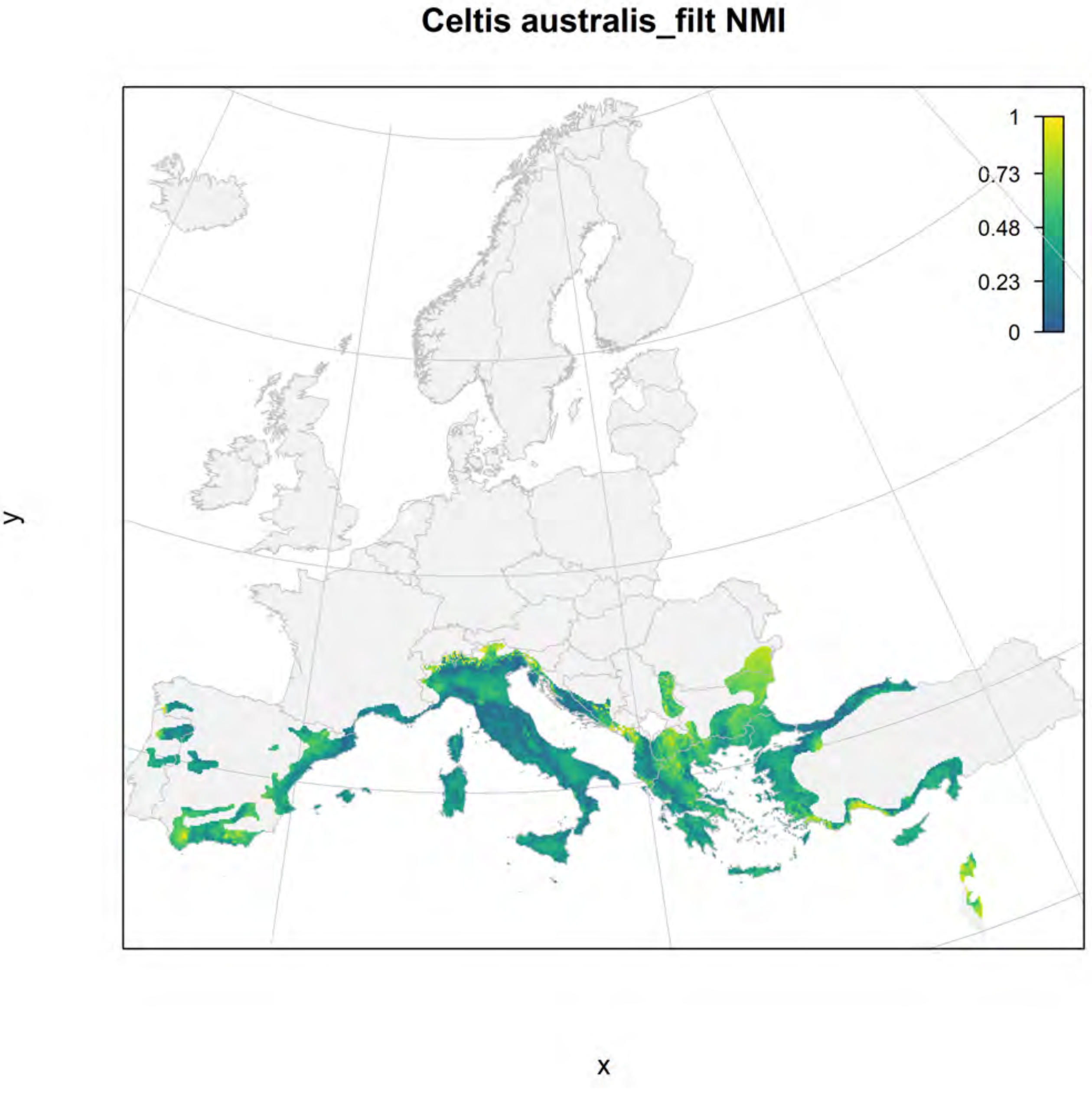
Celtis australis_filt NMI
1
0.73
0.48
0.23
0
y
x

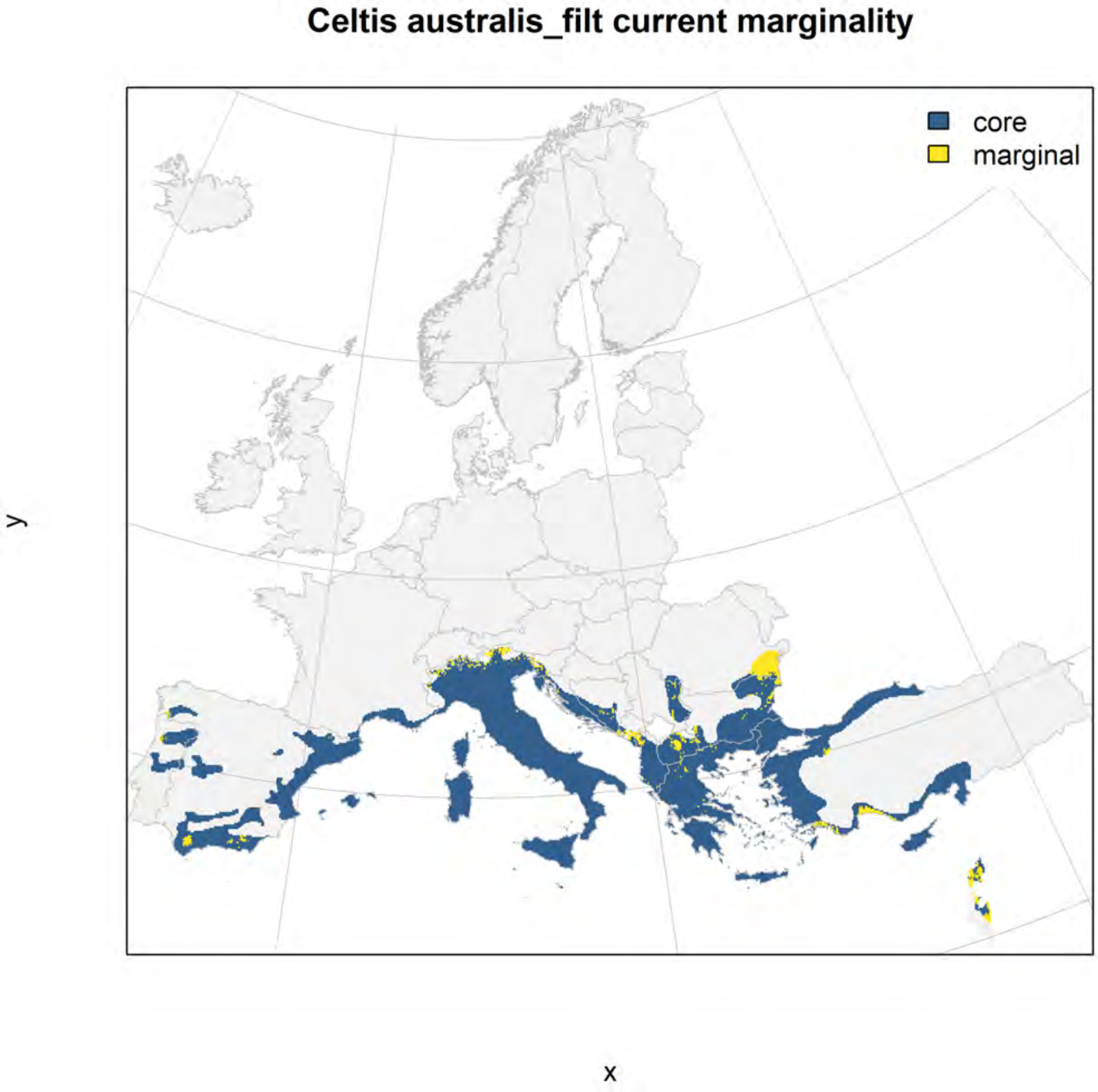
Celtis australis_filt current marginality
core
marginal
y
x

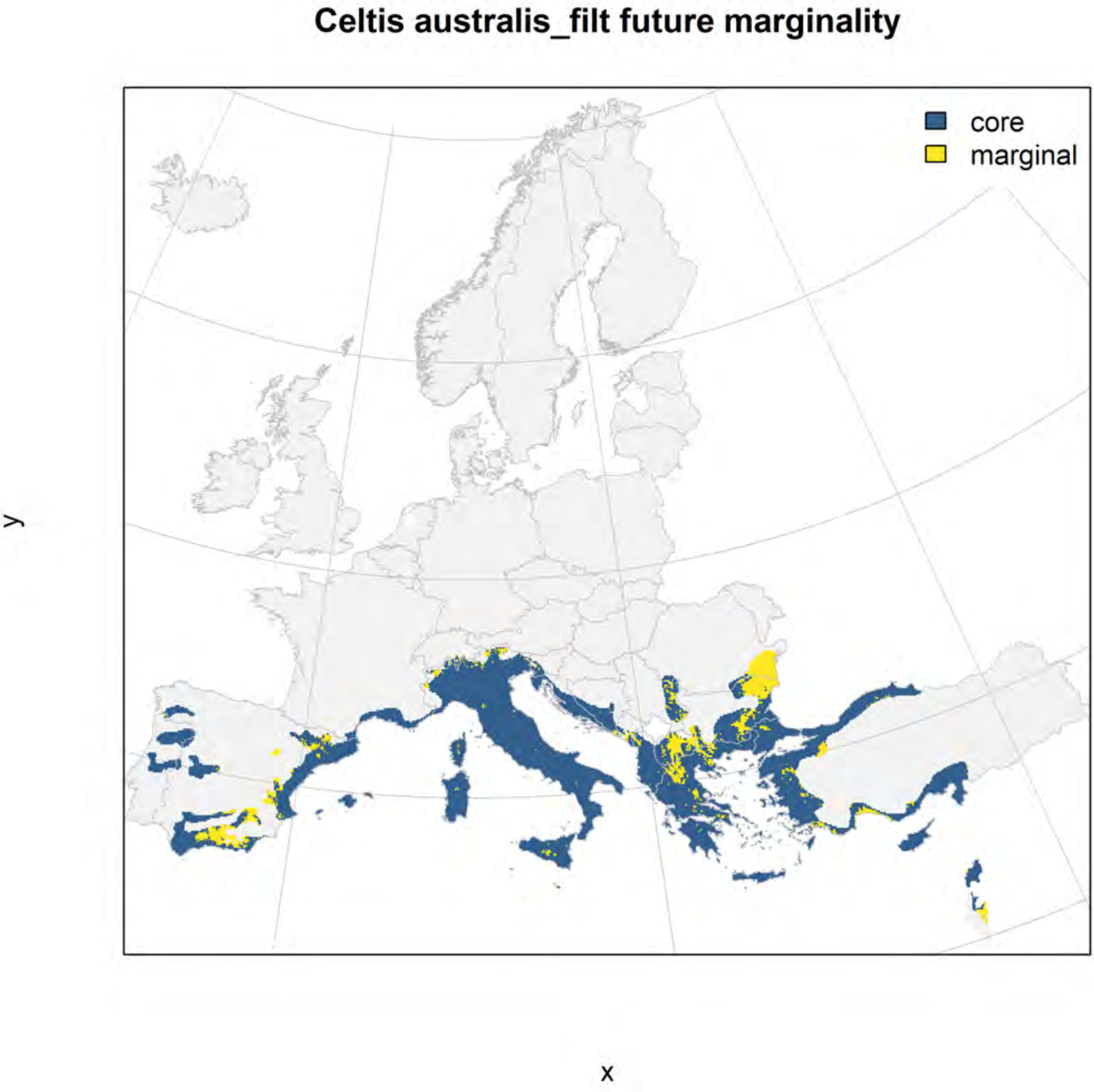
Celtis australis_filt future marginality
core
marginal
y
x

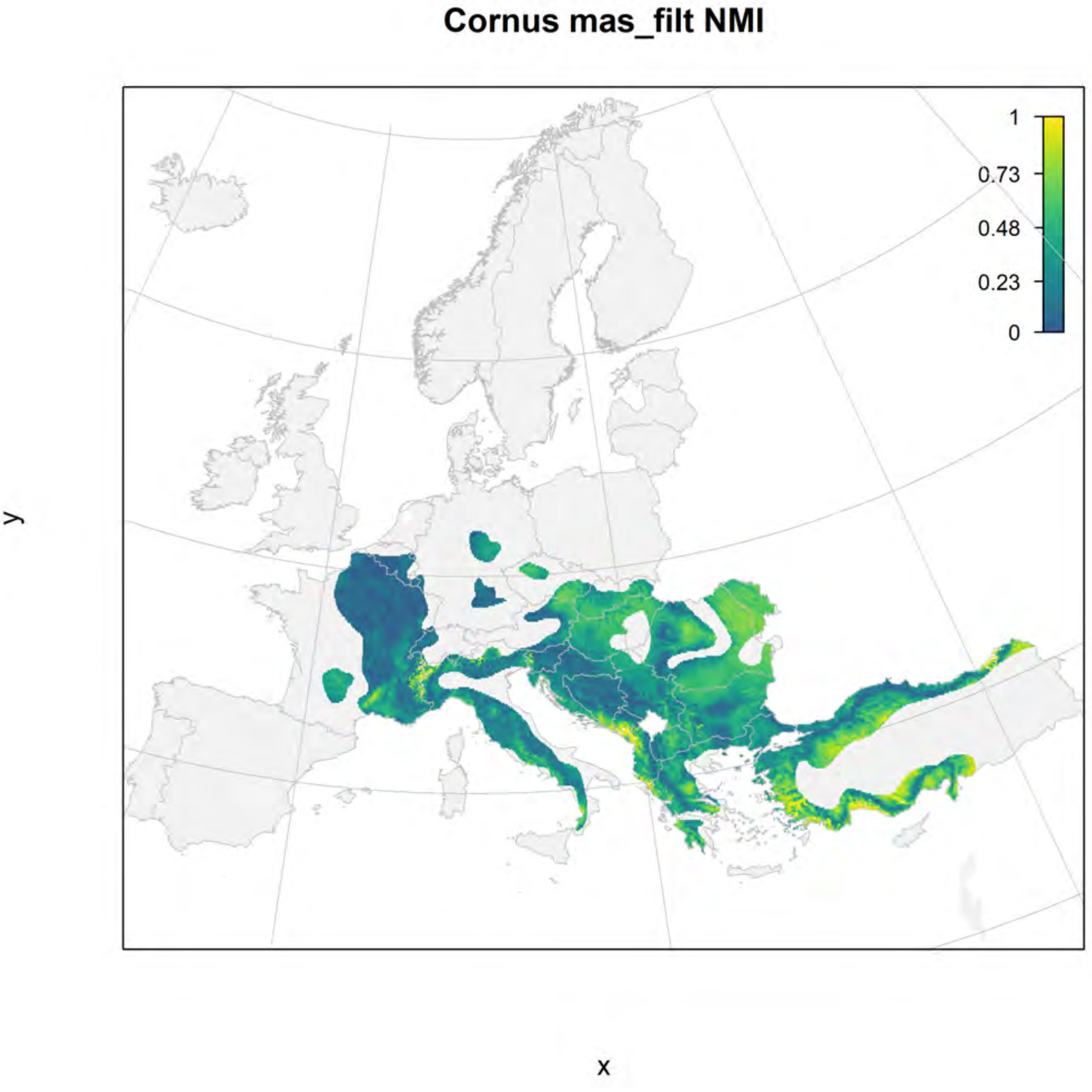
Cornus mas_filt NMI
1
0.73
0.48
0.23
0
y
x

### Cornus mas_filt current marginality

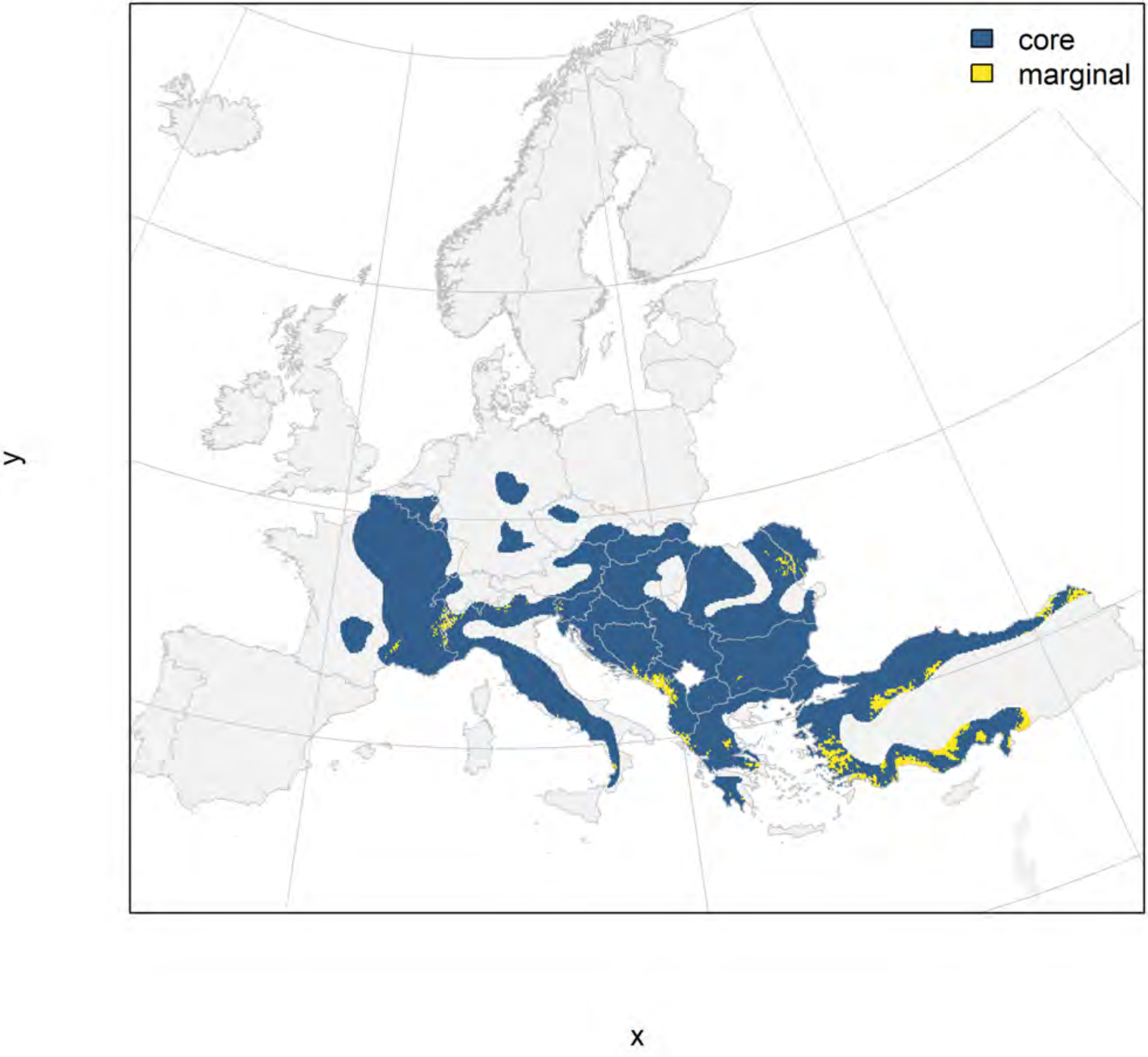

#### Cornus mas_filt future marginality

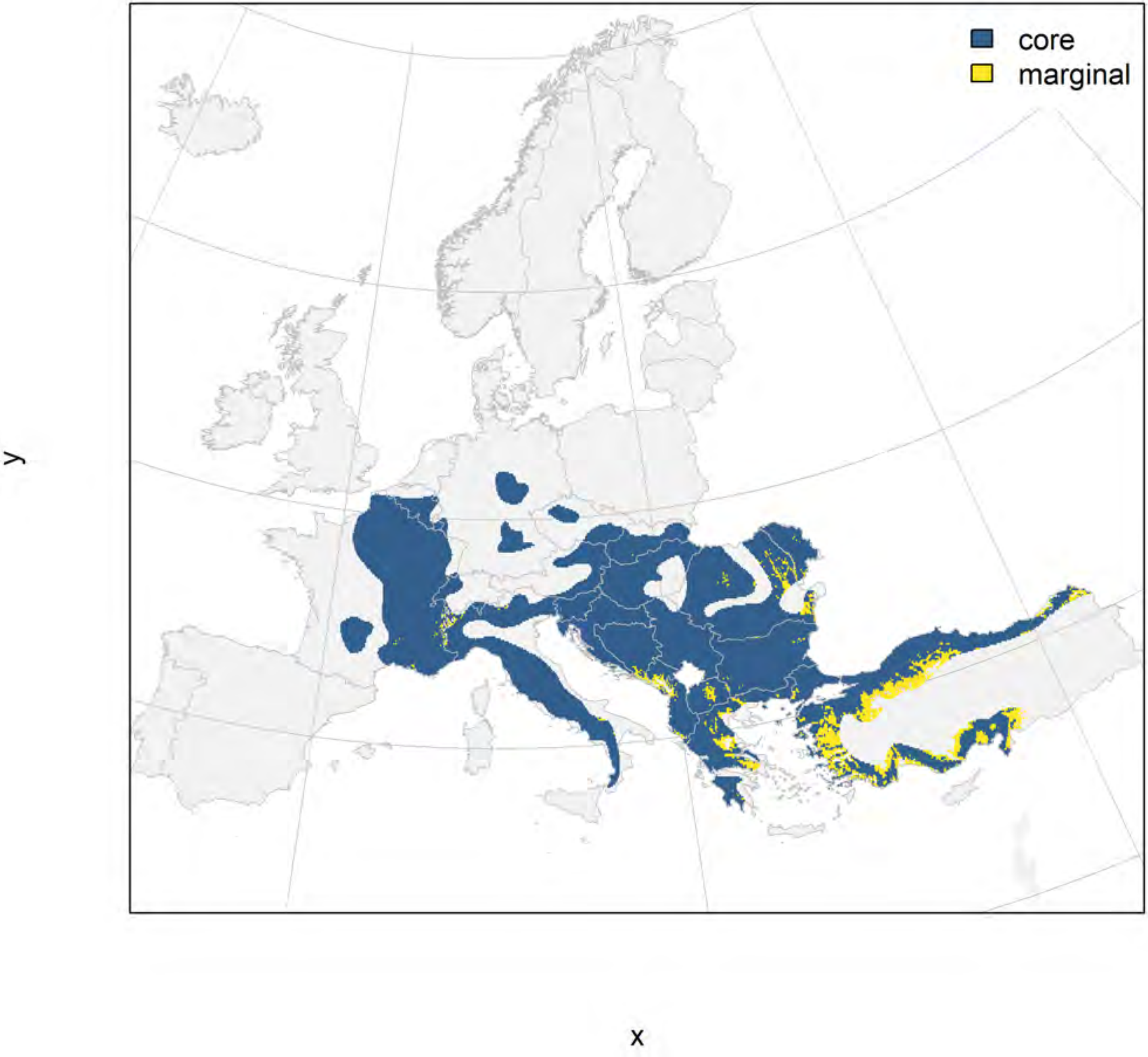

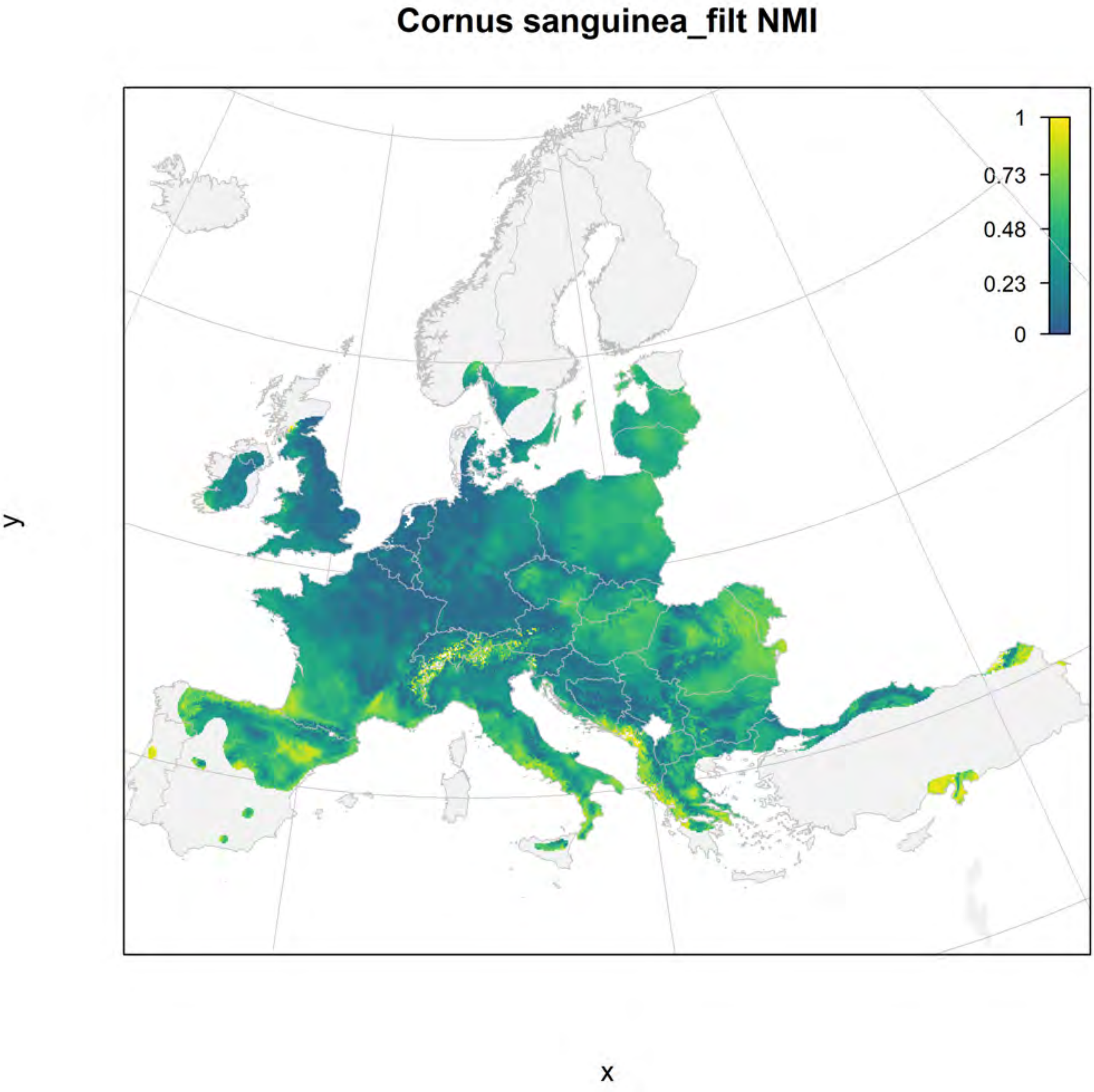

Cornus sanguinea_filt NMI
1
0.73
0.48
0.23
0
y
x

### Cornus sanguinea_filt current marginality

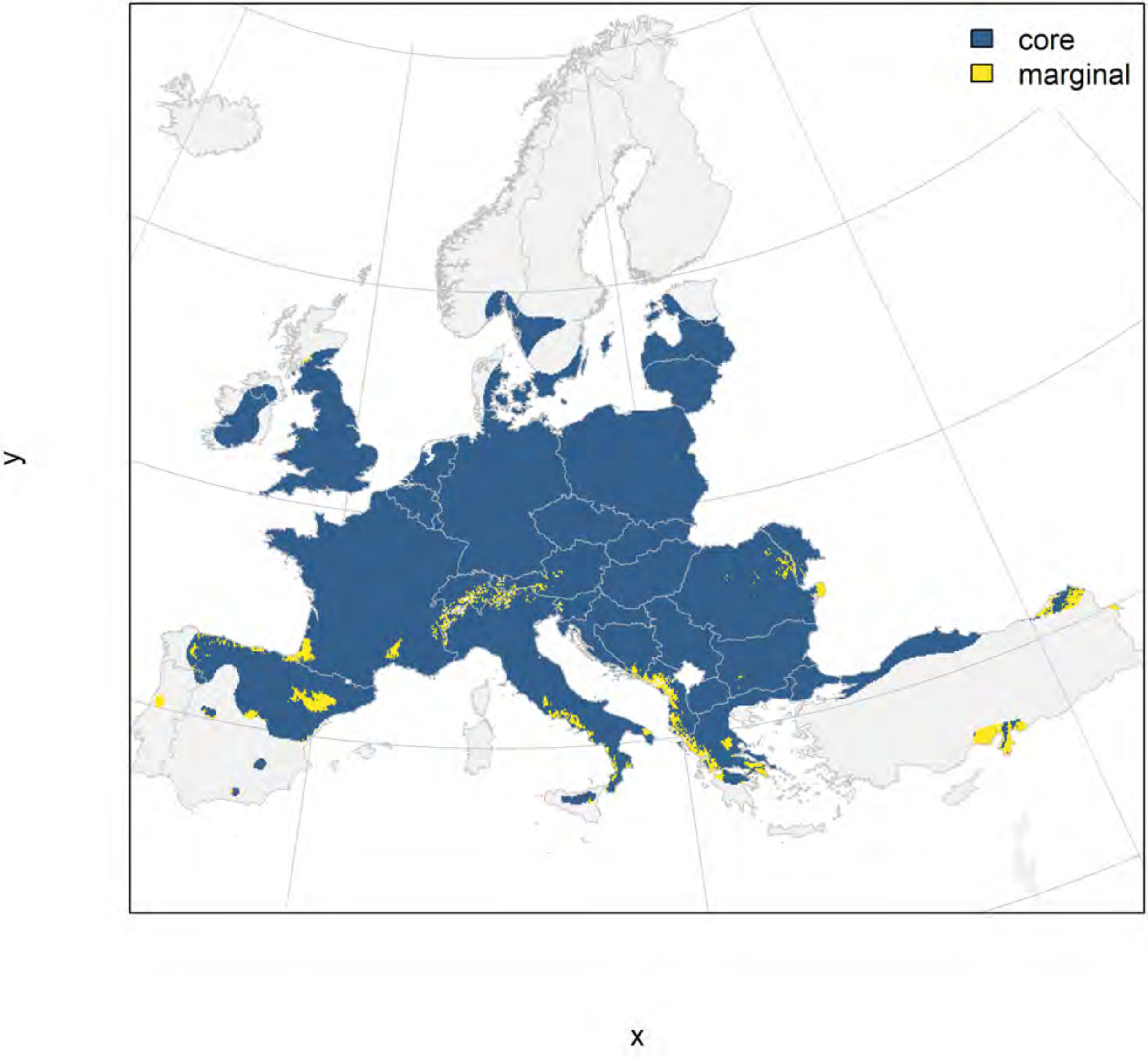

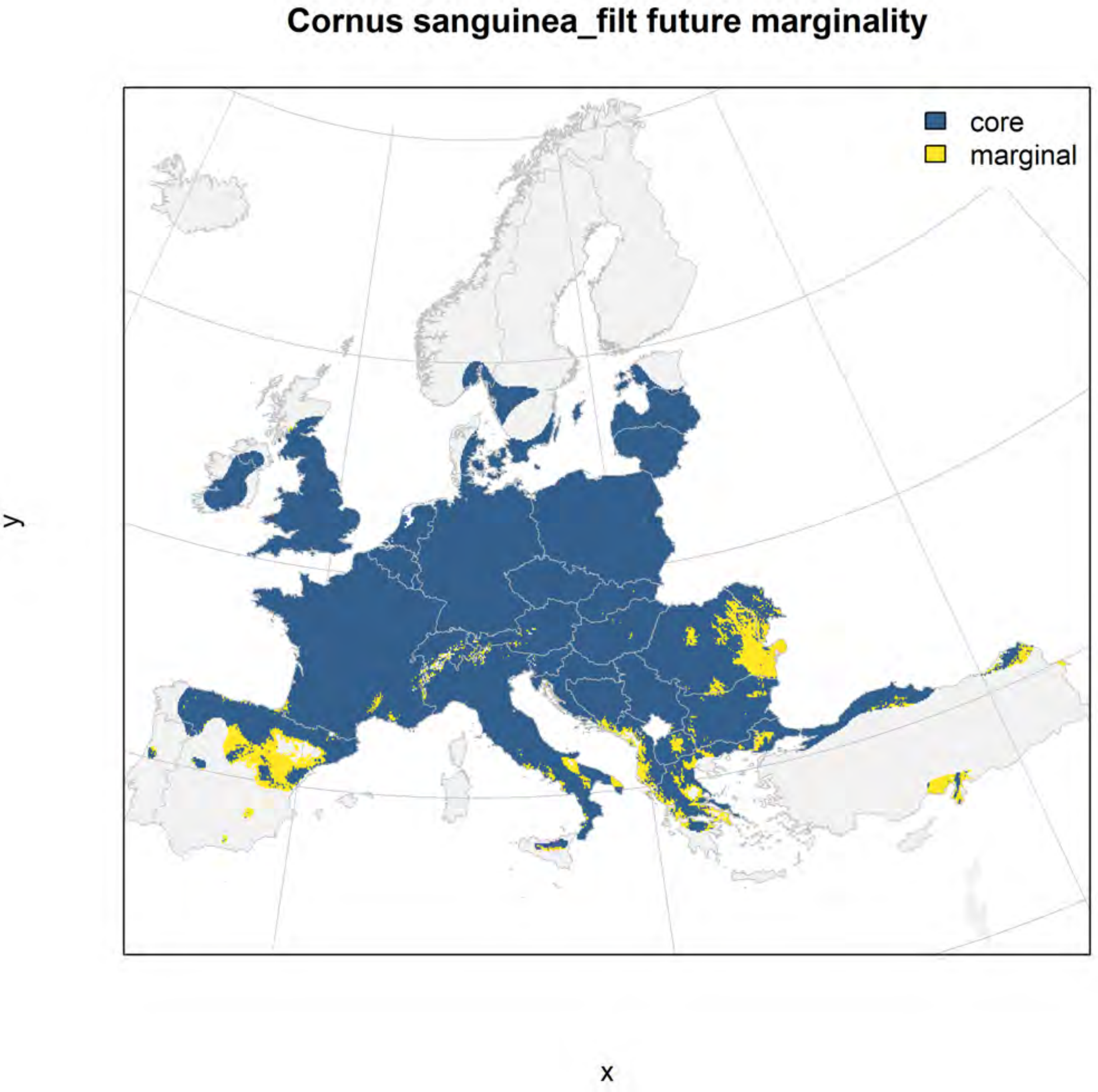
Cornus sanguinea_filt future marginality
core
marginal
y
x

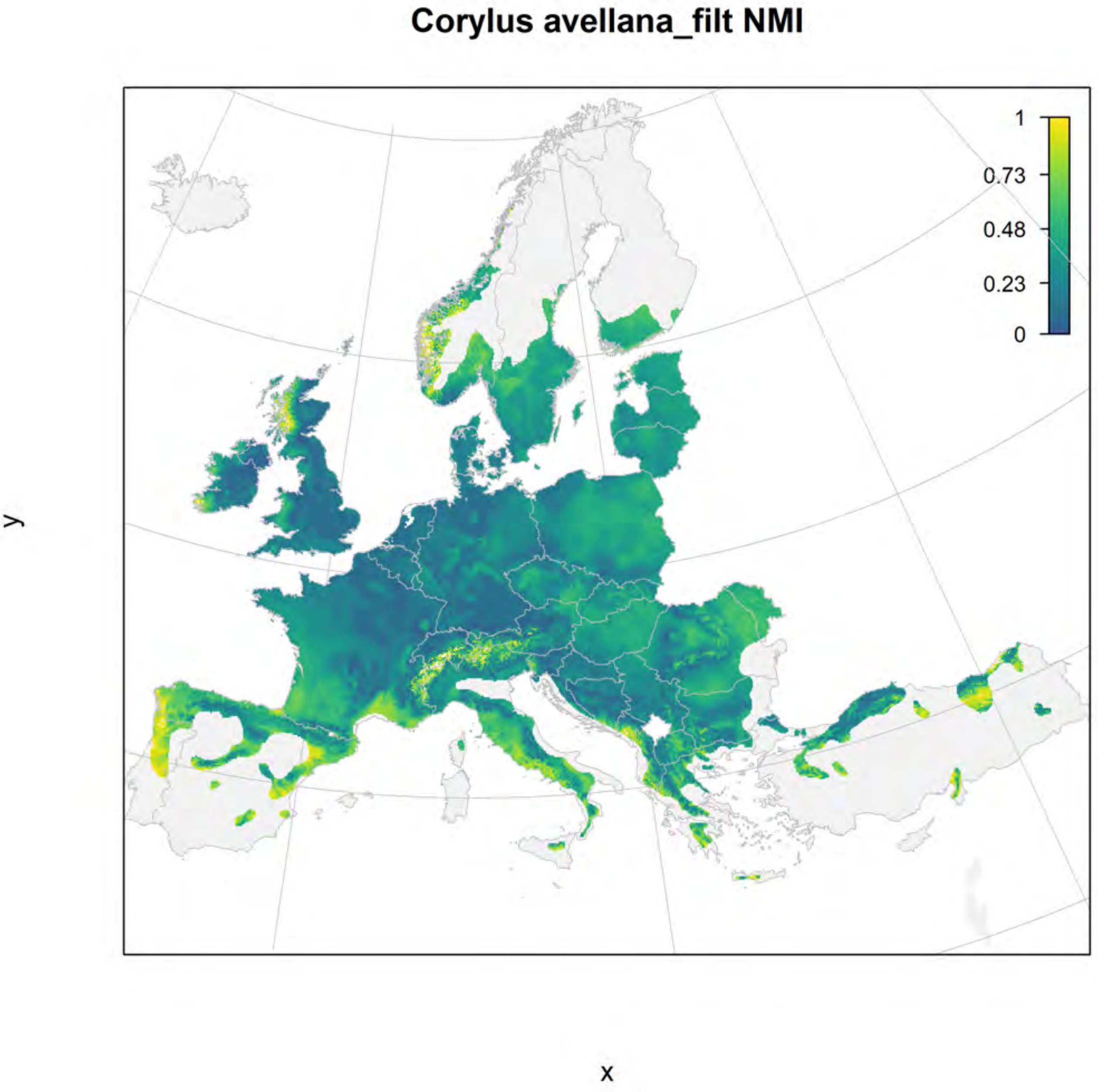

Corylus avellana_filt NMI
1
0.73
0.48
0.23
0
y
x

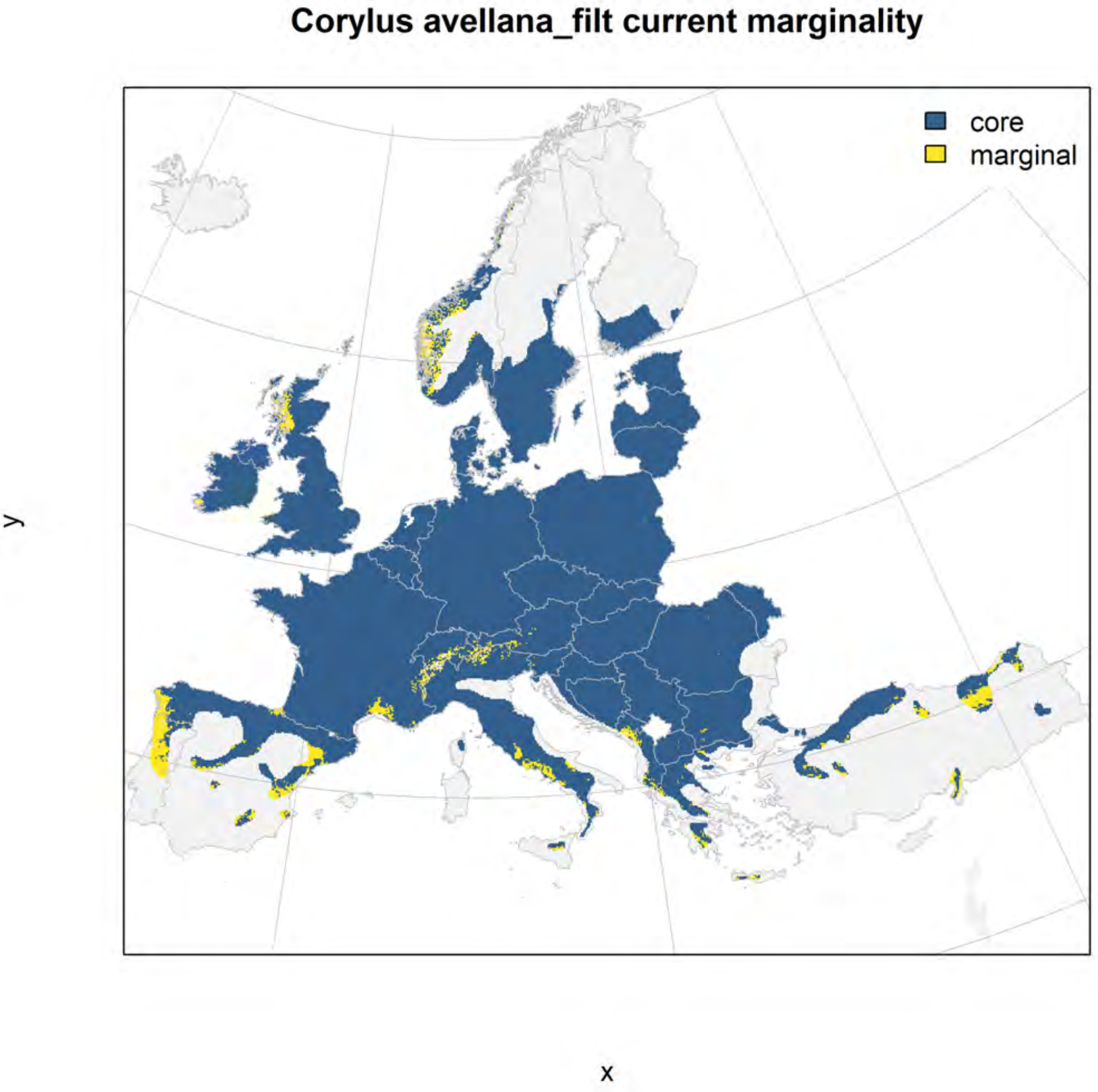
Corylus avellana_filt current marginality
core
marginal
y
x

#### Corylus avellana_filt future marginality

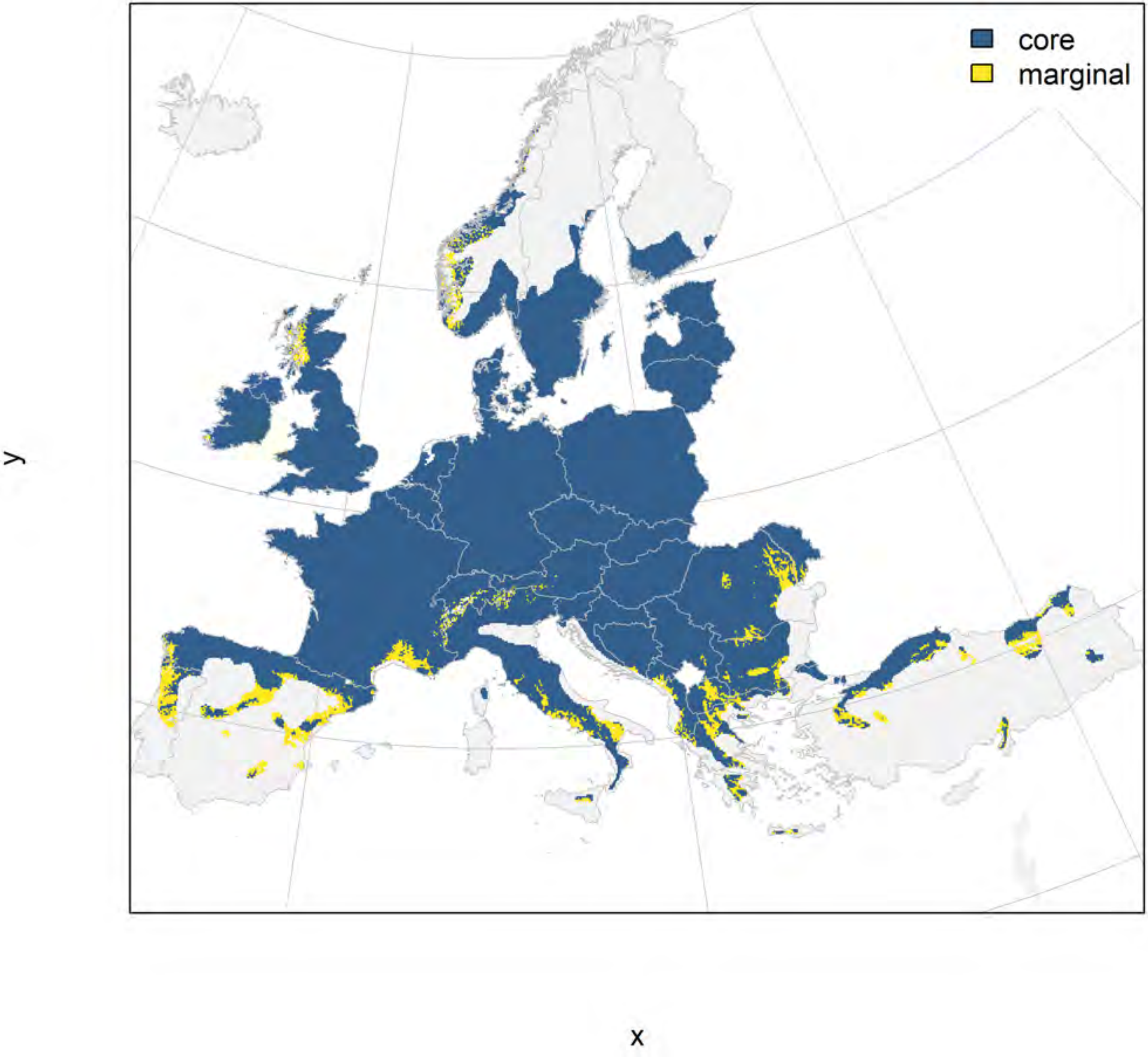

### Cupressus sempervirens_filt NMI

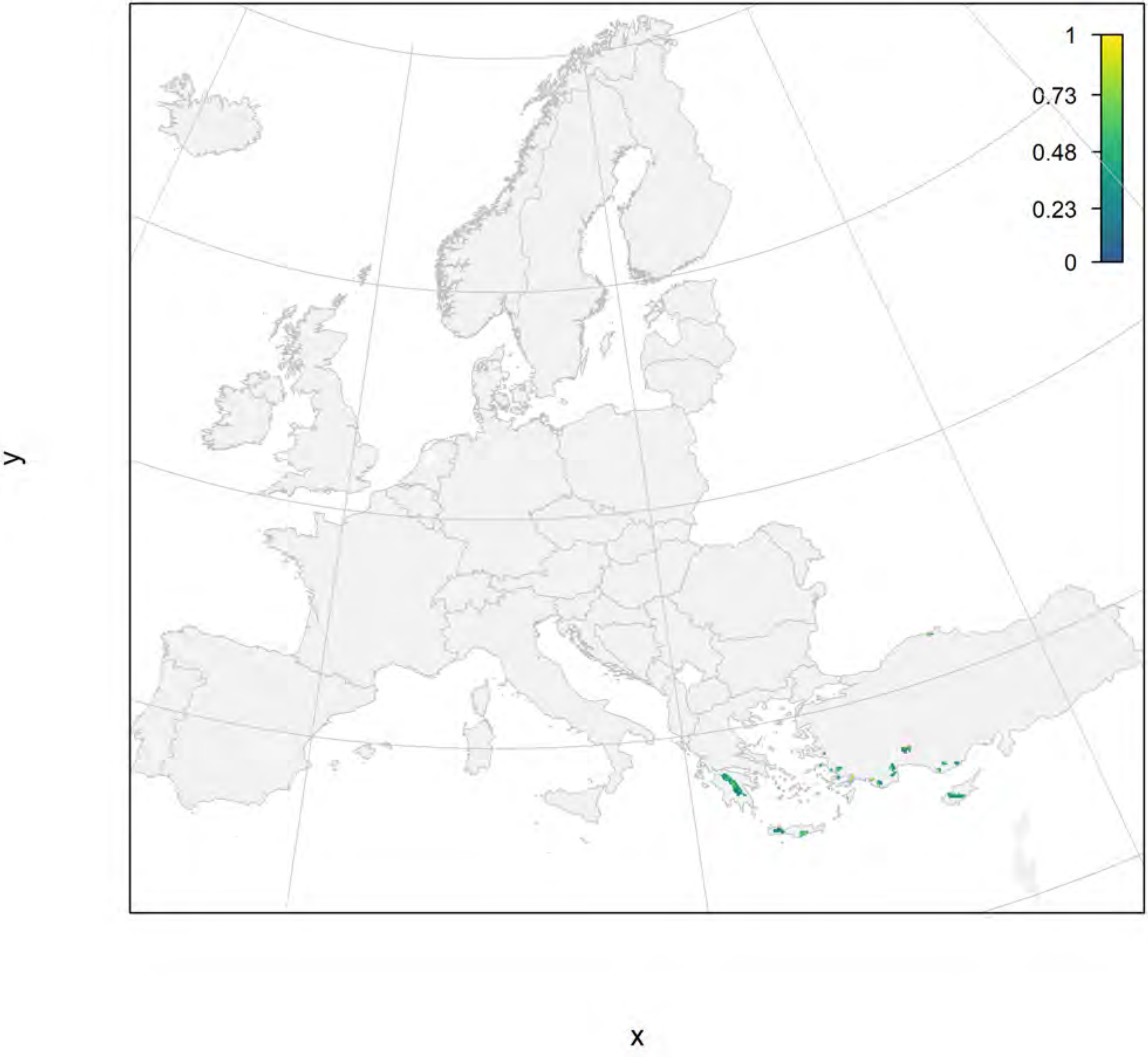

Cupressus sempervirens_filt current marginality

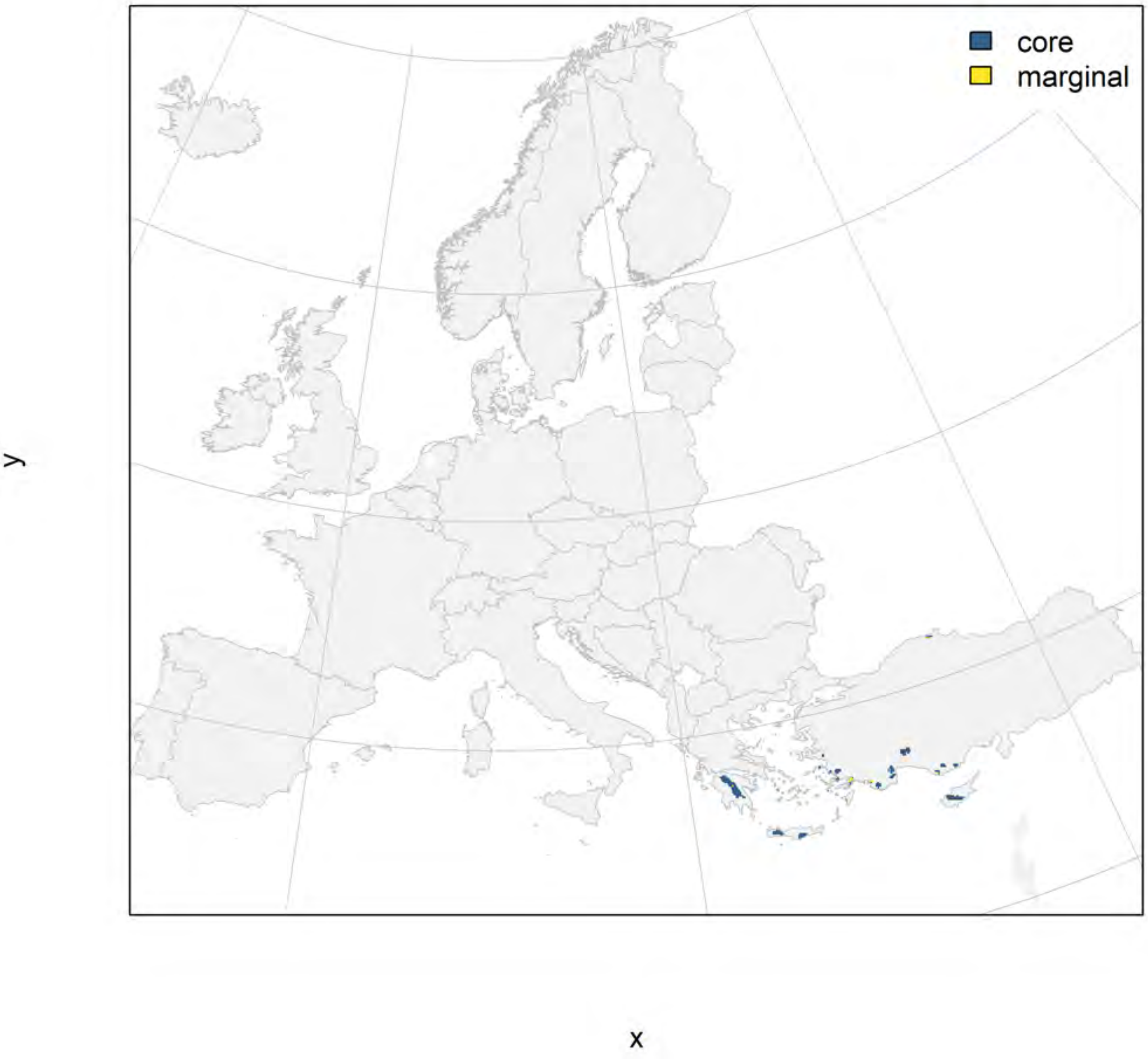

Cupressus sempervirens_filt future marginality

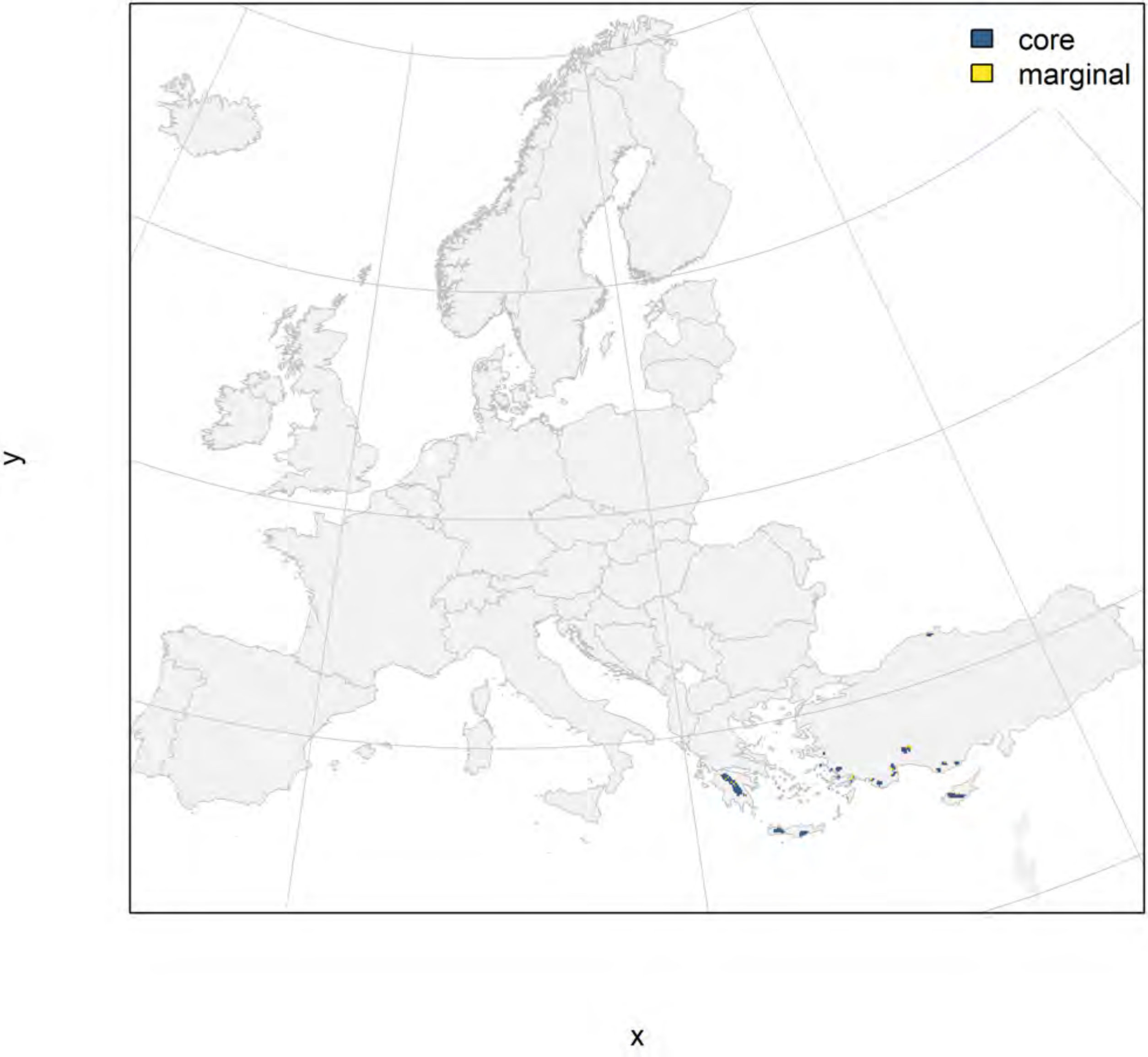

### Fagus sylvatica_filt NMI

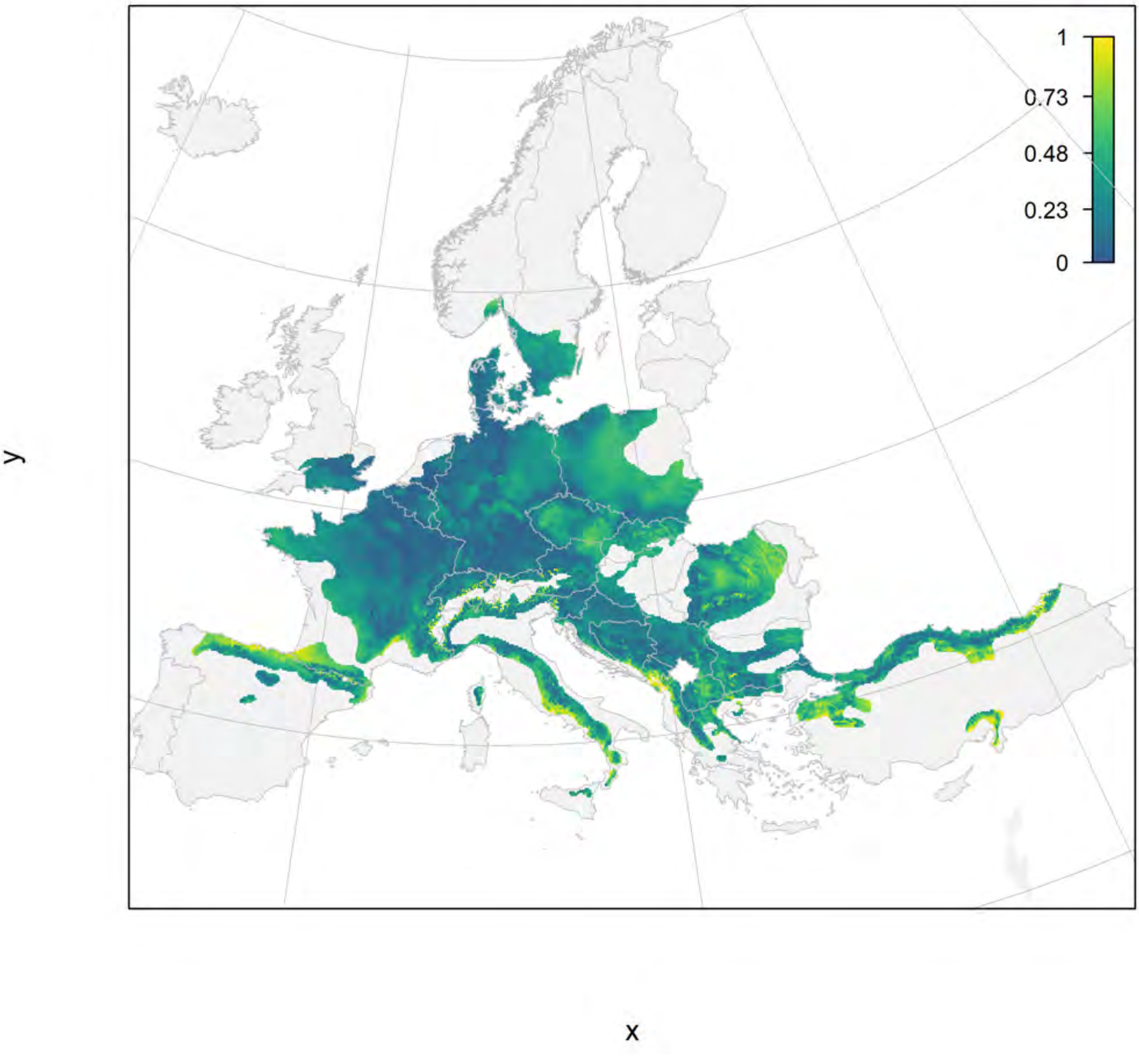

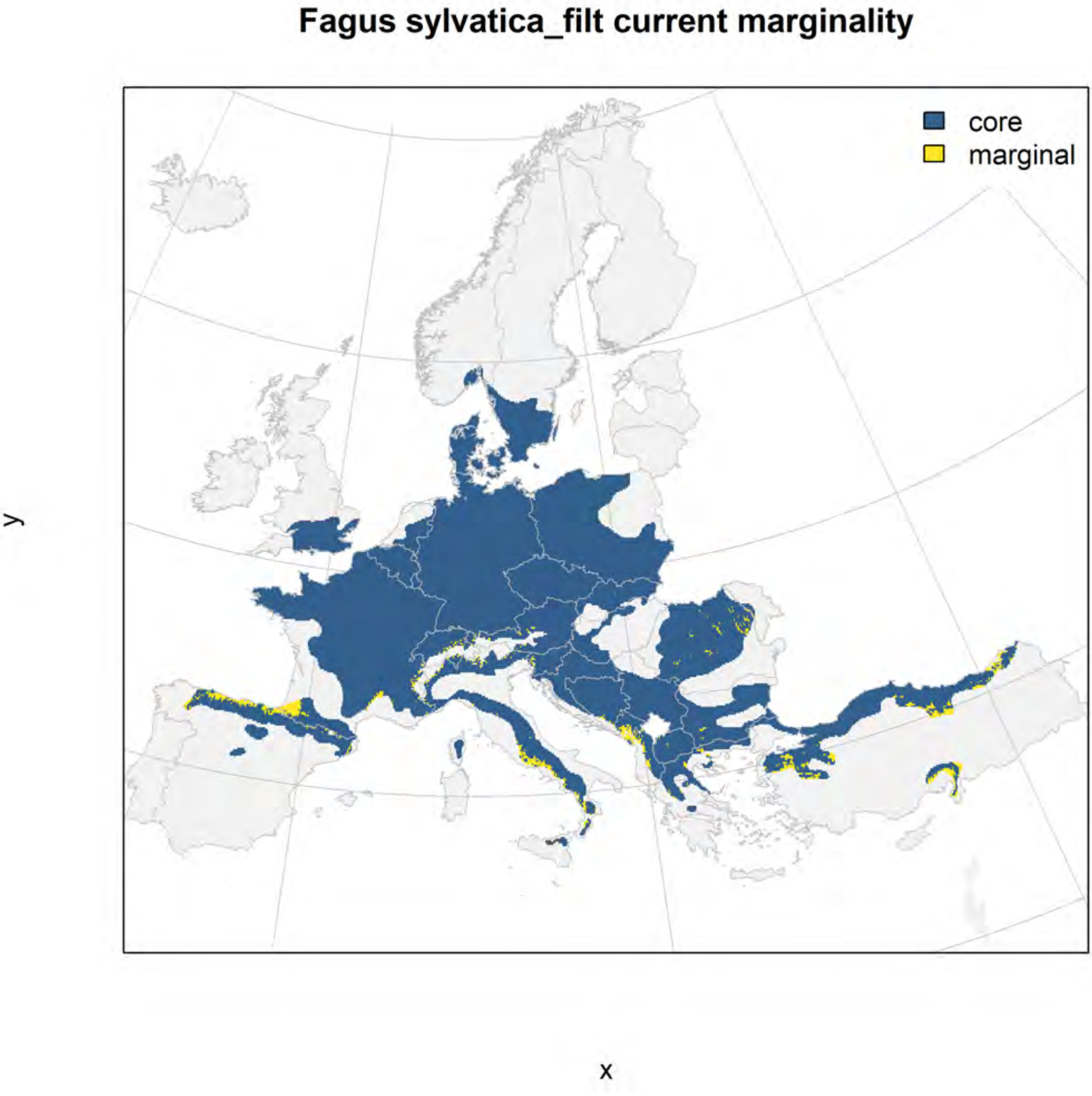
Fagus sylvatica_filt current marginality
core
marginal
y
x

#### Fagus sylvatica_filt future marginality

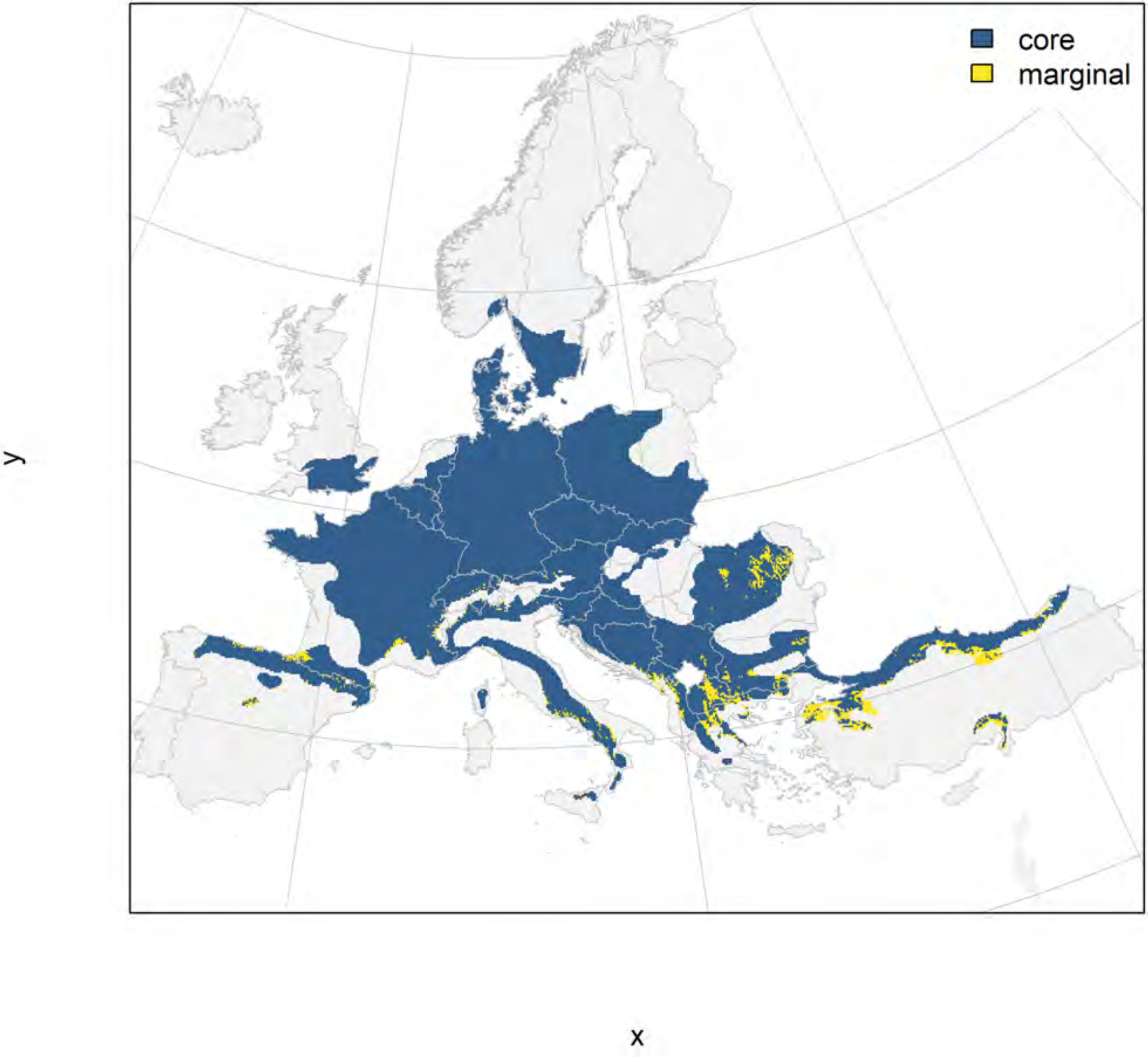

### Frangula alnus_filt NMI

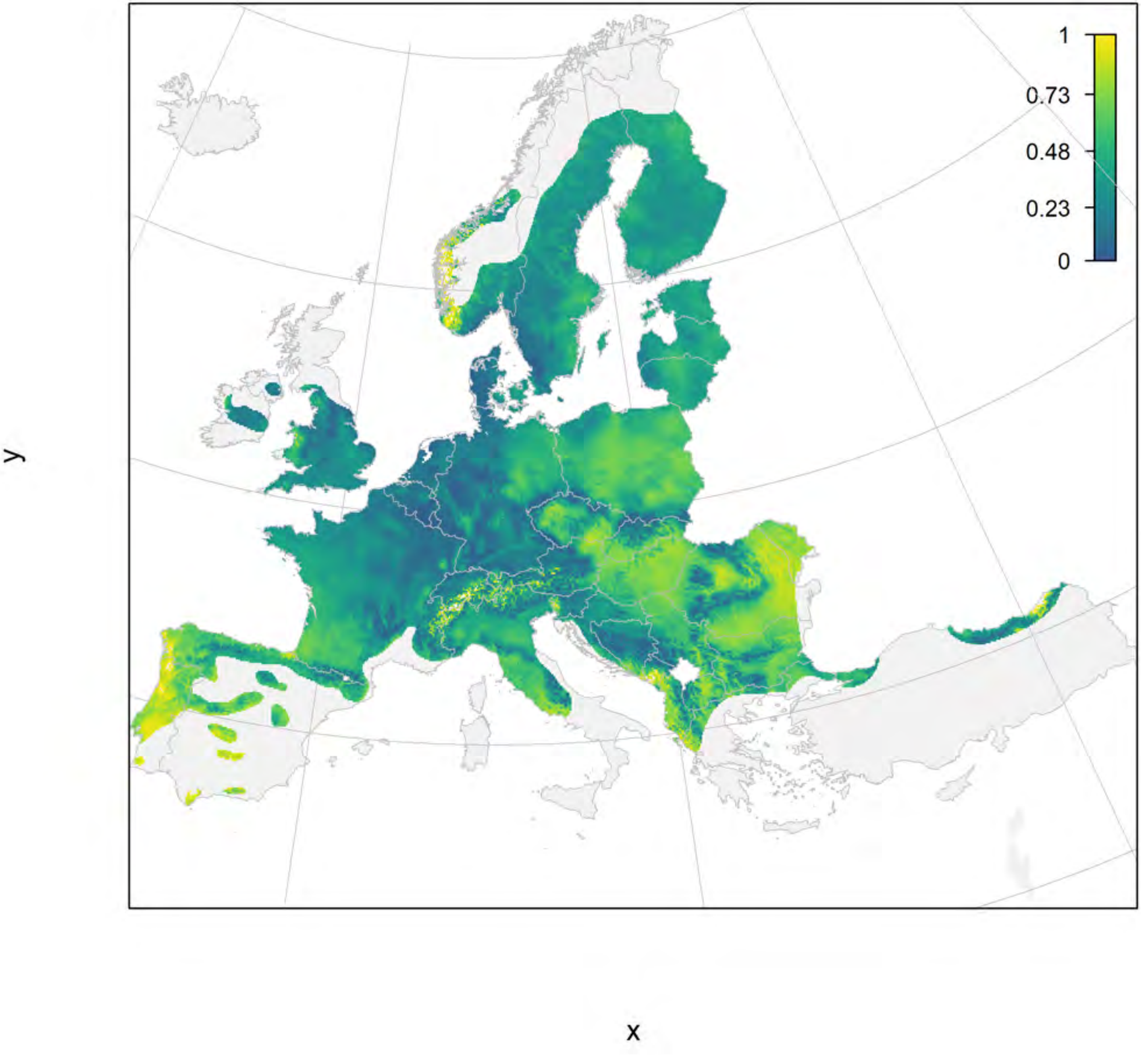

#### Frangula alnus_filt current marginality

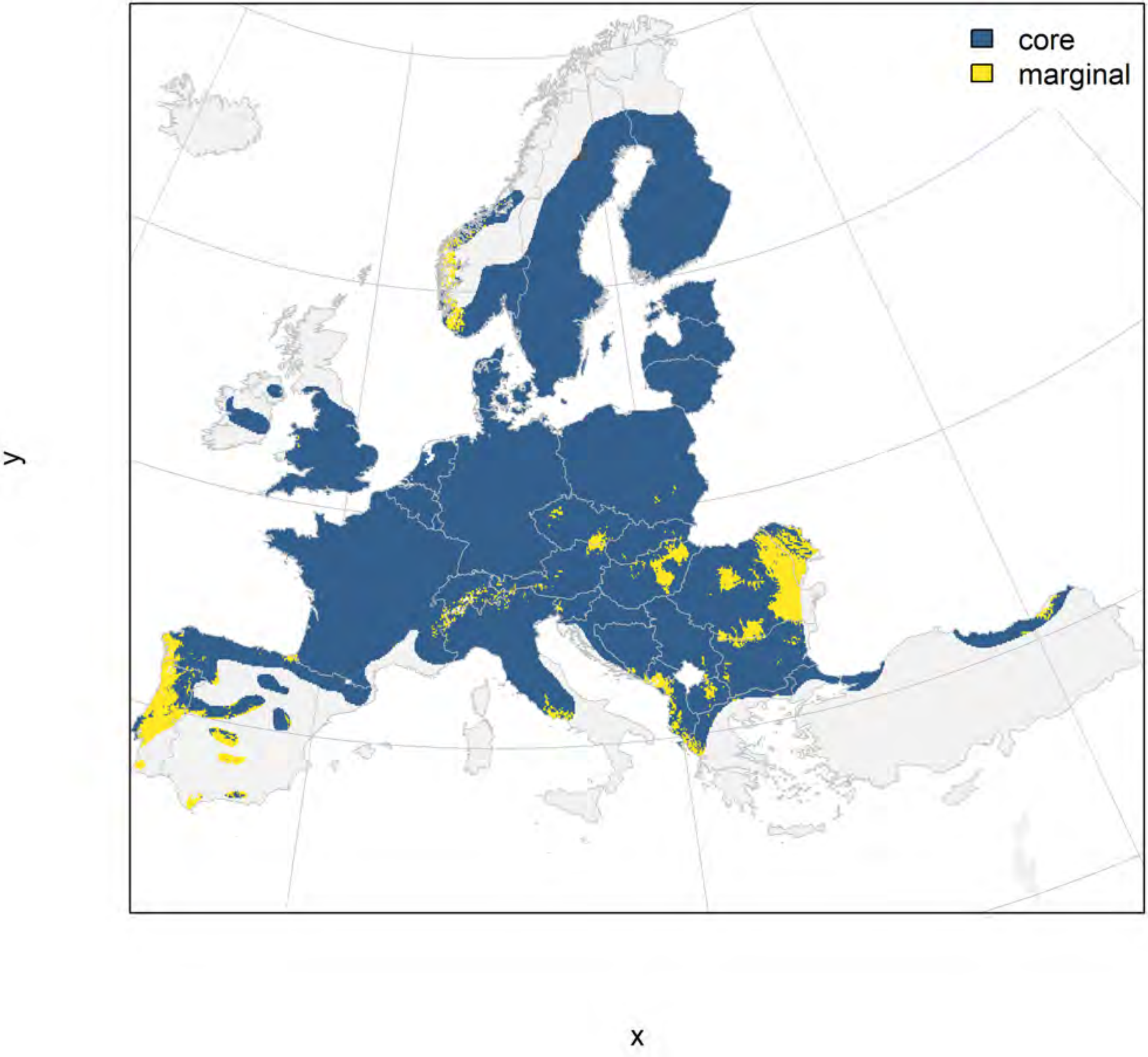

#### Frangula alnus_filt future marginality

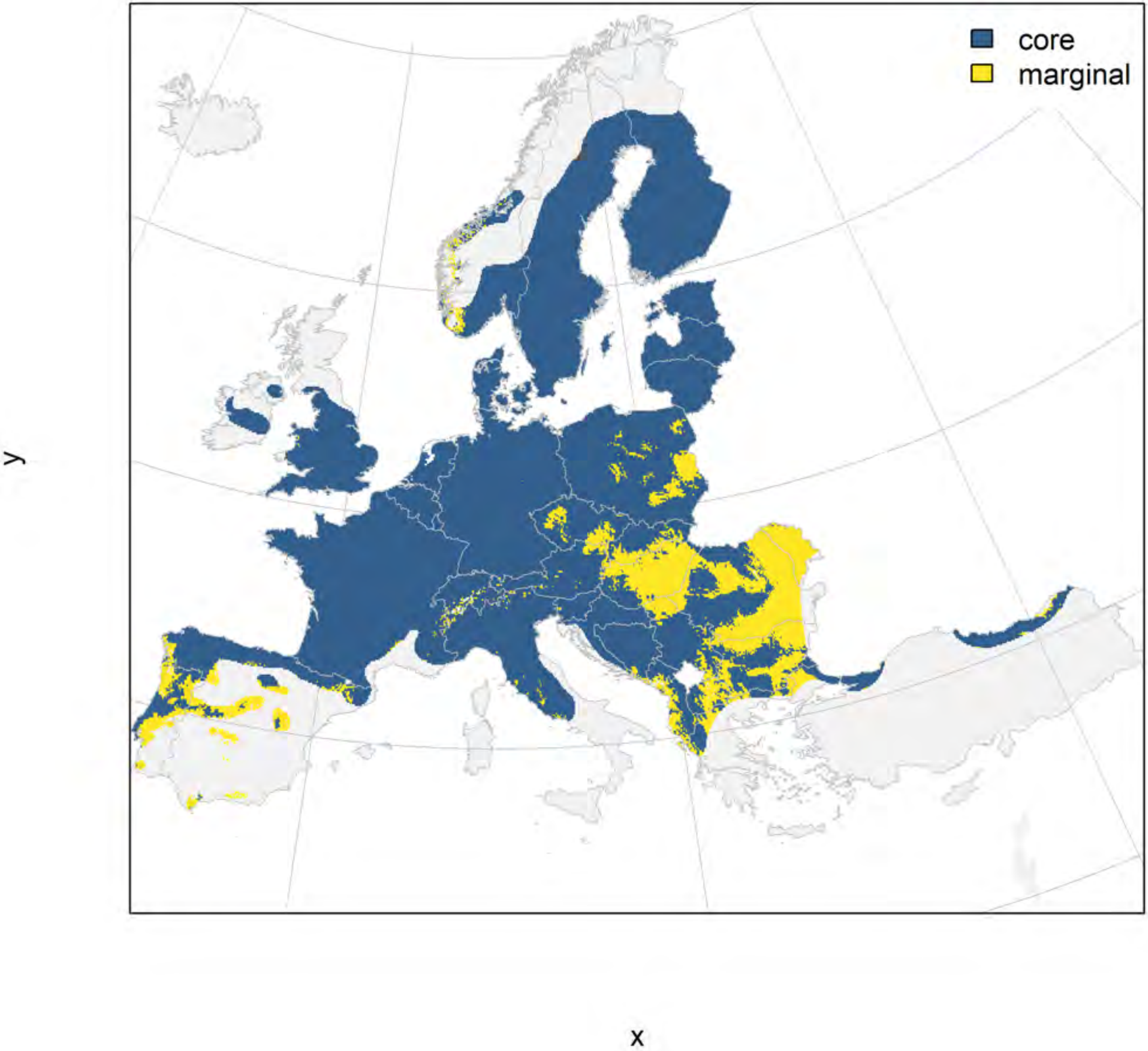

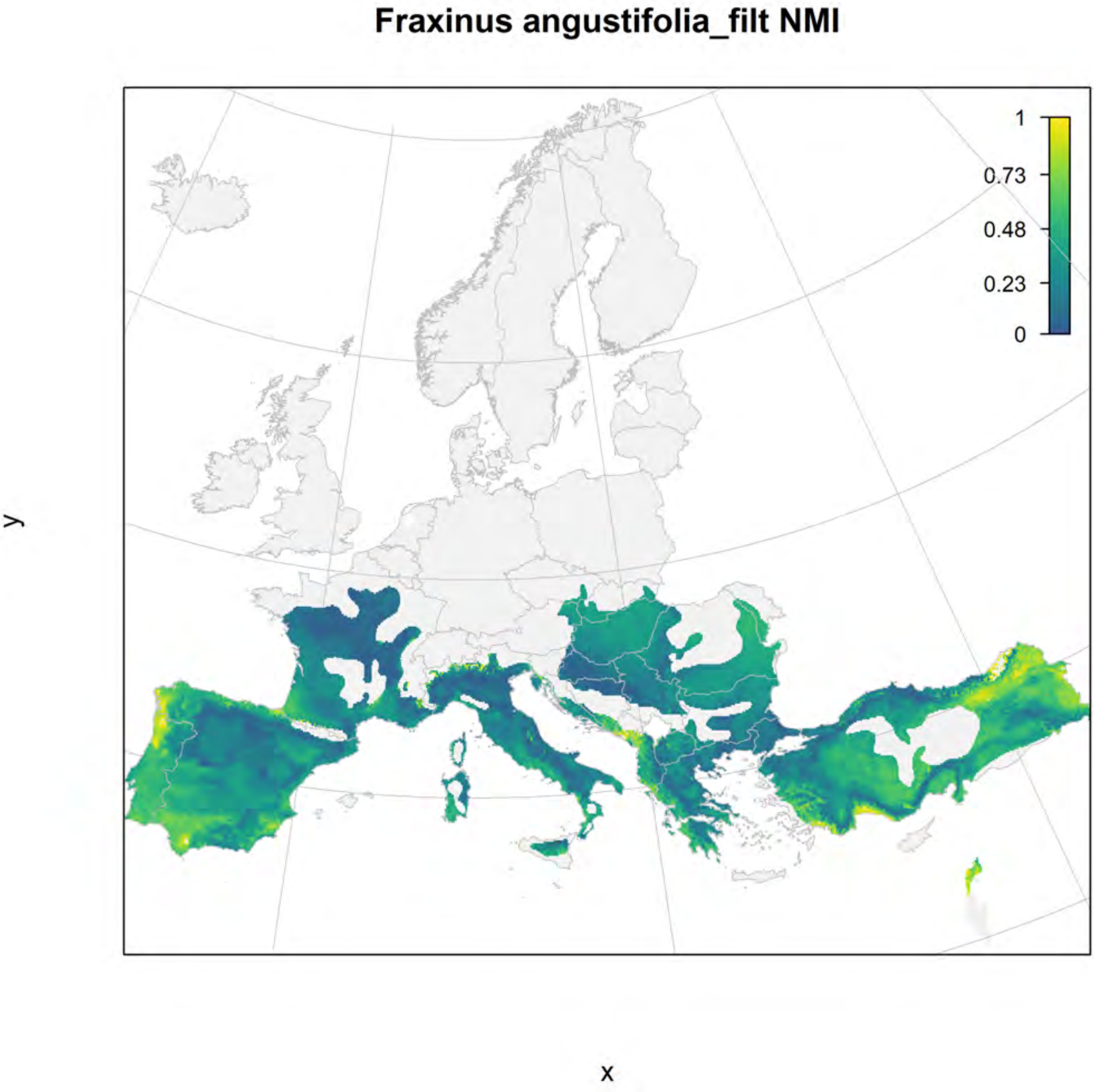
Fraxinus angustifolia_filt NMI
1
0.73
0.48
0.23
0
y
x

#### Fraxinus angustifolia_filt current marginality

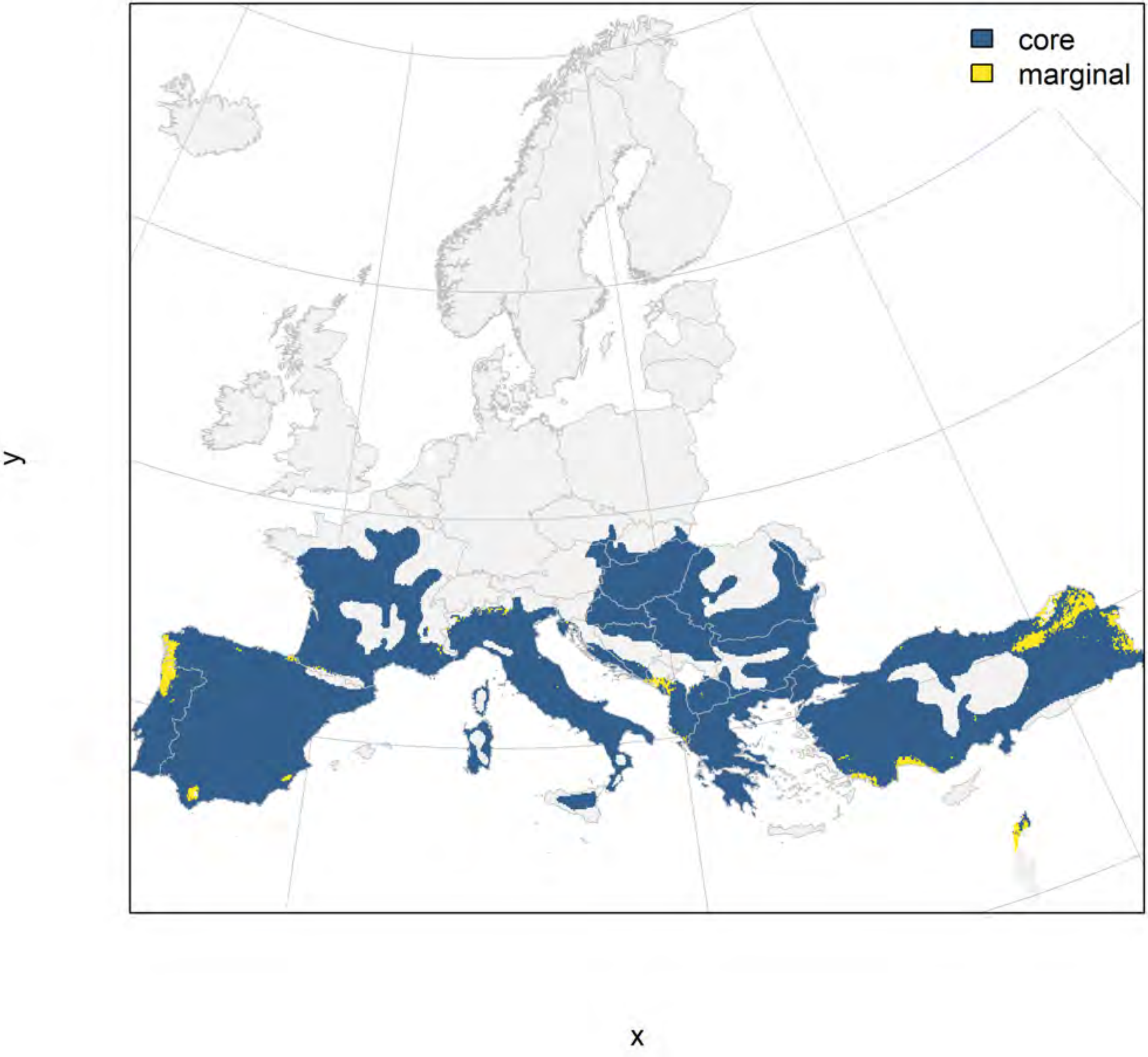

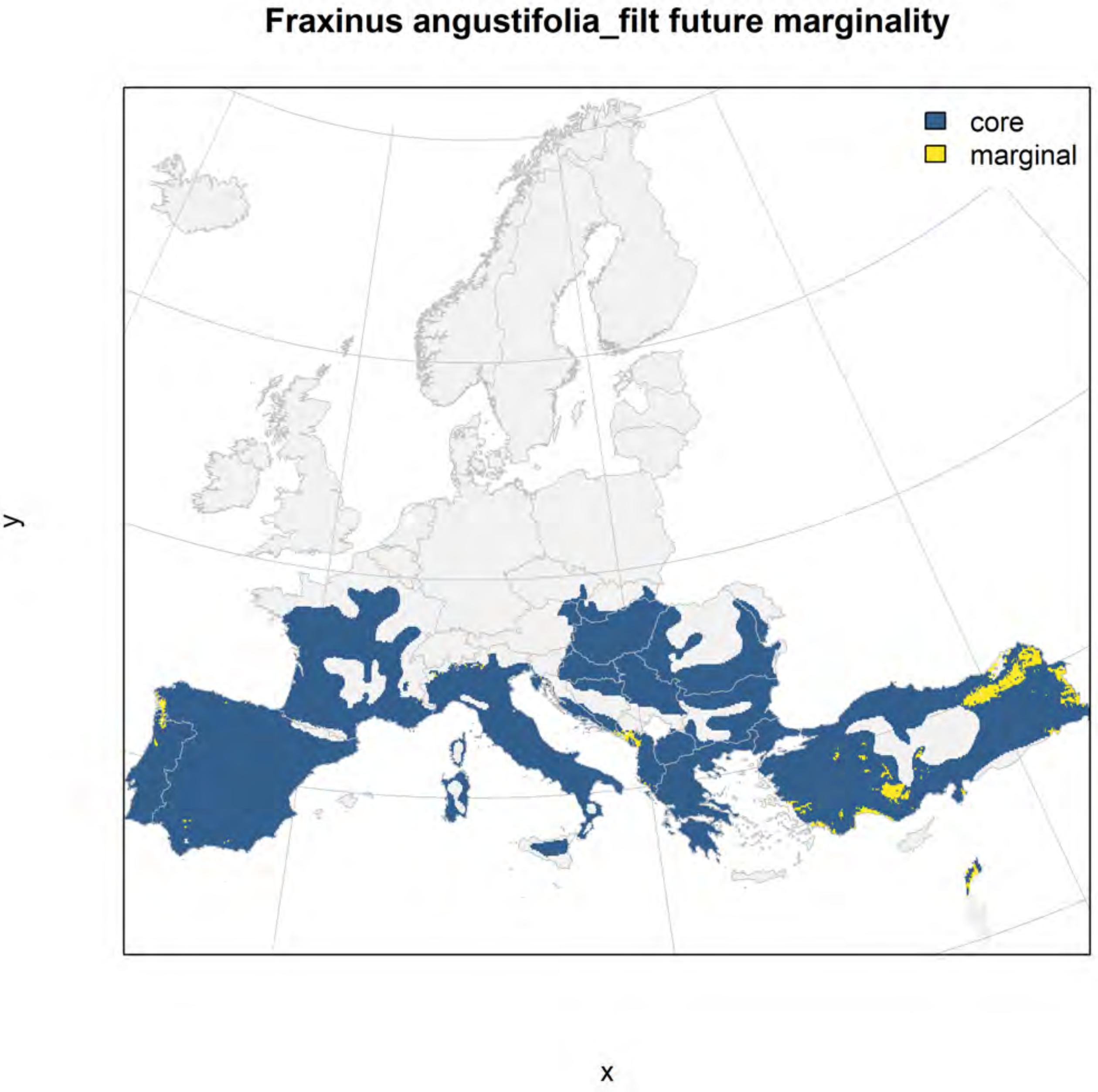
Fraxinus angustifolia_filt future marginality
core
marginal
y
x

Fraxinus excelsior_filt NMI

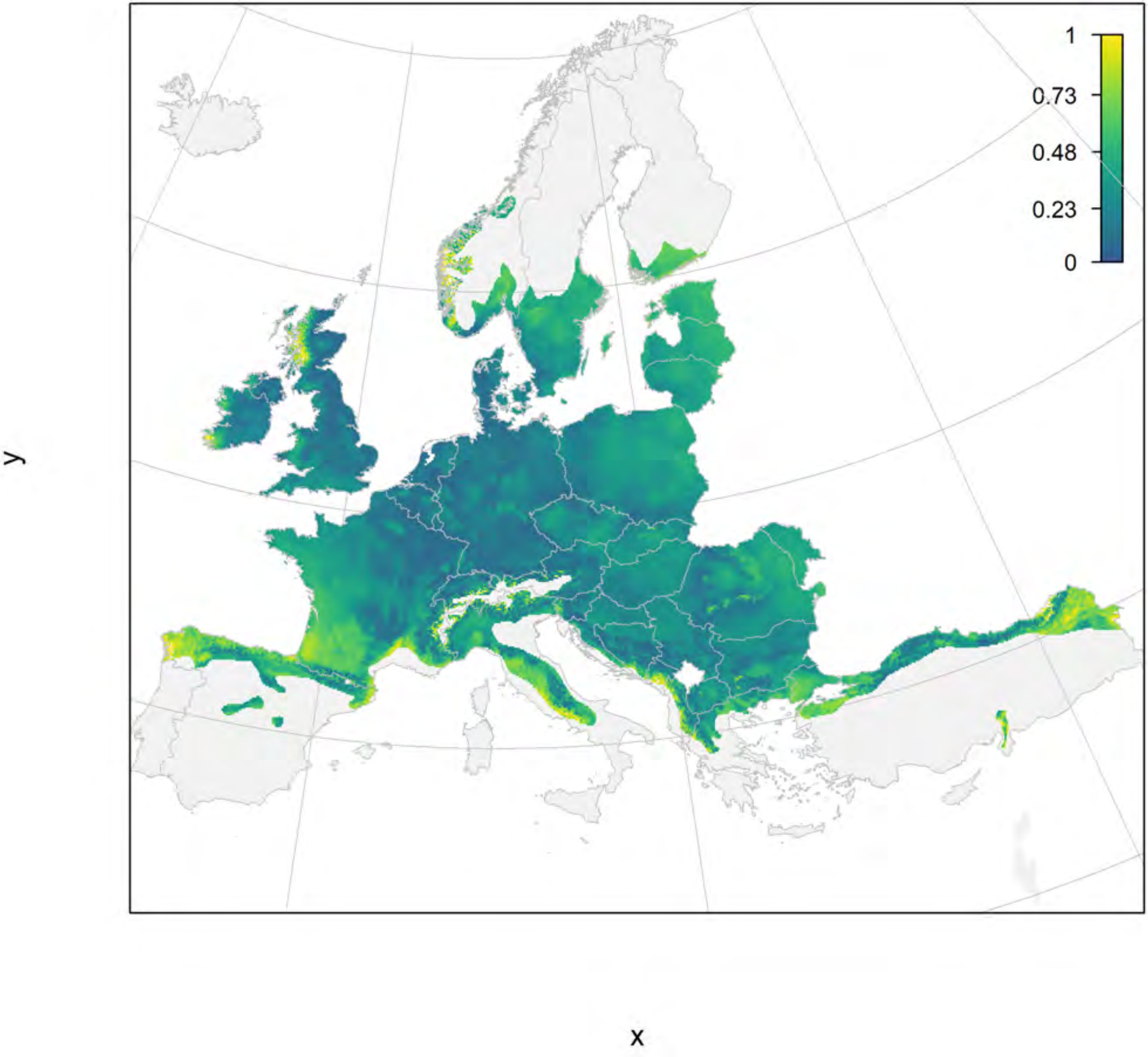

#### Fraxinus excelsior_filt current marginality

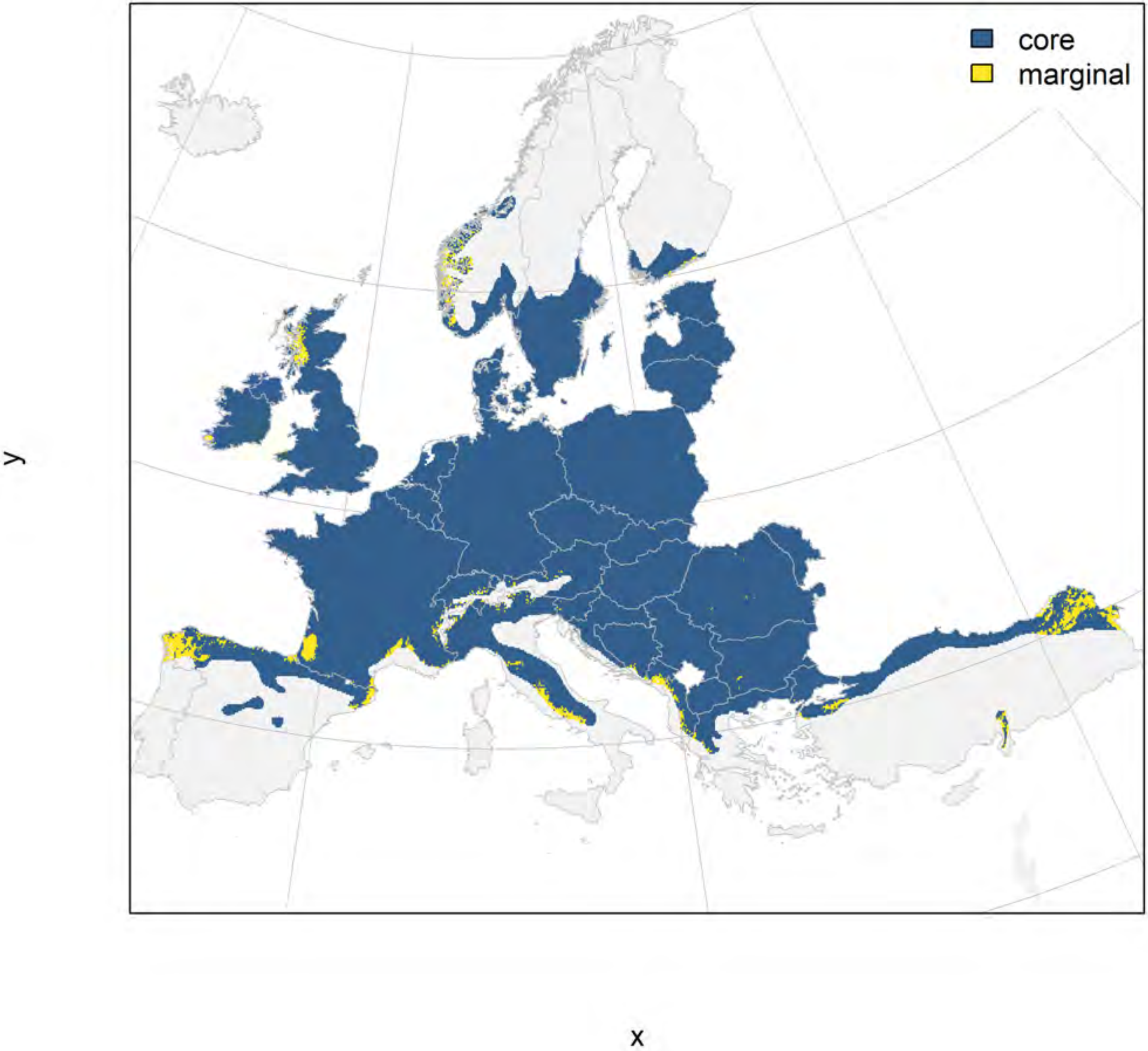

### Fraxinus excelsior_filt future marginality

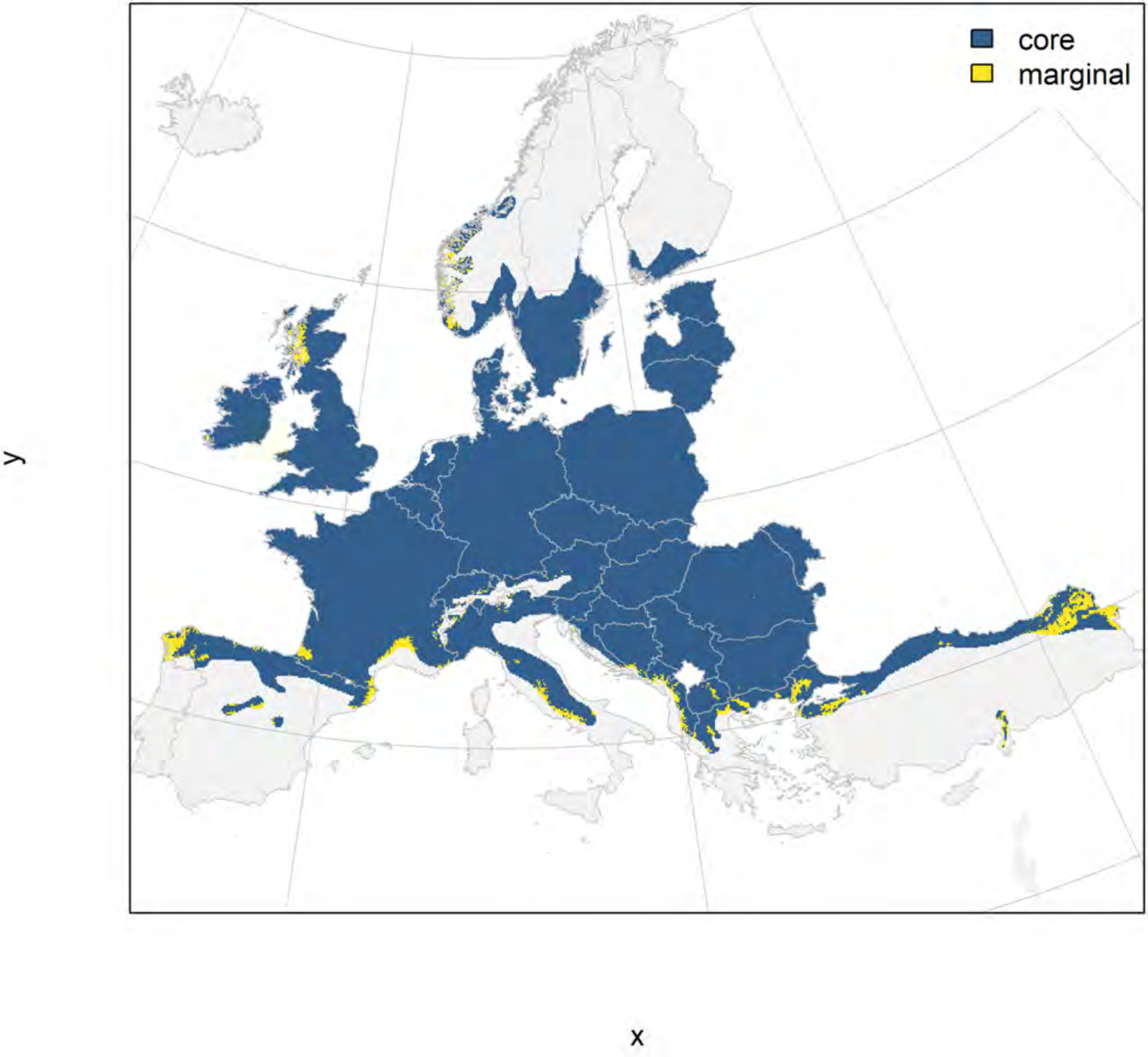

#### Ilex aquifolium_filt NMI

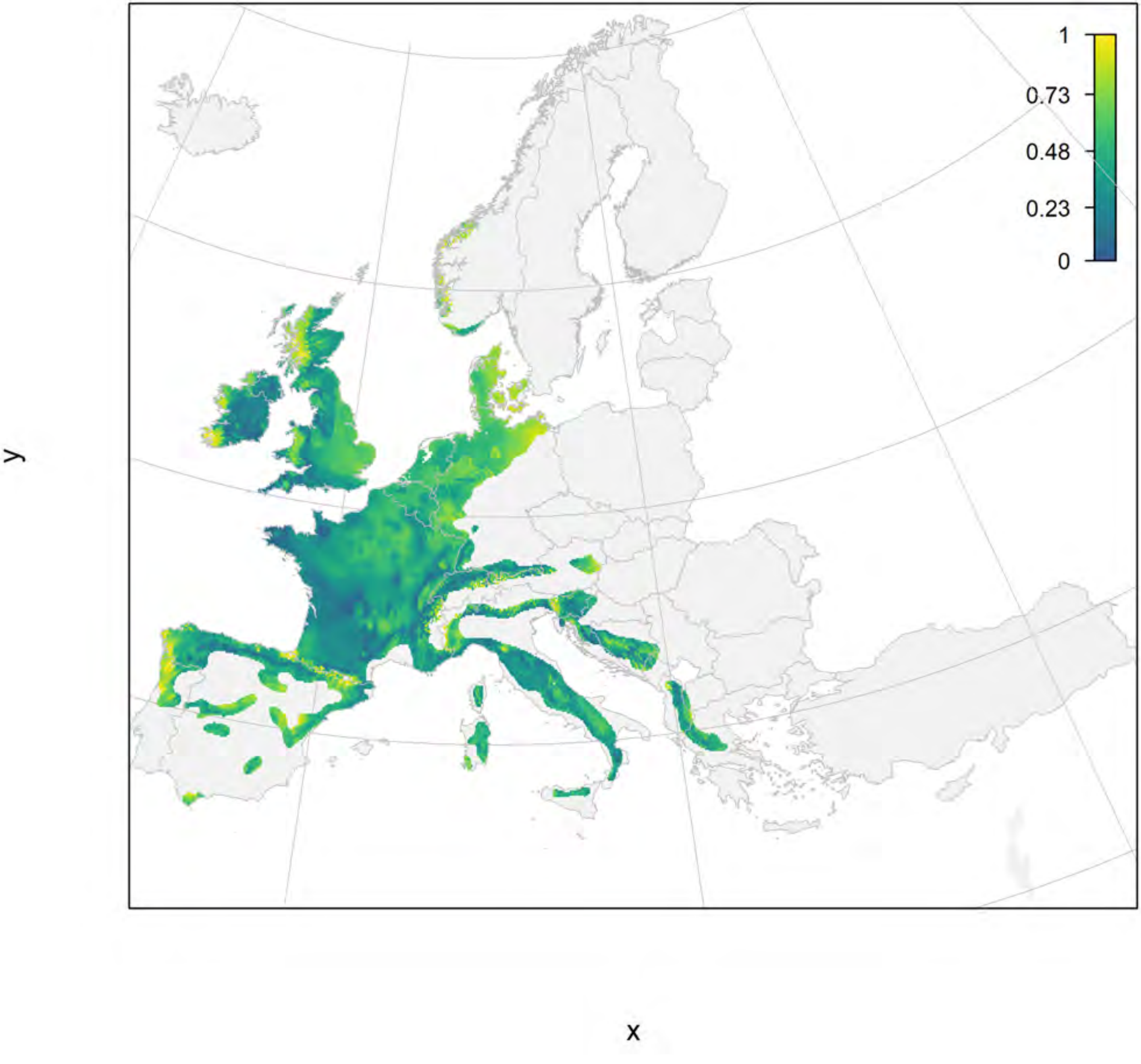

### Ilex aquifolium_filt current marginality

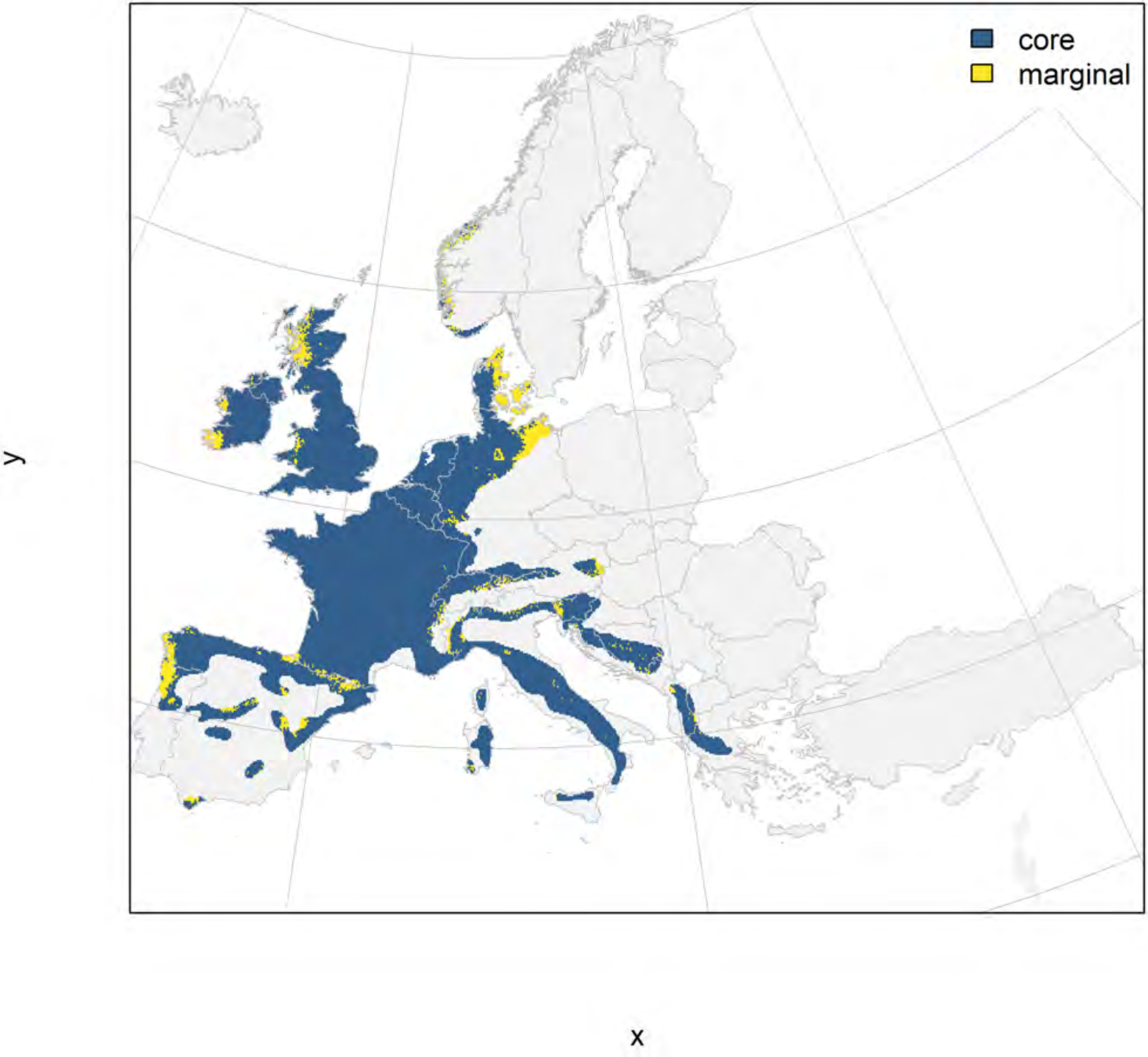

Ilex aquifolium_filt future marginality

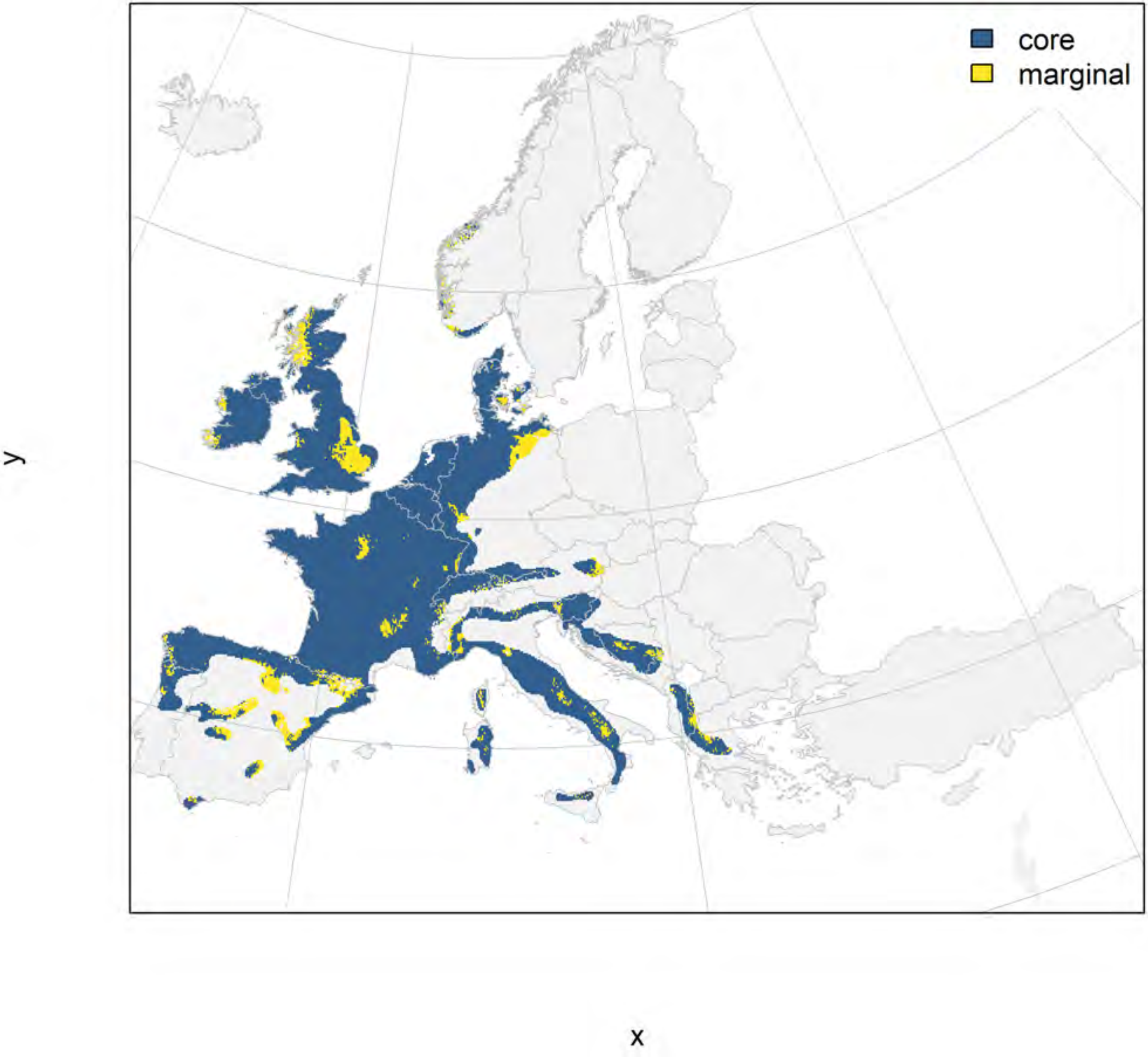

### Juglans regia_filt NMI

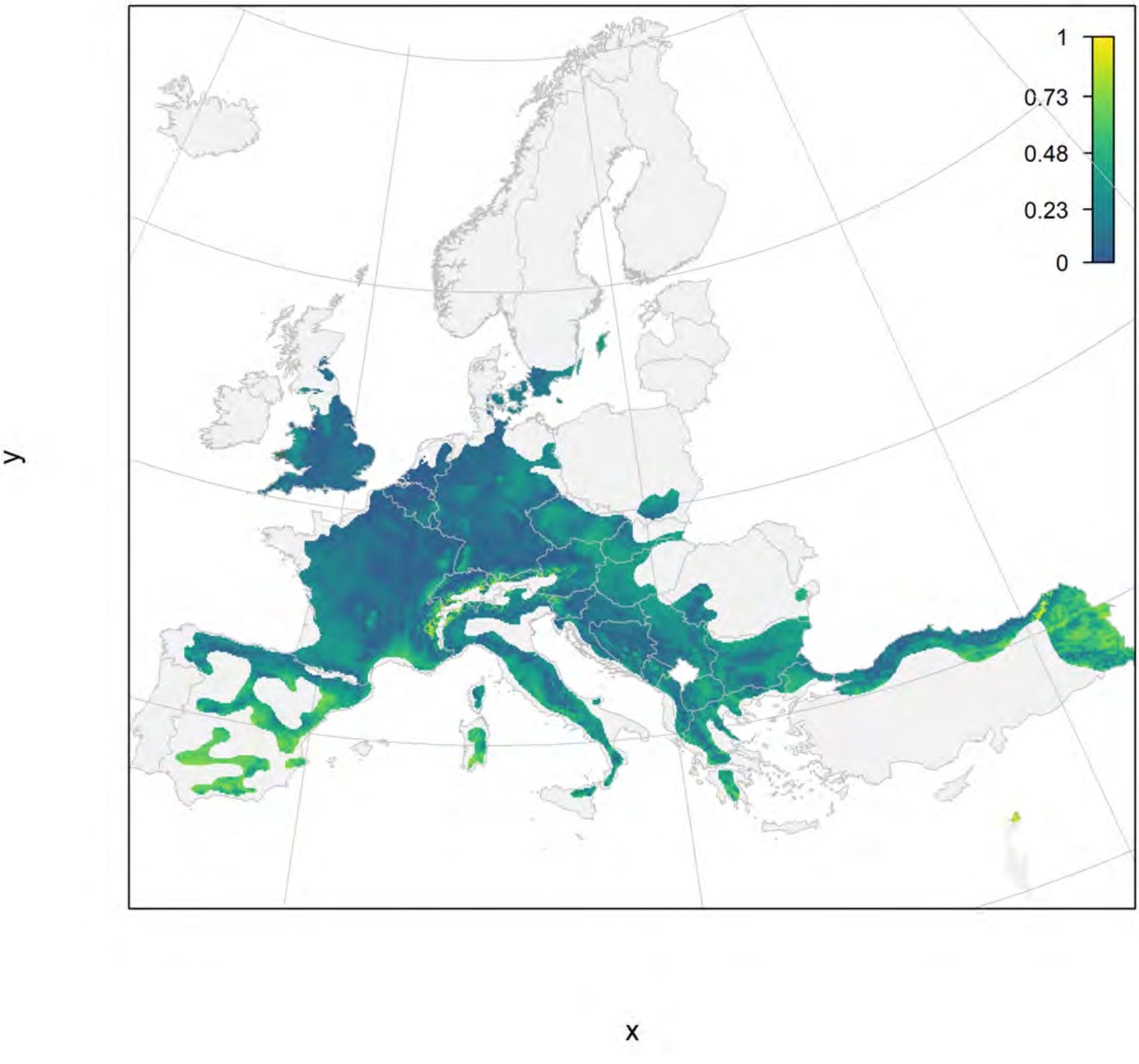

#### Juglans regia_filt current marginality

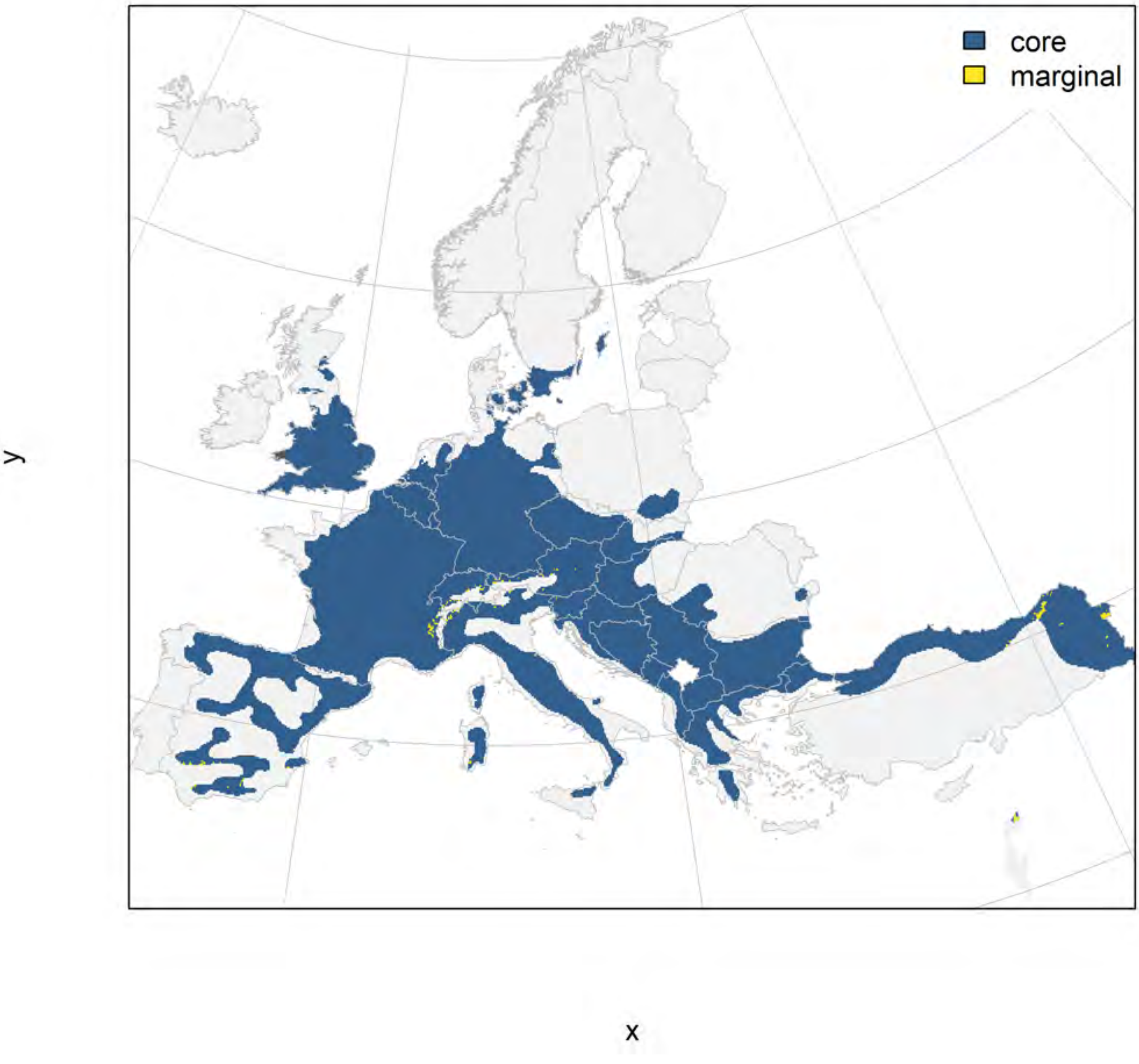

### Juglans regia_filt future marginality

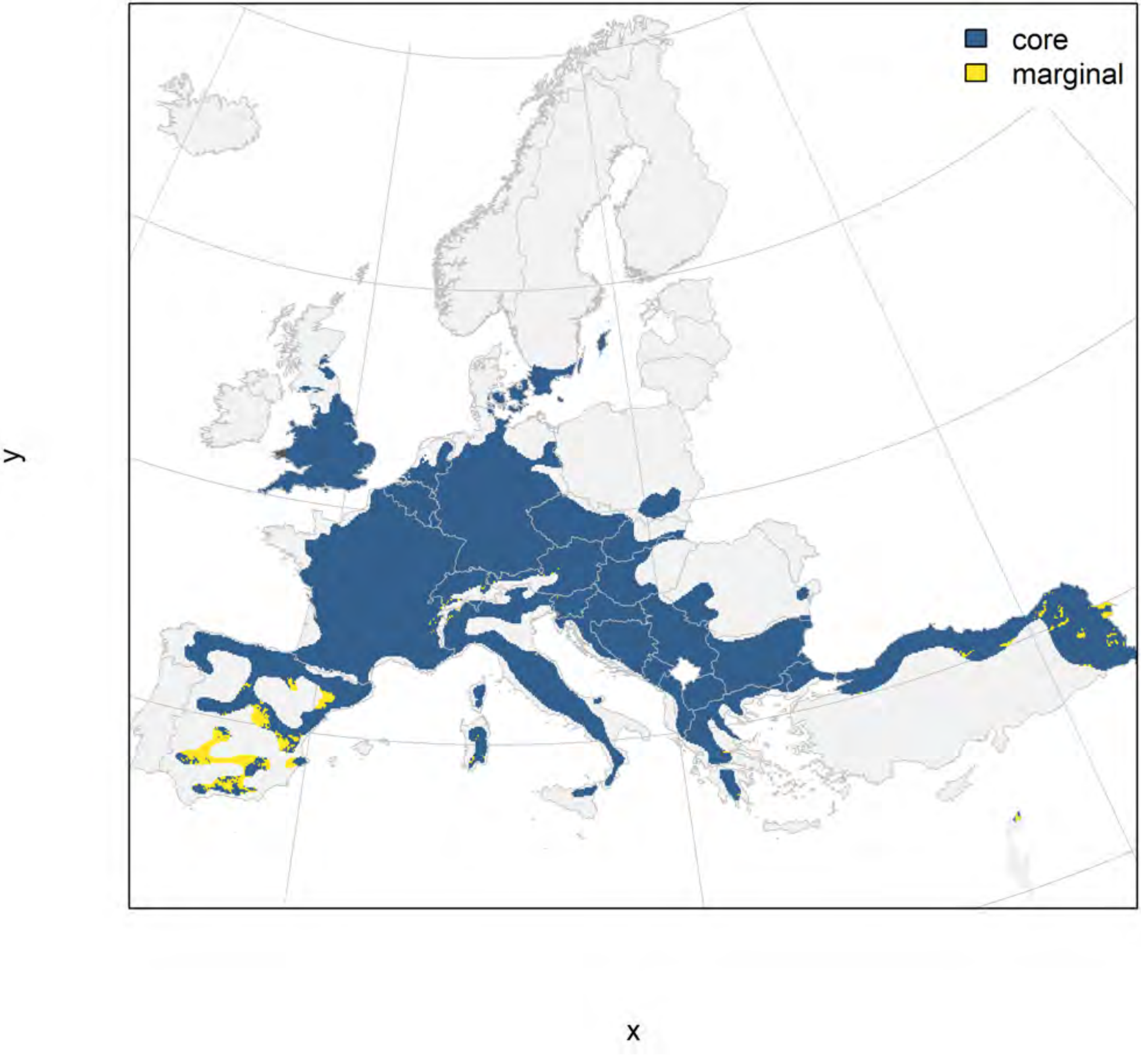

### Juniperus communis_filt NMI

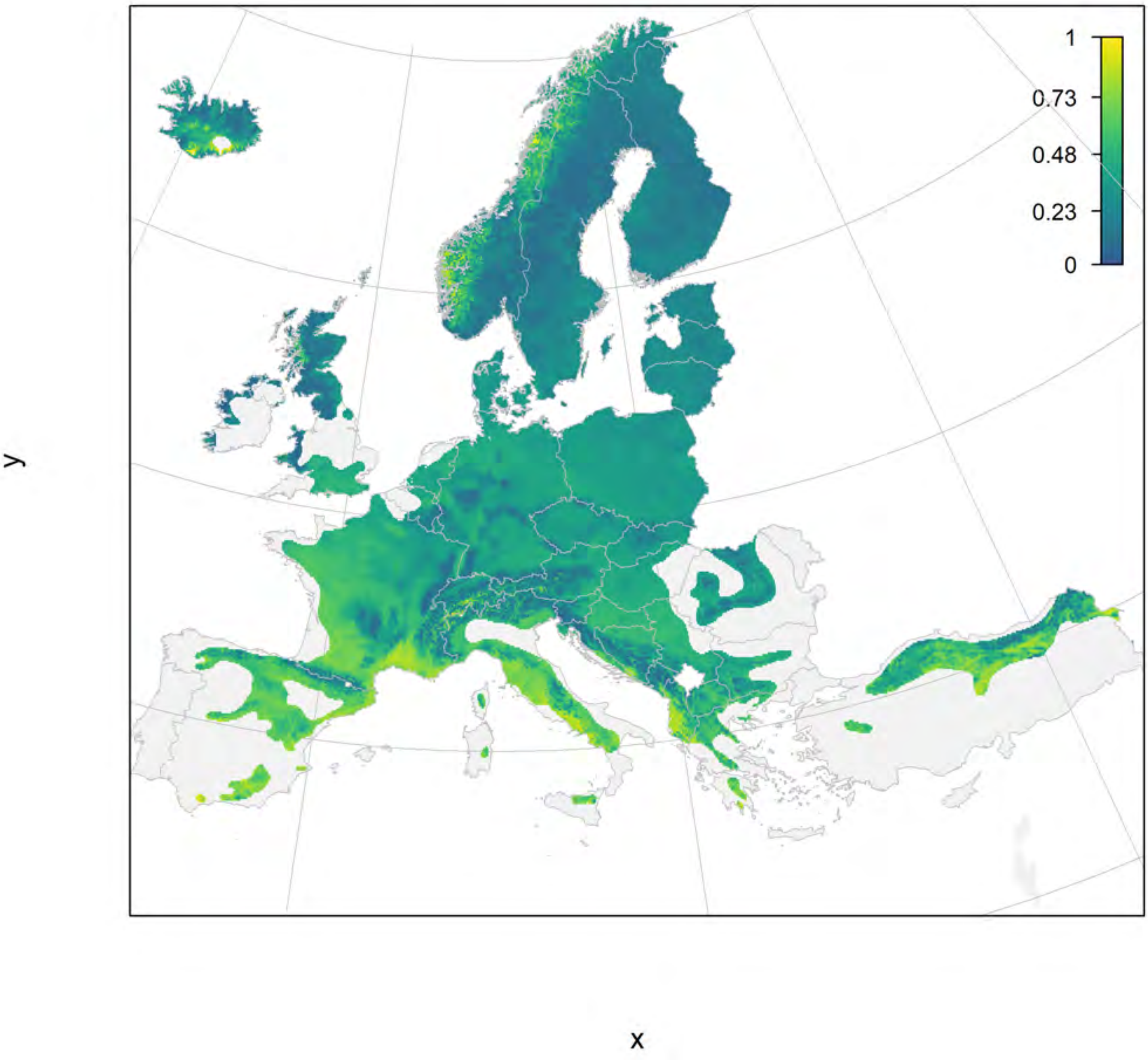

#### Juniperus communis_filt current marginality

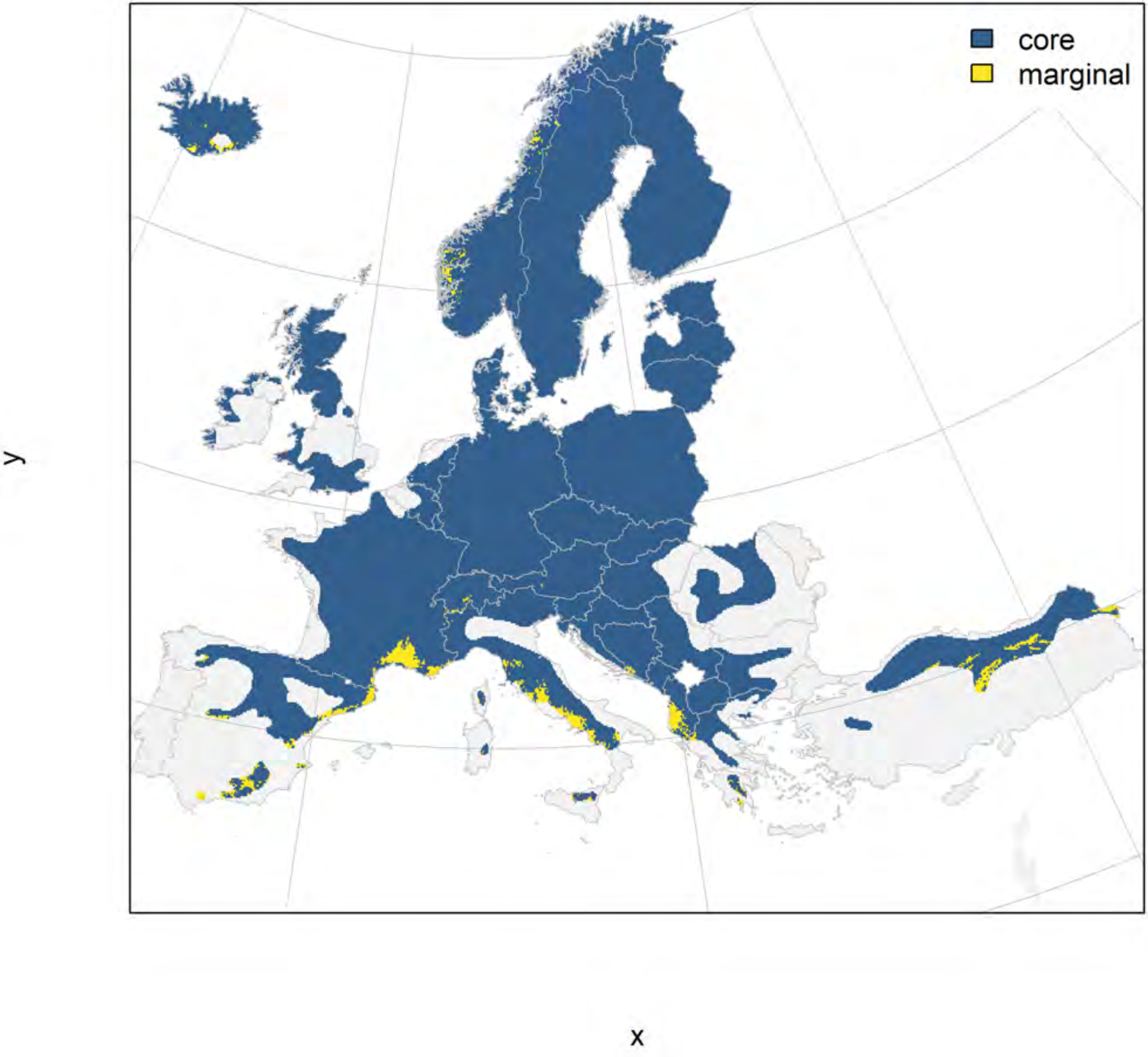

### Juniperus communis_filt future marginality

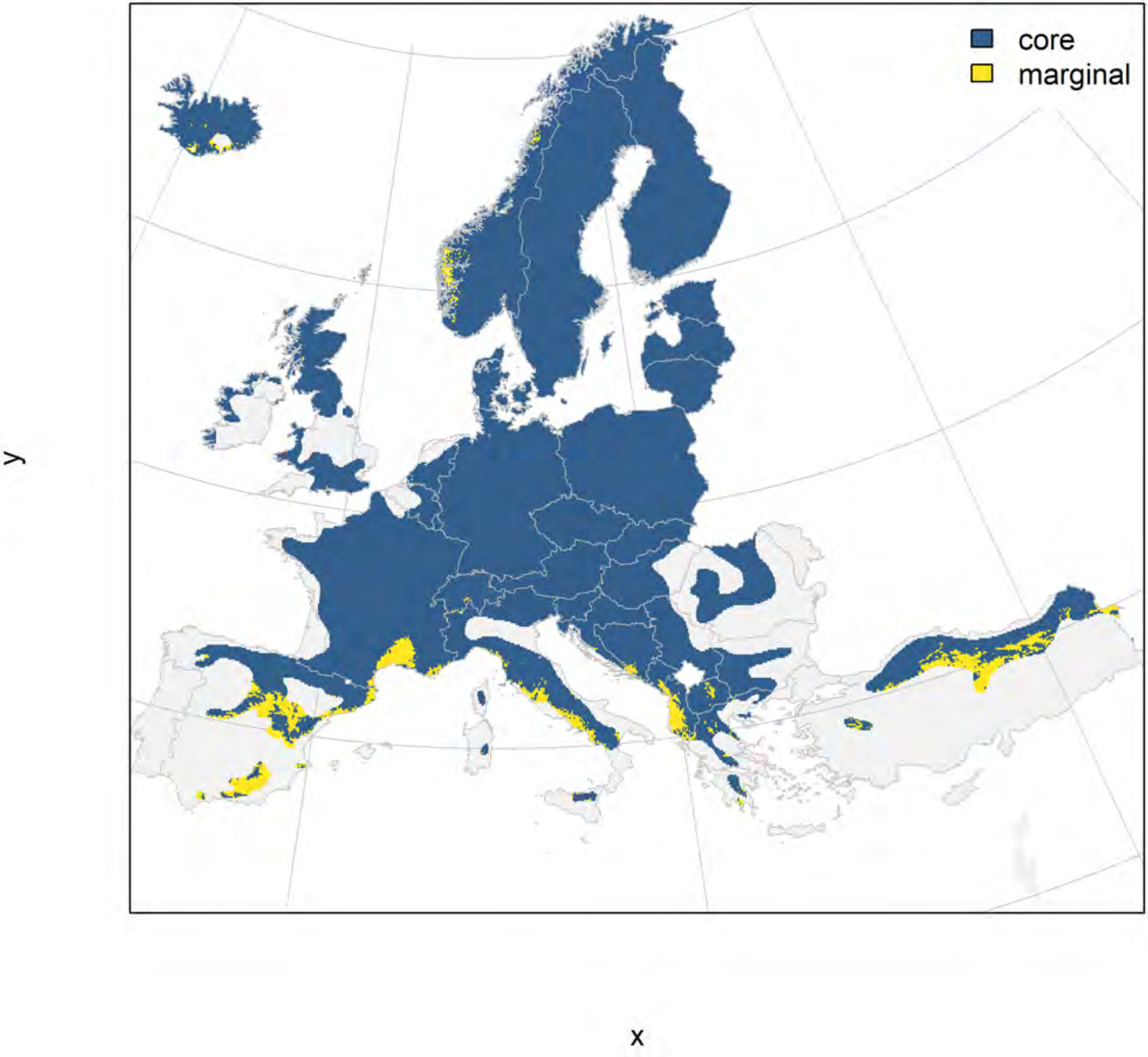

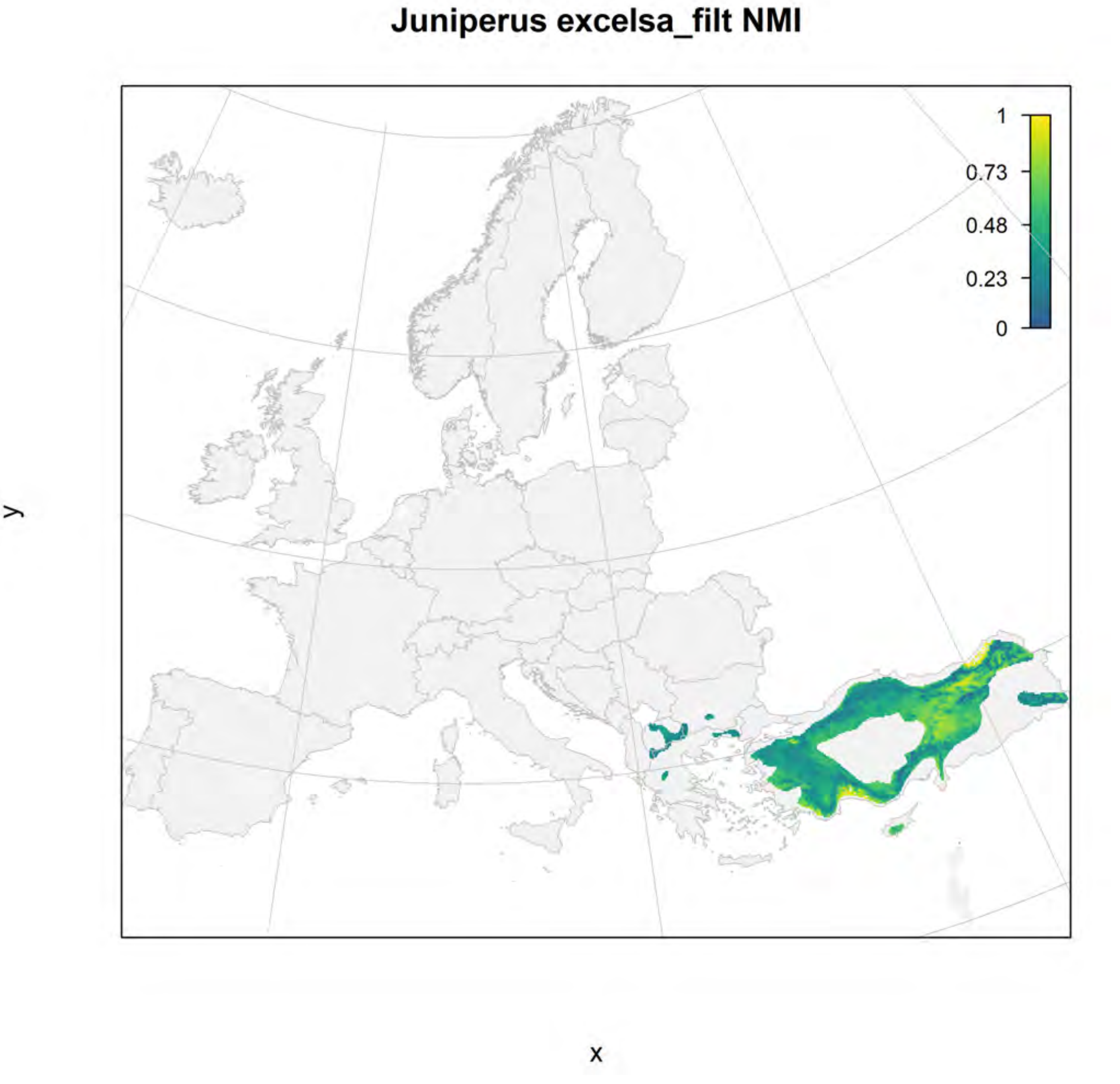

Juniperus excelsa_filt NMI
1
0.73
0.48
0.23
0
y
x

### Juniperus excelsa_filt current marginality

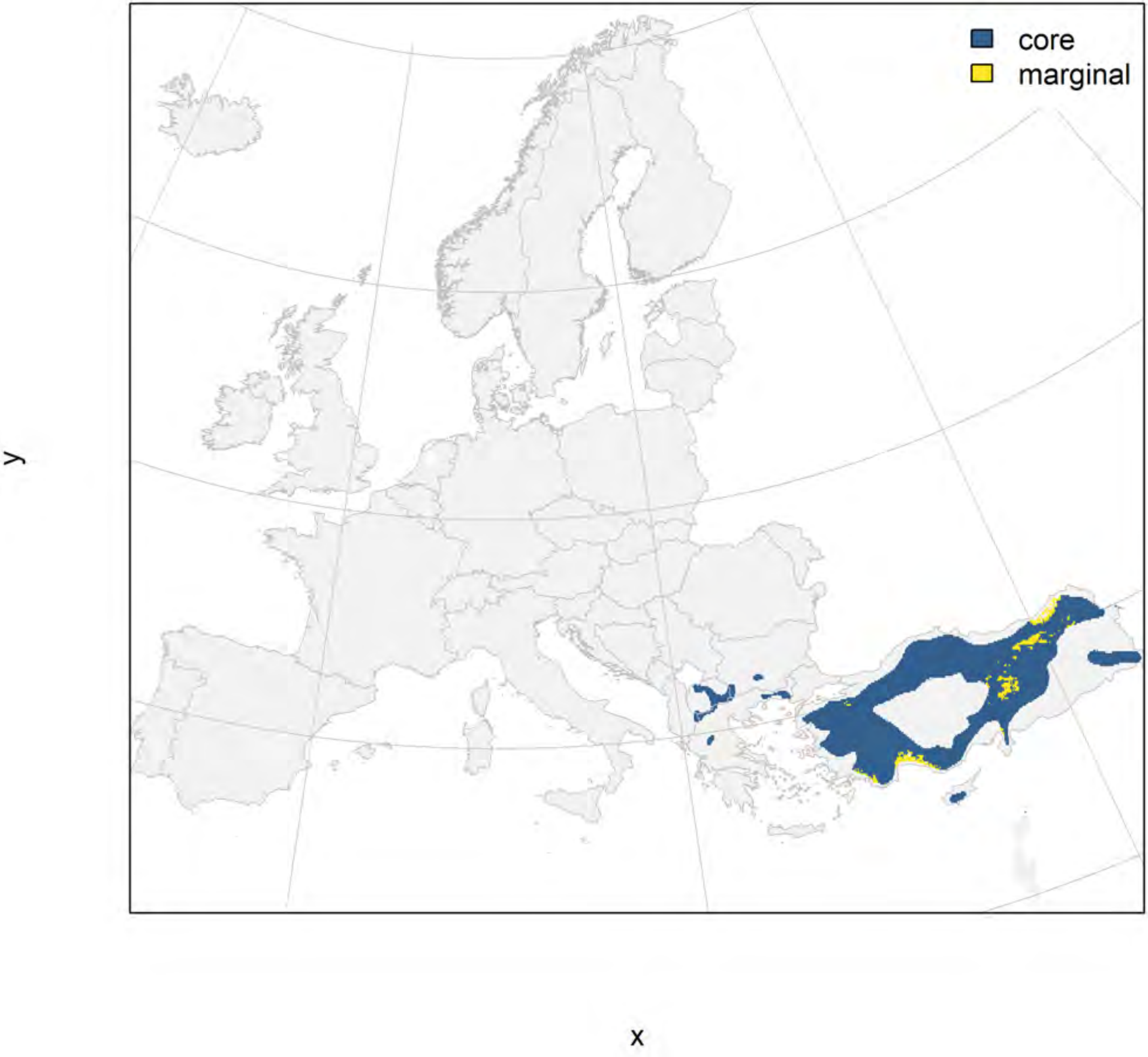

### Juniperus excelsa_filt future marginality

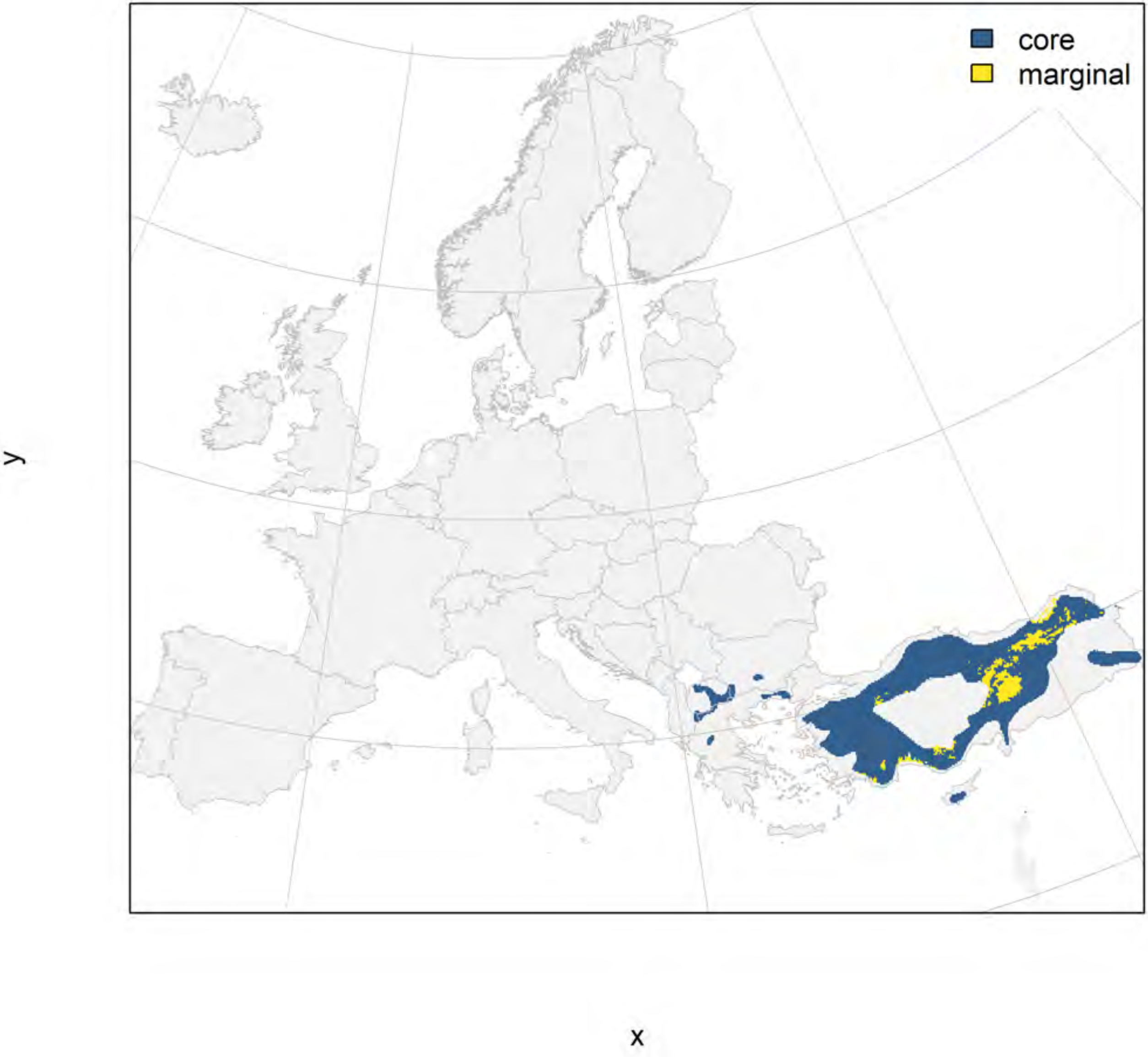

### Juniperus oxycedrus_filt NMI

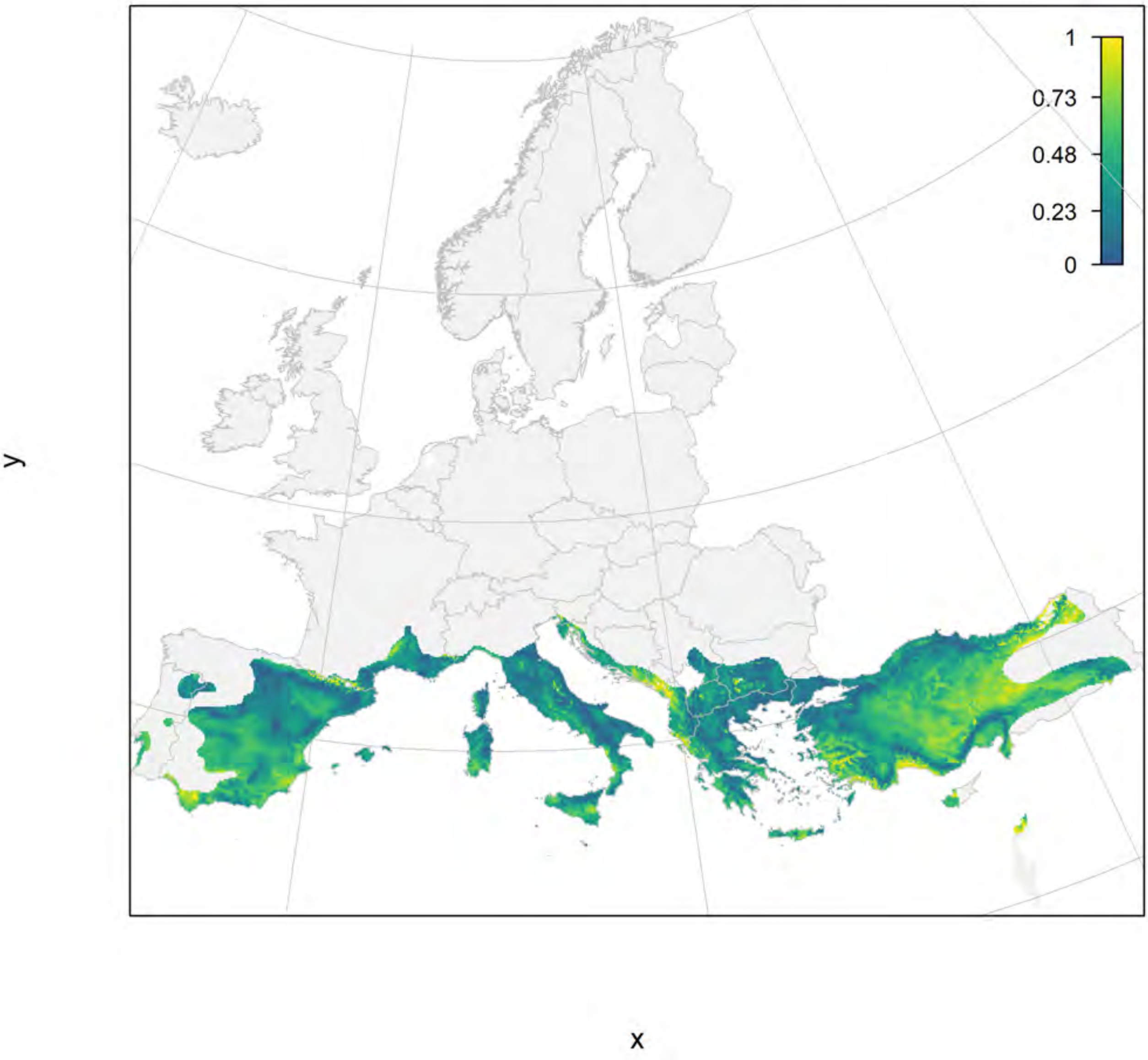

#### Juniperus oxycedrus_filt current marginality

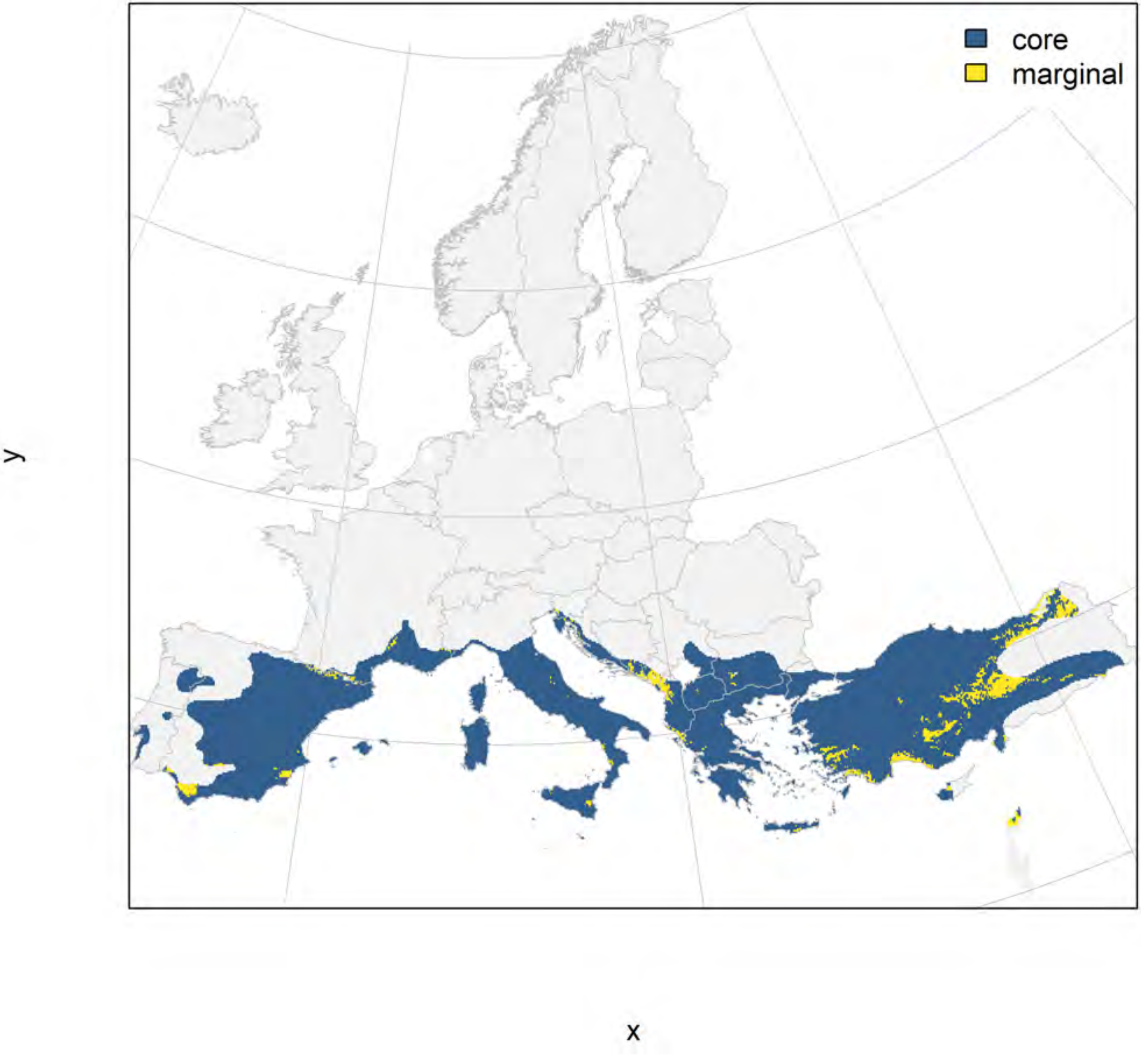

#### Juniperus oxycedrus_filt future marginality

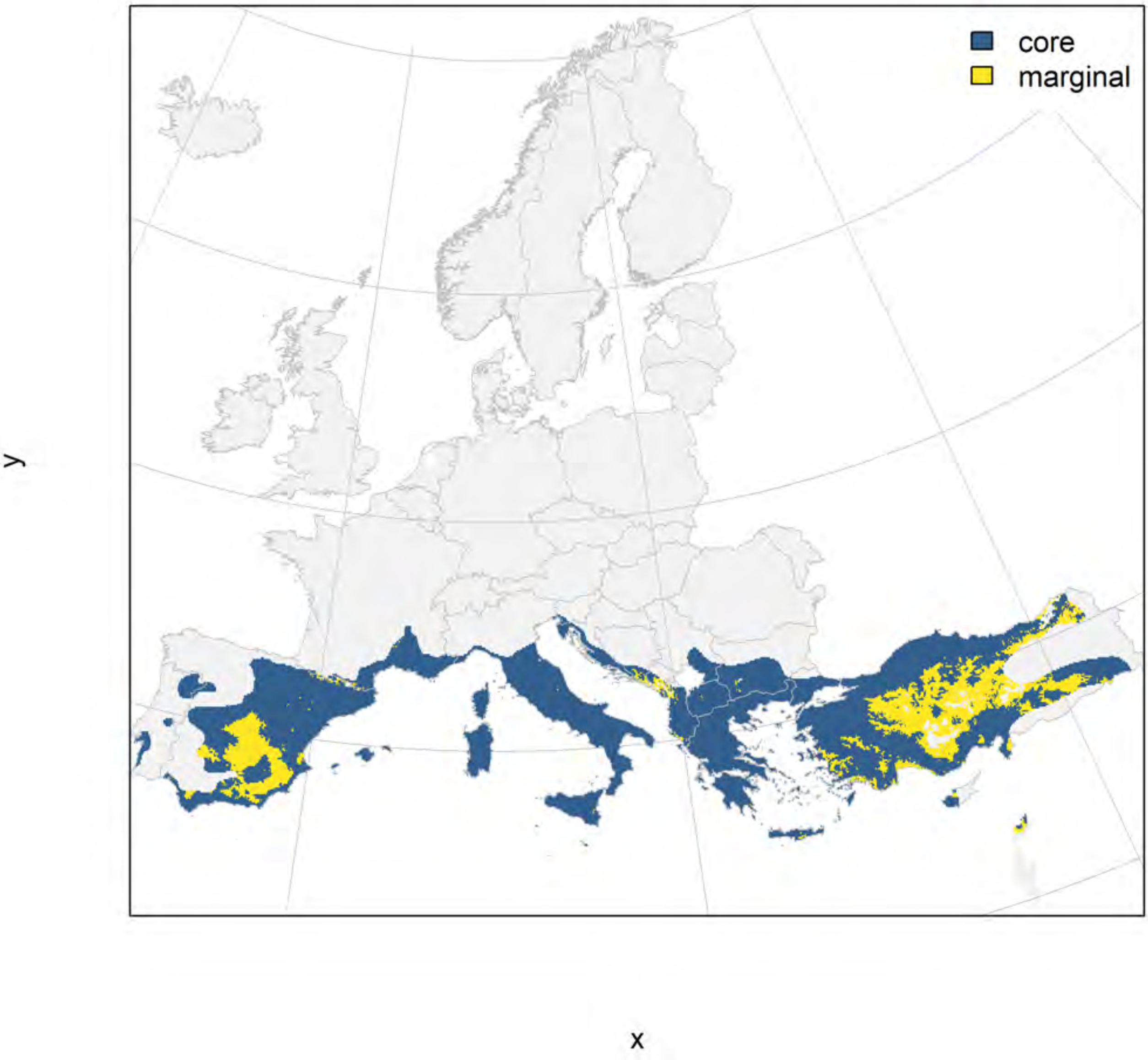

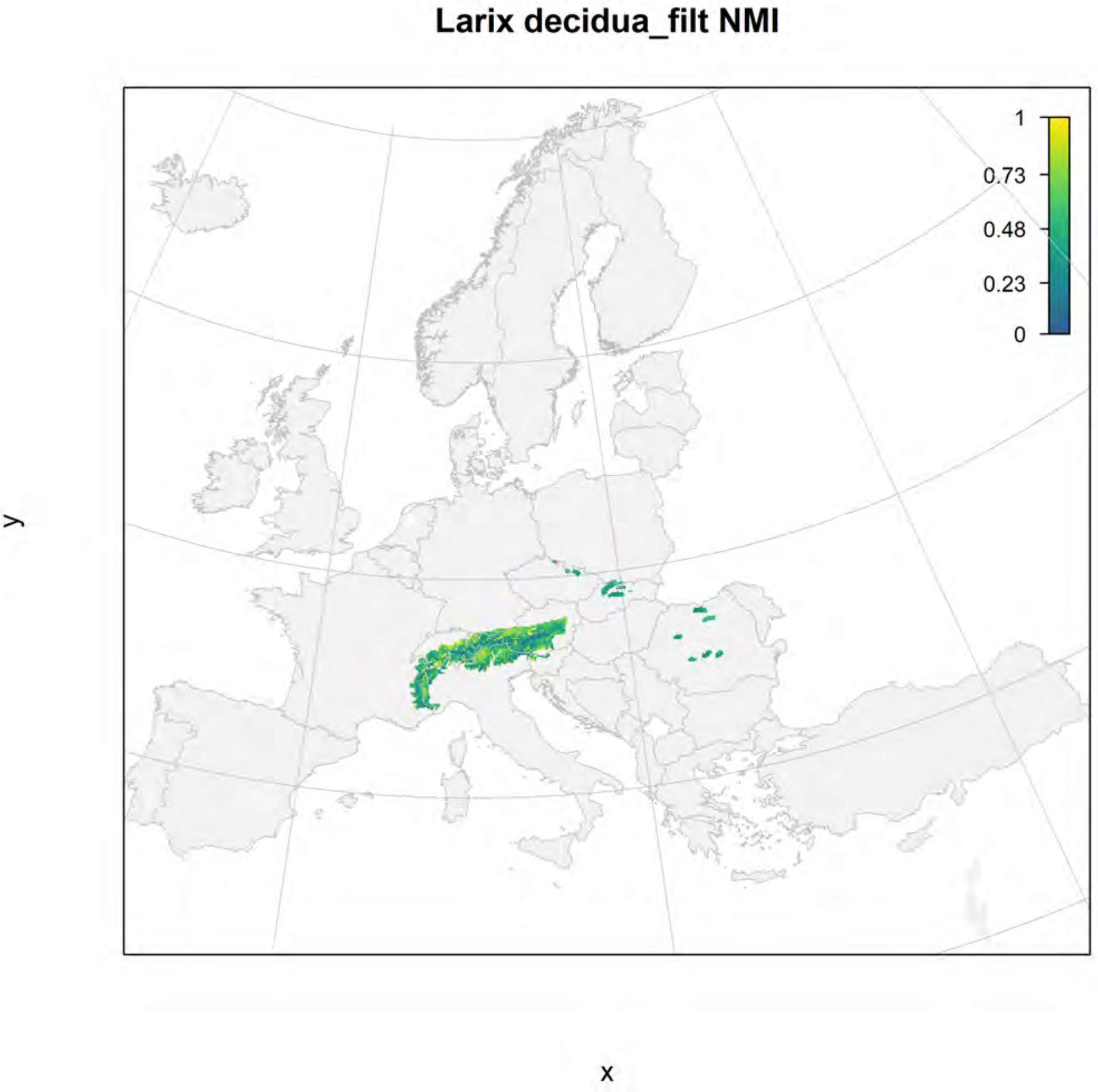
Larix decidua_filt NMI
1
0.73
0.48
0.23
0
y
x

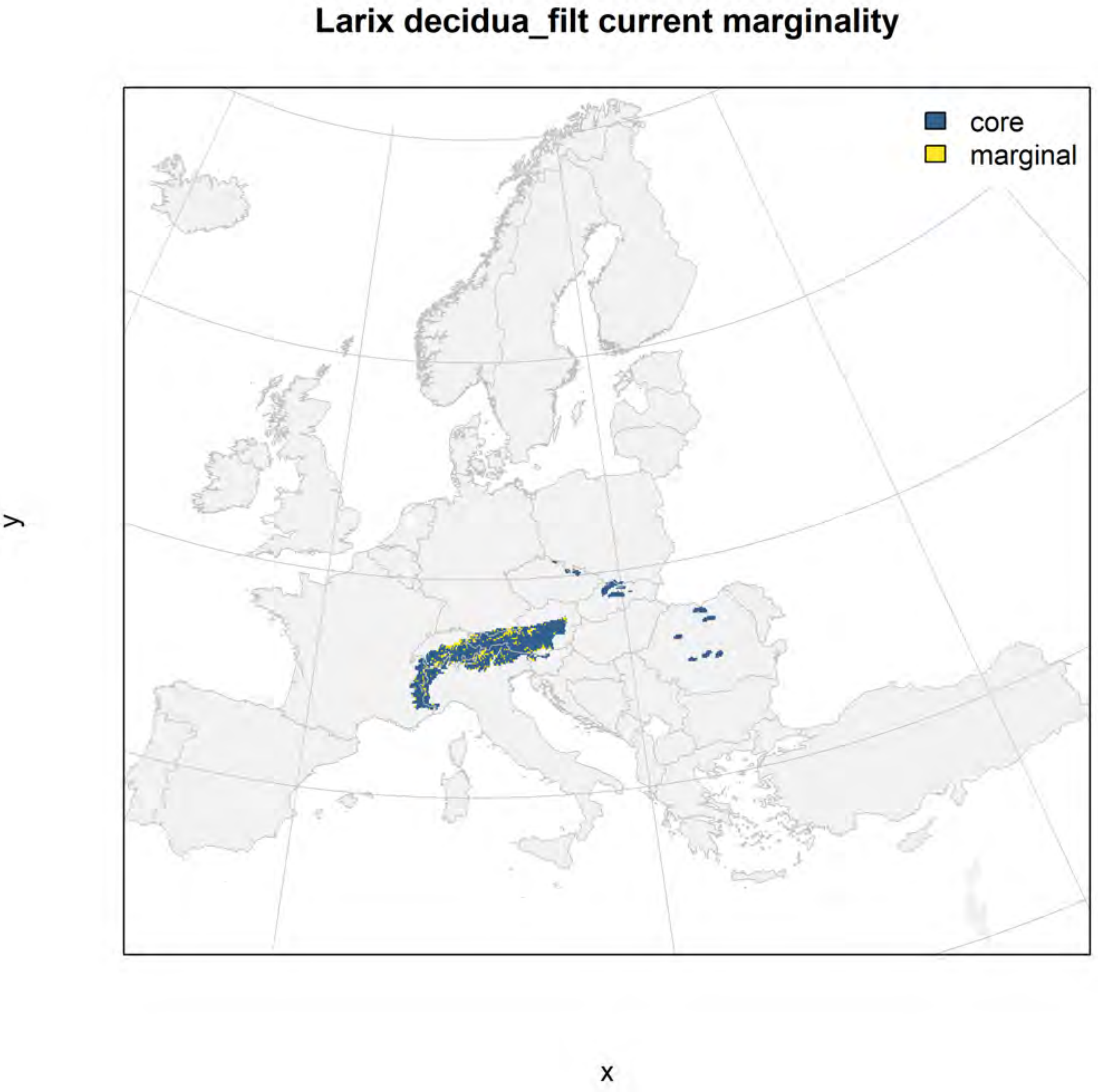
Larix decidua_filt current marginality
core
marginal
y
x

#### Larix decidua_filt future marginality

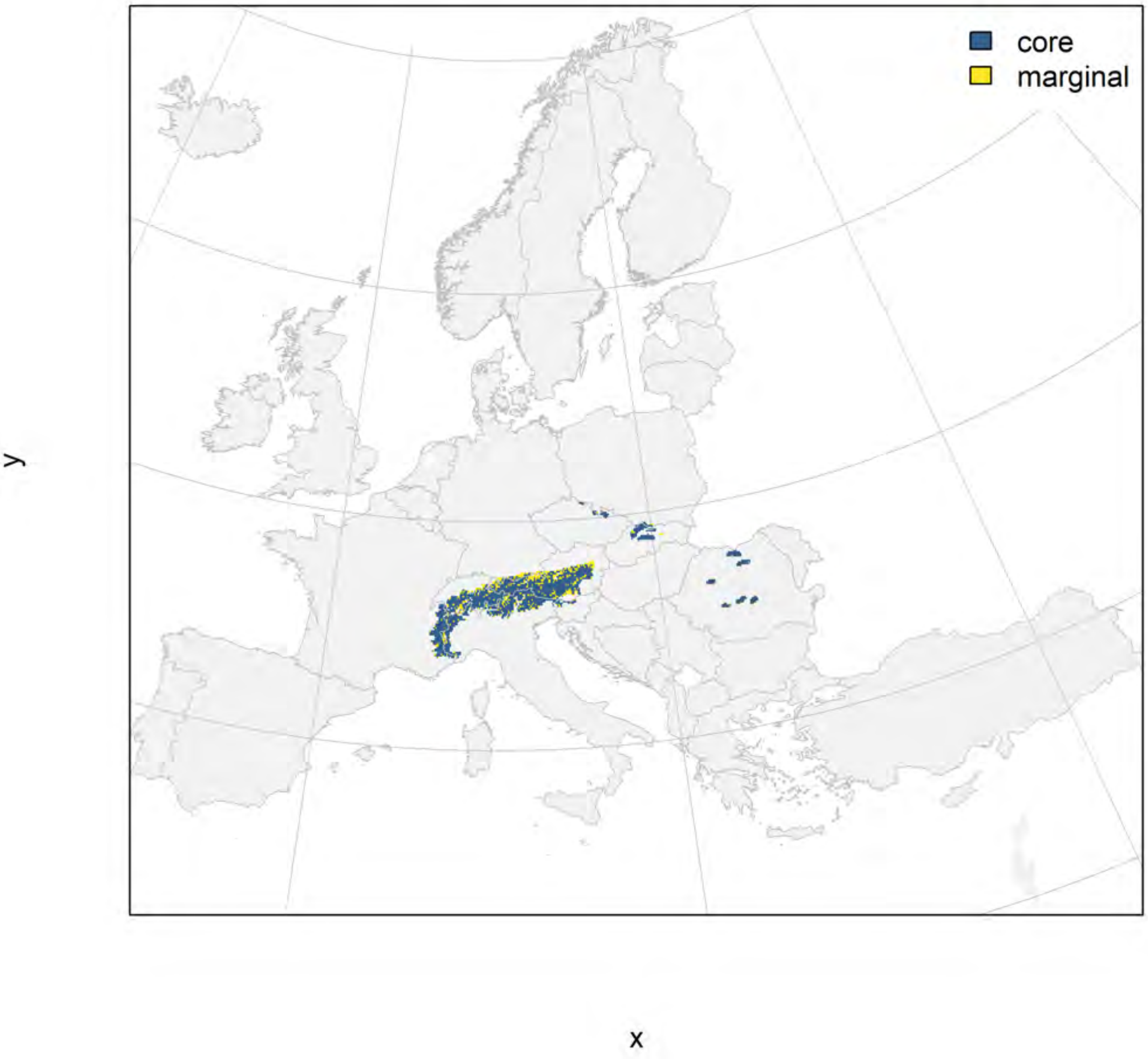

Liquidambar orientalis_filt NMI

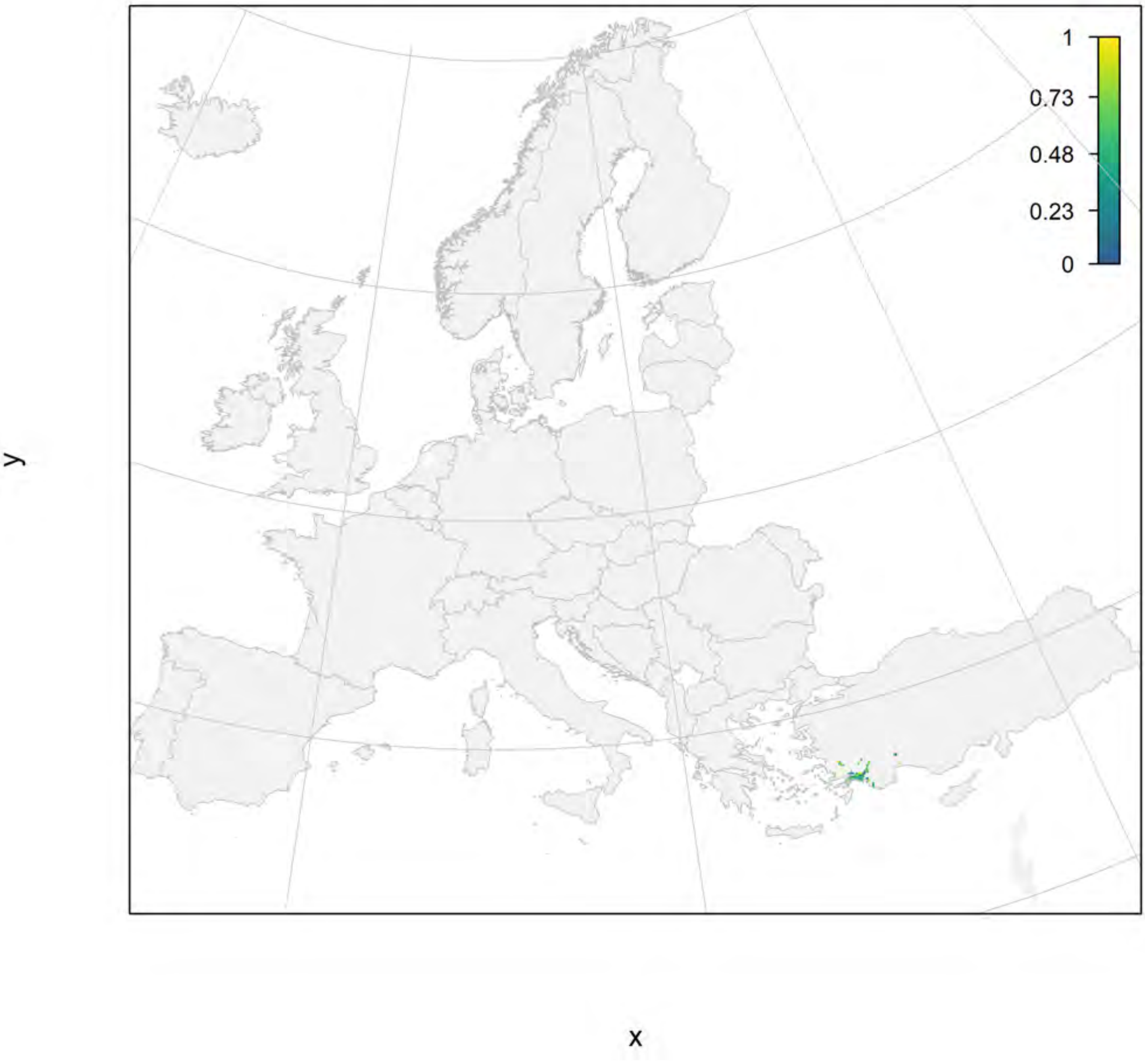

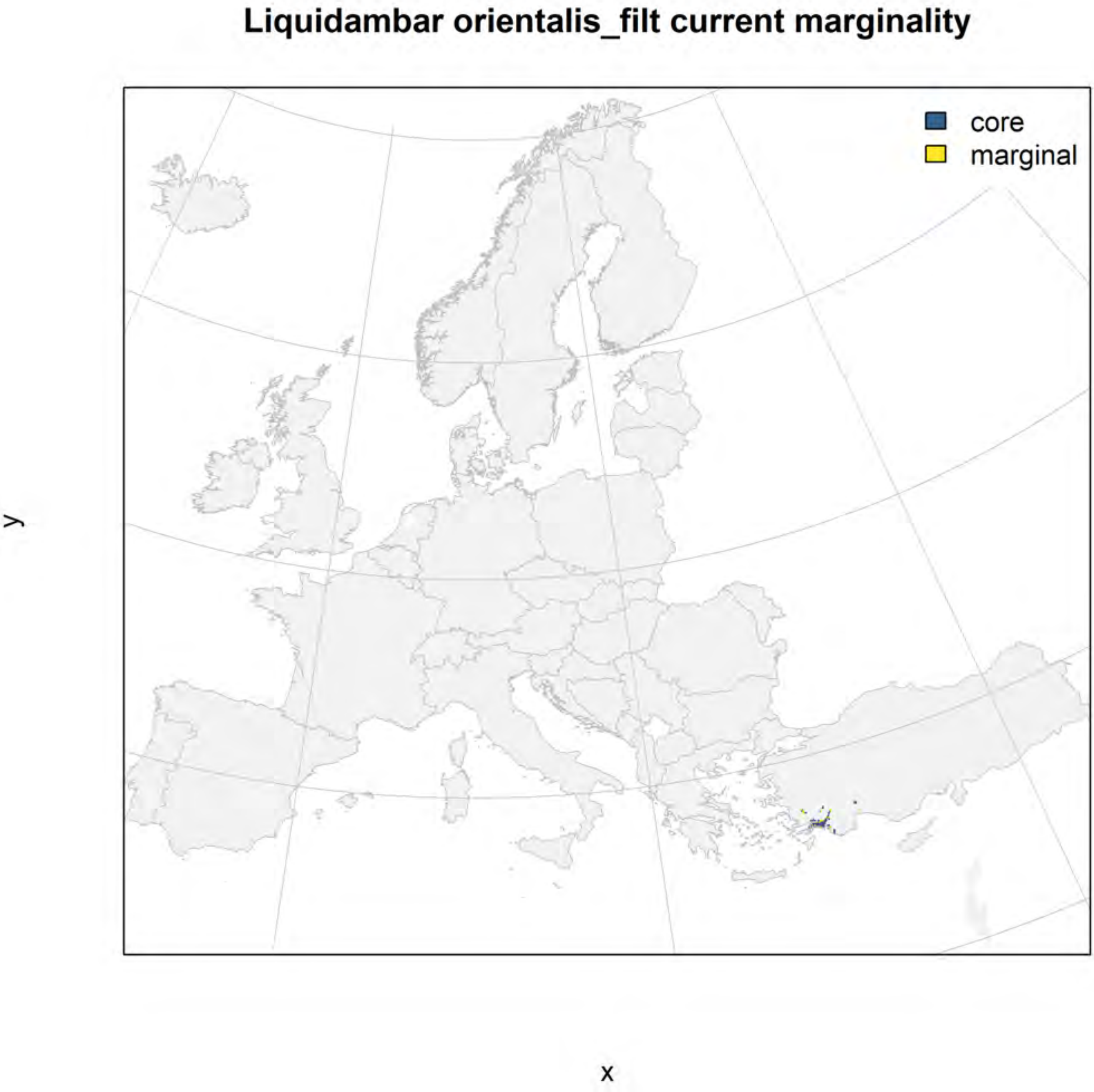

Liquidambar orientalis_filt current marginality
core
marginal
y
x

#### Liquidambar orientalis_filt future marginality

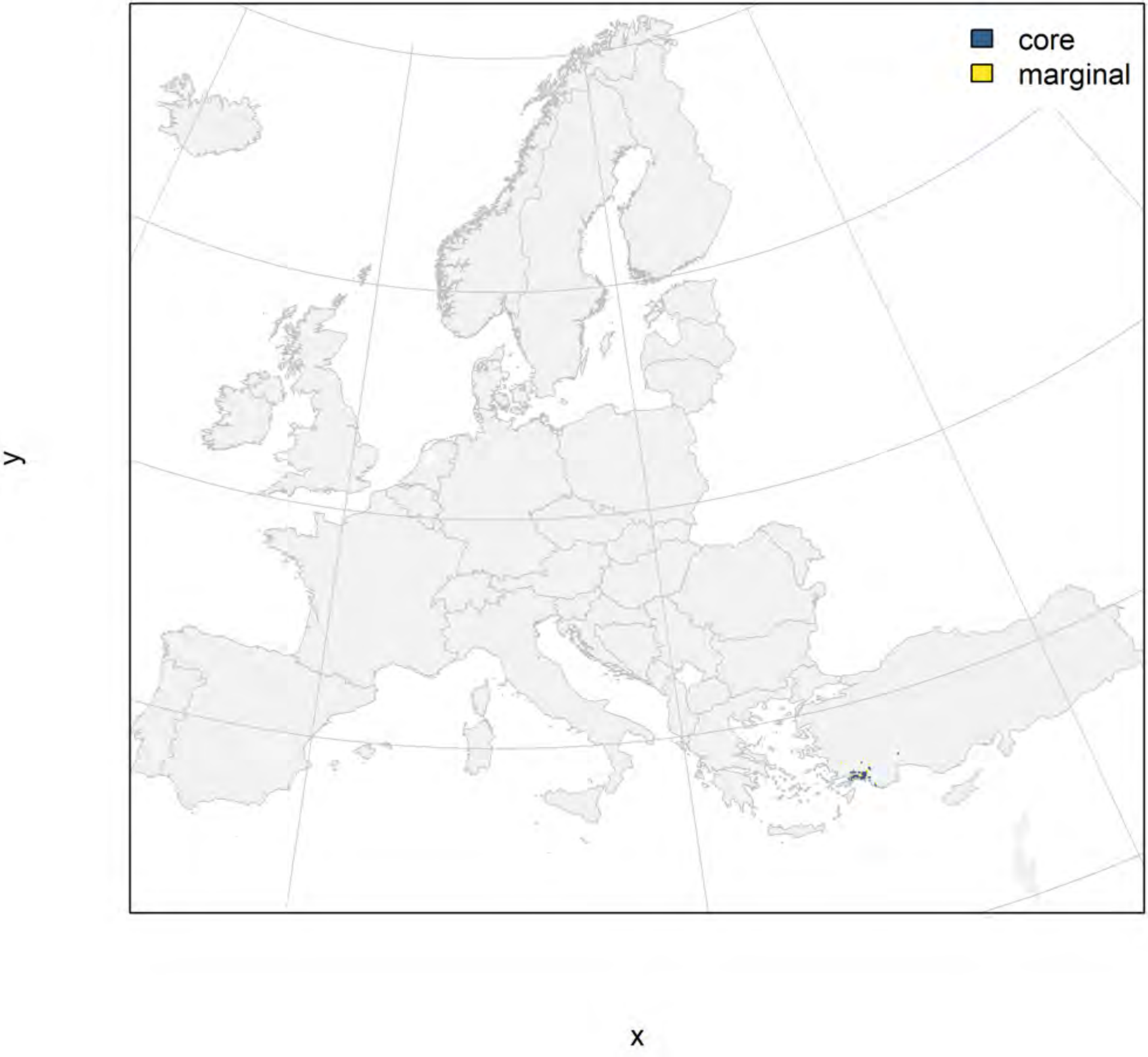

**Ostrya carpinifolia_filt NMI**

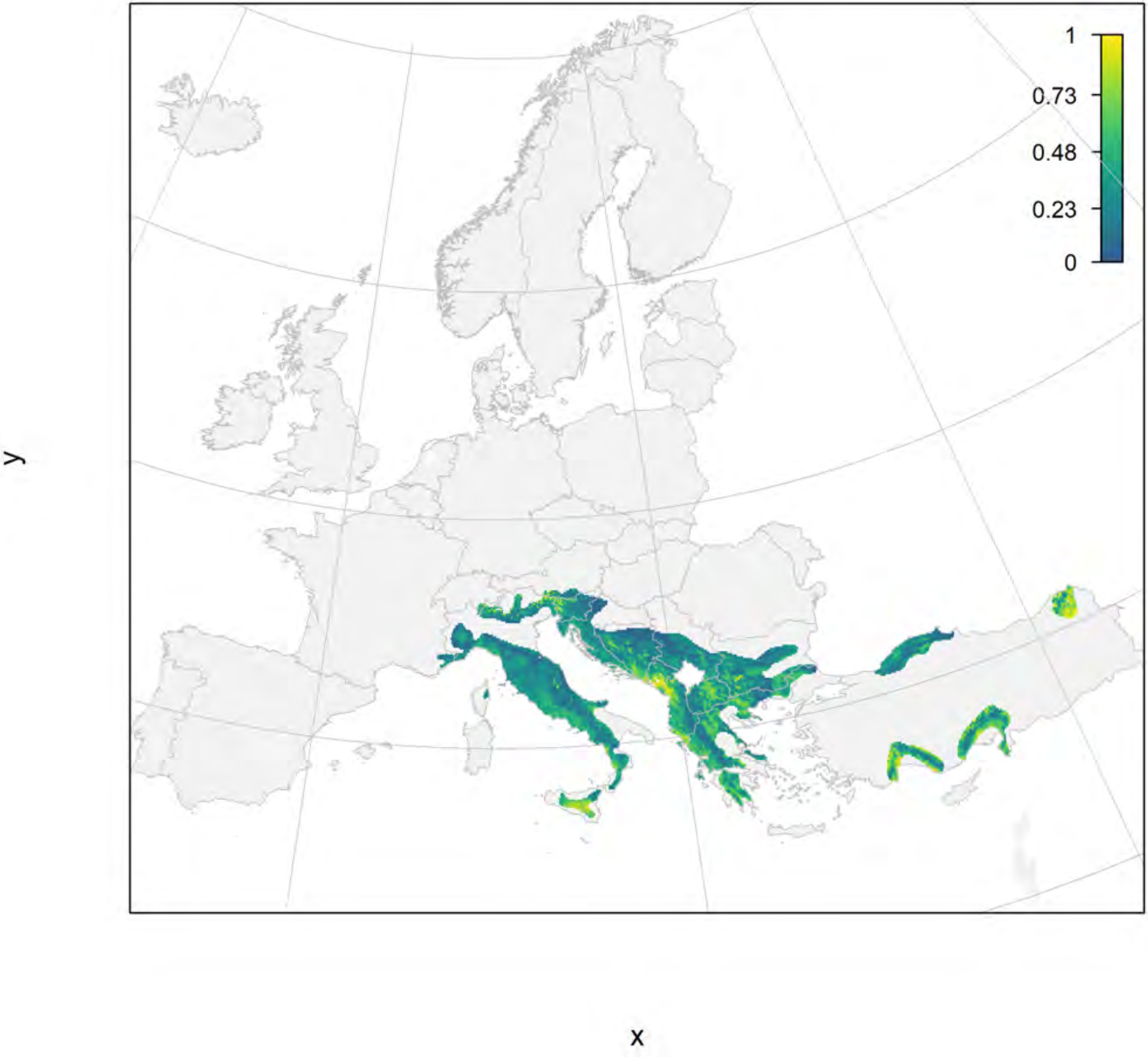

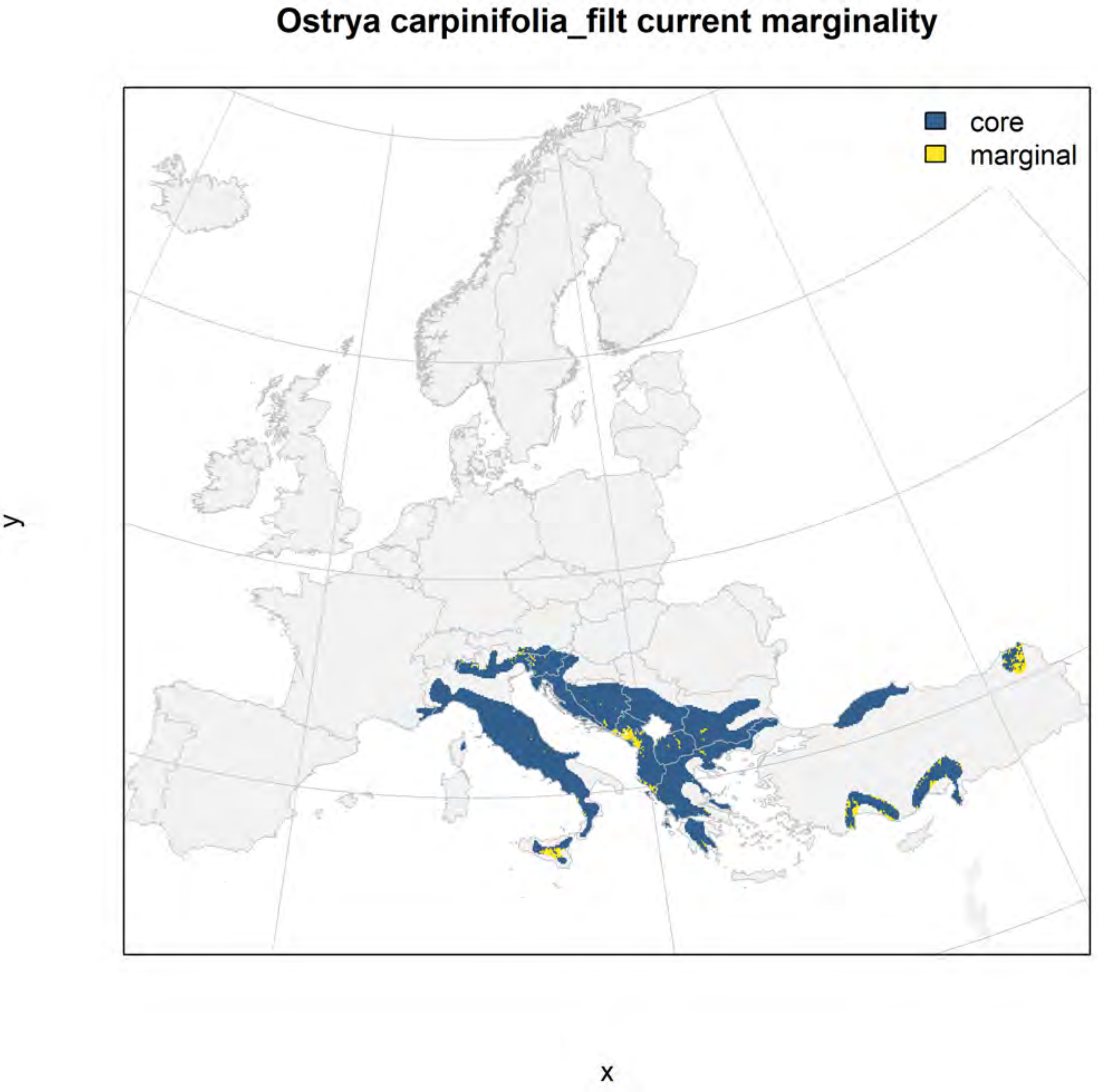
Ostrya carpinifolia_filt current marginality
core
marginal
y
x

#### Ostrya carpinifolia_filt future marginality

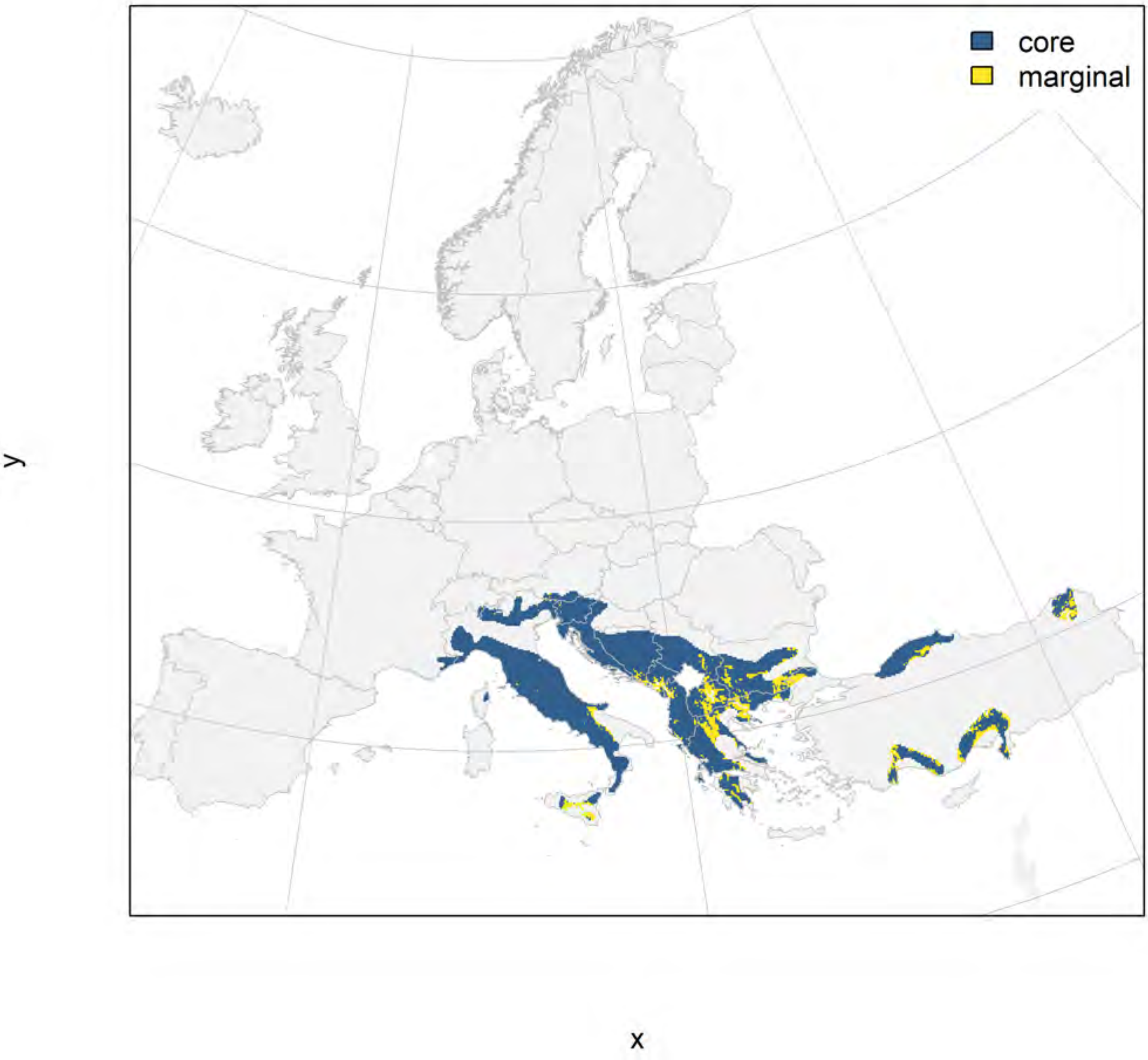

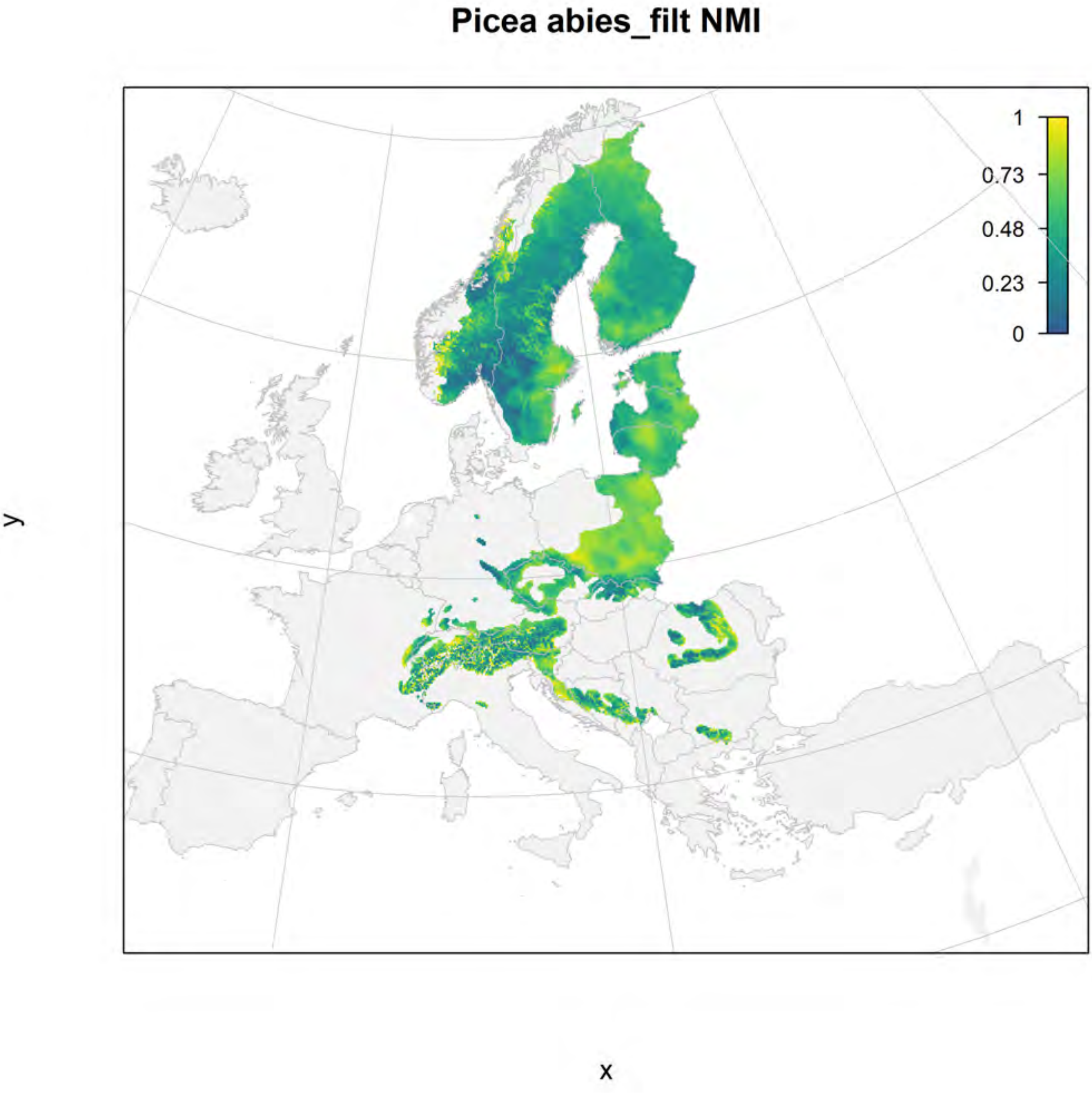
Picea abies_filt NMI
1
0.73
0.48
0.23
0
y
x

#### Picea abies_filt current marginality

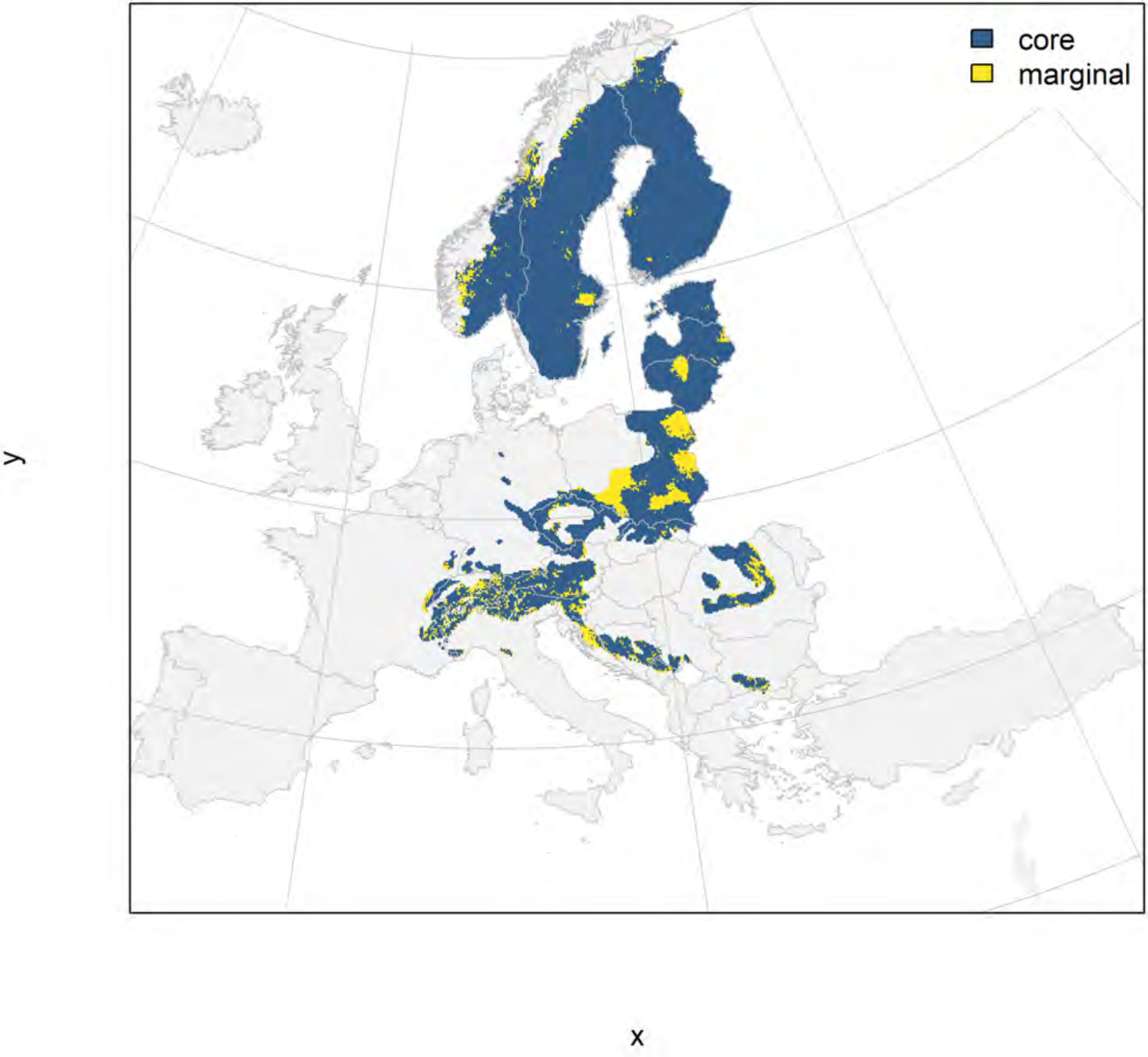

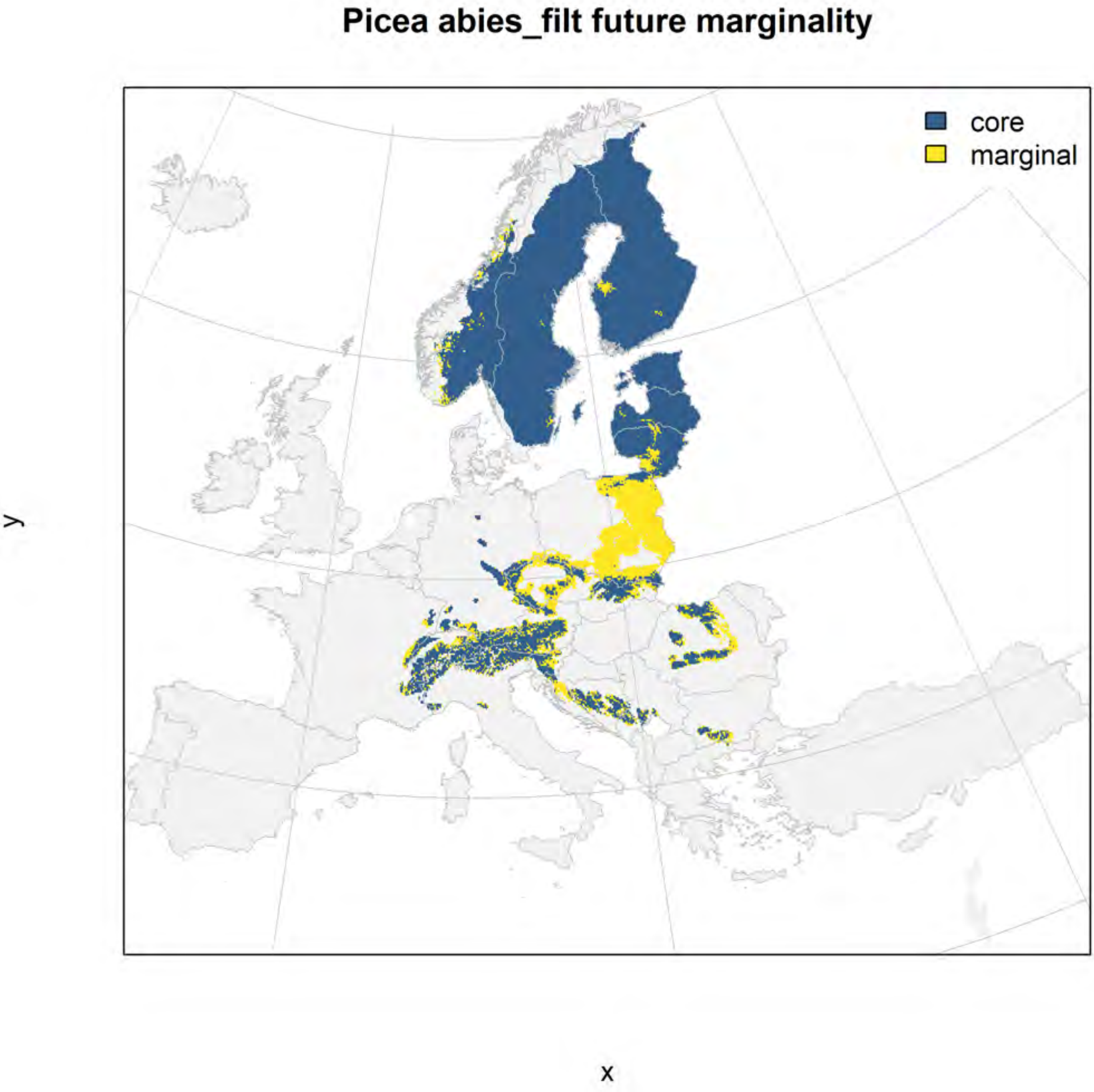
Picea abies_filt future marginality
core
marginal
y
x

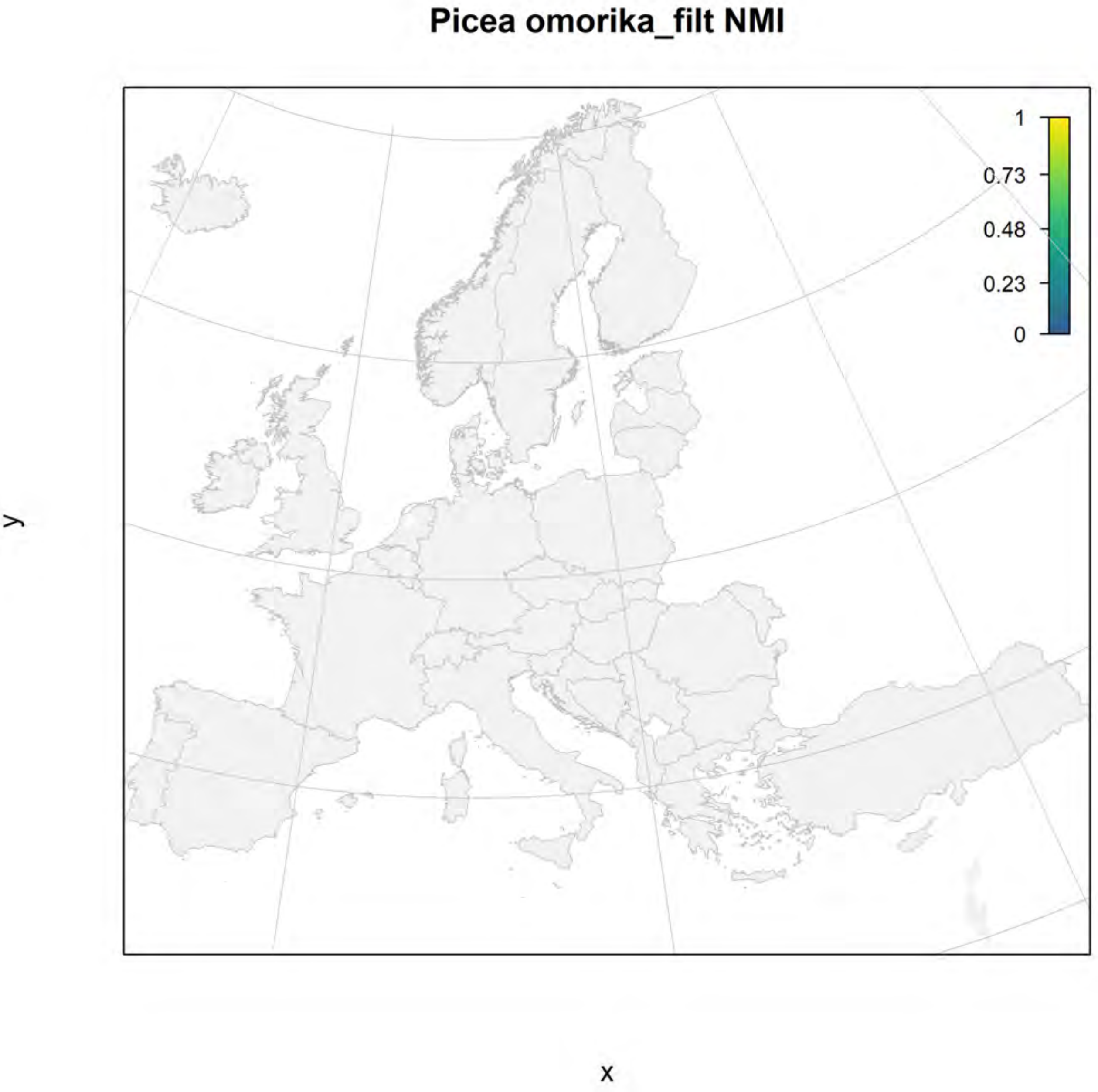
Picea omorika_filt NMI
1
0.73
0.48
0.23
0
y
x

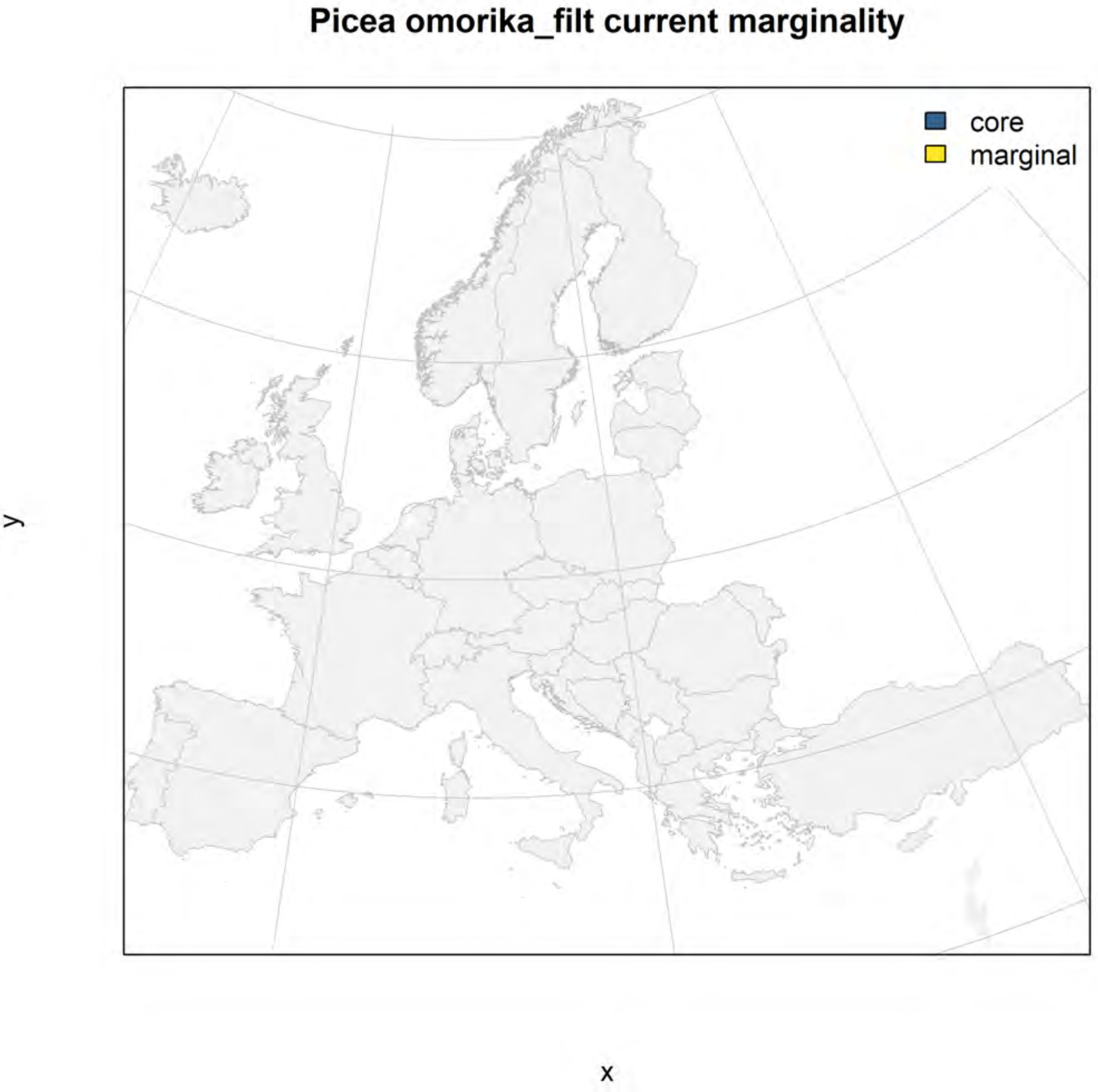
Picea omorika_filt current marginality
core
marginal
y
x

### Picea omorika_filt future marginality

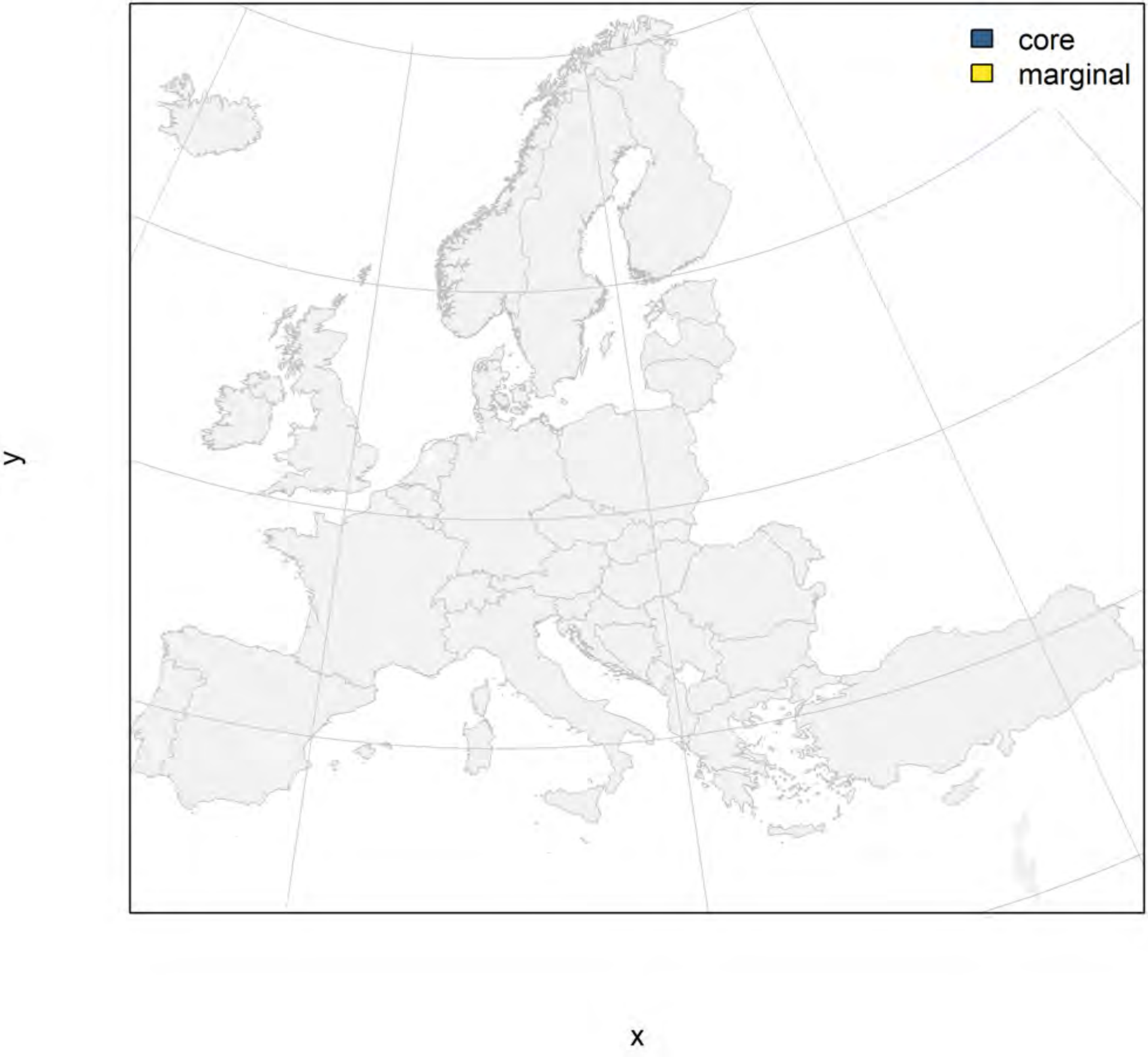

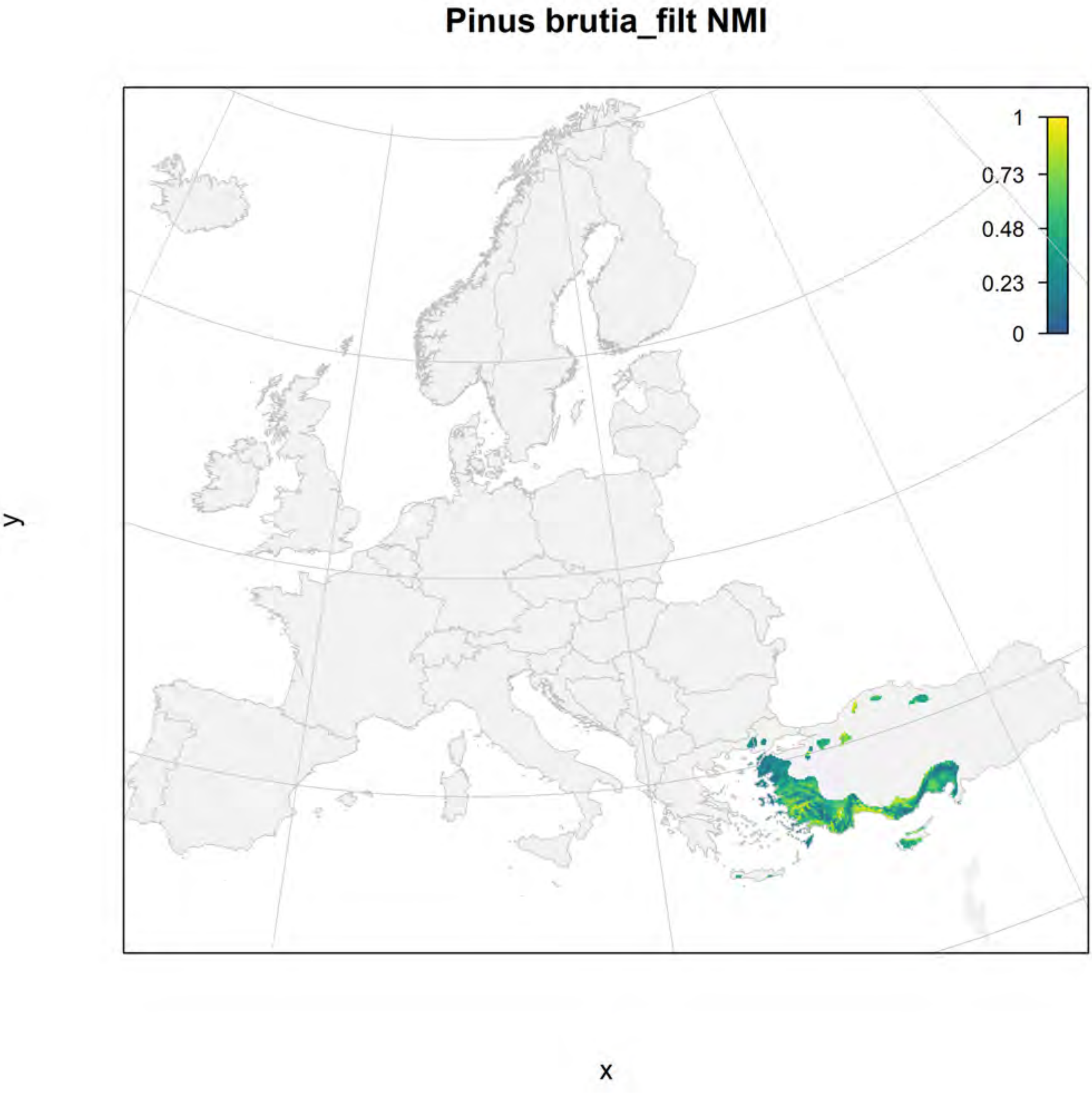
Pinus brutia_filt NMI
1
0.73
0.48
0.23
0
y
x

#### Pinus brutia_filt current marginality

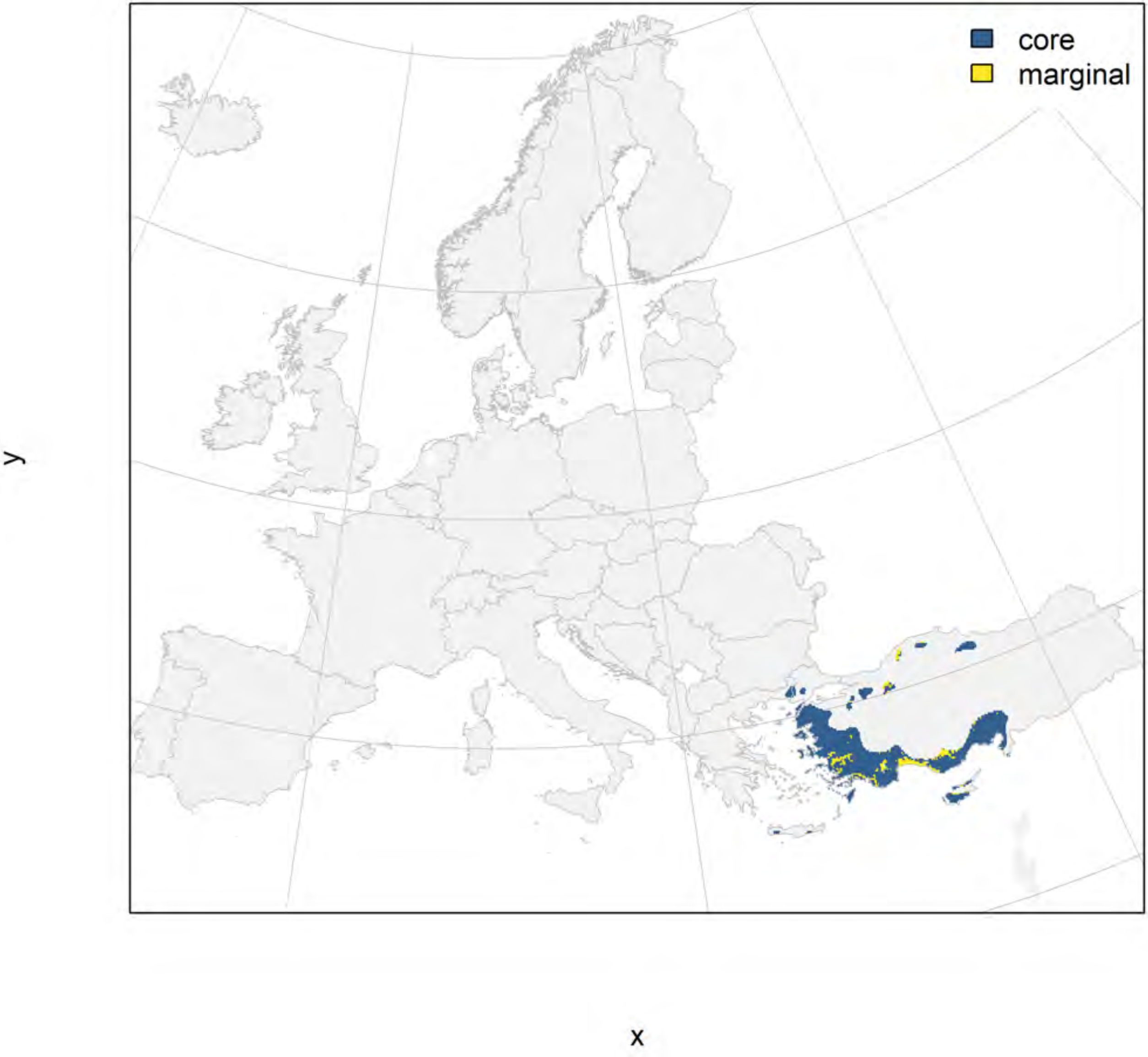

### Pinus brutia_filt future marginality

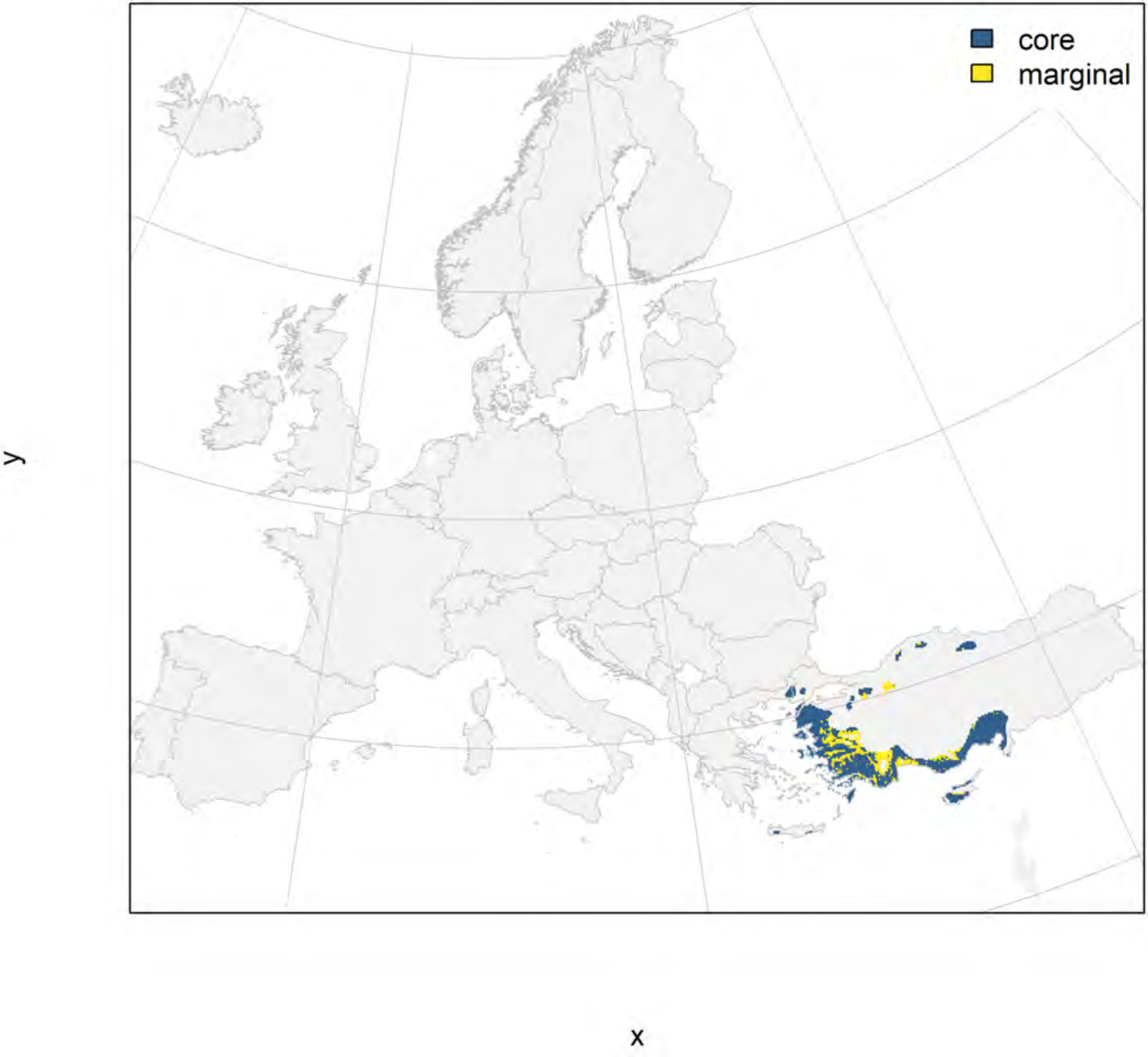

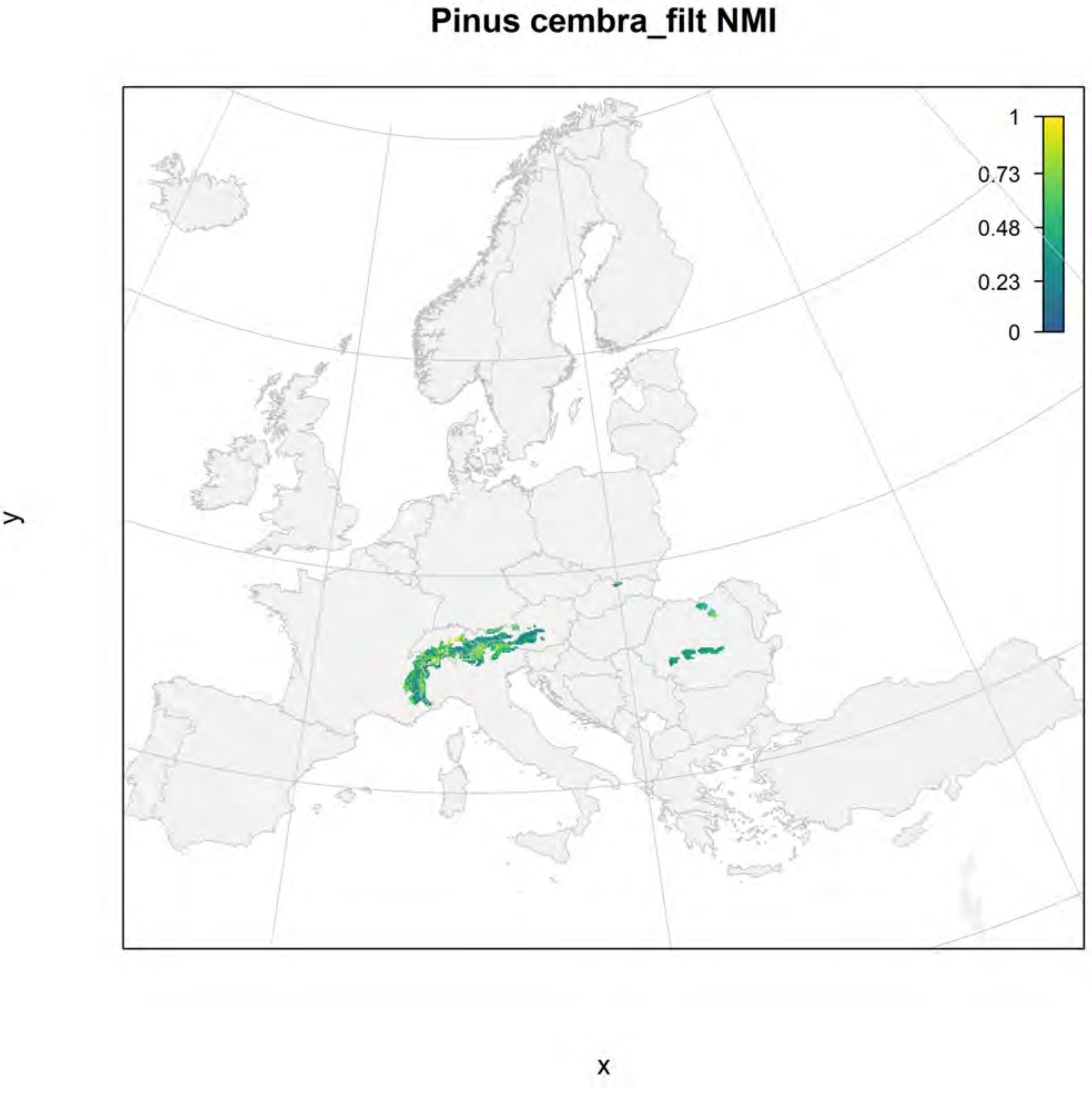
Pinus cembra_filt NMI
1
0.73
0.48
0.23
0
y
x

### Pinus cembra_filt current marginality

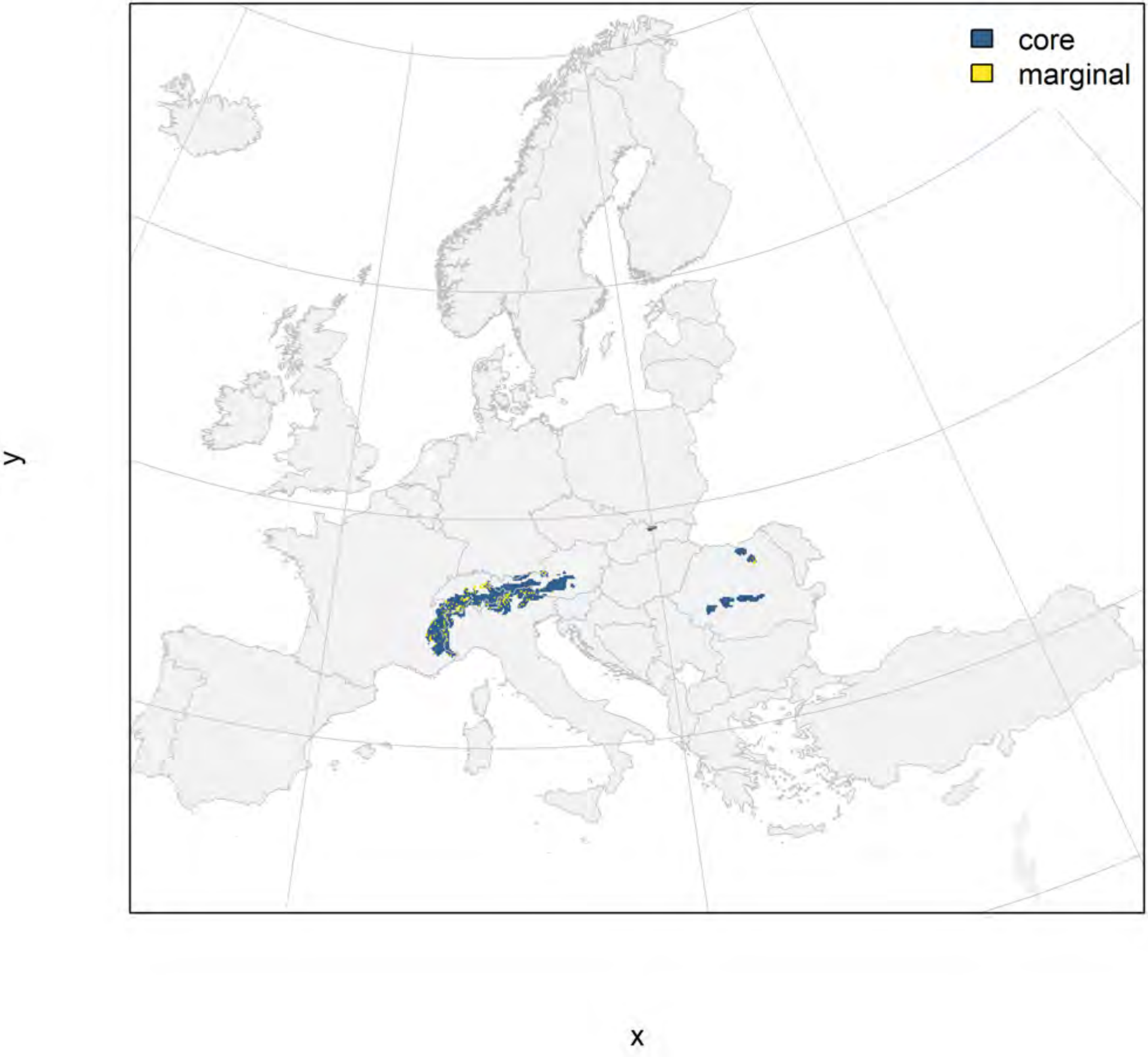

#### Pinus cembra_filt future marginality

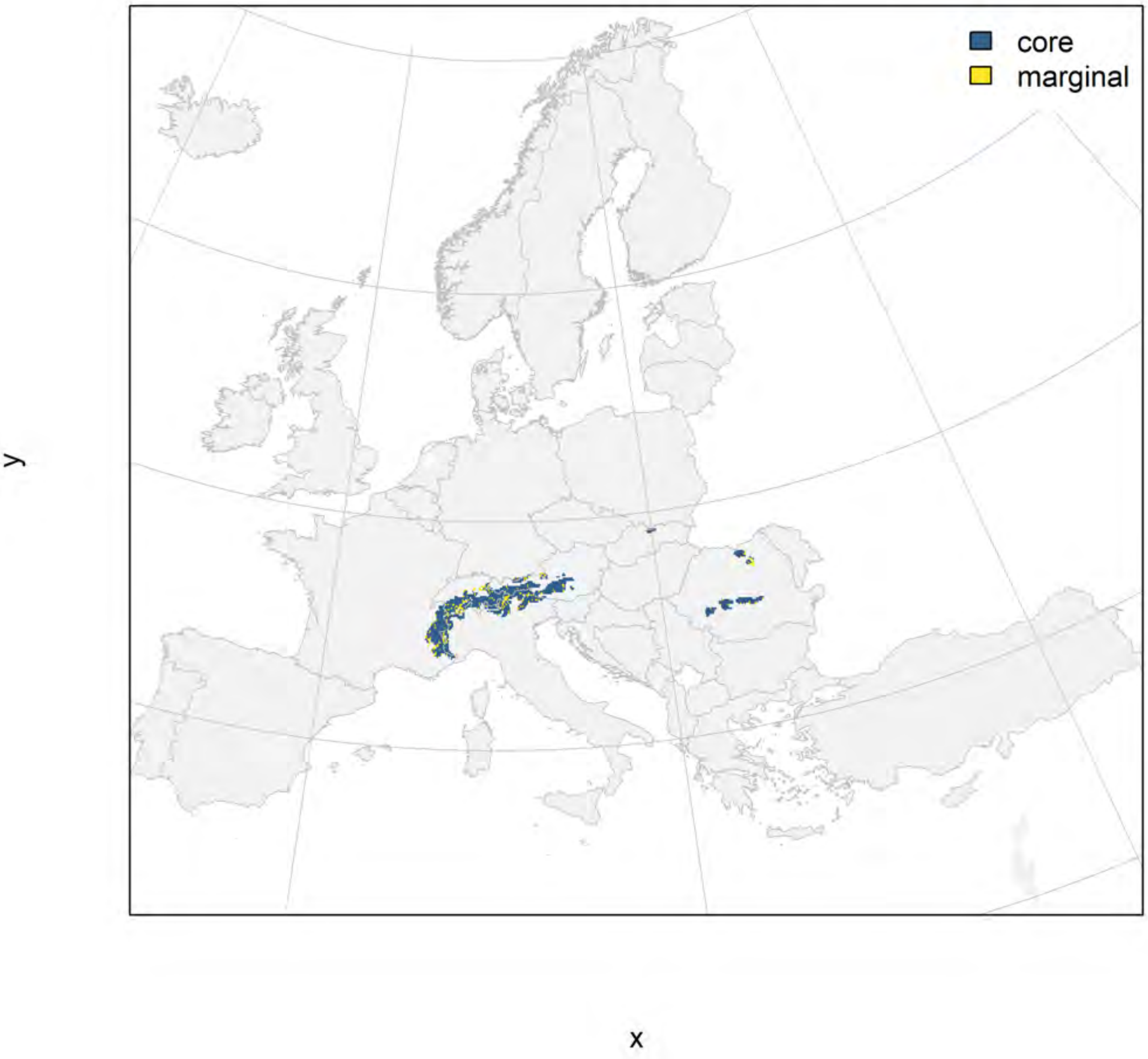

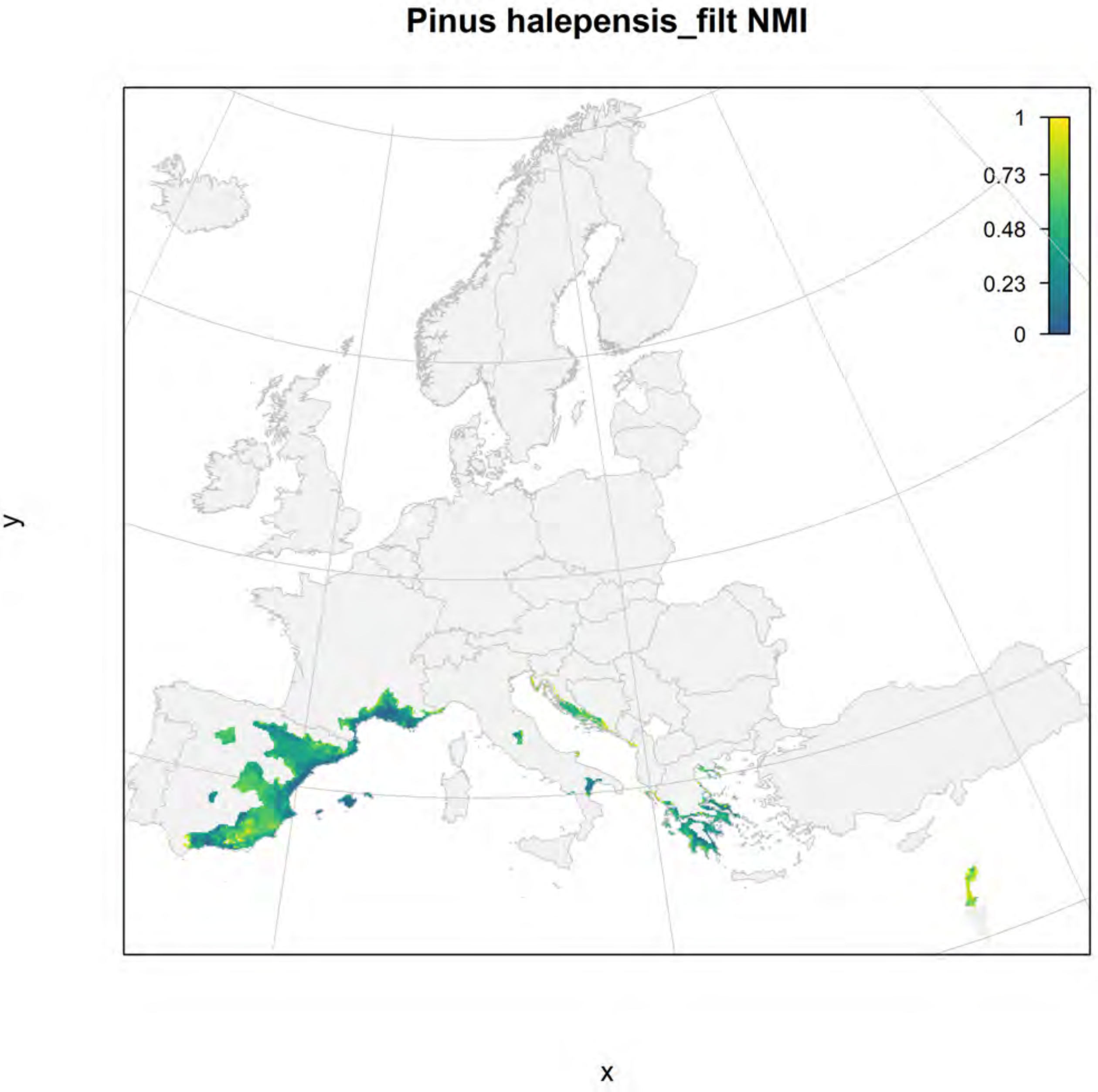

Pinus halepensis_filt NMI
1
0.73
0.48
0.23
0
y
x

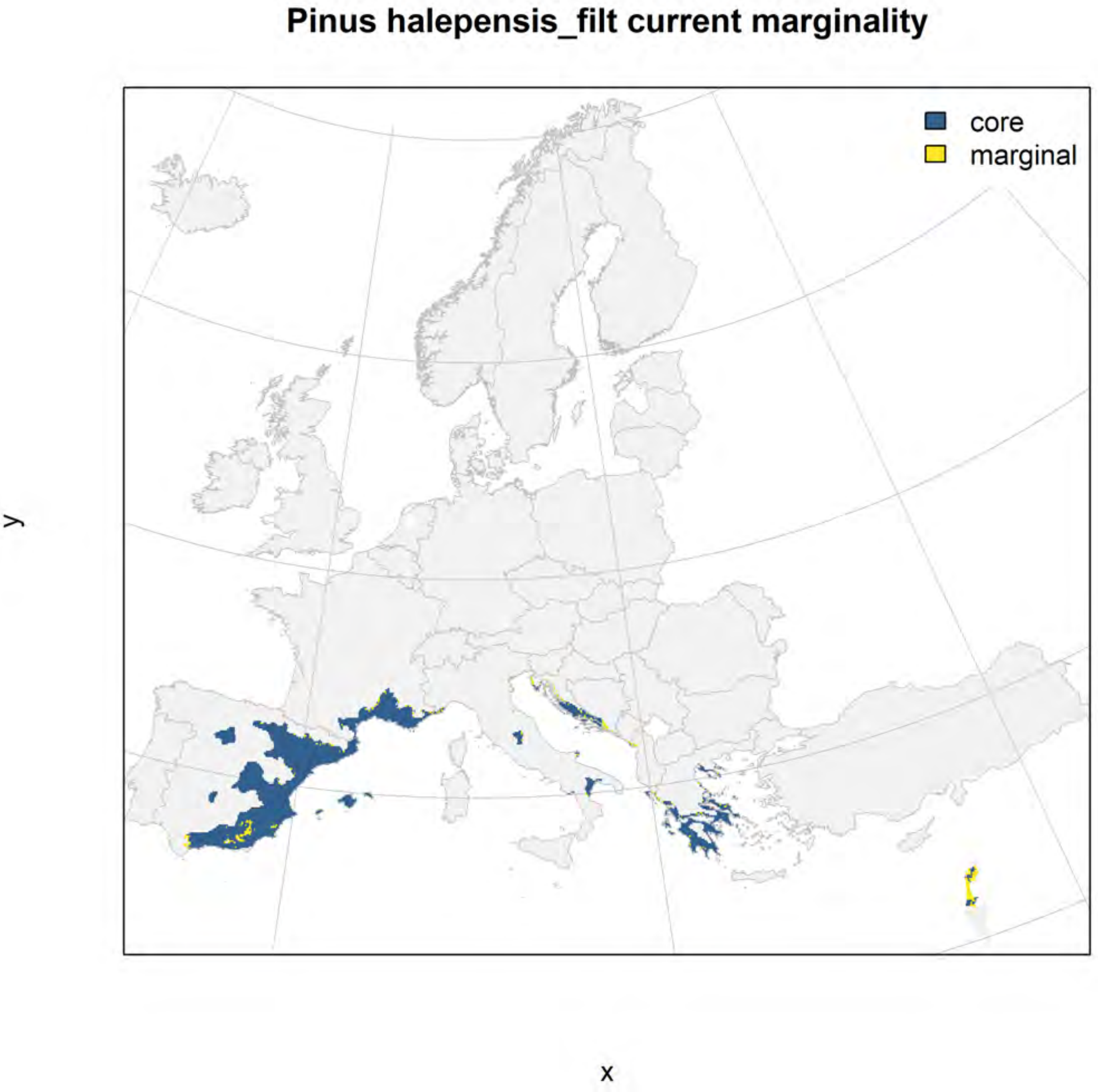
Pinus halepensis_filt current marginality
core
marginal
y
x

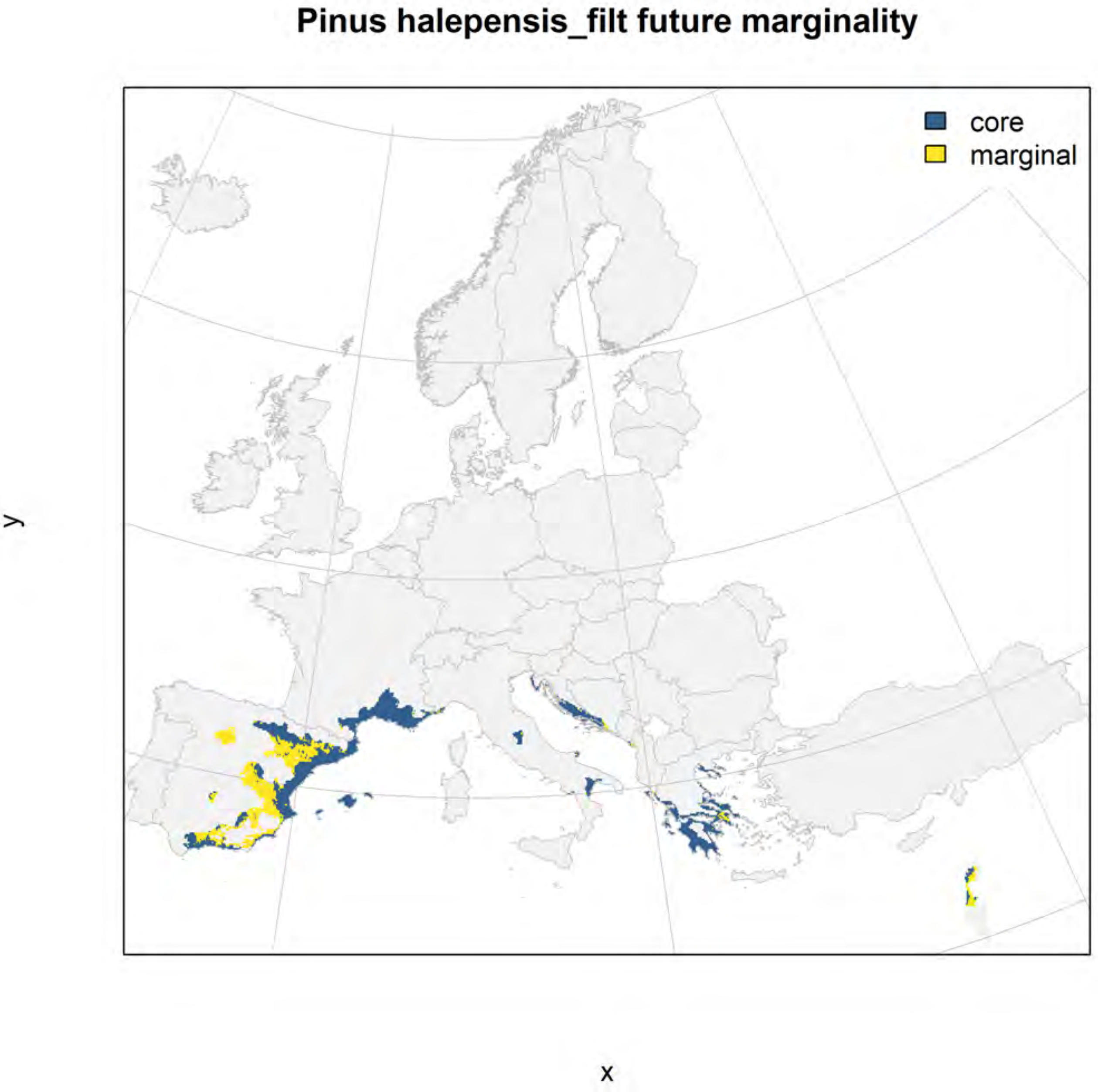
Pinus halepensis_filt future marginality
core
marginal
y
x

#### Pinus heldreichii_filt NMI

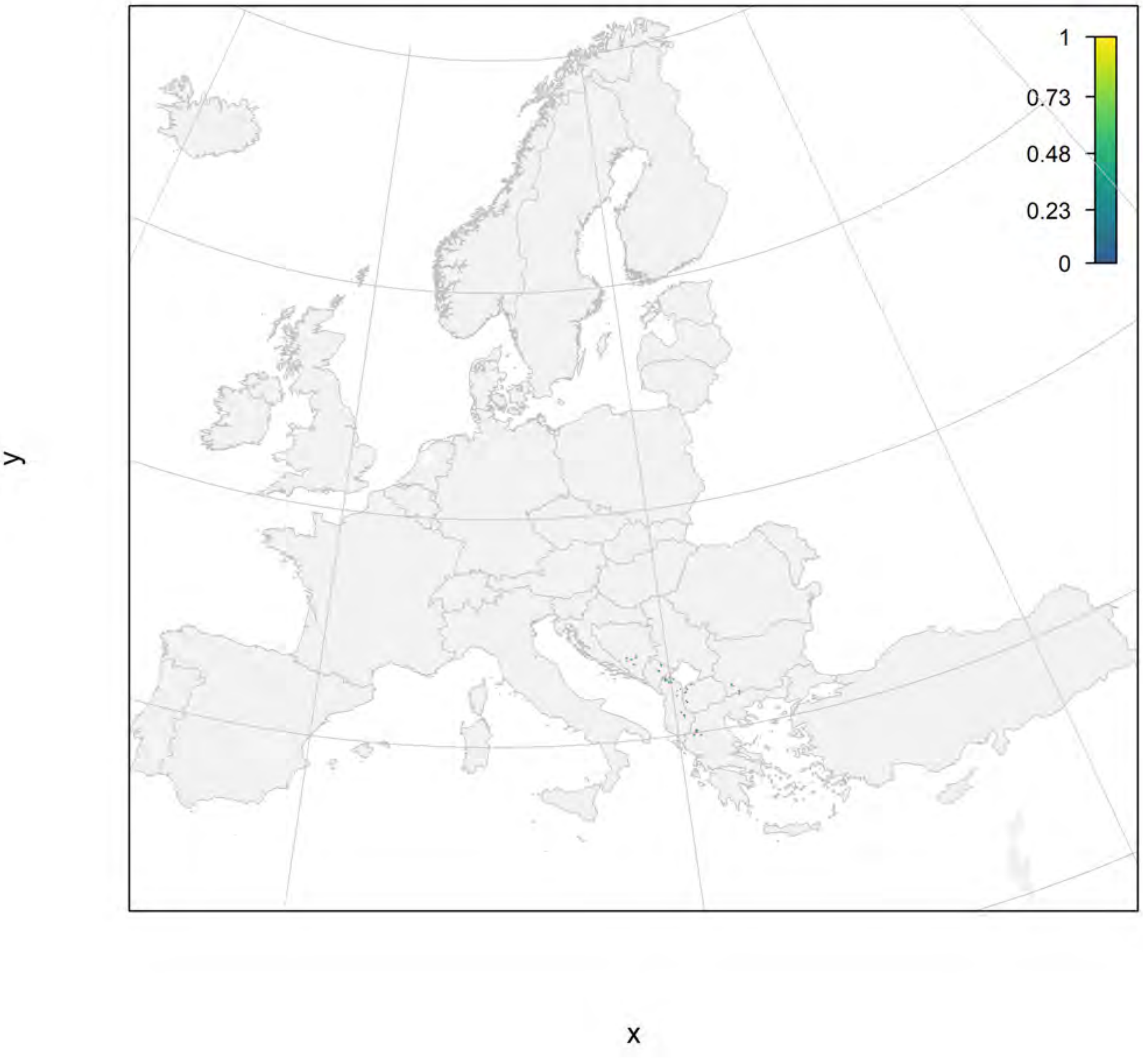

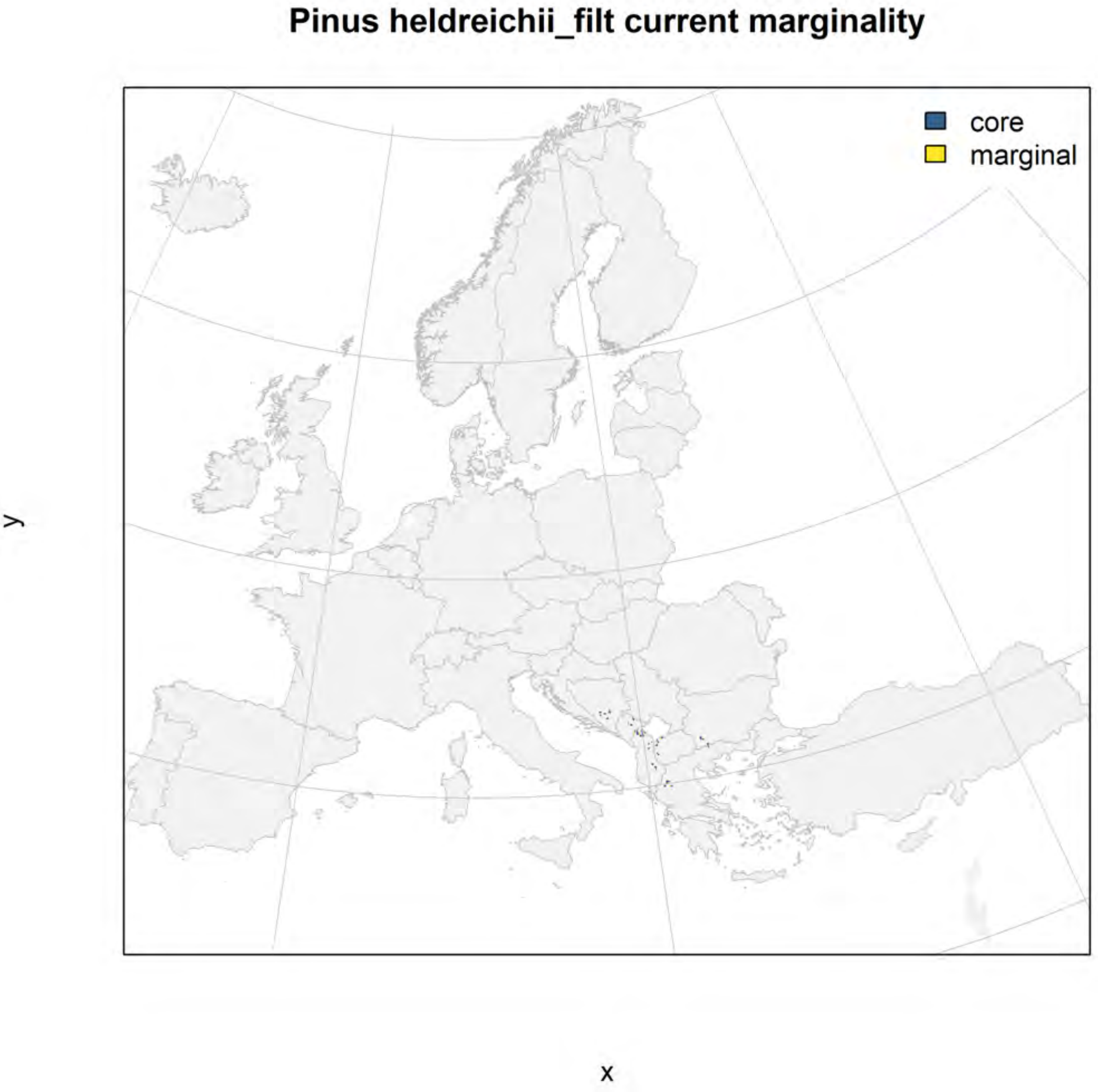
Pinus heldreichii_filt current marginality
core
marginal
y
x

Pinus heldreichii_filt future marginality

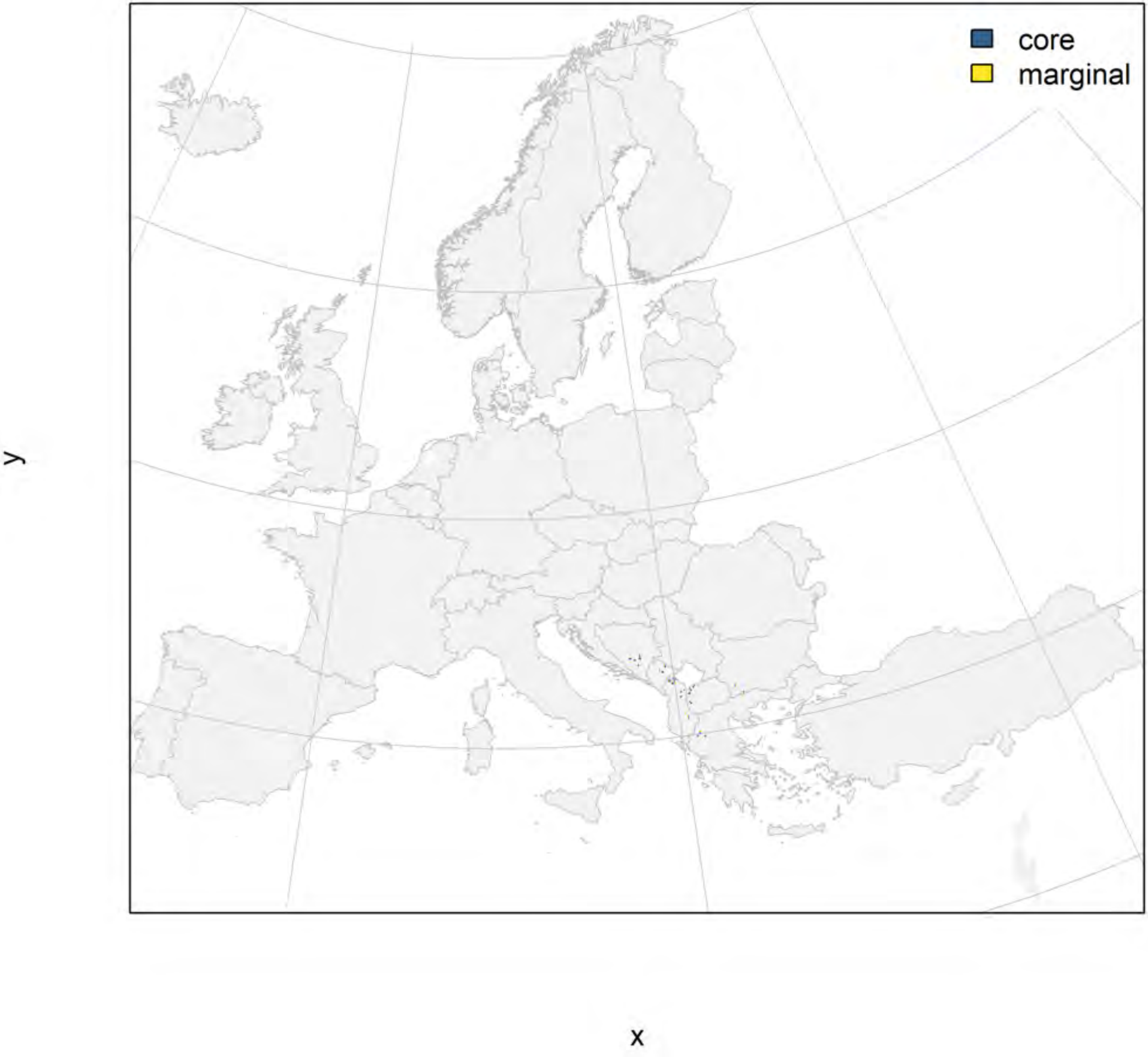

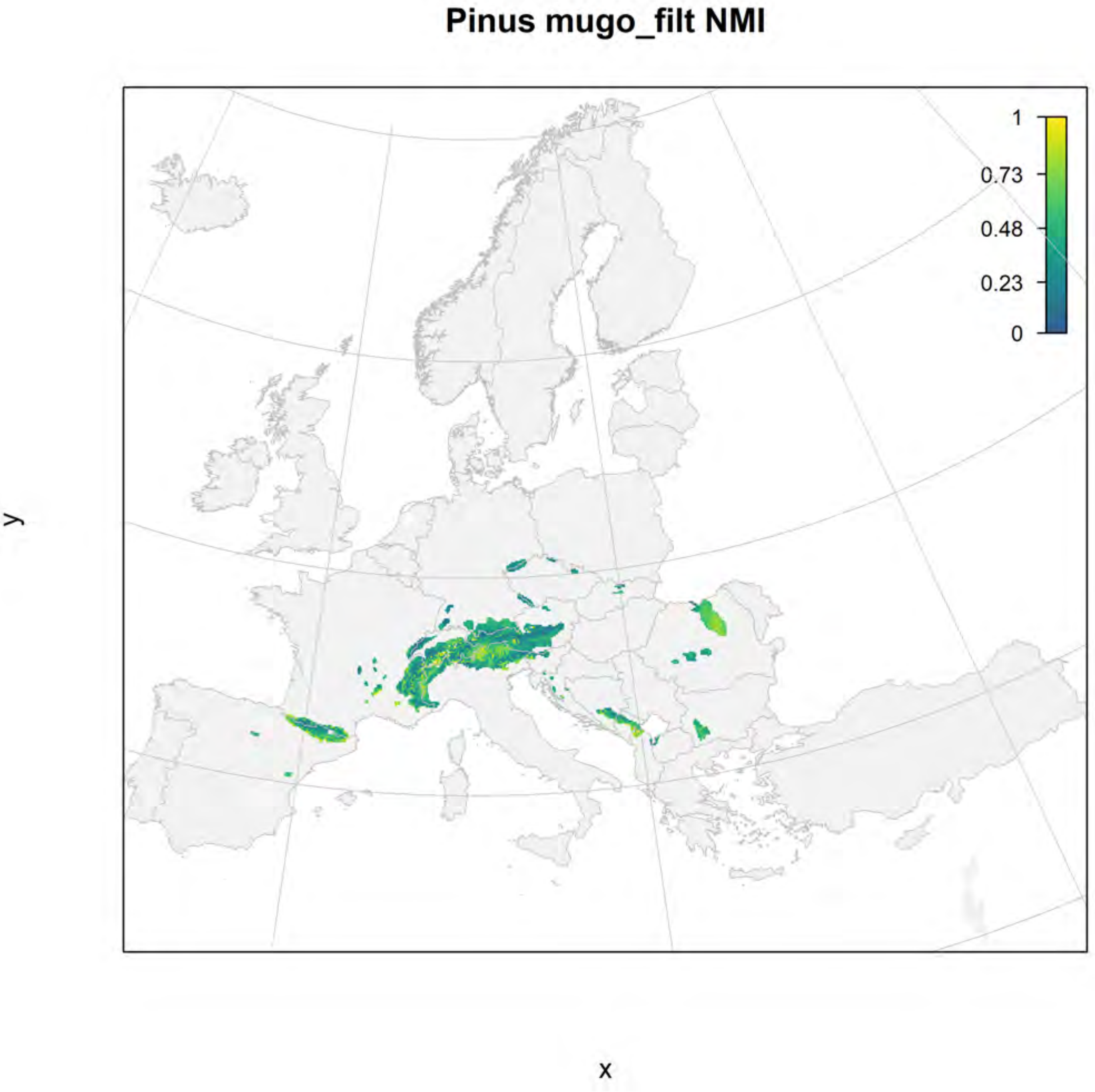
Pinus mugo_filt NMI
1
0.73
0.48
0.23
0
y
x

**Pinus mugo_filt current marginality**

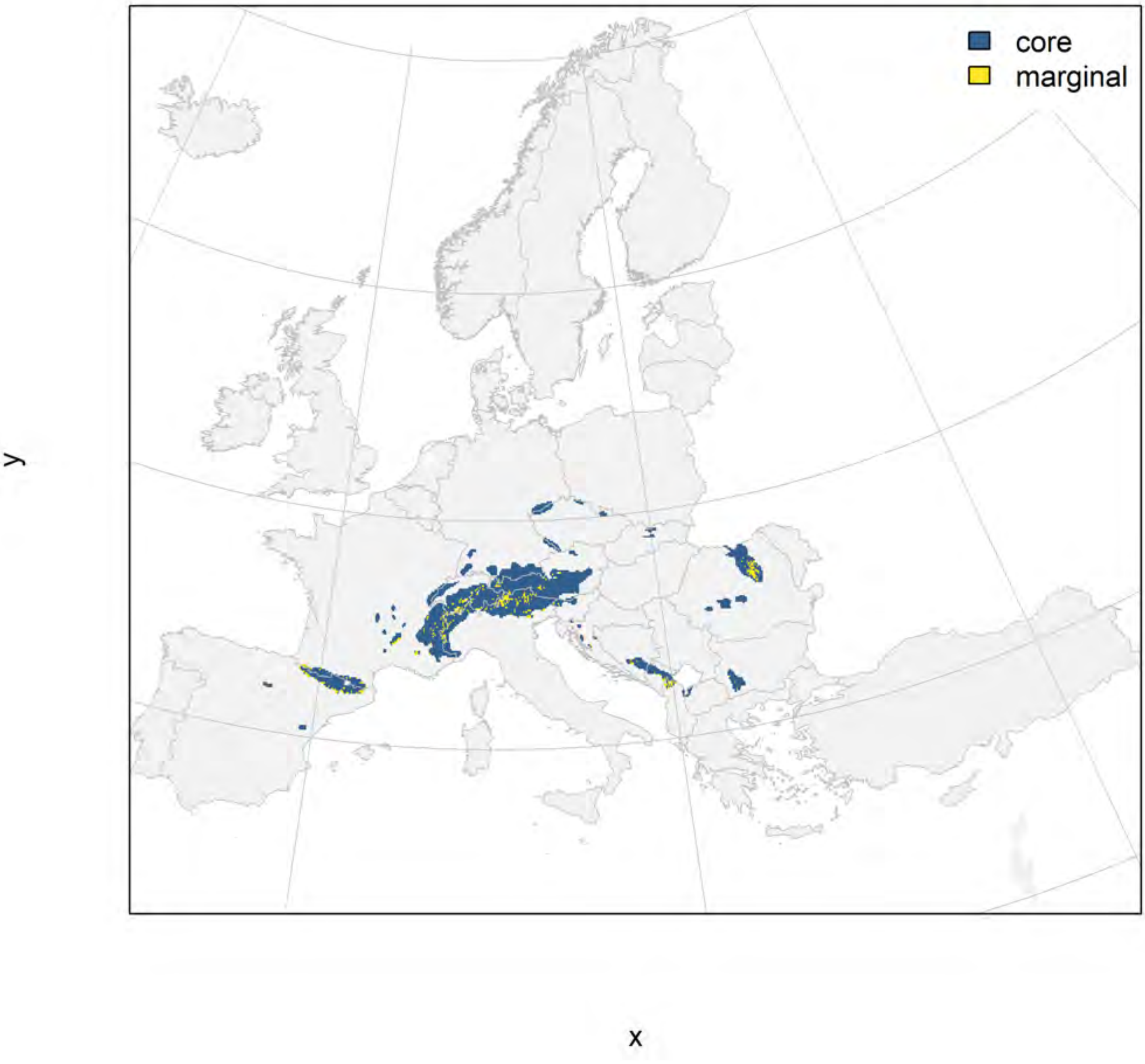

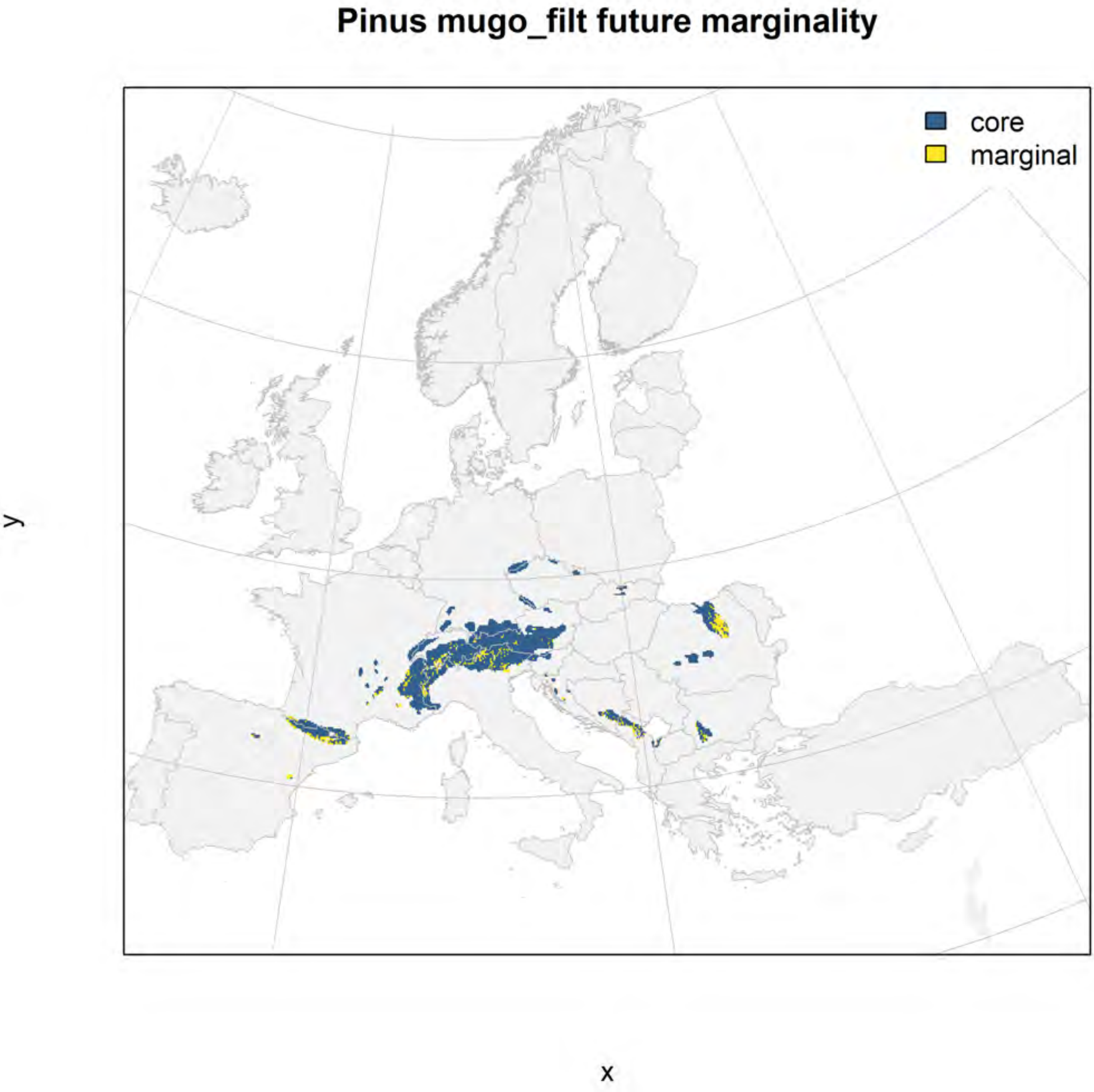
Pinus mugo_filt future marginality
core
marginal
y
x

### Pinus nigra_filt NMI

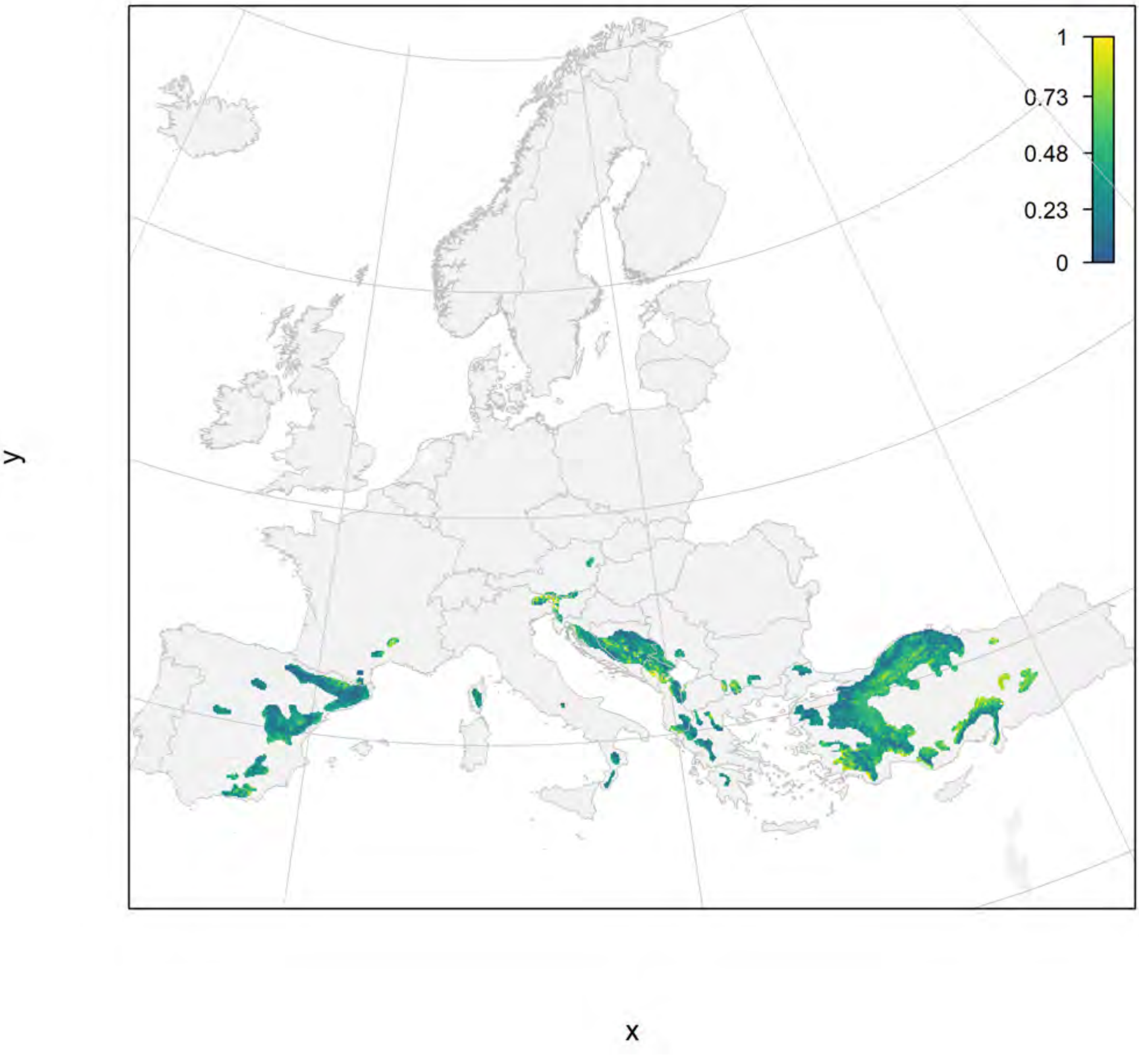

### Pinus nigra_filt current marginality

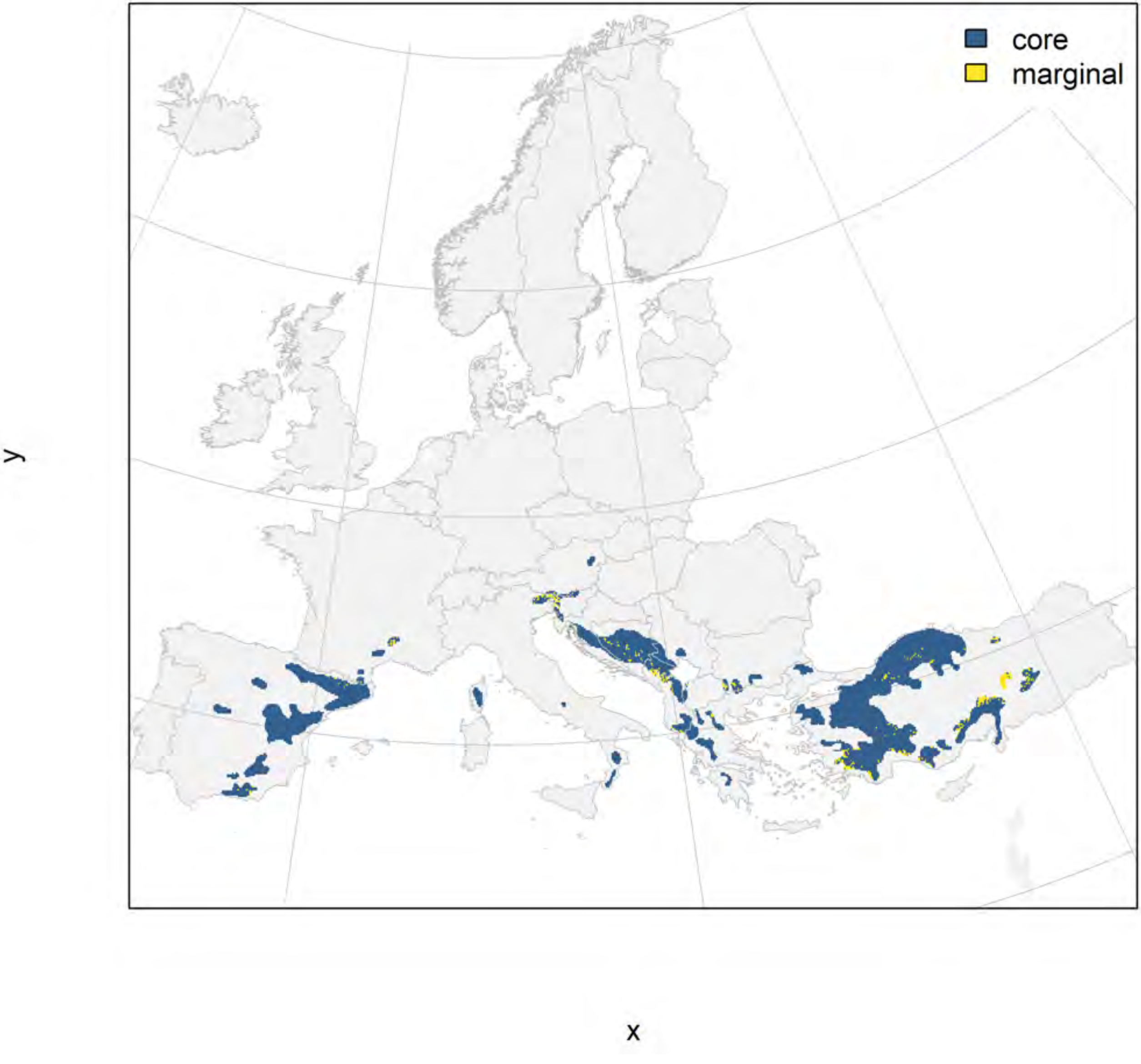

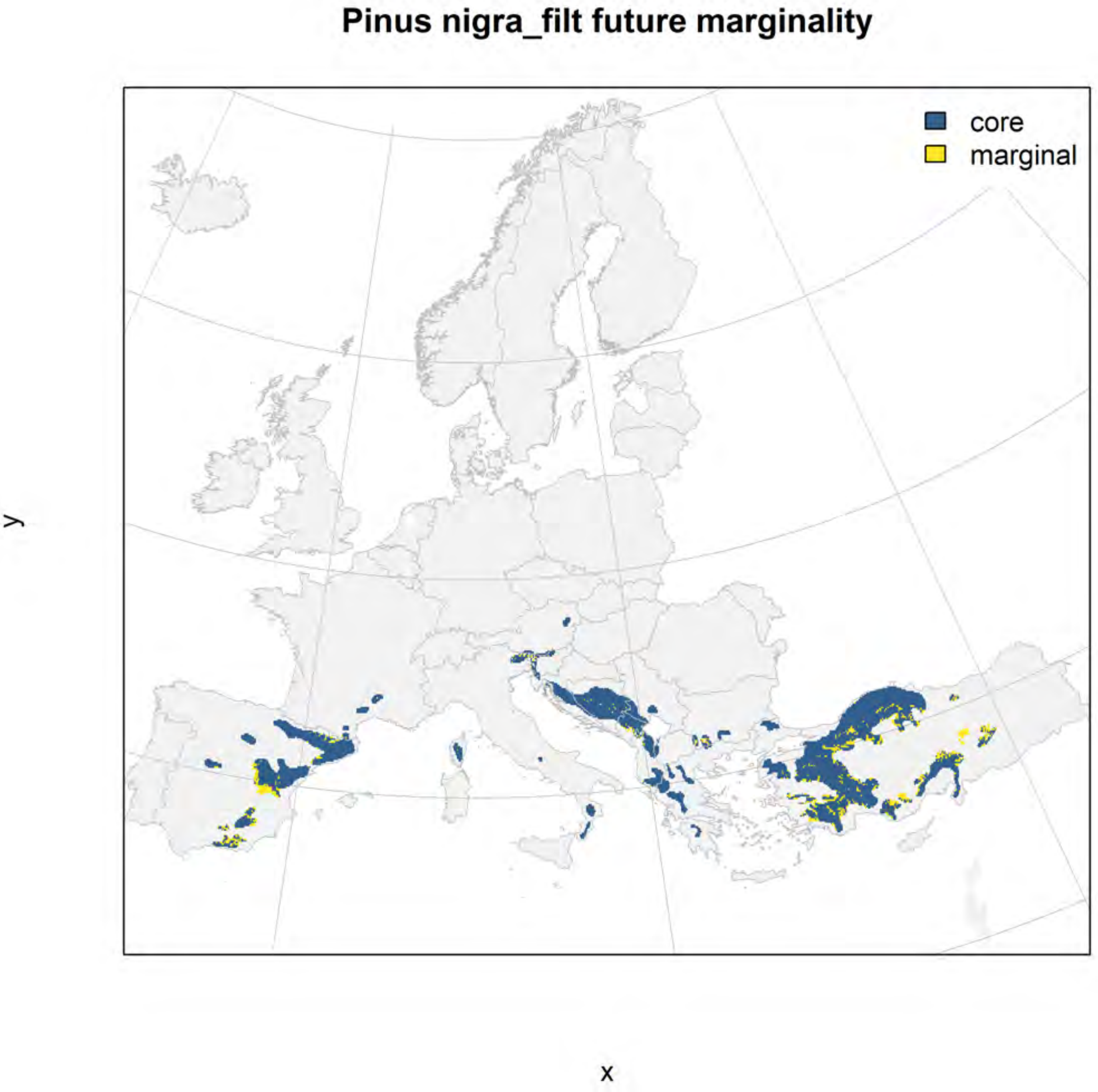
Pinus nigra_filt future marginality
core
marginal
y
x

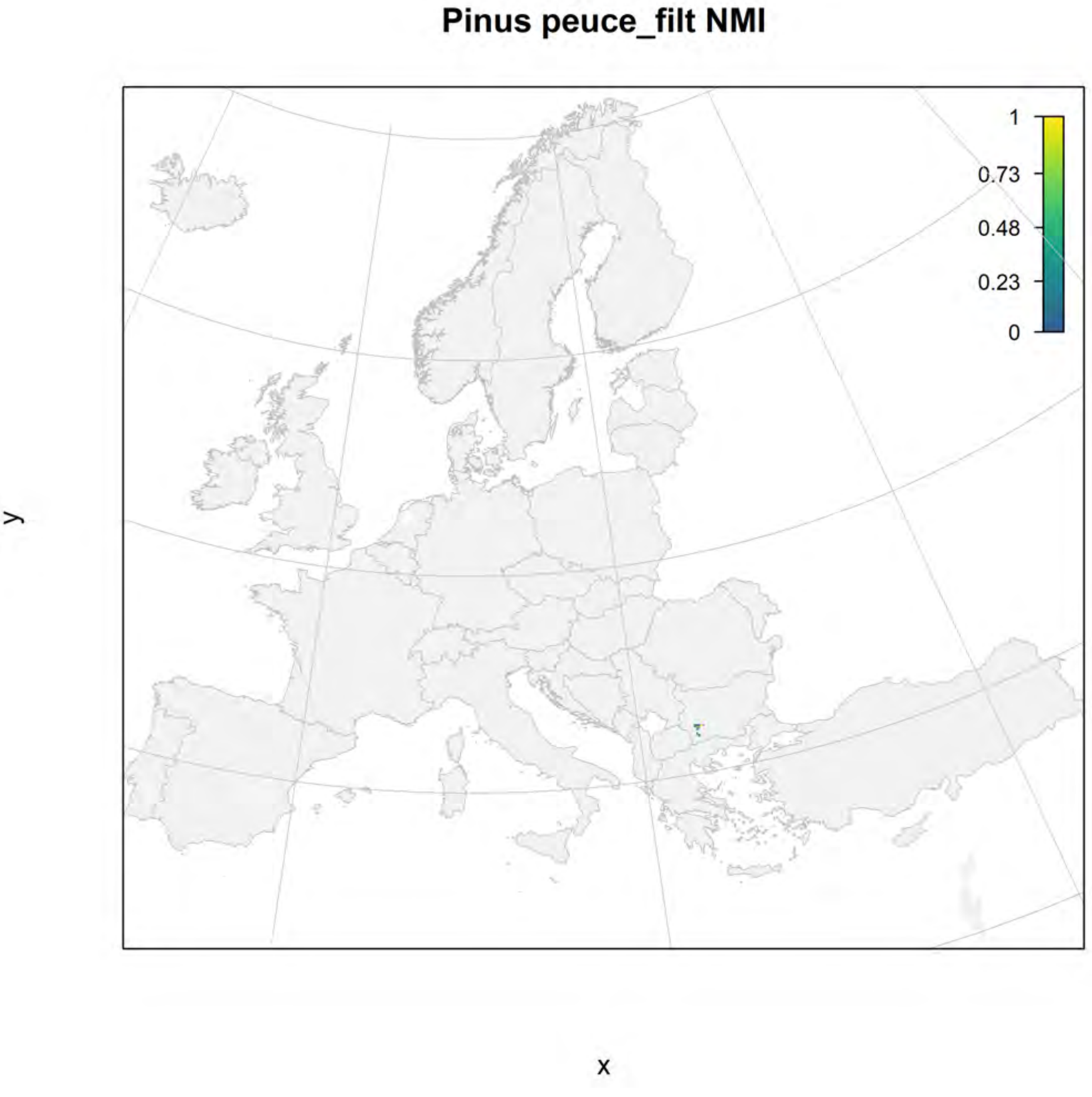
Pinus peuce_filt NMI
1
0.73
0.48
0.23
0
y
x

#### Pinus peuce_filt current marginality

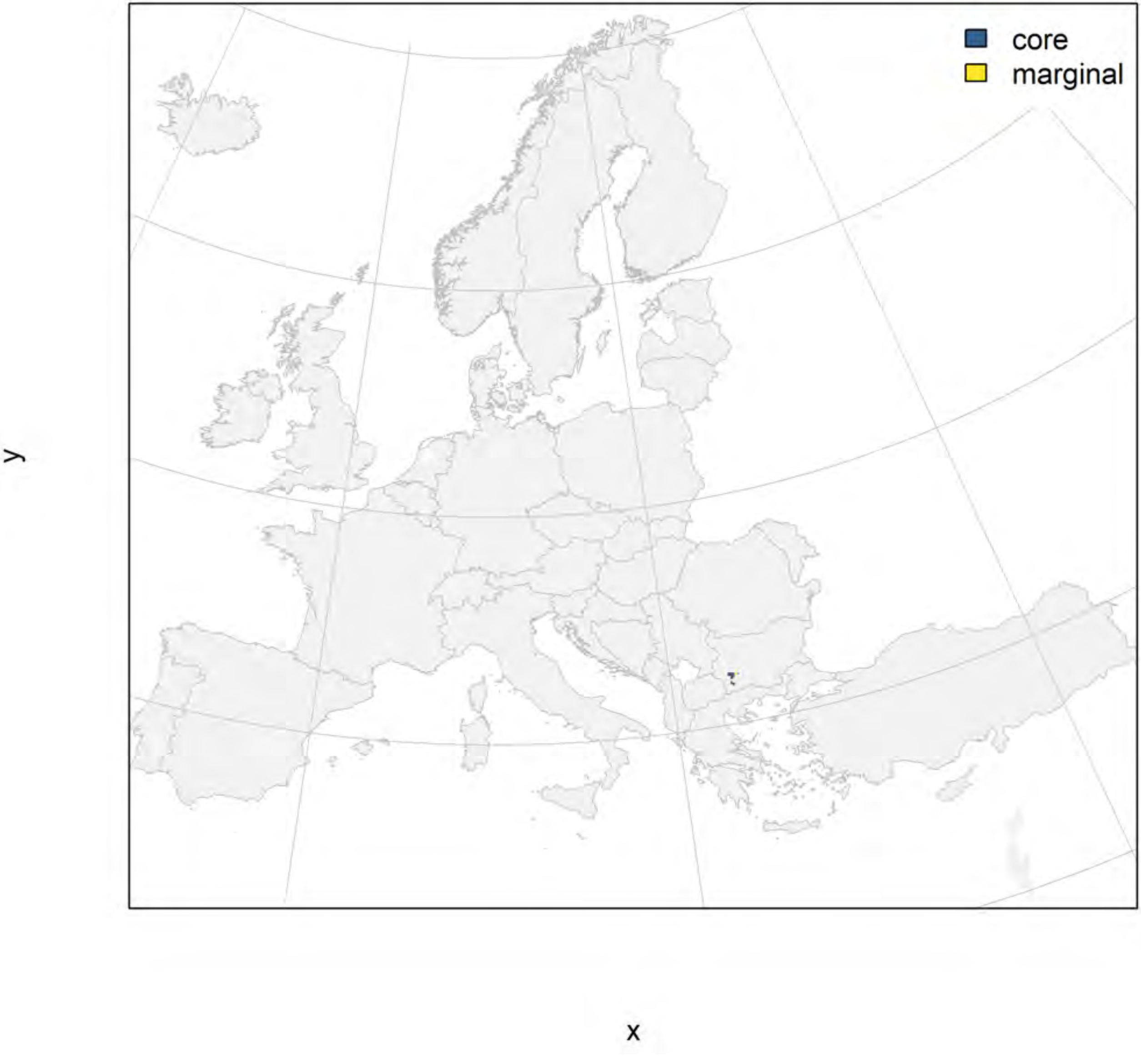

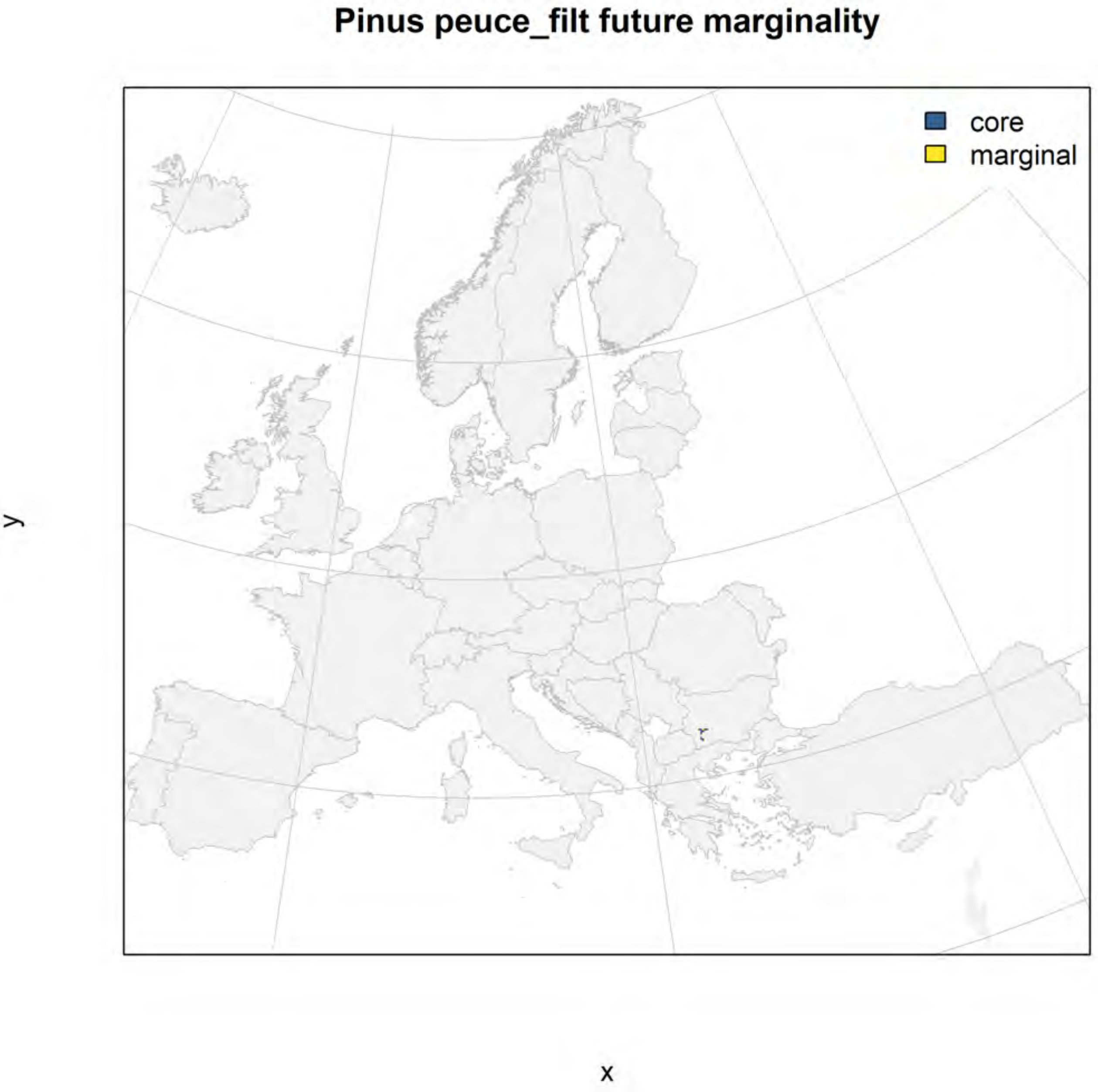
Pinus peuce_filt future marginality
core
marginal
y
x

### Pinus pinaster_filt NMI

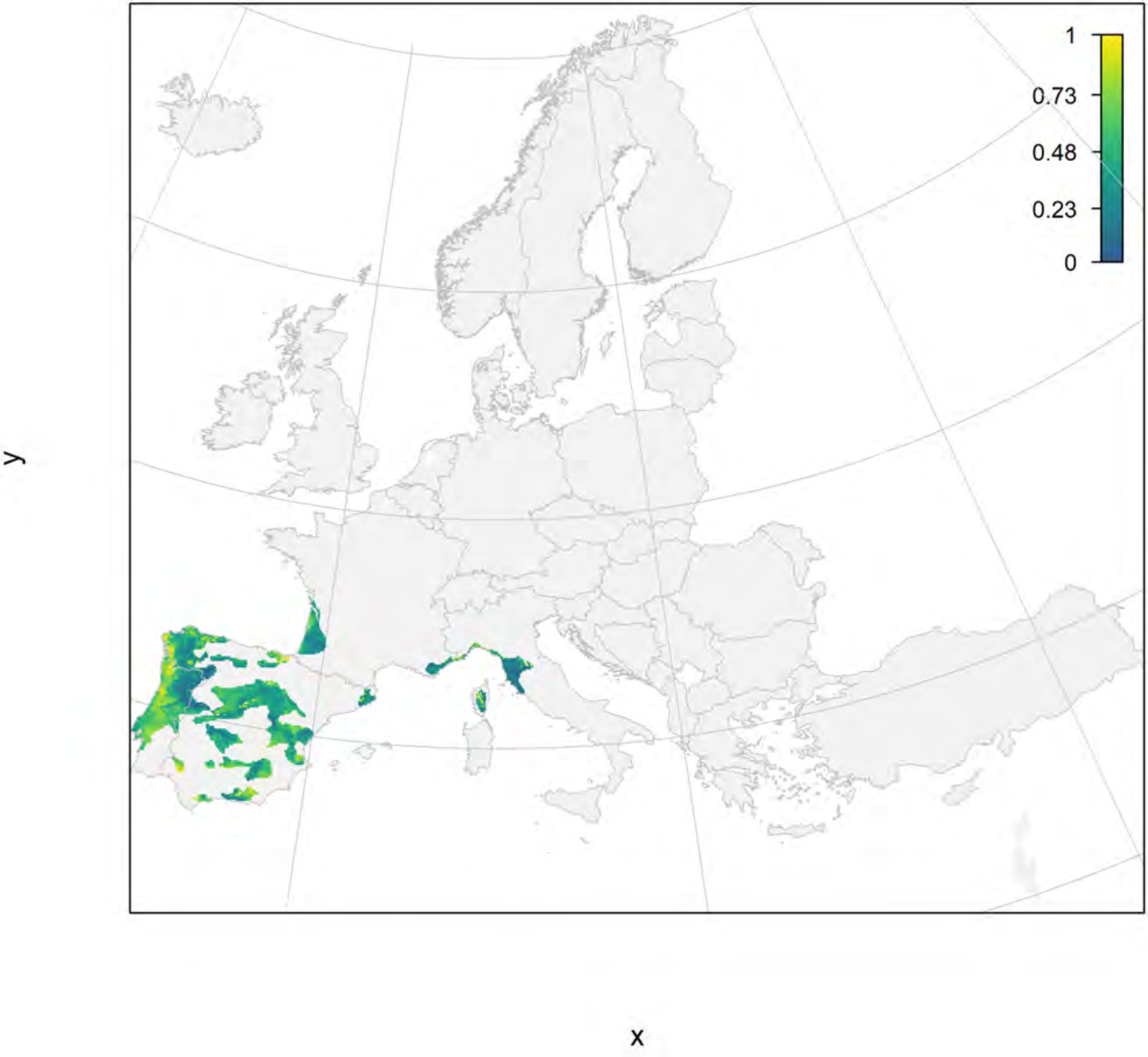

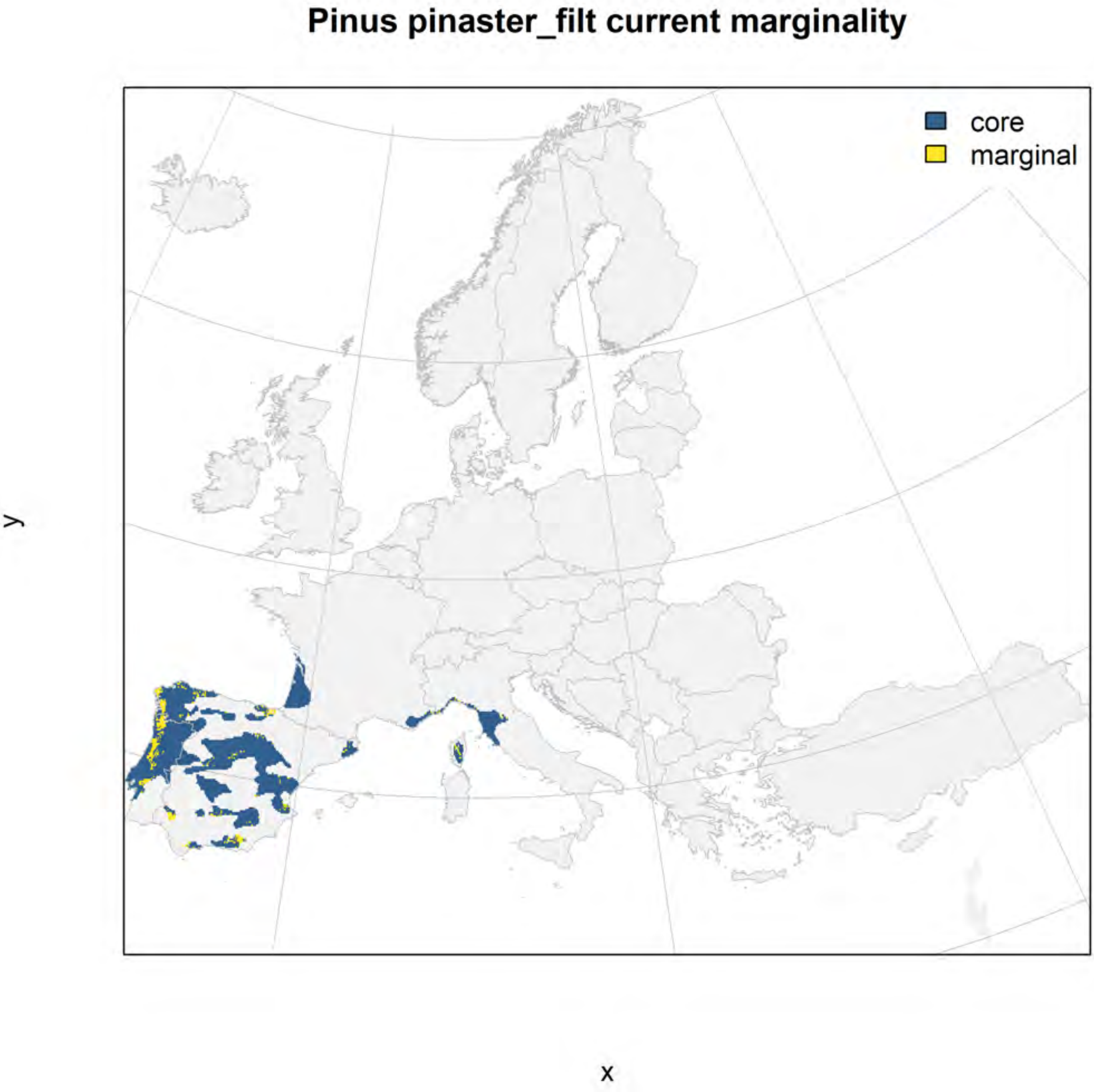
Pinus pinaster_filt current marginality
core
marginal
y
x

### Pinus pinaster_filt future marginality

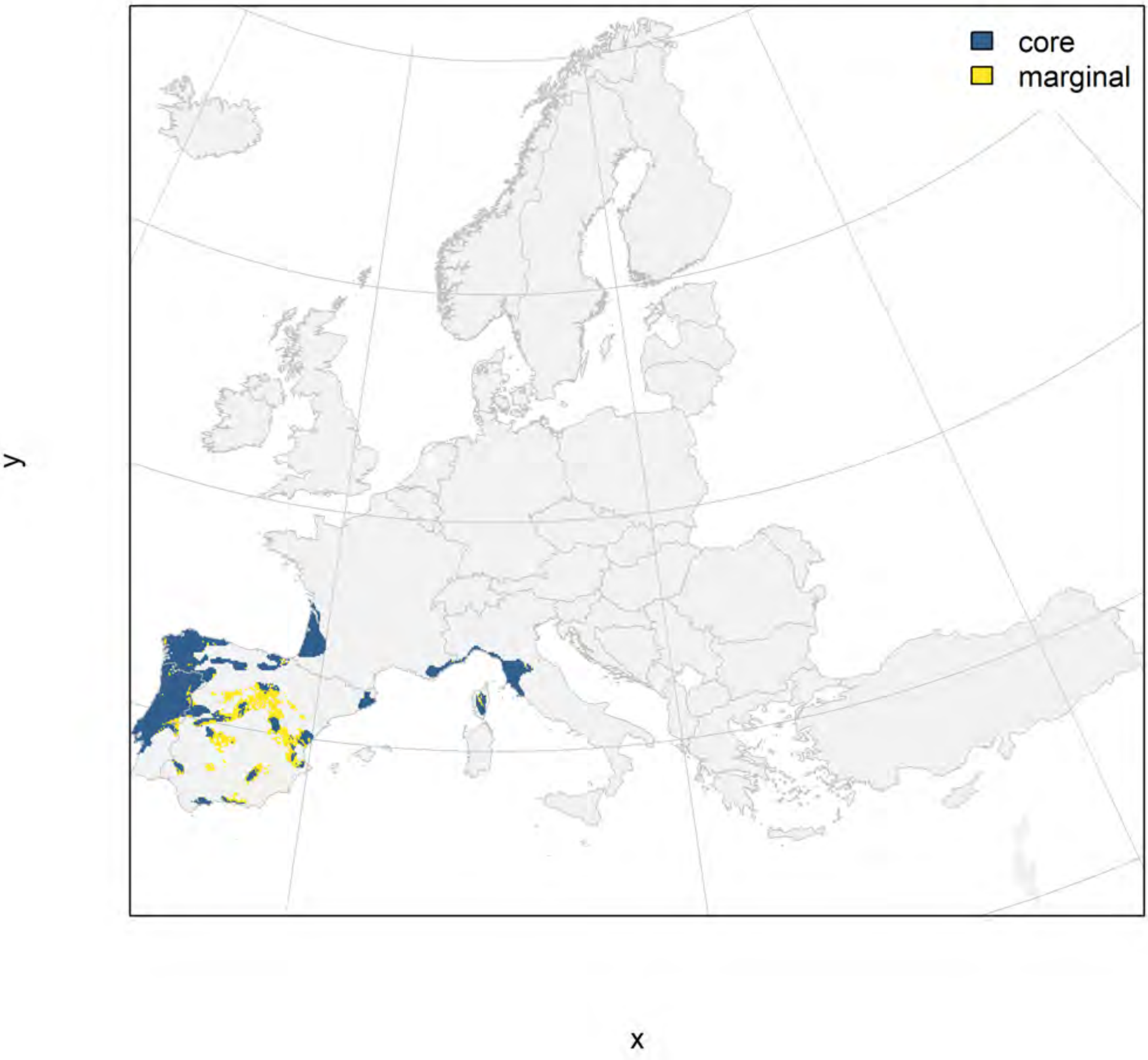

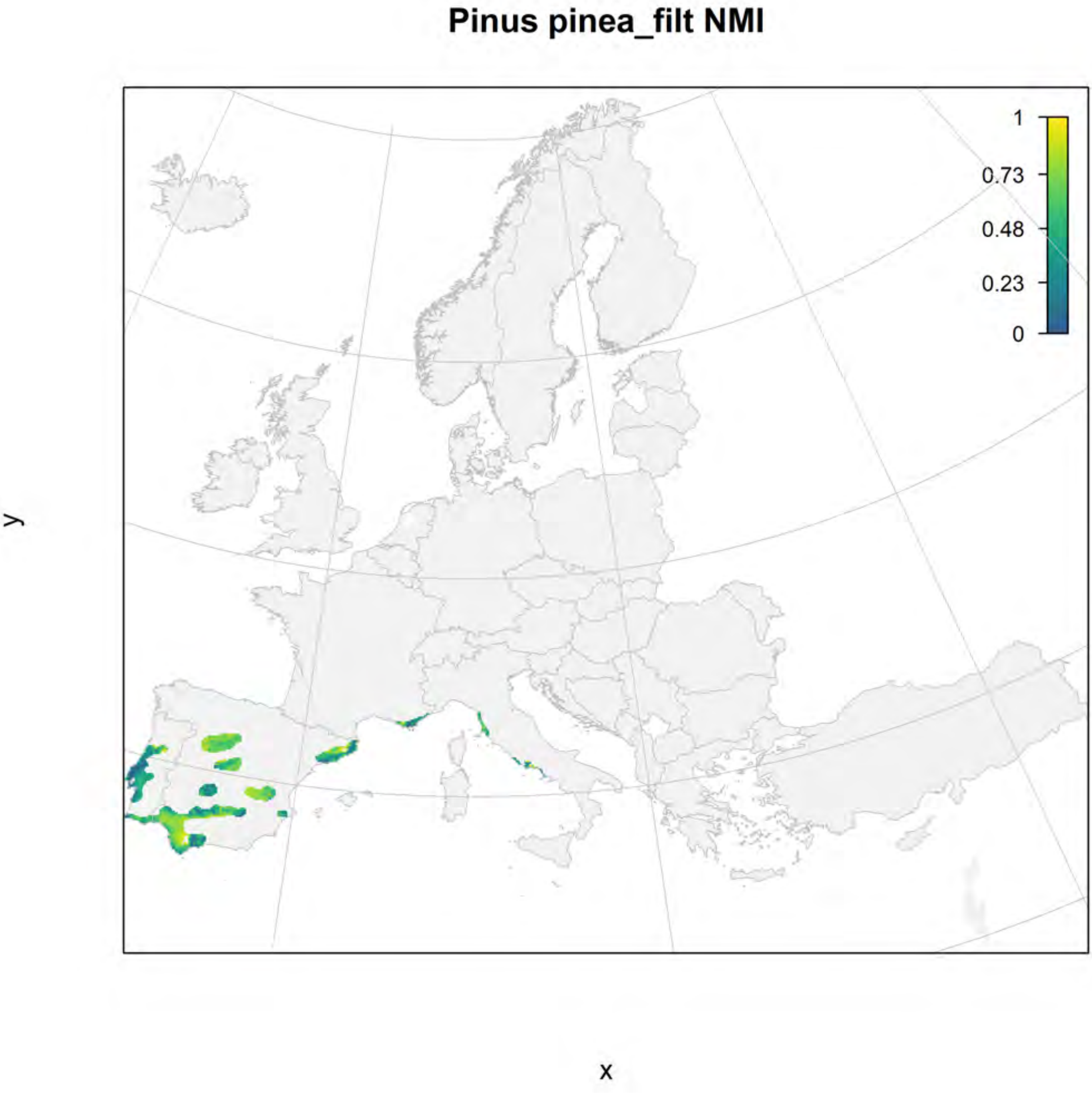
Pinus pinea_filt NMI
1
0.73
0.48
0.23
0
y
x

### Pinus pinea_filt current marginality

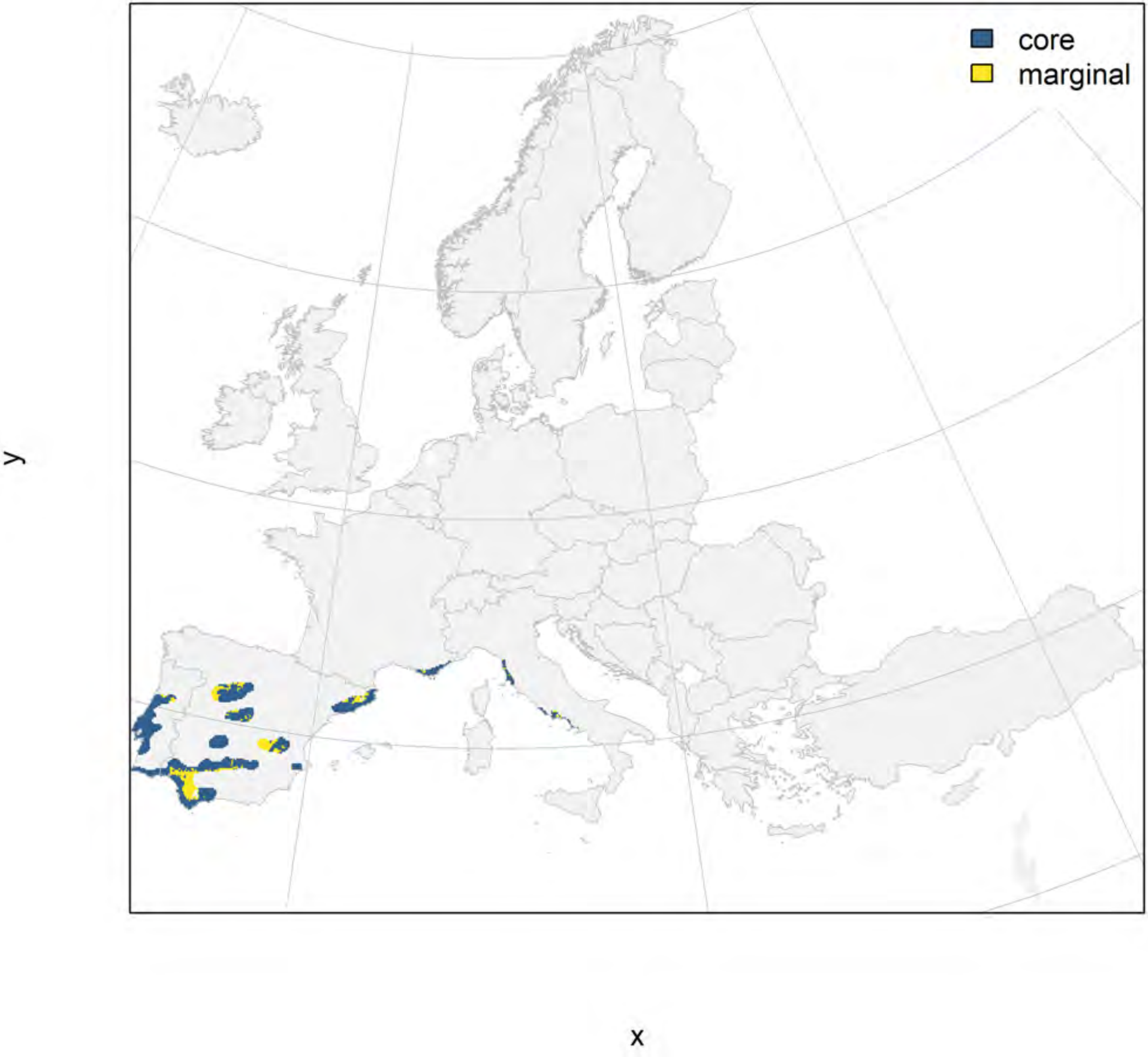

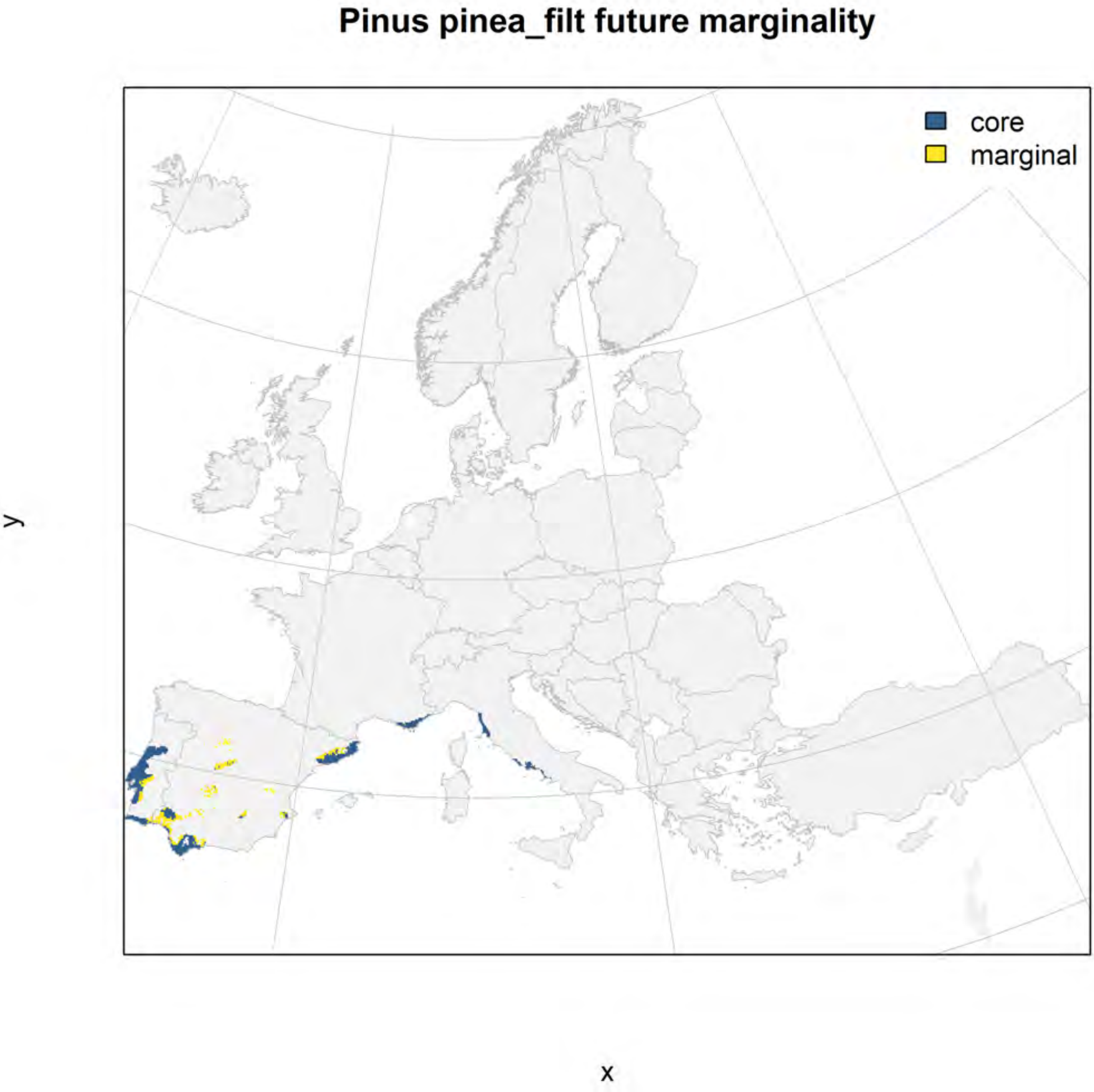
Pinus pinea_filt future marginality
core
marginal
y
x

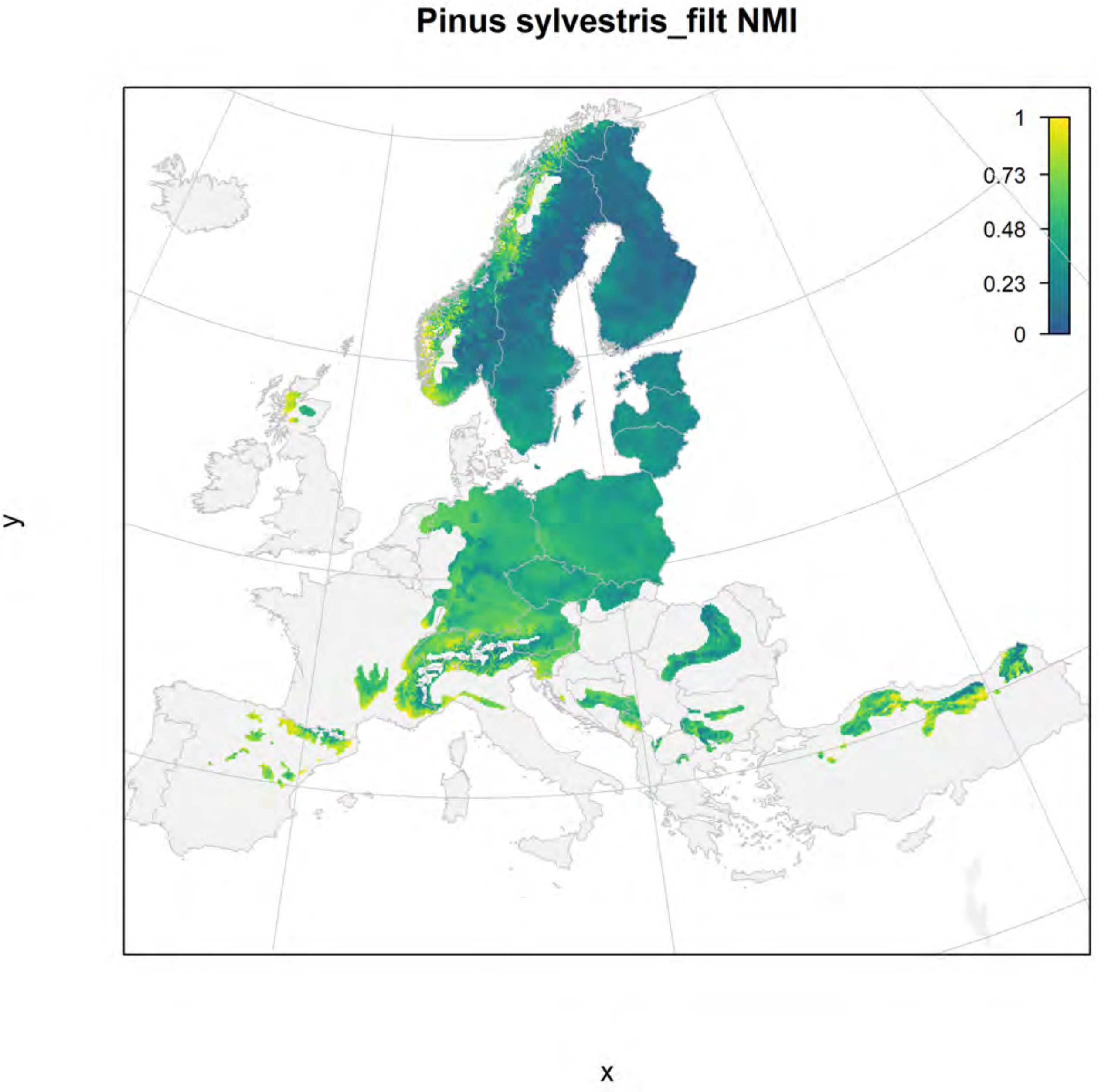
Pinus sylvestris_filt NMI
1
0.73
0.48
0.23
0
y
x

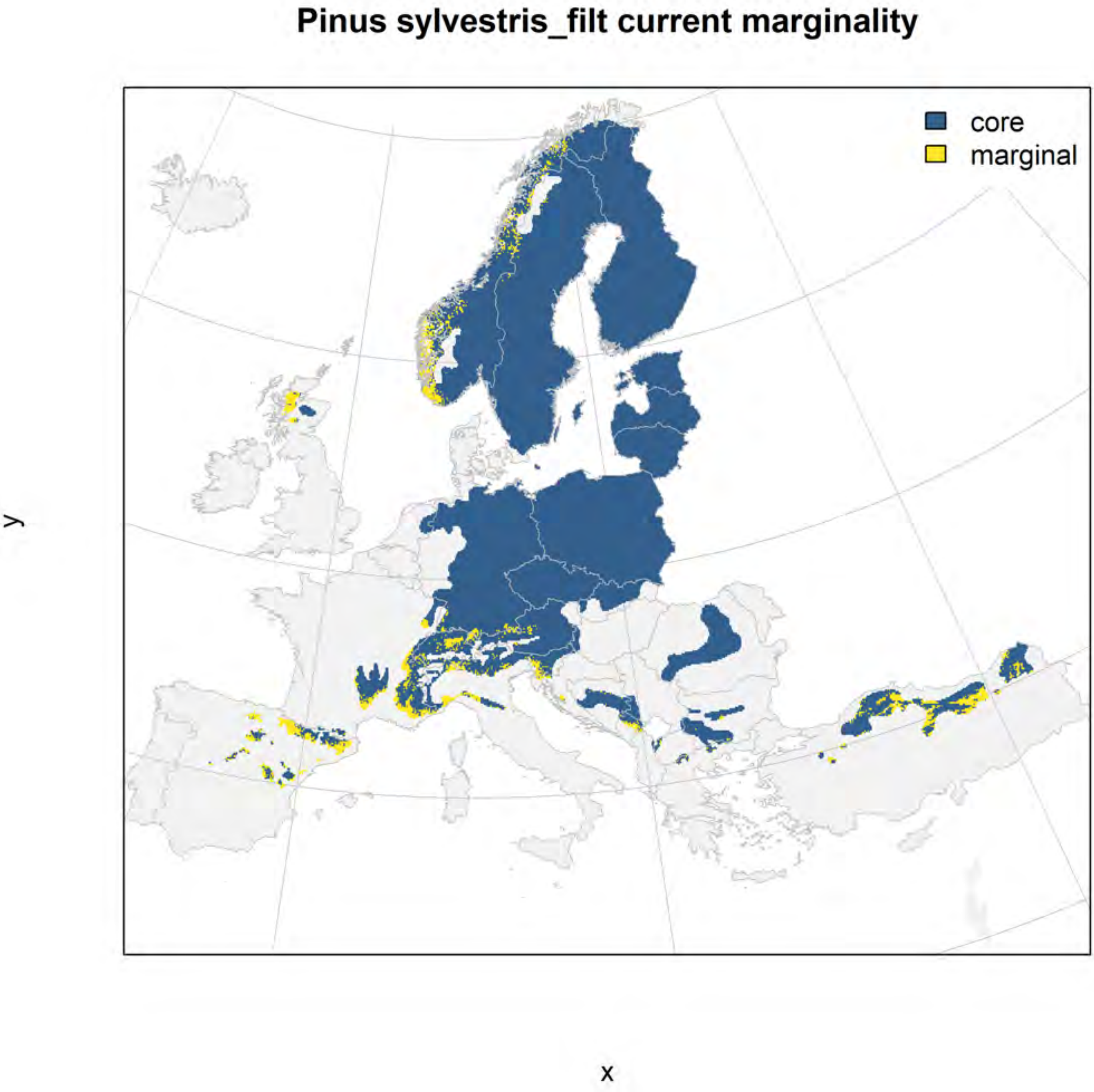
Pinus sylvestris_filt current marginality
core
marginal
y
x

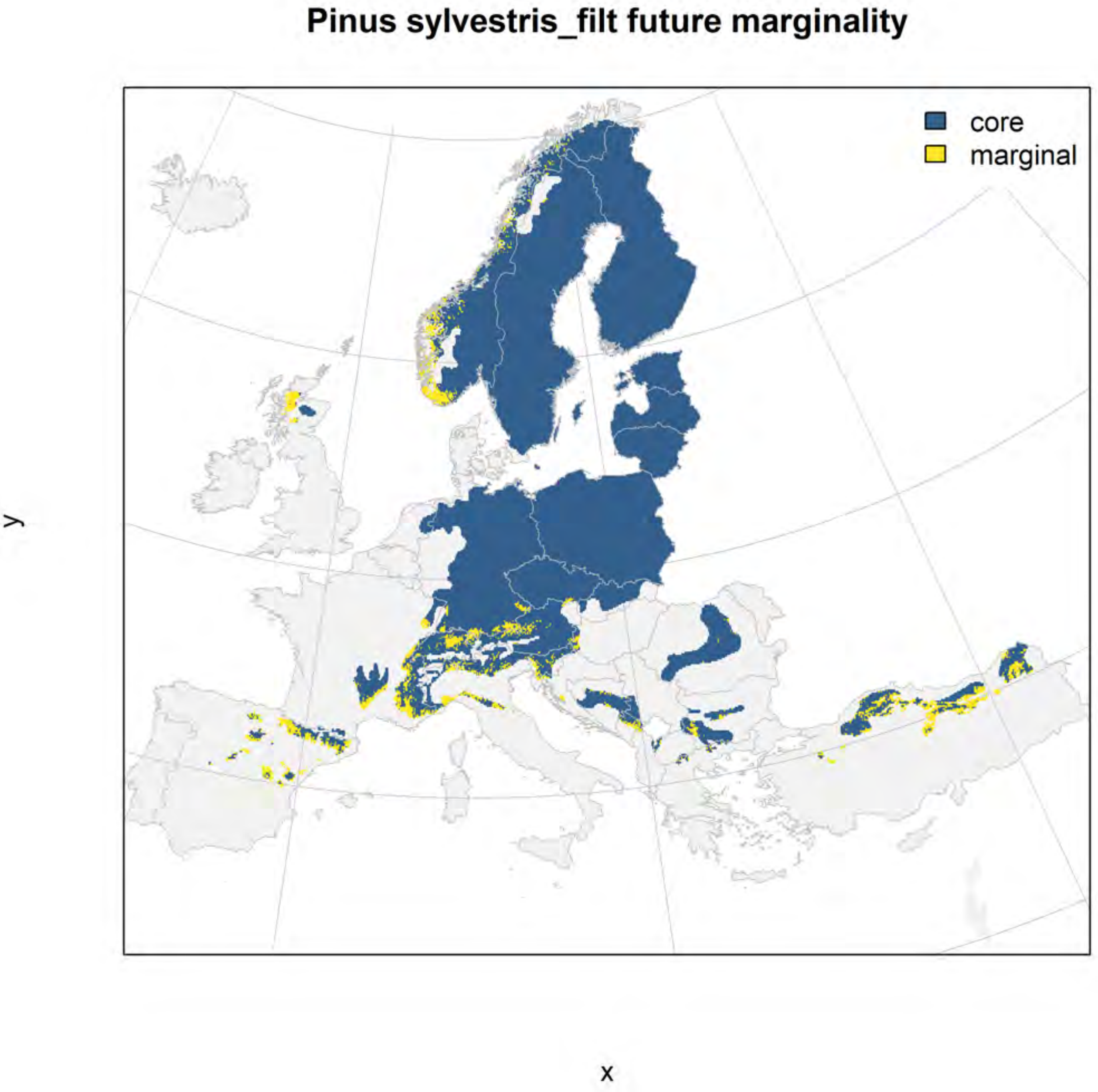
Pinus sylvestris_filt future marginality
core
marginal
y
x

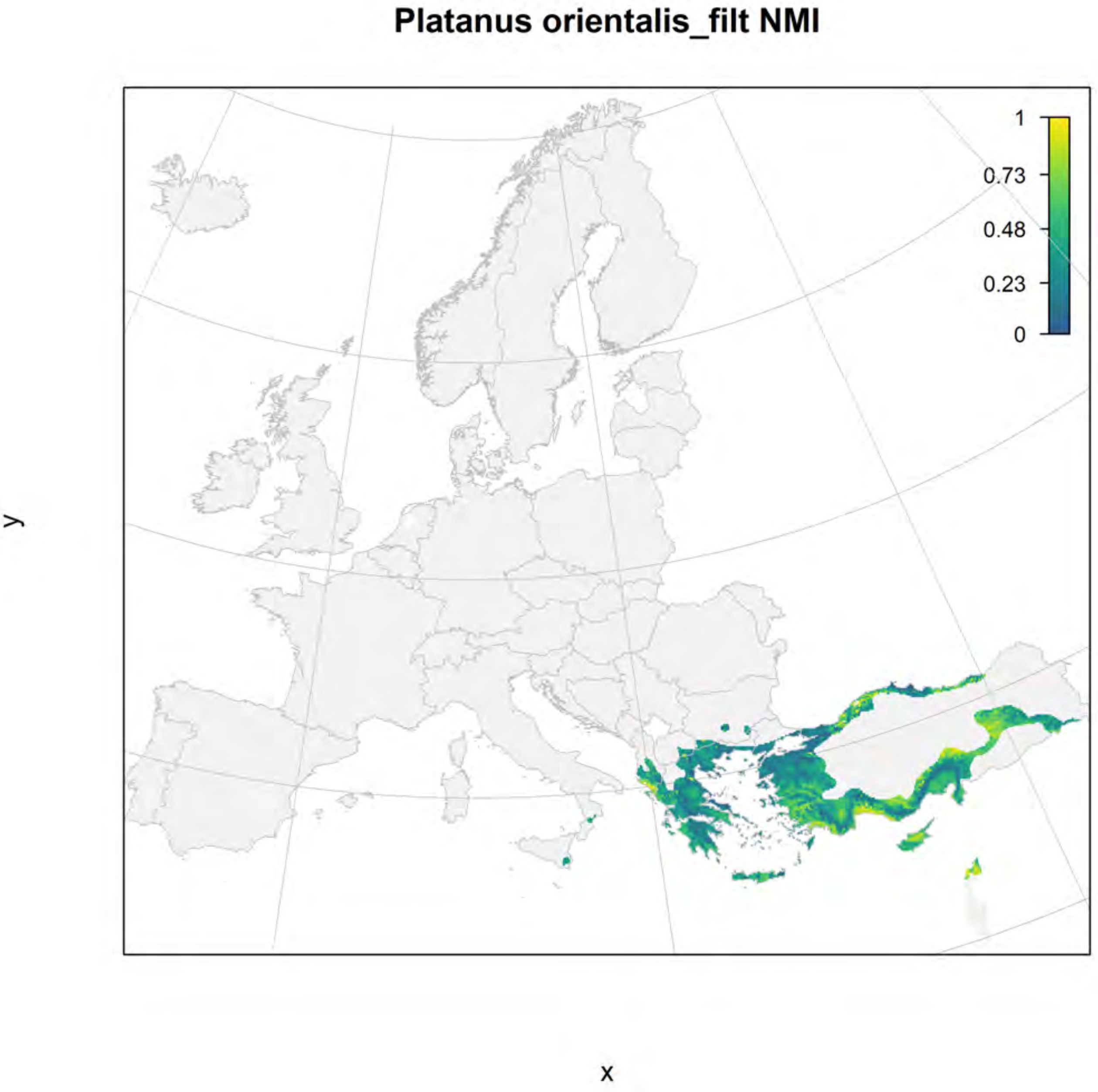

Platanus orientalis_filt NMI
1
0.73
0.48
0.23
0
y
x

### Platanus orientalis_filt current marginality

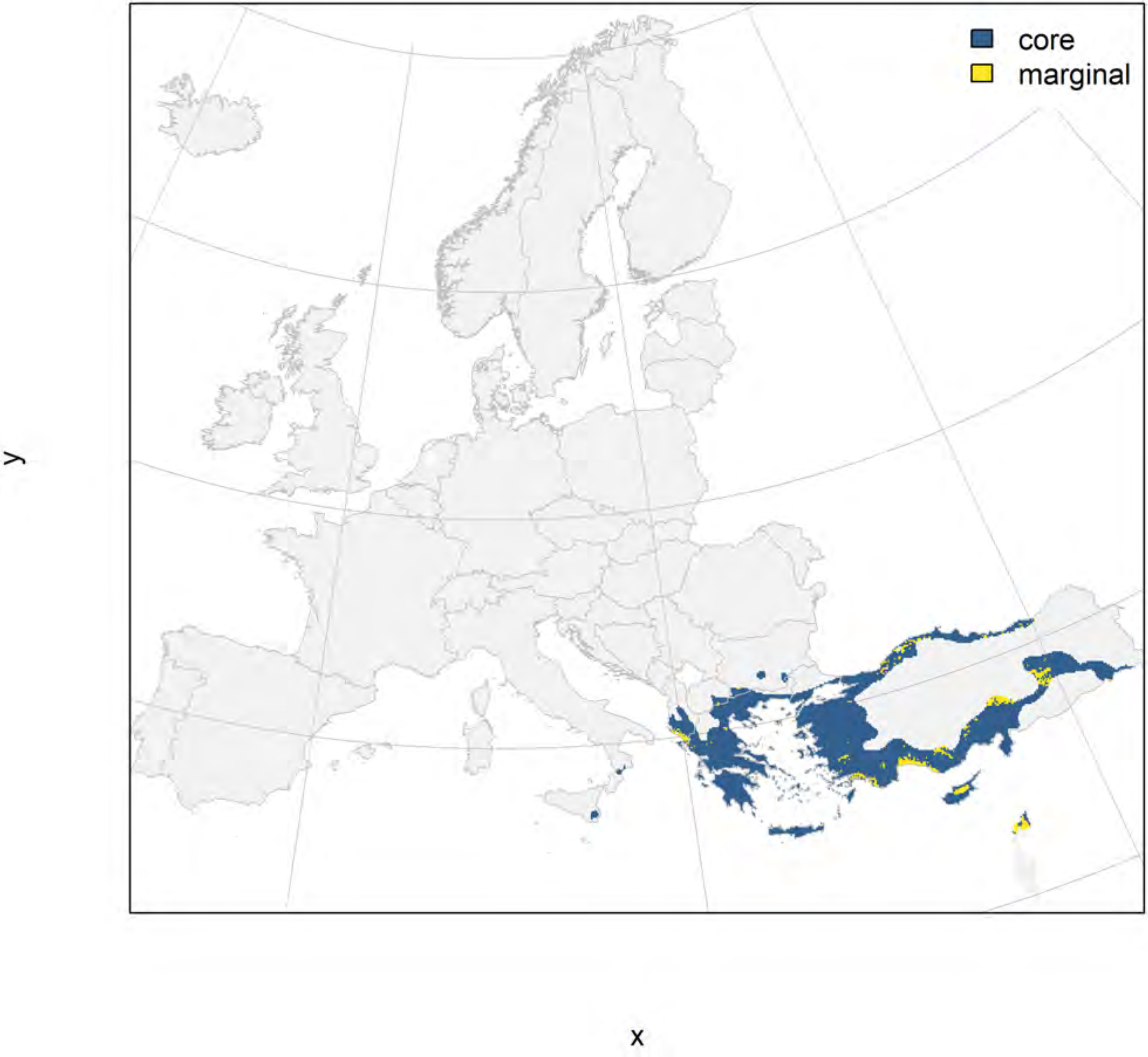

**Platanus orientalis_filt future marginality**

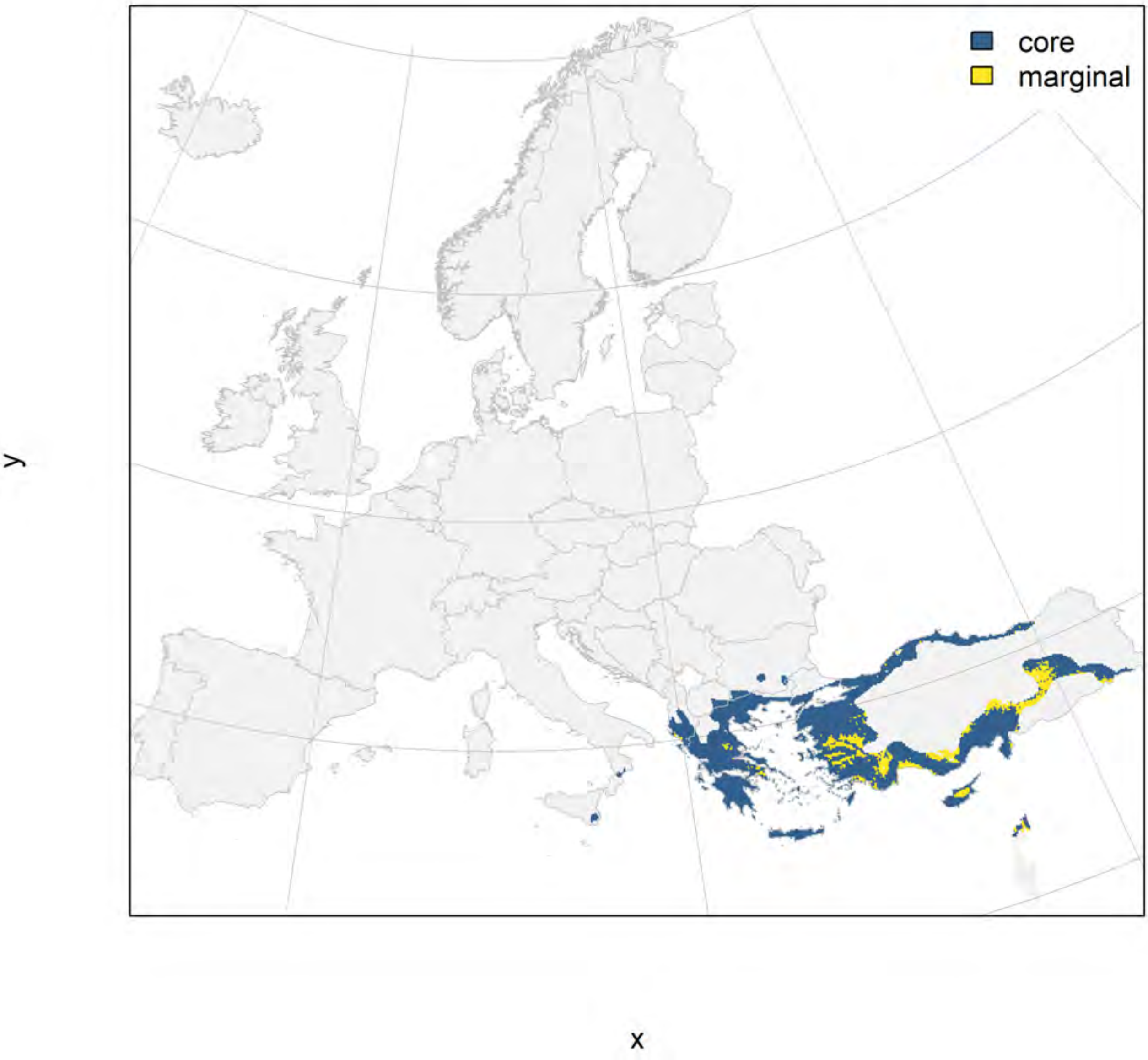

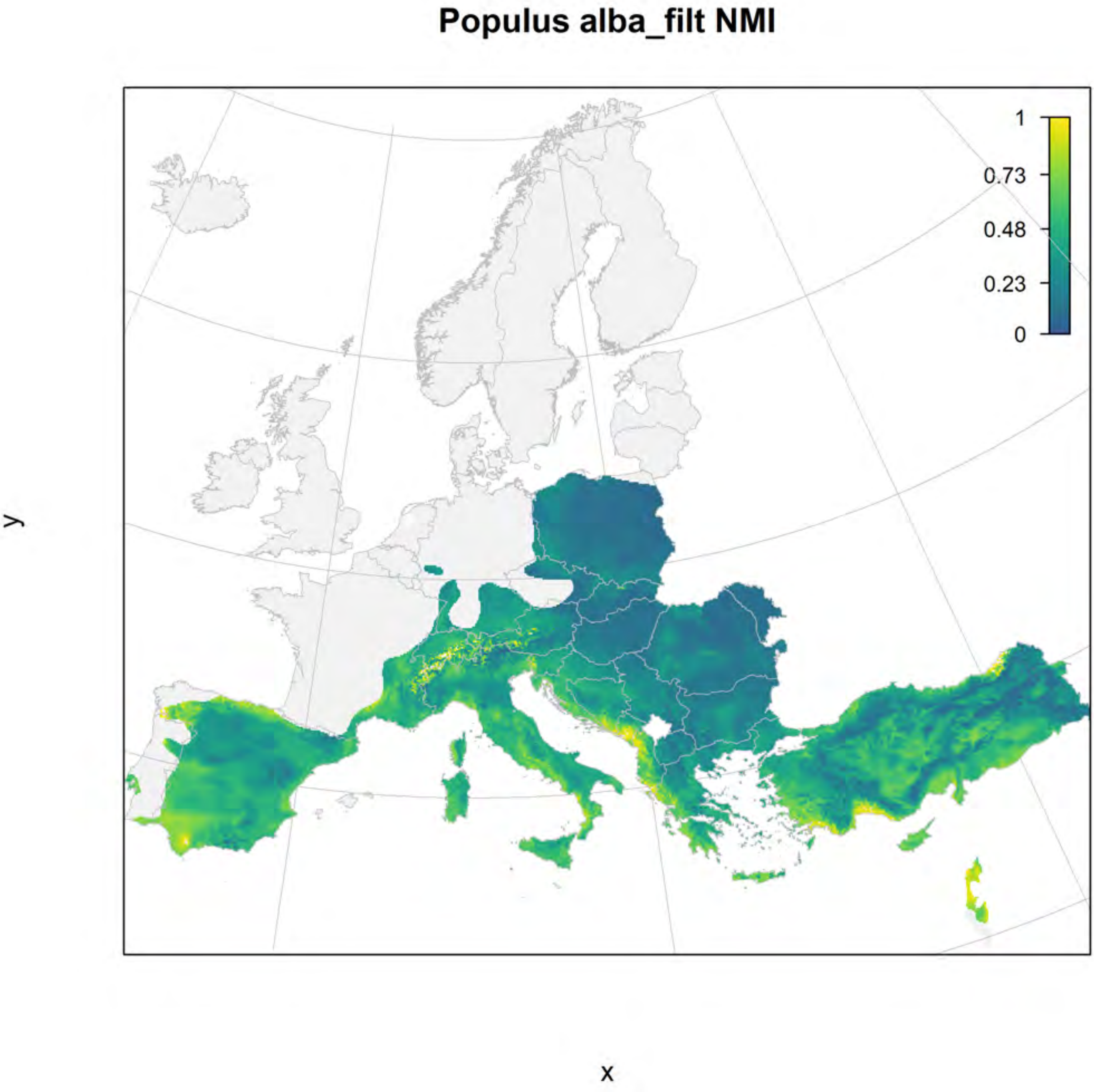
Populus alba_filt NMI
1
0.73
0.48
0.23
0
y
x

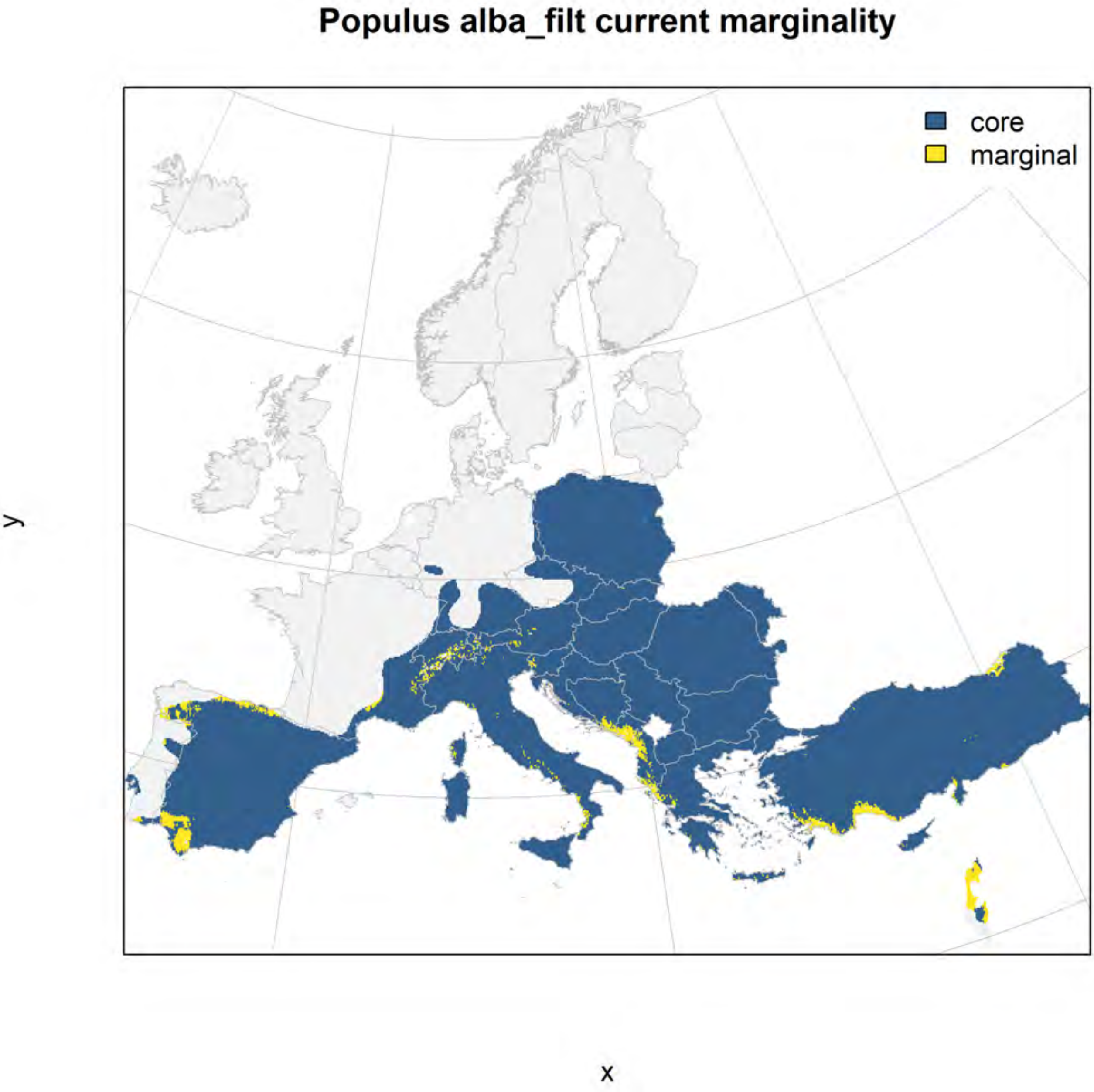
Populus alba_filt current marginality
core
marginal
y
x

#### Populus alba_filt future marginality

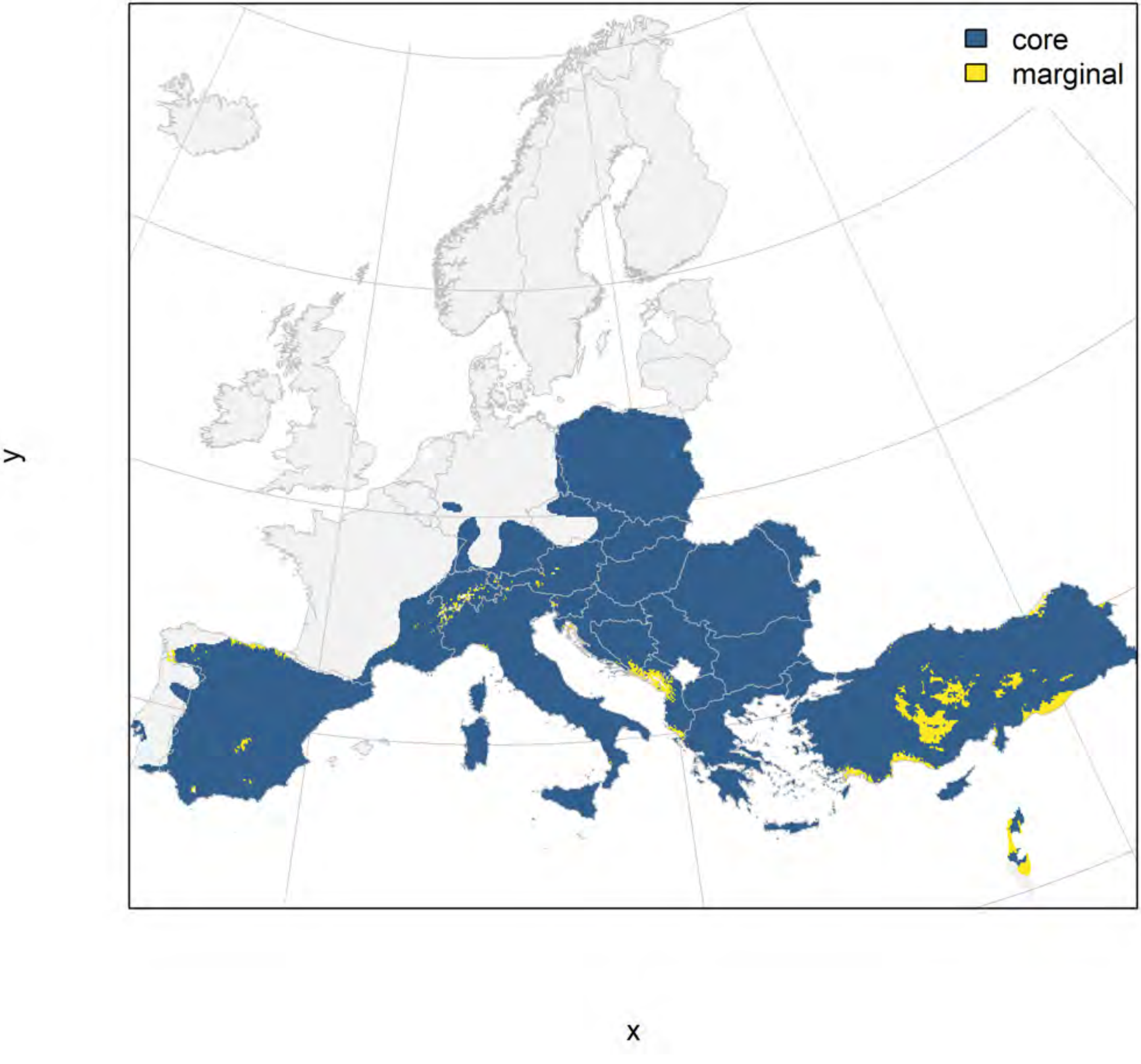

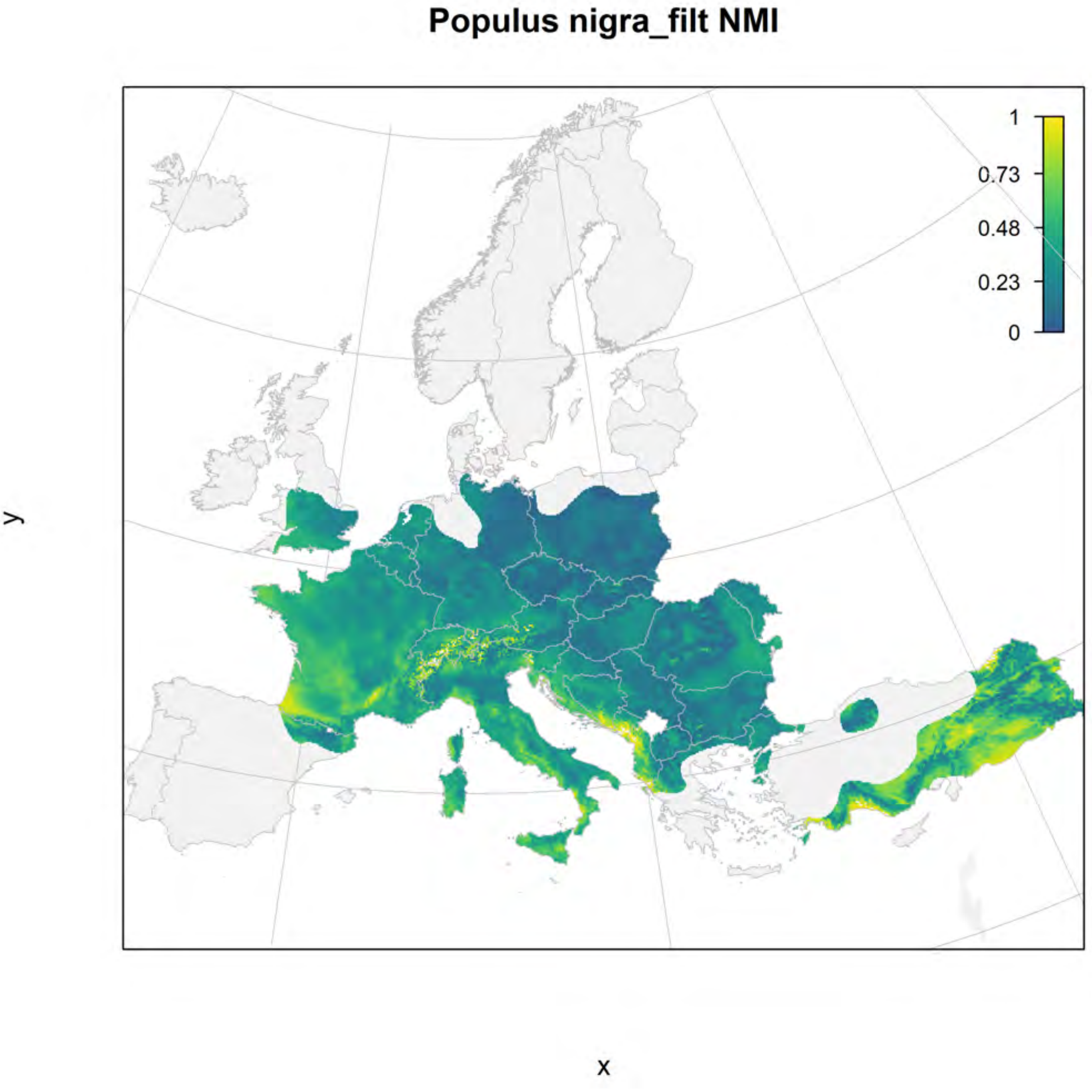
Populus nigra_filt NMI
1
0.73
0.48
0.23
0
y
x

#### Populus nigra_filt current marginality

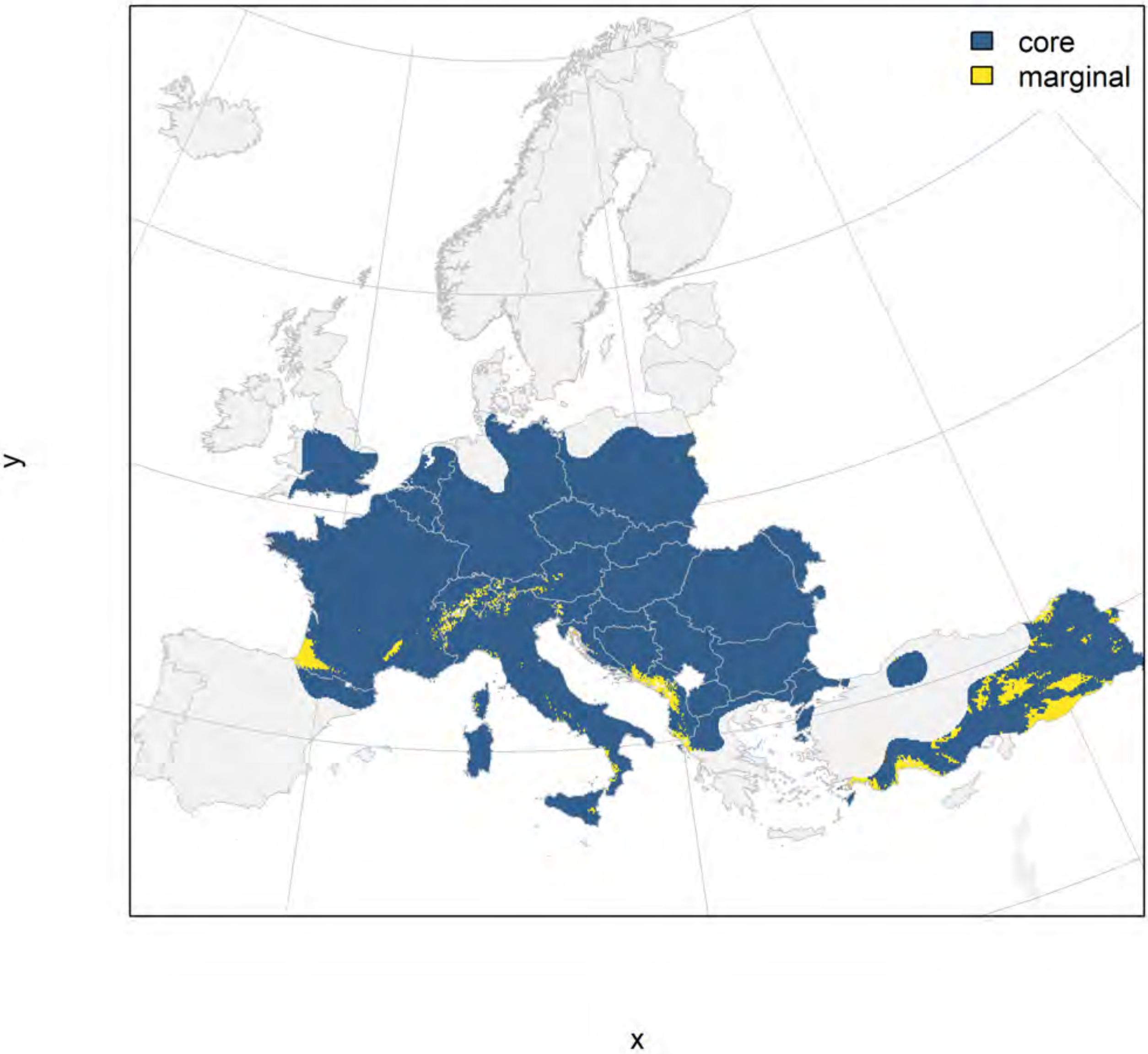

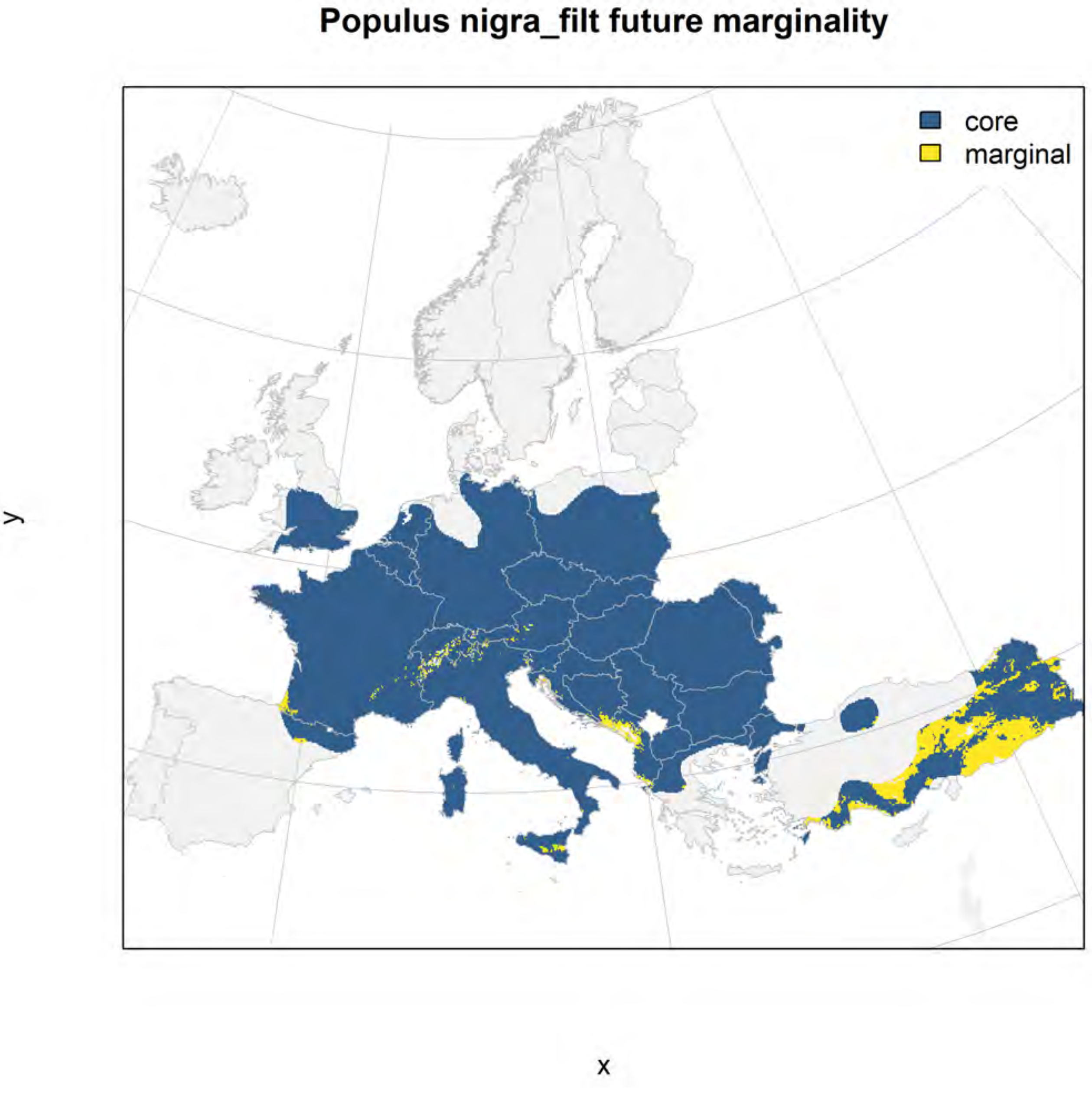
Populus nigra_filt future marginality
core
marginal
y
x

### Populus tremula_filt NMI

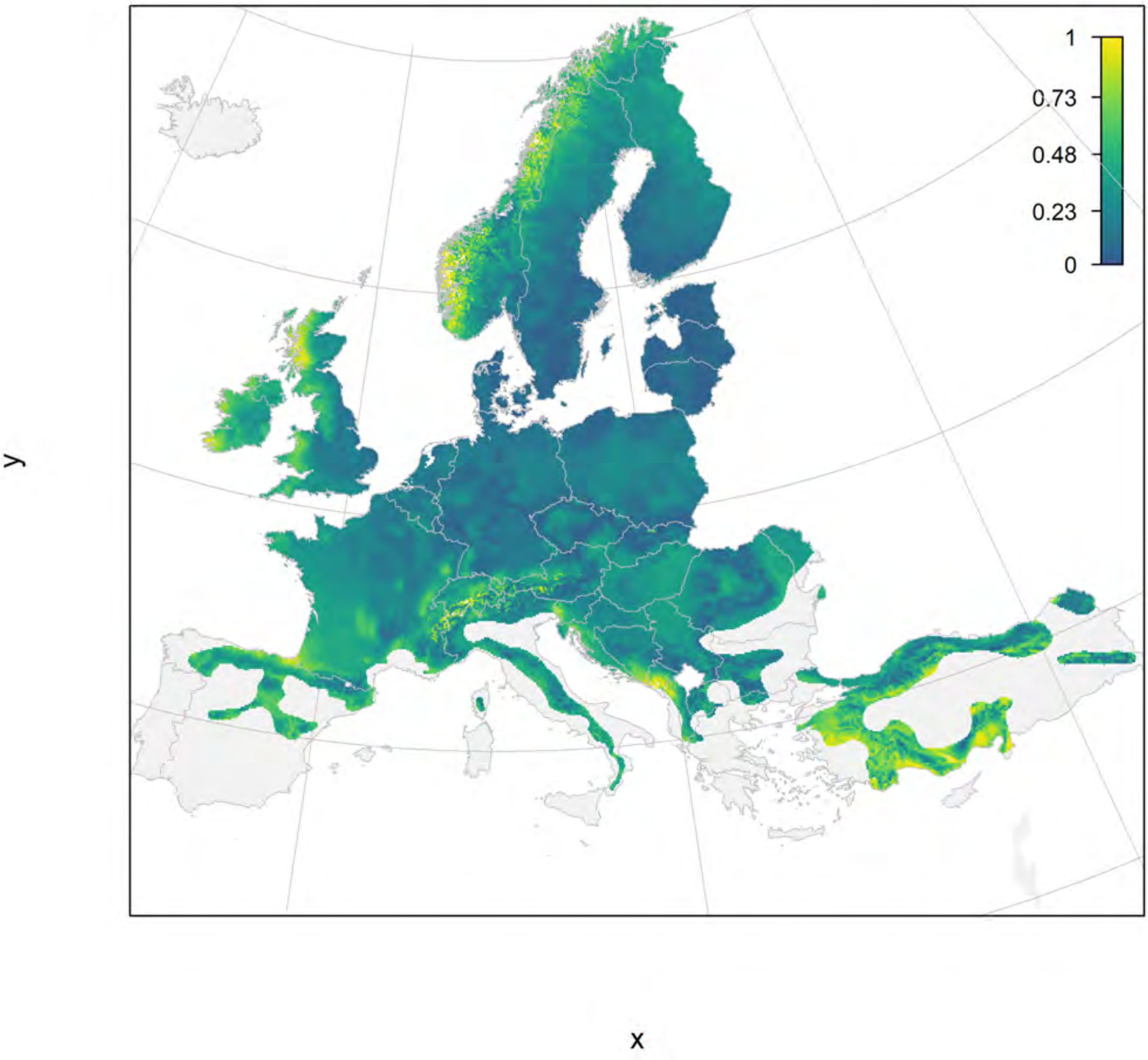

#### Populus tremula_filt current marginality

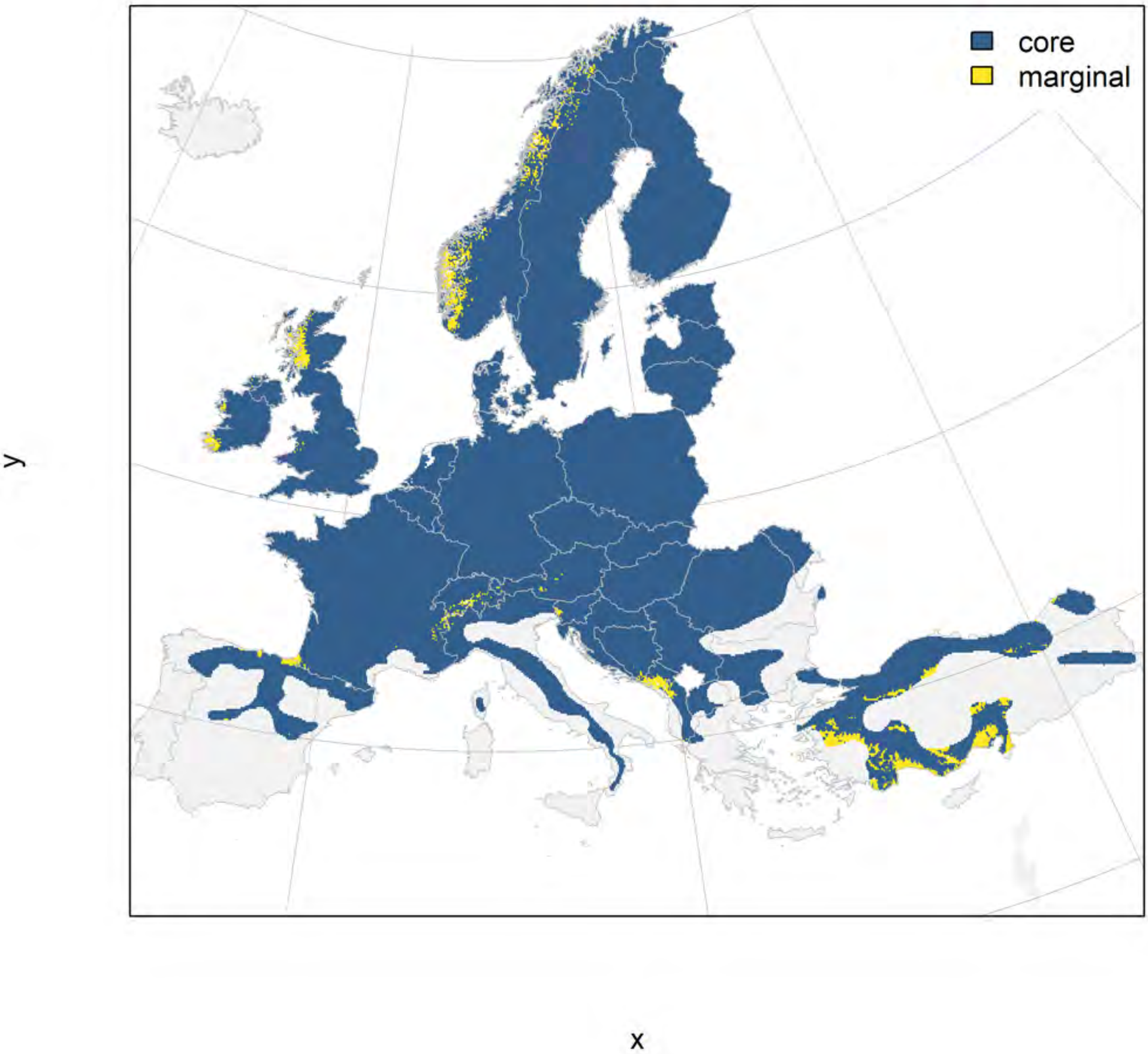

### Populus tremula_filt future marginality

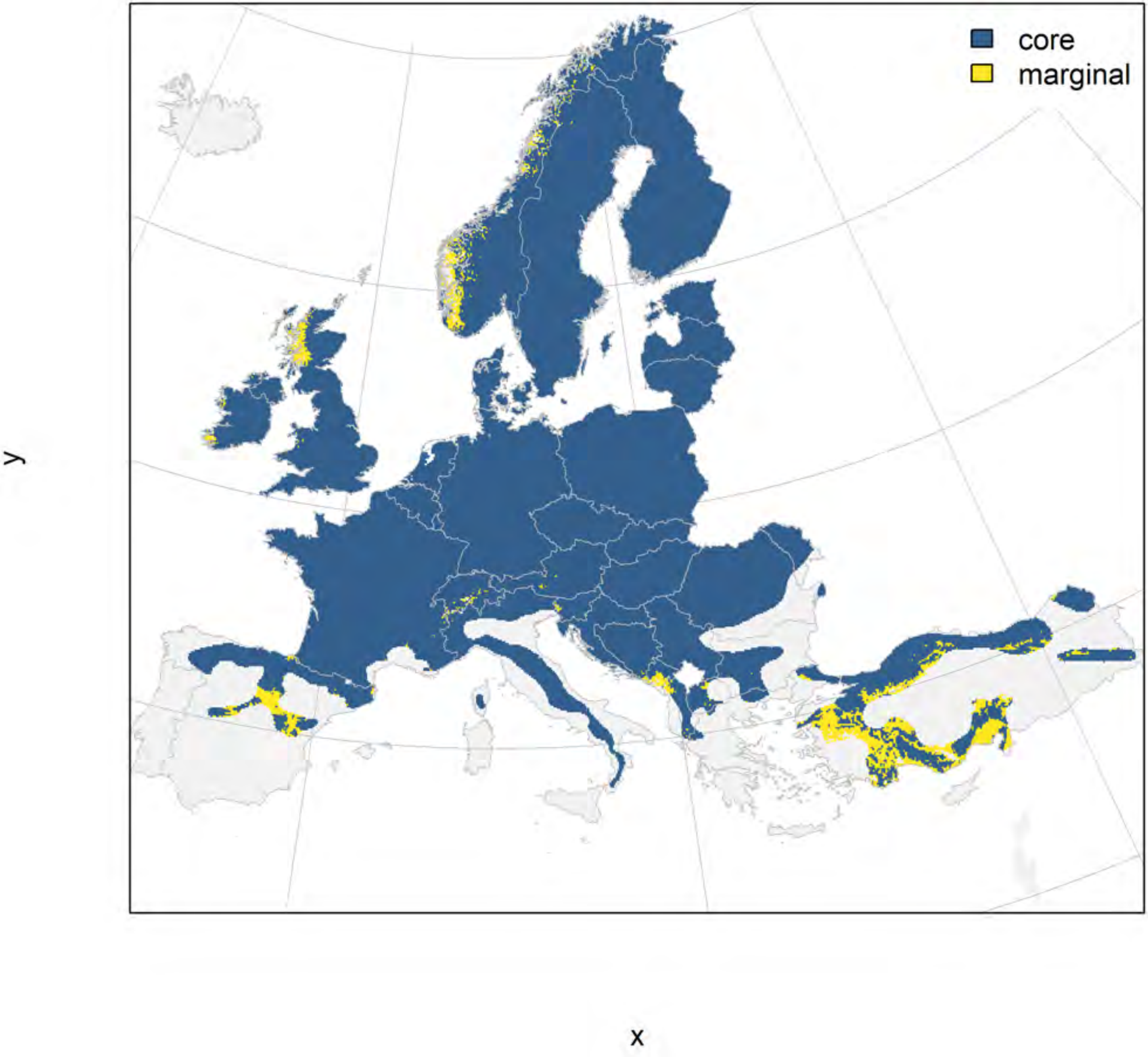

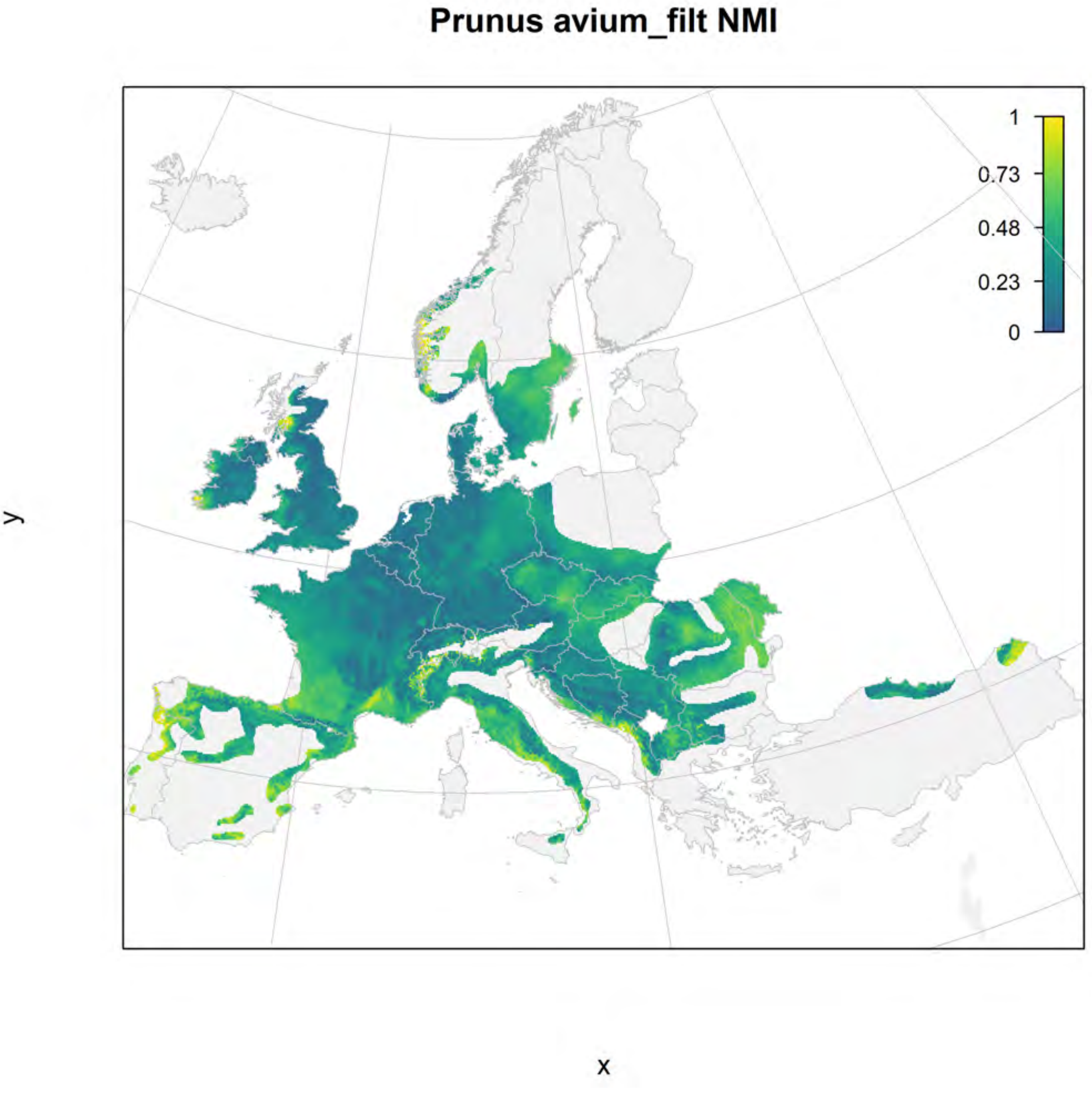
Prunus avium_filt NMI
1
0.73
0.48
0.23
0
y
x

### Prunus avium_filt current marginality

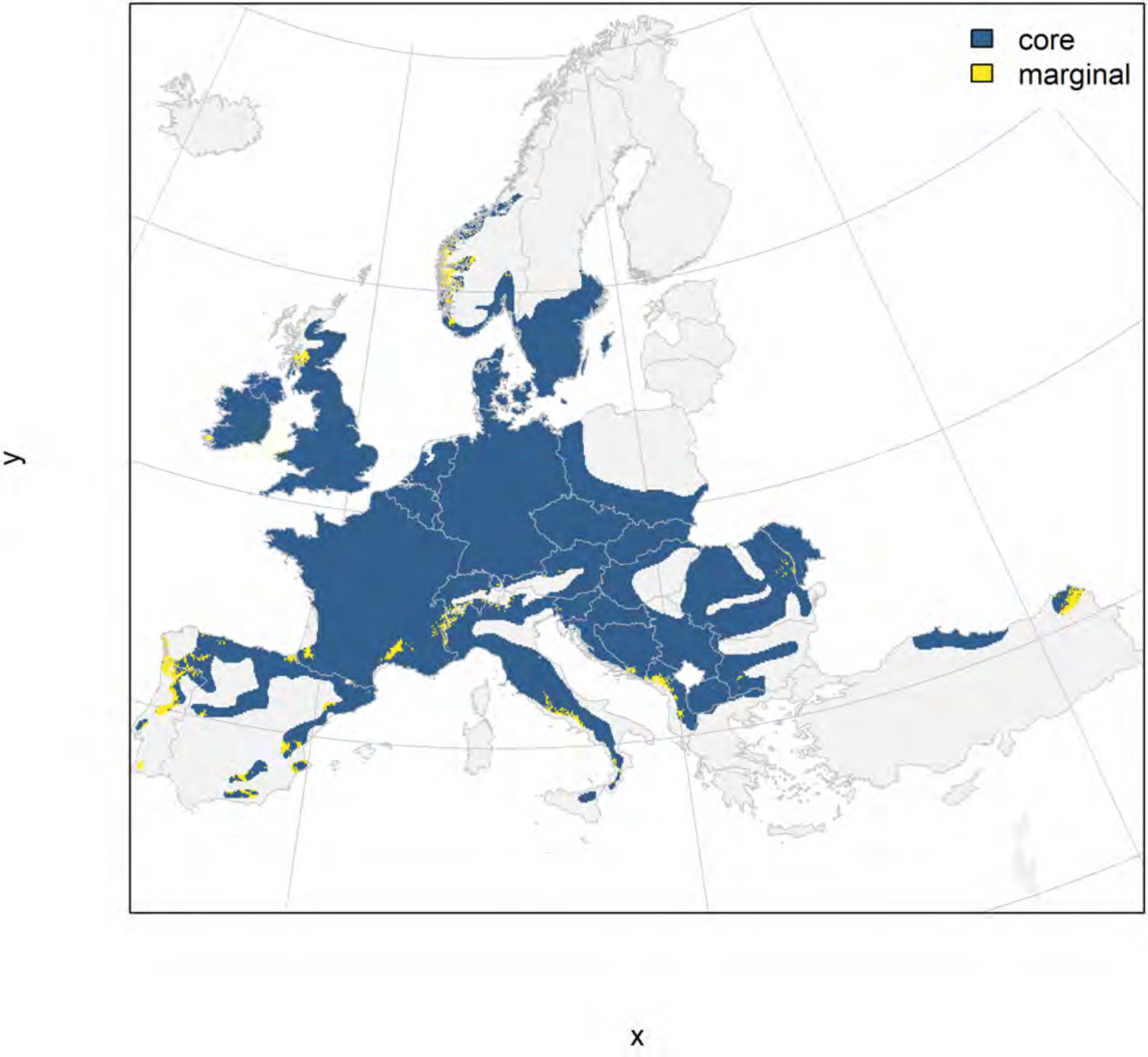

### Prunus avium_filt future marginality

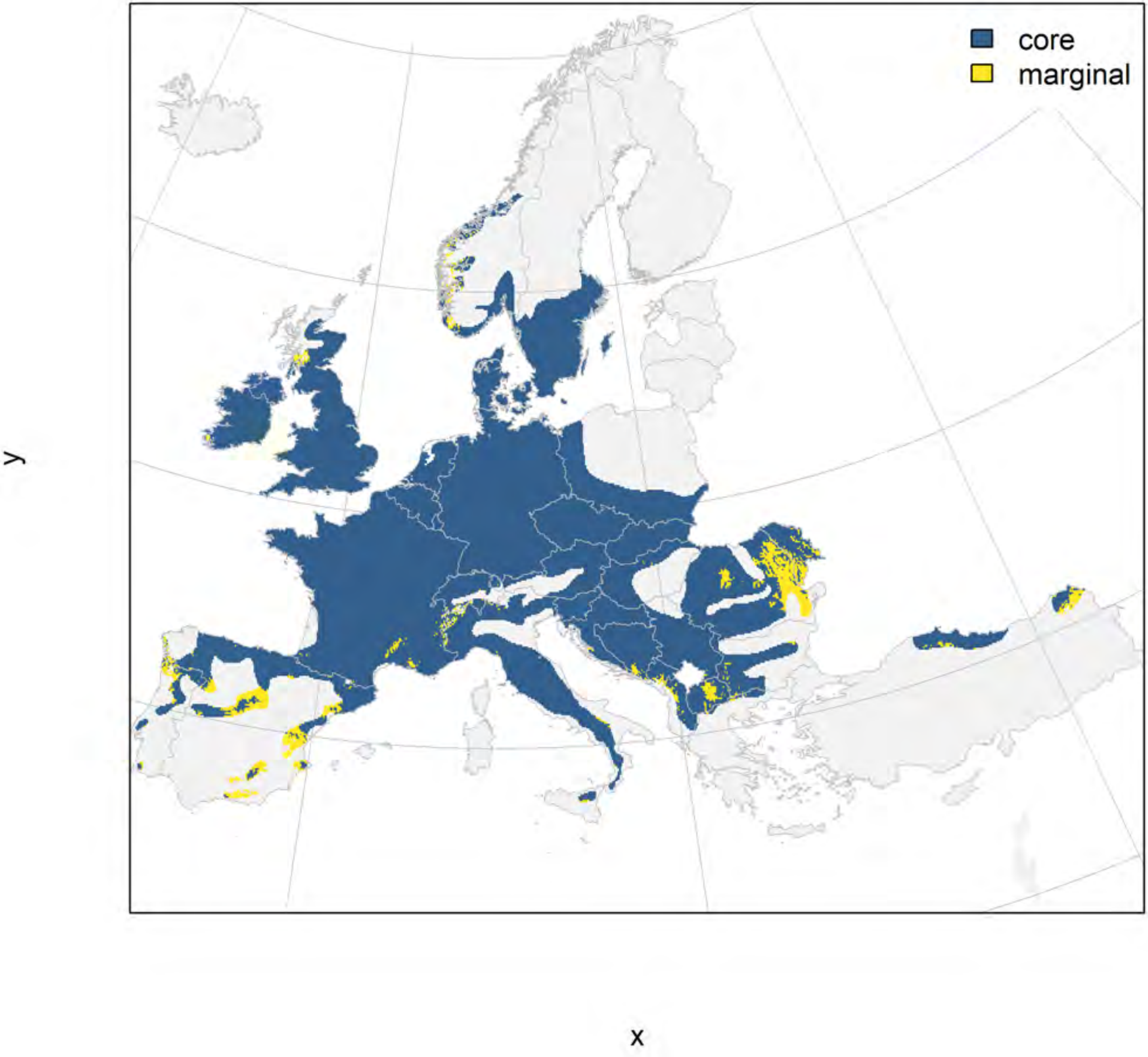

### Prunus padus_filt NMI

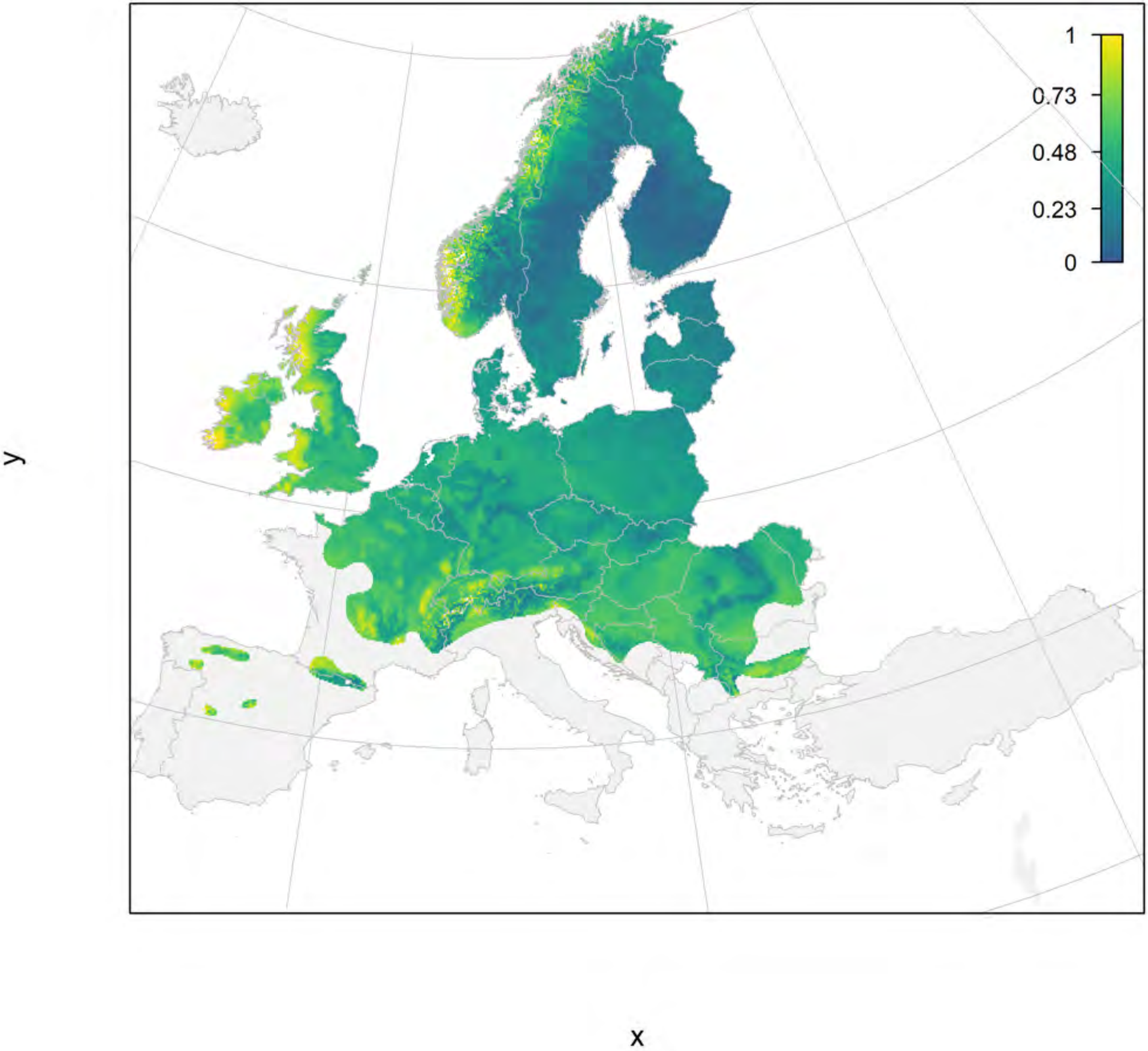

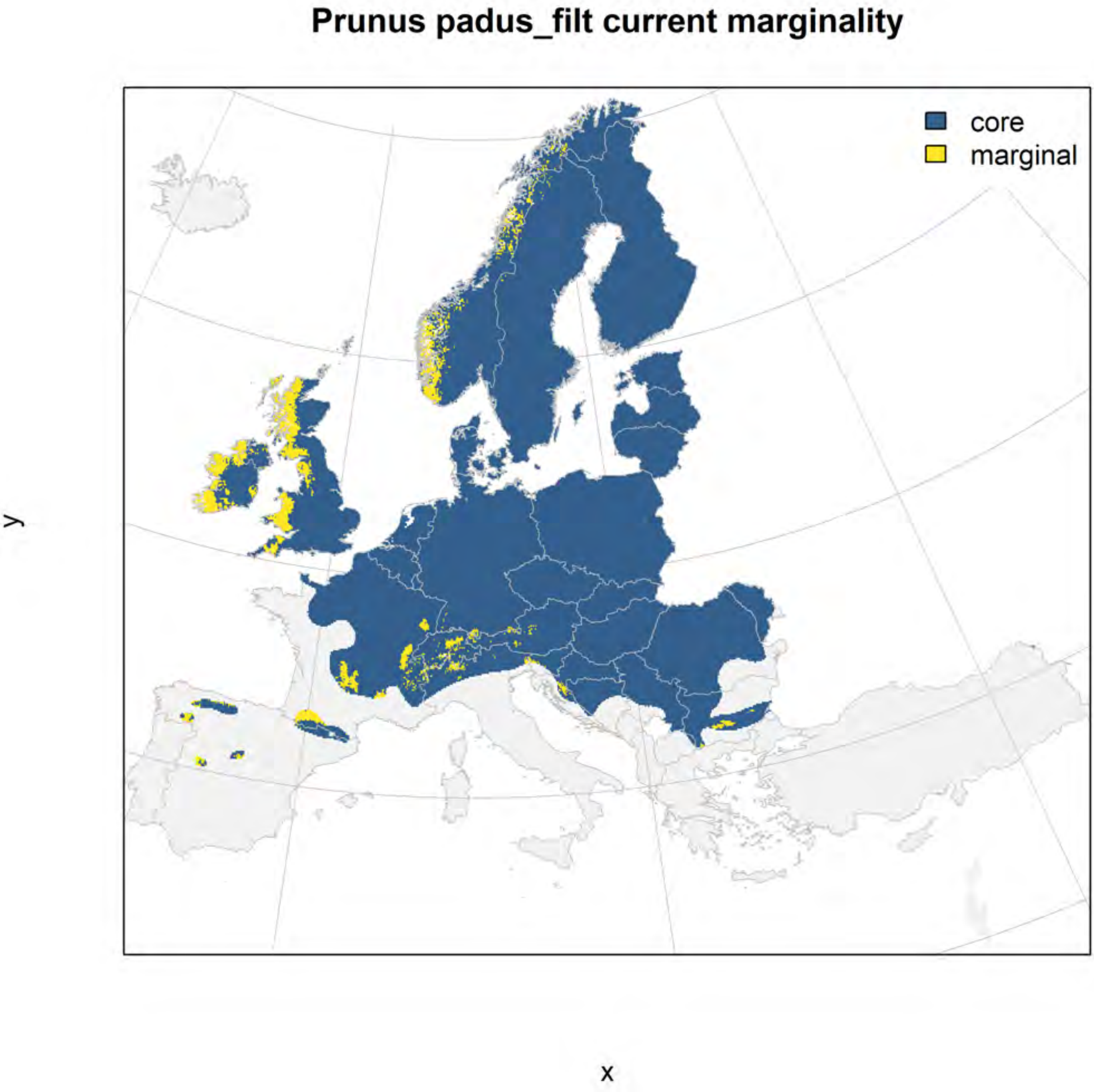
Prunus padus_filt current marginality
core
marginal
y
x

#### Prunus padus_filt future marginality

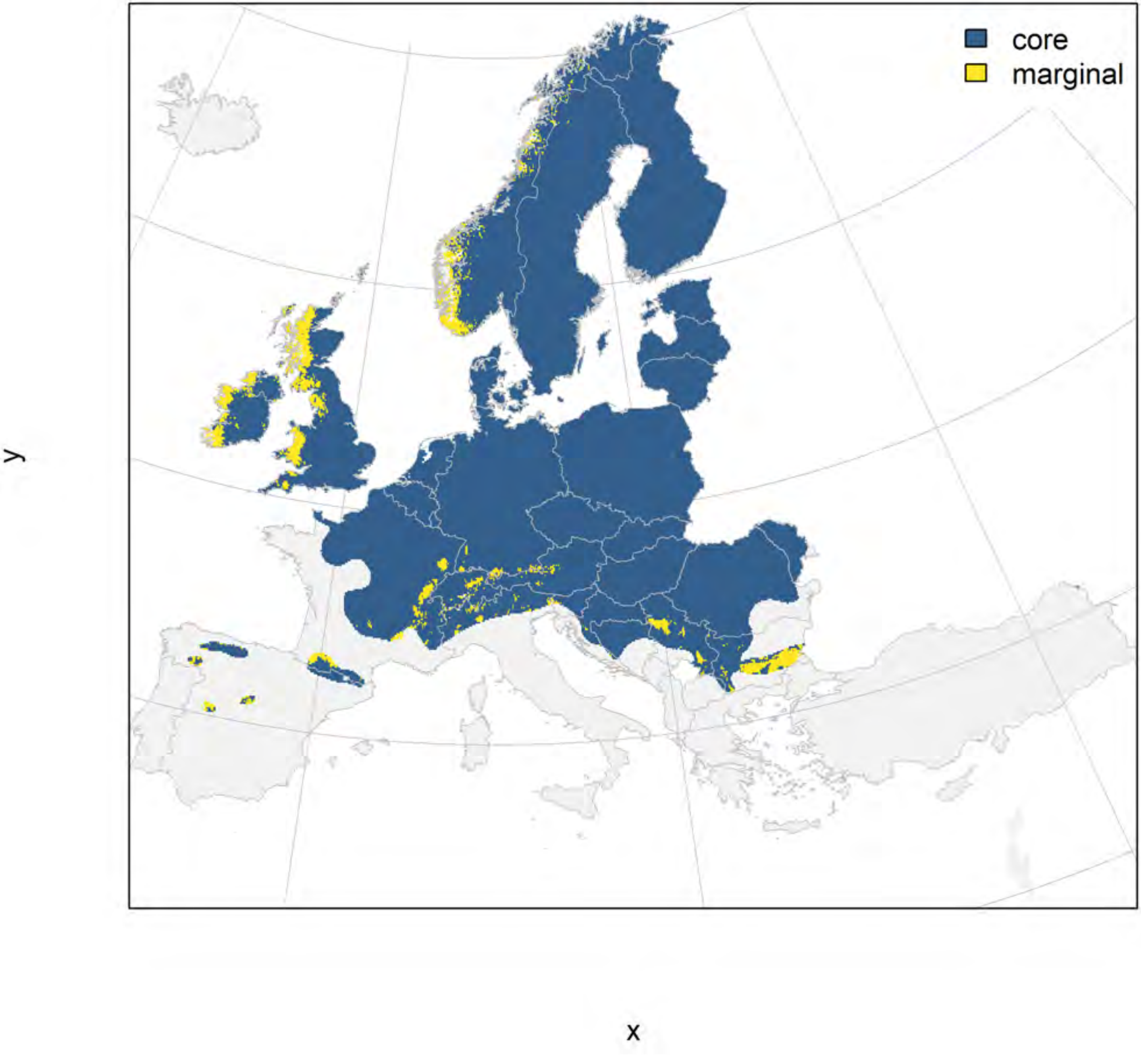

### Prunus spinosa_filt NMI

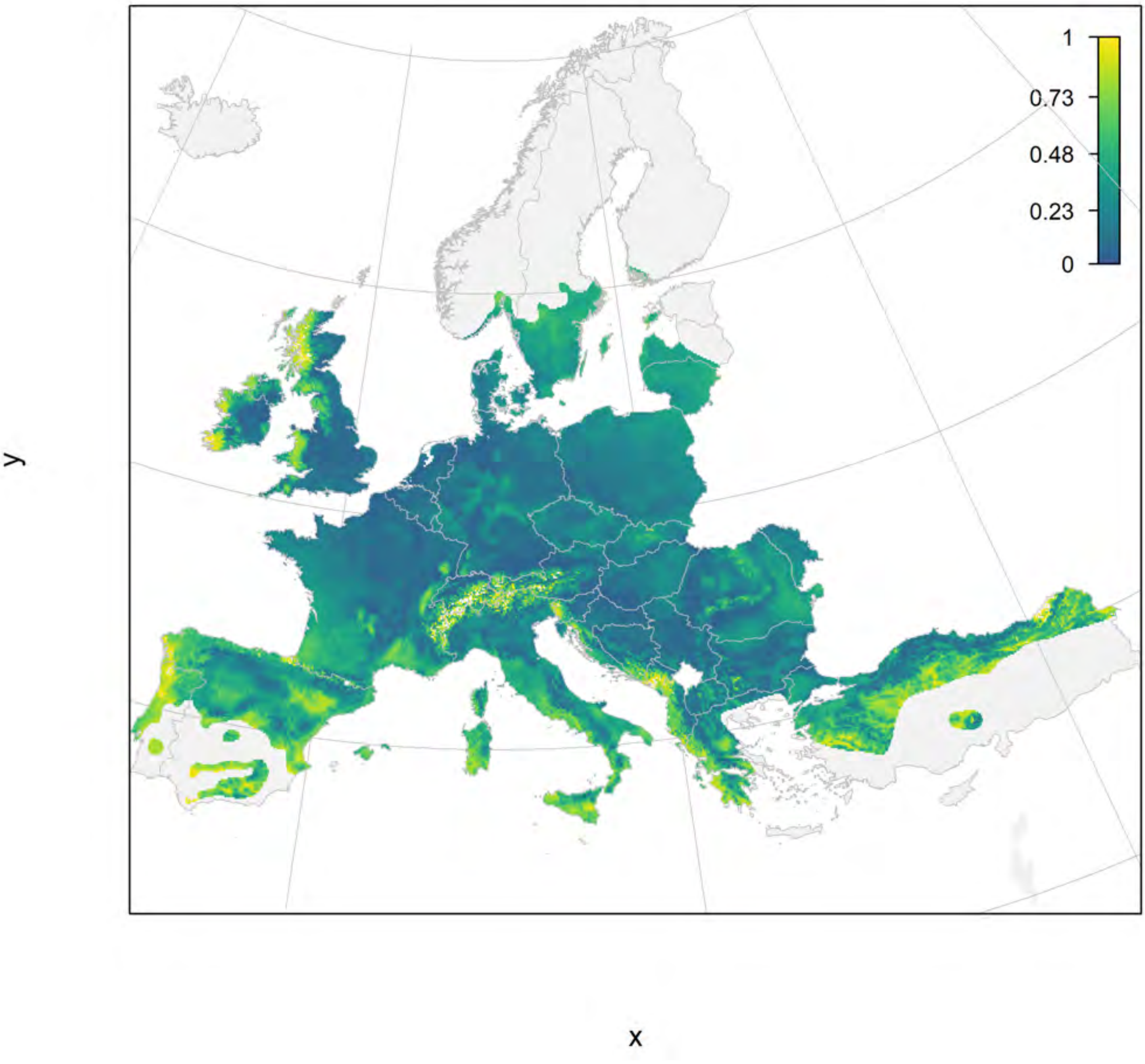

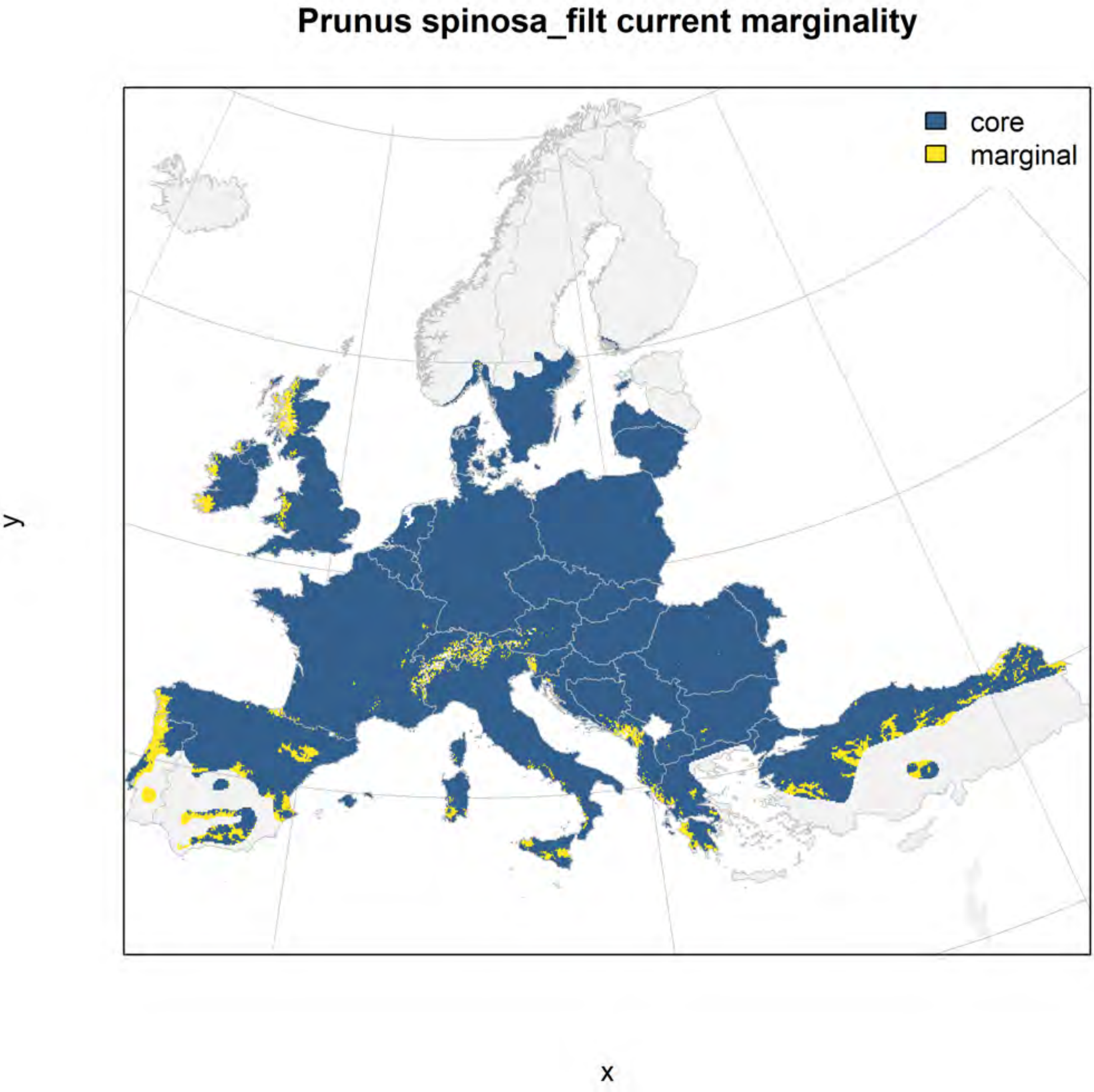
Prunus spinosa_filt current marginality
core
marginal
y
x

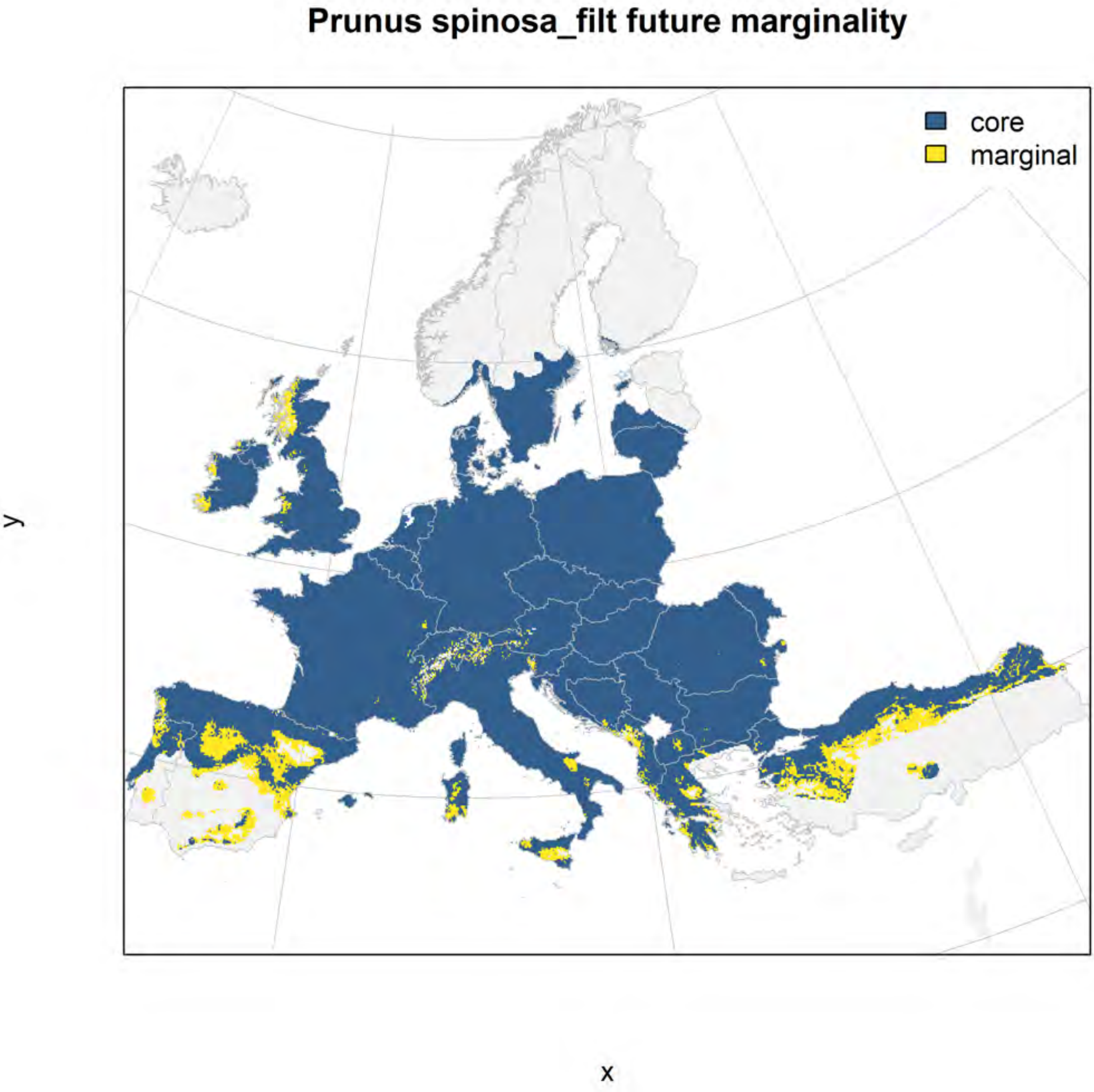

Prunus spinosa_filt future marginality
core
marginal
y
x

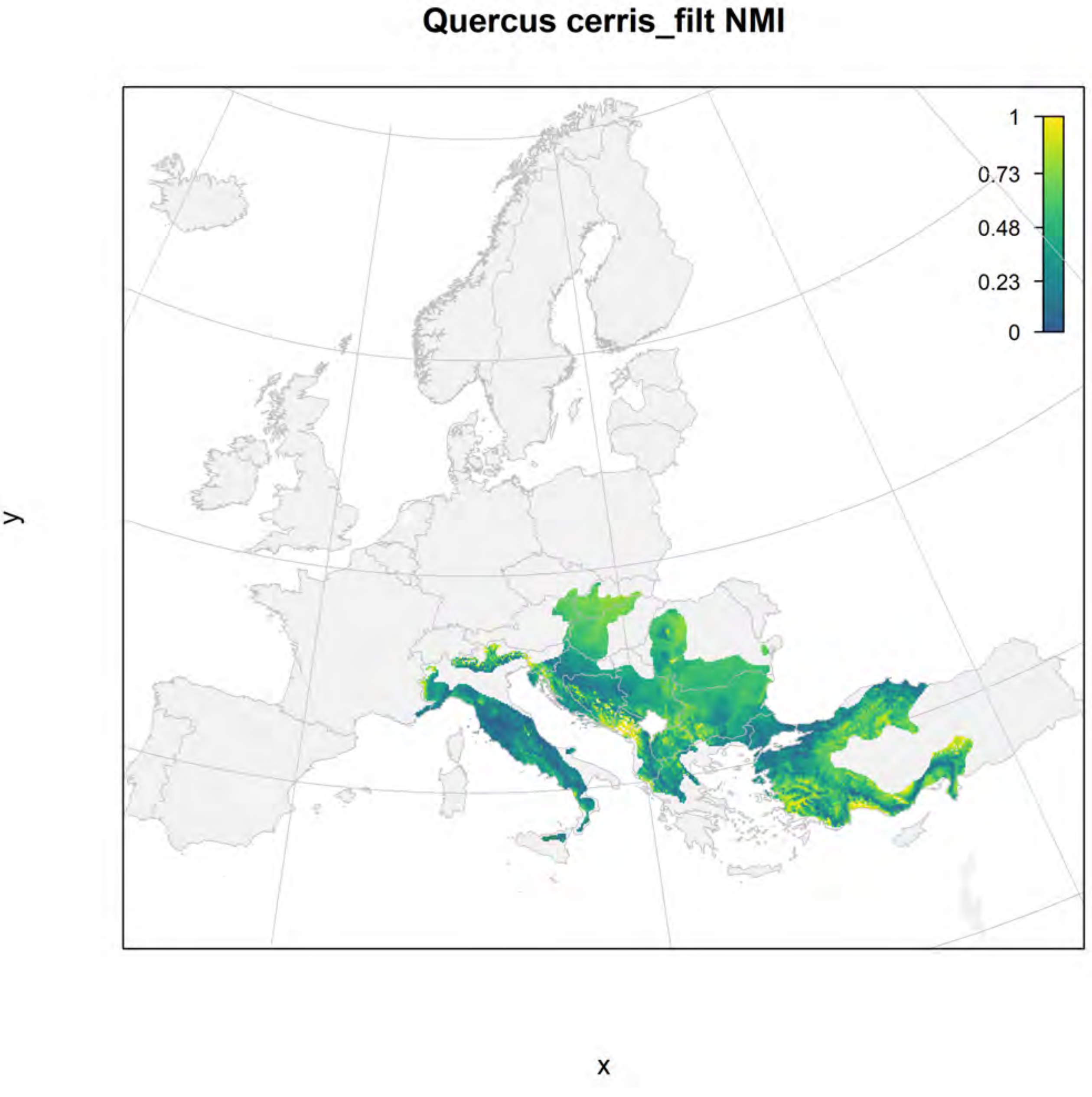
Quercus cerris_filt NMI
1
0.73
0.48
0.23
0
y
x

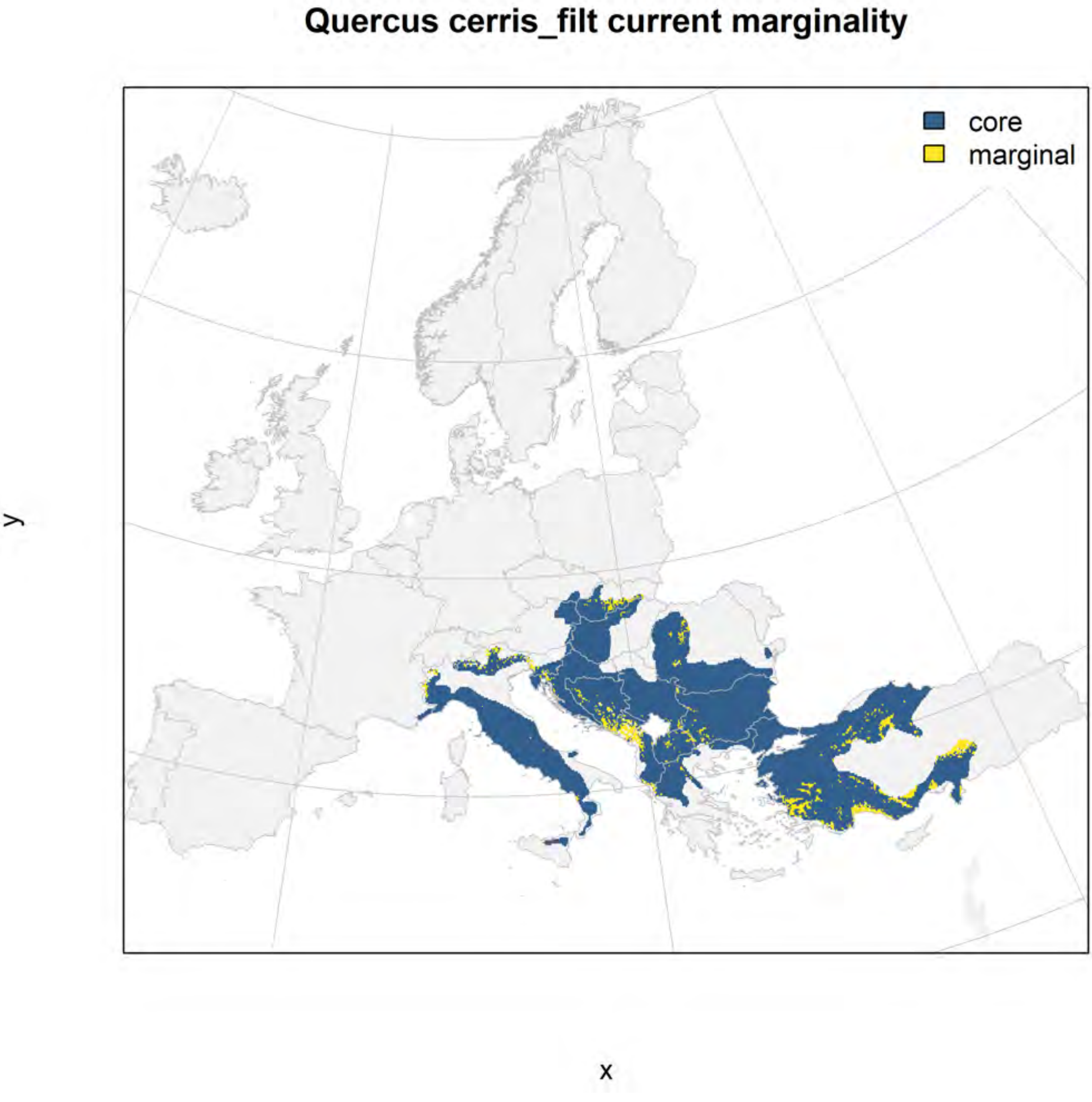
Quercus cerris_filt current marginality
core
marginal
y
x

### Quercus cerris_filt future marginality

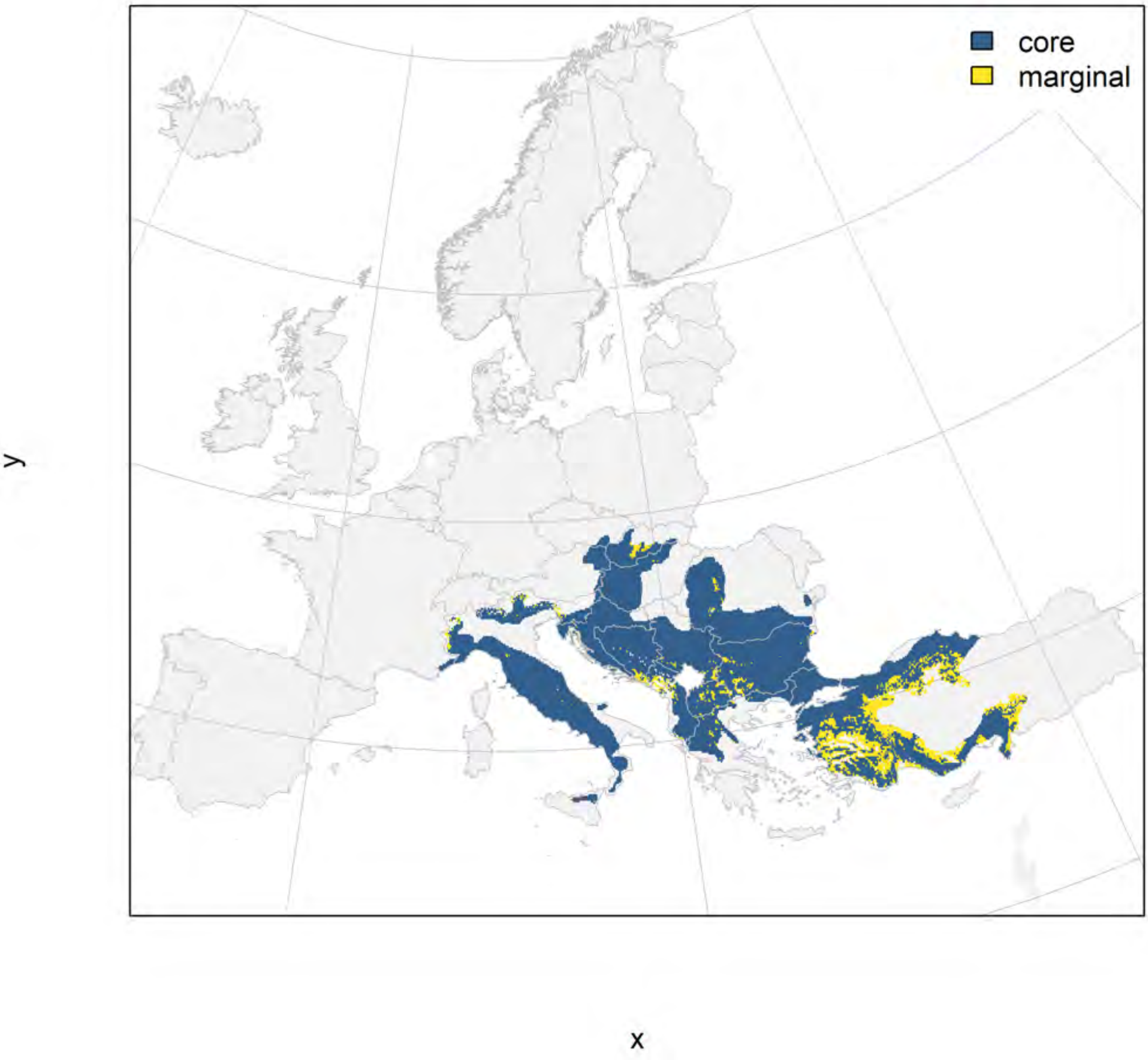

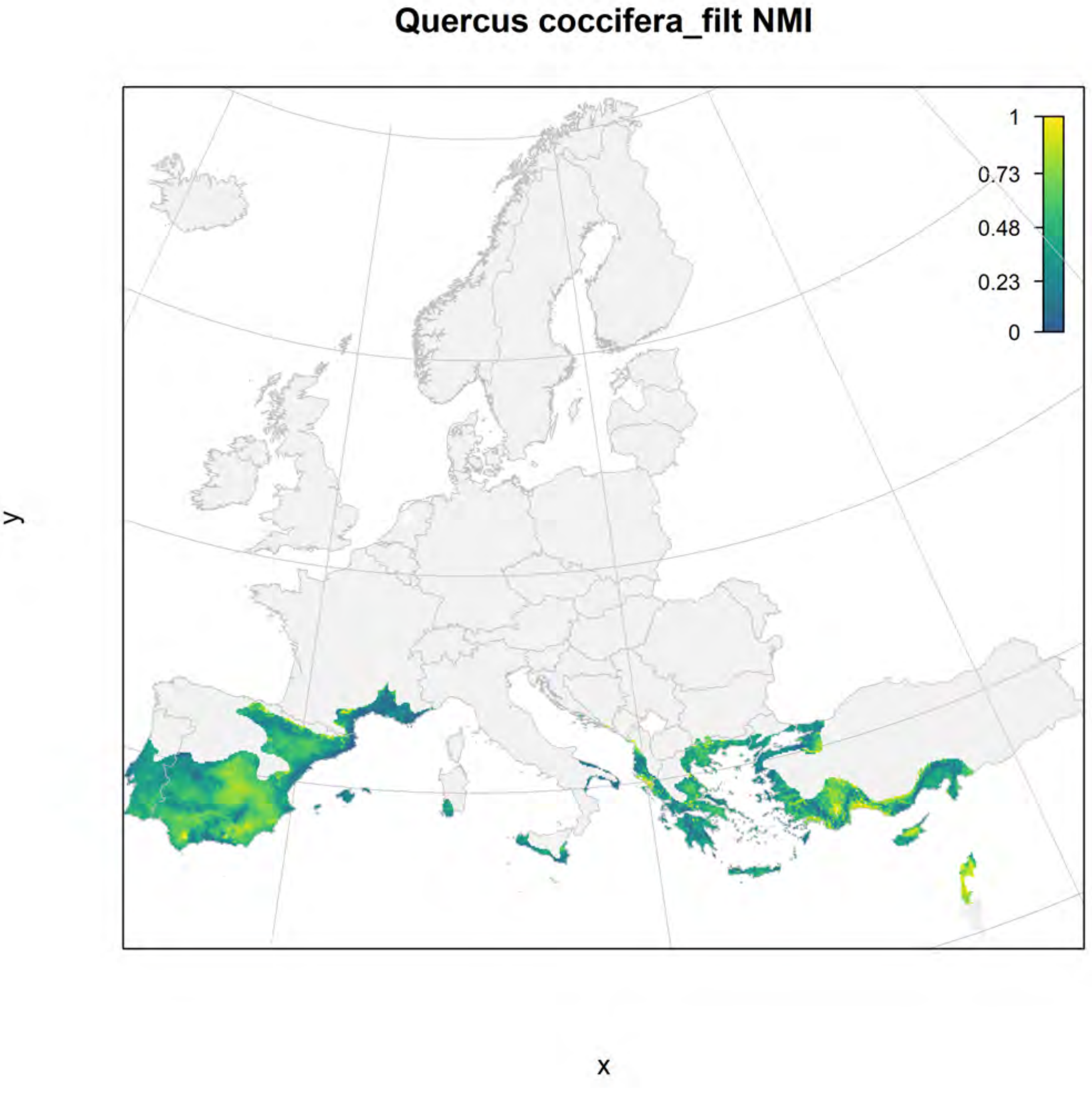
Quercus coccifera_filt NMI
1
0.73
0.48
0.23
0
y
x

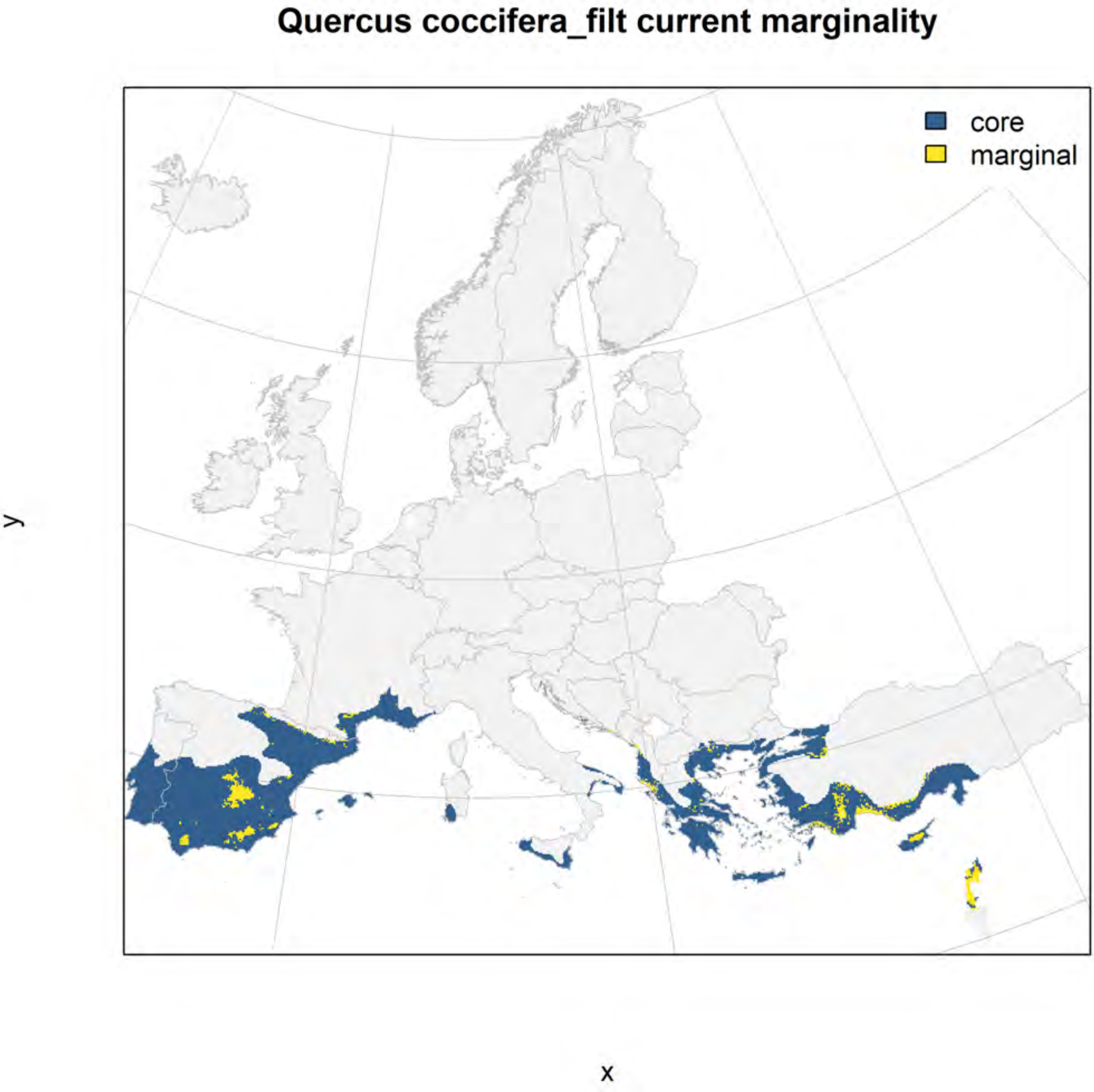
Quercus coccifera_filt current marginality
core
marginal
y
x

#### Quercus coccifera_filt future marginality

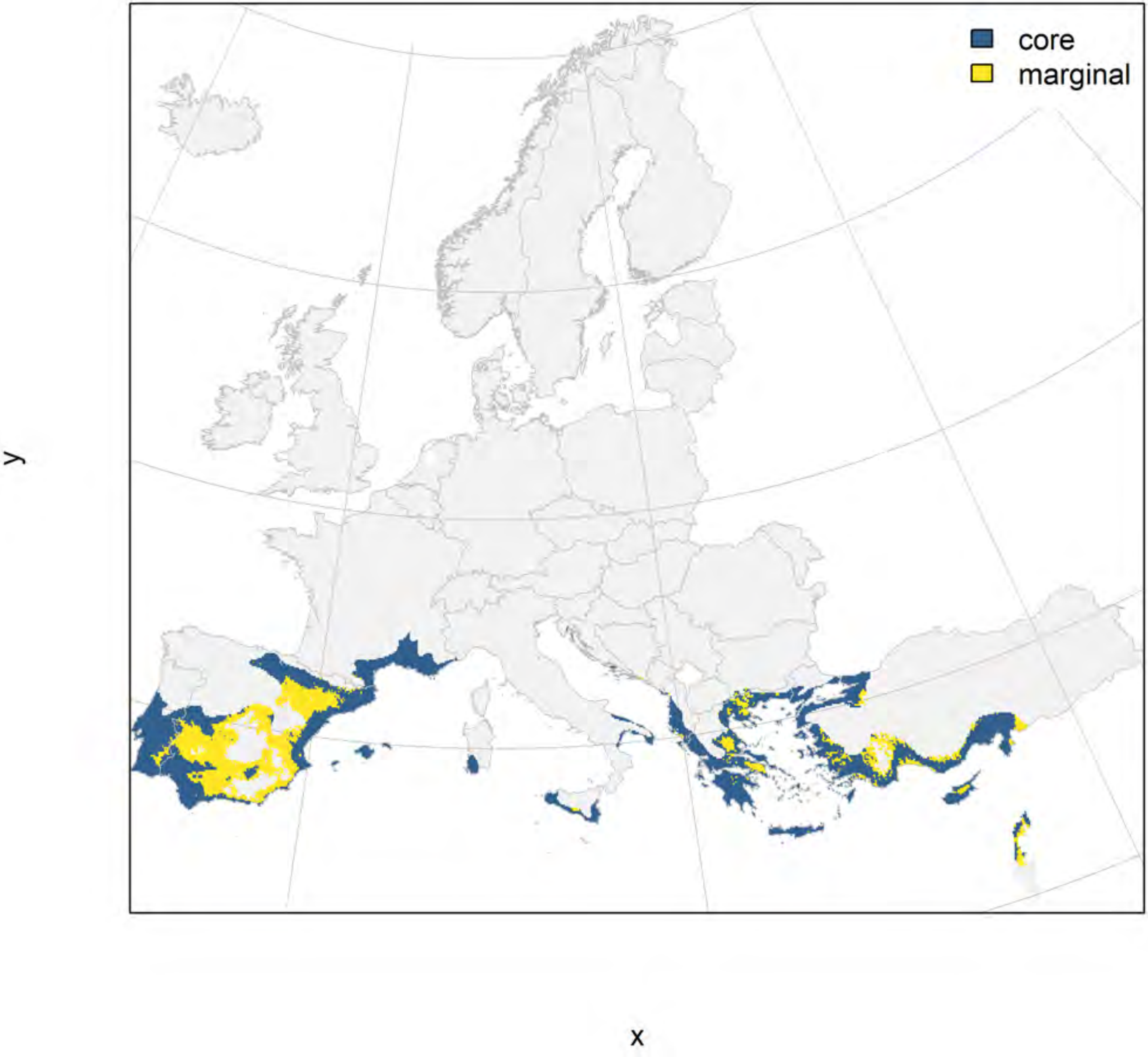

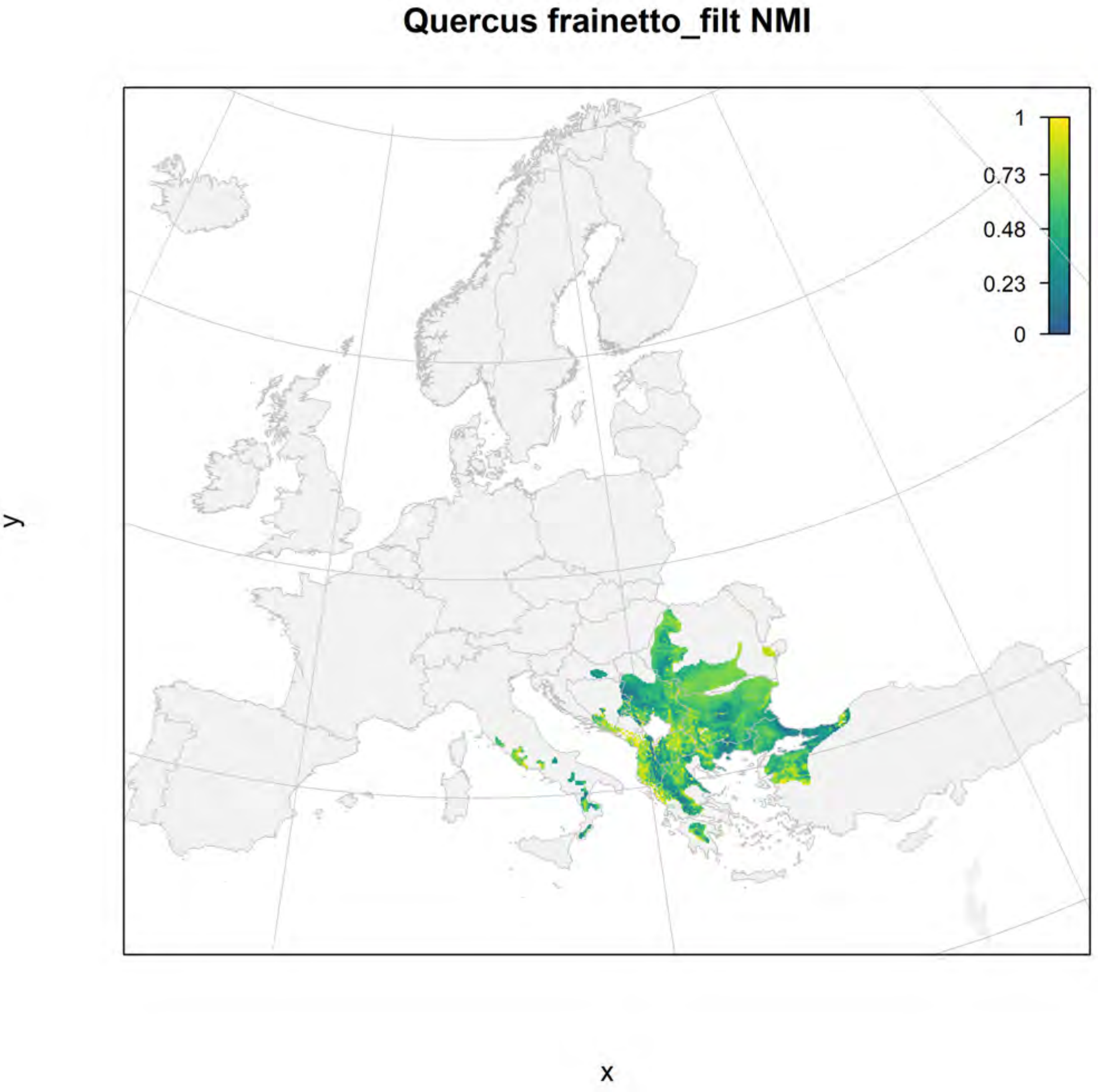
Quercus frainetto_filt NMI
1
0.73
0.48
0.23
0
y
x

### Quercus frainetto_filt current marginality

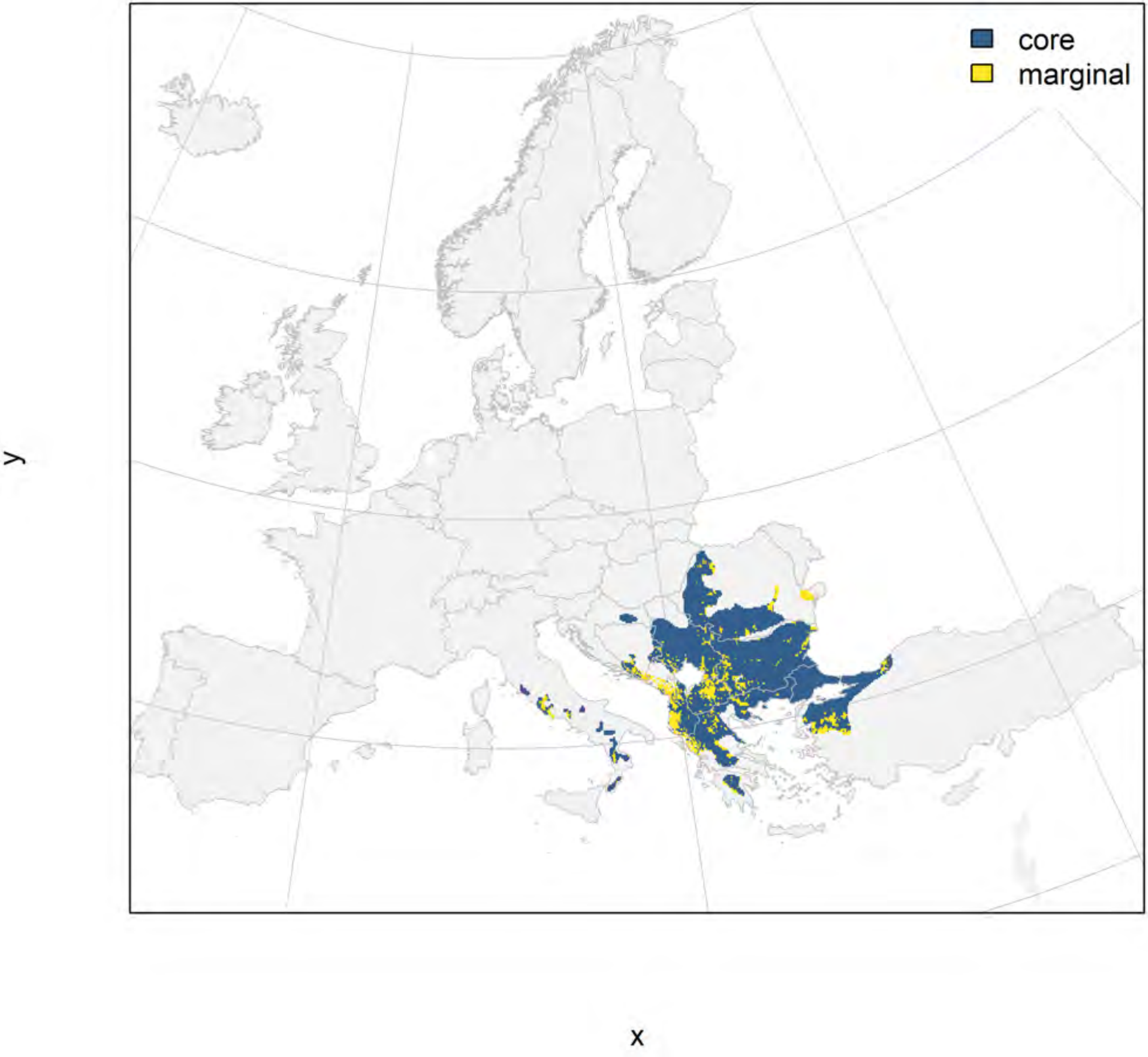

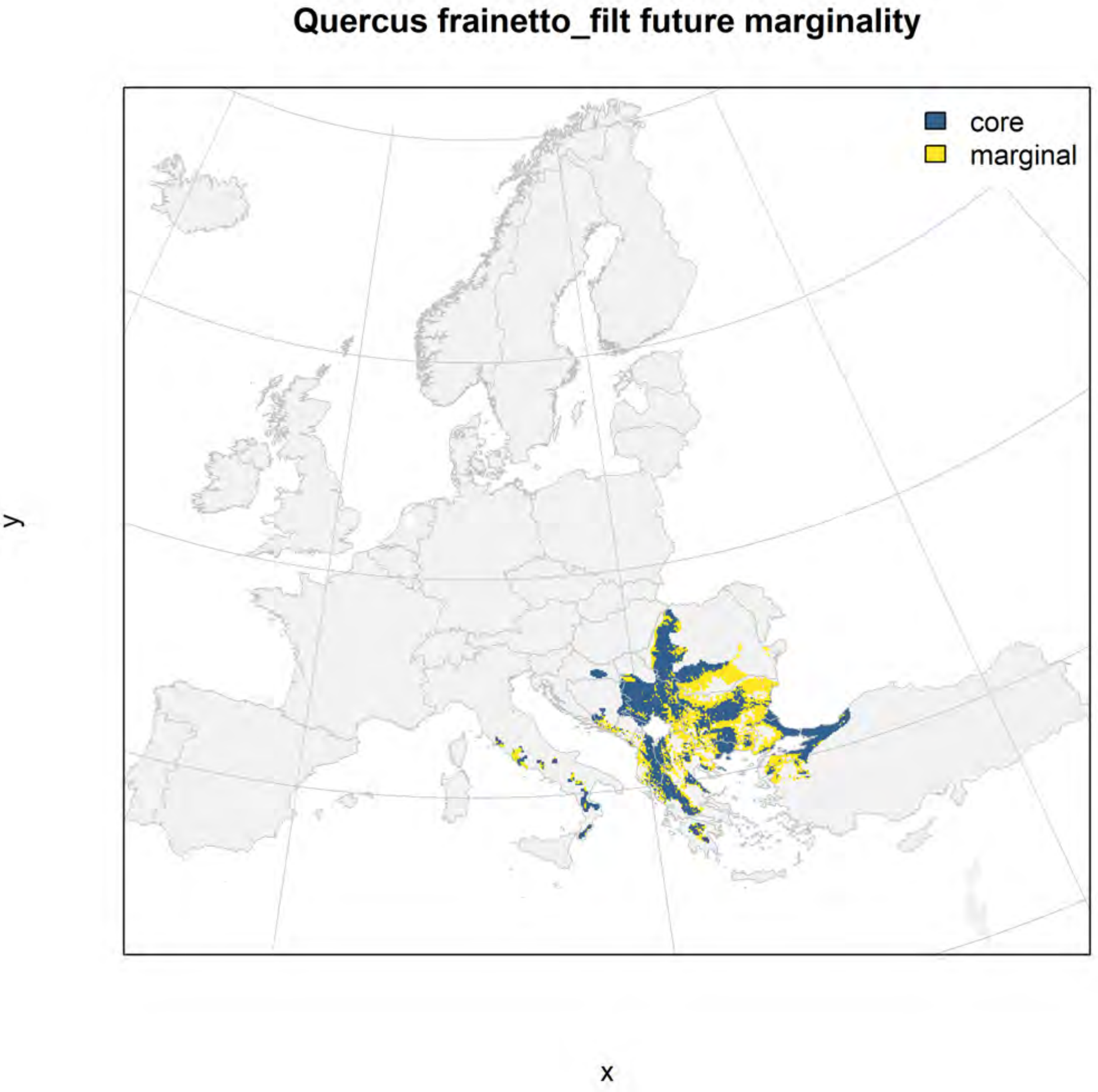
Quercus frainetto_filt future marginality
core
marginal
y
x

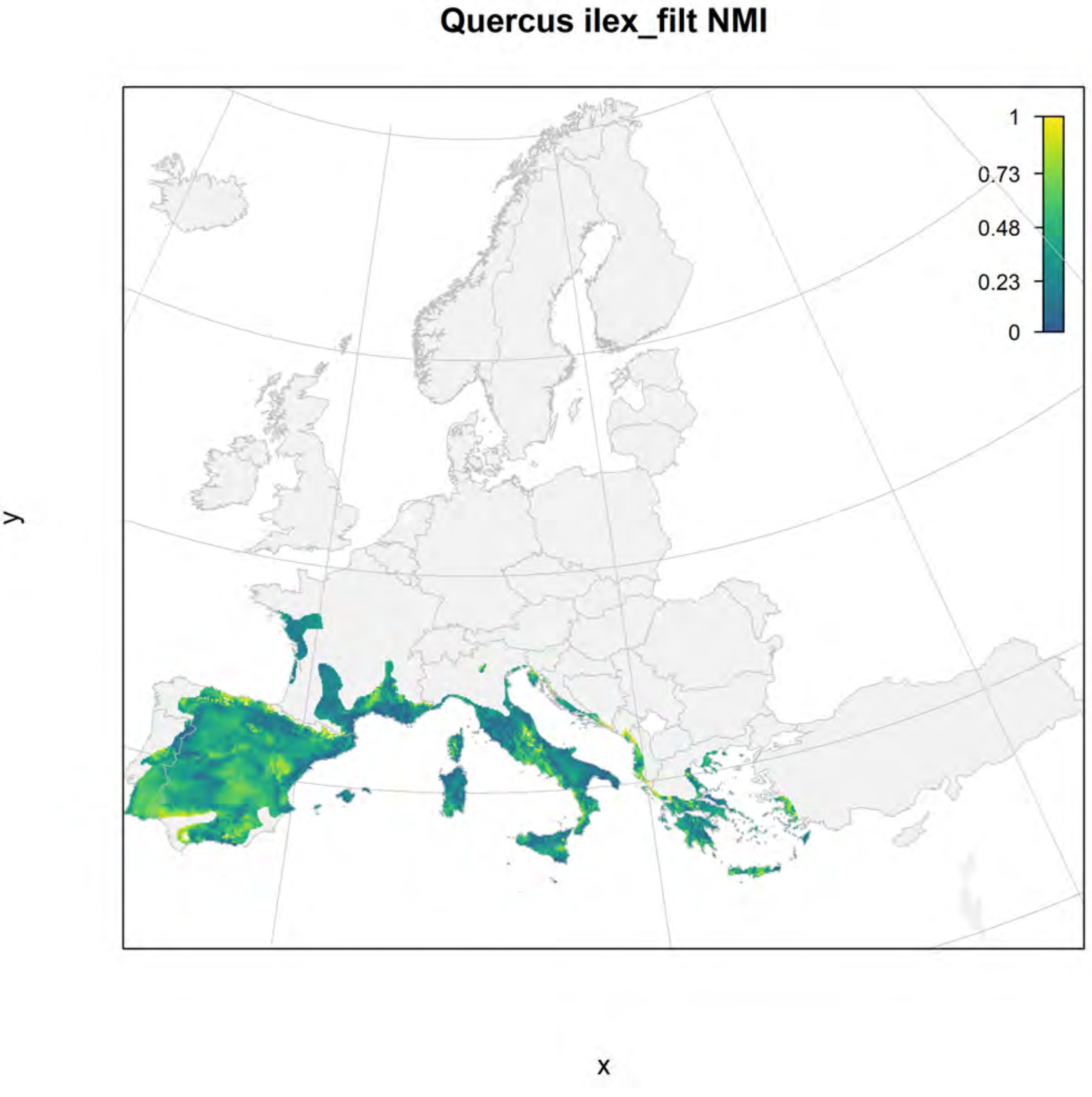

Quercus ilex_filt NMI
1
0.73
0.48
0.23
0
y
x

#### Quercus ilex_filt current marginality

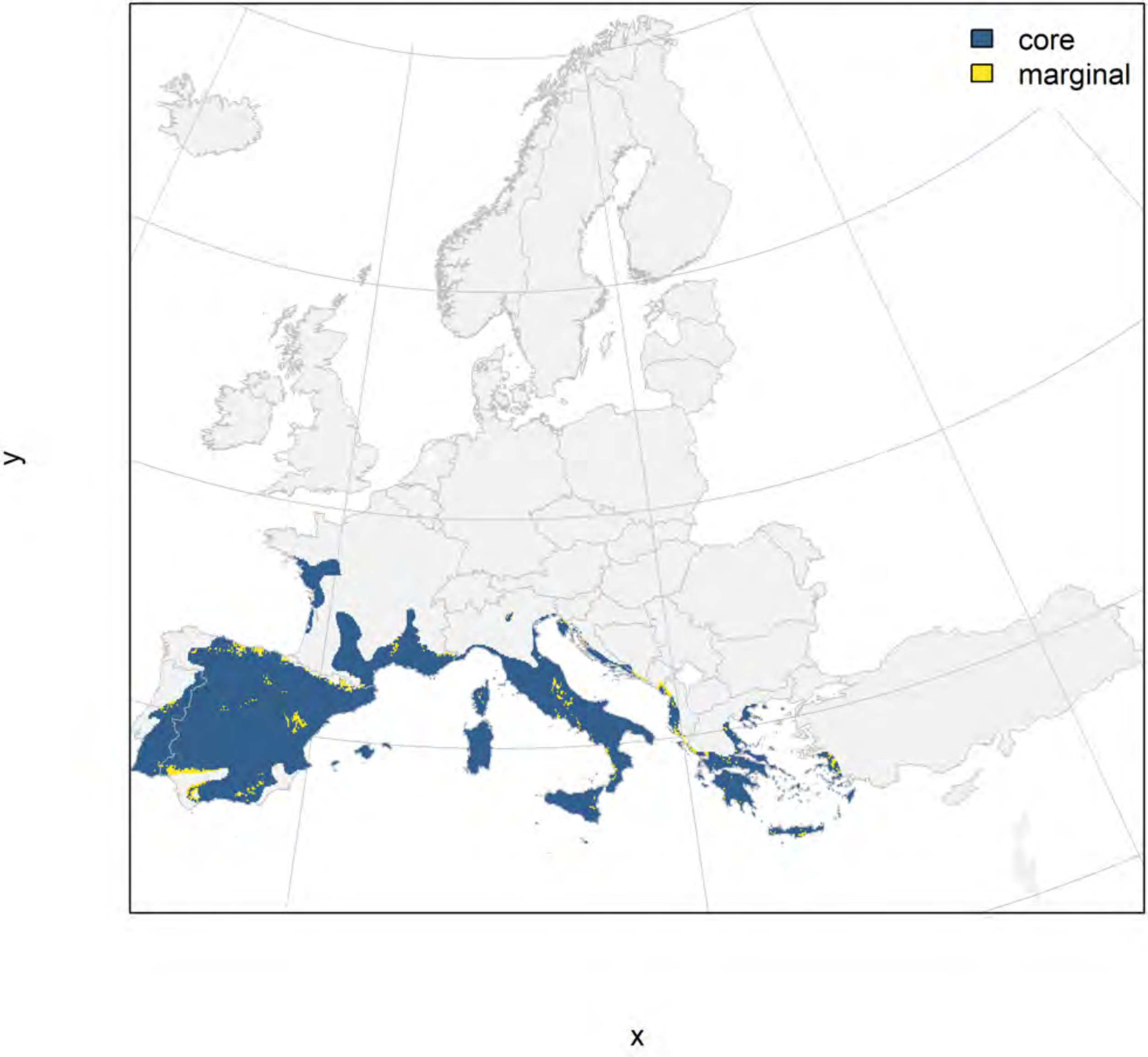

#### Quercus ilex_filt future marginality

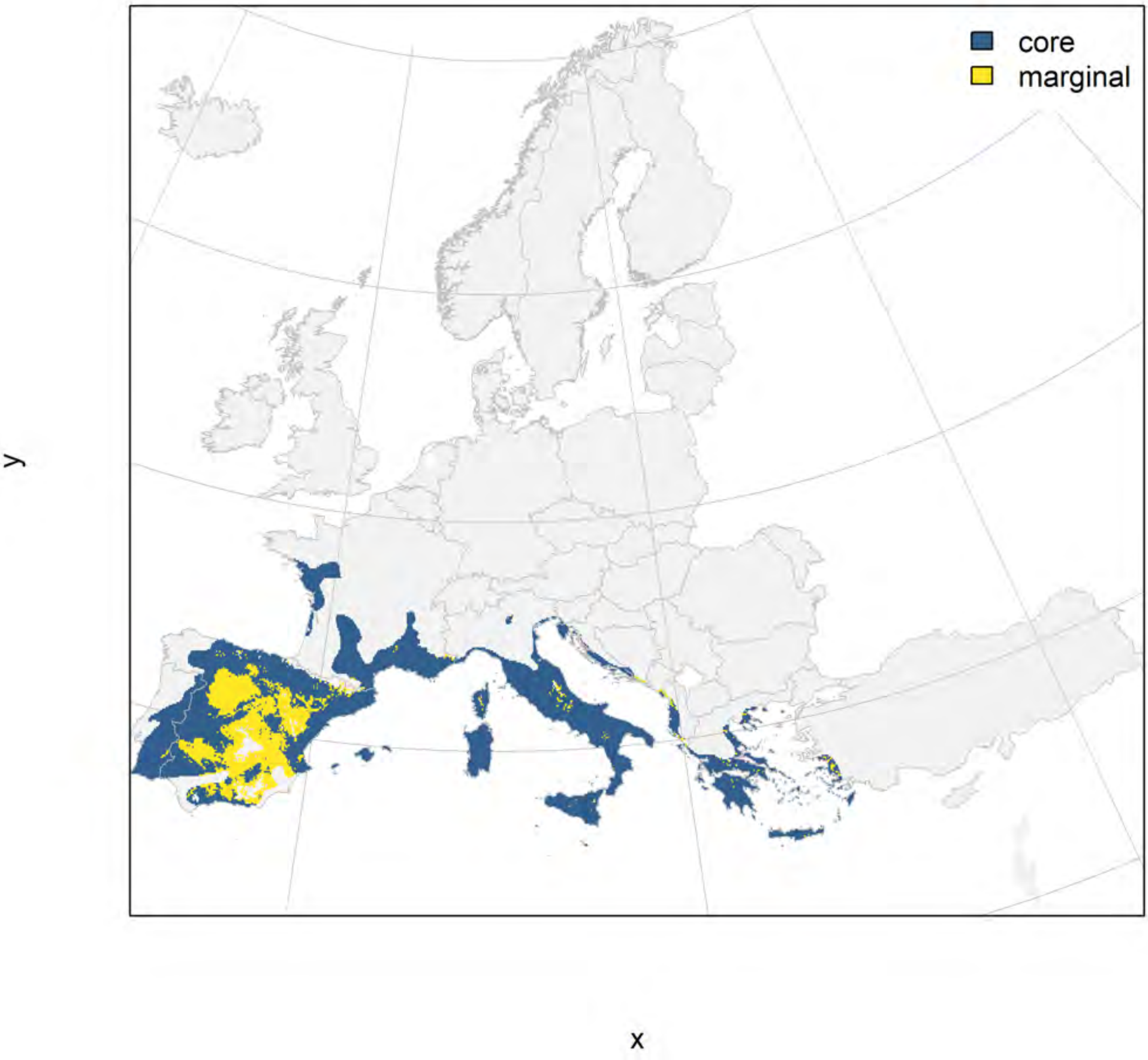

### Quercus petraea_filt NMI

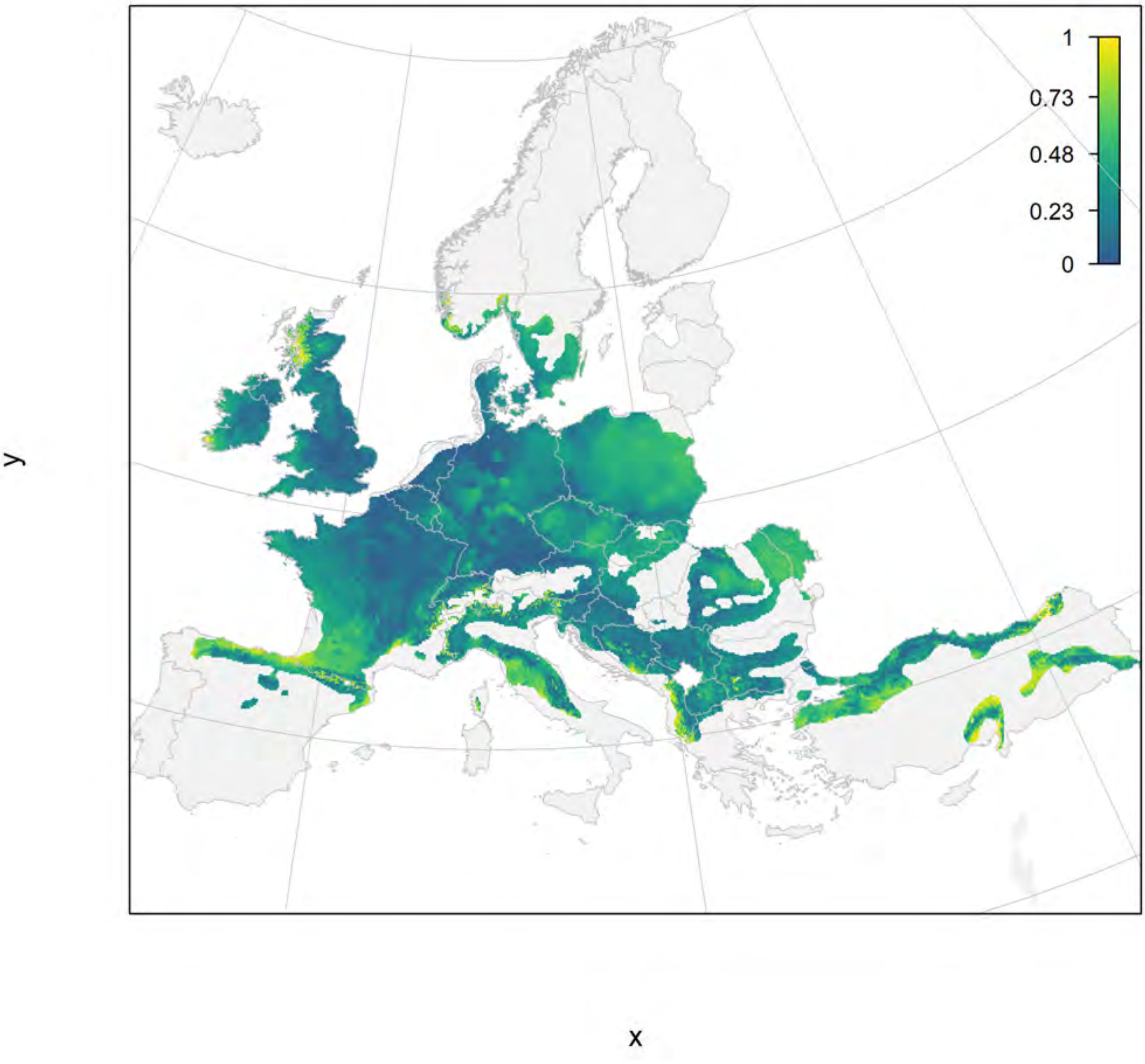

#### Quercus petraea_filt current marginality

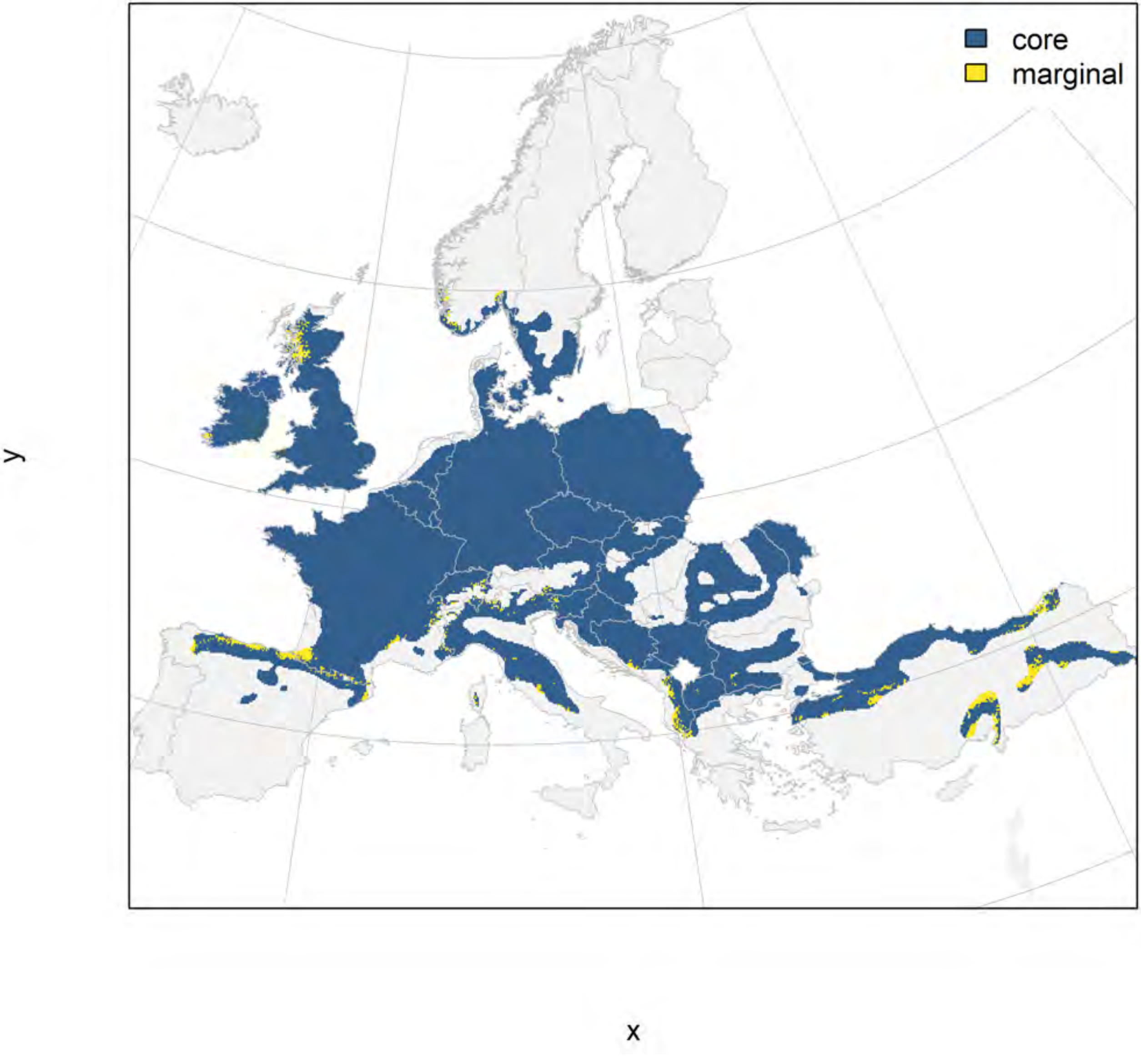

### Quercus petraea_filt future marginality

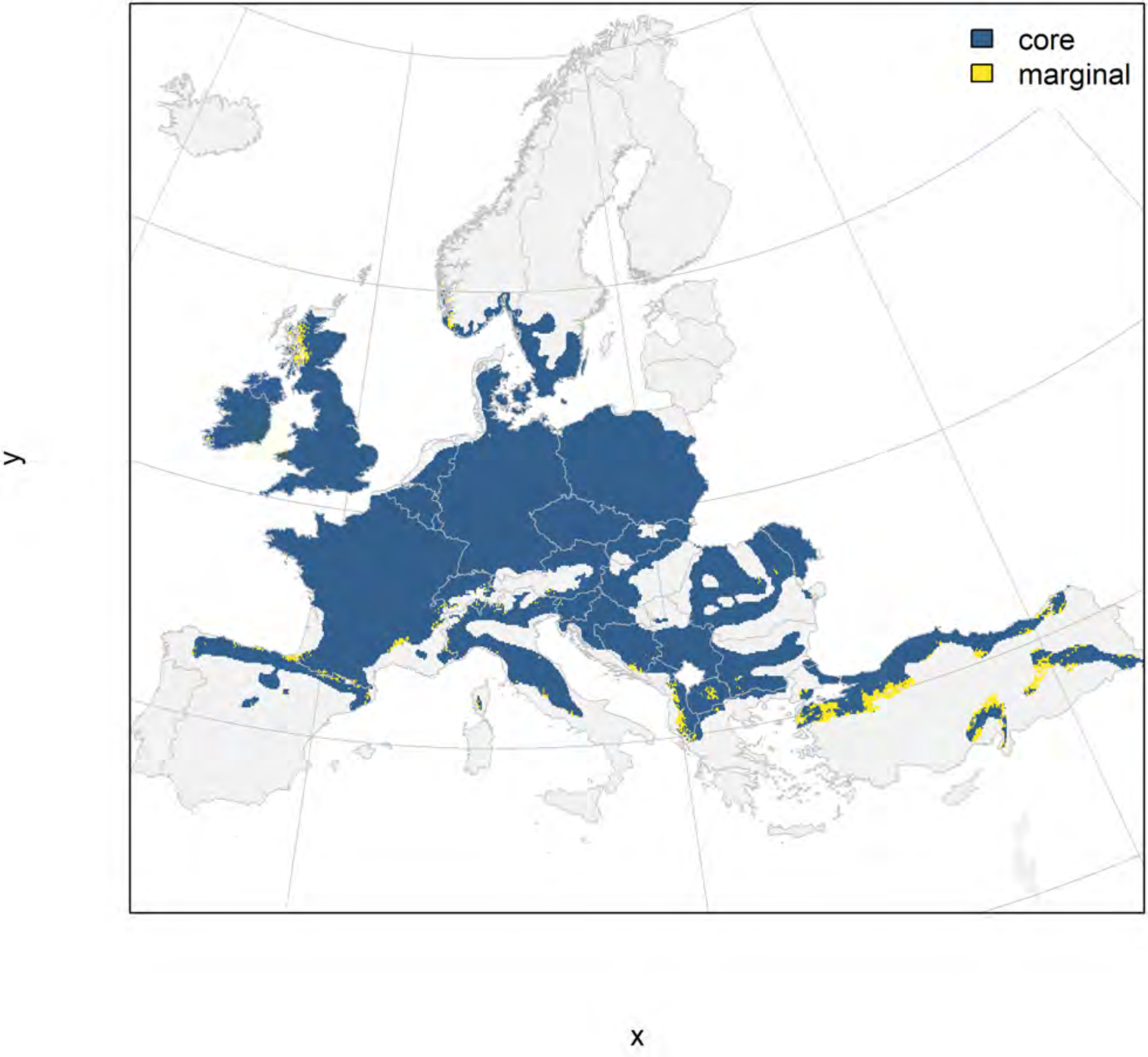

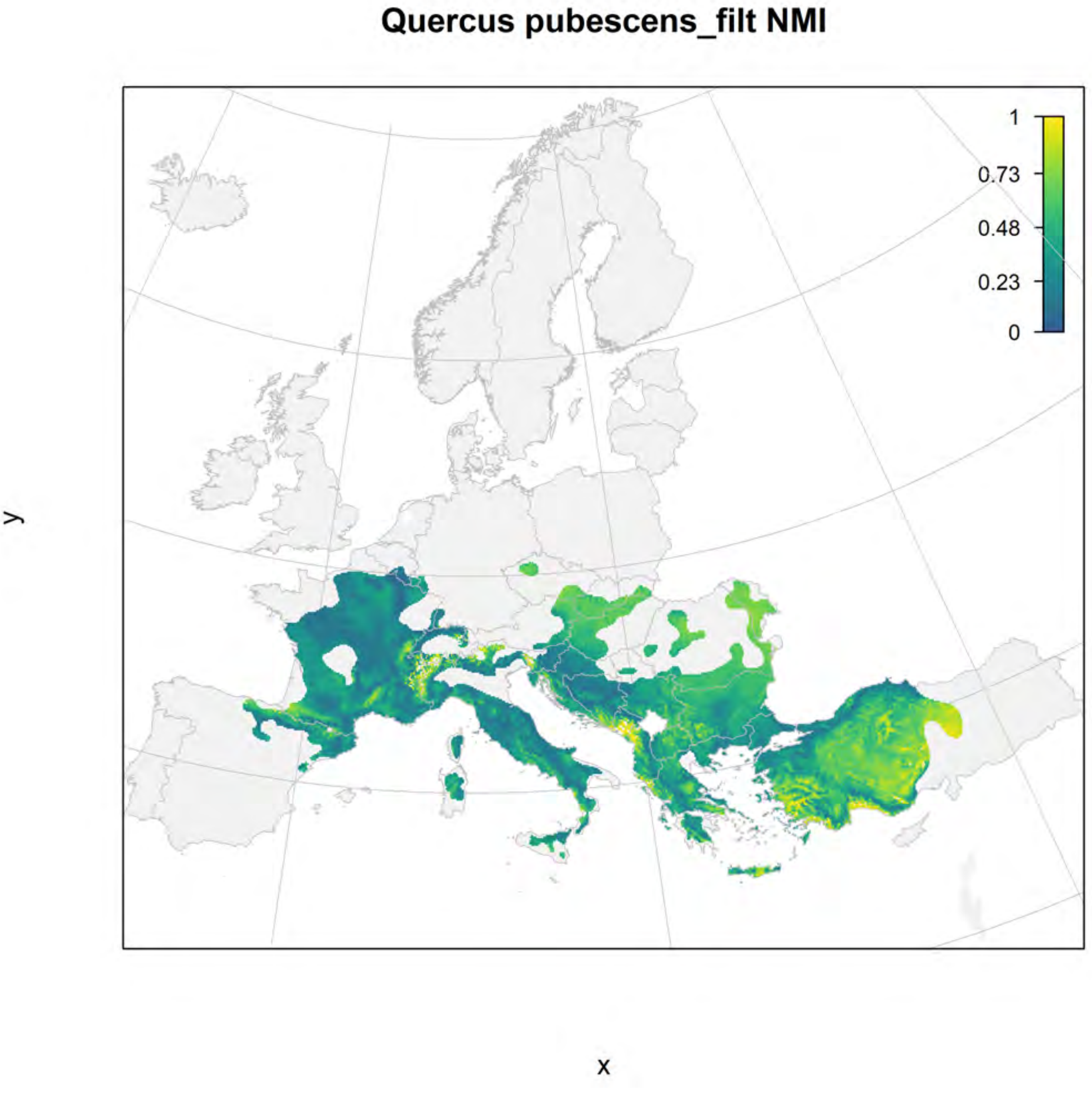
Quercus pubescens_filt NMI
1
0.73
0.48
0.23
0
y
x

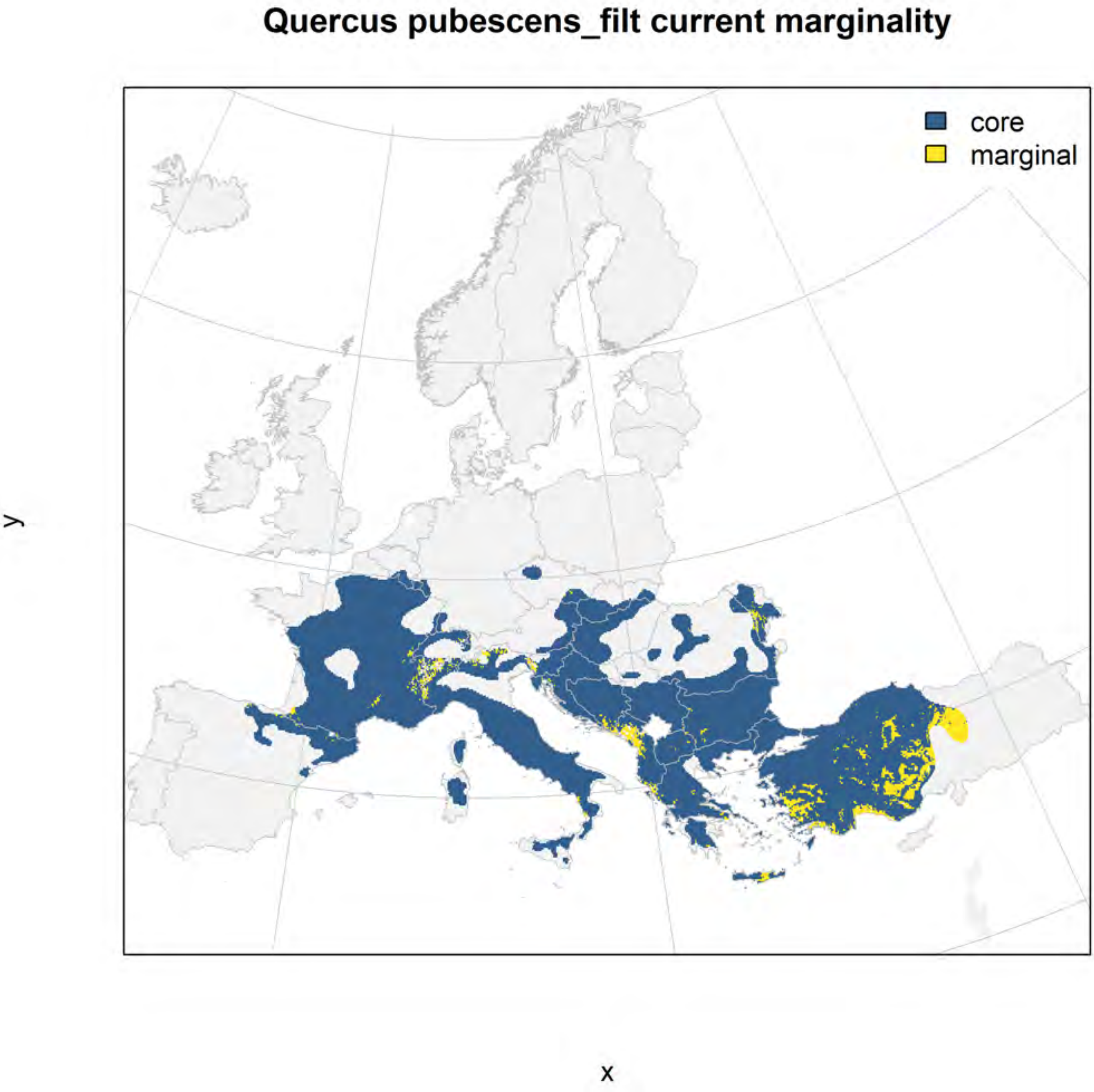
Quercus pubescens_filt current marginality
core
marginal
y
x

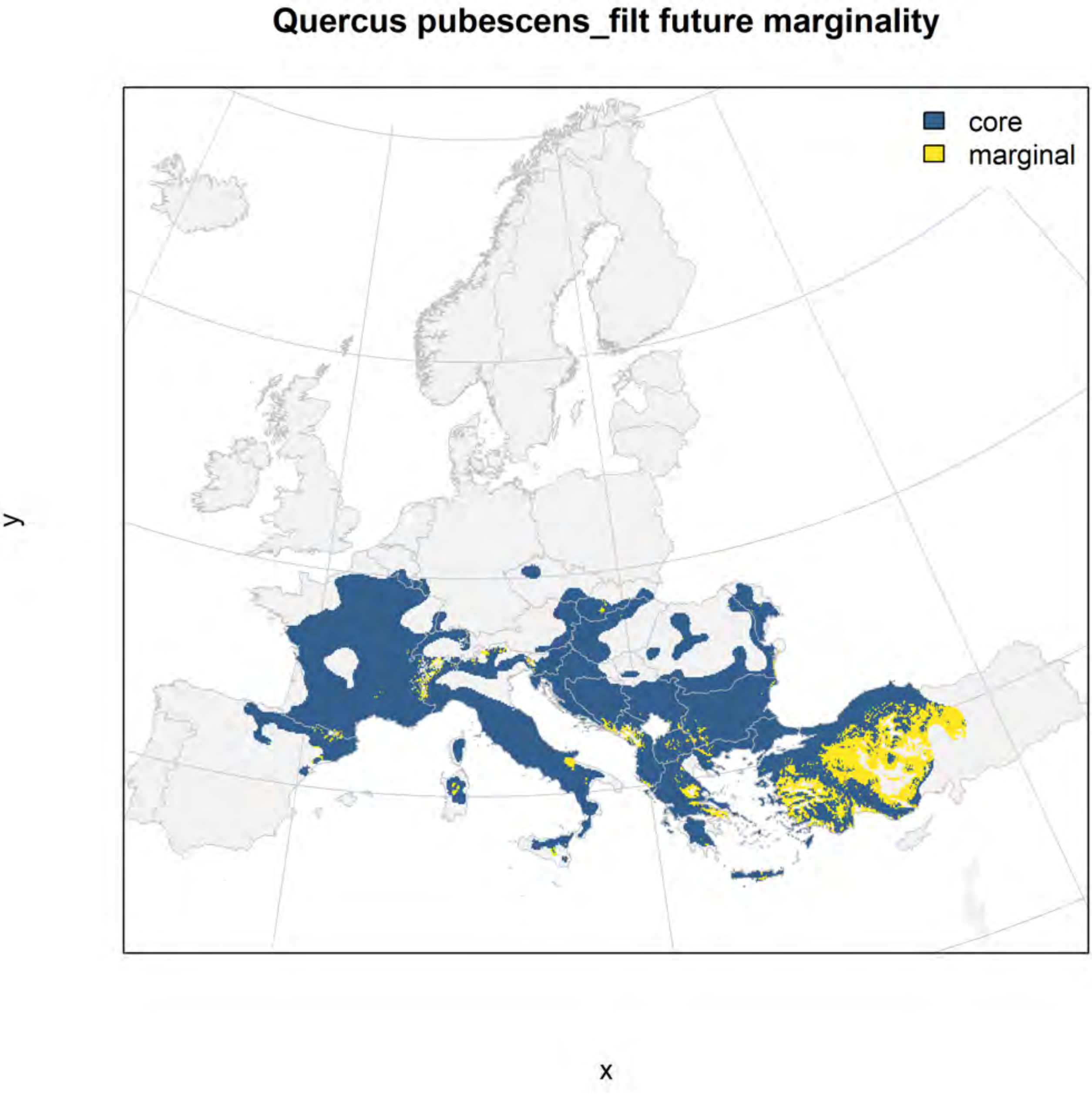
Quercus pubescens_filt future marginality
core
marginal
y
x

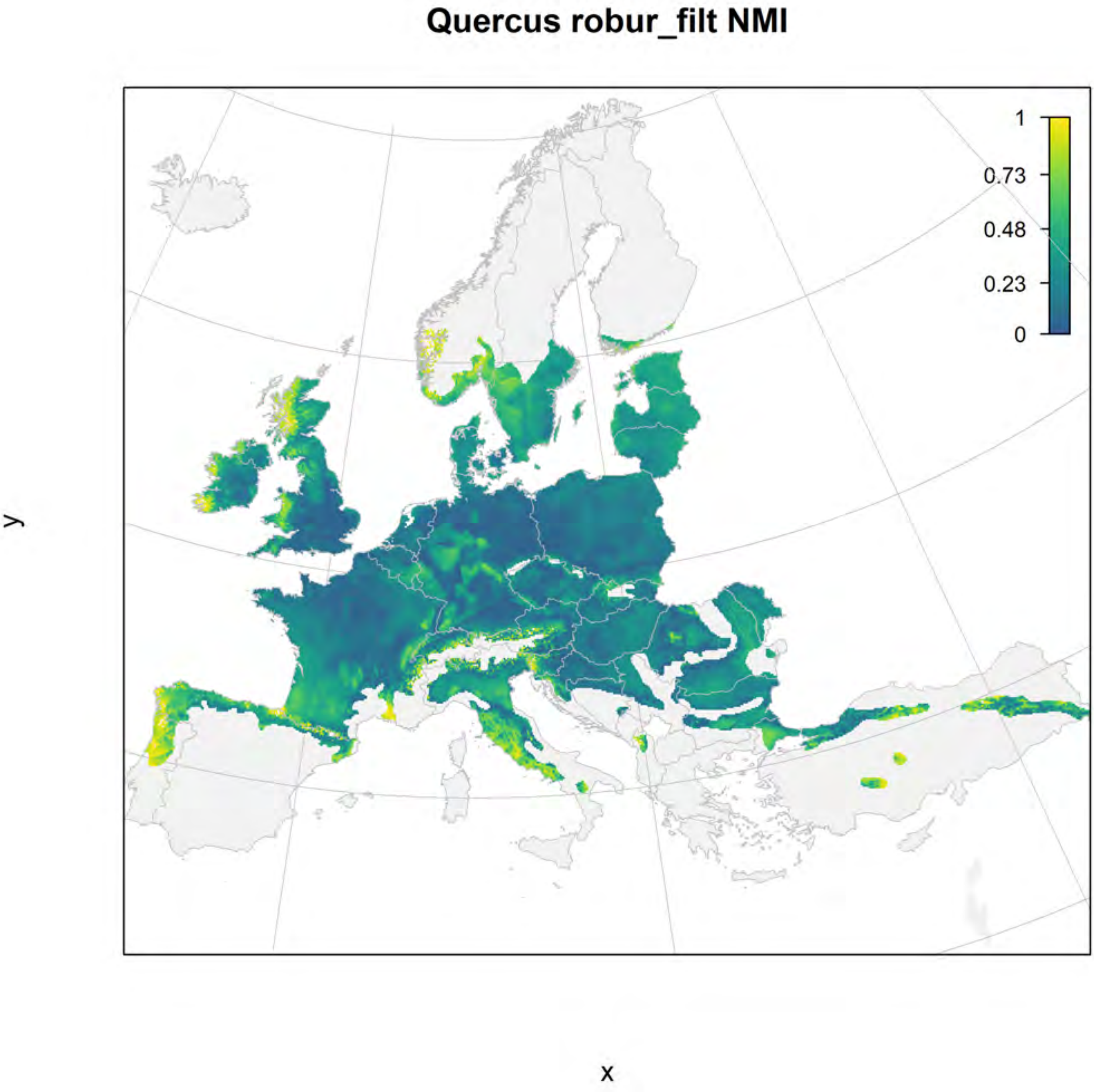
Quercus robur_filt NMI
1
0.73
0.48
0.23
0
y
x

#### Quercus robur_filt current marginality

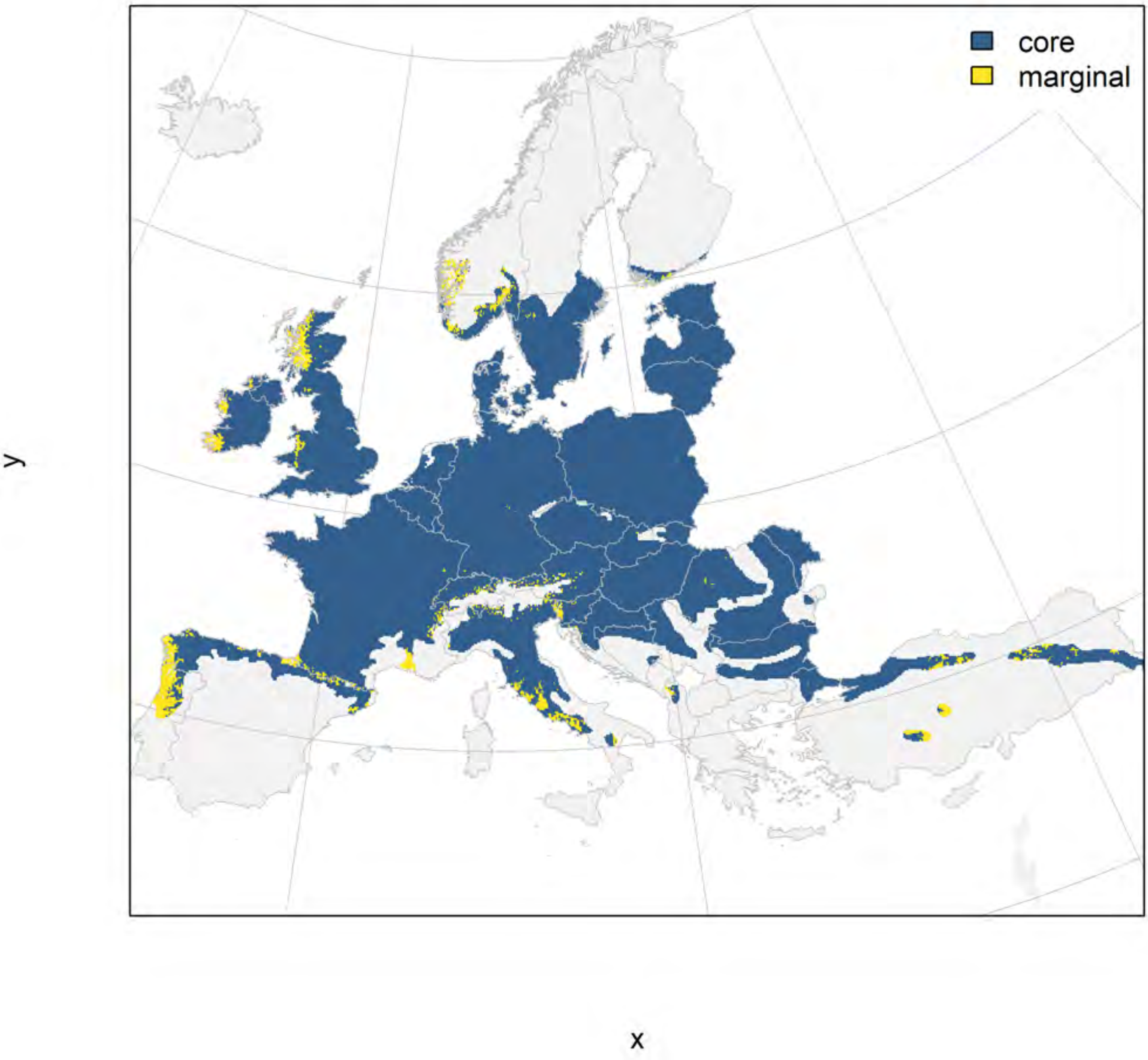

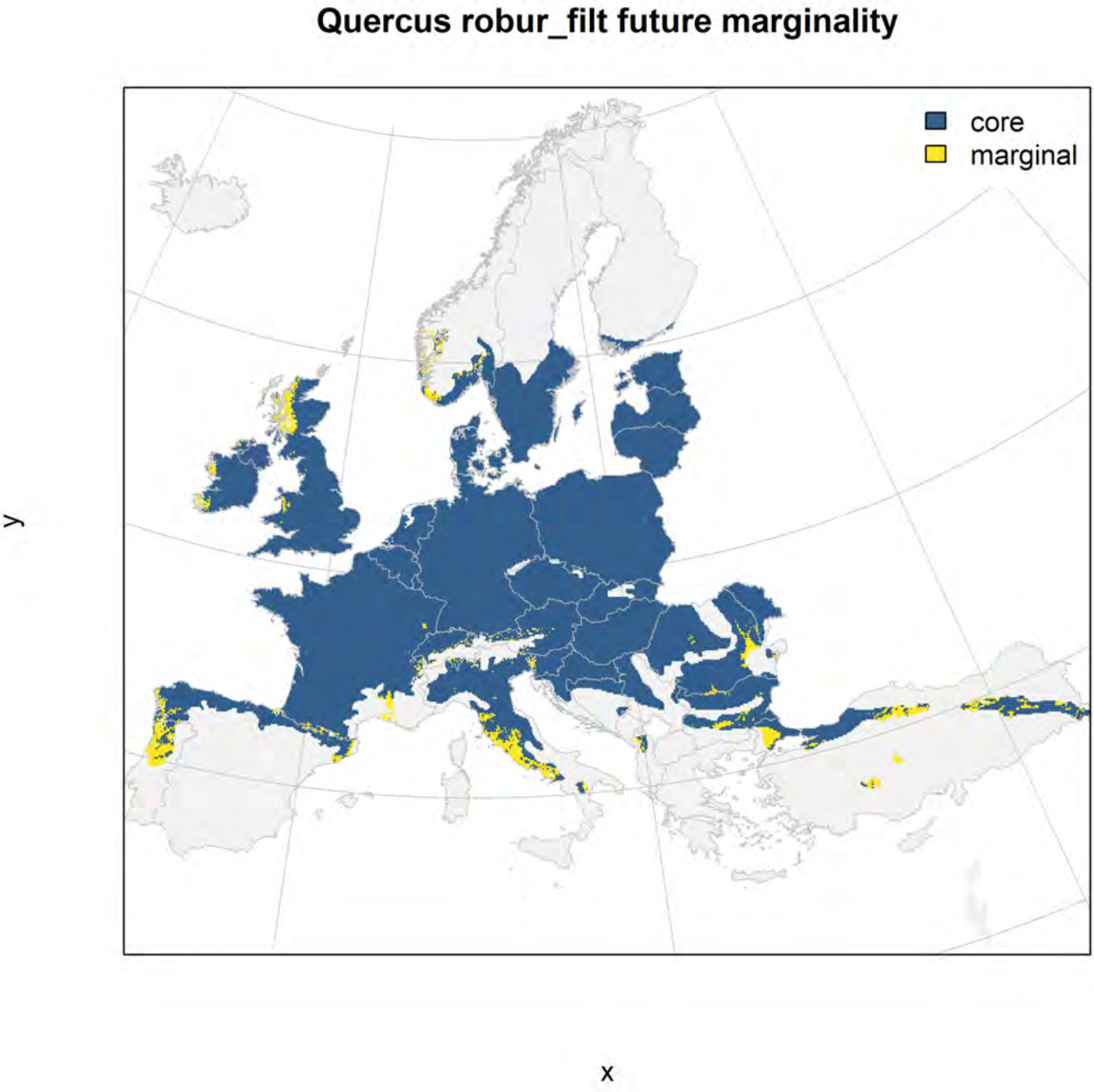

Quercus robur_filt future marginality
core
marginal
y
x

### Quercus suber_filt NMI

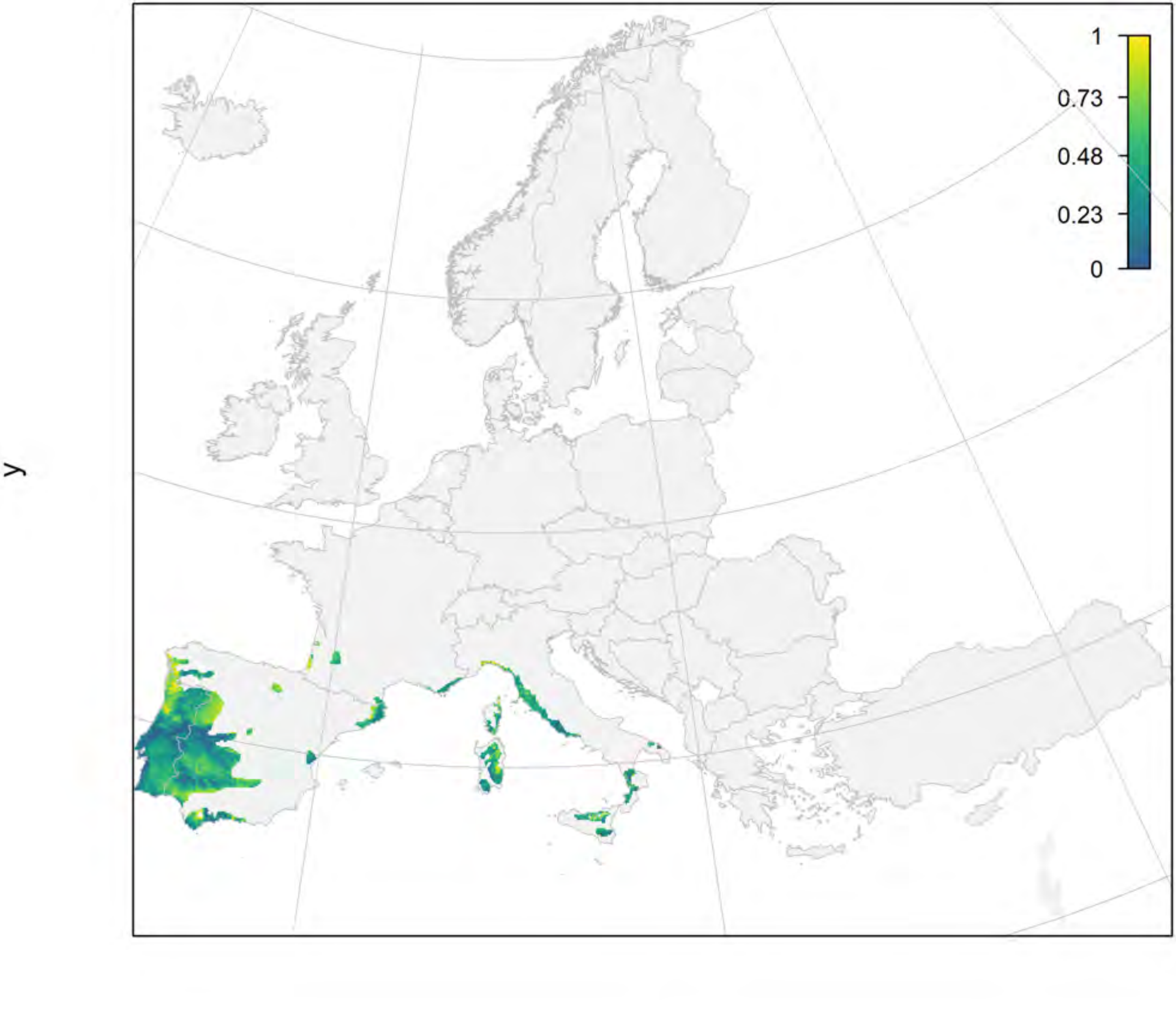

#### Quercus suber_filt current marginality

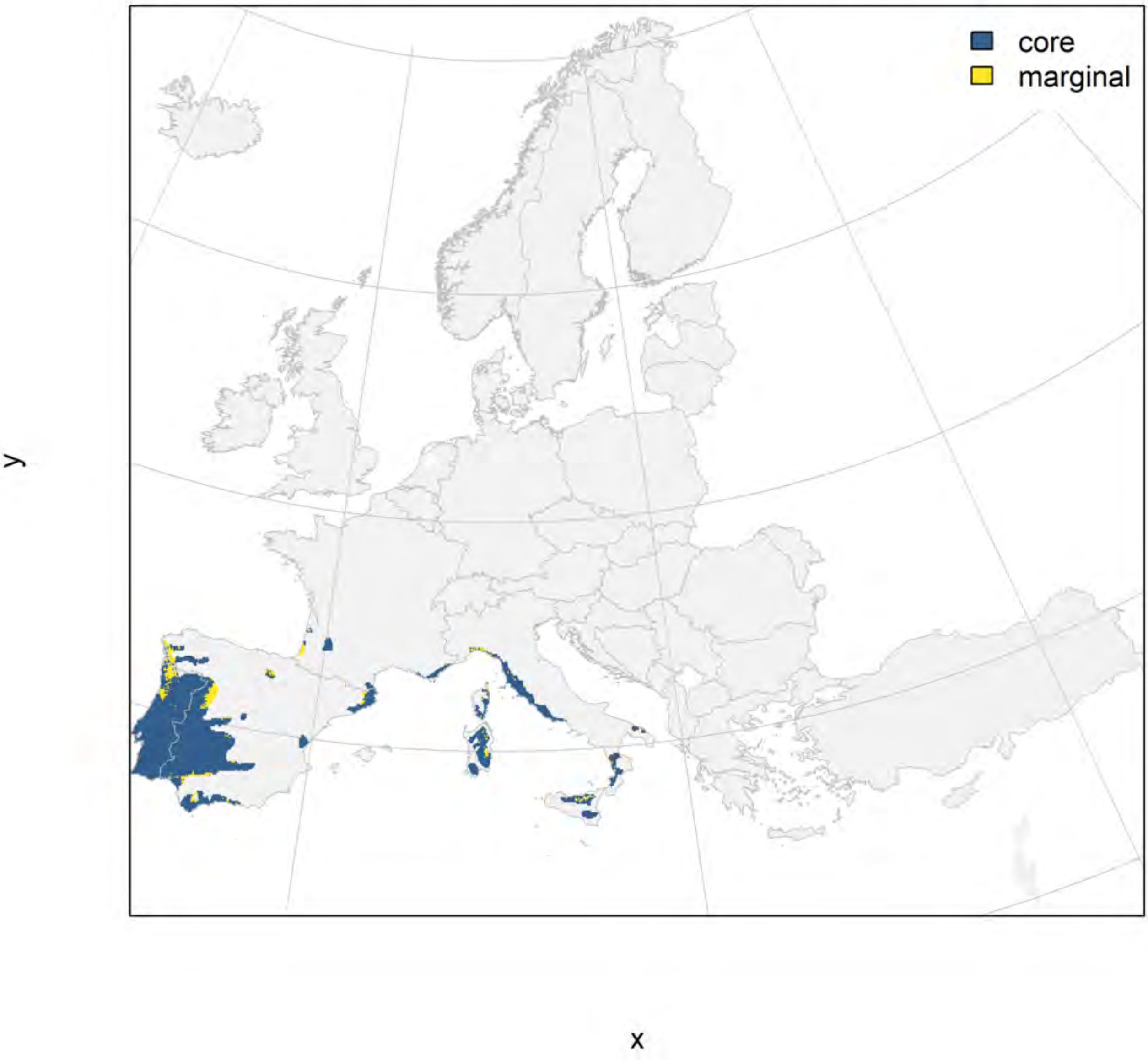

#### Quercus suber_filt future marginality

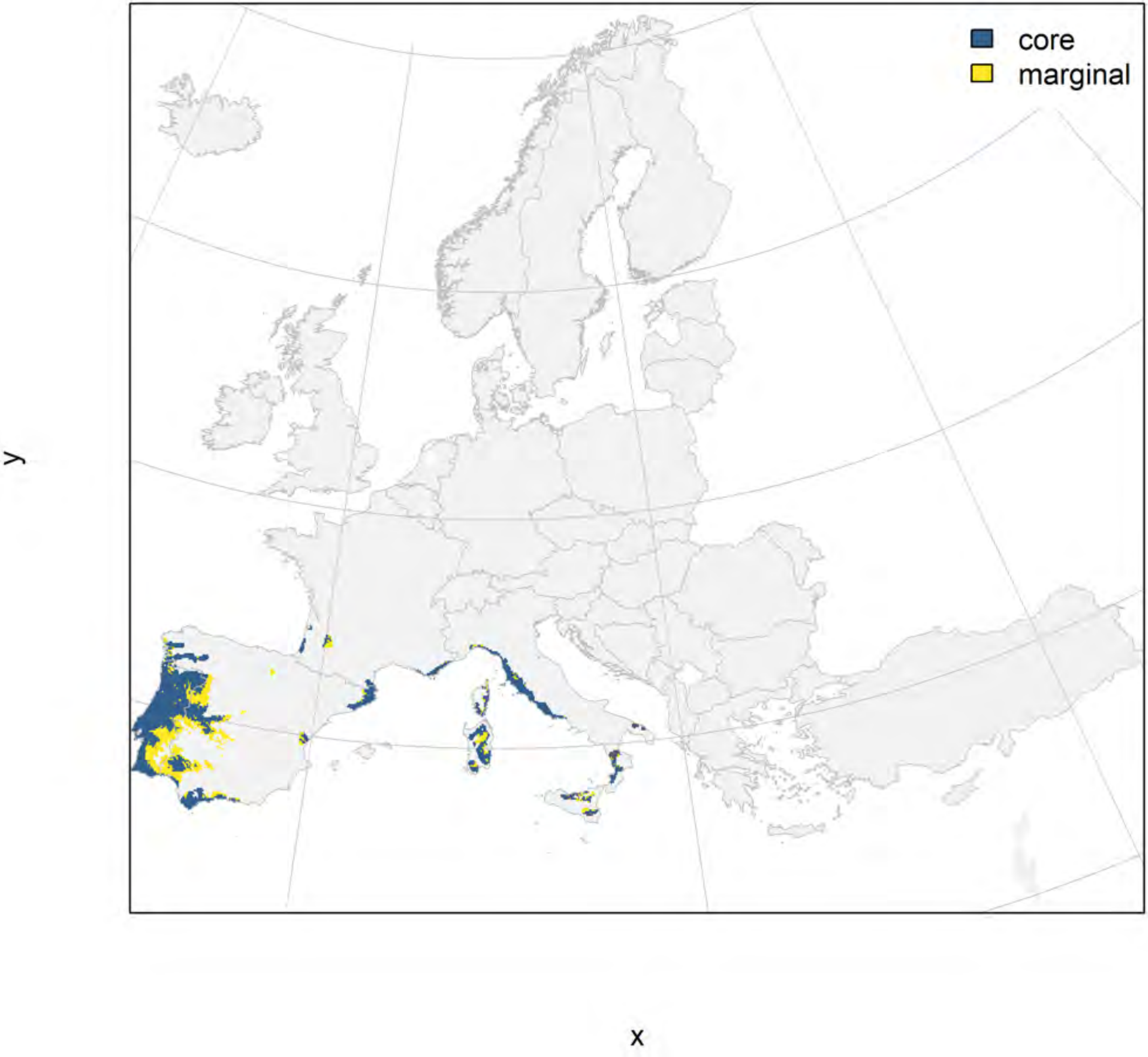

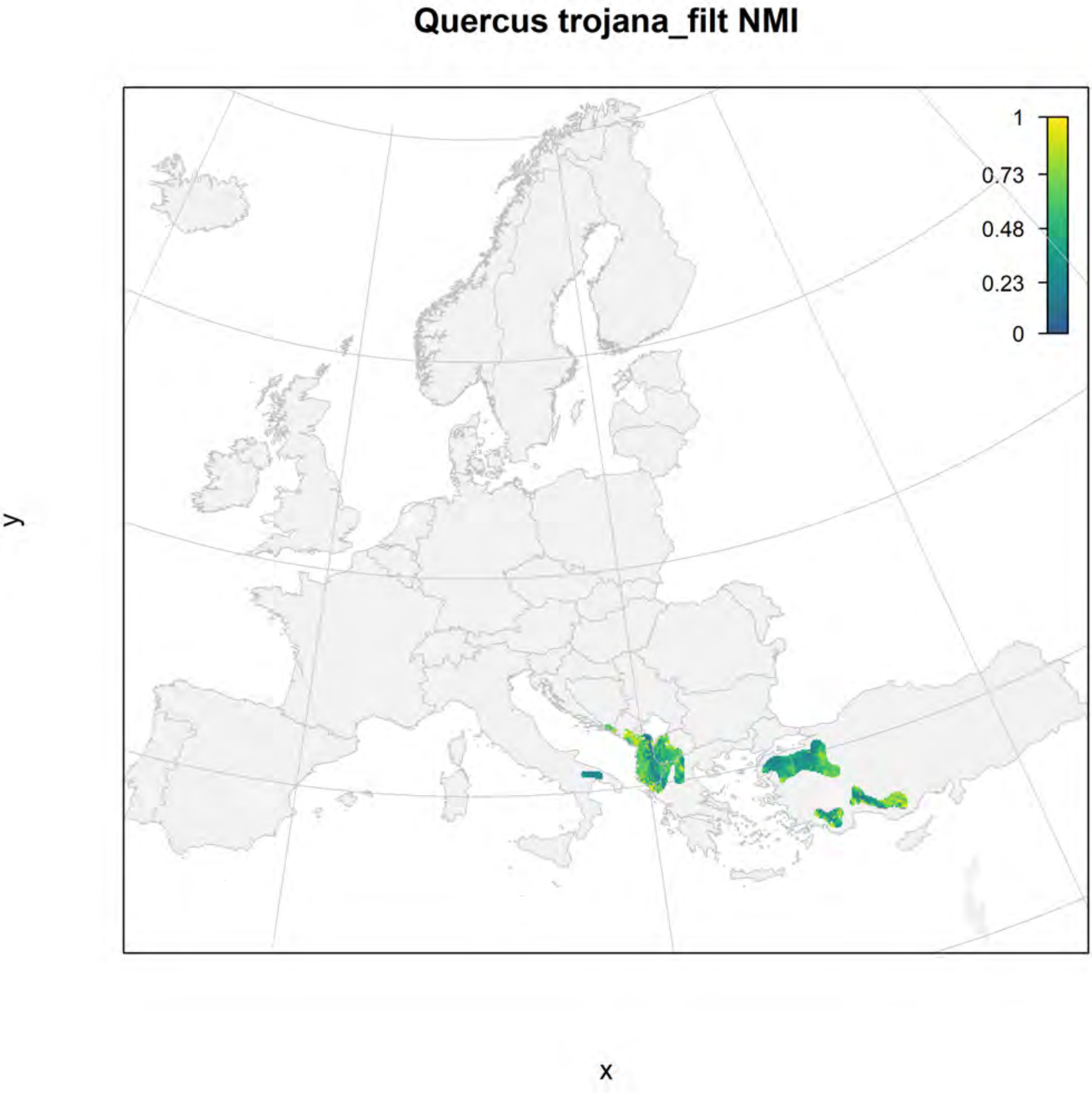
Quercus trojana_filt NMI
1
0.73
0.48
0.23
0
y
x

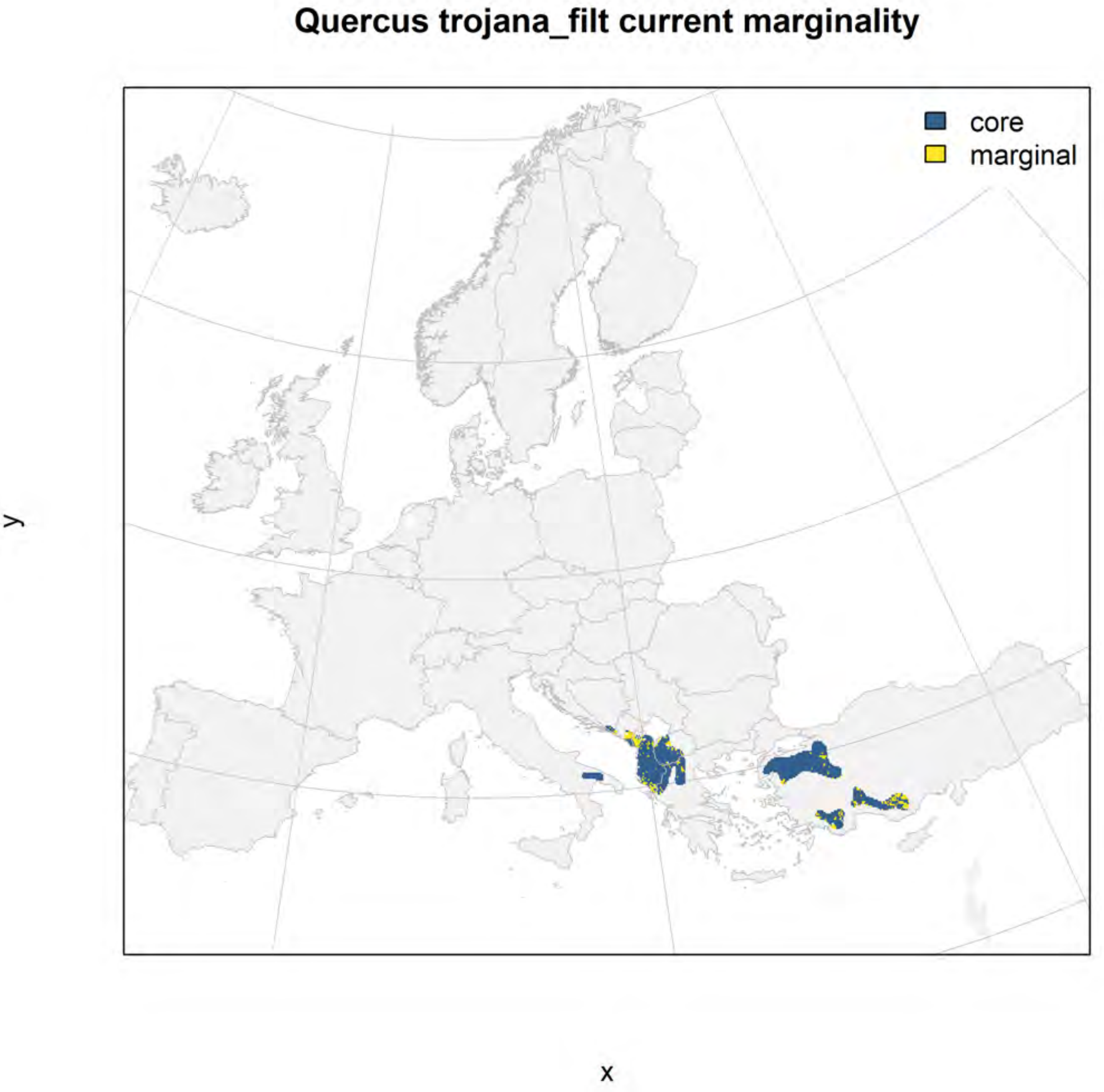
Quercus trojana_filt current marginality
core
marginal
y
x

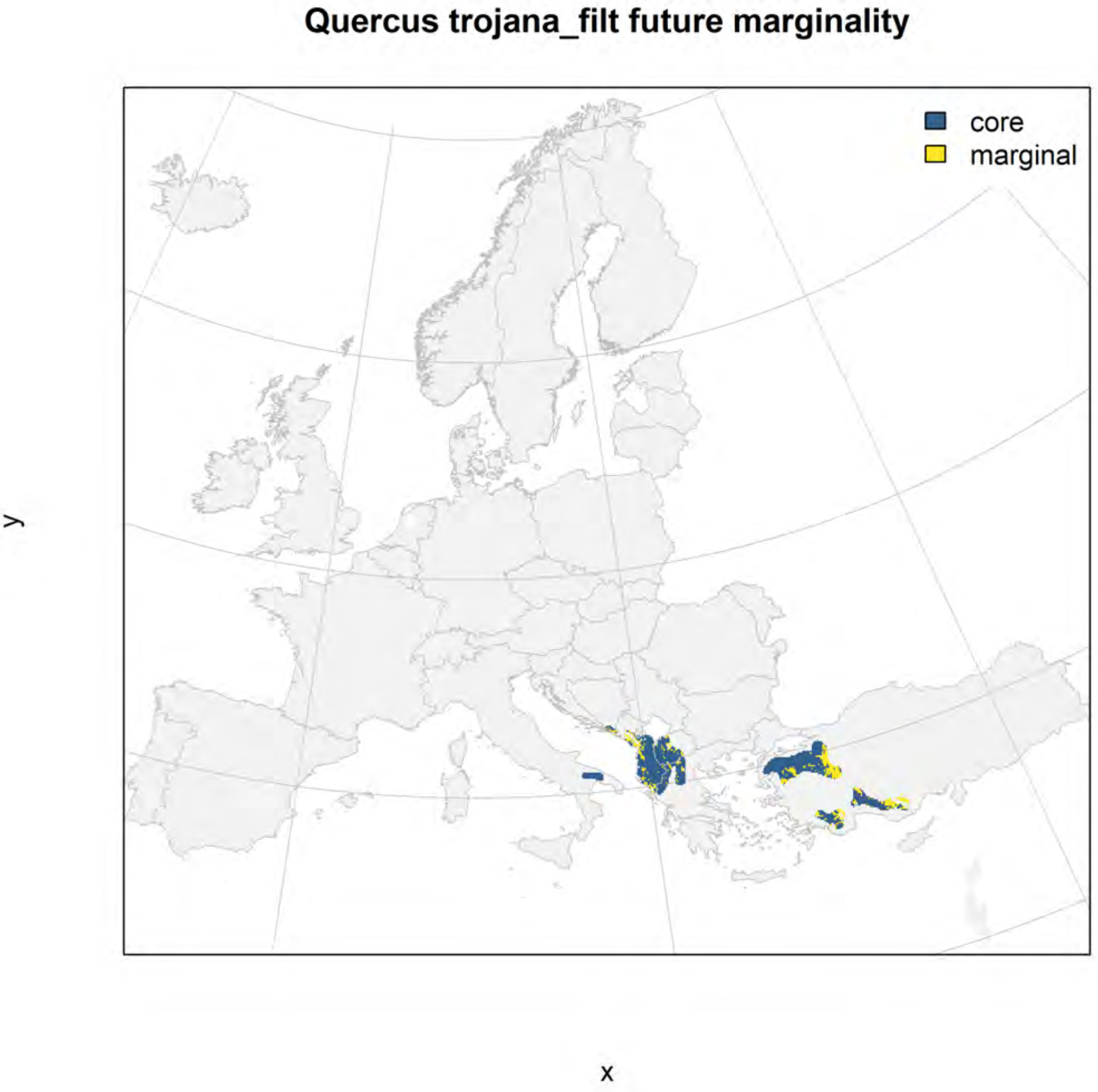
Quercus trojana_filt future marginality
core
marginal
y
x

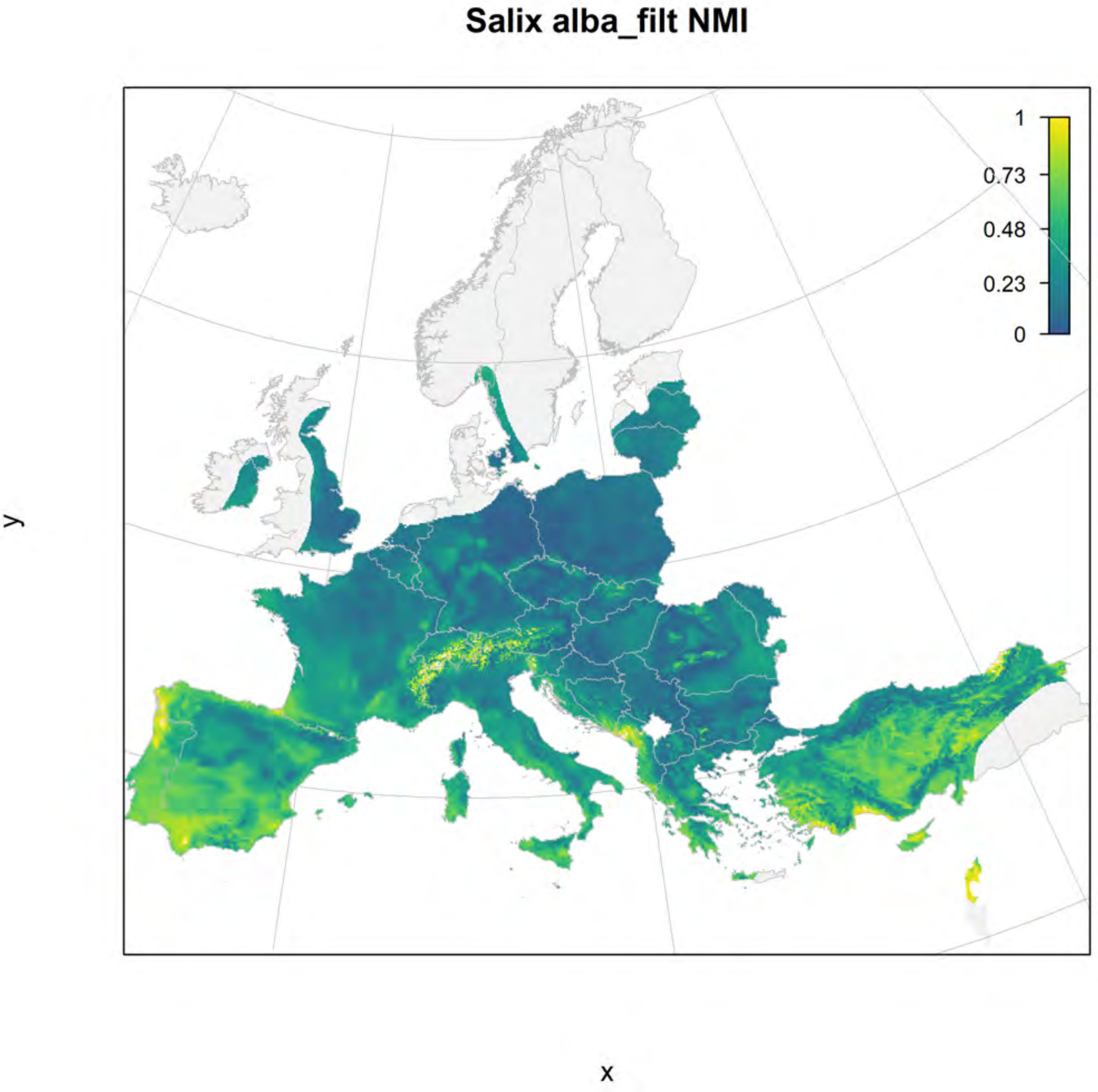
Salix alba_filt NMI
1
0.73
0.48
0.23
0
y
x

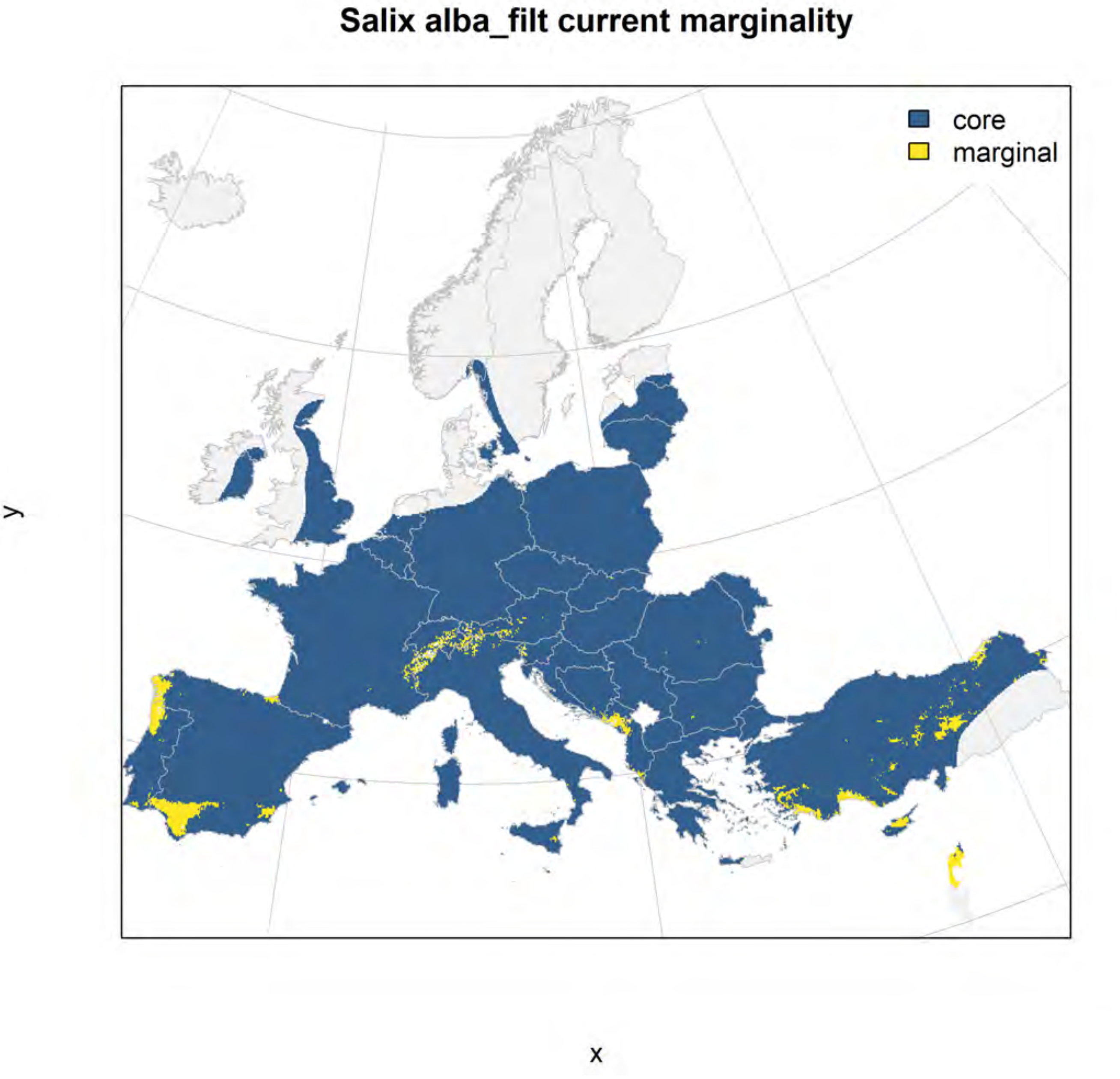
Salix alba_filt current marginality
core
marginal
y
x

#### Salix alba_filt future marginality

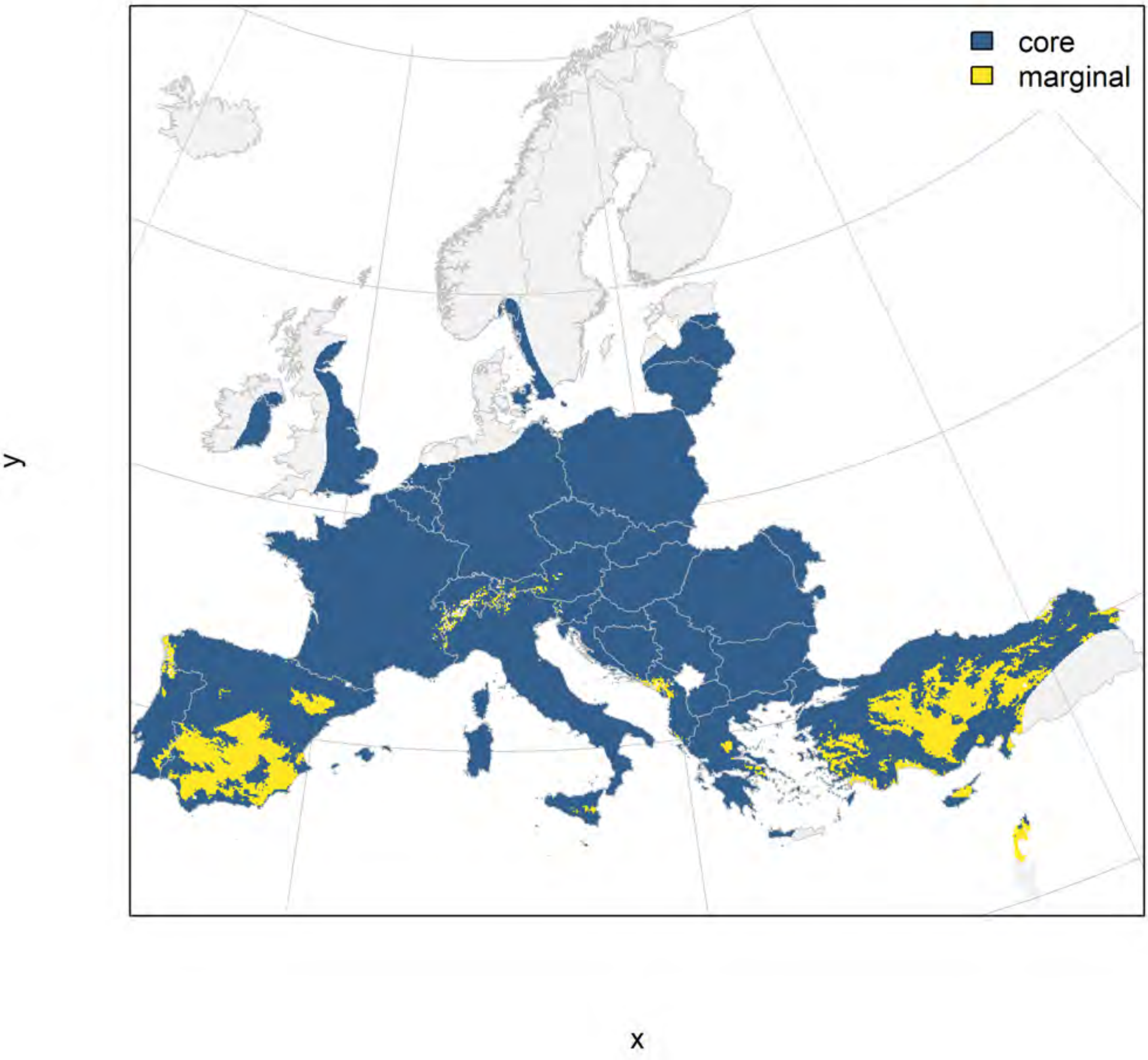

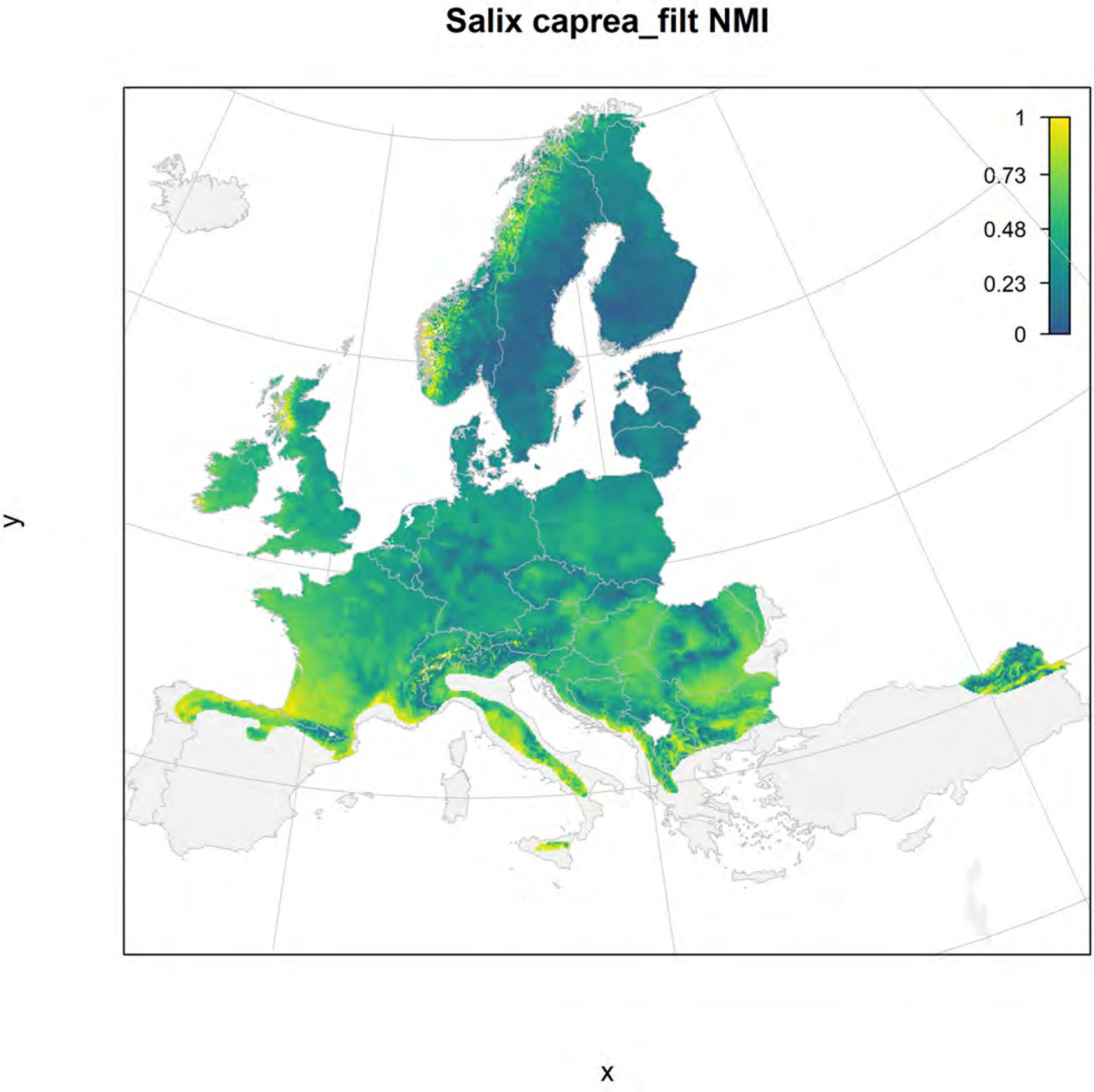
Salix caprea_filt NMI
1
0.73
0.48
0.23
0
y
x

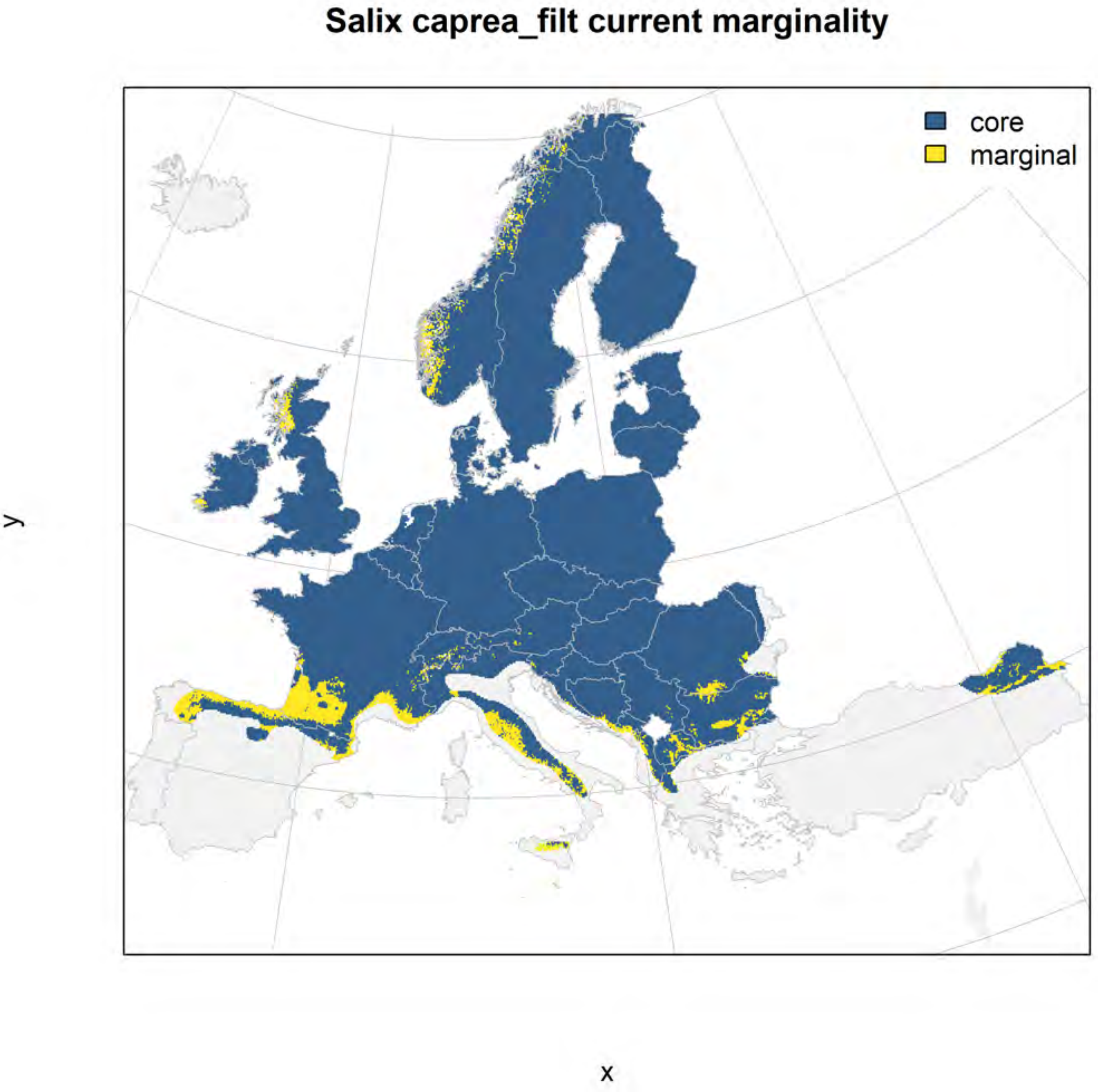
Salix caprea_filt current marginality
core
marginal
y
x

### Salix caprea_filt future marginality

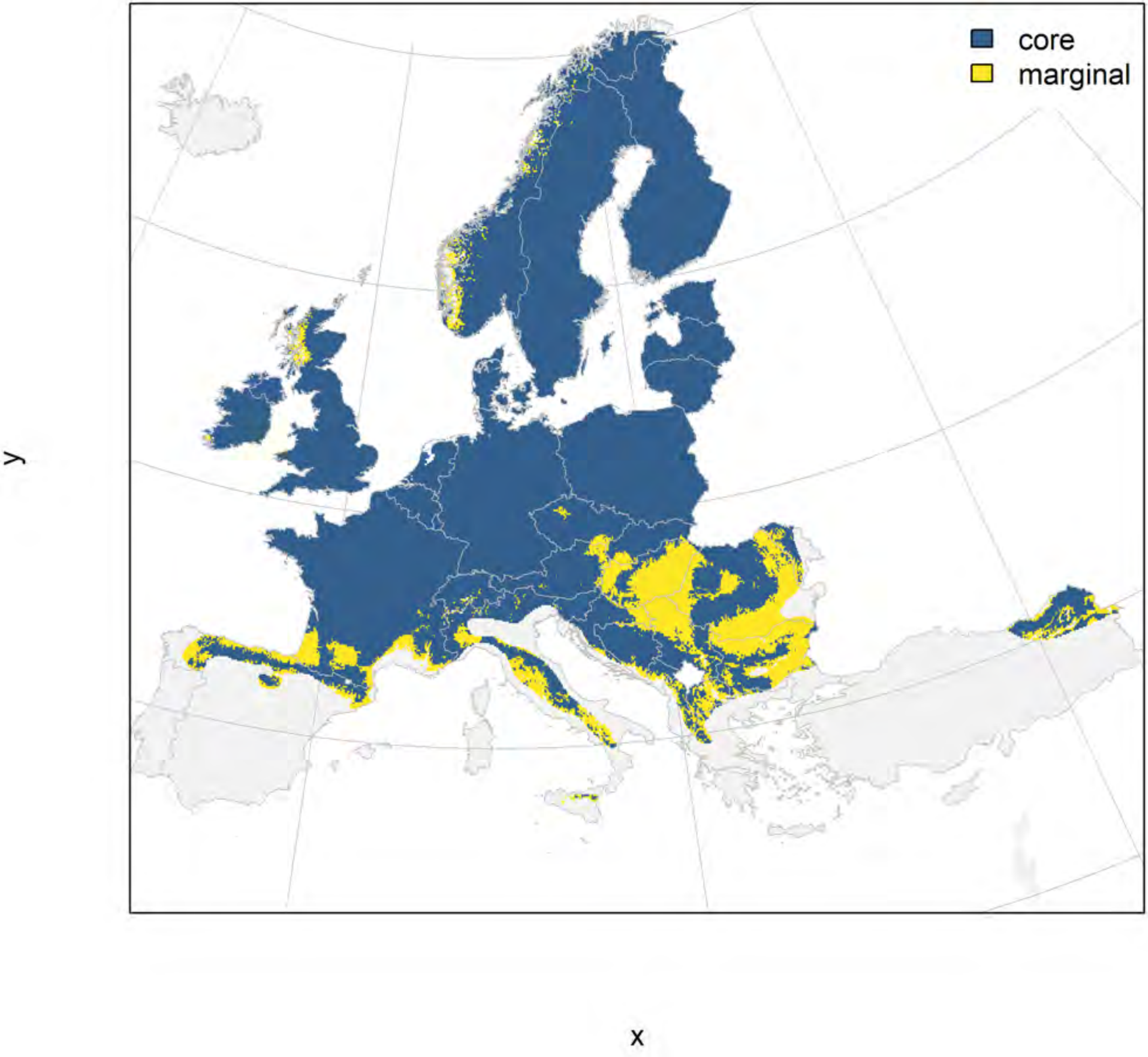

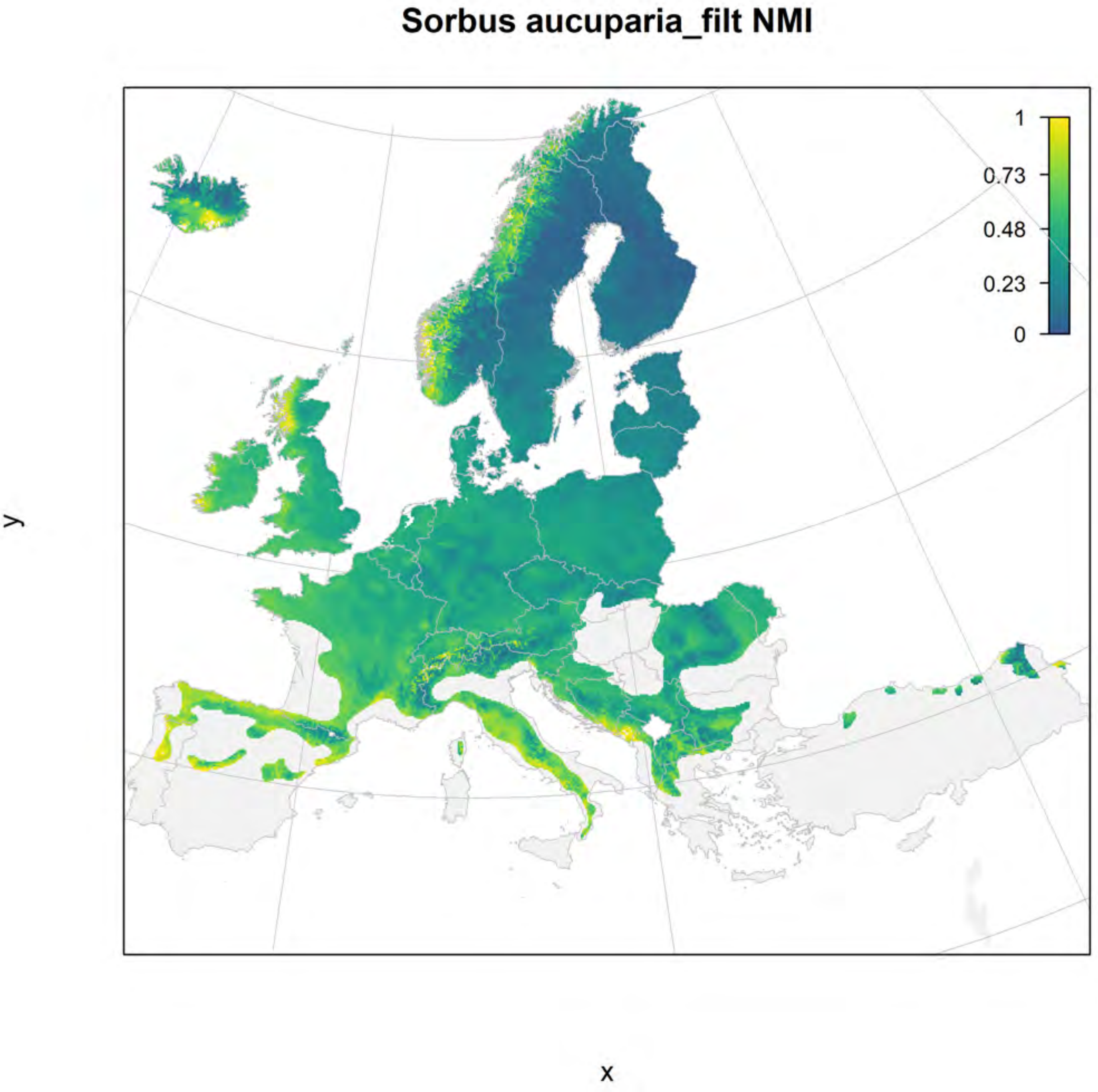

Sorbus aucuparia_filt NMI
1
0.73
0.48
0.23
0
y
x

#### Sorbus aucuparia_filt current marginality

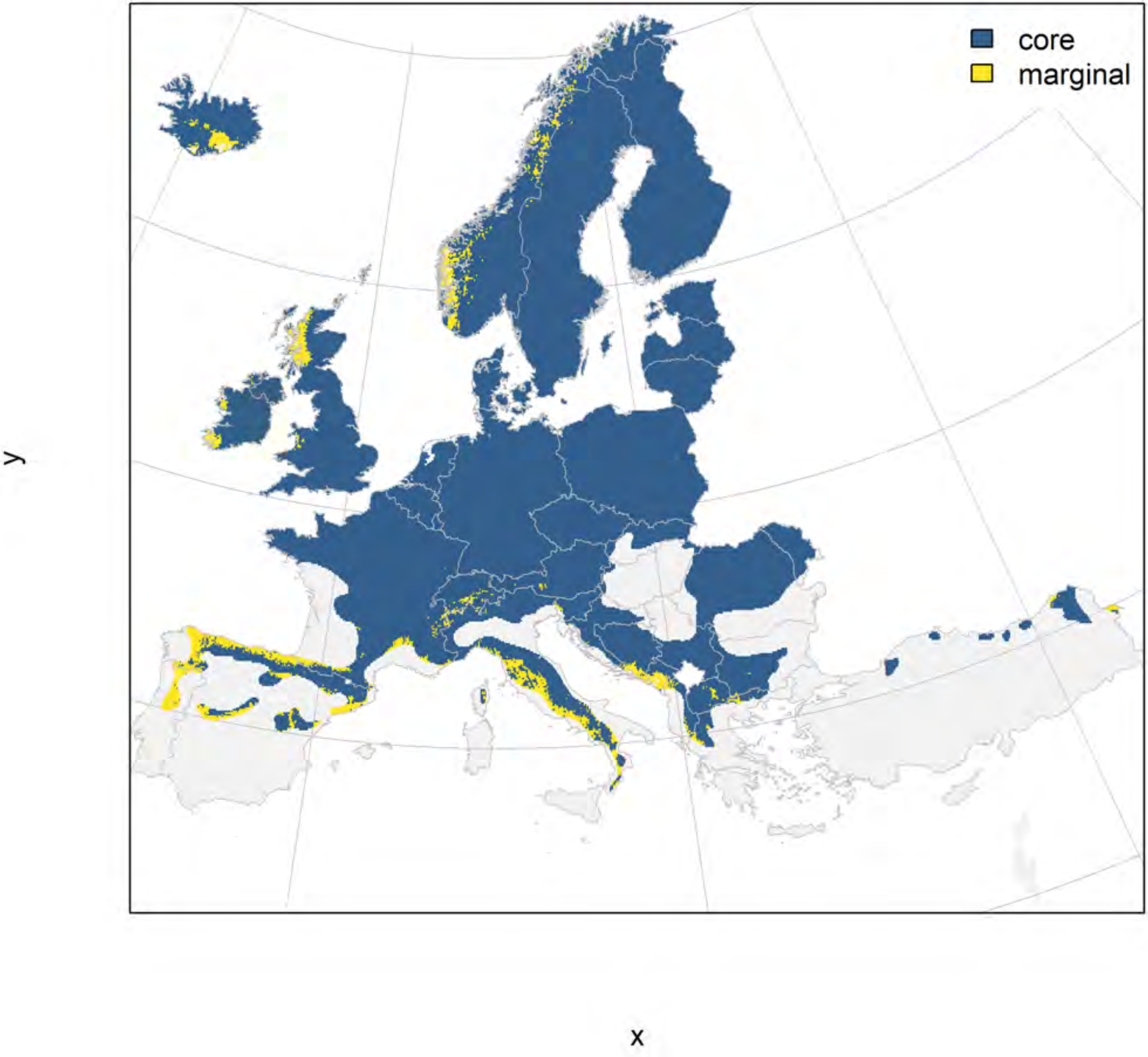

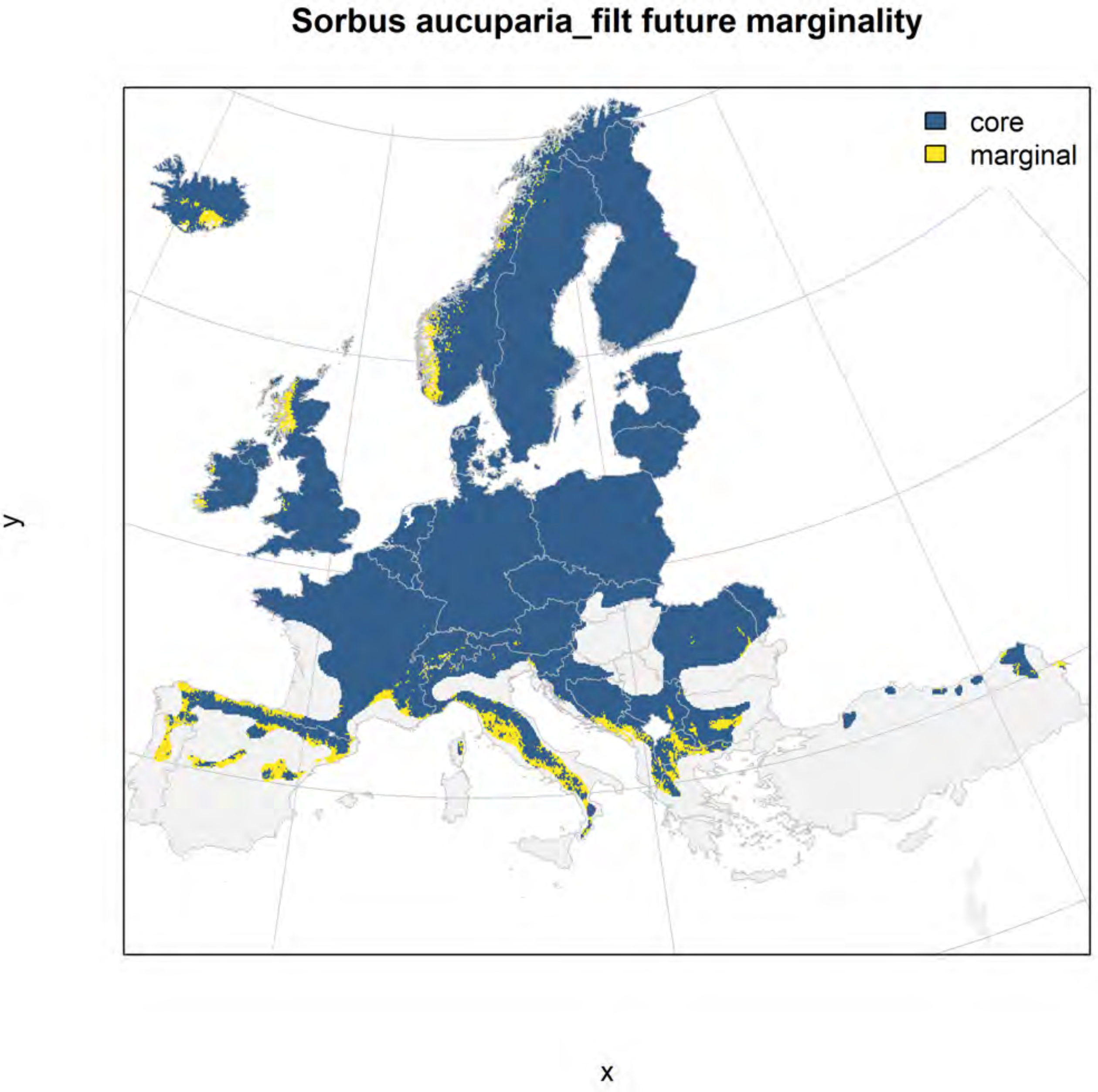
Sorbus aucuparia_filt future marginality
core
marginal
y
x

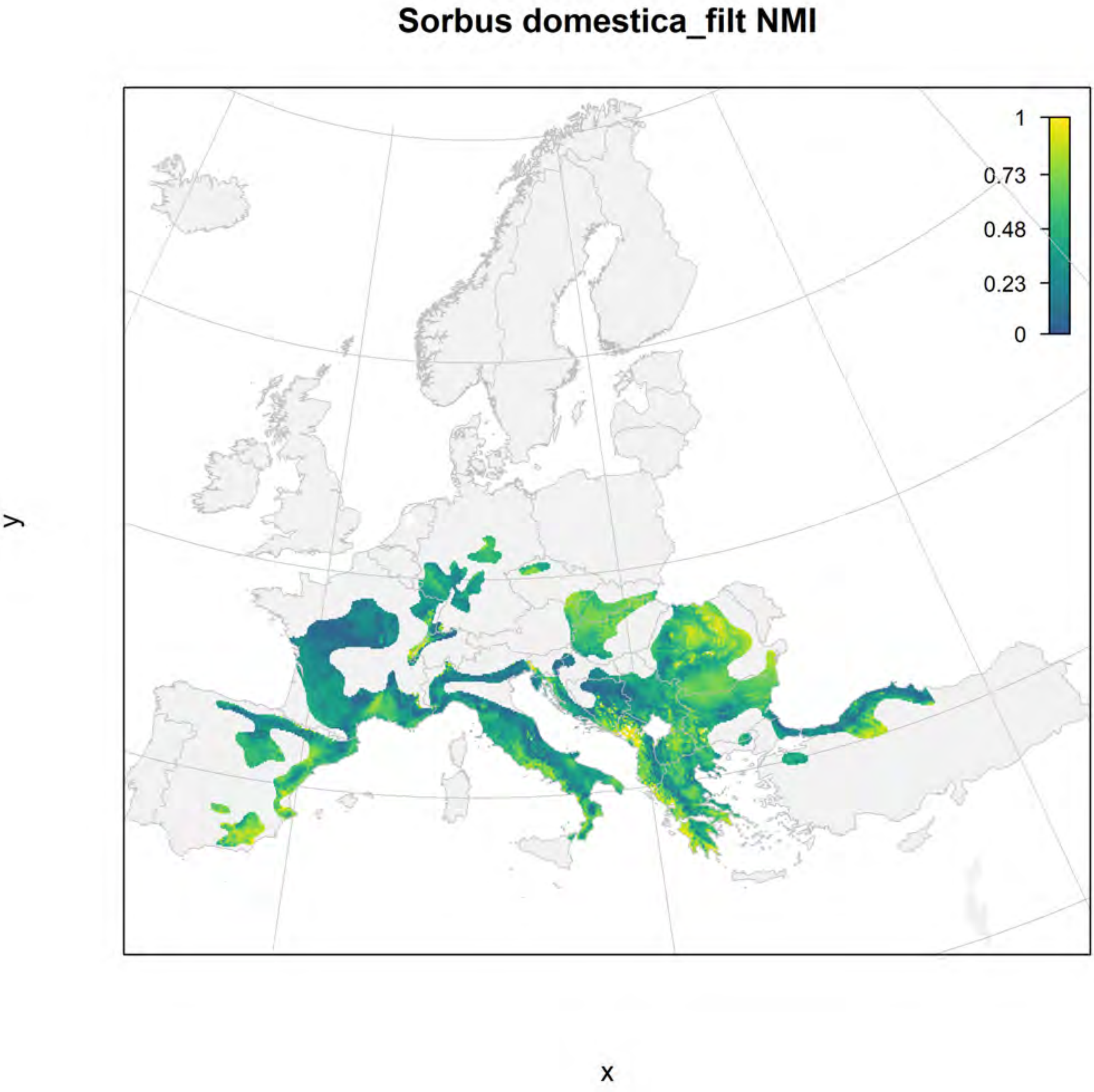

Sorbus domestica_filt NMI
1
0.73
0.48
0.23
0
y
x

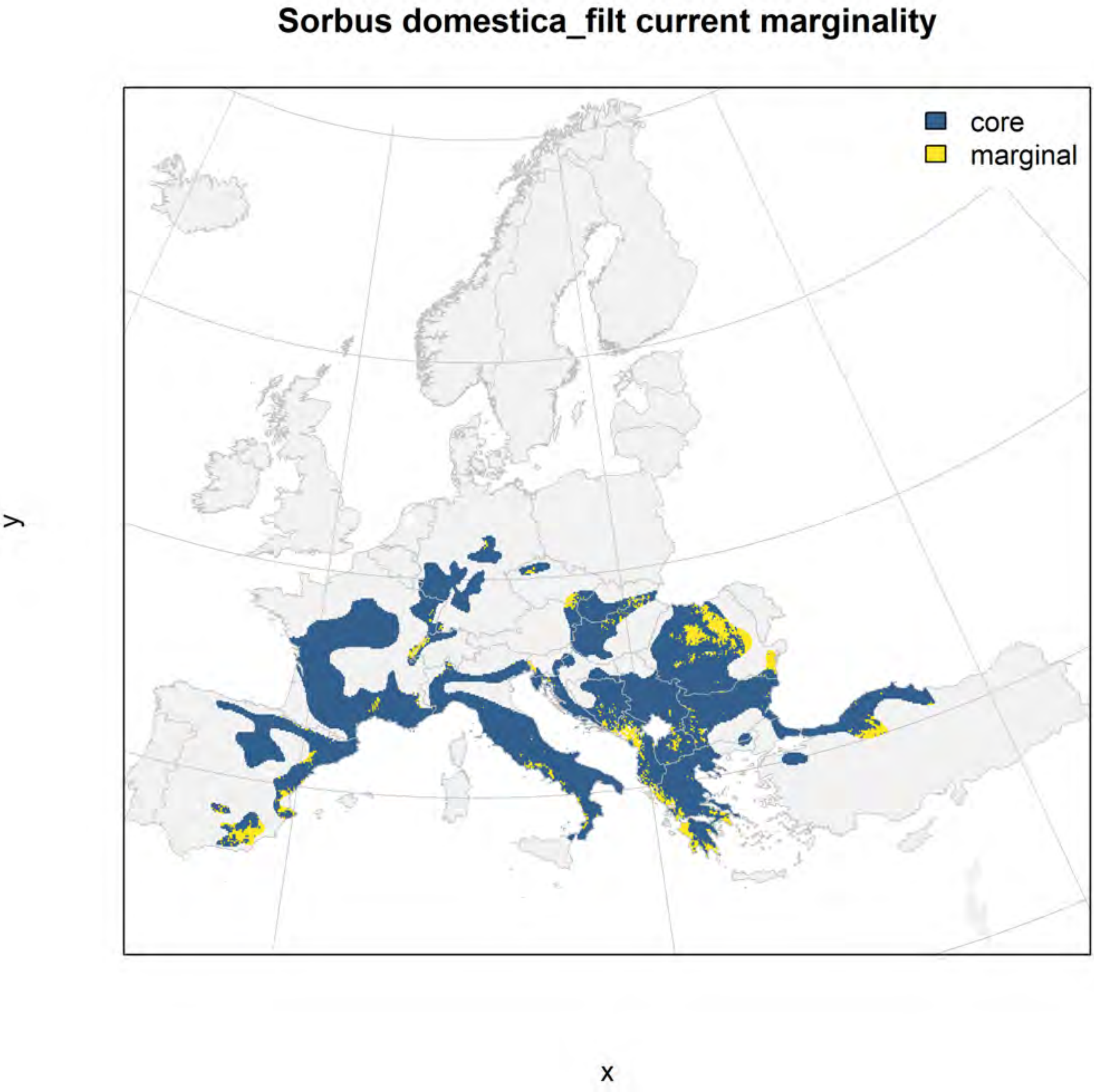

Sorbus domestica_filt current marginality
core
marginal
y
x

#### Sorbus domestica_filt future marginality

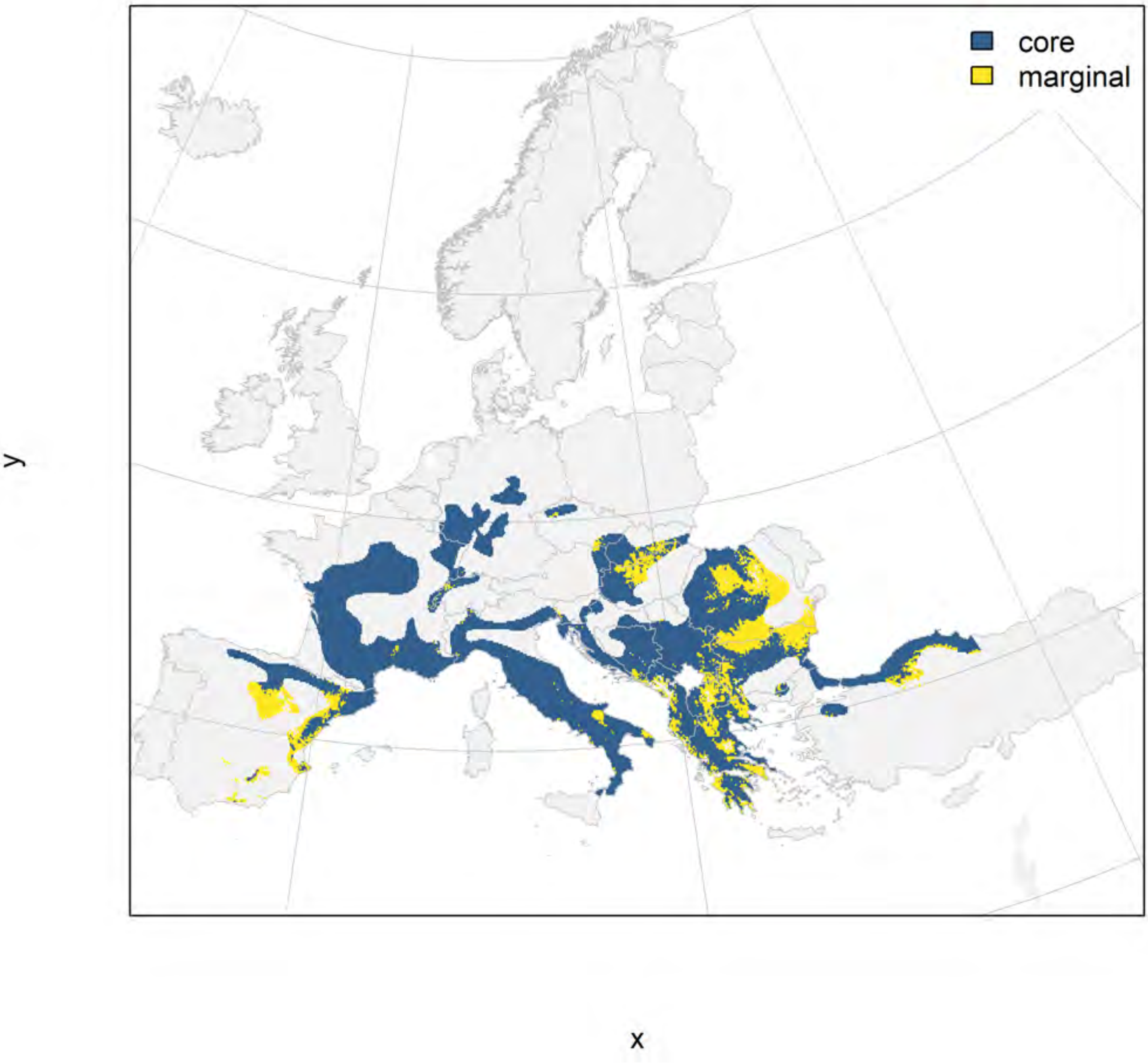

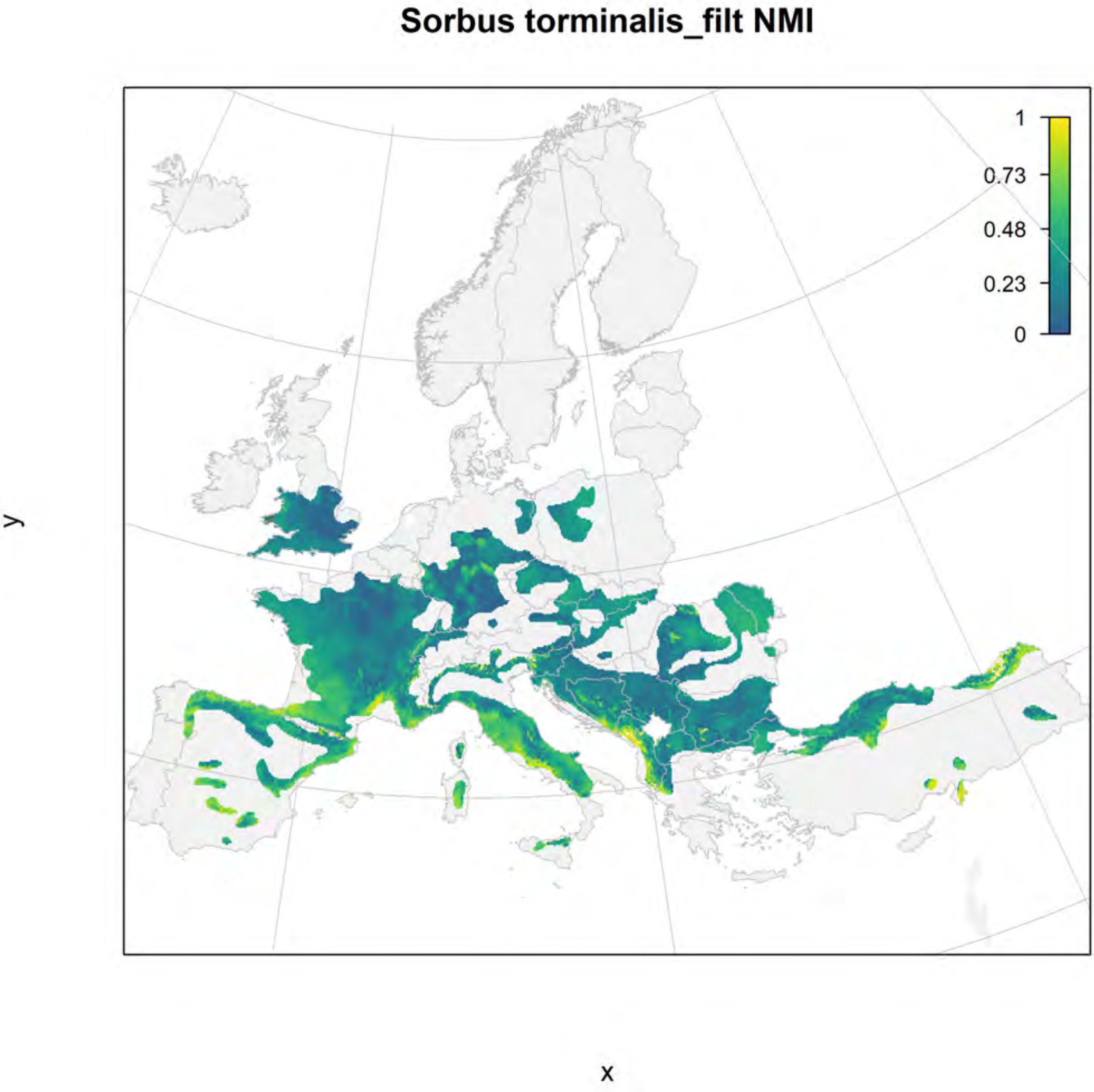
Sorbus torminalis_filt NMI
1
0.73
0.48
0.23
0
y
x

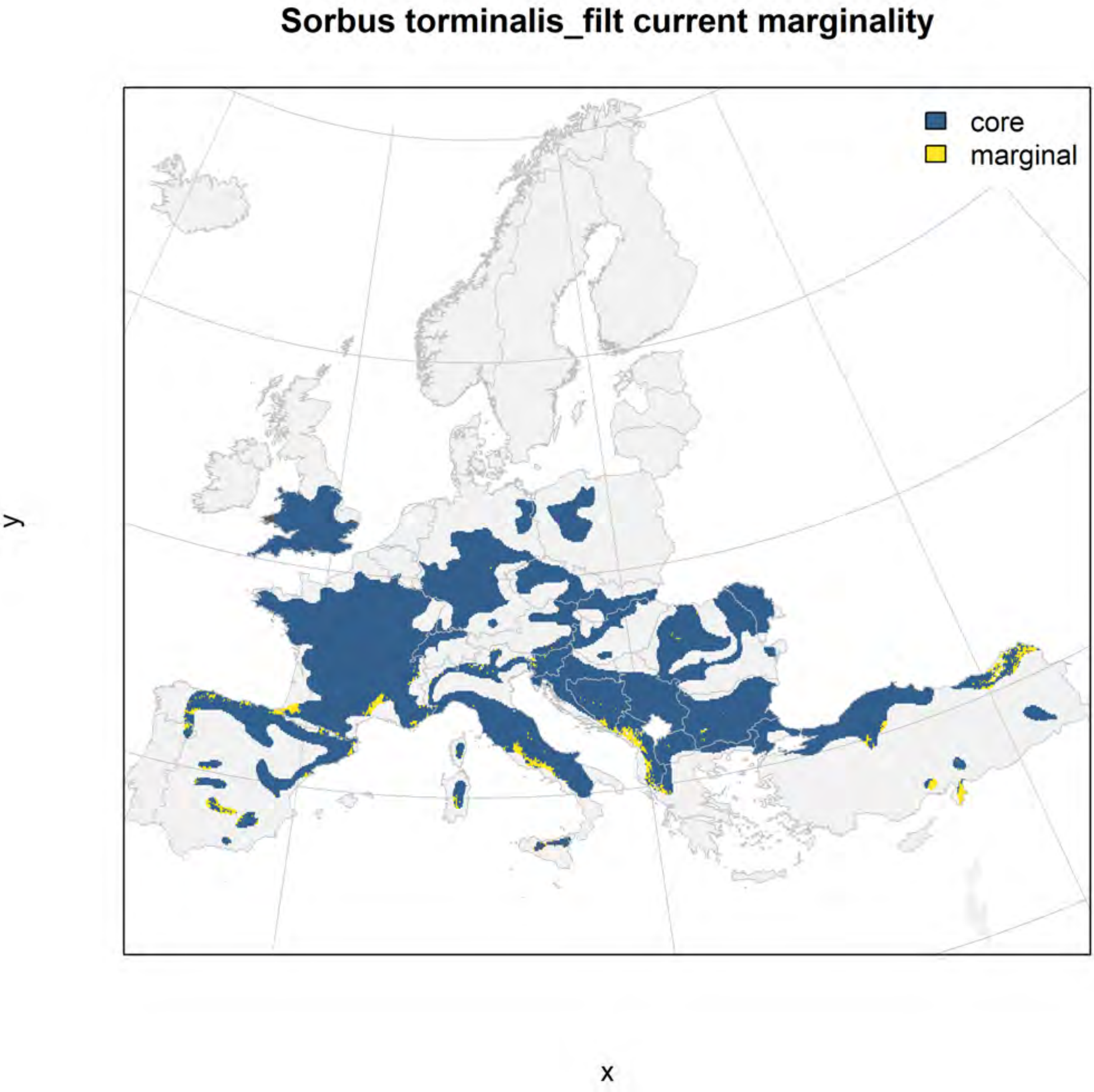
Sorbus torminalis_filt current marginality
core
marginal
y
x

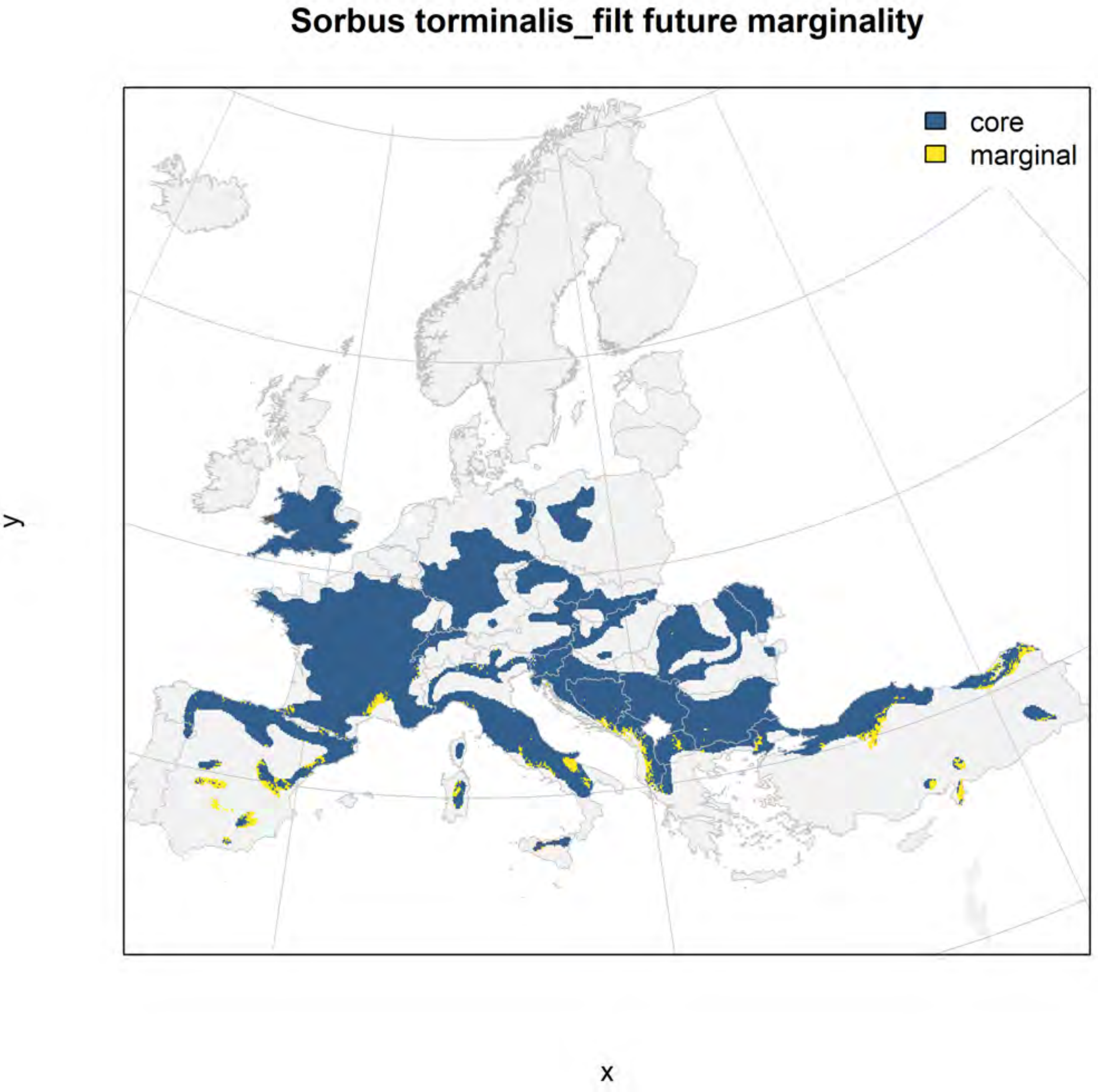
Sorbus torminalis_filt future marginality
core
marginal
y
x

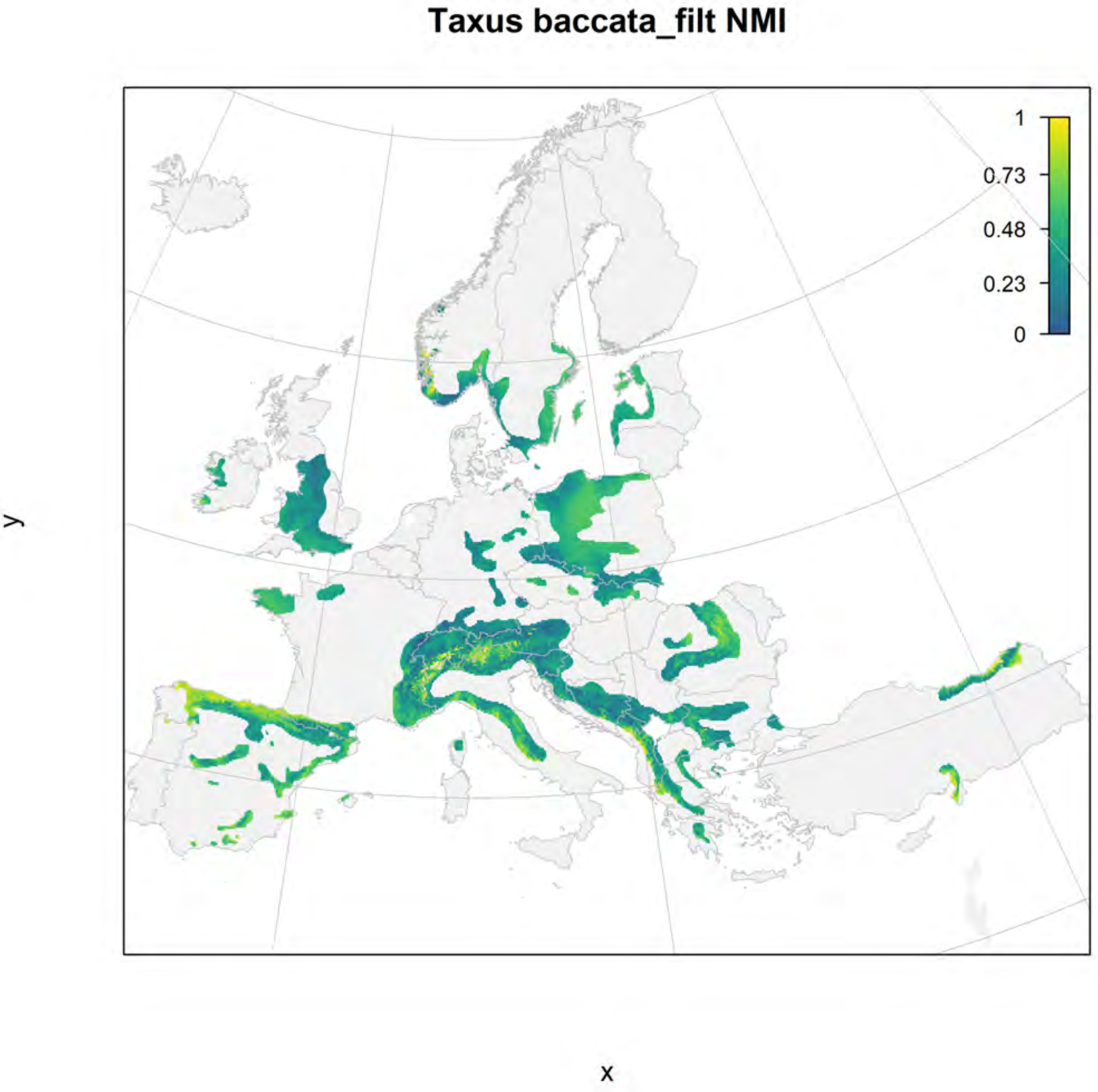
Taxus baccata_filt NMI
1
0.73
0.48
0.23
0
y
x

### Taxus baccata_filt current marginality

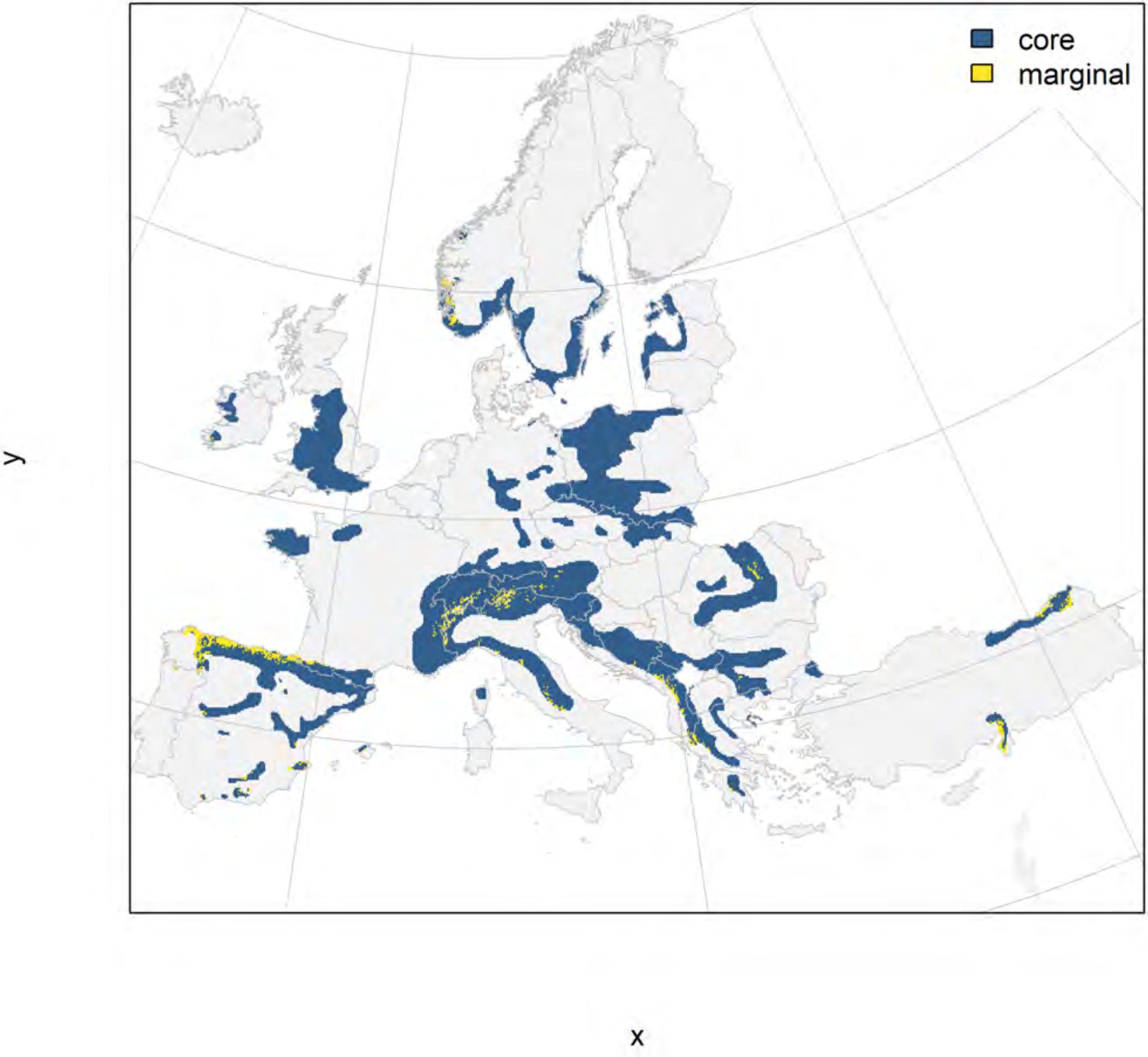

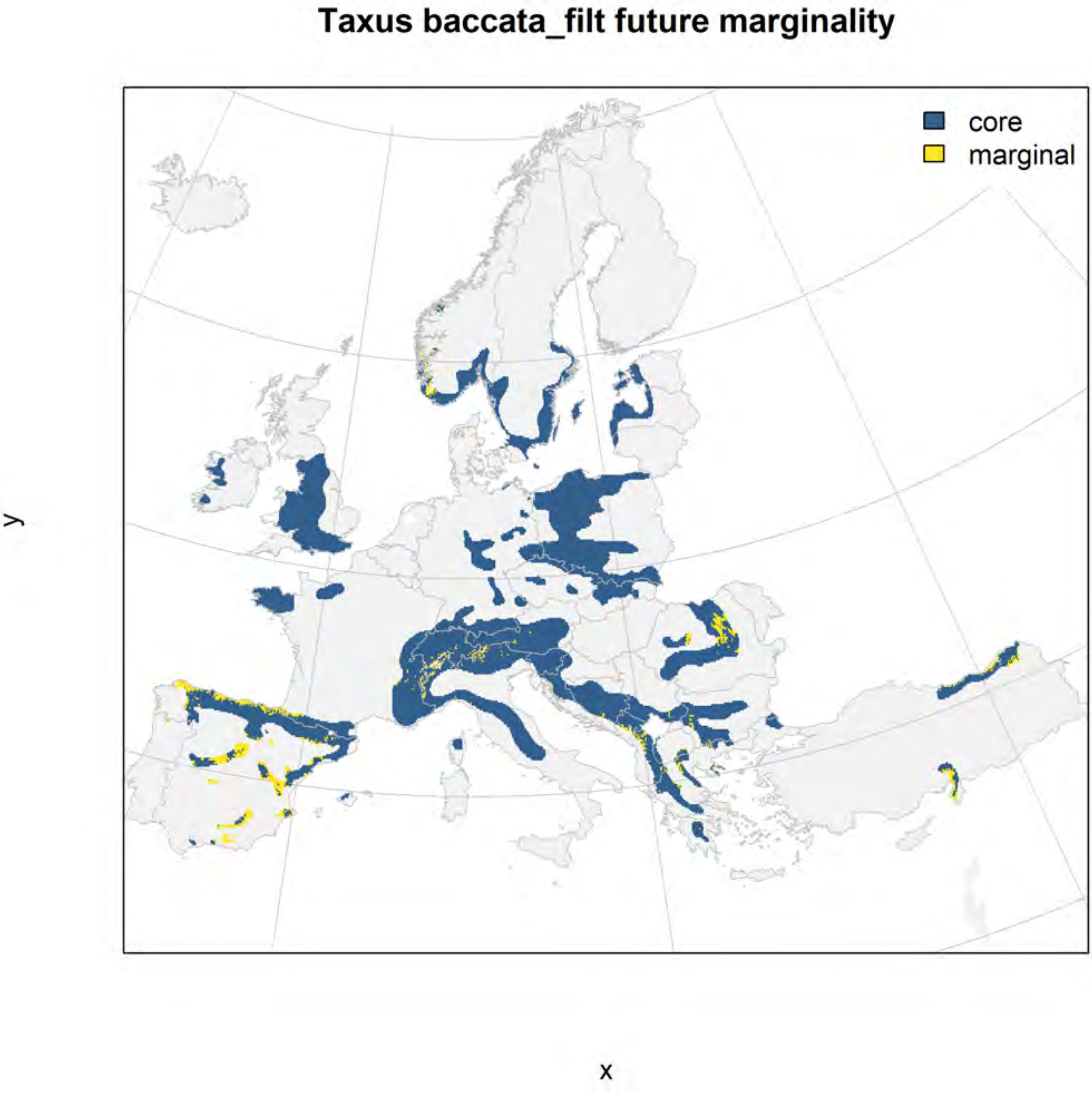
Taxus baccata_filt future marginality
core
marginal
y
x

### Tilia cordata_filt NMI

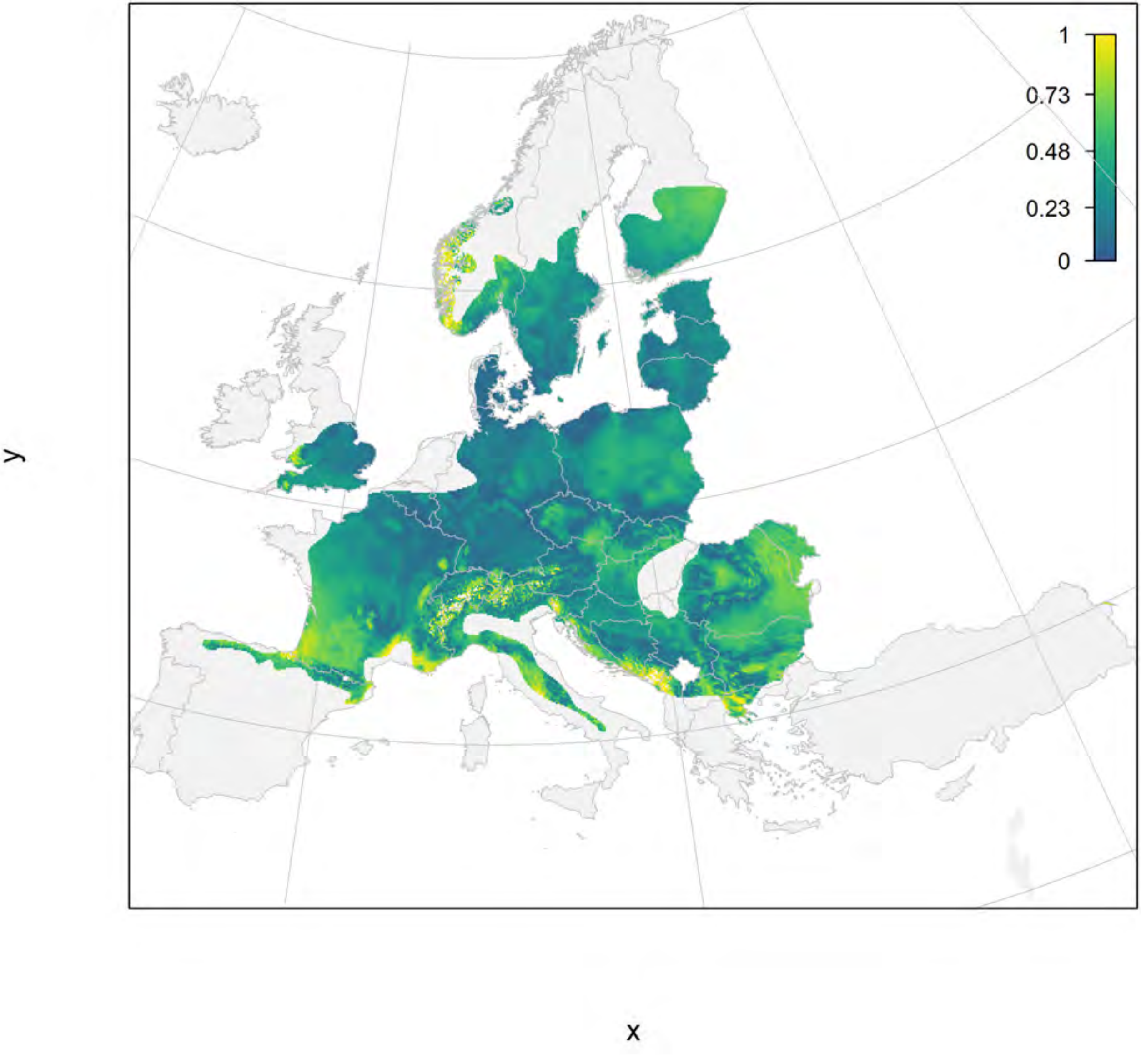

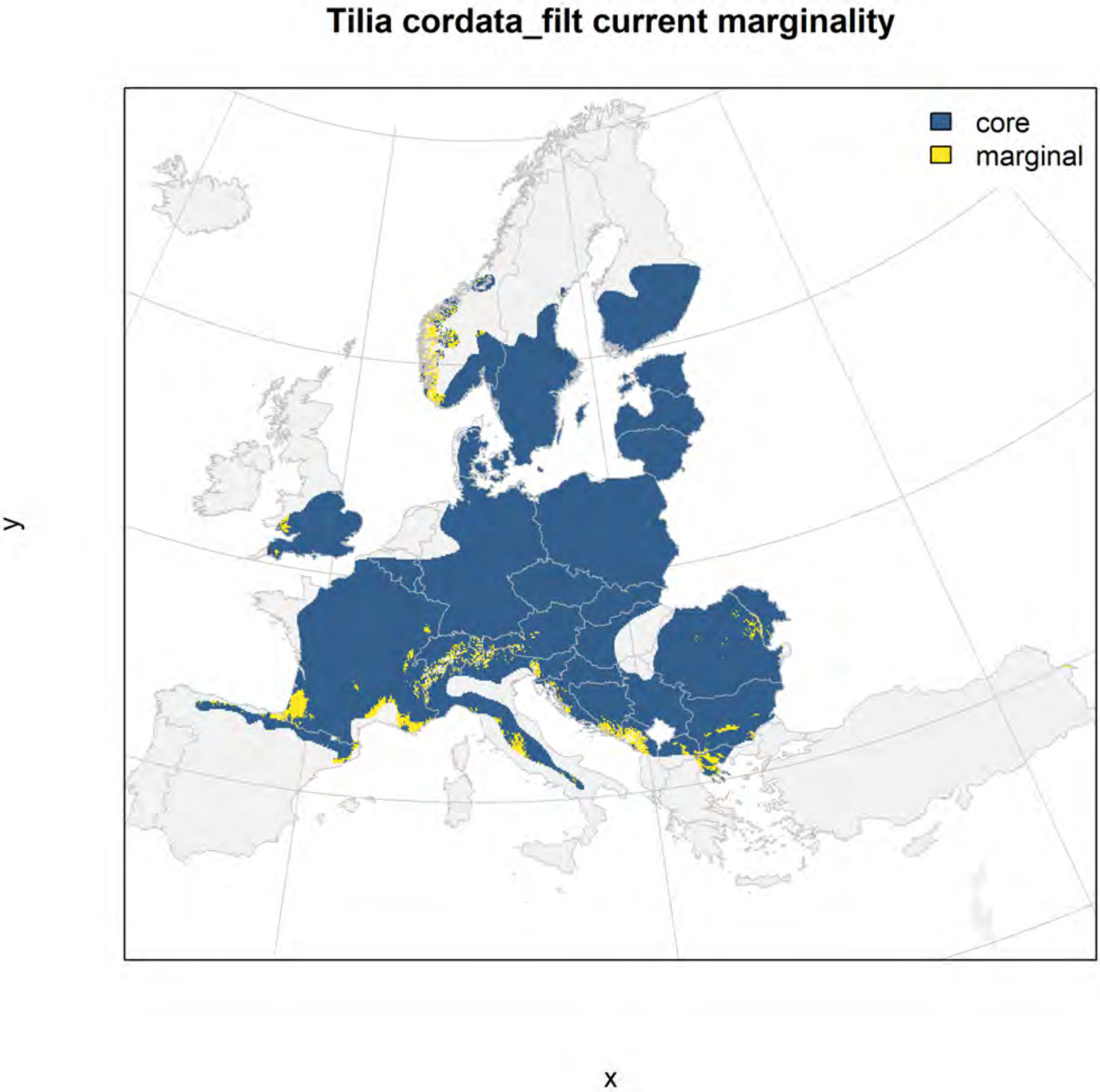
Tilia cordata_filt current marginality
core
marginal
y
x

Tilia cordata_filt future marginality

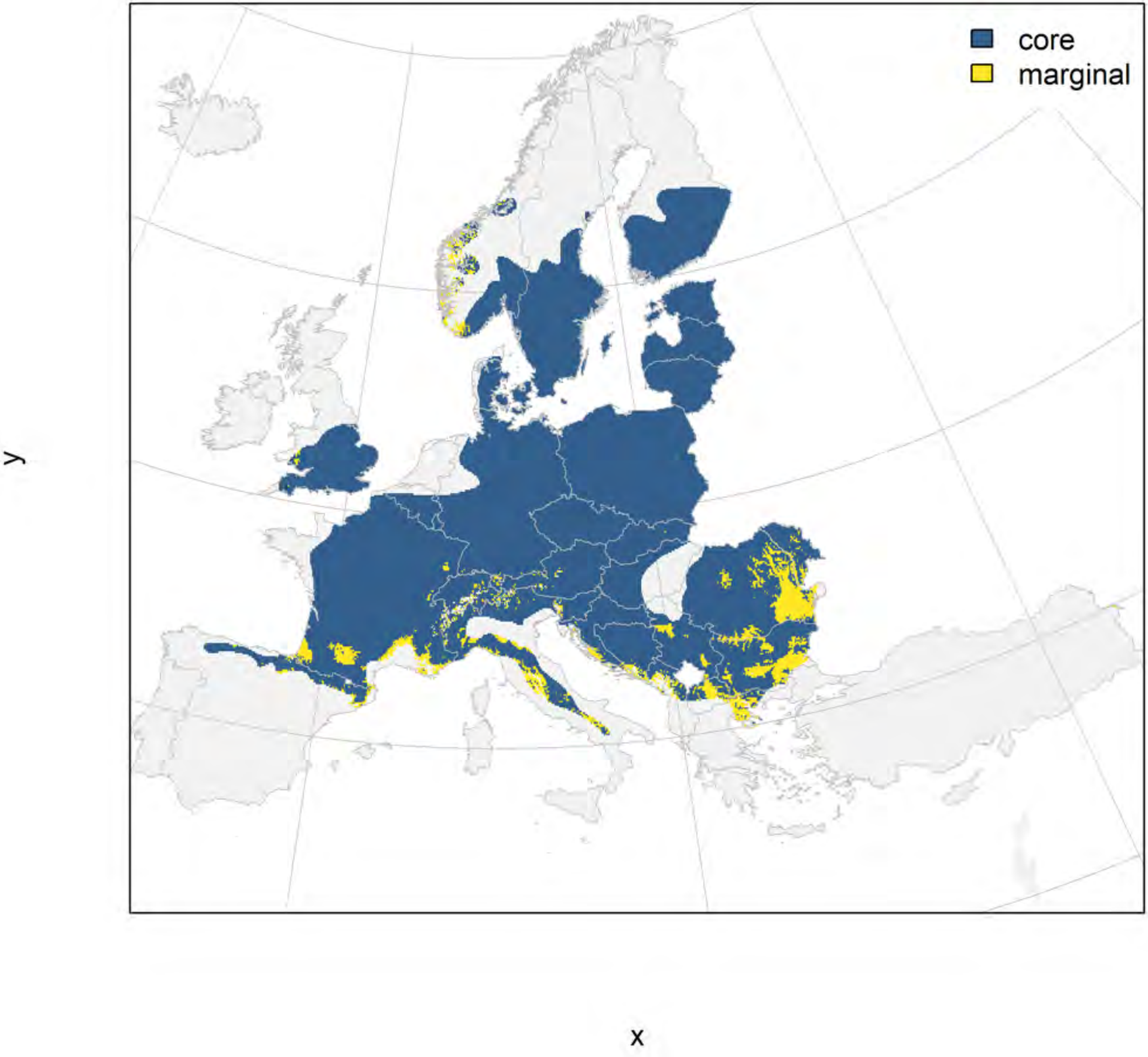

Tilia platyphyllos_filt NMI

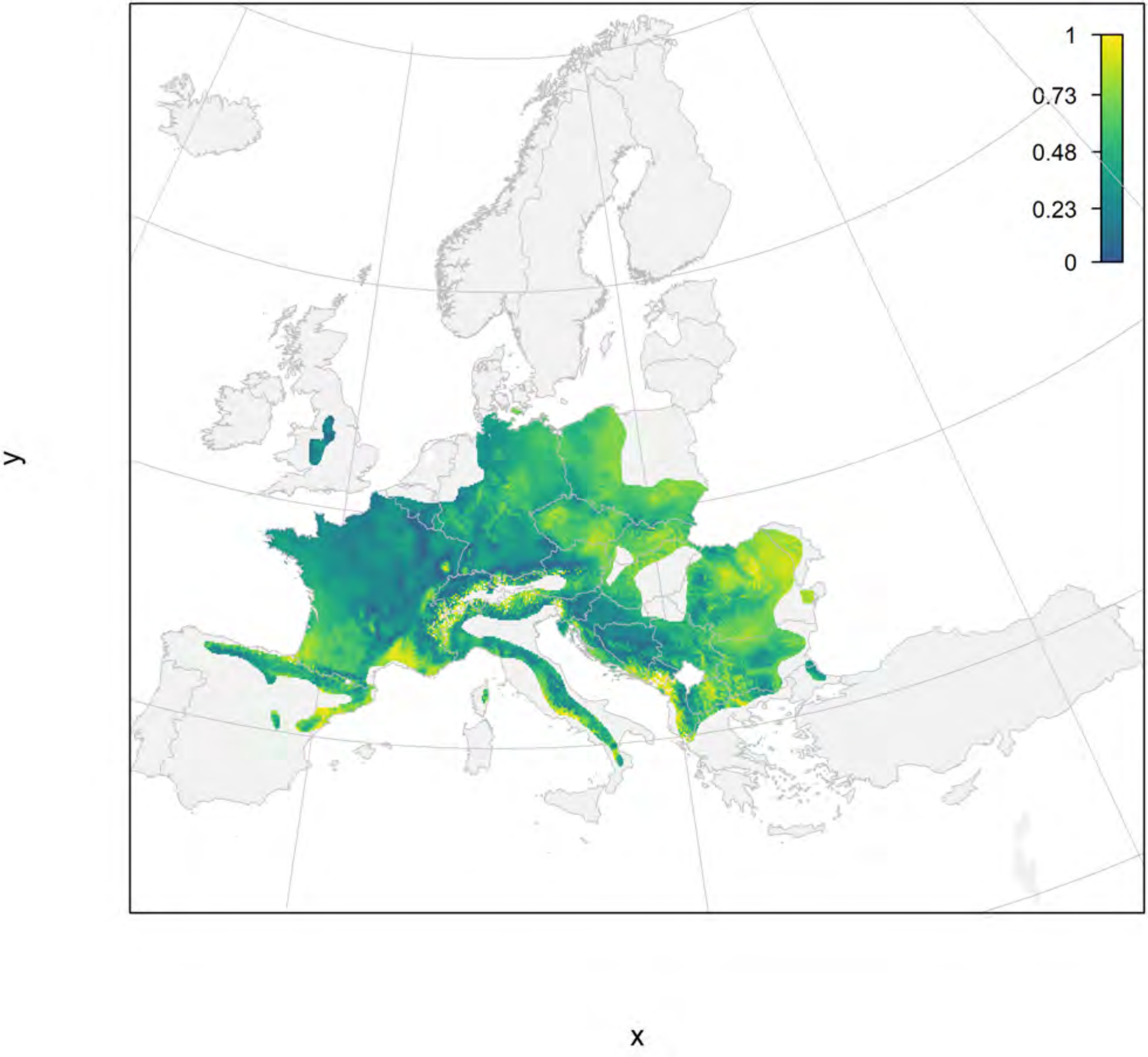

#### Tilia platyphyllos_filt current marginality

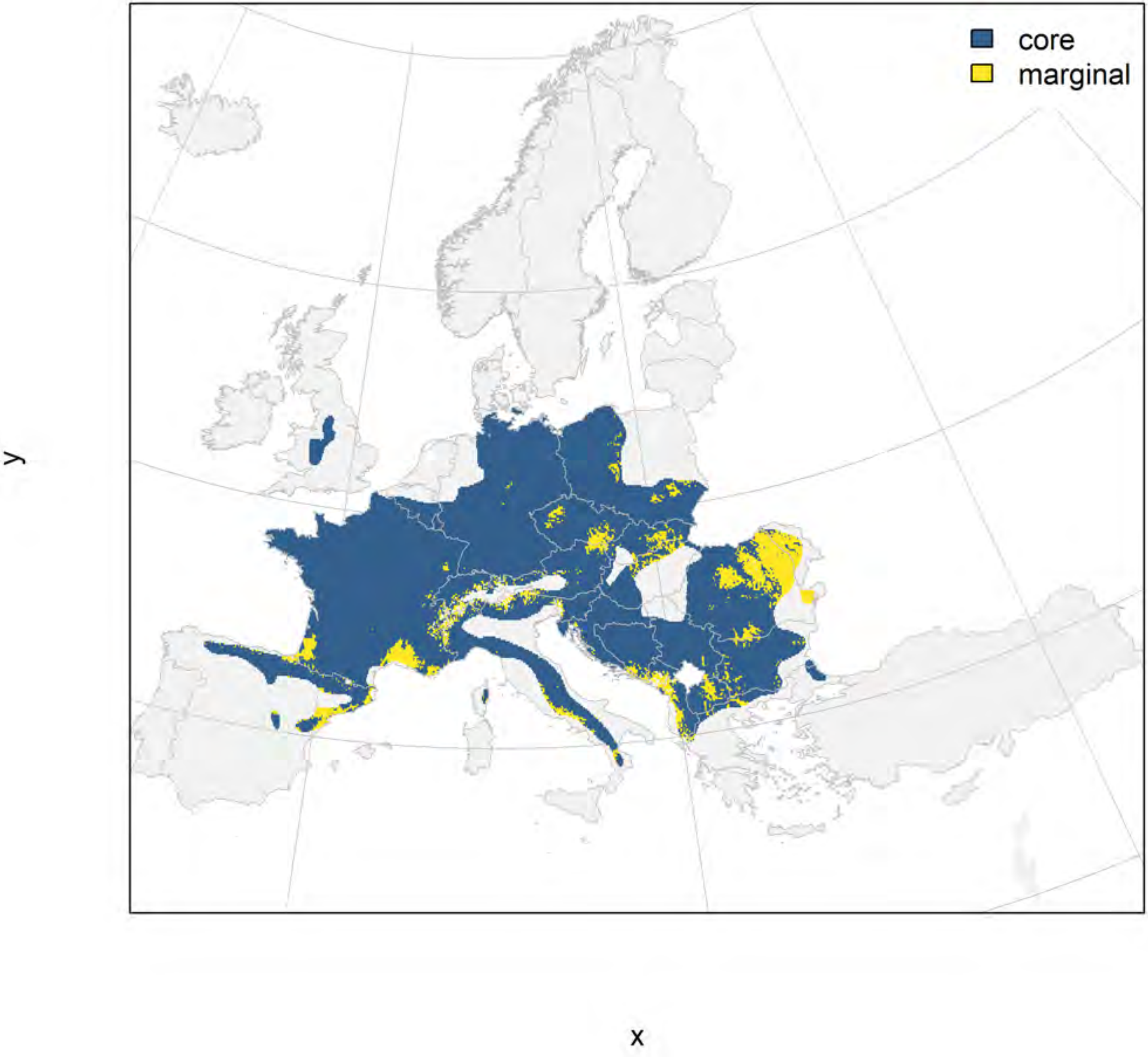

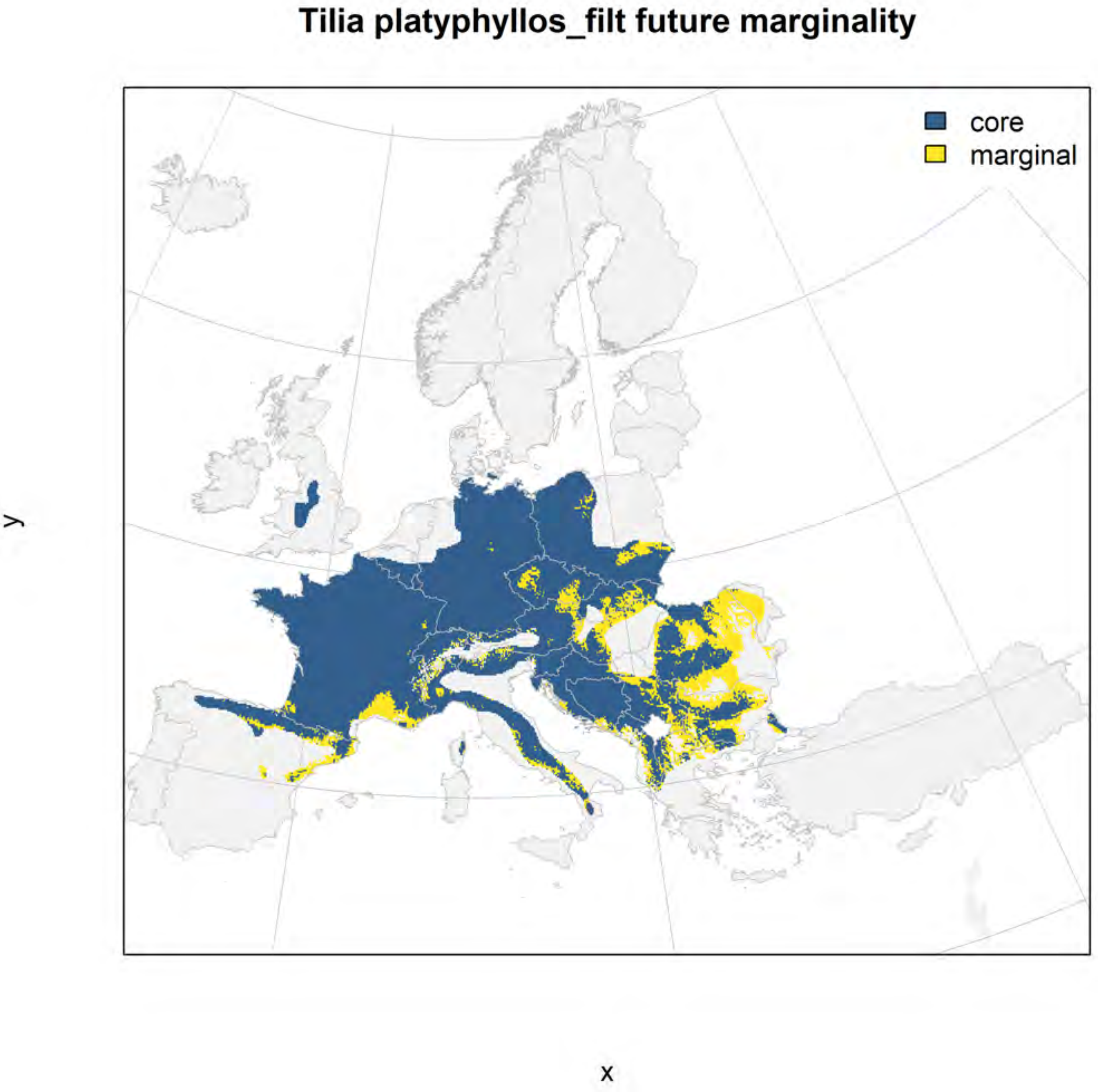
Tilia platyphyllos_filt future marginality
core
marginal
y
x

#### Tilia tomentosa_filt NMI

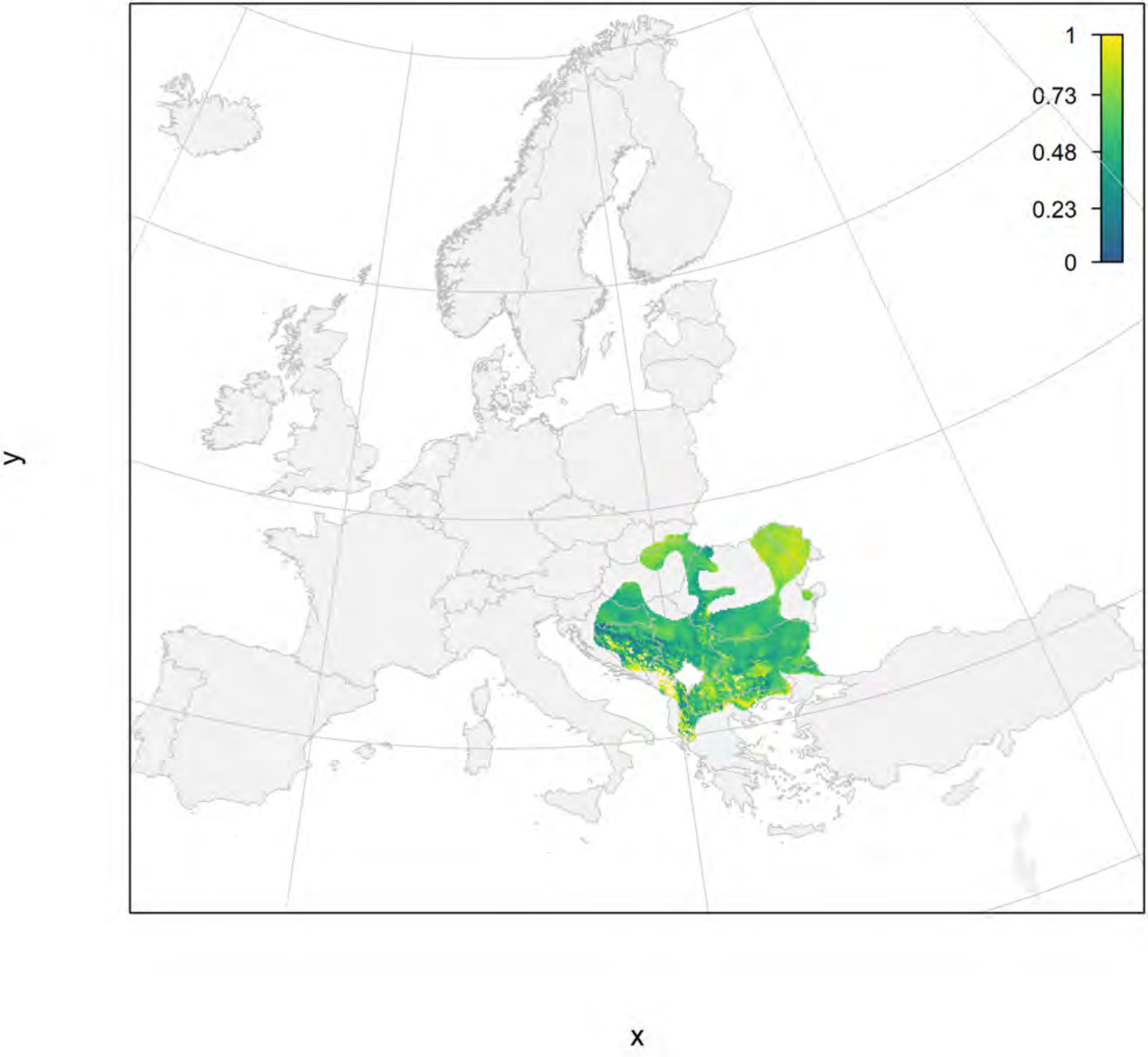

#### Tilia tomentosa_filt current marginality

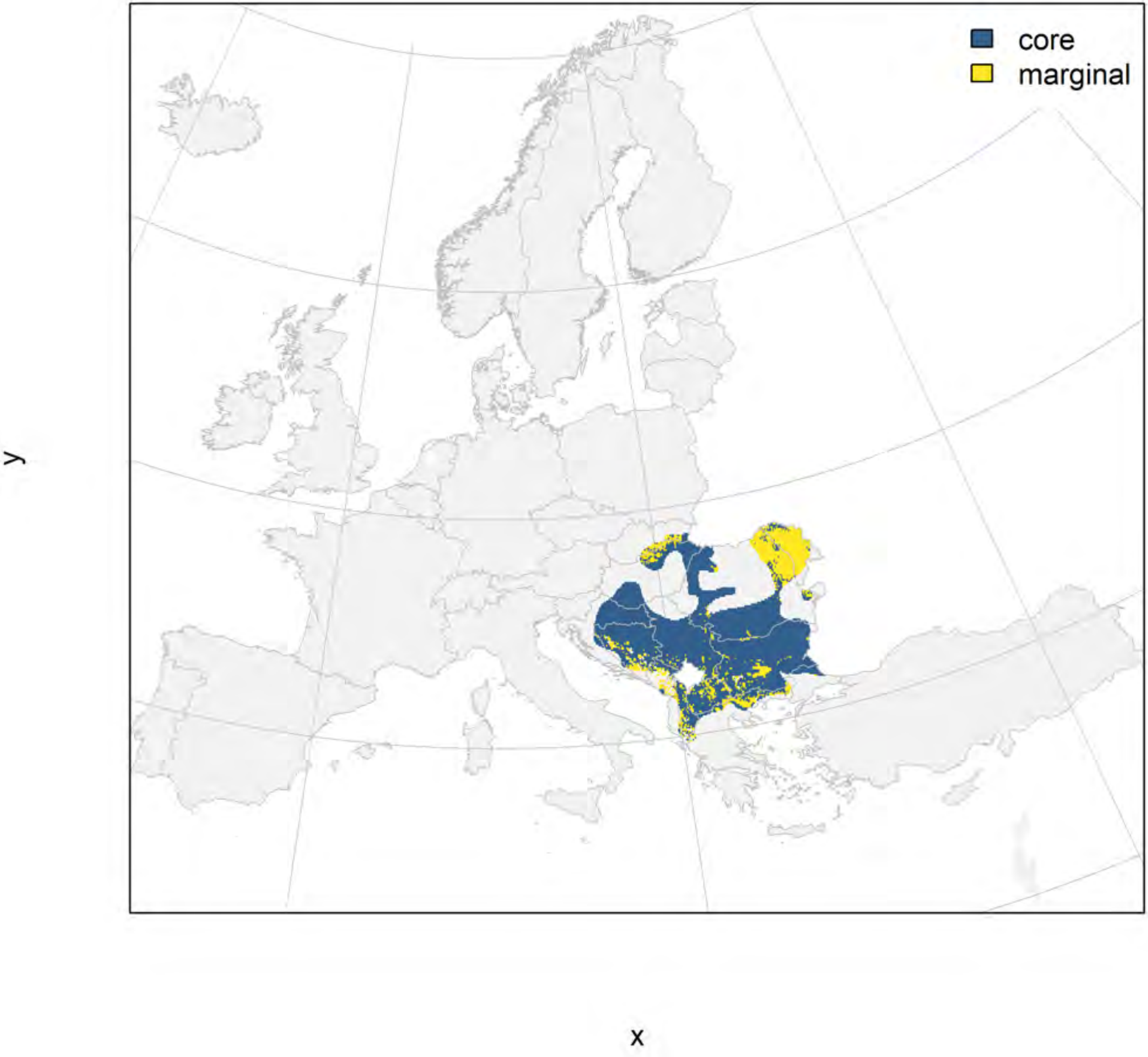

#### Tilia tomentosa_filt future marginality

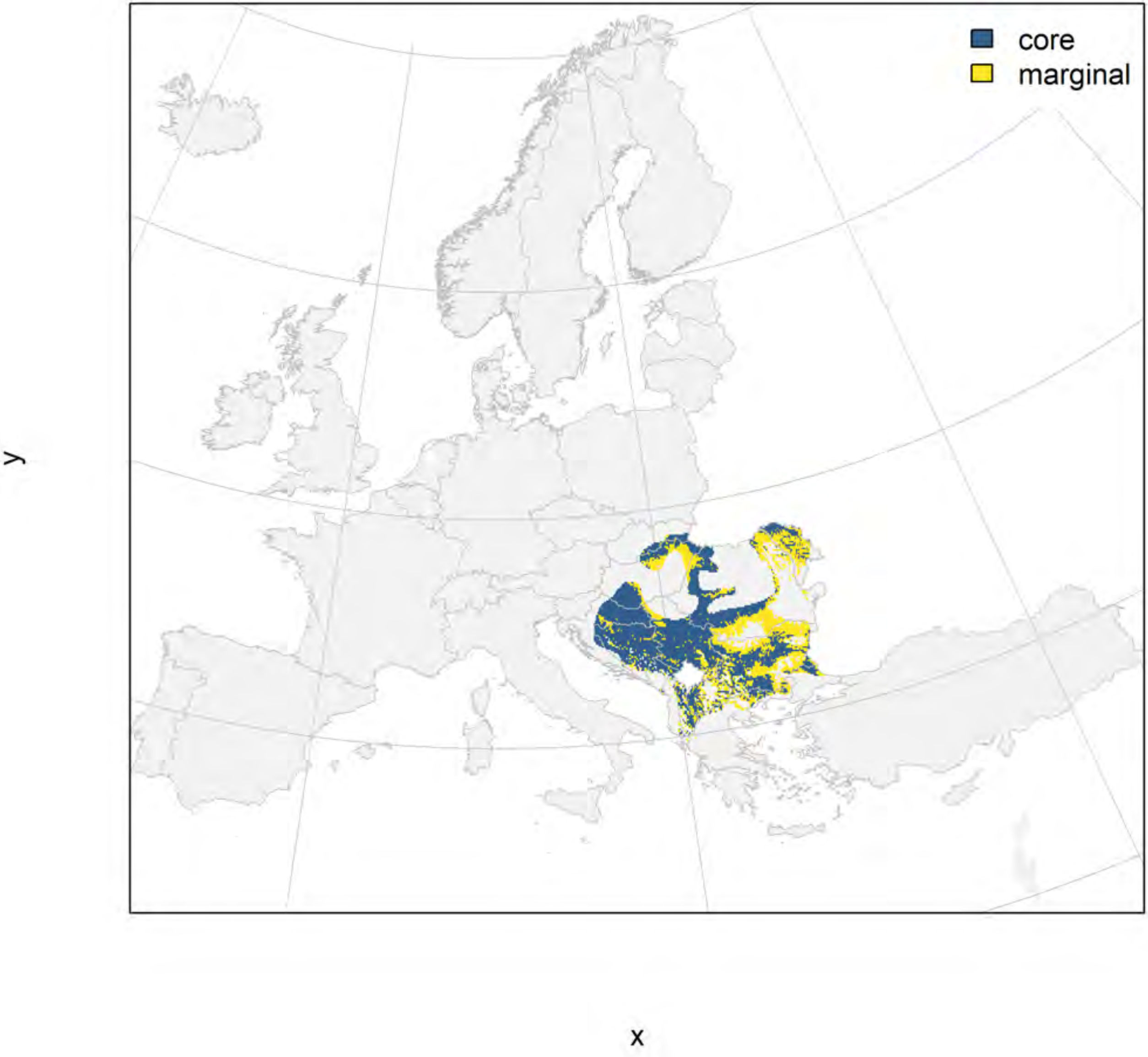

### Ulmus glabra_filt NMI

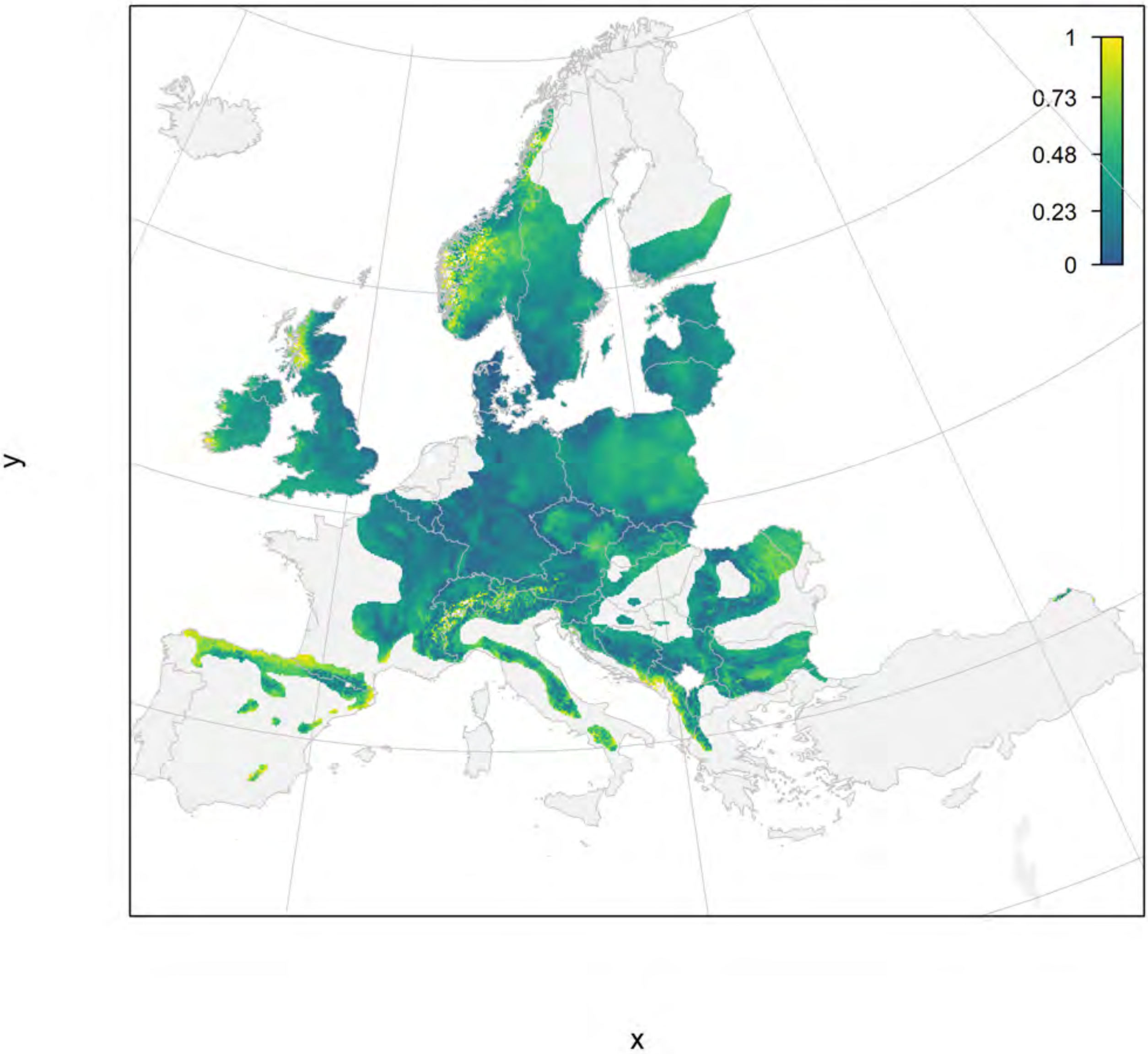

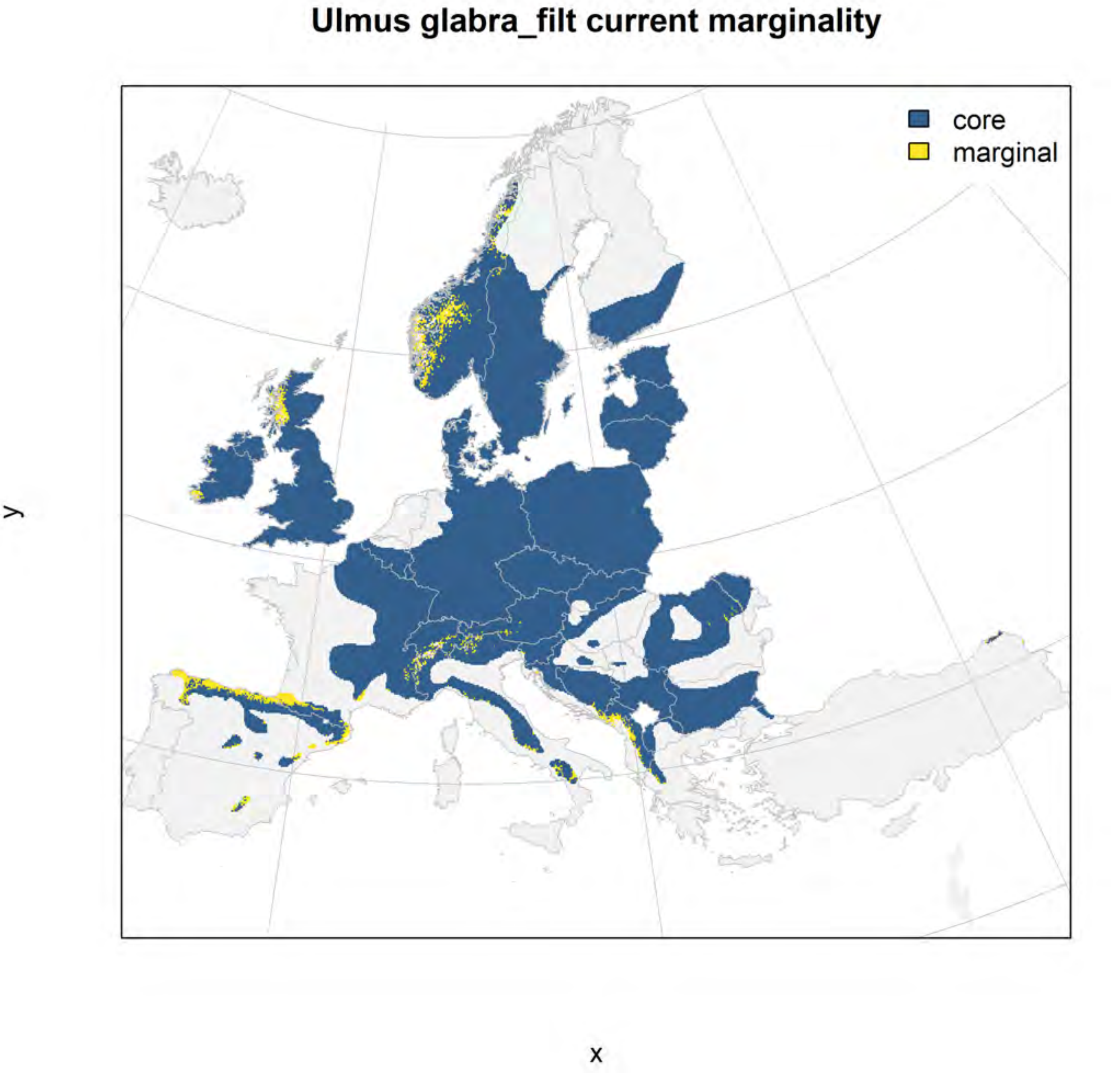
Ulmus glabra_filt current marginality
core
marginal
y
x

#### Ulmus glabra_filt future marginality

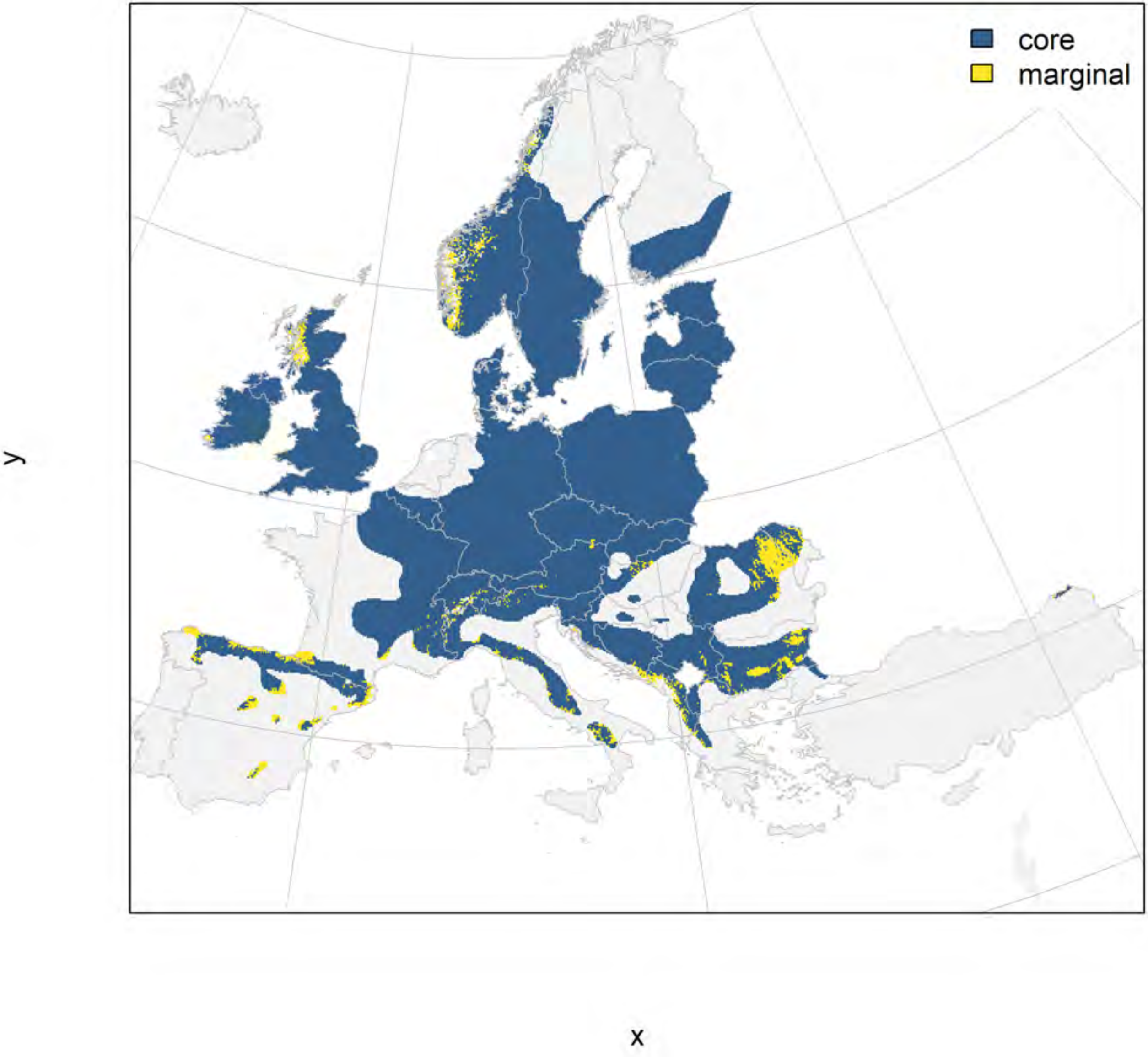

### Ulmus laevis_filt NMI

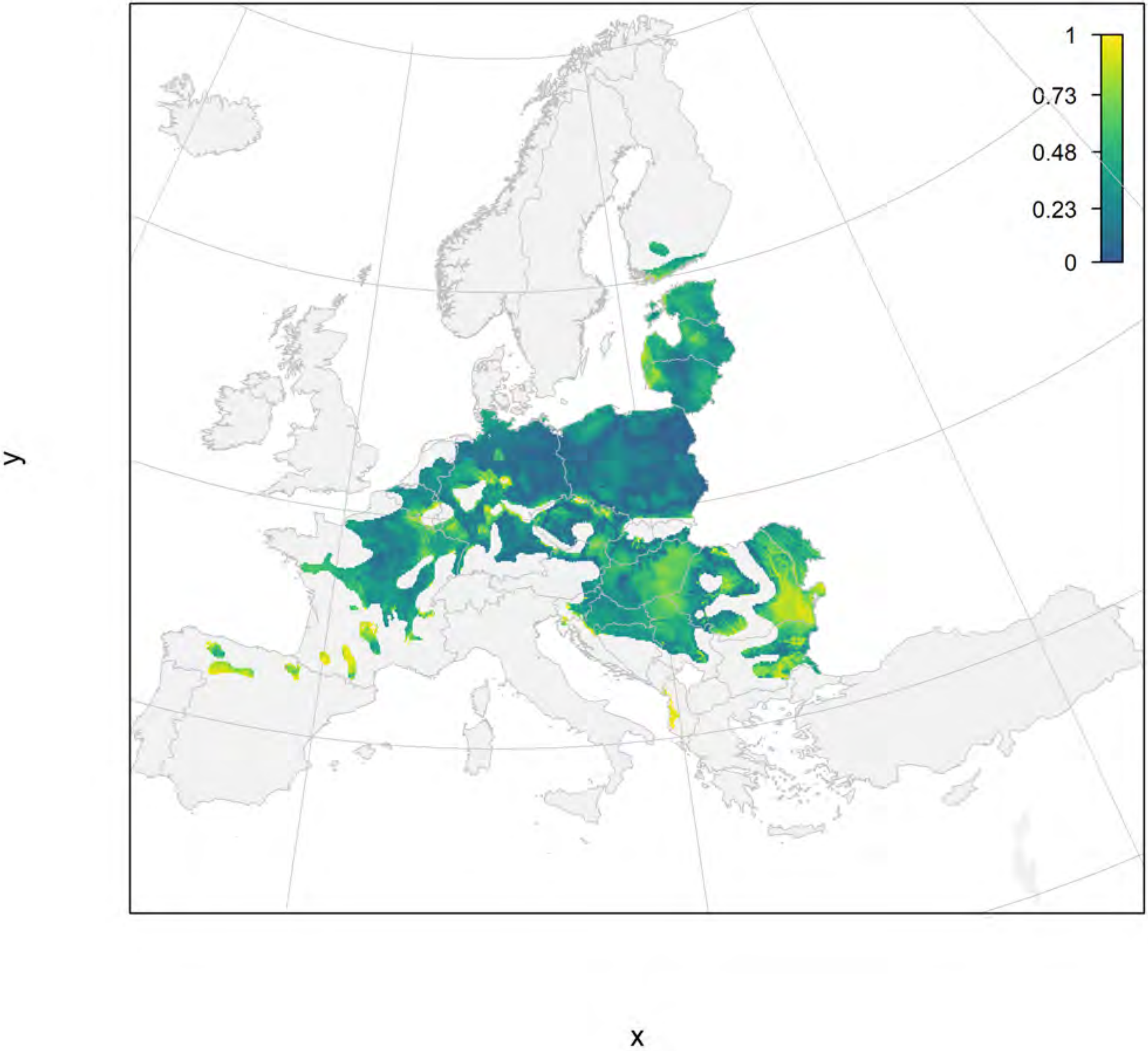

#### Ulmus laevis_filt current marginality

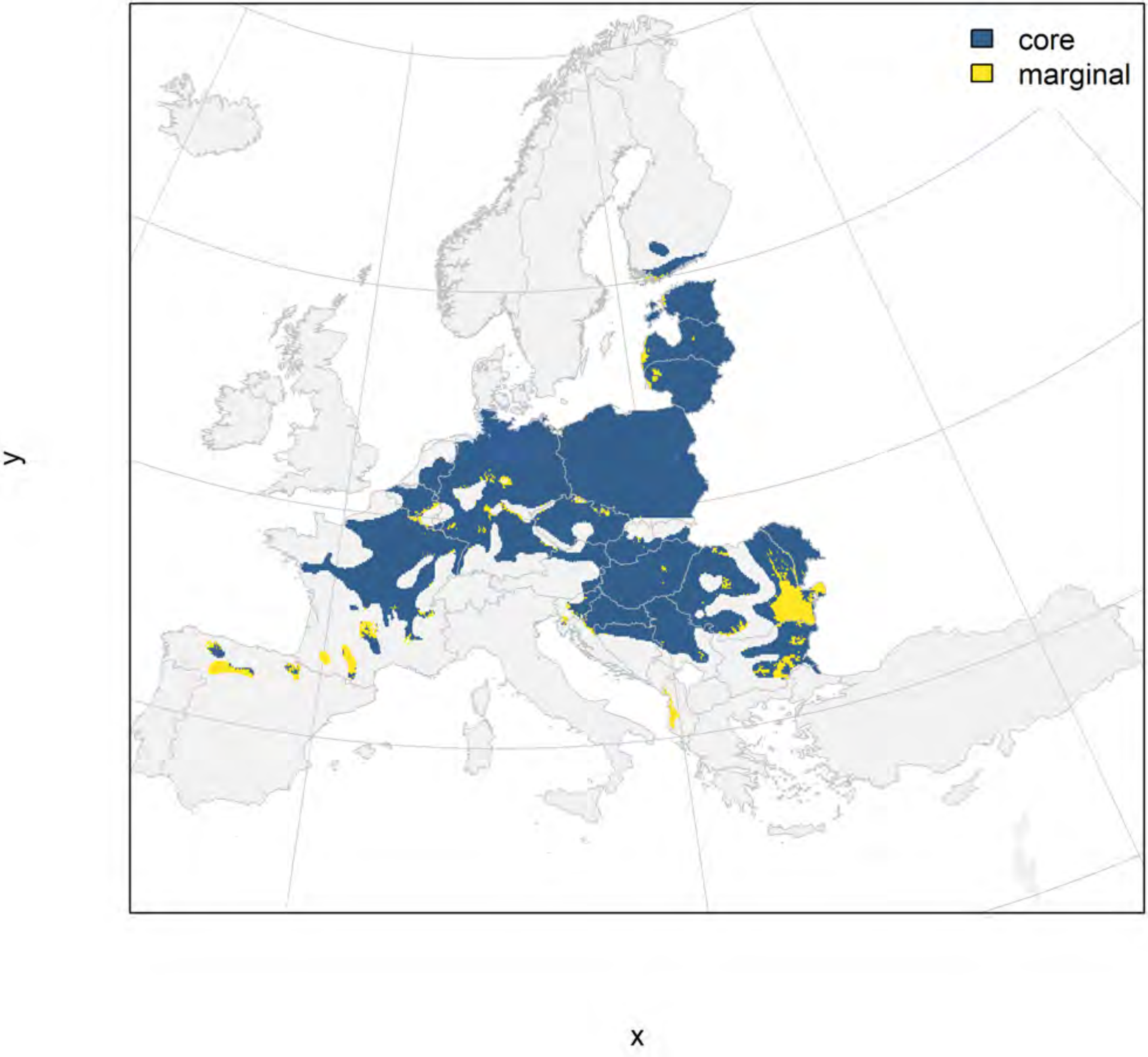

### Ulmus laevis_filt future marginality

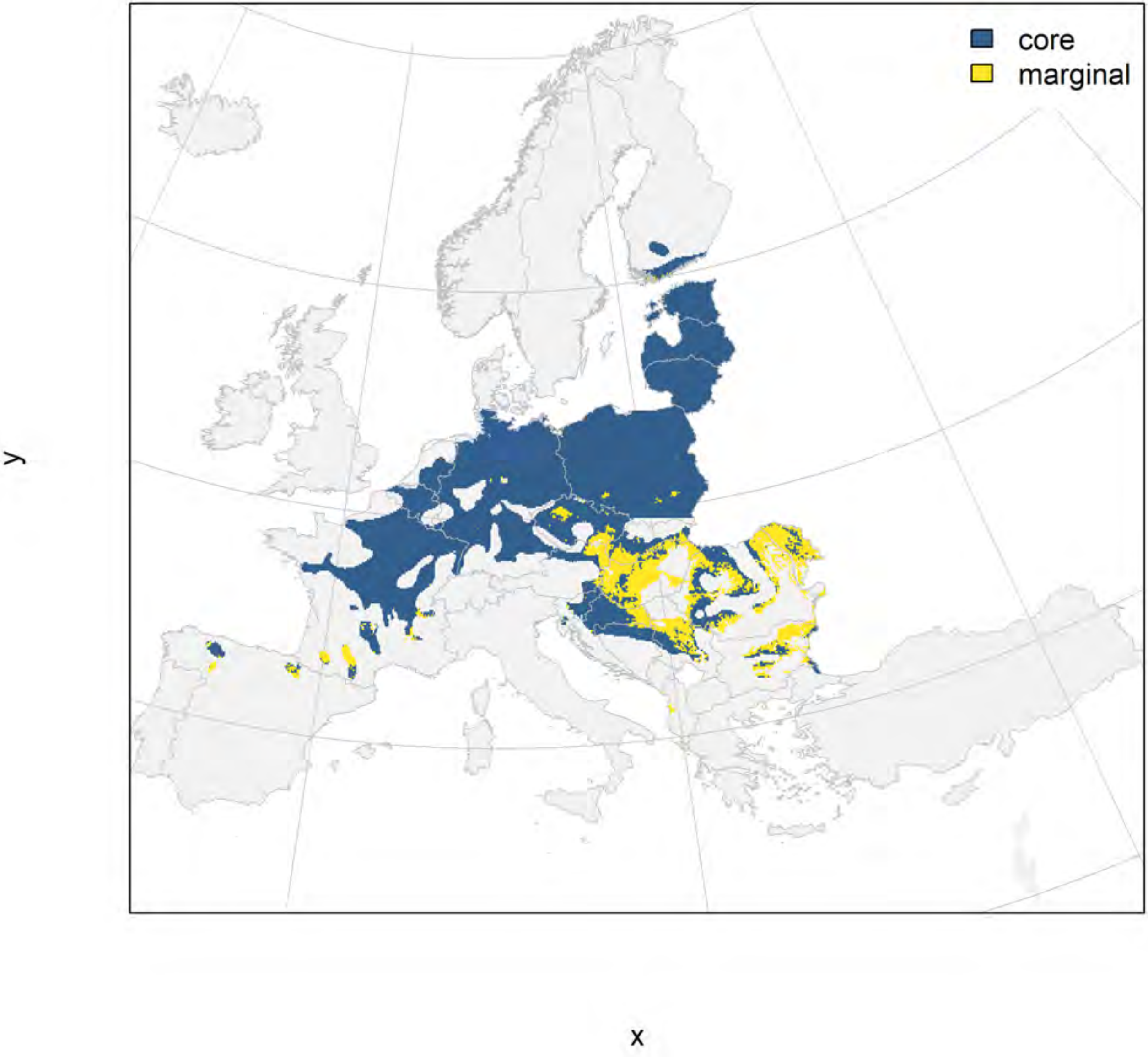

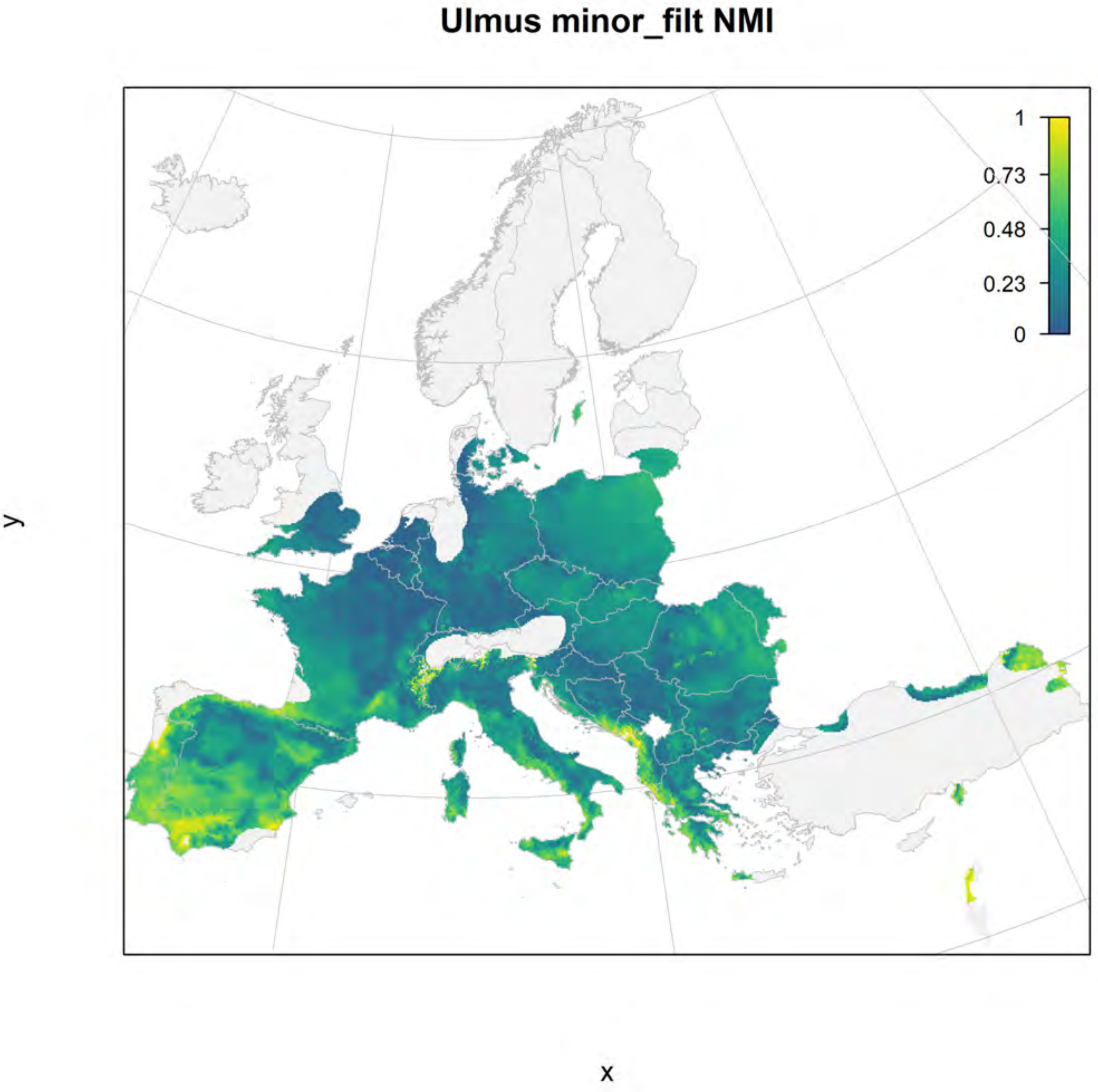
Ulmus minor_filt NMI
1
0.73
0.48
0.23
0
y
x

### Ulmus minor_filt current marginality

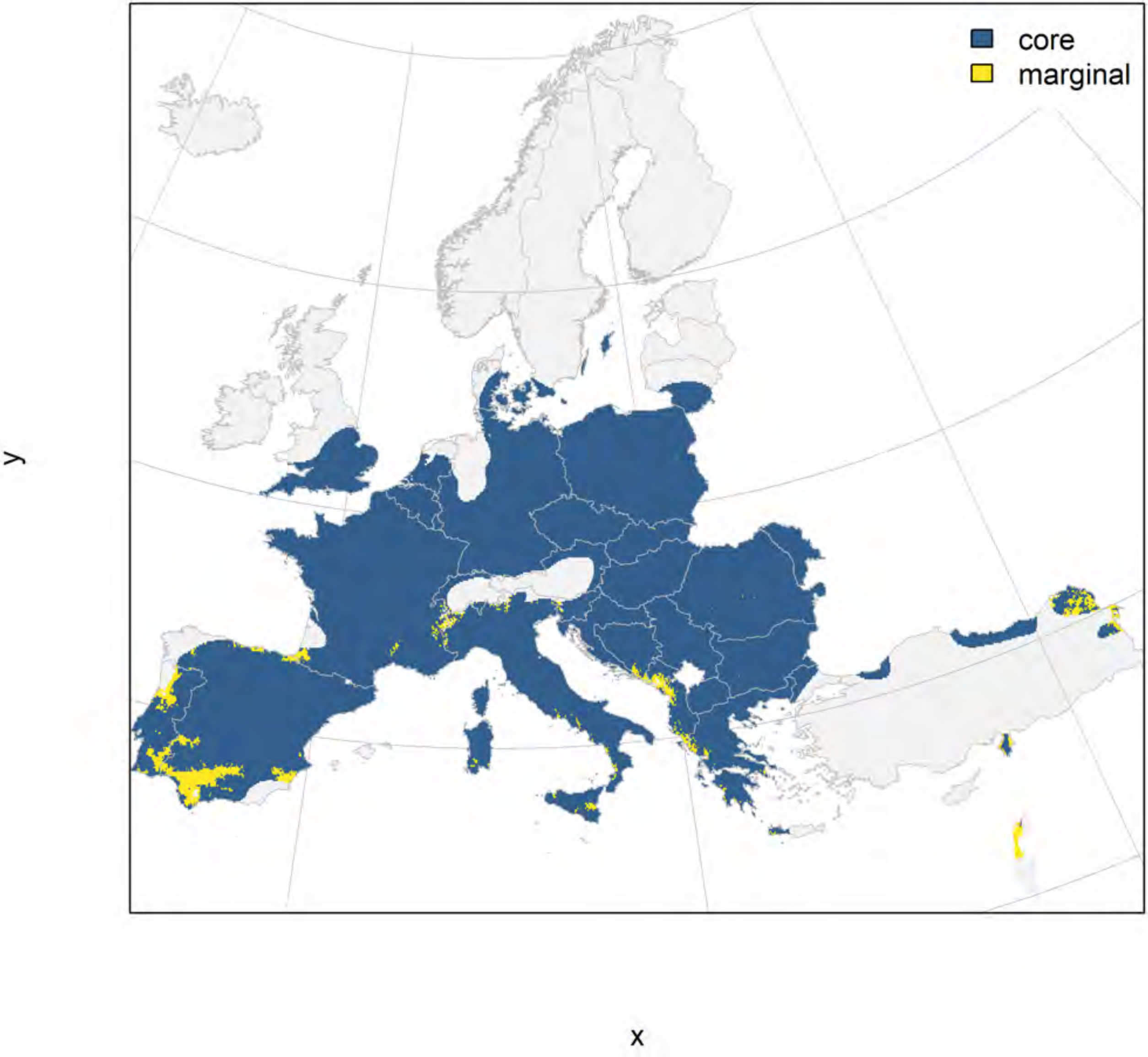

#### Ulmus minor_filt future marginality

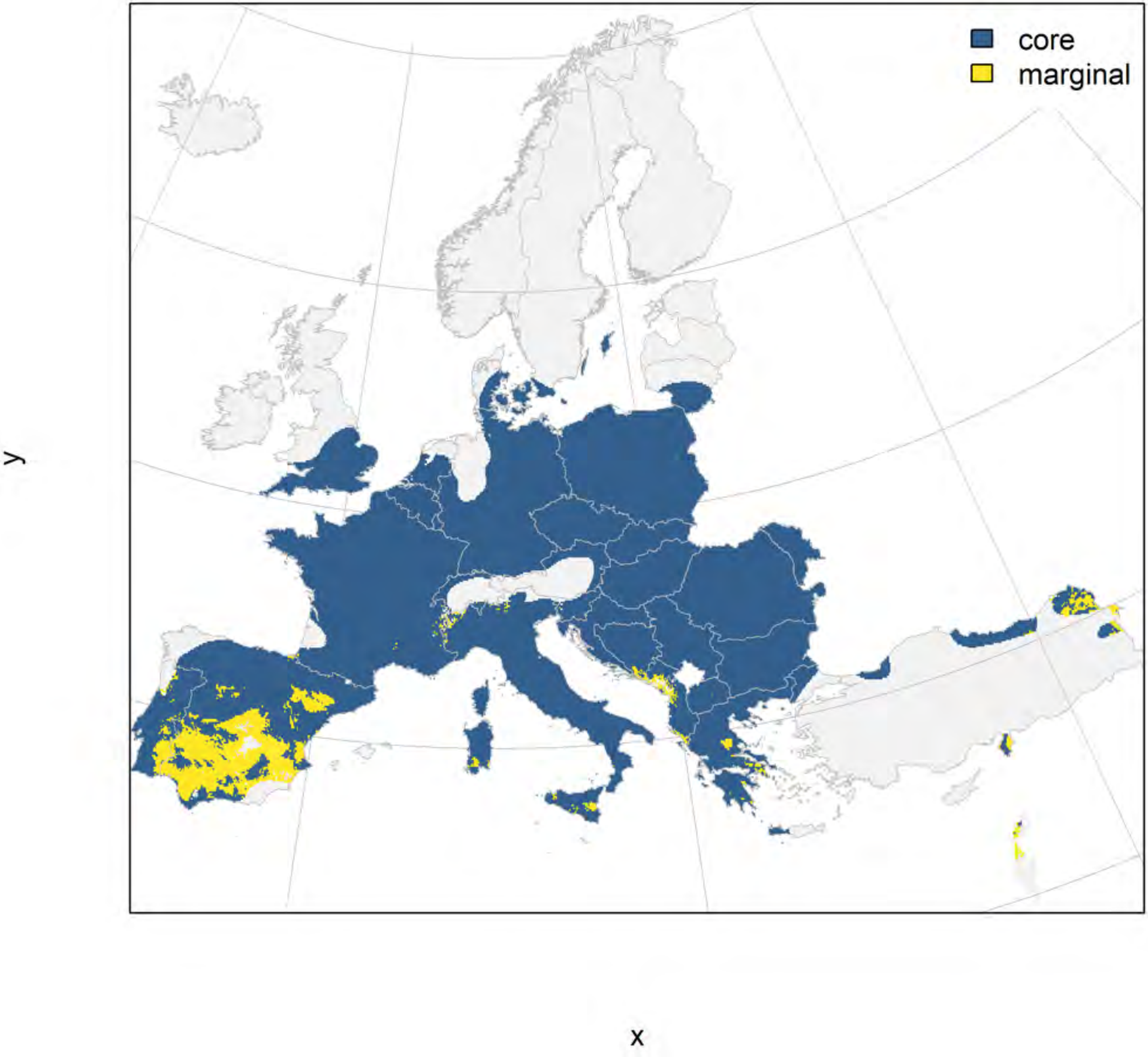

#### Aesculus hippocastanum_filt NMI

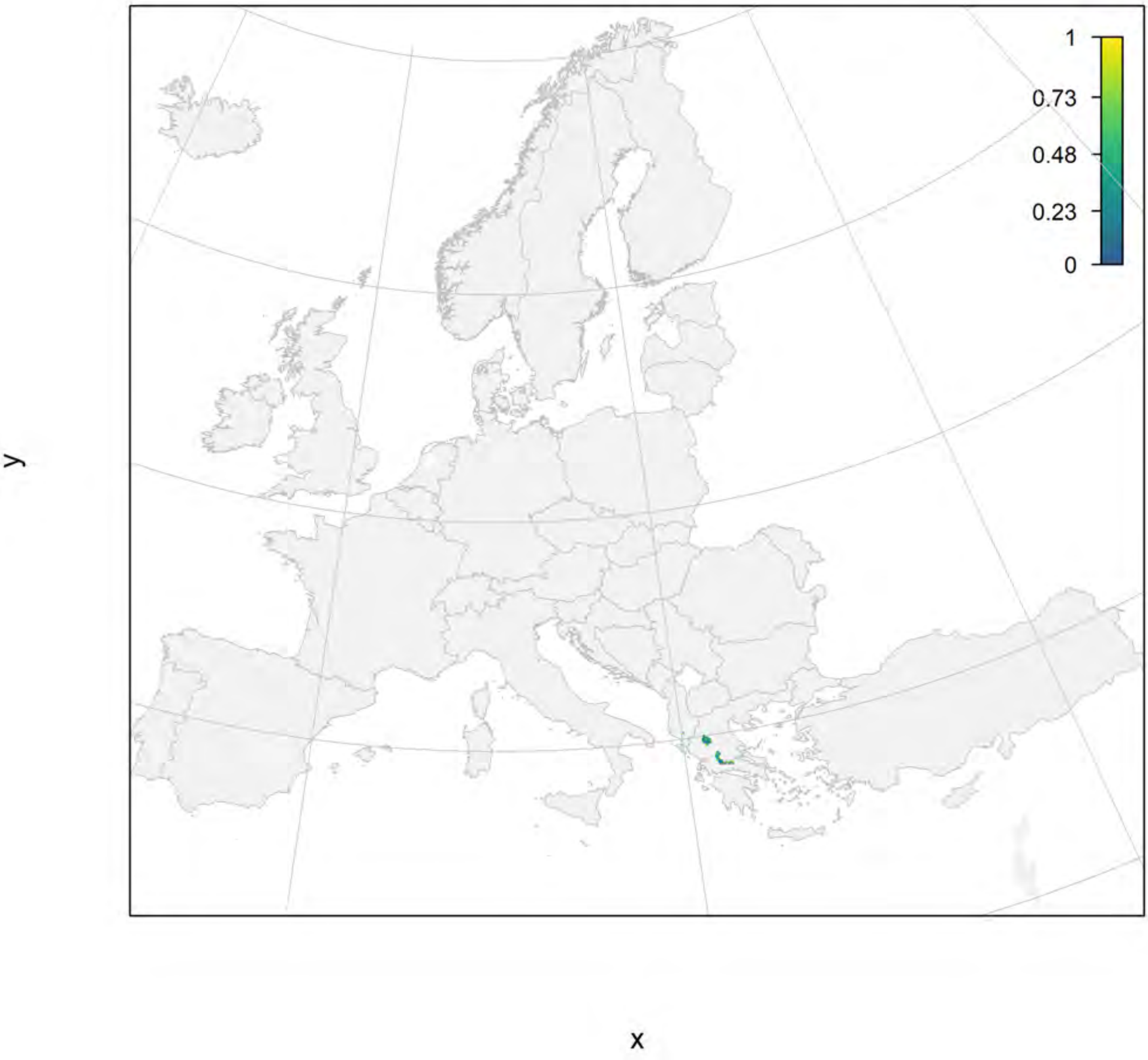

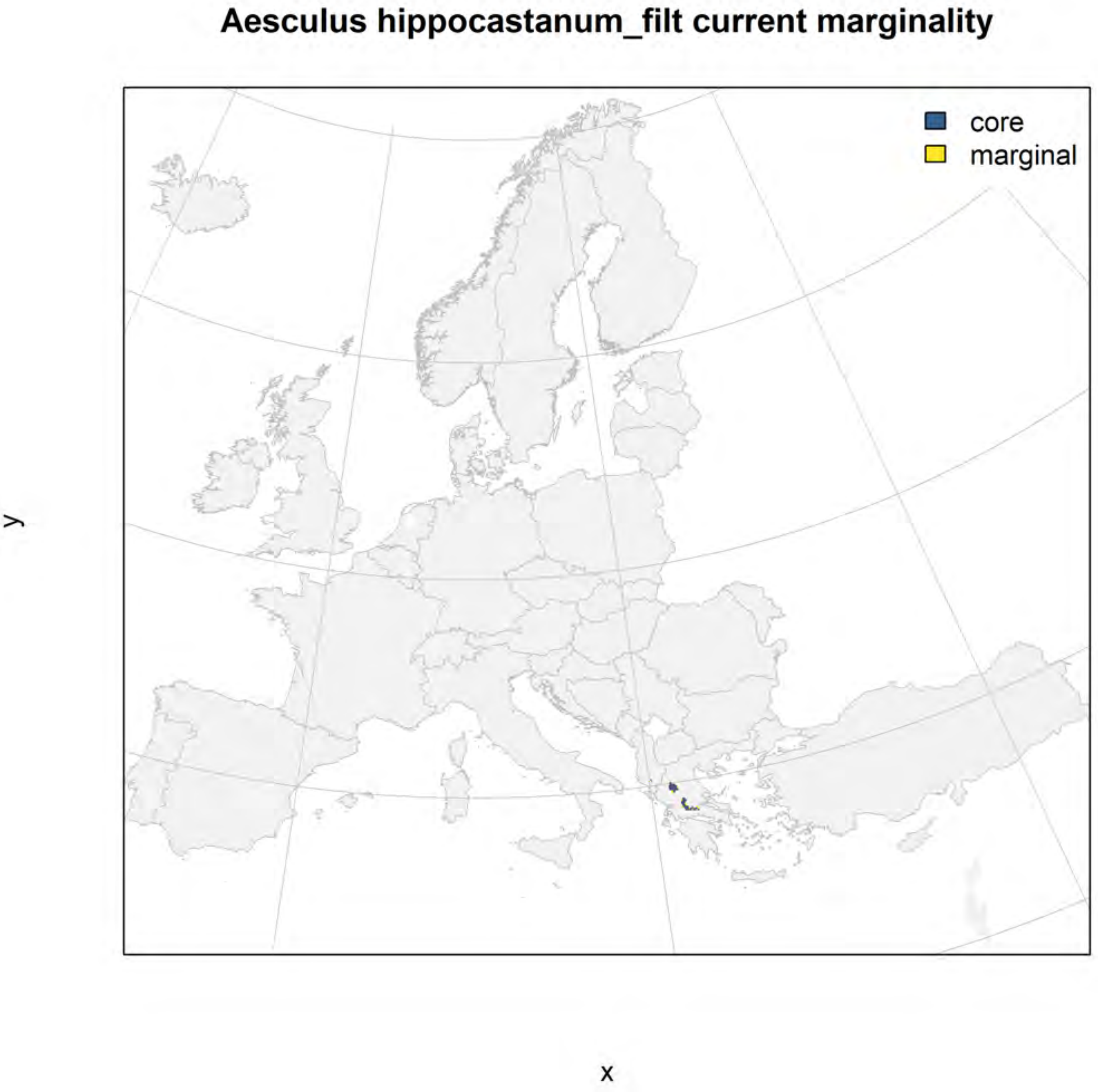
Aesculus hippocastanum_filt current marginality
core
marginal
y
x

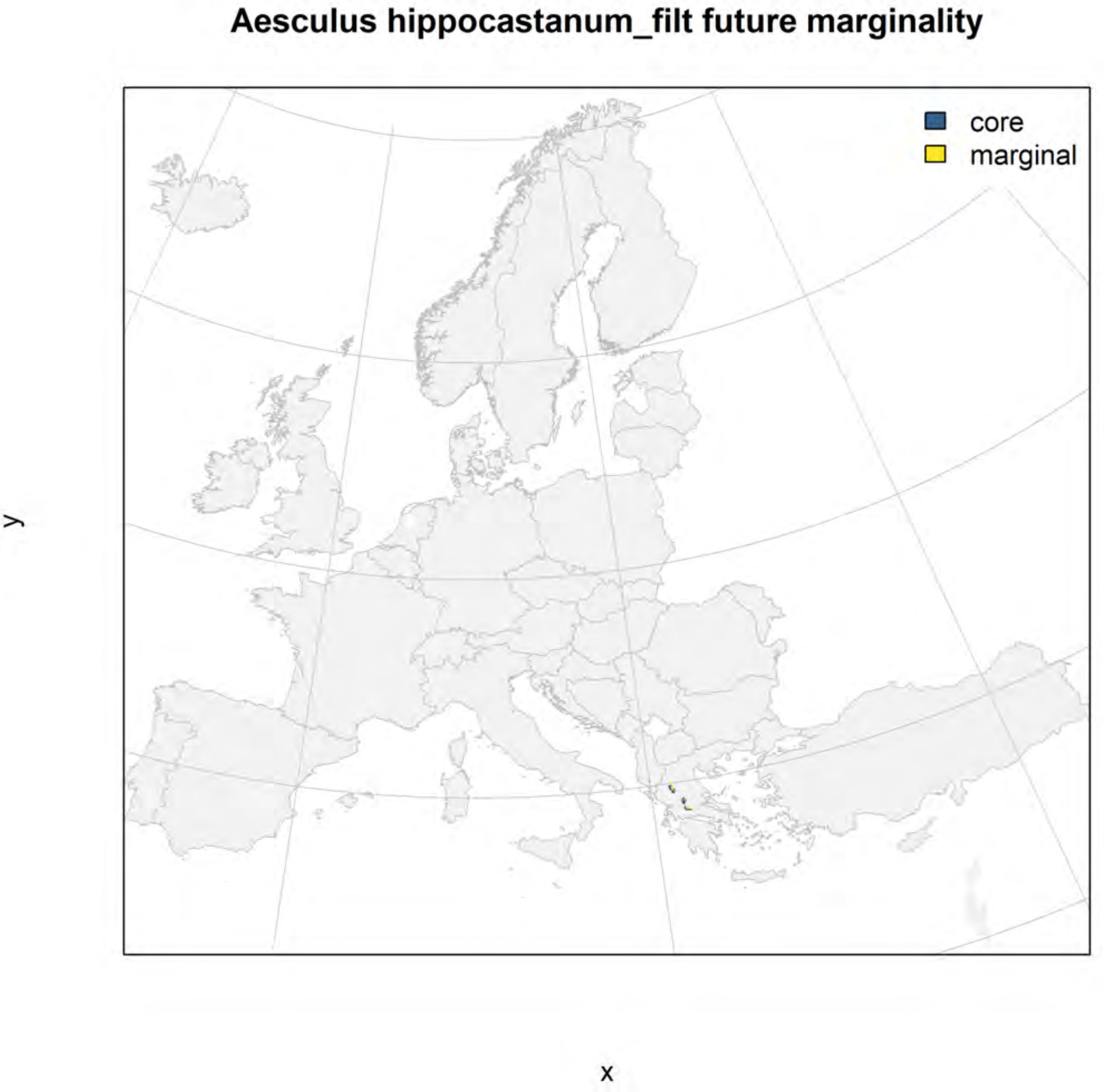
Aesculus hippocastanum_filt future marginality
core
marginal
y
x

### Arbutus unedo_filt NMI

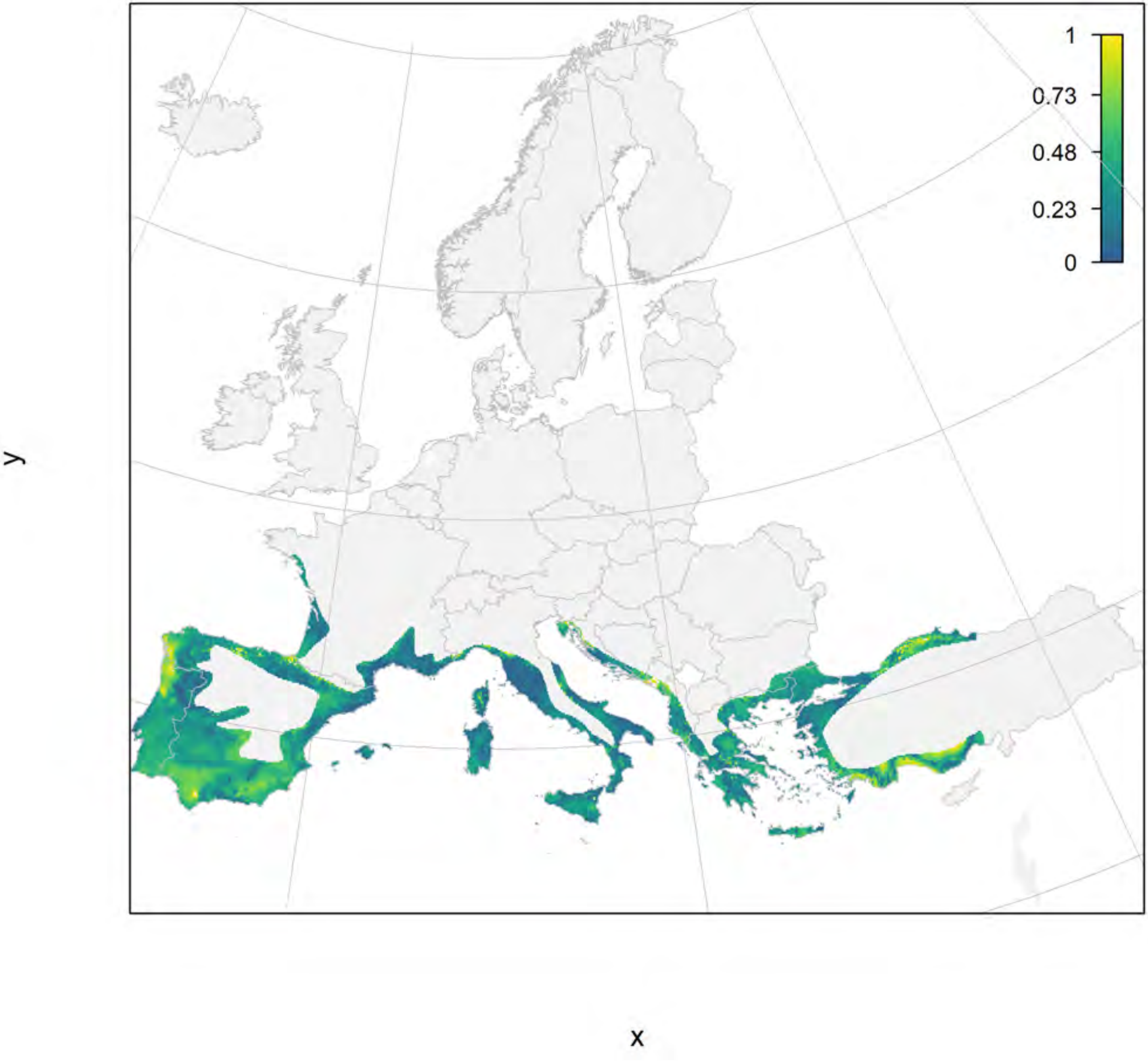

#### Arbutus unedo_filt current marginality

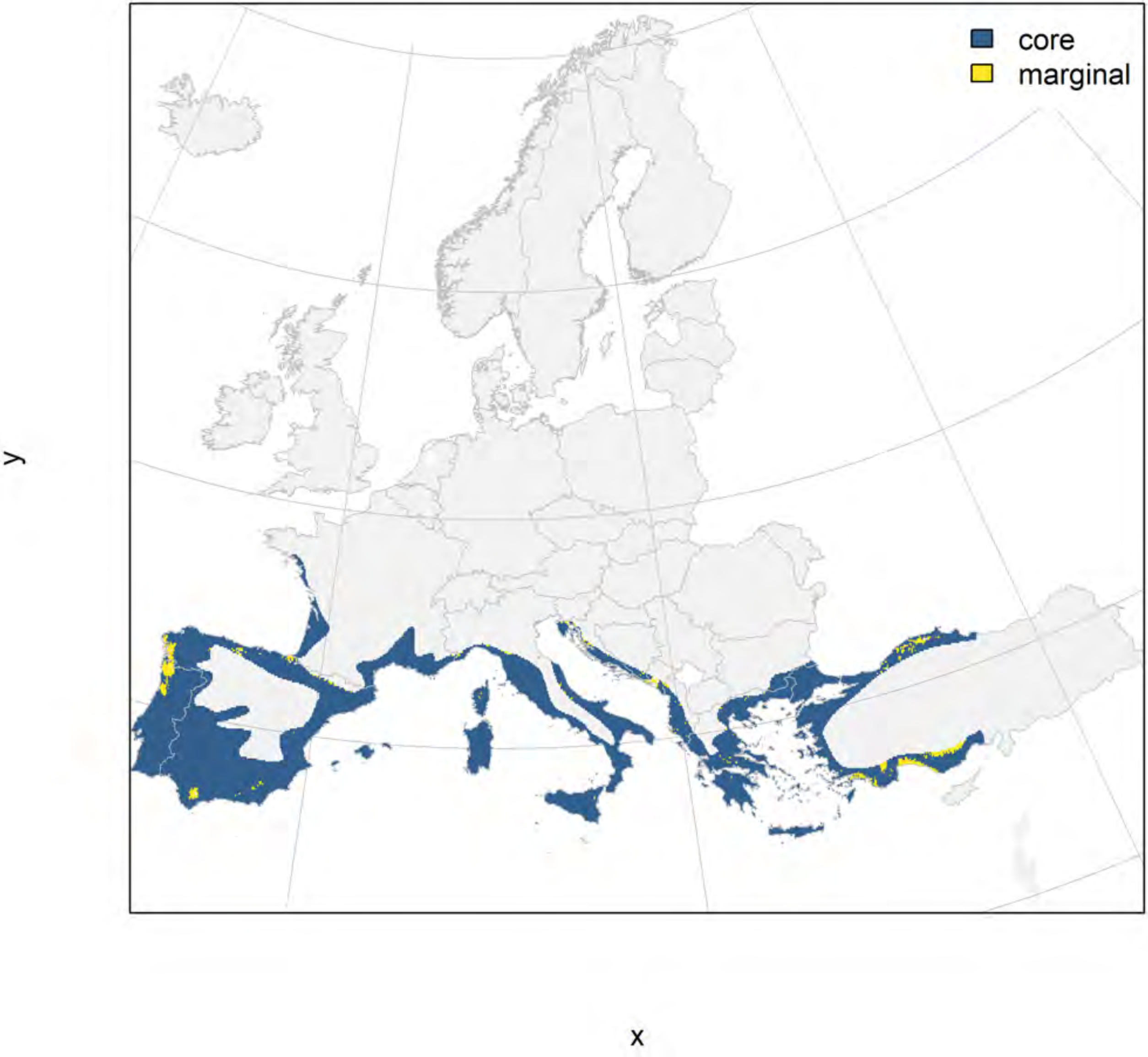

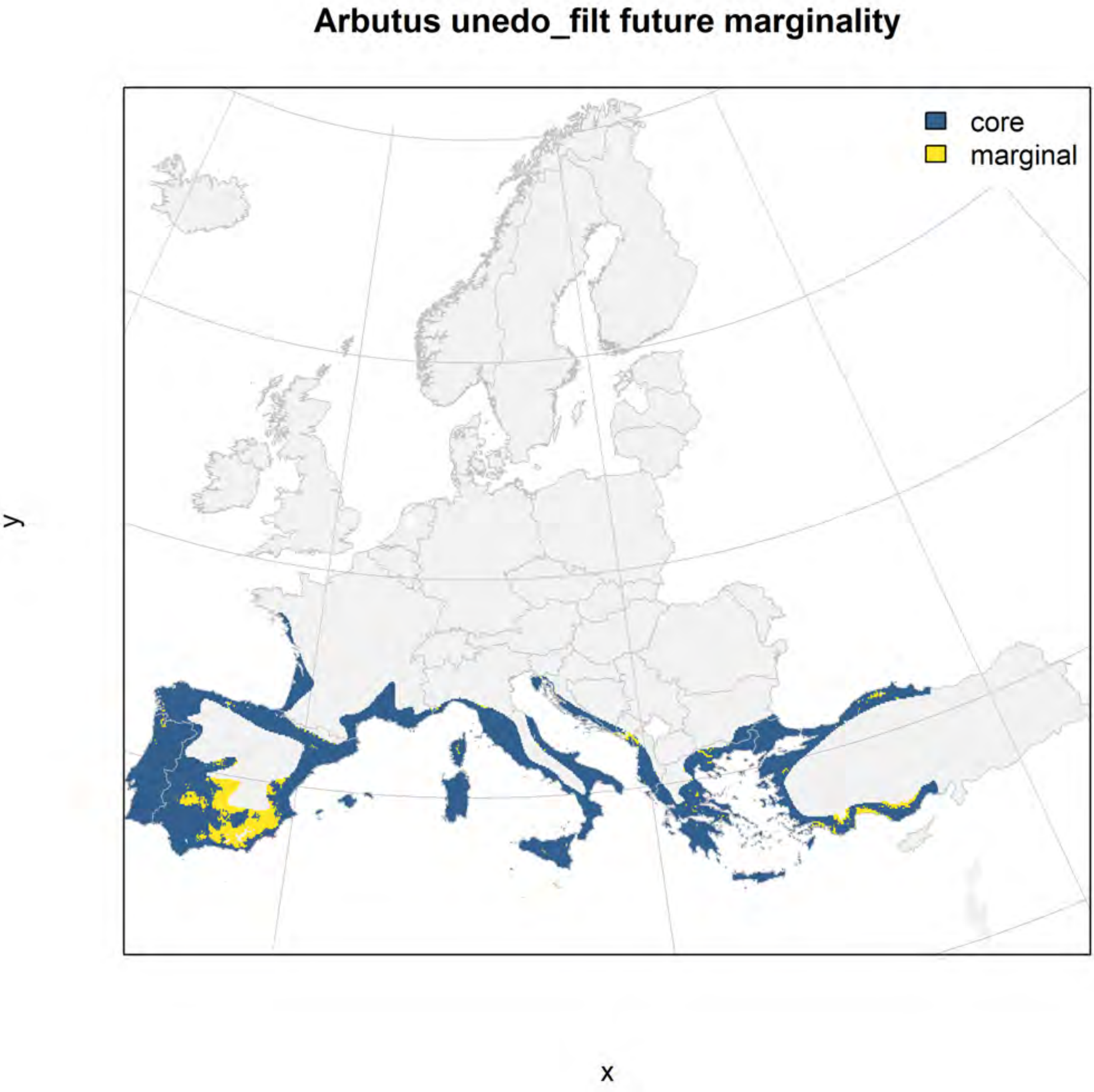

Arbutus unedo_filt future marginality
core
marginal
y
x

### Buxus sempervirens_filt NMI

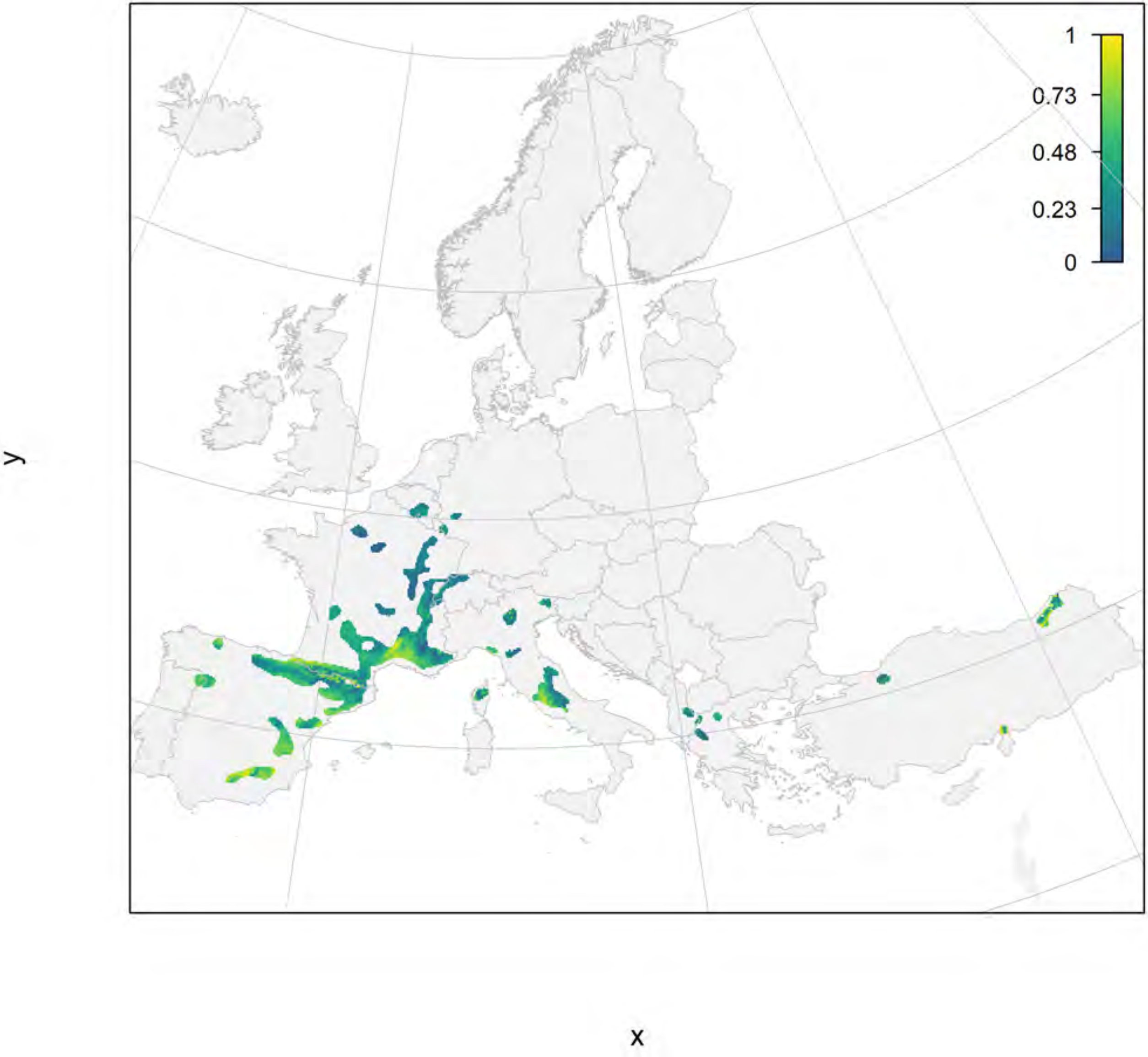

### Buxus sempervirens_filt current marginality

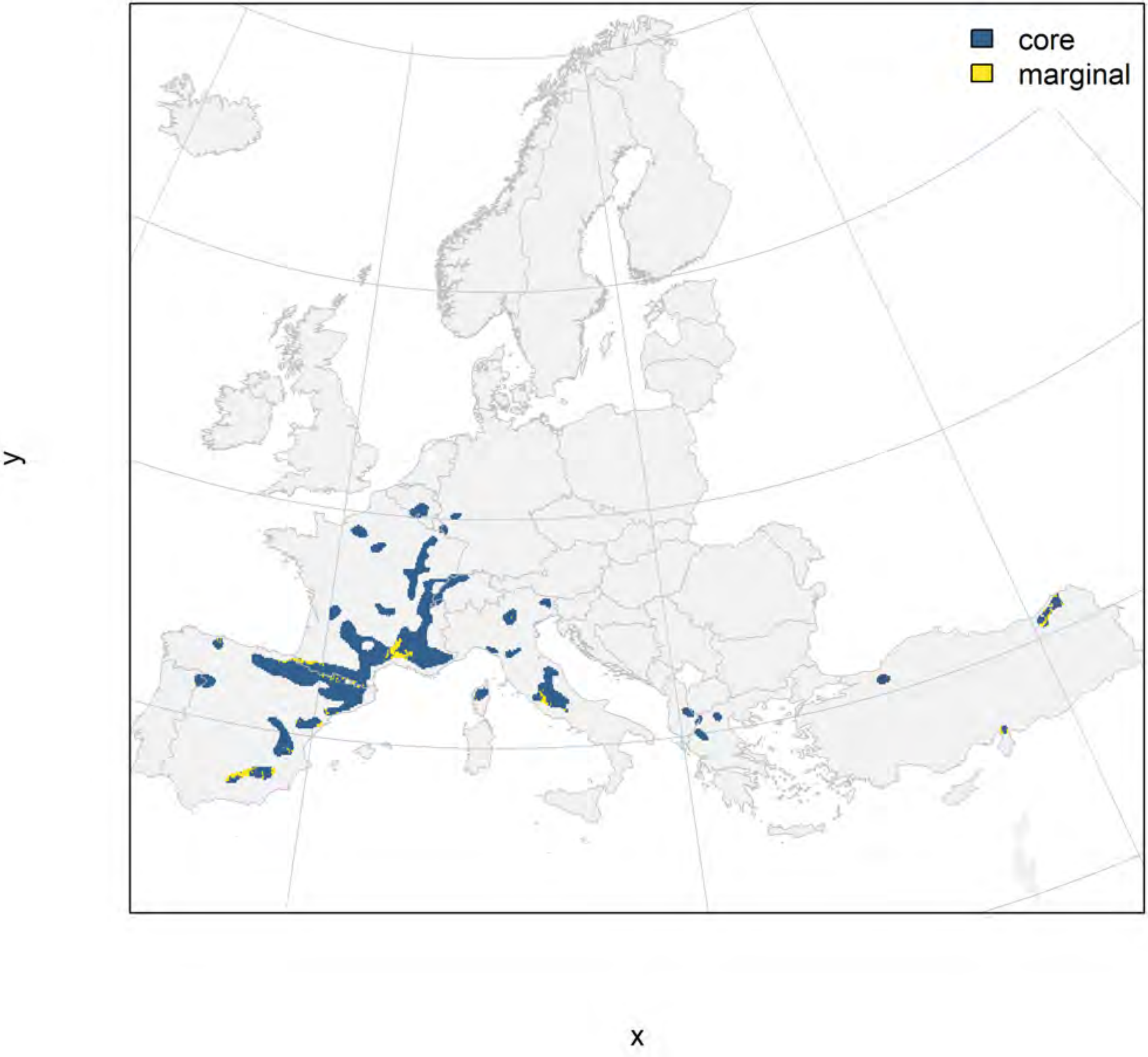

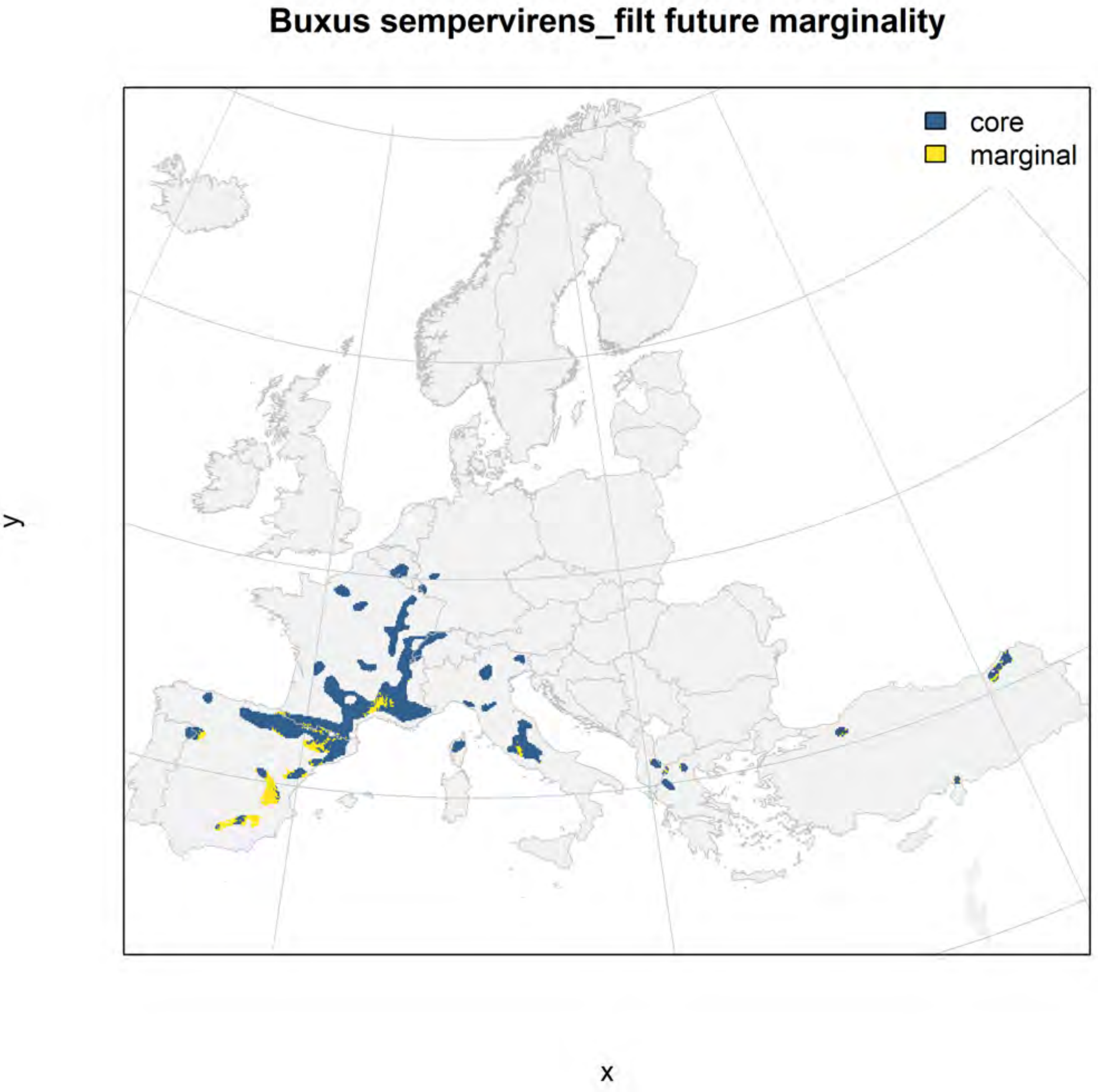
Buxus sempervirens_filt future marginality
core
marginal
y
x

### Carpinus orientalis_filt NMI

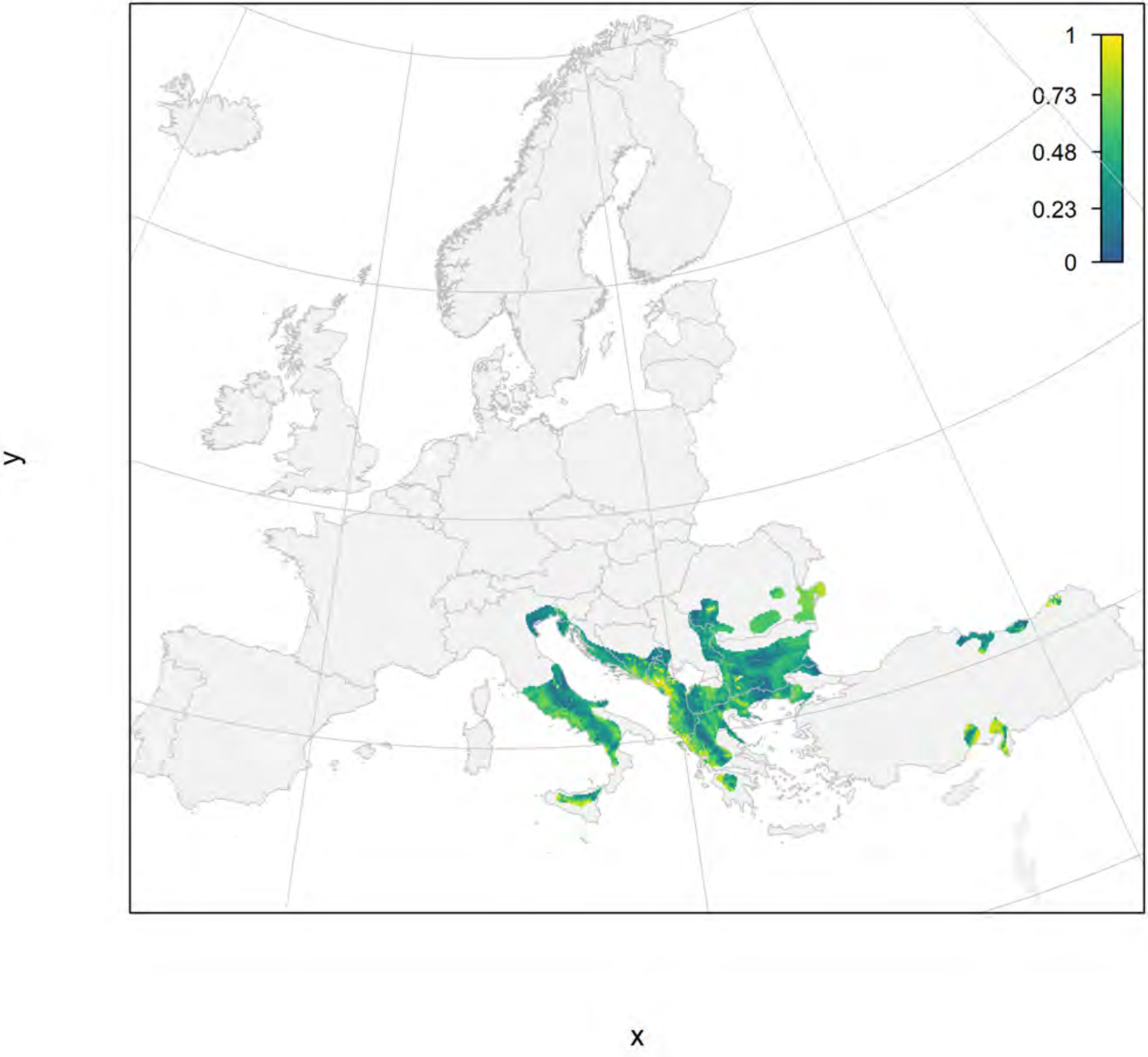

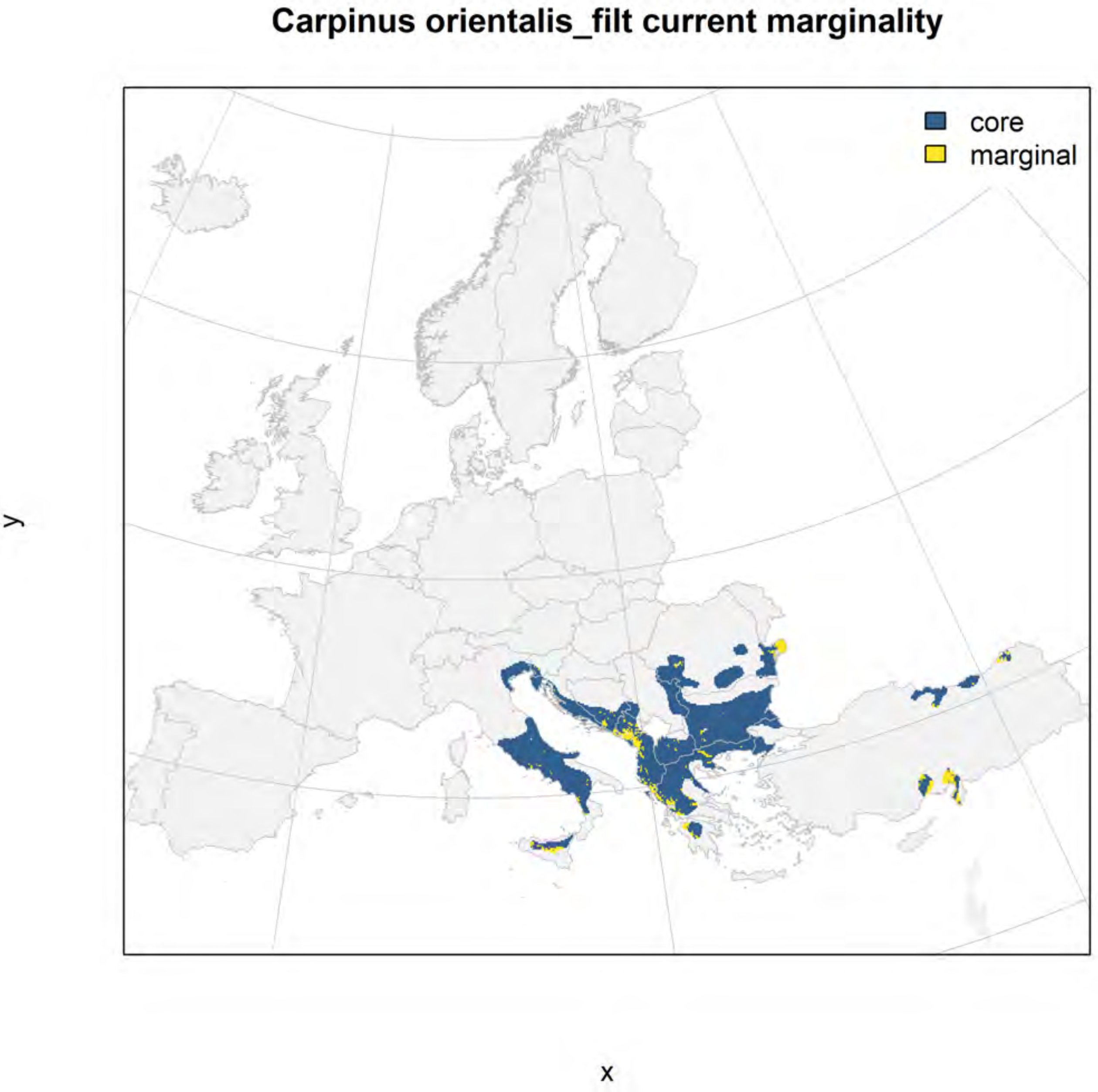
Carpinus orientalis_filt current marginality
core
marginal
y
x

#### Carpinus orientalis_filt future marginality

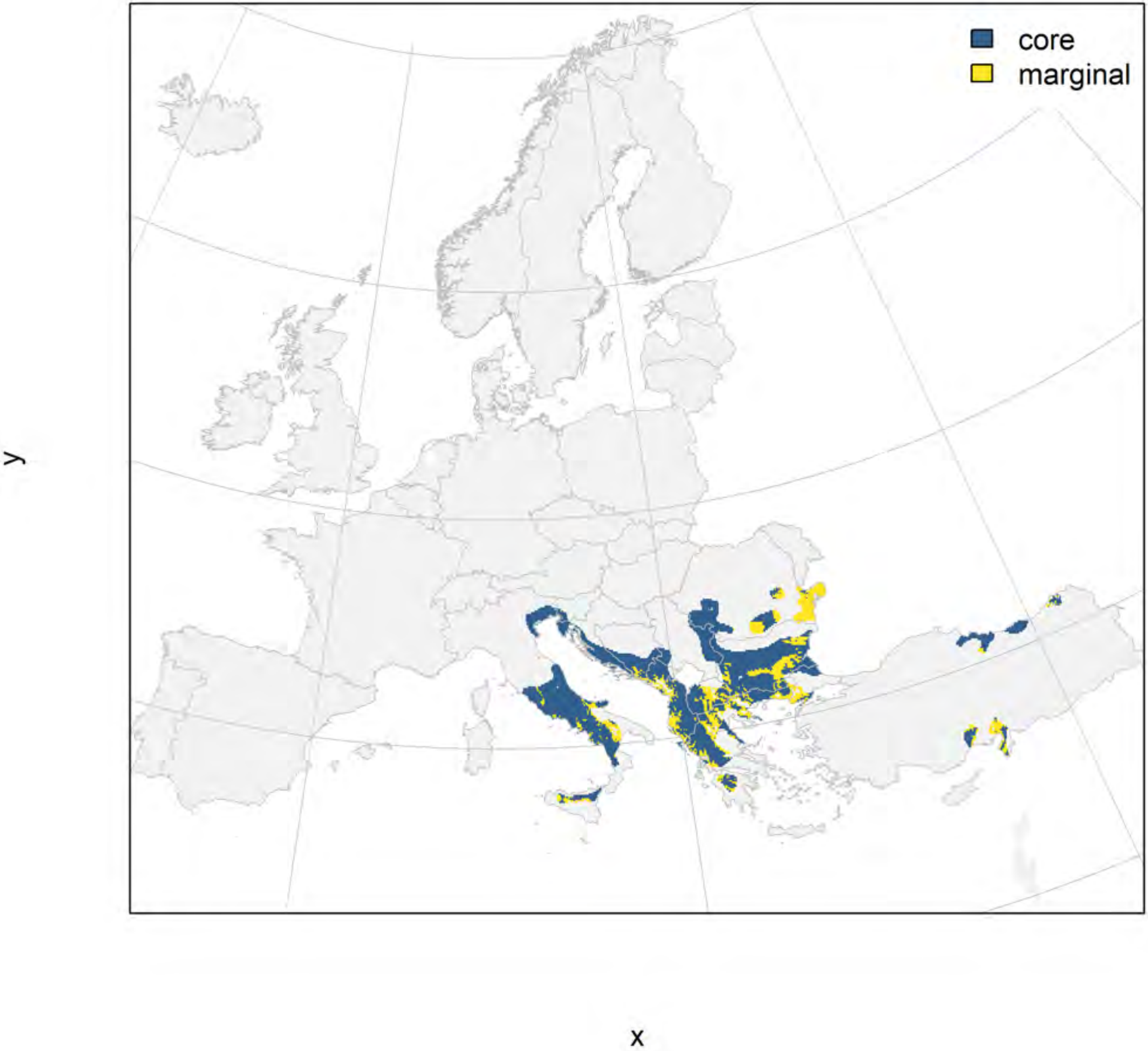

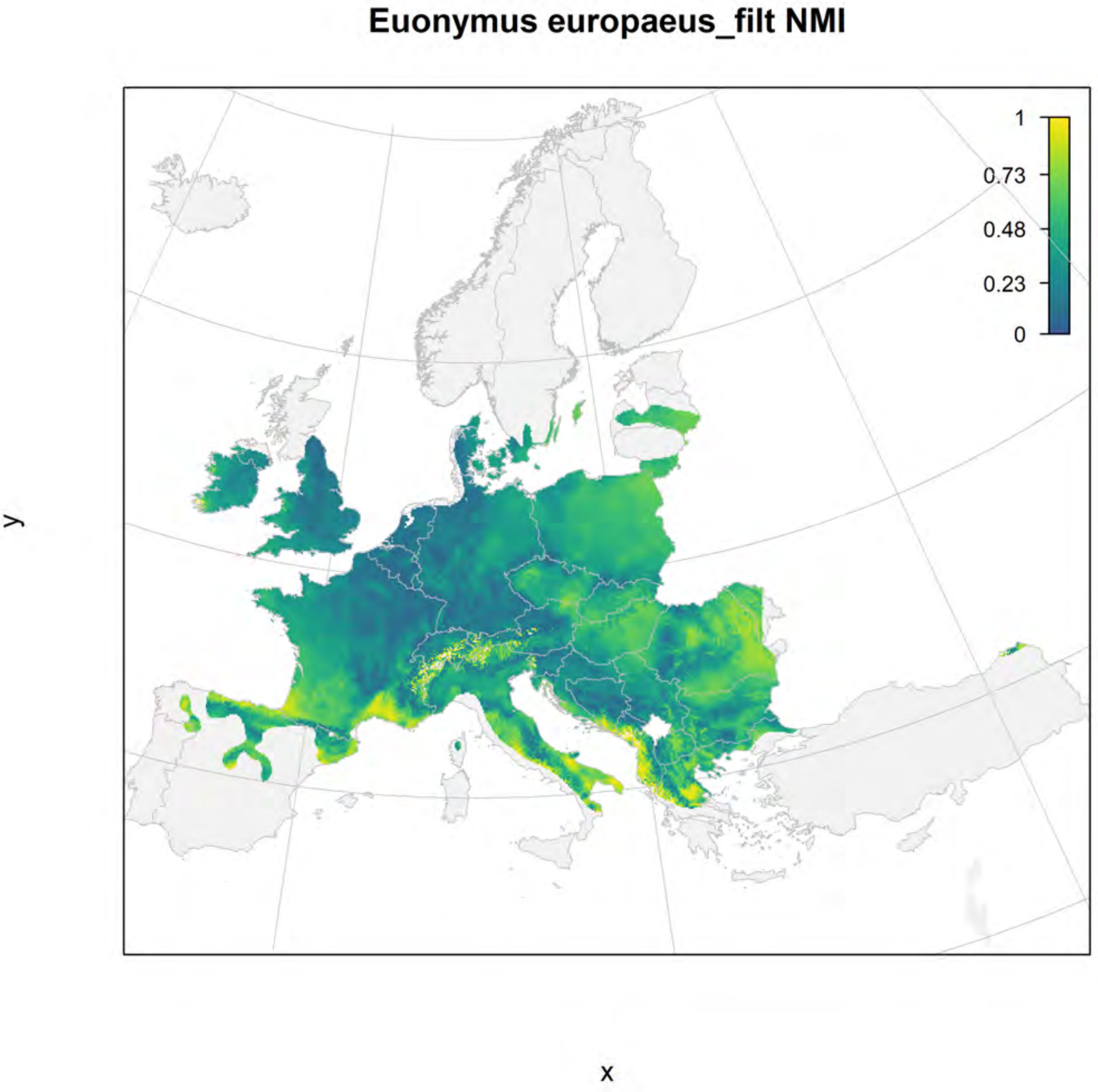
Euonymus europaeus_filt NMI
1
0.73
0.48
0.23
0
y
x

#### Euonymus europaeus_filt current marginality

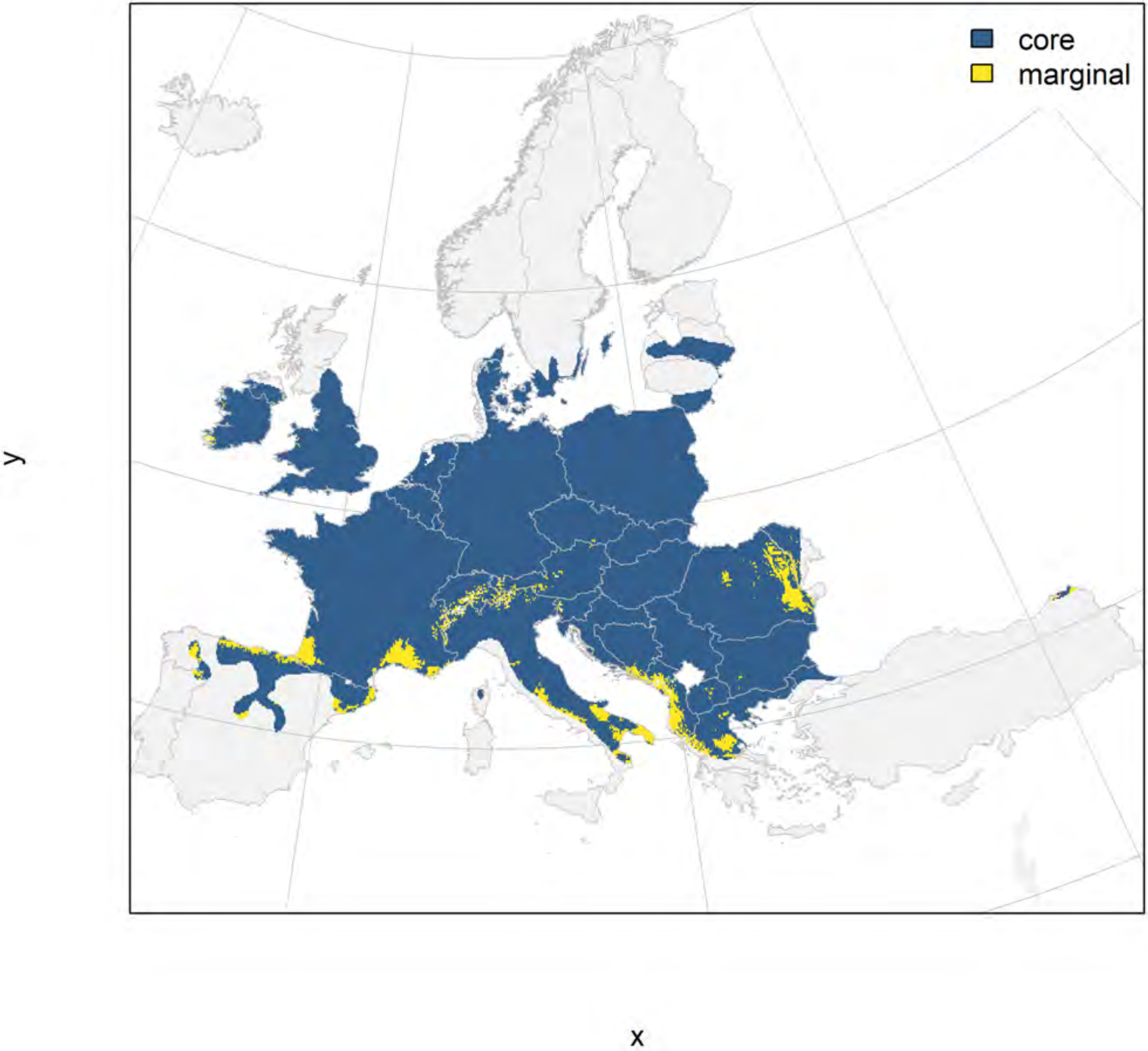

Euonymus europaeus_filt future marginality

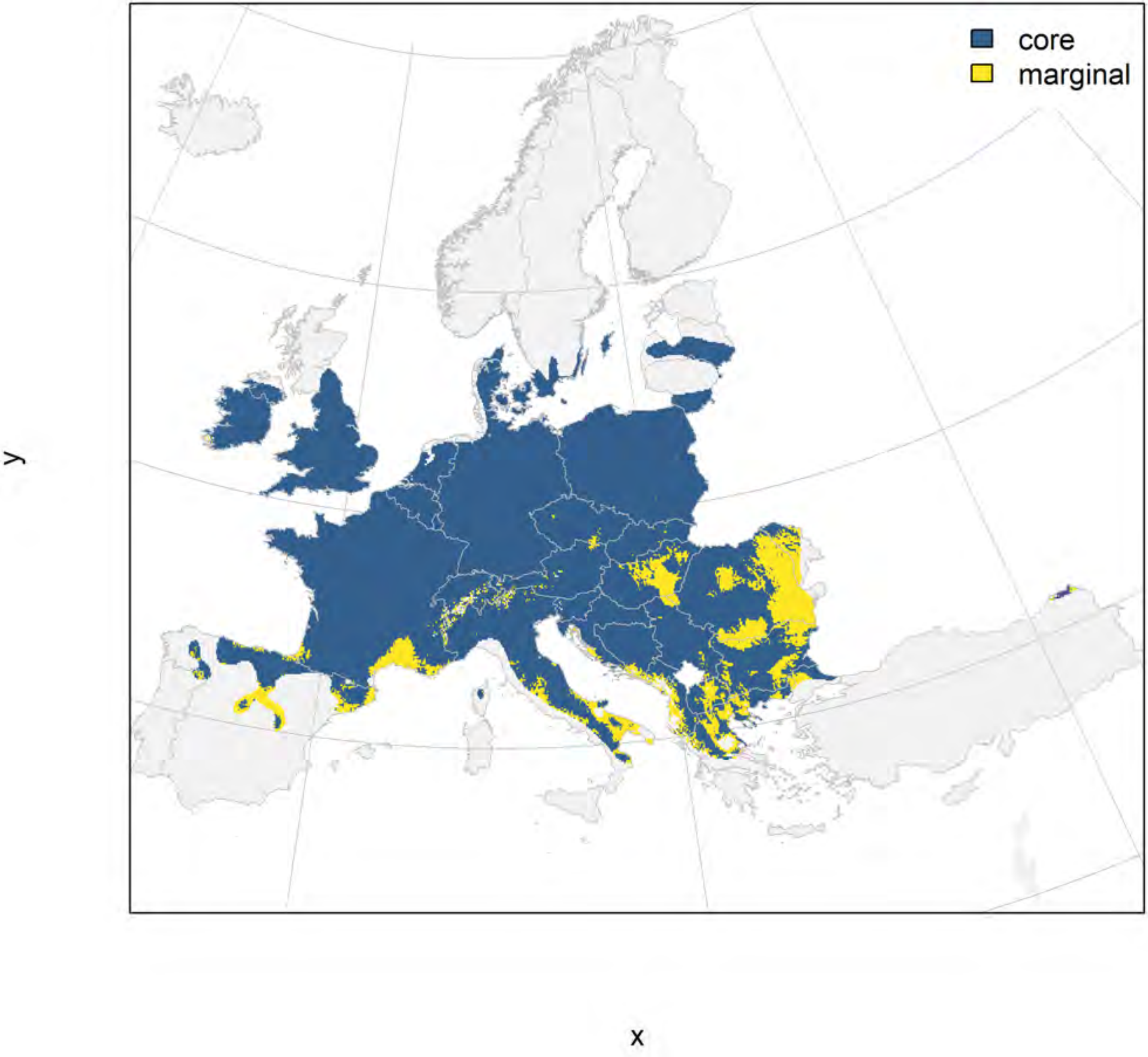

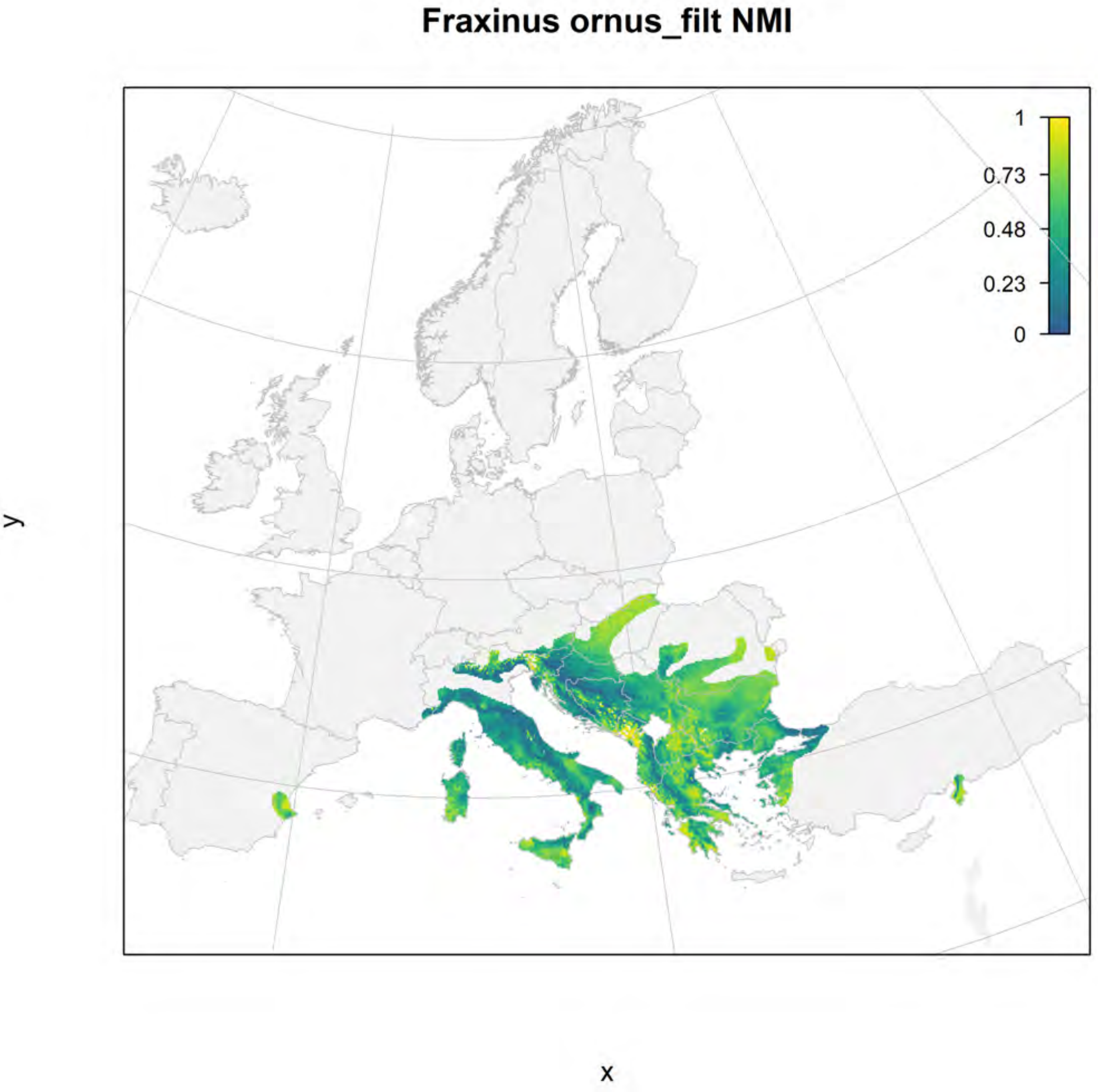

Fraxinus ornus_filt NMI
1
0.73
0.48
0.23
0
y
x

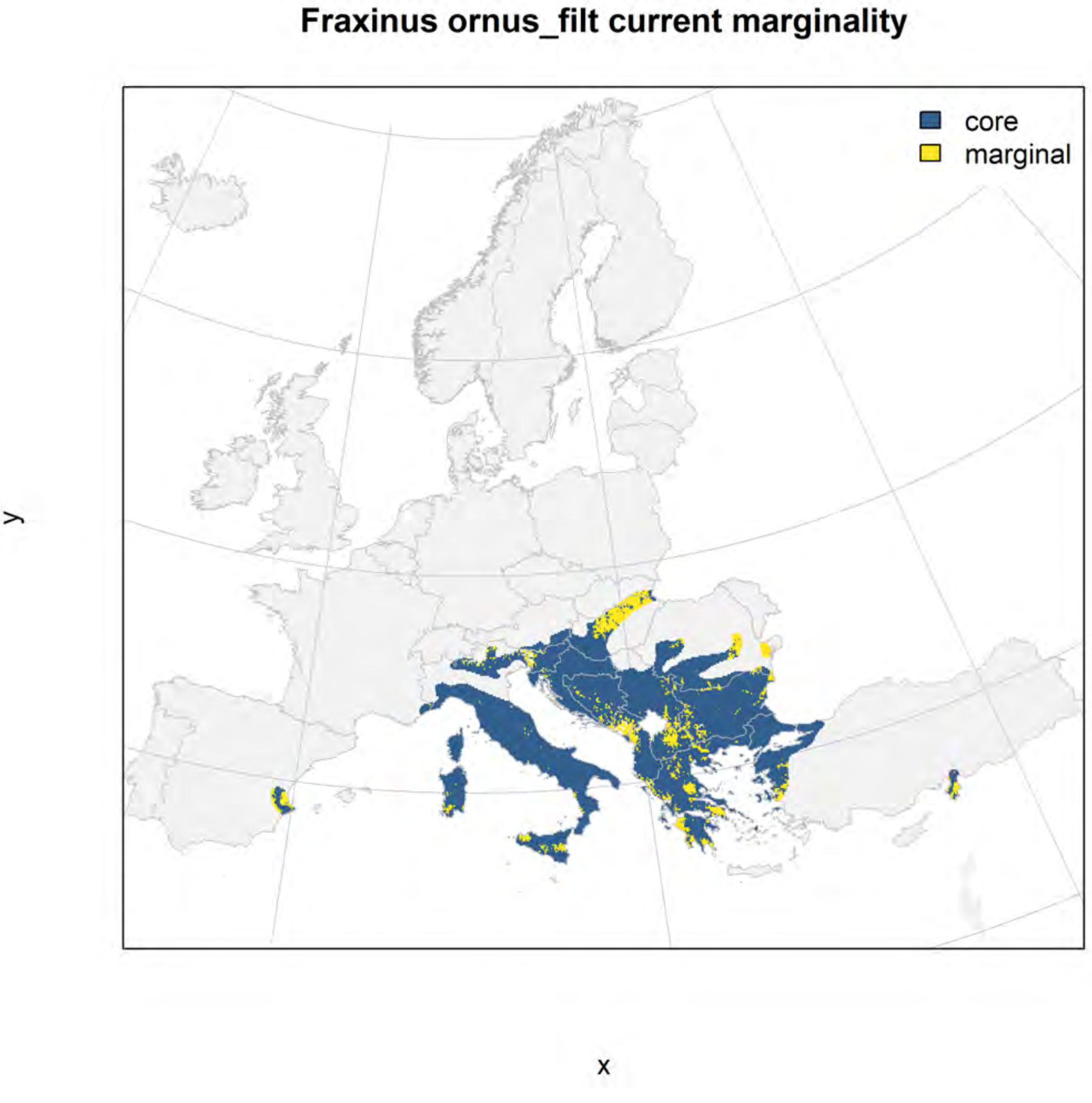
Fraxinus ornus_filt current marginality
core
marginal
y
x

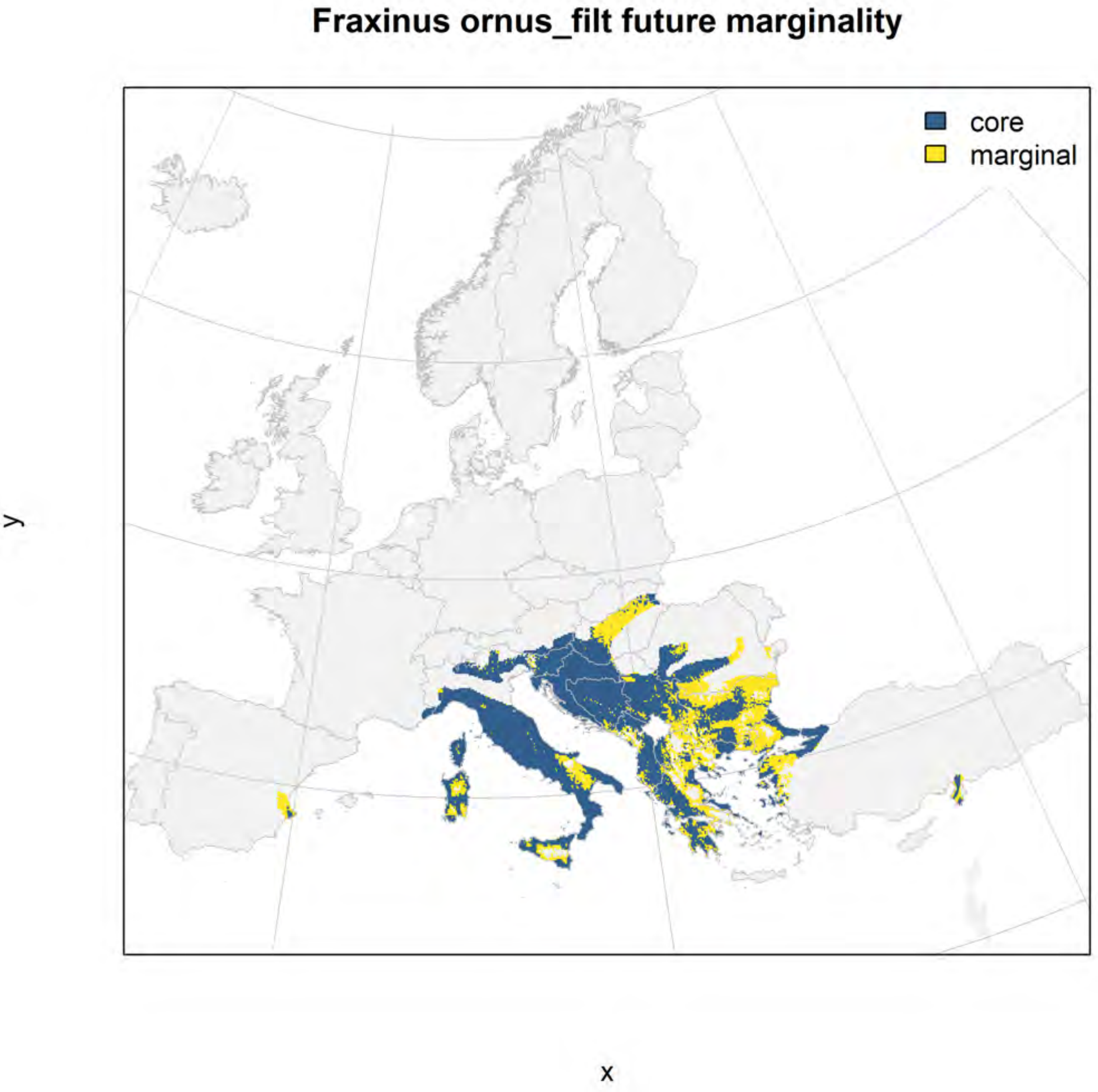
Fraxinus ornus_filt future marginality
core
marginal
y
x

#### Juniperus phoenicea_filt NMI

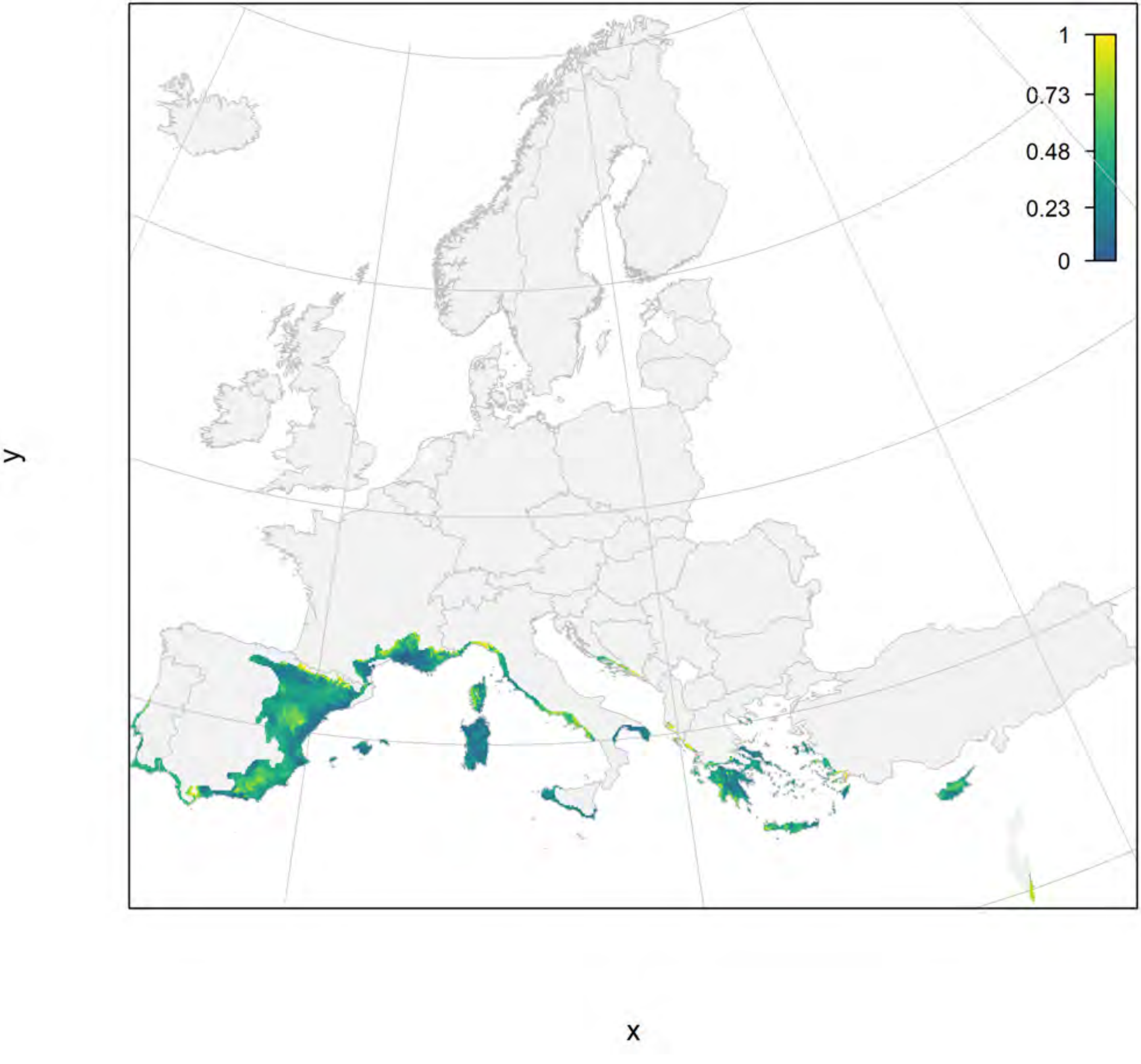

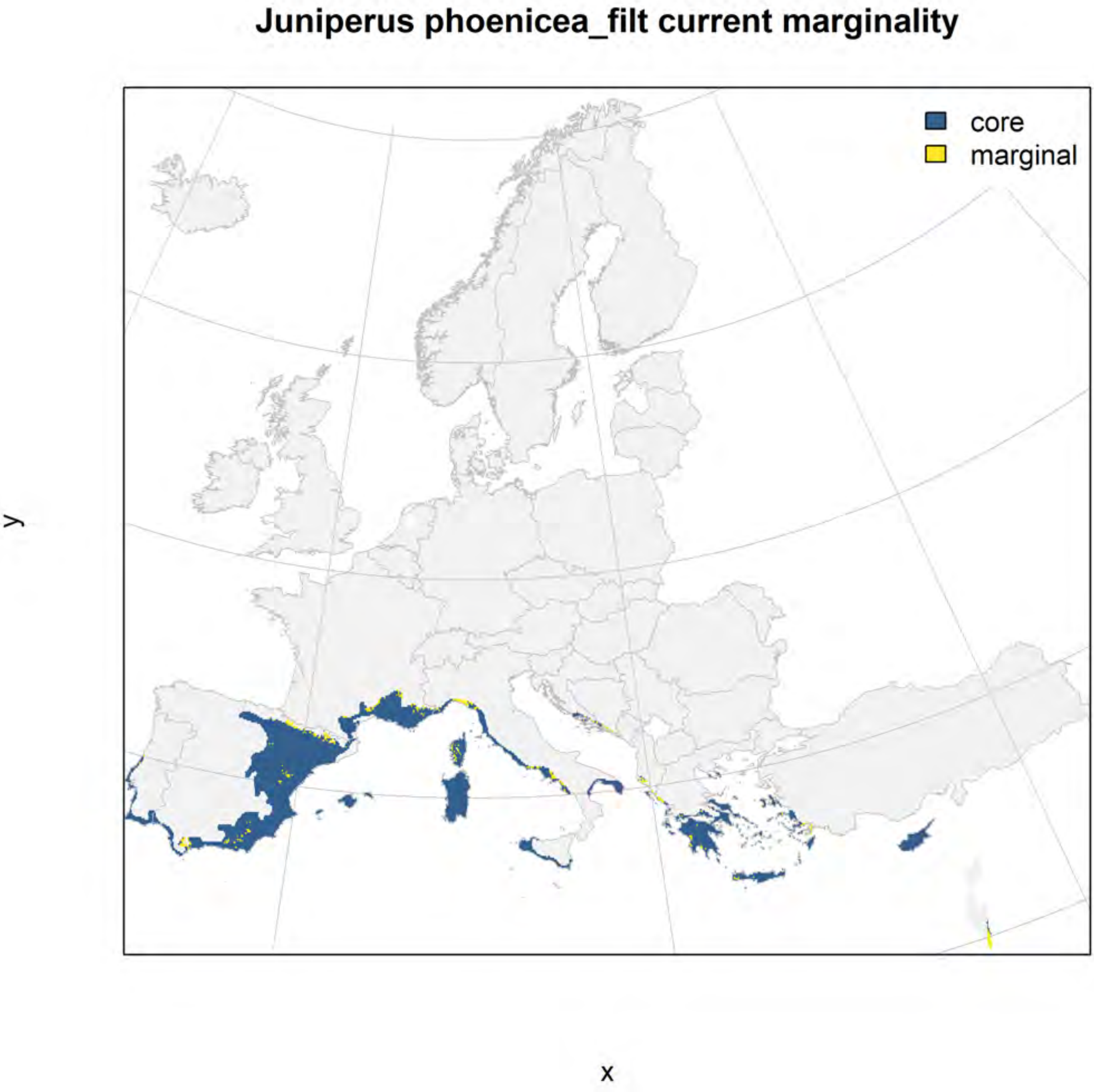
Juniperus phoenicea_filt current marginality
core
marginal
y
x

#### Juniperus phoenicea_filt future marginality

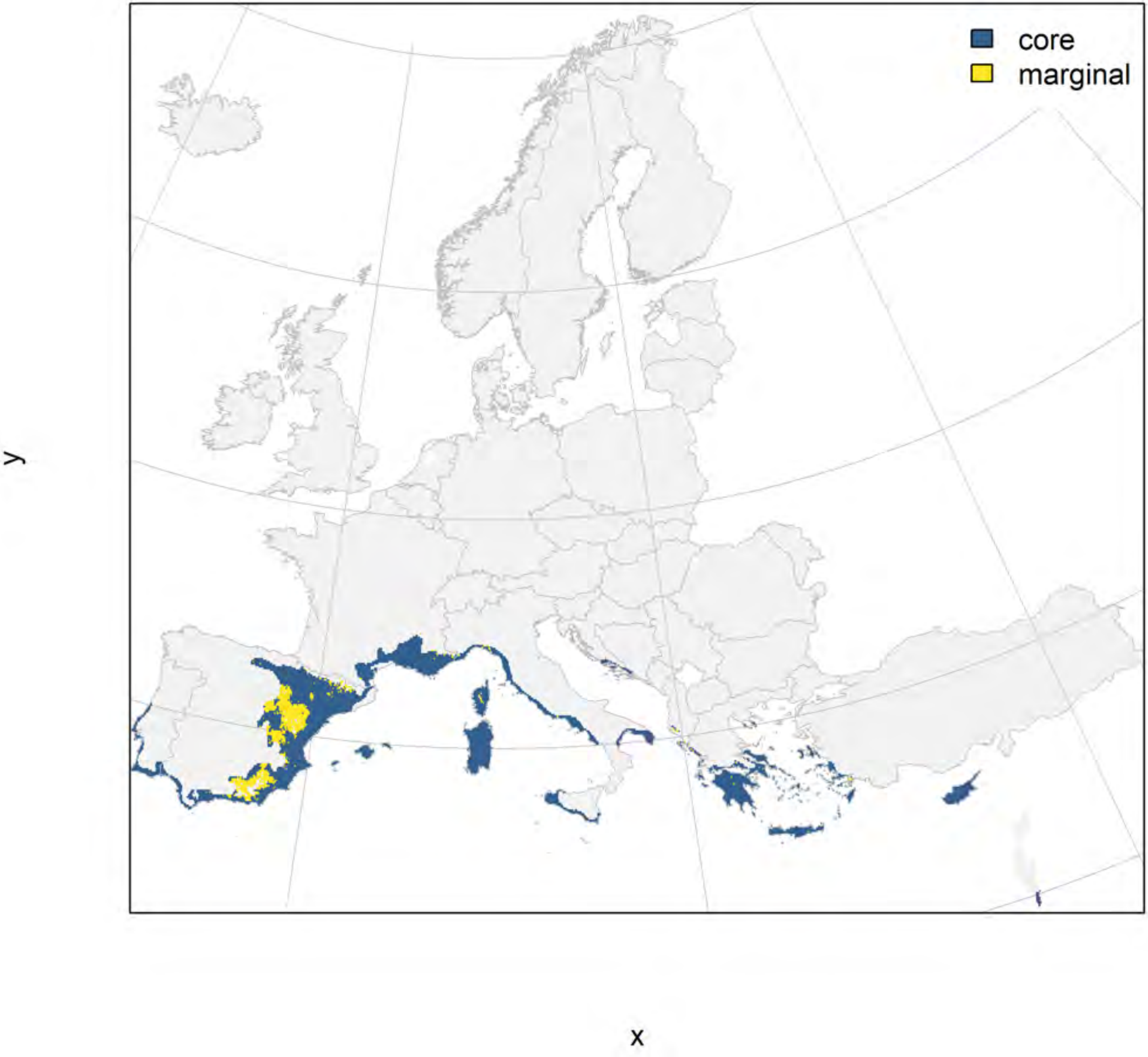

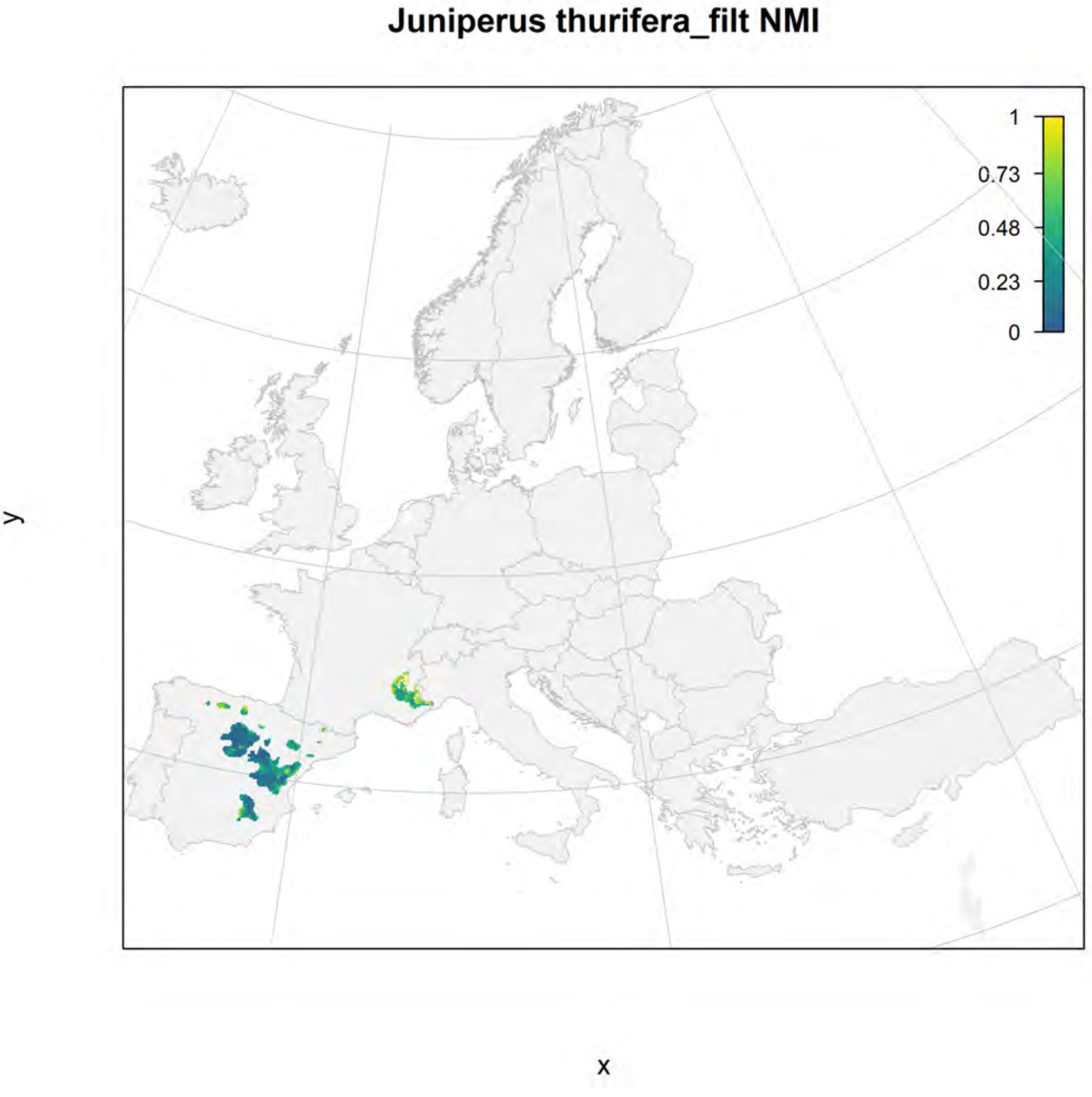
Juniperus thurifera_filt NMI
1
0.73
0.48
0.23
0
y
x

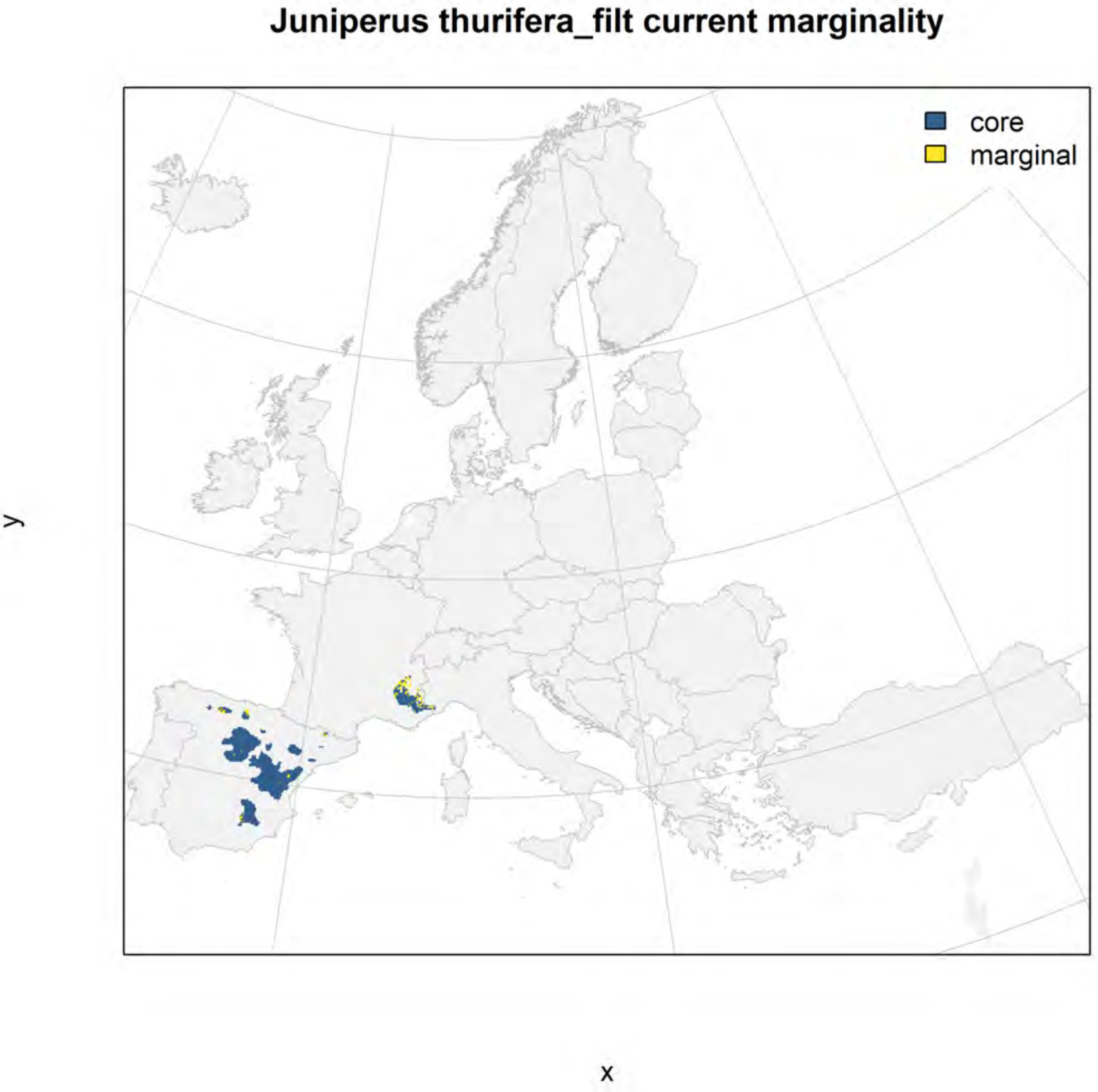
Juniperus thurifera_filt current marginality
core
marginal
y
x

#### Juniperus thurifera_filt future marginality

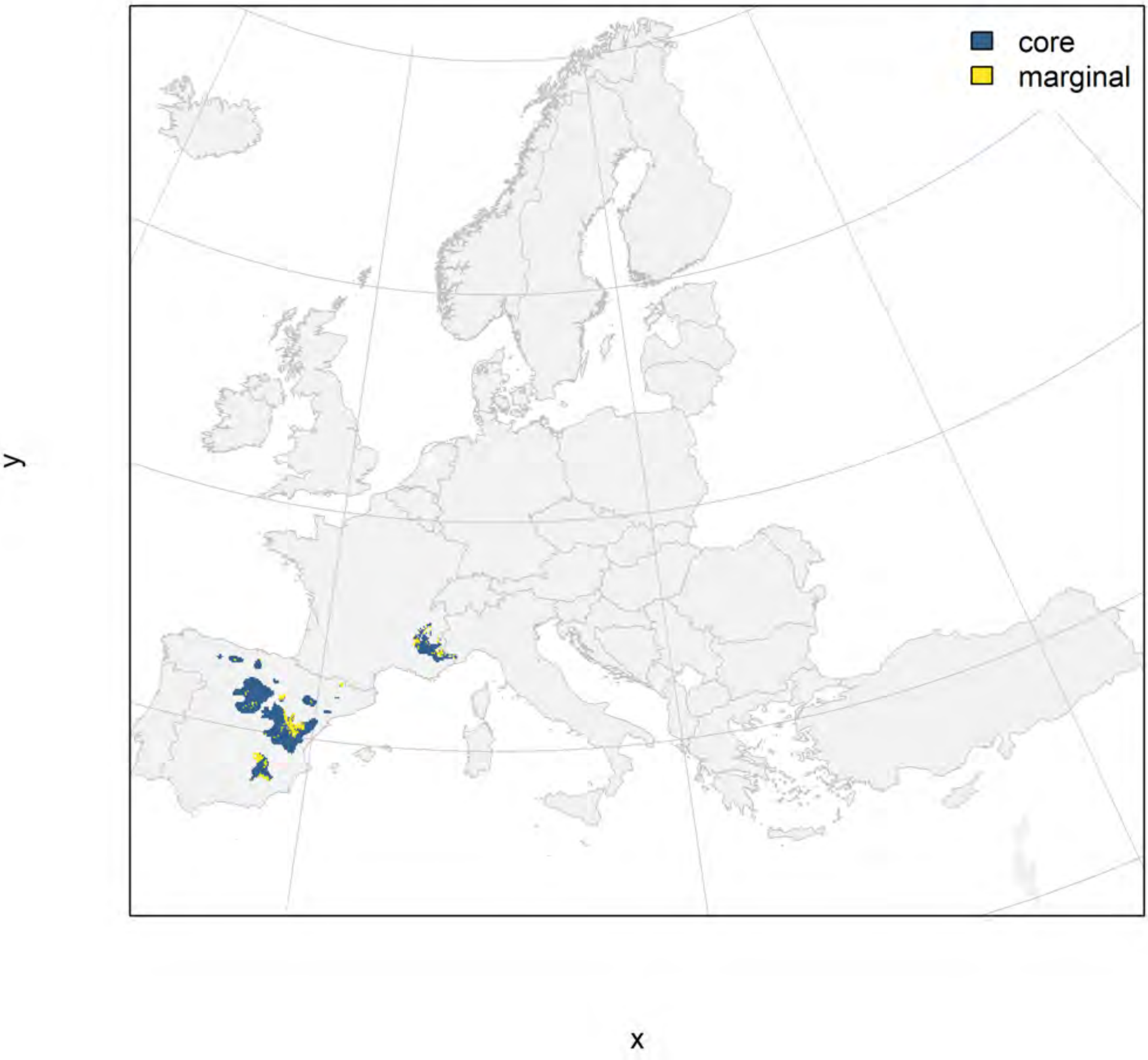

### Olea europaea_filt NMI

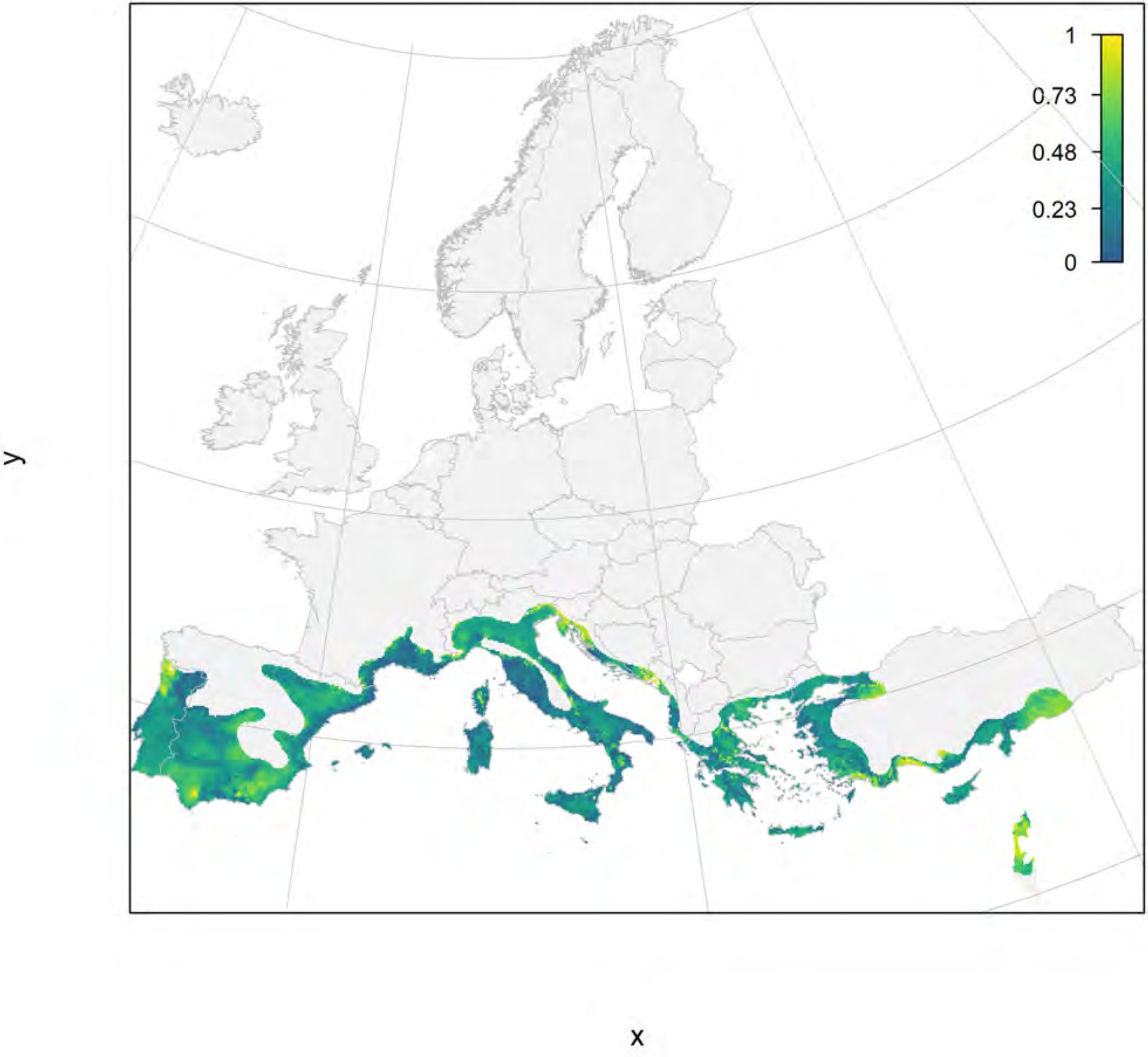

#### Olea europaea_filt current marginality

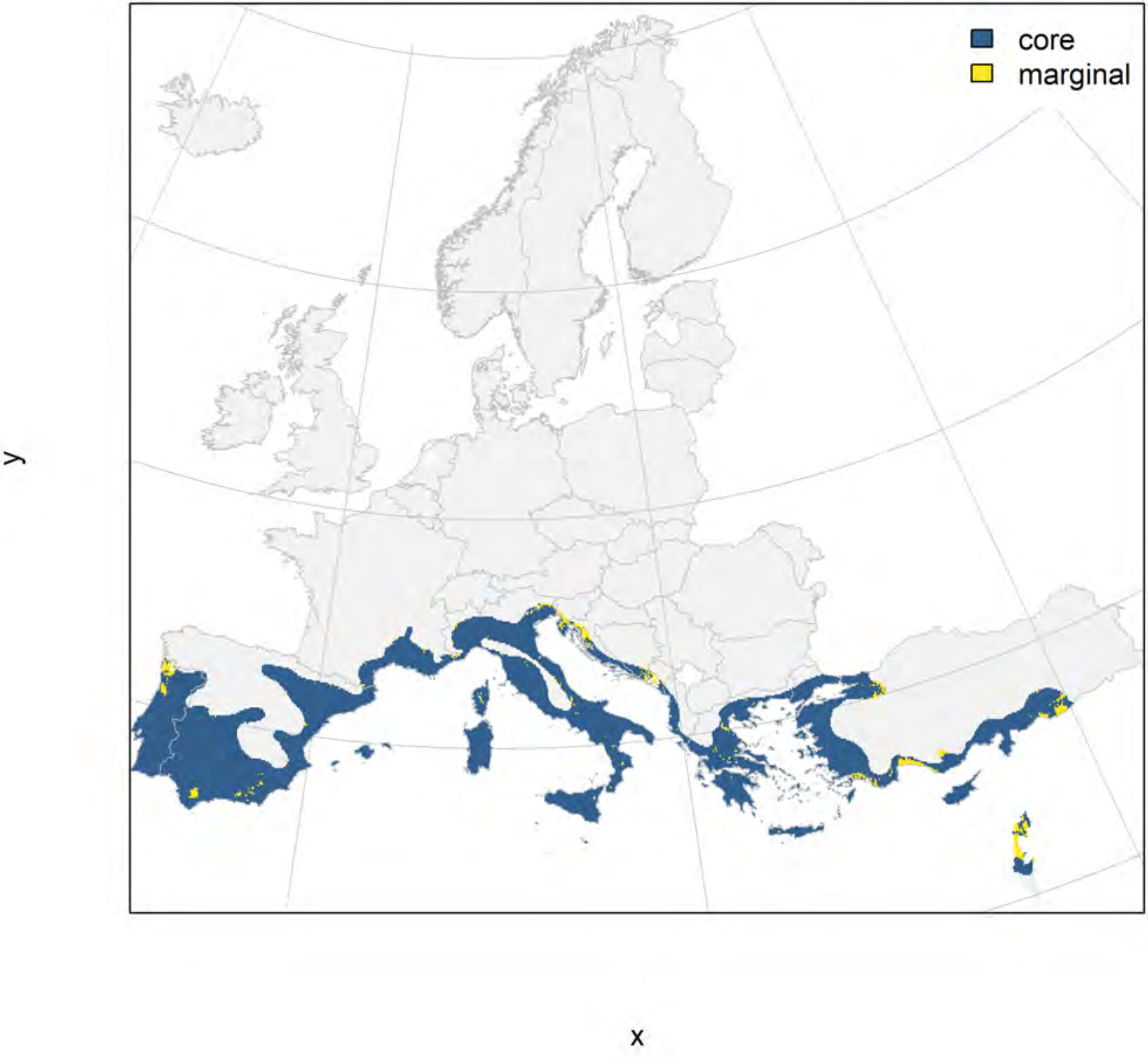

#### Olea europaea_filt future marginality

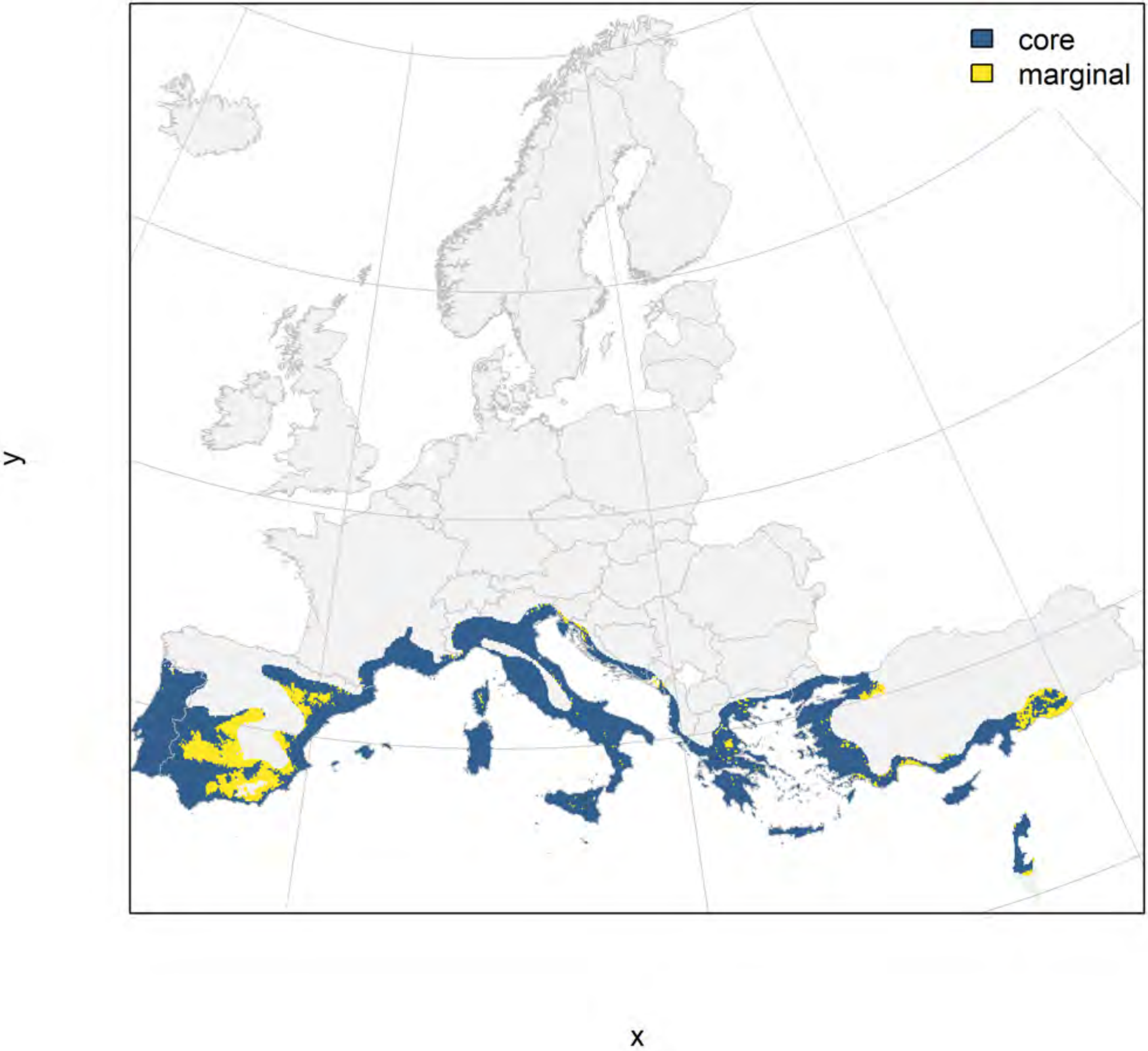

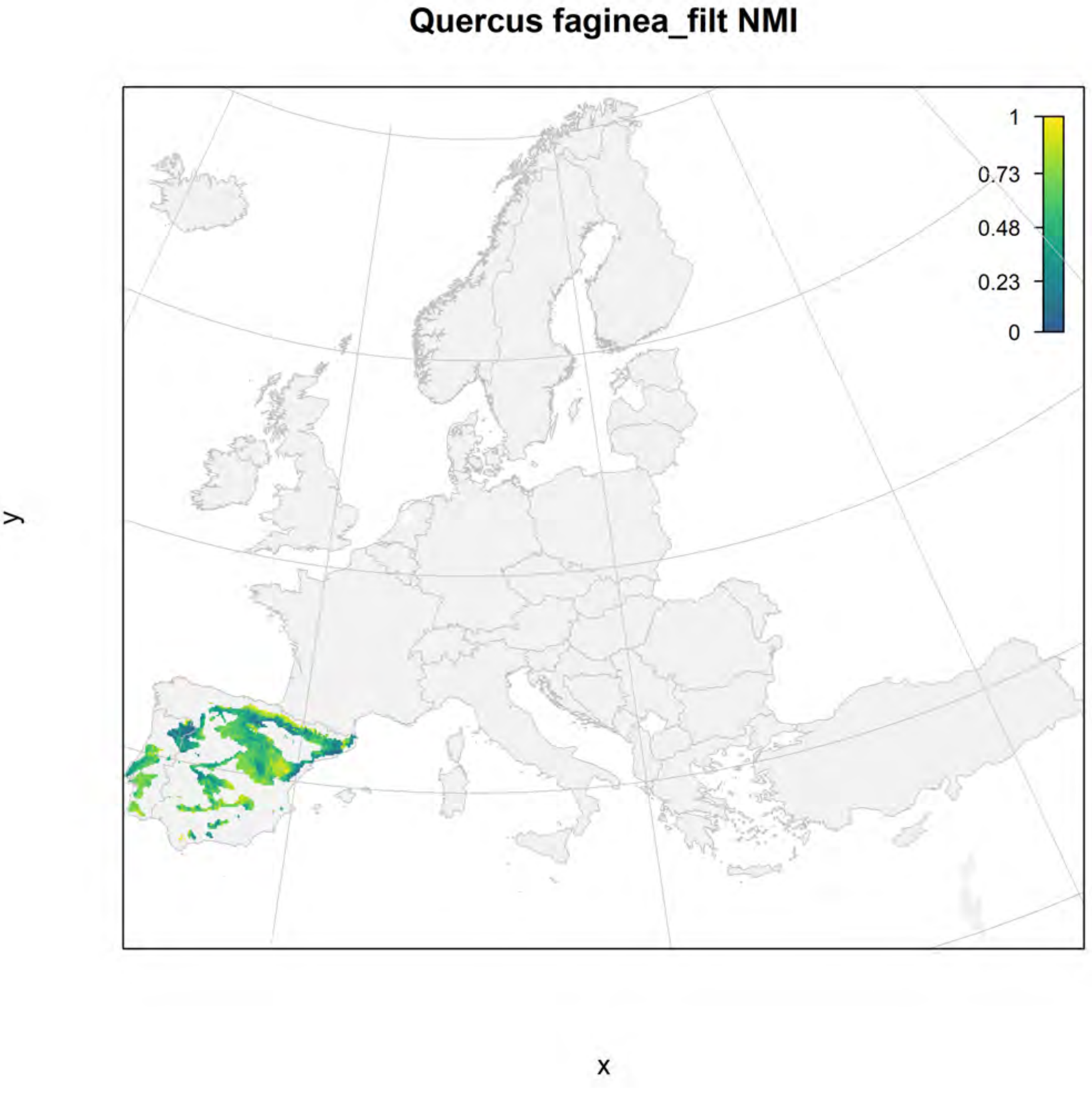
Quercus faginea_filt NMI
1
0.73
0.48
0.23
0
y
x

### Quercus faginea_filt current marginality

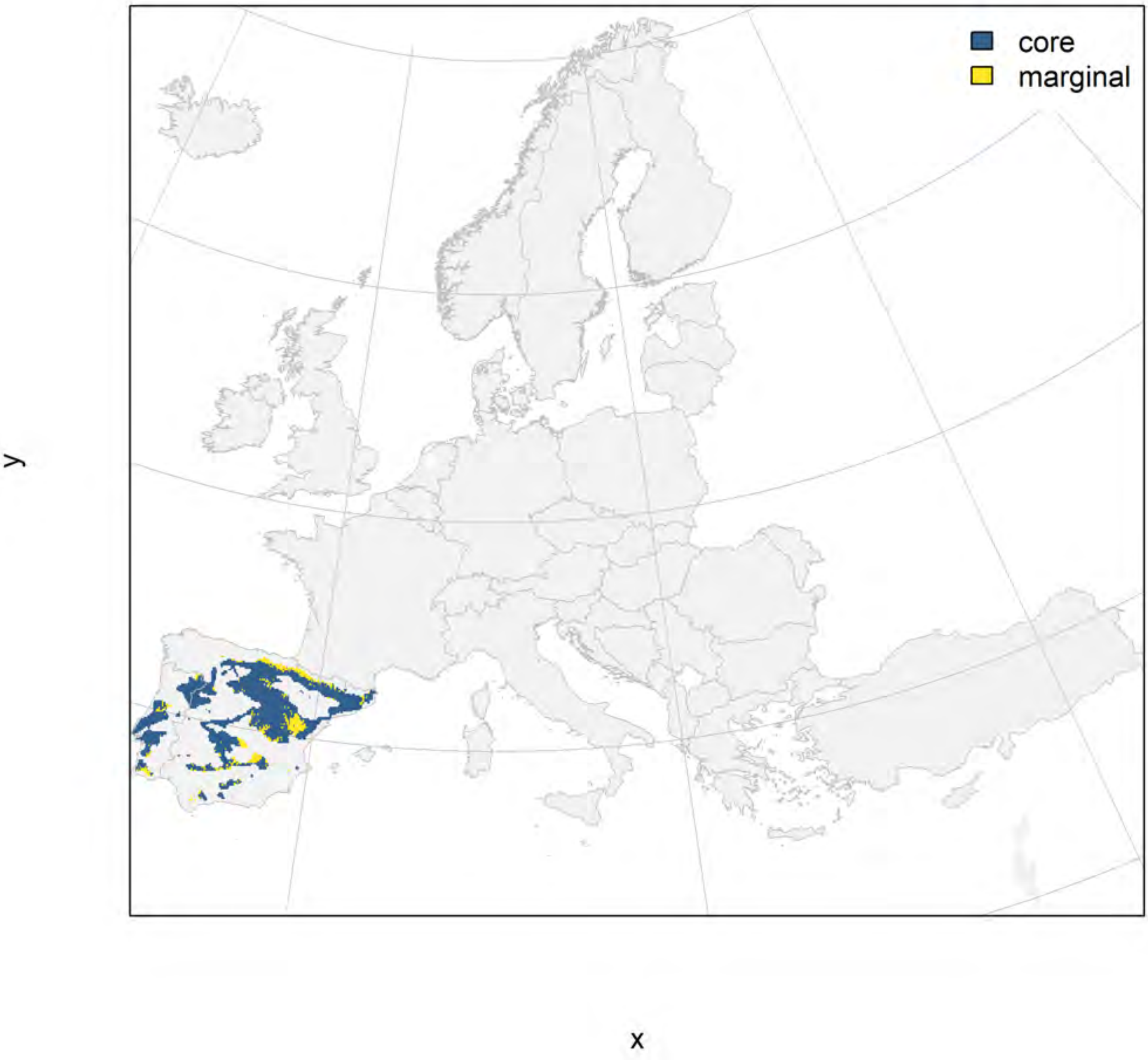

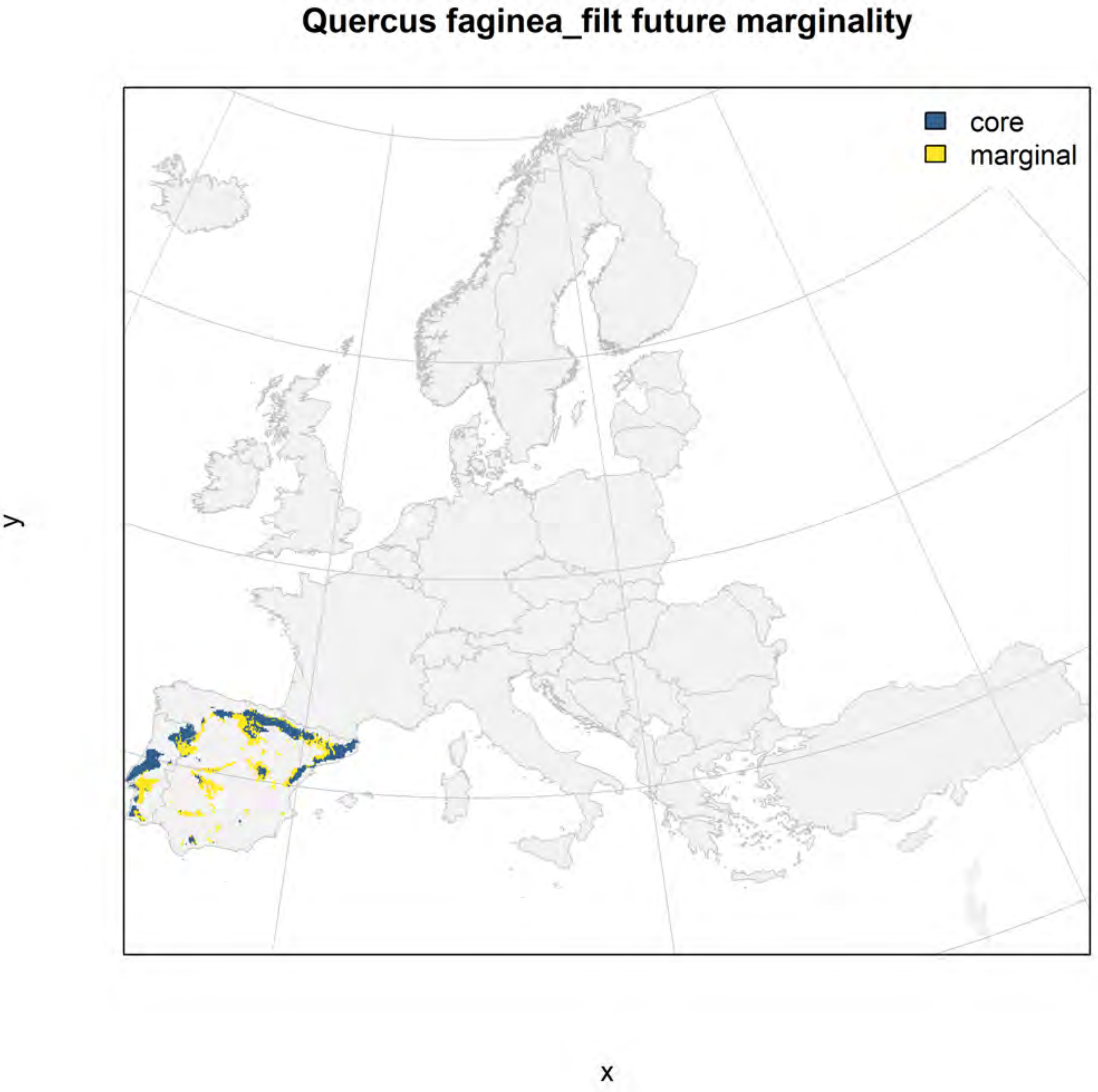
Quercus faginea_filt future marginality
core
marginal
y
x

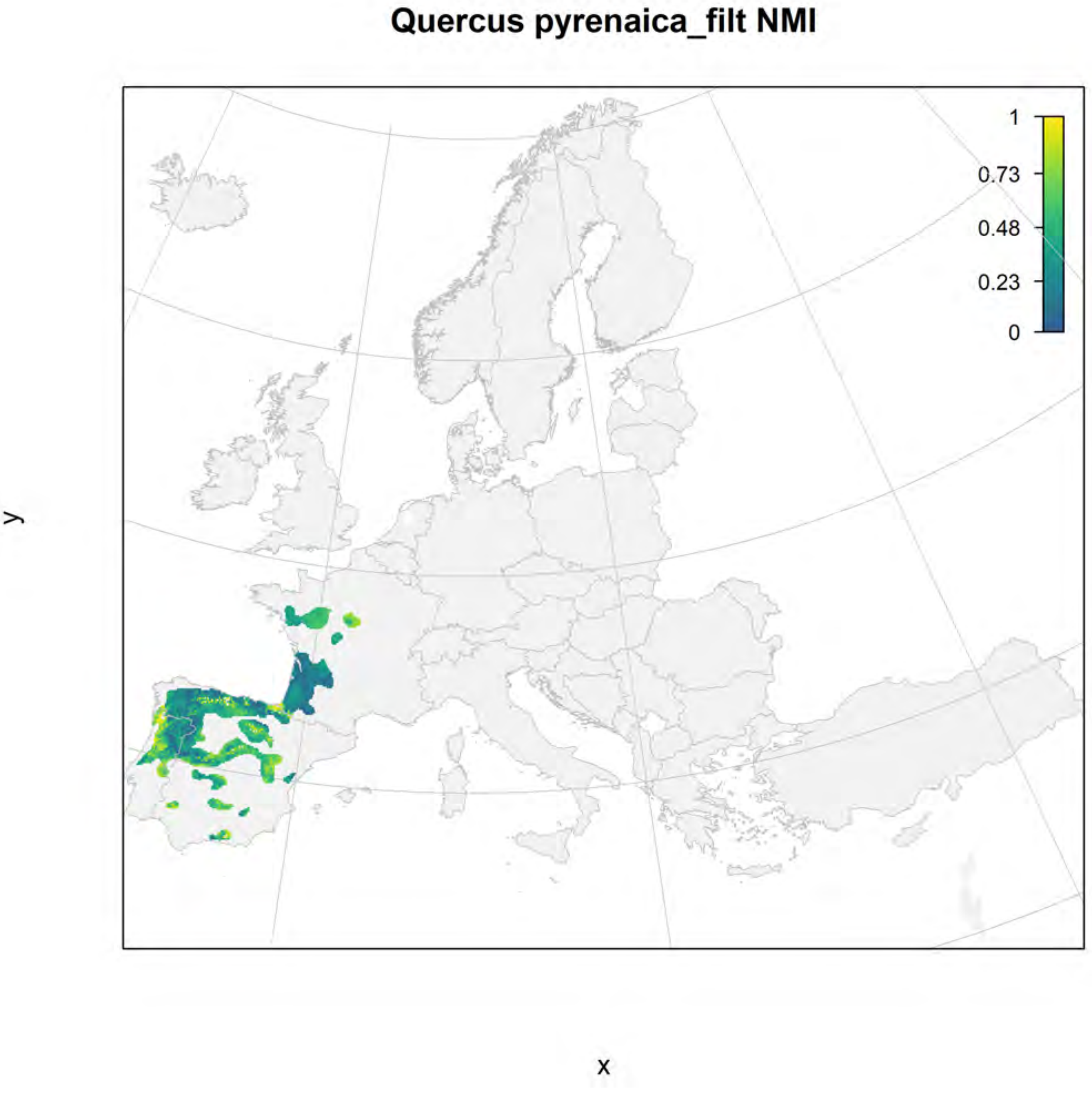
Quercus pyrenaica_filt NMI
1
0.73
0.48
0.23
0
y
x

#### Quercus pyrenaica_filt current marginality

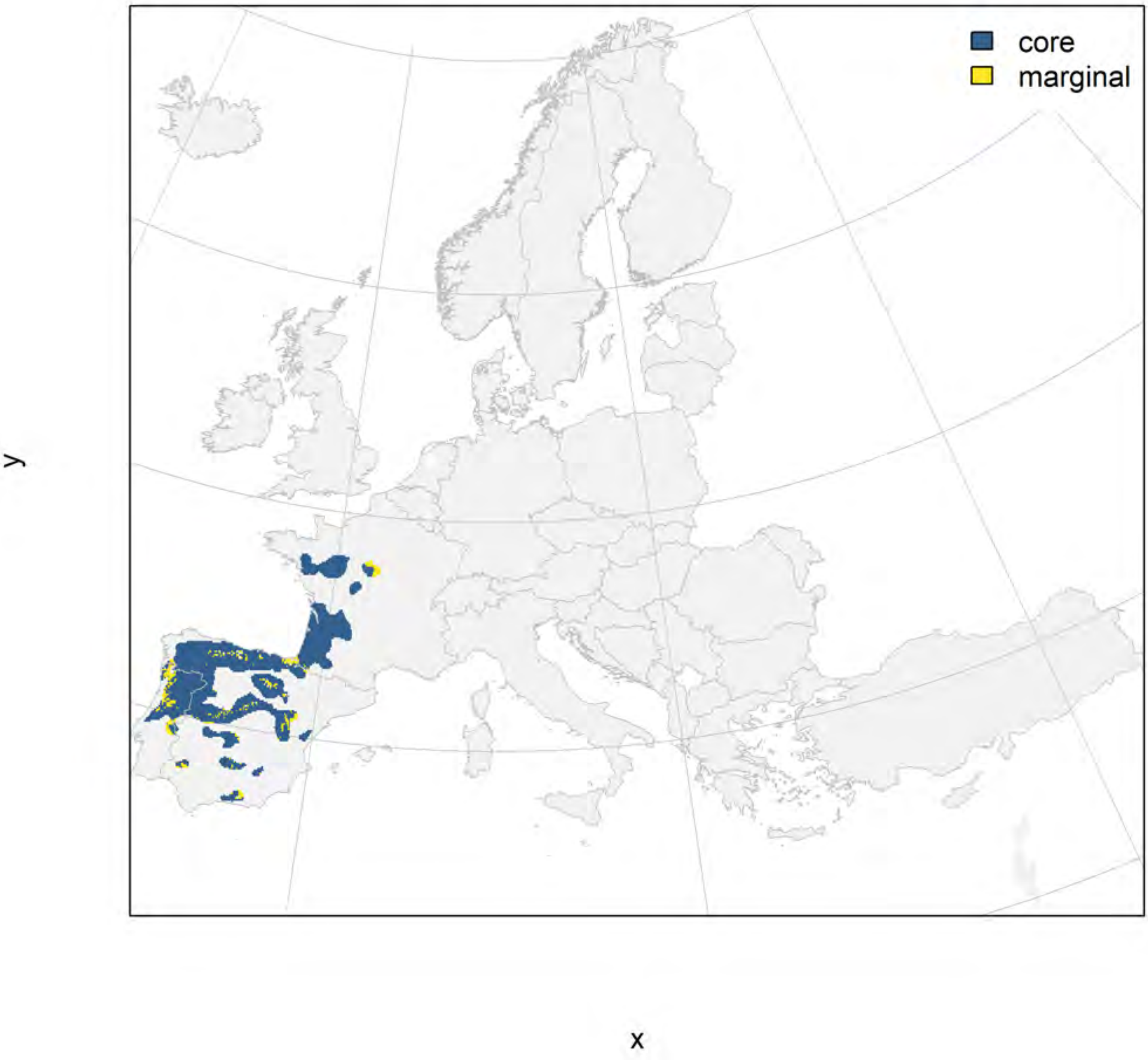

### Quercus pyrenaica_filt future marginality

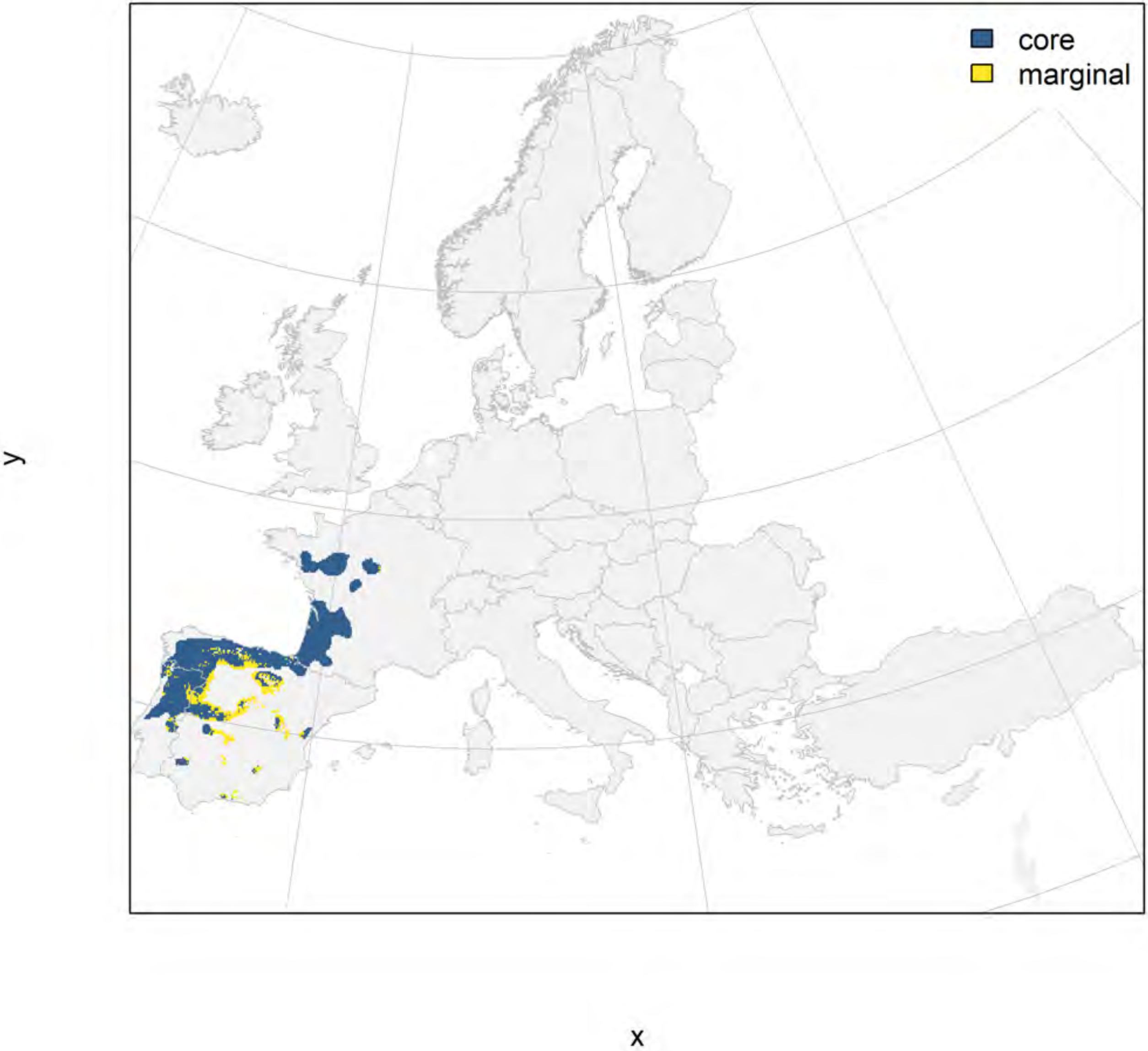

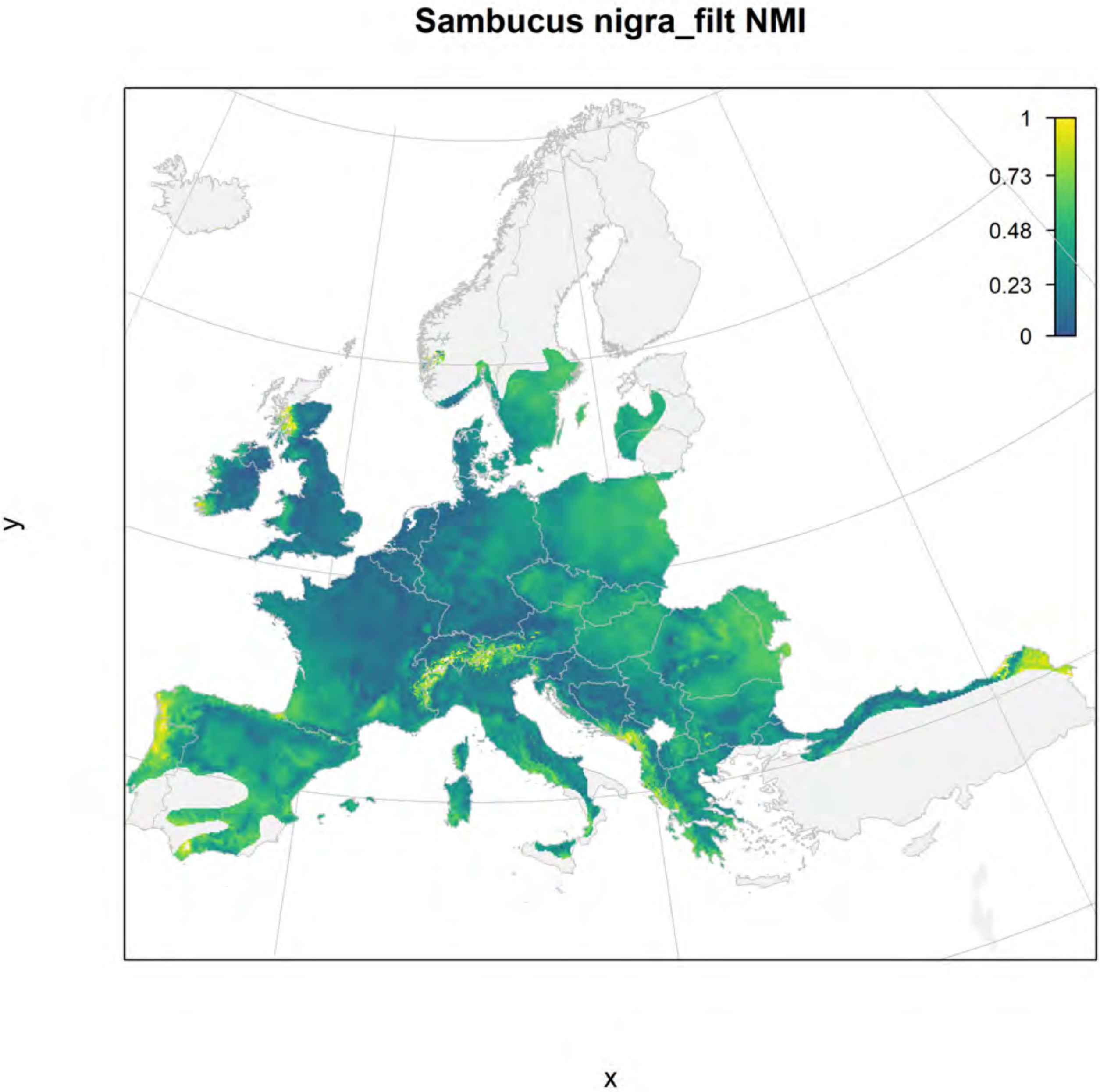
Sambucus nigra_filt NMI
1
0.73
0.48
0.23
0
y
x

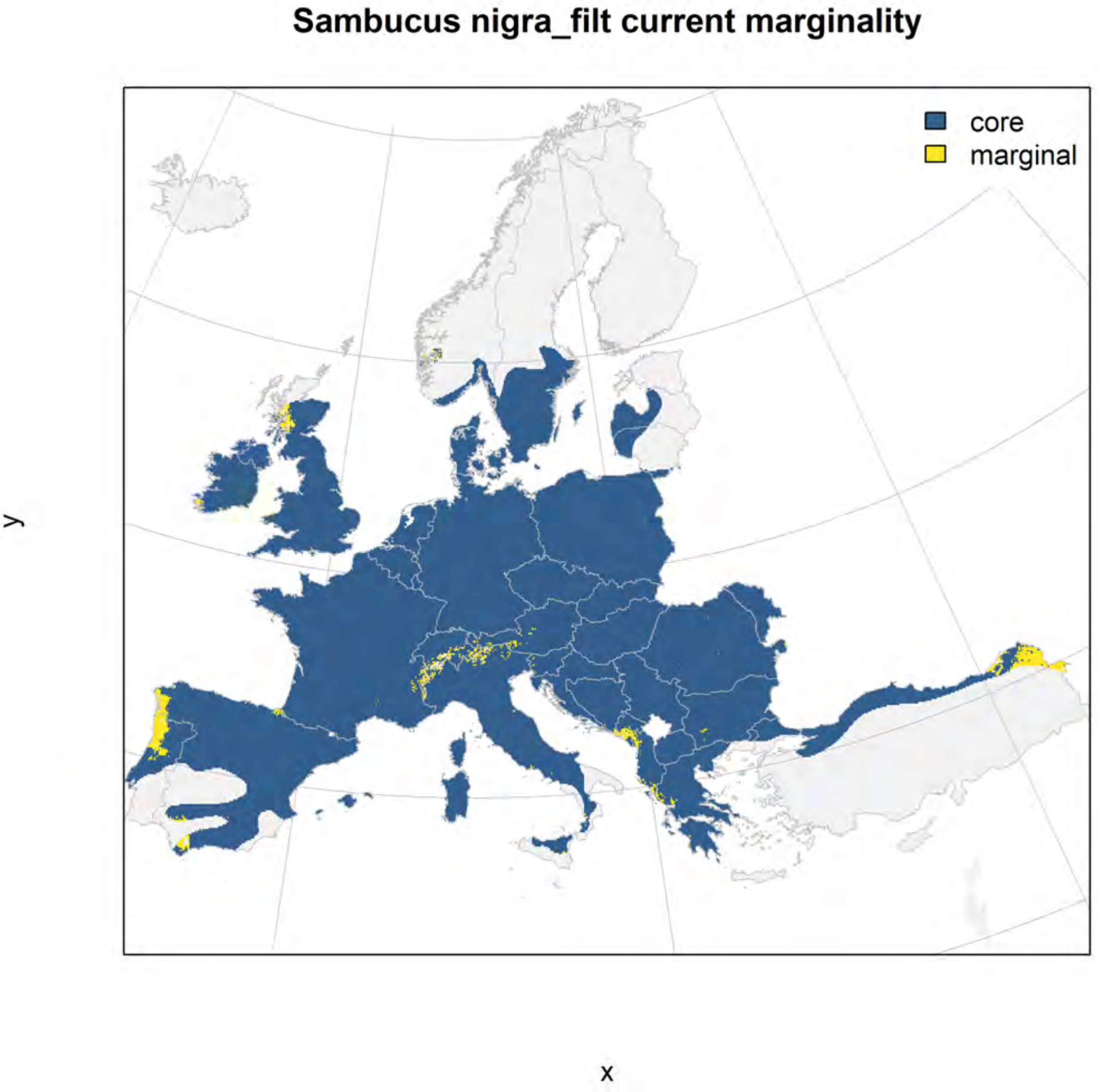
Sambucus nigra_filt current marginality
core
marginal
y
x

#### Sambucus nigra_filt future marginality

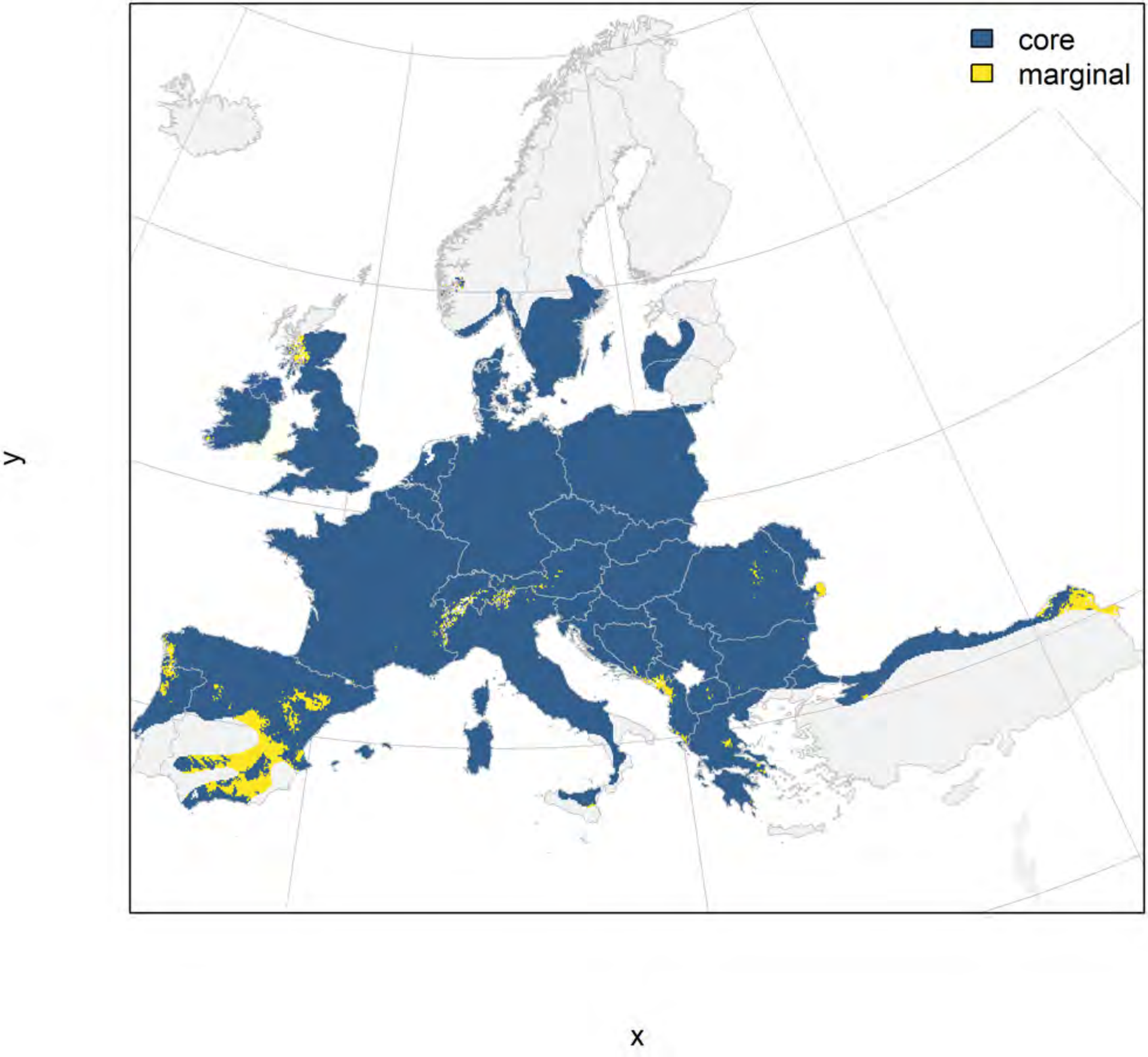

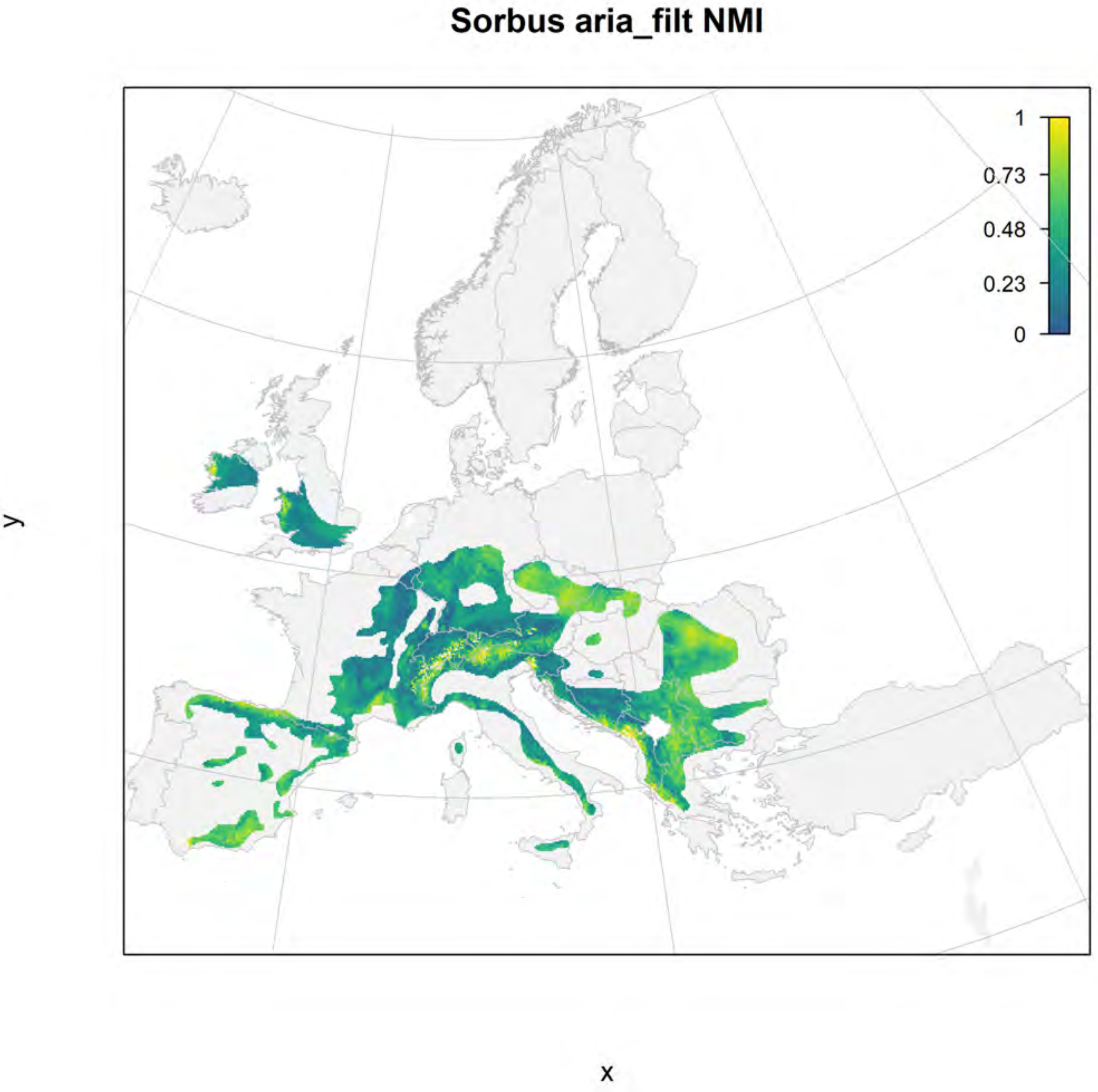
Sorbus aria_filt NMI
1
0.73
0.48
0.23
0
y
x

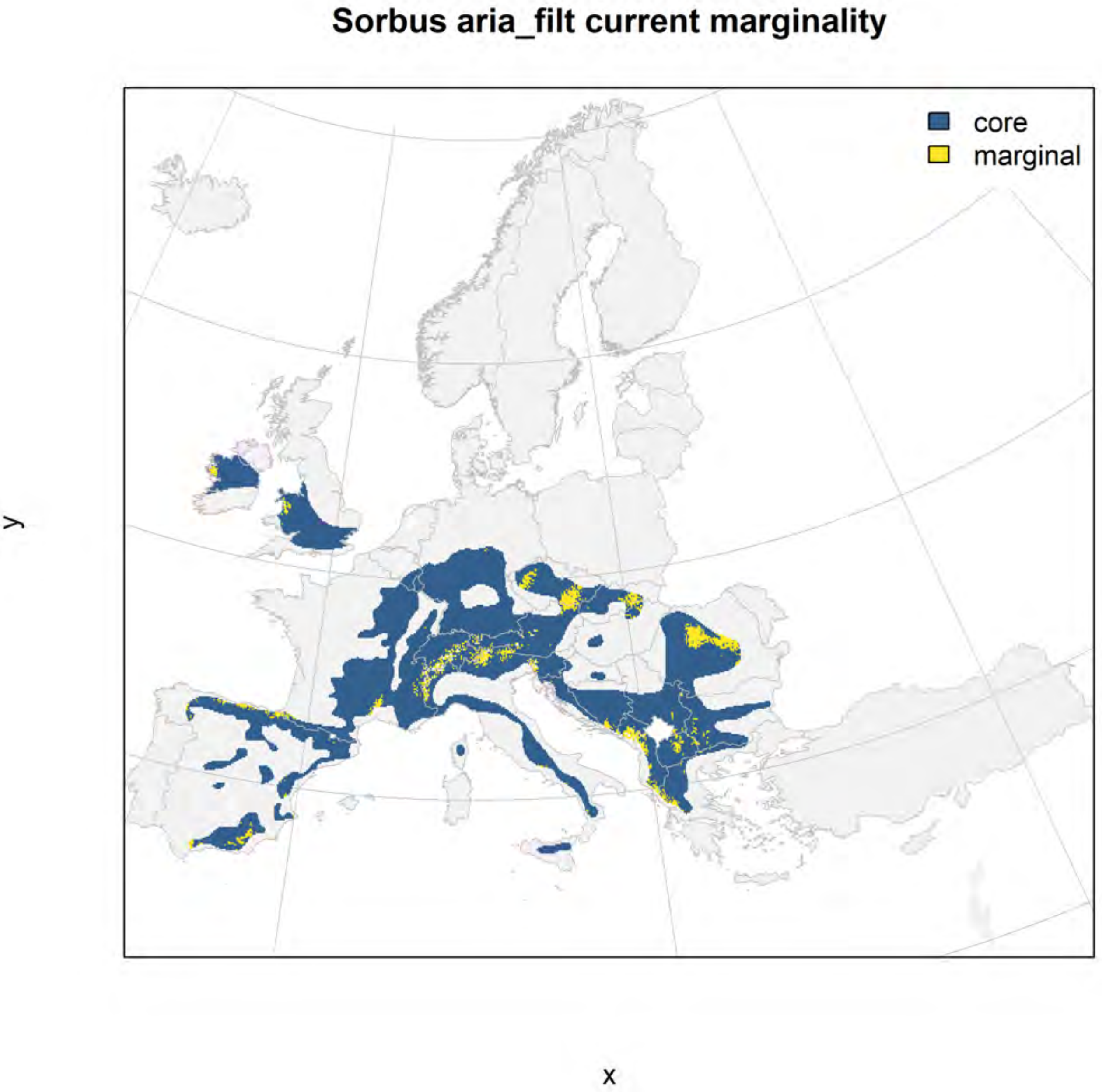
Sorbus aria_filt current marginality
core
marginal
y
x

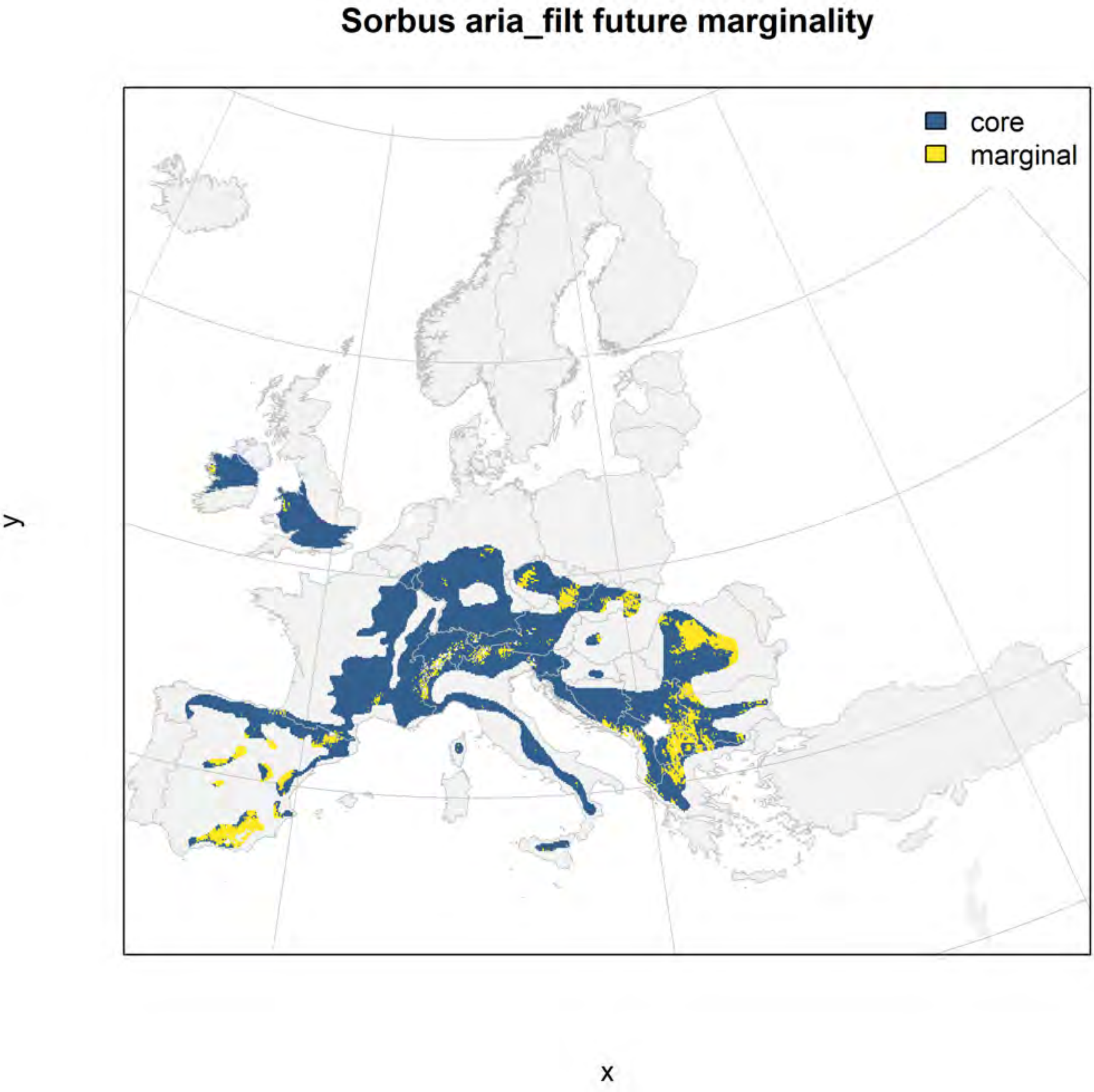
Sorbus aria_filt future marginality
core
marginal
y
x
