## Appendix S7 for "Conserving genetic diversity during climate change: Niche marginality and discrepant monitoring capacity in Europe"

### Extended Discussion

This study combines and reports two divergent types of data to reflect on the need and preparedness of European countries to monitor and address the effects of climate change on genetic diversity in species of conservation interest. The importance of genetic diversity in wild populations and their adaptive responses to changing environments is broadly recognized. Until now there has been no overview of historical and current efforts to monitor changes in PGD of species, to relate PGD monitoring to national characteristics, or to assess monitoring capacity for detecting potential genetic impacts of climate change. The overview we present here will assist future planning and implementation of PGD monitoring programs, and conservations actions more generally, by directing efforts at empirical genetic assessment, monitoring and management towards critical portions of environmental gradients. This will assist countries in meeting reporting requirements of the Directives of the European Union (EU) and the Convention on Biological Diversity by anticipating and accounting for ongoing habitat degradation due to climate change. In the future, programs for conserving genetic diversity should include monitoring across whole climate and other gradients. This will contribute to detection of genetic impacts of climate change in a wide variety of species of conservation interest and inform their management[1].

Prediction of effects of changing climate on adaptive potential of populations in the absence of species-specific understanding of the genetic architecture of adaptive traits is complicated[2]. Climate change may impact levels of additive genetic variation as well as genetic correlations and pleiotropy among adaptive life history traits[3]. For example, Etterson and Shaw[4] find in field experiments that genetic correlations among traits that confer adaptation limit responses to experimental conditions that simulate climate change. Further, the loss of range size when adaptation to changing climate fails increases extinction risk[5,6]. Despite these caveats, standing genetic variation has been important in adaptation to changing climate[7] and insufficient gene flow as climate change progresses can have detectable, detrimental demographic effects on populations[8]. While much monitoring will need to be done using demographic surrogates for empirically determined genetic variation[9], the loss of genetic diversity in populations at niche margins, whether through demographic processes in small populations or selection in large ones, should alert practitioners to a potential need for management action[10]. Well-planned management that accounts for the distribution of adaptive variation may reduce the need for more expensive management interventions, such as assisted gene flow, translocations and artificial colonization [11-13]. This suggests that understanding and prediction of climate-driven declines in environmental suitability and shifts of populations from or towards niche margin conditions are needed to provide a framework for effective monitoring actions nationally and across borders.  This will facilitate the detection and management of impacts of climate change on genetic diversity, adaptive potential, and the probability of population persistence.

Future evaluation of the extent, distribution and efficacy of genetic monitoring programs will need to account for and distinguish programs having the ability to predict and actively monitor changes in adaptive capacity[14,15]. Evidence suggests that relationships between decreases in environmental suitability, for example due to climate change, and functional genomic variation can predict demographic responses of species[8,16]. However, estimates of genetic offset due to climate change remain to be fully evaluated for usefulness to conservation efforts and their planning[17]. Within our dates of data collection, we did not sweep up such predictive studies as valid Category II monitoring. Many additional factors besides climate change per se, such as forest cover, landscape connectivity, and species interactions, may be highly relevant to the success of efforts to mitigate climate vulnerability and impacts on adaptive capacity[18,19], and to the management of climate change impacts generally.


1 Jensen, E. L. & Leigh, D. M. Using temporal genomics to understand contemporary climate change responses in wildlife. *Ecology and Evolution* **12** (2022). https://doi.org:10.1002/ece3.9340

2 Sgro, C. M., Lowe, A. J. & Hoffmann, A. A. Building evolutionary resilience for conserving biodiversity under climate change. *Evolutionary Applications* **4**, 326-337 (2011). https://doi.org:10.1111/j.1752-4571.2010.00157.x

3 Sgro, C. M. & Hoffmann, A. A. Genetic correlations, tradeoffs and environmental variation. *Heredity* **93**, 241-248 (2004). https://doi.org:10.1038/sj.hdy.6800532

4 Etterson, J. R. & Shaw, R. G. Constraint to adaptive evolution in response to global warming. *Science* **294**, 151-154 (2001). https://doi.org:10.1126/science.1063656

5 Thomas, C. D. *et al.* Extinction risk from climate change. *Nature* **427**, 145-148 (2004). https://doi.org:10.1038/nature02121

6 Urban, M. C. Accelerating extinction risk from climate change. *Science* **348**, 571-573 (2015). https://doi.org:10.1126/science.aaa4984

7 Lai, Y. T. *et al.* Standing genetic variation as the predominant source for adaptation of a songbird. *Proceedings of the National Academy of Sciences of the United States of America* **116**, 2152-2157 (2019). https://doi.org:10.1073/pnas.1813597116

8 Bay, R. A. *et al.* Genomic signals of selection predict climate-driven population declines in a migratory bird. *Science* **359**, 83-86 (2018). https://doi.org:10.1126/science.aan4380

9 Hoban, S. *et al.* Global Commitments to Conserving and Monitoring Genetic Diversity Are Now Necessary and Feasible. *Bioscience* **71**, 964-976 (2021). https://doi.org:10.1093/biosci/biab054

10 Fady, B. *et al.* Evolution-based approach needed for the conservation and silviculture of peripheral forest tree populations. *Forest Ecology and Management* **375**, 66-75 (2016). https://doi.org:10.1016/j.foreco.2016.05.015

11 Aitken, S. N. & Bemmels, J. B. Time to get moving: assisted gene flow of forest trees. *Evolutionary Applications* **9**, 271-290 (2016). https://doi.org:10.1111/eva.12293

12 Cook, C. N. & Sgro, C. M. Conservation practitioners' understanding of how to manage evolutionary processes. *Conservation Biology* **33**, 993-1001 (2019). https://doi.org:10.1111/cobi.13306

13 Van Rossum, F., Hardy, O. J., Le Pajolec, S. & Raspe, O. Genetic monitoring of translocated plant populations in practice. *Molecular Ecology* **29**, 4040-4058 (2020). https://doi.org:10.1111/mec.15550

14 Barbosa, S. *et al.* Integrative approaches to guide conservation decisions: Using genomics to define conservation units and functional corridors. *Molecular Ecology* **27**, 3452-3465 (2018). https://doi.org:10.1111/mec.14806
15 Capblancq, T., Fitzpatrick, M. C., Bay, R. A., Exposito-Alonso, M. & Keller, S. R. in *Annual Review of Ecology, Evolution, and Systematics, Vol 51, 2020* Vol. 51 *Annual Review of Ecology Evolution and Systematics* (ed D. J. Futuyma) 245-269 (2020).
16 Fitzpatrick, M. C., Chhatre, V. E., Soolanayakanahally, R. Y. & Keller, S. R. Experimental support for genomic prediction of climate maladaptation using the machine learning approach Gradient Forests. *Molecular Ecology Resources* **21**, 2749-2765 (2021). https://doi.org:10.1111/1755-0998.13374
17 Rellstab, C., Dauphin, B. & Exposito-Alonso, M. Prospects and limitations of genomic offset in conservation management. *Evolutionary Applications* **14**, 1202-1212 (2021). https://doi.org:10.1111/eva.13205
18 Williams, S. E., Shoo, L. P., Isaac, J. L., Hoffmann, A. A. & Langham, G. Towards an Integrated Framework for Assessing the Vulnerability of Species to Climate Change. *Plos Biology* **6**, 2621-2626 (2008). https://doi.org:10.1371/journal.pbio.0060325
19 Elmhagen, B. *et al.* Homage to Hersteinsson and Macdonald: climate warming and resource subsidies cause red fox range expansion and Arctic fox decline. *Polar Research* **36** (2017). https://doi.org:10.1080/17518369.2017.1319109
